# Screening of clustered regulatory elements reveals functional cooperating dependencies in Leukemia

**DOI:** 10.1101/680165

**Authors:** Salima Benbarche, Cécile K Lopez, Eralda Salataj, Cécile Thirant, Marie-Charlotte Laiguillon, Zakia Aid, Séverine Lecourt, Marion Antonini, Bryan Pardieu, Arnaud Petit, Alexandre Puissant, Julie Chaumeil, Thomas Mercher, Camille Lobry

**Author notes:** These authors contributed equally to the work (Julie Chaumeil, Thomas Mercher, Camille Lobry).

## Abstract

In the recent years, massively parallel sequencing approaches identified hundreds of mutated genes in cancer(*1*) providing an unprecedented amount of information about mechanisms of cancer cell maintenance and progression. However, while (it is widely accepted that) transformation processes result from oncogenic cooperation between deregulated genes and pathways, the functional characterization of candidate key players is mostly performed at the single gene level which is generally inadequate to identify these oncogene circuitries. In addition, studies aimed at depicting oncogenic cooperation involve the generation of challenging mouse models or the deployment of tedious screening pipelines. Genome wide mapping of epigenomic modifications on histone tails or binding of factors such as MED1 and BRD4 allowed identification of clusters of regulatory elements, also termed Super-Enhancers (SE)(*2*). Functional annotation of these regions revealed their high relevance during normal tissue development and cancer ontogeny(*3*). An interesting paradigm of the tumorigenic function of these SE regions comes from *ETO2-GLIS2*-driven acute megakaryoblastic leukemia (AMKL) in which the fusion protein ETO2-GLIS2 is sufficient to promote an aberrant transcriptional network by the rewiring of SE regions(*4*). We thus hypothesized that important regulatory regions could control simultaneously expression of genes cooperating in functional modules to promote cancer development. In an effort to identify such modules, we deployed a genome-wide CRISPRi-based screening approach and nominated SE regions that are functionally linked to leukemia maintenance. In particular, we pinpointed a novel SE region regulating the expression of both tyrosine kinases KIT and PDGFRA. Whereas the inhibition of each kinase alone affected modestly cancer cell growth, combined inhibition of both receptors synergizes to impair leukemia cell growth and survival. Our results demonstrate that genome-wide screening of regulatory DNA elements can identify co-regulated genes collaborating to promote cancer and could open new avenues to the concept of combined gene inhibition upon single hit targeting.

---

Super-Enhancers (SE) are clusters of regulatory elements characterized by high intensity of enhancer-related histone tail modifications such as histone H3 lysin 27 acetylation or lysine 4 mono-methylation (H3K27ac and H3K4me1 respectively) and binding of enhancer-associated factors such as the mediator complex, in particular MED1, or bromodomain-containing proteins such as BRD4(*5*). These clustered regulatory regions shape the transcriptional identity of specific tissues and cell types and their landscape often shifts in disease conditions particularly in cancer cells where they can control oncogene expression(*6, 7*). Whether these regions control expression of one or several genes and what are the genes directly controlled by these regions is mainly unknown. Additionally, whether these regions contribute to transformation or disease progression remains to be characterized. We hypothesized that SE could control and regulate expression of clusters of genes that might collaborate in cancer progression and/or maintenance and that impeding with their activity could reveal this synergistic action. We therefore designed a screening methodology to unbiasedly identify fundamental SE involved in cancer cell growth and survival (Fig. 1A). To this end, we used the CRISPRi methodology that relies on the use of a deactivated Cas9 fused to a KRAB domain(*8, 9*). Properly targeted dCas9-KRAB fusion can trigger the formation of local heterochromatin and impede with enhancer activity(*10*).

**Fig. 1.**
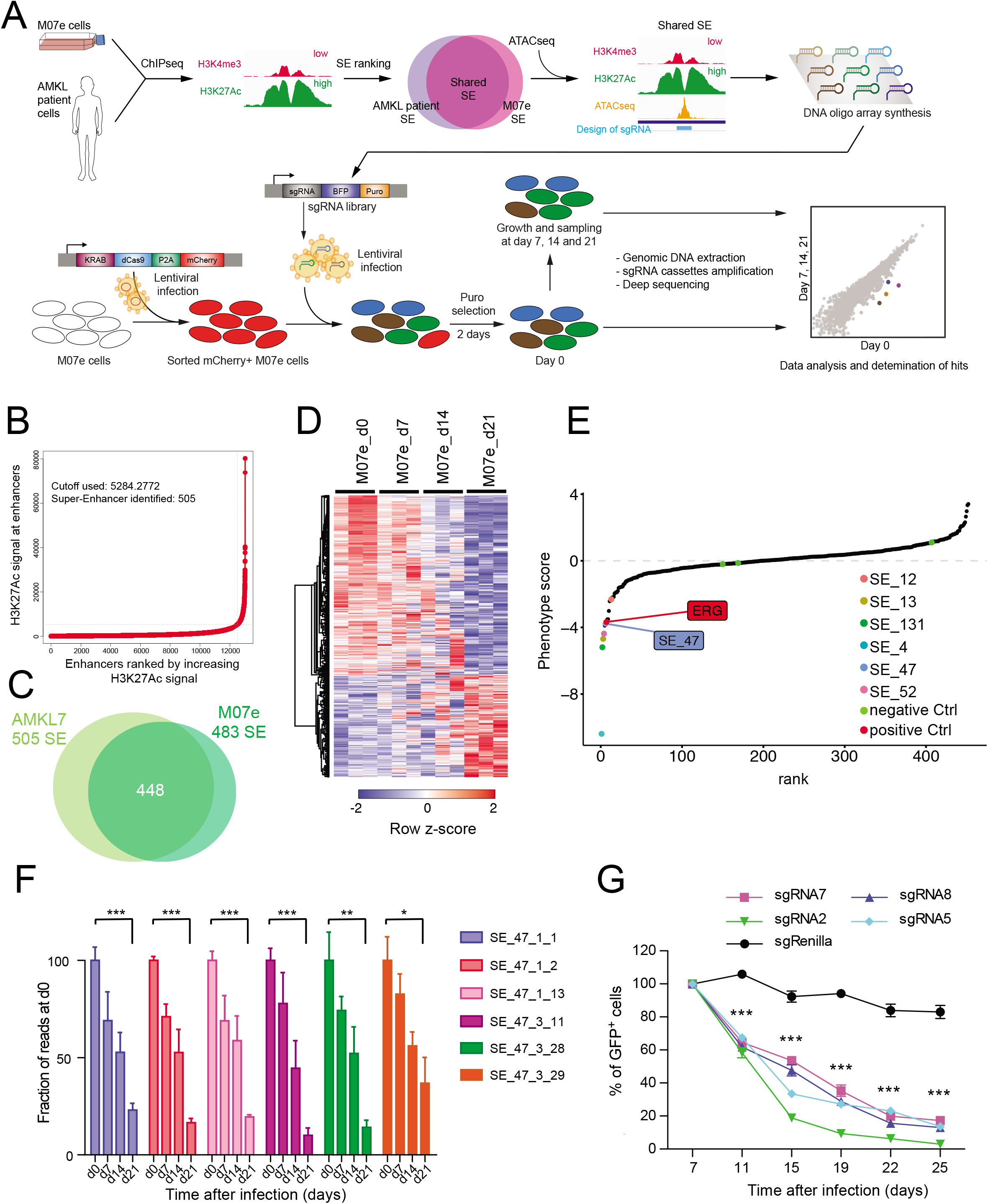
Genome wide CRISPRi screen identifies essential Super Enhancers for leukemia growth. **(A)** Schematic illustration of super enhancers (SE) screening strategy using CRISPRi in AMKL. **(B)** Distribution of H3K27Ac ChIP-seq density across enhancers, the 505 enhancers located above the tangent with high level of signal represent Super Enhancers **(C)** Venn diagram depicting the overlap between SE in M07e and AMKL7 cells. **(D)** Heatmap representation of 474significantly enriched and 278 significantly depleted sgRNAs among three replicates SE screening experiments in M07e cells. Representation of sgRNA with a minimum of 40x coverage and p-value < 0,05. **(E)** Pooled negative-selection screening depicting changes in representation of all SE ranked by the average of their depletion or enrichment score of all sgRNAs across the three replicates at day 21 compared to day 0. Significantly depleted SE are highlighted in color. Position of SE_47 and ERG positive control are shown. Negative controls are marked in green. **(F)** Bargraph representing variations of normalized counts of sgRNA targeting SE_47. Data are represented as ratio of counts at day 0. Data are represented as mean± SEM, statistical significance is determined using Student’s *t*-test, * p<0.05, ** p<0.01, *** p<0.001 **(G)** Percentage of GFP^+^ M07e cells following CRISPRi targeting of SE_47 with indicated sgRNAs compared to control sgRenilla and normalized to day 7 after infection. Mean ± SEM, n=3, significance is determined using Student’s *t*-test, *** p<0.001.

To functionally screen SE, we decided to use the ETO2-GLIS2 (EG) fusion driven model of acute megakaryoblastic leukemia (AMKL)(*11*). AMKL driven by this fusion have poor prognosis and EG fusion was recently shown to be the major oncogenic driver and strongly associated with SE(*4*). To define which SE to target, we performed H3K27ac ChIP-sequencing in AMKL patient-derived cells and called SE using ROSE algorithm(*5, 6*). In total, 505 SE were identified in these patient derived cells (Fig. 1B), of which 448 overlapped with previously defined(*4*) in the ETO2-GLIS2-expressing patient-derived M07e cell line (Fig. 1C). Subsequently, the design of sgRNAs targeting open chromatin regions, as defined by ATAC-sequencing peaks within these SE, led to a library of 7087 sgRNAs (Table S1). Screening was performed in triplicate in M07e cells stably expressing the dCas9-KRAB fusion. Analysis of biological replicates showed that the representation of 474 sgRNAs targeting 265 SE was significantly decreased over time whereas 278 sgRNAs targeting 185 SE were significantly upregulated (Fig. 1D, Fig. S1A). Maximum likelihood of enrichment analysis revealed that six SE markedly altered M07e cells growth upon CRISPRi inhibition (Fig. 1E, Fig. S1, B and C). Three sgRNAs targeting ERG which is required for ETO2-GLIS2^+^ AMKL cell growth(*4*), were used as positive controls and were significantly depleted over time (Fig. S1D) whereas sgRNAs targeting genes not present in the human genome (Luciferase, Renilla, mouse Lin28) showed no significant variation (Fig. S1E). Among top hits of the screen we decided to investigate SE_47 (Fig. 1E), located 5’ to the *KIT* gene on chromosome 4 (Fig. S1F) because of its location and because six sgRNAs targeting the two main H3K27ac peaks of this SE were significantly underrepresented (Fig. 1F). To validate these results, we designed 4 additional sgRNAs targeting these two regions of SE_47 (Fig S2A) and performed independent CRISPRi experiments in M07e cells stably expressing the dCas9-KRAB fusion. Expression of these 4 sgRNAs reproduced the loss of representation phenotype observed in the screen, confirming that CRISPRi targeting of SE_47 impairs M07e cell growth or survival (Fig. 1G).

To identify which genes are regulated by this SE, we performed RNA-sequencing analysis after short kinetics of CRISPRi inhibition (48h after transduction), in order to limit secondary inhibitory effects. This analysis showed that short term inhibition of this SE induces very few transcriptional changes and mainly *KIT* and *PDGFRA* proximal genes were significantly downregulated (Fig. 2A). The above results were confirmed by performing quantitative PCR with four independent sgRNAs that showed significant inhibition of *KIT* and *PDGFRA* expression when compared to non-targeting control (Fig. 2B). To control for the on-target specificity of our CRISPR guides, we performed ATAC-sequencing on M07e cells expressing each of these sgRNAs and revealed that CRISPRi induces significant loss of chromatin accessibility only at sgRNA-targeted regions without affecting additional region of the same locus or other regions of the genome (Fig. S2A) suggesting that CRISPRi does not induce unspecific spreading of heterochromatin. Additionally, we performed CRISPRi using 2 independent sgRNAs in HEL-5J20 cell line lacking the activity of this enhancer and expressing low level of *KIT*. CRISPRi expression in this cell line did not induce neither inhibition of *KIT* expression nor impaired growth of the cells suggesting that observed phenotypes in M07e cells are on target (Fig. S2, B and C).

**Fig. 2.**
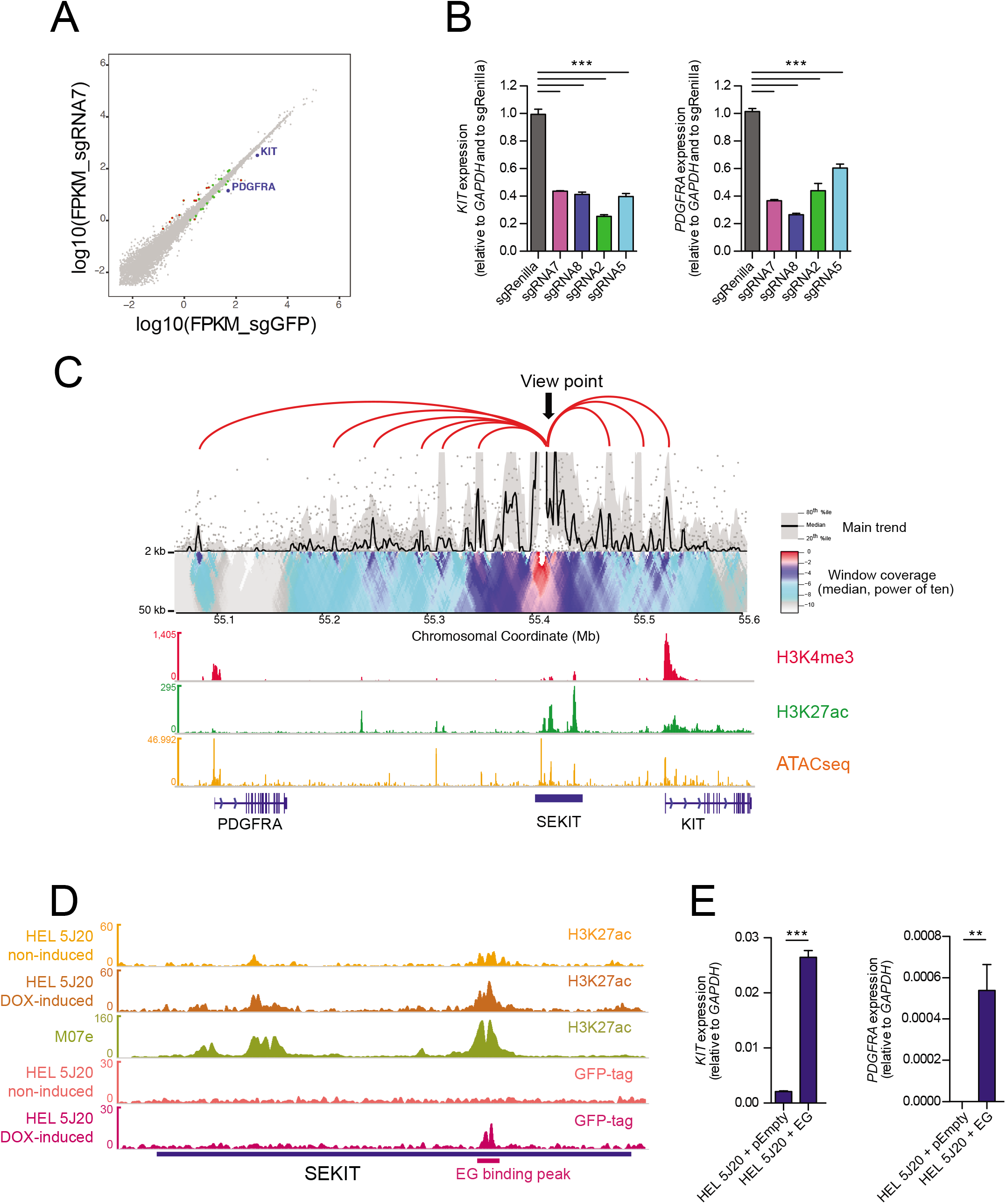
SEKIT controls *KIT* and *PDGFRA* expression and is induced by ETO2-GLIS2 fusion. **(A)** Scatter plot showing genes whose expression is significantly altered following short kinetic (48h post transduction) CRISPRi targeting of SEKIT with sgRNA7 compared to control sgGFP in M07e cells. Green dots p<0.05, red dots p<0.05 and >2-fold change. *KIT* and *PDGFRA* genes are highlighted. **(B)** qPCR showing *KIT* and *PDGFRA* expression following CRISPRi targeting of SEKIT with indicated sgRNAs compared to control sgRenilla in M07e cells. Mean ± SEM, n=3, significance is determined using Student’s *t*-test, *** p<0.001. **(C)** 4C-seq domainogram with viewpoint on SEKIT showing cis-interactions with proximal elements around SEKIT locus (top panel) and gene tracks showing normalized read density histograms of H3K4me3 ChIP-seq, H3K27ac ChIP-seq and ATAC-seq (bottom panel) in M07e cells. Read densities are shown as unique reads per million. **(D)** Gene track showing normalized read density histograms of H3K27Ac ChIP-seq analysis at SEKIT locus upon ETO2-GLIS2 (EG) expression induction in HEL 5J20 cells compared to non-induced cells and to M07e cells (orange, brown and green tracks respectively) and read densities of GFP ChIP-seq in non-induced versus doxycycline induced 5J20 cells (coral and magenta track respectively) showing EG binding in SEKIT locus. Read densities are shown as unique reads per million. **(E)** qPCR showing *KIT* and *PDGFRA* expression upon doxycycline-induced ETO2-GLIS2 expression in HEL 5J20 cells compared cells transduced with empty vector. Mean ± SEM, representative of two independent experiments in triplicate, significance is determined using Student’s *t*-test, ** p<0.01, *** p<0.001.

To investigate direct regulation of proximal *KIT* and *PDGFRA* genes by SE_47 we conducted chromatin conformation capture experiments(*12, 13*) (4C-sequencing). Using a bait located in SE_47, we identified directly interacting proximal regions including *KIT* and *PDGFRA* promoters (Fig. 2C). These results demonstrated that SE_47 is in close physical contact with *KIT* and *PDGFRA* and controls their expression. Therefore, the aforementioned SE_47 was named “SEKIT” thereafter.

We next wondered whether this SE is normally found in other hematopoietic cells expressing the *KIT* gene. To this end, we performed ChIP-sequencing analyses for H3K27ac in cell lacking ETO2-GLIS2, including human CD34^+^ cord blood hematopoietic stem and progenitor cells (HSPC) expressing high level of *KIT*, the AML1-ETO fusion expressing Kasumi-1 cell line also highly expressing *KIT* and the HEL 5J20 cell line expressing low level of *KIT*. We then compared profiles obtained with ETO2-GLIS2^+^ M07e cell line, AMKL7 patient cells and KIT^neg^ CD14^+^ monocytes from ENCODE dataset. This analysis showed that ETO2-GLIS2 negative cell types have no H3K27ac peaks located in SEKIT region and that instead, *KIT* expressing cells show H3K27ac peaks located on the 3’ of *KIT* (Fig. S3), in a region previously described as a *KIT* enhancer(*14*). These results showed that SEKIT is not active in wild type HSPC, suggesting that SEKIT activity is controlled by *ETO2-GLIS2* expression. To address this question, we generated a stable HEL 5J20 cell line expressing doxycyclin-inducible GFP-tagged ETO2-GLIS2 fusion (Fig. S4A). We performed ChIP-sequencing from the GFP-tag and H3K27ac upon induction of the fusion expression and observed a marked increase in H3K27ac compared to non-induced cells, with a profile similar to M07e, and identified a peak of ETO2-GLIS2 binding within SEKIT (Fig. 2D). Quantitative PCR analyses showed that induction of *ETO2-GLIS2* expression was accompanied by a strong upregulation of *KIT* and *PDGFRA* expression (Fig. 2E). Additionally, *ETO2-GLIS2* inhibition in M07e cells using a Nervy Homology Region 2 (NHR2)-interfering peptide (NC128)(*4, 15*) showed inhibition of *KIT* and *PDGFRA* expression (Fig. S4B) further indicating that the fusion activity is required to induce SEKIT activity and proximal *KIT* and *PDGFRA* gene expression.

Taken together, these results demonstrate that ETO2-GLIS2 fusion binds and induces activation of an aberrant *de novo* enhancer that regulates concomitantly *KIT* and *PDGFRA* expression.

To assess the functional consequences of SEKIT on KIT and PDGFRA protein levels, we performed flow cytometry with antibodies coupled with fluorophores to quantify KIT and PDGFRA receptors cell surface expression. The results indicated that CRISPRi inhibition of SEKIT induced a significant loss of KIT and PDGFRA cell surface protein expression (Fig. S5, A and B).

To probe mechanisms responsible for the loss of representation phenotype observed upon CRISPRi inhibition of SEKIT, we performed cell cycle analysis and observed a significant increase of cell in G0 phase and a significant decrease of cells in G1 and S phase (Fig. 3, A and B). To gain further insight into this growth inhibition mechanism, we performed RNA-sequencing experiments upon CRISPRi inhibition after longer kinetics (96h post infection). These transcriptomic analyses showed a higher number of genes significantly modulated upon SEKIT inhibition (Fig. S6A). Gene Set Enrichment Analysis (GSEA) showed that several genesets related to cytokine signaling where negatively enriched in SEKIT inhibited M07e cells, including genesets related to KIT and PDGF pathways (Fig. S6, B and C and Table S2). Interestingly, genesets related to AP1 (FOS/JUN) and AKT activity were also significantly downregulated upon SEKIT inhibition suggesting that these pathways are important downstream mediators of KIT and PDGFRA. Taken together, these data showed that SEKIT activity is required for proper growth of M07e cells through activation of KIT and PDGFRA signaling.

**Fig. 3.**
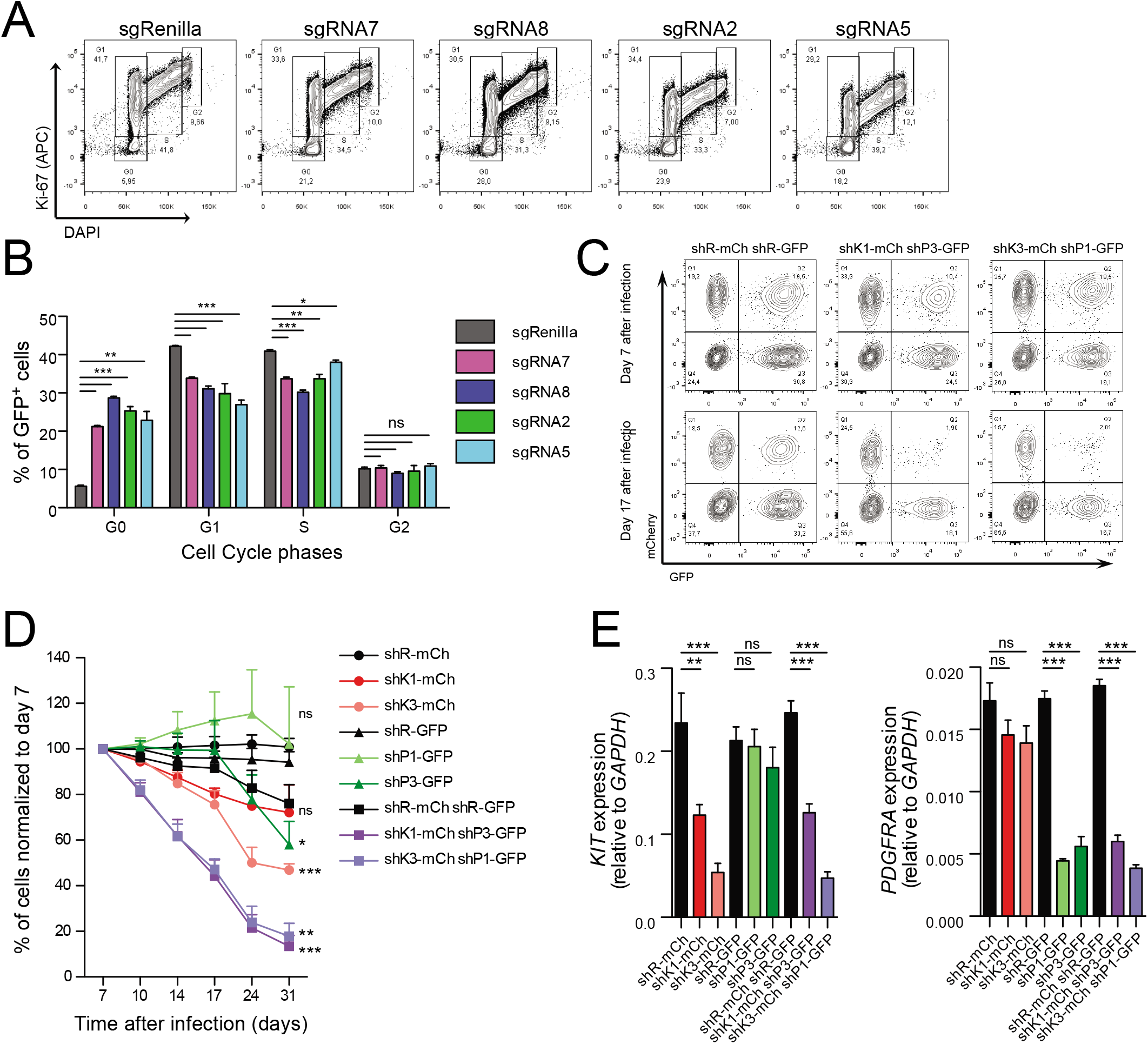
*KIT* and *PDGFRA* expression is required for M07e cell growth. **(A)** Representative flow cytometry analysis showing Ki-67 (APC) and DAPI staining gated on transduced GFP^+^ M07e cells following CRISPRi targeting of SEKIT with indicated sgRNAs and compared to control sgRenilla. **(B)** Quantification of cell cycle phases as analyzed in panel (a). Mean ± SEM, n=3, significance is determined using Student’s *t*-test, *** p<0.001. **(C-E)** Two independent shRNAs targeting either KIT (shK) or PDGFRA (shP) were expressed in M07e cells separately or in combination. shK are expressed with mCherry (mCh) in cells while shP are expressed with GFP. Corresponding shRNAs targeting Renilla (shR) were used as control. Cells are maintained in the presence of GM-CSF. **(C)** Representative flow cytometry analysis of shRNA expressing cells at day 7 and day 17 after infection. **(D)** Percentage of shRNA expressing M07e cells normalized to day 7 after infection. Mean ± SEM, n=4, 2 significance is determined using Student’s *t*-test, *p < 0.05, ** p<0.01. **(E)** qPCR showing *KIT* and *PDGFRA* expression at day 4 after infection. Mean ± SEM, n=3, significance is determined using Student’s *t*-test, **p < 0.01, *** p<0.001.

To decipher which of these receptors are essential for cell growth, shRNA against either *KIT*, or *PDGFRA* or both were transduced in M07e cells (Fig. S7A). Inhibition of these genes separately only modestly affected M07e cells proliferation but combined inhibition of both genes recapitulated growth inhibition to a similar extend as CRISPRi targeting of SEKIT (Fig. 3, C to E). These results suggest a functional collaboration of these two neighboring and co-regulated genes in cell growth.

M07e is a cytokine dependent cell line(*16*) that is generally cultured in the presence of Granulocyte-macrophage colony-stimulating factor (GM-CSF). The finding that KIT and PDGFRA expressions are required under GM-CSF stimulation raised the possibility that GM-CSF receptor activity could be dependent on *KIT* and *PDGFRA*. Canonical GM-CSF receptor is composed of a ligand-specific subunit α and a common β subunit encoded by *CSFR2A* and *CSFR2B* genes respectively(*16*). Off-target effects on GM-CSF receptor of shRNA targeting *KIT* and *PDGFRA* was ruled out as they did not impact *CSFR2A* and *CSFR2B* expression (Fig. S7D). Additionally, M07e cells can be cultured in the presence of SCF or PDGF alone. Under SCF or PDGF stimulation only shRNAs targeting *KIT* or *PDGFRA* respectively impaired cell growth, further demonstrating that shRNA silencing is on target (Fig. S7, B and C). It was previously reported that the β subunit can interact with KIT(*17*) and other growth factor receptors(*18, 19*). Additionally, we found that M07e cells express very low levels of *CSFR2A* compared to *CSFR2B* (40x less) (Fig. S7E). Furthermore, it was shown that KIT and PDGFRA are able to interact together in gastrointestinal stromal tumors(*20, 21*). These data suggest that functional interactions between CSFR2B, KIT and PDGFRA lead to survival and proliferative signals in leukemic cells.

To confirm that SEKIT activity is also important in AMKL patient cells, we transduced AMKL7 patient-derived cells to express the SEKIT-targeting CRISPRi system. Initially, we targeted the open region of the major H3K27ac peak in common with the M07e cell line using the sgSEKIT2 (Fig. S8A) and performed RNA-sequencing and quantitative PCR shortly after CRISPRi transduction (48h post infection). Similar to M07e cells, very few transcriptional changes were significant, including inhibition of *KIT* and *PDGFRA* (Supplemental Fig. S8, B and C). We next xenografted these transduced cells into immuno-compromised NSG mice to follow leukemia progression using non-invasive bioluminescent imaging (Fig. 4A). We observed a significant delay in leukemia progression in recipients transplanted with AMKL7 cells targeted for SEKIT when compared to animals transplanted with non-targeted control (Fig. 4, B and C). Analyses of hematopoietic tissue infiltration after 12 weeks showed a reduced infiltration of leukemic cells in bone marrow, spleen and liver (Fig. S9, A and B). These results demonstrate that SEKIT inhibition can impair AMKL progression *in vivo*.

**Fig. 4.**
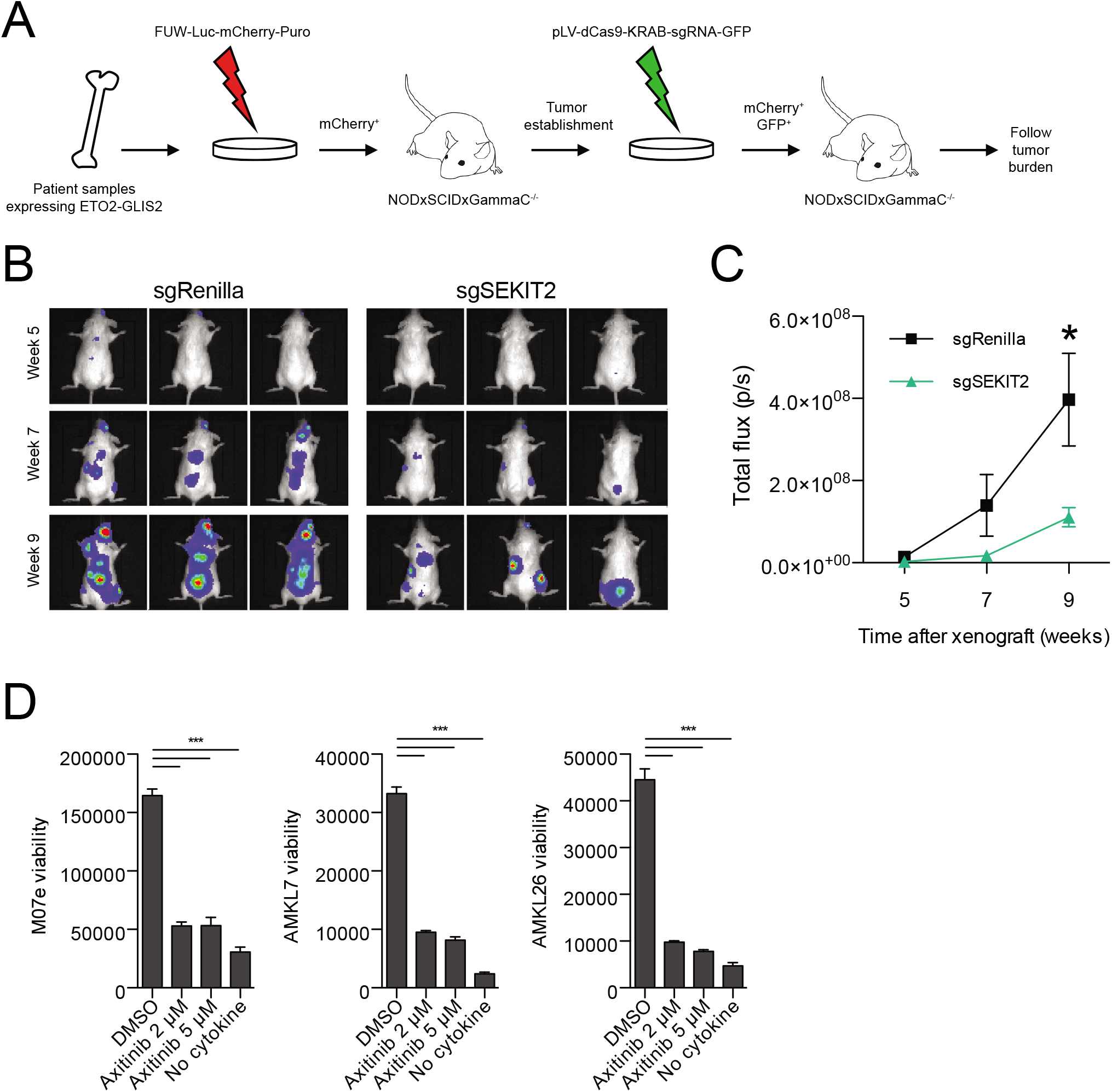
SEKIT targeting inhibits AMKL progression *in vivo*. **(A)** Schematic illustration describing CRISPRi targeting of SEKIT *in vivo* using patient derived xenograft model. **(B)** Representative bioluminescent imaging of NSG recipient mice transplanted with sgRNA2- or sgRenilla-transduced AMKL7^luc^mCherry^+^ patient cells at indicated post-transplant time **(C)** Quantification of bioluminescence *in vivo* as analyzed in panel b. Mean ± SEM, n=5, statistical significance is determined using Student’s *t*-test, * p<0.05. **(D)** Viability of M07e cell line, AMKL7 and AMKL26 patient cells treated with Axitinib- or vehicle (DMSO) for 96h. Negative control without cytokines is shown. Mean ± SEM, n=3, significance is determined using Student’s *t*-test, *** p<0.001.

Altogether these data indicate that ETO2-GLIS2^+^ leukemia growth is dependent on, at least, two tyrosine kinase receptors. These results introduced the exciting possibility that combined targeting of both receptors using dual specificity Tyrosine Kinase inhibitors (TKI) could be efficient for the treatment of this disease. To test this hypothesis, we treated M07e cell line as well as two AMKL patient-derived samples with increasing doses of five different TKI that can efficiently inhibit KIT and PDGFRA. Among those, Avapritinib and most particularly Axitinib potently blocked patient cell growth (Fig. 4E and Supplemental Fig. S10, A to C). Axitinib is a second-generation inhibitor of EGFR, KIT and PDGFRA(*22*) and is currently approved for clinical usage against refractory renal cell carcinoma in USA and Europe, raising the possibility of extension of its usage for these leukemia patients.

In this study, we sought to design a genome wide methodology to functionally screen SE in cancer cells in order to identify essential gene expression modules required for cancer cell maintenance. Using this approach, we pinpointed several enhancers whose activity is required for growth of a dismal prognosis subgroup of pediatric leukemia associated with fusion oncogenes. Specifically, we identified a *de novo* SE, induced by ETO2-GLIS2, controlling the expression of proximal neighboring genes *KIT* and *PDGFRA* which are required for leukemia cell growth. Therefore, it reveals that strong fusion oncogenes may lead, through SE-mediated regulation of functionally synergistic genes, to activation of signaling pathways essential for cancer development. Emergence and progression of diseases exhibiting strong and unique driver mutations may depend on the rewiring of a modest number of SE regions that would be manageable to annotate using our screening methodology in order to better understand the biology involved in the transformation potential of these oncogenes and narrow down the list of gene candidates with promising therapeutic values. This approach could be particularly amenable to studying other poorly-druggable oncogenes involving transcriptional and epigenetic remodeling factors, including fusion oncogenes (*e.g*. MLL-fusions(*23*) and NUP98-fusions(*24*)) in hematopoietic malignancies, but also in others cancers, including ependymoma(*25*) and H3.3 mutants glioma(*26*).

Together, we believe that systematic screening of essential SE can reveal coordinated regulation of genes modules involved in cancer cell transformation and cancer progression and uncover novel therapeutic approaches.

## Acknowledgments

This work was supported by the ATIP-Avenir program from French government (to CL and JC), The Fondation ARC (to CL and MCL) and the Fondation Gustave Roussy, PAIR-Pédiatrie/CONECT-AML (COllaborative Network for Children and Teenagers with Acute Myeloblastic Leukemia: INCa-ARC-LIGUE_11905), Association Laurette Fugain (ALF-2015/13 to TM), Fondation pour la Recherche Médicale (to CKL), Institut National du Cancer (PLBIO-2018-169 to TM)

## Author Contributions

CL and SB designed research, performed experiments and analyzed data. MCL helped optimizing CRISPRi screening. SL helped with shRNA experiments. MA helped with CRISPRi experiments. ES and JC performed 4C-seq experiments. AP provided primary patient samples. CKL and TM performed in vivo experiments. BP and AP helped with in vivo experiments. CT and ZA performed inducible fusion experiments and ChIP-seq. CL, SB, JC and TM co-wrote the paper. CL, JC and TM supervised the research.

## Materials & Correspondence

All correspondence and material request should be addressed to either Dr. Camille Lobry or Dr. Thomas Mercher.

## Tables

See Supplemental Table files.

## Methods

### Cells and culture

M07e, HEL-5J20 and AMKL patient cells were obtained as previously described(*1*). Kasumi-1 cells were obtained from ATCC. CD34+ cord blood cells were purchased from ABCellBio. M07e cells were cultured in MEM alpha medium supplemented with 20% FBS, 100 U/ml penicillin, 100 U/ml streptomycin (all from Gibco) and 5 ng/ml GM-CSF (PeproTech). HEL-5J20 cells were cultured in MEM alpha medium supplemented with 10% FBS, 100 U/ml penicillin and 100 U/ml streptomycin. Kasumi-1 cells were cultured in RPMI (Gibco) supplemented with 10% FBS, 100 U/ml penicillin and 100 U/ml streptomycin. 293T cells were cultured in DMEM medium (Gibco) supplemented with 10% FBS, 100 U/ml penicillin and 100 U/ml streptomycin. AMKL patient cells maintained in immunodeficient mice were cultured in StemSpan™ Serum-Free Expansion Medium (STEMCELL Technologies) supplemented with 100 U/ml penicillin, 100 U/ml streptomycin and 10 ng/ml each of human IL3, IL6, SCF, GMCSF, TPO, FLT3 and IGF1 (PeproTech).

The generation of doxycycline-inducible expression of ETO2-GLIS2 (and variants) HEL-5J20 cell lines was performed as follows. The cDNA encoding ETO2-GLIS2-GFP was cloned in a doxycycline (DOX) inducible LT3-GEPIR lentiviral vector (gift from J. Zuber, Austria) using BamHI/BglII and EcoRI. HEL-5J20 cells were transduced with lentiviral particles produced from LT3-GFP or LT3-ETO2-GLIS2-GFP vectors, maintained in RPMI medium supplemented with 10% FBS and induced with Doxycycline (500ng/ml). Twenty-four hours after induction, GFP+ cells were single-cell sorted in 96-well plates to obtain clones. Validation of the selected clones was performed by western blotting on protein extracted from cells induced with Doxycycline for 40 hours.

### Plasmids, lentiviral and retroviral gene transfer

pHR-SFFV-KRAB-dCas9-P2A-mCherry (Addgene #60954) was used to generate stable M07e expressing dCas9-KRAB fusion protein cell line. sgRNA library used in the screen was cloned into pU6-sgRNA-EF1Alpha-puro-T2A-BFP (Addgene #60955). The same vectors were used for screen validation or pLV-hU6-sgRNA-hUbC-dCas9-KRAB-T2a-GFP (Addgene #71237) was used. shRNAs targeting KIT, PDGFRA or Renilla were cloned into pLMP-Puro-IRES-GFP or pLMP-Puro-IRES-mCherry (gift from I. Aifantis, New York University).

Viral particles were produced in 293T cells by co-transfecting plasmids of interest along with a lentivirus packaging plasmid (psPAX2, Addgene #12260) and a VSV envelope expression plasmid (pMD2.G, Addgene #12259) or along with pCL-10A1 retrovirus packaging vector using calcium phosphate method. For transductions, cells were spinoculated twice with 293T supernatants harvested 24h and 48h after transfection and supplemented with 4 μg/mL Polybrene for 90 minutes at 2300 rpm and 30°C. Efficiency of knock down was checked on homogeneous cell populations with respect to BFP, GFP or mCherry expression after cell sorting or puromycin selection (1 μg/mL for 48h) by RT-qPCR.

### CRISPRi screen

SE in M07e cells were identified using H3K4me3 and H3K27Ac ChIP-seq by keeping H3K27Ac peaks overlapping low enrichment or no H3K4me3 peaks and not overlapping gene Transcription Start Sites. SE were ranked using ROSE algorithm(*2, 3*) based on H3K27Ac signal. 7394 sgRNAs were designed in chromatin accessible sites, as defined by ATAC-seq peaks (window centered on peak center and extended +/− 750bp) located in 448 SE using CLD(*4*). Non-targeting sgRNAs were included in the pool (1 sgRNA targeting Renilla, 1 sgRNA targeting Luciferase and 3 sgRNA targeting mouse Lin28) as negative controls and 9 sgRNA targeting a window of −100bp top +500 bp around *ERG* TSS were used as positive controls. Oligos were synthesized and ordered as a pool of 73-mers (GGAGAACCACCTTGTTGG(X)_20_GTTTTAGAGCTAGAAATAGCAAGTTAAAATAAGGC, where X is the protospacer sequence) from CustomArray Inc. Double stranded sgRNA was completed by PCR using the 73-mer oligo and the following reverse oligo: CCTAGTACTCGAGAAAAAAAGCACCGACTCGGTGCCACTTTTTCAAGTTGATACGGACTAGCCTTATTTTAACTTGCTATTTCTAGCTCTAAAAC. Library was then amplified using the following primers: Lib_amp_FOR: GGAGAACCACCTTGTTGG, Lib_amp_REV: CCTAGTACTCGAGAAAAAAAGCACC. Pooled sgRNA library was cloned into pU6-sgRNA-EF1Alpha-puro-T2A-BFP using BstXI and XhoI restriction sites. Polyclonal population of M07e dCas9-KRAB expressing cells was sorted by flow cytometry using mCherry expression, amplified, then infected with sgRNA library in triplicate. 50 million cells were infected with 15% transduction efficiency to achieve an effective multiplicity of infection of less than one sgRNA per cell and to ensure library coverage of 1000x. Two days after infection cells were treated with 1 μg/ml puromycin. Two days after puromycin selection, dead cells were removed by Ficoll gradient and 8 million BFP^+^ cells were sampled (day 0 of the screen). More than 16 million BFP^+^ cells were maintained in the presence of 0.5 μg/ml puromycin at each passage to preserve library representation for the next samplings. 8 million BFP^+^ cells sampled at day 0, 7, 14 and 21 of the screen were lysed in 50 mM Tris-HCl, 50 mM EDTA, 100 mM NaCl, 1% SDS supplemented with 0.1 mg/mL RNAse A (Thermo Scientific) and 0.2 mg/mL Proteinase K (Invitrogen) for 12 hours at 37°C. Genomic DNA was then isolated by phenol-chlorophorm-isoamyl alcohol extraction using PhaseLock tubes (5PRIME) followed by 100% ethanol precipitation in the presence of 0.1 M sodium acetate and 20 μg/mL glycogen. Genomic DNA was then digested with PstI (New England Biolabs) for 12 hours at 37°C, run on 1% agarose gel to isolate 700 bp to 1500 bp fraction that contain the sgRNA cassettes using E.Z.N.A. Gel Extraction Kit (Omega Bio-tek). Deep sequencing libraries were generated by PCR amplification of sgRNA cassettes using sgRNA_P5_seq: AATGATACGGCGACCACCGAGATCTACACTCTTTCCCTACACGACGCTCTTCCGATCTTTGGAGAACCACCTTGTTGG and sgRNA_P7_barcode_seq: CAAGCAGAAGACGGCATACGAGATXXXXXXGTGACTGGAGTTCAGACGTGTGCTCTTCC GATCTGCCTAATGGATCCTAGTACTCGAG where XXXXXX represents six nucleotide TruSeq indexing barcode for Illumina sequencing. We obtained at least 4 μg of 700-1500 bp DNA fraction from 8 million BFP^+^ cells that were used in total as template in multiple parallel 100 μL PCR reaction, each containing 1 μg template, 1x Phusion HF Buffer, 2U Phusion Hot Start II High-Fidelity DNA Polymerase (Thermo Scientific), 0.2 mM dNTP, 3% DMSO and 0.125 μM of each primer, which were run using the following cycling parameters: 98°C for 2 min; 29 cycles of 98°C for 30 s, 58°C for 15 s, 72°C for 15 s; 72°C for 10 min. PCR products (286 bp) were combined for each sample, purified on 1% agarose gel using E.Z.N.A. Gel Extraction Kit, further purified using 1.4 volume of AMPure XP beads and sequenced on the Illumina HiSeq 4000 Sequencer (50bp single end) at a coverage of more than 8 million reads per sample.

### Quantitative real-time PCR

For mRNA quantification, total RNA was isolated using E.Z.N.A.^®^ MicroElute Total RNA Kit (Omega Bio-tekkit) and transcribed into complementary DNA using High-Capacity cDNA Reverse Transcription Kit (Applied Biosystems). Real-time PCR reactions were carried out using 1x GoTaq™ qPCR Master Mix (Promega) and 0,25 μM of forward and reverse primers (Table S5) on a QuantStudio™ 7 Flex Real-Time PCR System (Applied Biosystems).

### Chromatin immunoprecipitation and sequencing

The ChIP protocol was described previously(*1*). Briefly, cells were fixed with 1% formaldehyde, lysed at a concentration of 20.10^6^ cells per mL, and finally sonicated (30-min cycle on Covaris apparatus; KBioscience). Sheared chromatin was immunoprecipitated overnight using the anti-H3K27Ac antibody (ActiveMotif, #39133). For GFP-ChIP, GFP-Trap agarose beads (Chromotek) were used. Enriched DNA from ChIP and input DNA fragments were end-repaired, extended with an ‘A’ base on the 3’end, ligated with indexed paired-end adaptors (NEXTflex, Bioo Scientific) using the Bravo Platform (Agilent), size-selected after 4 cycles of PCR with AMPure XP beads (Beckman Coulter) and amplified by PCR for 10 cycles more. Libraries were single-end sequenced (50bp) using Illumina HiSeq 4000 (Illumina, San Diego, CA).

Sequencing reads were aligned to the human hg19 version of the genome using Bowtie2(*5*) and peaks were called using MACS2(*6*) using default options. Normalized bigwig files for gene track representations were generated using Deeptools2(*7*) with the --normalizeUsing RPKM option.

### ATAC-seq

ATAC-seq was performed as previously described(*8*) with some modifications. Briefly, 50000 cells were spun at 500g for 5 min, washed with cold PBS, lysed in cold lysis buffer (10 mM Tris-HCl, pH 7.4, 10 mM NaCl, 3 mM MgCl2 and 0.1% IGEPAL CA-630) and immediately spun at 500g for 10 min. The pelleted nuclei were resuspended in 50 μl transposase reaction mix (Nextera Tn5 Transposase, Illumina) for 30 min at 37 °C. Transposed DNA was purified using a DNA Clean & Concentrator-5 kit (Zymo Research) in 10 μl nuclease-free H_2_O and amplified with NEBNext^®^ High-Fidelity 2X PCR Master Mix and 1.25 M of custom Nextera PCR primers as previously described(*8*), using the following PCR conditions: 72 °C for 5 min; 98 °C for 30 s then 12 cycles of 98 °C for 10 s, 63 °C for 30 s and 72 °C for 1 min. Libraries were purified with AMPure XP beads (Beckman Coulter) then subjected to high-throughput paired-end sequencing (50bp) using the Illumina HiSeq 4000 sequencer (Illumina, San Diego, CA).

Sequencing reads were aligned to the human hg19 version of the genome using Bowtie2(*5*) and accessibility peaks were called using MACS2(*6*) using default options. Bedgraph files normalized for the depth of sequencing of aligned reads were generated using bedtools.

### RNA sequencing

RNA from M07e and AMKL7 cells subjected to CRISPRi inactivation of KIT super enhancer, puromycin selected or GFP sorted, was extracted using the RNeasy Plus Mini Kit (Qiagen). Poly(A)-selected, first-stranded Illumina libraries were prepared with a modified TruSeq protocol using dUTP method(*9*). Two biological replicates per cell type were prepared. AMPure XP size selected libraries were amplified by PCR (maximum 16 cycles) and purified with AMPure XP beads and paired-end sequenced (50bp) on the Illumina HiSeq 4000 sequencer.

Sequencing reads were aligned to the hg19 version of the human genome using Tophat2. Differential expression analysis was done using CuffDiff(*10*).

Gene Set Enrichment Analysis (GSEA)(*11*) was performed on a pre-ranked list of genes after filtering out non expressed genes (genes with median FPKM expression below 1 in both compared conditions). Genes were ranked using their Log2 fold change of expression between compared conditions. GSEA was carried out using MSigDB genesets from C2 common pathways, C3 transcription factors, C4 cancer modules, C5 molecular functions, C6 and Hallmark collections.

### Cell viability assay

M07e and AMKL patient cells were plated in 96-well plates (2.10^5^ cells/well) in their respective media and treated with kinase inhibitors Axitinib, Amumavatinib, Telatinib, Avapritinib, Imatinib (all from Selleckchem and tested at 2 μM or 5 μM in 0.0005% DMSO). Cell viability was measured after 96h treatment as ATP luminescence signal readout using CellTiter-Glo Assay (Promega) according to manufacturer’s instructions.

### *In vivo* xenotransplantation assays

The use of human patient samples in this study was approved by the Internal Review Board of Gustave Roussy and samples were obtained with the informed consent of the patient in accordance with Good Clinical Practice rules and national ethics recommendations. Mice were maintained in pathogen-free conditions and all experiments were approved by the Gustave Roussy institute animal care and use committee (Comité d’Ethique #26, projects: 2012-017 and 2017122111548235).

AMKL patient-derived xenografts have been described previously (Thiollier et al., 2012). For bioluminescence follow-up of human AMKL cells, human AMKL7 or AMKL26 patient cells were transduced with a FUW-Luciferase-mCherry-Puromycin lentiviral vector (a kind gift from A. L. Kung, Dana Farber Cancer Institute, Boston), sorted on mCherry expression and then amplified in NSG recipients.

mCherry^+^ AMKL cells were further used for transduction with pLV-hU6-sgRNA-hUbC-dCas9-KRAB-T2a-GFP lentiviral vector for CRISPRi inactivation of KIT super enhancer (sgRNA2) or negative control (sgRenilla). 1.10^6^ AMKL7^luc^mCherry^+^GFP^+^ blasts expressing sgRNA2 or sgRenilla were intravenously injected into 6-10 weeks-old female NSG mice (The Jackson Laboratory). Recipients were monitored weekly by bioluminescence. To this aim, mice received 150 mg/kg D-Luciferin (Beetle luciferin, E1605; Promega, Charbonnières, France) intraperitoneally, were anesthetized with 3% isoflurane, and imaged 1 to 5 min with an IVIS50 system (Perkin Elmer, Courtaboeuf, France). Bioluminescence intensity is expressed as photons per second (p/s).

### Flow cytometry

Cells were analyzed on a BD LSR II or a BD Fortessa flow cytometers using FACS Diva software (BD Biosciences). FACS data were analyzed on Flow Jo version 10 (LLC).

To assess cell proliferation after CRISPRi inactivation of KIT super enhancer or shRNA knock down of KIT and PDGFRA expression, infected cells were mixed with uninfected cells in equal ratio. Percentages of infected cells were followed over time using BFP, GFP or mCherry expression, normalized to first day of follow up and compared to negative controls targeting Renilla or GFP.

For cell cycle analysis, 1×10^6^ M07e cells transduced with variants of pLV-hU6-sgRNA-hUbC-dCas9-KRAB-T2a-GFP lentiviral vector were fixed with 2% paraformaldehyde for 15 min, permeabilized with PBS-0.5% bovine serum albumin (BSA)-0.1% triton for 15 min, incubated in PBS-0.5% BSA containing mouse anti-human Ki-67 (Alexa Fluor 647, clone B56, #558615, BD Pharmingen) for 30 min at 4°C, washed twice with PBS, incubated in 500 uL of PBS-0.5% BSA containing 5 ug/mL RNAse A (Thermo Scientific) and 1 ug/mL DAPI (#564907, BD Pharmingen) for 15 min then analyzed gating on GFP^+^ cells. cKIT and PDGFRA expression was assessed using mouse anti-human CD117 (PE-Cyanine7, clone 104D2, #25-1178-42, eBioscience) and CD140a (PE, clone αR1, #556002, BD Pharmingen) respectively. Cells were stained in PBS-2% FBS-2mM EDTA for 30 min at 4°C, washed then analyzed.

12 weeks after xenotransplantation, mice were euthanized for assessment of chimerism in bone marrow, spleen and liver. Cells from diseased mice organs were collected, subjected to red blood cell lysis, washed, resuspended in PBS-2% FBS-2mM EDTA containing 1 ug/mL DAPI then analyzed.

### 4C-seq

The protocol was performed as previously described with minor modifications(*12*). Briefly, 10-15*10^6^ cultured M07e and HEL 5J20 cells were fixed with 2% formaldehyde (28908, Thermo Scientific) for 10 min at RT (tumbling). Quenching of the cross linking was performed with the addition of freshly prepared 1M glycine (500046, Sigma). Tubes were transferred directly on ice and centrifuged for 5 min 300g at 4°C. Cells were washed with 1xPBS and centrifuged for 5 min 300g at 4°C, and pellets were frozen in liquid nitrogen and stored at −80°C. Cells were then vigorously resuspended in 1ml of fresh ice-cold lysis buffer (50mM Tris pH 8, 150mM NaCl, 0.5% NP-40, 1 tablet of complete protease inhibitor (04693159001, Roche)) and transferred to 9 ml of pre-chilled lysis buffer and incubated for 20 min on ice. After 5 min of centrifugation at 2500 g at 4°C nuclei were resuspended in 10 ml lysis buffer, incubated for 10 min on ice, centrifuged at 2500 g at 4°C. Nuclei were then resuspended in 100 μL 0.5% SDS solution and incubated for 10 min at 62°C (under agitation-900rpm). 25 μL 20% Triton and 315 μL of sterile water were added to the nuclei and incubated for 15 min at 37°C (under agitation 900 rpm). After adding 50 μL DpnII buffer (R0543M, NEB), 2 μL BSA (NEB, 20 mg/mL) and 400 units of DpnII (R0543M, NEB), samples were incubated for 4 h at 37°C (under agitation-900 rpm). At this point 5 μl of the sample were taken as the “undigested control”. Another 400 Units of DpnII were added and incubated O/N at 37°C (under agitation-900 rpm). 5 μl of the sample were taken as the “digested control”. Efficiency of chromatin digestion was verified after DNA extraction from 5 μl “undigested” and “digested” controls and loading in a 1.5 % agarose gel. After verification of chromatin digestion (smear between 200 bp and 2 kb), DpnII was deactivated by 20 min incubation at 62°C (under agitation 600 rpm). Ligation of DNA ends between the cross-linked DNA fragments was performed by addition of 992 μL of ligation master mix (150 μL of 10x ligation buffer (B0202S, NEB), 759 μL sterile water, 75 μL of 20% triton, 8 μL of BSA (20 mg/mL B9000S, NEB,) and 100 Units of ligase (M0202M, NEB)). Samples were incubated overnight at 16°C followed by 30 min at RT. 100 μl of the ligated sample were tested as “ligated control”, on a 1.5 % agarose gel. Samples were reverse cross-linked overnight at 65°C (under agitation-900 rpm) in presence of 100 μl of 20mg/ml Proteinase K (P2308, Sigma) and 60 μL 20% SDS. Samples were transferred at 37°C and incubated with 30 μl RNase A (10mg/ml, R4642 Sigma) for 45 min. DNA was purified with phenol chloroform (PCI), ethanol precipitated, resuspended in 200 μl H_2_0 and incubated for 1 h at 37°C. Efficiency of extraction and purification was verified on a 1.5% agarose gel. The DpnII-ligated 3C template was diluted to a concentration of 100 ng/μL in 1X Cut smart buffer (B7204S, NEB), and digested overnight at 37°C with 1 Unit of NlaIII enzyme (R0125L, NEB)/μg of DNA (under agitation-600 rpm). NlaIII was inactivated at 65°C for 20 min and DNA fragmentation was tested on 1.5% agarose gel. Samples were transferred to 50ml falcon tubes and DNA was ligated at 16°C in 12.6 ml sterile water, 1.4ml 10x ligation buffer (B0202S, NEB), and 200U ligase (M0202M, NEB). After 30min incubation at RT, samples were phenol chloroform extracted, ethanol precipitated, resuspended in 200 μl of sterile water and purified using Qia quick PCR Purification Kit (Qiagen). DNA concentration of each digested sample was calculated using the Qubit^®^ brDNA HS assay kit (Invitrogen). Specific primers used to PCR-amplify the 4C DNA for *SEKIT VIEWPOINT*: PEAK1-1-DpnII-F-2, 5’-GGGTAAGCAAAGGTTAGGAA-3’ and PEAK1-1-NlaIII-R-2, 5’-ATTAGCCCACTCTCTCACAT-3’. PCR reactions were performed using the Expand Long Template PCR system (11759060001, Roche) with the following PCR conditions: 94°C for 2 min; 30 cycles of 94°C for 15s, 55°C for 1min and 68°C for 3min; followed by a final step of 68°C for 7min. The 4C library was sequenced on a NextSeq™ 500 Illumina sequencer (75bp, single-end).

Data was analyzed as previously described(*13*).

### Data Availability

M07e H3K27Ac and H3K4me3 Chip-seq are available at ArrayExpress database (E-MTAB-4367). M07e and AMKL7 ATAC-seq and RNA-seq data, AMKL7 and inducible HEL 5J20 ChIP-seq and M07e 4C-seq data will be available at Gene Expression Omnibus database under accession code GSE131462.

### Statistical analyses

Results are expressed as means ± standard error of the mean. Statistical significance was calculated by two-tailed unpaired *t*-test on two experimental conditions with P < 0.05 considered statistically significant. Statistical significance levels are denoted as follows: ***P < 0.001; ** P< 0.01; * P< 0.05. No statistical methods were used to predetermine sample size.

