## Supplementary material for "Screening of clustered regulatory elements reveals functional cooperating dependencies in Leukemia": Fig. S1 to S10

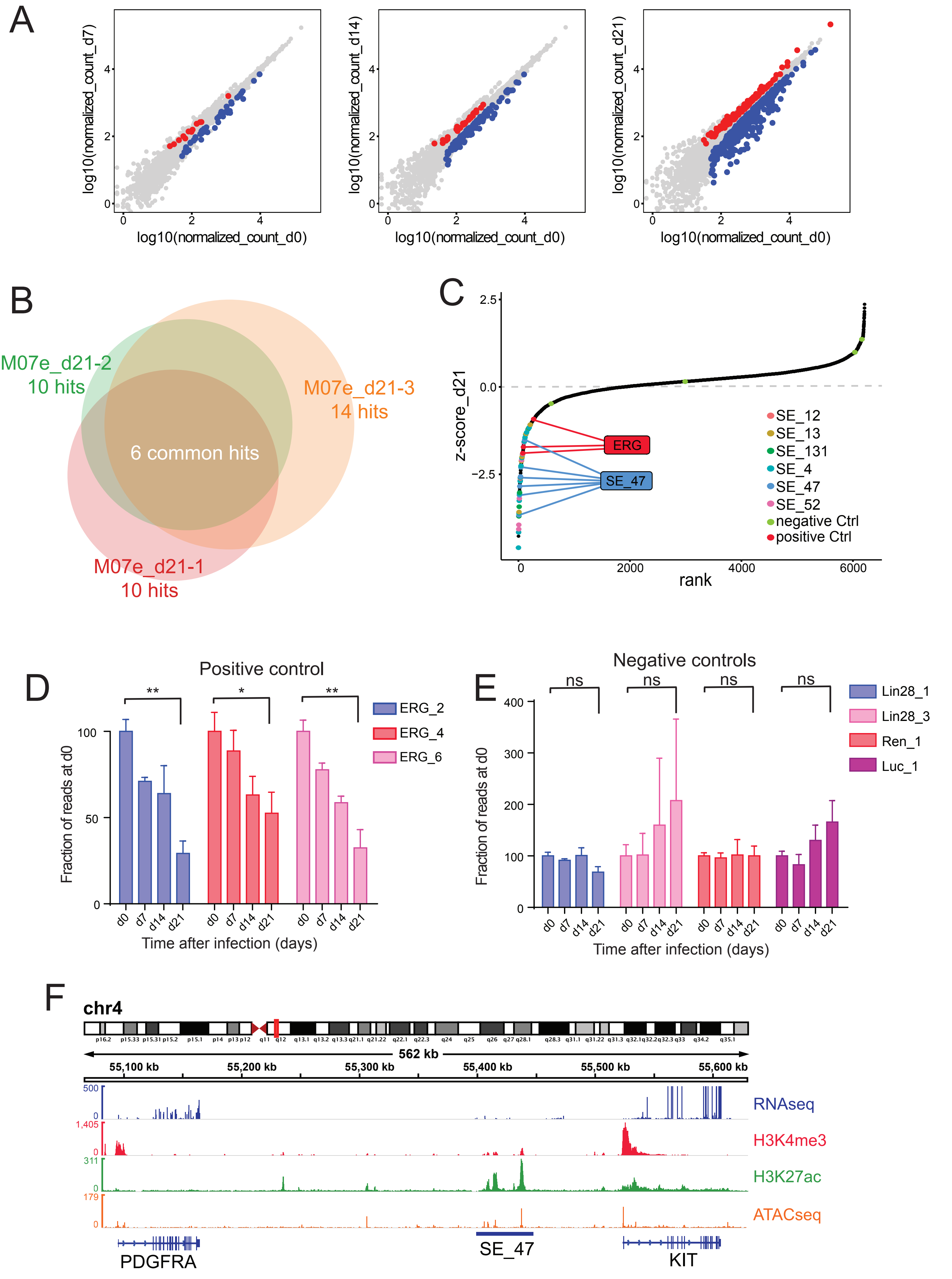
A
log10(normalized_count_d7)
log10(normalized_count_d14)
log10(normalized_count_d21)
log10(normalized_count_d0)
B
M07e_d21-2
10 hits
M07e_d21-3
14 hits
6 common hits
M07e_d21-1
10 hits
C
z-score_d21
rank
ERG
SE_47
SE_12
SE_13
SE_131
SE_4
SE_47
SE_52
negative Ctrl
positive Ctrl
D
Positive control
Fraction of reads at d0
ERG_2
ERG_4
ERG_6
Time after infection (days)
E
Negative controls
ns
Fraction of reads at d0
Lin28_1
Lin28_3
Ren_1
Luc_1
Time after infection (days)
F
chr4
562 kb
55,100 kb
55,200 kb
55,300 kb
55,400 kb
55,500 kb
55,600 kb
RNAseq
H3K4me3
H3K27ac
ATACseq
PDGFRA
SE_47
KIT

**Fig. S1. CRISPRi screening of Super Enhancer in ETO2-GLIS2$^+$ AMKL cell line M07e**

**(A)** Scatter plots showing significantly depleted (blue) or enriched (red) sgRNA at day 7, 14 and 21 compared to day 0. Only sgRNA with normalized read coverage over 40x at day 0 and with p-value < 0,05 are highlighted. Dots represent average of normalized read counts across the 3 replicates.

**(B)** Venn Diagram showing overlap of significant (FDR < 0,25) hits between the 3 replicates of the screen as determined by maximum likelihood of enrichment.

**(C)** Dot plot depicting changes in representation of the 6471 sgRNAs covered over 40x during 21 days of culture. sgRNA z-scores were calculated by subtracting to the log2 fold change of average normalized counts at d21 compared to d0 (Log2FC(d21/d0)) the average of all sgRNA Log2FC(d21/d0) and dividing by the standard deviation of all Log2FC(d21/d0) and are plotted in ascending order. Positive control sgRNAs targeting ERG gene are marked in red; negative control sgRNAs are marked in green. sgRNAs against common hits and significantly depleted are colored and sgRNA against SE_47 and ERG are highlighted.

**(D)** Bargraph representing variations of normalized counts of sgRNA targeting ERG, positive control of the screen. Data are represented as ratio of counts at day 0. Data are represented as mean± SEM, n=3, statistical significance is determined using Student's *t*-test, * p<0.05, ** p<0.01.

**(E)** Bargraph representing variations of normalized counts of sgRNA targeting Luciferas, Renilla and Lin28, hegative controls of the screen. Data are represented as ratio of counts at day 0. Data are represented as mean± SEM, n=3, statistical significance is determined using Student's *t*-test, ns=not significant.

**(F)** Gene track of SE_47 locus on chromosome 4, including proximal genes *KIT* and *PDGFRA,* showing normalized read density histograms of H3K4me3 ChIP-seq, H3K27ac ChIP-seq and ATAC-seq (bottom panel) in M07e cells. Read densities are shown as unique reads per million.

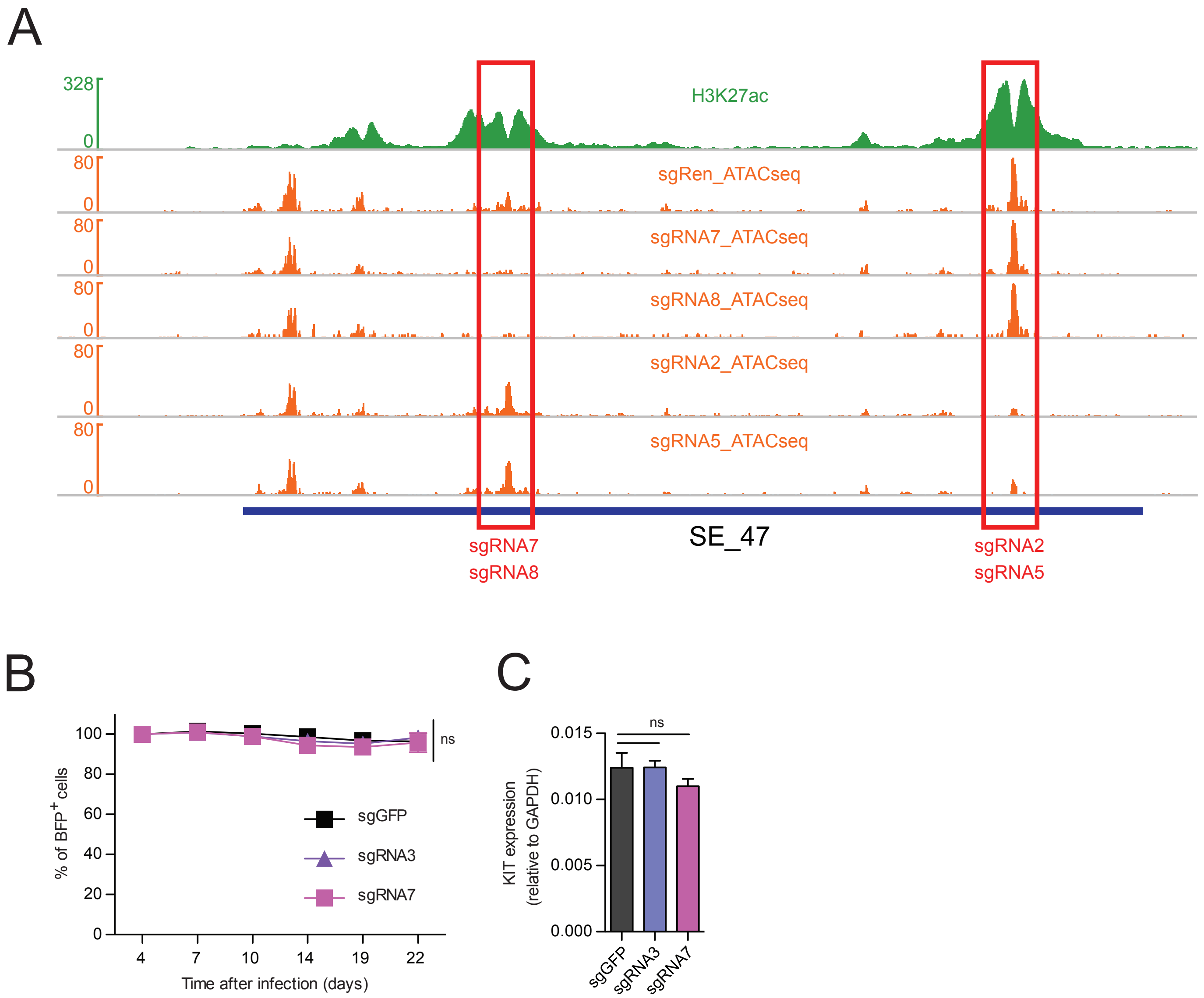

A
328
0
H3K27ac
80
0
sgRen_ATACseq
sgRNA7_ATACseq
sgRNA8_ATACseq
sgRNA2_ATACseq
sgRNA5_ATACseq
SE_47
sgRNA7
sgRNA8
sgRNA2
sgRNA5
B
% of BFP+ cells
100
80
60
40
20
0
4
7
10
14
19
22
Time after infection (days)
ns
sgGFP
sgRNA3
sgRNA7
C
KIT expression
(relative to GAPDH)
0.015
0.010
0.005
0.000
ns
sgGFP
sgRNA3
sgRNA7

**Fig. S2. CRISPRi targeting of SE is on target**

**(A)** Gene track showing normalized read density histograms of ATAC-seq following CRISPRi targeting of SE_47 with indicated sgRNAs compared to control sgRenilla (orange tracks) and H3K27ac ChIP-seq (green) in M07e cells. Targeted regions and location of sgRNAs are highlighted.

**(B)** Percentage of $BFP^{+}$ HEL 5J20 cells following CRISPRi targeting of SE_47 locus with indicated sgRNAs compared to control sgGFP and normalized to day 4 after infection. Mean ± SEM, n=3, statistical significance is determined using Student's *t*-test, ns=not significant.

**(C)** Quantitative PCR showing KIT expression in HEL 5J20 cells following CRISPRi targeting of SE_47 locus with indicated sgRNAs compared to control sgGFP. Mean ± SEM, n=3, statistical significance is determined using Student's *t*-test, ns=not significant.

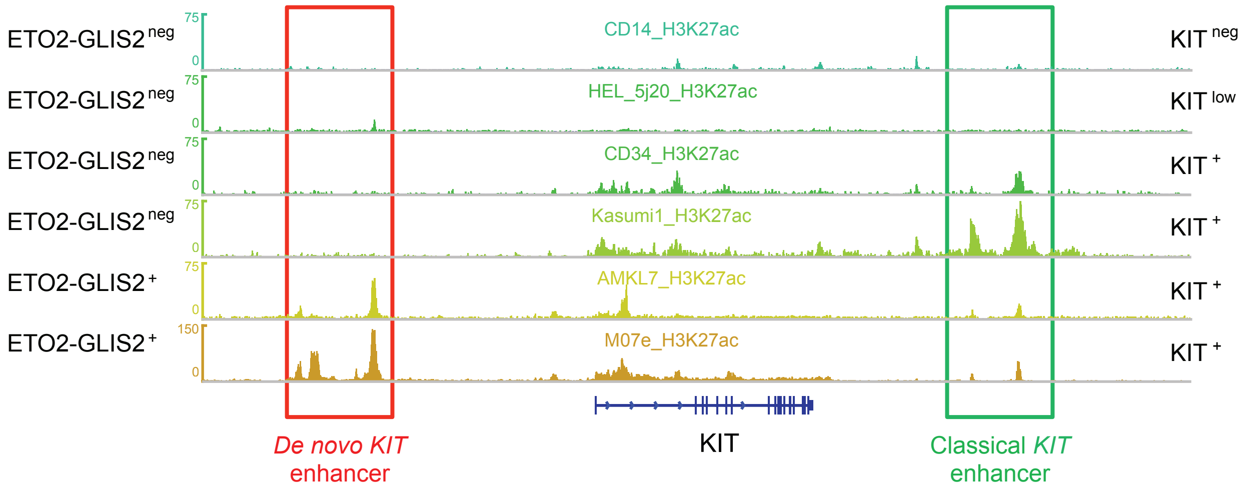
ETO2-GLIS2 neg
ETO2-GLIS2 neg
ETO2-GLIS2 neg
ETO2-GLIS2 neg
ETO2-GLIS2 +
ETO2-GLIS2 +
CD14_H3K27ac
HEL_5j20_H3K27ac
CD34_H3K27ac
Kasumi1_H3K27ac
AMKL7_H3K27ac
M07e_H3K27ac
KIT neg
KIT low
KIT +
KIT +
KIT +
KIT +
KIT
De novo KIT enhancer
Classical KIT enhancer

**Fig. S3. SEKIT is a *de novo* enhancer in hematopoietic cells**

Gene track showing normalized read density histograms of H3K27ac ChIP-seq around *KIT* gene in cells primary hematopoietic cells (CD34+ HSPC and CD14+ monocytes), AML cell lines (Kasumi-1, HEL 5J20) and AMKL cell line M07e and AMKL7 patient cells differentially expressing *KIT* and *ETO2-GLIS2*. Classical and de novo KIT enhancers are highlighted. Read densities are shown as unique reads per million.

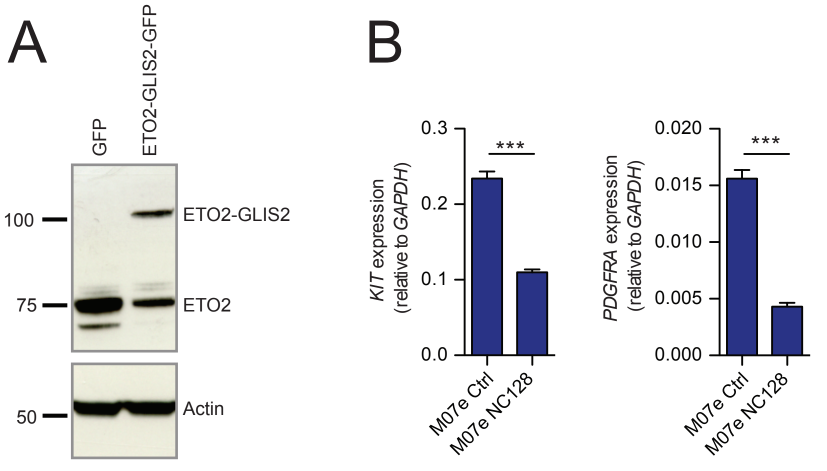
A
GFP
ETO2-GLIS2-GFP
100
75
50
ETO2-GLIS2
ETO2
Actin
B
KIT expression
(relative to GAPDH)
0.3
0.2
0.1
0.0
***
M07e Ctrl
M07e NC128
PDGFRA expression
(relative to GAPDH)
0.020
0.015
0.010
0.005
0.000
***
M07e Ctrl
M07e NC128

**Fig. S4. SEKIT is induced by ETO2-GLIS2**

**(A)** Western blot showing ETO2 and ETO2-GLIS2 expression in an individual clone of HEL 5J20 cells upon doxycycline induction of ETO2-GLIS2 expression compared to empty vector (GFP).

**(B)** qPCR showing *KIT* and *PDGFRA* expression in M07e cells expressing NC128 or control peptides. Mean ± SEM, n=3, statistical significance is determined using Student's *t*-test, *** $p<0.001$

A

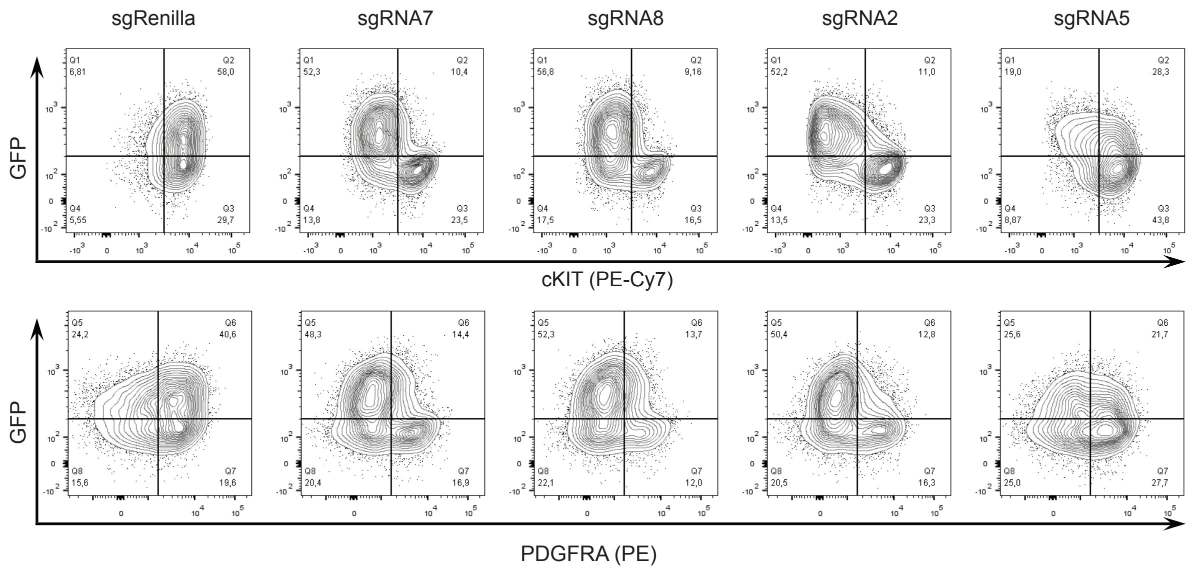
sgRenilla
sgRNA7
sgRNA8
sgRNA2
sgRNA5
GFP
Q1 6,81
Q2 58,0
Q4 5,55
Q3 29,7
Q1 52,3
Q2 10,4
Q4 13,8
Q3 23,5
Q1 56,8
Q2 9,16
Q4 17,5
Q3 16,5
Q1 52,2
Q2 11,0
Q4 13,5
Q3 23,3
Q1 19,0
Q2 28,3
Q4 8,87
Q3 43,8
cKIT (PE-Cy7)
GFP
Q5 24,2
Q6 40,6
Q8 15,6
Q7 19,6
Q5 48,3
Q6 14,4
Q8 20,4
Q7 16,9
Q5 52,3
Q6 13,7
Q8 22,1
Q7 12,0
Q5 50,4
Q6 12,8
Q8 20,5
Q7 16,3
Q5 25,6
Q6 21,7
Q8 25,0
Q7 27,7
PDGFRA (PE)

B

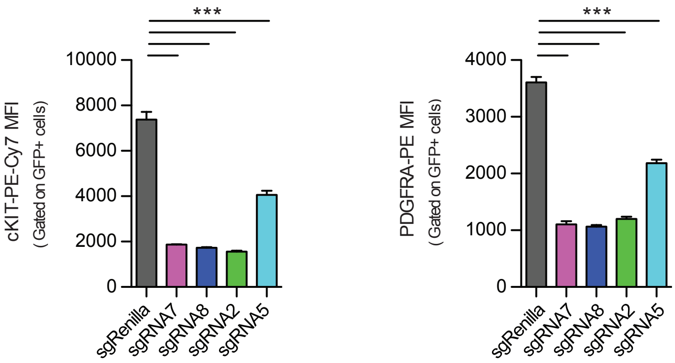
***
cKIT-PE-Cy7 MFI
( Gated on GFP+ cells)
10000
8000
6000
4000
2000
0
sgRenilla
sgRNA7
sgRNA8
sgRNA2
sgRNA5
***
PDGFRA-PE MFI
( Gated on GFP+ cells)
4000
3000
2000
1000
0
sgRenilla
sgRNA7
sgRNA8
sgRNA2
sgRNA5

**Fig. S5. SEKIT inactivation inhibits KIT and PDGFRA protein expression**

**(A)** Representative flow cytometry analyses showing KIT (higher panel) and PDGFRA (lower panel) protein expression at cell surface in M07e cells following CRISPRi targeting of SEKIT locus with indicated sgRNAs compared to control sgRenilla.

**(B)** Quantification of KIT and PDGFRA protein expression shown as mean fluorescent intensities of PE-Cy7 and PE respectively gated on transduced $GFP^+$ cells. Mean ± SEM, n=3, significance is determined using Student's *t*-test, *** p<0.001.

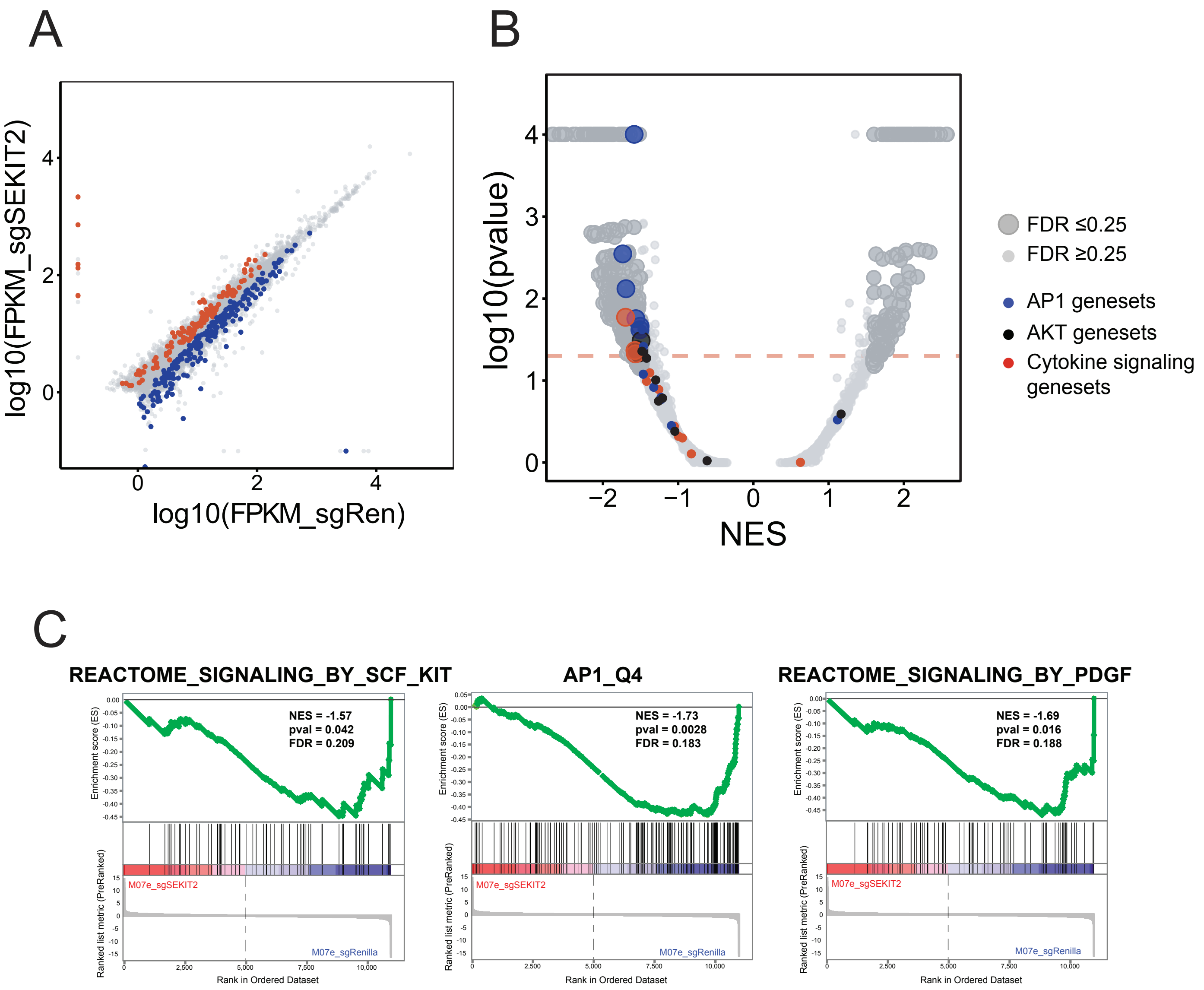
A
log10(FPKM_sgSEKIT2)
log10(FPKM_sgRen)
B
log10(pvalue)
NES
FDR ≤0.25
FDR ≥0.25
AP1 genesets
AKT genesets
Cytokine signaling genesets
C
REACTOME_SIGNALING_BY_SCF_KIT
NES = -1.57
pval = 0.042
FDR = 0.209
AP1_Q4
NES = -1.73
pval = 0.0028
FDR = 0.183
REACTOME_SIGNALING_BY_PDGF
NES = -1.69
pval = 0.016
FDR = 0.188
Enrichment score (ES)
Ranked list metric (PreRanked)
M07e_sgSEKIT2
M07e_sgRenilla
Rank in Ordered Dataset

**Fig. S6. SEKIT inhibition induces downregulation of pro-proliferative transcriptional programs**

**(A)** Scatter plot showing significantly depleted (blue) or enriched (red) genes 4 days transduction with sgSEKIT2. Dots represent average of log10FPKM across the 2 replicates.

**(B)** Volcano plot representing Gene Set Enrichment Analysis (GSEA) of expression changes in M07e cells expressing sgSEKIT2 versus sgRenilla. Normalized Enrichment Scores (NES) are plotted against P values. Red dotted line represent threshold of pvalue < 0,05, genesets with FDR > 0,25 are represented as small dots and genesets with FDR < 0,25 are represented as large dots.

**(C)** GSEA profiles of representative genesets negatively correlated in sgSEKIT2.

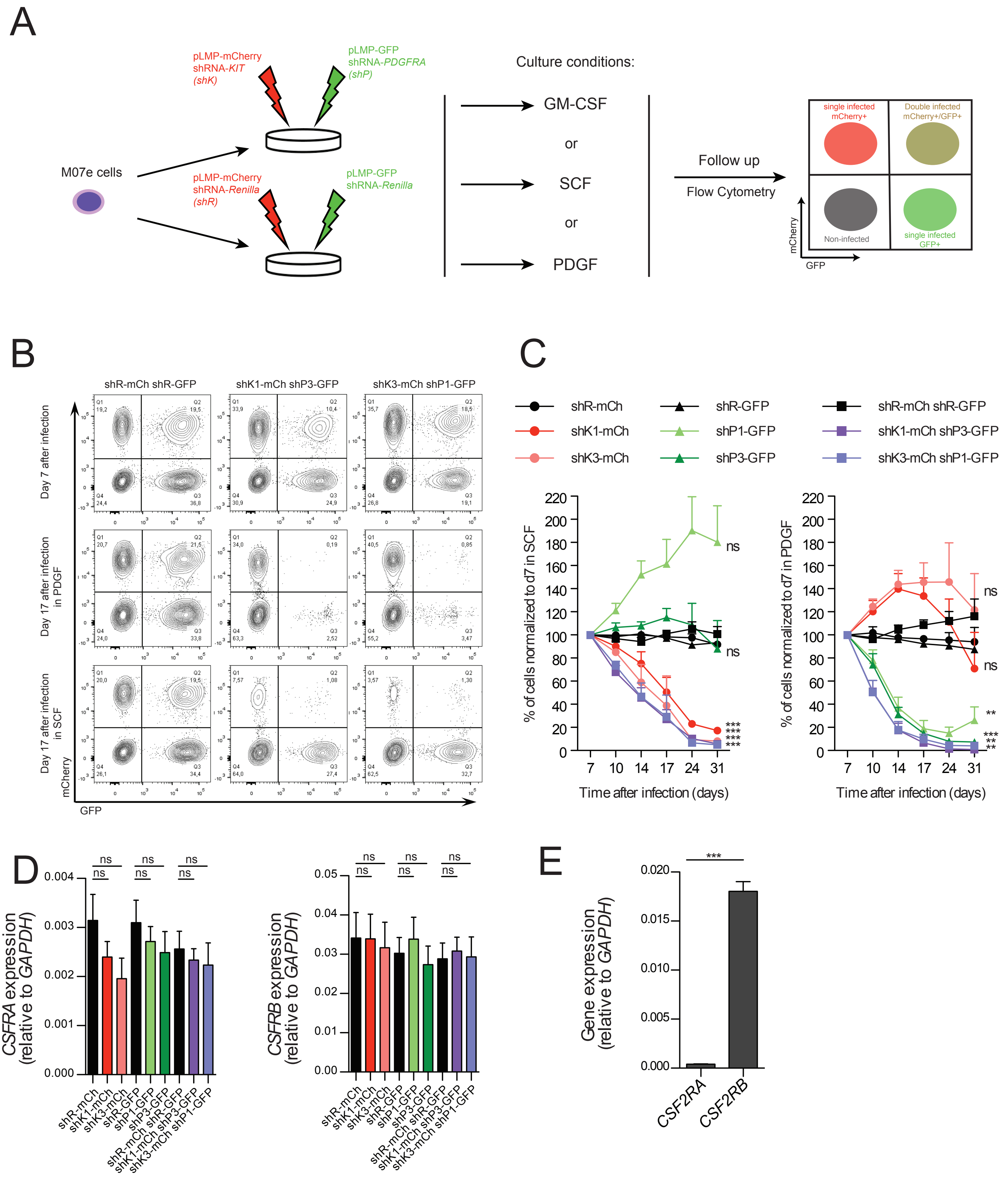
A
M07e cells
pLMP-mCherry
shRNA-KIT
(shK)
pLMP-GFP
shRNA-PDGFRA
(shP)
pLMP-mCherry
shRNA-Renilla
(shR)
pLMP-GFP
shRNA-Renilla
Culture conditions:
GM-CSF
or
SCF
or
PDGF
Follow up
Flow Cytometry
single infected mCherry+
Double infected mCherry+/GFP+
Non-infected
single infected GFP+
mCherry
GFP
B
shR-mCh shR-GFP
shK1-mCh shP3-GFP
shK3-mCh shP1-GFP
Day 7 after infection
Day 17 after infection in PDGF
Day 17 after infection in SCF
mCherry
GFP
C
shR-mCh
shK1-mCh
shK3-mCh
shR-GFP
shP1-GFP
shP3-GFP
shR-mCh shR-GFP
shK1-mCh shP3-GFP
shK3-mCh shP1-GFP
% of cells normalized to d7 in SCF
% of cells normalized to d7 in PDGF
Time after infection (days)
D
CSFRA expression (relative to GAPDH)
CSFRB expression (relative to GAPDH)
E
Gene expression (relative to GAPDH)
CSF2RA
CSF2RB

**Fig. S7. shRNA targeting of KIT and PDGFRA is on target**

**(A-C)** Two independent shRNAs targeting either KIT (shK) or PDGFRA (shP) were expressed in M07e cells separately or in combination. shK are expressed with mCherry (mCh) in cells while shP are expressed with GFP. Corresponding shRNAs targeting Renilla (shR) were used as control. 7 days after transduction, cells were washed and maintained in the presence of PDGF-AA or SCF as indicated.

**(A)** Schematic depicting experimental design

**(B)** Representative flow cytometry analyses of shRNA expressing cells at day 7 and day 17 after infection.

**(C)** Percentage of M07e cells expressing indicated shRNAs normalized to day 7 after infection. Mean ± SEM, n=3, significance is determined using Student's *t*-test, statistical significance is determined using Student's *t*-test, * $p<0.05$, ** $p<0.01$, *** $p<0.001$, ns=not significant.

**(D)** Quantitative PCR showing *CSF2RA* and *CSF2RB* expression at day 4 after infection. Mean ± SEM, n=3, significance is determined using Student's *t*-test, statistical significance is determined using Student's *t*-test, ns=not significant.

**(E)** Quantitative PCR comparing *CSF2RA* and *CSF2RB* expression in M07e cells. Mean ± SEM, n=3, significance is determined using Student's *t*-test, ***$p < 0.001$

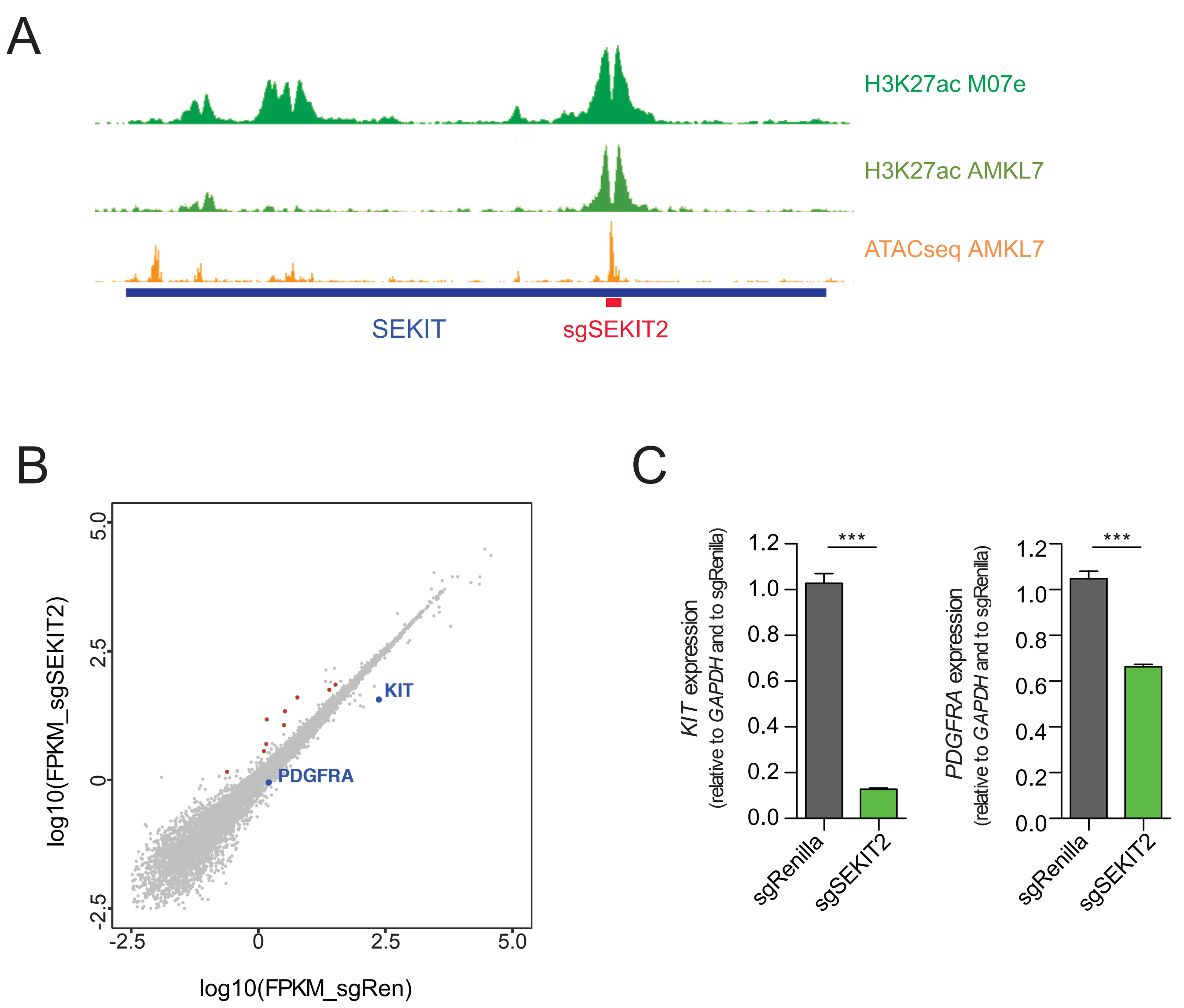

A
H3K27ac M07e
H3K27ac AMKL7
ATACseq AMKL7
SEKIT
sgSEKIT2
B
log10(FPKM_sgSEKIT2)
5.0
2.5
0
-2.5
KIT
PDGFRA
-2.5
0
2.5
5.0
log10(FPKM_sgRen)
C
KIT expression
(relative to GAPDH and to sgRenilla)
1.2
1.0
0.8
0.6
0.4
0.2
0.0
***
sgRenilla
sgSEKIT2
PDGFRA expression
(relative to GAPDH and to sgRenilla)
1.2
1.0
0.8
0.6
0.4
0.2
0.0
***
sgRenilla
sgSEKIT2

**Fig. S8. Targeting of SE_KIT in AMKL patient cells**

**(A)** Gene track showing normalized read density histograms of H3K27ac ChIP-seq and ATAC-seq (bottom panel) in AMKL7 patient cells (kaki and orange respectively) and H3K27ac ChIP-seq in M07e cells (green) at SEKIT locus. Read densities are shown as unique reads per million. Location of sgSEKIT2 is shown in red.

**(B)** Scatter plot showing genes whose expression is significantly altered following CRISPRi targeting of SEKIT with sgRNA2 compared to control sgRenilla in AMKL7 patient cells. Dots represent average of $\log_{10}$FPKM across the 2 replicates. Green dots $p<0.05$, red dots $p<0.05$ and >2-fold change. Statistical significance is determined using Student's *t*-test. *KIT* and *PDGFRA* genes are highlighted.

**(C)** qPCR showing KIT and PDGFRA expression following CRISPRi targeting of SE_KIT with sgRNA2 compared to control sgRenilla in AMKL7 patient cells. Mean ± SEM, n=6, Student's *t*-test, ***$p < 0.001$.

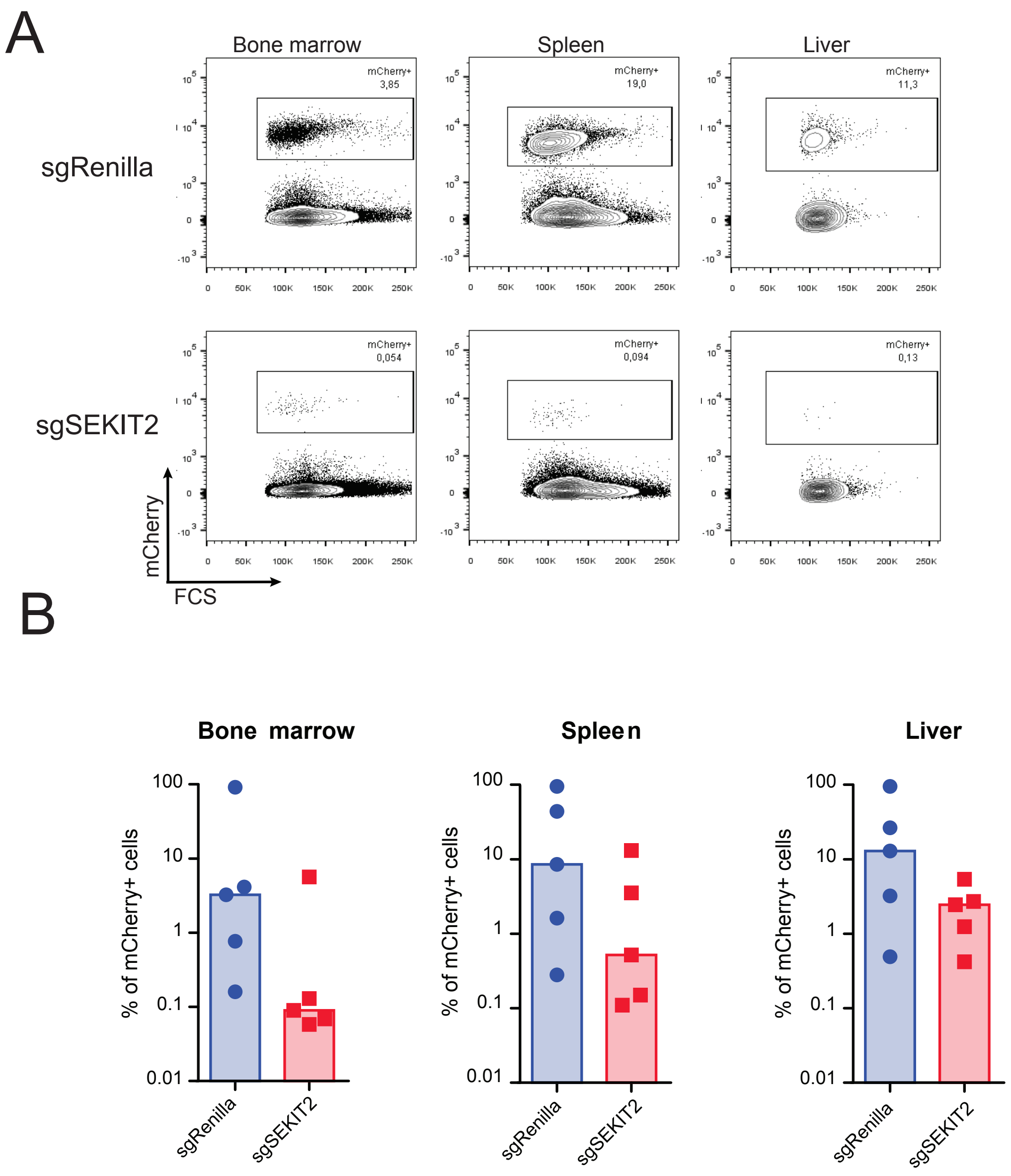

A
Bone marrow
Spleen
Liver
sgRenilla
sgSEKIT2
mCherry+
3,85
19,0
11,3
0,054
0,094
0,13
mCherry
FCS
B
Bone marrow
Spleen
Liver
% of mCherry+ cells
100
10
1
0.1
0.01
sgRenilla
sgSEKIT2

**Fig. S9. Inhibition of SEKIT impairs AMKL progression in vivo**

**(A)** Representative flow cytometry analyses of sgRNA2- or sgRenilla-transduced AMKL7$^{luc}$mCherry$^{+}$ cells in the bone marrow, spleen and liver of xenografted mice.

**(B)** Quantification of mCherry$^{+}$ cells as analyzed in panel **a**. Mean ± SEM, n=5.

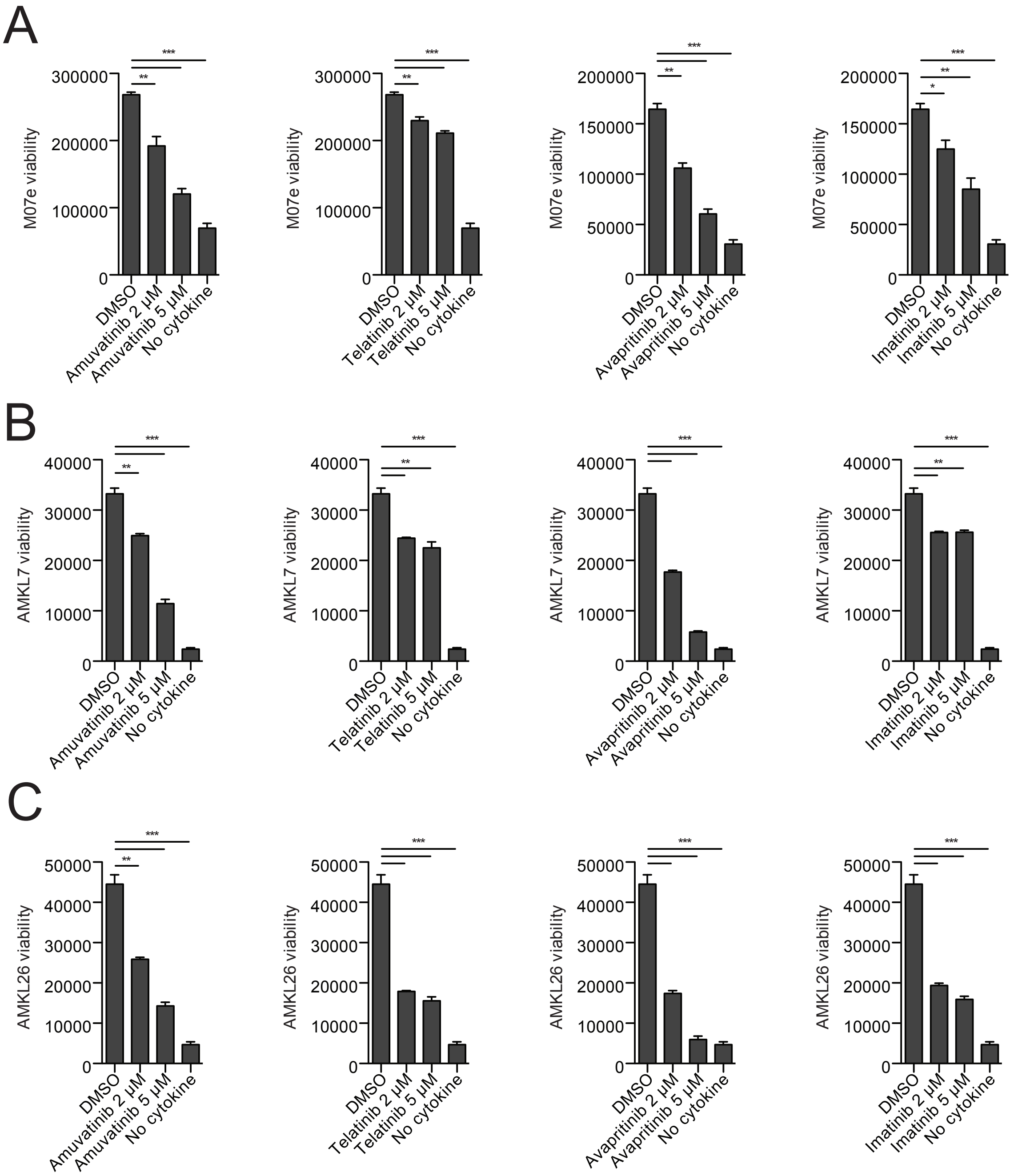

A
M07e viability
DMSO
Amuvatinib 2 μM
Amuvatinib 5 μM
No cytokine
Telatinib 2 μM
Telatinib 5 μM
Avapritinib 2 μM
Avapritinib 5 μM
Imatinib 2 μM
Imatinib 5 μM
B
AMKL7 viability
C
AMKL26 viability

**Fig. S10. Tyrosine kinase inhibitors affect AMKL cell growth**

**(A-C)** Viability of M07e cells **(A)**, AMKL7 **(B)** and AMKL26 **(C)** patient cells after 96h treatment with indicated kinase inhibitors or vehicle (DMSO). Negative control of culture without cytokines (No cytokine) is shown. Mean ± SEM, n=3, ***p < 0.001, *p < 0.05, **p < 0.01, and ***p < 0.001.
