## Supplementary material for "Screening of clustered regulatory elements reveals functional cooperating dependencies in Leukemia": Table S1

**Table S1: sgRNA composition and normalized counts of CRISPRi screen replicates**

| sgRNA | Gene | Peak | Protospacer_sequence | M07e-J0-1 | M07e-J0-2 | M07e-J0-3 | M07e-J7-1 | M07e-J7-2 | M07e-J7-3 | M07e-J14-1 | M07e-J14-2 | M07e-J14-3 | M07e-J21-1 | M07e-J21-2 | M07e-J21-3 |
|---|---|---|---|---|---|---|---|---|---|---|---|---|---|---|---|
| ERG_1 | ERG | NA | CAATAAACTTGATCGCATTA | 808.4975238994709 | 931.4337956787913 | 853.2232093162461 | 843.2479012390274 | 749.8403204996906 | 1051.7004733802482 | 856.6238477606594 | 924.5557137639489 | 1200.5560956196296 | 765.4572920682373 | 1044.6686185811568 | 875.1534835968739 |
| ERG_2 | ERG | NA | TGGCCAGCACTATTAAGGTA | 250.35245042999705 | 203.8760668653148 | 205.40558742798515 | 165.26617817493286 | 155.90739337122278 | 69.3048087894727 | 177.56650110616684 | 174.89039575289422 | 147.40555611897818 | 68.5266528137279 | 35.21354894093787 | 89.37243798005211 |
| ERG_3 | ERG | NA | GGATGAGATTGTGCTAAGTC | 739.1358060314199 | 815.5042674612592 | 883.8365420579169 | 765.1693918650435 | 851.6573937217136 | 521.518686140782 | 813.5774232500735 | 836.6378391422237 | 613.5978269771923 | 789.5145212475247 | 805.2164857827793 | 626.7677468730927 |
| ERG_4 | ERG | NA | GAAGAGGTTTCTATTGGTCC | 346.80858934025565 | 483.7059625628058 | 501.6636462183484 | 297.9996441107057 | 399.8441312989863 | 357.7860753756528 | 291.63952605921946 | 191.90675858290555 | 483.06399113689235 | 301.07986821350664 | 126.76877618737633 | 272.76003799106815 |
| ERG_5 | ERG | NA | CATTATGGCCAGCACTATTA | 1337.3806226433608 | 1496.0905494969427 | 1441.7892194464343 | 1444.4524234187045 | 1453.0144824392873 | 1386.9624858993225 | 1296.7735383814002 | 1400.06851951371 | 1791.0663053733674 | 1390.9452507297112 | 1358.069204155504 | 1283.713200077112 |
| ERG_6 | ERG | NA | GATGAGATTGTGCTAAGTCT | 734.8006986646667 | 885.461741385632 | 910.4997673490496 | 627.2306919710051 | 721.2042686559965 | 468.67376943880913 | 558.527358024852 | 458.4964429197497 | 620.7017091997936 | 210.68300705497197 | 450.73342644400475 | 161.33466076918498 |
| ERG_7 | ERG | NA | TTTCCTTACCTTAATAGTGC | 17.340429467012783 | 14.990887269508443 | 28.63827901640178 | 20.820935833062407 | 6.363567076376441 | 4.331550549342044 | 22.599372868057596 | 3.78141396222474 | 0.0 | 8.748083337922711 | 1.173784964697929 | 0.0 |
| ERG_8 | ERG | NA | AGCACTATTAAGGTAAGGAA | 304.541292514412 | 454.72358050842274 | 424.63655093285394 | 338.3402072872641 | 329.8448934588455 | 268.5561340592067 | 247.51694093586892 | 259.0268564123947 | 373.8418019643965 | 349.1943265720816 | 186.63180938697073 | 170.62010887100857 |
| ERG_9 | ERG | NA | ACCTCATCAAAACTACTTTC | 748.8897976066146 | 814.5048749766254 | 711.0193410968717 | 684.4882655119267 | 771.0522107542787 | 709.5079799822267 | 670.4480617523753 | 827.1843042366619 | 859.5697489347643 | 739.2130420544692 | 767.6553669124456 | 356.32907090748046 |
| Lin28_1 | Lin28 | NA | TTAGCTATCAACCATCACGC | 500.7049008599941 | 543.6695116408396 | 421.6739703449503 | 425.52787608821296 | 443.328506320892 | 609.8823173473597 | 356.2091628250983 | 513.3269453720085 | 475.07212363646585 | 408.97289604788676 | 362.69955409166005 | 233.29688355831783 |
| Lin28_2 | Lin28 | NA | TTCTATTGCATTTGTCCCGT | 0.0 | 3.9975699385355847 | 0.0 | 0.0 | 4.242378050917627 | 0.0 | 0.0 | 0.0 | 0.0 | 0.0 | 0.0 | 0.0 |
| Lin28_3 | Lin28 | NA | TATATTTACACAACTAGGCC | 24.926867358830876 | 52.9678016855965 | 54.31397744489992 | 54.65495656178882 | 71.05983235287025 | 2.598930329605226 | 24.75169409358689 | 184.3439306584561 | 8.879852778251697 | 38.63736807582531 | 230.0618530807941 | 5.803405063639747 |
| Luc_1 | Luc | NA | TTAATCAGAGACTTCAGGCG | 35.76463577571386 | 47.97083926242702 | 38.51354764274722 | 44.24448864525762 | 39.24199697098805 | 55.44384703157816 | 31.208657770174778 | 72.79221877282625 | 17.759705556503395 | 45.198430579267345 | 100.9455069640219 | 56.873369623669525 |
| Ren_1 | Ren | NA | TAGATAAGCATTATAATTCC | 721.7953765644071 | 867.4726766622219 | 877.9113808821096 | 648.0516278040674 | 803.9306406488903 | 1329.7860186480075 | 620.9446735652016 | 559.6492664092615 | 920.840733104701 | 978.32731995769 | 514.1178145376929 | 980.7754557551173 |
| SE_1_1_1 | SE_1 | SE_1_1 | CTTAGTTTTGAACAAGTACA | 61.77527997623304 | 68.95808143973883 | 63.20171920861082 | 74.17458390528482 | 45.60556404736449 | 70.17111889934111 | 67.79811860417279 | 34.97807915057884 | 126.98189472899928 | 12.393118062057175 | 15.259204541073078 | 55.71268861094157 |
| SE_1_1_4 | SE_1 | SE_1_1 | GTTGAGTTATCACACATCAT | 213.50403781259487 | 297.8189604209011 | 261.6946185981542 | 234.2355281219521 | 276.81516782237514 | 316.2031901019692 | 148.51016456152135 | 160.71009339455145 | 207.78855501108973 | 105.70600699989943 | 129.1163461167722 | 240.26096963468552 |
| SE_1_1_6 | SE_1 | SE_1_1 | AATGTAGTGTAGCTACAGTG | 285.0333093640226 | 339.7934447755247 | 340.6967676089177 | 337.0388987976977 | 358.48094530253945 | 368.1817966940737 | 276.5732774805144 | 274.1525122612937 | 276.1634214036278 | 384.9156668685993 | 194.84830413985622 | 330.7940886274656 |
| SE_1_1_8 | SE_1 | SE_1_1 | TTAGTTTATATCTTATGGCA | 6.5026610501297935 | 11.992709815606755 | 18.763010390056337 | 6.506542447832002 | 5.3029725636470335 | 0.0 | 2.152321225529295 | 3.78141396222474 | 47.063219724734 | 0.7290069448268927 | 5.868924823489645 | 17.410215190919242 |
| SE_1_1_9 | SE_1 | SE_1_1 | GATTGTCTGTGAAATTTCTG | 27.094421042207472 | 60.962941562667666 | 62.21419234597627 | 22.12224432262881 | 27.575457330964575 | 38.11764483420998 | 16.142409191469714 | 19.852423301679885 | 81.69464555991561 | 13.851131951710961 | 28.170839152750297 | 13.928172152735392 |
| SE_1_2_18 | SE_1 | SE_1_2 | TGGGGAGGCTGAAGTCAGGA | 470.3591492927217 | 576.649463633758 | 614.2417085586864 | 484.08675811870097 | 557.8727136956679 | 480.80211097696684 | 410.01719346333067 | 497.25593603255334 | 499.9357114155706 | 556.961305847746 | 534.0721589375577 | 713.8188228276889 |
| SE_1_2_6 | SE_1 | SE_1_2 | GCTCTGCCCACCTTTGGCAA | 0.0 | 0.9993924846338962 | 5.925161175807264 | 5.205233958265602 | 0.0 | 0.0 | 1.0761606127646475 | 0.0 | 0.0 | 10.93510417240339 | 0.0 | 1.1606810127279494 |
| SE_1_2_7 | SE_1 | SE_1_2 | CCAGTGCCGTTGCCAAAGGT | 548.3910818942793 | 415.74727360770083 | 385.13547642747216 | 579.0822778570482 | 375.45045750621 | 290.2138868059169 | 615.5638705013783 | 548.3050245225874 | 298.36305334925703 | 294.5188057100646 | 536.4197288669536 | 322.66932153836996 |
| SE_1_3_11 | SE_1 | SE_1_3 | ATGTTACCCCTGCTTCTGAA | 615.5852460789538 | 705.5710941515307 | 665.5931054156827 | 594.697979731845 | 617.2660064085147 | 646.2673419618329 | 606.9545855992611 | 652.2939084837676 | 681.0847080919052 | 683.0795073027984 | 382.65389849152484 | 748.6392532095274 |
| SE_1_3_12 | SE_1 | SE_1_3 | ACTGGCCTCATGTACTAAGG | 1566.0575362395919 | 1613.0194701991084 | 1466.4773910122979 | 1471.779901699599 | 1570.7404733522515 | 1448.4705036999794 | 1571.1944946363853 | 1783.8820366795212 | 1490.0392961906348 | 1522.1665007985519 | 1381.5449034494625 | 1669.0592963027914 |
| SE_1_3_13 | SE_1 | SE_1_3 | AATTTATCCACTGTACAGAG | 7.586437891818092 | 3.9975699385355847 | 4.93763431317272 | 7.807850937398403 | 0.0 | 0.8663101098684087 | 13.990087965940416 | 0.0 | 0.0 | 0.7290069448268927 | 0.0 | 0.0 |
| SE_1_3_14 | SE_1 | SE_1_3 | CTACTGACCACTCTGTACAG | 93.2048083851937 | 144.91191027191493 | 145.16644880727796 | 100.20075369661284 | 99.69588419656424 | 64.97325824013066 | 59.18883370205561 | 50.10373499947781 | 150.0695119524537 | 127.57621534470621 | 120.8998513638867 | 48.74860253457388 |
| SE_1_3_15 | SE_1 | SE_1_3 | ATGTGTTGTGAAATTCTGAT | 340.30592829012585 | 452.72479553915497 | 529.3143983721156 | 439.84226947344337 | 570.5998478484208 | 622.8769689953858 | 322.84818382939426 | 438.6440196180698 | 372.9538166865713 | 268.2745556962965 | 278.1870366334092 | 360.97179495839225 |
| SE_1_4_10 | SE_1 | SE_1_4 | CTTCTATGCAGTTTTCTAAG | 15.172875783636185 | 22.986027146579612 | 36.53849391747813 | 20.820935833062407 | 11.666539640023474 | 51.11229648223612 | 24.75169409358689 | 47.26767452780925 | 2.6639558334755096 | 45.92743752409424 | 21.128129364562724 | 54.55200759821362 |
| SE_1_4_11 | SE_1 | SE_1_4 | GTAAGACTCTGAGAGCCTTG | 166.90163361999802 | 289.82382054382987 | 271.5698872244996 | 213.41459228888968 | 260.90625013143404 | 112.62031428289313 | 175.41417988063753 | 128.56807471564116 | 414.6891247443543 | 122.47316673091797 | 124.42120625798047 | 392.3101823020469 |
| SE_1_4_15 | SE_1 | SE_1_4 | AGTTCCTCACAGAATCCAGA | 7.586437891818092 | 33.97934447755247 | 22.713117840594514 | 26.02616979132801 | 13.787728665482287 | 0.8663101098684087 | 11.837766740411123 | 18.9070698111237 | 5.327911666951019 | 8.748083337922711 | 43.43004369382337 | 10.446129114551544 |
| SE_1_4_20 | SE_1 | SE_1_4 | GGGGTACACAGCATGAACTG | 670.8578650050571 | 739.5504386290831 | 736.6950395253698 | 705.309201344989 | 714.8407015796201 | 1094.1496687638003 | 723.1799317778431 | 861.2170298966845 | 789.418911986576 | 774.20537540616 | 687.8379893129865 | 928.5448101823596 |
| SE_1_4_4 | SE_1 | SE_1_4 | TCTAGAAAAGTGATACACCA | 493.118462968176 | 580.6470335722937 | 654.7303099267027 | 514.0168533787282 | 554.6909301574798 | 759.7539663545945 | 482.11995451856205 | 616.3704758426326 | 659.7730614241011 | 564.2513752960149 | 590.4138372430583 | 796.2271747313733 |
| SE_1_5_16 | SE_1 | SE_1_5 | TTAGGAAATCCTGTTTGTGA | 184.24206308701082 | 233.8578414043317 | 270.5823603618651 | 193.89496494539367 | 279.9969513605634 | 128.21389626052448 | 126.9869523062284 | 343.16331707189516 | 214.89243723369108 | 112.99607644816837 | 369.74226387984766 | 225.1721164692222 |
| SE_1_5_19 | SE_1 | SE_1_5 | GATTTCCTAAAGAGGTCTTA | 4.335107366753196 | 3.9975699385355847 | 5.925161175807264 | 5.205233958265602 | 25.454268305505764 | 7.796790988815679 | 6.456963676587884 | 27.415251226129364 | 29.303514168230603 | 0.7290069448268927 | 93.90279717583432 | 9.285448101823595 |
| SE_1_5_4 | SE_1 | SE_1_5 | CTGAAGACTGAGCCTGAAGG | 240.59845885480235 | 322.80377253674845 | 300.2081662409014 | 320.1218884333345 | 335.1478660224925 | 256.42779252104896 | 256.1262258379861 | 180.56251669623134 | 266.39558334755094 | 178.6067014825887 | 166.67746498710594 | 170.62010887100857 |
| SE_1_5_5 | SE_1 | SE_1_5 | GCTTGGCCACCTGTCTGAGA | 1483.6904962712813 | 1767.9253053173622 | 1661.020182951303 | 1694.3036534154535 | 1564.376906275875 | 1532.502584357215 | 1532.452712576858 | 1345.2380170614513 | 2032.5983009418137 | 1818.1433203982704 | 1225.4315031446379 | 2327.1654305195384 |
| SE_1_5_6 | SE_1 | SE_1_5 | GCCAAAGAAATGGCATCACA | 1394.8207952528408 | 1206.2667289531128 | 1275.8847065238308 | 1405.4131687317124 | 1377.7122720354994 | 1322.8555377690602 | 1301.0781808324589 | 1219.5060028174787 | 1390.584945074216 | 1901.9791190533629 | 1164.3946849803456 | 2003.3354279684406 |
| SE_1_6_10 | SE_1 | SE_1_6 | TCTGAAATGGAAGTGTCTGT | 1543.2982225641376 | 1651.9957770998303 | 1758.7853423521228 | 1607.1159846145044 | 1531.4984763812633 | 1972.5881201703667 | 1597.0223493427368 | 1206.271053949692 | 1759.9868206494866 | 1891.0440148809596 | 1523.5728841779119 | 1615.6679697173056 |
| SE_1_6_15 | SE_1 | SE_1_6 | GAGAAGGAAAAGCTTTACCA | 306.7088461977886 | 486.7041400167074 | 510.5513879820593 | 377.3794619742561 | 464.54039657548014 | 518.0534457013084 | 358.3614840506276 | 416.9008893352776 | 392.48949279872505 | 430.8431049269353 | 423.7363722559524 | 478.2005772439152 |
| SE_1_6_22 | SE_1 | SE_1_6 | AACCATGTAGACCTAACCAT | 623.1716839707718 | 595.6379208418022 | 597.4537518938992 | 723.5275201989186 | 685.1440552231968 | 748.4919349263051 | 682.2858284927865 | 742.1024900866053 | 619.8137239219685 | 597.7856947580519 | 704.2709788187574 | 420.1665266075177 |
| SE_1_6_24 | SE_1 | SE_1_6 | TCCAGGAGACTCAGAGCAAG | 550.5586355776559 | 511.68895213255485 | 522.4017103336738 | 450.2527373899745 | 440.1467227827038 | 637.6042408631488 | 426.1596026548004 | 663.6381503704418 | 617.149768088493 | 446.8812571788852 | 712.4874735716429 | 269.2779949528843 |
| SE_1_6_4 | SE_1 | SE_1_6 | AGAGTTCTGCTGCAGAGTGA | 31.429528408960667 | 12.992102300240651 | 13.825376076883616 | 20.820935833062407 | 2.1211890254588135 | 6.93048087894727 | 39.817942672291956 | 23.633837263904624 | 65.71091055906257 | 5.103048613788249 | 12.911634611677218 | 66.15881772549312 |
| SE_1_7_12 | SE_1 | SE_1_7 | GATGTGATCATTAGGCGTGA | 3304.4355903076234 | 2964.198109424136 | 2980.356071431054 | 3245.463372978603 | 2860.42340083121 | 2571.208406089437 | 3563.167788863748 | 3010.9508674214494 | 3234.0423818392683 | 2609.1158555354486 | 3452.101581176609 | 3526.1489166675105 |
| SE_1_7_13 | SE_1 | SE_1_7 | ATTTGGCTCCACGTATCAAA | 89.95347786012881 | 103.93681840192521 | 118.50322351614528 | 89.79028578008163 | 149.54382629484635 | 186.25667362170788 | 100.08293698711222 | 149.36585150787724 | 193.580790565887 | 123.93118062057175 | 401.4344579266917 | 207.76190127830296 |
| SE_1_7_21 | SE_1 | SE_1_7 | ATTCCTTTAGGACTAACTCC | 2.167553683376598 | 9.993924846338961 | 12.837849214249072 | 9.109159426964803 | 2.1211890254588135 | 0.8663101098684087 | 12.913927353175769 | 5.67212094333711 | 25.751573056929924 | 168.4006042550122 | 0.0 | 0.0 |
| SE_1_7_6 | SE_1 | SE_1_7 | GCTTCCAGCAGGAGGAATAG | 760.8113428651858 | 927.4362257402556 | 1021.1027759641186 | 974.6800586852339 | 1003.3224090420188 | 946.0106399763023 | 758.6932319990765 | 1241.249133100271 | 991.8795553307147 | 1249.5179034332941 | 1081.0559524867926 | 511.8603266130257 |
| SE_1_7_9 | SE_1 | SE_1_7 | AATGGTGCCAGGCTTCCAGC | 432.42695983363126 | 692.5789918512901 | 631.0296652234737 | 489.29199207696655 | 581.2057929757149 | 473.8716300980196 | 413.2456753016246 | 667.4195643326666 | 633.133503089346 | 363.77446546861944 | 576.3284176666832 | 704.5333747258653 |
| SE_10_1_10 | SE_10 | SE_10_1 | TTACCGGGAGAGCTCCCAGA | 50.93751155935005 | 79.95139877071169 | 114.55311606560711 | 49.44972260352321 | 56.21150917465856 | 20.791442636841808 | 51.655709412703075 | 76.57363273505099 | 58.607028336461205 | 32.076305572383276 | 39.90868879972959 | 8.124767089095647 |
| SE_10_1_16 | SE_10 | SE_10_1 | GGACGGGGTGTCCCAAAGGA | 365.2327956489567 | 527.6792318866972 | 612.2666548334173 | 392.9951638490529 | 423.1772105790333 | 422.7593336157835 | 278.7255987060437 | 387.59493112803585 | 569.1985630859339 | 407.514882158233 | 323.96465025662843 | 408.5597164802382 |
| SE_10_1_17 | SE_10 | SE_10_1 | GGGACCACAAGATTTGCACC | 1058.849974329468 | 967.4119251256116 | 952.963422442335 | 1113.9200670688388 | 1011.807165143854 | 878.4384514065664 | 1278.478807964401 | 861.2170298966845 | 1232.5235656213356 | 792.4305490268323 | 768.8291518771435 | 678.9983924458504 |
| SE_10_1_24 | SE_10 | SE_10_1 | CTTCAGGAAGTCCCGTCCCG | 356.5625809154503 | 440.7320857235482 | 383.1604227022031 | 340.94282426639694 | 412.57126545173924 | 262.49196329012784 | 223.84140745504666 | 426.3544242408394 | 209.56452556674006 | 172.77464592397357 | 342.74520969179525 | 94.01516203096391 |
| SE_10_1_27 | SE_10 | SE_10_1 | TCTTCTCACCCTATGGACAG | 184.24206308701082 | 227.86148649652833 | 207.38064115325426 | 147.04785932100324 | 201.5129574185873 | 245.16576109275968 | 94.70213392328898 | 199.46958650735505 | 318.78671473923595 | 203.39293760670304 | 132.63770101086598 | 536.2346278803126 |
| SE_10_2_1 | SE_10 | SE_10_2 | CTTCATTCTCAAAATAACCT | 47.686181034285156 | 56.96537162413208 | 44.43870881855448 | 66.36673296788642 | 36.06021343279983 | 64.97325824013066 | 40.8941032850566 | 41.59555358447214 | 49.72717555820951 | 49.5724722482287 | 18.780559435166865 | 33.659749369110536 |
| SE_10_2_14 | SE_10 | SE_10_2 | TCTGCAGAGTGCTTGCGGCA | 622.0879071290835 | 143.91251778728105 | 305.1458005540741 | 486.68937509783376 | 103.93826224748186 | 1127.0694529387997 | 830.7959930543078 | 229.72089820515296 | 716.604119204912 | 45.198430579267345 | 107.98821675220947 | 100.9792481073316 |
| SE_10_2_18 | SE_10 | SE_10_2 | GGGCTAGGCCAGGTGCACAG | 674.109195530122 | 784.5231004376085 | 803.8468661845188 | 814.6191144685666 | 753.0221040378788 | 698.2459485539374 | 737.1700197437835 | 632.4414851820877 | 897.7531158812467 | 526.3430141650165 | 707.7923337128512 | 851.9398633423149 |
| SE_10_2_19 | SE_10 | SE_10_2 | ACAGTCTTCCTCTTTCTAGG | 167.98541046168634 | 241.8529812814029 | 247.86924252127056 | 166.56748666449926 | 211.05830803315195 | 216.5775274671022 | 95.77829453605362 | 80.35504669727572 | 149.18152667462851 | 34.26332640686395 | 50.47275348201095 | 136.96035950189804 |
| SE_10_2_28 | SE_10 | SE_10_2 | AGACAGTCTGCAGAGCTCAG | 253.60378095506195 | 90.94471610168455 | 160.96687860943067 | 167.86879515406565 | 57.27210368738796 | 643.6684116322277 | 337.9144324080993 | 145.5844375456525 | 657.1091055906256 | 26.244250013768134 | 84.5125174582509 | 289.0095721692594 |
| SE_10_3_13 | SE_10 | SE_10_3 | CCCGCTATGCGGCAACCAGC | 6.5026610501297935 | 22.986027146579612 | 11.850322351614528 | 5.205233958265602 | 12.727134152752882 | 1.7326202197368175 | 0.0 | 5.67212094333711 | 1.7759705556503396 | 6.561062503442034 | 0.0 | 55.71268861094157 |
| SE_10_3_17 | SE_10 | SE_10_3 | GTTCAGGGAGAAGAGAGGGC | 164.73407993662144 | 249.84812115847404 | 201.45547997744697 | 166.56748666449926 | 251.3608995168694 | 147.27271867762948 | 120.52998862964051 | 106.82494443284891 | 166.94123223113192 | 104.97700005507254 | 99.77172199932397 | 64.99813671276517 |
| SE_10_3_36 | SE_10 | SE_10_3 | TGCACAAGCCCAACACCAAG | 704.4549470973943 | 730.5559062673781 | 770.2709528549443 | 806.8112635311683 | 850.5967992089842 | 719.0373911907792 | 743.6269834203714 | 863.1077368777969 | 811.6185439322052 | 763.9992781785835 | 847.4727445119048 | 578.0191443385188 |
| SE_10_3_42 | SE_10 | SE_10_3 | CTCAGGCTGCACTCACTCCG | 1954.049645564003 | 1624.0127875300814 | 1687.6834082424357 | 1823.133193882527 | 1826.3437509200385 | 1936.2030955558935 | 2037.1720399634776 | 1799.00769252842 | 1443.864061743726 | 2067.4636955290675 | 2081.120742409428 | 2012.6208760702643 |
| SE_10_3_6 | SE_10 | SE_10_3 | AGATGCACTGTGAGAGTGAG | 2507.8596116667236 | 2457.5061197147506 | 2627.8089814705218 | 2415.2285566352393 | 2516.790778706882 | 2213.4223307137845 | 2550.5006522522144 | 2556.235838463924 | 2643.5321720855304 | 2655.7723000043698 | 3041.276843532334 | 1895.3920937847415 |
| SE_10_4_15 | SE_10 | SE_10_4 | TAGCGGCTATTATGTATAGA | 1106.536155363753 | 1240.246073430665 | 1343.0365331829798 | 1100.9069821731748 | 1349.0762201918053 | 1237.0908368920877 | 1133.1971252411738 | 1882.1987996973644 | 1722.6914389808294 | 2233.677278949599 | 1786.500716270248 | 1232.6432355170823 |
| SE_10_4_17 | SE_10 | SE_10_4 | ACTAGTTCAAGAAAAGTTAG | 46.60240419259685 | 55.96597913949819 | 51.35139685699629 | 57.25757354092162 | 51.96913112374093 | 63.24063802039384 | 66.72195799140815 | 47.26767452780925 | 45.28724916908366 | 48.84346530340181 | 159.63475519891836 | 23.21362025455899 |
| SE_10_4_24 | SE_10 | SE_10_4 | CTCACCACCCAGCAGAGCGA | 958.0587280524562 | 874.4684240546592 | 952.963422442335 | 922.6277191025779 | 992.7164639147247 | 649.7325824013066 | 935.1835724924787 | 1247.8666075341641 | 867.5616164351909 | 1057.0600699989943 | 1293.511031097118 | 835.6903291641236 |
| SE_10_4_33 | SE_10 | SE_10_4 | TCGGTCTTCCCAACTCCCGT | 1951.8820918806264 | 1788.912547494674 | 1758.7853423521228 | 1758.067769404207 | 1514.5289641775928 | 1550.6950966644515 | 1775.6650110616683 | 1603.3195199832899 | 1807.0500403742205 | 1770.7578689845222 | 1652.689230294684 | 1561.115962119092 |
| SE_10_4_5 | SE_10 | SE_10_4 | GCAAGAGACAGCTTGTTGAG | 4163.870625766444 | 4618.192671493234 | 4517.935396553039 | 4524.649618222375 | 4447.072791874402 | 5012.470295698613 | 4302.490129833061 | 4455.4510009913 | 4616.635459413058 | 4262.503606402842 | 4808.997000367415 | 4730.935807879122 |
| SE_10_5_11 | SE_10 | SE_10_5 | TCACACTGGAAAATGAACGG | 1752.4671530099793 | 2033.7637062299787 | 2029.367702713988 | 1797.107024091199 | 1938.7667692693556 | 1903.283311380894 | 1660.515825495851 | 1901.105869508488 | 2211.083341784673 | 2079.127806646298 | 1759.5036620821957 | 2157.706002661258 |
| SE_10_5_26 | SE_10 | SE_10_5 | CTCCAGCCCCCAGAAAGCAT | 105.126353643765 | 87.94653864778286 | 115.54064292824165 | 68.96934994701923 | 123.02896347661118 | 19.9251325269734 | 54.88419125099702 | 95.48070254617468 | 89.68651306034215 | 33.53431946203706 | 58.68924823489645 | 89.37243798005211 |
| SE_10_5_28 | SE_10 | SE_10_5 | GAAGCACCAATGCTTTCTGG | 668.6903113216804 | 738.5510461444493 | 741.6326738385426 | 558.2613420239858 | 626.8113570230794 | 616.812798226307 | 595.11681885885 | 823.4028902744371 | 902.1930422703725 | 502.28578498572904 | 833.3873249355296 | 437.57674179843696 |
| SE_10_5_6 | SE_10 | SE_10_5 | GCCTCGACAAAAGTGCTGCT | 2654.169485294644 | 2462.50308213792 | 2501.4055430533 | 2577.8921178310393 | 2423.4584615866943 | 2034.962448080892 | 2466.560124456572 | 2442.793419597182 | 2599.1329081942717 | 3165.348154438368 | 1798.2385659172273 | 2555.8195900269448 |
| SE_10_5_8 | SE_10 | SE_10_5 | CTTTTGTCGAGGCCTCACAC | 37.93218945909046 | 53.96719417023039 | 48.38881626909266 | 37.737946197425615 | 55.15091466192915 | 78.83421999802519 | 21.52321225529295 | 59.557269905039654 | 44.39926389125849 | 37.17935418617152 | 80.99116256415711 | 104.46129114551545 |
| SE_10_6_1 | SE_10 | SE_10_6 | TGAGCCGAGAATAGACTGAA | 1469.6013973293334 | 1733.94596083981 | 1636.3320113854395 | 1656.5657072180277 | 1473.165778181146 | 1646.855518859845 | 1659.4396648830864 | 1434.1012451737327 | 1688.0600131456476 | 2383.852709583939 | 1332.2459349321496 | 2014.9422380957203 |
| SE_10_6_10 | SE_10 | SE_10_6 | AGGGAAACTGAGGCTCAAAG | 4198.55148470047 | 4676.157435602 | 4641.376254382357 | 4529.85485218064 | 4628.434453551131 | 4529.06925439204 | 4552.159391994459 | 4830.756336742105 | 5044.64436332479 | 4382.060745354452 | 4554.285663027965 | 4177.29096480789 |
| SE_10_6_15 | SE_10 | SE_10_6 | AGAATTCCAAGGGTAGACAA | 9.75399157519469 | 31.980559508284678 | 38.51354764274722 | 14.314393385230405 | 12.727134152752882 | 46.78074593289407 | 18.294730416999005 | 12.289595377230405 | 24.863587779104755 | 2.9160277793075706 | 7.042709788187574 | 104.46129114551545 |
| SE_10_6_18 | SE_10 | SE_10_6 | CAAGCAACTCCAGAATCCTA | 448.68361245895574 | 613.6269855652123 | 524.3767640589429 | 510.112927910029 | 611.9630338448677 | 660.9946138295959 | 509.02396983767824 | 622.9879502765259 | 693.5165019814576 | 609.4498058752822 | 671.4049998072154 | 591.9473164912542 |
| SE_10_6_27 | SE_10 | SE_10_6 | GGGATTCTTGTGACTTGCAG | 28.17819788389577 | 62.96172653193546 | 55.301504307534465 | 42.943180155691216 | 25.454268305505764 | 105.68983340394587 | 34.43713960846872 | 12.289595377230405 | 57.71904305863604 | 26.244250013768134 | 3.521354894093787 | 111.42537722188314 |
| SE_10_7_2 | SE_10 | SE_10_7 | AAGGGGCAGGAACATAGGAG | 3316.3571355661948 | 3833.669571055626 | 3721.9887452695966 | 3405.5243171952698 | 3154.2080808572555 | 3189.753824535481 | 3168.216843979122 | 3070.508137326489 | 3492.446097686393 | 3403.733425396762 | 3655.166380069351 | 2732.243103961593 |

| | | | | | | | | | | | | | | | |
|---|---|---|---|---|---|---|---|---|---|---|---|---|---|---|---|
| SE_10_7_20 | SE_10 | SE_10_7 | CTTGGTAGCAGGCTGAAGGG | 2077.600205516469 | 2129.7053847548327 | 2169.5965172080932 | 2032.6438607027176 | 2077.7046504369077 | 2024.5667267624713 | 2022.1057913847726 | 1916.231525357387 | 2050.358006498317 | 1706.6052578397557 | 1916.7908473517182 | 1713.1651747864535 |
| SE_10_7_22 | SE_10 | SE_10_7 | GGGTTCTGTCTGGATCCCTA | 6.5026610501297935 | 4.996962423169481 | 7.900214901076352 | 5.205233958265602 | 8.484756101835254 | 0.0 | 1.0761606127646475 | 62.393330376708214 | 0.0 | 2.187020834480678 | 153.7658303754287 | 0.0 |
| SE_10_7_30 | SE_10 | SE_10_7 | GCAGACTGAAGCACACCAGT | 7848.711887506661 | 7316.552380004754 | 7142.781797435657 | 8134.479368279569 | 7086.892534057896 | 6239.16541127228 | 8129.317268824147 | 7149.708449076427 | 7807.166562638893 | 7298.088524662023 | 7755.197261759217 | 6510.259800391068 |
| SE_10_7_7 | SE_10 | SE_10_7 | ATGGGCTTGCAGGTTAAAAG | 2146.96192338452 | 2672.3755039110383 | 2719.6489796955343 | 2222.634900179412 | 2515.730184194153 | 2736.6736370743033 | 2099.5893555038274 | 2442.793419597182 | 2720.7868912563204 | 2187.7498414255047 | 2428.5610919600153 | 2922.594790048977 |
| SE_10_8_29 | SE_10 | SE_10_8 | TGGAAGCCCCAAGCAGGCAC | 143.05854310285545 | 150.9082651797183 | 203.43053370271608 | 149.65047630013606 | 165.45274398578746 | 36.38502461447317 | 116.22534617858193 | 95.48070254617468 | 103.0062922277197 | 217.97307650324092 | 70.42709788187574 | 179.90555697283216 |
| SE_10_8_32 | SE_10 | SE_10_8 | CCTCAACTGGTTTTACTCTG | 2044.0031234241317 | 2433.520700083537 | 2362.1642554218292 | 2412.6259396561063 | 2503.0030500414 | 2865.753843444696 | 1839.1584872147826 | 2187.5479771470123 | 2474.814969298748 | 2742.52412643877 | 1726.6376830706536 | 2707.868802694306 |
| SE_10_8_4 | SE_10 | SE_10_8 | TGTGTGTGTGTGTATGGTCT | 4526.935867732024 | 3886.6373727412224 | 3960.970246027156 | 4096.519125155029 | 3919.9573190478873 | 3266.855424313769 | 4421.943957849937 | 3867.4411298653527 | 4224.145966614333 | 3683.6720922102886 | 3688.032359080893 | 3433.2944356492744 |
| SE_10_8_44 | SE_10 | SE_10_8 | ACTGCCGCATGGGTATCACA | 3331.530011349831 | 3280.006134568447 | 3306.2399361004536 | 3323.5418823525865 | 3499.9618920070425 | 3552.737760570344 | 3191.8923774599443 | 3363.5677193989063 | 3366.352188235219 | 2754.1882375560003 | 3393.412332941713 | 3027.056081194492 |
| SE_10_8_5 | SE_10 | SE_10_8 | TGTGAGCTGCAGCCCTGCCT | 3420.399712368271 | 684.5838519742189 | 1456.6021223859525 | 2760.0753063703355 | 668.1745430195263 | 6933.946119386744 | 4339.079590667058 | 1796.1716320567516 | 5773.680276419254 | 249.3203751307973 | 980.1104455227708 | 1866.3750684665426 |
| SE_100_1_15 | SE_100 | SE_100_1 | ATTCTAGATGAGATTCTAGG | 0.0 | 0.9993924846338962 | 5.925161175807264 | 0.0 | 0.0 | 0.0 | 0.0 | 0.0 | 0.0 | 0.0 | 0.0 | 0.0 |
| SE_100_1_16 | SE_100 | SE_100_1 | AGGACAACAGACACTGGCTG | 10882.20326739221 | 10921.361072079217 | 10718.61656703534 | 10811.270931317655 | 9937.770584274542 | 10374.063565674194 | 10803.576391544297 | 9925.266297349386 | 10134.575975818663 | 10264.41778316265 | 9041.665583068147 | 10518.091337340678 |
| SE_100_1_3 | SE_100 | SE_100_1 | AATCTGAAGAAACCAGAGGA | 6.5026610501297935 | 14.990887269508443 | 15.800429802152705 | 9.109159426964803 | 1.0605945127294067 | 51.978606592104526 | 2.152321225529295 | 10.398888396118036 | 0.0 | 25.515243068941242 | 15.259204541073078 | 74.28358481458876 |
| SE_100_1_4 | SE_100 | SE_100_1 | ATCAAATCTGAAGAAACCAG | 47.686181034285156 | 73.95504386290831 | 73.07698783495626 | 36.43663770785921 | 47.72675307282331 | 3.465240439473635 | 13.990087965940416 | 145.5844375456525 | 40.84732277995781 | 32.076305572383276 | 43.43004369382337 | 2.321362025455899 |
| SE_100_1_7 | SE_100 | SE_100_1 | TCTCATTGAGAGCTAAAGCA | 166.90163361999802 | 158.9034050567895 | 147.14150253254707 | 140.54131687317124 | 147.42263726938754 | 38.98395494407839 | 173.26185865510826 | 249.57332150683285 | 205.12459917761421 | 106.43501394472632 | 117.3784964697929 | 221.69007343103834 |
| SE_100_2_0 | SE_100 | SE_100_2 | GTTTTGTGACATGCAGGATG | 83.45081680999901 | 104.9362108865591 | 92.82752508764715 | 65.06542447832003 | 85.90815553108195 | 161.133680435524 | 60.26499431482026 | 74.68292575393862 | 35.51941111300679 | 45.198430579267345 | 113.85714157569912 | 112.5860582346111 |
| SE_100_2_10 | SE_100 | SE_100_2 | TGCAACCACCAACTGAAAGG | 33.59708209233727 | 42.973876839257535 | 38.51354764274722 | 50.751031093089615 | 19.090701229129323 | 41.582885273683615 | 29.056336544645482 | 61.44797688615203 | 52.391131391685015 | 28.431270848248815 | 57.515463270198524 | 52.23064557275772 |
| SE_100_2_26 | SE_100 | SE_100_2 | GGTTGGCAAGATGACTAAGG | 344.64103565687907 | 321.8043800521146 | 313.04601545515044 | 394.2964723386193 | 237.57317085138712 | 449.61494702170415 | 393.87478427186096 | 272.2618052801813 | 238.86803973497067 | 368.8775140824077 | 231.235638045492 | 286.68821014380353 |
| SE_100_2_28 | SE_100 | SE_100_2 | GGGGTTGGCAAGATGACTAA | 193.9960546622055 | 205.8748518345826 | 236.018920169656 | 239.44076208021767 | 218.48246962225778 | 96.16042219539337 | 161.42409191469713 | 184.3439306584561 | 261.0676716805999 | 183.70975009637695 | 208.93372371623136 | 94.01516203096391 |
| SE_100_3_11 | SE_100 | SE_100_3 | CATCAAAGATGCTAGATGGA | 68.27794102636283 | 64.96051150120326 | 37.526020780112674 | 102.80337067574564 | 112.42301834931712 | 85.76470087697247 | 100.08293698711222 | 95.48070254617468 | 56.831057780810866 | 38.63736807582531 | 158.4609702342204 | 105.6219721582434 |
| SE_100_3_20 | SE_100 | SE_100_3 | GATGCTGTGACAACTTTCAG | 1405.6585636697237 | 1601.0267603835016 | 1691.633515692974 | 1359.8673715968885 | 1448.7721043883696 | 1405.154998206559 | 1447.4360241684508 | 1340.5112496086704 | 1591.2696178627043 | 1202.8614589643728 | 1649.1678754005902 | 1338.2652076753257 |
| SE_100_3_5 | SE_100 | SE_100_3 | TAATGTCCCATAATGCCCAA | 1225.751607949466 | 1174.286169444828 | 1297.610297501791 | 1210.2168952967525 | 1204.835366460606 | 1431.1443015026111 | 1277.4026473516365 | 1354.691551967013 | 1272.4829031234683 | 1358.868945157328 | 872.1222287705613 | 1087.5581089260886 |
| SE_100_3_7 | SE_100 | SE_100_3 | GGTGGATTCAGAAAGAGTAA | 30.34575156727237 | 79.95139877071169 | 59.25161175807264 | 57.25757354092162 | 61.51448173830559 | 41.582885273683615 | 55.960351863761666 | 16.071009339455145 | 61.27098416993672 | 8.748083337922711 | 1.173784964697929 | 113.74673924733905 |
| SE_100_3_9 | SE_100 | SE_100_3 | TCATTACCACCACAGCCATT | 3895.0939690277464 | 3526.8560782730196 | 3458.319072946173 | 4056.17856197847 | 3681.323553683771 | 3280.7163860716637 | 3775.171429578383 | 3683.0971992068967 | 3723.322269920937 | 4035.7824465616777 | 3663.3828748222363 | 3764.08852427674 |
| SE_100_4_11 | SE_100 | SE_100_4 | TGTAAACTCTGGCAGCCTTG | 1969.222521347639 | 2013.7758565373008 | 2003.69200428549 | 1868.678991017351 | 1857.1009917891913 | 1863.433046326947 | 1716.4761773596126 | 1768.7563808306222 | 1662.3084400887178 | 1640.2656258605084 | 1582.2621324128083 | 1331.301121598958 |
| SE_100_4_12 | SE_100 | SE_100_4 | GCATTGTGCCATGGATCAAG | 2843.8304325900963 | 2230.644025702856 | 2238.7233975925114 | 2671.58632907982 | 2473.3064036849764 | 2145.8501421440483 | 3101.494885987714 | 1988.078390639657 | 2123.172799279981 | 2360.5244873494785 | 2362.829133936931 | 1541.3843849027169 |
| SE_100_4_18 | SE_100 | SE_100_4 | TTTATTTACACATTTGCGGG | 2.167553683376598 | 3.9975699385355847 | 4.93763431317272 | 2.602616979132801 | 8.484756101835254 | 0.0 | 0.0 | 22.68848377334844 | 0.0 | 18.225173620672315 | 31.69219404684083 | 1.1606810127279494 |
| SE_100_4_4 | SE_100 | SE_100_4 | TTTGGTGTACTTCCAAACCT | 20.59175999207768 | 19.987849692677923 | 22.713117840594514 | 13.013084895664004 | 27.575457330964575 | 0.8663101098684087 | 8.60928490211718 | 41.59555358447214 | 26.639558334755094 | 19.683187510326103 | 82.16494752885504 | 37.14179240729438 |
| SE_100_4_5 | SE_100 | SE_100_4 | GCAAGCTCCTTTTCTCCAGA | 716.3764923559655 | 757.5395033524933 | 779.1586946186552 | 706.6105098345554 | 655.4474088667733 | 618.5454184460439 | 642.4678858204945 | 829.0750112177742 | 773.4351769857229 | 668.4993684062606 | 467.16641594977574 | 330.7940886274656 |
| SE_101_1_0 | SE_101 | SE_101_1 | AACAGGGTGAAGATTTGTGG | 725.046707089472 | 728.5571212981104 | 744.5952544264462 | 642.8463938458018 | 800.7488571107021 | 822.9946043749883 | 803.8919777351916 | 907.5393509339376 | 666.8769436467026 | 877.7243615715788 | 996.5434350285417 | 968.0079646151098 |
| SE_101_1_16 | SE_101 | SE_101_1 | AAGTGAAGACGATGTCTACG | 674.109195530122 | 676.5887120971477 | 632.0171920861081 | 766.4707003546099 | 613.0236283575971 | 706.9090496526215 | 700.5805589097855 | 671.2009782948913 | 630.4695472558706 | 726.819923992412 | 502.3799648907136 | 663.9095392803871 |
| SE_101_1_23 | SE_101 | SE_101_1 | ATGACATCAGTGCCCCCAAG | 1731.8753930179016 | 1818.894322033691 | 1735.0846976488938 | 1724.2337486754807 | 1731.9508392871212 | 1605.2726335861614 | 1938.16526358913 | 1915.2861718668307 | 1688.0600131456476 | 2111.933119163508 | 2122.2032161738557 | 1872.1784735301824 |
| SE_101_1_36 | SE_101 | SE_101_1 | GATTCTGAATGGTCTCCAGG | 10.837768416882989 | 23.98541963121351 | 22.713117840594514 | 3.9039254686992013 | 60.45388722557618 | 23.390372966447035 | 9.685445514881827 | 4.726767452780925 | 22.199631945629246 | 6.561062503442034 | 1.173784964697929 | 25.534982280014887 |
| SE_101_1_8 | SE_101 | SE_101_1 | CGAAGTCTCGTGGGGAAGAG | 2248.83694650322 | 2543.453873393266 | 2446.1040387457656 | 1946.757500391335 | 2293.0053365209774 | 2160.5774140118115 | 2004.8872215805382 | 1810.3519344150943 | 2023.718448163562 | 1588.5061327777992 | 1406.194387708119 | 1122.3785393079272 |
| SE_101_2_26 | SE_101 | SE_101_2 | GGCACAGCTTTGGAGTTGGA | 1163.976327973233 | 1414.140365756963 | 1402.2881449410525 | 1387.1948498777829 | 1255.7439030716175 | 1426.8127509532692 | 1153.6441768837021 | 1339.565896118114 | 1202.3320661752798 | 1312.2125006884069 | 1300.5537408853054 | 1541.3843849027169 |
| SE_101_2_36 | SE_101 | SE_101_2 | CTTCCTGGAGGGTCTGCGGA | 119.21545258571288 | 132.9192004563082 | 121.46580410404891 | 118.41907255054244 | 136.81669214209347 | 76.23528966841997 | 109.76838250199404 | 192.85211207346174 | 249.5238630688727 | 142.88536118607095 | 78.64359263476125 | 127.67491140007444 |
| SE_101_2_45 | SE_101 | SE_101_2 | CTTGGTGCTGGCAGAAGCCG | 383.6570019576578 | 468.71507529329733 | 425.62407779548846 | 387.78992989078733 | 457.1162349863743 | 228.7058690052599 | 330.3813081187468 | 378.141396222474 | 264.6196127919006 | 395.8507710410027 | 299.3151659979719 | 264.6352709019725 |
| SE_101_2_8 | SE_101 | SE_101_2 | GCTGCCAGTGGAAAGTCTGA | 1678.770327775175 | 2002.782539206328 | 1985.9165207580681 | 1704.7141213319846 | 1917.5548790147675 | 2396.2137638960185 | 1581.9561007640318 | 2535.438061671688 | 2124.060784557806 | 2049.2385219083953 | 2891.032368050999 | 2274.934784946781 |
| SE_101_2_9 | SE_101 | SE_101_2 | GCGAGCTCAGATCCTGGCAA | 5410.213993707988 | 5271.795356443802 | 5305.981832935405 | 5728.359971071295 | 5509.7884936292685 | 5552.181494146632 | 5702.575087039867 | 5475.487417301423 | 5572.995603630766 | 6317.574183869851 | 5806.7142203606545 | 6744.7173649621145 |
| SE_101_3_13 | SE_101 | SE_101_3 | TCTGAGGTATGAGGAGGTTA | 923.3778691184307 | 948.4234679175675 | 988.5143894971786 | 959.0643568104372 | 961.9592230455719 | 817.7967437157779 | 873.8424175648937 | 1023.8178302723484 | 949.2562619951065 | 821.590826819908 | 1025.88805914599 | 819.4407949859323 |
| SE_101_3_14 | SE_101 | SE_101_3 | GTGACTGGACATTTTCTCCC | 743.470913398173 | 899.4532361705066 | 888.7741763710897 | 814.6191144685666 | 826.2031254162079 | 799.6042314085413 | 754.3885895480179 | 870.6705648022464 | 1082.454053668882 | 1182.4492645092198 | 793.4786361358 | 1100.325600066096 |
| SE_101_3_18 | SE_101 | SE_101_3 | TATAAATGAAAATGTCAGGA | 21.675536833765978 | 25.984204600481302 | 37.526020780112674 | 10.410467916531204 | 31.8178353818822 | 24.256683076315444 | 4.30464245105859 | 24.57919075446081 | 28.415528890405433 | 15.309145841364746 | 11.73784964697929 | 2.321362025455899 |
| SE_101_3_4 | SE_101 | SE_101_3 | AGTCCAAATCCCTCTCCTGA | 161.48274941155654 | 235.8566263735995 | 237.00644703229057 | 175.67664609146405 | 223.78544218590483 | 275.486614938154 | 164.65257375299106 | 136.13090264009065 | 134.08577695160065 | 293.0607918204108 | 188.97937931636656 | 133.47831646371418 |
| SE_101_3_9 | SE_101 | SE_101_3 | TCCCAGTCTGAGGTATGAGG | 124.63433679415438 | 166.89854493386065 | 159.97935174679614 | 127.52823197750725 | 148.48323178211695 | 55.44384703157816 | 110.8445431147587 | 162.60080037566382 | 125.20592417334893 | 61.23658336545898 | 143.20176569314734 | 109.10401519642724 |
| SE_101_4_21 | SE_101 | SE_101_4 | CAGAGTGCAAGGAAGTGGGA | 2867.673523107239 | 2587.427142717157 | 2458.941887960015 | 2611.726138559766 | 2641.9409312089524 | 2045.358169399313 | 2929.3091879453705 | 2991.0984441197693 | 2713.683009033719 | 2622.237980542333 | 2829.995549886707 | 1940.6586532811314 |
| SE_101_4_24 | SE_101 | SE_101_4 | AGACAGGAACATGTTCCTGT | 152.81253467805016 | 148.90948021045054 | 159.97935174679614 | 174.37533760189766 | 177.11928362581094 | 127.34758615065608 | 188.3281072338133 | 146.52979103620868 | 119.87801250639792 | 128.3052222895331 | 109.1620017169074 | 145.08512659099367 |
| SE_101_4_31 | SE_101 | SE_101_4 | AATAAATGACCCTAAAACAG | 13.005322100259587 | 15.990279754142339 | 10.862795488979984 | 1.3013084895664004 | 29.69664635642339 | 30.320853845394307 | 22.599372868057596 | 17.961716320567515 | 17.759705556503395 | 0.7290069448268927 | 2.347569929395858 | 1.1606810127279494 |
| SE_101_4_4 | SE_101 | SE_101_4 | TTGAAAGGCACACCTCTCGG | 481.1969177096047 | 570.6531087259547 | 568.8154728774973 | 689.6934994701923 | 676.6592991213615 | 615.0801780065702 | 585.4313733439682 | 542.6329035792502 | 539.0070636398781 | 793.1595559716592 | 645.5817305838609 | 565.2516531985113 |
| SE_101_4_8 | SE_101 | SE_101_4 | GTCCTGTCTTGTCTTCACCT | 272.027987263763 | 259.842046004813 | 289.3453707519214 | 243.34468754891688 | 204.6947409567755 | 388.97323933091553 | 167.88105559128502 | 214.595242356254 | 252.18781890234823 | 185.16776398603074 | 240.62591776307545 | 540.8773519312244 |
| SE_101_5_0 | SE_101 | SE_101_5 | CACCCTACTCCATCCCACAG | 668.6903113216804 | 820.5012298844288 | 785.0838557944625 | 784.6890192085394 | 724.3860521941848 | 1191.176401069062 | 685.5143103310804 | 705.2337039549141 | 754.7874861513943 | 958.6441324473639 | 784.0883564182166 | 1133.9853494352067 |
| SE_101_5_15 | SE_101 | SE_101_5 | CCTCTGCATGGGGTTTTCCA | 70.44549470973944 | 133.9185929409421 | 125.4159115545871 | 84.58505182181602 | 111.36242383658771 | 74.50266944868315 | 55.960351863761666 | 90.75393509339376 | 190.02884945458632 | 107.8930278343801 | 129.1163461167722 | 112.5860582346111 |
| SE_101_5_17 | SE_101 | SE_101_5 | GCACTATCTGCAGATAGCTG | 150.64498099467355 | 51.968409200962604 | 80.97720273603261 | 70.27065843658562 | 34.999618920070425 | 415.8288527368362 | 168.95721620404964 | 70.90151179171387 | 322.3386558505366 | 21.141201399979888 | 76.29602270536539 | 95.17584304369186 |
| SE_101_5_4 | SE_101 | SE_101_5 | TCTACTGGCCACATATAAAA | 114.88034521895969 | 178.8912547494674 | 140.22881449410525 | 140.54131687317124 | 143.1802592184699 | 119.5507951618404 | 129.1392735317577 | 94.5353490556185 | 207.78855501108973 | 201.20591677222237 | 76.29602270536539 | 83.56903291641235 |
| SE_101_5_5 | SE_101 | SE_101_5 | AGATTTCTACTGGCCAGCGT | 41.18351998415536 | 64.96051150120326 | 84.92731018657079 | 57.25757354092162 | 58.33269820011737 | 163.73261076512924 | 36.58946083399801 | 45.37696754669688 | 93.23845417164283 | 135.59529173780203 | 62.210603128990236 | 104.46129114551545 |
| SE_102_1_18 | SE_102 | SE_102_1 | GTAATGACAGACAGAGCCCC | 552.7261892610325 | 607.6306306574089 | 574.7406340533046 | 533.5364807222242 | 589.6905490775501 | 706.0427395427531 | 471.3583483909156 | 556.813205937593 | 667.7649289245277 | 502.28578498572904 | 794.6524211004979 | 1241.928683618906 |
| SE_102_1_26 | SE_102 | SE_102_1 | TGGTAGTGAGGAGTACTGAG | 2.167553683376598 | 5.9963549078033775 | 0.987526862634544 | 7.807850937398403 | 8.484756101835254 | 0.0 | 1.0761606127646475 | 0.0 | 0.0 | 0.0 | 0.0 | 0.0 |
| SE_102_1_36 | SE_102 | SE_102_1 | TGGAGATGTGGTTTTCAAGA | 235.17957464636086 | 350.7867621064976 | 336.74666015837954 | 249.85122999674888 | 296.9664635642339 | 255.56148241118058 | 227.06988929334062 | 282.66069367629933 | 334.770449740089 | 279.93866681352677 | 269.9705418805237 | 104.46129114551545 |
| SE_102_1_4 | SE_102 | SE_102_1 | TATTGGTGCCCCTACTCCCA | 1114.1225932555712 | 1166.2910295677568 | 1156.393956145051 | 1220.6273632132836 | 1012.8677596565834 | 1312.4598164506392 | 1154.7203374964668 | 1297.970342533642 | 963.4640264403092 | 1618.3954175157016 | 1258.29748215618 | 749.7999342222554 |
| SE_102_2_21 | SE_102 | SE_102_2 | GGTGTACTCACATGAGTTCT | 699.0360628889528 | 759.5382883217611 | 754.4705230527917 | 767.7720088441763 | 853.7785827471724 | 490.33152218551936 | 707.0375225863734 | 635.2775456537563 | 848.9139256008623 | 621.1139169925125 | 606.8468267488294 | 601.2327645930778 |
| SE_102_2_22 | SE_102 | SE_102_2 | TGAGTTCTCGGACCATGTCC | 95.37236206857031 | 112.93135076363026 | 104.67784743926167 | 84.58505182181602 | 116.66539640023474 | 138.6096175789454 | 69.95043982970209 | 134.24019565897828 | 89.68651306034215 | 176.41968064810803 | 90.38144228174053 | 141.60308355280984 |
| SE_102_2_23 | SE_102 | SE_102_2 | GAGTTCTCGGACCATGTCCC | 187.49339361207572 | 169.89672238776234 | 172.81720096104522 | 205.60674135149128 | 219.5430641349872 | 68.4384986796043 | 193.70891029763655 | 199.46958650735505 | 175.8210850093836 | 137.0533056274558 | 328.65979011542015 | 94.01516203096391 |
| SE_102_2_25 | SE_102 | SE_102_2 | GTCCCGGGTGATGAGCTGGA | 451.93494298402067 | 562.6579688488836 | 588.5660101301883 | 438.54096098387697 | 515.4489331864917 | 656.6630632802538 | 435.84504816968223 | 418.79159631639 | 565.6466219746331 | 485.5186252547105 | 550.5051484433287 | 420.1665266075177 |
| SE_102_2_26 | SE_102 | SE_102_2 | GACCATCCAGCTCATCACCC | 7938.66536536679 | 7597.381668186878 | 7518.042005236784 | 7076.5155662620855 | 6892.803738228415 | 7721.422009257127 | 7004.7294284850905 | 5374.334593811912 | 5981.468831430344 | 5992.437086477058 | 5871.272393419041 | 5754.656461105174 |
| SE_102_3_29 | SE_102 | SE_102_3 | GTGAGGTAAGCTTCCAGCCC | 5777.614343040322 | 5389.723669630602 | 5392.884196847245 | 5803.835863466146 | 4937.067456755389 | 5205.657450199268 | 5659.528662529281 | 5265.61894239795 | 5390.958621676606 | 5628.662621008438 | 5069.577262530355 | 5748.853056041534 |
| SE_102_3_30 | SE_102 | SE_102_3 | CTTTGGCTTTGGCTTTGGTG | 127.88566731921927 | 154.90583511825392 | 196.51784566427426 | 144.44524234187045 | 146.36204275665813 | 144.67378834802426 | 131.291594757287 | 69.0108048106015 | 109.22218917249589 | 145.07238202055163 | 77.46980767006332 | 273.9207190037961 |
| SE_102_3_31 | SE_102 | SE_102_3 | AGCCAAAGAAAAGGCGGACA | 1820.7450940363422 | 2027.7673513221753 | 1928.6399627252645 | 2033.9451691922839 | 2119.067836433355 | 2404.010554884834 | 2156.6258679803536 | 2264.1216098820632 | 2329.18538373542 | 2469.875529073512 | 2197.325453914523 | 2676.5304153506513 |
| SE_102_3_32 | SE_102 | SE_102_3 | GCCAAAGAAAAGGCGGACAT | 6488.571951187845 | 6849.836089680724 | 6310.296652234736 | 6804.542091942708 | 7140.982854207095 | 7006.71616861569 | 6600.0930380855825 | 7258.424100490389 | 6639.465922298795 | 7187.279469048334 | 7359.631728656015 | 6923.462240922218 |
| SE_102_3_33 | SE_102 | SE_102_3 | ACCCATGTCCGCCTTTTCTT | 141.97476626116716 | 172.89489984166403 | 179.729888999487 | 167.86879515406565 | 214.24009157134017 | 276.35292504802237 | 180.79498294446077 | 218.37665631847872 | 185.58892306546048 | 294.5188057100646 | 240.62591776307545 | 467.7544481293636 |
| SE_102_4_11 | SE_102 | SE_102_4 | GAAAAGGCGGACATGGGTGG | 1231.1704921579076 | 1517.0777916742545 | 1429.9388970948198 | 1229.7365226402485 | 1376.65167752277 | 1310.7271962309023 | 1148.2633738198788 | 1601.4288130021773 | 1231.6355803435106 | 1006.0295838611119 | 1571.6980677305269 | 1142.1101165243022 |
| SE_102_5_10 | SE_102 | SE_102_5 | GCCTCTGGTCAGGAGCACAG | 490.9509092847994 | 549.665866548643 | 587.5784832675537 | 530.9338637430914 | 527.1154728265152 | 653.1978228407802 | 420.77879959097714 | 611.6437083898517 | 634.0214883671712 | 454.17132662715414 | 630.3225260427879 | 544.3593949694083 |
| SE_102_5_11 | SE_102 | SE_102_5 | GTCTCTCAATGGGCAGTAGA | 28.17819788389577 | 68.95808143973883 | 50.36386999436174 | 57.25757354092162 | 61.51448173830559 | 46.78074593289407 | 29.056336544645482 | 43.48626056558451 | 98.56636583859384 | 30.61829168272949 | 89.2076573170426 | 169.4594278582806 |
| SE_102_5_12 | SE_102 | SE_102_5 | TGTGCTCCTGACCAGAGGCA | 974.3153806777807 | 1056.3578562580283 | 1019.1277222388494 | 955.1604313417379 | 922.7172260745839 | 693.9143980045953 | 772.6833199650168 | 675.9277457476722 | 1055.8144953341268 | 992.1784519094009 | 726.5728931480181 | 643.0172810512839 |
| SE_102_5_30 | SE_102 | SE_102_5 | GTAGGGATAACAAGGAGTGG | 2266.1773759702332 | 2383.5510758518426 | 2465.8545759984563 | 2200.512655856783 | 2282.3993913936833 | 2444.7271300486495 | 2142.6357800144133 | 2151.624544505877 | 2444.6234698526923 | 1638.8076119708546 | 2286.5331112315657 | 2202.972562157648 |
| SE_102_5_4 | SE_102 | SE_102_5 | ATTGTTATCTAACTCCTTCA | 27.094421042207472 | 19.987849692677923 | 24.688171565863602 | 16.917010364363207 | 8.484756101835254 | 50.2459863723677 | 34.43713960846872 | 12.289595377230405 | 26.639558334755094 | 0.7290069448268927 | 18.780559435166865 | 53.39132658548567 |
| SE_103_1_17 | SE_103 | SE_103_1 | CTTCTTGTGTGCGGTCTGAG | 224.34180622947787 | 289.82382054382987 | 237.00644703229057 | 210.81197530975686 | 317.11775930609264 | 312.73794966249557 | 331.4574687315114 | 364.90644735468743 | 412.0251689108788 | 323.67908350314036 | 328.65979011542015 | 649.9813671276517 |
| SE_103_1_25 | SE_103 | SE_103_1 | GTCCTCTGCAGAGAGACCTT | 0.0 | 0.0 | 1.975053725269088 | 2.602616979132801 | 3.1817835381882205 | 29.454543735525895 | 0.0 | 0.0 | 8.879852778251697 | 0.0 | 0.0 | 12.767491140007444 |
| SE_103_1_38 | SE_103 | SE_103_1 | GCAAGCTGTGGGAACCACAC | 3178.717476671781 | 2784.3074621900346 | 2808.5263973326432 | 3197.314958864646 | 2622.850229979823 | 2572.0747161993054 | 2942.2231152985464 | 2757.5961319523917 | 2771.402052092355 | 2552.2533138389513 | 2049.428548362584 | 2193.6871140558246 |
| SE_103_1_40 | SE_103 | SE_103_1 | TTTGTCAGCAGTGAAGGAGC | 231.92824412129596 | 316.8074176289451 | 294.28300506509413 | 222.52375171585447 | 306.51181417879855 | 356.9197652657844 | 167.88105559128502 | 195.6881725451303 | 301.91499446055775 | 304.7249029376411 | 272.3181118099195 | 510.69964560029774 |
| SE_103_1_5 | SE_103 | SE_103_1 | GCGACCATAGAAAATATTAC | 391.2434398494759 | 487.70353250134133 | 575.7281609159392 | 486.68937509783376 | 607.7206557939501 | 393.30478988025754 | 359.43764466339223 | 639.058959615981 | 469.7442119695148 | 479.6865696960954 | 902.6406378527074 | 464.2724050911798 |
| SE_103_2_1 | SE_103 | SE_103_2 | GTAATAAGAAGTCCAGAGAT | 65.02661050129794 | 157.9040125721556 | 160.96687860943067 | 104.10467916531204 | 168.63452752397566 | 175.86095230328698 | 104.3875794381708 | 91.69928858394995 | 99.45435111641902 | 145.80138896537852 | 57.515463270198524 | 175.26283292192036 |
| SE_103_2_15 | SE_103 | SE_103_2 | GTACTGTGATCTGCAGCACC | 191.8285009788289 | 41.97448435462364 | 85.91483704920533 | 115.81645557140963 | 24.393673792776355 | 316.2031901019692 | 217.3844437784588 | 99.26211650839943 | 303.69096501620805 | 12.393118062057175 | 26.997054188052367 | 99.81856709460365 |
| SE_103_2_17 | SE_103 | SE_103_2 | TTTCTTCACACAAGGAAGGA | 8.670214733506391 | 5.9963549078033775 | 5.925161175807264 | 2.602616979132801 | 12.727134152752882 | 0.0 | 16.142409191469714 | 2.836060471668555 | 16.871720278678225 | 0.7290069448268927 | 0.0 | 0.0 |

| | | | | | | | | | | | | | | | |
|---|---|---|---|---|---|---|---|---|---|---|---|---|---|---|---|
| SE_103_2_29 | SE_103 | SE_103_2 | CATGGGTTCCCACCTAGTGG | 535.3857597940197 | 629.6172653193546 | 650.7802024761645 | 490.59330056653295 | 659.689786917691 | 716.438460861174 | 418.6264783654479 | 830.0203647083305 | 552.3268428072556 | 641.5261114476656 | 1082.2297374514906 | 255.34982280014887 |
| SE_103_2_7 | SE_103 | SE_103_2 | TTTGGCGAGGGTGTTAGCAG | 377.154340907528 | 450.7260105698872 | 409.8236479933358 | 413.81609966211535 | 335.1478660224925 | 343.92511361775826 | 428.3119238803297 | 394.21240556192913 | 593.1741655872135 | 508.84684748917107 | 380.306328562129 | 182.22691899828806 |
| SE_103_3_14 | SE_103 | SE_103_3 | GCTACGGCTTAGAAAGGCTG | 2853.5844241652912 | 2784.3074621900346 | 2774.9504840030686 | 2680.695488506785 | 2773.454650787399 | 2717.614814657198 | 2739.9049200987924 | 2292.4822145987487 | 2712.795023755894 | 3691.6911686033845 | 2366.350488831025 | 2353.8610938122815 |
| SE_103_3_3 | SE_103 | SE_103_3 | TATAATTCCAGCATGACACC | 346.80858934025565 | 313.8092401750434 | 364.39741231214674 | 385.18731291165454 | 269.3910062332693 | 447.88232680196734 | 354.056841599569 | 311.96665188354103 | 264.6196127919006 | 331.6981598962362 | 226.5404981867003 | 408.5597164802382 |
| SE_103_3_5 | SE_103 | SE_103_3 | CATGCTGGAATTATAGGCAC | 3947.115257428785 | 3852.6580282636696 | 3751.614551148633 | 3849.2705121374124 | 3537.0826999525716 | 4173.015799236125 | 3535.187612931867 | 4068.80142335382 | 3805.0169154808527 | 2840.211057045574 | 4001.43294465524 | 3116.4285191745444 |
| SE_103_3_6 | SE_103 | SE_103_3 | GCTGGAATTATAGGCACAGG | 775.984218648822 | 831.4945472154017 | 885.811595783186 | 843.2479012390274 | 866.5057168999253 | 693.048087894727 | 768.3786775139583 | 589.9005781070595 | 719.2680750383876 | 864.6022365646946 | 660.840935124934 | 1036.4881443660588 |
| SE_103_3_9 | SE_103 | SE_103_3 | AGCTACTGAGAGATAAACTG | 95.37236206857031 | 205.8748518345826 | 214.29332919169605 | 128.82954046707366 | 171.8163110621639 | 118.68448505197199 | 76.40740350628997 | 30.25131169779792 | 355.19411113006794 | 140.69834035159028 | 10.564064682281362 | 276.24208102925195 |
| SE_103_4_0 | SE_103 | SE_103_4 | AGAGTGTTTGTGCTTACACG | 657.8525429047975 | 777.5273530451713 | 816.6847153987679 | 791.1955616563714 | 674.5381100959027 | 656.6630632802538 | 562.8320004759106 | 657.9660294271048 | 605.6059594767657 | 527.0720211098434 | 863.9057340176757 | 459.629681040268 |
| SE_103_4_1 | SE_103 | SE_103_4 | TTTCATGTATGATGTCTAAA | 34.680858934025565 | 37.97691441608806 | 15.800429802152705 | 33.834020728726415 | 22.272484767317543 | 9.529411208552496 | 47.35106696164449 | 6.617474433893295 | 1.7759705556503396 | 2.9160277793075706 | 21.128129364562724 | 1.1606810127279494 |
| SE_103_4_11 | SE_103 | SE_103_4 | TTTCCCACAGAGCCATTTGC | 18.42420630870108 | 40.97509186998975 | 34.56344019220904 | 39.03925468699201 | 14.848323178211695 | 2.598930329605226 | 22.599372868057596 | 25.524544245016994 | 25.751573056929924 | 57.59154864132452 | 16.432989505771005 | 9.285448101823595 |
| SE_103_4_14 | SE_103 | SE_103_4 | TCTAACCTCCCTTGAAGCAA | 539.7208671607729 | 469.7144677779312 | 494.75095817990655 | 504.90769395176335 | 464.54039657548014 | 307.5400890032851 | 430.464245105859 | 624.8786572576383 | 380.9456841869978 | 379.0836113099842 | 571.6332778078914 | 424.8092506584295 |
| SE_103_4_18 | SE_103 | SE_103_4 | CTCTACTGTAGAGGTCAGCG | 82.36703996831072 | 54.96658665486429 | 55.301504307534465 | 97.59813671748003 | 38.181402458258646 | 24.256683076315444 | 109.76838250199404 | 89.80858160283758 | 183.81295250981015 | 99.1449444964574 | 107.98821675220947 | 47.58792152184593 |
| SE_104_1_0 | SE_104 | SE_104_1 | TGGGCACTGTGAGGACAGCG | 441.09717456713764 | 453.7241880237889 | 417.72386289441215 | 321.4231969229009 | 487.8734758555271 | 276.35292504802237 | 299.172650348572 | 346.9447310341199 | 413.8011394665291 | 223.07612511702916 | 285.22974642159676 | 586.1439114276145 |
| SE_104_1_12 | SE_104 | SE_104_1 | CAGAGCCTCAGTTTCCCCAG | 296.9548546225939 | 331.7983048984535 | 341.6842944715522 | 347.4493667142289 | 364.8445123789159 | 456.5454279006514 | 293.79184728474877 | 353.5622054680132 | 193.580790565887 | 325.86610433762104 | 255.88512230414852 | 175.26283292192036 |
| SE_104_1_2 | SE_104 | SE_104_1 | ACGAATGTCGACAACAAAAT | 3116.942196695548 | 3244.028005121627 | 3354.628752369546 | 3314.432722925622 | 3384.357090119537 | 2867.486463664433 | 3337.1740601831716 | 2864.4210763852407 | 3698.458682141832 | 3770.423918644689 | 3285.4241161895034 | 3160.534397658206 |
| SE_104_1_35 | SE_104 | SE_104_1 | ATTTTCCAGGGTGAACAGAA | 2.167553683376598 | 5.9963549078033775 | 4.93763431317272 | 13.013084895664004 | 3.1817835381882205 | 3.465240439473635 | 0.0 | 12.289595377230405 | 0.0 | 6.561062503442034 | 0.0 | 1.1606810127279494 |
| SE_104_1_39 | SE_104 | SE_104_1 | AATCATTGTCAAGATCAGAG | 41.18351998415536 | 46.97144677779312 | 47.40128940645811 | 44.24448864525762 | 30.757240869152795 | 298.0106777947326 | 20.4470516425283 | 25.524544245016994 | 103.0062922277197 | 50.30147919305559 | 15.259204541073078 | 174.10215190919243 |
| SE_104_2_15 | SE_104 | SE_104_2 | TCTGGTTTCCGGCTGAGCCT | 183.15828624532253 | 231.8590564350639 | 204.41806056535063 | 217.31851775758886 | 200.45236290585788 | 211.37966680789174 | 259.35470767628004 | 214.595242356254 | 348.97821418529173 | 293.7897987652377 | 292.2724562097843 | 220.52939241831038 |
| SE_104_2_2 | SE_104 | SE_104_2 | TTACTGCCTGGGCCAGTCTG | 174.48807151181612 | 271.8347558204198 | 293.2954782024596 | 206.90804984105768 | 211.05830803315195 | 314.4705698822324 | 172.1856980423436 | 283.6060471668555 | 289.48320057100534 | 111.53806255851458 | 295.7938111038781 | 355.1683898947525 |
| SE_104_2_4 | SE_104 | SE_104_2 | TGACACGGATGAGAAACGAG | 3286.011383998922 | 3405.929587632318 | 3408.942729814446 | 2953.970271315729 | 3172.2381875736555 | 2629.2511834506204 | 2963.7463275538394 | 2981.6449092142075 | 2998.7262832155984 | 3961.4237381893345 | 2950.8954012505937 | 2331.8081545704504 |
| SE_104_2_5 | SE_104 | SE_104_2 | GACACGGATGAGAAACGAGA | 1717.7862940759537 | 1910.8384306200096 | 1847.662759989232 | 1782.7926307059686 | 1549.5285830976632 | 1912.8127225894464 | 1896.1949996913088 | 1720.5433528122567 | 1707.5956892578015 | 2405.7229179287456 | 1872.1870186931967 | 2162.34872671217 |
| SE_104_2_9 | SE_104 | SE_104_2 | CACAGTTTCACACATCCTGG | 815.0001849496008 | 902.4514136244082 | 779.1586946186552 | 818.5230399372658 | 737.1131863469377 | 931.2833681085394 | 787.7495685437219 | 868.779857821134 | 831.1542200443589 | 604.346757261494 | 1037.6259087929693 | 1081.7547038624489 |
| SE_104_3_10 | SE_104 | SE_104_3 | GAGTAGAAGCATCTTGTGGG | 331.63571355661946 | 444.7296556620838 | 448.337195636083 | 392.9951638490529 | 391.3593751971511 | 362.11762592499485 | 356.2091628250983 | 460.3871499008621 | 196.24474639936253 | 518.3239377719207 | 403.7820278560876 | 412.04175951842205 |
| SE_104_3_12 | SE_104 | SE_104_3 | CACTTTCTATCGGTTCAGAG | 85.61837049337561 | 117.92831318679976 | 92.82752508764715 | 115.81645557140963 | 108.18064029839948 | 19.9251325269734 | 86.0928490211718 | 140.85767009287156 | 191.80482001023668 | 316.38901405487144 | 192.50073421046037 | 32.49906835638259 |
| SE_104_3_21 | SE_104 | SE_104_3 | AAATGAATGAATTCAAGGCG | 21.675536833765978 | 28.982382054382988 | 22.713117840594514 | 31.23140374959361 | 24.393673792776355 | 20.791442636841808 | 19.370891029763655 | 25.524544245016994 | 15.095749723027886 | 5.103048613788249 | 32.86597901154201 | 44.10587848366208 |
| SE_104_3_25 | SE_104 | SE_104_3 | AGCTCCTCTTCCCTTCCCAG | 60.69150313454474 | 73.95504386290831 | 82.9522564613017 | 83.28374333224963 | 99.69588419656424 | 94.42780197565655 | 52.73187002546773 | 184.3439306584561 | 93.23845417164283 | 127.57621534470621 | 98.59793703462604 | 185.70896203647192 |
| SE_104_3_32 | SE_104 | SE_104_3 | TCTCCCGTCCAGCTGCTCCT | 355.47880407376203 | 406.75274124599576 | 399.9483793669903 | 379.9820789533889 | 375.45045750621 | 408.89837185788895 | 379.88469630592056 | 305.34917744964775 | 325.00261168401215 | 247.8623612411435 | 277.01325166871123 | 320.34795951291403 |
| SE_104_4_10 | SE_104 | SE_104_4 | TGTGGATGACAAAACGGACG | 2870.924853632304 | 2754.3256876510177 | 2726.5616677339763 | 2901.917931733073 | 2793.6059465292574 | 2626.6522531210153 | 2813.0838417667883 | 3209.4751004382483 | 2344.281133458448 | 2991.8445015695675 | 2950.8954012505937 | 2471.0898760978043 |
| SE_104_4_28 | SE_104 | SE_104_4 | ACTCCAGAGATTAAACACGG | 39.01596630077876 | 63.961119016569356 | 79.98967587339807 | 36.43663770785921 | 133.63490860390525 | 33.78609428486794 | 54.88419125099702 | 29.305958207241734 | 79.03068972644012 | 110.0800486688608 | 31.692194046844083 | 125.35354937461854 |
| SE_104_4_29 | SE_104 | SE_104_4 | GGAACTCCAGAGATTAAACA | 26.010644200519174 | 56.96537162413208 | 55.301504307534465 | 50.751031093089615 | 27.575457330964575 | 36.38502461447317 | 23.675533480822246 | 44.431614056140695 | 40.84732277995781 | 45.92743752409424 | 63.384388093688166 | 46.42724050911798 |
| SE_104_4_30 | SE_104 | SE_104_4 | GAACACTGCAGAGTGACAGT | 253.60378095506195 | 46.97144677779312 | 87.88989077447442 | 227.72898567412008 | 40.30259148371746 | 698.2459485539374 | 279.80175931880837 | 73.73757226338243 | 277.051406681453 | 11.664111117230282 | 86.86008738764674 | 67.31949873822107 |
| SE_104_4_8 | SE_104 | SE_104_4 | TTTTGTCATCCACAGACCAG | 31.429528408960667 | 32.97995199291857 | 38.51354764274722 | 41.641871666124814 | 54.09032014919974 | 44.18181560328885 | 30.13249715741013 | 31.196665188354107 | 16.871720278678225 | 24.057229179287457 | 32.86597901154201 | 34.820430381838484 |
| SE_105_1_13 | SE_105 | SE_105_1 | CTTCAGCATAGGAATTTGTA | 34.680858934025565 | 61.96233404730156 | 70.11440724705263 | 46.847105624390416 | 72.12042686559965 | 45.91443582302566 | 55.960351863761666 | 31.196665188354107 | 26.639558334755094 | 59.04956253097831 | 8.216494752885502 | 61.51609367458132 |
| SE_105_1_14 | SE_105 | SE_105_1 | GGTGGATGCAGAGTCCCACA | 494.2022398098643 | 497.6974573476803 | 579.6782683664774 | 501.00376848306416 | 539.842606979268 | 710.3742900920952 | 513.3286122887368 | 583.2831036731661 | 466.19227085821416 | 419.17899327546326 | 581.0235575254749 | 805.5126228331969 |
| SE_105_1_15 | SE_105 | SE_105_1 | GTGGATGCAGAGTCCCACAG | 231.92824412129596 | 366.7770418606399 | 341.6842944715522 | 248.5499215071825 | 326.6631099206573 | 370.78072702367893 | 263.65935012733866 | 263.7536238651756 | 300.1390239049074 | 653.1902225648959 | 350.9617044446808 | 444.5408278748046 |
| SE_105_1_24 | SE_105 | SE_105_1 | AAGGCGTTGGTGAAGCCACT | 3028.0724956771073 | 3126.099691934827 | 2831.2395151732376 | 2873.289144962612 | 2995.1189039478445 | 2425.6683076315444 | 2903.481333239019 | 2798.2463320463075 | 2920.5835787669835 | 2719.1959042043095 | 2599.933696805913 | 2944.6477292908075 |
| SE_105_1_8 | SE_105 | SE_105_1 | TCGTCGGATGTCAACACTTA | 8282.22262418198 | 7623.36587278736 | 7892.314686175276 | 7829.973181721031 | 7687.18902826274 | 7196.438082676871 | 7617.064817148174 | 8051.5756790670275 | 7541.6589645691665 | 7316.3136982826945 | 8102.637611309804 | 7607.1033574189805 |
| SE_105_2_11 | SE_105 | SE_105_2 | TGAGGTGATTGTGAACGCCA | 11320.049111434282 | 10337.715861053022 | 10413.470766481267 | 10800.860463401124 | 10413.977520490045 | 9508.619765915655 | 10911.19245282076 | 10418.740819419714 | 9433.06760633678 | 10798.050866775933 | 11092.267916395429 | 9660.348068934723 |
| SE_105_2_12 | SE_105 | SE_105_2 | TGATTGTGAACGCCAAGGAA | 857.2674817754445 | 902.4514136244082 | 948.0257881291623 | 879.6845389468867 | 967.2621956092189 | 585.6256342710443 | 998.6770486455929 | 749.6653180110548 | 761.0033830961705 | 907.6136463094814 | 820.4756903238524 | 728.9076759931522 |
| SE_105_2_14 | SE_105 | SE_105_2 | TGTGAACGCCAAGGAATGGG | 5632.38824625409 | 5492.661095547894 | 5507.437312912852 | 5405.635465658827 | 5453.576984454609 | 5279.293809538083 | 5275.339323772302 | 4228.566163257816 | 5083.715715549097 | 4842.793134485048 | 4019.039719125709 | 3856.943005294976 |
| SE_105_2_24 | SE_105 | SE_105_2 | TGACATCCAGGAAATGATAA | 726.1304839311603 | 824.4987998229643 | 846.3105212778042 | 662.3660211892978 | 761.506860139714 | 648.8662722914381 | 682.2858284927865 | 706.1790574454702 | 552.3268428072556 | 607.2627850408015 | 630.3225260427879 | 745.1572101713435 |
| SE_105_2_25 | SE_105 | SE_105_2 | GACATCCAGGAAATGATAAG | 793.3246481158347 | 1043.3657539577875 | 1059.6163236068658 | 949.9551973834723 | 940.7473327909838 | 694.7807081144638 | 913.6603602371857 | 1009.6375279140055 | 1066.470318668029 | 749.4191392820456 | 1138.5714157569912 | 667.391582318571 |
| SE_105_3_11 | SE_105 | SE_105_3 | GATGAGCTCTGACCACGTGT | 9486.29869529768 | 8264.975847922322 | 8383.115536904645 | 9165.115692016157 | 7967.185979623304 | 8456.919292535405 | 9993.227450132517 | 8293.586172649411 | 8573.497857402015 | 8412.011136357514 | 7829.145714535187 | 9011.527382819799 |
| SE_105_3_15 | SE_105 | SE_105_3 | GCCTCTTCAGGAATCCCCAG | 31.429528408960667 | 62.96172653193546 | 64.18924607124536 | 26.02616979132801 | 54.09032014919974 | 11.262031428289314 | 17.21856980423436 | 18.9070698111237 | 45.28724916908366 | 144.34337507572474 | 9.390279717583432 | 24.37430126728694 |
| SE_105_3_16 | SE_105 | SE_105_3 | TACAGAATCCTCCATCCTGC | 1146.6358985062202 | 1234.2497185228617 | 1295.6352437765217 | 1181.5881085262915 | 1281.1981713771233 | 861.1122492091982 | 993.2962455817697 | 965.2059138578649 | 1379.9291217403138 | 1141.6248755989138 | 1101.0102968866574 | 1192.019400071604 |
| SE_105_3_7 | SE_105 | SE_105_3 | GGGGGCATGCAGGATCAAAA | 463.8564882425919 | 591.6403509032665 | 640.904933849819 | 566.0691929613841 | 635.2961131249147 | 799.6042314085413 | 474.5868302292095 | 718.4686528227006 | 563.8706514189828 | 509.57585443399796 | 926.116337146666 | 710.336779789505 |
| SE_105_3_8 | SE_105 | SE_105_3 | AGAGTATCATAGATGCCAGA | 150.64498099467355 | 220.86573910409106 | 199.4804262521779 | 216.01720926802247 | 259.84565561870465 | 240.83421054341764 | 200.16587397422444 | 208.92312141291688 | 231.7641575123693 | 465.10643079955753 | 160.80854016361627 | 301.77706330926685 |
| SE_105_4_12 | SE_105 | SE_105_4 | TCAGGGGGATGGAGGCAAAG | 6202.4548649821345 | 6137.269248136757 | 6295.483749295218 | 5959.992882214114 | 6159.9329299323945 | 5821.603938315707 | 5947.939706750207 | 5490.613073150323 | 5731.056983083646 | 4736.358120540322 | 5360.675933775442 | 4599.778853440864 |
| SE_105_4_24 | SE_105 | SE_105_4 | TGGACGTCAGAGACACACAG | 528.8830987438898 | 432.73694584647706 | 481.9131089656575 | 512.7155448891617 | 333.02667699703375 | 408.89837185788895 | 447.6828149100933 | 599.3541130126213 | 484.8399616925427 | 352.11035435138916 | 680.7952795247988 | 357.48975192020845 |
| SE_105_4_29 | SE_105 | SE_105_4 | ACCCCAGCTCCTGTTAGTAA | 133.30455152766078 | 179.89064723410132 | 166.89203978523796 | 93.69421124878083 | 153.786204345764 | 212.24597691776015 | 112.99686434028798 | 129.51342820619735 | 121.65398306204825 | 165.48457647570464 | 97.42415206992811 | 164.8167038073688 |
| SE_105_4_30 | SE_105 | SE_105_4 | CAGCTTCTAGGTGCTCGGGA | 231.92824412129596 | 268.8365783665181 | 316.99612290568865 | 249.85122999674888 | 251.3608995168694 | 168.0641613144713 | 200.16587397422444 | 315.7480658457658 | 227.32423112324346 | 267.5455487514696 | 397.9131030325979 | 176.42351393464833 |
| SE_105_4_31 | SE_105 | SE_105_4 | GAGTTGGTGCAGGTGCAGGA | 775.984218648822 | 845.4860420002761 | 761.3832110912334 | 873.1779964990546 | 816.6577748016432 | 906.1603749223556 | 824.33902937772 | 761.9549133882851 | 765.4433094852964 | 975.4112921783824 | 569.2857078784956 | 851.9398633423149 |
| SE_106_1_10 | SE_106 | SE_106_1 | TCCCACCAGTATGTGTGCGA | 416.1703072083068 | 374.7721817377111 | 326.87139153203407 | 396.89908931775216 | 376.5110520189394 | 388.97323933091553 | 485.348436356856 | 397.9938195241539 | 348.97821418529173 | 336.8012085100244 | 318.0957254331388 | 552.4841620585039 |
| SE_106_1_13 | SE_106 | SE_106_1 | TTCAGCTAGCAGCTGCCACG | 49.85373471766175 | 58.96415659339988 | 53.32645058226538 | 42.943180155691216 | 106.05945127294068 | 26.85561340592067 | 39.817942672291956 | 82.2457536783881 | 67.4868811147129 | 116.64111117230283 | 34.03976397623994 | 54.55200759821362 |
| SE_106_1_25 | SE_106 | SE_106_1 | CCAACCAAACTCCATCCCAC | 2816.7360115478887 | 2711.3518108117605 | 2582.3827457893326 | 2730.145211110308 | 2643.0015257216814 | 2481.978464772991 | 2718.3817078434995 | 2370.0012008243557 | 2507.6704245782794 | 2534.028140218279 | 2587.022062194236 | 2060.2087975921104 |
| SE_106_1_36 | SE_106 | SE_106_1 | GCCAACAAATTGCCTCCAGG | 389.0758861660993 | 449.7266180852533 | 437.474400147103 | 452.8553543691074 | 422.1166160663039 | 285.8823362565749 | 468.12986655262165 | 427.29977773139564 | 494.6077997486196 | 483.33160442022984 | 407.30338275018136 | 543.1987139566803 |
| SE_106_1_6 | SE_106 | SE_106_1 | CATGCACAATGGCAAAACAG | 11868.44019332856 | 10235.777827620364 | 10630.726676260867 | 11264.126285686762 | 10512.61281017388 | 10970.951231373529 | 11718.312912394247 | 11763.03348299061 | 10535.94532139564 | 10253.482678990245 | 11214.341552724014 | 11371.19188169572 |
| SE_106_2_17 | SE_106 | SE_106_2 | TCTTCATCTTGGATCCAAGA | 61.77527997623304 | 63.961119016569356 | 75.05204156022535 | 65.06542447832003 | 76.36280491651729 | 6.064170769078861 | 68.87427921693744 | 38.75949311280358 | 39.95933750213264 | 145.80138896537852 | 39.90868879972959 | 54.55200759821362 |
| SE_106_2_21 | SE_106 | SE_106_2 | GCATGGTAATGAGAACCACA | 7195.194451968617 | 6454.076665765701 | 6263.882889690913 | 6717.354423141759 | 6547.049927078628 | 5773.090572163076 | 6707.709099362048 | 6566.425345403261 | 6452.989013955509 | 5904.227246153004 | 7773.977821194384 | 6269.998830756383 |
| SE_106_2_23 | SE_106 | SE_106_2 | AGCTAGACTGTGCAGAACCA | 1803.4046645693293 | 1817.8949295490572 | 1834.8249107749828 | 1902.5130117460774 | 1801.9500771272621 | 1393.0266566684013 | 1761.674923095728 | 1926.630413753505 | 1971.327316771877 | 1677.44498004668 | 2236.060357749555 | 1821.1085089701526 |
| SE_106_2_28 | SE_106 | SE_106_2 | CCCAAGCCAAGAATTCAGCT | 27.094421042207472 | 66.95929647047105 | 95.79010567555078 | 44.24448864525762 | 40.30259148371746 | 89.2299413164461 | 26.904015319116187 | 42.54090707502832 | 85.2465866712163 | 45.92743752409424 | 35.21354894093787 | 228.65415950740604 |
| SE_106_2_29 | SE_106 | SE_106_2 | GGTGGCCAGCAGTCTCTGGG | 197.2473851872704 | 223.86391655799275 | 208.36816801588878 | 171.77272062276487 | 207.87652449496372 | 368.1817966940737 | 209.85131948910626 | 287.38746112908024 | 200.68467278848837 | 484.7896183098836 | 252.36376741005475 | 590.7866354785263 |
| SE_106_3_12 | SE_106 | SE_106_3 | ATTGAAACATGGCTTGGGAA | 154.98008836142674 | 181.8894322033691 | 181.7049427247561 | 143.14393385230406 | 201.5129574185873 | 336.994632738811 | 200.16587397422444 | 213.64988886569782 | 176.7090702872088 | 201.20591677222237 | 196.02208910455414 | 436.41606078570896 |
| SE_106_3_14 | SE_106 | SE_106_3 | TCTGCATTGAAACATGGCTT | 1928.0390013634837 | 1737.9435307783456 | 1581.030507077905 | 1708.6180468006837 | 2062.856327258696 | 1760.3421432526065 | 1811.1783112829016 | 2044.7996000730282 | 1651.6526167548159 | 2022.9942718946272 | 2292.4020360550553 | 1800.2162507410496 |
| SE_106_3_16 | SE_106 | SE_106_3 | CTTCTGTCTGCATTGAAACA | 55.27261892610324 | 89.94532361705066 | 100.72773998872348 | 54.65495656178882 | 88.02934455654076 | 44.18181560328885 | 68.87427921693744 | 101.15282348951179 | 73.70277805948909 | 115.18309728264904 | 154.93961534012664 | 94.01516203096391 |
| SE_106_3_22 | SE_106 | SE_106_3 | TGTAAGGCCCTGGCAGGCTG | 576.569279778175 | 520.6834844942599 | 546.1023550369028 | 573.8770438987826 | 499.54001549555056 | 433.1550549342044 | 568.2128035397338 | 620.1518898048573 | 431.5608450230325 | 667.0413545166068 | 651.4506554073506 | 342.40089875474507 |
| SE_106_3_5 | SE_106 | SE_106_3 | AGAGAAGGGCAGGCTAACCG | 2832.992664173213 | 3079.128245157034 | 3130.4601545515047 | 2933.1493354826666 | 2912.3925319549508 | 3397.668250903899 | 2951.908560813428 | 3209.4751004382483 | 2972.086724880843 | 3998.603092375506 | 3780.7613712920293 | 3511.060063502047 |
| SE_107_1_12 | SE_107 | SE_107_1 | GCATGGATTCAGAAGCCAGG | 1325.4590773847895 | 1597.0291904449662 | 1546.4670668856959 | 1395.0027008151812 | 1729.8296502616624 | 2585.0693678473317 | 1232.2039016155213 | 1694.0734550766836 | 2025.4944187192123 | 1213.7965631367763 | 1639.777595683007 | 2904.0238938453294 |
| SE_107_1_17 | SE_107 | SE_107_1 | ACTCTAGCTGATATAATGAA | 48.769957875973454 | 91.94410858631845 | 73.07698783495626 | 58.55888203048802 | 88.02934455654076 | 84.89839076710406 | 62.417315540349556 | 95.48070254617468 | 34.63142583518162 | 68.5266528137279 | 38.73490383503166 | 98.6578860818757 |
| SE_107_1_18 | SE_107 | SE_107_1 | GCAGGGCAGAGCTACTTGCA | 27576.701736758765 | 23989.417201152042 | 23891.237387717523 | 26636.48347293465 | 25335.481720080068 | 25327.442372112797 | 27331.25108238375 | 25862.035441145552 | 24126.555998509863 | 24044.8360612254 | 24533.279547151415 | 24055.11398878675 |
| SE_107_1_19 | SE_107 | SE_107_1 | ATAGCTAGCCAAAAGTTGGT | 267.69287989700985 | 227.86148649652833 | 249.84429624653964 | 218.6198262471553 | 196.20998485494025 | 191.45453428091832 | 227.06988929334062 | 219.3220098090349 | 276.1634214036278 | 179.3357084274156 | 194.84830413985622 | 132.31763545098624 |
| SE_107_1_20 | SE_107 | SE_107_1 | CCTCAAAACCAATATCAGCA | 2177.3076749517927 | 2331.5826666508797 | 2502.3930699159346 | 2223.9362086689785 | 2530.5785073723646 | 2570.3420959795685 | 2370.7818299205182 | 2734.907648179043 | 2276.7942523437355 | 2593.0777027492572 | 2953.2429711799896 | 2469.9291950850766 |
| SE_107_2_1 | SE_107 | SE_107_2 | AGGCATGCAGACACTTTAAA | 3167.8797082548977 | 3591.816589774223 | 3563.9844472480695 | 3389.908615320473 | 3811.776678749488 | 3724.267162324289 | 3155.302916625946 | 3743.5998226024926 | 3595.4523899141122 | 3323.5426614658036 | 3799.5419307271964 | 4200.504585062449 |
| SE_107_2_14 | SE_107 | SE_107_2 | GCAAAGAGCCAACCACCTAA | 1722.1214014427069 | 1828.88824688003 | 1764.71050352793 | 1661.7709411762933 | 1867.7069369164853 | 1389.5614162289276 | 1831.62536292543 | 2024.001823280792 | 1753.7709237047104 | 2300.745917873673 | 1757.1560921527998 | 1270.9457089371047 |
| SE_107_2_25 | SE_107 | SE_107_2 | CCAGTCCTGCTAGAAATTCA | 235.17957464636086 | 265.8384009126164 | 252.80687683444327 | 296.6983356211393 | 346.814405662516 | 244.29945098289127 | 212.00364071463557 | 397.9938195241539 | 327.66656751748764 | 166.21358342053153 | 178.4153146340852 | 133.47831646371418 |
| SE_107_2_5 | SE_107 | SE_107_2 | GACTCTTTCAGCCAAGCAAG | 41.18351998415536 | 59.96354907803377 | 41.47612823065085 | 66.36673296788642 | 32.87842989461161 | 59.7753975809202 | 54.88419125099702 | 34.97807915057884 | 40.84732277995781 | 87.48083337922712 | 34.03976397623994 | 131.15695443825828 |
| SE_107_2_6 | SE_107 | SE_107_2 | ACTCTTTCAGCCAAGCAAGA | 31.429528408960667 | 39.975699385355846 | 55.301504307534465 | 24.724861301761607 | 39.24199697098805 | 51.11229648223612 | 26.904015319116187 | 3.78141396222474 | 76.3667338929646 | 18.225173620672315 | 0.0 | 15.088853165463343 |
| SE_107_3_1 | SE_107 | SE_107_3 | GATTCCTGTATTTCCAACTG | 4.335107366753196 | 7.9951398770711695 | 15.800429802152705 | 0.0 | 7.4241615891058474 | 39.8502650539468 | 2.152321225529295 | 0.0 | 0.8879852778251698 | 0.7290069448268927 | 0.0 | 17.410215190919242 |
| SE_107_3_13 | SE_107 | SE_107_3 | CCTCTGGGGCAAACTTCCAG | 73.69682523480432 | 94.94228604022014 | 91.8399982250126 | 66.36673296788642 | 86.96875004381135 | 86.63101098684088 | 54.88419125099702 | 128.56807471564116 | 128.75786528464963 | 57.59154864132452 | 139.68041079905356 | 103.3006101327875 |
| SE_107_3_25 | SE_107 | SE_107_3 | CCCTGGGACAGAGCAGCTGA | 592.8259324034995 | 634.6142277425241 | 539.189666998461 | 701.4052758762898 | 546.2061740556445 | 441.81815603288845 | 640.3155645949653 | 669.310271313779 | 536.3431078064026 | 850.0220976681568 | 505.9013197848074 | 416.68448356933385 |
| SE_107_3_6 | SE_107 | SE_107_3 | GAGTTCACCACAGCTCAGCA | 999.2422480366116 | 871.4702466007575 | 910.4997673490496 | 866.6714540512227 | 809.2336132125373 | 1113.2084911809052 | 984.6869606796524 | 970.878034801202 | 1020.2950842211201 | 625.487958661474 | 961.3298860876039 | 1140.9494355115744 |

| | | | | | | | | | | | | | | | |
|---|---|---|---|---|---|---|---|---|---|---|---|---|---|---|---|
| SE_107_3_7 | SE_107 | SE_107_3 | TCACAAATCCTCGCCAGCAA | 2815.6522347062005 | 2845.2704037527024 | 2879.6283314423304 | 3006.022610898385 | 3128.75381255175 | 3439.251136177583 | 2780.799023383849 | 2525.9845267661262 | 3195.859014892786 | 2609.844862480276 | 2758.3946670401333 | 2858.7573343489394 |
| SE_107_4_28 | SE_107 | SE_107_4 | CGCTTTGGTTCCATGCTGGC | 281.78197883895774 | 419.7448435462364 | 467.10020602613935 | 346.1480582246625 | 379.69283555712764 | 329.1978417499953 | 290.56336544645484 | 401.7752334863786 | 416.46509530000463 | 477.4995488616147 | 434.30043693823376 | 421.32720762024564 |
| SE_107_4_29 | SE_107 | SE_107_4 | CAGCCAGCATGGAACCAAAG | 1682.02165830024 | 1654.993954553732 | 1648.182333737054 | 1889.4999268504134 | 1730.8902447743917 | 1573.2191595210302 | 1566.8898521853266 | 1885.034860169033 | 1556.6381920275226 | 1829.0784245706736 | 1463.7098509783175 | 1602.9004785772981 |
| SE_107_4_30 | SE_107 | SE_107_4 | GATCCGCTTTGGTTCCATGC | 419.42163773337165 | 367.7764343452738 | 329.8339721199377 | 361.7637600994593 | 284.239329411481 | 251.22993186183854 | 334.68595056980536 | 342.21796358133895 | 458.2004033577876 | 352.11035435138916 | 346.26656458588906 | 365.61451900930405 |
| SE_107_4_31 | SE_107 | SE_107_4 | GGAACCAAAGCGGATCAGAG | 411.83519984155356 | 455.72297299305666 | 485.86321641619566 | 404.7069402551505 | 556.8121191829385 | 292.81281713552215 | 492.8815606462085 | 333.7097821663333 | 535.4551225285774 | 542.3811669512081 | 458.9499211968903 | 347.04362280565687 |
| SE_107_4_33 | SE_107 | SE_107_4 | TAGCCCTCTCTGATCCGCTT | 473.6104798177866 | 565.6561463027853 | 557.9526773885174 | 494.4972260352322 | 600.2964942048442 | 448.7486369118357 | 459.5205816505045 | 609.7530014087394 | 520.3593728055495 | 718.0718406544893 | 366.22090898575385 | 538.5559899057686 |
| SE_107_5_3 | SE_107 | SE_107_5 | ATATCCCTGACCCAAGAAGA | 115.96412206064798 | 135.9173779102099 | 135.29118018093254 | 71.57196692615203 | 102.87766773475245 | 116.08555472236677 | 101.15909759987686 | 130.45878169675353 | 105.67024806119521 | 48.114458358574915 | 134.98527094026184 | 117.2287822855229 |
| SE_107_5_4 | SE_107 | SE_107_5 | TATCCCTGACCCAAGAAGAG | 208.0851536041534 | 194.88153450360974 | 207.38064115325426 | 221.22244322628808 | 250.30030500414 | 101.35828285460381 | 202.31819519975372 | 298.7317030157545 | 373.8418019643965 | 179.3357084274156 | 338.0500698330036 | 164.8167038073688 |
| SE_107_5_5 | SE_107 | SE_107_5 | TCACTGTGATTCAGTCAGGG | 387.992109324411 | 523.6816619481616 | 600.4163324818028 | 501.00376848306416 | 387.1169971462335 | 393.30478988025754 | 467.053705939857 | 415.9555358447214 | 423.56897752260596 | 617.468882268378 | 645.5817305838609 | 461.95104306572387 |
| SE_107_5_6 | SE_107 | SE_107_5 | CCATTTTGTCTGAAGTGACA | 203.7500462374002 | 225.86270152726053 | 231.0812858564833 | 187.38842249756166 | 216.36128059679896 | 136.8769973592086 | 220.61292561675273 | 86.02716764061283 | 279.7153625149285 | 368.8775140824077 | 220.67157336321065 | 251.86777976196504 |
| SE_107_5_8 | SE_107 | SE_107_5 | TCAACTTATCTTGCTCCCAG | 408.5838693164887 | 459.72054293159226 | 449.32472249871756 | 390.3925468699201 | 425.2983996044921 | 490.33152218551936 | 323.9243444421589 | 437.69866612751366 | 499.9357114155706 | 147.9884097998592 | 396.7393180679 | 441.05878483662076 |
| SE_107_6_11 | SE_107 | SE_107_6 | AATTTTCATGATGCACACCA | 17.340429467012783 | 11.992709815606755 | 11.850322351614528 | 7.807850937398403 | 24.393673792776355 | 0.8663101098684087 | 15.066248578705064 | 7.56282792444948 | 29.303514168230603 | 78.73275004130441 | 0.0 | 0.0 |
| SE_107_6_17 | SE_107 | SE_107_6 | ACAATGTTCTACCTGTAAGG | 314.2952840896067 | 312.8098476904095 | 310.0834348672468 | 327.92973937073293 | 271.5121952587281 | 606.4170769078861 | 345.44755669745183 | 502.9280569758904 | 260.17968640277473 | 191.72882648947277 | 375.6111887033373 | 597.750721554894 |
| SE_107_6_4 | SE_107 | SE_107_6 | CTACAGAGATTAGCAGCTGA | 283.9495325223343 | 370.7746117991755 | 349.5845093726286 | 409.91217421341617 | 376.5110520189394 | 394.171099990126 | 410.01719346333067 | 260.91756339350707 | 364.0739639083196 | 539.4651391719005 | 245.32105762186717 | 403.9169924293264 |
| SE_107_6_5 | SE_107 | SE_107_6 | TGCATAATGTGGCATCGAGG | 3220.9847734976242 | 3075.1306752184987 | 3064.29585475499 | 3262.380383342966 | 3088.4512210680323 | 2961.047955530221 | 3114.40881334089 | 2760.4321924240603 | 3058.2212968298845 | 2718.4668972594827 | 2995.499229909115 | 2888.9350406798662 |
| SE_107_6_6 | SE_107 | SE_107_6 | AATTGGGGATGATAGAGTCA | 69.36171786805113 | 77.9526138014439 | 87.88989077447442 | 74.17458390528482 | 84.84756101835254 | 13.86096175789454 | 62.417315540349556 | 157.8740329228829 | 136.74973278507613 | 88.20984032405401 | 71.60088284657367 | 34.820430381838484 |
| SE_107_7_10 | SE_107 | SE_107_7 | ATAATTCATTTCAAGAAACA | 1.083776841688299 | 7.9951398770711695 | 5.925161175807264 | 2.602616979132801 | 8.484756101835254 | 0.0 | 0.0 | 0.0 | 0.0 | 0.0 | 0.0 | 0.0 |
| SE_107_7_13 | SE_107 | SE_107_7 | TAAGTAGGCGCGATGACACG | 11874.94285437869 | 9977.93456658482 | 9893.044109872862 | 11010.371130221314 | 9850.80183423073 | 10660.812212040637 | 11888.34628921106 | 9808.987818010975 | 9168.447993544878 | 9055.72426863966 | 9209.51683301995 | 8720.196448625084 |
| SE_107_7_14 | SE_107 | SE_107_7 | ATGAATTATTCGGAACGCAG | 109.46146101051819 | 149.90887269508443 | 154.05419057098888 | 136.63739140447206 | 155.90739337122278 | 173.26202197368175 | 119.45382801687587 | 193.79746556401793 | 191.80482001023668 | 399.49580576513716 | 132.63770101086598 | 257.67118482560477 |
| SE_107_7_2 | SE_107 | SE_107_7 | TATACATAAGATACTGGAGG | 534.3019829523314 | 611.6282005959445 | 713.9819216847753 | 517.9207788474274 | 704.2347564523261 | 629.8074498743332 | 598.345300697144 | 596.5180525409528 | 702.3963547597093 | 942.6059796611722 | 728.920463077414 | 711.497460802233 |
| SE_107_7_8 | SE_107 | SE_107_7 | CTGTTGAGAGAAGTCCTGAA | 151.72875783636184 | 134.91798542557598 | 162.94193233469977 | 199.10019890365928 | 173.9375000876227 | 140.34223779868222 | 118.37766740411122 | 81.30040018783191 | 99.45435111641902 | 215.7860556687602 | 100.9455069640219 | 121.87150633643469 |
| SE_108_1_0 | SE_108 | SE_108_1 | GTACTCCTGTCTCTATCAGT | 97.53991575194691 | 134.91798542557598 | 124.42838469195254 | 105.40598765487843 | 151.66501532030517 | 121.28341538157723 | 78.55972473181926 | 106.82494443284891 | 188.252878898936 | 129.7632361791869 | 98.59793703462604 | 46.42724050911798 |
| SE_108_1_16 | SE_108 | SE_108_1 | GCCTCCTTCTCTAGACGTGG | 66.11038734298623 | 131.9198079716743 | 133.31612645566344 | 71.57196692615203 | 138.9378811675523 | 121.28341538157723 | 55.960351863761666 | 63.33868386726439 | 221.99631945629244 | 172.77464592397357 | 70.42709788187574 | 96.3365240564198 |
| SE_108_1_19 | SE_108 | SE_108_1 | ACGTCTAGAGAAGGAGGCGT | 6514.582595388365 | 6898.806321427785 | 6981.814918826226 | 6157.791772628207 | 6157.8117409069355 | 6443.614597201225 | 6746.450881421575 | 6359.392930971457 | 6622.594202020116 | 5727.078558560069 | 6165.892419558221 | 6471.957326971046 |
| SE_108_1_21 | SE_108 | SE_108_1 | GGAACAAGTCAAGCACACAA | 968.8964964693392 | 923.4386558017201 | 865.0735316678606 | 1004.6101539452611 | 833.6272870053137 | 1379.1656949105068 | 954.5544635222423 | 981.27692319732 | 671.3168700358284 | 1034.4608547093608 | 1150.3092654039704 | 697.5692886494976 |
| SE_108_1_34 | SE_108 | SE_108_1 | GATAGAAGACAGAGTGGCTG | 640.5121134377847 | 672.5911421586121 | 642.8799875750882 | 611.6149900962082 | 714.8407015796201 | 708.6416698723583 | 823.2628687649553 | 611.6437083898517 | 575.4144600307101 | 738.4840351096423 | 873.2960137352592 | 554.8055240839599 |
| SE_108_2_26 | SE_108 | SE_108_2 | GGAGAATACGGGGCACAGAA | 512.6264461185654 | 697.5759542744595 | 727.8072977616589 | 506.20900244132974 | 685.1440552231968 | 654.930443060517 | 578.9744096673803 | 619.2065363143012 | 623.3656650332692 | 764.7282851234104 | 581.0235575254749 | 778.816959540454 |
| SE_108_2_28 | SE_108 | SE_108_2 | AGACACATAGGGAGAATACG | 769.4815575986922 | 826.4975847922321 | 884.8240689205514 | 797.7021041042035 | 763.6280491651729 | 514.5882052618348 | 821.110547539426 | 811.1132948972067 | 1012.3032167206935 | 826.6938754336962 | 892.0765731704261 | 701.0513316876815 |
| SE_108_2_4 | SE_108 | SE_108_2 | GCAGCCCTCTGAAGCAACAG | 1279.940450033881 | 1404.146440910624 | 1433.889004545358 | 1281.7888622229045 | 1546.3467995594751 | 1289.9357535940605 | 1311.8397869601054 | 1513.5109383804522 | 1223.643712843084 | 1509.0443757916678 | 1423.801162178588 | 806.6733038459248 |
| SE_108_2_5 | SE_108 | SE_108_2 | GCCCTCTGAAGCAACAGCGG | 201.5824925540236 | 249.84812115847404 | 197.50537252690881 | 132.73346593577284 | 276.81516782237514 | 157.66843999605038 | 164.65257375299106 | 135.18554914953447 | 152.7334677859292 | 45.92743752409424 | 157.2871852695225 | 119.5501443109788 |
| SE_108_3_0 | SE_108 | SE_108_3 | CTCAAGATGGGAGGGAACAA | 134.38832836934907 | 190.88396456507417 | 158.00429802152703 | 94.99551973834723 | 108.18064029839948 | 207.9144263684181 | 105.46374005093546 | 142.74837707398393 | 38.1833669464823 | 39.36637502065221 | 203.06479889274172 | 154.37057469281729 |
| SE_108_3_2 | SE_108 | SE_108_3 | CTCCCAGCCTCTTAAACTTC | 20.59175999207768 | 17.98906472341013 | 24.688171565863602 | 7.807850937398403 | 29.69664635642339 | 47.64705604276248 | 4.30464245105859 | 16.071009339455145 | 31.96747000170611 | 19.683187510326103 | 21.128129364562724 | 84.7297139291403 |
| SE_108_3_3 | SE_108 | SE_108_3 | AACCCGAAGTTTAAGAGGCT | 108.37768416882989 | 152.90705014898612 | 133.31612645566344 | 93.69421124878083 | 142.1196647057405 | 58.04277736118338 | 132.36775537005164 | 124.78666075341643 | 139.41368861855165 | 147.2594028550323 | 131.46391604616804 | 215.88666836739858 |
| SE_108_3_5 | SE_108 | SE_108_3 | ATGTAAACCCGAAGTTTAAG | 523.4642145354484 | 764.5352507449306 | 727.8072977616589 | 668.8725636371298 | 694.6894058377615 | 747.6256248164367 | 487.5007575823853 | 830.9657181988866 | 653.5571644793249 | 863.8732296198677 | 1299.3799559206075 | 413.20244053115 |
| SE_108_3_6 | SE_108 | SE_108_3 | AGAGTACAACTTGTTTTTGA | 651.3498818546676 | 857.4787518158829 | 863.0984779425914 | 801.6060295729027 | 813.475991263455 | 1173.850198871694 | 764.0740350628997 | 784.6433971616335 | 778.7630886526739 | 1093.510417240339 | 739.4845277596953 | 1328.9797595735022 |
| SE_109_1_25 | SE_109 | SE_109_1 | GCAGCTCAACGAGCTGCTGG | 0.0 | 0.9993924846338962 | 0.0 | 0.0 | 0.0 | 0.0 | 0.0 | 0.0 | 0.0 | 0.0 | 0.0 | 0.0 |
| SE_109_1_26 | SE_109 | SE_109_1 | GGTGCAGCTCAACGAGCTGC | 0.0 | 0.9993924846338962 | 0.0 | 0.0 | 0.0 | 0.0 | 0.0 | 0.0 | 0.0 | 0.0 | 0.0 | 0.0 |
| SE_109_1_27 | SE_109 | SE_109_1 | GAGCTGCACCAGCGACAGCC | 2004.987157123353 | 2075.7381905846023 | 2146.883399367499 | 1976.6875956513622 | 1924.9790406038733 | 2131.9891803861537 | 1802.5690263807844 | 1673.2756782844474 | 1775.9705556503395 | 2255.547487294406 | 1903.8792127400409 | 2386.360162168664 |
| SE_109_1_29 | SE_109 | SE_109_1 | GGGAGCGAAGCCCGGTTCGC | 630.75812186259 | 745.5467935368865 | 691.2688038441809 | 710.5144353032547 | 777.4157778306552 | 399.36896064933643 | 643.5440464332592 | 739.2664296149367 | 718.3800897605623 | 842.7320282198879 | 759.4388721595601 | 559.4482481348716 |
| SE_109_1_30 | SE_109 | SE_109_1 | GGCGAACCGGGCTTCGCTCC | 79.11570944324582 | 103.93681840192521 | 132.3285995930289 | 111.91253010271043 | 62.575076251035 | 77.96790988815678 | 118.37766740411122 | 85.08181415005664 | 96.7903952829435 | 129.7632361791869 | 129.1163461167722 | 52.23064557275772 |
| SE_109_2_0 | SE_109 | SE_109_2 | GCTGAGACAAGAGTTCTCGT | 1547.6333299308908 | 1905.84146819684 | 1913.8270597857463 | 1566.775421437946 | 2114.825458382437 | 1833.112192481553 | 1499.091733581154 | 1686.510627152234 | 1552.1982656383968 | 1331.166681253906 | 1392.1089681317437 | 1622.6320557936733 |
| SE_109_2_1 | SE_109 | SE_109_2 | TCTTCCAGCCAGTCTCTAAC | 474.69425665947495 | 560.6591838796157 | 541.1647207237302 | 487.99068358740016 | 591.811738103009 | 745.0266944868315 | 328.2289868932175 | 472.6767452780925 | 838.2581022669602 | 524.1559933305358 | 268.79675691582577 | 506.05692154938595 |
| SE_109_2_20 | SE_109 | SE_109_2 | CTGCACTGTTTGACCCTCAC | 2040.751792899067 | 1648.9975996459286 | 1666.9453441271103 | 2248.66106997074 | 1690.5876532906743 | 1769.8715544611591 | 2162.006671044177 | 1614.663761869964 | 1396.8008420189922 | 1982.169882984321 | 2051.77611829198 | 1850.1255342883514 |
| SE_109_2_3 | SE_109 | SE_109_2 | CCTAACATCCTGTTAGAGAC | 352.22747354869716 | 421.7436285155042 | 423.6490240702194 | 320.1218884333345 | 410.4500764262804 | 434.8876751539412 | 292.7156866719841 | 351.6714984869008 | 472.4081678029903 | 393.663750206522 | 539.9410837610474 | 348.20430381838486 |
| SE_109_2_8 | SE_109 | SE_109_2 | CTTTGTAGAATTTCTTCCAA | 8.670214733506391 | 4.996962423169481 | 9.87526862634544 | 9.109159426964803 | 2.1211890254588135 | 0.0 | 0.0 | 1.89070698111237 | 13.319779167377547 | 0.7290069448268927 | 15.259204541073078 | 6.964086076367696 |
| SE_109_3_14 | SE_109 | SE_109_3 | GGAAACACAGTTCCTTCCCA | 235.17957464636086 | 341.7922297447925 | 300.2081662409014 | 296.6983356211393 | 292.72408551331625 | 293.67912724539053 | 342.2190748591579 | 265.64433084628797 | 310.79484723880944 | 406.0568682685792 | 269.9705418805237 | 313.38387343654637 |
| SE_109_3_18 | SE_109 | SE_109_3 | AACTGTGTTTCCAGCCACCG | 104.0425768020767 | 166.89854493386065 | 154.05419057098888 | 150.95178478970246 | 125.15015250207 | 105.68983340394587 | 159.27177068916782 | 138.96696311175918 | 103.89427750554486 | 158.19450702743572 | 107.98821675220947 | 263.4745898892445 |
| SE_109_3_23 | SE_109 | SE_109_3 | AGAAGAGCTGGGGTGGGCAT | 2102.5270728753 | 2333.5814516201476 | 2314.762966015371 | 2413.927248145673 | 2417.0948945103178 | 2250.6736654381257 | 2200.748453103704 | 2096.7940420536183 | 1871.8729656554578 | 2782.6195084042492 | 2216.10601334969 | 1846.6434912501675 |
| SE_109_3_35 | SE_109 | SE_109_3 | TTGCTCACCCACCAACAGCT | 39.01596630077876 | 22.986027146579612 | 36.53849391747813 | 46.847105624390416 | 38.181402458258646 | 6.93048087894727 | 88.2451702467011 | 4.726767452780925 | 47.95120500255917 | 49.5724722482287 | 7.042709788187574 | 12.767491140007444 |
| SE_109_3_5 | SE_109 | SE_109_3 | CTTATCAGCAGGAGACCAGT | 671.9416418467454 | 576.649463633758 | 587.5784832675537 | 664.9686381684306 | 600.2964942048442 | 519.7860659210453 | 642.4678858204945 | 654.18461546488 | 575.4144600307101 | 343.36227101346645 | 786.4359263476125 | 421.32720762024564 |
| SE_109_4_18 | SE_109 | SE_109_4 | AAGTATTGCTGGAGGGACCA | 215.67159149597148 | 242.85237376603678 | 279.470102125576 | 232.9342196323857 | 239.69435987684594 | 196.65239494012877 | 222.76524684228204 | 231.61160518626534 | 283.26730362622914 | 234.01122928943255 | 217.15021846911688 | 413.20244053115 |
| SE_109_4_19 | SE_109 | SE_109_4 | ATTTAAGCCAGGGAGTGCTG | 114.88034521895969 | 97.94046349412183 | 79.00214901076352 | 81.98243484268323 | 71.05983235287025 | 100.4919727447354 | 80.71204595734856 | 176.7811027340066 | 103.89427750554486 | 96.95792366197672 | 169.0250349165018 | 15.088853165463343 |
| SE_109_4_2 | SE_109 | SE_109_4 | GCTGACCCCCCACCTTCCAA | 807.4137470577826 | 1024.3772967497437 | 988.5143894971786 | 949.9551973834723 | 1021.3525157584187 | 1341.0480500762967 | 844.7860810202483 | 1091.8832815923936 | 1087.781965335833 | 1321.6895909711563 | 1076.360812628001 | 1402.102663375363 |
| SE_109_4_22 | SE_109 | SE_109_4 | GGTTAACTAATGGCACCTCG | 603.6637008203825 | 700.5741317283612 | 682.38106208047 | 654.5581702518995 | 712.7195125541614 | 703.4438092131479 | 697.3520770714915 | 928.3371277261737 | 870.2255722686664 | 944.7930004956529 | 805.2164857827793 | 844.9757772659472 |
| SE_109_4_34 | SE_109 | SE_109_4 | CATTAATGTGAACATGACTG | 1194.3220795405055 | 956.4186077946387 | 987.526862634544 | 1056.6624935279171 | 1012.8677596565834 | 827.3261549243304 | 1206.3760469091699 | 938.7360161222917 | 642.0133558675977 | 1713.1663203431976 | 1410.8895275669106 | 677.8377114331224 |
| SE_11_1_10 | SE_11 | SE_11_1 | GCAGAAAGCCCAATTCCATC | 105.126353643765 | 71.95625889364052 | 84.92731018657079 | 88.48897729051524 | 94.3929116329172 | 62.37432791052543 | 112.99686434028798 | 175.8357492434504 | 60.382998892111544 | 65.61062503442034 | 315.7481555037429 | 62.67677468730927 |
| SE_11_1_15 | SE_11 | SE_11_1 | GCCAACCTGCAGGGGCCAGA | 63.94283365960963 | 31.980559508284678 | 36.53849391747813 | 59.86019052005442 | 13.787728665482287 | 151.60426922697152 | 77.48356411905462 | 7.56282792444948 | 149.18152667462851 | 4.374041668961356 | 4.695139858791716 | 54.55200759821362 |
| SE_11_1_16 | SE_11 | SE_11_1 | AGTGCTGAGCCAACCTGCAG | 196.1636083455821 | 68.95808143973883 | 116.52816979087619 | 187.38842249756166 | 75.30221040378788 | 582.1603938315707 | 325.00050505492356 | 156.9286794323267 | 435.1127861343332 | 20.412194455152996 | 111.50957164630326 | 191.51236710011165 |
| SE_11_1_21 | SE_11 | SE_11_1 | CAGAGTCCCCTCCATGACCA | 1804.4884414110177 | 1721.9532510242032 | 1666.9453441271103 | 1634.4434628953989 | 1858.1615863019206 | 1875.561387865105 | 1736.923229002141 | 2124.2092932797477 | 2084.1014470556734 | 1490.0901952261686 | 1839.3210396816548 | 1983.6038507520657 |
| SE_11_1_23 | SE_11 | SE_11_1 | AGTGGTGCCTTGGTCATGGA | 1231.1704921579076 | 1029.374259172913 | 1027.0279371399258 | 1146.4527793079988 | 1046.8067840639244 | 1240.5560773315613 | 1347.3530871813387 | 1129.697421214641 | 1332.8659020155799 | 1811.5822578948282 | 1289.989676203024 | 1320.8549924844065 |
| SE_11_2_11 | SE_11 | SE_11_2 | CTAAATTATGGTAGATTAGG | 70.44549470973944 | 57.964764108765976 | 69.12688038441809 | 61.16149900962082 | 50.90853661101153 | 85.76470087697247 | 46.27490634887984 | 77.51898622560716 | 61.27098416993672 | 37.17935418617152 | 39.90868879972959 | 257.67118482560477 |
| SE_11_2_19 | SE_11 | SE_11_2 | ATTAAAATGGACCTTCCAAG | 13.005322100259587 | 15.990279754142339 | 25.675698428498144 | 39.03925468699201 | 23.33307928004695 | 0.0 | 15.066248578705064 | 62.393330376708214 | 19.535676112153734 | 9.477090282749604 | 8.216494752885502 | 25.534982280014887 |
| SE_11_2_20 | SE_11 | SE_11_2 | TCTTTTATCACTAGTGGCTG | 16.256652625324485 | 32.97995199291857 | 37.526020780112674 | 42.943180155691216 | 32.87842989461161 | 3.465240439473635 | 17.21856980423436 | 17.01636283001133 | 20.423661389978903 | 45.92743752409424 | 2.347569929395858 | 3.482043038183848 |
| SE_11_2_3 | SE_11 | SE_11_2 | AGAGACATTGAAGGGAGTGA | 1279.940450033881 | 1347.181069286492 | 1414.138467292667 | 1176.382874568026 | 1258.9256866098058 | 1515.1763821598468 | 1279.554968577166 | 1223.2874167797033 | 1104.6536856145112 | 1030.8158199852262 | 1001.2385748873335 | 1176.9305469061408 |
| SE_11_2_7 | SE_11 | SE_11_2 | GTAGTAAGAAGCAAGTGAAG | 1108.7037090471297 | 1248.2412133077364 | 1231.4459977052763 | 1205.0116613384869 | 1304.5312506571704 | 1536.834134906557 | 1290.3165747048124 | 1569.286794323267 | 1070.0222597793295 | 1876.4638759844217 | 1672.6435746945488 | 1564.5980051572758 |
| SE_11_3_0 | SE_11 | SE_11_3 | GATGGAAAGGAGTAAAAACA | 943.9696291105083 | 1119.3195827899638 | 1126.7681502660148 | 1020.225855820058 | 1074.382241394889 | 1440.6737127111637 | 955.630624135007 | 944.4081370656288 | 1214.7638600648322 | 1081.8463061231087 | 917.8998423937805 | 1568.0800481954598 |
| SE_11_3_23 | SE_11 | SE_11_3 | TAGAGGGCAATGTTAACGCA | 12210.913675302063 | 11055.27966502016 | 10893.408821721656 | 10873.733738816842 | 10621.854044985008 | 9091.92460306895 | 11394.388567952088 | 11041.728769696241 | 10475.562322503529 | 9790.563269025168 | 10386.823152611974 | 9548.92269171284 |
| SE_11_3_25 | SE_11 | SE_11_3 | TTACCTGCTCAGAAGCCACC | 503.95623138505897 | 457.72175796232443 | 505.61375366888655 | 571.2744269196498 | 500.60061000828 | 437.4866054835464 | 570.3651247652632 | 593.6819920692842 | 622.477679755444 | 763.9992781785835 | 534.0721589375577 | 1093.3615139897283 |
| SE_11_3_6 | SE_11 | SE_11_3 | AATGCCAGTTGACAATTCTG | 322.9654988231131 | 323.80316502138237 | 333.7840795704759 | 339.6415157768305 | 331.9660824843043 | 689.5828474552534 | 323.9243444421589 | 429.190484712508 | 398.70538974350126 | 262.44250013768135 | 300.48895096266983 | 500.2535164857462 |
| SE_11_3_7 | SE_11 | SE_11_3 | CCTACCACAGAATTGTCAAC | 2128.5377170758193 | 2287.6093973269885 | 2225.8855483782622 | 2065.1765729418776 | 2304.671876161001 | 1895.4865203920783 | 2043.6290036400655 | 2184.7119166753437 | 2459.71921957572 | 1838.5555148534233 | 2625.7569660292675 | 2236.6323115267587 |
| SE_11_4_12 | SE_11 | SE_11_4 | AGTCTTCTGAAAGGAGGTGG | 53.105065242726646 | 39.975699385355846 | 57.27655803280356 | 61.16149900962082 | 49.84794209828212 | 18.192512307236584 | 76.40740350628997 | 66.17474433893295 | 80.80666028209045 | 58.320555586151414 | 50.47275348201095 | 20.89225822910309 |
| SE_11_4_16 | SE_11 | SE_11_4 | GATAAACTCAAAGTGAAGGG | 134.38832836934907 | 150.9082651797183 | 172.81720096104522 | 174.37533760189766 | 155.90739337122278 | 120.41710527170882 | 128.06311291899306 | 295.8956425440859 | 172.26914389808294 | 250.77838902045107 | 228.88806811609615 | 189.19100507465575 |
| SE_11_4_17 | SE_11 | SE_11_4 | GCCTTTGTATTTCCTTTCAG | 20.59175999207768 | 59.96354907803377 | 33.5759133295745 | 40.34056317655841 | 60.45388722557618 | 44.18181560328885 | 15.066248578705064 | 55.775855942814914 | 75.47874861513944 | 3.6450347241344634 | 32.86597901154201 | 1.1606810127279494 |
| SE_11_4_2 | SE_11 | SE_11_4 | AATCTTATATACGTATCTGA | 2.167553683376598 | 1.9987849692677924 | 4.93763431317272 | 5.205233958265602 | 0.0 | 27.72192351578908 | 0.0 | 0.0 | 0.0 | 0.0 | 0.0 | 10.446129114551544 |
| SE_11_4_3 | SE_11 | SE_11_4 | CTAGATTGTGAATTCCTAGA | 166.90163361999802 | 272.83414830505365 | 298.2331125156323 | 274.5760912985105 | 328.7842989461161 | 459.1443582302566 | 150.66248578705066 | 279.82463320463074 | 265.50759806972576 | 442.50721550992387 | 369.74226387984766 | 485.16466332028284 |
| SE_11_5_12 | SE_11 | SE_11_5 | ACACTGTGTTGATAGGGCTA | 1674.4352204084219 | 1499.0887269508444 | 1692.6210425556085 | 1492.6008375326612 | 1460.4386440283931 | 2011.5720751144452 | 1484.025485002449 | 1338.620542627558 | 1610.805293974858 | 1052.686028330033 | 1253.6023422973883 | 1260.4995798225532 |
| SE_11_5_2 | SE_11 | SE_11_5 | GGATTTGTAGGGCGACCTGC | 207.0013767624651 | 247.84933618920624 | 257.744511147616 | 234.2355281219521 | 293.7846800260457 | 222.64169823618104 | 196.93739213593048 | 241.06514009182717 | 207.78855501108973 | 297.4348334893722 | 147.89690555193906 | 237.93960760922963 |
| SE_11_5_4 | SE_11 | SE_11_5 | GGTCGCCCTACAAATCCCAA | 300.2061851476588 | 397.7582088842907 | 386.12300329010674 | 307.1088035376705 | 306.51181417879855 | 352.58821471644234 | 266.8878319656326 | 403.665940467491 | 262.8436422362503 | 572.2704516891107 | 368.56847891514974 | 395.79222534023074 |
| SE_11_5_9 | SE_11 | SE_11_5 | TTACCCCTAAGCTAAGCAAA | 55.27261892610324 | 47.97083926242702 | 94.80257881291622 | 79.37981786355043 | 53.02972563647034 | 192.32084439078673 | 72.10276105523138 | 99.26211650839943 | 3.551941111300679 | 19.683187510326103 | 21.128129364562724 | 15.088853165463343 |
| SE_110_1_1 | SE_110 | SE_110_1 | ATCTTCTCTTGATTTTACTG | 6.5026610501297935 | 12.992102300240651 | 9.87526862634544 | 19.519627343496005 | 9.545350614564661 | 0.0 | 7.533124289352532 | 7.56282792444948 | 8.879852778251697 | 5.103048613788249 | 2.347569929395858 | 1.1606810127279494 |
| SE_110_1_2 | SE_110 | SE_110_1 | ACTGAGCTGTGCTAACTTGG | 3775.8785164420333 | 3545.8445354810638 | 3564.9719741107037 | 3592.9127396928316 | 3686.626526247418 | 3465.240439473635 | 3906.4630243356705 | 4134.030814202197 | 3572.364772690658 | 2768.0393695077114 | 3656.340165034049 | 3206.9616381673245 |

| | | | | | | | | | | | | | | | |
|---|---|---|---|---|---|---|---|---|---|---|---|---|---|---|---|
| SE_110_1_4 | SE_110 | SE_110_1 | TCAAGGCTTCTCCTTCAAGA | 352.22747354869716 | 449.7266180852533 | 445.37461504817935 | 386.48862140122094 | 362.7233233534571 | 319.66843054144283 | 321.7720232166296 | 415.9555358447214 | 498.15974085992025 | 587.5795975304754 | 225.36671322200237 | 537.3953088930406 |
| SE_110_1_7 | SE_110 | SE_110_1 | GCTCTATTCTTTTCTCCCAG | 28.17819788389577 | 65.95990398583714 | 90.85247136237805 | 57.25757354092162 | 48.78734758555271 | 134.27806702960336 | 40.8941032850566 | 15.12565584889896 | 79.03068972644012 | 94.77090282749604 | 52.820323411406804 | 52.23064557275772 |
| SE_110_1_8 | SE_110 | SE_110_1 | CTGAAAAGCTGGAATGTTGG | 606.9150313454473 | 670.5923571893443 | 605.3539667949755 | 571.2744269196498 | 714.8407015796201 | 1014.4491386559066 | 576.8220884418511 | 542.6329035792502 | 667.7649289245277 | 564.2513752960149 | 331.007360044816 | 561.7696101603275 |
| SE_110_2_15 | SE_110 | SE_110_2 | AGACATCCTTTGGGACAGAA | 200.4987157123353 | 257.8432610355452 | 237.00644703229057 | 187.38842249756166 | 217.42187510952837 | 448.7486369118357 | 140.9770402721688 | 198.52423301679886 | 207.78855501108973 | 286.4997293169688 | 321.61708032723254 | 342.40089875474507 |
| SE_110_2_22 | SE_110 | SE_110_2 | GTAAACATAGAGAGGAAGAA | 60.69150313454474 | 113.93074324826416 | 112.57806234033802 | 58.55888203048802 | 101.81707322202305 | 102.22459296447224 | 49.50338818717378 | 111.55171188562983 | 175.8210850093836 | 111.53806255851458 | 42.25625872912545 | 301.77706330926685 |
| SE_110_2_23 | SE_110 | SE_110_2 | GGCACTCAGTAAACATAGAG | 172.32051782843953 | 201.87728189604704 | 237.99397389492512 | 189.99103947669445 | 244.99733244049295 | 226.97324878552308 | 201.24203458698906 | 165.43686084733238 | 180.26101139850945 | 107.8930278343801 | 109.1620017169074 | 217.04734938012655 |
| SE_110_2_6 | SE_110 | SE_110_2 | TGGCTGCTATTTTCTCTGTG | 30.34575156727237 | 44.97266180852533 | 38.51354764274722 | 33.834020728726415 | 68.93864332741144 | 12.128341538157722 | 19.370891029763655 | 49.15838150892162 | 25.751573056929924 | 21.87020834480678 | 39.90868879972959 | 13.928172152735392 |
| SE_110_2_9 | SE_110 | SE_110_2 | GAACATGCAGATAATACAAG | 1076.1904037964807 | 1188.2776642297026 | 1135.6558920297257 | 1171.1776406097604 | 1176.199314616912 | 1203.3047426072196 | 1103.0646280837636 | 1297.970342533642 | 1253.8352122891397 | 569.3544239098031 | 1151.4830503686683 | 1455.4939899608487 |
| SE_111_1_1 | SE_111 | SE_111_1 | GAAAAAATGGAACAGAGAGG | 76.94815575986922 | 98.93985597875572 | 99.74021312608895 | 118.41907255054244 | 91.21112809472898 | 66.70587845986748 | 81.7882065701132 | 64.28403735782058 | 47.95120500255917 | 24.78623612411435 | 48.12518355261509 | 80.08698987822851 |
| SE_111_1_10 | SE_111 | SE_111_1 | TCAGGTATCAGCAGCAGGGA | 3261.0845166400914 | 3056.1422180104546 | 3121.572412787794 | 3313.1314144360554 | 2995.1189039478445 | 2987.9035689361417 | 3285.5183507704687 | 3200.9669190232426 | 3218.0586468384154 | 2521.6350221562216 | 3246.6892123544717 | 2745.0105951016003 |
| SE_111_1_14 | SE_111 | SE_111_1 | GGTCTGAGTCCCTAAAGACA | 108.37768416882989 | 125.92345306387092 | 142.20386821937433 | 114.51514708184324 | 138.9378811675523 | 187.12298373157628 | 89.32133085946575 | 83.19110716894428 | 155.39742361940472 | 56.13353475167074 | 65.73195802308402 | 76.60494684004466 |
| SE_111_1_7 | SE_111 | SE_111_1 | GGTCTCTTCTTGTCGAAGAG | 0.0 | 0.0 | 0.987526862634544 | 0.0 | 0.0 | 0.0 | 0.0 | 1.89070698111237 | 0.0 | 0.0 | 0.0 | 0.0 |
| SE_111_1_9 | SE_111 | SE_111_1 | CTTCGACAAGAAGAGACCTA | 304.541292514412 | 306.81349278260615 | 303.170746828805 | 329.2310478602993 | 397.7229422735275 | 424.4919538355203 | 341.1429142463933 | 394.21240556192913 | 214.89243723369108 | 309.09894460660246 | 403.7820278560876 | 269.2779949528843 |
| SE_111_2_10 | SE_111 | SE_111_2 | GAGGTGAACCTGCTTCTTAG | 1913.9499024215359 | 2087.730900400209 | 2104.4197442742134 | 1968.8797447139639 | 2066.038110796884 | 2246.3421148887837 | 1937.0891029763654 | 1725.2701202650376 | 2148.924372336911 | 1863.3417509775377 | 1778.2842215173625 | 1819.9478279574248 |
| SE_111_2_12 | SE_111 | SE_111_2 | TTTAAAGCCCGGCCTTTCCT | 2.167553683376598 | 1.9987849692677924 | 9.87526862634544 | 3.9039254686992013 | 14.848323178211695 | 0.0 | 7.533124289352532 | 0.0 | 0.8879852778251698 | 27.702263903421922 | 0.0 | 0.0 |
| SE_111_2_15 | SE_111 | SE_111_2 | AGCCCGGCCTTTCCTAGGCG | 169.06918730337463 | 227.86148649652833 | 201.45547997744697 | 126.22692348794084 | 233.33079280046948 | 215.71121735723378 | 105.46374005093546 | 145.5844375456525 | 322.3386558505366 | 120.28614589643729 | 34.03976397623994 | 227.4934784946781 |
| SE_111_2_3 | SE_111 | SE_111_2 | GCTGGGGTCTGAGCAATGGC | 1752.4671530099793 | 1762.9283428941928 | 1798.2864168575047 | 1748.9586099772423 | 1881.4946655819676 | 1691.037334463134 | 1363.4954963728082 | 2015.4936418657865 | 1542.43042758232 | 1361.0559659918085 | 2322.9204451372016 | 1764.235139346483 |
| SE_111_2_33 | SE_111 | SE_111_2 | AGAGAATCTGTGTGTCCTAG | 436.7620672003845 | 544.6689041254734 | 562.89031170169 | 442.44488645257616 | 574.8422258993385 | 317.935810321706 | 461.6729028760338 | 552.086438484812 | 804.5146617096038 | 337.5302154548513 | 674.9263547013092 | 468.9151291420916 |
| SE_111_3_0 | SE_111 | SE_111_3 | TGACGCTTAAAACAACAAAA | 1519.455132046995 | 1956.8104849131687 | 2138.9831844664222 | 1808.8188004972967 | 2114.825458382437 | 2269.732487855231 | 1602.4031524065601 | 2137.444242147534 | 2539.6378945799856 | 2562.4594110665275 | 2052.9499032566778 | 1963.8722735356905 |
| SE_111_3_23 | SE_111 | SE_111_3 | TTTAACCCAGCAGCCCCTAG | 9.75399157519469 | 30.98116702365078 | 27.650752153767232 | 10.410467916531204 | 12.727134152752882 | 3.465240439473635 | 8.60928490211718 | 30.25131169779792 | 8.879852778251697 | 116.64111117230283 | 39.90868879972959 | 1.1606810127279494 |
| SE_111_3_25 | SE_111 | SE_111_3 | GCTTTAACCCAGCAGCCCCT | 263.3577725302566 | 332.79769738308744 | 327.8589183946686 | 297.9996441107057 | 329.8448934588455 | 366.44917647433687 | 263.65935012733866 | 352.61685197745703 | 283.26730362622914 | 529.259041944324 | 392.0441782091083 | 345.88294179292893 |
| SE_111_3_26 | SE_111 | SE_111_3 | ATGCTCGGCCCACAGAGCAT | 1896.6094729545232 | 1610.0212927452067 | 1501.040831204507 | 1778.8887052372695 | 1447.71150987564 | 1321.9892276591918 | 1953.2315121678353 | 1448.2815475320754 | 1250.283271177839 | 1842.2005495775577 | 1252.4285573326904 | 1669.0592963027914 |
| SE_111_3_6 | SE_111 | SE_111_3 | AGAGTCAGAGCTGAAAGTGA | 844.2621596751849 | 895.455666231971 | 969.7513791071223 | 879.6845389468867 | 1006.504192580207 | 823.8609144848567 | 1024.5049033519444 | 818.6761228216562 | 647.3412675345488 | 1008.9456116404194 | 1009.455069640219 | 791.5844506804615 |
| SE_111_4_10 | SE_111 | SE_111_4 | GGGCCCACATGTGCACTCGG | 371.7354566990865 | 345.78979968332806 | 388.0980570153758 | 361.7637600994593 | 441.2073172954332 | 177.5935725230238 | 363.74228711445085 | 426.3544242408394 | 269.9475244588516 | 270.46157653077717 | 481.2518355261509 | 479.3612582566431 |
| SE_111_4_17 | SE_111 | SE_111_4 | TTTCTTTCTCCTCTCTCAGG | 0.0 | 0.0 | 1.975053725269088 | 0.0 | 0.0 | 0.0 | 0.0 | 0.0 | 0.0 | 0.0 | 0.0 | 0.0 |
| SE_111_4_18 | SE_111 | SE_111_4 | TTAAGGGATCCTCCTGAGAG | 46.60240419259685 | 53.96719417023039 | 62.21419234597627 | 45.545797134824014 | 113.48361286204653 | 84.89839076710406 | 48.42722757440914 | 132.3494886778659 | 39.95933750213264 | 8.019076393095819 | 126.76877618737633 | 26.695663292742836 |
| SE_111_4_30 | SE_111 | SE_111_4 | AAGAAAGCTCTCAAGAGGAG | 162.56652625324483 | 192.88274953434197 | 183.6799964500252 | 131.43215744620645 | 274.6939787969163 | 55.44384703157816 | 156.0432888508739 | 284.5514006574117 | 175.8210850093836 | 21.87020834480678 | 181.936669528179 | 167.1380658328247 |
| SE_111_4_5 | SE_111 | SE_111_4 | TGTGCACTCGGAGGATGGGG | 1964.8874139808859 | 2281.613042419185 | 2097.5070562357714 | 1941.5522664330695 | 2382.0952755902476 | 2237.6790137901 | 1873.5956268232512 | 1792.3902180945267 | 2221.739165118575 | 2107.5590774945467 | 1532.9631638954954 | 2403.770377359583 |
| SE_111_5_12 | SE_111 | SE_111_5 | GCTCTGGGCTTTCTTAGGAG | 29.26197472558407 | 74.95443634754221 | 85.91483704920533 | 37.737946197425615 | 28.63605184369398 | 6.064170769078861 | 10.761606127646475 | 2.836060471668555 | 58.607028336461205 | 8.748083337922711 | 5.868924823489645 | 5.803405063639747 |
| SE_111_5_16 | SE_111 | SE_111_5 | TTTTCATTATGAAAGAGCTG | 3.2513305250648967 | 0.9993924846338962 | 0.0 | 2.602616979132801 | 0.0 | 0.0 | 7.533124289352532 | 15.12565584889896 | 3.551941111300679 | 0.7290069448268927 | 1.173784964697929 | 0.0 |
| SE_111_5_17 | SE_111 | SE_111_5 | CTAAAACTTCCCTCTCCCAG | 337.054597765061 | 461.71932790086004 | 449.32472249871756 | 365.6676855681585 | 553.6303356447503 | 366.44917647433687 | 351.90452037403975 | 608.8076479181832 | 459.0883886356128 | 529.988048889151 | 566.9381379490998 | 524.6278177530331 |
| SE_111_5_4 | SE_111 | SE_111_5 | GATGGAAAGCAAAGGACTAG | 2146.96192338452 | 2248.6330904262663 | 2044.180605653506 | 2169.2812521071896 | 2075.583461411449 | 2406.6094852144392 | 2227.65246842282 | 2422.9409962955024 | 2510.334380411755 | 2147.654459460026 | 2387.478618195588 | 2591.800701421511 |
| SE_111_5_9 | SE_111 | SE_111_5 | GAGTCCATATGATTCTGCTG | 254.68755779675024 | 252.84629861237573 | 302.1832199661705 | 284.9865592150417 | 257.72446659324584 | 223.50800834604945 | 270.1163138039265 | 363.9610938641312 | 288.59521529318016 | 192.45783343429966 | 411.9985226089731 | 264.6352709019725 |
| SE_111_6_13 | SE_111 | SE_111_6 | GCCAGGGCAGGCCTACCTGG | 1428.4178773451779 | 1390.1549461257496 | 1364.76212416094 | 1307.8150320142324 | 1481.6505342829812 | 1670.245891826292 | 1288.164253479283 | 1081.4843931962757 | 1496.255193135411 | 1699.3151883914868 | 1145.6141255451787 | 1781.6453545374025 |
| SE_111_6_2 | SE_111 | SE_111_6 | CTAGCACAGGGCCTGCCACA | 1170.4789890233628 | 997.3936996646283 | 1053.6911624310585 | 1130.837077433202 | 970.4439791474072 | 1123.6042124993262 | 1100.9123068582344 | 1343.347310080339 | 1056.702480611952 | 801.9076393095819 | 1032.9307689341776 | 908.8132329659844 |
| SE_111_6_20 | SE_111 | SE_111_6 | GGGGCTGGTGAGAAAATGAG | 236.26335148804915 | 271.8347558204198 | 253.79440369707783 | 249.85122999674888 | 246.05792695322236 | 511.98927493222953 | 238.90765603375175 | 301.567763487423 | 295.69909751578155 | 257.3394515238931 | 272.3181118099195 | 432.93401774752516 |
| SE_111_6_3 | SE_111 | SE_111_6 | TAAACTGAGTCACAGTGAGG | 441.09717456713764 | 506.6919897093854 | 470.062786614043 | 455.45797134824016 | 466.66158560093896 | 393.30478988025754 | 408.94103285056605 | 470.7860382969801 | 617.149768088493 | 379.0836113099842 | 487.12076034964053 | 393.4708633147749 |
| SE_111_6_4 | SE_111 | SE_111_6 | AATATTGAGTATCTGCCATG | 37.93218945909046 | 40.97509186998975 | 24.688171565863602 | 52.05233958265602 | 28.63605184369398 | 37.251334724341575 | 36.58946083399801 | 34.03272566002266 | 29.303514168230603 | 59.7785694758052 | 15.259204541073078 | 73.12290380186082 |
| SE_112_1_1 | SE_112 | SE_112_1 | ACACCTCCTTGTTTAAGCAA | 364.1490188072684 | 368.7758268299077 | 376.24773466376126 | 314.9166544750689 | 395.6017532480687 | 674.8555755874904 | 368.04692956550946 | 271.3164517896251 | 381.833669464823 | 387.10268770308 | 264.10161705703405 | 1084.0760658879049 |
| SE_112_1_10 | SE_112 | SE_112_1 | TTCACAGTACTATTGCCTGC | 107.2939073271416 | 95.94167852485404 | 104.67784743926167 | 109.30991312357764 | 81.66577748016432 | 49.379676262499295 | 105.46374005093546 | 101.15282348951179 | 151.84548250810403 | 29.160277793075707 | 37.56111887033373 | 77.76562785277261 |
| SE_112_1_2 | SE_112 | SE_112_1 | ACACCATTGCTTAAACAAGG | 635.0932292293431 | 867.4726766622219 | 865.0735316678606 | 603.8071391588098 | 797.5670735725139 | 977.1978039315651 | 564.98432170144 | 752.5013784827232 | 934.1605122720786 | 543.110173896035 | 619.7584613605065 | 964.525921576926 |
| SE_112_1_4 | SE_112 | SE_112_1 | TCTATCTTGTCAAGGCTGGA | 218.92292202103638 | 266.83779339725027 | 213.30580232906152 | 212.11328379932328 | 198.33117388039906 | 300.60960812433785 | 163.5764131402264 | 175.8357492434504 | 172.26914389808294 | 177.8776945377618 | 98.59793703462604 | 270.4386759656122 |
| SE_112_1_7 | SE_112 | SE_112_1 | GGTCCATTAAGATAAGGAAA | 28.17819788389577 | 59.96354907803377 | 42.463655093285396 | 23.423552812195208 | 51.96913112374093 | 48.51336615263089 | 29.056336544645482 | 21.743130282792254 | 43.51127861343332 | 45.198430579267345 | 30.518409082146157 | 52.23064557275772 |
| SE_112_2_1 | SE_112 | SE_112_2 | AGATAACTCACTGAACCACA | 2169.7212370599746 | 2389.547430759646 | 2417.465759729364 | 2195.3074218985175 | 2311.0354432373774 | 2946.320683662458 | 2253.4803231291717 | 2173.3676747886693 | 2518.3262479121813 | 2131.616306673834 | 2220.8011532084815 | 3449.543969827466 |
| SE_112_2_15 | SE_112 | SE_112_2 | TCAAATGGTTGGAGACAGGC | 775.984218648822 | 1011.3851944495029 | 1034.9281520410022 | 1007.2127709243939 | 1010.7465706311247 | 1043.037372281564 | 769.454838126723 | 804.4958204633134 | 1156.1568317283711 | 1227.6476950884871 | 645.5817305838609 | 1000.5070329714924 |
| SE_112_2_2 | SE_112 | SE_112_2 | GATAACTCACTGAACCACAG | 210.25270728753 | 284.8268581206604 | 305.1458005540741 | 291.4931016628737 | 299.0876525896927 | 333.52939229933736 | 190.4804284593426 | 166.38221433788857 | 207.78855501108973 | 366.690493247927 | 167.85124995180385 | 351.68634685656866 |
| SE_112_2_3 | SE_112 | SE_112_2 | GTACTAGTGGGTAGTGGCCA | 886.5294565010286 | 933.432580648059 | 927.2877240138369 | 880.9858474364531 | 946.0503053546308 | 804.8020920677517 | 815.7297444756027 | 866.8891508400217 | 729.9238983722896 | 1102.9875075230887 | 1007.1074997108232 | 875.1534835968739 |
| SE_112_2_7 | SE_112 | SE_112_2 | ACTGGGTTAAAGTAAGTCAC | 710.9576081475241 | 853.4811818773474 | 917.4124553874914 | 750.854998479813 | 675.5987046086321 | 1245.7539379907716 | 729.6368954544309 | 578.5563362203852 | 725.4839719831638 | 769.1023267923717 | 807.5640557121752 | 992.3822658823967 |
| SE_112_3_14 | SE_112 | SE_112_3 | ACACCACATGAGAGGACACG | 3718.4383438325535 | 3380.9447755164706 | 3217.3625184633443 | 3483.602826569254 | 3624.051449996383 | 3752.855395949947 | 3679.3931350423295 | 3362.6223659083503 | 2986.294489326046 | 3046.5200224315845 | 3210.301878448836 | 2866.882101438035 |
| SE_112_3_2 | SE_112 | SE_112_3 | GGACACTCAGATAAGCAACT | 747.8060207649263 | 864.4744992083203 | 897.6619181348005 | 879.6845389468867 | 779.536966856114 | 740.6951439374894 | 818.9582263138967 | 599.3541130126213 | 840.9220581004358 | 966.6632088404597 | 511.77024460829705 | 484.0039823075549 |
| SE_112_3_21 | SE_112 | SE_112_3 | TCTTAAGGCTGGGTACATGG | 785.7382102240167 | 868.4720691468558 | 907.537186761146 | 834.1387418120627 | 776.3551833179257 | 647.1336520717014 | 587.5836945694975 | 749.6653180110548 | 811.6185439322052 | 540.1941461167274 | 804.0427008180814 | 798.5485367568292 |
| SE_112_4_13 | SE_112 | SE_112_4 | CAGCCAAAATGAGAGACAGG | 1912.8661255798477 | 2012.7764640526668 | 1948.3904999779554 | 1863.4737570590855 | 2136.0373486370254 | 2351.165638182861 | 1938.16526358913 | 2372.8372612960243 | 2128.500710946932 | 2012.7881746670507 | 2392.1737580543795 | 1962.7115925229625 |
| SE_112_4_19 | SE_112 | SE_112_4 | TTATAAAAGACTTCAAACAA | 4.335107366753196 | 10.993317330972857 | 10.862795488979984 | 0.0 | 8.484756101835254 | 0.0 | 6.456963676587884 | 0.0 | 9.767838056076867 | 21.141201399979888 | 1.173784964697929 | 2.321362025455899 |
| SE_112_4_20 | SE_112 | SE_112_4 | ACTCCGCACCAGATACAGAG | 373.90301038246315 | 407.7521337306296 | 378.22278838903037 | 389.09123838035373 | 347.8750001752454 | 418.4277830664414 | 313.1627383145124 | 233.5023121673777 | 300.1390239049074 | 156.00748619295504 | 316.9219404684408 | 350.5256658438407 |
| SE_112_4_21 | SE_112 | SE_112_4 | AAGGTTCTTTCACTGCTGAG | 124.63433679415438 | 174.89368481093183 | 165.9045129226034 | 110.61122161314404 | 190.90701229129323 | 84.03208065723565 | 147.4340039487567 | 137.07625613064684 | 214.00445195586593 | 123.93118062057175 | 248.84241251596094 | 263.4745898892445 |
| SE_112_4_4 | SE_112 | SE_112_4 | GGAGCAAGATAAAAGAGCCC | 244.93356622155557 | 362.77947192210434 | 380.1978421142995 | 232.9342196323857 | 433.78315570632736 | 282.41709581710126 | 246.44078032310426 | 501.037349994778 | 426.2329333560815 | 319.305041834179 | 523.5080942552763 | 145.08512659099367 |
| SE_112_5_14 | SE_112 | SE_112_5 | GATGTTTAAGAAAGACCAAA | 58.52394945116814 | 94.94228604022014 | 68.13935352178353 | 70.27065843658562 | 40.30259148371746 | 136.01068724934018 | 39.817942672291956 | 39.70484660335977 | 37.29538166865713 | 97.68693060680361 | 38.73490383503166 | 92.85448101823596 |
| SE_112_5_15 | SE_112 | SE_112_5 | ACAGAGTATGACATATTAGG | 1516.2038015219302 | 1599.0279754142339 | 1834.8249107749828 | 1673.482717582391 | 1807.2530496909092 | 1880.7592485243154 | 1585.1845826023257 | 1971.0620278096458 | 1533.5505748040682 | 1730.662487019043 | 1379.1973335200666 | 1396.2992583117232 |
| SE_112_5_2 | SE_112 | SE_112_5 | TATTCCACATCAAATCCACC | 458.4376040341504 | 492.7004949245108 | 429.5741852460267 | 420.3226421299473 | 426.3589941172215 | 372.51334724341575 | 537.004145769559 | 449.98826150474406 | 386.27359585394885 | 185.89677093085763 | 417.8674474324627 | 316.86591647473017 |
| SE_112_5_3 | SE_112 | SE_112_5 | ACTGTCTACTGTGCAACAAA | 589.5746018784346 | 724.5595513595747 | 690.2812769815463 | 705.309201344989 | 715.9012960923495 | 665.3261643789378 | 613.4115492758491 | 895.2497555567072 | 798.2987647648276 | 408.97289604788676 | 665.5360749837257 | 973.8113696787495 |
| SE_113_1_1 | SE_113 | SE_113_1 | TATGGCTTGAGGGAGAGGTT | 10288.293558147021 | 10508.611975925418 | 10846.995059177832 | 9696.04955575925 | 10155.19245938407 | 9537.207999541311 | 10016.90298361334 | 9817.49599942598 | 10503.089866116108 | 7949.820733337265 | 7739.938057218144 | 8672.608527103239 |
| SE_113_1_13 | SE_113 | SE_113_1 | GAAGAGAGAAGAGTCTTGGG | 5597.707387320064 | 7949.16782277801 | 7584.206305033298 | 6493.529362936338 | 7927.943982652316 | 7402.619888825552 | 5474.429037133762 | 6551.299689554362 | 7737.015725690704 | 6353.295524166369 | 5412.322472222151 | 6207.322056069073 |
| SE_113_1_14 | SE_113 | SE_113_1 | AAGTGAGTAACTCTTACAGC | 196.1636083455821 | 250.84751364310793 | 223.18107095540694 | 251.15253848631528 | 171.8163110621639 | 217.4438375769706 | 227.06988929334062 | 214.595242356254 | 214.89243723369108 | 303.2668890479873 | 392.0441782091083 | 384.1854152129513 |
| SE_113_1_21 | SE_113 | SE_113_1 | GTAATCATAAAGCCAATGAT | 342.47348197350243 | 344.7904071986942 | 370.322573487954 | 304.5061865585377 | 349.99618920070424 | 369.0481068039421 | 366.9707689527448 | 248.62796801627667 | 253.07580418017338 | 193.91584732395344 | 254.7113373394506 | 276.24208102925195 |
| SE_113_1_8 | SE_113 | SE_113_1 | GCCCACATAGCATTTCCTTA | 589.5746018784346 | 656.6008624044698 | 608.3165473828791 | 592.0953627527122 | 806.0518296743492 | 564.8341916342025 | 648.9248494970824 | 656.0753224459924 | 770.7712211522473 | 833.9839448819652 | 563.4167830550059 | 435.25537977298103 |
| SE_113_2_13 | SE_113 | SE_113_2 | GCGAGTCCTACAGCCTCAGA | 1716.7025172342655 | 1856.8712364497792 | 1777.5483527421793 | 1840.0502042468902 | 1871.9493149674029 | 1739.5507006157648 | 1740.151710840435 | 1452.0629614943002 | 2097.4212262230512 | 1831.2654454051544 | 1629.2135310007254 | 2070.6549267066616 |
| SE_113_2_17 | SE_113 | SE_113_2 | GCTGTAGGACTCGCAGTCAG | 2045.08690026582 | 2137.700524631904 | 2266.3741497462784 | 2218.730974710713 | 2277.0964188300363 | 1604.406323476293 | 2254.5564837419365 | 2448.465540540519 | 2476.5909398543986 | 2911.653737638609 | 2680.92485937007 | 2134.492382406699 |
| SE_113_2_19 | SE_113 | SE_113_2 | CTTGGGCTCTCAATCTTCCA | 632.9256755459666 | 620.6227329576495 | 608.3165473828791 | 575.178352388349 | 606.6600612812207 | 574.363602842755 | 735.0176985182542 | 555.8678524470367 | 522.1353433611998 | 797.5335976406205 | 609.1943966782252 | 827.565562075028 |
| SE_113_2_25 | SE_113 | SE_113_2 | GTTTAAACTTCTCTGAAGGG | 4.335107366753196 | 5.9963549078033775 | 7.900214901076352 | 0.0 | 0.0 | 17.326202197368175 | 4.30464245105859 | 1.89070698111237 | 16.871720278678225 | 1.4580138896537853 | 0.0 | 42.945197470934126 |
| SE_113_2_28 | SE_113 | SE_113_2 | AACTTCTCTGAAGGGAGGTG | 24.926867358830876 | 25.984204600481302 | 38.51354764274722 | 31.23140374959361 | 31.8178353818822 | 75.36897955855156 | 15.066248578705064 | 15.12565584889896 | 43.51127861343332 | 2.9160277793075706 | 4.695139858791716 | 138.12104051462597 |
| SE_113_3_0 | SE_113 | SE_113_3 | TTACACTATGAAGTACACAT | 132.22077468597246 | 130.9204154870404 | 139.2412876314707 | 145.74655083143685 | 158.0285823966816 | 111.75400417302473 | 118.37766740411122 | 94.5353490556185 | 90.57449833816732 | 51.759493082709376 | 62.210603128990236 | 68.48017975094902 |
| SE_113_3_12 | SE_113 | SE_113_3 | CCCAGCCGAGGGAATCCCGT | 419.42163773337165 | 374.7721817377111 | 396.9857987790867 | 468.4710562439042 | 384.99580812077465 | 381.17644834209983 | 435.84504816968223 | 303.4584704685354 | 374.7297872422217 | 497.1827363719408 | 260.58026216294024 | 228.65415950740604 |
| SE_113_3_32 | SE_113 | SE_113_3 | AGAGCTGCCATGCTATCAGA | 1383.9830268359576 | 1525.0729315513256 | 1401.300618078418 | 1513.4217733657238 | 1436.0449702356168 | 1455.4009845789267 | 1519.5387852236822 | 1259.2108494208385 | 1347.9616517386078 | 2382.394695694285 | 1116.2695014277306 | 1747.9856051682918 |
| SE_113_3_39 | SE_113 | SE_113_3 | ATTGTAGCAAAAGTTCTGAG | 68.27794102636283 | 124.92406057923702 | 126.40343841722164 | 96.29682822791364 | 129.39253055298764 | 168.0641613144713 | 79.63588534458391 | 103.04353047062416 | 107.44621861684554 | 110.80905561368769 | 88.03387235234467 | 119.5501443109788 |
| SE_113_3_5 | SE_113 | SE_113_3 | TTATAAACCACCTTATAGGT | 1.083776841688299 | 2.9981774539016888 | 2.962580587903632 | 2.602616979132801 | 1.0605945127294067 | 1.7326202197368175 | 0.0 | 0.0 | 0.0 | 0.0 | 0.0 | 1.1606810127279494 |
| SE_113_4_10 | SE_113 | SE_113_4 | TTCAGGGTGTCAGATCACAT | 272.027987263763 | 233.8578414043317 | 213.30580232906152 | 279.7813252567761 | 205.7553354695049 | 367.3154865842053 | 410.01719346333067 | 139.91231660231537 | 244.1959514019217 | 598.5147017028788 | 169.0250349165018 | 139.28172152735394 |
| SE_113_4_11 | SE_113 | SE_113_4 | ATTCAGGGTGTCAGATCACA | 119.21545258571288 | 180.8900397187352 | 225.15612468067604 | 170.47141213319847 | 129.39253055298764 | 123.01603560131404 | 94.70213392328898 | 195.6881725451303 | 164.2772763976564 | 115.18309728264904 | 53.994108376104734 | 120.71082532370674 |
| SE_113_4_12 | SE_113 | SE_113_4 | CACCCTGAATAATTCACACC | 702.2873934140177 | 767.5334281988323 | 707.0692336463335 | 692.296116449325 | 654.386814354044 | 1005.7860375572226 | 650.0010101098471 | 534.1247221642445 | 694.4044872592827 | 764.7282851234104 | 336.8762848683056 | 689.444521560402 |
| SE_113_4_13 | SE_113 | SE_113_4 | GACCTGGTGTGAATTATTCA | 182.0745094036342 | 259.842046004813 | 237.00644703229057 | 209.51066682019047 | 309.69359771698674 | 188.8556039513131 | 153.8909676253446 | 311.96665188354103 | 320.5626852948863 | 218.7020834480678 | 399.0868879972959 | 152.0492126673614 |
| SE_113_4_14 | SE_113 | SE_113_4 | GGACCTGGTGTGAATTATTC | 186.4096167703874 | 266.83779339725027 | 306.13332741670865 | 232.9342196323857 | 286.3605184369398 | 165.46523098486605 | 224.91756806781132 | 294.0049355629735 | 206.0125844554394 | 241.30129873770147 | 233.58320797488787 | 191.51236710011165 |
| SE_114_1_10 | SE_114 | SE_114_1 | GATGTGCGTGTTTCAATGGA | 2932.700133608537 | 2777.3117147975977 | 2827.2894077226997 | 2951.3676543365964 | 2735.27324832914 | 2632.716423890094 | 2952.9847214261927 | 2456.0283684649685 | 2699.475244588516 | 2687.1195986319262 | 3018.9749292030733 | 3049.109020436323 |

| | | | | | | | | | | | | | | | |
|---|---|---|---|---|---|---|---|---|---|---|---|---|---|---|---|
| SE_114_1_20 | SE_114 | SE_114_1 | TGAGTGGTCATTTAGAGCAG | 1393.7370184111523 | 1626.011572499349 | 1604.731151781134 | 1522.5309327926884 | 1572.8616623777102 | 1499.5828001822156 | 1411.9227239472175 | 1521.0737663049017 | 1516.67885452539 | 1734.3075217431776 | 1818.1929103170921 | 1261.660260835281 |
| SE_114_1_26 | SE_114 | SE_114_1 | ACTCACCAAGAGCAAGGCCT | 1075.1066269547925 | 910.4465535014795 | 842.360413827266 | 975.9813671748003 | 915.293064485478 | 825.5935347045935 | 980.3823182285938 | 840.4192531044484 | 927.0566300494772 | 852.9381254474644 | 1029.4094140400837 | 673.1949873822107 |
| SE_114_1_8 | SE_114 | SE_114_1 | GCTGGCAAACAGAATTGAGG | 2042.9193465824435 | 2479.492754376696 | 2439.1913507073236 | 2343.6565897090873 | 2331.186738979236 | 2252.4062856578626 | 2100.665516116592 | 2322.7335262965466 | 2294.553957900239 | 3645.0347241344634 | 2324.0942301018995 | 2099.6719520248607 |
| SE_114_1_9 | SE_114 | SE_114_1 | AGCTTCCTGAAAGCTGCCCA | 94.288585226882 | 118.92770567143364 | 135.29118018093254 | 117.11776406097604 | 161.21036593486983 | 34.65240439473635 | 106.5399006637001 | 52.93979547114636 | 130.53383584029996 | 46.65644446892113 | 89.2076573170426 | 106.78265317097134 |
| SE_114_2_12 | SE_114 | SE_114_2 | AATGCTGGACCTTCCTACTG | 528.8830987438898 | 601.6342757496055 | 624.1169771850318 | 671.4751806162626 | 556.8121191829385 | 707.77535976249 | 587.5836945694975 | 683.4905736721217 | 627.8055914223951 | 501.55677804090215 | 629.14874107809 | 388.8281392638631 |
| SE_114_2_19 | SE_114 | SE_114_2 | GATTGGGGATTGGCGGGTGA | 716.3764923559655 | 872.4696390853913 | 831.4976183382861 | 897.9028578008163 | 824.0819363907491 | 1003.1871072276173 | 936.2597331052433 | 1069.1947978190453 | 905.7449833816731 | 1262.6400284401782 | 975.415305663979 | 1362.6395089426126 |
| SE_114_2_2 | SE_114 | SE_114_2 | GGCCTCTTTTAGAACACACA | 2.167553683376598 | 11.992709815606755 | 11.850322351614528 | 1.3013084895664004 | 5.3029725636470335 | 12.128341538157722 | 0.0 | 0.0 | 0.0 | 4.374041668961356 | 0.0 | 4.642724050911798 |
| SE_114_2_27 | SE_114 | SE_114_2 | TATGGGTGACAATAGAACAA | 1379.6479194692045 | 1372.1658814023394 | 1368.712231611478 | 1421.0288706065094 | 1366.0457323954759 | 1766.4063140216854 | 1518.4626246109176 | 1343.347310080339 | 1487.3753403571593 | 1797.0021189982904 | 1461.3622810489217 | 1765.395820359211 |
| SE_114_2_6 | SE_114 | SE_114_2 | TGACAGGCTGGGCTGCCACA | 1605.0735025403708 | 1459.1130275654884 | 1460.5522298364906 | 1536.8453261779189 | 1534.6802599194516 | 1551.56140677432 | 1803.6451869935493 | 1324.440240269215 | 1478.4954875789076 | 1328.9796604194253 | 1691.4241341297156 | 1925.5698001156682 |
| SE_114_3_19 | SE_114 | SE_114_3 | ACTGCTTTCAGACCAGCCCT | 688.1982944720698 | 722.560766390307 | 658.6804173772408 | 722.2262117093522 | 639.5384911758323 | 565.700501744071 | 744.703144033136 | 738.3210761243805 | 662.4370172575767 | 861.6862087853871 | 744.179667618487 | 625.6070658603647 |
| SE_114_3_20 | SE_114 | SE_114_3 | AAGGGCTGGTCTGAAAGCAG | 3090.9315524950284 | 3492.8767337954673 | 3533.3711145063985 | 3203.8215013124777 | 3493.598324930666 | 3223.5399188203487 | 3030.4682855452475 | 4157.6646514661015 | 3561.708949356756 | 3321.355640631323 | 4266.708346676972 | 3998.546088847786 |
| SE_114_3_21 | SE_114 | SE_114_3 | GGGTCTGTAACTTACTGAAC | 69.36171786805113 | 73.95504386290831 | 72.08946097232172 | 81.98243484268323 | 124.08955798934059 | 99.625662634867 | 34.43713960846872 | 69.0108048106015 | 152.7334677859292 | 26.973256958595027 | 42.25625872912545 | 29.017025318198737 |
| SE_114_3_24 | SE_114 | SE_114_3 | AGCGAGGAAAGAGTAGAACA | 2461.257207474127 | 2683.368821242011 | 2676.1977977396145 | 2593.507819705836 | 2653.6074708489755 | 2634.449044109831 | 2381.543436048165 | 2590.268564123947 | 2886.840138209627 | 2682.745556962965 | 2879.29451840402 | 2323.6833874813547 |
| SE_114_3_25 | SE_114 | SE_114_3 | AAGGACAGTAGTGAGCAGCG | 6517.83392591343 | 6410.10339644181 | 6488.051487508954 | 6502.638522363303 | 6530.080414874958 | 7439.871223549894 | 6799.182751447042 | 5988.814362673432 | 6606.6104670192635 | 5629.391627953265 | 6611.930706143435 | 6929.265645985858 |
| SE_114_4_0 | SE_114 | SE_114_4 | GCAGAGGCTCAGGGAGGGAG | 5807.960094607593 | 7178.636217125276 | 6576.928905146064 | 4769.295614260857 | 5053.732853155623 | 3519.817976395345 | 4017.307567450429 | 2984.480969685876 | 4714.313839973826 | 2694.4096680801954 | 2110.4653665268766 | 2092.707865948493 |
| SE_114_4_11 | SE_114 | SE_114_4 | AACATGCTAAGATTACAGGT | 500.7049008599941 | 573.6512861798564 | 518.4516028831356 | 567.3705014509505 | 524.9942838010563 | 620.2780386657806 | 468.12986655262165 | 721.3047132943692 | 619.8137239219685 | 346.278298792774 | 1081.0559524867926 | 528.109860791217 |
| SE_114_4_12 | SE_114 | SE_114_4 | CCACAGATGGTGGTTACAGT | 264.4415493719449 | 225.86270152726053 | 237.99397389492512 | 260.2616979132801 | 200.45236290585788 | 237.36897010394398 | 238.90765603375175 | 90.75393509339376 | 312.57081779445974 | 218.7020834480678 | 134.98527094026184 | 247.22505571105322 |
| SE_114_4_24 | SE_114 | SE_114_4 | GCATCCAGACCCCTGAAAGG | 240.59845885480235 | 390.7624614918534 | 349.5845093726286 | 300.6022610898385 | 466.66158560093896 | 567.4331219638077 | 216.30828316569415 | 261.8629168840632 | 385.38561057612367 | 252.23640291010486 | 411.9985226089731 | 220.52939241831038 |
| SE_114_4_5 | SE_114 | SE_114_4 | ACTCAGATAATTCACAGCAG | 251.43622727168534 | 189.88457208044028 | 218.24343664223423 | 232.9342196323857 | 199.39176839312847 | 225.24062856578627 | 212.00364071463557 | 230.66625169570915 | 239.75602501279585 | 439.5911877306163 | 200.71722896334586 | 388.8281392638631 |
| SE_114_5_38 | SE_114 | SE_114_5 | CGCAGCGTGACAGTTGTAAG | 12480.77410888245 | 10459.641744178358 | 10687.015707431035 | 11771.636596617658 | 10863.669593887313 | 11687.389692234703 | 12722.370764103662 | 10396.997689136922 | 9394.884239390296 | 11451.24108934083 | 10458.424035458547 | 11706.628694374098 |
| SE_114_5_39 | SE_114 | SE_114_5 | GCGTGACAGTTGTAAGAGGC | 7775.015062271857 | 8694.714616314897 | 8657.648004717048 | 7652.995227140001 | 7949.155872906904 | 8074.01022397357 | 7241.484763293312 | 8062.919920953702 | 8101.977674876849 | 9001.77775472247 | 7589.693581736809 | 7697.636476411761 |
| SE_114_5_40 | SE_114 | SE_114_5 | AGGCAGGCAAGAGTATTTAA | 3346.702887133467 | 3690.7564457529784 | 3808.891109181436 | 3737.357982034702 | 3717.3837671165707 | 4447.636104064411 | 3475.998779229811 | 4512.172210424671 | 3669.1551679736017 | 4236.259356389073 | 5047.275348201095 | 3959.0829344150357 |
| SE_114_5_41 | SE_114 | SE_114_5 | GGCAGGCAAGAGTATTTAAT | 2591.310428476723 | 2872.254000837818 | 2996.1565012332067 | 3054.171025012342 | 3392.841846221372 | 3837.753786717051 | 2690.401531911619 | 3267.141663362175 | 3432.9510840721064 | 3251.370973927941 | 3875.837953432562 | 3648.020423003945 |
| SE_114_5_43 | SE_114 | SE_114_5 | GAGTATTTAATGGGCATTGA | 40.09974314246706 | 110.93256579436247 | 99.74021312608895 | 85.88636031138243 | 106.05945127294068 | 149.0053388973663 | 91.47365208499504 | 48.213028018365435 | 147.40555611897818 | 131.22125006884067 | 161.98232512831422 | 183.38760001101602 |
| SE_114_6_4 | SE_114 | SE_114_6 | TGGCACAGCCCTGGTCGAAG | 203.7500462374002 | 249.84812115847404 | 194.54279193900518 | 217.31851775758886 | 204.6947409567755 | 143.80747823815585 | 218.46060439122343 | 205.14170745069214 | 348.97821418529173 | 278.480652923873 | 173.7201747752935 | 123.03218734916264 |
| SE_114_6_5 | SE_114 | SE_114_6 | GGATACAACCCCTTCGACCA | 177.73940203688102 | 169.89672238776234 | 164.91698605996885 | 148.34916781056964 | 182.42225618945795 | 140.34223779868222 | 201.24203458698906 | 215.5405958468102 | 121.65398306204825 | 142.88536118607095 | 165.503680022408 | 103.3006101327875 |
| SE_114_6_51 | SE_114 | SE_114_6 | TCTGGCTTGCCAACTCACTT | 1915.0336792632243 | 1645.0000297073932 | 1533.629217671447 | 1767.1769288311718 | 1683.1634917015685 | 1767.2726241315538 | 2302.9837113163458 | 2348.2580705415635 | 1895.8485681567374 | 1714.6243342328514 | 2516.59496431236 | 1701.5583646591738 |
| SE_114_6_52 | SE_114 | SE_114_6 | CCACTGGGACCCAAGTGAGT | 459.52138087583876 | 559.6597913949819 | 547.0898818995374 | 508.8116194204626 | 589.6905490775501 | 776.2138584420942 | 447.6828149100933 | 414.064828863609 | 504.37563780469645 | 680.1634795234909 | 255.88512230414852 | 896.045741825977 |
| SE_114_6_7 | SE_114 | SE_114_6 | CGAAGGGGTTGTATCCCTGA | 4170.373286816574 | 4539.240665207157 | 4527.810665179384 | 4342.466429683078 | 4395.103660750661 | 4695.400795486776 | 4099.095774020542 | 4357.134237973457 | 5214.249551389397 | 4602.220842692173 | 4359.437358888108 | 4895.752511686491 |
| SE_115_1_17 | SE_115 | SE_115_1 | TCTTGCCTATGCCATTCACC | 4284.169855193845 | 3528.8548632422876 | 3610.398209791893 | 4039.261551614107 | 3297.3883400757254 | 3899.261804517708 | 3973.1849823270786 | 3412.726100907828 | 3076.8689876642134 | 3837.492557568763 | 3331.2017298127225 | 5100.03236992661 |
| SE_115_1_23 | SE_115 | SE_115_1 | TGGAGGTGATCCCTGCCACA | 1072.939073271416 | 1011.3851944495029 | 1009.252453612504 | 1129.5357689436355 | 1016.0495431947717 | 800.4705415184096 | 1180.5481922028182 | 1101.3368164979556 | 1008.7512756093929 | 1093.510417240339 | 879.1649385587489 | 1530.9382557881654 |
| SE_115_1_26 | SE_115 | SE_115_1 | AAAGGATGACTGCATCCCTG | 1416.4963320866066 | 1447.1203177498817 | 1412.163413567398 | 1488.6969120639621 | 1349.0762201918053 | 874.1069008572244 | 1589.4892250533844 | 1578.740329228829 | 1136.6211556162173 | 1441.2467299227667 | 1536.4845187895892 | 1043.4522304424265 |
| SE_115_1_29 | SE_115 | SE_115_1 | CATACAGATGGAAAACCCCT | 1596.4032878068642 | 1906.840860681474 | 2161.696302307017 | 1817.9279599242614 | 1773.314025283568 | 2029.7645874216817 | 1562.5852097342681 | 1828.313650735662 | 1979.3191842723033 | 3313.3365642382273 | 1744.2444575411225 | 2851.793248272572 |
| SE_115_1_32 | SE_115 | SE_115_1 | ACAGGTGGAGAAGCTAACCA | 828.0055070498604 | 1007.3876245109674 | 976.6640671455641 | 994.19968602873 | 988.4740858638071 | 854.181768330251 | 867.3854538883058 | 782.7526901805212 | 1039.830760333274 | 850.7511046129837 | 762.9602270536539 | 935.5088962587272 |
| SE_115_2_0 | SE_115 | SE_115_2 | GGCTGGCTAATGAATCACCC | 786.821987065705 | 768.5328206834662 | 817.6722422614025 | 761.2654663963442 | 826.2031254162079 | 583.0267039414391 | 684.4381497183158 | 656.0753224459924 | 929.7205858829527 | 435.94615300648184 | 774.6980767006331 | 625.6070658603647 |
| SE_115_2_1 | SE_115 | SE_115_2 | CCTGGTCTCTCAAACTTCCT | 506.1237850684356 | 561.6585763642497 | 496.72601190517565 | 540.0430231700561 | 473.0251526773154 | 551.8395399861763 | 446.6066542973287 | 397.9938195241539 | 491.94384391514404 | 360.85843768931187 | 403.7820278560876 | 438.7374228111649 |
| SE_115_2_12 | SE_115 | SE_115_2 | ATTATGGCCCATTTGGCCCA | 36.84841261740216 | 24.984812115847404 | 36.53849391747813 | 41.641871666124814 | 39.24199697098805 | 16.459892087499767 | 51.655709412703075 | 29.305958207241734 | 64.82292528123739 | 8.019076393095819 | 1.173784964697929 | 56.873369623669525 |
| SE_115_2_15 | SE_115 | SE_115_2 | TAGTCCTCATGCAAGACCCA | 657.8525429047975 | 580.6470335722937 | 660.65547110251 | 702.7065843658562 | 587.5693600520914 | 511.12296482236115 | 645.6963676587885 | 519.9444198059017 | 670.4288847580032 | 625.487958661474 | 315.7481555037429 | 601.2327645930778 |
| SE_115_2_4 | SE_115 | SE_115_2 | GAAGAGGTTTGCTCAAGGAA | 560.3126271528505 | 773.5297831066356 | 774.2210603054825 | 704.0078928554226 | 755.1432930633376 | 556.1710905355184 | 643.5440464332592 | 710.9058248982511 | 1016.7431431098194 | 753.0641740061801 | 728.920463077414 | 624.4463848476368 |
| SE_115_3_1 | SE_115 | SE_115_3 | CTAGGCTTGAGGAAGCTAAA | 471.44292613441 | 459.72054293159226 | 489.81332386673387 | 527.0299382743922 | 477.26753072823306 | 442.68446614275683 | 454.13977858668125 | 477.40351273087344 | 401.36934557697674 | 293.7897987652377 | 518.8129543964847 | 417.8451645820618 |
| SE_115_3_12 | SE_115 | SE_115_3 | TCCCTCACAGAGTTGTTGTA | 141.97476626116716 | 179.89064723410132 | 200.46795311481245 | 143.14393385230406 | 180.30106716399914 | 131.67913669999814 | 112.99686434028798 | 115.33312584785457 | 117.21405667292241 | 96.95792366197672 | 203.06479889274172 | 184.54828102374395 |
| SE_115_3_2 | SE_115 | SE_115_3 | TTACAGAGGCTGGCAGACCA | 408.5838693164887 | 518.6846995249921 | 505.61375366888655 | 370.87291952642414 | 572.7210368738796 | 718.1710810809109 | 425.08344204203576 | 402.72058697693484 | 507.9275789159971 | 518.3239377719207 | 417.8674474324627 | 410.8810785056941 |
| SE_115_3_8 | SE_115 | SE_115_3 | TGTCCACATGGCTAAAGCTT | 955.8911743690796 | 876.4672090239269 | 952.963422442335 | 845.8505182181602 | 936.5049547400662 | 915.689786130908 | 965.3160696498887 | 1135.3695421579782 | 926.1686447716521 | 846.3770629440223 | 1240.690707685711 | 643.0172810512839 |
| SE_115_3_9 | SE_115 | SE_115_3 | ATACCGAAGCTTTAGCCATG | 303.4575156727237 | 240.85358879676897 | 308.10838114197776 | 256.35777244458086 | 238.63376536411653 | 283.28340592696964 | 322.84818382939426 | 375.30533575080545 | 252.18781890234823 | 151.63344452399366 | 178.4153146340852 | 385.3460962256792 |
| SE_115_4_14 | SE_115 | SE_115_4 | GGAACTGTGCTATTAATACT | 228.67691359623106 | 246.84994370457235 | 277.49504840030687 | 303.2048780689713 | 253.4820885423282 | 271.1550643888119 | 273.34479564222045 | 187.17999113012462 | 264.6196127919006 | 303.2668890479873 | 292.2724562097843 | 366.77520002203204 |
| SE_115_4_17 | SE_115 | SE_115_4 | GACTGGAGTTGGCTTCCAAG | 761.8951197068741 | 807.5091275841881 | 805.8219199097879 | 847.1518267077267 | 762.5674546524434 | 814.3315032763043 | 687.6666315566098 | 645.6764340498744 | 904.856998103848 | 661.2092989579917 | 539.9410837610474 | 734.711081056792 |
| SE_115_4_33 | SE_115 | SE_115_4 | GTAGGAATTTTCCCACAGAA | 97.53991575194691 | 201.87728189604704 | 186.64257703792882 | 123.62430650880805 | 209.99771352042254 | 178.4598826328922 | 102.23525821264151 | 193.79746556401793 | 116.32607139509724 | 210.68300705497197 | 66.90574298778195 | 234.4575645710458 |
| SE_115_4_34 | SE_115 | SE_115_4 | CTGTGGGAAAATTCCTACCG | 82.36703996831072 | 121.92588312533533 | 139.2412876314707 | 89.79028578008163 | 137.87728665482288 | 117.81817494210358 | 107.61606127646475 | 192.85211207346174 | 173.1571291759081 | 222.34711817220227 | 173.7201747752935 | 71.96222278913287 |
| SE_115_4_5 | SE_115 | SE_115_4 | TGGGGTACATATGACCCAAA | 122.46678311077778 | 118.92770567143364 | 131.34107273039436 | 111.91253010271043 | 110.3018293238583 | 177.5935725230238 | 97.93061576158291 | 71.84686528227006 | 120.76599778422309 | 51.759493082709376 | 92.72901221113639 | 141.60308355280984 |
| SE_115_5_11 | SE_115 | SE_115_5 | GCGGGAAAGCAGACATAGCA | 5519.675454718506 | 5129.881623625789 | 5341.532799990248 | 5293.722935556117 | 5458.879957018256 | 5948.085214356494 | 5984.5291675842045 | 5465.088528905306 | 5971.700993374267 | 4787.388606678204 | 4986.238530036802 | 5342.614701586751 |
| SE_115_5_14 | SE_115 | SE_115_5 | TCCCTTTGACAGACTCCAAC | 69.36171786805113 | 80.95079125534559 | 107.6404280271653 | 78.07850937398402 | 64.69626527649382 | 64.10694813026224 | 55.960351863761666 | 78.46433971616335 | 87.02255722686664 | 137.0533056274558 | 106.81443178751154 | 125.35354937461854 |
| SE_115_5_15 | SE_115 | SE_115_5 | TTTATATCTTGACAAATGGG | 0.0 | 0.9993924846338962 | 0.0 | 2.602616979132801 | 0.0 | 36.38502461447317 | 0.0 | 0.0 | 0.8879852778251698 | 0.0 | 0.0 | 3.482043038183848 |
| SE_115_5_16 | SE_115 | SE_115_5 | GAAATGGAAGAAAAGCACAA | 211.3364841292183 | 261.8408309740808 | 235.0313933070215 | 269.37085734024487 | 260.90625013143404 | 277.2192351578908 | 213.0798013274002 | 334.6551356568895 | 316.12275890576046 | 206.3089653860106 | 187.80559435166865 | 208.9225822910309 |
| SE_115_5_2 | SE_115 | SE_115_5 | ATGAGAGCAAGGGTTAGAAG | 1666.8487825166037 | 1761.9289504095589 | 1800.2614705827739 | 1743.7533760189765 | 1718.163110621639 | 1638.1924177611609 | 1725.0854622617298 | 1764.0296133778413 | 1489.1513109128098 | 1441.2467299227667 | 1605.7378317067669 | 1080.594022849721 |
| SE_116_1_12 | SE_116 | SE_116_1 | TAATCCGCCATGGATACCGA | 471.44292613441 | 540.6713341869379 | 505.61375366888655 | 525.7286297848258 | 501.6612045210094 | 612.4812476769649 | 490.7292394206792 | 621.0972432954136 | 487.5039175260182 | 698.3886531441632 | 640.8865907250693 | 1377.728362108076 |
| SE_116_1_13 | SE_116 | SE_116_1 | CTAATCCGCCATGGATACCG | 128.96944416090756 | 137.91616287947767 | 149.11655625781614 | 132.73346593577284 | 155.90739337122278 | 86.63101098684088 | 117.30150679134657 | 134.24019565897828 | 69.26285167036325 | 172.04563897914667 | 124.42120625798047 | 193.83372912556754 |
| SE_116_1_29 | SE_116 | SE_116_1 | CTGAGGGTAAAAGAGCATGG | 1536.7955615140079 | 1533.0680714283967 | 1452.6520149354142 | 1782.7926307059686 | 1770.1322417453798 | 2131.9891803861537 | 1726.1616228744945 | 1882.1987996973644 | 1789.2903348177172 | 3308.233515624439 | 2500.1619748065887 | 3174.4625698109417 |
| SE_116_1_6 | SE_116 | SE_116_1 | GCAGTTTTTATTTTGATGAG | 42.26729682584366 | 71.95625889364052 | 106.65290116453076 | 57.25757354092162 | 98.63528968383483 | 91.82887164605133 | 55.960351863761666 | 87.9178746217252 | 94.126439449468 | 131.22125006884067 | 72.7746678112716 | 69.64086076367697 |
| SE_116_1_8 | SE_116 | SE_116_1 | ATTCCAGTGCACTCATCCCT | 187.49339361207572 | 152.90705014898612 | 97.76515940081985 | 203.00412437235846 | 113.48361286204653 | 74.50266944868315 | 140.9770402721688 | 186.23463763956843 | 114.5501008394469 | 116.64111117230283 | 278.1870366334092 | 218.20803039285448 |
| SE_116_2_10 | SE_116 | SE_116_2 | GGTATTTCCGCCCTCTTGGA | 14.089098941947885 | 29.981774539016886 | 18.763010390056337 | 23.423552812195208 | 20.15129574185873 | 60.641707690788614 | 27.980175931880833 | 80.35504669727572 | 28.415528890405433 | 48.114458358574915 | 43.43004369382337 | 0.0 |
| SE_116_2_12 | SE_116 | SE_116_2 | ACACCAACCCTCCAAGAGGG | 358.73013459882696 | 464.7175035547617 | 473.0253672019466 | 342.24413275596334 | 499.54001549555056 | 321.40105076117965 | 378.80853569315593 | 578.5563362203852 | 626.9176061445698 | 570.8124377994569 | 294.6200261391802 | 434.0946987602531 |
| SE_116_2_20 | SE_116 | SE_116_2 | CTCTGAGCAGATTAGAACAG | 145.22609678623206 | 153.90644263362 | 154.05419057098888 | 167.86879515406565 | 203.6341464440461 | 269.4224441690751 | 199.0897133614598 | 211.75918188458544 | 103.0062922277197 | 412.6179307720212 | 338.0500698330036 | 414.3631215438779 |
| SE_116_2_24 | SE_116 | SE_116_2 | CTCAGATTCCTGGAGCACCG | 119.21545258571288 | 119.92709815606754 | 111.59053547770347 | 135.33608291490566 | 143.1802592184699 | 353.4545248263108 | 173.26185865510826 | 187.17999113012462 | 152.7334677859292 | 262.44250013768135 | 273.4918967746175 | 229.814840520134 |
| SE_116_2_28 | SE_116 | SE_116_2 | GACATCAATGAGATTGACGA | 1148.8034521895968 | 1140.3068249672756 | 1105.0425592880547 | 1094.4004397253427 | 1189.9870432823943 | 899.2298940434083 | 1099.8361462454698 | 1176.9650957424503 | 919.0647625490507 | 1003.1135560818043 | 1216.0412234270545 | 1045.7735924678825 |
| SE_116_3_0 | SE_116 | SE_116_3 | TGTCAATGAGGAATTACCCC | 1274.5215658254394 | 1038.3687915346181 | 1181.0821277109146 | 1165.9724066514948 | 1211.1989335369824 | 1146.994585465773 | 1373.18094188769 | 1124.0253002713039 | 1173.9165372848745 | 1806.47920928104 | 998.8910049579376 | 1154.8776076643096 |
| SE_116_3_20 | SE_116 | SE_116_3 | GCAGGGTGCCTACATACTGT | 8428.5324978099 | 7092.6884634467615 | 6934.413629419768 | 8043.3877740099215 | 7510.0697446369295 | 7198.170702896608 | 8504.89732267901 | 6988.053002191319 | 6987.556151206261 | 7943.98867777865 | 7032.145723505293 | 8183.961820744771 |
| SE_116_3_33 | SE_116 | SE_116_3 | CAGTCCACCGAGAAGAAGAG | 612.3339155538889 | 715.5650189978696 | 737.6825663880044 | 642.8463938458018 | 742.4161589105847 | 604.6844566881493 | 668.2957405268461 | 695.7801690493521 | 723.7080014275134 | 497.1827363719408 | 593.9351921371521 | 550.162800033048 |
| SE_116_3_5 | SE_116 | SE_116_3 | GATAGCAATGGGAAGTTGCG | 893.0321175511583 | 863.4751067236863 | 816.6847153987679 | 878.3832304573203 | 819.8395583398315 | 729.4331125092001 | 955.630624135007 | 828.129657727218 | 836.48213171131 | 646.6291600614538 | 907.3357777114992 | 371.41792407294383 |
| SE_116_3_9 | SE_116 | SE_116_3 | CGTACTATCTGGCAGCAGAA | 4960.446604407344 | 5357.743110122317 | 5469.91129213274 | 5314.543871389179 | 5590.393676596703 | 5676.063839857814 | 5098.8489832789 | 6331.032326254771 | 5807.42371697661 | 5767.902947470375 | 6015.647944076886 | 5644.391764896018 |
| SE_116_4_2 | SE_116 | SE_116_4 | ACTCACATGTTACTGCACAG | 1309.202424759465 | 1424.1342906033021 | 1197.870084375702 | 1305.2124150350996 | 1344.8338421408878 | 1207.6362931565618 | 1187.0051558794062 | 1220.4513563080347 | 1413.6725622976703 | 1209.422521467815 | 1058.754038157532 | 1366.1215519807965 |
| SE_116_4_4 | SE_116 | SE_116_4 | GTACAGATACACAAGCTGTA | 461.68893455921534 | 523.6816619481616 | 520.4266566084048 | 495.7985345247986 | 621.5083844594324 | 589.9571848203864 | 573.5936066035571 | 557.7585594281492 | 597.6140919763393 | 515.4079099926131 | 605.6730417841313 | 532.7525848421288 |
| SE_116_4_44 | SE_116 | SE_116_4 | TACCTCCCTGATGCCCCACT | 1116.2901469389478 | 1026.3760817190114 | 1004.3148192993312 | 1171.1776406097604 | 916.3536589982074 | 830.791395363804 | 1304.3066626707528 | 1062.5773233851519 | 1303.5623878473493 | 1207.2355006333341 | 860.384379123582 | 951.7584304369185 |
| SE_116_4_7 | SE_116 | SE_116_4 | ATTGTTCGTTTTCGTCGAAA | 114.88034521895969 | 81.9501837399795 | 121.46580410404891 | 131.43215744620645 | 101.81707322202305 | 152.47057933683993 | 205.54667703804768 | 134.24019565897828 | 58.607028336461205 | 87.48083337922712 | 113.85714157569912 | 89.37243798005211 |
| SE_116_4_8 | SE_116 | SE_116_4 | ATAGCTCCTGGGATGCTTCA | 128.96944416090756 | 94.94228604022014 | 73.07698783495626 | 55.95626505135522 | 78.4839939419761 | 89.2299413164461 | 138.82471904663953 | 58.61191641448347 | 65.71091055906257 | 43.74041668961356 | 29.344624117448227 | 17.410215190919242 |
| SE_117_1_2 | SE_117 | SE_117_1 | GCATGAGCCTAGGATATGGA | 2584.807767426593 | 2547.4514433318013 | 2479.67995207534 | 2704.11904131898 | 2339.6714950810715 | 1840.9089834703686 | 2638.7458224989155 | 2410.651400918272 | 2433.0796612409654 | 3102.653557183255 | 2056.4712581507715 | 2153.063278610346 |
| SE_117_1_21 | SE_117 | SE_117_1 | ACAATACTCCTAAAAGAAGG | 239.51468201311405 | 226.86209401189444 | 241.94408134546327 | 213.41459228888968 | 187.725228753105 | 96.16042219539337 | 144.20552211046277 | 224.04877726181584 | 147.40555611897818 | 206.3089653860106 | 278.1870366334092 | 80.08698987822851 |
| SE_117_1_24 | SE_117 | SE_117_1 | GGAACTTGTCCGCTTCTCGG | 197.2473851872704 | 421.7436285155042 | 433.5242926965648 | 264.1656233819793 | 359.5415398152689 | 292.81281713552215 | 255.05006522522146 | 335.60048914744567 | 392.48949279872505 | 529.980048889151 | 284.05596145689884 | 615.1609367458132 |
| SE_117_1_5 | SE_117 | SE_117_1 | TGCTGCAGCCTCTTTCACAG | 309.9601767228535 | 80.95079125534559 | 183.6799964500252 | 301.9035695794049 | 73.18102137832906 | 699.1122586638058 | 451.98745736115194 | 255.24544245016995 | 652.6691792014998 | 26.973256958595027 | 131.46391604616804 | 181.06623798556012 |
| SE_117_1_7 | SE_117 | SE_117_1 | ACACAATTATACCATCACTG | 1342.7995068518023 | 1389.1555536411156 | 1532.6416908088124 | 1452.2602743561029 | 1392.5605952137112 | 1775.935725230238 | 1335.5153204409276 | 1315.9320588542096 | 1618.7971614752846 | 1415.0024799089986 | 1373.328408696577 | 1479.8682912281356 |
| SE_117_2_0 | SE_117 | SE_117_2 | CACTCACAGAATTATTCGAA | 651.3498818546676 | 938.4295430712285 | 1069.4915922332111 | 772.9772428024419 | 974.6863571983248 | 952.9411208552496 | 782.3687654798987 | 1130.6427747051973 | 936.8244681055542 | 1140.16686170926 | 1352.2002793320144 | 1048.0949544933383 |
| SE_117_2_10 | SE_117 | SE_117_2 | ATTTGCTTACCAATTACATG | 8.670214733506391 | 14.990887269508443 | 24.688171565863602 | 18.218318853929606 | 11.666539640023474 | 38.98395494407839 | 2.152321225529295 | 8.508181415005666 | 16.871720278678225 | 2.187020834480678 | 1.173784964697929 | 1.1606810127279494 |

| | | | | | | | | | | | | | | | |
|---|---|---|---|---|---|---|---|---|---|---|---|---|---|---|---|
| SE_117_2_12 | SE_117 | SE_117_2 | TTACATGTGGATGCTGAAGA | 329.4681598732429 | 382.76732161478225 | 358.4722511363395 | 257.6590809341473 | 405.1471038626334 | 261.62565318025946 | 262.583189514574 | 437.69866612751366 | 277.93939195927817 | 306.1829168272949 | 289.9248862803885 | 372.57860508567177 |
| SE_117_2_21 | SE_117 | SE_117_2 | CAGTGCCTACCTGCCTAAGA | 211.3364841292183 | 155.9052276028878 | 166.89203978523796 | 160.06094421666725 | 215.30068608406958 | 106.55614351381428 | 195.86123152316583 | 132.3494886778659 | 193.580790565887 | 185.89677093085763 | 161.98232512831422 | 175.26283292192036 |
| SE_117_2_8 | SE_117 | SE_117_2 | ATATGTGGCTTGGGCTACAG | 267.69287989700985 | 347.7885846525959 | 348.59698250999406 | 330.5323563498657 | 357.4203507898101 | 283.28340592696964 | 264.7355107401033 | 351.6714984869008 | 297.47506807143185 | 169.1296111998391 | 549.3313634786308 | 340.0795367292892 |
| SE_118_1_15 | SE_118 | SE_118_1 | CGAGGCATCAGACCTTACCA | 25707.18668484645 | 23005.015603787655 | 23778.659325377186 | 25087.926370350633 | 23631.10633812391 | 24268.8114178536 | 26116.265750572464 | 24925.190132004373 | 22822.997610662514 | 25438.69733973442 | 24475.764083881215 | 21187.07120633599 |
| SE_118_1_21 | SE_118 | SE_118_1 | TTTCCAGGCTCCATGACAAG | 22.759313675454276 | 57.964764108765976 | 34.56344019220904 | 15.615701874796805 | 22.272484767317543 | 1.7326202197368175 | 7.533124289352532 | 8.508181415005666 | 43.51127861343332 | 31.347298627556384 | 4.695139858791716 | 1.1606810127279494 |
| SE_118_1_27 | SE_118 | SE_118_1 | CTAGGAAGCCACATCGTCCA | 322.9654988231131 | 327.800734959918 | 331.8090258452068 | 360.4624516098929 | 224.84603669863424 | 280.68447559736444 | 403.5602297867428 | 304.4038239590916 | 304.5789502940323 | 463.64841690990374 | 291.0986712450864 | 233.29688355831783 |
| SE_118_1_4 | SE_118 | SE_118_1 | TTAATCGTATAGCCTCCAGT | 57.440172609479845 | 81.9501837399795 | 79.98967587339807 | 62.46280749918722 | 108.18064029839948 | 114.35293450262995 | 67.79811860417279 | 36.868786131691216 | 34.63142583518162 | 26.973256958595027 | 15.259204541073078 | 35.98111139456643 |
| SE_118_1_6 | SE_118 | SE_118_1 | AGTGCTCTAAAGAAGTACTG | 87.78592417675222 | 144.91191027191493 | 149.11655625781614 | 93.69421124878083 | 141.0590701930111 | 129.0802063703929 | 108.6922218892294 | 88.86322811228139 | 129.64585056247478 | 72.90069448268926 | 152.59204541073078 | 208.9225822910309 |
| SE_118_2_0 | SE_118 | SE_118_2 | ACCATTAATCCAACCCAGAG | 83.45081680999901 | 139.91494784874547 | 147.14150253254707 | 134.03477442533924 | 214.24009157134017 | 74.50266944868315 | 99.00677637434757 | 185.28928414901227 | 136.74973278507613 | 258.7974654135469 | 210.1075086809293 | 292.4916152074432 6 |
| SE_118_2_1 | SE_118 | SE_118_2 | TTGCATTGCCTTACTAAAAT | 94.288585226882 | 139.91494784874547 | 137.26623390620162 | 134.03477442533924 | 116.66539640023474 | 397.6363404295996 | 95.77829453605362 | 190.96140509234937 | 152.7334677859292 | 178.6067014825887 | 261.75404712763816 | 113.74673924733905 |
| SE_118_2_17 | SE_118 | SE_118_2 | CATCCAAGACGCACCATGAG | 1738.3780540680314 | 1639.0036747995898 | 1642.2571725612468 | 1720.3298232067814 | 1689.527058777945 | 1684.1068535841866 | 1759.5226018701985 | 1729.0515342272624 | 1332.8659020155799 | 1877.9218898740755 | 1298.2061709559096 | 1691.1122355446223 |
| SE_118_2_20 | SE_118 | SE_118_2 | GAATCTGGAGACAGGAACTG | 76.94815575986922 | 103.93681840192521 | 110.60300861506893 | 135.33608291490566 | 43.48437502190568 | 102.22459296447224 | 73.17892166799602 | 108.71565141396128 | 67.4868811147129 | 116.64111117230283 | 123.24742129328254 | 38.30247342002233 |
| SE_118_2_38 | SE_118 | SE_118_2 | GCACTTCAGGAACCACGATA | 891.94834070947 | 911.4459459861133 | 1008.2649267498695 | 841.946592749461 | 852.717988234443 | 1027.4437903039327 | 783.4449260926633 | 850.8181415005665 | 882.6573661582188 | 899.5945699163856 | 1104.5316517807512 | 1166.4844177915893 |
| SE_118_3_13 | SE_118 | SE_118_3 | GGTAGTTGACAACCAAACAC | 614.5014692372655 | 613.6269855652123 | 565.8528922895937 | 641.5450853562354 | 534.539634415621 | 693.048087894727 | 570.3651247652632 | 648.5124945215429 | 441.3286830791094 | 673.6024170200488 | 566.9381379490998 | 529.270541803945 |
| SE_118_3_18 | SE_118 | SE_118_3 | GTACAGTAGAGGTTTAACTA | 248.18489674662044 | 282.82807315139263 | 290.3328976145559 | 279.7813252567761 | 265.1486281823517 | 316.2031901019692 | 232.45069235716386 | 258.08150292183853 | 269.0595391810264 | 158.9235139722626 | 323.9646502566284 3 | 341.24021774201714 |
| SE_118_3_19 | SE_118 | SE_118_3 | CGTACAGTAGAGGTTTAACT | 13197.150601238416 | 13051.06645683405 | 13329.637591841076 | 13058.630692798828 | 13229.85595178662 | 13795.122189544541 | 12025.01868703217 | 13136.632104768747 | 13430.777327105692 | 12800.632944215407 | 14053.727382328305 | 13936.296919824488 |
| SE_118_3_24 | SE_118 | SE_118_3 | TGACTCCCAACGACGAACAG | 887.6132333427169 | 910.4465535014795 | 921.3625628380296 | 744.3484560319811 | 894.0811742308899 | 722.5026316302528 | 835.1006355053664 | 916.0475323489433 | 864.8976606017154 | 475.312528027134 | 833.3873249355296 | 591.9473164912542 |
| SE_118_3_3 | SE_118 | SE_118_3 | TGTCGAGACTGAACATTCAG | 1959.4685297724445 | 1917.8341780124467 | 1970.1160909559153 | 1963.6745107556983 | 2056.4927601823197 | 1790.662997098001 | 1891.8903572402503 | 1997.5319255452189 | 1934.919920381045 | 1816.6853065086166 | 2153.8954102207 | 2306.2731722904355 |
| SE_118_4_1 | SE_118 | SE_118_4 | AATCAAGGAGGGTTTTATAG | 5.4188842084414945 | 13.991494784874547 | 6.912688038441808 | 10.410467916531204 | 16.969512203670508 | 0.0 | 12.913927353175769 | 1.89070698111237 | 12.431793889552377 | 11.664111117230282 | 1.173784964697929 | 0.0 |
| SE_118_4_12 | SE_118 | SE_118_4 | AAGGTATTGAGGCAGCCAAA | 939.6345217437552 | 899.4532361705066 | 756.4455767780607 | 1070.9768869131476 | 897.2629577690781 | 1267.4116907374819 | 1058.9420429604131 | 885.7962206511454 | 1070.0222597793295 | 841.2740143302341 | 872.1222287705613 | 1049.2556355060663 |
| SE_118_4_3 | SE_118 | SE_118_4 | TAAGGAATCAGAGATACTGA | 1095.6983869468702 | 1251.239390761638 | 1261.0718035843126 | 1220.6273632132836 | 1284.3799549153116 | 1066.4277452480112 | 1084.7698976667646 | 1159.948732912439 | 991.8795553307147 | 1318.044556247022 | 1150.3092654039704 | 1413.7094735026424 |
| SE_118_4_4 | SE_118 | SE_118_4 | CTATCTGTCACCTCTTCACA | 467.10781876765685 | 422.74302100013807 | 462.1625717129666 | 455.45797134824016 | 526.0548783137857 | 412.3636122973626 | 468.12986655262165 | 467.9499778253116 | 579.8543864198359 | 758.1672226199684 | 563.4167830550059 | 559.4482481348716 |
| SE_118_4_7 | SE_118 | SE_118_4 | TTCCCTCACTGCTATCTCAG | 8.670214733506391 | 9.993924846338961 | 5.925161175807264 | 16.917010364363207 | 12.727134152752882 | 21.65775274671022 | 8.60928490211718 | 12.289595377230405 | 21.311646667804077 | 0.0 | 9.390279717583432 | 1.1606810127279494 |
| SE_119_1_16 | SE_119 | SE_119_1 | GCTACCACACCCAAGAGACT | 1751.383376168291 | 1957.8098773978027 | 1793.348782544332 | 1787.9978646642342 | 1832.7073179964148 | 1486.5881485341895 | 1739.0755502276704 | 1859.510315924016 | 1989.0870223283803 | 2528.196084659664 | 2084.642097303522 | 1710.8438127609975 |
| SE_119_1_25 | SE_119 | SE_119_1 | GGAGGGAGGGTAGGACACAT | 10538.646008577018 | 8624.757142390525 | 8464.092739640677 | 9590.643568104371 | 8315.060979798549 | 8709.88184461698 | 10073.939496089864 | 7817.128013409094 | 8563.730019345938 | 7502.939476158379 | 7214.082393033472 | 7858.971137180945 |
| SE_119_1_30 | SE_119 | SE_119_1 | ATATAGCTCCCAAAGCCTGC | 101.8750231187001 | 175.89307729556572 | 164.91698605996885 | 115.81645557140963 | 120.90777445115236 | 72.77004922894633 | 89.32133085946575 | 148.42049801732105 | 118.99002722857276 | 171.31663203431978 | 154.93961534012664 | 11.606810127279495 |
| SE_119_1_38 | SE_119 | SE_119_1 | GGGGATTGTGGGCCACTCCA | 184.24206308701082 | 198.87910444214535 | 222.1935440927724 | 171.77272062276487 | 167.57393301124625 | 170.66309164407653 | 165.7287343657557 | 199.46958650735505 | 258.40371584712443 | 246.40434735148972 | 197.19587406925208 | 189.19100507465575 |
| SE_119_1_40 | SE_119 | SE_119_1 | GGAGTGGCCCACAATCCCCT | 949.3885133189499 | 1094.3347706741163 | 973.7014865576605 | 918.7237936338787 | 880.2934455654076 | 816.064123496041 | 965.3160696498887 | 813.0040018783191 | 913.7368508820997 | 1506.8573549571872 | 890.9027882057281 | 1373.0856380571643 |
| SE_119_2_17 | SE_119 | SE_119_2 | TGGAAATTCAGACAAGAAGA | 35.76463577571386 | 32.97995199291857 | 47.40128940645811 | 33.834020728726415 | 27.575457330964575 | 12.128341538157722 | 12.913927353175769 | 27.415251226129364 | 63.93494000341222 | 8.019076393095819 | 38.73490383503166 | 32.49906835638259 |
| SE_119_2_18 | SE_119 | SE_119_2 | GAGCACAAGTGTGCTCCGGA | 3.2513305250648967 | 0.0 | 0.0 | 0.0 | 0.0 | 0.0 | 2.152321225529295 | 0.0 | 0.0 | 0.0 | 0.0 | 0.0 |
| SE_119_2_4 | SE_119 | SE_119_2 | CTAACAGCAGCTCATCCAGG | 1401.3234563029705 | 1592.0322280217965 | 1561.279969825214 | 1543.351868625751 | 1527.2560983303458 | 1720.4918781986598 | 1298.9258596069294 | 1358.472965929238 | 1485.599369801509 | 1451.4528271503432 | 1370.980838767181 | 1626.114098831857 |
| SE_119_2_5 | SE_119 | SE_119_2 | GCTGCTCACGCCTCACCCCG | 829.0892838915487 | 874.4684240546592 | 854.2107361788806 | 835.4400503016291 | 877.1116620272194 | 739.8288338276211 | 832.9483142798372 | 728.8675412188186 | 885.3213219916943 | 1256.807972881563 | 536.4197288669536 | 998.1856709460366 |
| SE_119_2_7 | SE_119 | SE_119_2 | GGAGGTGTTTCATCGCTGCA | 343.5572588151908 | 71.95625889364052 | 175.77978154894885 | 318.8205799437681 | 84.84756101835254 | 523.2513063605189 | 442.3020118462701 | 186.23463763956843 | 582.5183422533114 | 32.076305572383276 | 105.64064682281361 | 253.02846077469297 |
| SE_119_3_11 | SE_119 | SE_119_3 | TTTCCGAGGGAAGTTTGCAG | 28.17819788389577 | 29.981774539016886 | 32.58838646693995 | 66.36673296788642 | 57.27210368738796 | 51.11229648223612 | 23.675533480822246 | 29.305958207241734 | 26.639558334755094 | 82.37778476543888 | 15.259204541073078 | 2.321362025455899 |
| SE_119_3_12 | SE_119 | SE_119_3 | TGGGCCACCATGATGGCACG | 160.39897256986825 | 151.90765766435223 | 148.12902939518162 | 149.65047630013606 | 76.36280491651729 | 159.4010602157872 | 118.37766740411122 | 163.54615386622 | 167.8292175089571 | 96.95792366197672 | 97.42415206992811 | 105.6219721582434 |
| SE_119_3_18 | SE_119 | SE_119_3 | GCCCACAAGTGCTAGCCTTG | 922.2940922767424 | 954.4198228253708 | 941.1131000907204 | 1011.1166963930931 | 931.2019821764192 | 1021.3796195348539 | 871.6900963393645 | 814.8947088594315 | 1030.062922277197 | 987.8044102404396 | 937.8541867936453 | 869.3500785332342 |
| SE_119_3_21 | SE_119 | SE_119_3 | ATTCTCCGTTTCCTTTCAGA | 3.2513305250648967 | 2.9981774539016888 | 5.925161175807264 | 3.9039254686992013 | 4.242378050917627 | 0.0 | 1.0761606127646475 | 0.0 | 11.543808611727208 | 11.664111117230282 | 1.173784964697929 | 0.0 |
| SE_119_3_38 | SE_119 | SE_119_3 | AGAGCTCTCAGCTTTCATAA | 33.59708209233727 | 84.94836119388117 | 81.96472959866715 | 35.13532921829281 | 90.15053358199957 | 11.262031428289314 | 44.12258512335055 | 17.961716320567515 | 17.759705556503395 | 39.36637502065221 | 25.823269223354437 | 17.410215190919242 |
| SE_119_4_19 | SE_119 | SE_119_4 | GGAATTTTCACCCCCGCCAG | 9.75399157519469 | 7.9951398770711695 | 12.837849214249072 | 9.109159426964803 | 26.51486281823517 | 0.8663101098684087 | 17.21856980423436 | 103.98888396118035 | 0.0 | 2.187020834480678 | 43.43004369382337 | 12.767491140007444 |
| SE_119_4_28 | SE_119 | SE_119_4 | CTAGAACACAAAGGGCTGGA | 856.1837049337562 | 954.4198228253708 | 979.6266477334677 | 943.4486549356403 | 953.4744669437366 | 687.8502272355165 | 821.110547539426 | 848.9274345194541 | 881.7693808803936 | 774.9343823509869 | 551.6789334080266 | 662.7488582676591 |
| SE_119_4_35 | SE_119 | SE_119_4 | TACGAAAACATGGAAACCCG | 5788.452111457204 | 6878.818471735108 | 6945.276424908749 | 6217.6519631482615 | 6327.506862943641 | 6212.309797866359 | 5097.772822666135 | 5762.8748784305035 | 7455.524392620126 | 7070.6383578760315 | 5211.605243258805 | 6303.658580125493 |
| SE_119_4_36 | SE_119 | SE_119_4 | GCGCGCGTCGGAAGGAACGC | 7510.573512899911 | 6214.222469453566 | 5850.109134247039 | 6164.298315076039 | 5918.11738103009 | 5357.261719426239 | 7433.04135236542 | 5414.039440415271 | 6100.458858658916 | 5240.8309263605315 | 5313.724535187524 | 5558.50136995415 |
| SE_119_4_5 | SE_119 | SE_119_4 | GTAGAAGTTTGTTTTTGAGA | 115.96412206064798 | 220.86573910409106 | 264.6571991860578 | 196.49758192452646 | 240.75495438957532 | 291.08019691578534 | 147.4340039487567 | 183.3985771678999 | 212.22848140021557 | 555.5032919580922 | 213.62886357502308 | 331.95476964019355 |
| SE_119_5_0 | SE_119 | SE_119_5 | AAGAATACTTTACAACACCT | 55.27261892610324 | 89.94532361705066 | 138.25376076883617 | 65.06542447832003 | 136.81669214209347 | 97.02673230526177 | 43.0464245105859 | 29.305958207241734 | 114.5501008394469 | 87.48083337922712 | 19.954344399864794 | 62.67677468730927 |
| SE_119_5_13 | SE_119 | SE_119_5 | CCTTCAGTCGGAGGGTAGAG | 951.5560670023265 | 1132.3116850902043 | 1179.1070739856457 | 1073.5795038922804 | 1205.8959609733354 | 963.3368421736706 | 898.5941116584806 | 705.2337039549141 | 1190.7882575635526 | 985.6173894059589 | 596.282762066548 | 986.578860818757 |
| SE_119_5_22 | SE_119 | SE_119_5 | GGGGGCTGAGATTTGCCCAA | 281.78197883895774 | 257.8432610355452 | 265.64472604869235 | 268.0695488506785 | 257.72446659324584 | 158.5347501059188 | 306.70577463792455 | 257.1361494312823 | 284.1552889040543 | 344.82028490312024 | 331.007360044816 | 134.63899747644214 |
| SE_119_5_4 | SE_119 | SE_119_5 | GCATTACTCTTCCAAGCAAG | 747.8060207649263 | 618.6239479883817 | 573.7531071906701 | 685.7895740014931 | 653.3262198413146 | 564.8341916342025 | 739.3223409693128 | 860.2716764061283 | 602.9420036432903 | 744.3160906682574 | 1007.1074997108232 | 314.5445544492743 |
| SE_119_5_6 | SE_119 | SE_119_5 | GTATTCTTAGCAGGCTGTGG | 40.09974314246706 | 47.97083926242702 | 79.98967587339807 | 31.23140374959361 | 56.21150917465856 | 72.77004922894633 | 57.03651247652632 | 37.8141396222474 | 27.527543612580263 | 100.60295838611118 | 68.07952795247988 | 67.31949873822107 |
| SE_119_6_16 | SE_119 | SE_119_6 | CTGGACACTGTCCAAATCCG | 299.1224083059705 | 377.7703591916128 | 379.2103152516649 | 337.0388987976977 | 358.48094530253945 | 225.24062856578627 | 231.3745317443992 | 233.5023121673777 | 255.7397600136489 | 514.6789030477862 | 246.49484258656508 | 273.9207190037961 |
| SE_119_6_17 | SE_119 | SE_119_6 | AGGAGTAGAGAGAATTCTAA | 326.21682934817795 | 471.713252747199 | 512.5264417073283 | 485.38806660826737 | 461.35861303729195 | 558.7700208651237 | 374.5038932420973 | 471.7313917875363 | 507.03959363817194 | 770.5603406820255 | 400.2606729619938 | 652.3027291531076 |
| SE_119_6_18 | SE_119 | SE_119_6 | TGGAGCTTGCCCTTCCACGG | 574.4017260947984 | 699.5747392437273 | 719.9070828605826 | 657.1607872310323 | 696.8105948632202 | 952.9411208552496 | 612.3353886630844 | 643.785727068762 | 857.793778379114 | 628.4039864407815 | 673.7525697366112 | 733.5504000440641 |
| SE_119_6_25 | SE_119 | SE_119_6 | GGGCTCTTCAGTTCACCAGC | 13.005322100259587 | 9.993924846338961 | 16.78795666478725 | 13.013084895664004 | 44.54496953463509 | 0.8663101098684087 | 8.60928490211718 | 33.08737216946648 | 8.879852778251697 | 8.019076393095819 | 5.868924823489645 | 81.24767089095646 |
| SE_119_6_39 | SE_119 | SE_119_6 | GCTTTTGTGCACCCTACGAC | 293.703524097529 | 398.75760136892455 | 425.62407779548846 | 382.58469593252175 | 432.7225611935979 | 359.5186955953896 | 294.8680078975134 | 262.80827037461944 | 401.36934557697674 | 346.278298792774 | 142.02798072844942 | 443.3801468620767 |
| SE_119_7_1 | SE_119 | SE_119_7 | GAATAGATAAATGAGCACAA | 27.094421042207472 | 84.94836119388117 | 58.264084895438096 | 52.05233958265602 | 106.05945127294068 | 6.064170769078861 | 29.056336544645482 | 79.40969320671954 | 32.855455279531284 | 66.33963197924723 | 98.59793703462604 | 31.338387343654635 |
| SE_119_7_10 | SE_119 | SE_119_7 | AGCATCAGAGCACAAAACAG | 1426.2503236618013 | 1404.146440910624 | 1493.1406163034305 | 1449.65765737697 | 1612.1036593486983 | 1307.2619557914288 | 1474.340039487567 | 1281.899333194187 | 1443.864061743726 | 1656.3037786467 | 1558.7864331188498 | 1745.6642431428359 |
| SE_119_7_11 | SE_119 | SE_119_7 | CAGCATCAGAGCACAAAACA | 2632.493948460878 | 2501.4793890386422 | 2589.2954338277746 | 2335.8487387716887 | 2720.424925150928 | 3015.6254924519308 | 2587.0901130862126 | 2980.6995557236514 | 2926.7994757117594 | 2920.401820976532 | 2433.256231818807 | 3270.7990938673615 |
| SE_119_7_25 | SE_119 | SE_119_7 | CCTGGGACACCTCTTCCCCA | 3581.8824617798277 | 4298.387076410388 | 3989.608525043558 | 3436.7557209448637 | 3784.2012214185233 | 2730.6094663052245 | 3103.6472072132433 | 2180.9305027131186 | 3511.0937885207213 | 1740.8685842466195 | 2250.14577732593 | 1777.0026304864905 |
| SE_119_7_6 | SE_119 | SE_119_7 | TAGTGGCCAGACACCTTGAG | 3628.484865972425 | 3589.817804804955 | 3367.466601583795 | 3691.812184899878 | 3566.7793463089947 | 3636.76984122758 | 3722.4395595529154 | 4059.3478884482583 | 3505.7658768537704 | 4369.667627292394 | 3267.8173417190346 | 3479.7216761583923 |
| SE_12_1_11 | SE_12 | SE_12_1 | GTTGTTGGTAGTTATCAGAC | 70.44549470973944 | 93.94289355558624 | 74.06451469759081 | 83.28374333224963 | 96.51410065837601 | 19.9251325269734 | 59.18883370205561 | 60.50262339559584 | 54.167101947335354 | 11.664111117230282 | 90.38144228174053 | 47.58792152184593 |
| SE_12_1_12 | SE_12 | SE_12_1 | CATTGGACAGAGACCAGAAC | 1621.3301551656953 | 1694.969653939088 | 1661.020182951303 | 1514.72308185529 | 1739.375000876227 | 1774.203105010501 | 1622.8502040490885 | 1979.5702092246513 | 1482.9354139680336 | 2152.757508073814 | 2061.166398009563 | 1804.8589747919614 |
| SE_12_1_13 | SE_12 | SE_12_1 | GATGGAGACAGAAAATCCTG | 3820.3133669512536 | 4687.150752932973 | 4667.051952810855 | 4274.798388225626 | 4654.949316369366 | 4678.940903399275 | 4054.973188897192 | 5281.689951737406 | 4967.389644154 | 4477.560655126775 | 5622.42998090308 | 4075.1510356878302 |
| SE_12_1_30 | SE_12 | SE_12_1 | TTTCCTAGTAGGAAGAACAT | 13.005322100259587 | 3.9975699385355847 | 11.850322351614528 | 22.12224432262881 | 25.454268305505764 | 0.8663101098684087 | 18.294730416999005 | 0.0 | 4.439926389125849 | 0.0 | 0.0 | 0.0 |
| SE_12_1_8 | SE_12 | SE_12_1 | TCCAAAGAGGAGCCTCCCAA | 59.60772629285644 | 80.95079125534559 | 85.91483704920533 | 97.59813671748003 | 89.08993906927017 | 15.593581977631358 | 59.18883370205561 | 101.15282348951179 | 139.41368861855165 | 96.95792366197672 | 117.3784964697929 | 15.088853165463343 |
| SE_12_2_16 | SE_12 | SE_12_2 | GGTTGTTTAGTCTACAAGGA | 32.51330525064897 | 20.98724217731182 | 32.58838646693995 | 33.834020728726415 | 20.15129574185873 | 1.7326202197368175 | 12.913927353175769 | 1.89070698111237 | 31.07948472388094 | 0.0 | 17.606774470468935 | 0.0 |
| SE_12_2_5 | SE_12 | SE_12_2 | TTAGCGTAAAAATTAACTGG | 255.77133463843853 | 387.7642840379517 | 381.185368976934 | 331.8336648394321 | 352.11737822616305 | 369.0481068039421 | 247.51694093586892 | 298.7317030157545 | 301.91499446055775 | 197.5608820480879 | 323.9646502566284 3 | 259.9925468510607 |
| SE_12_2_6 | SE_12 | SE_12_2 | CAGGCTCCTAAACCTAAACG | 208.0851536041534 | 317.806810113579 | 331.8090258452068 | 157.45832723753446 | 120.90777445115236 | 116.95186483223517 | 69.95043982970209 | 69.0108048106015 | 205.12459917761421 | 6.561062503442034 | 31.692194046844083 | 13.928172152735392 |
| SE_12_2_7 | SE_12 | SE_12_2 | TTCTGTTTGCTGCTGCCACA | 1.083776841688299 | 3.9975699385355847 | 1.975053725269088 | 0.0 | 0.0 | 0.0 | 7.533124289352532 | 0.0 | 0.0 | 0.0 | 0.0 | 0.0 |
| SE_12_2_9 | SE_12 | SE_12_2 | AGGACTACACCTAAGCAGCG | 2596.729312685164 | 3456.898604348647 | 3260.8137004192645 | 2321.5343453864584 | 2560.2751537287877 | 1437.20847227169 | 1669.1251103979682 | 830.0203647083305 | 2560.9495412477895 | 621.8429239373394 | 613.8895365370169 | 530.4312228166729 |
| SE_12_3_1 | SE_12 | SE_12_3 | TCCTTGCGATGTCAGTAAGG | 457.3538271924621 | 401.75577882282624 | 454.26235681189024 | 352.6546006724945 | 461.35861303729195 | 346.5240439473635 | 453.06361797391656 | 446.2068475425193 | 404.9212866882774 | 430.11409744786664 | 224.19292825730446 | 300.6163822965389 |
| SE_12_3_12 | SE_12 | SE_12_3 | ATTTTGCTTAGCGTTGGCAA | 21.675536833765978 | 29.981774539016886 | 43.451181955919935 | 36.43663770785921 | 31.8178353818822 | 43.31550549342044 | 22.599372868057596 | 26.46989773557318 | 24.863587779104755 | 21.141201399979888 | 70.42709788187574 | 49.909283547301825 |
| SE_12_3_2 | SE_12 | SE_12_3 | TCCTCCTTACTGACATCGCA | 15.172875783636185 | 7.9951398770711695 | 16.78795666478725 | 33.834020728726415 | 3.1817835381882205 | 0.8663101098684087 | 24.75169409358689 | 24.57919075446081 | 1.7759705556503396 | 15.309145841364746 | 1.173784964697929 | 0.0 |
| SE_12_3_27 | SE_12 | SE_12_3 | GGCCCCAAACCACTGCCACA | 87.78592417675222 | 140.91434033337936 | 125.4159115545871 | 96.29682822791364 | 108.18064029839948 | 57.176467251314975 | 107.61606127646475 | 52.93979547114636 | 110.11017445032105 | 256.6104445790662 | 43.43004369382337 | 100.9792481073316 |
| SE_12_3_5 | SE_12 | SE_12_3 | TATTTGGAGGTAAAAAACAG | 26.010644200519174 | 42.973876839257535 | 54.31397744489992 | 67.66804145745283 | 42.42378050917627 | 32.919784174999535 | 34.43713960846872 | 13.23494886778659 | 28.415528890405433 | 13.122125006884067 | 37.56111887033373 | 23.21362025455899 |
| SE_12_4_1 | SE_12 | SE_12_4 | ACACTGAAGAGAAATAGCTG | 926.6291996434956 | 1241.245465915299 | 1184.0447082988183 | 992.8983775391636 | 1083.9275920094537 | 719.9037013006476 | 801.7396565096624 | 720.3593598038129 | 1269.8189472899928 | 669.9573822959144 | 731.2680330068098 | 793.9058127059175 |
| SE_12_4_2 | SE_12 | SE_12_4 | ACTCTGGTAACTTTTGGCCT | 148.47742731129696 | 218.86695413482326 | 210.3432217411579 | 192.59365645582727 | 178.17987813854035 | 119.5507951618404 | 162.50025252746178 | 185.28928414901227 | 275.2754361258026 | 185.16776398603074 | 84.5125174582509 | 175.26283292192036 |
| SE_12_4_20 | SE_12 | SE_12_4 | GGGCCATGTGAAATACCGAG | 448.68361245895574 | 550.6652590332768 | 600.4163324818028 | 422.9252591090801 | 470.9039636518566 | 241.70052065328605 | 317.467380765571 | 344.1086705624513 | 461.7523444690883 | 120.28614589643729 | 80.99116256415711 | 259.9925468510607 |
| SE_12_4_22 | SE_12 | SE_12_4 | TGGTGCACATCTGCAGAAAG | 0.0 | 0.9993924846338962 | 0.987526862634544 | 3.9039254686992013 | 2.1211890254588135 | 3.465240439473635 | 1.0761606127646475 | 0.0 | 1.7759705556503396 | 0.7290069448268927 | 3.521354894093787 | 0.0 |
| SE_12_4_25 | SE_12 | SE_12_4 | GCACCAACAACAACGTTTAG | 1364.4750436855684 | 1462.1112050193901 | 1461.539756699125 | 1570.6793469066454 | 1473.165778181146 | 1013.5828285460382 | 1465.7307545854499 | 1345.2380170614513 | 1199.6681103418043 | 2283.2497511978277 | 1427.3225170726816 | 1283.713200077112 |
| SE_12_5_1 | SE_12 | SE_12_5 | GCAATGCCTCTCACATGCAA | 1934.5416624136135 | 1525.0729315513256 | 1503.0158849297761 | 1817.9279599242614 | 1459.3780495156636 | 1096.7485990934053 | 1777.8173322871976 | 1688.4013341333464 | 1603.7014117522567 | 1447.8077924262088 | 1601.0426918479752 | 1198.9834861479717 |
| SE_12_5_17 | SE_12 | SE_12_5 | GCTCTGACCGACATCTGAAG | 267.69287989700985 | 335.7958748369891 | 317.9836497683232 | 356.55852614119374 | 336.20846053522195 | 188.8556039513131 | 290.56336544645484 | 322.3655402796591 | 342.76231724051553 | 293.0607918204108 | 385.00146842092073 | 213.5653063419427 |
| SE_12_5_19 | SE_12 | SE_12_5 | GCTCGACAAAAAGCGCCCCT | 283.9495325223343 | 433.73633833111097 | 369.3350466253195 | 347.4493667142289 | 409.389481913551 | 389.8395494407839 | 266.8878319656326 | 492.5291685797724 | 560.3187103076822 | 363.77446546861944 | 508.24888971420324 | 435.25537977298103 |

| | | | | | | | | | | | | | | | |
|---|---|---|---|---|---|---|---|---|---|---|---|---|---|---|---|
| SE_12_5_23 | SE_12 | SE_12_5 | ACAGAGCCCAGGAGTATCAG | 928.7967533268721 | 1117.320797820696 | 1086.2795488979984 | 925.2303360817107 | 1041.5038115002774 | 1311.5935063407708 | 957.7829453605362 | 1107.0089374412926 | 1259.1631239560907 | 1052.686028330033 | 1177.3063195920229 | 974.9720506914775 |
| SE_12_5_26 | SE_12 | SE_12_5 | GCTCTGCCAGTAGAACAGGG | 99.7074694353235 | 93.94289355558624 | 116.52816979087619 | 104.10467916531204 | 107.12004578567009 | 243.43314087302286 | 67.79811860417279 | 81.30040018783191 | 129.64585056247478 | 102.78997922059186 | 53.994108376104734 | 171.7807898837365 |
| SE_12_6_6 | SE_12 | SE_12_6 | GCACTATTAAGGTAAGGAAA | 370.6516798573982 | 489.7023174706091 | 529.3143983721156 | 337.0388987976977 | 509.0853661101152 | 583.8930140513074 | 334.68595056980536 | 468.89533131586774 | 538.1190783620528 | 367.4195001927539 | 237.10456286898167 | 579.1798253512468 |
| SE_120_1_11 | SE_120 | SE_120_1 | GTTAAAAGTAGTGCACACCT | 382.57322511596954 | 448.72722560061936 | 421.6739703449503 | 412.5147919254896 | 441.2073172954332 | 473.00531998815114 | 326.0766656676882 | 494.41987556088475 | 473.2961530808155 | 564.2513752960149 | 554.0265033374225 | 422.4878886329736 |
| SE_120_1_19 | SE_120 | SE_120_1 | CTACCCCCAAGGCTGATAAG | 82.36703996831072 | 96.94107100948793 | 112.57806234033802 | 72.87327541571842 | 125.15015250207 | 94.42780197565655 | 102.23525821264151 | 136.13090264009065 | 75.47874861513944 | 98.4159375516305 | 316.9219404684408 | 54.55200759821362 |
| SE_120_1_3 | SE_120 | SE_120_1 | ATACGGCTACTGAGGACATG | 2821.0711189146423 | 2813.289844244418 | 2703.8485498933815 | 2734.049136579007 | 2697.0918458708816 | 2899.539937729564 | 2689.325371298854 | 2626.191996765082 | 2658.6279218085583 | 2828.5469459283436 | 2474.3387055832345 | 2598.764787497879 |
| SE_120_1_4 | SE_120 | SE_120_1 | TTGAAAACATGCTAACAAGG | 210.25270728753 | 166.89854493386065 | 178.74236213685248 | 119.72038104010883 | 206.8159299822343 | 79.7005301078936 | 123.75847046793446 | 190.96140509234937 | 245.08393667974687 | 24.057229179287457 | 65.73195802308402 | 46.42724050911798 |
| SE_120_1_7 | SE_120 | SE_120_1 | ACATTCACCATAGCCAGCCG | 2559.880900067762 | 2591.424712655693 | 2627.8089814705218 | 2769.1844657973 | 2442.549162815824 | 2432.598788510492 | 2774.3420597072613 | 2682.913206198453 | 3038.685620717731 | 3569.2180018724666 | 2555.3298681473916 | 2822.776222954373 |
| SE_120_2_14 | SE_120 | SE_120_2 | TCACAGGTGTGCTAGTCAAG | 784.6544333823284 | 740.5498311137171 | 753.4829961901571 | 869.2740710303555 | 659.689786917691 | 775.3475483322258 | 882.4517024670109 | 605.9715874465146 | 713.0521780936114 | 733.380986495854 | 669.0574298778196 | 986.578860818757 |
| SE_120_2_21 | SE_120 | SE_120_2 | CATGGTATAGCAGGACCGTG | 4279.8347478270925 | 4055.5347026443505 | 3973.808095241405 | 3954.676499792291 | 3968.74466663344 | 3799.6361418828405 | 4122.771307501364 | 3324.8082262861026 | 3809.4568418699782 | 3856.446738134262 | 3506.0956895527142 | 3610.8786305966505 |
| SE_120_2_6 | SE_120 | SE_120_2 | GGTCACACTAGAGTTCCATA | 204.8338230790885 | 264.8390084279825 | 284.40773643874866 | 209.51066682019047 | 255.60327756778702 | 335.2620125190742 | 306.70577463792455 | 143.69373056454012 | 370.2898608530958 | 602.1597364270133 | 147.89690555193906 | 316.86591647473017 |
| SE_120_2_7 | SE_120 | SE_120_2 | GATAATCATATCCAAGTAAA | 18.42420630870108 | 20.98724217731182 | 30.613332741670867 | 23.423552812195208 | 22.272484767317543 | 50.2459863723677 | 17.21856980423436 | 14.180302358342775 | 55.94307250298569 | 104.97700005507254 | 0.0 | 18.57089620364719 |
| SE_120_2_8 | SE_120 | SE_120_2 | AATGCCCTCCTAAATTTGGG | 128.96944416090756 | 157.9040125721556 | 153.06666370835433 | 145.74655083143685 | 112.42301834931712 | 133.41175691973496 | 109.76838250199404 | 121.95060028174787 | 156.28540889722987 | 263.90051402733513 | 65.73195802308402 | 92.85448101823596 |
| SE_120_3_0 | SE_120 | SE_120_3 | GAAGCTTTAAAAGGTGTAAG | 82.36703996831072 | 139.91494784874547 | 117.51569665351074 | 81.98243484268323 | 115.60480188750533 | 116.08555472236677 | 93.62597331052433 | 71.84686528227006 | 124.31793889552377 | 72.90069448268926 | 91.55522724643846 | 102.13992912005955 |
| SE_120_3_14 | SE_120 | SE_120_3 | GGTATCAGTTCCCTCCCTCA | 115.96412206064798 | 129.9210230024065 | 136.27870704356707 | 114.51514708184324 | 82.72637199289373 | 86.63101098684088 | 93.62597331052433 | 74.68292575393862 | 104.78226278337003 | 85.29381254474644 | 151.41826044603283 | 225.1721164692222 |
| SE_120_3_19 | SE_120 | SE_120_3 | TTTCTGAGTCAAATACCGTT | 2.167553683376598 | 8.994532361705065 | 12.837849214249072 | 1.3013084895664004 | 20.15129574185873 | 0.0 | 0.0 | 1.89070698111237 | 0.0 | 0.7290069448268927 | 0.0 | 0.0 |
| SE_120_3_20 | SE_120 | SE_120_3 | CTAGTGTGCAGGGTGCTTCG | 1294.029548975829 | 1371.1664889177055 | 1458.5771761112214 | 1285.6927876916036 | 1462.559833053852 | 1156.5239966743256 | 1103.0646280837636 | 1278.117919231962 | 1334.6418725712301 | 1250.246910378121 | 946.0706815465307 | 1528.6168937627094 |
| SE_120_3_6 | SE_120 | SE_120_3 | ATATTGCATCCCGATCACAT | 267.69287989700985 | 293.8213904823655 | 290.3328976145559 | 283.68525072547527 | 356.3597562770807 | 195.78608483026036 | 212.00364071463557 | 224.04877726181584 | 307.2429061275087 | 569.3544239098031 | 326.31222018602426 | 133.47831646371418 |
| SE_120_4_1 | SE_120 | SE_120_4 | GCTGCCTCCCTTCTATGCTG | 1473.9365046960866 | 1246.2424283384685 | 1295.6352437765217 | 1565.4741129483798 | 1259.9862811225353 | 1360.97318260327 | 1508.7771790960358 | 1356.5822589481254 | 1486.4873550793343 | 1113.1936047506651 | 1523.5728841779119 | 1398.6206203371792 |
| SE_120_4_17 | SE_120 | SE_120_4 | CCCTTGTTCCTATCCAACAG | 82.36703996831072 | 139.91494784874547 | 149.11655625781614 | 87.18766880094883 | 108.18064029839948 | 296.2780575749958 | 80.71204595734856 | 104.93423745173654 | 180.26101139850945 | 209.22499316531818 | 119.72606639918877 | 213.5653063419427 |
| SE_120_4_19 | SE_120 | SE_120_4 | GCCCCTTGTTCCTATCCAAC | 23.843090517142578 | 33.97934447755247 | 24.688171565863602 | 20.820935833062407 | 59.39329271284678 | 29.454543735525895 | 19.370891029763655 | 94.5353490556185 | 17.759705556503395 | 5.832055558615141 | 2.347569929395858 | 9.285448101823595 |
| SE_120_4_2 | SE_120 | SE_120_4 | GTGTTTTGAATGCAGAGTGA | 139.80721257779055 | 188.8851795958064 | 182.69246958739063 | 130.13084895664005 | 161.21036593486983 | 204.44918592894447 | 160.34793130193248 | 152.2019119795458 | 144.74160028550267 | 63.42360419993966 | 58.68924823489645 | 170.62010887100857 |
| SE_120_4_34 | SE_120 | SE_120_4 | GCCCTGAATGACCAACCCTA | 817.1677386329774 | 741.549223598351 | 747.5578350143498 | 839.3439757703283 | 894.0811742308899 | 650.598892511175 | 870.6139357265998 | 876.3426857455835 | 700.620384204059 | 1126.3157297575492 | 698.4020539952678 | 1002.8283949969483 |
| SE_121_1_2 | SE_121 | SE_121_1 | GCTGGGATAAGATTACTTTG | 1219.2489468993363 | 1486.0966246506036 | 1437.839111995896 | 1335.1425102951268 | 1415.893674493758 | 1503.048040621689 | 1183.7766740411123 | 1295.1342820619734 | 1594.821558974005 | 1443.4337507572475 | 1198.4344489565856 | 943.6336633478229 |
| SE_121_1_22 | SE_121 | SE_121_1 | ACACAGTTCATAATGATGGT | 754.308681815056 | 867.4726766622219 | 893.7118106842623 | 663.6673296788642 | 768.9310217288199 | 812.5988830565674 | 647.8486888843178 | 888.6322811228139 | 972.3438792185609 | 713.6977989855279 | 788.7834962770083 | 1038.8095063915148 |
| SE_121_1_29 | SE_121 | SE_121_1 | TCTGCAGAAAACAGATACGG | 49.85373471766175 | 33.97934447755247 | 53.32645058226538 | 65.06542447832003 | 57.27210368738796 | 139.4759276888138 | 110.8445431147587 | 18.9070698111237 | 156.28540889722987 | 3.6450347241344634 | 38.73490383503166 | 42.945197470934126 |
| SE_121_1_3 | SE_121 | SE_121_1 | GCGGCACAGGTGGAATCACG | 5574.948073644609 | 5038.936907524105 | 5045.274741199885 | 5327.556956284843 | 5116.307929406658 | 5487.208235906501 | 5533.6178708358175 | 4965.94188589164 | 4597.987768578729 | 6060.963739290785 | 5098.921886647804 | 5635.106316794195 |
| SE_121_1_4 | SE_121 | SE_121_1 | CTAATGAAAAGGGTGCAGTA | 247.10111990493215 | 397.7582088842907 | 333.7840795704759 | 264.1656233819793 | 351.0567837134336 | 325.73260131052166 | 273.34479564222045 | 419.73694980694614 | 443.1046536347597 | 467.2934516340382 | 551.6789334080266 | 193.83372912556754 |
| SE_121_2_10 | SE_121 | SE_121_2 | GCACAGCGGATGCTGCCAGG | 3774.794739600345 | 3742.7248549539413 | 3750.6270242859982 | 3988.5105205210175 | 3787.3830049567114 | 3465.240439473635 | 4184.11246242895 | 3407.999333455047 | 3526.189538243749 | 3523.2905643483723 | 4104.726021548658 | 2568.587081166952 |
| SE_121_2_17 | SE_121 | SE_121_2 | AGCCCAAGAGGTGAAAACAA | 259.02266516350346 | 269.83597085115196 | 276.50752153767235 | 230.33160265325287 | 253.4820885423282 | 504.1924839434139 | 257.20238645075074 | 294.95028905352973 | 239.75602501279585 | 280.66767375835366 | 267.6229719511278 | 377.22132913658356 |
| SE_121_2_2 | SE_121 | SE_121_2 | GGAGGGTCCGTGTTGCCCCA | 4098.8440152651465 | 3820.6774687553852 | 3654.8369186104474 | 3949.4712658340254 | 3768.2923037275823 | 3691.3473781492894 | 4417.639315398877 | 3484.5729661900978 | 3848.528194094286 | 3486.1112101622007 | 3619.952831128413 | 3935.8693141604767 |
| SE_121_2_24 | SE_121 | SE_121_2 | ACAGTGGAAGGAAAGAACCC | 1710.1998561841356 | 1848.8760965727079 | 1813.0993197970229 | 1901.211703256511 | 2078.7652449496372 | 2037.5613784104974 | 1880.0525904998392 | 2015.4936418657865 | 1805.27406981857 | 2791.367591742172 | 1998.9557948805732 | 1768.877863397395 |
| SE_121_2_8 | SE_121 | SE_121_2 | GGCTAATGTCACATGCACAG | 1315.705085809595 | 1339.1859294094209 | 1281.809867699638 | 1379.3869989403845 | 1396.8029732646287 | 1179.9143696407727 | 1422.684330074864 | 1349.9647845142322 | 1116.1974942262384 | 1457.2848827089583 | 1119.7908563218243 | 1104.9683241170078 |
| SE_121_3_11 | SE_121 | SE_121_3 | CCTCGGGTGCCACAGACACA | 1971.3900750310156 | 1828.88824688003 | 1763.7229766652956 | 1972.7836701826632 | 1829.5255344582267 | 1405.154998206559 | 2213.66238045688 | 2097.7393955441744 | 1749.3309973155845 | 1834.9104801292888 | 1940.2665466456767 | 2053.2447115157424 |
| SE_121_3_12 | SE_121 | SE_121_3 | GCAATAGGTAGAGATGTTGA | 1335.2130689599842 | 1480.1002697428003 | 1492.153089440796 | 1483.4916781056966 | 1473.165778181146 | 1200.7058122776145 | 1400.0849572068064 | 1784.8273901700773 | 1680.9561309230464 | 1781.6929731569257 | 1784.1531463408521 | 1486.8323773045033 |
| SE_121_3_15 | SE_121 | SE_121_3 | TGAGGGGAGGGGACTTCAGA | 2594.561759001788 | 2974.1920342704752 | 2946.780158101479 | 2843.359049702585 | 2713.0007635618226 | 2504.5025276295696 | 2452.5700364906315 | 2781.2299692162965 | 2664.8438187533347 | 3361.451022596802 | 2471.9911356538387 | 2214.5793722849276 |
| SE_121_3_22 | SE_121 | SE_121_3 | AGTCCACTGTAGCAAAGGGA | 202.6662693957119 | 315.8080251443112 | 259.7195648728851 | 234.2355281219521 | 240.75495438957532 | 243.43314087302286 | 176.4903404934022 | 291.168875091305 | 275.2754361258026 | 209.95400011014507 | 315.7481555037429 | 256.51050381287683 |
| SE_121_3_3 | SE_121 | SE_121_3 | GGGCAGGCAGATATGCTGCT | 2074.348874991404 | 1766.9259128327285 | 1704.471364907223 | 2115.9276040349673 | 2078.7652449496372 | 1859.1014957776051 | 2498.8449428395115 | 2056.1438419597025 | 1888.7446859341362 | 1285.9682506746387 | 2135.114850785533 | 1483.3503342663194 |
| SE_121_4_19 | SE_121 | SE_121_4 | GAGCCTCACATGGAACCCAG | 374.98678722415144 | 391.7618539764873 | 362.42235858687764 | 343.54544124552973 | 419.9954270408451 | 462.60959866973025 | 388.4939812080377 | 432.97189867473276 | 383.60964002047336 | 344.09127795829335 | 388.52282331501453 | 821.7621570113882 |
| SE_121_4_23 | SE_121 | SE_121_4 | TGTGAGGCTCCGCCACCAGG | 42.26729682584366 | 85.94775367851507 | 84.92731018657079 | 70.27065843658562 | 110.3018293238583 | 174.12833208355016 | 57.03651247652632 | 159.76473990399526 | 36.40739639083196 | 162.56854869639707 | 230.0618530807941 | 34.820430381838484 |
| SE_121_4_24 | SE_121 | SE_121_4 | TCTAGGGTTGACGGAACAAA | 959.1425048941445 | 872.4696390853913 | 884.8240689205514 | 851.0557521764259 | 791.2035064961374 | 857.6470087697246 | 812.5012626373089 | 841.3646065950046 | 668.6529142023528 | 350.6523404617354 | 812.2591955709669 | 782.2990025786379 |
| SE_121_4_27 | SE_121 | SE_121_4 | TGTTCCGTCAACCCTAGAGG | 120.29922942740117 | 165.89915244922676 | 166.89203978523796 | 126.22692348794084 | 230.14900926228125 | 63.24063802039384 | 95.77829453605362 | 214.595242356254 | 224.66027528976795 | 148.7174167446861 | 123.24742129328254 | 127.67491140007444 |
| SE_121_4_3 | SE_121 | SE_121_4 | AGCTCAAGTGATCCACCCGC | 1376.3965889441397 | 1490.0941945891393 | 1595.8434100174231 | 1307.8150320142324 | 1317.2583848099232 | 1391.2940364486644 | 1143.9587313688203 | 1156.1673189502142 | 1243.1793889552378 | 723.9038962131044 | 1246.5596325092006 | 1267.4636658989207 |
| SE_122_1_11 | SE_122 | SE_122_1 | TTTTCTTTAACGTCCTCTGG | 0.0 | 4.996962423169481 | 1.975053725269088 | 0.0 | 0.0 | 0.0 | 0.0 | 0.0 | 0.0 | 0.0 | 0.0 | 0.0 |
| SE_122_1_14 | SE_122 | SE_122_1 | GTTTCCATAGGGAGAGACAG | 26.010644200519174 | 42.973876839257535 | 35.55096705484358 | 14.314393385230405 | 40.30259148371746 | 56.31015714144657 | 22.599372868057596 | 37.8141396222474 | 83.47061611556596 | 72.90069448268926 | 53.994108376104734 | 39.46315443275028 |
| SE_122_1_17 | SE_122 | SE_122_1 | TTCAGAGGCCTGCAAAACCA | 37.93218945909046 | 54.96658665486429 | 62.21419234597627 | 58.55888203048802 | 42.42378050917627 | 31.187163955262715 | 38.74178205952731 | 68.06545132004533 | 88.79852778251698 | 56.13353475167074 | 35.21354894093787 | 124.19286836189059 |
| SE_122_1_18 | SE_122 | SE_122_1 | AGAGGCCTGCAAAACCACGG | 891.94834070947 | 965.4131401563437 | 877.9113808821096 | 988.9944520704644 | 1073.3216468821597 | 1146.994585465773 | 970.696872713712 | 1080.5390397057195 | 907.5209539373235 | 1314.3995215228874 | 1447.2768614725464 | 1299.9627342553033 |
| SE_122_1_19 | SE_122 | SE_122_1 | GAAGTCAGAGTTGTAAAGGA | 246.01734306324386 | 368.7758268299077 | 359.459777998974 | 338.3402072872641 | 383.9352136080452 | 317.935810321706 | 293.79184728474877 | 331.8190751852209 | 487.5039175260182 | 456.3583474616348 | 232.40942301018995 | 153.20989368008932 |
| SE_123_1_0 | SE_123 | SE_123_1 | TGAGATGGCTCAGAGCAAGG | 2878.511291524122 | 2643.3931218566554 | 2873.703170266523 | 2540.1541716336137 | 2763.909300172834 | 2495.8394265308857 | 2584.9377918606833 | 2586.487150161722 | 2552.069688469538 | 1800.647153722425 | 2275.9690465492845 | 2711.3508457324897 |
| SE_123_1_10 | SE_123 | SE_123_1 | GGGCTGATACCAGGAGAAGA | 238.43090517142576 | 314.8086326596773 | 319.9587034935923 | 286.2878677046081 | 334.0872715097631 | 597.7539758092021 | 283.0302411571023 | 301.567763487423 | 356.97008168571824 | 524.8850002753627 | 186.63180938697073 | 289.0095721692594 |
| SE_123_1_19 | SE_123 | SE_123_1 | AGTTCTTTATGGACTGTAGG | 0.0 | 7.9951398770711695 | 13.825376076883616 | 10.410467916531204 | 14.848323178211695 | 0.8663101098684087 | 6.456963676587884 | 0.0 | 0.8879852778251698 | 0.0 | 1.173784964697929 | 0.0 |
| SE_123_1_2 | SE_123 | SE_123_1 | GGCCACCTCTCTGATCCCCA | 686.0307407886933 | 860.4769292697846 | 803.8468661845188 | 713.1170522823875 | 901.5053358199957 | 644.534721742096 | 694.1235952331976 | 888.6322811228139 | 1036.2788192219732 | 691.0985836958943 | 1022.3667042518962 | 543.1987139566803 |
| SE_123_2_1 | SE_123 | SE_123_2 | GGCAGTGTCATAACTAAGCG | 1041.5095448624552 | 978.4052424565843 | 988.5143894971786 | 880.9858474364531 | 1010.7465706311247 | 852.4491481105142 | 958.8591059733009 | 1094.7193420640622 | 799.1867500426528 | 936.7739241025571 | 746.5272375478829 | 971.4900076532937 |
| SE_123_2_16 | SE_123 | SE_123_2 | GTTCCTGAGCTGCTGAGTGG | 109.46146101051819 | 148.90948021045054 | 121.46580410404891 | 118.41907255054244 | 139.9984756802817 | 85.76470087697247 | 137.74855843387488 | 105.87959094229272 | 191.80482001023668 | 207.0379723308375 | 117.3784964697929 | 125.35354937461854 |
| SE_123_2_18 | SE_123 | SE_123_2 | ATTCATTGTATTCATAACAG | 7.586437891818092 | 5.9963549078033775 | 8.887741763710896 | 6.506542447832002 | 3.1817835381882205 | 0.0 | 0.0 | 0.0 | 0.0 | 0.0 | 0.0 | 0.0 |
| SE_123_2_3 | SE_123 | SE_123_2 | GAGGCAGTGTCATAACTAAG | 8223.698674730813 | 7576.394426009567 | 7809.362429713974 | 7992.636742916831 | 7913.095659474104 | 6714.769661590036 | 8000.177995292389 | 7040.992797662466 | 7888.8612081988085 | 6944.520156420979 | 7241.079447221524 | 6909.534068769483 |
| SE_123_2_4 | SE_123 | SE_123_2 | TTATGACACTGCCTCCTAAG | 230.84446727960767 | 266.83779339725027 | 328.8464452573032 | 243.34468754891688 | 252.42149402959882 | 166.33154109473446 | 231.3745317443992 | 200.41493999791123 | 285.9312594597047 | 165.48457647570464 | 72.7746678112716 | 229.814840520134 |
| SE_123_3_10 | SE_123 | SE_123_3 | GAGATAGTGACCACACCATG | 3558.039371262685 | 3278.0073495991796 | 3135.3977888646773 | 3417.2360936013674 | 3198.7530503918906 | 3923.518487594023 | 3534.111452319102 | 3217.9832818532536 | 3448.9348190729593 | 2679.1005222388303 | 3202.0853836959504 | 3148.9275875309268 |
| SE_123_3_12 | SE_123 | SE_123_3 | CAGAGCTTGAAGGTACCCCA | 1063.1850816962212 | 1126.3153301824009 | 1022.090302826753 | 1154.2606302453971 | 1027.716082834795 | 1451.935744139453 | 1107.3692705348224 | 1296.0796355525297 | 998.983437553316 | 1139.4378547644333 | 1386.2400433082541 | 1533.2596178136212 |
| SE_123_3_22 | SE_123 | SE_123_3 | GGTTCAACATGAATTGACCC | 44.434850509220254 | 54.96658665486429 | 61.22666548334173 | 48.1484141395682 | 41.363185996446866 | 15.593581977631358 | 65.6457973786435 | 56.7212094333711 | 39.95933750213264 | 145.07238202055163 | 30.518409082146157 | 31.338387343654635 |
| SE_123_3_27 | SE_123 | SE_123_3 | GCTGGACTCTACCTCTTGAT | 5909.835117726294 | 5881.424772070479 | 5859.984402873384 | 5786.918853101783 | 5749.482853506114 | 6968.59852378148 | 5417.392524657235 | 5686.301245695453 | 6214.120974220538 | 4714.487912195515 | 5798.497725607769 | 5454.0400788086345 |
| SE_123_3_29 | SE_123 | SE_123_3 | CTGCTGCTCCTCCCATCAAG | 312.12773040623006 | 395.75942391502286 | 344.64687505945585 | 426.82918457777936 | 405.1471038626334 | 359.5186955953896 | 372.351572016568 | 299.67705650631063 | 233.54012806801964 | 271.91959042043095 | 247.66862755126303 | 370.25724306021584 |
| SE_123_4_11 | SE_123 | SE_123_4 | ATAAATAGCATAGGCTCCTA | 1.083776841688299 | 7.9951398770711695 | 12.837849214249072 | 5.205233958265602 | 18.030106716399914 | 6.93048087894727 | 13.990087965940416 | 2.836060471668555 | 11.543808611727208 | 5.103048613788249 | 0.0 | 2.321362025455899 |
| SE_123_4_12 | SE_123 | SE_123_4 | AGATGCTCAGTAATCACACG | 1865.1799445455624 | 1951.8135224899993 | 1906.9143717473046 | 2032.6438607027176 | 1652.4062508324157 | 1978.6522909394455 | 2222.271665358997 | 1687.4559806427903 | 2013.0626248296599 | 2326.2611609426144 | 1542.3534436130788 | 1767.717182384667 |
| SE_123_4_14 | SE_123 | SE_123_4 | TGACTGCAACAGGCAAGCAG | 4115.1006678904705 | 3662.7734561832294 | 3638.0489619456603 | 4334.65857874568 | 3944.350992840664 | 3745.058604961131 | 4280.966917577768 | 4197.369498069462 | 3848.528194094286 | 4242.091411947688 | 4790.216440932249 | 3595.7897774311873 |
| SE_123_4_17 | SE_123 | SE_123_4 | GGCATTGGTGACCCAGAATA | 174.48807151181612 | 243.85176625067066 | 210.3432217411579 | 192.59365645582727 | 292.72408551331625 | 202.71656570920763 | 214.15596194016484 | 189.070698111237 | 238.86803973497067 | 301.07986821350664 | 214.80264853972102 | 311.06251141109044 |
| SE_123_4_5 | SE_123 | SE_123_4 | GAGCATCTGAAACGTGGCTG | 1882.5203740125753 | 2038.7606686531483 | 2080.7190995709843 | 1950.6614258600343 | 2150.885671815237 | 2244.609494669047 | 1936.0129423636008 | 2227.252823750372 | 2148.924372336911 | 2234.4062858944258 | 2028.3004189980213 | 3173.301888798214 |
| SE_123_5_0 | SE_123 | SE_123_5 | TGCCAAGGATAAGCACCCAC | 590.6583787201229 | 544.6689041254734 | 596.4662250312646 | 592.0953627527122 | 534.539634415621 | 641.9357914124909 | 674.752704203434 | 363.9610938641312 | 562.9826661411577 | 560.6063405718804 | 494.16347013782814 | 321.50864052564197 |
| SE_123_5_14 | SE_123 | SE_123_5 | GCTCCGACCTGCCGTCTGGC | 395.5785472162291 | 330.79891241381966 | 359.459777998974 | 348.7506752037953 | 289.54230197512805 | 630.6737599842015 | 443.3781724590348 | 446.2068475425193 | 469.7442119695148 | 208.4959862204913 | 260.58026216294024 | 646.4993240894678 |
| SE_123_5_21 | SE_123 | SE_123_5 | ACCCATGATTATTCACCACC | 625.3392376541485 | 665.5953947661749 | 685.3436426683736 | 671.4751806162626 | 718.0224851178084 | 956.4063612947232 | 516.5570941270308 | 486.85704763643525 | 824.0503378217576 | 263.17150708250824 | 643.2341606544651 | 827.565562075028 |
| SE_123_5_28 | SE_123 | SE_123_5 | ATCCAGCAAGTCCTAAAAGT | 151.72875783636184 | 202.87667438068092 | 202.44300684008152 | 188.68973098712806 | 207.87652449496372 | 330.9304619697321 | 178.6426617189315 | 156.9286794323267 | 278.8273772371033 | 152.36245146882055 | 122.07363632858463 | 428.29129369661337 |
| SE_123_5_9 | SE_123 | SE_123_5 | ACTGCAACCTCAAAACTCTT | 1257.1811363584268 | 1465.1093824732918 | 1423.0262090563779 | 1512.1204648761573 | 1383.0152445991464 | 1184.2459201901147 | 1468.9592364237437 | 1390.6149846081482 | 1299.1224614582234 | 1312.9415076332336 | 1321.681870249868 | 985.4181798060291 |
| SE_124_1_29 | SE_124 | SE_124_1 | AAGCTCTGTAGCACTGTGAC | 322.9654988231131 | 343.7910147140603 | 367.3599929000504 | 296.6983356211393 | 397.7229422735275 | 301.47591823420623 | 239.98381664651637 | 192.85211207346174 | 388.9375516874244 | 193.91584732395344 | 326.31222018602426 | 228.65415950740604 |
| SE_124_1_30 | SE_124 | SE_124_1 | GCAGGAAGTTGACTGGACCA | 9291.218863793787 | 9025.513528728716 | 8914.404989002029 | 9357.709348471986 | 8937.62995877071 | 9460.106399763023 | 9505.726692550132 | 8224.57536783881 | 9315.853549663856 | 10092.372144183502 | 8605.017576200518 | 9302.858317014514 |
| SE_124_1_31 | SE_124 | SE_124_1 | AGCTCTGTAGCACTGTGACT | 299.1224083059705 | 362.77947192210434 | 389.08558387801037 | 290.1917931733073 | 364.8445123789159 | 499.8609333940718 | 177.56650110616684 | 248.62796801627667 | 260.17968640277473 | 217.244069558414 | 227.71428315139823 | 242.58233166014142 |
| SE_124_1_32 | SE_124 | SE_124_1 | GCTCTGTAGCACTGTGACTG | 744.5546902398613 | 906.4489835629438 | 880.8739614700132 | 834.1387418120627 | 726.5072412196437 | 879.3047615164348 | 658.6102950119642 | 778.0259227277403 | 687.3006050366814 | 914.9037157577503 | 907.3357777114992 | 604.7148076312617 |
| SE_124_1_33 | SE_124 | SE_124_1 | CTCTGTAGCACTGTGACTGG | 128.96944416090756 | 203.8760668653148 | 181.7049427247561 | 153.55440176883525 | 253.4820885423282 | 319.66843054144283 | 112.99686434028798 | 246.7372610351643 | 150.95749723027888 | 66.33963197924723 | 435.4742219029317 | 315.70523546200224 |
| SE_124_2_13 | SE_124 | SE_124_2 | GGAACAGAATGGAAGAGCTG | 770.5653344403805 | 957.4180002792725 | 846.3105212778042 | 805.5099550416019 | 911.0506864345604 | 676.5881958072272 | 719.9514499395492 | 903.7579369717129 | 851.5778814343378 | 1083.3043200127624 | 1333.4197198968475 | 766.0494684004466 |
| SE_124_2_21 | SE_124 | SE_124_2 | ACAGACTGTTGAGGAACAGG | 1757.8860372184208 | 1841.8803491802707 | 1901.9767374341318 | 1719.0285147172149 | 1823.1619673818502 | 1779.4009656697115 | 1786.4266171893148 | 1778.209915736184 | 1941.135817325821 | 2767.3103625628846 | 1912.0957074929263 | 2171.6341748139935 |
| SE_124_2_3 | SE_124 | SE_124_2 | CTATCCATGGTGACTCCCCT | 256.8551114801268 | 314.8086326596773 | 273.5449409497687 | 313.61534598550253 | 250.30030500414 | 133.41175691973496 | 285.1825623826316 | 399.88452650526625 | 324.11462640618697 | 289.41575709627637 | 369.74226387984766 | 450.3442329384444 |
| SE_124_2_5 | SE_124 | SE_124_2 | TCATTTCCCCGATAACCTCA | 826.9217302081721 | 777.5273530451713 | 842.360413827266 | 866.6714540512227 | 790.142911983408 | 438.3529155934148 | 660.7626162374936 | 964.2605603673087 | 673.0928405914786 | 818.6747990406004 | 1279.4256115207427 | 916.93800005508 |

| | | | | | | | | | | | | | | | |
|---|---|---|---|---|---|---|---|---|---|---|---|---|---|---|---|
| SE_124_2_7 | SE_124 | SE_124_2 | GGGAAATGAAGGTTGAAACA | 338.13837460674927 | 395.75942391502286 | 353.53461682316674 | 321.4231969229009 | 493.17644841917416 | 528.4491670197293 | 372.351572016568 | 416.9008893352776 | 418.241065855655 | 278.480652923873 | 340.3976397623994 | 515.3423696512095 |
| SE_124_4_10 | SE_124 | SE_124_4 | GAAGAGGAGGAGTGTCTCAA | 3858.2455564103443 | 4181.458155708222 | 4147.612823065085 | 3807.628640471288 | 4163.8940569756505 | 4467.561236591384 | 3691.230901782741 | 4012.0802139204493 | 4500.30938801796 | 5065.869259602077 | 4092.9881719016785 | 4549.869569893562 |
| SE_124_4_12 | SE_124 | SE_124_4 | ATGAGGCATTAAAGAAGAGG | 9097.222809131581 | 11038.289992781383 | 11054.375700331086 | 9831.385638674155 | 10498.825081508397 | 10361.935224136038 | 8777.165957708465 | 11257.26936554305 | 10540.385247784765 | 11024.04301967227 | 11284.768650605889 | 10632.998757600744 |
| SE_124_4_4 | SE_124 | SE_124_4 | GGTGATGCAAGGTAGGTGTG | 3611.144436505412 | 3220.0425854904133 | 3303.27735551255 | 3348.2667436543484 | 3042.845657020668 | 2585.0693678473317 | 3697.687865459329 | 3390.9829706250357 | 2714.570994311544 | 2999.8635779626634 | 3556.568443034725 | 2881.9709546034983 |
| SE_124_4_6 | SE_124 | SE_124_4 | GGTGAGCTGGAGGTGATGCA | 5927.175547193307 | 5835.45271777732 | 5842.208919345962 | 5518.8493042511045 | 5695.392533356914 | 5871.849924688075 | 5714.412853780278 | 5634.306803714862 | 5846.495069200918 | 5611.89546127742 | 4497.943984722464 | 4343.268349627987 |
| SE_124_4_9 | SE_124 | SE_124_4 | GAGGAGGAGTGTCTCAAGGG | 4754.529004486567 | 6353.138024817678 | 6424.849768300343 | 5179.207788474274 | 6576.746573435052 | 5480.277755027553 | 4866.398290921736 | 6416.114140404828 | 6142.194166716699 | 8109.473254254354 | 6750.43733197779 | 5557.340688941422 |
| SE_125_1_14 | SE_125 | SE_125_1 | GGACTTGGTGATGTATTAGG | 537.5533134773963 | 666.5947872508087 | 679.4184814925663 | 468.4710562439042 | 591.811738103009 | 668.7914048184116 | 469.2060271653863 | 606.9169409370708 | 494.6077997486196 | 716.6138267648355 | 685.4904193835905 | 401.5956304038705З |
| SE_125_1_19 | SE_125 | SE_125_1 | GGCAGTTCTCTAGAGACAAG | 63.94283365960963 | 110.93256579436247 | 128.37849214249073 | 89.79028578008163 | 121.96836896388177 | 94.42780197565655 | 103.31141882540615 | 77.51898622560716 | 82.5826308377408 | 23.328222234460565 | 130.29013108147012 | 15.088853165463343 |
| SE_125_1_6 | SE_125 | SE_125_1 | GTAAAAGCTGTGGTTCTCTG | 20.59175999207768 | 25.984204600481302 | 33.5759133295745 | 19.519627343496005 | 36.06021343279983 | 52.844916701972934 | 38.74178205952731 | 56.7212094333711 | 37.29538166865713 | 17.496166675845423 | 172.54638981059557 | 15.088853165463343 |
| SE_125_1_8 | SE_125 | SE_125_1 | TCTGGTCTCAGGATCCTTGA | 227.59313675454277 | 176.89246978019963 | 204.41806056535063 | 216.01720926802247 | 213.17949705861076 | 338.7272529585478 | 161.42409191469713 | 305.34917744964775 | 172.26914389808294 | 168.4006042550122 | 220.67157336321065 | 147.40648861644956 |
| SE_125_1_9 | SE_125 | SE_125_1 | GGAGGCCTATACAAATGGCA | 2074.348874991404 | 2086.731507915575 | 2073.8064115325424 | 2238.2506020542087 | 2206.036586477166 | 2145.8501421440483 | 2106.046319180415 | 2543.946243086694 | 2022.8304628857368 | 3058.1841335488148 | 2003.6509347393649 | 2048.601987464831 |
| SE_125_2_1 | SE_125 | SE_125_2 | CCAAACGGAGAGAATCTGAT | 950.4722901606382 | 1206.2667289531128 | 1233.4210514305455 | 1100.9069821731748 | 1247.2591469697823 | 1149.5935157953784 | 975.0015151647706 | 1211.9431748930292 | 1393.2489009076914 | 1370.5330562745582 | 1160.8733300862518 | 1425.3162836299218 |
| SE_125_2_10 | SE_125 | SE_125_2 | CTATTTACATGGACTTAAGG | 27.094421042207472 | 42.973876839257535 | 50.36386999436174 | 22.12224432262881 | 43.48437502190568 | 70.17111889934111 | 29.056336544645482 | 76.57363273505099 | 20.423661389978903 | 37.17935418617152 | 53.994108376104734 | 119.5501443109788 |
| SE_125_2_18 | SE_125 | SE_125_2 | TGATGATTCTTGGTAAATAA | 226.50935991285448 | 263.8396159433486 | 291.3204244771905 | 253.7551554654481 | 285.2999239242104 | 517.18713559144 | 248.59310154863357 | 249.57332150683285 | 412.91315418870397 | 398.0377918754834 | 345.09277962119114 | 321.50864052564197 |
| SE_125_2_20 | SE_125 | SE_125_2 | AGAGGCACAGGAAACCACCG | 6044.223446095643 | 6043.326354581171 | 5860.971929736019 | 6267.101685751784 | 5827.96684744809 | 6184.58787435057 | 6335.35752734548 | 7128.910672284191 | 5830.511334200065 | 6703.218857683278 | 7158.914499692669 | 5365.82832184131 |
| SE_125_2_7 | SE_125 | SE_125_2 | TCCTAAGAAGAGAACCTGCG | 355.47880407376203 | 460.7199354162261 | 429.5741852460267 | 421.6239506195137 | 537.7214179538092 | 441.81815603288845 | 359.43764466339223 | 323.3108937702153 | 450.2085358573611 | 416.9919724409826 | 190.1531642810645 | 533.9132658548567 |
| SE_125_3_14 | SE_125 | SE_125_3 | CTAAGAAACAAAACCAAAGG | 118.13167574402458 | 190.88396456507417 | 201.45547997744697 | 119.72038104010883 | 179.24047265126973 | 130.81282659012973 | 87.16900963393644 | 190.96140509234937 | 184.70093778763533 | 175.69067370328113 | 254.7113373394506 | 76.60494684004466 |
| SE_125_3_15 | SE_125 | SE_125_3 | AGTTGCTTGTTCCTTCCAGT | 147.39365046960864 | 206.8742443192165 | 208.36816801588878 | 158.75963572710086 | 113.48361286204653 | 149.8716490072347 | 131.291594757287 | 120.0598933006355 | 242.41998084627136 | 141.42734729641717 | 107.98821675220947 | 27.856344305470785 |
| SE_125_3_2 | SE_125 | SE_125_3 | GTAGTACCAACTGAAACTAA | 399.9136545829823 | 382.76732161478225 | 408.83612113070126 | 363.0650685890257 | 374.38986299348056 | 492.93045251512456 | 319.61970199110027 | 416.9008893352776 | 529.2392255838012 | 388.5607015927338 | 522.3343092905784 | 405.07767344205433 |
| SE_125_3_3 | SE_125 | SE_125_3 | GTGAAGAAGAAAAGAACAGG | 378.2381177492163 | 478.70900013963626 | 476.97547465248476 | 361.7637600994593 | 513.3277441610329 | 442.68446614275683 | 383.1131781442145 | 720.3593598038129 | 539.0070636398781 | 544.5681877856888 | 530.5508040434639 | 649.9813671276517 |
| SE_125_3_9 | SE_125 | SE_125_3 | GTCATTTGTATCTTTTTGAG | 71.52927155142773 | 132.9192004563082 | 124.42838469195254 | 88.48897729051524 | 98.63528968383483 | 136.01068724934018 | 66.72195799140815 | 142.74837707398393 | 98.56636583859384 | 81.64877782061198 | 171.37260484589763 | 71.96222278913287 |
| SE_125_4_1 | SE_125 | SE_125_4 | AAAATCGTGGATGCTCTTAG | 30.34575156727237 | 53.96719417023039 | 73.07698783495626 | 29.93009526002721 | 15.9089176909411 | 28.588233625657487 | 29.056336544645482 | 24.57919075446081 | 22.199631945629246 | 8.019076393095819 | 2.347569929395858 | 13.928172152735392 |
| SE_125_4_12 | SE_125 | SE_125_4 | TGAGATTTAGAGTCTAAGGA | 88.86970101844051 | 118.92770567143364 | 118.50322351614528 | 124.92561499837444 | 155.90739337122278 | 122.14972549144564 | 149.586325174286 | 152.2019119795458 | 129.64585056247478 | 75.08771531716994 | 154.93961534012664 | 53.39132658548567 |
| SE_125_4_13 | SE_125 | SE_125_4 | CTAGTGATGGGAAGAGTCGA | 490.9509092847994 | 551.6646515179107 | 588.5660101301883 | 497.099843014365 | 592.8723326157384 | 978.0641140414334 | 516.5570941270308 | 600.2994665031774 | 518.5834022498991 | 578.8315141925527 | 326.31222018602426 | 603.5541266185337 |
| SE_125_4_20 | SE_125 | SE_125_4 | GCTGACATCTGGATAATTAC | 3733.61121961619 | 3447.9040719869417 | 3499.795201176824 | 3570.7904953702027 | 3456.4775169851364 | 3778.844699245999 | 3841.8933875697917 | 4272.0524238234 | 3505.7658768537704 | 3107.0275988522167 | 4209.192883406774 | 3359.0108508346857 |
| SE_125_4_24 | SE_125 | SE_125_4 | ATAATTACAGGATGGTTGGG | 71.52927155142773 | 125.92345306387092 | 117.51569665351074 | 85.88636031138243 | 63.6356707637644 | 220.90907801644423 | 71.02660044246673 | 43.48626056558451 | 109.22218917249589 | 43.74041668961356 | 68.07952795247988 | 224.01143545649424 |
| SE_125_5_11 | SE_125 | SE_125_5 | TGAACAGCAGTCAGGAAACA | 3695.679030157099 | 3908.624007403168 | 3974.79562210404 | 3868.7901394809087 | 3691.929498811065 | 3745.058604961131 | 3658.9460833998014 | 3841.916585620336 | 3908.023207708572 | 3930.805446506605 | 4109.421161407449 | 3867.3891344095277 |
| SE_125_5_16 | SE_125 | SE_125_5 | TCAGTCATGCCTATTCCAGG | 1354.7210521103736 | 1327.193219593814 | 1381.550080825727 | 1362.4699885760213 | 1287.5617384534999 | 1207.6362931565618 | 1265.5648806112254 | 1417.0848823437213 | 1254.723197566965 | 1527.2695494123402 | 1459.0147111195258 | 1311.5695443825828 |
| SE_125_5_5 | SE_125 | SE_125_5 | TAAGGAGAGCCTCTGCCAAG | 4593.046255075011 | 4862.044437743905 | 4914.9211953321255 | 4522.0470012432415 | 5136.459225148516 | 5611.956891727552 | 4559.6925162838115 | 4941.362695137179 | 5344.783387229697 | 6102.5171351459185 | 4705.703923473998 | 5679.212195277856 |
| SE_125_5_7 | SE_125 | SE_125_5 | CTATCCTTACCTCTTGGCAG | 65.02661050129794 | 101.9380334326574 | 95.79010567555078 | 63.7641159887536 2 | 41.363185996446866 | 72.77004922894633 | 87.16900963393644 | 77.51898622560716 | 39.07135222430747 | 170.5876250894929 | 58.68924823489645 | 66.15881772549312 |
| SE_126_1_4 | SE_126 | SE_126_1 | TCTTTCCTTTTCAGCAATAC | 19.50798315038938 | 23.98541963121351 | 15.800429802152705 | 27.32747828089441 | 38.181402458258646 | 62.37432791052543 | 4.30464245105859 | 20.79777679223607 | 49.72717555820951 | 13.122125006884067 | 28.170839152750297 | 11.606810127279495 |
| SE_126_1_5 | SE_126 | SE_126_1 | TTACTCCAGTATTGCTGAAA | 43.351073667531956 | 16.989672238776237 | 23.700644703229056 | 10.410467916531204 | 11.666539640023474 | 30.320853845394307 | 37.665621446762664 | 7.56282792444948 | 29.303514168230603 | 2.187020834480678 | 11.73784964697929 | 6.964086076367696 |
| SE_126_1_6 | SE_126 | SE_126_1 | CTGCAAAATGGTTTAGGAAG | 250.35245042999705 | 325.80194999065014 | 337.73418702101407 | 300.6022610898385 | 381.81402458258646 | 301.47591823420623 | 281.9540805443376 | 276.9885727329622 | 349.8661994631169 | 282.12568764800744 | 176.06774470468935 | 560.6089291475996 |
| SE_126_1_7 | SE_126 | SE_126_1 | ACATTTCTGCAAAATGGTTT | 237.34712832973747 | 371.7740042838094 | 349.5845093726286 | 338.3402072872641 | 279.9969513605634 | 343.92511361775826 | 261.50702890180935 | 260.91756339350707 | 326.77858223966246 | 209.95400011014507 | 387.34903835031656 | 459.629681040268 |
| SE_126_1_9 | SE_126 | SE_126_1 | ACCATTTTGCAGAAATGTAA | 93.2048083851937 | 106.9349958558269 | 91.8399982250126 | 102.80337067574564 | 102.87766773475245 | 58.04277736118338 | 65.6457973786435 | 86.02716764061283 | 39.95933750213264 | 48.114458358574915 | 167.85124995180385 | 58.034050636397474 |
| SE_126_2_13 | SE_126 | SE_126_2 | GGTCTGTCCTTCAGACAAGG | 60.69150313454474 | 57.964764108765976 | 66.16429979651446 | 50.751031093089615 | 68.93864332741144 | 22.524062856578627 | 61.3411549275849 | 47.26767452780925 | 72.81479278166393 | 9.477090282749604 | 41.08247376442752 | 53.39132658548567 |
| SE_126_2_14 | SE_126 | SE_126_2 | CCTGCCCTGGAGTTTCAGTA | 1013.3313469785595 | 1198.2715890760414 | 1140.5935263428983 | 1154.2606302453971 | 1471.0445891556872 | 1672.8448221558972 | 1052.4850792838251 | 1321.6041797975467 | 1095.7738328362595 | 1295.4453409573882 | 1442.5817216137548 | 1121.2178582951992 |
| SE_126_2_15 | SE_126 | SE_126_2 | AGGCAAGGTTACCATCCTCA | 5998.704818744734 | 5837.451502746588 | 5619.027848390556 | 6082.3158802333555 | 6121.751527474135 | 5437.828559644002 | 6083.535943958552 | 5719.388617864919 | 5758.584526696226 | 6234.467392159586 | 5622.42998090308 | 5580.554309195981 |
| SE_126_2_21 | SE_126 | SE_126_2 | GCACAAAGGAGGTCACCAGA | 2023.411363432054 | 2150.692626932145 | 2183.4218932849767 | 1993.6046060157255 | 2406.4889493830237 | 1866.0319766565524 | 2050.0859673166533 | 1992.805158092438 | 2055.685918165268 | 1743.0556050811003 | 1596.3475519891836 | 2735.725146999777 |
| SE_126_2_24 | SE_126 | SE_126_2 | GAGACACTGGAAGATTTAAG | 503.95623138505897 | 688.5814219127544 | 649.7926756135299 | 553.0561080657202 | 655.4474088667733 | 580.4277736118338 | 443.3781724590348 | 484.9663406553229 | 591.3981950315631 | 626.2169656063008 | 536.4197288669536 | 371.4179240729438 3 |
| SE_126_3_10 | SE_126 | SE_126_3 | GTTGGAGTGGGAGGAAGCAG | 588.4908250367463 | 803.5115576456525 | 794.9591244208079 | 697.5013504075906 | 765.7492381906317 | 679.1871261368325 | 643.5440464332592 | 556.813205937593 | 793.8588383757018 | 480.41557664092227 | 373.26361877394146 | 614.0002557330853 |
| SE_126_3_12 | SE_126 | SE_126_3 | GGAGAAGCTAAGATAAGCAA | 1590.984403598423 | 1777.9192301637013 | 1809.1492123464848 | 1842.652821226023 | 1639.679116679663 | 2372.8233909295714 | 1892.966517853015 | 1726.2154737555938 | 2033.4862862196387 | 2122.8682233359114 | 1666.7746498710592 | 1832.7153190974323 |
| SE_126_3_26 | SE_126 | SE_126_3 | CTTGTAATCCCAGCGACTAG | 627.5067913375251 | 696.5765617898256 | 713.9819216847753 | 735.2392966050162 | 509.0853661101152 | 707.77535976249 | 653.229491948141 | 638.1136061254249 | 676.6447817027794 | 1197.7584103505847 | 816.9543354297587 | 818.2801139732044 |
| SE_126_3_5 | SE_126 | SE_126_3 | TCTGCTACAAGAGTGCAGTA | 387.992109324411 | 385.76549906868394 | 436.48687328446846 | 392.9951638490529 | 417.87423801538625 | 495.5293828447298 | 389.5701418208024 | 254.30008895961376 | 278.8273772371033 | 204.85095149635683 | 75.12223774066746 | 529.270541803945 |
| SE_126_3_7 | SE_126 | SE_126_3 | GCACTCTTGTAGCAGAAGAG | 299.1224083059705 | 380.7685366455145 | 432.5367658339303 | 296.6983356211393 | 313.9359757679044 | 404.5668213085469 | 253.9739046124568 | 397.0484660335977 | 328.5545527953128 | 228.9081806756443 | 487.12076034964053 | 244.90369368559732 |
| SE_127_1_10 | SE_127 | SE_127_1 | TGAGGGGCAGAAAGAGAGGC | 1661.4298983081621 | 2465.501259591822 | 2463.8795222731874 | 1889.4999268504134 | 2303.6112816482714 | 1945.732506764446 | 1594.8700281172075 | 1992.805158092438 | 2455.2792931865943 | 1853.1356537499612 | 1782.9793613761542 | 1794.41284567741 |
| SE_127_1_12 | SE_127 | SE_127_1 | AATGGAGGCGAAGACAGACG | 1596.4032878068642 | 1559.052276028878 | 1617.569000995383 | 1454.8628913352356 | 1383.0152445991464 | 1414.6844094151115 | 1487.2539668407428 | 1278.117919231962 | 1506.9110164693132 | 1344.2888062607901 | 1847.5375344345402 | 2303.9518102649795 |
| SE_127_1_13 | SE_127 | SE_127_1 | TTCTGCCCCTCACAGCAACA | 49.85373471766175 | 35.97812944682026 | 18.763010390056337 | 20.820935833062407 | 45.60556404736449 | 11.262031428289314 | 47.35106696164449 | 23.633837263904624 | 21.311646667804077 | 43.01140974478667 | 9.390279717583432 | 39.46315443275028 |
| SE_127_1_26 | SE_127 | SE_127_1 | ACATAGTTACTAGTCTTAAG | 259.02266516350346 | 357.7825094989348 | 429.5741852460267 | 355.25721765162734 | 400.90472581171576 | 523.2513063605189 | 313.1627383145124 | 361.1250333924627 | 326.77858223966246 | 481.87359053057605 | 197.19587406925208 | 678.9983924458504 |
| SE_127_1_8 | SE_127 | SE_127_1 | ATAAAACCAGAAACACTGGG | 65.02661050129794 | 80.95079125534559 | 62.21419234597627 | 55.95626505135522 | 33.939024407341016 | 82.29946043749882 | 47.35106696164449 | 36.868786131691216 | 48.83919028038434 | 74.35870837234305 | 52.820323411406804 | 29.017025318198737 |
| SE_128_1_1 | SE_128 | SE_128_1 | GACTGCCAGTATCATGAAGT | 1536.7955615140079 | 1423.1348981186682 | 1474.3776059133743 | 1517.3256988344228 | 1694.830031341592 | 1627.79669644274 | 1557.204406670445 | 1297.0249890430857 | 1453.631899799803 | 1635.891584191547 | 1101.0102968866574 | 1416.0308355280984 |
| SE_128_1_15 | SE_128 | SE_128_1 | GGCTGGTTCTTACCAACTGT | 98.6236925936352 | 107.93438834046079 | 79.98967587339807 | 80.68112635311682 | 85.90815553108195 | 32.05347406513112 | 75.33124289352533 | 153.14726547010198 | 51.50314611385985 | 33.53431946203706 | 415.5198775030669 | 37.1417924072943 8 |
| SE_128_1_2 | SE_128 | SE_128_1 | TTGGAAATAATAGAGAACAA | 65.02661050129794 | 106.9349958558269 | 83.93978332393624 | 53.35364807222242 | 97.57469517110542 | 109.1550738434195 | 38.74178205952731 | 146.52979103620868 | 102.11830694989453 | 67.06863892407412 | 118.55228143449084 | 207.76190127830296 |
| SE_128_1_23 | SE_128 | SE_128_1 | TATTTCTTGACTTATAGACA | 1.083776841688299 | 1.9987849692677924 | 4.93763431317272 | 0.0 | 1.0605945127294067 | 5.197860659210452 | 1.0761606127646475 | 0.0 | 0.0 | 5.103048613788249 | 0.0 | 0.0 |
| SE_128_1_9 | SE_128 | SE_128_1 | TACACCCTCACGTATCTAAT | 947.2209596355732 | 816.5036599458932 | 824.5849302998442 | 853.6583691555587 | 833.6272870053137 | 797.8716111888044 | 814.6535838628381 | 825.2935972555495 | 867.5616164351909 | 545.2971947305157 | 838.0824647943214 | 887.9209747368814 |
| SE_129_1_17 | SE_129 | SE_129_1 | ATAATAAACAAGGAAATAAA | 15.172875783636185 | 31.980559508284678 | 24.688171565863602 | 37.737946197425615 | 28.63605184369398 | 0.0 | 12.913927353175769 | 21.743130282792254 | 4.439926389125849 | 0.7290069448268927 | 39.90868879972959 | 0.0 |
| SE_129_1_22 | SE_129 | SE_129_1 | ATGATGCTATGCCACCAAAG | 1210.57873216583 | 1306.2059774165023 | 1227.4958902547382 | 1234.941756598514 | 1355.4397872681818 | 1374.8341443611646 | 1305.3828232835174 | 1324.440240269215 | 1372.8252395177126 | 1014.7776671990346 | 1596.3475519891836 | 1104.9683241170078 |
| SE_129_1_23 | SE_129 | SE_129_1 | GTGTGGAAGAATAAACCAGA | 260.10644200519175 | 370.7746117991755 | 383.1604227022031 | 292.7944101524401 | 391.3593751971511 | 266.8235138394699 | 275.49711686774975 | 253.35473546905757 | 621.5896944776189 | 255.15243068941243 | 196.02208910455414 | 236.7789265965017 |
| SE_129_1_26 | SE_129 | SE_129_1 | GAGAAGGGCCCCATTAAAGG | 110.54523785220648 | 187.88578711117248 | 202.44300684008152 | 110.61122161314404 | 193.02820131675202 | 247.7646914223649 | 106.5399006637001 | 206.08706094124832 | 109.22218917249589 | 112.26706950334147 | 99.77172199932397 | 293.6522962201712 |
| SE_129_1_8 | SE_129 | SE_129_1 | CAAATTCTAGCCCCACCAGG | 62.859056817921335 | 75.95382883217611 | 73.07698783495626 | 57.25757354092162 | 75.30221040378788 | 67.57218856973589 | 44.12258512335055 | 49.15838150892162 | 63.04695472558706 | 136.32429868262892 | 11.73784964697929 | 62.67677468730927 |
| SE_129_2_12 | SE_129 | SE_129_2 | CTCTGCCTAGAGATCAATAC | 4.335107366753196 | 21.986634661945715 | 22.713117840594514 | 7.807850937398403 | 10.605945127294067 | 3.465240439473635 | 0.0 | 9.45353490556185 | 1.7759705556503396 | 2.9160277793075706 | 0.0 | 9.285448101823595 |
| SE_129_2_13 | SE_129 | SE_129_2 | GTTGGCCAGTATTGATCTCT | 631.8418987042783 | 641.6099751349614 | 651.7677293387991 | 625.9293834814387 | 675.5987046086321 | 678.320816026964 | 693.047434620433 | 593.6819920692842 | 554.9907986407311 | 770.5603406820255 | 671.4049998072154 | 689.444521560402 |
| SE_129_2_14 | SE_129 | SE_129_2 | CAATACTGGCCAACACATGA | 99.7074694353235 | 103.93681840192521 | 71.10193410968716 | 124.92561499837444 | 102.87766773475245 | 33.78609428486794 | 114.07302495305264 | 77.51898622560716 | 60.382998892111544 | 191.72882648947277 | 37.56111887033373 | 116.06810127279495 |
| SE_129_2_15 | SE_129 | SE_129_2 | AATACTGGCCAACACATGAT | 326.21682934817795 | 384.76610658405 | 359.459777998974 | 305.8074950481041 | 357.4203507898101 | 457.4117380105198 | 338.99059302086397 | 305.34917744964775 | 332.99447918443866 | 505.2018127650366 | 217.15021846911688 | 676.6770304203945 |
| SE_129_2_4 | SE_129 | SE_129_2 | GCACTGGACAGATAGCAAAA | 1475.0202815377747 | 1619.0158251069117 | 1436.8515851332616 | 1616.2251440414693 | 1656.6486288833332 | 1649.4544491894503 | 1655.1350224320279 | 1636.4068921527562 | 1325.7620197929784 | 1909.2691885016318 | 1625.6921761066317 | 1251.2141317207295 |
| SE_129_3_16 | SE_129 | SE_129_3 | GCCGGCCCCGGGAGCCTTTA | 9.75399157519469 | 12.992102300240651 | 31.60085960430541 | 29.93009526002721 | 20.15129574185873 | 51.11229648223612 | 7.533124289352532 | 19.852423301679885 | 41.73530805778298 | 18.225173620672315 | 18.780559435166865 | 17.410215190919242 |
| SE_129_3_27 | SE_129 | SE_129_3 | TGAAGCTTCATCTGTTCTTA | 639.4283365960963 | 772.5303906220017 | 868.0361122557642 | 718.322286240653 | 831.5060979798549 | 595.1550454795968 | 669.3719011396107 | 691.9987550871274 | 764.5553242074711 | 910.529674088789 | 697.2282690305699 | 712.6581418149609 |
| SE_129_3_28 | SE_129 | SE_129_3 | ACACAGGGCATTATGTGGTG | 5920.672886143177 | 5367.737034968656 | 5557.801182907214 | 5638.569685291213 | 5280.700078879716 | 4508.277811755199 | 5806.962666478038 | 5374.334593811912 | 5264.8647122254315 | 6042.7385656701135 | 5841.927769301593 | 4974.6788205519915 |
| SE_129_3_40 | SE_129 | SE_129_3 | AAAGTGAAGGACGATATTAG | 238.43090517142576 | 347.7885846525959 | 363.4098854495122 | 305.8074950481041 | 251.3608995168694 | 239.9679004335492 | 298.0964897358073 | 367.74250782635596 | 321.4506705727115 | 258.7974654135469 | 335.7024999036077 | 220.52939241831038 |
| SE_129_3_9 | SE_129 | SE_129_3 | GGCAAGGGTCCCCAGCCCCT | 44.434850509220254 | 32.97995199291857 | 29.62580587903632 | 45.545797134824014 | 59.39329271284678 | 51.11229648223612 | 51.655709412703075 | 13.23494886778659 | 46.17523444690883 | 44.46942363444045 | 16.432989505771005 | 35.98111139456643 |
| SE_129_4_17 | SE_129 | SE_129_4 | TTAATAGGCATTTTTAAGCT | 36.84841261740216 | 48.970231747060915 | 44.43870881855448 | 46.847105624390416 | 47.72675307282331 | 45.91443582302566 | 31.208657770174778 | 71.84686528227006 | 80.80666028209045 | 21.87020834480678 | 83.33873249355297 | 41.78451645820618 |
| SE_129_4_6 | SE_129 | SE_129_4 | CAGCAGGAGACGAGCGGCTG | 1948.6307613555614 | 2411.5340654215915 | 2564.607262261911 | 2264.2767718455366 | 2358.7621963102006 | 2378.88756169865 | 1875.7479480487805 | 1949.3188975268536 | 2259.922532065057 | 2463.31446657007 | 2423.8659521012232 | 2603.40751154879 1 |
| SE_129_4_7 | SE_129 | SE_129_4 | ACAGCAGGAGACGAGCGGCT | 3969.874571104239 | 4456.291088982543 | 4484.359483223465 | 4268.291845777793 | 4507.5266790999785 | 5000.341954160455 | 4171.198535075773 | 5227.804802775703 | 4280.977024395143 | 4823.838953919549 | 4857.122183920031 | 4625.313835720878 |
| SE_129_4_8 | SE_129 | SE_129_4 | CACAGCAGGAGACGAGCGGC | 2632.493948460878 | 2735.337230442974 | 2754.212419887743 | 2816.0315714216904 | 2879.514102060339 | 2987.0372588262735 | 2580.633149409625 | 3015.67763487423 | 2772.29003737018 | 2467.6885082390318 | 2555.3298681473916 | 2787.9557925725344 |
| SE_13_1_12 | SE_13 | SE_13_1 | TCAACTCCACAGAGTGCAGA | 335.9708209233727 | 372.77339676844326 | 331.8090258452068 | 322.7245054124673 | 274.6939787969163 | 128.21389626052448 | 222.76524684228204 | 120.0598933006355 | 237.9800544571455 | 103.51898616541875 | 82.16494752885504 | 99.81856709460365 |
| SE_13_1_21 | SE_13 | SE_13_1 | GGATGAAGAGGAAAAGCAGG | 727.2142607728485 | 1168.2898145370245 | 1041.840840079444 | 487.99068358740016 | 674.5381100959027 | 434.02136504407275 | 407.86487223780136 | 207.0324144318045 | 902.1930422703725 | 188.0837917653383 | 56.341678305500594 | 67.31949873822107 |
| SE_13_1_3 | SE_13 | SE_13_1 | ACAGGAAGAGACACACTCAT | 5896.829795626034 | 5811.467298146106 | 5717.78053465401 | 5258.587606337825 | 5169.3376550431285 | 5064.448902290717 | 5420.621006495529 | 4730.54886674315 | 5753.256615029275 | 4301.140974478667 | 4813.692140226207 | 4358.35720279345 |
| SE_13_1_34 | SE_13 | SE_13_1 | ATGTCACCAGGGCAAGGCAG | 1881.436597170887 | 2255.6288378187037 | 2199.2223230871296 | 1518.6270073239893 | 1504.9836135630283 | 1198.1068819480092 | 1248.3463108069911 | 1183.5825701763436 | 1715.587556758228 | 451.98430579267347 | 1207.824728674169 | 1189.6980380461482 |
| SE_13_1_38 | SE_13 | SE_13_1 | GTCCCTCTGAGGAAGCCACA | 78.03193260155751 | 134.91798542557598 | 135.29118018093254 | 66.3667329678864 2 | 112.42301834931712 | 38.11764483420998 | 55.960351863761666 | 50.10373499947781 | 84.35860139339113 | 83.10679171026577 | 58.68924823489645 | 5.803405063639747 |
| SE_13_2_18 | SE_13 | SE_13_2 | CTTCCTTGAGAGAGGCAAGC | 28.17819788389577 | 71.95625889364052 | 83.93978332393624 | 50.751031093089615 | 60.45388722557618 | 72.77004922894633 | 25.827854706351538 | 28.36060471668555 | 7.991867500426528 | 17.496166675845423 | 34.03976397623994 | 3.482043038183848 |
| SE_13_2_19 | SE_13 | SE_13_2 | CTATCTCCCGGACAAGCCCA | 118.13167574402458 | 124.92406057923702 | 125.4159115545871 | 70.27065843658562 | 69.99923784014085 | 124.74865582105086 | 129.1392735317577 | 79.40969320671954 | 140.30167389637683 | 98.4159375516305 | 52.820323411406804 | 131.15695443825828 |

| | | | | | | | | | | | | | | | |
|---|---|---|---|---|---|---|---|---|---|---|---|---|---|---|---|
| SE_13_2_20 | SE_13 | SE_13_2 | GGCTTGCCTCTCTCAAGGAA | 84.53459365168732 | 137.91616287947767 | 145.16644880727796 | 85.88636031138243 | 129.39253055298764 | 19.05882241710499 | 34.43713960846872 | 95.48070254617468 | 173.1571291759081 | 18.954180565499207 | 57.515463270198524 | 17.410215190919242 |
| SE_13_2_25 | SE_13 | SE_13_2 | TTTCCCCCCAATTTCCACAT | 16.256652625324485 | 29.981774539016886 | 19.75053725269088 | 5.205233958265602 | 5.3029725636470335 | 6.93048087894727 | 2.152321225529295 | 0.0 | 22.199631945629246 | 2.9160277793075706 | 1.173784964697929 | 0.0 |
| SE_13_2_26 | SE_13 | SE_13_2 | ATACCCATGTGGAAATTGGG | 44.434850509220254 | 66.95929647047105 | 43.451181955919935 | 29.93009526002721 | 60.45388722557618 | 79.7005301078936 | 23.675533480822246 | 123.84130726286024 | 54.167101947335354 | 29.160277793075707 | 305.18409082146155 | 19.73157721637514 |
| SE_13_3_15 | SE_13 | SE_13_3 | TTGGCATTACAGGCCTCATG | 134.38832836934907 | 193.88214201897586 | 181.7049427247561 | 130.13084895664005 | 173.9375000876227 | 228.7058690052599 | 119.45382801687587 | 169.21827480955713 | 213.11646667804075 | 206.3089653860106 | 96.25036710523018 | 345.88294179292893 |
| SE_13_3_16 | SE_13 | SE_13_3 | AGGTGGCTCCAGTTAAACCA | 50.93751155935005 | 49.96962423169481 | 67.151826659149 | 39.03925468699201 | 74.24161589105847 | 186.25667362170788 | 38.74178205952731 | 34.97807915057884 | 30.191499446055772 | 14.580138896537854 | 29.344624117448227 | 174.10215190919243 |
| SE_13_3_20 | SE_13 | SE_13_3 | TCCATTTTCCATTAGCAAGG | 101.8750231187001 | 114.93013573289807 | 110.60300861506893 | 105.40598765487843 | 118.78658542569356 | 201.85025559933922 | 96.85445514881827 | 67.12009782948914 | 65.71091055906257 | 37.17935418617152 | 63.384388093688166 | 78.92630886550056 |
| SE_13_3_21 | SE_13 | SE_13_3 | TTTTCCATTTTCCATTAGCA | 0.0 | 0.9993924846338962 | 0.0 | 0.0 | 0.0 | 0.0 | 0.0 | 0.0 | 0.0 | 0.0 | 0.0 | 0.0 |
| SE_13_3_9 | SE_13 | SE_13_3 | CCAAAAGATTTGACCCCCCG | 127.88566731921927 | 152.90705014898612 | 182.69246958739063 | 143.14393385230406 | 218.48246962225778 | 129.94651648026132 | 72.10276105523138 | 197.57887952624267 | 271.72349501450196 | 172.04563897914667 | 53.994108376104734 | 134.63899747644214 |
| SE_13_4_0 | SE_13 | SE_13_4 | CATGAGAGGTGAGTACATGC | 9066.877057564308 | 8113.068190257969 | 8006.867802240883 | 7850.794117554094 | 7007.34794560319 | 6104.0210341328075 | 7538.505092416355 | 4858.171587968235 | 6674.097348133976 | 4003.7061409892945 | 3582.391712258079 | 4440.765554697135 |
| SE_13_4_1 | SE_13 | SE_13_4 | TCACCTCTCATGAAGCACTG | 535.3857597940197 | 464.7175053547617 | 491.7883775920029 | 459.3618968169393 | 580.1451984629855 | 444.4170863624937 | 477.8153120675035 | 593.6819920692842 | 610.0458858658916 | 345.54929184794713 | 322.79086529193046 | 276.24208102925195 |
| SE_13_4_2 | SE_13 | SE_13_4 | CACCTCTCATGAAGCACTGT | 475.77803350116324 | 501.69502728621586 | 458.21246426242845 | 373.47553650555693 | 550.4485521065621 | 369.91441691381056 | 404.63639039950743 | 279.82463320463074 | 411.1371836330536 | 331.6981598962362 | 309.8792306802533 | 248.38573672378118 |
| SE_13_4_3 | SE_13 | SE_13_4 | TACCCACAGTGCTTCATGAG | 483.36447139298133 | 552.6640440025446 | 546.1023550369028 | 480.1828326500018 | 433.78315570632736 | 203.58287581907607 | 401.4079085612135 | 249.57332150683285 | 405.8092719661026 | 199.0188959377417 | 349.78791947998286 | 97.49720506914775 |
| SE_13_4_4 | SE_13 | SE_13_4 | ATGAAGCACTGTGGGTACGA | 2345.293085413479 | 2303.5996770811307 | 2210.0851185761094 | 2213.5257407524473 | 2053.3109766441316 | 2063.5506817065498 | 2346.0301358269317 | 2059.925255921927 | 2084.1014470556734 | 1501.7543063433989 | 1879.2297284813844 | 1539.063022877261 |
| SE_130_1_10 | SE_130 | SE_130_1 | TGCAGCTGAAGGCAGAAGAG | 1705.8647488173824 | 1541.0632113054678 | 1475.3651327760087 | 1814.0240344555623 | 1332.106707988135 | 1721.358188308528 | 1827.3207204743715 | 1701.636283001133 | 1743.1151003708082 | 1908.540181556805 | 1683.2076393768302 | 2384.038800143208 |
| SE_130_1_11 | SE_130 | SE_130_1 | ATTGAGACTGAGTAGAAGAG | 87.78592417675222 | 117.92831318679976 | 91.8399982250126 | 134.03477442533924 | 135.75609762936406 | 51.978606592104526 | 101.15909759987686 | 180.56251669623134 | 213.11646667804075 | 24.057229179287457 | 329.83357508011807 | 52.23064557275772 |
| SE_130_1_15 | SE_130 | SE_130_1 | GATGTACAAAGAAGATCAGT | 1189.9869721737523 | 1460.1124200501224 | 1286.7475020128109 | 1405.4131687317124 | 1391.5000007009817 | 1027.4437903039327 | 1358.1146933089851 | 1366.9811473442435 | 996.3194817198405 | 1625.6854869639706 | 1359.2429891202019 | 1087.5581089260886 |
| SE_130_1_24 | SE_130 | SE_130_1 | GTTAGCCTCCTACTCAGGAG | 76.94815575986922 | 101.9380334326574 | 63.20171920861082 | 163.96486968536647 | 108.18064029839948 | 145.54009845789267 | 86.0928490211718 | 66.17474433893295 | 187.36489362111084 | 75.08771531716994 | 63.384388093688166 | 185.70896203647192 |
| SE_130_1_8 | SE_130 | SE_130_1 | TGAAAGTGAGAGGCATCTGG | 628.5905681792134 | 740.5498311137171 | 677.4434277672972 | 508.8116194204626 | 619.3871954339735 | 771.8823078927521 | 644.6202070460239 | 664.583503860998 | 660.6610467019264 | 807.739694868197 | 744.179667618487 | 803.1912608077411 |
| SE_130_2_10 | SE_130 | SE_130_2 | GACTGTGAGACTTATACCGG | 534.3019829523314 | 585.6439959954631 | 617.20428914659 | 573.8770438987826 | 606.6600612812207 | 576.0962230624918 | 526.2425396419126 | 654.18461546488 | 688.1885903145065 | 717.3428337096624 | 691.3593442070802 | 792.7451316931895 |
| SE_130_2_12 | SE_130 | SE_130_2 | ACACAAGCCAGAAATCTGGC | 293.703524097529 | 389.7630690072195 | 379.2103152516649 | 321.4231969229009 | 324.5419208951985 | 476.4705604276248 | 266.8878319656326 | 361.1250333924627 | 427.1209186339067 | 279.2096598686999 | 264.10161705703405 | 615.1609367458132 |
| SE_130_2_19 | SE_130 | SE_130_2 | CACGGGAGAAAGATAAAGAC | 4836.896044454878 | 5852.442390016096 | 5950.8368742357625 | 4889.015995300966 | 6018.873859739383 | 6097.090553253861 | 4462.838061134993 | 5729.787506261037 | 6344.654810060838 | 7443.889913627401 | 5522.658258903756 | 6181.787073789059 |
| SE_130_2_4 | SE_130 | SE_130_2 | ACACTGAGAGATTTCCAGCT | 618.8365766040187 | 880.4647789624626 | 807.796973635057 | 594.697979731845 | 826.2031254162079 | 1051.7004733802482 | 523.0140578036187 | 595.5726990503965 | 740.5797217061915 | 493.53770164780633 | 586.8924823489646 | 800.8698987822851 |
| SE_130_2_9 | SE_130 | SE_130_2 | GTTTCCTGTCCATATCAACA | 36.84841261740216 | 38.97630690072195 | 66.16429979651446 | 55.95626505135522 | 18.030106716399914 | 91.82887164605133 | 19.370891029763655 | 36.868786131691216 | 25.751573056929924 | 37.17935418617152 | 56.341678305500594 | 9.285448101823595 |
| SE_130_3_12 | SE_130 | SE_130_3 | GCTATTAAAGAAAGGACAAG | 158.23141888649164 | 198.87910444214535 | 174.7922546863143 | 204.30543286192486 | 274.6939787969163 | 189.7219140611815 | 204.47051642528302 | 287.38746112908024 | 91.46248361599248 | 350.6523404617354 | 255.88512230414852 | 105.6219721582434 |
| SE_130_3_17 | SE_130 | SE_130_3 | ATACGATAATTAGCCAGGCG | 356.5625809154503 | 479.7083926242702 | 459.199991125063 | 373.47553650555693 | 441.2073172954332 | 377.7112079026262 | 355.1330022123337 | 293.05958207241736 | 385.38561057612367 | 311.2859654410832 | 363.873339056358 | 335.4368126783774 |
| SE_130_3_4 | SE_130 | SE_130_3 | ATGCATGTCAAGAAGTAGAA | 434.59451351700784 | 540.6713341869379 | 486.85074327883024 | 486.68937509783376 | 411.5106709390098 | 406.2994415282837 | 475.6629908419742 | 333.7097821663333 | 630.4695472558706 | 492.07968775815255 | 417.8674474324627 | 499.0928354730183 |
| SE_130_3_7 | SE_130 | SE_130_3 | TGTGAGAGAGAGCATGATGA | 5926.091770351619 | 5471.673853370582 | 5388.934089396706 | 5724.456045602596 | 5582.969515007597 | 5927.293771719653 | 5757.459278290864 | 5207.007025983467 | 5341.231446118396 | 6000.456162870153 | 5237.428512482159 | 5687.336962366952 |
| SE_131_1_10 | SE_131 | SE_131_1 | GATAATGTCTCCGCGATGGA | 174.48807151181612 | 294.8207829669994 | 307.1208542793432 | 149.65047630013606 | 203.6341464440461 | 118.68448505197199 | 77.48356411905462 | 79.40969320671954 | 171.38115862025776 | 16.76715973101853 | 84.5125174582509 | 13.928172152735392 |
| SE_131_1_11 | SE_131 | SE_131_1 | GATGAACTAACCATCCATCG | 985.1531490946637 | 1447.1203177498817 | 1425.001262781647 | 595.9992882214113 | 617.2660064085147 | 494.6630727348614 | 471.3583483909156 | 247.68261452572048 | 576.3024453085352 | 97.68693060680361 | 170.1988198811997 | 181.06623798556012 |
| SE_131_1_2 | SE_131 | SE_131_1 | TGGTATGGCAGCATAGGAAA | 2.167553683376598 | 7.9951398770711695 | 5.925161175807264 | 2.602616979132801 | 0.0 | 0.0 | 0.0 | 0.0 | 0.8879852778251698 | 0.0 | 0.0 | 5.803405063639747 |
| SE_131_1_6 | SE_131 | SE_131_1 | TCAAAGATAATGAAGACTGG | 216.75536833765977 | 424.7418059694059 | 413.77375544387394 | 210.81197530975686 | 146.36204275665813 | 72.77004922894633 | 164.65257375299106 | 83.19110716894428 | 244.1959514019217 | 16.76715973101853 | 39.90868879972959 | 47.58792152184593 |
| SE_131_1_8 | SE_131 | SE_131_1 | CATTATCTTTGAAAAACACT | 166.90163361999802 | 184.8876096572708 | 158.9918248841616 | 110.61122161314404 | 147.42263726938754 | 181.05881296249743 | 163.5764131402264 | 94.5353490556185 | 161.6133205641809 | 71.44268059303548 | 59.86303319959438 | 142.76376456553777 |
| SE_131_2_10 | SE_131 | SE_131_2 | TCATTGTTGGGTCTCCCACA | 1381.815473152581 | 1695.969046423722 | 1789.3986750937938 | 1497.8060714909268 | 1499.6806409993812 | 1911.0801023697097 | 1471.1115576492732 | 2009.8215209224493 | 1607.2533528635572 | 1269.930097888447 | 1343.9837845791287 | 1787.4487596010422 |
| SE_131_2_11 | SE_131 | SE_131_2 | AGAATTTCCTTCGTGATTAC | 9.75399157519469 | 22.986027146579612 | 24.688171565863602 | 24.724861301761607 | 30.757240869152795 | 34.65240439473635 | 19.370891029763655 | 22.68848377334844 | 9.767838056076867 | 2.187020834480678 | 14.085419576375148 | 3.482043038183848 |
| SE_131_2_2 | SE_131 | SE_131_2 | CCACGACCTTGTCTTCAGCC | 1377.4803657858279 | 1166.2910295677568 | 1195.8950306504328 | 1141.2475453497332 | 1168.7751530278063 | 942.5453995368287 | 1169.7865860751717 | 1074.8669187623823 | 1293.7945497912724 | 887.9304587991553 | 843.951389617811 | 1205.9475722243394 |
| SE_131_2_25 | SE_131 | SE_131_2 | CATTTTACAGAAGGTAACTG | 63.94283365960963 | 109.93317330972857 | 114.55311606560711 | 54.65495656178882 | 68.93864332741144 | 51.978606592104526 | 71.02660044246673 | 67.12009782948914 | 52.391131391685015 | 32.076305572383276 | 45.7776136232192 | 52.23064557275772 |
| SE_131_2_4 | SE_131 | SE_131_2 | GTTATGTGTCAAAATCAGCT | 207.0013767624651 | 220.86573910409106 | 244.90666193336693 | 193.89496494539367 | 181.36166167672854 | 161.133680435524 | 180.79498294446077 | 223.10342377125966 | 362.29799335266927 | 220.1600973377216 | 100.9455069640219 | 121.87150633643469 |
| SE_132_1_26 | SE_132 | SE_132_1 | GCTGGGGCTCTGGGACACCT | 2339.8742012050375 | 2248.6330904262663 | 2195.272215636591 | 2393.1063123126105 | 2272.854040779119 | 2236.812703680231 | 2397.6858452396345 | 2079.777679223607 | 2109.853020112603 | 2213.994091439273 | 1846.3637494698423 | 2630.1031748415335 |
| SE_132_1_30 | SE_132 | SE_132_1 | AGTCCAGGAAAGCCTGAAGG | 89.95347786012881 | 93.94289355558624 | 105.66537430189621 | 74.17458390528482 | 126.21074701479941 | 103.95721318420905 | 58.112673089290965 | 109.66100490451745 | 135.86174750725098 | 29.8892847379026 | 79.81737759945918 | 214.72598735467065 |
| SE_132_1_36 | SE_132 | SE_132_1 | TGGCCTGGCTCAGTGCCAAG | 60.69150313454474 | 28.982382054382988 | 41.47612823065085 | 59.86019052005442 | 62.575076251035 | 51.11229648223612 | 49.50338818717378 | 27.415251226129364 | 42.62329333560815 | 67.79764586890101 | 28.170839152750297 | 34.820430381838484 |
| SE_132_1_7 | SE_132 | SE_132_1 | CCTTGTCCTTAAGAAAGATG | 431.34318299194297 | 444.7296556620838 | 405.87354054279757 | 435.9383440047441 | 475.14634170277424 | 346.5240439473635 | 404.63639039950743 | 440.5347265991822 | 606.493944754591 | 451.2552988478466 | 489.4683302790364 | 394.6315443275028 |
| SE_132_1_8 | SE_132 | SE_132_1 | TTAAGGCACTCTAGCAGGCA | 1107.6199322054415 | 1057.3572487426623 | 1054.678689293693 | 1104.8109076418739 | 1081.806402983995 | 798.7379212986729 | 1030.9618670285322 | 922.6650067828366 | 1280.4747706238948 | 939.6899518818647 | 955.4609612641142 | 1920.9270760647564 |
| SE_132_2_14 | SE_132 | SE_132_2 | AGGCATGGTTGCCAACATGA | 2452.5869927406206 | 2455.507334745483 | 2386.852426987693 | 2374.8879934586807 | 2392.7012207175417 | 2618.8554621321996 | 2554.805294703273 | 2365.274433371575 | 2308.7617223454413 | 2659.4173347285046 | 2879.29451840402 | 2461.804427995981 |
| SE_132_2_15 | SE_132 | SE_132_2 | GGAGGATCAATAATTTTAAA | 184.24206308701082 | 405.75334876136185 | 417.72386289441215 | 330.5323563498657 | 393.4805642226099 | 406.2994415282837 | 287.33488360816085 | 480.239573202542 | 404.9212866882774 | 668.4993684062606 | 659.6671501602361 | 646.4993240894678 |
| SE_132_2_16 | SE_132 | SE_132_2 | TCAAGTTGAGGTTTTCTGAG | 329.4681598732429 | 387.7642840379517 | 381.185368976934 | 301.9035695794049 | 382.87461909531584 | 523.2513063605189 | 306.70577463792455 | 354.5075589585694 | 340.9863466848652 | 293.7897987652377 | 458.9499211968903 | 248.38573672378118 |
| SE_132_2_19 | SE_132 | SE_132_2 | GCAAATGCCCTATTGTATAG | 14.089098941947885 | 22.986027146579612 | 26.66322529113269 | 20.820935833062407 | 16.969512203670508 | 58.04277736118338 | 12.913927353175769 | 0.945353490556185 | 34.63142583518162 | 20.412194455152996 | 1.173784964697929 | 38.30247342002233 |
| SE_132_2_6 | SE_132 | SE_132_2 | ACAGGAGTTGCACATGCAGA | 2746.2905168381494 | 2758.3232575895536 | 2734.4618826350525 | 3082.799811782803 | 2805.272486169281 | 3340.491783652584 | 2841.0640176986694 | 3050.655714024809 | 2955.215004602165 | 2437.070216556302 | 3220.865943131117 | 4097.203974929662 |
| SE_132_3_18 | SE_132 | SE_132_3 | AGCAGGGTAAGGAAGGGTGA | 2911.024596774771 | 2585.4283577478895 | 2533.99392952024 | 2571.385575383207 | 2705.5766019727166 | 3120.449015746008 | 3003.564270226131 | 2769.885727329622 | 2677.2756126428867 | 4255.942543899399 | 3182.1310392960854 | 3443.740564763826 |
| SE_132_3_2 | SE_132 | SE_132_3 | ATATTGGTGTCAGGAAGTAG | 127.88566731921927 | 168.89732990312845 | 187.63010390056337 | 114.51514708184324 | 212.11890254588135 | 186.25667362170788 | 104.3875794381708 | 229.72089820515296 | 140.30167389637683 | 95.49990977232294 | 169.0250349165018 | 60.35541266185337 |
| SE_132_3_21 | SE_132 | SE_132_3 | TGAGATCCTGAGGTATGAAG | 731.5493681396017 | 880.4647789624626 | 842.360413827266 | 735.2392966050162 | 880.2934455654076 | 879.3047615164348 | 790.9780503820159 | 847.0367275383418 | 817.8344408769814 | 930.941868543942 | 746.5272375478829 | 851.9398633423149 |
| SE_132_3_33 | SE_132 | SE_132_3 | CACATCCCCCTCCTTTAAAA | 1074.0228501131041 | 1652.9951695844643 | 1640.2821188359776 | 1242.7496075359124 | 1463.6204275665814 | 1775.0694151203695 | 1190.2336377177 | 1229.9048912135968 | 1603.7014117522567 | 1269.20109094362 | 1289.989676203024 | 1526.2955317372534 |
| SE_132_3_6 | SE_132 | SE_132_3 | ATGCAGAGAAAACAAGTAGG | 476.8618103428515 | 791.5188478300457 | 715.9569754100444 | 597.3005967109779 | 655.4474088667733 | 688.716537345385 | 427.23576326756506 | 806.3865274444258 | 864.8976606017154 | 426.4690627237322 | 807.5640557121752 | 922.7414051187199 |
| SE_132_4_14 | SE_132 | SE_132_4 | ACCAGGCTCAGAGACACACA | 3625.23353544736 | 3243.0286126369933 | 3460.2941266714424 | 3590.310122713699 | 3366.326983403137 | 3671.422245622316 | 3282.2898689321746 | 3395.7097380778164 | 3348.5924826787154 | 2792.0965986869987 | 3074.1428225438763 | 3707.2151546530704 |
| SE_132_4_19 | SE_132 | SE_132_4 | TGTGAGTTATTACTTCCCCC | 2288.9366896456872 | 2468.4994370457234 | 2653.4846798990197 | 2502.416225436188 | 2367.246952412036 | 2518.363489387464 | 2281.460499061053 | 1941.756069602404 | 2759.8582434806276 | 2759.2912861697887 | 1760.6774470468936 | 2747.3319571270563 |
| SE_132_4_27 | SE_132 | SE_132_4 | CATGGAAGAGAAGGGAGCAG | 5569.529189436168 | 5732.515291860029 | 5398.8093580230525 | 5393.92368925273 | 5675.241237615055 | 5607.62534117821 | 5235.52138110001 | 4995.247844098882 | 5443.349753068291 | 5129.292863802017 | 4602.410846580579 | 5335.650615510383 |
| SE_132_4_28 | SE_132 | SE_132_4 | ACTTATCCAGATTCAAACCT | 197.2473851872704 | 231.8590564350639 | 248.8567693839051 | 255.0564639550145 | 282.1181403860222 | 251.22993186183854 | 198.01355274869513 | 140.85767009287156 | 234.42811334584482 | 425.0110488340784 | 194.84830413985622 | 261.15322786378863 |
| SE_132_4_9 | SE_132 | SE_132_4 | CTAGCCAACGACCAGCCTGT | 139.80721257779055 | 122.92527560996923 | 130.3535458677598 | 135.33608291490566 | 109.24123481112889 | 129.0802063703929 | 187.25194662104866 | 94.5353490556185 | 85.2465866712163 | 104.24799311024564 | 100.9455069640219 | 450.3442329384444 |
| SE_133_1_13 | SE_133 | SE_133_1 | TAGCCCAGGGAGACTCACCA | 1425.1665468201131 | 1534.0674639130307 | 1574.1178190394633 | 1324.7320423785957 | 1503.9230190502988 | 1266.5453806276137 | 1239.737025904874 | 1284.7353936658553 | 1356.8415045168595 | 853.6671323922913 | 1112.7481465336368 | 972.6506886660217 |
| SE_133_1_3 | SE_133 | SE_133_1 | AGAGTTTCTGATTCTGTAAG | 10.837768416882989 | 12.992102300240651 | 27.650752153767232 | 18.218318853929606 | 26.51486281823517 | 2.598930329605226 | 8.60928490211718 | 15.12565584889896 | 57.71904305863604 | 21.141201399979888 | 0.0 | 0.0 |
| SE_133_1_34 | SE_133 | SE_133_1 | AATGCAGTGAGTCTCTCAAG | 337.054597765061 | 348.78797713722975 | 379.2103152516649 | 308.4101120272369 | 422.1166160663039 | 402.83420108881006 | 262.583189514574 | 380.9774566941426 | 233.54012806801964 | 244.94633346183593 | 372.0898338092435 | 156.69193671827318 |
| SE_133_1_6 | SE_133 | SE_133_1 | GCTTAGCTATCAAGCTCCAC | 1499.9471488966058 | 1295.2126600855295 | 1325.261049655558 | 1503.0113054491926 | 1469.9839946429577 | 1364.4384230427438 | 1454.9691484578034 | 1387.7789241364796 | 1284.9146970130207 | 1349.3918548745783 | 1160.8733300862518 | 856.5825873932267 |
| SE_133_1_7 | SE_133 | SE_133_1 | GCATTCAGTGAGAACGATGA | 1634.3354772659548 | 1652.9951695844643 | 1610.6563129569413 | 1620.1290695101686 | 1691.6482478034038 | 1689.304714243397 | 1449.58834539398 | 1257.3201424397262 | 1587.7176767514036 | 984.888382461132 | 1351.0264943673162 | 1043.4522304424265 |
| SE_133_2_1 | SE_133 | SE_133_2 | TCTAAAAGTAACATGATGCA | 1064.2688585379096 | 1182.2813093218992 | 1120.8429890902075 | 1097.0030567044755 | 1042.5644060130069 | 950.3421905256444 | 1068.627488475295 | 976.5501557445391 | 905.7449833816731 | 1046.8539727714178 | 826.344615147342 | 838.0116911895794 |
| SE_133_2_13 | SE_133 | SE_133_2 | GTAGAAATCTCTCATTGCAC | 458.4376040341504 | 484.70535504743964 | 501.6636462183484 | 425.52787608821296 | 427.4195886299509 | 615.9464881164386 | 498.26236371003176 | 296.8409960346421 | 511.4795200272978 | 287.2287362617957 | 302.83652089206566 | 752.1212962477113 |
| SE_133_2_21 | SE_133 | SE_133_2 | TTAAATCCCACAAGAATGCA | 17.340429467012783 | 29.981774539016886 | 38.51354764274722 | 24.724861301761607 | 37.120807945529236 | 42.44919538355203 | 1.0761606127646475 | 22.68848377334844 | 25.751573056929924 | 0.7290069448268927 | 32.86597901154201 | 10.446129114551544 |
| SE_133_2_22 | SE_133 | SE_133_2 | GCTTGGCAGAGGCATCACCA | 4910.592869689683 | 4854.0492978668335 | 4609.775394778051 | 4816.142719885248 | 5186.307167246799 | 4344.54520099007 | 4890.073824402558 | 5096.400667588393 | 4781.800721088539 | 3854.9887242446084 | 4533.157533663402 | 3985.7785977077783 |
| SE_133_2_9 | SE_133 | SE_133_2 | GCATGTGTAGAAAGGTCCCA | 2937.03524097529 | 3234.034080275288 | 3185.761658859039 | 2965.6820477218266 | 3158.4504589081735 | 3346.555954421663 | 2827.073929732729 | 3027.9672302514605 | 3400.9836140704 | 2768.768376452538 | 2872.2518086158325 | 2623.1390887651655 |
| SE_134_1_10 | SE_134 | SE_134_1 | AAACAATCATCAGTTTACAG | 19.50798315038938 | 33.97934447755247 | 28.63827901640178 | 33.834020728726415 | 33.939024407341016 | 14.727271867762948 | 13.990087965940416 | 76.57363273505099 | 16.871720278678225 | 87.48083337922712 | 24.64948425865651 | 4.642724050911798 |
| SE_134_1_19 | SE_134 | SE_134_1 | AGATCTGTATCTAGGTTGGG | 101.8750231187001 | 168.89732990312845 | 187.63010390056337 | 147.04785932100324 | 154.84679885849337 | 133.41175691973496 | 136.67239782111022 | 133.2948421684221 | 190.02884945458632 | 225.99215289633673 | 137.3328408696577 | 69.64086076367697 |
| SE_134_1_5 | SE_134 | SE_134_1 | AATGGGCTGTGAGCTGTGCA | 1787.1480119440048 | 1730.9477833859082 | 1727.1844827478176 | 1892.1025438295462 | 1663.0121959597097 | 1563.6897483124778 | 1737.9993896149056 | 1174.1290352707817 | 1673.852248700445 | 1511.9604035709754 | 989.5007252403542 | 1511.2066785717902 |
| SE_134_1_8 | SE_134 | SE_134_1 | TTTGGCCTCCCAAAAAGTGC | 193.9960546622055 | 188.8851795958064 | 177.75483527421792 | 157.45832723753446 | 168.63452752397566 | 79.7005301078936 | 195.86123152316583 | 264.6989773557318 | 188.252878898936 | 214.32804177910643 | 223.01914329260651 | 85.89039494186825 |
| SE_134_1_9 | SE_134 | SE_134_1 | TACTATTACCCATTTAGCAG | 184.24206308701082 | 170.89611487239625 | 174.7922546863143 | 148.34916781056964 | 138.9378811675523 | 439.2192257032832 | 179.71882233169612 | 192.85211207346174 | 108.33420389467071 | 255.88143763423932 | 144.37555065784528 | 194.9944101382955 |
| SE_134_2_11 | SE_134 | SE_134_2 | ACCAACATCAACACACACAG | 3535.280057587231 | 3090.121562488007 | 3127.497573963601 | 3145.26261928199 | 3313.2972577666665 | 2316.513233788125 | 3020.7828400303656 | 2930.5958207241733 | 2924.135519878284 | 2456.7534040666283 | 2976.718670473948 | 2610.3715976251583 |
| SE_134_2_24 | SE_134 | SE_134_2 | GACAAGTTCCTACAAGTGAA | 209.16893044584168 | 182.888824688003 | 214.29332919169605 | 270.6721658298113 | 200.45236290585788 | 401.9678909789416 | 231.3745317443992 | 266.5896843368442 | 278.8273772371033 | 227.45016678599052 | 95.07658214053225 | 674.3556683949387 |
| SE_134_2_4 | SE_134 | SE_134_2 | GAGGATCACTTGATGCTAGG | 1149.8872290312852 | 1260.2339231233432 | 1195.8950306504328 | 1184.1907255054243 | 1171.9569365659945 | 1088.0854979947214 | 982.5346394541232 | 1252.593374986945 | 1058.4784511676023 | 935.3159102129033 | 1933.2238368574892 | 730.0683570058802 |
| SE_134_2_6 | SE_134 | SE_134_2 | AAAGATGTAGATAAAACAGG | 80.19948628493412 | 104.9362108865591 | 98.75268626345441 | 74.17458390528482 | 69.99923784014085 | 126.48127604078768 | 77.48356411905462 | 23.633837263904624 | 82.5826308377408 | 36.45034724134463 | 52.82032341140680 4 | 54.55200759821362 |
| SE_134_2_7 | SE_134 | SE_134_2 | ACAGAGGAGACAGATCAAGG | 2398.3981506562054 | 2747.329940258581 | 2699.8984424428436 | 2624.7392234554295 | 2879.514102060339 | 2528.759210705885 | 2301.907550703581 | 2967.4646068558645 | 2304.321795956316 | 2291.9978345357504 | 2707.9219135581225 | 1991.7286178411612 |
| SE_135_1_10 | SE_135 | SE_135_1 | TCAGTGTTAGCGTCCACAGG | 1697.194534083876 | 1745.9386706554167 | 1746.9350200005083 | 1493.9021460222277 | 1765.8898636944623 | 2025.4330368723397 | 1450.664506006745 | 1368.8718543253558 | 1736.899203426032 | 1533.1016049709553 | 1478.9690555193906 | 1382.3710861589877 |
| SE_135_1_25 | SE_135 | SE_135_1 | AGGGTAGTCAGGGATCAGAG | 2275.9313675454277 | 2717.348165719564 | 2672.247690289076 | 2456.870428301364 | 2451.033918917659 | 2211.6897104940476 | 2325.583084184403 | 2134.6081816758656 | 2388.6803973497067 | 3267.409126714133 | 2310.0088105255245 | 2156.5453216485303 |
| SE_135_1_5 | SE_135 | SE_135_1 | GATGGAGGCGGAAGGAGAGA | 43379.251865415856 | 45403.39996940254 | 44826.806875569855 | 45977.83155336006 | 45955.560236565194 | 50667.87939587362 | 47478.05391395072 | 55090.47466216168 | 48197.17692451674 | 54585.85300780324 | 62981.779850796775 | 55900.718935003504 |
| SE_135_1_6 | SE_135 | SE_135_1 | TGAGCCTGTCTCCATACTGG | 2964.1296620174976 | 2979.1889966936446 | 2893.453707519214 | 2885.00092136871 | 2726.7884922273047 | 2756.5987696012767 | 2862.5872299539624 | 2824.716229781881 | 2717.2349501450194 | 2660.875348618158 | 2297.097175913847 | 2698.5833545924825 |

| | | | | | | | | | | | | | | | |
|---|---|---|---|---|---|---|---|---|---|---|---|---|---|---|---|
| SE_135_1_7 | SE_135 | SE_135_1 | GCACAGAGATTGCAGAGAAG | 2545.7918011258143 | 2561.442938116676 | 2642.62188441004 | 2588.3025857475704 | 2973.9070136932564 | 2552.149583672332 | 2676.4114439456785 | 3242.5624726077144 | 2451.7273520752938 | 3123.065751638408 | 3766.6759517156543 | 2561.6229950905845 |
| SE_135_2_11 | SE_135 | SE_135_2 | TGAAACAAAGAAACACTGGG | 304.541292514412 | 514.6871295864565 | 504.626226806252 | 321.4231969229009 | 394.5411587353393 | 777.9464786618311 | 224.91756806781132 | 194.7428190545741 | 439.55271252345904 | 398.76679882031027 | 241.7997027277734 | 417.8451645820618 |
| SE_135_2_21 | SE_135 | SE_135_2 | GGATGCTTCAGAGTGAGCGA | 451.93494298402067 | 603.6330607188733 | 610.2916011081483 | 514.0168533787282 | 671.3563265577145 | 380.31013823223145 | 515.4809335142661 | 574.7749222581605 | 447.54458002388554 | 710.0527642613935 | 481.2518355261509 | 400.43494939114254 |
| SE_135_2_23 | SE_135 | SE_135_2 | CGAGTTCCTTCATCTGTAAA | 226.50935991285448 | 321.8043800521146 | 357.48472427370496 | 240.7420705697841 | 320.29954284428084 | 332.6630821894689 | 146.35784333599204 | 303.4584704685354 | 454.64846224648693 | 397.3087849306565 | 463.645061055682 | 327.31204558928175 |
| SE_135_2_8 | SE_135 | SE_135_2 | GTTTAAGTCTTTTAGCACAT | 19.50798315038938 | 32.97995199291857 | 25.675698428498144 | 10.410467916531204 | 67.87804881468203 | 92.69518175591973 | 23.675533480822246 | 22.68848377334844 | 92.35046889381766 | 6.561062503442034 | 9.390279717583432 | 64.99813671276517 |
| SE_135_2_9 | SE_135 | SE_135_2 | GTGAACTCTGCCACAAACAG | 605.831254503759 | 637.6124051964258 | 621.1543965971282 | 676.6804145745282 | 640.5990856885617 | 548.3742995467027 | 744.703144033136 | 614.4797688615203 | 613.5978269771923 | 782.2244517992558 | 1007.1074997108232 | 690.6052025731299 |
| SE_135_3_19 | SE_135 | SE_135_3 | CTAATACCATTTACCCACCG | 300.2061851476588 | 320.8049875674807 | 299.22063937826687 | 351.3532921829281 | 306.51181417879855 | 378.57751801249464 | 321.7720232166296 | 412.17412188249665 | 561.2066955855073 | 490.62167386849876 | 572.8070627725893 | 436.41606078570896 |
| SE_135_3_21 | SE_135 | SE_135_3 | ATAGTAACAAAGCTTAAAGA | 39.01596630077876 | 37.97691441608806 | 41.47612823065085 | 22.12224432262881 | 29.69664635642339 | 5.197860659210452 | 21.52321225529295 | 5.67212094333711 | 35.51941111300679 | 8.748083337922711 | 10.564064682281362 | 44.10587848366208 |
| SE_135_3_7 | SE_135 | SE_135_3 | TGGACCATCCATTTGAACTG | 508.2913387518122 | 441.7314782081821 | 474.01289406458113 | 543.9469486387554 | 557.8727136956679 | 544.909059107229 | 474.5868302292095 | 549.2503780131435 | 471.5201825251652 | 402.41183354444473 | 622.1060312899024 | 618.642979783997 |
| SE_135_3_9 | SE_135 | SE_135_3 | TTTGTGCTTGAAAAGAGCAC | 47.686181034285156 | 45.972054293159225 | 49.376343131727204 | 76.77720088441762 | 34.999618920070425 | 86.63101098684088 | 52.73187002546773 | 50.10373499947781 | 45.28724916908366 | 18.954180565499207 | 16.432989505771005 | 12.767491140007444 |
| SE_135_4_10 | SE_135 | SE_135_4 | GTTTCCATAATTCAACTCAG | 1.083776841688299 | 3.9975699385355847 | 9.87526862634544 | 5.205233958265602 | 2.1211890254588135 | 0.0 | 11.837766740411123 | 0.945353490556185 | 1.7759705556503396 | 48.84346530340181 | 0.0 | 0.0 |
| SE_135_4_15 | SE_135 | SE_135_4 | ACTCAAGCAATGAGCCCAGT | 407.5000924748004 | 502.6944197708498 | 470.062786614043 | 465.8684392647714 | 466.66158560093896 | 601.2192162486757 | 475.6629908419742 | 423.5183637691709 | 618.0377533663182 | 392.20573631686824 | 338.0500698330036 | 424.8092506584295 |
| SE_135_4_2 | SE_135 | SE_135_4 | CAGCACCAACAAGATCCTGA | 2.167553683376598 | 6.9957473924372735 | 5.925161175807264 | 3.9039254686992013 | 20.15129574185873 | 0.0 | 0.0 | 9.45353490556185 | 1.7759705556503396 | 2.9160277793075706 | 0.0 | 0.0 |
| SE_135_4_20 | SE_135 | SE_135_4 | AAAAATAGCCTAGTGCCAGT | 71.52927155142773 | 75.95382883217611 | 62.21419234597627 | 85.88636031138243 | 69.99923784014085 | 77.96790988815678 | 59.18883370205561 | 79.40969320671954 | 72.81479278166393 | 115.91210422747594 | 70.42709788187574 | 89.37243798005211 |
| SE_135_4_23 | SE_135 | SE_135_4 | TTTACAATAGCCAAACCCAC | 95.37236206857031 | 127.92223803313871 | 124.42838469195254 | 88.48897729051524 | 73.18102137832906 | 72.77004922894633 | 54.88419125099702 | 48.213028018365435 | 156.28540889722987 | 37.17935418617152 | 139.68041079905356 | 103.3006101327875 |
| SE_136_1_1 | SE_136 | SE_136_1 | GTGCCACATACAAGACCAGG | 896.2834480762232 | 859.4775367851507 | 875.9363271568405 | 893.9989323321171 | 933.3231712018779 | 799.6042314085413 | 993.2962455817697 | 757.2281459355042 | 831.1542200443589 | 1495.9222507847837 | 677.2739246307051 | 879.7962076477856 |
| SE_136_1_15 | SE_136 | SE_136_1 | GTAGTGGGTGACTTGTCCTA | 1265.851351091933 | 1272.22663293895 | 1329.2111571060964 | 1254.4613839420101 | 1274.8346043007468 | 1885.9571091835257 | 1345.2007659558094 | 1270.5550913075126 | 1299.1224614582234 | 1547.6817438674932 | 1501.2709698486512 | 1894.2314127720135 |
| SE_136_1_16 | SE_136 | SE_136_1 | GCTCTCTTCCTGCTGCACAG | 69.36171786805113 | 86.94714616314897 | 100.72773998872348 | 76.77720088441762 | 75.30221040378788 | 45.048125713157255 | 117.30150679134657 | 125.7320142439726 | 87.02255722686664 | 173.50365286880046 | 95.07658214053225 | 30.177706330926686 |
| SE_136_1_19 | SE_136 | SE_136_1 | ACCAATCTCAGTTTGCACAG | 848.597267041938 | 821.5006223690626 | 908.5247136237805 | 843.2479012390274 | 721.2042686559965 | 855.9143885499878 | 768.3786775139583 | 620.1518898048573 | 832.9301906000093 | 702.0336878682976 | 543.4624386551411 | 756.764020298623 |
| SE_136_1_8 | SE_136 | SE_136_1 | GAACATTTAACACTACAAAA | 108.37768416882989 | 175.89307729556572 | 188.61763076319792 | 156.15701874796804 | 152.72560983303458 | 115.21924461249836 | 162.50025252746178 | 298.7317030157545 | 197.13273167718768 | 91.85487504818848 | 158.4609702342204 | 154.37057469281729 |
| SE_136_2_1 | SE_136 | SE_136_2 | GTTTGGTCTCGTACAACAAA | 47.686181034285156 | 63.961119016569356 | 65.1767729338799 | 63.76411598875362 | 84.84756101835254 | 38.11764483420998 | 44.12258512335055 | 21.743130282792254 | 26.639558334755094 | 43.01140974478667 | 15.259204541073078 | 48.74860253457388 |
| SE_136_2_11 | SE_136 | SE_136_2 | ACTTTGGTCACCTAGACCAG | 207.0013767624651 | 168.89732990312845 | 182.69246958739063 | 175.67664609146405 | 237.57317085138712 | 213.97859713749696 | 133.4439159828163 | 129.51342820619735 | 190.02884945458632 | 146.5303959102054 | 191.32694924576242 | 197.3157721637514 |
| SE_136_2_14 | SE_136 | SE_136_2 | TCAGGGCCAATGCCAGCCCA | 4337.274920436572 | 3302.9921617150267 | 3352.653698644277 | 4014.5366903123454 | 3584.8094530253948 | 4049.9997636348107 | 4444.543330717994 | 3321.026812323878 | 3520.8616265767982 | 3173.3672308314635 | 2537.7230936769224 | 3128.035329301824 |
| SE_136_2_26 | SE_136 | SE_136_2 | TAAGGTAGTGTGACCAGCCA | 6808.286119485893 | 5776.48856118392 | 5979.475153252164 | 6366.001130958831 | 6159.9329299323945 | 6333.593213247937 | 6423.60269759218 | 6386.808182197586 | 5360.76712223055 | 6711.966941021201 | 6095.465321676345 | 5399.4880712104205 |
| SE_136_2_7 | SE_136 | SE_136_2 | GCTAGGCCCTGAAATTCCAG | 60.69150313454474 | 135.9173779102099 | 113.56558920297256 | 72.87327541571842 | 130.45312506571702 | 213.11228702762855 | 49.50338818717378 | 23.633837263904624 | 260.17968640277473 | 131.22125006884067 | 76.29602270536539 | 121.87150633643469 |
| SE_136_3_22 | SE_136 | SE_136_3 | AGTCGAGACACAGCTCTGAA | 378.2381177492163 | 451.72540305452105 | 444.3870881855448 | 377.3794619742561 | 453.9344514481861 | 608.149697127623 | 364.81844772721547 | 455.6603824480812 | 300.1390239049074 | 249.3203751307973 | 461.2974911262861 | 291.3309341947153 |
| SE_136_3_28 | SE_136 | SE_136_3 | TCAGACAGTCACCAACCTCA | 5530.513223135389 | 4831.063270720254 | 5010.711301007676 | 4976.203664101915 | 4761.008767642307 | 6684.448807744642 | 5436.763415686999 | 4592.527257121947 | 4395.52712523459 | 4815.090870581626 | 4885.293023072781 | 5156.90573955028 |
| SE_136_3_3 | SE_136 | SE_136_3 | ATAACTGTATGTCTCTAACC | 109.46146101051819 | 88.94593113241676 | 94.80257881291622 | 101.50206218617923 | 120.90777445115236 | 84.89839076710406 | 122.6823098551698 | 159.76473990399526 | 80.80666028209045 | 29.8892847379026 | 174.89395973999143 | 100.9792481073316 |
| SE_136_3_6 | SE_136 | SE_136_3 | GCATGGAGTTAGCATACTGC | 9067.960834405996 | 7545.413258985916 | 7636.545228752929 | 7673.816162973063 | 7195.073174356296 | 7529.101164866341 | 8368.2249248579 | 7421.024900866052 | 6900.533593979394 | 7114.3787745656455 | 7570.913022301642 | 6858.464104209454 |
| SE_137_1_1 | SE_137 | SE_137_1 | TACCAGAAGTAATTAAGTAA | 114.88034521895969 | 174.89368481093183 | 174.7922546863143 | 174.37533760189766 | 137.87728665482288 | 165.46523098486605 | 172.1856980423436 | 149.36585150787724 | 257.51573056929925 | 149.44642368951298 | 193.67451917515828 | 175.26283292192036 |
| SE_137_1_18 | SE_137 | SE_137_1 | TTAGAACTGCTGAGAGCATG | 549.4748587359676 | 513.6877371018227 | 502.6511730809829 | 556.9600335344194 | 425.2983996044921 | 649.7325824013066 | 523.0140578036187 | 400.82987999582247 | 449.3205505795359 | 416.2629654961557 | 462.471276090984 | 446.86218990026055 |
| SE_137_1_2 | SE_137 | SE_137_1 | TGGTAGTCCTAATCTCTGTA | 3.2513305250648967 | 3.9975699385355847 | 1.975053725269088 | 3.9039254686992013 | 3.1817835381882205 | 0.0 | 1.0761606127646475 | 20.79777679223607 | 1.7759705556503396 | 2.187020834480678 | 0.0 | 1.1606810127279494 |
| SE_137_1_5 | SE_137 | SE_137_1 | ACTGGAGAGTGGAAAGTTGG | 19454.878085146654 | 21634.848507354585 | 21375.018941724706 | 20542.455816295198 | 21215.072038126324 | 21361.474689135222 | 20242.581126103018 | 20724.03921997269 | 22319.50995813564 | 21801.681691993053 | 20284.177974944912 | 21999.547915245552 |
| SE_137_1_9 | SE_137 | SE_137_1 | TTTTGAACTGAGGGTTTTGG | 5.4188842084414945 | 31.980559508284678 | 40.488601368016305 | 37.737946197425615 | 13.787728665482287 | 26.85561340592067 | 1.0761606127646475 | 14.180302358342775 | 39.95933750213264 | 3.6450347241344634 | 0.0 | 42.945197470934126 |
| SE_138_1_24 | SE_138 | SE_138_1 | GGACAGCGGCCACGGGCCTG | 217.8391451793481 | 218.86695413482326 | 245.89418879600146 | 192.59365645582727 | 329.8448934588455 | 317.0695002118376 | 166.80489497852037 | 318.58412631743437 | 290.3711858488305 | 152.36245146882055 | 239.45213279837753 | 522.3064557275773 |
| SE_138_1_32 | SE_138 | SE_138_1 | CAGGGATACTTGGGGTTGCT | 4645.067543476049 | 4733.122807226132 | 4669.027006536125 | 4778.404773687823 | 4660.252288933013 | 4029.208320997969 | 4541.397785866812 | 4443.16140561407 | 5131.666920551656 | 5080.449398498615 | 4036.646493596178 | 4418.712615455303 |
| SE_138_1_36 | SE_138 | SE_138_1 | AACCCCAAGTATCCCTGGTG | 18.42420630870108 | 30.98116702365078 | 22.713117840594514 | 32.53271223916001 | 24.393673792776355 | 0.8663101098684087 | 23.675533480822246 | 0.945353490556185 | 10.655823333902038 | 34.992333351690846 | 45.77761362321923 | 1.1606810127279494 |
| SE_138_1_42 | SE_138 | SE_138_1 | ATCCCTGGTGGGGCAGAGCG | 67.19416418467453 | 103.93681840192521 | 107.6404280271653 | 83.28374333224963 | 129.39253055298764 | 19.05882241710499 | 62.417315540349556 | 93.58999556506231 | 78.14270444861494 | 248.5913681859704 | 77.46980767006332 | 23.21362025455899 |
| SE_138_1_50 | SE_138 | SE_138_1 | GGCCAGCGCGGCCCGTCCCA | 3.2513305250648967 | 10.993317330972857 | 10.862795488979984 | 10.410467916531204 | 6.363567076376441 | 0.0 | 5.380803063823238 | 1.89070698111237 | 0.0 | 8.019076393095819 | 23.47569929395858 | 0.0 |
| SE_138_2_10 | SE_138 | SE_138_2 | CTTGCAAGTTCAAGACTGTG | 1540.0468920390726 | 1454.116065142319 | 1436.8515851332616 | 1648.7578562806293 | 1454.0750769520166 | 1345.3796006256387 | 1515.2341427726237 | 1517.2923523426768 | 1525.5587073036418 | 1560.8038688743773 | 1467.2312058724112 | 1547.1877899663566 |
| SE_138_2_25 | SE_138 | SE_138_2 | TGCTGCCTGAGCTCCAACCG | 1383.9830268359576 | 1427.1324680572038 | 1367.7247047488436 | 1503.0113054491926 | 1355.4397872681818 | 978.0641140414334 | 1492.6347699045662 | 1446.390840550963 | 1381.7050922959643 | 1541.120681364051 | 1012.9764245343127 | 1231.4825545043543 |
| SE_138_2_31 | SE_138 | SE_138_2 | GCTCCACTATATACTGTAGG | 107.2939073271416 | 163.900367479959 | 149.11655625781614 | 114.51514708184324 | 130.45312506571702 | 227.8395588953915 | 55.960351863761666 | 134.24019565897828 | 229.98818695671898 | 39.36637502065221 | 120.8998513638867 | 284.3668481183476 |
| SE_138_2_32 | SE_138 | SE_138_2 | CTACCTCCTACAGTATATAG | 33.59708209233727 | 59.96354907803377 | 62.21419234597627 | 58.55888203048802 | 51.96913112374093 | 136.8769973592086 | 18.294730416999005 | 27.415251226129364 | 63.04695472558706 | 74.35870837234305 | 1.173784964697929 | 25.534982280014887 |
| SE_138_2_5 | SE_138 | SE_138_2 | AGGACTTTAGACTTATGGGA | 126.80189047753097 | 194.88153450360974 | 187.63010390056337 | 193.89496494539367 | 208.93711900769313 | 129.0802063703929 | 154.96712823810924 | 242.95584707293955 | 144.74160028550267 | 271.19058347560406 | 293.4462411744823 | 224.01143545649424 |
| SE_138_3_10 | SE_138 | SE_138_3 | ATAGGACTGTACCCTTTTAC | 1436.004315236996 | 1641.0024597688575 | 1572.142765314194 | 1359.8673715968885 | 1321.5007628608407 | 1367.9036634822173 | 1445.2837029429215 | 1195.872165553574 | 1186.348331174427 | 1067.9951741713978 | 1025.88805914599 | 1262.820941848009 |
| SE_138_3_2 | SE_138 | SE_138_3 | GTTGGGACGCTTTAATGTCA | 393.4109935328525 | 463.7181128701278 | 446.3621419108139 | 480.1828326500018 | 461.35861303729195 | 297.1443676848642 | 442.3020118462701 | 559.6492664092615 | 410.24919835522843 | 381.27063214446486 | 583.3711274548707 | 205.44053925284706 |
| SE_138_3_3 | SE_138 | SE_138_3 | ATTGATACAACTTTATGGCA | 53.105065242726646 | 55.96597913949819 | 55.301504307534465 | 61.16149900962082 | 65.75685978922321 | 34.65240439473635 | 24.75169409358689 | 127.62272122508497 | 43.51127861343332 | 10.206097227576498 | 71.60088284657367 | 26.695663292742836 |
| SE_138_3_6 | SE_138 | SE_138_3 | GCAGATTCGAACATAACAAC | 2035.3329086906253 | 2102.7217876697177 | 2216.0102797519166 | 1976.6875956513622 | 2179.521723658931 | 1754.2779724835277 | 2009.1918640315969 | 1533.363361682132 | 2053.0219623317926 | 1962.486695473995 | 1711.3784785295804 | 2020.74564315936 |
| SE_138_3_9 | SE_138 | SE_138_3 | TCAGTGTGTTAACATACAAT | 1826.1639782447837 | 1739.9423157476133 | 1658.0576023633994 | 1842.652821226023 | 1789.2229429745091 | 2286.1923799427304 | 1959.688475844423 | 2043.854246582472 | 1809.713996207696 | 1757.6357439776382 | 1755.982307188102 | 1591.2936684500187 |
| SE_139_1_13 | SE_139 | SE_139_1 | CAAGAGGTAACAGAGGAGCA | 5392.873564240976 | 5305.774700921354 | 5114.401621584304 | 5695.827258832135 | 5895.844896262772 | 7055.22953476832 | 6097.526031924493 | 6167.486172388551 | 5495.740884459976 | 6702.489850738451 | 6552.06767294384 | 7549.069306782583 |
| SE_139_1_26 | SE_139 | SE_139_1 | TCTTGCAGACAAGGAAAGTG | 617.7527997623304 | 643.6087601042292 | 623.1294503223972 | 696.2000419180242 | 601.3570887175737 | 661.8609239394642 | 658.6102950119642 | 535.0700756548007 | 570.9745336415841 | 823.7778476543887 | 347.440349550587 | 817.1194329604764 |
| SE_139_1_3 | SE_139 | SE_139_1 | CAGCCTAGATGAGCAGAAGT | 1102.201047997 | 1142.3056099365433 | 1172.1943859472037 | 1241.448299046346 | 1140.1391011841122 | 1042.1710621716957 | 1239.737025904874 | 1039.8888396118034 | 1224.5316981209091 | 1054.1440422196868 | 967.1988109110936 | 1523.9741697117977 |
| SE_139_1_31 | SE_139 | SE_139_1 | CTTTCCTTGTCTGCAAGAAG | 32.51330525064897 | 16.989672238776237 | 21.725590977959968 | 29.93009526002721 | 36.06021343279983 | 20.791442636841808 | 43.0464245105859 | 28.36060471668555 | 1.7759705556503396 | 2.187020834480678 | 16.432989505771005 | 110.26469620915519 |
| SE_139_1_32 | SE_139 | SE_139_1 | ACTGAGAGCCCTTTTCAGCT | 670.8578650050571 | 1032.3724366268148 | 973.7014865576605 | 759.9641579067778 | 902.5659303327251 | 684.3849867960429 | 575.7459278290864 | 919.828946311168 | 800.074735320478 | 961.5601602266714 | 909.683347640895 | 1067.8265317097134 |
| SE_139_2_16 | SE_139 | SE_139_2 | GACACATGCCAAGAATTCCA | 2363.71729172218 | 2381.5522908825747 | 2345.376298757042 | 2295.5081755951305 | 2411.7919219466708 | 2015.9036256637871 | 2304.05987192911 | 1921.9036463007242 | 2625.772466529027 | 1872.0898343154604 | 2299.444745843243 | 2068.333564681206 |
| SE_139_2_19 | SE_139 | SE_139_2 | ACTGCCTCACCCCAGCTGGG | 405.3325387914238 | 418.7454510616025 | 445.37461504817935 | 460.66320530650574 | 243.93673792776355 | 341.32618328815306 | 390.64630243356703 | 340.3272566002266 | 440.4406978012842 | 458.5453682961155 | 417.8674474324627 | 582.6618683894307 |
| SE_139_2_24 | SE_139 | SE_139_2 | AAATCCTTCAGACACAGCAA | 15.172875783636185 | 39.975699385355846 | 38.51354764274722 | 41.641871666124814 | 30.757240869152795 | 0.0 | 31.208657770174778 | 18.9070698111237 | 21.311646667804077 | 10.93510417240339 | 11.73784964697929 | 9.285448101823595 |
| SE_139_2_26 | SE_139 | SE_139_2 | GCTCACAGAAATGACCTCCG | 1692.859426717123 | 2125.7078148162973 | 2097.5070562357714 | 1785.3952476851014 | 2107.4012967933313 | 2071.3474726953655 | 1609.9362766959125 | 1811.2972879056504 | 2079.661520666548 | 1841.471542632731 | 1692.5979190944136 | 1741.0215190919241 |
| SE_139_2_35 | SE_139 | SE_139_2 | TTGTAGTGAGGGGAGCACAA | 112.71279153558308 | 158.9034050567895 | 171.82967409841066 | 147.04785932100324 | 160.14977142214042 | 184.52405340197106 | 88.2451702467011 | 190.96140509234937 | 71.92680750383876 | 158.19450702743572 | 185.45802442227279 | 143.92444557826573 |
| SE_139_3_14 | SE_139 | SE_139_3 | TTTTGGGTCCTGTTGCTGGA | 44.434850509220254 | 53.96719417023039 | 40.488601368016305 | 41.641871666124814 | 55.15091466192915 | 30.320853845394307 | 22.599372868057596 | 36.868786131691216 | 27.527543612580263 | 38.63736807582531 | 12.911634611677218 | 47.58792152184593 |
| SE_139_3_18 | SE_139 | SE_139_3 | TCTAGATGCAGCTCACAGGG | 518.0453303270069 | 598.6360982957038 | 661.6429979651446 | 582.9862033257474 | 613.0236283575971 | 634.1390004236752 | 443.3781724590348 | 641.8950200876496 | 861.3457194904147 | 801.9076393095819 | 386.17525338561865 | 672.0343063694827 |
| SE_139_3_19 | SE_139 | SE_139_3 | GGATGTACGAGCACAGACCC | 2902.3543820412647 | 2338.578414043317 | 2276.249418372624 | 2640.3549253302267 | 2378.913492052059 | 2034.0961379710236 | 3126.246580081301 | 2669.6782573306664 | 2217.299238729449 | 2504.8678624252034 | 2329.963154925389 | 1844.3221292247117 |
| SE_139_3_28 | SE_139 | SE_139_3 | GAGCACCCAGACTTGGCTGA | 777.0679954905103 | 875.467816539293 | 940.125573228086 | 821.1256569163987 | 982.1105187874307 | 1160.8555472236676 | 801.7396565096624 | 1007.7468209328932 | 1314.2182111812513 | 937.502931047384 | 947.2444665112288 | 1533.2596178136212 |
| SE_139_3_4 | SE_139 | SE_139_3 | AGTGGAACAATGGGATACAG | 916.8752080683008 | 961.4155702178082 | 969.7513791071223 | 880.9858474364531 | 845.2938266453372 | 708.6416698723583 | 834.0244748926018 | 951.9709649900783 | 1046.9346425558751 | 783.6824656889096 | 1201.9558038506793 | 836.8510101768516 |
| SE_139_4_1 | SE_139 | SE_139_4 | TTTATCTGCACTTCTCTCCA | 46.60240419259685 | 51.968409200962604 | 33.5759133295745 | 45.545797134824014 | 49.84794209828212 | 6.93048087894727 | 65.6457973786435 | 63.33868386726439 | 1.7759705556503396 | 13.851131951710961 | 146.72312058724114 | 30.177706330926686 |
| SE_139_4_22 | SE_139 | SE_139_4 | AGAGGCCCAAAACTGCCAGC | 1217.0813932159597 | 1547.0595662132712 | 1507.9535192429487 | 1331.2385848264275 | 1425.4390251083228 | 1349.7111511749808 | 1258.031756321873 | 1612.7730548888517 | 1416.3365181311458 | 1633.7045633570665 | 1788.8482861996438 | 1107.2896861424638 |
| SE_139_4_24 | SE_139 | SE_139_4 | GGTTGTGTCCTCCTCCTCGG | 360.89768828220355 | 465.71689783939564 | 455.2498836745248 | 379.9820789533889 | 303.33003064061035 | 597.7539758092021 | 292.7156866719841 | 221.21271679014728 | 386.27359585394885 | 152.36245146882055 | 105.64064682281361 | 131.15695443825828 |
| SE_139_4_25 | SE_139 | SE_139_4 | CCTTGACTGATAGCCAAAGG | 779.235549173887 | 797.5152027378491 | 807.796973635057 | 783.387710718973 | 861.2027443362783 | 581.2940837217022 | 703.8090407480795 | 1055.0144954607024 | 682.8606786475556 | 534.3620905581123 | 1129.1811360394076 | 634.8925139621883 |
| SE_139_4_4 | SE_139 | SE_139_4 | TGCCACCTCCCAGAGTGCAG | 643.7634439628496 | 566.6555387874191 | 610.2916011081483 | 507.5103109308962 | 672.4169210704439 | 752.8234854756472 | 695.1997558459623 | 885.7962206511454 | 737.9157658727161 | 423.5530349444246 | 731.2680330068098 | 648.8206861149238 |
| SE_14_1_10 | SE_14 | SE_14_1 | ATCCTGGCGTCGGCCGTGCT | 398.829877741294 | 406.75274124599576 | 391.0606376032794 | 456.75927983780656 | 426.3589941172215 | 466.9411492190723 | 416.4741571399186 | 338.43654961911426 | 506.15160836034676 | 473.12550719265334 | 470.68777084386954 | 358.6504329329364 |
| SE_14_1_11 | SE_14 | SE_14_1 | GATCCTGGCGTCGGCCGTGC | 692.533401838823 | 633.6148352578901 | 629.0546114982045 | 478.88152416043533 | 516.509527699221 | 678.320816026964 | 542.3849488333823 | 536.0154291453568 | 792.9708530978767 | 804.8236670888895 | 721.8777532892263 | 750.9606152349833 |
| SE_14_1_13 | SE_14 | SE_14_1 | GGGGCTGCTGCTCATGATCC | 1098.9497174719352 | 1044.3651464424215 | 1154.418902419782 | 1207.6142783176197 | 940.7473327909838 | 1023.1122397545907 | 1277.4026473516365 | 1161.8394398935513 | 940.3764092168548 | 1258.9949937160436 | 1416.7584523904004 | 1124.699901333383 |
| SE_14_1_8 | SE_14 | SE_14_1 | CTGGGCGAGGAGCGTGAGGA | 1871.6826055956922 | 1816.8955370644233 | 1779.5234064674482 | 1767.1769288311718 | 1875.1310985055911 | 2043.6255491795762 | 1924.1751756231897 | 2091.121921110281 | 1640.9967934209137 | 1791.1700634396752 | 2074.0780326212407 | 2216.9007343103835 |
| SE_14_1_9 | SE_14 | SE_14_1 | CGTGAGGAAGGAACCCAGCA | 28651.808363713557 | 26361.974959672913 | 25920.605090431513 | 26718.465907777332 | 25848.8094642411 | 24488.854185760178 | 26177.606905500048 | 23791.711296827507 | 24344.11639157703 | 19471.046489381475 | 21831.226558416784 | 21425.01081394522 |
| SE_14_10_2 | SE_14 | SE_14_10 | GTGAAATGTTTAGTTCTGTA | 314.2952840896067 | 416.7466660923347 | 414.7612823065085 | 379.9820789533889 | 468.7827746263978 | 415.8288527368362 | 283.0302411571023 | 453.7696754669688 | 339.21037612921486 | 578.8315141925527 | 441.3431467264213 | 438.7374228111649 |
| SE_14_10_3 | SE_14 | SE_14_10 | AGATGAGCTCAACAGAAAAC | 1105.452378522065 | 1272.22663293895 | 1223.5457828042001 | 1102.208290662741 | 1229.2290402533824 | 995.3903162388016 | 1071.8559703135888 | 1091.8832815923936 | 1069.1342745015045 | 863.8732296198677 | 897.9454979939158 | 1241.928683618906 |
| SE_14_10_5 | SE_14 | SE_14_10 | TTTGACAGAAGAGAGAAATA | 24.926867358830876 | 47.97083926242702 | 38.51354764274722 | 42.943180155691216 | 62.575076251035 | 72.77004922894633 | 21.52321225529295 | 37.8141396222474 | 31.07948472388094 | 26.244250013768134 | 3.521354894093787 | 18.57089620364719 |
| SE_14_10_6 | SE_14 | SE_14_10 | TTTGGATTGAGTAATTCACG | 115.96412206064798 | 99.93924846338962 | 115.54064292824165 | 132.73346593577284 | 109.24123481112889 | 65.83956834999907 | 74.25508228076067 | 115.33312584785457 | 237.09206917932033 | 99.1449444964574 | 196.02208910455414 | 16.249534178191293 |
| SE_14_11_12 | SE_14 | SE_14_11 | TGAGAGAATCACTTGCGCCT | 3411.729497634765 | 3604.8086920744636 | 3603.4855217534514 | 3734.755365055569 | 3572.0823188726417 | 3193.2190649749546 | 3272.6044234172928 | 3719.020631848032 | 3342.376585733939 | 3963.6107590238153 | 3770.197306609748 | 3508.7387014765914 |
| SE_14_11_15 | SE_14 | SE_14_11 | TGAAGAATACACTGGCCGGG | 390.15966300778763 | 649.6051150120326 | 628.06708463557 | 461.96451379607214 | 564.2362807720444 | 784.0106494309099 | 463.82522410156304 | 634.3321921632001 | 687.3006050366814 | 525.6140072201896 | 484.7731904202447 | 922.7414051187199 |
| SE_14_11_20 | SE_14 | SE_14_11 | CTTTATTAAGAGTTATCGGC | 0.0 | 6.9957473924372735 | 5.925161175807264 | 3.9039254686992013 | 1.0605945127294067 | 0.0 | 3.228481838293942 | 7.56282792444948 | 0.0 | 0.0 | 0.0 | 0.0 |

| | | | | | | | | | | | | | | | |
|---|---|---|---|---|---|---|---|---|---|---|---|---|---|---|---|
| SE_14_11_22 | SE_14 | SE_14_11 | GTATCCACATTCCTGAAAAT | 57.440172609479845 | 92.94350107095235 | 77.02709528549444 | 76.77720088441762 | 90.15053358199957 | 27.72192351578908 | 59.18883370205561 | 71.84686528227006 | 83.47061611556596 | 86.75182643440023 | 57.515463270198524 | 132.31763545098624 |
| SE_14_11_3 | SE_14 | SE_14_11 | GTAATCGCAGCTACTAAGGA | 605.831254503759 | 739.5504386290831 | 764.345791679137 | 663.6673296788642 | 655.4474088667733 | 803.9357819578833 | 733.9415379054896 | 678.7638062193408 | 598.5020772541644 | 392.93474326169513 | 551.6789334080266 | 1224.5184684279866 |
| SE_14_12_10 | SE_14 | SE_14_12 | ATTTATACATATTATCTGAA | 0.0 | 1.9987849692677924 | 0.0 | 0.0 | 0.0 | 6.064170769078861 | 0.0 | 0.945353490556185 | 14.207764445202717 | 0.7290069448268927 | 0.0 | 0.0 |
| SE_14_12_12 | SE_14 | SE_14_12 | ACAATACATGAAGAATTCAC | 619.920353445707 | 743.5480085676188 | 855.1982630415151 | 745.6497645215475 | 709.5377290159731 | 836.8555661328828 | 577.8982490546157 | 680.6545132004532 | 808.0666028209044 | 941.8769727163453 | 656.1457952661424 | 790.4237696677336 |
| SE_14_12_16 | SE_14 | SE_14_12 | TTATAGTCTCATTTTACTAA | 14.089098941947885 | 11.992709815606755 | 11.850322351614528 | 26.02616979132801 | 5.3029725636470335 | 25.989303296052263 | 21.52321225529295 | 1.89070698111237 | 23.087617223454416 | 29.160277793075707 | 21.128129364562724 | 0.0 |
| SE_14_12_20 | SE_14 | SE_14_12 | ACTAAGGGCAGAAGGAAGGA | 1225.751607949466 | 1296.2120525701635 | 1316.3733078918472 | 1132.1383859227683 | 1368.1669214209346 | 1790.662997098001 | 1197.7667620070526 | 1246.921254043608 | 1245.8433447887132 | 1027.8997922059186 | 1018.8453493578024 | 1351.032698815333 |
| SE_14_12_6 | SE_14 | SE_14_12 | TTAGAAATGGTGTTTCTCCT | 7.586437891818092 | 16.989672238776237 | 9.87526862634544 | 15.615701874796805 | 14.848323178211695 | 0.8663101098684087 | 7.533124289352532 | 11.34424188667422 | 20.423661389978903 | 21.141201399979888 | 46.95139858791716 | 1.1606810127279494 |
| SE_14_2_0 | SE_14 | SE_14_2 | GTATTCCCAAAAACATCTCC | 17.340429467012783 | 34.97873696218637 | 39.50107450538176 | 40.34056317655841 | 36.06021343279983 | 128.21389626052448 | 23.675533480822246 | 11.34424188667422 | 30.191499446055772 | 37.908361130998415 | 3.521354894093787 | 1.1606810127279494 |
| SE_14_2_10 | SE_14 | SE_14_2 | TTCTTATTAGGATCTAGCAC | 3.2513305250648967 | 0.9993924846338962 | 3.950107450538176 | 0.0 | 0.0 | 0.0 | 0.0 | 0.0 | 0.0 | 0.0 | 0.0 | 0.0 |
| SE_14_2_14 | SE_14 | SE_14_2 | ATAAGAACAAACCAAATAAG | 121.38300626908948 | 170.89611487239625 | 148.12902939518162 | 128.82954046707366 | 190.90701229129323 | 95.29411208552496 | 160.34793130193248 | 228.77554471459678 | 64.82292528123739 | 127.57621534470621 | 234.7569929395858 | 140.44240254008187 |
| SE_14_2_19 | SE_14 | SE_14_2 | GTAGTCTTATTTTTTAGTGA | 6.5026610501297935 | 32.97995199291857 | 22.713117840594514 | 15.615701874796805 | 37.120807945529236 | 38.11764483420998 | 5.380803063823238 | 20.79777679223607 | 42.62329333560815 | 2.187020834480678 | 9.390279717583432 | 24.37430126728694 |
| SE_14_2_4 | SE_14 | SE_14_2 | GGAAGACAGAGGATCCAAAA | 333.80326723999605 | 481.70717759353795 | 469.0752597514084 | 369.57161103685775 | 478.32812524096244 | 485.1336615263089 | 266.8878319656326 | 452.8243219764126 | 554.102813362906 | 395.8507710410027 | 458.9499211968903 | 513.0210076257537 |
| SE_14_3_1 | SE_14 | SE_14_3 | AGTCCAGTCAGACCTGCTAG | 304.541292514412 | 332.79769738308744 | 359.459777998974 | 356.55852614119374 | 443.328506320892 | 521.518686140782 | 318.54354137833565 | 536.0154291453568 | 370.2898608530958 | 239.1142779032208 | 529.377019078766 | 897.206422838705 |
| SE_14_3_10 | SE_14 | SE_14_3 | GAAGGAAGAGGCTTTGTAGA | 1375.3128121024513 | 1966.8044097595077 | 1979.0038327196262 | 1601.9107506562389 | 2095.734757153308 | 1820.1175408335268 | 1395.7803147557477 | 1881.2534462068081 | 1917.1602148245415 | 1798.4601328879442 | 1876.8821585519886 | 2401.4490153341276 |
| SE_14_3_11 | SE_14 | SE_14_3 | GTACGTTTCAAACAGTTGAG | 1085.9443953716755 | 1136.30925502874 | 1133.6808383044565 | 1067.0729614444483 | 1228.168445740653 | 710.3742900920952 | 1122.4355191135273 | 1124.0253002713039 | 1225.4196833987344 | 1459.471903543439 | 1029.4094140400837 | 901.8491468896167 |
| SE_14_3_6 | SE_14 | SE_14_3 | GTACCATTGATAAATATGGC | 104.0425768020767 | 139.91494784874547 | 153.06666370835433 | 137.93869989403845 | 129.39253055298764 | 85.76470087697247 | 100.08293698711222 | 169.21827480955713 | 166.94123223113192 | 249.3203751307973 | 107.98821675220947 | 116.06810127279495 |
| SE_14_3_8 | SE_14 | SE_14_3 | ATCCGACTTGTGGTTTACCA | 115.96412206064798 | 133.9185929409421 | 115.54064292824165 | 118.41907255054244 | 151.66501532030517 | 84.89839076710406 | 118.37766740411122 | 103.98888396118035 | 138.5257033407265 | 244.21732651700904 | 122.07363632858463 | 90.53311899278006 |
| SE_14_4_1 | SE_14 | SE_14_4 | GCAGGTCACCTTATAAAGAA | 1368.8101510523215 | 1697.9678313929896 | 1685.7083545171668 | 1626.6356119580005 | 1817.8589948182032 | 1965.6576392914194 | 1390.3995116919245 | 1736.614362151712 | 1815.9298931524722 | 1851.6776398603074 | 2016.5625693510422 | 1725.932665926461 |
| SE_14_4_13 | SE_14 | SE_14_4 | GGACACCCAACTGGAATGCA | 277.4468714722045 | 289.82382054382987 | 297.24558565299776 | 252.45384697588167 | 203.6341464440461 | 213.97859713749696 | 346.5237173102165 | 223.10342377125966 | 323.2266411283618 | 225.26314595150984 | 145.5493356225432 | 574.5371013003349 |
| SE_14_4_15 | SE_14 | SE_14_4 | AATGTGGATGAGCCAAGAGA | 309.9601767228535 | 401.75577882282624 | 458.21246426242845 | 326.62843088116654 | 387.1169971462335 | 263.3582733999963 | 320.69586260386495 | 500.09199650422187 | 317.89872946141077 | 341.1752501789858 | 393.21796317380625 | 107.94333418369929 |
| SE_14_4_31 | SE_14 | SE_14_4 | GGAAAAGGTTTGGACAGCAA | 542.9721976858377 | 645.6075450734969 | 693.2438575694499 | 685.7895740014931 | 550.4485521065621 | 621.1443487756491 | 622.0208341779662 | 589.9005781070595 | 713.9401633714365 | 962.2891671714983 | 750.0485924419767 | 886.7602937241534 |
| SE_14_5_14 | SE_14 | SE_14_5 | GCAATATCTTCCGTATCAGC | 846.4297133585615 | 933.432580648059 | 934.2004120522787 | 723.5275201989186 | 796.5064790597845 | 570.0320522934129 | 818.9582263138967 | 642.8403735782058 | 685.5246344810311 | 970.3082435645941 | 772.3505067712373 | 783.4596835913659 |
| SE_14_5_15 | SE_14 | SE_14_5 | ACACATTGTCCAATGCAGAG | 62.859056817921335 | 85.94775367851507 | 95.79010567555078 | 85.88636031138243 | 77.42339942924669 | 100.4919727447354 | 88.2451702467011 | 109.66100490451745 | 31.07948472388094 | 6.561062503442034 | 78.64359263476125 | 102.13992912005955 |
| SE_14_5_20 | SE_14 | SE_14_5 | GGAGTGAACTGGGAAAGCCA | 610.1663618705123 | 673.590534643246 | 658.6804173772408 | 636.3398513979698 | 638.4778966631029 | 433.1550549342044 | 666.1434193013168 | 682.5452201815656 | 686.4126197588563 | 414.8049516065019 | 1032.9307689341776 | 788.1024076422776 |
| SE_14_5_3 | SE_14 | SE_14_5 | TTGAAGGTCTGGAAGAAGAA | 84.53459365168732 | 122.92527560996923 | 102.70279371399258 | 113.21383859227684 | 133.63490860390525 | 187.12298373157628 | 132.36775537005164 | 58.61191641448347 | 225.54826056759313 | 57.59154864132452 | 90.38144228174053 | 48.74860253457388 |
| SE_14_5_6 | SE_14 | SE_14_5 | GGTCTAACACAAGCAGAGAT | 282.86575568064603 | 270.83536333578587 | 310.0834348672468 | 343.54544124552973 | 290.60289648785744 | 205.31549603881288 | 342.2190748591579 | 203.25100046957976 | 275.2754361258026 | 537.2781183374199 | 173.7201747752935 | 270.4386759656122 |
| SE_14_6_0 | SE_14 | SE_14_6 | CATCCATCATCCATACCTGA | 2117.699948658936 | 1706.9623637546947 | 1962.215876054839 | 1858.2685231008197 | 1850.7374247128148 | 2326.0426449966776 | 1819.7875961850189 | 1986.1876836585448 | 1973.1032873275274 | 1568.822945267473 | 2183.240034338148 | 1608.7038836409379 |
| SE_14_6_1 | SE_14 | SE_14_6 | GAGAATAGCTTGAATCAAGG | 263.3577725302566 | 287.8250355745621 | 324.89633780676496 | 309.7114205168033 | 318.178353818822 | 510.2566547124927 | 314.23889892727703 | 211.75918188458544 | 377.39374307569716 | 368.8775140824077 | 369.74226387984766 | 298.295020271083 |
| SE_14_6_18 | SE_14 | SE_14_6 | GTATGGCCGAGTGCGAAAGA | 1449.0096373372558 | 1577.041340752288 | 1645.2197531491504 | 1383.2909244090836 | 1475.2869672066047 | 1161.721857333536 | 1459.2737909088619 | 1599.538106021065 | 1648.9886609213404 | 1484.9871466123802 | 2362.829133936931 | 1326.6583975480462 |
| SE_14_6_2 | SE_14 | SE_14_6 | GTAGGGTGAATTGAAAACCA | 1709.1160793424474 | 1973.8001571519449 | 1929.627489587899 | 1867.3776825277846 | 1998.1600619822023 | 2153.646933132864 | 1608.860116083148 | 1915.2861718668307 | 2055.685918165268 | 1938.4294662947075 | 2624.5831810645695 | 2013.7815570829923 |
| SE_14_6_9 | SE_14 | SE_14_6 | AATCTGGGAAAACTTCCAGT | 7.586437891818092 | 9.993924846338961 | 3.950107450538176 | 11.711776406097604 | 20.15129574185873 | 11.262031428289314 | 10.761606127646475 | 19.852423301679885 | 10.655823333902038 | 0.7290069448268927 | 48.12518355261509 | 0.0 |
| SE_14_7_2 | SE_14 | SE_14_7 | CCACAGCCCAAGCTATACGT | 196.1636083455821 | 242.85237376603678 | 268.607306636596 | 231.63291114281927 | 219.5430641349872 | 466.0748391092039 | 258.2785470635154 | 246.7372610351643 | 209.56452556674006 | 238.3852709583939 | 356.8306292681704 | 220.52939241831038 |
| SE_14_7_3 | SE_14 | SE_14_7 | TTGGTACCATTAGAGTGGGG | 287.2008630473992 | 318.8062025982129 | 356.4971974110704 | 329.2310478602993 | 286.3605184369398 | 297.1443676848642 | 447.6828149100933 | 289.2781681101926 | 388.0495664095992 | 468.751465523692 | 301.66273592736775 | 342.40089875474507 |
| SE_14_7_4 | SE_14 | SE_14_7 | ATACAAGTCCGAAATCCAGA | 76.94815575986922 | 126.92284554850481 | 173.80472782367974 | 96.29682822791364 | 117.72599091296415 | 155.06950966644516 | 71.02660044246673 | 221.21271679014728 | 134.9737622294258 | 217.244069558414 | 106.81443178751154 | 221.69007343103834 |
| SE_14_7_6 | SE_14 | SE_14_7 | ACAGGCTGAATAAGGCTTGG | 5942.348422976943 | 5824.459400446347 | 5802.707844840581 | 6017.250455755036 | 6087.812503066794 | 5678.6627701874195 | 6417.145733915593 | 5508.57478947089 | 4913.222542206664 | 6617.9250451385315 | 5265.59935163491 | 6442.940301652848 |
| SE_14_7_8 | SE_14 | SE_14_7 | CCACAGGCTGAATAAGGCTT | 320.7979451397365 | 302.81592284407054 | 360.4473048616086 | 374.7768449951233 | 325.60251540792785 | 190.58822417104992 | 308.8580958634538 | 539.7968431075816 | 221.99631945629244 | 299.62185432385286 | 721.8777532892263 | 222.85075444376628 |
| SE_14_8_12 | SE_14 | SE_14_8 | ACACACATGTGCCAGACACG | 2895.8517209911347 | 2540.455695939364 | 2569.5448965750834 | 2978.695132617491 | 2518.911967732341 | 2576.4062667486473 | 2969.1271306176623 | 2325.5695867682152 | 2315.8656045680427 | 1926.0363482326504 | 1880.4035134460823 | 2857.5966533362116 |
| SE_14_8_15 | SE_14 | SE_14_8 | AAAGGAATGTTGGCCCAGCG | 1012.2475701368712 | 1024.3772967497437 | 1068.5040653705767 | 1002.0075369661283 | 1141.1996956968417 | 1038.7058217322221 | 1067.5513278625303 | 1107.0089374412926 | 1065.5823333902038 | 833.2549379371383 | 1044.6686185811568 | 983.0968177805731 |
| SE_14_8_25 | SE_14 | SE_14_8 | ACTTCACTTCCAGAGGACAG | 276.36309463051623 | 318.8062025982129 | 330.8214989825723 | 281.0826337463425 | 342.5720276115984 | 186.25667362170788 | 190.4804284593426 | 436.75331263695745 | 246.8599072353972 | 177.8776945377618 | 457.7761362321923 | 152.0492126673614 |
| SE_14_8_8 | SE_14 | SE_14_8 | AGGGTGGCTGATCAGCTCTG | 4613.638015067088 | 4298.387076410388 | 4458.683784794966 | 5011.338993320208 | 4682.524773700331 | 3641.96770188679 | 4948.186497491849 | 4410.074033444603 | 5151.20259666381 | 4724.694009423091 | 4571.8924374984335 | 3388.0278761528843 |
| SE_14_8_9 | SE_14 | SE_14_8 | ATAAACATGGTAACCCACTG | 123.55055995246607 | 167.89793741849456 | 125.4159115545871 | 137.93869989403845 | 158.0285823966816 | 183.65774329210265 | 130.21543414452233 | 231.61160518626534 | 124.31793889552377 | 29.160277793075707 | 309.8792306802533 | 233.29688355831783 |
| SE_14_9_0 | SE_14 | SE_14_9 | GTTTTACTTCACCCTCTCAT | 4.335107366753196 | 18.98845720804403 | 21.725590977959968 | 10.410467916531204 | 6.363567076376441 | 24.256683076315444 | 19.370891029763655 | 1.89070698111237 | 2.6639558334755096 | 5.832055558615141 | 2.347569929395858 | 0.0 |
| SE_14_9_13 | SE_14 | SE_14_9 | GGAAGAGTTTAATAGACACA | 172.32051782843953 | 285.8262506052943 | 307.1208542793432 | 230.33160265325287 | 298.02705807696327 | 238.2352802138124 | 172.1856980423436 | 319.5294798079905 | 356.08209640789306 | 266.8165418066427 | 315.7481555037429 | 277.40276204197994 |
| SE_14_9_19 | SE_14 | SE_14_9 | TGCGCCAAATTGCTTCTGGG | 9526.398438440148 | 10950.343454133601 | 10941.797637990749 | 10187.94416481535 | 11063.061362280441 | 10910.30952368274 | 9605.809629537243 | 11123.029169884072 | 11099.815972814622 | 11524.870790768346 | 11916.264961613375 | 11072.896861424637 |
| SE_14_9_3 | SE_14 | SE_14_9 | AAGGTATTGTAACCACCTGA | 1044.7608753875202 | 1148.3019648443467 | 1137.6309457549946 | 1251.8587669628773 | 1107.2606712895006 | 1130.5346933782735 | 1226.8230985516982 | 1236.52236564749 | 1046.04665727805 | 1450.7238202055164 | 1314.6391604616806 | 993.5429468951247 |
| SE_14_9_5 | SE_14 | SE_14_9 | GTTTTACTGCTGCCATGTGA | 62.859056817921335 | 80.95079125534559 | 73.07698783495626 | 81.98243484268323 | 73.18102137832906 | 54.57753692170975 | 54.88419125099702 | 66.17474433893295 | 82.5826308377408 | 11.664111117230282 | 52.820323411406804 | 102.13992912005955 |
| SE_140_1_0 | SE_140 | SE_140_1 | ATTGTGATGCCAGTATGCCC | 799.8273091659646 | 730.5559062673781 | 708.0567605089681 | 910.9159426964803 | 823.0213418780196 | 763.2192067940681 | 775.9118018033108 | 1031.3806581967979 | 744.1316628174923 | 886.4724449095015 | 1005.9337147461251 | 517.6637316766654 |
| SE_140_1_12 | SE_140 | SE_140_1 | CTGTTTGTCTCACATATGTA | 58.52394945116814 | 72.95565137827442 | 84.92731018657079 | 81.98243484268323 | 84.84756101835254 | 273.7539947184172 | 92.54981269775968 | 68.06545132004533 | 99.45435111641902 | 61.23658336545898 | 63.384388093688166 | 123.03218734916264 |
| SE_140_1_3 | SE_140 | SE_140_1 | GGAGTCTGACTTGGATCAGT | 396.6623240579174 | 532.6761943098667 | 563.8778385643246 | 504.90769395176335 | 529.236661851974 | 456.5454279006514 | 441.22585123350547 | 522.7804802775703 | 443.1046536347597 | 339.717236289332 | 488.2945453143385 | 633.7318329494603 |
| SE_140_1_5 | SE_140 | SE_140_1 | ATACCAACGCCATGGAAGAA | 98.6236925936352 | 121.92588312533533 | 122.45333096668347 | 100.20075369661284 | 115.60480188750533 | 71.03742900920952 | 67.79811860417279 | 146.52979103620868 | 84.35860139339113 | 99.1449444964574 | 172.54638981059557 | 88.21175696732416 |
| SE_140_1_6 | SE_140 | SE_140_1 | CGTCTCTTAGCATCTTTAAG | 268.77665673869814 | 416.7466660923347 | 391.0606376032794 | 343.54544124552973 | 409.389481913551 | 300.60960812433785 | 234.60301358269314 | 528.4526012209075 | 388.9375516874244 | 480.41557664092227 | 528.2032341140681 | 359.8111139456643 |
| SE_140_2_0 | SE_140 | SE_140_2 | ATTTGCCTAAAGTCACACAG | 71.52927155142773 | 86.94714616314897 | 87.88989077447442 | 35.13532921829281 | 30.757240869152795 | 48.51336615263089 | 57.03651247652632 | 67.12009782948914 | 63.93494000341222 | 20.412194455152996 | 1.173784964697929 | 90.53311899278006 |
| SE_140_2_11 | SE_140 | SE_140_2 | ATGCAGCTGCCAGGGATCCG | 2109.02973392543 | 2067.743050707531 | 1979.0038327196262 | 2114.626295545401 | 2081.9470284878253 | 2080.0105737940494 | 2271.775053546171 | 2314.225344881541 | 1842.5694514872273 | 1546.9527369226662 | 1814.6715554229984 | 1728.2540279519167 |
| SE_140_2_12 | SE_140 | SE_140_2 | GCATCAGCTATTATAAGAGT | 1002.4935785616765 | 1167.2904220523908 | 1246.2589006447945 | 1136.0423113914676 | 1305.5918451698997 | 1524.7057933683993 | 1053.5612398965898 | 1333.8937751747771 | 1267.1549914565173 | 960.8311532818445 | 1297.0323859912116 | 1145.5921595624861 |
| SE_140_2_7 | SE_140 | SE_140_2 | GTTCCTTCATCTACAAAGTG | 8.670214733506391 | 10.993317330972857 | 17.77548352742179 | 10.410467916531204 | 12.727134152752882 | 21.65775274671022 | 7.533124289352532 | 0.0 | 11.543808611727208 | 20.412194455152996 | 0.0 | 5.803405063639747 |
| SE_140_2_9 | SE_140 | SE_140_2 | ACAAAACTAAAATTCAGCAT | 668.6903113216804 | 706.5704866361646 | 740.645146975908 | 641.5450853562354 | 716.9618906050789 | 786.6095797605151 | 702.7328801353148 | 570.0481548053796 | 711.276207537961 | 693.2856045303749 | 610.3681816429231 | 949.4370684114626 |
| SE_141_1_17 | SE_141 | SE_141_1 | GTTTATCTACTTGCAAGTGG | 15.172875783636185 | 15.990279754142339 | 10.862795488979884 | 16.917010364363207 | 15.9089176909411 | 2.598930329605226 | 4.30464245105859 | 11.34424188667422 | 7.991867500426528 | 37.908361130998415 | 10.564064682281362 | 9.285448101823595 |
| SE_141_1_2 | SE_141 | SE_141_1 | GAATGCGTAGAAACCTTGTT | 472.5267029760983 | 628.6178728347207 | 655.7178367893372 | 486.68937509783376 | 661.8109759431499 | 573.4972927328865 | 377.73237508039125 | 657.0206759365486 | 594.0621508650386 | 583.934562806341 | 640.8865907250693 | 687.123159534946 |
| SE_141_1_6 | SE_141 | SE_141_1 | TTCAGCAGAACATTTAGGAA | 32.51330525064897 | 50.9690167163287 | 43.451181955919935 | 40.34056317655841 | 44.54496953463509 | 74.50266944868315 | 36.58946083399801 | 53.88514896170255 | 89.68651306034215 | 11.664111117230282 | 73.94845277596953 | 125.35354937461854 |
| SE_141_1_7 | SE_141 | SE_141_1 | GAAGTATCTTTCCTTCCACA | 53.105065242726646 | 96.94107100948793 | 111.59053547770347 | 102.80337067574564 | 119.84717993842297 | 65.83956834999907 | 121.60614924240517 | 84.13646065950047 | 142.07764445202716 | 66.33963197924723 | 88.03387235234467 | 81.24767089095646 |
| SE_141_1_8 | SE_141 | SE_141_1 | GCTATTGCCAGCCTCACAGA | 141.97476626116716 | 119.92709815606754 | 150.1040831204507 | 149.65047630013606 | 136.81669214209347 | 173.26202197368175 | 88.2451702467011 | 105.87959094229272 | 218.44437834499178 | 164.75556953087775 | 100.9455069640219 | 225.1721164692222 |
| SE_141_2_0 | SE_141 | SE_141_2 | AGTTGTGCTCTTGCAGCGGG | 216.75536833765977 | 272.83414830505365 | 259.7195648728851 | 256.35777244458086 | 270.45160074599875 | 192.32084439078673 | 243.21229848481033 | 212.70453537514163 | 322.3386558505366 | 647.3581670062807 | 193.67451917515828 | 152.0492126673614 |
| SE_141_2_17 | SE_141 | SE_141_2 | TGAAAGGACTTGCACTGCAG | 411.83519984155356 | 61.96233404730156 | 122.45333096668347 | 274.5760912985105 | 59.39329271284678 | 399.36896064933643 | 469.2060271653863 | 138.021609621203 | 371.17784613092095 | 24.057229179287457 | 77.46980767006332 | 121.87150633643469 |
| SE_141_2_19 | SE_141 | SE_141_2 | TCACTCATCCCACTGGAAGG | 590.6583787201229 | 686.5826369434867 | 735.7075126627353 | 508.8116194204626 | 661.8109759431499 | 505.9251041631507 | 514.4047729015015 | 386.6495776374797 | 509.7035494716475 | 275.5646251445654 | 392.0441782091083 | 247.22505571105322 |
| SE_141_2_5 | SE_141 | SE_141_2 | CCTCCACTAAATGTCTACCA | 545.1397513692143 | 620.6227329576495 | 646.8300950256263 | 609.0123731170754 | 543.0243905174563 | 656.6630632802538 | 530.5471820929712 | 860.2716764061283 | 606.493944754591 | 402.41183354444473 | 899.1192829586137 | 1007.4711190478602 |
| SE_141_2_8 | SE_141 | SE_141_2 | TTTTGACATTTAAATACACA | 2.167553683376598 | 4.996962423169481 | 5.925161175807264 | 6.506542447832002 | 2.1211890254588135 | 25.989303296052263 | 0.0 | 0.0 | 0.0 | 10.206097227576498 | 0.0 | 0.0 |
| SE_142_1_11 | SE_142 | SE_142_1 | GCTGTGGCCCTCCTGACATG | 119.21545258571288 | 91.94410858631845 | 104.67784743926167 | 111.91253010271043 | 90.15053358199957 | 154.20319955657675 | 218.46060439122343 | 126.67736773452879 | 81.69464555991561 | 121.01515284126418 | 56.341678305500594 | 87.05107595459621 |
| SE_142_1_14 | SE_142 | SE_142_1 | CATCAGTGATTACAACCCAA | 1571.4764204480334 | 1505.0850818586478 | 1574.1178190394633 | 1543.351868625751 | 1558.0133391994984 | 1788.930376878264 | 1343.0484447302802 | 1747.958604038386 | 1503.3590753580124 | 1706.6052578397557 | 1712.5522634942783 | 1716.6472178246372 |
| SE_142_1_17 | SE_142 | SE_142_1 | ACTGATGCACCATTGTTAGG | 2471.0111990493215 | 2086.731507915575 | 2184.4094201476114 | 2415.2285566352393 | 1996.0388729567435 | 1877.2940080848416 | 2518.2158338692752 | 2018.329702337455 | 1972.2153020497021 | 2611.3028763699294 | 2115.160506385668 | 2442.0728507796057 |
| SE_142_1_27 | SE_142 | SE_142_1 | AGAAACAGCGAGGTTCCCAA | 96.4561389102586 | 136.91677039484378 | 128.37849214249073 | 130.13084895664005 | 108.18064029839948 | 335.2620125190742 | 89.32133085946575 | 175.8357492434504 | 105.67024806119521 | 161.1105348067433 | 299.3151659979719 | 76.60494684004466 |
| SE_142_1_9 | SE_142 | SE_142_1 | ACAGGAATCGGGGAATGCAG | 1713.4511867092006 | 2103.7211801543513 | 1890.1264150825173 | 1950.6614258600343 | 1702.2541929306979 | 1351.4437713947177 | 1703.562250006437 | 1342.4019565897827 | 1637.444852309613 | 1360.3269590469818 | 1405.020602743421 | 1341.7472507135096 |
| SE_142_2_1 | SE_142 | SE_142_2 | CTAGGGGGTTATTGAAAAGG | 5104.588924351888 | 6932.785665905338 | 7101.305669205006 | 5734.866513519127 | 7093.256101134272 | 6271.218885337411 | 4985.852118938612 | 6771.567052853953 | 6579.970908684508 | 7450.450976130843 | 6363.088293627473 | 7191.579554862375 |
| SE_142_2_12 | SE_142 | SE_142_2 | CCCAAAAGGCTAAATCCCAC | 688.1982944720698 | 579.6476410876597 | 525.3642909215774 | 572.5757354092162 | 531.3578508774327 | 736.3635933881474 | 581.1267308929097 | 539.7968431075816 | 473.2961530808155 | 555.5032919580922 | 706.6185487481533 | 572.2157392748791 |
| SE_142_2_22 | SE_142 | SE_142_2 | GCAATTTCAAGAGCTTCAGG | 35.76463577571386 | 101.9380334326574 | 112.57806234033802 | 49.44972260352321 | 88.02934455654076 | 20.791442636841808 | 27.980175931880833 | 117.22383282896693 | 99.45435111641902 | 88.9388472688809 | 130.29013108147012 | 33.659749369110536 |
| SE_142_2_31 | SE_142 | SE_142_2 | GTTTGGTTCTTAACACATAG | 9.75399157519469 | 15.990279754142339 | 8.887741763710896 | 10.410467916531204 | 42.42378050917627 | 0.0 | 8.60928490211718 | 18.9070698111237 | 46.17523444690883 | 15.309145841364746 | 36.3873339056358 | 1.1606810127279494 |
| SE_142_2_6 | SE_142 | SE_142_2 | GCATGTCCAAGGCATGTCCA | 4706.842823452283 | 4034.547460467039 | 3977.7582026919436 | 4410.134471140531 | 3795.867761058547 | 4067.325965832179 | 4997.689885679023 | 3737.9277016591554 | 3934.6627660433273 | 4713.758905250688 | 4008.4756544434276 | 3594.6290964184595 |
| SE_143_1_13 | SE_143 | SE_143_1 | GCTATGCTGCAAACAGTCCT | 2662.83970002815 | 2529.462378608391 | 2374.0145777734438 | 2895.411389285241 | 2518.911967732341 | 2245.4758047789155 | 3057.3723008643633 | 2746.2518900657174 | 2577.821261526468 | 2981.6384043419907 | 3217.3445882370233 | 3229.014577409155 |
| SE_143_1_16 | SE_143 | SE_143_1 | ATATCCCTATTTTCTAGACG | 29.26197472558407 | 11.992709815606755 | 14.81290293951816 | 44.24448864525762 | 29.69664635642339 | 1.7326202197368175 | 29.056336544645482 | 24.57919075446081 | 99.45435111641902 | 5.832055558615141 | 51.646538446708874 | 2.321362025455899 |
| SE_143_1_17 | SE_143 | SE_143_1 | TTTTAAGGTGGCAAATTTGG | 11.921545258571289 | 17.98906472341013 | 7.900214901076352 | 1.3013084895664004 | 8.484756101835254 | 0.0 | 2.152321225529295 | 0.945353490556185 | 3.551941111300679 | 0.0 | 28.170839152750297 | 1.1606810127279494 |
| SE_143_1_29 | SE_143 | SE_143_1 | AATTATGTGTGGTAACCAAA | 5.4188842084414945 | 20.98724217731182 | 13.825376076883616 | 14.314393385230405 | 15.9089176909411 | 8.663101098684088 | 8.60928490211718 | 43.48626056558451 | 0.8879852778251698 | 26.244250013768134 | 28.170839152750297 | 1.1606810127279494 |
| SE_143_1_6 | SE_143 | SE_143_1 | TTAATCCATCCTGCACCTGA | 115.96412206064798 | 119.92709815606754 | 124.42838469195254 | 104.10467916531204 | 153.786204345764 | 250.36362175197013 | 116.22534617858193 | 122.89595377230405 | 100.34233639424419 | 74.35870837234305 | 124.42120625798047 | 109.10401519642724 |

| | | | | | | | | | | | | | | | |
|---|---|---|---|---|---|---|---|---|---|---|---|---|---|---|---|
| SE_143_2_14 | SE_143 | SE_143_2 | TGACCTGCACTCAACCTAGG | 2718.1123189542536 | 2371.558366036236 | 2509.3057579543765 | 2520.6345442901174 | 2566.638720805164 | 2463.7859524657542 | 2875.501157307138 | 2361.49301940935 | 2561.837526525615 | 2528.196084659664 | 1986.044160268896 | 3317.2263343764794 |
| SE_143_2_15 | SE_143 | SE_143_2 | CATCTACCAAGGGAAAGGGT | 702.2873934140177 | 738.5510461444493 | 727.8072977616589 | 654.5581792815895 | 724.3860521941848 | 593.42242525986 | 598.345300697144 | 472.6767452780925 | 692.6285167036324 | 419.17899327546326 | 347.440349550587 | 557.1268861094158 |
| SE_143_2_19 | SE_143 | SE_143_2 | AGAGGCACAAATGAGCACAA | 5555.44009049422 | 5681.5462751437 | 5293.1439837211565 | 5931.364095443653 | 5592.514865622162 | 6057.240288199914 | 5577.740455959168 | 5595.547310602059 | 5902.438141703904 | 6174.688822683781 | 5708.116283326029 | 6227.053633285449 |
| SE_143_2_35 | SE_143 | SE_143_2 | GGAGTGGGCAGCACAATGGG | 2693.185451595423 | 3041.151330740946 | 2789.7633869425867 | 2774.3896997555657 | 3215.722562595561 | 2699.4223023499617 | 2652.735910464856 | 3602.742152509621 | 2798.04161042711 | 2598.9097583078724 | 3516.6597542349955 | 2617.335683701526 |
| SE_143_2_6 | SE_143 | SE_143_2 | ACAATTATAATATAGTTGCA | 26.010644200519174 | 55.96597913949819 | 38.51354764274722 | 42.943180155691216 | 36.06021343279983 | 6.064170769078861 | 7.533124289352532 | 17.01636283001133 | 11.543808611727208 | 12.393118062057175 | 32.86597901154201 | 2.321362025455899 |
| SE_143_3_2 | SE_143 | SE_143_3 | GGTTACTTCTAGGTTACGGA | 34.680858934025565 | 12.992102300240651 | 14.81290293951816 | 22.12224432262881 | 18.030106716399914 | 32.919784174999535 | 17.21856980423436 | 3.78141396222474 | 15.983735000853056 | 5.103048613788249 | 1.173784964697929 | 31.338387343654635 |
| SE_143_3_3 | SE_143 | SE_143_3 | TATTGCTGAGTAACGTTCCG | 1401.3234563029705 | 1166.2910295677568 | 1243.2963200568909 | 1400.2079347734468 | 1104.0788877513123 | 1535.9678247966888 | 1463.5784333599206 | 1144.82307706354 | 1397.6888272968172 | 1050.4990074955524 | 1326.3770101086598 | 1261.660260835281 |
| SE_143_3_5 | SE_143 | SE_143_3 | AATTCTTTAGAGATTCATTC | 0.0 | 0.0 | 0.987526862634544 | 0.0 | 0.0 | 0.0 | 0.0 | 0.0 | 0.0 | 0.0 | 0.0 | 0.0 |
| SE_143_3_6 | SE_143 | SE_143_3 | ACTGCTAGCAATCTACACTA | 614.5014692372655 | 776.5279605605373 | 730.7698783495625 | 730.0340626467506 | 573.7816313866091 | 585.6256342710443 | 622.0208341779662 | 728.8675412188186 | 961.6880558846589 | 758.8962295647952 | 515.2915995023908 | 587.3045924403424 |
| SE_143_3_8 | SE_143 | SE_143_3 | TCAAATACTCATTTTCACAG | 158.23141888649164 | 175.89307729556572 | 161.95440547206522 | 139.24000838360485 | 155.90739337122278 | 269.4224441690751 | 153.8909676253446 | 148.42049801732105 | 220.2203489006421 | 267.5455487154696 | 147.89690555193906 | 204.2798582401191 |
| SE_144_1_18 | SE_144 | SE_144_1 | CGCTGTAGACAGGGCTGGAA | 4886.74977917254 | 4937.998266576081 | 5065.0252784525765 | 5315.8451798787455 | 5538.424545472962 | 4677.208283179539 | 5449.677430401745 | 5524.645798810345 | 5349.223313618822 | 6667.497517386761 | 5649.427035091133 | 5854.475028199777 |
| SE_144_1_19 | SE_144 | SE_144_1 | AGACAGGGCTGGAAGGGCTG | 2449.3356622155557 | 2546.4520508471674 | 2578.4326383387943 | 2779.5949337138313 | 2539.0632634741996 | 2501.037287190096 | 3001.411949000602 | 2809.590573932982 | 2978.3026218256196 | 3446.0158281967215 | 3352.3298591772855 | 2661.441562185188 |
| SE_144_1_34 | SE_144 | SE_144_1 | GGAAACGAGCACGAAATCTT | 1321.1239700180363 | 1807.9010047027182 | 1711.3840529456647 | 1486.0942950848294 | 1620.5884154505336 | 1717.8929478690545 | 1292.4688959303417 | 1314.0413518730973 | 1681.8441162008717 | 1447.8077924262088 | 1316.9867303910764 | 1481.0289722408634 |
| SE_144_1_42 | SE_144 | SE_144_1 | CCGGCCAACTCCTTCCTCCG | 479.0293640262281 | 506.6919897093854 | 520.4266566084048 | 540.0430231700561 | 656.5080033795027 | 518.9197558111769 | 556.3750367993227 | 486.85704763643525 | 465.304285580389 | 550.4002433443039 | 543.4624386551411 | 509.5389645875698 |
| SE_144_1_7 | SE_144 | SE_144_1 | ACTCACTGCCCCCAGGCCCG | 1152.0547827146618 | 1206.2667289531128 | 1127.7556771286493 | 1335.1425102951268 | 1626.95198252691 | 1770.7378645710273 | 1446.3598635556862 | 1646.8057805488743 | 1193.4522133970281 | 1727.7464592397355 | 1453.1457862960362 | 1470.582843126312 |
| SE_144_2_22 | SE_144 | SE_144_2 | GGAGGCAGGAACTCGCCCAG | 4189.881269966963 | 4527.24795539155 | 4626.563351442839 | 4572.7980323336331 | 4721.766770671319 | 5275.8285690986095 | 4334.774948216 | 4365.642419388462 | 4451.470197737576 | 4156.797599402942 | 3845.3195443504155 | 5431.987139566803 |
| SE_144_2_27 | SE_144 | SE_144_2 | GCAGCGCCGGGCGCGGGCCT | 376.07056406583973 | 305.81410029797223 | 307.1208542793432 | 330.5323563498657 | 337.2690550479513 | 422.7593336157835 | 372.351572016568 | 333.7097821663333 | 380.05769890917264 | 308.36993766177557 | 341.57142472709734 | 499.0928354730183 |
| SE_144_2_30 | SE_144 | SE_144_2 | CGGGCCTGGGTGCAGTGTGG | 4187.713716283587 | 3719.7388278073618 | 3809.878636044071 | 4463.488119212753 | 4047.228660575416 | 4272.641461870991 | 4492.970558292403 | 4509.336149953003 | 3910.6871635420475 | 4783.014565009243 | 4546.069168275079 | 4657.812904077261 |
| SE_144_2_59 | SE_144 | SE_144_2 | GTGCTGGCGTGACTCACCGG | 2783.1389294555175 | 2677.3724663342077 | 2771.0003765525307 | 2965.6820477218266 | 2731.030870278222 | 3163.7645212394286 | 3027.2398037069534 | 3145.1910630804277 | 2758.082272924977 | 3855.717731189435 | 3688.032359080893 | 3455.3473748911056 |
| SE_144_2_8 | SE_144 | SE_144_2 | GATCTTGTGGATGAAAGCGA | 257.9388883218151 | 305.81410029797223 | 304.15827368143955 | 283.68525072547527 | 330.9054879715749 | 245.6577338374951 | 261.50702890180935 | 335.60048914744567 | 364.9619491861448 | 229.6371876204712 | 483.5994054555468 | 275.081400016524 |
| SE_145_1_10 | SE_145 | SE_145_1 | AATAGGGTGTGGGTTCTGAG | 110.54523785220648 | 127.92223803313871 | 143.19139508200888 | 126.22692348794084 | 154.84679885849337 | 149.0053388973663 | 132.36775537005164 | 132.3494886778659 | 197.13273167718768 | 168.40060425550122 | 111.50957164630326 | 200.79781520193526 |
| SE_145_1_12 | SE_145 | SE_145_1 | GTACCTAGAGACAACTCAAT | 422.6729682584366 | 464.7175053547617 | 404.88601368016305 | 428.13049306734575 | 547.2667685683739 | 285.01602614670645 | 343.2952354719225 | 518.0537128247894 | 397.8174044656761 | 371.79354186171525 | 438.9955767970255 | 450.3442329384444 |
| SE_145_1_13 | SE_145 | SE_145_1 | GATCAAAGCATGCAACCTTG | 1663.5974519915387 | 1657.9921320076337 | 1606.7062055064032 | 1754.1638439355079 | 1796.6471045636151 | 2719.347434876935 | 1574.4229764746792 | 1904.8872834707129 | 1913.6082737132408 | 1802.8341745569055 | 1829.9307599640713 | 2447.8762558432454 |
| SE_145_1_2 | SE_145 | SE_145_1 | AAGGAGCTCTAACTTTGCCA | 1252.8460289916736 | 1348.180461771126 | 1350.9367480840563 | 1319.52680842033 | 1401.0453513155462 | 1191.176401069062 | 1396.8564753685125 | 1410.4674079098281 | 1241.4034183995873 | 1316.5865423573682 | 1338.1148597556391 | 1013.2745241114999 |
| SE_145_1_3 | SE_145 | SE_145_1 | TTTGTGCCCTCTGTAAGCAG | 247.10111990493215 | 264.8390084279825 | 242.93160820809783 | 247.2486130176161 | 252.42149402959882 | 285.8823362565749 | 235.6791741954578 | 423.5183637691709 | 219.33236362281693 | 161.1105348067433 | 720.7039683245284 | 277.40276204197994 |
| SE_145_2_0 | SE_145 | SE_145_2 | ACTCTAAGCAACTTGAGTGG | 119.21545258571288 | 179.89064723410132 | 204.41806056535063 | 109.30991312357764 | 167.57393301124625 | 197.51870504998718 | 105.46374005093506 | 202.30564697902358 | 310.79484723880944 | 83.83579865509266 | 149.070690516637 | 134.63899747644214 |
| SE_145_2_15 | SE_145 | SE_145_2 | GGAAATATCTCAGTAAGTAC | 82.36703996831072 | 81.9501837399795 | 87.88989077447442 | 100.20075369661284 | 99.69588419656424 | 80.566840217762 | 115.14918556581728 | 67.12009782948914 | 136.74973278507613 | 185.89677093085763 | 176.06774470468935 | 59.19473164912542 |
| SE_145_2_20 | SE_145 | SE_145_2 | TCATCTGGTGAGCTGTGTGT | 2023.411363432054 | 1673.9824117617761 | 1806.1863017585811 | 1882.9933844025813 | 2045.8868150550256 | 1923.2084439078674 | 2040.4005218017717 | 1841.5485996034483 | 2005.9587426070586 | 1430.3116257503634 | 2525.9852440299433 | 1712.0044937737255 |
| SE_145_2_4 | SE_145 | SE_145_2 | TTACACGGCAGAAATCATAC | 703.371170255706 | 732.5546912366459 | 782.1212752065588 | 599.9032136901106 | 638.4778966631029 | 627.2085195447279 | 505.7954879993843 | 817.7307693311001 | 810.73055865438 | 401.68282659961784 | 833.3873249355296 | 703.3726937131373 |
| SE_145_2_8 | SE_145 | SE_145_2 | GTTAGTTGCAAGATAAGAAG | 82.36703996831072 | 106.9349958558269 | 105.66537430189621 | 115.81645557140963 | 111.36242383658771 | 119.5507951618404 | 92.54981269775968 | 63.33868386726439 | 104.78226278337003 | 48.84346530340181 | 91.55522724643846 | 358.6504329329364 |
| SE_146_1_17 | SE_146 | SE_146_1 | GGAGGTGGCTAACTAGGAGC | 395.5785472162291 | 592.6397433879005 | 672.5057934541245 | 425.52787608821296 | 564.2362807720444 | 562.2352613045973 | 322.84818382939426 | 493.4745220703286 | 787.6429414309256 | 458.5453682961155 | 389.69660827971525 | 443.3801468620767 |
| SE_146_1_21 | SE_146 | SE_146_1 | GCAGCCATGTGATTGGTTAG | 2070.013767624651 | 1935.8232427358569 | 1970.1160909559153 | 2130.2419974201975 | 1833.7679125091443 | 2340.7699168644403 | 2150.1689043037654 | 1769.7017343211783 | 2044.1421095535409 | 2424.677098494245 | 2196.1516689498253 | 2215.7400532976553 |
| SE_146_1_28 | SE_146 | SE_146_1 | GATAATCTCCTGGACTTGGT | 92.12103154350541 | 165.89915244922676 | 122.45333096668347 | 134.03477442533924 | 154.84679885849337 | 102.22459296447224 | 150.66248578705066 | 195.6881725451303 | 80.80666028209045 | 340.4462432341589 | 125.5949912226784 | 211.2439443164868 |
| SE_146_1_5 | SE_146 | SE_146_1 | TGATGGGAGAACTTGTTAAA | 753.2249049733678 | 1047.3633238963232 | 1058.6287967442313 | 878.3832304573203 | 960.8986285328425 | 1419.882270074322 | 850.1668840840715 | 1309.3145844203161 | 843.5860139339113 | 862.415215730214 | 1293.511031097118 | 1417.1915165408263 |
| SE_146_1_6 | SE_146 | SE_146_1 | GTTACAGTGGACAAAAGGGA | 360.89768828220355 | 507.69138219401924 | 454.26235681189024 | 460.66320530650574 | 407.2682928880922 | 411.49730218749414 | 331.4574687315114 | 575.7202757487166 | 380.9456841869978 | 376.1675835306766 | 367.39469395045177 | 348.20430381838486 |
| SE_147_1_11 | SE_147 | SE_147_1 | AAGGCCATGCTGCCAGACAG | 3662.081948064762 | 3462.8949592564504 | 3472.144449023057 | 3444.563571882262 | 3555.112806668971 | 3385.539909365741 | 3564.2439494765126 | 3459.993775435637 | 3819.224679926055 | 2690.7646333560606 | 3061.2311879321987 | 4466.300536977149 |
| SE_147_1_12 | SE_147 | SE_147_1 | TTATCTAGGGATTGAGGGCG | 63.94283365960963 | 68.95808143973883 | 77.02709525849444 | 46.847105624390416 | 29.69664635642339 | 25.12299318618385 | 45.19874573611519 | 55.775855942814914 | 61.27098416993672 | 2.187020834480678 | 88.03387235234467 | 17.410215190919242 |
| SE_147_1_17 | SE_147 | SE_147_1 | AGGAAGGGTGGCAGGAACAA | 5435.140861066819 | 5689.541415020771 | 5354.370649204498 | 5548.7793995111315 | 5481.152441785574 | 5376.3205418433445 | 5463.667431006115 | 5424.4383288113895 | 5228.457315834599 | 4407.575988423393 | 4561.328372816152 | 4859.771400291925 |
| SE_147_1_18 | SE_147 | SE_147_1 | ACTCTTGGGGACAGCGAGAA | 79.11570944324582 | 144.91191027191493 | 130.3535458677598 | 100.20075369661284 | 75.30221040378788 | 149.0053388973663 | 93.62597331052433 | 103.04353047062416 | 91.46248361599248 | 37.17935418617152 | 143.20176569314734 | 62.67677468730927 |
| SE_147_1_19 | SE_147 | SE_147_1 | GTGGCAGGAACAAAGGAGAA | 204.8338230790885 | 234.8572338889656 | 237.00644703229057 | 209.51066682019047 | 241.81554890230473 | 193.18715450065514 | 187.25194662104866 | 153.14726547010198 | 229.1002016788938 | 179.3357084274156 | 97.42415206992811 | 126.51423038734649 |
| SE_147_2_10 | SE_147 | SE_147_2 | GCGTAGGGCAAGAAGAGGCA | 15720.183088688776 | 15528.560426241478 | 15422.207013763675 | 14323.502544657369 | 14535.44779695652 | 12658.52332539719 | 14001.925732680827 | 11964.393776479077 | 12736.37283984641 | 9555.82303279091 | 10797.647890256249 | 9770.612765143878 |
| SE_147_2_25 | SE_147 | SE_147_2 | GGAAAGCCTGTTCTAGAGAG | 145.22609678623206 | 307.81288526724 | 280.4576289882105 | 175.67664609146405 | 259.84565561870465 | 286.74864636644327 | 153.8909676253446 | 176.7811027340066 | 229.98818695671898 | 129.7632361791869 | 118.55228143449084 | 92.85448101823596 |
| SE_147_2_27 | SE_147 | SE_147_2 | CATGGCTCTGAGCAAACTGA | 5558.691421019285 | 5133.879195364325 | 5294.13151058179 | 5448.5786458145185 | 5092.45896568755 | 5336.470276789398 | 5346.365924214769 | 5518.028324376452 | 4675.242487749736 | 5071.7013151606925 | 5567.262087562277 | 5586.3577142596205 |
| SE_147_2_4 | SE_147 | SE_147_2 | AGCCTCGGCGACCACCTCTG | 418.33786089168336 | 476.7102151703685 | 527.3393446468465 | 408.6108657238497 | 388.17759165896285 | 877.5721412966981 | 312.0865777017478 | 638.1136061254249 | 562.0946808633324 | 430.11409744786664 | 461.2974911262861 | 1245.4107266570898 |
| SE_147_2_8 | SE_147 | SE_147_2 | TTCCAGGATGACCACTTCAG | 37.93218945909046 | 47.97083926242702 | 57.526020780112674 | 58.55888203048802 | 30.757240869152795 | 37.251334724341575 | 21.52321225529295 | 38.75949311280358 | 32.855455279531284 | 42.282402799959776 | 16.432989505771005 | 90.53311899278006 |
| SE_147_3_27 | SE_147 | SE_147_3 | CTCATGCCCACAGCCAGGTA | 134.38832836934907 | 118.92770567143364 | 147.14150253254707 | 128.82954046707366 | 164.39214947305805 | 61.50801780065702 | 112.99686434028798 | 165.43686084733238 | 165.1652616754816 | 111.53806255851458 | 138.50662583435562 | 5.803405063639747 |
| SE_147_3_28 | SE_147 | SE_147_3 | TCTCATGCCCACAGCCAGGT | 1107.6199322054415 | 1049.362108865591 | 1013.2025610630421 | 1258.3653094107092 | 1104.0788877513123 | 952.9411208552496 | 1063.2466854114716 | 1509.7295244182274 | 933.2725269942534 | 1052.686028300033 | 1240.690707685711 | 749.7999342222554 |
| SE_147_3_29 | SE_147 | SE_147_3 | TACCTGGCTGTGGGCATGAG | 2622.7399568856836 | 2786.3062471593025 | 2662.3724216627306 | 2536.2502461649146 | 2783.0000014019263 | 2577.2725768858516 | 2210.4338986185858 | 2891.83632761137 | 2357.600912625826 | 2304.390995297808 | 2709.0956985228204 | 2662.602243197916 |
| SE_147_3_30 | SE_147 | SE_147_3 | CGCCTCTCATGCCCACAGCC | 1268.0189047753097 | 1382.1598062486785 | 1401.300618078418 | 1391.0987753464822 | 1486.9535068466282 | 1719.6255680887914 | 1340.8961235047507 | 1587.2485106438346 | 1278.6988000682445 | 1100.071479743781 | 1656.210585188778 | 1334.783164637142 |
| SE_147_3_31 | SE_147 | SE_147_3 | CTGGCTGTGGGCATGAGAGG | 736.9682523480433 | 722.560766390307 | 609.3040742455137 | 762.5667748859106 | 689.3864332741144 | 704.3101193230164 | 756.5409107735471 | 800.7144065010887 | 504.37563780469645 | 423.5530349444246 | 867.4270889117696 | 653.4634101658355 |
| SE_147_4_1 | SE_147 | SE_147_4 | GGGACGTCAATAACAGTCTA | 1761.1373677434858 | 1458.1136350808545 | 1432.9014776827235 | 1573.2819638857782 | 1787.1017539490504 | 1532.502584357215 | 1872.5194662104866 | 1501.2213430032218 | 1269.8189472899928 | 1431.0406326951902 | 1879.2297284813844 | 1822.2691899828806 |
| SE_147_4_14 | SE_147 | SE_147_4 | AAAAATGTACAGATCAGCAG | 37.93218945909046 | 26.983597085115196 | 25.67569842498144 | 37.737946197425615 | 47.7627507282331 | 75.36897955855156 | 34.43713960846872 | 30.25131169779792 | 27.527543612580263 | 16.76715973101853 | 7.042709788187574 | 0.0 |
| SE_147_4_5 | SE_147 | SE_147_4 | GAAATGGTCCCAGCTCTCCA | 55.27261892610324 | 104.9362108865591 | 95.79010567555078 | 57.25757354092162 | 76.36280491651729 | 88.3636312065777 | 43.0464245105859 | 80.35504669727572 | 111.88614500597139 | 75.08771531716994 | 30.518409082146157 | 77.76562785277261 |
| SE_147_4_6 | SE_147 | SE_147_4 | GCTCTGCTCTCCCACATTGG | 207.0013767624651 | 287.8250355745621 | 279.470102125576 | 286.2878677046081 | 323.48132638246904 | 265.9572037296015 | 225.99372868057597 | 171.10898179066947 | 323.2266411283618 | 441.0492016202701 | 160.80854016361627 | 572.2157392748791 |
| SE_147_4_7 | SE_147 | SE_147_4 | GCTCTCCCACATTGGTGGCG | 48.76995785973454 | 77.95261380144339 | 109.61548175243439 | 55.95626505135522 | 71.05983235287025 | 78.83421999802519 | 67.79811860417279 | 94.5353490556185 | 145.62958556332785 | 42.282402799959776 | 41.08247376442752 | 83.56903291641235 |
| SE_148_1_1 | SE_148 | SE_148_1 | GTTGCCACTCTGACAAGTGA | 332.71949039830776 | 291.8226055130977 | 260.7070917355196 | 297.9996441107057 | 323.48132638246904 | 239.1015903236808 | 291.63952605921946 | 474.56745225920486 | 270.8355097366768 | 352.11035435138916 | 266.4491869864299 | 399.2742683784146 |
| SE_148_1_12 | SE_148 | SE_148_1 | GTAGGTCTTTGTAGTCCAGT | 667.6065344799921 | 627.6184803500868 | 644.8550413003572 | 545.2482571283218 | 568.478658822962 | 382.90906856183665 | 721.0276105523138 | 636.2228991443125 | 553.2148280850807 | 1105.1745283575692 | 522.3343092905784 | 755.6033392858951 |
| SE_148_1_16 | SE_148 | SE_148_1 | GGGCTAGGTACAAAGCACAG | 291.5359704141524 | 339.7934447755247 | 413.77375544387394 | 378.6807704638225 | 565.2968752847738 | 355.1871450460476 | 363.74228711445085 | 450.9336149953003 | 324.11462640618697 | 656.1062503442034 | 539.9410837610474 | 263.4745898892445 |
| SE_148_1_25 | SE_148 | SE_148_1 | CTCAGTAAGTAGAAGCACCT | 450.8511661423323 | 480.7077851089041 | 384.14794956483763 | 501.00376848306416 | 519.6913112374093 | 369.0481068039421 | 379.88469630592056 | 499.1466430136657 | 452.87249169083657 | 494.9957155374601 | 697.2282690305699 | 728.9076759931522 |
| SE_148_1_26 | SE_148 | SE_148_1 | CCTCAAGCTAGATACCAACA | 1753.5509298516677 | 1663.988486915437 | 1732.1221170609901 | 1945.4561919017688 | 1807.2530496909092 | 2136.3207309354957 | 2019.9534701592434 | 2066.5427303558204 | 1820.369819541598 | 2234.4062858944258 | 2047.0809784331882 | 2888.9350406798662 |
| SE_148_2_14 | SE_148 | SE_148_2 | TCTGTGATGTTATTTGGAGG | 352.22747354869716 | 334.7964823523552 | 377.23526152639585 | 325.3271223916001 | 309.69359771698674 | 269.4224441690751 | 352.9806809868044 | 285.49675414796786 | 350.75418474094204 | 166.94259036535843 | 397.9131030325979 | 436.41606078570896 |
| SE_148_2_3 | SE_148 | SE_148_2 | GAACTGTTCCAGGAAGCCAG | 261.19021884688004 | 296.81956793626716 | 256.75698428498146 | 261.5630064028465 | 306.51181417879325 | 284.1064017698669 | 207.97776792235607 | 260.17968640277473 | 312.743979330737 | 187.3547848205114 | 102.11929192871983 | 321.50864052564197 |
| SE_148_2_6 | SE_148 | SE_148_2 | TCACAGCAGAGGGAACCTGG | 3062.7533546111326 | 3104.1130572728816 | 3078.1212308318736 | 2972.1885901696587 | 2791.4847575037984 | 2750.534598832198 | 3156.379077238711 | 3123.447932797635 | 3375.2320410134703 | 3680.756064430981 | 3339.418224565608 | 2617.335683701526 |
| SE_148_2_7 | SE_148 | SE_148_2 | CCAGACACATGGATTTGTGG | 402.0812082663589 | 411.7497036691652 | 425.62407779548846 | 404.7069402551505 | 398.78353678625695 | 402.8342018810006 | 377.73237508039125 | 533.1793686736884 | 503.4876525268713 | 724.6329031579313 | 684.3166344188926 | 224.01143545649424 |
| SE_148_2_8 | SE_148 | SE_148_2 | GAGGAATCACATTTTCCCCC | 1497.7795952132292 | 1364.1707415252683 | 1471.4150253254707 | 1569.3780384170789 | 1408.4695129046522 | 1821.8501610532635 | 1638.992613240558 | 1639.2429526244248 | 1716.4755420360532 | 1710.25029256389 | 1618.6494663184442 | 2026.5490482229998 |
| SE_148_3_13 | SE_148 | SE_148_3 | GTCGAATCATGTCGATAAAA | 256.8551114801268 | 365.777649376006 | 379.2103152516649 | 229.03029416368648 | 317.11775930609264 | 389.8395494407839 | 328.2289868932175 | 255.24544245016995 | 302.8029797383829 | 392.20573631686824 | 427.25772175004615 | 421.32720762024564 |
| SE_148_3_14 | SE_148 | SE_148_3 | TCGAATCATGTCGATAAAAT | 565.7315113612921 | 754.5413258895781 | 779.1586946186552 | 680.5843400432275 | 748.7797259869611 | 658.3959683499906 | 652.1533313553764 | 540.7421965981379 | 809.8425733765548 | 933.8578963232495 | 833.3873249355296 | 598.911402567622 |
| SE_148_3_15 | SE_148 | SE_148_3 | TCATGTCGATAAAATGGGCT | 2144.7943697011438 | 2004.7813241755957 | 2064.9186697688315 | 2053.46479653578 | 1936.6455802438968 | 1983.8501515986559 | 1871.443305597722 | 1798.062339037864 | 2441.959514019217 | 1408.4414174055566 | 1910.9219225282284 | 2692.7799495288427 |
| SE_148_3_16 | SE_148 | SE_148_3 | ATGAATCTCACTGTTATGTG | 248.18489674662044 | 207.87363680385042 | 193.55526507637063 | 206.90804984105768 | 267.2698172078105 | 288.4812665861801 | 227.06988929334062 | 331.8190751852209 | 269.94754245588516 | 142.15635424124406 | 406.12959778548344 | 249.54641773650914 |
| SE_148_3_17 | SE_148 | SE_148_3 | TATGAATCTCACTGTTATGT | 584.1557176699931 | 612.6275930805783 | 489.81332386673387 | 542.645640149189 | 506.9641770846564 | 429.68981449473074 | 521.9378979108541 | 403.665940467491 | 668.6529142023528 | 954.2700907784025 | 385.00146842092073 | 768.3708304259026 |
| SE_148_4_0 | SE_148 | SE_148_4 | GTATGTAGTCATCTTCTCAA | 216.75536833765977 | 282.82807315139263 | 320.9462303562268 | 251.15253848631528 | 307.572408691528 | 248.6310015322333 | 229.2222105188699 | 212.70453537514163 | 353.4181405744176 | 313.47298627556387 | 154.93961534012664 | 452.6655949639003 |
| SE_148_4_10 | SE_148 | SE_148_4 | TGGACATGAGAGCGCGATGGA | 1354.7210521103736 | 1399.1494784874546 | 1381.550080825727 | 1253.1600754524436 | 1278.016387838935 | 1526.4384135881362 | 1332.2868386026337 | 1046.5061340456969 | 1257.3871534004404 | 1069.4531880610516 | 1124.485996180616 | 1486.8323773045033 |
| SE_148_4_11 | SE_148 | SE_148_4 | AGGTGACTATGCAGTGCTGA | 2638.996609511008 | 2766.3183974666244 | 2743.3496243987634 | 2713.228200745945 | 2841.3326996020805 | 2478.5132243335174 | 2667.802159043561 | 2728.29017374515 | 2336.2892659580216 | 2212.536077549619 | 2360.4815640075353 | 2415.3771874868626 |
| SE_148_4_17 | SE_148 | SE_148_4 | CTACATTTCCTACGACTCCT | 137.63965889441397 | 129.9210230024065 | 147.14150253254707 | 134.03477442533924 | 174.99809460035212 | 236.50265999407557 | 132.36775537005164 | 180.56251669623134 | 92.35648367881766 | 187.3547848205114 | 102.11929192871983 | 381.86405318749536 |
| SE_148_4_19 | SE_148 | SE_148_4 | GAGGTAAACGACTAACTGCA | 546.2235282109026 | 145.91130275654885 | 278.48257526629414 | 529.632555253525 | 98.63528968383483 | 1277.807412055903 | 860.928490211718 | 288.33281461963645 | 963.4640264403092 | 39.36637502065221 | 197.19587406925208 | 176.42351393464833 |
| SE_148_5_11 | SE_148 | SE_148_5 | GAGCTGGAGCTAAATCTGAA | 229.76069643791938 | 338.7940522908908 | 336.74666015837954 | 273.2747828089441 | 259.84565561870465 | 533.6470276789398 | 227.06988929334062 | 299.67705650631063 | 220.2203489006421 | 187.3547848205114 | 292.2724562097843 | 301.77706330926685 |
| SE_148_5_12 | SE_148 | SE_148_5 | AGTCAGTTAACAATTGGAAT | 100.7912462770118 | 199.87849692677923 | 196.51784566427426 | 173.07402911233126 | 233.33079280046948 | 161.99999054539242 | 162.50025252746178 | 218.37665631847872 | 225.54826056759313 | 141.42734729641717 | 191.32694924576242 | 226.33279748195014 |
| SE_148_5_3 | SE_148 | SE_148_5 | TTATGACTCACATATTGAAA | 72.61304839311603 | 75.95382883217611 | 67.151826659149 | 92.39290275921444 | 102.87766773475245 | 10.395721318420904 | 83.94052779564251 | 54.83050245225873 | 126.98189472899928 | 51.030486137882484 | 122.07363632858463 | 69.64086076367697 |
| SE_148_5_6 | SE_148 | SE_148_5 | TCAATATGTGAGTCATAAGG | 345.72481249856736 | 418.7454510616025 | 410.811748559703 | 317.5192714542017 | 461.35861303729195 | 540.83420979493345 | 239.98381664651637 | 323.3108937702153 | 357.85080669635434 | 435.94615300648184 | 388.52282331501453 | 421.32720762024564 |
| SE_148_5_9 | SE_148 | SE_148_5 | GGATTTGTAAACAGAATCAT | 68.27794102636283 | 103.93681840192521 | 79.98967587339807 | 65.06542447832003 | 108.18064029839948 | 61.50801780065702 | 77.48356411905462 | 83.19110716894428 | 205.12459917761421 | 81.64877782061198 | 89.2076573170426 | 157.85261773100112 |
| SE_148_6_0 | SE_148 | SE_148_6 | GAGCAAAGTGAGTATCACTG | 1120.625254305701 | 1151.3001422982484 | 1082.3294414474603 | 1171.1776406097604 | 1111.5030493404183 | 1174.7165089815621 | 1250.4986320325204 | 1000.1839930084437 | 1239.627447843937 | 1356.6819243228472 | 1119.7908563218243 | 1370.7642760317083 |

| | | | | | | | | | | | | | | | |
|---|---|---|---|---|---|---|---|---|---|---|---|---|---|---|---|
| SE_148_6_1 | SE_148 | SE_148_6 | AGCAAAGTGAGTATCACTGT | 873.524134400769 | 704.5717016668968 | 757.4331036406953 | 951.2565058730387 | 730.7496192705612 | 846.3849773414353 | 1023.4287427391797 | 862.1623833872408 | 553.2148280850807 | 667.7703614614337 | 855.6892392647902 | 1088.7187899388166 |
| SE_148_6_2 | SE_148 | SE_148_6 | TTGCTTGTGATCACCTCAGT | 422.6729682584366 | 317.806810113579 | 364.39741231214674 | 389.09123838035373 | 341.51143309886896 | 400.2352707592048 | 389.5701418208024 | 403.665940467491 | 270.8355097366768 | 273.37760431008473 | 304.01030585676364 | 526.949179778489 |
| SE_148_6_3 | SE_148 | SE_148_6 | GCTGTGACAGGATCCTACTG | 1727.5402856511485 | 1676.9805892156778 | 1706.4464186324922 | 1750.2599184668086 | 1694.830031341592 | 1498.7164900723471 | 1673.4297528490267 | 1599.538106021065 | 1981.0951548279538 | 2287.6237928667892 | 2142.1575605737203 | 1862.893025428359 |
| SE_148_6_4 | SE_148 | SE_148_6 | GTGTGAATGCAGGCTGTGAC | 2824.322449439707 | 2430.5225226296357 | 2549.7943593223927 | 2545.3594055918793 | 2626.032013518011 | 2494.106806311149 | 2489.1594973246297 | 2793.519564593527 | 2243.938797064204 | 2501.2228277010686 | 2887.5110131569054 | 2646.352709019725 |
| SE_149_1_17 | SE_149 | SE_149_1 | GTTGCATGTGGAGTCTAAAA | 139.80721257779055 | 271.8347558204198 | 282.4326827134796 | 162.66356119580004 | 240.75495438957532 | 394.171099990126 | 136.67239782111022 | 244.84655405405192 | 311.6828325166346 | 244.21732651700904 | 133.8114859755639 | 356.32907090748046 |
| SE_149_1_19 | SE_149 | SE_149_1 | TTTAGCTGTTAGAAATTCAA | 5.4188842084414945 | 14.990887269508443 | 16.78795666478725 | 6.506542447832002 | 14.848323178211695 | 5.197860659210452 | 1.0761606127646475 | 3.78141396222474 | 0.8879852778251698 | 6.561062503442034 | 8.216494752885502 | 25.534982280014887 |
| SE_149_1_23 | SE_149 | SE_149_1 | CGAGAATAGCTTGAACCTGG | 22974.985266950247 | 24374.183307736093 | 24157.86964062885 | 23659.089648806726 | 25103.211521792327 | 24720.158985095044 | 22793.081778355234 | 25251.337086246258 | 24668.231017983217 | 24468.389096169823 | 24823.2044334318 | 26405.49303956085 |
| SE_149_1_5 | SE_149 | SE_149_1 | AGGAAGCATGAAATGAGCTG | 3537.4476112706075 | 3634.7904666134805 | 3573.8597158744146 | 3376.8955304248093 | 3494.658919443395 | 3162.8982111295604 | 3470.6179761659882 | 3486.4636731712103 | 3425.847201849505 | 3774.79796031365 | 3637.559605598882 | 3398.474005267436 |
| SE_149_1_9 | SE_149 | SE_149_1 | CTGAGCCAGTATATACCAAG | 203.7500462374002 | 267.8371858818842 | 196.51784566427426 | 208.20935833062407 | 220.6036586477166 | 124.74865582105086 | 185.09962539551935 | 201.36029348846742 | 236.20408390149515 | 66.33963197924723 | 95.07658214053225 | 308.74114938563457 |
| SE_149_2_12 | SE_149 | SE_149_2 | TACTCTTCTTTATGACCACG | 162.56652625324483 | 203.8760668653148 | 215.2808560543306 | 227.72898567412008 | 218.48246962225778 | 208.7807364782865 | 133.4439159828163 | 158.81938641343908 | 163.38929111983123 | 75.08771531716994 | 190.1531642810645 | 285.52752913107554 |
| SE_149_2_17 | SE_149 | SE_149_2 | AGCCAGCAAGAGTTAGGCAG | 2592.394205318411 | 2478.4933618920627 | 2404.6279105151148 | 2886.302229858276 | 2469.064025634059 | 2332.9731258756246 | 3012.173555128248 | 2355.820898466013 | 2275.018281788085 | 3042.145980762623 | 2599.933696805913 | 2998.0390558762933 |
| SE_149_2_2 | SE_149 | SE_149_2 | TATGCTTCCTTCCTGACAAG | 877.8592417675221 | 793.5176327993136 | 800.8842855966152 | 851.0557521764259 | 792.2641010088669 | 781.4117191013047 | 815.7297444756027 | 583.2831036731661 | 869.3375869908413 | 886.4724449095015 | 652.6244403720485 | 1630.756822882769 |
| SE_149_2_22 | SE_149 | SE_149_2 | CTTTCCATGCAAAACTGTGG | 100.7912462770118 | 106.9349958558269 | 113.56558920297256 | 115.81645557140963 | 163.33155496032865 | 146.40640856776108 | 86.0928490211718 | 90.75393509339376 | 142.07764445202716 | 78.00374309647752 | 109.1620017169074 | 276.24208102925195 |
| SE_149_3_11 | SE_149 | SE_149_3 | TGTGTCCTCAGCCATAGGAG | 457.3538271924621 | 383.76671409941616 | 463.15009857560113 | 343.54544124552973 | 427.4195886299509 | 346.5240439473635 | 362.6661265016862 | 425.40907075028326 | 312.57081779445974 | 180.79372231706938 | 389.69660827971245 | 558.2875671221436 |
| SE_149_3_15 | SE_149 | SE_149_3 | TGTCAGCGATGGGAATATAG | 872.4403575590807 | 829.4957622461338 | 814.7096616734988 | 770.3746258233091 | 703.1741619395967 | 710.3742900920952 | 825.4151899904846 | 821.5121832933248 | 671.3168700358284 | 1180.2622436747392 | 820.4756903238524 | 616.3216177585412 |
| SE_149_3_16 | SE_149 | SE_149_3 | AGAGGCATCTCAGACTTCTG | 1322.2077468597247 | 1524.0735390666916 | 1620.5315815832867 | 1557.6662620109814 | 1623.7701989887216 | 1605.2726335861614 | 1533.5288731896226 | 1558.8879059271492 | 1617.0211909196341 | 1487.9031743916878 | 2079.9469574447303 | 2009.1388330320806 |
| SE_15_1_16 | SE_15 | SE_15_1 | GGTGGGTGGGTTACCTGGCG | 604.7474776620708 | 915.4435159246489 | 885.811595783186 | 667.5712551475634 | 837.8696650562314 | 814.3315032763043 | 447.6828149100933 | 834.7471321611114 | 912.8488656042746 | 687.4535489717598 | 741.8320976890911 | 821.7621570113882 |
| SE_15_1_21 | SE_15 | SE_15_1 | CAGCAGCAGCAGCAGCAGGC | 2275.9313675454277 | 2981.187781662912 | 2965.5431684915357 | 2417.831173614372 | 2988.755336871468 | 3331.8286825539 | 2382.6195966609293 | 3085.633793175388 | 2996.062327382123 | 2595.993730528565 | 2956.7643260740833 | 2989.9142887871976 |
| SE_15_1_22 | SE_15 | SE_15_1 | GCAGCAGCAGGCTGGCGAGC | 9419.104531113006 | 9843.016581159243 | 9838.730132427962 | 9809.263394351527 | 10060.799547751152 | 9201.07967691237 | 9400.262952499195 | 9374.125212355131 | 9246.590697993493 | 9689.960310639057 | 9169.608144220221 | 10846.564063942687 |
| SE_15_1_23 | SE_15 | SE_15_1 | CAGCAGCAGGCTGGCGAGCA | 5124.096907502278 | 4759.107011826613 | 4638.413673794454 | 5073.801800819395 | 4924.340322602636 | 5030.66280800585 | 5027.822382836433 | 4801.450378534863 | 4347.575920232031 | 4831.129023367817 | 5185.781974035451 | 5112.799861066617 |
| SE_15_1_24 | SE_15 | SE_15_1 | AGGGCGAGCAGTGCGCCCCC | 857.2674817754445 | 851.4823969080795 | 842.360413827266 | 923.9290275921443 | 832.5666924925843 | 1129.668383268405 | 907.2033965605979 | 722.2500667849254 | 1027.3989664437215 | 737.0260212199885 | 765.3077969830497 | 890.2423367623372 |
| SE_15_3_11 | SE_15 | SE_15_3 | GCACTAGATAAGGAGATGGA | 1330.877961593231 | 1404.146440910624 | 1445.7393268969724 | 1350.7582121699236 | 1579.2252294540867 | 1289.9357535940605 | 1157.9488193347606 | 1288.5168076280802 | 1268.0429767343423 | 1280.1361951160236 | 897.9454979939158 | 1763.0744583337553 |
| SE_15_3_17 | SE_15 | SE_15_3 | GTTTGAATAATGCTATGGGG | 14.089098941947885 | 20.98724217731182 | 23.700644703229056 | 23.423552812195208 | 20.15129574185873 | 1.7326202197368175 | 17.21856980423436 | 34.97807915057884 | 21.311646667804077 | 0.0 | 1.173784964697929 | 4.642724050911798 |
| SE_15_3_26 | SE_15 | SE_15_3 | CTAGTCACCCCATTGGTAAA | 172.32051782843953 | 281.8286806667587 | 251.81934997180872 | 243.34468754891688 | 212.11890254588135 | 311.87163955262713 | 205.54667703804768 | 201.36029348846742 | 267.2835686253761 | 166.94259036535843 | 184.28423945757487 | 374.8999671111277 |
| SE_15_3_27 | SE_15 | SE_15_3 | CTGGCTTTGTAGTATCAGCA | 149.56120415298525 | 168.89732990312845 | 190.592684488467 | 178.27926307059687 | 201.5129574185873 | 318.8021204315744 | 157.11944946363852 | 274.1525122612937 | 158.06137945288023 | 96.95792366197672 | 381.4801135268269 | 141.60308355280984 |
| SE_15_3_9 | SE_15 | SE_15_3 | GCTACAATGTAAAAGCGTGG | 1416.4963320866066 | 1589.0340505678948 | 1602.7560980558649 | 1478.286444147431 | 1412.7118909555697 | 1586.2138111690563 | 1348.4292477941033 | 1402.9045799853786 | 1418.112488686796 | 1290.3422923436 | 1455.493356225432 | 1456.6546709735765 |
| SE_15_4_1 | SE_15 | SE_15_4 | GATAAGACTAAAAGTTAACA | 31.429528408960667 | 41.97448435462364 | 57.27655803280356 | 61.16149900962082 | 63.6356707637644 | 33.78609428486794 | 26.904015319116187 | 39.70484660335977 | 86.13457194904147 | 45.198430579267345 | 39.90868879972959 | 29.017025318198737 |
| SE_15_4_18 | SE_15 | SE_15_4 | TTGAGAACAATGCAACAGCA | 995.9909175115467 | 1004.3894470570657 | 1007.2773998872349 | 1008.5140794139603 | 932.2625766891485 | 999.7218667881436 | 984.6869606796524 | 884.8508671605891 | 800.074735320478 | 1476.9680702192845 | 826.344615147342 | 847.2971392914031 |
| SE_15_4_2 | SE_15 | SE_15_4 | ACAGGGAGAAAGTTACAGAG | 831.2568375749253 | 1239.2466809460313 | 1240.3337394689872 | 893.9989323321171 | 1103.018293238583 | 1453.6683643591898 | 823.2628687649553 | 987.8943976312133 | 1430.5442825763485 | 1303.4644173504842 | 985.9793703462603 | 1418.352197553554 |
| SE_15_4_21 | SE_15 | SE_15_4 | AGAAGGAACAGCAAGTAAAA | 535.3857597940197 | 706.5704866361646 | 623.1294503223972 | 648.0516278040674 | 629.9931405612676 | 550.973229876308 | 500.41468493556107 | 449.0429080141879 | 854.2418372678134 | 492.07968775815255 | 438.9955767970255 | 508.3782835748419 |
| SE_15_4_27 | SE_15 | SE_15_4 | GAATTGATAGAAAAGGGGAC | 8.670214733506391 | 19.987849692677923 | 17.77548352742179 | 23.423552812195208 | 22.272484767317543 | 19.05882241710499 | 11.837766740411123 | 24.57919075446081 | 14.207764445202717 | 53.94651391719005 | 1.173784964697929 | 37.14179240729438 |
| SE_15_5_10 | SE_15 | SE_15_5 | ACTCTTGAAATGAAAAGGAG | 22.759313675454276 | 34.97873696218637 | 35.55096705484358 | 33.834020728726415 | 21.211890254588134 | 38.11764483420998 | 10.761606127646475 | 13.23494886778659 | 41.73530805778298 | 0.7290069448268927 | 7.042709788187574 | 9.285448101823595 |
| SE_15_5_14 | SE_15 | SE_15_5 | GCTGATTTGTAAGCCCAGGA | 96.4561389102586 | 191.88335704970808 | 157.0167711588925 | 137.93869989403845 | 193.02820131675202 | 239.9679004335492 | 145.28168272322742 | 99.26211650839943 | 176.7090702872088 | 147.2594028550323 | 99.77172199932397 | 99.81856709460365 |
| SE_15_5_19 | SE_15 | SE_15_5 | GTGAGCAGATCCCATCTAAG | 1102.201047997 | 1251.239390761638 | 1161.3315904582237 | 1229.7365226402485 | 1310.8948177335467 | 1086.3528777749846 | 1170.8627466879364 | 1815.0787018678752 | 996.3194817198405 | 1143.0828894885676 | 2490.7716950890053 | 1157.1989696897656 |
| SE_15_5_7 | SE_15 | SE_15_5 | TCATATGCTGGACTTCTACC | 1383.9830268359576 | 1381.1604137640445 | 1441.7892194464343 | 1363.7712970655878 | 1251.5015250207 | 1323.7218478789284 | 1362.4193357600436 | 1638.2975991338685 | 1534.4385600818935 | 1141.6248755989138 | 1714.8998334236744 | 1232.6432355170823 |
| SE_15_5_9 | SE_15 | SE_15_5 | CTAAACCAGGAACAACATCG | 1152.0547827146618 | 1220.2582237379872 | 1196.8825575130675 | 961.66697378957 | 1235.592607329759 | 1308.9945760111657 | 972.8491939392413 | 1379.2707427214739 | 1367.4973278507614 | 1000.9265352473236 | 1375.675978625973 | 1231.4825545043543 |
| SE_150_1_0 | SE_150 | SE_150_1 | GTAGAATGCGAAGCATCCCG | 364.1490188072684 | 519.684092009626 | 482.900635828292 | 324.0258139020337 | 439.0861282699744 | 666.1924744888063 | 351.90452037403975 | 281.7153401857431 | 349.8661994631169 | 228.1791737308174 | 221.8453583279086 | 478.2005772439152 |
| SE_150_1_10 | SE_150 | SE_150_1 | CTCAACCCACAGCCTCCCCA | 719.6278228810305 | 759.5382883217611 | 798.9092318713462 | 818.5230399372658 | 880.2934455654076 | 868.9090401980139 | 732.8653772927249 | 640.9496665970935 | 639.3494000341223 | 998.010507468016 | 958.982316158208 | 338.9188557165612 |
| SE_150_1_26 | SE_150 | SE_150_1 | CCAGCCTCAGTGTAGAGAAG | 741.3033597147964 | 725.5589438442087 | 684.356115805739 | 598.6019052005443 | 749.8403204996906 | 1304.6630254618235 | 716.7229681012552 | 618.261182823745 | 961.6880558846589 | 1221.0866325850452 | 708.9661186775492 | 674.3556683949387 |
| SE_150_1_3 | SE_150 | SE_150_1 | CCAACGATATGATCACCTCA | 2102.5270728753 | 1873.8609086885554 | 1849.637813714501 | 1970.1810532035302 | 1729.8296502616624 | 1745.6148713848436 | 2053.3144491549474 | 1575.9042687571605 | 1949.1276848262478 | 1871.3608273706334 | 1841.6686096110507 | 2103.1539950630445 |
| SE_150_1_7 | SE_150 | SE_150_1 | TCTCCCAGCCTGAGAATCCA | 227.59313675454277 | 224.86330904262664 | 214.29332919169605 | 171.77272062276487 | 307.572408691528 | 248.6310015322333 | 205.54667703804768 | 316.693419336322 | 436.88875668998355 | 182.98074315155006 | 286.4035313862947 | 189.19100507465575 |
| SE_150_2_14 | SE_150 | SE_150_2 | TCACACAGATAGCAAAGCTG | 2857.9195315320444 | 2665.379756518601 | 2621.8838202947145 | 2626.040531944996 | 2791.4847575037984 | 2554.7485140019376 | 2604.308682890447 | 2957.0657184597467 | 2510.334380411755 | 2096.6239733221432 | 3364.0677088242646 | 2365.467903939561 |
| SE_150_2_19 | SE_150 | SE_150_2 | TGGGTTCCAGAAGAGGGGCA | 4.335107366753196 | 5.9963549078033775 | 1.975053725269088 | 1.3013084895664004 | 3.1817835381882205 | 16.459892087499767 | 0.0 | 0.0 | 23.087617223454416 | 0.0 | 0.0 | 1.1606810127279494 |
| SE_150_2_35 | SE_150 | SE_150_2 | GGGAACTCAAGGGGCCCCAA | 3.2513305250648967 | 1.9987849692677924 | 5.925161175807264 | 6.506542447832002 | 0.0 | 0.8663101098684087 | 3.228481838293942 | 0.0 | 0.8879852778251698 | 0.0 | 3.521354894093787 | 0.0 |
| SE_150_2_6 | SE_150 | SE_150_2 | GCTTCCTGCACCCTCCACAC | 1813.158656144524 | 1773.9216602251656 | 1847.662759989232 | 1728.1376741441798 | 1725.5872722107447 | 1419.0159599644535 | 1670.2012710107329 | 1793.335571585083 | 1600.149470640956 | 1990.917966322244 | 1781.8055764114563 | 1416.0308355280984 |
| SE_150_2_9 | SE_150 | SE_150_2 | TTAGGACCAACGCCACCAGG | 389.0758861660993 | 322.80377253674845 | 347.6094556473595 | 344.8467497350961 | 370.147484942563 | 705.1764294328847 | 375.58005385486194 | 291.168875091305 | 388.0495664095992 | 252.23640291010486 | 458.9499211968903 | 593.1079975039821 |
| SE_150_3_1 | SE_150 | SE_150_3 | TCTCCTGGAACATCCTGTAG | 30.34575156727237 | 66.95929647047105 | 46.41376254382357 | 39.03925468699201 | 51.96913112374093 | 13.86096175789454 | 63.4934761531142 | 158.81938641343908 | 15.095749723027886 | 66.33963197924723 | 332.1811450095139 | 37.14179240729438 |
| SE_150_3_10 | SE_150 | SE_150_3 | GCAGAGGCTGTGAGCTCGGG | 3289.262714523987 | 3732.730930107602 | 3740.7517556596526 | 3723.0435886494715 | 4163.8940569756505 | 3835.1548563874453 | 3371.6111997916405 | 4118.905158353298 | 4281.865009672969 | 4405.388967588912 | 4366.480068676296 | 4264.342040762486 |
| SE_150_3_11 | SE_150 | SE_150_3 | GCAAGATGATTGTGACTCGT | 6773.605260551868 | 5362.740072545487 | 5421.522475863647 | 6464.900576165877 | 5736.755719353361 | 4860.866026471642 | 7159.6965567232 | 5330.8483332463275 | 5144.098714441208 | 5288.945384719106 | 5706.942498361331 | 5037.355595239301 |
| SE_150_3_12 | SE_150 | SE_150_3 | TACATAGGATGTGTATGGCG | 3045.41292514412 | 2734.33783795834 | 2928.017147711423 | 2877.1930704313113 | 2881.635291085798 | 3288.5131770604794 | 2941.1469546857816 | 2926.8144067619487 | 2648.8600837524814 | 2329.177188721922 | 3098.7923068025325 | 3347.4040407074062 |
| SE_150_3_14 | SE_150 | SE_150_3 | GCAGACCCTGGAGTGCCTGA | 1784.9804582606282 | 1970.8019796980432 | 1898.0266299835937 | 1907.718245704343 | 2031.0384918768139 | 1632.9945571019505 | 1858.5293782445463 | 2128.936060732529 | 1451.8559292441525 | 2309.494001211596 | 2058.8188280801673 | 2090.386503923037 |
| SE_150_4_0 | SE_150 | SE_150_4 | GGAGGTTTCTCACCGTGGTA | 252.52000411337366 | 317.806810113579 | 336.74666015837954 | 278.4800167672097 | 306.51181417879855 | 278.9518553776276 | 301.3249715741013 | 321.4201867891029 | 173.1571291759081 | 236.19825012391323 | 267.6229719511278 | 82.4083519036844 |
| SE_150_4_12 | SE_150 | SE_150_4 | TCTGCCCGAGAAGGGGGTGA | 982.9855954112871 | 1167.2904220523908 | 1160.3440635955892 | 1099.6056736836083 | 1121.0483999549829 | 1010.1175881065645 | 1160.1011405602899 | 1038.9434861212474 | 1183.6843753409514 | 1708.7922786742363 | 1031.7569839694797 | 786.9417266295497 |
| SE_150_4_16 | SE_150 | SE_150_4 | TGGGCTCTGGACATTGTCAA | 81.28326312662242 | 56.96537162413208 | 70.11440724705263 | 68.96934994701923 | 85.90815553108195 | 144.67378834802426 | 58.112673089290965 | 113.4424188667422 | 95.90241000511834 | 21.87020834480678 | 104.46686185811568 | 26.695663292742836 |
| SE_150_4_29 | SE_150 | SE_150_4 | TGTGCCGAGGAACTGGGCAG | 3191.7227987720403 | 3154.082681504576 | 3232.1754214028624 | 2912.3283996496043 | 2880.5746965730686 | 1956.1282280828668 | 2714.077065392441 | 1846.2753670562292 | 2580.4852173599434 | 1663.593848094969 | 1528.2680240367035 | 1193.180081084332 |
| SE_150_4_41 | SE_150 | SE_150_4 | GTGATCGTCTCTCTGCAGGA | 102.95879996038839 | 19.987849692677923 | 51.35139685699629 | 130.13084895664005 | 36.06021343279983 | 253.82886219144376 | 149.586325174286 | 57.66656292392729 | 208.6765402889149 | 7.290069448268927 | 36.3873339056358 | 71.96222278913287 |
| SE_150_5_10 | SE_150 | SE_150_5 | GGTGTCATCTAAATGCAATA | 146.30987362792035 | 200.87788941141312 | 224.1685978180415 | 179.58057156016326 | 186.6646342403756 | 41.582885273683615 | 174.33801926787288 | 119.11453981007931 | 227.32423112324346 | 167.67159731018532 | 75.12223774066746 | 196.15509115102344 |
| SE_150_5_18 | SE_150 | SE_150_5 | GCTGAGGCCCAGACATTGCA | 10.837768416882989 | 35.97812944682026 | 30.613332741670867 | 33.834020728726415 | 23.33307928004695 | 3.465240439473635 | 17.21856980423436 | 20.79777679223607 | 45.28724916908366 | 34.992333351690846 | 21.128129364562724 | 1.1606810127279494 |
| SE_150_5_20 | SE_150 | SE_150_5 | GCCAGGCATAAGCTCACAGA | 2876.3437378407452 | 2924.2224100387803 | 2993.193920645303 | 3086.703737251502 | 3201.934833930079 | 2799.0479649848285 | 2724.8386715200872 | 2953.284304497522 | 3216.282676282765 | 3975.2748701410455 | 2305.313670666733 | 3393.831281216524 |
| SE_150_5_21 | SE_150 | SE_150_5 | GCAATGTCTGGGCCTCAGCT | 1598.5708414902408 | 1865.8657688114843 | 1820.0120078354646 | 1905.1156287252102 | 1891.0400161965322 | 1461.4651553480055 | 1867.1386631466635 | 1464.3525568715306 | 1772.4186145390388 | 2273.0436539702514 | 1536.4845187895892 | 2057.8874355666544 |
| SE_150_5_5 | SE_150 | SE_150_5 | GCAAGCAGCAGCACTCCATG | 13005.322100259587 | 11697.889032639754 | 11516.538272044052 | 13087.25947956929 | 12585.01448804714 | 12306.801420790614 | 13527.338902451618 | 13430.63704033172 | 12324.347670935531 | 13921.116618414342 | 15069.051376792013 | 12324.110993145367 |
| SE_150_6_18 | SE_150 | SE_150_6 | GAGCAGGTCCGGGCTCCTCA | 21.675536833765978 | 54.96658665486429 | 62.21419234597627 | 29.93009526002721 | 114.54420737477592 | 46.78074593289407 | 4.30464245105859 | 82.2457536783881 | 44.39926389125849 | 2.9160277793075706 | 95.07658214053225 | 46.42724050911798 |
| SE_150_6_22 | SE_150 | SE_150_6 | GCCGGGAAAGAATACACTCA | 1611.5761635905005 | 1613.0194701991084 | 1675.8330858908212 | 1758.067769404207 | 1685.2846807270273 | 1788.930376878264 | 1559.3567278959742 | 2069.378790827489 | 1720.027483147354 | 1086.949354736897 | 2022.4314941745317 | 1482.1896532535914 |
| SE_150_6_23 | SE_150 | SE_150_6 | CTTTTCTACCTGAGGAGAGA | 1.083776841688299 | 6.9957473924372735 | 8.887741763710896 | 1.3013084895664004 | 14.848323178211695 | 12.128341538157722 | 0.0 | 0.0 | 0.0 | 0.0 | 1.173784964697929 | 6.964086076367696 |
| SE_150_6_5 | SE_150 | SE_150_6 | CAAGAAGCTAGGGTACTGCG | 3159.2094935213913 | 3040.1519382563124 | 3437.5810088308476 | 2939.6558779304987 | 2587.8506110597523 | 3166.3634515690337 | 2603.232522277682 | 2591.2139176145033 | 2901.0479026548296 | 2393.3297998666885 | 2693.836493981747 | 2604.5681925615186 |
| SE_150_6_9 | SE_150 | SE_150_6 | GTACAGTGGCAGGCAAGCAT | 4635.313551900855 | 3813.681721362948 | 3904.681214856987 | 4462.186810723187 | 4119.349087441015 | 3334.4276128835054 | 4571.530283024223 | 4528.2432197641265 | 4390.199213567639 | 3907.4772242721447 | 4128.201720842617 | 4563.797742046298 |
| SE_151_1_10 | SE_151 | SE_151_1 | CCCCACTGAGGGACACGGAA | 59.60772629285644 | 50.9690167163287 | 53.32645058226538 | 41.641871666124814 | 90.15053358199957 | 19.9251325269734 | 30.13249715741013 | 78.46433971616335 | 50.615160836034676 | 12.393118062057175 | 152.59204541073078 | 109.10401519642724 |
| SE_151_1_16 | SE_151 | SE_151_1 | GTTAAGCAGTGAACATCACG | 1010.0800164534946 | 986.4003823336556 | 1027.0279371399258 | 966.8722077478355 | 1107.2606712895006 | 873.240590747356 | 980.3823182285938 | 900.9218765000443 | 1025.622995888071 | 1360.3269590469818 | 672.5787847719133 | 797.3878557441012 |
| SE_151_1_22 | SE_151 | SE_151_1 | CAGTTCCCAGAGTGAAACCA | 306.7088461977886 | 370.7746117991755 | 340.6967676089177 | 281.0826337463425 | 319.23894833155146 | 272.02137449868036 | 303.47729279963056 | 311.96665188354103 | 330.3305233509632 | 400.95381965479095 | 288.75110131569056 | 402.75631141659846 |
| SE_151_1_8 | SE_151 | SE_151_1 | GAGTTCACCACAGAACCGCT | 480.1131408679164 | 630.6166578039885 | 697.1939650199881 | 528.3312467639586 | 579.0846039502561 | 690.4491575651217 | 399.2555873356842 | 441.4800800897384 | 890.6492336586452 | 605.8047711511477 | 407.30338275018136 | 820.6014759986602 |
| SE_151_1_9 | SE_151 | SE_151_1 | AAAAGAGTAGAGTAATACAT | 34.680858934025565 | 46.97144677779312 | 37.526020780112674 | 22.12224432262881 | 64.69626527649382 | 42.44919538355203 | 41.970263897821255 | 24.57919075446081 | 41.73530805778298 | 26.244250013768134 | 28.170839152750297 | 126.51423038734649 |
| SE_151_2_47 | SE_151 | SE_151_2 | TACTGGAGGTTGCGCCTCGG | 2308.4446727960767 | 2224.647670795053 | 2328.5883420922546 | 2076.8883493479752 | 2288.76295847006 | 1890.288659732868 | 2197.5199712654103 | 2499.5146290305534 | 2420.647867351413 | 2025.1812927291078 | 2554.1560831826937 | 1956.9081874593228 |
| SE_151_2_51 | SE_151 | SE_151_2 | TCCAGTACGGCGTCGGCCCG | 1417.580108928295 | 1316.1999022628413 | 1290.697609463349 | 1331.2385848264275 | 1539.9832324830986 | 1408.6202386460327 | 1177.3197103645243 | 1410.4674079098281 | 1356.8415045168595 | 1227.6476950884871 | 1196.0868790271898 | 1460.1367140117604 |
| SE_151_2_52 | SE_151 | SE_151_2 | CCCGCGGGCCGACGCCGTAC | 2.167553683376598 | 14.990887269508443 | 7.900214901076352 | 3.9039254686992013 | 16.969512203670508 | 14.727271867762948 | 3.228481838293942 | 0.0 | 1.7759705556503396 | 0.0 | 4.695139858791716 | 29.017025318198737 |
| SE_151_2_54 | SE_151 | SE_151_2 | GTCGGCCCGCGGGCCACTCC | 21.675536833765978 | 22.986027146579612 | 15.800429802152705 | 36.43663770785921 | 19.090701229129323 | 51.11229648223612 | 12.913927353175769 | 6.617474433893295 | 11.543808611727208 | 22.599215289633673 | 0.0 | 20.89225822910309 |
| SE_151_2_55 | SE_151 | SE_151_2 | TGGCGCCCGGAGTGGCCCGC | 19.50798315038938 | 29.981774539016886 | 31.60085960430541 | 14.314393385230405 | 30.757240869152795 | 24.256683076315444 | 9.685445514881827 | 30.25131169779792 | 30.191499446055772 | 13.851131951710961 | 19.954344399864794 | 25.534982280014887 |
| SE_151_3_2 | SE_151 | SE_151_3 | AATAAAAGCCCGTGTCACAG | 19.50798315038938 | 42.973876839257535 | 42.463655093285396 | 45.545797134824014 | 50.90853661101153 | 29.454543735525895 | 29.056336544645482 | 23.633837263904624 | 54.167101947335354 | 91.85487504818848 | 84.5125174582509 | 13.928172152735392 |
| SE_151_3_28 | SE_151 | SE_151_3 | TGGCTTCTGTGGACGCAGGA | 2.167553683376598 | 6.9957473924372735 | 6.912688038441808 | 18.218318853929606 | 22.272484767317543 | 5.197860659210452 | 10.761606127646475 | 16.071009339455145 | 9.767838056076867 | 0.0 | 49.29896851731302 | 25.534982280014887 |
| SE_151_3_3 | SE_151 | SE_151_3 | GGTCCCATCTGGTCCTCACA | 88.86970101844051 | 135.9173779102099 | 149.11655625781614 | 118.41907255054244 | 156.9679878839522 | 70.17111889934111 | 72.10276105523138 | 153.14726547010198 | 68.37486639253807 | 74.35870837234305 | 210.1075086809293 | 111.42537722188314 |
| SE_151_3_36 | SE_151 | SE_151_3 | GACCCTGCGGAGTCTCTGCG | 44.434850509220254 | 83.94896870924728 | 80.97720273603261 | 81.98243484268323 | 75.30221040378788 | 91.82887164605133 | 46.27490634887984 | 38.75949311280358 | 43.51127861343332 | 172.04563897914667 | 17.606774470468935 | 32.49906835638259 |
| SE_151_3_7 | SE_151 | SE_151_3 | CTGTACTGAGAGGTTCCGCG | 298.0386314642822 | 331.7983048984535 | 298.2331125156323 | 238.13945359065127 | 327.7237044333867 | 200.9839454894708 | 287.33488360816085 | 323.3108937702153 | 142.96562972985234 | 473.12550719265334 | 133.8114859755639 | 133.47831646371418 |

| | | | | | | | | | | | | | | | |
|---|---|---|---|---|---|---|---|---|---|---|---|---|---|---|---|
| SE_151_4_0 | SE_151 | SE_151_4 | GCAGAAGTTCAAAAAATGCA | 1551.968437297644 | 2004.7813241755957 | 1986.9040476207026 | 1666.976175134559 | 2031.0384918768139 | 2203.892919505232 | 1591.6415462789137 | 2135.553535166422 | 2211.083341784673 | 2123.597230280738 | 1834.6258998228632 | 2026.5490482229998 |
| SE_151_4_1 | SE_151 | SE_151_4 | TTATGAGTACATTCCTTAGT | 114.88034521895969 | 142.91312530264716 | 106.65290116453076 | 101.50206218617923 | 81.66577748016432 | 80.566840217762 | 80.71204595734856 | 130.45878169675353 | 152.7334677859292 | 72.17168753786237 | 157.2871852695225 | 109.10401519642724 |
| SE_151_4_3 | SE_151 | SE_151_4 | TTAGTGGGTGACTTCCAGAG | 192.9122778205172 | 265.8384009126164 | 261.6946185981542 | 257.6590809341473 | 252.42149402959882 | 272.88768460854874 | 171.10953742957895 | 416.9008893352776 | 354.30612585224276 | 206.3089653860106 | 462.471276090984 | 366.77520002203204 |
| SE_151_4_6 | SE_151 | SE_151_4 | TTTAAATCTTGACTTCCATG | 4.335107366753196 | 2.9981774539016888 | 6.912688038441808 | 1.3013084895664004 | 5.3029725636470335 | 0.0 | 1.0761606127646475 | 8.508181415005666 | 22.199631945629246 | 0.0 | 0.0 | 19.73157721637514 |
| SE_151_4_7 | SE_151 | SE_151_4 | CTTGTAATCCCAACAGTGGG | 109.46146101051819 | 135.9173779102099 | 97.76515940081985 | 122.32299801924164 | 97.57469517110542 | 233.03741955460194 | 86.0928490211718 | 44.431614056140695 | 60.382998892111544 | 100.60295838611118 | 31.692194046844083 | 99.81856709460365 |
| SE_152_1_10 | SE_152 | SE_152_1 | GACATAGCAGAGAAGACTGA | 2135.040378125949 | 2384.550468336476 | 2243.661031905684 | 2383.9971528856454 | 2339.6714950810715 | 2792.9837942157496 | 2205.053095554763 | 2531.6566477094634 | 2348.721059847574 | 2835.1080084317855 | 2387.478618195588 | 2595.282744459695 |
| SE_152_1_11 | SE_152 | SE_152_1 | TGACATAGCAGAGAAGACTG | 2264.0098222868564 | 2288.608789811622 | 2347.351352482311 | 2235.647985075076 | 1934.5243912184378 | 1918.010583248657 | 2290.06978396317 | 2068.433437336933 | 2368.256735959728 | 2442.1732651700904 | 1954.351966222052 | 2158.866683673986 |
| SE_152_1_16 | SE_152 | SE_152_1 | AGTTGTTCCTATTATCCCTA | 50.93751155935005 | 23.98541963121351 | 34.56344019220904 | 81.98243484268323 | 43.48437502190568 | 39.8502650539468 | 66.72195799140815 | 59.557269905039654 | 13.319779167377547 | 129.7632361791869 | 25.823269223354437 | 23.21362025455899 |
| SE_152_1_5 | SE_152 | SE_152_1 | ACAGGCAGAGATTCACCCTT | 10107.302825585075 | 9958.946109376775 | 9398.293151692955 | 9938.0929348186 | 9345.958846171532 | 10589.774783031427 | 10440.91026504261 | 9661.51267348421 | 9280.33413855085 | 9452.30404662549 | 10078.117706896419 | 11349.13894245389 |
| SE_152_1_7 | SE_152 | SE_152_1 | GTATCTTAGAGGAAACTCAT | 26.010644200519174 | 41.97448435462364 | 35.55096705484358 | 24.724861301761607 | 48.78734758555271 | 33.78609428486794 | 7.533124289352532 | 7.56282792444948 | 72.81479278166393 | 19.683187510326103 | 3.521354894093787 | 13.928172152735392 |
| SE_152_2_0 | SE_152 | SE_152_2 | ATCTTTGTGAATATAAGCAG | 13.005322100259587 | 26.983597085115196 | 17.77548352742179 | 22.12224432262881 | 20.15129574185873 | 1.7326202197368175 | 13.990087965940416 | 0.945353490556185 | 7.991867500426528 | 32.076305572383276 | 36.3873339056358 | 1.1606810127279494 |
| SE_152_2_1 | SE_152 | SE_152_2 | TCAGTTGCTGGGTTTTCAGT | 813.9164081079125 | 998.3930921492623 | 1000.3647118487931 | 797.7021041042035 | 928.0201986382309 | 810.8662628368305 | 731.7892166799603 | 1350.9101380047885 | 812.5065292100304 | 1439.788716033113 | 1126.8335661100118 | 786.9417266295497 |
| SE_152_2_12 | SE_152 | SE_152_2 | TTTCAGATTAGGGTATCAGA | 1.083776841688299 | 8.994532361705065 | 5.925161175807264 | 3.9039254686992013 | 5.3029725636470335 | 0.0 | 0.0 | 0.0 | 31.07948472388094 | 0.0 | 3.521354894093787 | 0.0 |
| SE_152_2_8 | SE_152 | SE_152_2 | TCAAATAATGTAATTTTGAA | 30.34575156727237 | 37.97691441608806 | 46.41376254382357 | 74.17458390528482 | 50.90853661101153 | 49.379676262499295 | 46.27490634887984 | 33.08737216946648 | 79.91867500426528 | 16.76715973101853 | 49.29896851731302 | 45.26655949639003 |
| SE_152_2_9 | SE_152 | SE_152_2 | GCAGACCCTACAGCCACCCT | 5749.4361451564255 | 4940.996444029983 | 4950.472162386969 | 5331.460881753543 | 5216.003813603223 | 4704.930206695328 | 5674.594911107986 | 5639.0335711676435 | 4919.43843915144 | 5159.18214853992 | 5419.365182010339 | 5318.240400319464 |
| SE_153_1_10 | SE_153 | SE_153_1 | GCAGGATTCAGCATACGAGG | 3020.486057785289 | 3375.947813093301 | 3285.501871985128 | 3313.1314144360554 | 3209.3589955191846 | 2864.021223224959 | 3178.978450106769 | 3881.6214322236956 | 3510.2058032428963 | 3835.305536734282 | 4062.469762819532 | 3830.247342002233 |
| SE_153_1_12 | SE_153 | SE_153_1 | CAAGGCAAGGAGGAGAGTGG | 44681.951629125186 | 41424.818488075 | 41321.086513217226 | 40171.39307291478 | 39962.140645131316 | 36048.029981734355 | 40244.10227494676 | 34392.90533992457 | 38366.29191371429 | 32399.984655886416 | 29436.179344694665 | 30109.226151175735 |
| SE_153_1_13 | SE_153 | SE_153_1 | AATGCCAGTTGCACTCTCAG | 435.6782903586962 | 442.730870692816 | 438.46192700973756 | 451.554045879541 | 520.7519057501387 | 534.5133377888081 | 483.19611513132674 | 454.71502895752496 | 437.7767419678087 | 291.60277793075704 | 602.1516868900376 | 391.14950128931895 |
| SE_153_1_2 | SE_153 | SE_153_1 | AGAAGAGTCATCACAGACAT | 1124.9603616724542 | 1151.3001422982484 | 1115.9053547770347 | 1189.39595946369 | 1064.8368907803244 | 1567.1549887519514 | 1343.0484447302802 | 1239.3584261191586 | 1097.5498033919098 | 1336.9987368125212 | 1503.618539778047 | 1173.4485038679568 |
| SE_153_1_6 | SE_153 | SE_153_1 | GCCATTGAAAGAGTTAGCAT | 1272.3540121420629 | 1463.110597504024 | 1456.6021223859525 | 1366.3739140447206 | 1311.9554122462762 | 1669.3795817164237 | 1417.3035270110406 | 1193.9814585724616 | 1565.5180448057743 | 1412.815459074518 | 1001.2385748873335 | 1616.8286507300336 |
| SE_153_2_17 | SE_153 | SE_153_2 | TCTTAGAGTGAGTGATCCAA | 373.90301038246315 | 419.7448435462364 | 464.1376254382357 | 334.4362818185649 | 433.78315570632736 | 781.4117191013047 | 433.6927269441529 | 456.60573593863734 | 465.304285580389 | 376.1675835306766 | 312.2268006096491 | 496.77147344756236 |
| SE_153_2_19 | SE_153 | SE_153_2 | TGATCCAAGGGAACATAAGG | 282.86575568064603 | 304.8147078133383 | 334.77160643311043 | 281.0826337463425 | 284.239329411481 | 349.1229742769687 | 209.85131948910626 | 194.7428190545741 | 237.09206917932033 | 242.75931262735526 | 257.05890726884644 | 279.7241240674358 |
| SE_153_2_2 | SE_153 | SE_153_2 | ACTAGATTAATAAGTCCTAG | 30.34575156727237 | 40.97509186998975 | 42.463655093285396 | 55.95626505135522 | 29.69664635642339 | 51.978606592104526 | 37.665621446762664 | 65.22939084837677 | 32.855455279531284 | 10.93510417240339 | 36.3873339056358 | 2.321362025455899 |
| SE_153_2_30 | SE_153 | SE_153_2 | GTAGCTGAGGTTGAGGGAAA | 565.7315113612921 | 587.642780964731 | 587.5784832675537 | 532.2351722326578 | 713.7801070668908 | 654.0641329506486 | 564.98432170144 | 642.8403735782058 | 519.4713875277243 | 489.163659978845 | 642.0603756897672 | 567.5730152239672 |
| SE_153_2_8 | SE_153 | SE_153_2 | AGTGTGGCTGGAGTAAAGCC | 1858.6772834954327 | 1905.84146819684 | 1855.5629748903082 | 1812.7227259659958 | 1835.889101534603 | 1901.5506911611571 | 1818.7114355722542 | 1673.2756782844474 | 1997.078889828807 | 1992.3759802118977 | 1502.4447548133492 | 1969.6756785993302 |
| SE_153_3_14 | SE_153 | SE_153_3 | TTGAGTGGAGATAAACGCTG | 680.6118565802517 | 700.5741317283612 | 733.7324589374662 | 720.9249032197858 | 694.6894058377615 | 957.2726714045916 | 679.0573466544926 | 931.1731881978423 | 789.418911986576 | 578.8315141925527 | 1070.4918878045112 | 878.6355266350578 |
| SE_153_3_18 | SE_153 | SE_153_3 | AAGAGCACAGGGGAGTTAGG | 14820.648310087488 | 16289.098107047874 | 16019.660765657572 | 15009.292118658863 | 15594.981715173197 | 15068.598051051102 | 14738.019591811848 | 15208.846956067904 | 15813.241827510623 | 15531.492959536949 | 14678.180983547603 | 15333.75685914894 |
| SE_153_3_7 | SE_153 | SE_153_3 | GCAGGGGCTAGAGAGTGGGT | 12840.588020322966 | 12114.635698732089 | 11987.58858552073 | 12751.521889261157 | 12068.50496034792 | 11622.416433994571 | 13353.000883183746 | 12013.552157987999 | 12184.045997039155 | 11121.000943334247 | 11551.21783759232 | 10889.509261413621 |
| SE_153_3_8 | SE_153 | SE_153_3 | AGTGGAAAGAACTTGCTGCC | 1701.5296414506292 | 1568.046808390583 | 1642.2571725612468 | 1823.133193882527 | 1498.6200464866517 | 1591.411671828267 | 1839.1584872147826 | 1859.510315924016 | 1702.2677775908505 | 1459.471903543439 | 1955.5257511867499 | 1133.9853494352067 |
| SE_153_3_9 | SE_153 | SE_153_3 | TCCGTTGGAATGGAAAGCCG | 1124.9603616724542 | 1054.3590712887606 | 1103.0675055627858 | 1182.889417015858 | 1035.140244423901 | 1146.1282753559049 | 1043.8757943817081 | 1242.1944865908272 | 1082.454053668882 | 940.4189588266916 | 1437.8865817549631 | 894.885060813249 |
| SE_153_4_10 | SE_153 | SE_153_4 | CACTTCAGCCTGGGTGACAC | 1558.471098347774 | 1335.1883594708852 | 1290.697609463349 | 1277.8849367542052 | 1196.3506103587708 | 1669.3795817164237 | 1438.8267392663336 | 1526.745887248239 | 1226.3076686765594 | 1495.193243839957 | 1462.5360660136196 | 1621.4713747809453 |
| SE_153_4_11 | SE_153 | SE_153_4 | AGGGGAACAGTGCCTTTCAG | 6254.476153383173 | 6378.122836933526 | 6400.16159673448 | 6267.101685751784 | 6357.203509300064 | 6093.625312814387 | 6058.784249864965 | 5811.087906448869 | 5846.495069200918 | 6722.902045193604 | 5408.801117328057 | 5635.106316794195 |
| SE_153_4_13 | SE_153 | SE_153_4 | ACTCCAACTTCTGCAGACAG | 65.02661050129794 | 10.993317330972857 | 23.700644703229056 | 45.545797134824014 | 4.242378050917627 | 57.176467251314975 | 112.99686434028798 | 9.45353490556185 | 76.3667338929646 | 2.9160277793075706 | 21.128129364562724 | 23.21362025455899 |
| SE_153_4_18 | SE_153 | SE_153_4 | CGAGGATCACTTGAACCCAG | 9886.212349880663 | 9596.166637454671 | 9445.694441099413 | 9301.75308342063 | 9850.80183423073 | 8890.940657579478 | 9306.636979188672 | 9410.993998486822 | 8900.276439641677 | 10220.677366473035 | 8779.91153594051 | 8851.353403063342 |
| SE_153_4_9 | SE_153 | SE_153_4 | AGGATCTTCCTGTGTCACCC | 566.8152882029804 | 569.6537162413208 | 555.9776236632483 | 514.0168533787282 | 551.5091466192915 | 542.3101287776238 | 592.9644976333208 | 461.3325033914183 | 634.9094736449964 | 471.66749330299956 | 502.3799648907136 | 738.1931240949758 |
| SE_153_5_11 | SE_153 | SE_153_5 | GCTGTTTCAGAGCAGAGCAC | 390.15966300778763 | 401.75577882282624 | 396.9857987790867 | 476.27890718130254 | 401.96532032444514 | 315.33687999210076 | 510.1001304504429 | 348.8354380152323 | 641.1253705897726 | 389.28970853756067 | 458.9499211968903 | 215.88666836739858 |
| SE_153_5_21 | SE_153 | SE_153_5 | GGAAGTGTCATGGAGAGCTG | 830.173060733237 | 1097.332948128018 | 992.4644969477167 | 998.1036114974291 | 1132.7149395950064 | 1233.625596452614 | 774.8356411905462 | 909.4300579150499 | 1216.5398306204827 | 1174.430188116124 | 1035.2783388635735 | 1609.8645646536659 |
| SE_153_5_5 | SE_153 | SE_153_5 | GGAACCCTTTATTACCAGGG | 18.42420630870108 | 9.993924846338961 | 13.825376076883616 | 18.218318853929606 | 4.242378050917627 | 3.465240439473635 | 12.913927353175769 | 34.97807915057884 | 87.9105425046918 | 86.02281948957334 | 3.521354894093787 | 2.321362025455899 |
| SE_153_5_6 | SE_153 | SE_153_5 | TGGCTGGACTGGCTGAGTGG | 102.95879996038839 | 90.94471610168455 | 99.74021312608895 | 100.20075369661284 | 90.15053358199957 | 218.310147686839 | 100.08293698711222 | 87.9178746217252 | 134.08577695160065 | 188.8127987101652 | 57.515463270198524 | 87.05107595459621 |
| SE_153_5_7 | SE_153 | SE_153_5 | AAAGCCCCCCTGGTAATAAA | 13.005322100259587 | 34.97873696218637 | 43.451181955919935 | 18.218318853929606 | 13.787728665482287 | 25.12299318618385 | 20.4470516425283 | 21.743130282792254 | 15.983735000853056 | 29.160277793075707 | 2.347569929395858 | 27.856344305470785 |
| SE_153_6_10 | SE_153 | SE_153_6 | GATGAAATGTGGTCACTTGA | 880.0267954508987 | 937.4301505865947 | 946.0507344038932 | 922.6277191025779 | 916.3536589982074 | 1040.4384419519588 | 1105.2169493092929 | 1135.3695421579782 | 1033.6148633884977 | 992.9074588542278 | 1333.4197198968475 | 1114.2537722188315 |
| SE_153_6_3 | SE_153 | SE_153_6 | CTTCCAGTGCAATATCAGAA | 166.90163361999802 | 244.85115873530455 | 221.20601723013786 | 231.63291114281927 | 224.84603669863424 | 272.02137449868036 | 175.41417988063753 | 291.168875091305 | 251.29983362452305 | 406.7858752134061 | 177.2415296693873 | 119.5501443109788 |
| SE_153_6_4 | SE_153 | SE_153_6 | TGCAGAGGCTTTTAACACAG | 910.3725470181711 | 1153.298927267516 | 1244.2838469195256 | 1037.142866184421 | 1183.623476206018 | 1068.1603654677479 | 856.6238477606594 | 1285.6807471564116 | 1125.9653322823153 | 873.3503199026175 | 960.156101122906 | 1483.3503342663194 |
| SE_153_6_6 | SE_153 | SE_153_6 | TCCTGCCTCTTGGCACTGAA | 327.30060618986624 | 339.7934447755247 | 380.1978421142995 | 297.9996441107057 | 316.0571647933632 | 485.9999716361773 | 292.7156866719841 | 398.9391730147101 | 357.8580669635434 | 279.93866681352677 | 444.8645016205151 | 746.3178911840715 |
| SE_153_6_7 | SE_153 | SE_153_6 | GGGTGGTGATGCTTGTCACA | 289.36841673077583 | 343.7910147140603 | 342.6718213341868 | 339.6415157768305 | 291.6634910005869 | 288.4812665861801 | 301.3249715741013 | 219.3220098090349 | 372.9538166865713 | 218.7020834480678 | 153.7658303754287 | 82.4083519036844 |
| SE_154_1_0 | SE_154 | SE_154_1 | TTTTCATTAAATATTCTCAA | 0.0 | 0.0 | 0.0 | 0.0 | 0.0 | 0.0 | 0.0 | 0.0 | 0.0 | 0.0 | 1.173784964697929 | 0.0 |
| SE_154_1_13 | SE_154 | SE_154_1 | GTTCTCGGCTGAGAGAAACT | 8.670214733506391 | 28.982382054382988 | 29.62580587903632 | 10.410467916531204 | 23.33307928004695 | 32.919784174999535 | 11.837766740411123 | 6.617474433893295 | 13.319779167377547 | 59.04956253097831 | 0.0 | 52.23064557275772 |
| SE_154_1_17 | SE_154 | SE_154_1 | TGCCCCTCCGGTGAAAACAG | 54.188842084414944 | 89.94532361705066 | 92.82752508764715 | 89.79028578008163 | 110.3018293238583 | 58.04277736118338 | 49.50338818717378 | 84.13646065950047 | 36.40739639083196 | 91.85487504818848 | 77.46980767006332 | 40.62383544547823 |
| SE_154_1_19 | SE_154 | SE_154_1 | CTTCCTCTGTTTTCACCGGA | 59.60772629285644 | 46.97144677779312 | 69.12688038441809 | 68.96934994701923 | 104.99885676021127 | 101.35828285460381 | 32.28481838293943 | 125.7320142439726 | 121.65398306204825 | 53.21750697236316 | 89.2076573170426 | 54.55200759821362 |
| SE_154_1_4 | SE_154 | SE_154_1 | AGTCTCCTATCCAGGAAACA | 29.26197472558407 | 63.961119016569356 | 27.650752153767232 | 48.14841411395682 | 53.02972563647034 | 34.65240439473635 | 45.19874573611519 | 18.9070698111237 | 61.27098416993672 | 27.702263903421922 | 4.695139858791716 | 152.0492126673614 |
| SE_154_3_11 | SE_154 | SE_154_3 | GCCAGGGAGAAACAACCACG | 4994.043686499681 | 5722.52136701369 | 5550.888494868772 | 4834.361038739177 | 5429.183310661833 | 5615.422132167026 | 4976.16667342373 | 6213.808493425804 | 5379.414813064878 | 5186.1554054985145 | 5444.014666268995 | 5304.312228166729 |
| SE_154_3_13 | SE_154 | SE_154_3 | GGTTTATTTAATTTTCCACG | 3.2513305250648967 | 2.9981774539016888 | 4.93763431317272 | 1.3013084895664004 | 2.1211890254588135 | 0.8663101098684087 | 1.0761606127646475 | 0.0 | 0.0 | 5.832055558615141 | 0.0 | 0.0 |
| SE_154_3_14 | SE_154 | SE_154_3 | TGTTGCACAACCATGCCAGG | 2185.977889685299 | 1907.840253166108 | 1990.8541550712407 | 2262.9754633559705 | 2226.1878822190247 | 1681.5079232545813 | 2268.546571707877 | 1899.2151625273757 | 2020.1665070522613 | 2882.4934598455334 | 1484.8379803428802 | 2162.34872671217 |
| SE_154_3_4 | SE_154 | SE_154_3 | AGAACAGAAATAGTAGCTGA | 244.93356622155557 | 344.7904071986942 | 309.0959080046123 | 268.0695488506785 | 319.23894833155146 | 320.5347406513112 | 189.40426784657797 | 332.76442867577714 | 298.36305334925703 | 209.22499316531818 | 197.19587406925208 | 306.41978736017865 |
| SE_154_3_5 | SE_154 | SE_154_3 | TAGAACAGAAATAGTAGCTG | 359.81391144051526 | 517.6853070403582 | 500.6761193557138 | 464.56713077520493 | 505.903582571927 | 598.6202859190704 | 316.39122015280634 | 518.0537128247894 | 429.78487446738217 | 493.53770164780633 | 530.5508040434639 | 471.23649116754746 |
| SE_155_1_15 | SE_155 | SE_155_1 | GAGCAGCTACTCATGAACTG | 2015.8249255402359 | 2007.7795016294974 | 2027.392648988719 | 1794.5044071120662 | 2013.008385160414 | 2194.3635082966794 | 1843.463129665841 | 1875.581325263471 | 1995.3029192731565 | 1189.010327012662 | 2041.2120536096986 | 2036.9951773375512 |
| SE_155_1_16 | SE_155 | SE_155_1 | GATATGGTAGTGGACAGCAG | 701.2036165723293 | 638.6117976810597 | 659.6679442398754 | 836.7413587911955 | 585.4481710266325 | 760.6202764644629 | 670.4480617523753 | 599.3541130126213 | 640.2373853119474 | 564.2513752960149 | 673.7525697366112 | 678.9983924458504 |
| SE_155_1_20 | SE_155 | SE_155_1 | AACCCCATAACAACTCCTTG | 49.85373471766175 | 78.9520062860778 | 87.88989077447442 | 66.36673296788642 | 84.84756101835254 | 35.51871450460476 | 50.57954879993843 | 114.38777235729839 | 79.03068972644012 | 29.160277793075707 | 65.73195802308402 | 59.19473164912542 |
| SE_155_1_24 | SE_155 | SE_155_1 | ATGGTTCTTTACCATCCCCA | 0.0 | 3.9975699385355847 | 4.93763431317272 | 9.109159426964803 | 0.0 | 0.0 | 3.228481838293942 | 0.0 | 0.0 | 29.160277793075707 | 0.0 | 1.1606810127279494 |
| SE_155_1_6 | SE_155 | SE_155_1 | GATGTAATTAAGTTGCAGGG | 171.23674098675122 | 274.8329332743215 | 308.10838114197776 | 206.90804984105768 | 221.664253160446 | 168.93047142433971 | 177.56650110616684 | 253.35473546905757 | 348.09022890746655 | 257.3394515238931 | 350.9617044446808 | 396.95290635295873 |
| SE_155_2_10 | SE_155 | SE_155_2 | AATGTTCTAGGTCTGCAGGA | 14.089098941947885 | 4.996962423169481 | 2.962580587903632 | 7.807850937398403 | 5.3029725636470335 | 3.465240439473635 | 24.75169409358689 | 1.89070698111237 | 55.94307250298569 | 5.832055558615141 | 0.0 | 16.249534178191293 |
| SE_155_2_11 | SE_155 | SE_155_2 | GGGCTGTACGATGAACCAGG | 312.12773040623006 | 356.78311701430096 | 281.445155850845 | 260.2616979132801 | 275.75457330964576 | 199.251325269734 | 294.8680078975134 | 319.5294798079905 | 302.8029797383829 | 109.3510417240339 | 383.8276834562228 | 174.10215190919243 |
| SE_155_2_16 | SE_155 | SE_155_2 | GAAAGGTAGAGAGAACCAAG | 373.90301038246315 | 442.730870692816 | 415.74880916914304 | 352.6546006724945 | 356.3597562770807 | 334.3957024092058 | 348.67603853574576 | 428.2451312219518 | 549.6628869737801 | 352.11035435138916 | 610.3681816429231 | 407.39903546751026 |
| SE_155_2_2 | SE_155 | SE_155_2 | CATGTCCACGTGTTTGTGAA | 9388.758779545733 | 8379.905983655219 | 8082.907370663743 | 8817.666325301929 | 8561.118906751772 | 7842.705424638704 | 8863.258806729636 | 8542.214140665688 | 8005.187279593905 | 8339.839448819652 | 8941.893861068824 | 7713.886010589952 |
| SE_155_2_25 | SE_155 | SE_155_2 | CCAAAGGGAAAAGTAAACCC | 955.8911743690796 | 1221.2576162226212 | 1248.2339543700637 | 1005.9114624348275 | 946.0503053546308 | 1248.352868320377 | 891.0609873691282 | 976.5501557445391 | 1350.6256075720833 | 1121.2126811437608 | 1002.4123598520314 | 826.4048810623 |
| SE_155_3_10 | SE_155 | SE_155_3 | TCATCCAGAGAACATCTATG | 849.6810438836263 | 862.4757142390524 | 843.3479406899006 | 908.3133257173475 | 947.1108998673602 | 758.0213461348576 | 908.2795571733625 | 1091.8832815923936 | 1117.0854795040636 | 653.1902225648959 | 1139.7452007216891 | 582.6618683894307 |
| SE_155_3_17 | SE_155 | SE_155_3 | AAGACCATACTGGTACAAAG | 369.56790301570993 | 337.7946598062569 | 342.6718213341868 | 424.22656759864657 | 309.69359771698674 | 312.73794966249557 | 336.83827179533466 | 425.40907075028326 | 364.9619491861448 | 457.8163613512886 | 576.3284176666832 | 500.2535164857462 |
| SE_155_3_18 | SE_155 | SE_155_3 | ACTCCTGGTTTTGCTCCAAG | 42.26729682584366 | 77.9526138014439 | 74.06451469759081 | 28.62878677046081 | 47.72675307282331 | 76.23528966841997 | 51.655709412703075 | 24.57919075446081 | 85.2465866712163 | 26.244250013768134 | 15.259204541073078 | 78.92630886550056 |
| SE_155_3_4 | SE_155 | SE_155_3 | CCTTCCATCCCCATGCAGAA | 769.4815575986922 | 937.4301505865947 | 957.9010567555077 | 966.8722077478355 | 924.8384151000427 | 1236.2245267822193 | 755.4647501607825 | 934.954602160067 | 1123.3013764488398 | 799.7206184751012 | 1049.3637584399485 | 1902.356179861109 |
| SE_155_3_7 | SE_155 | SE_155_3 | ACAGGGCCTTGTAAATACCA | 2035.3329086906253 | 2301.600892111863 | 2238.7233975925114 | 2143.2550823158617 | 2363.004574361118 | 2255.8715260973363 | 2014.57266709542 | 2033.455358186354 | 2246.6027528976797 | 2022.9942718946272 | 1885.098653304874 | 2579.0332102815037 |
| SE_155_4_0 | SE_155 | SE_155_4 | AGACTTAAGAGGGAACAAGA | 235.17957464636086 | 294.8207829669994 | 256.75698428498146 | 295.3970271315729 | 241.81554890230473 | 136.01068724934018 | 187.25194662104866 | 402.72058697693484 | 227.32423112324346 | 320.7630557238328 | 562.242998090308 | 157.85261773100112 |
| SE_155_4_19 | SE_155 | SE_155_4 | CAGAAGGGACAAATAACAAA | 1153.13855955635 | 1501.087511920112 | 1558.3173892373104 | 1315.6228829516308 | 1348.015625679076 | 1239.6897672216928 | 1300.0020202196943 | 1488.9317476259914 | 1524.6707220258165 | 1793.357084274156 | 1510.6612495662346 | 1344.0686127389654 |
| SE_155_4_3 | SE_155 | SE_155_4 | ATACAACGGAAAGTTAACGA | 287.2008630473992 | 288.824428059196 | 314.033542317785 | 281.0826337463425 | 278.93635684783396 | 323.99998109078484 | 279.80175931880837 | 435.8079591464013 | 182.0369819541598 | 210.68300705497197 | 291.0986712450864 | 182.22691899828806 |
| SE_155_4_5 | SE_155 | SE_155_4 | ATTTAACCTTCTGTCTACCA | 19.50798315038938 | 27.982989569749094 | 33.5759133295745 | 19.519627343496005 | 18.030106716399914 | 0.0 | 12.913927353175769 | 15.12565584889896 | 22.199631945629246 | 16.038152786191638 | 9.390279717583432 | 27.856344305470785 |
| SE_155_4_9 | SE_155 | SE_155_4 | TTAGGAATCAAGGAGGCTGA | 373.90301038246315 | 381.76792913014833 | 392.048164465914 | 378.6807704638225 | 284.239329411481 | 269.4224441690751 | 337.9144324080993 | 371.5239217885807 | 341.87433196269035 | 408.97289604788676 | 516.4653844670888 | 210.08326330375886 |
| SE_156_1_12 | SE_156 | SE_156_1 | GCTGTGCGTTTGGTCGAGGA | 2696.436782120488 | 2479.492754376696 | 2607.0709173551963 | 2853.769517619116 | 2740.576220892787 | 2080.0105737940494 | 2836.759375247611 | 2629.0280572367506 | 2473.0389987430976 | 2604.7418138664875 | 3034.2341337441467 | 2669.566329274284 |
| SE_156_1_19 | SE_156 | SE_156_1 | CTGCTCCATATGTATCCCTC | 164.73407993662144 | 175.89307729556572 | 174.7922546863143 | 126.22692348794084 | 161.21036593486983 | 90.09625142631451 | 137.74855843387488 | 79.40969320671954 | 148.29354139680336 | 121.74415978609107 | 36.3873339056358 | 41.78451645820618 |
| SE_156_1_2 | SE_156 | SE_156_1 | GTTGGTGCTTCAAAGATGAG | 209.16893044584168 | 216.86816916555549 | 257.744511147616 | 193.89496494539367 | 160.14977142214042 | 282.41709581710126 | 185.09962539551935 | 104.93423745173654 | 244.1959514019217 | 131.22125006884067 | 68.07952795247988 | 717.3008658658728 |
| SE_156_1_33 | SE_156 | SE_156_1 | AGTCAACCCTGGTTATCATG | 356.5625809154503 | 394.760031430389 | 393.0356913285485 | 389.09123838035373 | 429.5407776554097 | 589.0908747105179 | 416.4741571399186 | 312.91200537409725 | 362.29799335266927 | 523.4269863857089 | 556.3740732668184 | 692.9265645985859 |
| SE_156_1_34 | SE_156 | SE_156_1 | CTGTCACCCCATGATAACCA | 455.18627350908554 | 396.7588163996568 | 453.2748299492557 | 473.67629020216975 | 463.47980206275076 | 365.5828663644685 | 446.6066542973287 | 500.09199650422187 | 409.36121307740325 | 547.4842155649964 | 447.21207154991095 | 285.52752913107554 |
| SE_156_2_12 | SE_156 | SE_156_2 | GGAGCTATGAATATGCATGG | 505.0400082267473 | 551.6646515179107 | 526.351817784212 | 514.0168533787282 | 552.5697411320209 | 669.6577149282799 | 615.5638705013783 | 560.5946198998178 | 554.9907986407311 | 545.2971947305157 | 433.1266519735358 | 459.629681040268 |

| | | | | | | | | | | | | | | | |
|---|---|---|---|---|---|---|---|---|---|---|---|---|---|---|---|
| SE_156_2_14 | SE_156 | SE_156_2 | GTCTGTTTCATTTTGCCTGG | 52.02128840103835 | 88.94593113241676 | 68.13935352178353 | 49.44972260352321 | 93.3323171201878 | 62.37432791052543 | 54.88419125099702 | 26.46989773557318 | 66.59889583688773 | 25.515243068941242 | 28.170839152750297 | 81.24767089095646 |
| SE_156_2_18 | SE_156 | SE_156_2 | GGCGAAACCGTGATAAAGGT | 2431.9952327485425 | 2429.5231301450017 | 2611.021024805734 | 2490.7044490300905 | 2527.396723834176 | 2401.411624555229 | 2311.5929962184628 | 2532.6020012000195 | 2234.1709590081273 | 2469.875529073512 | 2548.287158359204 | 2394.4849292577596 |
| SE_156_2_6 | SE_156 | SE_156_2 | GCTCTGGTTTCTTTACCTGC | 89.95347786012881 | 81.9501837399795 | 112.57806234033802 | 126.22692348794084 | 76.36280491651729 | 115.21924461249836 | 146.35784333599204 | 62.393330376708214 | 82.5826308377408 | 364.50347241344633 | 17.606774470468935 | 33.659749369110536 |
| SE_156_2_7 | SE_156 | SE_156_2 | GTAAATTTCTATATTTTAAT | 0.0 | 0.0 | 1.975053725269088 | 0.0 | 6.363567076376441 | 0.0 | 0.0 | 0.0 | 22.199631945629246 | 0.0 | 14.085419576375148 | 0.0 |
| SE_157_1_2 | SE_157 | SE_157_1 | CTGCAGTGTTCTCAGAGAGA | 23.843090517142578 | 13.991494784874547 | 17.77548352742179 | 14.314393385230405 | 4.242378050917627 | 114.35293450262995 | 53.808030638232374 | 21.743130282792254 | 90.57449833816732 | 0.7290069448268927 | 8.216494752885502 | 11.606810127279495 |
| SE_157_1_29 | SE_157 | SE_157_1 | AATTGCCATGGGGCTTCACG | 30.34575156727237 | 38.97630690072195 | 33.5759133295745 | 18.218318853929606 | 28.63605184369398 | 63.24063802039384 | 22.599372868057596 | 5.67212094333711 | 55.94307250298569 | 22.599215289633673 | 7.042709788187574 | 44.10587848366208 |
| SE_157_1_3 | SE_157 | SE_157_1 | CTAGGAAGCAGCAGAGGGCA | 1685.2729888253048 | 1718.9550735703015 | 1772.6107184290065 | 1752.8625354459414 | 1760.5868911308153 | 1722.2244984183965 | 1750.9133169680815 | 1715.8165853594758 | 2257.2585762315816 | 2148.3834664048527 | 1955.5257511867499 | 1944.1406963193153 |
| SE_157_1_4 | SE_157 | SE_157_1 | TGACCTGCTTCATGCACACG | 9014.85576916327 | 7180.635002094544 | 7014.403305293166 | 7794.837852502738 | 6927.803357148485 | 5753.165439636103 | 8441.403846525895 | 7534.467319732795 | 6064.051462268085 | 7069.180343986378 | 6550.893887979142 | 5810.369149716115 |
| SE_157_1_43 | SE_157 | SE_157_1 | GGAGGAGCTTCATAGCTCAA | 1386.1505805193342 | 1534.0674639130307 | 1529.6791102209088 | 1323.4307338890292 | 1595.1341471450278 | 1524.7057933683993 | 1228.9754197772274 | 1467.1886173431992 | 1831.0256428755001 | 1128.5027505920298 | 1432.0176569314733 | 1316.2122684334947 |
| SE_157_2_2 | SE_157 | SE_157_2 | GCAGTGGCTGGGATAACAAG | 4180.127278391769 | 4316.376141133797 | 4395.482065586356 | 4566.291489888499 | 4348.437502190568 | 4570.6521396657245 | 4155.056125884304 | 4904.493909005488 | 4496.757446906659 | 4812.174842802318 | 4432.21202669938 | 3902.209564791366 |
| SE_157_2_23 | SE_157 | SE_157_2 | CTAGGGGATGACGGACTGGT | 5441.643522116949 | 4828.065093266352 | 4707.540554178871 | 5211.740500713434 | 4975.248859213647 | 4894.652120756509 | 5703.651247652631 | 5169.192886361219 | 5506.396707793878 | 6101.788128201091 | 4239.71129248892 | 4814.50484079535 |
| SE_157_2_27 | SE_157 | SE_157_2 | AGGACAAGACAAGATTCTAG | 2271.5962601786746 | 2261.625192726507 | 2202.184903675033 | 2273.3859312725017 | 2503.0030500414 | 2406.6094852144392 | 2546.196009801156 | 2217.7992888448102 | 2309.6497076232667 | 2205.9750150461773 | 2082.294527374126 | 2826.258265992557 |
| SE_157_2_4 | SE_157 | SE_157_2 | TTCCTAGTGTGGGGTCAGCG | 34.680858934025565 | 31.980559508284678 | 46.41376254382357 | 19.519627343496005 | 13.787728665482287 | 0.8663101098684087 | 5.380803063823238 | 0.0 | 54.167101947335354 | 6.561062503442034 | 1.173784964697929 | 6.964086076367696 |
| SE_157_2_42 | SE_157 | SE_157_2 | GCAGTGCCCAGCACATAGTT | 265.5253262136332 | 215.86877668092157 | 211.33074860379241 | 235.53683661151848 | 235.4519818259283 | 93.56149186578814 | 291.63952605921946 | 229.72089820515296 | 88.79852778251698 | 78.00374309647752 | 164.32989505771008 | 56.873369623669525 |
| SE_157_3_0 | SE_157 | SE_157_3 | AGAGAGAGCAACTCGCCCTG | 3920.0208363865772 | 4257.4119845403975 | 4160.450672279334 | 4285.208856142157 | 4117.227898415557 | 3949.5077908900753 | 4316.480217799001 | 4429.926456746283 | 4577.56410718875 | 4627.736085761115 | 4509.681834369443 | 4795.933944591887 |
| SE_157_3_1 | SE_157 | SE_157_3 | AGAGCAACTCGCCCTGTGGA | 2589.142874793346 | 3037.1537608024105 | 3045.5328443649337 | 2818.6341884008234 | 2839.211510576622 | 3343.0907139821893 | 2866.891872405021 | 3133.8468211937534 | 2885.9521529318017 | 3565.572967148332 | 3058.883618002803 | 2821.615541941645 |
| SE_157_3_2 | SE_157 | SE_157_3 | CACTGTATGCTCCTTCCACA | 2371.303729613998 | 2140.6987020858055 | 2044.180605653506 | 2465.9795877283286 | 2438.3067847649063 | 2532.2244511453587 | 2306.2121931546394 | 2512.74957789834 | 1657.868513699592 | 2102.4560288807584 | 2522.4638891358495 | 2165.8307697503537 |
| SE_157_3_4 | SE_157 | SE_157_3 | CTGTGGAAGGAGCATACAGT | 865.9376965089508 | 595.6379208418022 | 626.0920309103009 | 761.2654663963442 | 854.8391772599018 | 672.2566452578852 | 801.7396565096624 | 736.4303691432681 | 537.2310930842277 | 893.7625143577704 | 814.6067655003627 | 632.5711519367325 |
| SE_157_3_8 | SE_157 | SE_157_3 | ATACAGTGGGAGATGGGGAC | 339.22215144843756 | 369.7752193145416 | 392.048164465914 | 344.8467497350961 | 377.5716465316688 | 419.29409317630984 | 404.63639039950743 | 441.4800800897384 | 364.0739639083196 | 312.0149723859101 | 291.0986712450864 | 600.0720835803498 |
| SE_157_4_2 | SE_157 | SE_157_4 | GGACCTGGACAAGAGCCAAA | 487.6995787597345 | 578.6482486030259 | 565.8528922895937 | 638.9424683771026 | 634.2355186121853 | 439.2192257032832 | 651.0771707226118 | 635.2775456537563 | 745.0196480953174 | 696.2016323096825 | 577.5022026313811 | 450.3442329384444 |
| SE_157_4_26 | SE_157 | SE_157_4 | ATGTTGCTTGGATCTCCCCA | 749.9735744483029 | 770.5316056527339 | 814.7096616734988 | 783.387710718973 | 869.6875004381135 | 719.0373911907792 | 691.9712740076683 | 658.911382917661 | 608.2699153102413 | 1097.1554519644735 | 1069.3181028398133 | 832.2082861259397 |
| SE_157_4_29 | SE_157 | SE_157_4 | GTAGCCCCATACAAGCAGGT | 156.06386520311503 | 147.91008772581662 | 174.7922546863143 | 182.18318853929605 | 137.87728665482288 | 77.10159977828837 | 160.34793130193248 | 224.04877726181584 | 92.35046889381766 | 65.61062503442034 | 228.88806811609615 | 95.17584304369186 |
| SE_157_4_3 | SE_157 | SE_157_4 | AAGCACTTCCTCTCAGACCA | 343.5572588151908 | 417.7460585769686 | 438.46192700973756 | 366.9689940577249 | 473.0251526773154 | 209.64704658815492 | 278.7255987060437 | 311.96665188354103 | 391.6015075208999 | 255.88143763423932 | 318.0957254331388 | 323.8300025510979 |
| SE_157_4_6 | SE_157 | SE_157_4 | GGTATACCCTCACAGTCACA | 242.76601253817896 | 204.87545934994873 | 246.881715658636 | 269.37085734024487 | 217.42187510952837 | 267.6898239493383 | 267.9639925783972 | 166.38221433788857 | 177.59705556503397 | 102.78997922059186 | 165.503680022408 | 276.24208102925195 |
| SE_157_5_0 | SE_157 | SE_157_5 | GCCAAAAAGACTATGCTAGA | 544.055974527526 | 832.4939397000355 | 719.9070828605826 | 696.2000419180242 | 772.1128052670081 | 811.732572946699 | 583.2790521184389 | 802.6051134822011 | 821.3863819882821 | 1255.3499589919093 | 788.7834962770083 | 636.0531949749163 |
| SE_157_5_1 | SE_157 | SE_157_5 | AAATTCCTGAGAGACTAGAG | 6.5026610501297935 | 6.9957473924372735 | 5.925161175807264 | 6.506542447832002 | 0.0 | 0.0 | 0.0 | 0.0 | 1.7759705556503396 | 0.0 | 0.0 | 1.1606810127279494 |
| SE_157_5_22 | SE_157 | SE_157_5 | GGTGCCCGAGCCCCGGTGCG | 230.84446727960767 | 310.8110627211417 | 346.62192878472496 | 288.8904846837409 | 293.7846800260457 | 518.9197558111769 | 167.88105559128502 | 233.5023121673777 | 243.3079661240965 | 278.480652923873 | 322.79086529193046 | 492.12874939665056 |
| SE_157_5_31 | SE_157 | SE_157_5 | CAAATGGGTCTCCTGGTGCG | 875.6916880841455 | 1122.3177602438655 | 1190.9573963372602 | 857.5622946242579 | 904.687119358184 | 869.7753503078824 | 859.8523295989534 | 945.353490556185 | 910.184909770799 | 990.7204380197471 | 843.951389617811 | 791.5844506804615 |
| SE_157_5_40 | SE_157 | SE_157_5 | TGGGGCGACAGATAGCACAT | 2052.673338157638 | 1832.8858168185657 | 1705.4588917698575 | 1949.3601173704678 | 1941.9485528075438 | 1877.2940080848416 | 1960.7646364571876 | 2004.149399979112 | 1674.7402339782702 | 1853.1356537499612 | 2014.2149994216463 | 1938.3372912556756 |
| SE_157_6_0 | SE_157 | SE_157_6 | CATCAGCCTTCACTTCCGGA | 787.9057639073933 | 789.520062860778 | 785.0838557944625 | 809.413880510301 | 847.415015670796 | 835.122945913146 | 654.3056525609056 | 604.0808804654022 | 655.3331350349753 | 716.6138267648355 | 572.8070627725893 | 436.41606078570896 |
| SE_157_6_1 | SE_157 | SE_157_6 | CATCCGGAAGTGAAGGCTGA | 1821.8288708780306 | 1833.8852093031994 | 1811.1242660717537 | 1794.5044071120662 | 2059.6745437205077 | 1612.2031144651087 | 1616.3932403725005 | 1588.1938641343909 | 1934.919920381045 | 1735.7655356328314 | 1460.1884960842237 | 2575.5511672433 |
| SE_157_6_2 | SE_157 | SE_157_6 | ATCCGGAAGTGAAGGCTGAT | 405.3325387914238 | 335.7958748369891 | 400.9359062296249 | 379.9820789533889 | 407.2682928880922 | 517.18713559144 | 291.63952605921946 | 250.518674997389 | 450.2085358573611 | 252.23640291010486 | 248.8424125159609 | 735.87176206952 |
| SE_157_6_56 | SE_157 | SE_157_6 | CTGGGCCTGGAGAGGATGGC | 455.18627350908554 | 451.72540305452105 | 381.185368976934 | 404.7069402551505 | 556.8121191829385 | 331.79677207960054 | 335.76211118257004 | 487.80240112699147 | 382.7216547426482 | 317.1180209969833 | 564.5905680197038 | 290.17025318198733 |
| SE_157_6_57 | SE_157 | SE_157_6 | GTCACTGGGCCTGGAGAGGA | 1207.327401640765 | 1491.093587073773 | 1490.178035715527 | 1259.6666179002757 | 1257.8650920970765 | 641.0694813026224 | 885.6801843053049 | 676.8730992382285 | 1395.0248714633417 | 394.3927571513489 | 334.5287149389098 | 219.36871140558245 |
| SE_158_1_12 | SE_158 | SE_158_1 | GCTGAACCGTATTTGGGTGG | 3756.370533291644 | 3790.6956942163683 | 3952.082504263445 | 3484.9041350588204 | 3603.9001542545243 | 4193.807241872967 | 3680.4692956550944 | 4180.35313523945 | 3831.6564738156076 | 3763.862856141247 | 4441.602306416963 | 4077.472397713286 |
| SE_158_1_15 | SE_158 | SE_158_1 | CCAAACGAGCTGCAAAATGA | 1463.0987362792034 | 1374.1646663716072 | 1439.8141657211652 | 1275.2823197750724 | 1381.954650086417 | 1428.545371173006 | 1457.1214696833326 | 1663.8221433788856 | 1197.0041545083288 | 1440.51772297794 | 2241.9292825730445 | 1157.1989696897656 |
| SE_158_1_17 | SE_158 | SE_158_1 | CTGAGCGTATTCAATAAGCA | 777.0679954905103 | 846.4854344849101 | 797.9217050087116 | 840.6452842598947 | 820.9001528525608 | 725.9678720697265 | 751.1601077097239 | 994.5118720651067 | 704.1723253153597 | 392.93474326169513 | 789.9572812417063 | 766.0494684004466 |
| SE_158_1_8 | SE_158 | SE_158_1 | AAGTTACCAGGAGAATTCAG | 68.27794102636283 | 101.9380334326574 | 107.6404280271653 | 61.16149900962082 | 59.39329271284678 | 59.7753975809202 | 55.960351863761666 | 141.80302358342774 | 146.517570841153 | 25.515243068941242 | 203.06479889274172 | 30.177706330926686 |
| SE_158_1_9 | SE_158 | SE_158_1 | ACTGCTATTCCAGCAAAACA | 372.8192335407748 | 442.730870692816 | 426.61160465812304 | 422.9252591090801 | 422.1166160663039 | 207.0481162585497 | 284.1064017698669 | 239.1744331107148 | 239.75602501279585 | 521.9689724960551 | 286.4035313862947 | 372.57860508567177 |
| SE_158_2_11 | SE_158 | SE_158_2 | ACAGGAGCCTGCAGAACAAG | 617.7527997623304 | 127.92223803313871 | 228.11870526857967 | 439.84226947344337 | 126.21074701479941 | 907.8929951420923 | 751.1601077097239 | 229.72089820515296 | 916.4008067155752 | 37.908361130998415 | 219.49778839851274 | 280.88480508016374 |
| SE_158_2_12 | SE_158 | SE_158_2 | TTGGCAGTCAGCTATGTAGG | 549.4748587359676 | 474.7114302011007 | 475.00042092721566 | 537.4404061909233 | 394.5411587353393 | 447.88232680196734 | 572.5174459907925 | 664.583503860998 | 465.304285580389 | 1012.5906463645539 | 748.8748074772788 | 352.8470278692966 |
| SE_158_2_13 | SE_158 | SE_158_2 | CTTGGCAGTCAGCTATGTAG | 1599.6546183319292 | 1849.8754890573418 | 1912.8395329231118 | 1616.2251440414693 | 1857.1009917891913 | 1557.625577543399 | 1541.0619974789752 | 1657.2046689449924 | 1908.2803620462898 | 1463.1169382675735 | 1714.8998334236744 | 1783.9667165628582 |
| SE_158_2_18 | SE_158 | SE_158_2 | GAGCTGAGTTAGAGTGCCAA | 985.1531490946637 | 1048.3627163809572 | 1037.8907326289059 | 899.2041662903827 | 1056.3521346784892 | 963.3368421736706 | 851.2430446968361 | 1163.7301468746637 | 739.6917364283664 | 1046.124965826591 | 784.0883564182166 | 796.2271747313733 |
| SE_158_2_7 | SE_158 | SE_158_2 | CAAAACTAGGAACAGACGGG | 485.5320250763579 | 600.6348832649716 | 524.3767640589429 | 559.5626505135522 | 667.1139485067969 | 760.6202764644629 | 397.1032661101549 | 477.40351273087344 | 574.5264747528848 | 435.2171460616549 | 550.5051484433287 | 678.9983924458504 |
| SE_158_3_10 | SE_158 | SE_158_3 | GAGAGTAAATGGGTTACCAG | 580.9043871449283 | 743.5480085676188 | 850.2606287283425 | 631.1346174397042 | 646.9626527649381 | 837.7218762427513 | 569.2889641524985 | 700.506936502133 | 800.9627205983031 | 669.9573822959144 | 505.9013197848074 | 828.7262430877558 |
| SE_158_3_11 | SE_158 | SE_158_3 | GGTTACCAGAGGAAGAGCTA | 110.54523785220648 | 148.90948021045054 | 133.31612645566344 | 127.52823197750725 | 89.08993906927017 | 47.64705604276248 | 105.46374005093546 | 190.01605160179318 | 64.82292528123739 | 95.49990977232294 | 79.81737759945918 | 31.338387343654635 |
| SE_158_3_5 | SE_158 | SE_158_3 | CTTAAAGAGCTTTCACATTC | 251.43622727168534 | 273.83354078968756 | 245.89418879600146 | 239.44076208021767 | 346.814405662516 | 255.56148241118058 | 209.85131948910626 | 299.67705650631063 | 263.7316275140754 | 161.1105348067433 | 312.2268006096491 | 244.90369368559732 |
| SE_158_3_6 | SE_158 | SE_158_3 | TTTCACATTCAGGTCTCTAC | 33.59708209233727 | 28.982382054382988 | 41.47612823065085 | 22.12224432262881 | 40.30259148371746 | 53.71122681184134 | 38.74178205952731 | 78.46433971616335 | 12.431793889552377 | 5.832055558615141 | 139.68041079905356 | 155.53125570554522 |
| SE_158_3_7 | SE_158 | SE_158_3 | CATTTACTCTCAAACTATCG | 268.77665673869814 | 293.8213904823655 | 271.5698872244996 | 296.6983356211393 | 390.2987806844217 | 212.24597691776015 | 320.69586260386495 | 573.8295687676043 | 421.79300696695566 | 185.89677093085763 | 1112.7481465336368 | 160.17397975645702 |
| SE_158_4_15 | SE_158 | SE_158_4 | GTCCTCAAAGACTCTTACAA | 18.42420630870108 | 27.982989569749094 | 36.53849391747813 | 42.943180155691216 | 51.96913112374093 | 37.251334724341575 | 3.228481838293942 | 18.9070698111237 | 14.207764445202717 | 2.187020834480678 | 14.085419576375148 | 9.285448101823595 |
| SE_158_4_22 | SE_158 | SE_158_4 | TGTTCTTGAGGACTGACCAA | 123.55055995246607 | 125.92345306387092 | 91.8399982250126 | 135.33608291490566 | 161.21036593486983 | 42.44919538355203 | 109.76838250199404 | 115.33312584785457 | 140.30167389637683 | 14.580138896537854 | 140.85419576375148 | 135.79967848917008 |
| SE_158_4_29 | SE_158 | SE_158_4 | TGTAAACTAGAGGCCACACA | 1289.6944416090757 | 1456.1148501115867 | 1425.001262781647 | 1353.3608291490564 | 1490.1352903848165 | 1264.8127604078768 | 1287.0880928665183 | 1690.2920411144587 | 1567.2940153614247 | 1561.532875819204 | 1797.0647809525294 | 1361.4788279298846 |
| SE_158_4_3 | SE_158 | SE_158_4 | TTAGAAGTAGTATTTCTTGG | 95.37236206857031 | 106.9349958558269 | 99.74021312608895 | 130.13084895664005 | 101.81707322202305 | 103.09090307434064 | 154.96712823810924 | 193.79746556401793 | 134.9737622294258 | 238.3852709583939 | 82.1649475288504 | 106.78265317097134 |
| SE_158_4_32 | SE_158 | SE_158_4 | ACATAAATAAAGAATGGCTG | 1011.1637932951829 | 1420.1367206647665 | 1376.6124465125545 | 1137.3436198810339 | 1425.4390251083228 | 1450.2031239197163 | 978.2299970030646 | 1336.7298356464455 | 1347.9616517386078 | 1544.7657160881856 | 1935.571406786885 | 1196.662124122516 |
| SE_158_5_4 | SE_158 | SE_158_5 | AAGCTCTCCATGCTCATAGA | 100.7912462770118 | 94.94228604022014 | 128.37849214249073 | 72.87327541571842 | 85.90815553108195 | 173.26202197368175 | 78.55972473181926 | 151.2565584889896 | 96.7903952829435 | 55.404527806843845 | 177.2415296693873 | 194.9944101382955 |
| SE_158_5_5 | SE_158 | SE_158_5 | TGAGCATGGAGAGCTTAGCT | 2709.4421042207473 | 2808.292881821248 | 2735.449409497687 | 2680.695488506785 | 2877.3929130348806 | 2493.24049620128 | 2525.748958158628 | 2586.487150161722 | 2394.8962942944827 | 2389.684765142554 | 2347.5699293958583 | 2624.2997697778937 |
| SE_158_5_7 | SE_158 | SE_158_5 | CTTGGACAGCCAGCTGATGA | 74.78060207649263 | 119.92709815606754 | 130.3535458677598 | 70.27065843658562 | 118.78658542569356 | 53.71122681184134 | 79.63588534458391 | 161.65544688510764 | 47.95120500255917 | 124.66018756539864 | 116.20471150509498 | 117.2287822855229 |
| SE_158_5_8 | SE_158 | SE_158_5 | TCGATGTGTCCCTCATCAGC | 6203.538641823823 | 5705.531694774913 | 6127.604182647346 | 5738.770438987826 | 5623.272106491315 | 5631.015714144657 | 6015.737825354379 | 5445.236105603625 | 5779.008188086205 | 6155.005635173455 | 6218.71274296928 | 6030.898542134425 |
| SE_158_5_9 | SE_158 | SE_158_5 | TCGAATCAAGATACATGACA | 920.1265385933658 | 1104.3286955204553 | 1181.0821277109146 | 987.693143580898 | 1191.0476377951238 | 1062.096194698669 | 822.1867081521907 | 759.1188529166166 | 1292.018579235622 | 774.9343823509869 | 480.078050561453 | 1130.5033063970227 |
| SE_159_1_20 | SE_159 | SE_159_1 | GAGCCTGGAGTGAGGCGCAG | 2182.726559160234 | 2662.3815790646995 | 2601.145756179389 | 2354.0670576256184 | 2548.6086140887646 | 1910.2137922598413 | 1940.3175848146593 | 2093.9579815819498 | 2557.397600136489 | 1843.6585634672115 | 1809.9764155642065 | 2064.851521643022 |
| SE_159_1_23 | SE_159 | SE_159_1 | GCAGCAGCGGCAACAAGTGC | 28630.13282687979 | 23340.811478624644 | 23138.741918390002 | 25460.100598366625 | 22623.541551030976 | 19082.21279007144 | 26755.505154554667 | 19434.57705885405 | 22856.74105121987 | 14271.03995193125 | 16817.990974191925 | 15170.100836354299 |
| SE_159_1_24 | SE_159 | SE_159_1 | GCAGCGGCAACAAGTGCCGG | 799.8273091659646 | 989.3985597875572 | 992.4644969477167 | 897.9028578008163 | 1041.5038115002774 | 886.2352423953821 | 775.9118018033108 | 770.4630948032908 | 1212.0999042313567 | 839.0869934957534 | 642.0603756897672 | 591.9473164912542 |
| SE_159_1_41 | SE_159 | SE_159_1 | CGCAGCTCACGCTCCATGCG | 4034.9011816055367 | 3266.014639783573 | 3312.165097276261 | 3812.8338744295534 | 3160.571647933632 | 3029.4864542098253 | 4071.1155980886615 | 3687.823966659678 | 2957.8789604356407 | 3976.7328840306996 | 3399.2812577652026 | 3116.4285191745444 |
| SE_159_1_43 | SE_159 | SE_159_1 | GCATGGAGCGTGAGCTGCGG | 31565.000514171705 | 26694.772657056 | 26816.291954841043 | 29412.174481179783 | 27525.609388866294 | 26879.870088996988 | 32524.80219958594 | 30157.72170223286 | 26984.09662255126 | 29745.670369771702 | 31551.33985108033 | 29231.751305553407 |
| SE_159_2_0 | SE_159 | SE_159_2 | TTAATCGTCGTAGGGAGCCC | 109.46146101051819 | 156.9046200875217 | 141.2163413567398 | 81.98243484268323 | 133.63490860390525 | 140.34223779868222 | 86.0928490211718 | 119.11453981007931 | 118.99002722857276 | 43.74041668961356 | 63.384388093688166 | 129.99627342553035 |
| SE_159_2_13 | SE_159 | SE_159_2 | TGGAAATAGCACCAGCCTGG | 126.80189047753097 | 180.8900397187352 | 204.41806056535063 | 178.27926307059687 | 191.9676068040226 | 124.74865582105086 | 138.82471904663953 | 133.2948421684221 | 150.95749723027888 | 190.99981954464587 | 154.93961534012664 | 45.26655949639003 |
| SE_159_2_24 | SE_159 | SE_159_2 | TGCAAAGTGCCTCAACCGCA | 7299.2370287706935 | 6458.074235704237 | 6269.80805086672 | 6558.594787414659 | 6300.992000125406 | 6123.079856549913 | 6622.69241095364 | 6376.409293801468 | 5558.787839185563 | 5410.689544505197 | 5381.8040631400045 | 5267.170435759434 |
| SE_159_2_3 | SE_159 | SE_159_2 | GGCTAGGAAGACAACAGAGA | 414.0027535249302 | 424.7418059694059 | 459.199991125063 | 524.4273212952594 | 489.9946648809859 | 484.2673514164405 | 527.3187002546773 | 511.4362383908961 | 452.87249169083657 | 457.8163613512886 | 850.9940994059986 | 377.22132913658356 |
| SE_159_2_4 | SE_159 | SE_159_2 | AGGCTAGGAAGACAACAGAG | 1556.3035446643971 | 1820.893107002959 | 1825.9371690112719 | 1694.3036534154535 | 1687.405869752486 | 1687.5720940236602 | 1608.860116083148 | 1607.1009339455145 | 1755.5468942603607 | 1417.1895007434794 | 1309.9440206028887 | 1557.633919080908 |
| SE_159_3_19 | SE_159 | SE_159_3 | GGTCGCCGGGCTGAGCCGCG | 332.71949039830776 | 426.7405909386737 | 468.08773288877387 | 387.78992989078733 | 460.2980185245625 | 331.79677207960054 | 392.79862365909634 | 611.6437083898517 | 456.4244328021373 | 432.3011182823473 | 774.6980767006331 | 335.4368126783774 |
| SE_159_3_23 | SE_159 | SE_159_3 | CACGTCCCGGCCGCTCCGCG | 6381.278043860704 | 7907.193338423386 | 8150.059197322892 | 7222.262117093523 | 8100.820888227208 | 8150.245513641989 | 6804.563554510866 | 7829.417608786324 | 7901.2930020883605 | 8360.251643274805 | 8513.462348954079 | 7314.61742211538 |
| SE_159_3_35 | SE_159 | SE_159_3 | GGAGCAGCTCCAGCTCGGCG | 1800.1533340442645 | 2039.760061137782 | 2112.3199591752896 | 1781.4913222164023 | 1833.7679125091443 | 1833.9785025914214 | 1821.939917410548 | 2035.3460651674663 | 1823.0337753750737 | 2075.4827719221635 | 1770.067726764477 | 2183.240984941273 |
| SE_159_4_12 | SE_159 | SE_159_4 | CAGAAGTTAAAGAACTCAAA | 171.23674098675122 | 296.81956793626716 | 234.04386644438694 | 244.64599603848328 | 241.81554890230473 | 176.7272624131554 | 156.0432888508739 | 219.3220098090349 | 166.94123223113192 | 250.04938207562418 | 385.00146842092073 | 301.77706330926685 |
| SE_159_4_19 | SE_159 | SE_159_4 | CTTCCAAAAGGCCTTCGCAT | 176.65562519519273 | 179.89064723410132 | 190.592684488467 | 208.20935833062407 | 266.2092226950811 | 153.33688944670834 | 206.6228376508123 | 313.8573588646534 | 83.47061611556596 | 325.13709739279415 | 274.6656817393154 | 181.06623798556012 |
| SE_159_4_42 | SE_159 | SE_159_4 | TATTTCTGCAAATTTCCAGA | 5.4188842084414945 | 13.991494784874547 | 13.825376076883616 | 10.410467916531204 | 9.545350614564661 | 28.588233625657487 | 5.380803063823238 | 16.071009339455145 | 7.103882222601358 | 5.103048613788249 | 41.08247376442752 | 82.4083519036844 |
| SE_159_4_5 | SE_159 | SE_159_4 | TGTTGCTTACTCTACCCCAG | 407.5000924748004 | 520.6834844942599 | 494.75095817990655 | 551.7547995761538 | 432.7225611935979 | 600.3529061388073 | 434.7688875569176 | 449.0429080141879 | 474.18413835864067 | 719.5298545441431 | 577.5022026313811 | 618.642979783997 |
| SE_159_4_8 | SE_159 | SE_159_4 | TGTTAATATATGCTATCTGA | 204.8338230790885 | 308.8122777518739 | 299.22063937826687 | 197.79889041409285 | 260.90625013143404 | 343.92511361775826 | 203.39435581251837 | 132.3494886778659 | 335.6584350179142 | 381.27063214446486 | 145.5493356225432 | 306.41978736017865 |
| SE_16_1_36 | SE_16 | SE_16_1 | GAGTCAGAGGGAAACCGGGG | 270.9442104220747 | 371.7740042838094 | 378.22278838903037 | 262.8643148924129 | 316.0571647933632 | 329.1978417499953 | 237.8314954209871 | 387.59493112803585 | 430.67285974520735 | 449.7972849581928 | 314.574370539045 | 407.39903546751026 |
| SE_16_1_37 | SE_16 | SE_16_1 | CCTGAGTCAGAGGGAAACCG | 1593.1519572817995 | 1599.0279754142339 | 1573.1302921768286 | 1525.1335497718212 | 1568.6192843267925 | 1281.2726524953764 | 1412.9988845599821 | 1625.062650266082 | 1522.8947514701663 | 1245.1438617643328 | 1841.6686096110507 | 1619.1500127554896 |
| SE_16_1_40 | SE_16 | SE_16_1 | TTTCCCTCTGACTCAGGCTG | 18.42420630870108 | 20.98724217731182 | 19.75053725269088 | 13.013084895664004 | 32.87842989461161 | 0.8663101098684087 | 18.294730416999005 | 34.03272566002266 | 7.991867500426528 | 8.019076393095819 | 77.46980767006332 | 1.1606810127279494 |
| SE_16_1_49 | SE_16 | SE_16_1 | CTAGTCCGCAGGCCCCACAC | 1145.552121664532 | 1314.2011172935736 | 1235.3961051558147 | 1117.823992537538 | 1358.62157080637 | 1114.941111400642 | 1120.283197887998 | 1275.2818587602935 | 1372.8252395177126 | 1413.5444660193448 | 1216.0412234270545 | 1582.008220348195 |

| | | | | | | | | | | | | | | | |
|---|---|---|---|---|---|---|---|---|---|---|---|---|---|---|---|
| SE_16_1_55 | SE_16 | SE_16_1 | ACAGGAGATGAGATGAGGGC | 2032.0815781655604 | 2293.605752234792 | 2269.336730334182 | 1955.8666598183 | 1939.8273637820848 | 1933.6041652262884 | 2064.0760552825936 | 2246.1598935614957 | 2117.84488761303 | 1976.3378274257059 | 2540.0706636063182 | 1761.9137773210273 |
| SE_16_2_0 | SE_16 | SE_16_2 | AGGCCGAGGCCTCCGCATCC | 0.0 | 0.0 | 0.0 | 1.3013084895664004 | 0.0 | 0.0 | 1.0761606127646475 | 0.0 | 0.0 | 0.0 | 0.0 | 0.0 |
| SE_16_2_1 | SE_16 | SE_16_2 | TCACCAGGATGCGGAGGCCT | 1807.7397719360827 | 1984.7934744829179 | 2521.156080305991 | 1840.0502042468902 | 2060.735138233237 | 2099.9357063210227 | 1712.1715349085541 | 2360.547665918794 | 1982.871125383604 | 1873.5478482051142 | 2378.088338478004 | 1749.1462861810198 |
| SE_16_2_2 | SE_16 | SE_16_2 | GAGGCCTCCGCATCCTGGTG | 415.0865303666185 | 441.7314782081821 | 407.8485942680667 | 383.88600442208815 | 457.1162349863743 | 291.08019691578534 | 372.351572016568 | 534.1247221642445 | 506.15160836034676 | 464.37742385473064 | 551.6789334080266 | 243.74301267286938 |
| SE_16_2_27 | SE_16 | SE_16_2 | GCCAGTGTCTCCACCCAGAG | 693.6171786805113 | 895.455666231971 | 918.399982250126 | 879.6845389468867 | 898.3235522818076 | 1142.6630349164311 | 723.1799317778431 | 783.6980436710774 | 786.7549561531005 | 1067.9951741713978 | 839.2562497590193 | 813.6373899222925 |
| SE_16_2_50 | SE_16 | SE_16_2 | GGCCCAGGAAGACGTCCCCA | 233.01202096298425 | 228.86087898116222 | 262.6821454607887 | 209.51066682019047 | 373.3292684807512 | 228.7058690052599 | 188.3281072338133 | 343.16331707189516 | 192.69280528806183 | 280.66767375835366 | 335.7024999036077 | 347.04362280565687 |
| SE_16_3_12 | SE_16 | SE_16_3 | ATTCACAGTGAGGGTCCTGG | 20.59175999207768 | 51.968409200962604 | 53.32645058226538 | 52.05233958265602 | 37.120807945529236 | 16.459892087499767 | 12.913927353175769 | 46.32232103725307 | 83.47061611556596 | 16.76715973101853 | 4.695139858791716 | 5.803405063639747 |
| SE_16_3_25 | SE_16 | SE_16_3 | GACACCGAAAGAACCCCAGT | 1484.7742731129695 | 1771.922875255898 | 1869.3883509671919 | 1332.539893315994 | 1457.2568604902049 | 822.1282942651198 | 892.1371479818928 | 724.1407737660377 | 1523.7827367479913 | 319.305041834179 | 361.52576912696213 | 235.61824558377373 |
| SE_16_3_33 | SE_16 | SE_16_3 | TCTATTCATCCAGCAGTGAA | 105.126353643765 | 188.8851795958064 | 160.96687860943067 | 145.74655083143685 | 144.2408537311993 | 201.85025559933922 | 143.1293614976981 | 88.86322811228139 | 122.54196833987343 | 155.27847924812812 | 106.81443178751154 | 153.20989368008932 |
| SE_16_3_34 | SE_16 | SE_16_3 | GCACCAACTGCCCGGCCTGG | 309.9601767228535 | 334.7964823523552 | 321.93375721886133 | 284.9865592150417 | 357.4203507898101 | 360.38500570525804 | 265.8116713528679 | 329.92836820410855 | 304.57895029403323 | 289.41575709627637 | 557.5478582315163 | 420.1665266075177 |
| SE_16_3_41 | SE_16 | SE_16_3 | GCTTGTGGGGAAGCAGACGT | 3758.5380869750206 | 4883.031679921217 | 4805.305713579692 | 3146.5639277715563 | 3475.568218214266 | 1775.935725230238 | 2403.066648303458 | 1445.4454870604068 | 3127.484148500248 | 425.0110488340784 | 764.1340120183518 | 691.7658835858579 |
| SE_16_4_19 | SE_16 | SE_16_4 | AGATGAGGCCGGGACCGAGG | 1456.5960752290737 | 1848.8760965727079 | 1763.7229766652956 | 1616.2251440414693 | 1831.6467234836855 | 1670.245891826292 | 1423.7604906876286 | 1910.5594044140498 | 1714.699571480403 | 1308.5674659642723 | 1521.225314248516 | 1254.6961747589132 |
| SE_16_4_22 | SE_16 | SE_16_4 | CCACCGCAGCCTCTCCCCCT | 3800.805383800864 | 3831.670786086358 | 3957.020138576618 | 4052.274636509771 | 4129.955032568309 | 4257.914190003229 | 3944.128645782433 | 4610.488973442514 | 4174.418791056123 | 3859.3627659135695 | 5323.114814905108 | 4063.544225560551 |
| SE_16_4_31 | SE_16 | SE_16_4 | TTTGGAGTGGAGCCCTCAGA | 49.85373471766175 | 85.94775367851507 | 84.92731018657079 | 63.76411598875362 | 67.87804881468203 | 37.251334724341575 | 76.40740350628997 | 93.58999556506231 | 43.51127861343332 | 193.91584732395344 | 152.59204541073078 | 24.37430126728694 |
| SE_16_4_33 | SE_16 | SE_16_4 | CGCGGTAGACTGCCGCCACT | 2811.3171273394473 | 2311.594816958202 | 2355.2515673833877 | 2754.87007241207 | 2690.728278794505 | 2221.2191217026 | 2526.825118771392 | 2890.8909741208136 | 2594.6929818051462 | 2383.1237026391123 | 3043.62441346173 | 2539.5700558487533 |
| SE_16_4_9 | SE_16 | SE_16_4 | GATGAGAGGAGATCTCCCCA | 1899.860803479588 | 2347.5729464050223 | 2326.6132883669857 | 2207.019198304615 | 2189.0670742734956 | 2021.1014863229975 | 2132.9503344995314 | 2178.0944422414505 | 2097.4212262230512 | 2033.9293760670305 | 2112.8129364562724 | 2334.1295165959064 |
| SE_16_5_13 | SE_16 | SE_16_5 | GGAGCCACAGGTGGATCCCG | 40.09974314246706 | 61.96233404730156 | 52.338923719630834 | 49.44972260352321 | 41.363185996446866 | 216.5775274671022 | 39.817942672291956 | 82.2457536783881 | 61.27098416993672 | 29.160277793075707 | 24.64948425865651 | 10.446129114551544 |
| SE_16_5_14 | SE_16 | SE_16_5 | TCAGCCTCGGGATCCACCTG | 3198.2254598221703 | 3014.167733655831 | 3094.909187496661 | 2571.385575383207 | 2825.4237819111395 | 2000.3100436861557 | 2520.3681550948045 | 2088.2858606386126 | 3157.675647946304 | 1401.1513479572877 | 1772.4152966938727 | 791.5844506804615 |
| SE_16_5_17 | SE_16 | SE_16_5 | CAGGGCAGCCAGTTCAGCCT | 23434.50664782609 | 19539.122467077304 | 19197.522209615538 | 22845.771842827726 | 19608.27135134127 | 20541.07901508984 | 23069.65505583575 | 19987.60885082942 | 18375.07935403624 | 21331.472212579705 | 21095.26338555118 | 20822.61736833941 |
| SE_16_5_20 | SE_16 | SE_16_5 | CTGTCCCTCAGTCTTGAAGC | 254.68755779675024 | 209.8724217731182 | 227.13117840594512 | 216.01720926802247 | 271.5121952587281 | 295.41174746512735 | 270.1163138039265 | 236.33837263904624 | 353.4181405744176 | 77.27473615165061 | 203.06479889274172 | 662.7488582676591 |
| SE_16_5_21 | SE_16 | SE_16_5 | CATGCCGGCTTCAAGACTGA | 4304.761615185923 | 5038.936907524105 | 5043.299687474617 | 4352.876897599609 | 4810.856709740589 | 5002.074574380192 | 3677.2408138168003 | 4506.500089481334 | 4772.920868310288 | 4169.190717464999 | 4860.643538814124 | 3758.2851192131 |
| SE_160_1_14 | SE_160 | SE_160_1 | GGGAAAGCAGAGTAAGTAGA | 369.56790301570993 | 472.7126452318329 | 450.3122493613521 | 454.15666285867377 | 494.23704293190355 | 395.9037202098628 | 450.9112967483873 | 599.3541130126213 | 455.5364475243121 | 879.9113824060595 | 738.3107427949974 | 312.2231924238184 |
| SE_160_1_15 | SE_160 | SE_160_1 | ATTACCTGCCCATCATTGAA | 214.5878146542832 | 199.87849692677923 | 227.13117840594512 | 217.31851775758886 | 190.90701229129323 | 168.0641613144713 | 137.74855843387488 | 134.24019565897828 | 226.43624584541828 | 344.09127795829335 | 150.24447548133492 | 155.53125570554522 |
| SE_160_1_20 | SE_160 | SE_160_1 | GGGTTAGTGTTCATAATCAA | 95.37236206857031 | 109.93317330972857 | 80.97720273603261 | 104.10467916531204 | 64.69626527649382 | 126.48127604078768 | 109.76838250199404 | 57.66656292392729 | 120.76599778422309 | 101.33196533093808 | 109.1620017169074 | 39.46315443275028 |
| SE_160_1_5 | SE_160 | SE_160_1 | CTCCATTCTGGAGTAACCAG | 11.921545258571289 | 8.994532361705065 | 13.825376076883616 | 11.711776406097604 | 14.848323178211695 | 30.320853845394307 | 17.21856980423436 | 3.78141396222474 | 15.095749723027886 | 220.1600973377216 | 2.347569929395858 | 4.642724050911798 |
| SE_160_1_8 | SE_160 | SE_160_1 | ACCAGGGGCACAATGCACGG | 3851.7428953602143 | 4205.4435753394355 | 4247.3530361911735 | 3710.030503753808 | 4404.649011365226 | 4211.999754180203 | 3708.449471586975 | 5060.477234947259 | 3938.214707154628 | 4171.3777382994795 | 5127.092725800554 | 4467.461217989878 |
| SE_160_2_10 | SE_160 | SE_160_2 | AGATTTAAAGACAAGACCAT | 67.19416418467453 | 126.92284554850481 | 107.6404280271653 | 74.17458390528482 | 93.3323171201878 | 63.24063802039384 | 72.10276105523138 | 58.61191641448347 | 59.49501361428638 | 75.08771531716994 | 102.11929192871983 | 70.80154177640492 |
| SE_160_2_14 | SE_160 | SE_160_2 | CATGGAGCTACCGTTTAGGT | 7457.468447657185 | 6580.999511314207 | 6571.003743970256 | 6917.755930534985 | 6880.076604075662 | 6921.817777848586 | 6845.457657795922 | 7163.88875143477 | 6494.724322013291 | 7481.069267813573 | 7823.276789711697 | 7366.842387784295 |
| SE_160_2_19 | SE_160 | SE_160_2 | GGTCTAGCTCTGTCACCACT | 205.91759992077678 | 256.84386855091134 | 224.1685978180415 | 174.37533760189766 | 212.11890254588135 | 53.71122681184134 | 231.3745317443992 | 196.6335260356865 | 157.17339417505505 | 696.2016323096825 | 131.46391604616804 | 284.3668481183476 |
| SE_160_2_3 | SE_160 | SE_160_2 | AACGGCATGGTTGAACCTGG | 7385.939176105757 | 6647.958807784677 | 6567.053636519718 | 7233.97389349962 | 6751.744668035403 | 7628.7268275012075 | 7548.1905379312375 | 6900.135127569594 | 7324.990556779826 | 7470.134163641169 | 8640.231125141456 | 8528.684081524972 |
| SE_160_2_4 | SE_160 | SE_160_2 | CAATATAATGTGCTTCCAGA | 175.57184835350444 | 185.8870021419047 | 202.44300684008152 | 173.07402911233126 | 160.14977142214042 | 196.65239494012877 | 195.86123152316583 | 253.35473546905757 | 195.35676112153735 | 77.27473615165061 | 165.503680022408 | 259.9925468510607 |
| SE_160_3_10 | SE_160 | SE_160_3 | ATTCTGCACGGTCTGAGCCA | 111.62901469389479 | 155.9052276028878 | 135.29118018093254 | 130.13084895664005 | 164.39214947305805 | 330.9304619697321 | 103.31141882540615 | 155.98332594177052 | 212.22848140021557 | 91.12586810336158 | 241.7997027277734 | 84.7297139291403 |
| SE_160_3_4 | SE_160 | SE_160_3 | TCACGTGCCACATTTGCTTG | 25958.622912118135 | 19759.988206181395 | 19845.339831503796 | 23083.911296418377 | 20854.469903798326 | 21303.431911774038 | 25746.066499781427 | 20798.722145726628 | 20603.922401377415 | 20701.61021224927 | 21472.048359219214 | 18372.41975047071 |
| SE_160_3_6 | SE_160 | SE_160_3 | CATCACGTGCCACATTTGCT | 1449.0096373372558 | 1277.2235953621193 | 1233.4210514305455 | 1462.670742272634 | 1153.9268298495945 | 1427.6790610631376 | 1326.9060355388103 | 1231.795598194709 | 1063.8063628345535 | 1388.0292229504037 | 1069.3181028398133 | 1944.1406963193153 |
| SE_160_3_8 | SE_160 | SE_160_3 | AAGCAAATGTGGCACGTGAT | 1947.5469845138732 | 1912.8372155892773 | 1839.7625450881555 | 1761.9716948729063 | 1857.1009917891913 | 1696.2351951223443 | 1861.7578600828401 | 1542.816896587694 | 2018.3905364966108 | 1468.2199868813618 | 1576.3932075893188 | 1685.3088304809826 |
| SE_160_3_9 | SE_160 | SE_160_3 | CGACAATGTAACATTCTGCA | 1945.3794308304966 | 424.7418059694059 | 873.9612734315715 | 1368.9765310238533 | 393.4805642226099 | 4076.8553770407316 | 2596.7755586010944 | 1039.8888396118034 | 3193.1950590593106 | 174.96166675845424 | 633.8438809368816 | 1599.4184355391144 |
| SE_160_4_15 | SE_160 | SE_160_4 | TTTGAGCCACAGCTGAAGAT | 275.27931778882794 | 263.8396159433486 | 266.6322529113269 | 322.7245054124673 | 201.5129574185873 | 333.52939229933736 | 253.9739046124568 | 242.95584707293955 | 237.9800544571455 | 228.1791737308174 | 225.36671322200237 | 458.46900002754 |
| SE_160_4_16 | SE_160 | SE_160_4 | GAGCCACAGCTGAAGATTGG | 1919.3687866299774 | 2072.740013130701 | 2018.504907225008 | 1941.5522664330695 | 1958.9180650112144 | 1636.459797541424 | 1998.4302579039504 | 2622.4105828028573 | 2243.938797064204 | 2028.0973205084154 | 2612.84533141759 | 2149.5812355721623 |
| SE_160_4_25 | SE_160 | SE_160_4 | ATACCACGCCCCTCTCCAGC | 1501.030925738294 | 1662.9890944308033 | 1800.2614705827739 | 1478.286444147431 | 1630.1337660650981 | 2018.5025559933924 | 1490.4824486790367 | 2063.706669884152 | 1745.7790562042837 | 1944.9905287981496 | 1770.067726764477 | 1537.902341864533 |
| SE_160_4_5 | SE_160 | SE_160_4 | TTATAGGTTACCCAGCAGGA | 23.843090517142578 | 34.97873696218637 | 34.56344019220904 | 13.013084895664004 | 10.605945127294067 | 0.0 | 5.380803063823238 | 4.726767452780925 | 28.415528890405433 | 33.53431946203706 | 7.042709788187574 | 3.482043038183848 |
| SE_160_4_6 | SE_160 | SE_160_4 | AAGAAATTATTAGGTAGACA | 7.586437891818092 | 6.9957473924372735 | 10.862795488979984 | 11.711776406097604 | 22.272484767317543 | 18.192512307236584 | 2.152321225529295 | 22.68848377334844 | 11.543808611727208 | 0.7290069448268927 | 5.868924823489645 | 15.088853165463343 |
| SE_161_1_13 | SE_161 | SE_161_1 | CTTCCTGGAGTATTTCAGCA | 71.52927155142773 | 127.92223803313871 | 125.4159115545871 | 59.86019052005442 | 160.14977142214042 | 111.75400417302473 | 57.03651247652632 | 151.2565584889896 | 103.0062922277197 | 52.48850002753627 | 134.98527094026184 | 103.3006101327875 |
| SE_161_1_14 | SE_161 | SE_161_1 | GATCCCTGCTGAAATACTCC | 107.2939073271416 | 152.90705014898612 | 162.94193233469977 | 134.03477442533924 | 151.66501532030517 | 38.11764483420998 | 139.90087965940418 | 97.37140952728706 | 331.21850862878836 | 167.67159731018532 | 113.85714157569912 | 139.28172152735394 |
| SE_161_1_5 | SE_161 | SE_161_1 | TTTCAGTTGAAGGTCTGGAA | 13.005322100259587 | 10.993317330972857 | 20.738064115325425 | 0.0 | 4.242378050917627 | 9.529411208552496 | 2.152321225529295 | 1.89070698111237 | 4.439926389125849 | 0.7290069448268927 | 1.173784964697929 | 16.249534178191293 |
| SE_161_1_7 | SE_161 | SE_161_1 | AACTCCTATTGTGATCTTAA | 6.5026610501297935 | 7.9951398770711695 | 7.900214901076352 | 10.410467916531204 | 14.848323178211695 | 4.331550549342044 | 9.685445514881827 | 1.89070698111237 | 15.095749723027886 | 17.496166675845423 | 0.0 | 15.088853165463343 |
| SE_161_1_8 | SE_161 | SE_161_1 | ACAGCCCTTAAGATCACAAT | 1555.219767822709 | 1288.2169126930921 | 1364.76212416094 | 1483.4916781056966 | 1377.7122720354994 | 1247.4865582105085 | 1439.9028998790984 | 1293.243575080861 | 1593.9335736961798 | 1042.4799311024565 | 1474.273915660599 | 1344.0686127389654 |
| SE_161_2_10 | SE_161 | SE_161_2 | CTCTTCTCTGCTGGAAGCAA | 2.167553683376598 | 10.993317330972857 | 7.900214901076352 | 5.205233958265602 | 5.3029725636470335 | 0.8663101098684087 | 1.0761606127646475 | 0.0 | 0.8879852778251698 | 0.0 | 0.0 | 0.0 |
| SE_161_2_17 | SE_161 | SE_161_2 | CTTATTGTAACAGATTTCAG | 61.77527997623304 | 83.94896870924728 | 93.81505195028168 | 61.16149900962082 | 99.69588419656424 | 84.89839076710406 | 19.370891029763655 | 53.88514896170255 | 79.03068972644012 | 53.21750697236316 | 45.77761362321923 | 38.30247342002233 |
| SE_161_2_2 | SE_161 | SE_161_2 | GCCGACTAAAAGGGACAAGA | 907.1212164931062 | 1158.2958896906857 | 1158.36900987032 | 1031.9376322261555 | 1083.9275920094537 | 932.1496782184078 | 928.7266088158908 | 915.1021788583871 | 1128.6292881157908 | 645.9001531166269 | 968.3725958757915 | 1145.5921595624861 |
| SE_161_2_25 | SE_161 | SE_161_2 | TTTCCAGAGCTAAAGAAGTG | 6.5026610501297935 | 13.991494784874547 | 8.887741763710896 | 6.506542447832002 | 7.4241615891058474 | 1.7326202197368175 | 1.0761606127646475 | 0.945353490556185 | 24.863587779104755 | 7.290069448268927 | 1.173784964697929 | 0.0 |
| SE_161_2_27 | SE_161 | SE_161_2 | CTGTCAGCCTGCATGTAAAG | 162.56652625324483 | 139.91494784874547 | 156.02924429625796 | 169.17010364363205 | 107.12004578567009 | 106.55614351381428 | 174.33801926787288 | 230.66625169570915 | 106.55823333902038 | 48.114458358574915 | 318.0957254331388 | 156.69193671827318 |
| SE_161_3_10 | SE_161 | SE_161_3 | AGCCTGCCGGATCATCTTGG | 389.0758861660993 | 530.6774093405988 | 488.8257970040993 | 521.8247043161266 | 518.6307167246799 | 525.8502366901241 | 365.89460833998015 | 501.98270348533424 | 564.758636696808 | 519.0529447167476 | 484.7731904202447 | 501.41419749847415 |
| SE_161_3_15 | SE_161 | SE_161_3 | CTAATTCAATCAAGAAGAAA | 18.42420630870108 | 35.97812944682026 | 19.75053725269088 | 7.807850937398403 | 33.939024407341016 | 9.529411208552496 | 12.913927353175769 | 5.67212094333711 | 0.8879852778251698 | 1.4580138896537853 | 7.042709788187574 | 17.410215190919242 |
| SE_161_3_19 | SE_161 | SE_161_3 | GAATTCCTCATCCTGTTAGT | 54.188842084414944 | 56.96537162413208 | 69.12688038441809 | 50.751031093089615 | 54.09032014919974 | 16.459892087499767 | 64.56963676587885 | 55.775855942814914 | 44.39926389125849 | 45.198430579267345 | 26.997054188052367 | 34.820430381838484 |
| SE_161_3_5 | SE_161 | SE_161_3 | AGCCTTTCAGTCTGAACACG | 37.93218945909046 | 48.970231747060915 | 42.463655093285396 | 52.05233958265602 | 84.84756101835254 | 26.85561340592067 | 36.58946083399801 | 41.59555358447214 | 70.15083694818841 | 138.5113195171096 | 99.77172199932397 | 94.01516203096391 |
| SE_161_3_9 | SE_161 | SE_161_3 | TTCCACCAAGATGATCCGGC | 3.2513305250648967 | 5.9963549078033775 | 4.93763431317272 | 0.0 | 0.0 | 0.0 | 0.0 | 0.0 | 0.0 | 0.0 | 0.0 | 1.1606810127279494 |
| SE_162_1_10 | SE_162 | SE_162_1 | GCAAGCATCAGGAAACCCCT | 7064.057454124332 | 7427.484945799117 | 7319.549105847241 | 7131.170522823874 | 7582.190171502529 | 5926.427461609784 | 7230.723157165667 | 7647.9097385995365 | 7527.451200123965 | 6981.699510607151 | 6609.583136214038 | 6411.601914309193 |
| SE_162_1_12 | SE_162 | SE_162_1 | ACTCCTGTATTATCTCCAGA | 20.59175999207768 | 22.986027146579612 | 25.675698428498144 | 5.205233958265602 | 45.60556404736449 | 71.90373911907793 | 19.370891029763655 | 17.961716320567515 | 99.45435111641902 | 24.78623612411435 | 2.347569929395858 | 83.56903291641235 |
| SE_162_1_13 | SE_162 | SE_162_1 | TTACCTCCAGTTGAGAACCG | 56.35639576779154 | 102.9374259172913 | 89.8649444997435 | 61.16149900962082 | 45.60556404736449 | 32.05347406513112 | 41.970263897821255 | 28.36060471668555 | 106.55823333902038 | 78.00374309647752 | 28.170839152750297 | 3.482043038183848 |
| SE_162_1_7 | SE_162 | SE_162_1 | ATCGCTTGAAACCAAGGCAG | 521.2966608520718 | 589.6415659339988 | 579.6782683664774 | 569.9731184300834 | 495.2976374446329 | 855.9143885499878 | 502.5670061610904 | 666.4742108421104 | 542.5590047511787 | 458.5453682961155 | 703.0971938540595 | 615.1609367458132 |
| SE_162_1_9 | SE_162 | SE_162_1 | TAACTTGCCTGTGGCTACCC | 2231.4965170362075 | 1995.7867918138907 | 2038.2554444776988 | 2191.4034964298185 | 2040.5838424913786 | 2191.7645779670743 | 2213.66238045688 | 2526.9298802566827 | 2184.4437834499176 | 1659.9488133708346 | 2763.089806898925 | 2079.9403748084856 |
| SE_162_2_16 | SE_162 | SE_162_2 | GCAAGGTGACTGGTGTTAGG | 405.3325387914238 | 384.76610658405 | 367.3599929000504 | 403.40563176558413 | 271.5121952587281 | 287.6149564763117 | 316.39122015280634 | 380.03210320358636 | 409.36121307740325 | 242.75931262735526 | 157.2871852695225 | 270.4386759656122 |
| SE_162_2_23 | SE_162 | SE_162_2 | TTTCCTAAAATAGCATCGCC | 7.586437891818092 | 22.986027146579612 | 27.650752153767232 | 16.917010364363207 | 6.363567076376441 | 61.50801780065702 | 37.665621446762664 | 0.945353490556185 | 33.74344055735645 | 145.07238202055163 | 4.695139858791716 | 9.285448101823595 |
| SE_162_2_24 | SE_162 | SE_162_2 | GCCTTGTGTCCCGCTTAGGA | 631.8418987042783 | 615.62577053448 | 615.2292354213209 | 624.6280749918722 | 673.4775155831733 | 880.1710716263033 | 686.590470943845 | 670.2556248043352 | 638.461414756297 | 902.5105976956931 | 953.1133913347184 | 817.1194329604764 |
| SE_162_2_3 | SE_162 | SE_162_2 | TTACCCAGCGGTAGGTGTGA | 180.99073256194592 | 241.8529812814029 | 229.10623213121423 | 171.77272062276487 | 251.3608995168694 | 156.80212988618197 | 154.96712823810924 | 204.19635396013595 | 270.8355097366768 | 121.01515284126418 | 80.99116256415711 | 218.20803039285448 |
| SE_162_2_8 | SE_162 | SE_162_2 | CTTCTGCTGTGGTTTCCTAA | 58.52394945116814 | 91.94410858631845 | 74.06451469759081 | 53.35364807222242 | 90.15053358199957 | 129.0802063703929 | 58.112673089290965 | 83.19110716894428 | 80.80666028209045 | 67.79764586890101 | 61.03681816429231 | 40.62383544547823 |
| SE_163_1_10 | SE_163 | SE_163_1 | ACTTTGGTCCTTCTCAAAGG | 30.34575156727237 | 47.97083926242702 | 38.51354764274722 | 36.43663770785921 | 30.757240869152795 | 1.7326202197368175 | 71.02660044246673 | 78.46433971616335 | 47.063219724734 | 25.515243068941242 | 91.55522724643846 | 12.767491140007444 |
| SE_163_1_17 | SE_163 | SE_163_1 | CCGCTGAAGTGAGGAAACAG | 1445.7583068121908 | 1516.0783991896205 | 1475.3651327760087 | 1456.1641998248022 | 1284.3799549153116 | 1193.7753313986673 | 1477.568521325861 | 1505.0027569654465 | 1355.9535192390342 | 1009.6746185852463 | 1669.122219800455 | 1348.7113367898774 |
| SE_163_1_19 | SE_163 | SE_163_1 | ACAATACATGTGTACAAGGA | 679.5280797385634 | 683.584459489585 | 717.9320291353135 | 710.5144353032547 | 690.4470277868438 | 882.7700019559085 | 718.8752893267845 | 600.2994665031774 | 532.7911666951019 | 582.4765489166872 | 671.4049998072154 | 677.8377114331224 |
| SE_163_1_20 | SE_163 | SE_163_1 | GTAAGGATGAGGGCATCAGT | 1656.0110140997208 | 1721.9532510242032 | 1684.720827654532 | 1609.7186015936372 | 1756.3445130798975 | 1302.0640951322184 | 1858.5293782445463 | 1562.6693198893738 | 1476.7195170232574 | 1659.9488133708346 | 1427.3225170726816 | 1027.2026962642353 |
| SE_163_1_32 | SE_163 | SE_163_1 | GGTTTCTTCTTACAAACACT | 1.083776841688299 | 5.9963549078033775 | 10.862795488979984 | 1.3013084895664004 | 18.030106716399914 | 0.0 | 1.0761606127646475 | 0.0 | 12.431793889552377 | 1.4580138896537853 | 0.0 | 16.249534178191293 |
| SE_164_1_1 | SE_164 | SE_164_1 | TCTCCTGTTAATGTGTTACA | 28.17819788389577 | 52.9678016855965 | 45.42623568118903 | 41.641871666124814 | 83.78696650562313 | 44.18181560328885 | 39.817942672291956 | 77.51898622560716 | 31.96747000170611 | 88.20984032405401 | 22.30191432926065 | 73.12290380186082 |
| SE_164_1_11 | SE_164 | SE_164_1 | CTCTACCACAGCAAGAGCCA | 8.670214733506391 | 28.982382054382988 | 22.713117840594514 | 11.711776406097604 | 13.787728665482287 | 0.8663101098684087 | 1.0761606127646475 | 14.180302358342775 | 12.431793889552377 | 2.187020834480678 | 0.0 | 1.1606810127279494 |
| SE_164_1_14 | SE_164 | SE_164_1 | TATAGATGTATAAAGACAAA | 22.759313675454276 | 63.961119016569356 | 47.40128940645811 | 50.751031093089615 | 45.60556404736449 | 38.11764483420998 | 58.112673089290965 | 74.68292575393862 | 26.639558334755094 | 16.76715973101853 | 56.341678305500594 | 33.659749369110536 |
| SE_164_1_7 | SE_164 | SE_164_1 | ACTTCTACCAGGGAGAGCAA | 114.88034521895969 | 170.89611487239625 | 140.22881449410525 | 115.81645557140963 | 135.75609762936406 | 161.99999054539242 | 99.00677637434757 | 99.26211650839943 | 103.89427750554486 | 96.22891671714983 | 99.77172199932397 | 105.6219721582434 |
| SE_164_1_8 | SE_164 | SE_164_1 | CCAGGGAGACCGCAGCAACA | 4228.897236267742 | 4004.565685928022 | 3970.8455146535016 | 4071.794263853267 | 3854.200459258664 | 3992.8232963834957 | 4103.400416471601 | 3765.342952885285 | 4463.901991627128 | 3430.706682355357 | 3104.661231626022 | 3815.1584888367697 |
| SE_164_2_15 | SE_164 | SE_164_2 | GCCTGGGAGATCAAAAGCCT | 1028.5042227621957 | 1414.140365756963 | 1310.44814671604 | 1156.86324722453 | 1362.8639488572876 | 1574.951779740767 | 1034.1903488668263 | 1460.5711429093058 | 1134.845185060567 | 1389.4872368400574 | 1397.9778929552335 | 962.2045595514701 |
| SE_164_2_20 | SE_164 | SE_164_2 | AGTCCCCAACAAAGGACCCG | 27.094421042207472 | 11.992709815606755 | 6.912688038441808 | 32.53271223916001 | 28.63605184369398 | 0.0 | 29.056336544645482 | 27.415251226129364 | 8.879852778251697 | 24.78623612411435 | 28.170839152750297 | 0.0 |
| SE_164_2_38 | SE_164 | SE_164_2 | GTGAATGCCAGTGTTCAGCA | 1250.678475308297 | 1312.2023323243056 | 1318.3483616171163 | 1302.6097980559668 | 1254.6833085588883 | 1383.4972454598487 | 1301.0781808324589 | 1432.2105381926203 | 1060.2544217232528 | 1910.7272023912856 | 989.5007252403542 | 1265.142303873465 |
| SE_164_2_39 | SE_164 | SE_164_2 | CCAGGAGCCCTGCTGAACAC | 1325.4590773847895 | 1058.356641227296 | 1058.6287967442313 | 1156.86324722453 | 1180.4416926678298 | 1126.2031428289313 | 1230.051580389992 | 1181.6918631952312 | 1093.109877002784 | 1301.2773965160034 | 1353.3740642967123 | 1188.5373570334202 |
| SE_164_3_10 | SE_164 | SE_164_3 | GAGAGTTGTCGCACTAGGGG | 733.7169218229784 | 954.4198228253708 | 923.3376165632986 | 972.0774417061011 | 1011.807165143854 | 942.5453995368287 | 811.4251020245442 | 733.5943086715996 | 955.4721589398827 | 936.7739241025571 | 629.14874107809 | 540.8773519312244 |

| | | | | | | | | | | | | | | | |
|---|---|---|---|---|---|---|---|---|---|---|---|---|---|---|---|
| SE_164_3_22 | SE_164 | SE_164_3 | CCTCCCAGTGTGCCAGGACA | 1045.8446522292084 | 1041.3669689885198 | 1001.3522387114276 | 1064.4703444653155 | 1023.4737047838776 | 1045.6363026111694 | 1137.5017676922323 | 1089.0472211207252 | 1145.501008394469 | 814.3007573716391 | 753.5699473360704 | 1493.7964633808708 |
| SE_164_3_23 | SE_164 | SE_164_3 | CTGGCATTAGGCCCAAACAA | 469.27537245103343 | 459.72054293159226 | 474.01289406458113 | 420.3226421299473 | 411.5106709390098 | 392.43847977038916 | 505.7954879993843 | 441.4800800897384 | 537.2310930842277 | 406.7858752134061 | 353.30927437407667 | 493.2894304093785 |
| SE_164_3_28 | SE_164 | SE_164_3 | CAGACATCATTACTTCCTCC | 3233.9900955978837 | 2880.2491407148887 | 2929.992201436692 | 2814.7302629321243 | 2875.2717240094216 | 2682.9624102624616 | 3389.9059302086393 | 2922.087639309168 | 2577.821261526468 | 2534.028140218279 | 2935.6361967095204 | 2381.7174381177524 |
| SE_165_1_10 | SE_165 | SE_165_1 | ATAAAGACGTGCTACTGCAT | 178.8231788785693 | 227.86148649652833 | 214.29332919169605 | 214.71590077845607 | 127.2713415275288 | 184.52405340197106 | 136.67239782111022 | 121.00524679119168 | 243.3079661240965 | 188.0837917653383 | 117.3784964697929 | 199.6371341892073 |
| SE_165_1_11 | SE_165 | SE_165_1 | GTGTGTCTTGATGGTAACAG | 229.76069043791938 | 278.830503212857 | 269.5948334992305 | 304.5061865585377 | 284.239329411481 | 146.40640856776108 | 235.6791741954578 | 416.9008893352776 | 234.42811334584482 | 337.5302154548513 | 692.5331291717781 | 279.7241240674358 |
| SE_165_1_25 | SE_165 | SE_165_1 | AACTGCCCTCCAGAGAAGCA | 248.18489674662044 | 289.82382054382987 | 299.22063937826687 | 238.13945359065127 | 276.81516782237514 | 252.96255208157535 | 299.172650348572 | 305.34917744964775 | 211.34049612239042 | 1007.4875977507656 | 261.75404712763816 | 331.95476964019355 |
| SE_165_1_28 | SE_165 | SE_165_1 | AGCAAGTCCTTGCTGACCCA | 344.64103565687907 | 309.81167023650784 | 261.6946185981542 | 307.1088035376705 | 305.45121966606916 | 307.5400890032851 | 261.50702890180935 | 339.3819031096704 | 279.7153625149285 | 344.82028490312024 | 350.9617044446808 | 286.68821014380353 |
| SE_165_1_38 | SE_165 | SE_165_1 | TGTTTCCCTACTTAGTACCA | 13.005322100259587 | 23.98541963121351 | 6.912688038441808 | 16.917010364363207 | 2.1211890254588135 | 9.529411208552496 | 36.58946083399801 | 0.0 | 1.7759705556503396 | 0.0 | 0.0 | 0.0 |
| SE_165_2_12 | SE_165 | SE_165_2 | TCGGACACAGAAGGCATGCA | 13740.122798924254 | 11718.876274817067 | 11619.241065758046 | 13415.189218940022 | 11752.447795554555 | 12272.149016395879 | 14148.28357601682 | 11794.230148178964 | 11778.236725073051 | 12288.141062002103 | 13053.662592405668 | 13683.268459049796 |
| SE_165_2_16 | SE_165 | SE_165_2 | GTGAGCCTCGACTTATCTAT | 656.7687660631092 | 726.5583363288425 | 723.8571903111208 | 728.7327541571842 | 753.0221040378788 | 893.1657232743294 | 719.9514499395492 | 676.8730992382285 | 599.3900625319897 | 820.8618198750811 | 805.2164857827793 | 774.1742354895423 |
| SE_165_2_18 | SE_165 | SE_165_2 | TAGCATGTAGCAGGTCTCTG | 1765.472475110239 | 2041.75884610705 | 2056.0309280051206 | 1948.0588088809016 | 2232.551449295401 | 1807.1228891855005 | 1700.333768168143 | 2671.568964311779 | 2037.9262126087647 | 1869.1738065361528 | 2219.6273682437836 | 2313.2372583668034 |
| SE_165_2_19 | SE_165 | SE_165_2 | TCATTTCTCCCAGGGAGTAG | 78.03193260155751 | 102.9374259172913 | 94.80257881291622 | 78.07850937398402 | 107.12004578567009 | 75.36897955855156 | 83.94052779564251 | 108.71565141396128 | 110.99815972814622 | 118.8281320067835 | 77.46980767006332 | 66.15881772549312 |
| SE_165_2_6 | SE_165 | SE_165_2 | TAGATGGTCTTGCTTGTCAG | 2612.9859653104886 | 2062.7460882843616 | 2161.696302307017 | 2482.896598092692 | 2230.430260269942 | 1907.614861930236 | 2532.2059218352156 | 2020.2204093185674 | 2040.5901684422402 | 2057.2575983014913 | 2454.3843611833695 | 2579.0332102815037 |
| SE_166_1_1 | SE_166 | SE_166_1 | GCTCCGGATCCCGCCTCGCT | 348.97614302363223 | 378.76975167624664 | 392.048164465914 | 400.80301478645134 | 386.05640263350404 | 401.9678909789416 | 376.6562144676266 | 328.9830147135524 | 468.85622669168964 | 333.15617378588996 | 226.5404981867003 | 407.39903546751026 |
| SE_166_1_15 | SE_166 | SE_166_1 | GGCCAGGCCGGGAGGTACCA | 644.8472208045379 | 874.4684240546592 | 902.5995524479732 | 884.8897729051523 | 844.2332321326078 | 913.9571659111712 | 694.1235952331976 | 521.8351267870141 | 1102.8777150588608 | 499.36975720642147 | 379.1325435974311 | 631.4104709240045 |
| SE_166_1_21 | SE_166 | SE_166_1 | GGGGCAAGGGTCATCTCGGA | 714.2089386725889 | 791.5188478300457 | 915.4374016622223 | 754.7589239485122 | 935.4443602273368 | 1009.2512779966962 | 846.9384022457775 | 697.6708760304646 | 1014.9671725541691 | 541.6521600063812 | 848.6465294766027 | 619.803660796725 |
| SE_166_1_28 | SE_166 | SE_166_1 | GATCAGAGGTCTGGGCTGTG | 838.8432754667433 | 1112.3238353975264 | 989.5019163598131 | 917.4224851443123 | 1058.473323703948 | 783.1443393210415 | 747.93162587143 | 770.4630948032908 | 1006.9753050537425 | 1059.247090833475 | 1218.3887933564504 | 763.7281063749907 |
| SE_166_1_32 | SE_166 | SE_166_1 | TTTAATAAGGATCAGAGGTC | 4.335107366753196 | 11.992709815606755 | 6.912688038441808 | 14.314393385230405 | 8.484756101835254 | 0.8663101098684087 | 7.533124289352532 | 0.0 | 0.8879852778251698 | 0.0 | 1.173784964697929 | 0.0 |
| SE_166_2_11 | SE_166 | SE_166_2 | AAGGAGGTCCAGAATCATTG | 2821.0711189146423 | 3196.0571658592 | 2992.2063937826683 | 2855.0708261086825 | 2942.0891783113743 | 2984.438328496668 | 2801.246075026377 | 3089.4152071376125 | 3129.2601190558985 | 3485.382203217374 | 3116.3990812730017 | 3188.390741963677 |
| SE_166_2_18 | SE_166 | SE_166_2 | ATATCTTCTCCTGTTTGAAA | 10.837768416882989 | 14.990887269508443 | 11.850322351614528 | 11.711776406097604 | 19.090701229129323 | 1.7326202197368175 | 7.533124289352532 | 1.89070698111237 | 0.8879852778251698 | 23.328222234460565 | 29.344624117448227 | 1.1606810127279494 |
| SE_166_2_19 | SE_166 | SE_166_2 | ACAGGCATCTGAACCACGGG | 3331.530011349831 | 3610.805046982267 | 3683.4751976268494 | 3496.615911464918 | 3548.749239592595 | 3559.6682414492916 | 3386.6774483703457 | 3902.4192090159318 | 3512.869759076372 | 4171.3777382994795 | 4392.303337899651 | 4097.203974929662 |
| SE_166_2_21 | SE_166 | SE_166_2 | TGTTACAGGCATCTGAACCA | 932.048083851937 | 1022.3785117804758 | 1055.6662161563277 | 871.8766880094882 | 1112.5636438531476 | 543.1764388874923 | 830.7959930543078 | 1161.8394398935513 | 1223.643712843084 | 1228.3767020333141 | 1022.3667042518962 | 916.93800005508 |
| SE_166_2_24 | SE_166 | SE_166_2 | CAGATTGTGAAGAAGAGGCA | 904.9536628097296 | 1035.3706140807165 | 989.5019163598131 | 935.640803998242 | 915.293064485478 | 777.9464786618311 | 811.4251020245442 | 768.5723878221784 | 853.3538519899881 | 570.08343085463 | 684.3166344188926 | 841.4937342277633 |
| SE_166_3_10 | SE_166 | SE_166_3 | GAAGAAATCTCTGAAGACCC | 92.12103154350541 | 158.9034050567895 | 165.9045129226034 | 135.33608291490566 | 172.8769055748933 | 96.16042219539337 | 85.01668840840715 | 178.67180971511897 | 160.72533528635574 | 185.16776398603074 | 273.4918967746175 | 150.88853165463343 |
| SE_166_3_15 | SE_166 | SE_166_3 | GTCTGTGCCTCAGAGCCCCA | 190.74472413714062 | 177.89186226483352 | 164.91698605996885 | 210.81197530975686 | 183.48285070218736 | 234.77003977433876 | 218.46060439122343 | 212.70453537514163 | 187.36489362111084 | 239.1142779032208 | 143.20176569314734 | 150.88853165463343 |
| SE_166_3_18 | SE_166 | SE_166_3 | TCTTCCAGAGACAAAGGCTG | 189.6609472954523 | 259.842046004813 | 316.00859604305407 | 217.31851775758886 | 204.6947409567755 | 279.818165487496 | 174.33801926787288 | 169.21827480955713 | 237.9800544571455 | 121.01515284126418 | 158.4609702342204 | 221.69007343103834 |
| SE_166_3_4 | SE_166 | SE_166_3 | TTCAGGTTCGTGCCAGATGG | 124.63433679415438 | 96.94107100948793 | 108.62795488979984 | 113.21383859227684 | 113.48361286204653 | 108.2887637335511 | 123.75847046793446 | 135.18554914953447 | 36.40739639083196 | 62.69459725511277 | 130.29013108147012 | 85.89039494186825 |
| SE_166_3_8 | SE_166 | SE_166_3 | AGTGGGGTCACGGGGCCCAG | 380.4056714325929 | 602.6336682342394 | 615.2292354213209 | 463.26582228563854 | 606.6600612812207 | 737.2299034980158 | 245.3646197103396 | 656.0753224459924 | 618.0377533663182 | 403.8698474340985 | 487.12076034964053 | 641.856600038556 |
| SE_167_1_11 | SE_167 | SE_167_1 | GTAGCTTAAACATAAGAAGG | 183.15828624532253 | 203.8760668653148 | 221.20601723013786 | 148.34916781056964 | 163.33155496032865 | 137.743307469077 | 120.52998862964051 | 153.14726547010198 | 157.17339417505505 | 112.26706950334147 | 45.77761362321923 | 227.4934784946781 |
| SE_167_1_3 | SE_167 | SE_167_1 | CAGCCCTTTGAACTAAAGGG | 43.351073667531956 | 30.98116702365078 | 49.376343131727204 | 42.943180155691216 | 98.63528968383483 | 25.989303296052263 | 44.12258512335055 | 43.48626056558451 | 79.91867500426528 | 40.82438891030599 | 21.128129364562724 | 1.1606810127279494 |
| SE_167_1_4 | SE_167 | SE_167_1 | GAGACTATACTATAGACAAA | 126.80189047753097 | 128.9216305177726 | 170.8421472357761 | 147.04785932100324 | 144.2408537311993 | 213.97859713749696 | 144.20552211046277 | 103.98888396118035 | 149.18152667462851 | 47.38545141374802 | 115.03092654039705 | 228.65415950740604 |
| SE_167_1_8 | SE_167 | SE_167_1 | AGATGGCTCCTTCAGAGGAG | 215.67159149597148 | 360.7806869528365 | 311.0709617298814 | 176.97795458103047 | 199.39176839312847 | 245.16576109275968 | 117.30150679134657 | 112.49706537618601 | 293.9231269601312 | 73.62970142751615 | 24.64948425865651 | 251.86777976196504 |
| SE_167_1_9 | SE_167 | SE_167_1 | TGTAACCCTGTAATGCCCCA | 651.3498818546676 | 768.5328206834662 | 705.0941799210644 | 714.4183607719539 | 780.5975613688433 | 742.4277641572263 | 647.8486888843178 | 723.1954202754815 | 813.3945144878555 | 902.5105976956931 | 888.5552182763323 | 331.95476964019355 |
| SE_167_2_13 | SE_167 | SE_167_2 | GAGCACACAGTATTCTGCCT | 2863.3384157404857 | 2830.279516483194 | 2862.8403747775433 | 2838.153815744319 | 2667.395199514458 | 2975.775227397984 | 2709.7724229413825 | 2802.9730994990887 | 2737.6586115349983 | 3064.7451960522567 | 2567.0677177943708 | 3559.8086660366207 |
| SE_167_2_2 | SE_167 | SE_167_2 | TGACGATGATAGCAACCACA | 9668.373204701315 | 8088.083378142122 | 8077.96973635057 | 8939.989323321171 | 8177.183693143726 | 8228.213423530146 | 9515.412138065012 | 7525.0137848272325 | 7861.333664586228 | 6699.573822959143 | 7964.130985475448 | 8130.570494159286 |
| SE_167_2_26 | SE_167 | SE_167_2 | TCTCCCTTTGAAATGAGTAA | 5.4188842084414945 | 16.989672238776237 | 17.77548352742179 | 3.9039254686992013 | 27.575457330964575 | 0.0 | 1.0761606127646475 | 20.79777679223607 | 24.863587779104755 | 0.0 | 4.695139858791716 | 2.321362025455899 |
| SE_167_2_4 | SE_167 | SE_167_2 | CATTGGTAAACAACAGTCAG | 1615.9112709572537 | 1617.017040137644 | 1687.6834082424357 | 1617.5264525310358 | 1524.0743147921576 | 1587.0801212789247 | 1564.7375309597974 | 1392.5056915892606 | 1357.7294897946847 | 2032.4713621773767 | 1386.2400433082541 | 1965.0329545484185 |
| SE_167_2_5 | SE_167 | SE_167_2 | CTTCAGCCAAGAAAGCATGA | 18.42420630870108 | 16.989672238776237 | 18.763010390056337 | 20.820935833062407 | 7.4241615891058474 | 0.8663101098684087 | 21.52321225529295 | 3.78141396222474 | 22.199631945629246 | 5.103048613788249 | 4.695139858791716 | 1.1606810127279494 |
| SE_167_3_2 | SE_167 | SE_167_3 | AGTTGGGCAGGAGAGATGAC | 721.7953765644071 | 729.5565137827442 | 756.4455767780607 | 878.3832304573203 | 886.657012641784 | 746.7593147065684 | 733.9415379054896 | 931.1731881978423 | 824.0503378217576 | 812.1137365371584 | 1074.0132426986052 | 649.9813671276517 |
| SE_167_3_4 | SE_167 | SE_167_3 | CCAAGAGGTTGGGAAAGAAG | 218.92292202103638 | 209.8724217731182 | 186.64257703792882 | 217.31851775758886 | 232.27019828774007 | 169.79678153420812 | 301.3249715741013 | 176.7811027340066 | 184.70093778763533 | 138.5113195171096 | 208.93372371623136 | 258.8318658383327 |
| SE_167_3_5 | SE_167 | SE_167_3 | ACAGGTGACAGGCATGGTAG | 1688.5243193503698 | 1442.1233553267123 | 1459.5647029738561 | 1519.9283158135556 | 1389.378811675523 | 1568.0212988618198 | 1702.4860893936723 | 1334.8391286653332 | 1562.8540889722988 | 1187.5523131230082 | 1520.0515292838181 | 1269.7850279243767 |
| SE_167_3_7 | SE_167 | SE_167_3 | CAGGACACACCATAATACCA | 13059.510942344003 | 11528.991702736626 | 11619.241065758046 | 12270.03774812159 | 12137.44360367533 | 11321.806825870233 | 12562.02283280173 | 12156.300535061982 | 11562.456302561535 | 10679.951741713978 | 11785.974830531906 | 10814.064995586305 |
| SE_167_3_8 | SE_167 | SE_167_3 | GACACACCATAATACCAGGG | 1232.2542689995958 | 1371.1664889177055 | 1429.9388970948198 | 1267.474468837674 | 1414.8330799810285 | 1506.5132810611628 | 1166.5581042368779 | 1174.1290352707817 | 1284.0267117351955 | 1218.1706048057376 | 1021.1929192871983 | 945.9550253732788 |
| SE_168_1_29 | SE_168 | SE_168_1 | GTTGTGTGTGTCACGAGGCG | 894.1158943928466 | 908.4477685322116 | 902.5995524479732 | 948.6538888939059 | 903.6265248454546 | 676.5881958072272 | 857.700008373424 | 831.9110716894428 | 1184.5723606187764 | 831.7969240474845 | 819.3019053591545 | 1323.1763545098624 |
| SE_168_1_33 | SE_168 | SE_168_1 | TGACACACACAACCCCTGGG | 2015.8249255402359 | 2114.7144974853245 | 2131.082969565346 | 2122.434146482799 | 2097.8559461787668 | 1882.4918687440522 | 2053.3144491549474 | 2802.9730994990887 | 2110.7410053904287 | 1930.4103899016118 | 2838.2120446395925 | 2090.386503923037 |
| SE_168_1_34 | SE_168 | SE_168_1 | TACGCAGGCAAAAGAGCCCG | 6727.002856359271 | 7732.299653612455 | 8014.76801714196 | 6976.3148125654725 | 8271.576604776643 | 8071.411293643964 | 6561.351256026056 | 7858.723566993566 | 8721.791398798818 | 9485.838366087528 | 7712.941003030091 | 9062.59734737983 |
| SE_168_1_46 | SE_168 | SE_168_1 | AGAGCCCCAGAGGCCTCTAA | 6.5026610501297935 | 13.991494784874547 | 20.738064115325425 | 15.615701874796805 | 4.242378050917627 | 45.048125713157255 | 12.913927353175769 | 1.89070698111237 | 15.095749723027886 | 17.496166675845423 | 18.780559435166865 | 1.1606810127279494 |
| SE_168_2_24 | SE_168 | SE_168_2 | CTTTACTGTGAAGAAGCCGT | 70.44549470973944 | 96.94107100948793 | 88.87741763710896 | 62.46280749918722 | 124.08955798934059 | 122.14972549144564 | 32.28481838293943 | 151.2565584889896 | 159.83735000853056 | 74.35870837234305 | 109.1620017169074 | 23.21362025455899 |
| SE_168_2_32 | SE_168 | SE_168_2 | TCCTGAGAAGAGCTGCCCAG | 880.0267954508987 | 908.4477685322116 | 948.0257881291623 | 893.9989323321171 | 1000.1406255038305 | 913.0908558013028 | 870.6139357265998 | 698.6162295210207 | 991.8795553307147 | 844.1900421095417 | 557.5478582315163 | 482.843301294827 |
| SE_168_2_35 | SE_168 | SE_168_2 | GACCTCCTCCTGGATCACCA | 425.92429878350146 | 387.7642840379517 | 417.72386289441215 | 313.61534598550253 | 314.9965702806338 | 488.59890196578255 | 470.28218777815096 | 417.84624282583377 | 394.26546335437536 | 512.4918822133055 | 474.20912573796335 | 503.7355595239301 |
| SE_168_2_5 | SE_168 | SE_168_2 | GCACAGCTCTGCGTTACCCG | 5477.4081578926625 | 5236.816619481616 | 5278.331080781638 | 5343.17265815964 | 5188.428356272258 | 4693.668175267038 | 5681.051874784574 | 4922.455625326055 | 5322.583755284068 | 5159.18214853992 | 5179.913049211961 | 5068.693982582955 |
| SE_168_2_7 | SE_168 | SE_168_2 | GGCCATTTCCGGCCTCCACA | 0.0 | 4.996962423169481 | 0.0 | 0.0 | 4.242378050917627 | 0.0 | 0.0 | 0.0 | 0.0 | 0.0 | 0.0 | 1.1606810127279494 |
| SE_168_3_1 | SE_168 | SE_168_3 | ACAGAGAGACAACCAGGTAA | 2232.5802938778957 | 2343.5753764664864 | 2134.0455501532497 | 2642.9575423093593 | 2715.121952587281 | 3142.973078602587 | 2829.2262509582583 | 2720.7273458207005 | 2520.990203745657 | 2974.348334893722 | 3108.1825865201163 | 2805.3660077634536 |
| SE_168_3_21 | SE_168 | SE_168_3 | GTATATCGAGGCCAAAGGAA | 134.38832836934907 | 141.91373281801324 | 169.85462037314159 | 149.65047630013606 | 151.66501532030517 | 151.60426922697152 | 101.15909759987686 | 161.65544688510764 | 110.99815972814622 | 154.54947230330123 | 98.59793703462604 | 308.74114938563457 |
| SE_168_3_30 | SE_168 | SE_168_3 | AGGCCTGGAGGCTAACACCA | 1204.0760711157002 | 1187.2782717450687 | 1190.9573963372602 | 1207.6142783176197 | 1309.8342232208174 | 1038.7058217322221 | 1303.2305020579881 | 1485.1503336637666 | 1159.7087728396718 | 858.7701810060795 | 1402.6730328140252 | 1260.4995798225532 |
| SE_168_3_34 | SE_168 | SE_168_3 | GCTCTGGCCCTCCAATCAGG | 27.094421042207472 | 37.97691441608806 | 40.488601368016305 | 14.314393385230405 | 20.15129574185873 | 46.78074593289407 | 27.980175931880833 | 10.398888396118036 | 38.1833669464823 | 4.374041668961356 | 7.042709788187574 | 0.0 |
| SE_168_3_40 | SE_168 | SE_168_3 | GATCCATGATACCAGCAGCA | 241.68223569649066 | 251.84690612774185 | 243.91913507073238 | 229.03029416368648 | 310.7541922297162 | 316.2031901019692 | 215.2321225529295 | 298.7317030157545 | 203.34862862196388 | 343.36227101346645 | 419.0412323971607 | 146.24580760372163 |
| SE_168_4_16 | SE_168 | SE_168_4 | GGGGGGGCACCTACAATAAT | 499.6211240183058 | 712.5668415439679 | 648.8051487508955 | 602.5058306692434 | 686.2046497359262 | 947.7432601960392 | 592.9644976333208 | 734.5396621621558 | 749.4595744844434 | 963.7471810611521 | 770.0029368418415 | 681.3197544713063 |
| SE_168_4_44 | SE_168 | SE_168_4 | CTTGGGGCTGTCCCCACCCA | 623.1716839707718 | 840.4890795771067 | 761.3832110912334 | 732.6366796258834 | 650.1444363031263 | 509.39034460262434 | 670.4480617523753 | 634.3321921632001 | 797.4107794870025 | 1077.4722644541473 | 497.6848250319219 | 665.070220293115 |
| SE_168_4_50 | SE_168 | SE_168_4 | TCAGGGGCCCACAGGACCGT | 2246.669392819844 | 2240.6379505491955 | 2286.1246869989695 | 2307.219952001228 | 2058.6139492077787 | 2053.1549603881285 | 2159.8543498186473 | 2052.3624279974774 | 1953.5676112153735 | 1502.4833132882259 | 2007.1722896334586 | 2332.9688355831786 |
| SE_168_4_6 | SE_168 | SE_168_4 | GGACTGGAGCATCTGCCAAG | 1020.9177848703775 | 1421.1361131494004 | 1297.610297501791 | 1270.0770858168069 | 1268.4710372243705 | 1270.8769311769556 | 1155.7964981092314 | 1245.9759005530518 | 1428.7683120206982 | 1490.0901952261686 | 985.9793703462603 | 1407.9060684390026 |
| SE_168_4_8 | SE_168 | SE_168_4 | GGCAGATGCTCCAGTCCCGT | 2370.2199527723096 | 2340.577199012585 | 2304.8876973890256 | 2330.643504813423 | 2107.4012967933313 | 2506.2351478493065 | 2321.2784417333446 | 2206.455046958136 | 2423.3118231848885 | 2087.8758899842205 | 2254.8409171847215 | 2329.4867925449944 |
| SE_168_5_18 | SE_168 | SE_168_5 | GGAAAGGAGGTGCCACCTCG | 1549.8008836142674 | 1505.0850818586478 | 1386.4877151388998 | 1445.753731908271 | 1411.6512964428405 | 1416.4170296348482 | 1474.340039487567 | 1544.7076035688062 | 1832.8016134311504 | 1390.9452507297112 | 1207.824728674169 | 1510.0459975590622 |
| SE_168_5_21 | SE_168 | SE_168_5 | TCAGGCCAGAAGCCCCTACA | 600.4123702953176 | 831.4945472154017 | 794.9591244208079 | 664.9686381684306 | 861.2027443362783 | 838.5881863526197 | 684.4381497183158 | 886.7415741417016 | 893.3131894921208 | 899.5945699163856 | 820.4756903238524 | 1244.2500456443618 |
| SE_168_5_32 | SE_168 | SE_168_5 | GGACACTGAAGATTCAGGAA | 368.48412617402164 | 481.70717759353795 | 521.4141834710392 | 514.0168533787282 | 615.144817383056 | 452.21387735130935 | 487.5007575823853 | 835.6924856516675 | 616.2617828106678 | 1139.4378547644333 | 765.3077969830497 | 648.8206861149238 |
| SE_168_5_33 | SE_168 | SE_168_5 | GGAGCCTCTGAACTTCTGTG | 165.81785677830973 | 264.8390084279825 | 265.64472604869235 | 200.40150739322567 | 229.08841474955184 | 409.7646819677573 | 148.51016456152135 | 358.2889729207941 | 272.61148029232714 | 324.40809044796725 | 346.26656458588906 | 177.58419494737626 |
| SE_168_5_38 | SE_168 | SE_168_5 | GAGAGAGCCGGAACAGGTGG | 2446.0843316904907 | 3010.1701637172955 | 3030.7199414254155 | 2616.9313725180314 | 2915.5743154931392 | 2600.662949824963 | 2689.325371298854 | 3028.912583742017 | 3103.508545998968 | 3855.717731189435 | 2493.119265018401 | 2594.122063446967 |
| SE_168_6_2 | SE_168 | SE_168_6 | GATACCACTCAGACAACTTT | 290.4521935724641 | 309.81167023650784 | 311.0709617298814 | 273.2747828089441 | 307.572408691528 | 202.71656570920763 | 255.05006522522146 | 288.33281461963645 | 360.5220227970189 | 198.2898889929148 | 404.9558128207855 | 271.59935697834015 |
| SE_168_6_28 | SE_168 | SE_168_6 | CATTTAATTAGCAGTCCGGT | 179.90695572025763 | 238.8548038275012 | 224.1685978180415 | 199.10019890365928 | 218.48246962225778 | 252.09624197170695 | 184.02346478275473 | 134.24019565897828 | 107.44621861684554 | 314.20199322039076 | 117.3784964697929 | 127.67491140007444 |
| SE_168_6_34 | SE_168 | SE_168_6 | CTAATTAAATGCAGAGCTGG | 74.78060207649263 | 110.93256579436247 | 83.93978332393624 | 85.88636031138243 | 130.45312506571702 | 116.95186483223517 | 85.01668840840715 | 132.3494886778659 | 103.89427750554486 | 32.80531251721017 | 210.1075086809293 | 194.9944101382955 |
| SE_168_6_37 | SE_168 | SE_168_6 | AAATGCAGAGCTGGGGGGAA | 115.96412206064798 | 169.89672238776234 | 147.14150253254707 | 106.70729614444484 | 168.63452752397566 | 159.4010602157872 | 130.21543414452233 | 161.65544688510764 | 168.71720278678225 | 104.24799311024564 | 240.62591776307545 | 121.87150633643469 |
| SE_168_6_8 | SE_168 | SE_168_6 | AAGCTGAAGCAGGAAGGGGA | 1027.4204459205073 | 1109.3256579436247 | 1008.2649267498695 | 1112.6187585792725 | 1061.6551072421362 | 1341.9143601861651 | 1164.4057830113486 | 1027.599244234573 | 982.1117172746377 | 1089.8653825162046 | 1196.0868790271898 | 973.8113696787495 |
| SE_169_1_0 | SE_169 | SE_169_1 | TAGCAATGAAAACTGTCAGA | 603.6637008203825 | 806.5097350995542 | 851.2481555909769 | 610.3136816066418 | 825.1425309034785 | 1044.769992501301 | 655.3818131736704 | 667.4195643326666 | 786.7549561531005 | 1090.5943894610314 | 841.6038196884151 | 1015.5958861369558 |
| SE_169_1_10 | SE_169 | SE_169_1 | TGGCCACTGTGATTAACCAG | 73.69682523480432 | 47.97083926242702 | 62.21419234597627 | 63.76411598875362 | 68.93864332741144 | 41.582885273683615 | 47.35106696164449 | 43.48626056558451 | 100.34233639424419 | 10.206097227576498 | 197.19587406925208 | 88.21175696732416 |
| SE_169_1_11 | SE_169 | SE_169_1 | AGTCCACTGGTTAATCACAG | 125.71811363584267 | 192.88274953434197 | 206.3931142906197 | 114.51514708184324 | 205.7553354695049 | 23.390372966447035 | 117.30150679134657 | 188.1253446206808 | 221.1083341784673 | 20.412194455152996 | 180.76288456348107 | 60.35541266185337 |
| SE_169_1_15 | SE_169 | SE_169_1 | TTAAAGCTAAACAAGCGACA | 101.8750231187001 | 140.91434033337936 | 114.55311606560711 | 114.51514708184324 | 151.66501532030517 | 225.24062856578627 | 75.33124289352533 | 123.84130726286024 | 262.8436422362503 | 86.75182643440023 | 204.23858385743966 | 162.4953417819129 |
| SE_169_1_8 | SE_169 | SE_169_1 | TAGAACATAGACCCAATGGG | 411.83519984155356 | 637.6124051964258 | 623.1294503223972 | 528.3312467639586 | 589.6905490775501 | 615.9464881164386 | 539.1564669950884 | 497.25593603255334 | 679.3087375362549 | 556.2322989029191 | 529.377019078766 | 789.2630886550056 |
| SE_169_2_13 | SE_169 | SE_169_2 | TCCCCAGGCTTTAGGCAGCA | 332.71949039830776 | 345.78979968332806 | 301.1956931035359 | 402.10432327601774 | 381.81402458258646 | 582.1603938315707 | 295.9441685102781 | 450.9336149953003 | 437.7767419678087 | 407.514882158233 | 413.172307573671 | 450.3442329384444 |
| SE_169_2_17 | SE_169 | SE_169_2 | TGTGAAAATGGACCATACTG | 2870.924853632304 | 2665.379756518601 | 2756.1874736130126 | 2675.4902545485193 | 3014.209605176974 | 2733.2083966348296 | 3082.1239949579503 | 2749.087950537386 | 2815.801315983613 | 2378.020654025324 | 3342.9395794597017 | 3094.375579932713 |
| SE_169_2_3 | SE_169 | SE_169_2 | CGATGGCATTCAAATTCCAG | 4312.3480530777415 | 5092.904101694335 | 5202.291512358778 | 4866.893750978337 | 4908.431404911695 | 5451.689521401896 | 4784.610084351622 | 5604.055492017064 | 5160.970434719887 | 6452.440468662827 | 5017.930724083647 | 5046.641043341124 |
| SE_169_2_5 | SE_169 | SE_169_2 | CATTCAAATTCCAGAGGAGG | 68.27794102636283 | 134.91798542557598 | 135.29118018093254 | 110.61122161314404 | 121.96836896388177 | 240.83421054341764 | 48.42722757440914 | 128.56807471564116 | 135.86174750725098 | 174.23265981362735 | 88.03387235234467 | 270.4386759656122 |

| | | | | | | | | | | | | | | | |
|---|---|---|---|---|---|---|---|---|---|---|---|---|---|---|---|
| SE_169_2_9 | SE_169 | SE_169_2 | TCTATTCTGGAAATCTAGGT | 29.26197472558407 | 27.982989569749094 | 37.526020780112674 | 23.423552812195208 | 27.575457330964575 | 25.989303296052263 | 29.056336544645482 | 13.23494886778659 | 2.6639558334755096 | 88.20984032405401 | 65.73195802308402 | 78.92630886550056 |
| SE_17_1_10 | SE_17 | SE_17_1 | AATTGACACTCACTATGCGG | 84.53459365168732 | 85.94775367851507 | 88.87741763710896 | 114.51514708184324 | 60.45388722557618 | 93.56149186578814 | 80.71204595734856 | 57.66656292392729 | 80.80666028209045 | 102.78997922059186 | 73.94845277596953 | 138.12104051462597 |
| SE_17_1_13 | SE_17 | SE_17_1 | GGCCATGTAACAGAGCACTG | 1085.9443953716755 | 1326.1938271091803 | 1158.36900987032 | 1145.1514708184325 | 1152.8662353368652 | 1126.2031428289313 | 1240.8131865176385 | 1141.0416631013154 | 1254.723197566965 | 1013.3196533093808 | 942.549326652437 | 1341.7472507135096 |
| SE_17_1_15 | SE_17 | SE_17_1 | ACTTGGCTGCTACATTCAGC | 2611.9021884688004 | 2309.596031988934 | 2365.126836009733 | 2446.459960384833 | 2192.2488578116836 | 1997.7111133565506 | 2314.821478056757 | 2141.2256561097593 | 2473.926984020923 | 2440.7152512804364 | 1964.9160309043332 | 1304.6054583062153 |
| SE_17_1_20 | SE_17 | SE_17_1 | GTAGTCCTCTCCTTAAAACC | 137.63965889441397 | 208.8730292884843 | 201.45547997744697 | 108.00860463401123 | 177.11928362581094 | 271.1550643888119 | 135.59623720834557 | 89.80858160283758 | 165.1652616754816 | 126.11820145505243 | 116.20471150509498 | 109.10401519642724 |
| SE_17_1_23 | SE_17 | SE_17_1 | CTTAGAAGTCTTCATTTGAA | 48.769957875973454 | 109.93317330972857 | 103.69032057662713 | 123.62430650880805 | 167.57393301124625 | 63.24063802039384 | 94.70213392328898 | 105.87959094229272 | 52.391131391685015 | 131.22125006884067 | 53.994108376104734 | 51.069964560029774 |
| SE_17_2_15 | SE_17 | SE_17_2 | GTCTTGCCCGATGCTCCCGG | 412.9189766832419 | 433.73633833111097 | 476.97547465248476 | 467.1697477543378 | 440.1467227827038 | 485.9999716361773 | 383.1131781442145 | 418.79159631639 | 517.695416972074 | 410.43090993754055 | 471.86155580856746 | 392.3101823020469 |
| SE_17_2_17 | SE_17 | SE_17_2 | TGTCCTCCGACTTCTTCTGA | 60.69150313454474 | 90.94471610168455 | 72.08946097232172 | 72.87327541571842 | 138.9378811675523 | 92.69518175591973 | 43.0464245105859 | 169.21827480955713 | 50.615160836034676 | 25.515243068941242 | 130.29013108147012 | 85.89039494186825 |
| SE_17_2_19 | SE_17 | SE_17_2 | CATAAGGCGTGAATTCCCCG | 3474.5885544526864 | 3097.1173098804443 | 3327.9655270784133 | 3259.777766363833 | 3109.663111322621 | 2850.1602614670646 | 3245.700408098177 | 3323.8628727955465 | 3341.488600456114 | 2727.2149805974054 | 3450.9277962119113 | 2836.704395107108 |
| SE_17_2_2 | SE_17 | SE_17_2 | AATTGCAGAAGTCAACGTGC | 460.60515771752705 | 537.6731567330362 | 554.9900968006137 | 404.7069402551505 | 579.0846039502561 | 444.4170863624937 | 422.93112081650645 | 525.6165407492389 | 565.6466219746331 | 312.0149723859101 | 495.33725510252606 | 619.803660796725 |
| SE_17_2_9 | SE_17 | SE_17_2 | ACTGAGATAGCCAAACGCCT | 956.9749512107679 | 1018.3809418419402 | 1109.9801936012275 | 1132.1383859227683 | 1056.3521346784892 | 1072.49191601709 | 1079.3890946029414 | 1081.4843931962757 | 1156.1568317283711 | 1724.1014245156011 | 1102.1840818513554 | 1095.6828760151843 |
| SE_17_3_11 | SE_17 | SE_17_3 | AGAGTCATACTGCTGCTCAC | 1439.255645762061 | 1444.12214029598 | 1427.9638433695507 | 1453.5615828456694 | 1426.499619621052 | 1408.6202386460327 | 1394.704154142983 | 1288.5168076280802 | 1440.3121206324254 | 1261.1820145505244 | 1301.7275258500033 | 2230.828906463119 |
| SE_17_3_20 | SE_17 | SE_17_3 | GGATCACAGCTATTGAACTG | 1101.1172711553118 | 1092.3359857048486 | 1121.830515952842 | 1093.0991312357764 | 1097.715320674936 | 1216.299394255246 | 1177.3197103645243 | 983.1676301784324 | 1137.5091408940425 | 1123.3997019782416 | 1024.714274181292 | 879.7962076477856 |
| SE_17_3_23 | SE_17 | SE_17_3 | GATGCATGTCAAAGAGGGGA | 1554.1359909810205 | 1560.051668513512 | 1701.5087843193194 | 1663.0722496658598 | 1525.1349093048868 | 1786.3314465486587 | 1684.1913589766732 | 1795.2262785661953 | 1603.7014117522567 | 1571.0099661019538 | 1354.5478492614102 | 1492.635782368143 |
| SE_17_3_24 | SE_17 | SE_17_3 | GGATGCATGTCAAAGAGGGG | 995.9909175115467 | 1155.297712236784 | 1125.78062340338 | 1054.0598765487844 | 1212.259528049712 | 1104.545390082221 | 994.3724061945343 | 960.479146405084 | 1187.236316452252 | 1040.292910267976 | 764.1340120183518 | 834.5296481513957 |
| SE_17_3_25 | SE_17 | SE_17_3 | ACAGGATGCATGTCAAAGAG | 2023.411363432054 | 2180.6744014711617 | 2083.681680158888 | 2240.8532190333417 | 2031.0384918768139 | 2376.288631369045 | 1877.90026927431 | 1998.4772790357752 | 1833.6895987089756 | 1923.1203204533429 | 2028.3004189980213 | 2206.454605195832 |
| SE_17_4_13 | SE_17 | SE_17_4 | CTTTGGCCTCACATTCTTGG | 67.19416418467453 | 94.94228604022014 | 72.08946097232172 | 68.96934994701923 | 81.66577748016432 | 17.326202197368175 | 111.92070372752333 | 41.59555358447214 | 119.87801250639792 | 91.12586810336158 | 172.54638981059557 | 38.30247342002233 |
| SE_17_4_15 | SE_17 | SE_17_4 | TTATCTCCACGCCAAGCAAC | 159.31519572817993 | 129.9210230024065 | 158.9918248841616 | 182.18318853929605 | 160.14977142214042 | 226.10693867565467 | 207.69899826357695 | 112.49706537618601 | 61.27098416993672 | 276.2936320893923 | 65.73195802308402 | 114.907420260067 |
| SE_17_4_2 | SE_17 | SE_17_4 | TTAAAACCTGCACAGAGGGC | 32.51330525064897 | 58.96415659339988 | 34.56344019220904 | 35.13532921829281 | 29.69664635642339 | 77.96790988815678 | 38.74178205952731 | 9.45353490556185 | 20.423661389978903 | 62.69459725511277 | 59.86303319959438 | 1.1606810127279494 |
| SE_17_4_23 | SE_17 | SE_17_4 | GTTAGGTTATATGGCAAGAG | 171.23674098675122 | 174.89368481093183 | 185.6550501752943 | 171.77272062276487 | 183.48285070218736 | 147.27271867762948 | 228.14604990610528 | 160.71009339455145 | 176.7090702872088 | 126.84720839987932 | 227.71428315139823 | 152.0492126673614 |
| SE_17_4_26 | SE_17 | SE_17_4 | CTTCCAGTCCAAACAACAGG | 160.39897256986825 | 169.89672238776234 | 157.0167711588925 | 121.02168952967524 | 139.9984756802817 | 225.24062856578627 | 131.291594757287 | 112.49706537618601 | 311.6828325166346 | 121.74415978609107 | 116.20471150509498 | 201.9584962146632 |
| SE_170_1_15 | SE_170 | SE_170_1 | CCACATTTCACTCCTCTAAA | 172.32051782843953 | 160.9021900260573 | 174.7922546863143 | 174.37533760189766 | 181.36166167672854 | 275.486614938154 | 192.6327496848719 | 256.19079594072616 | 287.70723001535504 | 180.79372231706938 | 152.59204541073078 | 492.12874939665056 |
| SE_170_1_23 | SE_170 | SE_170_1 | GCACCCAGAACTGGAGCACG | 1619.1626014823187 | 1666.9866643693388 | 1584.980614528443 | 1604.5133676353717 | 1702.2541929306979 | 1706.6309164407653 | 1657.2873436575571 | 1783.8820366795212 | 1441.2001059102506 | 1864.7997648671915 | 1193.739309097794 | 1860.571663402903 |
| SE_170_1_25 | SE_170 | SE_170_1 | GCAGACATGGACACAGGAAG | 5390.706010557598 | 4614.195101554698 | 4621.625717129666 | 5285.9150846187185 | 4872.371191478895 | 4226.727026047966 | 5393.716991176413 | 5021.7177418344545 | 4963.837703042699 | 5039.625009588309 | 4808.997000367415 | 5137.174162333904 |
| SE_170_1_35 | SE_170 | SE_170_1 | GCTGCAGTCATCACTGGGCA | 1380.7316963108929 | 341.7922297447925 | 620.1668697344936 | 1141.2475453497332 | 376.5110520189394 | 3077.1335102525877 | 1714.3238561340834 | 603.135526974846 | 2069.0056973326455 | 108.622034779207 | 367.39469395045177 | 896.045741825977 |
| SE_170_1_8 | SE_170 | SE_170_1 | CATCTATGATCTCAGGGGCA | 324.04927566480137 | 364.7782568913721 | 405.87354054279757 | 318.8205799347681 | 380.753430069857 | 427.9571947749939 | 333.60978995704073 | 437.69866612751366 | 464.4163003025638 | 400.95381965479095 | 487.12076034964053 | 278.5634430547079 |
| SE_170_2_12 | SE_170 | SE_170_2 | GCAGGCAGAGGGAGTACCAA | 12992.316778159327 | 12008.700095360897 | 11711.081063983058 | 12677.347305355874 | 12166.079655519025 | 12138.737259476144 | 12153.081799951164 | 11652.427124595537 | 12260.41273093212 | 10262.959769272995 | 10474.857024964318 | 12057.154360217939 |
| SE_170_2_14 | SE_170 | SE_170_2 | GCGAGAGCTTTGCAGGCAGA | 7170.267584609785 | 7488.4478873617845 | 7241.534483699112 | 7607.449430005177 | 7902.48971434681 | 8284.523580671592 | 8064.747632058268 | 7943.805381143623 | 7596.714051794327 | 8552.709476709104 | 7629.6022705365385 | 9021.973511934351 |
| SE_170_2_16 | SE_170 | SE_170_2 | GTATATTAAACAGATCCTAG | 9.75399157519469 | 20.98724217731182 | 15.800429802152705 | 19.519627343496005 | 4.242378050917627 | 0.0 | 4.30464245105859 | 27.415251226129364 | 7.103882222601358 | 33.53431946203706 | 11.73784964697929 | 46.42724050911798 |
| SE_170_2_25 | SE_170 | SE_170_2 | TTATGAAACACTAGCAAGCG | 180.99073256194592 | 189.88457208044028 | 201.45547997744697 | 161.36225270623365 | 224.84603669863424 | 181.92512307236584 | 126.9869523062284 | 116.27847933841076 | 158.94936473070538 | 159.6525209170895 | 105.64064682281361 | 453.8262759766282 |
| SE_170_2_5 | SE_170 | SE_170_2 | TCTAGGAAGTAAAAGAGAGA | 86.70214733506391 | 118.92770567143364 | 142.20386821937433 | 137.93869989403845 | 161.21036593486983 | 149.8716490072347 | 106.5399006637001 | 112.49706537618601 | 157.17339417505505 | 131.22125006884067 | 152.59204541073078 | 160.17397975645702 |
| SE_170_3_0 | SE_170 | SE_170_3 | ATCGACATATAGACACTCTG | 1766.5562519519272 | 1857.870628934413 | 1866.4257703792882 | 1970.1810532035302 | 1929.2214186547908 | 1770.7378645710273 | 1714.3238561340834 | 1888.8162741312576 | 2006.8467278848836 | 1768.5708481500417 | 1725.4638981059556 | 1851.2862153010794 |
| SE_170_3_18 | SE_170 | SE_170_3 | AGGCCCTCTTTTGACCTAAG | 54.188842084414944 | 100.93864094802352 | 81.96472959866715 | 59.86019052005442 | 117.72599091296415 | 177.5935725230238 | 38.74178205952731 | 76.57363273505099 | 228.21221640106864 | 101.33196533093808 | 75.12223774066746 | 97.49720506914775 |
| SE_170_3_2 | SE_170 | SE_170_3 | GAGGCACCCACATATCCACA | 9838.526168846378 | 9785.051817050478 | 9387.430356203975 | 9451.403559720766 | 9734.136437830495 | 8747.999489451191 | 9793.061576158292 | 9915.812762443824 | 9293.653917718228 | 9034.58306723968 | 9698.985163298988 | 9294.733549925419 |
| SE_170_3_29 | SE_170 | SE_170_3 | GCAGGTCATGACTCAGGGGC | 2842.746655748408 | 3614.8026169208024 | 3690.387885665291 | 3154.3717787089545 | 3778.8982488548763 | 2991.3688093756155 | 2595.6993979883296 | 3263.3602493999506 | 3643.4035949166714 | 4078.7938563064645 | 3355.851214071379 | 3086.2508128436175 |
| SE_170_3_6 | SE_170 | SE_170_3 | GCAAAGCCAGGTCACTGTTG | 1153.13855955635 | 1242.244858399933 | 1155.4064292824164 | 1212.8195122758852 | 1124.2301834931711 | 997.9892465684069 | 1224.6707773261687 | 1284.7353936658553 | 1108.205626725812 | 1306.3804451297917 | 973.0677357345832 | 1012.1138430987719 |
| SE_170_4_16 | SE_170 | SE_170_4 | TCTGTGGTTTTGGTTACCCA | 492.0346861264877 | 403.7545637920941 | 418.7113897570467 | 415.11740817168175 | 356.3597562770807 | 421.89302350591504 | 520.8617365780893 | 401.7752334863786 | 386.27359585394885 | 266.0875348618158 | 341.57142472709734 | 366.77520002203204 |
| SE_170_4_21 | SE_170 | SE_170_4 | TATTTACCAGGTAGAAGCCA | 213.50403781259487 | 254.84508358164354 | 253.79440369707783 | 271.9734743193777 | 294.8452745387751 | 304.0748485638115 | 189.40426784657797 | 253.35473546905757 | 206.0125844554394 | 388.5607015927338 | 289.9248862803885 | 244.90369368559732 |
| SE_170_4_27 | SE_170 | SE_170_4 | TAGTGGCCCCCAAAGTTCAA | 1304.8673173927118 | 1558.0528835442442 | 1563.2550235504832 | 1413.2210196691108 | 1638.6185221669334 | 1478.7913575453738 | 1253.7271138708143 | 1557.942552436593 | 1812.3779520411715 | 1667.9678897639303 | 1262.9926220149716 | 1454.3333089481207 |
| SE_170_4_3 | SE_170 | SE_170_4 | GTTGAACTCTATTATCACCA | 268.77665673869814 | 320.8049875674807 | 326.87139153203407 | 300.6022610898385 | 299.0876525896927 | 523.2513063605189 | 306.70577463792455 | 302.5131169779792 | 388.0495664095992 | 336.8012085100244 | 228.88806811609615 | 329.6334076147376 |
| SE_170_4_5 | SE_170 | SE_170_4 | TCCAAATGCCTCCTGCCCGG | 864.8539196672625 | 936.4307581019607 | 1057.6412698815966 | 864.0688370720899 | 1077.5640249330772 | 1146.994585465773 | 803.8919777351916 | 872.5612717833587 | 1163.2607139509723 | 1101.5294936334349 | 1190.2179542037 | 1174.6091848806848 |
| SE_171_1_12 | SE_171 | SE_171_1 | TCTCTCCATTCTAGTCCCAA | 41.18351998415536 | 28.982382054382988 | 38.51354764274722 | 37.737946197425615 | 16.969512203670508 | 58.04277736118338 | 33.36097899570407 | 14.180302358342775 | 74.59076333731426 | 30.61829168272949 | 42.25625872912545 | 19.73157721637514 |
| SE_171_1_13 | SE_171 | SE_171_1 | TATGTTCTATCCATTAACAG | 105.126353643765 | 158.9034050567895 | 118.50322351614528 | 102.80337067574564 | 147.42263726938754 | 94.42780197565655 | 119.45382801687587 | 264.6989773557318 | 77.25471917078977 | 126.11820145505243 | 157.2871852695225 | 8.124767089095647 |
| SE_171_1_24 | SE_171 | SE_171_1 | TATATGCTAGAGACTGATGG | 252.52000411337366 | 194.88153450360974 | 199.4804262521779 | 178.27926307059687 | 221.664253160446 | 116.08555472236677 | 241.05997725928103 | 80.35504669727572 | 328.5545527953128 | 193.91584732395344 | 79.81737759945918 | 321.50864052564197 |
| SE_171_1_7 | SE_171 | SE_171_1 | AAGAAACGTGACCAGCTGAA | 863.7701428255742 | 976.4064574873165 | 914.4498747995877 | 918.7237936338787 | 991.6558694019953 | 803.0694718480149 | 901.8225934967746 | 1136.3148956485343 | 1037.1668044997982 | 785.1404795785634 | 1039.9734787223651 | 766.0494684004466 |
| SE_171_1_8 | SE_171 | SE_171_1 | TCATCTGGATCATCAAGCAG | 901.7023322846646 | 861.4763217544186 | 912.4748210743187 | 990.2957605600308 | 826.2031254162079 | 1036.1068914026168 | 1061.0943641859424 | 795.9876390483078 | 840.9220581004358 | 1024.983764426611 | 662.014720089632 | 806.6733038459248 |
| SE_171_2_10 | SE_171 | SE_171_2 | GTGTAAGAGTCTGATCAAAG | 294.7873009392173 | 356.78311701430096 | 323.90881094413044 | 291.4931016628737 | 324.5419208951985 | 273.7539947184172 | 246.44078032310426 | 336.5458426380019 | 418.241065855655 | 210.68300705497197 | 284.05596145689884 | 437.57674179843696 |
| SE_171_2_11 | SE_171 | SE_171_2 | GGGAAGCATGACACTCAAAG | 861.6025891421976 | 1078.344490919974 | 905.5621330358769 | 1008.5140794139603 | 1084.988186522183 | 1500.449110292084 | 865.2331326627766 | 1126.8613607429725 | 1190.7882575635526 | 1094.239424185166 | 1225.4315031446379 | 1221.0364253898028 |
| SE_171_2_14 | SE_171 | SE_171_2 | CACCTTCCCTAATCACCCCA | 1301.615986867647 | 1333.1895745016175 | 1324.2735227929236 | 1310.4176489933652 | 1407.4089183919227 | 1915.4116529190517 | 1120.283197887998 | 1591.0299246060595 | 1203.220051453105 | 1240.0408131505444 | 1444.9292915431506 | 1475.2255671772236 |
| SE_171_2_18 | SE_171 | SE_171_2 | TTTTCTGGTACTCAGCCCTG | 3.2513305250648967 | 7.9951398770711695 | 2.962580587903632 | 16.917010364363207 | 6.363567076376441 | 82.29946043749882 | 13.990087965940416 | 0.0 | 5.327911666951019 | 12.393118062057175 | 1.173784964697929 | 1.1606810127279494 |
| SE_171_2_9 | SE_171 | SE_171_2 | GTCAAAATATCAGAAACACA | 338.13837460674927 | 388.7636765225856 | 408.83612113070126 | 359.1611431203265 | 396.6623477607981 | 450.48125713157253 | 308.8580958634538 | 287.38746112908024 | 405.8092719661026 | 145.07238202055163 | 151.41826044603283 | 842.6544152404913 |
| SE_172_1_19 | SE_172 | SE_172_1 | TAAGCCTGAAAATTACACAT | 2552.294462175944 | 2719.3469506888314 | 2599.1707024541197 | 2675.4902545485193 | 2814.8178367838455 | 2933.326032014432 | 2520.3681550948045 | 3143.300356099315 | 2631.988363473803 | 3074.222286335006 | 3740.8526824922997 | 3927.744547071381 |
| SE_172_1_25 | SE_172 | SE_172_1 | AGACCAAAGAAGAATCAAAA | 187.49339361207572 | 280.82928818212486 | 331.8090258452068 | 235.53683661151848 | 329.8448934588455 | 273.7539947184172 | 146.35784333599204 | 493.4745220703286 | 352.5301552965924 | 199.74790288256858 | 537.5935138316515 | 401.59563040387053 |
| SE_172_1_30 | SE_172 | SE_172_1 | TCTGGATTTCCCAAGTGCAA | 270.9442104220747 | 336.795267321623 | 307.1208542793432 | 383.88600442208815 | 347.8750001752454 | 349.1229742769687 | 363.74228711445085 | 355.45291244912556 | 288.59521529318016 | 394.3927571513489 | 211.28129364562722 | 324.9906835638258 |
| SE_172_1_33 | SE_172 | SE_172_1 | AGTGTTGGTCAATACTAGAG | 177.73940203688102 | 249.84812115847404 | 260.7070917355196 | 221.22244322628808 | 265.1486281823517 | 280.68447559736444 | 148.51016456152135 | 219.3220098090349 | 333.88246446226384 | 435.94615300648184 | 245.32105762186717 | 178.74487596010422 |
| SE_172_1_5 | SE_172 | SE_172_1 | TCCGCTTGTTCATCTTTGGG | 531.0506524272664 | 569.6537162413208 | 623.1294503223972 | 628.5320004605715 | 667.1139485067969 | 816.064123496041 | 462.7490634887984 | 678.7638062193408 | 734.3638247614155 | 613.0948405994167 | 885.0338633822385 | 438.7374228111649 |
| SE_172_2_18 | SE_172 | SE_172_2 | TTATGTCTCTGTGGGACCGG | 154.98008836142674 | 214.86938419628768 | 183.6799964500252 | 212.11328379932328 | 121.96836896388177 | 223.50800834604945 | 179.71882233169612 | 135.18554914953447 | 222.88430473411762 | 89.6678542137078 | 131.46391604616804 | 76.60494684004466 |
| SE_172_2_19 | SE_172 | SE_172_2 | AAGGATGTGTGCACTATGAG | 1999.5682729149114 | 1625.012180014715 | 1700.521257456685 | 2138.049848357596 | 2005.5842235713083 | 2094.737845661812 | 2324.5069235716387 | 2461.7004894083057 | 1801.7221287072696 | 3993.500043761718 | 2327.6155849959932 | 2797.2412406743583 |
| SE_172_2_2 | SE_172 | SE_172_2 | AGCACCTTCTAACTTTGAGA | 114.88034521895969 | 88.94593113241676 | 102.70279371399258 | 96.29682822791364 | 128.33193604025823 | 112.62031428289313 | 132.36775537005164 | 120.0598933006355 | 93.23845417164283 | 150.17543063433988 | 98.59793703462604 | 95.17584304369186 |
| SE_172_2_5 | SE_172 | SE_172_2 | CTAGCCTGTAAAATGGACTG | 675.1929723718102 | 701.5735242129952 | 723.8571903111208 | 707.9118183241219 | 672.4169210704439 | 615.0801780065702 | 633.8586009183773 | 660.8020898987734 | 597.6140919763393 | 850.0220976681568 | 705.4447637834553 | 595.4293595294381 |
| SE_172_2_7 | SE_172 | SE_172_2 | GGAAGGAGAGGGATACGGAG | 2313.863557004518 | 2503.4781740079097 | 2438.2038238446894 | 2447.7612688743993 | 2416.0342999975887 | 2394.4811436762816 | 2390.152720950282 | 2399.3071590315976 | 2456.1672784644197 | 3225.855730859 | 2281.837971372774 | 3311.4229293128396 |
| SE_172_3_13 | SE_172 | SE_172_3 | TTTACTCCTTAGCTGGACAG | 21.675536833765978 | 25.984204600481302 | 21.725590977959968 | 15.615701874796805 | 27.575457330964575 | 1.7326202197368175 | 8.60928490211718 | 23.633837263904624 | 0.8879852778251698 | 4.374041668961356 | 46.95139858791716 | 0.0 |
| SE_172_3_18 | SE_172 | SE_172_3 | CCACTTCAGCCTTAGTCATG | 337.054597765061 | 386.7648915533178 | 363.4098854495122 | 273.2747828089441 | 337.2690550479513 | 317.935810321706 | 280.877919931573 | 239.1744331107148 | 231.7641575123693 | 285.041715427315 | 179.58909959878315 | 401.59563040387053 |
| SE_172_3_25 | SE_172 | SE_172_3 | GGTGTCCCTAAACTTGAAAG | 66.11038734298623 | 83.94896870924728 | 95.79010567555078 | 68.96934994701923 | 71.05983235287025 | 85.76470087697247 | 76.40740350628997 | 83.19110716894428 | 102.11830694989453 | 52.48850002753627 | 58.68924823489645 | 116.06810127279495 |
| SE_172_3_3 | SE_172 | SE_172_3 | TCAAACCTATTGGATCGGAG | 351.14369670700887 | 309.81167023650784 | 322.9212840814959 | 365.6676855681585 | 374.38986299348056 | 337.86094284867943 | 330.3813081187468 | 511.4362383908961 | 307.2429061275087 | 225.99215289633673 | 355.6568443034725 | 554.8055240839599 |
| SE_172_3_8 | SE_172 | SE_172_3 | ACCAGCGGCTTCAAACCTAT | 1507.5335867884237 | 1978.7971195751145 | 2043.1930787908716 | 1735.9455250815781 | 1603.618903246863 | 1650.3207592993185 | 1547.5189611555631 | 1747.0132505478298 | 1817.7058637081225 | 2725.0279597629246 | 1518.8777443191202 | 1576.2048152845553 |
| SE_172_4_13 | SE_172 | SE_172_4 | TTTCAGAGTCTCTTGCCTGG | 7.586437891818092 | 12.992102300240651 | 14.81290293951816 | 13.013084895664004 | 28.63605184369398 | 1.7326202197368175 | 2.152321225529295 | 12.289595377230405 | 15.095749723027886 | 0.7290069448268927 | 51.646538446708874 | 15.088853165463343 |
| SE_172_4_27 | SE_172 | SE_172_4 | GGGTTGGAATCCAGAAAAGA | 70.44549470973944 | 63.961119016569356 | 59.25161175807264 | 76.77720088441762 | 88.02934455654076 | 94.42780197565655 | 74.25508228076067 | 87.9178746217252 | 86.13457194904147 | 99.87395144128429 | 102.11929192871983 | 6.964086076367696 |
| SE_172_4_34 | SE_172 | SE_172_4 | AGGCCTTCCCAGGAAAGCCA | 308.8763998811652 | 382.76732161478225 | 420.6864434823158 | 348.7506752037953 | 405.1471038626334 | 464.34221888946706 | 320.69586260386495 | 333.7097821663333 | 485.7279469703679 | 400.22481270996406 | 235.93077790428373 | 472.39717218027545 |
| SE_172_4_6 | SE_172 | SE_172_4 | GATGTTCTTCGAGAAGATGG | 136.55588205272565 | 129.9210230024065 | 148.12902939518162 | 180.88188004972966 | 158.0285823966816 | 101.35828285460381 | 163.5764131402264 | 95.48070254617468 | 141.189659174202 | 201.93492371704926 | 125.5949912226784 | 60.35541266185337 |
| SE_172_4_9 | SE_172 | SE_172_4 | AGCCTTCTCTCTTAATTGGT | 3.2513305250648967 | 14.990887269508443 | 24.688171565863602 | 6.506542447832002 | 13.787728665482287 | 37.251334724341575 | 3.228481838293942 | 9.45353490556185 | 18.647690834328564 | 0.0 | 9.390279717583432 | 49.909283547301825 |
| SE_172_5_12 | SE_172 | SE_172_5 | GGAGGTTTTCTGTTGTATGG | 1217.0813932159597 | 1943.8183826129282 | 2106.3947979994823 | 1517.3256988344228 | 1857.1009917891913 | 2254.1389058775994 | 1256.9555957091081 | 1974.8434417718704 | 2273.2423112324345 | 2122.8682233359114 | 2219.6273682437836 | 2661.441562185188 |
| SE_172_5_13 | SE_172 | SE_172_5 | GGTTTTCTGTTGTATGGCGG | 34.680858934025565 | 48.970231747060915 | 51.35139685699629 | 37.737946197425615 | 44.54496953463509 | 86.63101098684088 | 8.60928490211718 | 86.97252113116902 | 59.49501361428638 | 22.599215289633673 | 75.12223774066746 | 8.124767089095647 |
| SE_172_5_2 | SE_172 | SE_172_5 | TAGTTGGGAGAGAAGCAGCT | 1426.2503236618013 | 1437.1263929035426 | 1446.726853759607 | 1543.351868625751 | 1375.5910830100406 | 1270.0106210670872 | 1504.4725366449773 | 1430.319831211508 | 1404.7927095194186 | 1907.811174611978 | 1063.4491780163237 | 1425.3162836299218 |
| SE_172_5_22 | SE_172 | SE_172_5 | GCAGCTGTTCTCATTGGCCA | 3506.018082861647 | 3366.9532807315964 | 3478.069610198864 | 3561.681335943238 | 3708.8990110147356 | 3558.801931339423 | 3713.8302746507984 | 3117.7758118542984 | 3870.7278260399153 | 3537.1416963000834 | 3349.9822892478896 | 4020.5990280896167 |
| SE_172_5_8 | SE_172 | SE_172_5 | AGGCACACAGGCTCTCCAGG | 1550.8846604559558 | 1954.811699943901 | 1967.1535103680117 | 1899.9103947669446 | 1923.9184460911438 | 1982.9838414887877 | 1678.81055591285 | 2162.023432901995 | 2016.6145659409606 | 2008.4141329980894 | 2393.3475430190774 | 2193.6871140558246 |
| SE_173_1_10 | SE_173 | SE_173_1 | ATACAGAGCAAATCAAGGCA | 591.7421555618112 | 578.6482486030259 | 552.0275162127101 | 545.2482571283218 | 575.9028204120679 | 568.2994320736761 | 543.461109446147 | 527.5072477303512 | 549.6628869737801 | 489.163659978845 | 504.7275348201095 | 652.3027291531076 |
| SE_173_1_11 | SE_173 | SE_173_1 | ACCTGACTCACCCAAGGGCA | 1596.4032878068642 | 1657.9921320076337 | 1621.5191084459213 | 1693.002344925887 | 1710.7389490325331 | 2127.657629836812 | 1577.651458312973 | 1508.7841709276713 | 1542.43042758232 | 1514.876431350283 | 1144.4403405804808 | 2286.5415950740603 |
| SE_173_1_13 | SE_173 | SE_173_1 | ATTTCAGCAGAGGTGGGGGA | 454.10249666739725 | 752.5425409293239 | 685.3436426683736 | 480.1828326500018 | 655.4474088667733 | 521.518686140782 | 417.5503177526832 | 415.9555358447214 | 570.086548363759 | 489.89266692367187 | 544.636223619839 | 587.3045924403424 |
| SE_173_1_36 | SE_173 | SE_173_1 | GTGAACACACTGGACATGGG | 790.0733175907699 | 916.4429084092828 | 909.5122404864151 | 724.828828688485 | 969.3833846346778 | 572.6309826230182 | 794.2065322203098 | 1045.5609605551406 | 727.259942538814 | 639.3390906131849 | 600.9779019253397 | 791.5844506804615 |
| SE_173_1_6 | SE_173 | SE_173_1 | AATTGTGCTACACGAAACCA | 172.32051782843953 | 130.9204154870404 | 129.36601900512528 | 176.97795458103047 | 80.60518296743491 | 121.28341538157723 | 128.06311291899306 | 161.65544688510764 | 130.53383584029996 | 65.61062503442034 | 109.1620017169074 | 89.37243798005211 |

| | | | | | | | | | | | | | | | |
|---|---|---|---|---|---|---|---|---|---|---|---|---|---|---|---|
| SE_173_2_10 | SE_173 | SE_173_2 | AAACCAGGACTAATATTCAA | 79.11570944324582 | 95.94167852485404 | 89.8649444997435 | 80.68112635311682 | 98.63528968383483 | 144.67378834802426 | 90.39749147223039 | 103.98888396118035 | 98.56636583859384 | 87.48083337922712 | 86.86008738764674 | 62.67677468730927 |
| SE_173_2_11 | SE_173 | SE_173_2 | TTACTGTGTGAATTAATTGA | 68.27794102636283 | 75.95382883217611 | 84.92731018657079 | 46.847105624390416 | 82.72637199289373 | 92.69518175591973 | 31.208657770174778 | 108.71565141396128 | 46.17523444690883 | 4.374041668961356 | 82.16494752885504 | 15.088853165463343 |
| SE_173_2_15 | SE_173 | SE_173_2 | TGTGGCCACCATGGGCCCCA | 24.926867358830876 | 37.97691441608806 | 48.38881626909266 | 13.013084895664004 | 28.63605184369398 | 0.8663101098684087 | 21.5232122552929 | 99.26211650839943 | 15.983735000853056 | 14.580138896537854 | 85.68630242294881 | 3.482043038183848 |
| SE_173_2_17 | SE_173 | SE_173_2 | GTGCGCCAGCGCCTCCTACG | 286.1170862057109 | 283.8274656360265 | 274.53246781240324 | 324.0258139020337 | 317.11775930609264 | 304.0748485638115 | 329.3051475059821 | 308.1852379213163 | 155.39742361940472 | 433.0301252271742 | 165.503680022408 | 119.5501443109788 |
| SE_173_2_3 | SE_173 | SE_173_2 | TCAAGGTTAAAAAGGTAGAG | 164.73407993662144 | 214.86938419628768 | 254.78193055971235 | 171.77272062276487 | 181.36166167672854 | 187.12298373157628 | 151.73864639981528 | 254.30008895961376 | 221.1083341784673 | 116.64111117230283 | 475.38291070266126 | 243.74301267286938 |
| SE_173_3_0 | SE_173 | SE_173_3 | ACGAGTGTGAGAACAGTGAG | 12810.242268755694 | 11486.017825897368 | 11450.373972247538 | 11791.156223961154 | 11960.32432004952 | 11394.57687509918 | 12718.066121652604 | 11601.378036105503 | 11353.779762272621 | 12514.862221843267 | 11184.996928606566 | 12133.759307057984 |
| SE_173_3_2 | SE_173 | SE_173_3 | AGGGAACGTAGAGAGAAGAG | 2363.71729172218 | 2867.257038414648 | 2830.2519883106033 | 2179.691720023721 | 2596.335367161588 | 2091.272605222339 | 2089.9039099889455 | 1774.4285017739592 | 2565.3894676369155 | 1810.1242440051744 | 1410.8895275669106 | 1597.0970735136584 |
| SE_173_3_23 | SE_173 | SE_173_3 | AGCTCAGTCCTCCAGAGGCG | 123.55055995246607 | 186.8863946265386 | 177.75483527421792 | 169.17010364363205 | 144.2408537311993 | 208.7807364782865 | 147.4340039487567 | 189.070698111237 | 180.26101139850945 | 158.19450702743572 | 200.71722896334586 | 218.20803039285448 |
| SE_173_3_25 | SE_173 | SE_173_3 | GTATTTCCAACAGACTTCCC | 28.17819788389577 | 51.968409200962604 | 53.32645058226538 | 50.751031093089615 | 33.939024407341016 | 72.77004922894633 | 39.817942672291956 | 22.68848377334844 | 33.74344055735645 | 76.54572920682372 | 43.43004369382337 | 160.17397975645702 |
| SE_173_3_5 | SE_173 | SE_173_3 | CGTTCCCTAATCCGTTGTGA | 1792.5668961524464 | 1898.8457208044028 | 1964.190929780108 | 1584.9937402918758 | 1819.980183843662 | 1559.3581977631356 | 1631.4594889512055 | 1902.0512229990443 | 1415.4485328533206 | 1689.838098108737 | 1806.4550606701127 | 1868.6964304919986 |
| SE_173_4_10 | SE_173 | SE_173_4 | CTTCAGCACCTGAATCCGAG | 235.17957464636086 | 182.888824688003 | 198.49289938954334 | 209.51066682019047 | 203.6341464440461 | 90.09625142631451 | 243.21229848481033 | 212.70453537514163 | 85.2465866712163 | 104.24799311024564 | 295.7938111038781 | 165.97738482009677 |
| SE_173_4_13 | SE_173 | SE_173_4 | GCTACTTGGGAGACTCACGT | 829.0892838915487 | 1165.2916370831229 | 1167.2567516340312 | 908.3133257173475 | 972.565168172866 | 779.6790988815678 | 738.2461803565482 | 715.632592351032 | 816.9464555991563 | 1014.0486602542077 | 953.1133913347184 | 547.8414380075922 |
| SE_173_4_6 | SE_173 | SE_173_4 | TCCTCAACCACATGACCACT | 1373.1452584190747 | 1149.3013573289807 | 1139.6059994802638 | 1190.6972679532564 | 1237.7137963552177 | 1130.5346933782735 | 1164.4057830113486 | 1418.9755893248337 | 1096.6618181140848 | 763.9992781785835 | 1434.3652268608694 | 1044.6129114551545 |
| SE_173_4_7 | SE_173 | SE_173_4 | GTTGCATGCCACCATGCTCA | 1756.8022603767324 | 1418.1379356954988 | 1454.6270686606833 | 1661.7709411762933 | 1545.2862050467456 | 1401.6897577670852 | 1685.267519589438 | 1759.3028459250602 | 1138.3971261718677 | 1039.563903323149 | 1970.7849557278228 | 1830.3939570719763 |
| SE_173_4_9 | SE_173 | SE_173_4 | ACTGCGGCTGAGACACAGAA | 2229.3289633528307 | 2386.549253305744 | 2487.5801669764164 | 2583.097351789305 | 2507.2454280923175 | 2629.2511834506204 | 2479.474051809748 | 2257.50413544817 | 2647.084113196831 | 2919.672814031705 | 1982.5228053748021 | 2140.295787470339 |
| SE_174_1_0 | SE_174 | SE_174_1 | CCTGTAAGCAGTAACTTCAA | 1873.8501592790687 | 2275.6166875113818 | 2231.8107095540695 | 2043.0543286192487 | 2265.429879190013 | 2647.443695757857 | 1853.148575180723 | 1994.6958650735503 | 2697.699274032866 | 2733.7760431008473 | 2102.2488717739907 | 3235.978663485523 |
| SE_174_1_2 | SE_174 | SE_174_1 | CTTACAGGATCAGCTGACTT | 1969.222521347639 | 2158.6877668092156 | 2022.4550146755462 | 1954.5653513287334 | 2037.4020589531904 | 1892.0212799526046 | 1778.8934928999622 | 2147.8431305436525 | 1933.1439498253947 | 1569.5519522123 | 2143.331345538418 | 1692.2729165573503 |
| SE_174_1_3 | SE_174 | SE_174_1 | TTGCTGCTTGATTCTGATAA | 175.57184835350444 | 207.87363680385042 | 237.99397389492512 | 166.56748666449926 | 188.7858232658344 | 153.33688944670834 | 115.14918556581728 | 191.90675858290555 | 228.21221640106864 | 59.7785694758052 | 103.29307689341775 | 126.51423038734649 |
| SE_174_1_4 | SE_174 | SE_174_1 | AGAATCAAGCAGCAAATTAC | 605.831254503759 | 469.7144677779312 | 529.3143983721156 | 675.3791060849618 | 557.8727136956679 | 556.1710905355184 | 597.2691400843794 | 618.261182823745 | 485.7279469703679 | 614.5528544890705 | 638.5390207956734 | 581.5011873767027 |
| SE_174_1_5 | SE_174 | SE_174_1 | GTAGCAAGCTAGGAACTCTG | 1143.3845679811554 | 1174.286169444828 | 1231.4459977052763 | 1129.5357689436355 | 1154.987424362324 | 1249.2191784302454 | 953.4783029094776 | 1318.7681193258782 | 1457.1838409111035 | 1252.4339312126017 | 1315.8129454263785 | 1236.125278555266 |
| SE_174_2_1 | SE_174 | SE_174_2 | AGCAGCCTAGGGCACCTGCA | 220.00669886272468 | 58.96415659339988 | 121.46580410404891 | 139.24000838360485 | 30.757240869152795 | 626.3422094348595 | 309.9342564762185 | 156.9286794323267 | 339.21037612921486 | 10.93510417240339 | 61.03681816429231 | 172.94147089646447 |
| SE_174_2_20 | SE_174 | SE_174_2 | TGGCAGCCAAGAAGGAAAAA | 60.69150313454474 | 30.98116702365078 | 40.488601368016305 | 49.44972260352321 | 39.24199697098805 | 35.51871450460476 | 63.4934761531142 | 51.994441980590175 | 14.207764445202717 | 21.87020834480678 | 9.390279717583432 | 23.21362025455899 |
| SE_174_2_29 | SE_174 | SE_174_2 | TTTGGGGCATCTGGTGACAC | 641.595890279473 | 669.5929647047104 | 553.0150430753447 | 645.4490108249346 | 677.7198936340909 | 482.53473119670366 | 600.4976219226733 | 843.255313576117 | 586.070283364612 | 780.0374309647751 | 1274.730471661951 | 603.5541266185337 |
| SE_174_2_3 | SE_174 | SE_174_2 | ACCTGGCTCAGAAACTCTGG | 547.307305052591 | 647.6063300427647 | 674.4808471793935 | 597.3005967109779 | 600.2964942048442 | 995.3903162388016 | 553.1465549610288 | 604.0808804654022 | 784.0910003196249 | 468.751465523692 | 521.1605243258805 | 752.1212962477113 |
| SE_174_2_9 | SE_174 | SE_174_2 | CTACTGAGCCCCAAGAAAGG | 28.17819788389577 | 47.97083926242702 | 51.35139685699629 | 28.62878677046081 | 55.15091466192915 | 12.128341538157722 | 15.066248578705064 | 45.37696754669688 | 63.93494000341222 | 4.374041668961356 | 28.170839152750297 | 19.73157721637514 |
| SE_174_3_15 | SE_174 | SE_174_3 | TCTCAAGGAAAGCTGTTCCT | 755.3924586567443 | 684.5838519742189 | 817.6722422614025 | 783.387710718973 | 747.7191314742317 | 735.497283278279 | 790.9780503820159 | 1031.3806581967979 | 664.212987813227 | 580.2895280822065 | 859.2105941588841 | 851.9398633423149 |
| SE_174_3_23 | SE_174 | SE_174_3 | GATAAGAATGGGTTAACCAA | 124.63433679415438 | 144.91191027191493 | 176.76730841158337 | 109.30991312357764 | 159.089176909411 | 64.97325824013066 | 94.70213392328898 | 71.84686528227006 | 133.19779167377547 | 107.8930278343801 | 146.72312058724114 | 163.6560227946408 |
| SE_174_3_30 | SE_174 | SE_174_3 | GTCGTCTCCCACACACGTGA | 857.2674817754445 | 903.4508061090421 | 914.4498747995877 | 817.2217314476994 | 898.3235522818076 | 1135.7325540374839 | 815.7297444756027 | 904.703290462269 | 688.1885903145065 | 543.110173896035 | 1163.2209000156477 | 638.3745570003722 |
| SE_174_3_32 | SE_174 | SE_174_3 | TTTAGGGTCTAAGAAAGAAG | 31.429528408960667 | 20.98724217731182 | 21.725590977959968 | 49.44972260352321 | 12.727134152752882 | 7.796790988815679 | 77.48356411905462 | 23.633837263904624 | 3.551941111300679 | 43.74041668961356 | 7.042709788187574 | 3.482043038183848 |
| SE_174_3_6 | SE_174 | SE_174_3 | GACTCTGGTCACCTGCAGCG | 37.93218945909046 | 9.993924846338961 | 31.60085960430541 | 9.109159426964803 | 5.3029725636470335 | 149.8716490072347 | 17.21856980423436 | 33.08737216946648 | 103.89427750554486 | 3.6450347241344634 | 36.3873339056358 | 52.23064557275772 |
| SE_175_1_10 | SE_175 | SE_175_1 | CAGCTTGAGCAGGAAAAGGA | 894.1158943928466 | 972.408887548781 | 990.4894432224477 | 1081.3873548296788 | 1097.715320674936 | 735.497283278279 | 975.0015151647706 | 1090.9379281018375 | 859.5697489347643 | 826.6938754336962 | 1535.3107338248913 | 886.7602937241534 |
| SE_175_1_14 | SE_175 | SE_175_1 | TTTCTAGTATGAATCCAGGA | 1.083776841688299 | 1.9987849692677924 | 0.987526862634544 | 0.0 | 8.484756101835254 | 0.0 | 1.0761606127646475 | 5.67212094333711 | 0.0 | 0.7290069448268927 | 9.390279717583432 | 0.0 |
| SE_175_1_28 | SE_175 | SE_175_1 | CATCTTCCTGACCCCAGGGG | 208.0851536041534 | 291.8226055130977 | 286.38279016401776 | 238.13945359065127 | 292.72408551331625 | 284.1497160368381 | 189.40426784657797 | 224.04877726181584 | 340.9863466848652 | 116.64111117230283 | 221.8453583279086 | 234.4575645710458 |
| SE_175_1_35 | SE_175 | SE_175_1 | ACAGGATGCTGACTACTGAC | 4442.401274080337 | 3925.613679641944 | 3972.8205683787705 | 4039.261551614107 | 3918.896724535158 | 3475.6361607920558 | 4294.957005543708 | 4155.773944484989 | 3537.7333468554766 | 3150.76801554183 | 3833.5816947034364 | 3425.1696685601787 |
| SE_175_1_4 | SE_175 | SE_175_1 | GTTATGGAAGAAAACCAGGC | 36.84841261740216 | 62.96172653193546 | 68.13935352178353 | 48.14841411395682 | 45.60556404736449 | 71.90373911907793 | 18.294730416999005 | 12.289595377230405 | 84.35860139339113 | 62.69459725511277 | 50.47275348201095 | 117.2287822855229 |
| SE_175_2_10 | SE_175 | SE_175_2 | GCAGTGGTGGTGGTTCAGGT | 1145.552121664532 | 1157.2964972060518 | 1141.581053205533 | 1218.0247462341508 | 1038.3220279620891 | 808.2673325072253 | 1016.9717790625918 | 1001.1293464989999 | 1244.0673742330628 | 1323.1476048608101 | 902.6406378527074 | 843.8150962532193 |
| SE_175_2_19 | SE_175 | SE_175_2 | GGGAGTGTTCCAGGTAAAAG | 110.54523785220648 | 163.900367479959 | 171.82967409841066 | 144.44524234187045 | 215.30068608406958 | 175.86095230328698 | 131.291594757287 | 226.8848377334844 | 113.66211556162173 | 182.25173620672317 | 258.2326922335444 | 200.79781520193526 |
| SE_175_2_2 | SE_175 | SE_175_2 | TGAAGAGAAGAGCCTAACAA | 736.9682523480433 | 880.4647789624626 | 875.9363271568405 | 862.7675285825235 | 979.9893297619718 | 926.9518175591974 | 768.3786775139583 | 782.7526901805212 | 1032.7268781106725 | 906.8846393646545 | 907.3357777114992 | 658.1061342167474 |
| SE_175_2_30 | SE_175 | SE_175_2 | CCACAGTGTCTGGCAAATGA | 1004.6611322450531 | 959.4167852485404 | 1007.2773998872349 | 1085.291280298378 | 1022.4131102711481 | 1043.9036823914325 | 1244.0416683559324 | 1019.0910628195675 | 872.0015428243167 | 1054.1440422196868 | 1106.879221710147 | 1086.3974279133606 |
| SE_175_2_33 | SE_175 | SE_175_2 | AGGGTAAGACCTTAAGGGCA | 1960.5523066141327 | 1991.789221875355 | 2015.5423266371045 | 2255.167612418572 | 1944.0697418330026 | 1858.2351856677367 | 2162.006671044177 | 1963.4991998851963 | 1483.8233992458588 | 2024.452285784281 | 2125.7245710679495 | 1508.8853165463343 |
| SE_175_3_13 | SE_175 | SE_175_3 | AGCAGCTTTATGATACACGC | 649.182328171291 | 659.5990398583715 | 624.1169771850318 | 550.4534910865874 | 624.6901679976206 | 330.9304619697321 | 597.2691400843794 | 722.2500667849254 | 685.5246344810311 | 430.84310439269353 | 831.0397550061338 | 526.949179778489 |
| SE_175_3_14 | SE_175 | SE_175_3 | GCAGCTTTATGATACACGCA | 1353.6372752686852 | 1348.180461771126 | 1453.6395417980489 | 1314.3215744620645 | 1388.3182171627934 | 1304.6630254618235 | 1293.5450565431063 | 911.3207648961624 | 1181.0204195074757 | 1713.1663203431976 | 863.9057340176757 | 1080.594022849721 |
| SE_175_3_15 | SE_175 | SE_175_3 | TAAACGCTAAGGACTGTGGG | 6766.01882266005 | 6701.926001954907 | 6436.700090651958 | 6670.507317517368 | 6470.68712216211 | 6292.0103279742525 | 6377.327791243301 | 6656.233927006098 | 6589.738746740585 | 5914.4333433805805 | 6951.154560941136 | 6330.354243418236 |
| SE_175_3_2 | SE_175 | SE_175_3 | ATTGGGGTGCCTCCTTACCG | 514.7939998019419 | 475.71082268573457 | 522.4017103336738 | 566.0691929613841 | 538.7820124665386 | 647.9999621815697 | 634.934761531142 | 554.9224989564806 | 594.9501361428637 | 755.9802017854877 | 879.1649385587489 | 937.8302582841832 |
| SE_175_3_24 | SE_175 | SE_175_3 | GCTAAGAAAGACCTTAGAAG | 153.89631151973845 | 270.83536333578587 | 286.38279016401776 | 195.19627343496006 | 224.84603669863424 | 369.91441691381056 | 247.51694093586892 | 198.52423301679886 | 237.09206917932033 | 288.6867501514495 | 187.80559435166865 | 254.18914178742094 |
| SE_175_4_10 | SE_175 | SE_175_4 | CGTTTAGTTCCAAGGCCCCA | 727.2142607728485 | 988.3991673029233 | 1029.002990865195 | 809.413880510301 | 990.5952748892659 | 732.0320428388054 | 901.8225934967746 | 1064.4680303662644 | 1283.1387264573702 | 793.8885629164861 | 578.675987596079 | 1131.6639874097507 |
| SE_175_4_15 | SE_175 | SE_175_4 | TCTTCAGCCTGACTTCCCAG | 861.6025891421976 | 861.4763217544186 | 867.0485853931297 | 809.413880510301 | 888.7782016672428 | 1077.6897766763004 | 860.928490211718 | 797.8783460294202 | 785.8669708752752 | 720.25886148897 | 866.2533039470716 | 958.7225165132862 |
| SE_175_4_2 | SE_175 | SE_175_4 | ATAATGCTAAGTGAAAAACG | 106.21013048545329 | 158.9034050567895 | 156.02924429625796 | 131.43215744620645 | 219.5430641349872 | 97.02673230526177 | 153.8909676253446 | 141.80302358342774 | 220.2203489006421 | 166.94259036535843 | 107.98821675220947 | 107.94333418369929 |
| SE_175_4_23 | SE_175 | SE_175_4 | GTCTGCTGAGAGTAAGGAGA | 228.67691359623106 | 276.83171824358925 | 269.5948334992305 | 257.6590809341473 | 298.02705807696327 | 235.63634988420716 | 210.9274801018709 | 409.3380614108281 | 332.99447918443866 | 616.7398753235511 | 253.53755237475266 | 289.0095721692594 |
| SE_175_4_26 | SE_175 | SE_175_4 | GCCACAATCTCAGAAGCAGG | 4652.653981367867 | 4762.105189280515 | 4711.490661629409 | 4701.627572803405 | 4797.068981075106 | 5369.390060964397 | 4649.013847143277 | 4995.247844098882 | 5311.927931950166 | 4850.812210878144 | 4104.726021548658 | 4555.672974957201 |
| SE_176_1_1 | SE_176 | SE_176_1 | TGCCACTCATGATTATGTGG | 521.2966608520718 | 591.6403509032665 | 583.6283758170156 | 523.126012805693 | 568.478658822962 | 585.6256342710443 | 518.7094153525601 | 514.2722988625646 | 491.94384391514404 | 618.9268961580318 | 455.42856630279647 | 444.5408278748046 |
| SE_176_1_28 | SE_176 | SE_176_1 | AAGTGGCCTCCCTCAGTGCA | 981.9018185695988 | 918.4416933785506 | 926.3001971512023 | 981.1866011330659 | 849.5362046962548 | 821.2619841552515 | 991.1439243562403 | 725.086127256594 | 775.2111475413732 | 903.23960464052 | 708.9661186775492 | 725.4256329549684 |
| SE_176_1_47 | SE_176 | SE_176_1 | GTGCAGGAAGGGGAATCAGA | 797.659755482588 | 874.4684240546592 | 943.0881538159896 | 861.4662200929571 | 731.8102137832907 | 1549.8287865545833 | 766.226356288429 | 1053.12378847959 | 1049.5985983893506 | 675.7894378545295 | 853.3416693353944 | 1215.2330203261631 |
| SE_176_1_56 | SE_176 | SE_176_1 | ATGTGGTGTCCCCCACAGTG | 482.28069455129304 | 426.7405909386737 | 486.85074327883024 | 559.5626505135522 | 435.9043447317862 | 432.28874482433594 | 524.0902184163833 | 512.3815918814523 | 458.2004033577876 | 531.4460627788047 | 465.9926309850778 | 459.629681040268 |
| SE_176_1_7 | SE_176 | SE_176_1 | ACAAGGCTAACCGACCTCCA | 2339.8742012050375 | 2099.723610215816 | 2024.4300684008153 | 2259.071537887271 | 2386.337653641165 | 1572.3528494111617 | 2378.314954209871 | 2621.465229312301 | 2320.3055309571687 | 1994.5630010463783 | 2704.4005586640287 | 1606.3825216154821 |
| SE_176_2_17 | SE_176 | SE_176_2 | ATACCTTCTGGTGTAAACCT | 31.429528408960667 | 60.962941562667666 | 50.36386999436174 | 14.314393385230405 | 59.39329271284678 | 172.39571186381335 | 15.066248578705064 | 53.88514896170255 | 48.83919028038434 | 24.057229179287457 | 72.7746678112716 | 56.873369623669525 |
| SE_176_2_19 | SE_176 | SE_176_2 | GGTGCTCCTAAAATACCTTC | 53.105065242726646 | 56.96537162413208 | 95.79010567555078 | 65.06542447832003 | 69.99923784014085 | 62.37432791052543 | 17.21856980423436 | 44.431614056140695 | 123.4299536176986 | 50.30147919305559 | 10.564064682281362 | 23.21362025455899 |
| SE_176_2_24 | SE_176 | SE_176_2 | GGACCCATGAGTGAAGAAAG | 111.62901469389479 | 203.8760668653148 | 163.9294591973343 | 144.44524234187045 | 113.48361286204653 | 206.1818061486813 | 133.4439159828163 | 121.95060028174787 | 176.7090702872088 | 156.00748619295504 | 115.03092654039705 | 148.56716962917753 |
| SE_176_2_25 | SE_176 | SE_176_2 | AATTTGGCCCGACAACCCGG | 30.34575156727237 | 42.973876839257535 | 46.41376254382357 | 27.32747828089441 | 13.787728665482287 | 52.844916701972934 | 30.13249715741013 | 27.415251226129364 | 18.647690834328564 | 8.019076393095819 | 69.25331291717781 | 25.534982280014887 |
| SE_176_2_7 | SE_176 | SE_176_2 | GGTGTCGTGTGTTCCCTGAG | 709.8738313058358 | 863.4751067236863 | 968.7638522444877 | 854.9596776451251 | 873.9298784890311 | 1035.2405812927484 | 676.9050254289633 | 719.4140063132568 | 586.070283364612 | 959.3731393921908 | 890.9027882057281 | 950.5977494241906 |
| SE_176_3_1 | SE_176 | SE_176_3 | GTAGCCACTGACACTTGGAA | 993.82336382817 | 1151.3001422982484 | 1094.1797637990749 | 1038.4441746739876 | 1038.3220279620891 | 1052.5667834901167 | 1154.7203374964668 | 1257.3201424397262 | 956.3601442177079 | 1393.8612785090188 | 1212.5198685329608 | 882.1175696732415 |
| SE_176_3_13 | SE_176 | SE_176_3 | TTATACGTTTAGCAGAGACG | 91.0372547018171 | 92.94350107095235 | 86.90236391183987 | 92.39290275921444 | 107.12004578567009 | 6.93048087894727 | 78.55972473181926 | 63.33868386726439 | 165.1652616754816 | 48.114458358574915 | 85.68630242294881 | 0.0 |
| SE_176_3_16 | SE_176 | SE_176_3 | GAGGGCTCTCTACTTCCTGT | 1649.508353049591 | 1827.888854395396 | 1893.088995670421 | 1747.6573014876758 | 1681.0423026761098 | 1747.3474916045805 | 1769.2080473850804 | 1734.7236551705994 | 1467.8396642450057 | 2405.7229179287456 | 2435.603801748203 | 1510.0459975590622 |
| SE_176_3_32 | SE_176 | SE_176_3 | ATAAAACTGAGATAACGGCC | 71.52927155142773 | 96.94107100948793 | 117.51569665351074 | 87.18766880094883 | 109.24123481112889 | 144.67378834802426 | 62.417315540349556 | 84.13646065950047 | 170.4931733424326 | 132.67926395849446 | 78.64359263476125 | 132.31763545098624 |
| SE_176_3_8 | SE_176 | SE_176_3 | ATAGTGGCTGCTCTTAAGGG | 332.71949039830776 | 424.7418059694059 | 369.3350466253195 | 317.5192714542017 | 382.87461909531584 | 332.6630821894689 | 326.0766656676882 | 459.4417964103059 | 278.8273772371033 | 457.8163613512886 | 647.9293005132569 | 307.5804683729066 |
| SE_176_4_11 | SE_176 | SE_176_4 | AAGCTCTGAGACCAAGGCAG | 261.19021884688004 | 352.78554707576535 | 340.6967676089177 | 297.9996441107057 | 334.0872715097631 | 295.41174746512735 | 178.6426617189315 | 261.8629168840632 | 350.75418474094204 | 196.831875103261 | 211.28129364562722 | 311.06251141109044 |
| SE_176_4_21 | SE_176 | SE_176_4 | GGGGATCAGTTCCCAAGCCA | 318.63039145635986 | 521.6828769788938 | 502.6511730809829 | 350.0519836933617 | 433.78315570632736 | 391.57216966052073 | 402.4840691739781 | 368.6878613169122 | 508.8155641938223 | 754.5221878958339 | 374.4374037386394 | 393.4708633147749 |
| SE_176_4_32 | SE_176 | SE_176_4 | AGATGGGCAGGAAAAGGCCA | 3275.1736155820395 | 3215.045623067244 | 3258.838646693995 | 3353.471977612614 | 3141.480946704503 | 3223.5399188203487 | 3424.3430698171082 | 3061.054602420927 | 2793.6016840379843 | 4021.9313146099666 | 3430.9734518120467 | 3440.258521725642 |
| SE_176_4_43 | SE_176 | SE_176_4 | TGAAAACCCAGCACAGCCCA | 1240.9244837331023 | 1432.1294304803732 | 1504.990938655045 | 1385.8935413882164 | 1408.4695129046522 | 1831.379572261816 | 1277.4026473516365 | 1727.16082724615 | 1563.742074250124 | 1449.9948132606894 | 1433.1914418961715 | 1400.941982362635 |
| SE_176_4_9 | SE_176 | SE_176_4 | GGCCCAACACAGATAAGAAA | 76.94815575986922 | 132.9192004563082 | 132.3285995930289 | 61.16149900962082 | 85.90815553108195 | 19.05882241710499 | 69.95043982970209 | 64.28403735782058 | 207.78855501108973 | 26.973256958595027 | 240.62591776307545 | 152.0492126673614 |
| SE_176_5_39 | SE_176 | SE_176_5 | TTCGAGGACTTCCTGAAGGT | 124.63433679415438 | 156.9046200875217 | 166.89203978523796 | 117.11776406097604 | 145.30144824392872 | 29.454543735525895 | 153.8909676253446 | 181.50787018678753 | 161.6133205641809 | 168.4006042550122 | 165.503680022408 | 103.3006101327875 |
| SE_176_5_42 | SE_176 | SE_176_5 | CACCTTCGAGGACTTCCTGA | 374.98678722415144 | 417.7460585769686 | 422.66149720758483 | 335.7375903081313 | 407.2682928880922 | 654.930443060517 | 351.90452037403975 | 530.3433082020198 | 627.8055914223951 | 424.2820418892515 | 331.007360044816 | 364.4538379965761 |
| SE_176_5_45 | SE_176 | SE_176_5 | AGCCTGATCAGGTGTACGAG | 446.51605877557915 | 518.6846995249921 | 495.73848504254113 | 482.7854496291346 | 575.9028204120679 | 698.2459485539374 | 432.6165663313883 | 467.0046243347554 | 634.0214883671712 | 640.7971045028387 | 671.4049998072154 | 516.5030506639375 |
| SE_176_5_46 | SE_176 | SE_176_5 | GAGCCTGATCAGGTGTACGA | 817.1677386329774 | 645.6075450734969 | 793.9715975581734 | 819.8243484268323 | 765.7492381906317 | 808.2673325072253 | 707.0375225863734 | 746.8292575393862 | 676.6447817027794 | 622.5719308821664 | 758.2650871948622 | 727.7469949804243 |
| SE_176_5_47 | SE_176 | SE_176_5 | GGAGCCTGATCAGGTGTACG | 730.4655912979134 | 768.5328206834662 | 846.3105212778042 | 788.5929446772386 | 861.2027443362783 | 570.8983624032813 | 751.1601077097239 | 927.3917742356175 | 895.0891600477711 | 813.5717504268122 | 1225.4315031446379 | 553.6448430712319 |
| SE_177_1_16 | SE_177 | SE_177_1 | TGTCTCATAGTCTTCCATGA | 140.89098941947887 | 120.92649064070145 | 145.16644880727796 | 152.25309327926885 | 125.15015250207 | 105.68983340394587 | 154.96712823810924 | 130.45878169675353 | 115.43808611727208 | 152.36245146882055 | 113.85714157569912 | 177.58419494737626 |
| SE_177_1_17 | SE_177 | SE_177_1 | GTGTCTCATAGTCTTCCATG | 54.188842084414944 | 76.95322131681 | 65.1767729338799 | 72.87327541571842 | 90.15053358199957 | 67.57218856973589 | 52.73187002546773 | 65.22939084837677 | 58.607028336461205 | 20.412194455152996 | 42.25625872912545 | 68.48017975094902 |
| SE_177_1_5 | SE_177 | SE_177_1 | TTAGCCCCTGAAGAGGCTGC | 57.440172609479845 | 92.94350107095235 | 87.88989077447442 | 71.57196692615203 | 75.30221040378788 | 216.5775274671022 | 108.6922218892294 | 41.59555358447214 | 103.0062922277197 | 344.82028490312024 | 71.60088284657367 | 95.17584304369186 |
| SE_177_1_6 | SE_177 | SE_177_1 | GGTTCCGGCAGCCTCTTCAG | 15.172875783636185 | 46.97144677779312 | 33.5759133295745 | 39.03925468699201 | 31.8178353818822 | 7.796790988815679 | 8.60928490211718 | 28.36060471668555 | 2.6639558334755096 | 35.72134029651774 | 56.341678305500594 | 45.26655949639003 |
| SE_177_1_9 | SE_177 | SE_177_1 | AAGAGGCTGCCGGAACCCAG | 1289.6944416090757 | 1680.9781591542135 | 1621.5191084459213 | 1359.8673715968885 | 1741.496189901686 | 1649.4544491894503 | 1177.3197103645243 | 1567.3960873421547 | 1531.774604248418 | 1541.849688308878 | 1637.430025753611 | 1382.3710861589877 |
| SE_177_2_1 | SE_177 | SE_177_2 | ACACCTGACAACTTCCTCAG | 785.7382102240167 | 744.5474010522527 | 904.5746061732424 | 778.1824767607075 | 839.9908540816901 | 1061.2298845888006 | 828.6436718287786 | 836.6378391422237 | 818.7224261548065 | 958.6441324473639 | 1084.5773073808864 | 1017.9172481624116 |

| | | | | | | | | | | | | | | | |
|---|---|---|---|---|---|---|---|---|---|---|---|---|---|---|---|
| SE_177_2_12 | SE_177 | SE_177_2 | CCTGGATGTCATGACCACAG | 3223.152327181001 | 2714.349988265662 | 2871.728116541254 | 3046.3631740749433 | 2832.8479435002455 | 2573.8073364190423 | 3135.932025596183 | 2896.5630950641507 | 2601.7968640277477 | 4364.564578678606 | 2888.6847981216033 | 3295.1733951346487 |
| SE_177_2_19 | SE_177 | SE_177_2 | CTAGAGGTCAGAGAAGTAGC | 1395.904572094529 | 1312.2023323243056 | 1259.0967498590437 | 1510.819156386591 | 1465.7416165920401 | 1686.7057839137917 | 1306.458983896282 | 1692.1827480955712 | 1478.4954875789076 | 1346.4758270952707 | 1558.7864331188498 | 2090.386503923037 |
| SE_177_2_28 | SE_177 | SE_177_2 | CTTGATACTGACCAGAATGA | 352.22747354869716 | 389.7630690072195 | 452.2873030866212 | 451.554045879541 | 407.2682928880922 | 232.17110944473353 | 278.7255987060437 | 366.7971543357998 | 229.98818695671898 | 282.12568764800744 | 248.84241251596094 | 355.1683898947525 |
| SE_177_2_32 | SE_177 | SE_177_2 | TTAGGACGGTGACAGCACTC | 2534.954032708931 | 2191.6677188021345 | 2183.4218932849767 | 2468.5822047074616 | 2209.218370015354 | 1863.433046326947 | 2611.8418071797996 | 2266.957670353732 | 1989.0870223283803 | 2228.5742303358106 | 2642.1899555350383 | 2515.1957545814666 |
| SE_177_3_17 | SE_177 | SE_177_3 | GGTGCCCTCATTGCTCCATG | 438.92962088376106 | 497.6974573476803 | 483.8881626909265 5 | 422.9252591090801 | 506.9641770846564 | 565.700501744071 | 385.2654993697438 | 436.75331263695745 | 385.38561057612367 | 336.8012085100244 | 294.6200261391802 | 624.4463848476368 |
| SE_177_3_6 | SE_177 | SE_177_3 | TCCATAATGTAGGTCTCACG | 1931.2903318885487 | 1807.9010047027182 | 1819.0244809728301 | 1916.8274051313078 | 1636.4973331414747 | 1787.1977566585272 | 1604.5554736320894 | 2226.3074702598155 | 1548.646324527096 | 1994.5630010463783 | 2003.6509347393649 | 1562.27664313182 |
| SE_177_3_7 | SE_177 | SE_177_3 | ATTATCTGCAAATATCACAA | 19.50798315038938 | 11.992709815606755 | 18.763010390056337 | 9.10915942696 4803 | 11.666539640023474 | 23.390372966447035 | 7.533124289352532 | 0.0 | 18.647690834328564 | 5.103048613788249 | 7.042709788187574 | 11.606810127279495 |
| SE_177_3_8 | SE_177 | SE_177_3 | CTAGCTTCACACAGCCAGAA | 201.5824925540236 | 191.88335704970808 | 213.30580232906152 | 187.38842249756166 | 181.36166167672854 | 129.0802063703929 | 230.29837113163455 | 171.10898179066947 | 145.62958556332785 | 214.32804177910643 | 184.28423945757487 | 104.46129114551545 |
| SE_177_3_9 | SE_177 | SE_177_3 | GCTAGCTTCACACAGCCAGA | 1301.615986867647 | 1204.2679439838448 | 1218.6081484910274 | 1414.5223281586773 | 1297.1070890680644 | 1194.6416415085357 | 1385.0187086281012 | 1265.8283238547317 | 1029.1749369993718 | 1433.9566604744978 | 1184.3490293802104 | 873.9928025841459 |
| SE_177_4_1 | SE_177 | SE_177_4 | AGATATGGTTGGAAATCCTG | 234.09579780467257 | 256.84386855091134 | 279.470102125576 | 261.5630064028465 | 254.5426830550576 | 251.22993186183854 | 289.48720483369016 | 175.8357492434504 | 491.94384391514404 | 218.7020834480678 | 115.03092654039705 | 201.9584962146632 |
| SE_177_4_11 | SE_177 | SE_177_4 | TCTTCGAGATCATCTCTCCC | 523.4642145354484 | 613.6269855652123 | 660.65547110251 | 624.6280749918722 | 584.3875765139031 | 656.6630632802538 | 516.5570941270308 | 738.3210761243805 | 614.4858122550174 | 566.4383961304956 | 855.6892392647902 | 457.30831901481207 |
| SE_177_4_16 | SE_177 | SE_177_4 | CATGCGTGTGAAGTCGAAAT | 13646.91799053906 | 12054.672149654056 | 12607.755455255223 | 12807.478154312514 | 12337.895966581189 | 11664.865629378124 | 12768.645670452543 | 11094.668565167387 | 12549.895931503124 | 11209.210783658302 | 11013.624323760669 | 12491.249058978192 |
| SE_177_4_24 | SE_177 | SE_177_4 | AATGTGATAAAATTAAACTG | 35.76463577571386 | 90.94471610168455 | 64.18924607124536 | 39.03925468699201 | 53.02972563647034 | 32.05347406513112 | 59.18883370205561 | 67.12009782948914 | 31.07948472388094 | 25.515243068941242 | 51.646538446708874 | 54.55200759821362 |
| SE_177_4_25 | SE_177 | SE_177_4 | AGTTTTCCTAGTCTTTCCCT | 4.335107366753196 | 7.9951398770711695 | 5.925161175807264 | 1.3013084895664004 | 13.787728665482287 | 3.465240439473635 | 0.0 | 26.46989773557318 | 17.759705556503395 | 0.0 | 9.390279717583432 | 15.088853165463343 |
| SE_178_1_11 | SE_178 | SE_178_1 | GGTGGCCAGAGTAAGAAGGG | 174.48807151181612 | 249.84812115847404 | 285.39526330138324 | 182.18318853929605 | 214.24009157134017 | 187.12298373157628 | 146.35784333599204 | 308.1852379213163 | 412.91315418870397 | 176.41968064810803 | 138.50662583435562 | 395.79222534023074 |
| SE_178_1_28 | SE_178 | SE_178_1 | CAGGTTTTTCCCCAAAGCCA | 298.0386314642822 | 408.75152621526354 | 416.73633603177757 | 347.4493667142289 | 425.2983996044921 | 489.4652120756509 | 374.5038932420973 | 311.0212983929849 | 450.2085358573611 | 405.3278613237523 | 402.60824289138964 | 331.95476964019355 |
| SE_178_1_4 | SE_178 | SE_178_1 | CTTTACCCCAGAGCATTGGG | 24.926867358830876 | 36.977521931454156 | 40.488601368016305 | 23.423552812195208 | 26.51486281823517 | 13.86096175789454 | 9.685445514881827 | 29.305958207241734 | 82.5826308377408 | 2.187020834480678 | 0.0 | 0.0 |
| SE_178_1_6 | SE_178 | SE_178_1 | AATTGTGCTTACAGGAAGCG | 97.53991575194691 | 129.9210230024065 | 158.00429802152703 | 141.84262536273764 | 126.21074701479941 | 194.05346461052355 | 129.1392735317577 | 157.8740329228829 | 83.47061611556596 | 101.33196533093808 | 201.8910139280438 | 61.51609367458132 |
| SE_178_2_16 | SE_178 | SE_178_2 | TTCCCAATAGCAAAGTCTGA | 10.837768416882989 | 1.9987849692677924 | 7.900214901076352 | 5.20523395826 5602 | 3.1817835381882205 | 0.866310109868 4087 | 0.0 | 0.945353490556185 | 6.215896944776189 | 0.0 | 9.390279717583432 | 0.0 |
| SE_178_2_18 | SE_178 | SE_178_2 | CTCTTGGCTTCCCAGATGCA | 41.18351998415536 | 81.9501837399795 | 88.87741763710896 | 41.641871666124814 | 90.15053358199957 | 42.44919538355203 | 18.294730416999005 | 37.8141396222474 | 108.33420389467071 | 37.908361130998415 | 86.86008738764674 | 119.5501443109788 |
| SE_178_2_22 | SE_178 | SE_178_2 | GAAAGTTTCGAGAATGAAGG | 36.84841261740216 | 75.95382883217611 | 47.40128940645811 | 58.55888203048802 | 32.87842989461161 | 44.18181560328885 | 30.13249715741013 | 40.650200093915956 | 55.94307250298569 | 107.8930278343801 | 96.25036710523018 | 94.01516203096391 |
| SE_178_2_24 | SE_178 | SE_178_2 | GTGTTTGAAAAAGTAAGGCG | 40.09974314246706 | 38.97630690072195 | 30.613332741670867 | 26.02616979132801 | 33.939024407341016 | 91.82887164605133 | 46.27490634887984 | 28.36060471668555 | 86.13457194904147 | 59.04956253097831 | 50.47275348201095 | 23.21362025455899 |
| SE_178_2_7 | SE_178 | SE_178_2 | TCACCCATCCTGGAGTGCCA | 402.0812082663589 | 369.7752193145416 | 406.86106740543215 | 411.2134827029 8256 | 451.8132624227273 | 466.0748391092039 | 398.1794267229196 | 535.0700756548007 | 438.66472724563386 | 370.33552797206147 | 497.6848250319219 | 595.4293595294381 |
| SE_179_1_16 | SE_179 | SE_179_1 | GAAAGGGGAATAAAATAGGG | 145.22609678623206 | 291.8226055130977 | 237.00644703229057 | 217.31851775758886 | 254.5426830550576 | 394.171099990126 | 161.42409191469713 | 335.60048914744567 | 357.8580669635434 | 411.15991688236744 | 420.2150173618586 | 381.86405318749536 |
| SE_179_1_18 | SE_179 | SE_179_1 | CGTTCAGCAATTTTAGAACG | 1643.005691999461 | 2180.6744014711617 | 2041.2180250656024 | 1836.1462787781911 | 2118.0072419206253 | 2218.6201913729947 | 1650.8303799809692 | 1984.2969766774324 | 2557.397600136489 | 1959.5706676946875 | 2225.4962930672737 | 2182.080303928545 |
| SE_179_1_20 | SE_179 | SE_179_1 | CAAGACCTGTTGGAACAGCG | 7103.073420425111 | 6790.871933087325 | 6529.527615739605 | 6926.86508996195 | 6626.594515533333 | 6880.234892574902 | 6921.865061302213 | 6976.708760304646 | 6302.03151672523 | 6450.253447828346 | 7504.007279313861 | 7183.454787773279 |
| SE_179_1_28 | SE_179 | SE_179_1 | GCTCCTGATTCCAGGATGGA | 31.429528408960667 | 50.9690167163287 | 38.51354764274722 | 28.62878677046081 | 20.15129574185873 | 85.76470087697247 | 58.112673089290965 | 61.44797688615203 | 69.26285167036325 | 70.71367364820858 | 109.1620017169074 | 55.71268861094157 |
| SE_179_1_34 | SE_179 | SE_179_1 | CTTTAGAGTGAGCTGATACT | 62.859056817921335 | 86.94714616314897 | 73.07698783495626 | 75.4758923948 5123 | 101.81707322202305 | 71.90373911907793 | 61.3411549275849 | 89.80858160283758 | 192.69280528806183 | 62.69459725511277 | 25.823269223354437 | 94.01516203096391 |
| SE_179_2_0 | SE_179 | SE_179_2 | GCTGGCCCGCAGCAACCTCA | 4626.643337167348 | 4115.498251722384 | 4270.066154031769 | 4797.924401031319 | 4570.101755351014 | 4566.3205891163825 | 5023.517740385374 | 5034.952690702241 | 5232.0092569459 | 4818.0068983609335 | 4456.861510958037 | 5021.106061061109 |
| SE_179_2_13 | SE_179 | SE_179_2 | ACCCTCCCCCAGCTGCAGGG | 13.005322100259587 | 14.990887269508443 | 29.62580587903632 | 26.02616979132801 | 7.4241615891058474 | 117.81817494210358 | 47.35106696164449 | 17.961716320567515 | 164.2772763976564 | 1.4580138896537853 | 12.911634611677218 | 8.124767089095647 |
| SE_179_2_2 | SE_179 | SE_179_2 | GCATCTCCGTGAGGTTGCTG | 625.3392376541485 | 709.5686640900662 | 749.5328887396189 | 662.3660211892978 | 654.386814354044 | 622.8769689953858 | 583.2790521184389 | 520.889773296458 | 751.2355450400936 | 879.9113824060595 | 657.3195802308403 | 785.7810456168218 |
| SE_179_2_3 | SE_179 | SE_179_2 | CATTGAGAAGCATCTCCGTG | 2102.5270728753 | 1946.8165600668297 | 1962.215876054839 | 2184.8969539819864 | 2057.553354695049 | 2116.3955984085223 | 2206.1292561675273 | 2250.8866610142763 | 2183.5557981720926 | 2279.6047164736933 | 2212.584658455596 | 1980.1218077138817 |
| SE_179_2_5 | SE_179 | SE_179_2 | TCTCAATGTGGAGTACATGA | 1199.7409637489468 | 1164.292244598489 | 1184.0447082988183 | 1180.2868000367253 | 1270.5922262498293 | 890.5667929447242 | 1200.9952438453465 | 1165.6208538557762 | 1213.8758747870072 | 1186.8233061781812 | 1497.7496149545575 | 1122.3785393079272 |
| SE_179_3_10 | SE_179 | SE_179_3 | GGCTTCCCTCTTTCTTGTGG | 7.586437891818092 | 22.986027146579612 | 27.650752153767232 | 20.820935833062407 | 19.090701229129323 | 0.0 | 1.0761606127646475 | 25.524544245016994 | 23.975602501279585 | 2.187020834480678 | 9.390279717583432 | 44.10587848366208 |
| SE_179_3_19 | SE_179 | SE_179_3 | GCCTGAGGCTGGAGACTAGG | 1285.3593342423226 | 1368.1683114638038 | 1347.9741674961526 | 1393.701392325615 | 1368.1669214209346 | 1505.6469709512944 | 1319.3729112494577 | 1449.2269010226316 | 1164.1486992287976 | 1554.242806370935 | 1748.9395973999142 | 1301.1234152680313 |
| SE_179_3_3 | SE_179 | SE_179_3 | GAAACCTGTGGAAGCAAGAG | 195.07983150389381 | 251.84690612774185 | 236.018920169656 | 175.67664609146405 | 162.27096044759924 | 162.86630065526083 | 192.6327496848719 | 217.43130282792256 | 229.1002016788938 | 247.1333542963166 | 228.88806811609615 | 103.3006101327875 |
| SE_179_3_31 | SE_179 | SE_179_3 | GCTGGGCTGCAGGGCTGCAG | 733.7169218229784 | 155.9052276028878 | 333.7840795704759 | 722.2262117093522 | 153.786204345764 | 1412.0854790855062 | 1318.296750636693 | 355.45291244912556 | 1294.6825350690976 | 48.114458358574915 | 240.62591776307545 | 322.66932153836996 |
| SE_179_3_5 | SE_179 | SE_179_3 | TCTTGCTTCCACAGGTTTCT | 599.3285934536293 | 729.5565137827442 | 722.8696634484862 | 645.4490108249346 | 756.203887576067 | 771.8823078927521 | 553.1465549610288 | 808.2772344255382 | 708.6122517044855 | 1229.105708978141 | 579.849772560777 | 1230.3218734916263 |
| SE_179_4_11 | SE_179 | SE_179_4 | CATCCCCTGGAGACAGAGGA | 318.63039145635986 | 295.82017545163325 | 284.40773643874866 | 376.0781534846897 | 296.9664635642339 | 189.7219140611815 | 370.1992507910387 | 331.8190751852209 | 267.2835686253761 | 681.6214934131447 | 442.5169316911192 | 194.9944101382955 |
| SE_179_4_12 | SE_179 | SE_179_4 | GAACGGGGTGAGGGAAGTGA | 55711.548546987004 | 56975.36554897842 | 56172.503000378136 | 53759.65632096714 | 53958.8064296213 | 50587.31255565586 | 52971.853842114244 | 44492.11667953629 | 52387.579450573714 | 38049.05947135001 | 33624.244098736875 | 39414.40583021571 |
| SE_179_4_13 | SE_179 | SE_179_4 | AGGCGACTGCACCACACAAA | 2770.133607355292 | 3607.806869528365 | 3590.647672539202 | 3114.0312155323963 | 3450.1139499087603 | 2713.283264107856 | 2878.729639145432 | 3251.07065402272 | 3701.1226379753075 | 4278.541759189033 | 3561.2635828935167 | 3364.8142558983254 |
| SE_179_4_20 | SE_179 | SE_179_4 | AAAGCAACTGGACATCCCCT | 694.7009555221996 | 712.5668415439679 | 710.0318142342371 | 685.7895740014931 | 785.9005339324904 | 646.2673419618329 | 746.8554652586654 | 995.4572255556628 | 626.9176061445698 | 809.9267157026777 | 598.6303319959438 | 511.8603266130257 |
| SE_179_4_32 | SE_179 | SE_179_4 | GAGGGGAAGAGACATGGCCG | 6200.287311298758 | 6834.845202411216 | 6787.272126887221 | 6637.974605278208 | 6926.742762635756 | 5601.561170409131 | 6665.738835464226 | 6255.404047010276 | 7119.865957602211 | 7857.236851344249 | 6037.949858406147 | 6377.942164940082 |
| SE_179_5_27 | SE_179 | SE_179_5 | TGAGTAGGCCTTGTGCATGC | 6833.212986844725 | 5303.775915952087 | 5586.439461923615 | 6161.695698096906 | 5133.277441610328 | 4477.956957909805 | 6187.923523396723 | 4563.221298914705 | 5346.559357785347 | 4467.354557899198 | 4962.762830742844 | 4263.1813597497585 |
| SE_179_5_28 | SE_179 | SE_179_5 | GATCTTCTTCTCAGGTGAGT | 50.93751155935005 | 51.968409200962604 | 50.36386999436174 | 62.46280749918722 | 69.99923784014085 | 61.50801780065702 | 63.4934761531142 | 25.524544245016994 | 64.82292528123739 | 201.93492371704926 | 12.911634611677218 | 25.534982280014887 |
| SE_179_5_29 | SE_179 | SE_179_5 | AGGATGGAGATCTTCTTCTC | 1012.2475701368712 | 1216.2606537994516 | 1137.6309457549946 | 1086.5925887879444 | 1233.4714183043 | 1108.8769406315632 | 1028.809545803003 | 888.6322811228139 | 960.8000706068337 | 1018.422701923169 | 1203.1295888153772 | 1101.486281078824 |
| SE_179_5_30 | SE_179 | SE_179_5 | TTTGCAGGTCTTGAGCAGGA | 483.36447139298133 | 398.75760136892455 | 407.8485942680667 | 478.88152416043533 | 362.7233233534571 | 384.64168878157346 | 537.004145769559 | 364.90644735468743 | 349.8661994631169 | 705.6787225924321 | 335.7024999036077 | 391.14950128931895 |
| SE_179_5_31 | SE_179 | SE_179_5 | TGAGTTTGCAGGTCTTGAGC | 658.9363197464858 | 718.5631964517713 | 753.4829961901571 | 648.0516278040674 | 636.356707637644 | 529.3154771295978 | 845.8622416330129 | 479.2942197119858 | 474.18413835864067 | 801.178632364755 | 572.8070627725893 | 531.5919038294008 |
| SE_179_6_15 | SE_179 | SE_179_6 | CGTGCAGGGCAATTTCCAAG | 18220.456262463682 | 20106.777398349357 | 20307.502403216764 | 18828.63253553625 | 19316.607860340686 | 19562.148590938537 | 17960.0444664292 | 20685.279726859884 | 19857.126782726446 | 18057.502023362133 | 19241.856926293152 | 18805.353768218236 |
| SE_179_6_29 | SE_179 | SE_179_6 | CTGAATCCTCCACAAGAAAG | 94.288585226882 | 110.93256579436247 | 118.50322351614528 | 110.61122161314404 | 85.90815553108195 | 168.0641613144713 | 74.25508228076067 | 134.24019565897828 | 128.75786528464963 | 64.88161808959345 | 113.85714157569912 | 309.9018303983625 |
| SE_18_1_19 | SE_18 | SE_18_1 | ATAATTTATCGCCCAAACAG | 13.005322100259587 | 28.982382054382988 | 15.800429802152705 | 1.3013084895664004 | 13.787728665482287 | 0.0 | 12.913927353175769 | 0.0 | 12.431793889552377 | 0.0 | 5.868924823489645 | 0.0 |
| SE_18_1_2 | SE_18 | SE_18_1 | AGCTGAGCTATTCTTAGCCG | 361.98146512389184 | 471.713252747199 | 479.9380552403884 | 446.3488119212 7534 | 474.0857471900448 | 524.1176164703872 | 389.5701418208024 | 458.4964429197497 | 380.9456841869978 | 726.819923992412 | 401.4344579266917 | 285.52752913107554 |
| SE_18_1_31 | SE_18 | SE_18_1 | GCACAGGTGGGGATTGTCCG | 1095.6983869468702 | 1151.3001422982484 | 1130.718257716553 | 1077.4834293609795 | 1040.443216987548 | 1344.5132905157704 | 977.1538363903 | 1111.7357048940735 | 1125.9653322823153 | 933.8578963232495 | 1525.9204541073077 | 1303.4447772934873 |
| SE_18_1_6 | SE_18 | SE_18_1 | ACTCACAGTTGAGCAAGGGG | 409.667646158177 | 452.72479553915497 | 481.9131089656575 | 357.85983463076013 | 378.6322410443982 | 141.20854790855063 | 309.9342564762185 | 187.17999113012462 | 274.38745084797745 | 305.453909882468 | 241.7997027277734 | 142.76376456553777 |
| SE_18_1_9 | SE_18 | SE_18_1 | GACAGGACCCGTGAAATGCA | 3484.342546027881 | 3248.0255750601627 | 3197.6119812106535 | 3359.978520060446 | 3044.9668460461266 | 3361.283226289426 | 3741.810450582679 | 2967.4646068558645 | 2910.8157407109065 | 3381.863217051955 | 3267.8173417190346 | 3300.9768001982884 |
| SE_18_10_0 | SE_18 | SE_18_10 | GCTGTCTTAGCTGGCAACTA | 512.6264461185654 | 628.6178728347207 | 697.1939650199881 | 467.1697477543378 | 745.597942448773 | 580.4277736118338 | 527.3187002546773 | 1073.9215652718262 | 763.667338929646 | 573.7284655787645 | 1011.8026395696148 | 764.8887873877187 |
| SE_18_10_11 | SE_18 | SE_18_10 | AGACAGAAGTGGTTGGGCCA | 3965.5394637374857 | 4456.291088982543 | 4355.980991080974 | 4272.195771246493 | 4452.375764438049 | 4644.2884990045395 | 4163.6654107864215 | 4467.7405963685305 | 4653.930841081715 | 4057.6526549064847 | 3987.347525078865 | 4366.481969882546 |
| SE_18_10_12 | SE_18 | SE_18_10 | TCAGTCCCTTGCATGAGAAG | 800.9110860076529 | 819.5018373997949 | 871.9862197063023 | 766.4707003546099 | 843.1726376198784 | 813.4651931664358 | 767.3025169011936 | 952.9163184806345 | 882.6573661582188 | 915.6327227025772 | 1205.477158744773 | 968.0079646151098 |
| SE_18_10_20 | SE_18 | SE_18_10 | TTTGCTAGTGGAAGTCACTG | 99.7074694353235 | 106.9349958558269 | 111.59053547770347 | 75.47589239485123 | 85.90815553108195 | 111.75400417302473 | 92.54981269775968 | 71.84686528227006 | 82.5826308377408 | 159.6525209170895 | 91.55522724643846 | 265.7959519147004 |
| SE_18_10_23 | SE_18 | SE_18_10 | AGCAAGCCGGGCTTACCCAG | 404.24876194973547 | 701.5735242129952 | 677.4434277672972 | 566.0691929613841 | 655.4474088667733 | 732.0320428388054 | 403.5602297867428 | 513.3269453720085 | 812.5065292100304 | 815.029764316466 | 726.5728931480181 | 702.2120127004094 |
| SE_18_11_1 | SE_18 | SE_18_11 | AGCAAGTGTGACAAATTTGG | 482.28069455129304 | 491.70110243987693 | 487.83827014146476 | 428.13049306734575 | 610.9024393321383 | 300.60960812433785 | 458.4444210377398 | 731.7036016904872 | 459.97637391343795 | 330.2401460065824 | 862.7319490529778 | 581.5011873767027 |
| SE_18_11_10 | SE_18 | SE_18_11 | GCTTGAGGACCAGGTCAAAA | 1380.7316963108929 | 1596.0297979603322 | 1678.7956664787248 | 1443.151114929138 | 1432.8631866974285 | 2330.3741955460196 | 1467.883075810979 | 1883.1441531879204 | 1564.6300595279492 | 1394.5902854538456 | 2109.2915815621786 | 1912.8023089756607 |
| SE_18_11_15 | SE_18 | SE_18_11 | TGTCTTTGTGATCCCATGAG | 121.38300626908948 | 130.9204154870404 | 160.96687860943067 | 105.40598765487843 | 113.48361286204653 | 43.31550549342044 | 83.94052779564251 | 104.93423745173654 | 173.1571291759081 | 118.8281320067835 | 83.33873249355297 | 80.08698987822851 |
| SE_18_11_30 | SE_18 | SE_18_11 | TATATCCTCTGTACTGCCCA | 215.67159149597148 | 231.8590564350639 | 174.7922546863143 | 216.01720926802247 | 274.6939787969163 | 191.45453428091832 | 242.13613787204568 | 210.81382839402926 | 252.18781890234823 | 161.1105348067433 | 325.13843522132635 | 133.47831646371418 |
| SE_18_11_4 | SE_18 | SE_18_11 | GTGAAGTTCTCAGGATACAG | 108.37768416882989 | 158.9034050567895 | 123.440857829318 | 83.28374333224963 | 84.84756101835254 | 199.251325269734 | 125.91079169346375 | 96.42605603673087 | 101.23032167206935 | 164.02656258605086 | 171.37260484589763 | 146.24580760372163 |
| SE_18_12_14 | SE_18 | SE_18_12 | ATAGGATTGGAGGACAACTG | 1352.553498426997 | 1372.1658814023394 | 1406.2382523915908 | 1302.6097980559668 | 1280.1375768643938 | 1426.8127509532692 | 1305.3828232835174 | 1138.2056026296468 | 1611.6932792526832 | 971.7662574542479 | 1009.455069640219 | 1287.1952431152959 |
| SE_18_12_15 | SE_18 | SE_18_12 | TGAAAAGCAAGACCAAGTAT | 1984.3953971312753 | 1897.8463283197689 | 1822.9745884233682 | 2157.569475701092 | 1970.5846046512377 | 1897.2191406118152 | 2150.1689043037654 | 2355.820898466013 | 1987.31105177273 | 1788.9830426051947 | 2401.564037771963 | 2083.4224178466693 |
| SE_18_12_23 | SE_18 | SE_18_12 | TCTCATGTTATATTTTAAGG | 22.759313675454276 | 49.96962423169481 | 43.451181955919935 | 41.641871666124814 | 57.27210368738796 | 56.31015714144657 | 8.60928490211718 | 30.25131169779792 | 29.303514168230603 | 5.103048613788249 | 133.8114859755639 | 33.659749369110536 |
| SE_18_12_24 | SE_18 | SE_18_12 | ATACAGTCTGTGGCTCCCAA | 266.60910305532155 | 384.76610658405 | 362.42235858687764 | 314.9166544750689 | 319.23894833155146 | 375.9785876828894 | 191.55658907210724 | 370.57856829802455 | 409.36121307740325 | 299.62185432385286 | 352.1354894093787 | 445.70150888753255 |
| SE_18_12_3 | SE_18 | SE_18_12 | AGAAGTTTATATTCTTACTG | 1.083776841688299 | 7.9951398770711695 | 5.925161175807264 | 0.0 | 2.1211890254588135 | 0.8663101098684087 | 0.0 | 0.0 | 4.439926389125849 | 1.4580138896537853 | 0.0 | 0.0 |
| SE_18_13_12 | SE_18 | SE_18_13 | TGGAGTTTAGGAAAAGCAAT | 54.188842084414944 | 95.94167852485404 | 102.70279371399258 | 96.29682822791364 | 98.63528968383483 | 64.10694813026224 | 50.57954879993843 | 81.30040018783191 | 160.72533528635574 | 54.67552086201695 | 36.3873339056358 | 47.58792152184593 |
| SE_18_13_16 | SE_18 | SE_18_13 | GTGAACCTAACCTGACTCAG | 475.77803350116324 | 480.7077851089041 | 476.97547465248476 | 446.3488119212 7534 | 461.35861303729195 | 525.8502366901241 | 325.00050505492356 | 342.21796358133895 | 454.64846224648693 | 481.14458358574916 | 326.31222018602426 | 485.16466332028284 |
| SE_18_13_21 | SE_18 | SE_18_13 | AACGGCCTACTTCCTGGCAG | 2439.5816706403607 | 2547.4514433318013 | 2415.4907060040946 | 2499.813608457055 | 2183.7641017098485 | 2735.8073269644347 | 2567.719222056449 | 2197.001512052574 | 2577.821261526468 | 3109.214619686697 | 2311.1825954902224 | 2294.666362163156 |
| SE_18_2_13 | SE_18 | SE_18_2 | TCCTCCCTCTTTCTACCCAA | 14.089098941947885 | 10.993317330972857 | 8.887741763710896 | 24.724861301761607 | 13.787728665482287 | 32.919784174999535 | 25.827854706351538 | 9.45353490556185 | 52.391131391685015 | 5.103048613788249 | 17.606774470468935 | 35.98111139456643 |
| SE_18_2_15 | SE_18 | SE_18_2 | GACTTTCTCTTCAGTCCGGA | 5.4188842084414945 | 21.986634661945715 | 32.58838646693995 | 16.917010364363207 | 11.666539640023474 | 32.919784174999535 | 9.685445514881827 | 29.305958207241734 | 55.94307250298569 | 53.21750697236316 | 10.564064682281362 | 8.124767089095647 |
| SE_18_2_17 | SE_18 | SE_18_2 | GCACTTTTTGAGACTCCTTT | 34.680858934025565 | 98.93985597875572 | 101.71526685135804 | 63.76411598875362 | 59.39329271284678 | 94.42780197565655 | 33.36097899570407 | 81.30040018783191 | 115.43808611727208 | 170.5876250894929 | 72.7746678112716 | 80.08698987822851 |
| SE_18_2_5 | SE_18 | SE_18_2 | AATGATGGAAGAAAAGCCAA | 144.14231994454374 | 194.88153450360974 | 182.69246958739063 | 188.68973098712806 | 203.6341464440461 | 268.5561340592067 | 162.50025252746178 | 177.72645622456278 | 301.02700918273257 | 83.83579865509266 | 154.93961534012664 | 131.15695443825828 |
| SE_18_2_6 | SE_18 | SE_18_2 | GATGGAAGAAAAGCCAAGGG | 507.2075619101239 | 768.5328206834662 | 721.8821365858516 | 560.8639590031186 | 722.264863168726 | 501.59355361380864 | 538.0803063823238 | 897.1404625378195 | 641.1253705897726 | 754.5221878958339 | 782.9145714535186 | 640.6959190258281 |
| SE_18_3_10 | SE_18 | SE_18_3 | AAAGGCTCTTCTTCTTCGGG | 99.7074694353235 | 108.93378082509469 | 127.39096527985618 | 104.10467916531204 | 115.60480188750533 | 103.09090307434064 | 81.7882065701132 | 107.7702979234051 | 111.88614500597139 | 68.5266528137279 | 104.46686185811568 | 123.03218734916264 |
| SE_18_3_12 | SE_18 | SE_18_3 | CATACCCTGGTGAGAAGACA | 940.7182985854434 | 989.3985597875572 | 908.5247136237805 | 1042.3481001426867 | 873.9298784890311 | 1178.1817494210359 | 1071.8559703135888 | 924.5557137639489 | 783.2030150417997 | 855.8541532267719 | 963.6774560169997 | 908.8132329659844 |
| SE_18_3_14 | SE_18 | SE_18_3 | CTGCTCTTAGTGTAGACAGT | 140.89098941947887 | 115.92952821753195 | 126.40343841722164 | 157.45832723753446 | 142.1196647057405 | 123.01603560131404 | 136.67239782111022 | 71.84686528227006 | 110.99815972814622 | 112.26706950334147 | 91.55522724643846 | 153.20989368008932 |

| | | | | | | | | | | | | | | | |
|---|---|---|---|---|---|---|---|---|---|---|---|---|---|---|---|
| SE_18_3_22 | SE_18 | SE_18_3 | TTCCTGCTGCGGAGGAGCTA | 39.01596630077876 | 27.982989569749094 | 48.38881626909266 | 18.218318853929606 | 34.999618920070425 | 0.8663101098684087 | 20.4470516425283 | 25.524544245016994 | 18.647690834328564 | 48.84346530340181 | 21.128129364562724 | 68.48017975094902 |
| SE_18_3_38 | SE_18 | SE_18_3 | ATAAGGACAAAACCACAGTC | 428.0918524668781 | 593.6391358725343 | 596.4662250312646 | 520.5233958265602 | 580.1451984629855 | 609.8823173473597 | 469.2060271653863 | 423.5183637691709 | 503.4876525268713 | 624.0299447718202 | 701.9234088893616 | 719.6222278913286 |
| SE_18_4_17 | SE_18 | SE_18_4 | GAGTCTTCCCTCAAAGGCGA | 115.96412206064798 | 105.935603371193 | 122.45333096668347 | 97.59813671748003 | 94.3929116329172 | 223.50800834604945 | 132.36775537005164 | 227.8301912240406 | 63.04695472558706 | 84.56480559991955 | 130.29013108147012 | 58.034050636397474 |
| SE_18_4_2 | SE_18 | SE_18_4 | CCTTGAGAGTAATCCAGTGA | 883.2781259759636 | 989.3985597875572 | 1024.0653565520222 | 890.0950068634179 | 930.1413876636897 | 1191.176401069062 | 844.7860810202483 | 1306.4785239486478 | 1096.6618181140848 | 758.8962295647952 | 1076.360812628001 | 1342.9079317262374 |
| SE_18_4_30 | SE_18 | SE_18_4 | GGAGTTGGAGAAAAGCAAGA | 346.80858934025565 | 510.68955964792093 | 459.199991125063 | 372.17422801599054 | 514.3883386737623 | 518.0534457013084 | 279.80175931880837 | 515.2176523531208 | 554.102813362906 | 405.3278613237523 | 390.87039324441037 | 579.1798253512468 |
| SE_18_4_31 | SE_18 | SE_18_4 | AGATTCCTAGGAAATATGGG | 18.42420630870108 | 33.97934447755247 | 32.58838646693995 | 16.917010364363207 | 27.575457330964575 | 0.8663101098684087 | 23.675533480822246 | 18.9070698111237 | 31.07948472388094 | 8.019076393095819 | 10.564064682281362 | 0.0 |
| SE_18_4_8 | SE_18 | SE_18_4 | AAGTGTGTCGACAGTGGCAG | 1176.9816500734926 | 1125.3159376977671 | 1153.4313755571475 | 1293.500638629002 | 1180.4416926678298 | 1638.1924177611609 | 1254.803274483579 | 778.0259227277403 | 1362.1694161838104 | 1063.6211325024365 | 842.777604653113 | 1121.2178582951992 |
| SE_18_5_0 | SE_18 | SE_18_5 | GTCAGCTTCTAATTATCTAT | 62.859056817921335 | 89.94532361705066 | 91.8399982250126 | 75.47589239485123 | 111.36242383658771 | 59.7753975809202 | 74.25508228076067 | 123.84130726286024 | 96.7903952829435 | 34.992333351690846 | 170.1988198811997 | 39.46315443275028 |
| SE_18_5_1 | SE_18 | SE_18_5 | GCTTCTAATTATCTATTGGC | 23.843090517142578 | 69.95747392437274 | 75.05204156022535 | 15.615701874796805 | 77.42339942924669 | 151.60426922697152 | 26.904015319116187 | 58.61191641448347 | 47.95120500255917 | 15.309145841364746 | 30.518409082146157 | 129.99627342553035 |
| SE_18_5_18 | SE_18 | SE_18_5 | TCTAAGCACTCAAGATGGAG | 1639.7543614743963 | 1771.922875255898 | 1687.6834082424357 | 1552.4610280527156 | 1560.1345282249574 | 1564.5560584223463 | 1272.0218442878133 | 1487.041040644879 | 1918.936185380192 | 1891.7730218257864 | 1750.113382364612 | 1702.7190456719018 |
| SE_18_5_19 | SE_18 | SE_18_5 | CCAGCTCTAAGCACTCAAGA | 250.35245042999705 | 208.8730292884843 | 253.79440369707783 | 209.51066682019047 | 238.63376536411653 | 164.59892087499765 | 272.2686350294558 | 392.32169858081676 | 261.0676716805999 | 129.7632361791869 | 410.82473764427516 | 297.13433925835506 |
| SE_18_5_8 | SE_18 | SE_18_5 | CCTGTGCCTTCCCTTAGCAA | 836.6757217833667 | 826.4975847922321 | 818.659769124037 | 744.3484560319811 | 833.6272870053137 | 804.8020920677517 | 697.3520770714915 | 638.1136061254249 | 694.4044872592827 | 846.3770629440223 | 631.4963110074858 | 735.87176206952 |
| SE_18_6_10 | SE_18 | SE_18_6 | CATCAACTAAATGAAACATG | 2120.951279184001 | 2088.730292884843 | 1925.6773821373608 | 2398.311546270876 | 2313.156632262836 | 2164.9089645611534 | 2367.5533480822246 | 2597.8313920483965 | 1878.088862600234 | 2448.0053207287056 | 2272.4476916551907 | 2170.4734938012652 |
| SE_18_6_16 | SE_18 | SE_18_6 | TGGCTTTGTGGACTTCAGGT | 9.75399157519469 | 36.977521931454156 | 25.675698428498144 | 24.724861301761607 | 21.211890254588134 | 46.78074593289407 | 24.75169409358689 | 40.650200093915956 | 24.863587779104755 | 8.019076393095819 | 31.692194046844083 | 75.44426582731671 |
| SE_18_6_17 | SE_18 | SE_18_6 | TGTCTCATAGCCACACGACC | 1468.517620487645 | 1330.1913970477158 | 1338.0988988698073 | 1354.6621376386229 | 1474.2263726938754 | 1509.112211390768 | 1335.5153204409276 | 1228.9595377230405 | 1433.208238409824 | 1501.025299398572 | 1274.730471661951 | 1452.0119469226647 |
| SE_18_6_23 | SE_18 | SE_18_6 | GGGGGCTCTGAATTGTTACA | 67.19416418467453 | 110.93256579436247 | 101.71526685135804 | 104.10467916531204 | 115.60480188750533 | 75.36897955855156 | 66.72195799140815 | 91.69928858394995 | 57.71904305863604 | 142.88536118607095 | 138.50662583435562 | 13.928172152735392 |
| SE_18_6_6 | SE_18 | SE_18_6 | GCTCAGCTGGGGTGAACGCA | 3743.3652111913843 | 3448.9034644715757 | 3544.2339099953783 | 3542.161708599742 | 3400.266007810478 | 2896.07469729009 | 3642.803674208332 | 3667.9715433579977 | 2981.85456293692 | 2549.3372860596437 | 3610.5625514108297 | 3101.3396660090807 |
| SE_18_7_13 | SE_18 | SE_18_7 | CGCATCTCGGGCCTGCAGCG | 800.9110860076529 | 253.84569109700962 | 489.81332386673387 | 730.0340626467506 | 180.30106716399914 | 2242.0105643394418 | 1196.690601394288 | 479.2942197119858 | 1953.5676112153735 | 54.67552086201695 | 315.7481555037429 | 689.444521560402 |
| SE_18_7_19 | SE_18 | SE_18_7 | CGCGGAGCCGGCGTCCGCAG | 665.4389807966155 | 1083.3414533431435 | 944.075680678624 | 792.4968701459379 | 1011.807165143854 | 1035.2405812927484 | 674.752704203434 | 795.0422855577516 | 1071.79823033498 | 789.5145212475247 | 819.3019053591545 | 1345.2292937516934 |
| SE_18_7_2 | SE_18 | SE_18_7 | GTTTGCGCAGTACCCCCGGA | 252.52000411337366 | 378.76975167624664 | 427.59913152075757 | 305.8074950481041 | 431.66196668086855 | 444.4170863624937 | 267.9639925783972 | 509.54553140978373 | 406.69725724392777 | 274.8356181997385 | 362.69955409166005 | 465.4330861039077 |
| SE_18_7_21 | SE_18 | SE_18_7 | TGCGGACGCCGGCTCCGCGG | 134.38832836934907 | 236.8560188582334 | 247.86924252127056 | 175.67664609146405 | 244.99733244049295 | 214.84490724736537 | 132.36775537005164 | 184.3439306584561 | 174.04511445373328 | 398.76679882031027 | 164.32989505771008 | 270.4386759656122 |
| SE_18_7_3 | SE_18 | SE_18_7 | TGTTTGCGCAGTACCCCCGG | 320.7979451397365 | 426.7405909386737 | 525.3642909215774 | 351.3532921829281 | 511.20655513557404 | 356.05345515591597 | 229.2222105188699 | 477.40351273087344 | 302.8029797383829 | 472.39650024782645 | 500.0323949613178 | 401.59563040387053 |
| SE_18_8_15 | SE_18 | SE_18_8 | AATTAAAAAGGAAGGGCGGG | 14.089098941947885 | 10.993317330972857 | 15.800429802152705 | 18.218318853929606 | 33.939024407341016 | 6.93048087894727 | 15.066248578705064 | 10.398888396118036 | 20.423661389978903 | 66.33963197924723 | 18.780559435166865 | 0.0 |
| SE_18_8_18 | SE_18 | SE_18_8 | TTAACAAGCCAAAGGAAAAG | 48.769957875973454 | 95.94167852485404 | 116.52816979087619 | 74.17458390528482 | 106.05945127294068 | 129.0802063703929 | 88.2451702467011 | 67.12009782948914 | 48.83919028038434 | 128.3052222895331 | 57.515463270198524 | 55.71268861094157 |
| SE_18_8_20 | SE_18 | SE_18_8 | GTAAAAGAAACCCTTCCCCC | 361.98146512389184 | 432.73694584647706 | 417.72386289441215 | 343.54544124552973 | 463.47980206275076 | 381.17644834209983 | 279.80175931880837 | 491.5838150892162 | 634.9094736449964 | 492.07968775815255 | 441.3431467264213 | 713.8188228276889 |
| SE_18_8_3 | SE_18 | SE_18_8 | AGTTGGATGTCATTAAAATG | 117.04789890233629 | 136.91677039484378 | 131.34107273039436 | 122.32299801924164 | 151.66501532030517 | 130.81282659012973 | 112.99686434028798 | 121.95060028174787 | 134.9737622294258 | 126.84720839987932 | 29.344624117448227 | 183.38760001101602 |
| SE_18_8_6 | SE_18 | SE_18_8 | GAAAATTTAAAACTTCCCTG | 0.0 | 1.9987849692677924 | 5.925161175807264 | 2.602616979132801 | 15.9089176909411 | 0.0 | 2.152321225529295 | 3.78141396222474 | 0.0 | 21.87020834480678 | 1.173784964697929 | 0.0 |
| SE_18_9_10 | SE_18 | SE_18_9 | GAGCTGTAAGCCCTTAAAAG | 65.02661050129794 | 82.94957622461338 | 98.75268626345441 | 70.27065843658562 | 90.15053358199957 | 51.978606592104526 | 54.88419125099702 | 86.02716764061283 | 79.91867500426528 | 29.8892847379026 | 111.50957164630326 | 51.069964560029774 |
| SE_18_9_11 | SE_18 | SE_18_9 | CCAAGACTCAGACATTTTAG | 330.5519367149312 | 334.7964823523552 | 321.93375721886133 | 282.3839422359089 | 384.99580812077465 | 272.02137449868036 | 319.61970199110027 | 286.4421076385241 | 536.3431078064026 | 164.02656258605086 | 376.7849736680352 | 269.2779949528843 |
| SE_18_9_5 | SE_18 | SE_18_9 | TACTGAAGAATAAATCAAGG | 189.6609472954523 | 293.8213904823655 | 333.7840795704759 | 253.7551554654481 | 264.0880336696223 | 234.77003977433876 | 262.583189514574 | 258.08150292183853 | 278.8273772371033 | 185.16776398603074 | 258.2326922335444 | 358.6504329329364 |
| SE_18_9_6 | SE_18 | SE_18_9 | TATTAGCCTGGTAAAATAAG | 97.53991575194691 | 116.92892070216585 | 97.76515940081985 | 85.88636031138243 | 91.21112809472898 | 185.39036351183947 | 136.67239782111022 | 156.9286794323267 | 106.55823333902038 | 60.50757642063209 | 149.070690516637 | 157.85261773100112 |
| SE_18_9_7 | SE_18 | SE_18_9 | AGTACTCTGAAACTACCAAC | 34.680858934025565 | 79.95139877071169 | 48.38881626909266 | 61.16149900962082 | 54.09032014919974 | 78.83421999802519 | 35.513300221233365 | 84.13646065950047 | 135.86174750725098 | 73.62970142751615 | 32.86597901154201 | 71.96222278913287 |
| SE_180_1_14 | SE_180 | SE_180_1 | ACAGGCCCTGACAGATACCA | 2516.52982640023 | 2357.566871251361 | 2287.112213861604 | 2590.9052027267035 | 2263.308690164554 | 2967.1121262992997 | 2822.76928728167 | 2246.1598935614957 | 2196.87557733947 | 2863.5392792800344 | 2510.7260394888704 | 2930.719557138072 |
| SE_180_1_19 | SE_180 | SE_180_1 | TGGACCACAGAGATCTCCAA | 97.53991575194691 | 145.91130275654885 | 138.25376076883617 | 162.66356119580004 | 169.69512203670507 | 171.52940175394494 | 138.82471904663953 | 280.76998669518696 | 174.93309973155846 | 215.05704872393332 | 232.40942301018995 | 269.2779949528843 |
| SE_180_1_25 | SE_180 | SE_180_1 | CCTTCTCTGGGTTTTCCATG | 99.7074694353235 | 137.91616287947767 | 127.39096527985618 | 135.33608291490566 | 142.1196647057405 | 304.0748485638115 | 100.08293698711222 | 245.7919075446081 | 73.70277805948909 | 110.0800486688608 | 375.6111887033373 | 206.601220265575 |
| SE_180_1_37 | SE_180 | SE_180_1 | GTAGTGCTGTGCACATGCCG | 964.5613891025861 | 924.4380482863539 | 972.7139596950259 | 978.5839841539331 | 942.8685218164426 | 1172.117578651957 | 1109.5215917603516 | 854.5995554627913 | 892.4252042142956 | 586.1215836408217 | 1157.351975192158 | 994.7036279078527 |
| SE_180_1_5 | SE_180 | SE_180_1 | GGCCTCTCTGGCTCTGAGAA | 6.5026610501297935 | 11.992709815606755 | 17.77548352742179 | 2.602616979132801 | 11.666539640023474 | 25.12299318618385 | 3.228481838293942 | 0.0 | 1.7759705556503396 | 0.0 | 1.173784964697929 | 31.338387343654635 |
| SE_180_2_2 | SE_180 | SE_180_2 | TTCCTTTTCGTTTTTTAAAG | 0.0 | 1.9987849692677924 | 0.0 | 2.602616979132801 | 0.0 | 0.0 | 0.0 | 0.0 | 0.0 | 0.0 | 0.0 | 0.0 |
| SE_180_2_22 | SE_180 | SE_180_2 | GTTAGACCTGGAAACCCACA | 227.59313675454277 | 260.8414384894469 | 241.94408134546327 | 265.4669318715457 | 246.05792695322236 | 277.2192351578908 | 230.29837113163455 | 181.50787018678753 | 265.50759806972576 | 377.6255974203304 | 244.14727265716925 | 133.47831646371418 |
| SE_180_2_32 | SE_180 | SE_180_2 | GGGTGGCCTAGAGAAAGCTG | 261.19021884688004 | 268.8365783665181 | 232.06881271911786 | 221.22244322628808 | 239.69435987684594 | 319.66843054144283 | 205.54667703804768 | 175.8357492434504 | 215.78042251151626 | 112.26706950334147 | 234.7569929395858 | 286.68821014380353 |
| SE_180_2_40 | SE_180 | SE_180_2 | CAGCATTTAGAGCATGGCCT | 994.9071406698584 | 1002.3906620877979 | 959.8761104807768 | 1090.4965142566436 | 860.1421498235488 | 730.2994226190685 | 934.107411879714 | 1089.0472211207252 | 919.9527478268759 | 820.8618198750811 | 845.1251745825089 | 966.8472836023819 |
| SE_180_2_7 | SE_180 | SE_180_2 | CCCTGGATACTGGTACTTGG | 468.19159560934514 | 485.70474753207355 | 471.0503134766775 | 452.8553543691074 | 549.3879575938327 | 384.64168878157346 | 453.06361797391656 | 359.2343264113503 | 491.05585863731886 | 488.4346530340181 | 410.82473764427516 | 691.7658835858579 |
| SE_180_3_0 | SE_180 | SE_180_3 | GCTGAAATGTTTGAAGTCCA | 864.8539196672625 | 934.431973132693 | 961.8511642060458 | 852.3570606659923 | 877.1116620272194 | 1180.7806797506412 | 812.5012626373089 | 814.8947088594315 | 1046.9346425558751 | 1277.220167336716 | 832.2135399708317 | 749.7999342222554 |
| SE_180_3_19 | SE_180 | SE_180_3 | TCTCTGCAGAGACTGCAGCA | 23.843090517142578 | 2.9981774539016888 | 0.987526862634544 | 5.205233958265602 | 0.0 | 3.465240439473635 | 21.52321225529295 | 1.89070698111237 | 16.871720278678225 | 2.187020834480678 | 5.868924823489645 | 1.1606810127279494 |
| SE_180_3_31 | SE_180 | SE_180_3 | GATCTTCTGCAGGTATCACC | 34.680858934025565 | 15.990279754142339 | 14.81290293951816 | 16.917010364363207 | 7.4241615891058474 | 82.29946043749882 | 13.990087965940416 | 17.961716320567515 | 13.319779167377547 | 0.0 | 19.954344399864794 | 10.446129114551544 |
| SE_180_3_32 | SE_180 | SE_180_3 | GCCTACCCAAAGGGAGGCAG | 479.0293640262281 | 437.7339082696465 | 393.0356913285485 | 385.18731291165454 | 432.7225611935979 | 353.4545248263108 | 467.053705939857 | 682.5452201815656 | 443.1046536347597 | 550.4002433443039 | 930.8114770054577 | 705.6940557385933 |
| SE_180_3_4 | SE_180 | SE_180_3 | TTTGCTTATCAAGAAAAGAG | 8.670214733506391 | 13.991494784874547 | 11.850322351614528 | 16.917010364363207 | 2.1211890254588135 | 12.128341538157722 | 6.456963676587884 | 2.836060471668555 | 12.431793889552377 | 16.038152786191638 | 10.564064682281362 | 19.73157721637514 |
| SE_180_4_11 | SE_180 | SE_180_4 | CCTCCAGATGAAAACAGCCT | 664.3552039549272 | 737.5516536598154 | 798.9092318713462 | 727.4314456676178 | 842.1120431071489 | 640.2031711927541 | 652.1533313353764 | 852.7088484816788 | 666.8769436467026 | 882.0984032405402 | 901.4668528880095 | 1017.9172481624116 |
| SE_180_4_16 | SE_180 | SE_180_4 | GCTGGGGCTGGAAAGTTTAA | 760.8113428651858 | 889.4593113241676 | 889.7617032337241 | 866.6714540512227 | 895.1417687436193 | 778.8127887716995 | 817.8820657011321 | 1025.7085372534607 | 673.9808258693039 | 1148.185938102356 | 876.817368629353 | 666.230901305843 |
| SE_180_4_29 | SE_180 | SE_180_4 | CAGCCTCTGCTGATCCCTGT | 767.3140039153157 | 672.5911421586121 | 616.2167622839555 | 730.0340626467506 | 646.9626527649381 | 531.914407459203 | 752.2362683224886 | 743.0478435771614 | 697.0684430927582 | 768.3733198475448 | 683.1428494541947 | 560.6089291475996 |
| SE_180_4_30 | SE_180 | SE_180_4 | CTACCCACAGGGATCAGCAG | 237.34712832973747 | 291.8226055130977 | 344.64687505945585 | 303.2048780689713 | 336.20846053522195 | 281.5507857072328 | 238.90765603375175 | 222.15807028070347 | 225.54826056759313 | 239.1142779032208 | 181.936669528179 | 208.9225822910309 |
| SE_180_4_5 | SE_180 | SE_180_4 | GCGTGGTGTGTGAGCTGCCG | 27290.584650553054 | 24059.37467507642 | 24471.903182946637 | 26105.54960919156 | 25755.477147120913 | 27268.8433283279 | 27303.27090645187 | 25479.167277470297 | 25512.705017194952 | 26247.89504849227 | 24968.753769054347 | 27002.083080103017 |
| SE_180_5_14 | SE_180 | SE_180_5 | ATCAGTAGCAGGATGGGCCA | 2060.2597760494564 | 1991.789221875355 | 1941.4778119395135 | 2056.067413514913 | 1807.2530496909092 | 1855.6362553381316 | 2216.890862295174 | 2320.842819315434 | 2118.732872890855 | 1741.5975911914466 | 2733.7451827814766 | 1522.8134886990697 |
| SE_180_5_18 | SE_180 | SE_180_5 | TCCCTGGATCCCATGTCCCA | 156.06386520311503 | 136.91677039484378 | 152.07913684571977 | 180.88188004972966 | 138.9378811675523 | 167.1978512046029 | 180.79498294446077 | 192.85211207346174 | 144.74160028550267 | 66.33963197924723 | 120.8998513638867 | 385.3460962256792 |
| SE_180_5_19 | SE_180 | SE_180_5 | CTGGTGTGAGAGAGATGAGG | 292.6197472558407 | 314.8086326596773 | 312.0584885925159 | 307.1088035376705 | 326.6631099206573 | 184.52405340197106 | 283.0302411571023 | 409.3380614108281 | 395.15344863220054 | 283.5837015376612 | 224.19292825730446 | 441.05878483662076 |
| SE_180_5_32 | SE_180 | SE_180_5 | CCCTGCCATGAAATGAAGCA | 74.78060207649263 | 88.94593113241676 | 103.69032057662713 | 106.70729614444484 | 108.18064029839948 | 32.919784174999535 | 125.91079169346375 | 66.17474433893295 | 80.80666028209045 | 259.5264723583738 | 30.518409082146157 | 64.99813671276517 |
| SE_180_5_33 | SE_180 | SE_180_5 | GGACCCTACCTGTCTGCCCG | 431.34318299194297 | 512.6883446171887 | 552.0275162127101 | 417.72002515081454 | 553.6303356447503 | 287.6149564763117 | 386.3416599825084 | 574.7749222581605 | 436.0007714121584 | 375.4385765858497 | 566.9381379490998 | 424.8092506584295 |
| SE_181_1_11 | SE_181 | SE_181_1 | GTACCCAAGGAGTCTTCCCA | 32.51330525064897 | 59.96354907803377 | 51.35139685699629 | 29.93009526002721 | 36.06021343279983 | 97.89304241513018 | 33.36097899570407 | 38.75949311280358 | 83.47061611556596 | 57.59154864132452 | 32.86597901154201 | 116.06810127279495 |
| SE_181_1_20 | SE_181 | SE_181_1 | GTTACCTGTGTGCATTTTCG | 298.0386314642822 | 257.8432610355452 | 250.8318231091742 | 297.9996441107057 | 267.2698172078105 | 404.5668213085469 | 249.66926216139822 | 278.8792797140746 | 521.2473580833747 | 150.90443757916677 | 379.1325435974311 | 312.2231924238184 |
| SE_181_1_7 | SE_181 | SE_181_1 | TGTTGTGGCCTCTTTCCAGA | 1424.0827699784247 | 1826.8894619107623 | 1875.3135121429991 | 1635.7447713849654 | 1790.2835374872386 | 1868.6309069861577 | 1403.3134390451003 | 1646.8057805488743 | 1766.2027175942626 | 1837.0975009637696 | 1468.4049908371092 | 2114.760805190324 |
| SE_181_1_8 | SE_181 | SE_181_1 | TGAATTTATAGCCTGATGCA | 41.18351998415536 | 120.92649064070145 | 88.87741763710896 | 71.57196692615203 | 132.57431409117584 | 131.67913669999814 | 83.94052779564251 | 98.31676301784324 | 98.56636583859384 | 412.6179307720212 | 92.72901221113639 | 304.0984253347228 |
| SE_181_1_9 | SE_181 | SE_181_1 | CGTAGCAGCAGCAATTTCAA | 12365.893763663491 | 12013.697057784066 | 11580.727518115298 | 12073.540166197063 | 12070.626149373378 | 12464.469860786665 | 12292.982679610568 | 12571.310717416149 | 12276.396465932972 | 10901.569852941353 | 11963.216360201293 | 12775.615907096539 |
| SE_181_2_10 | SE_181 | SE_181_2 | GCTACCTACCTCATAGAAAA | 195.07983150389381 | 292.82199799773156 | 317.9836497683232 | 219.92113473672168 | 354.2385672516219 | 243.43314087302286 | 235.6791741954578 | 239.1744331107148 | 326.77858223966246 | 316.38901405487144 | 199.54344399864794 | 253.02846077469297 |
| SE_181_2_12 | SE_181 | SE_181_2 | TCACCTGCCAAGGGAACGGG | 1529.2091236221897 | 1472.1051298657292 | 1396.3629837652452 | 1483.4916781056966 | 1500.7412355121105 | 1315.925056890113 | 1421.6081694620993 | 1420.866296305946 | 1345.2976959051323 | 996.5524935783623 | 1415.5846674257025 | 1062.0231266460737 |
| SE_181_2_13 | SE_181 | SE_181_2 | ACTGTCACCTGCCAAGGGAA | 978.6504880445339 | 1048.3627163809572 | 1024.0653565520222 | 1041.0467916531204 | 947.1108998673602 | 1003.1871072276173 | 1134.2732858539384 | 883.905513670033 | 1132.1812292270915 | 1030.8158199852262 | 645.5817305838609 | 1057.380402595162 |
| SE_181_2_4 | SE_181 | SE_181_2 | TGAATCTCCCACCAAAAGTG | 30.34575156727237 | 32.97995199291857 | 32.58838646693995 | 22.12224432262881 | 50.90853661101153 | 38.11764483420998 | 33.36097899570407 | 21.743130282792254 | 69.26285167036325 | 36.45034724134463 | 41.08247376442752 | 12.767491140007444 |
| SE_181_2_6 | SE_181 | SE_181_2 | CGGACATCAGAGAATCTCTG | 2101.4432960336117 | 2473.496399468893 | 2496.4679087401273 | 2170.582560596756 | 2771.33346176194 | 2555.614824111806 | 1982.2878487124806 | 2736.7983551601556 | 2584.037158471244 | 2797.199647300787 | 3815.974920232967 | 2949.2904533417195 |
| SE_182_1_11 | SE_182 | SE_182_1 | GAGAGCGCCACGGAAGCCTG | 5619.38292415383 | 5614.586978673229 | 5490.649356248065 | 5509.74014482414 | 5619.029728440397 | 5433.49700909466 | 5133.286122887368 | 6313.070609934203 | 5618.282852799849 | 5480.674211208579 | 6383.042638027338 | 6711.057615593004 |
| SE_182_1_14 | SE_182 | SE_182_1 | TCAAAGGACCACGAGAGGCA | 299.1224083059705 | 394.760031430389 | 396.9857987790867 | 321.4231969229009 | 299.0876525896927 | 363.85024614473167 | 279.80175931880837 | 246.7372610351643 | 537.2310930842277 | 476.0415349719609 | 140.85419576375148 | 300.6163822965389 |
| SE_182_1_22 | SE_182 | SE_182_1 | TGAGCTACCGCACCCCGCCG | 251.43622727168534 | 307.81288526724 | 233.05633958175238 | 223.82506020542087 | 260.90625013143404 | 418.4277830664414 | 248.59310154863357 | 216.48594933736638 | 162.50130584200608 | 313.47298627556387 | 282.88217649220087 | 284.3668481183476 |
| SE_182_1_24 | SE_182 | SE_182_1 | TGGACTTCCGGTCCCTGCAA | 49.85373471766175 | 77.9526138014439 | 89.8649444997435 | 57.25757354092162 | 81.66577748016432 | 85.76470087697247 | 20.4470516425283 | 102.09817698006798 | 110.99815972814622 | 105.70600699989943 | 123.24742129328254 | 160.17397975645702 |
| SE_182_1_3 | SE_182 | SE_182_1 | GCTTGGCTTTGGTAACTAGG | 1592.068180440111 | 1713.958111147132 | 1812.1117929343884 | 1737.2468335711446 | 1615.2854428868866 | 1773.3367949006326 | 1940.3175848146593 | 1781.0459762078526 | 1759.9868206494866 | 1619.1244244605286 | 1738.375532717633 | 2575.55116724332 |
| SE_182_2_11 | SE_182 | SE_182_2 | GCAATCCTGCTGTTGTCAAT | 1551.968437297644 | 1709.9605412085964 | 1689.6584619677049 | 1493.9021460222277 | 1726.6478667234742 | 1811.4544397348427 | 1714.3238561340834 | 1709.1991109255825 | 2164.020122059939 | 1642.4526466949892 | 1691.4241341297156 | 1732.8967520028284 |
| SE_182_2_15 | SE_182 | SE_182_2 | ATTGCACAGGTCAAAGGAAG | 1238.7569300497257 | 1341.1847143786886 | 1323.2859959302891 | 1371.5791480029861 | 1289.6829274789586 | 1300.3314749124816 | 1374.2571025004547 | 1460.5711429093058 | 1159.7087728396718 | 1267.0140701091395 | 1079.8821675220947 | 1549.5091519918126 |
| SE_182_2_17 | SE_182 | SE_182_2 | GGCCATGTCCAGCTGAAGCA | 1400.239679461282 | 1316.1999022628413 | 1317.3608347544816 | 1385.8935413882164 | 1303.470656144441 | 1115.8074215105105 | 1300.0020202196943 | 1126.8613607429725 | 1337.3058284047056 | 1081.1172991782819 | 1045.8424035458547 | 1029.524058289691 |
| SE_182_2_3 | SE_182 | SE_182_2 | GGGCTCTGCAACAGACACAG | 98.6236925936352 | 125.92345306387092 | 109.61548175243439 | 161.36225270623365 | 83.78696650562313 | 52.844916701972934 | 99.00677637434757 | 276.9885727329622 | 64.82292528123739 | 107.8930278343801 | 413.172307573671 | 71.96222278913287 |
| SE_182_2_7 | SE_182 | SE_182_2 | GCTGTTGTCAATGGGCCCGG | 1020.9177848703775 | 1114.3226203667944 | 1225.520836529469 | 1212.8195122758852 | 1061.6551072421362 | 1355.7753219440597 | 1082.6175764412353 | 872.5612717833587 | 1139.2851114496927 | 1396.0482993434994 | 1625.6921761066317 | 1361.4788279298846 |
| SE_182_3_19 | SE_182 | SE_182_3 | CACACTTCTCTGTGTCCCAA | 122.46678311077778 | 172.89489984166403 | 165.9045129226034 | 156.15701874796804 | 138.9378811675523 | 181.05881296249743 | 138.82471904663953 | 167.32756782844476 | 126.09390945117411 | 516.8659238822669 | 111.50957164630326 | 96.3365240564198 |
| SE_182_3_4 | SE_182 | SE_182_3 | GTGTGTCAGCTGCTTCTCCA | 97.53991575194691 | 155.9052276028878 | 175.77978154894885 | 88.48897729051524 | 204.6947409567755 | 219.1764577967074 | 64.56963676587885 | 111.55171188562983 | 224.66027528976795 | 79.4617569861313 | 43.43004369382337 | 87.05107595459621 |
| SE_182_3_41 | SE_182 | SE_182_3 | AATGGGGGCCCCAAACCAGC | 292.6197472558407 | 604.6324532035072 | 566.8404191522283 | 363.0650685890257 | 564.2362807720444 | 492.93045251512456 | 234.60301358269314 | 457.55108942919355 | 553.2148280850807 | 430.11409744786664 | 312.2268006096491 | 601.2327645930778 |
| SE_182_3_6 | SE_182 | SE_182_3 | TCCGTCCAGCAGGTGCCGCA | 4840.147374979943 | 4274.401656779174 | 4179.21368266939 | 4714.6406576990685 | 4145.863950259251 | 4667.678871970987 | 4847.027399891972 | 4281.505958728962 | 4321.824347175101 | 4095.561016037483 | 4657.578739921382 | 4314.251324309788 |

| | | | | | | | | | | | | | | | |
|---|---|---|---|---|---|---|---|---|---|---|---|---|---|---|---|
| SE_182_3_8 | SE_182 | SE_182_3 | GCCCGTCTCCGCCCCACAGA | 78.03193260155751 | 101.9380334326574 | 108.62795488979984 | 62.46280749918722 | 74.24161589105847 | 77.96790988815678 | 20.4470516425283 | 101.15282348951179 | 137.6377180629013 | 47.38545141374802 | 102.11929192871983 | 97.49720506914775 |
| SE_182_4_1 | SE_182 | SE_182_4 | GGTAAACGATTCTCTTCCCA | 247.10111990493215 | 283.8274656360265 | 270.5823603618651 | 219.92113473672168 | 329.8448934588455 | 241.70052065328605 | 255.05006522522146 | 325.20160075132765 | 244.1959514019217 | 292.3317848753393 | 382.65389849152484 | 222.85075444376628 |
| SE_182_4_10 | SE_182 | SE_182_4 | ATACAGCCTCTAACAGCACT | 801.9948628493412 | 917.4423008939167 | 881.8614883326478 | 870.5753795199219 | 855.8997717726312 | 1218.898324584851 | 818.9582263138967 | 901.8672299906005 | 809.8425733765548 | 820.1328129302542 | 753.5699473360704 | 774.1742354895423 |
| SE_182_4_2 | SE_182 | SE_182_4 | GTAAACGATTCTCTTCCCAG | 107.2939073271416 | 112.93135076363026 | 145.16644880727796 | 96.29682822791364 | 117.72599091296435 | 133.41175691793496 | 77.48356411905462 | 86.02716764061283 | 62.15896944776188 | 91.12586810336158 | 134.98527094026184 | 106.78265317097134 |
| SE_182_4_20 | SE_182 | SE_182_4 | GTCTCACAGCACAAACCCCT | 1555.219767822709 | 1484.097839681336 | 1596.8309368800576 | 1517.3256988344228 | 1561.1951227376867 | 1649.4544491894503 | 1588.4130644406196 | 1509.7295244182274 | 1323.9860492373282 | 1737.952556467312 | 1471.926345731203 | 1912.8023089756607 |
| SE_182_4_27 | SE_182 | SE_182_4 | CCAAAGTCTGCACATGGCAT | 726.1304839311603 | 680.5862820356833 | 704.1066530584299 | 732.6366796258834 | 753.0221040378788 | 946.0106399763023 | 722.1037711650785 | 564.3760338620425 | 698.8444136484086 | 780.766437909602 | 559.8954281609122 | 760.2460633368069 |
| SE_183_1_1 | SE_183 | SE_183_1 | ATTGTGGATGACAAGAGGGA | 240.59845885480235 | 288.824428059196 | 262.6821454607887 | 326.62843088116654 | 276.81516782237514 | 170.66309164407653 | 257.2023856507574 | 309.1305914117376 | 332.99447918443866 | 191.72882648947277 | 212.45507861032516 | 287.84889115653147 |
| SE_183_1_10 | SE_183 | SE_183_1 | GGAGACTCTGGAGTGCGTAA | 622.0879071290835 | 777.5273530451713 | 788.0464363823661 | 676.6804145745282 | 928.0201986382309 | 895.7646536039347 | 654.3056525609056 | 741.1571365960491 | 707.7242664266603 | 670.6863892407413 | 1002.4123598520314 | 805.5126228331969 |
| SE_183_1_13 | SE_183 | SE_183_1 | TGCCAACTCACAAGGAGGCA | 720.7115997227188 | 698.5753467590935 | 641.8924607124536 | 711.8157437928211 | 628.9325460485383 | 801.336851628278 | 750.0839470969593 | 734.5396621621558 | 750.3475597622685 | 662.6673128476455 | 684.3166344188926 | 647.6600051021958 |
| SE_183_1_23 | SE_183 | SE_183_1 | AGGTAAGGCAGACAAATGGA | 3220.9847734976242 | 3756.7163497388156 | 3562.0093935228 | 3404.2230087057037 | 3304.8125016648314 | 3505.0907045275817 | 3457.7040488128123 | 3318.1907518522094 | 3729.5381668657133 | 3648.679758858598 | 3088.2282421202513 | 3964.8863394786754 |
| SE_183_1_31 | SE_183 | SE_183_1 | ACAGCTGGTTTCCATAGGCA | 609.082585028824 | 640.6105826503275 | 690.2812769815463 | 595.9992882214113 | 820.9001528525608 | 1039.5721318420906 | 559.6035186376167 | 845.1460205572293 | 549.6628869737801 | 251.50739595627796 | 832.2135399708317 | 627.9284278858206 |
| SE_183_2_17 | SE_183 | SE_183_2 | GGGCGCCGTGGATCAGGGGA | 1286.4431110840108 | 1588.034658083261 | 1449.6894343475105 | 1389.7974668569157 | 1419.0754580319463 | 1736.9517702861594 | 1418.3796876238055 | 1356.5822589481254 | 1427.880326742873 | 1496.6512577296107 | 1694.9454890238096 | 1295.3200102043916 |
| SE_183_2_19 | SE_183 | SE_183_2 | GTAATCATGTCTCTGCACCA | 606.9150313454473 | 627.6184803500868 | 650.7802024761645 | 642.8463938458018 | 651.2050308158557 | 624.6095892151227 | 552.0703943482641 | 553.9771454659244 | 668.6529142023528 | 445.42324328923144 | 666.7098599484237 | 517.6637316766654 |
| SE_183_2_2 | SE_183 | SE_183_2 | GCCGCAGTCTCACTGCACCC | 2440.6654474820493 | 1939.8208126743925 | 2049.118239966679 | 2239.551910543775 | 2066.038110796884 | 1939.6683359953672 | 2400.9143270779286 | 2090.176567619725 | 1839.0175103759266 | 2037.574410791165 | 1732.5066078941431 | 1773.5205874483067 |
| SE_183_2_20 | SE_183 | SE_183_2 | GATTACCCATCTGCAAAGAG | 82.36703996831072 | 88.94593113241676 | 86.90236391183987 | 113.21383859227684 | 80.60518296743491 | 42.44919538355203 | 72.10276105523138 | 126.67736773452879 | 102.11830694989453 | 86.75182643440023 | 76.29602270536539 | 8.124767089095647 |
| SE_183_2_6 | SE_183 | SE_183_2 | GCCAGCAGTAGCAACCCCCT | 2270.5124833369864 | 2178.6756165018937 | 2213.047699164013 | 2225.237517158545 | 2312.096037750107 | 2577.272576858516 | 2253.4803231291717 | 2261.2855494103947 | 2276.7942523437355 | 2107.5590774945467 | 2328.789369960691 | 2604.5681925615186 |
| SE_183_3_11 | SE_183 | SE_183_3 | GACTACACCAGATAAGTGG | 132.22077468597246 | 222.86452407335884 | 224.1685978180415 | 167.86879515406565 | 162.27096044759924 | 186.25667362170788 | 129.1392735317577 | 93.58999556506231 | 129.64585056247478 | 177.8776945377618 | 22.30191432926065 | 111.42537722188314 |
| SE_183_3_13 | SE_183 | SE_183_3 | GAGCCAGTAAAAGCCCCACG | 484.4482482346696 | 488.70292498597524 | 483.88816269092655 | 532.2351722326578 | 501.6612045210094 | 430.5561246045991 | 481.04379390579743 | 761.0095598977289 | 764.5553242074711 | 406.0568682685792 | 903.8144228174053 | 603.5541266185337 |
| SE_183_3_19 | SE_183 | SE_183_3 | GGACCTTAAGAGGAGAGAAG | 108.37768416882989 | 125.92345306387092 | 110.60300861506893 | 89.79028578008163 | 106.05945127294068 | 81.43315032763041 | 66.72195799140815 | 130.45878169675353 | 162.50130584200608 | 64.88161808959345 | 159.63475519891836 | 27.856344305470785 |
| SE_183_3_4 | SE_183 | SE_183_3 | ACTGAACCCTCCTCCAACAA | 593.9097092451879 | 638.6117976810597 | 623.1294503223972 | 687.0908824910595 | 643.7808692267499 | 584.7593241611759 | 644.6202070460239 | 757.2281459355042 | 749.4595744844434 | 688.1825559165867 | 595.10897710185 | 710.336779789505 |
| SE_183_4_12 | SE_183 | SE_183_4 | TCATGATGTTTGCAGCAGGG | 1672.2676667250453 | 1847.8767040880741 | 1798.2864168575047 | 1802.3122580494646 | 1746.7991624655333 | 1740.4170107256332 | 1537.8335156406813 | 1794.280925075639 | 1798.1701875959689 | 2179.730765032409 | 1973.1325256572186 | 2180.919622915817 |
| SE_183_4_14 | SE_183 | SE_183_4 | GCAGATACCGGGGTACATGA | 1096.7821637885586 | 1331.1907895323498 | 1306.4980392655018 | 1315.6228829516308 | 1462.559833053852 | 1347.978530955244 | 1038.4949913178848 | 1401.013873004266 | 1347.0736664607825 | 1398.964327122807 | 1210.1722986035647 | 1428.7983266681058 |
| SE_183_4_17 | SE_183 | SE_183_4 | AAGCTCTCAGATAGCTTAAA | 55.22761892610324 | 49.96962423169481 | 48.38881626909266 | 37.737946197425615 | 56.21150917465856 | 15.593581977631358 | 82.86436718287786 | 40.650200093915956 | 58.607028336461205 | 95.49990977232294 | 35.21354894093787 | 45.26655949639003 |
| SE_183_4_2 | SE_183 | SE_183_4 | ATAATTGTTCCAAAACAAGG | 15.172875783636185 | 25.984204600481302 | 16.78795666478725 | 7.807850937398403 | 7.4241615891058474 | 28.588233625657487 | 10.761606127646475 | 4.726767452780925 | 26.639558334755094 | 0.0 | 10.564064682281362 | 1.1606810127279494 |
| SE_183_4_3 | SE_183 | SE_183_4 | CTAGGATTCAAACTGCCCAG | 148.47742731129696 | 197.87971195751143 | 228.11870526857967 | 161.36225270623365 | 254.5426830550576 | 317.935810321706 | 206.6228376508123 | 296.8409960346421 | 86.13457194904147 | 562.7933614063611 | 264.10161705703405 | 112.5860582346111 |
| SE_183_5_11 | SE_183 | SE_183_5 | GTACCCTATGAAGTAAAAGA | 13.005322100259587 | 19.987849692677923 | 22.713117840594514 | 13.013084895664004 | 38.18140245825846 | 37.251334724341575 | 9.685445514881827 | 49.15838150892162 | 32.855455279531284 | 7.290069448268927 | 61.03681816429231 | 40.62383544547823 |
| SE_183_5_3 | SE_183 | SE_183_5 | CTTGTATTTAGACTTCCTTA | 222.1742525461013 | 198.87910444214535 | 195.5303188016397 | 195.19627343496006 | 203.6341464440461 | 231.30479933486512 | 162.50025252746178 | 227.8301912240406 | 166.94123223113192 | 85.29381254476444 | 292.2724562097843 | 121.87150633643469 |
| SE_183_5_6 | SE_183 | SE_183_5 | TTAACATCTTTGTTTCCTAA | 18.42420630870108 | 29.981774539016886 | 28.638279016401 78 | 37.737946197425615 | 37.120807945529236 | 0.8663101098684087 | 23.675533480822246 | 85.08181415005664 | 35.51941111300679 | 50.30147919305569 | 44.60382865853213 | 0.0 |
| SE_183_5_7 | SE_183 | SE_183_5 | ATAGCATGTGGCTTTTATGA | 138.72343573610226 | 254.84508358164354 | 249.84429624653964 | 169.17010364363205 | 208.93711900769313 | 274.62030482828555 | 156.0432888508739 | 200.41493999791123 | 387.161581131774 | 142.15635424124406 | 237.10456286898167 | 405.07767344205433 |
| SE_183_5_8 | SE_183 | SE_183_5 | TGGGAACTCCAAATAGCATG | 174.48807151181612 | 183.8882171726369 | 217.25590977959968 | 149.65047630013606 | 156.9679878839522 | 116.95186483223517 | 216.30828316569415 | 138.96696311175918 | 166.05324695330674 | 108.622034779207 | 228.88806811609615 | 51.069964560029774 |
| SE_184_1_11 | SE_184 | SE_184_1 | GTCAACAGATGTTTGCCGGG | 163.65030309493312 | 223.86391655799275 | 217.25590977959968 | 195.19627343496006 | 265.1486281823517 | 360.38500570525804 | 189.40426784527713 | 261.8629168840632 | 251.29983362452305 | 284.3127084824881 | 359.1781991975663 | 206.601220265575 |
| SE_184_1_12 | SE_184 | SE_184_1 | TTTGGGGGCCGTGTTTTGAA | 411.83519984155356 | 551.6646515179107 | 545.1148281742683 | 527.0299382743922 | 596.0541161539265 | 717.3047709710424 | 609.1069068247905 | 725.086127256594 | 851.5778814343378 | 748.6901323372188 | 663.1885050543299 | 601.2327645930778 |
| SE_184_1_14 | SE_184 | SE_184_1 | GGCCATATCCATTCAAAACA | 457.3538271924621 | 463.7181128701278 | 523.3892371963084 | 568.6718099405169 | 497.4188264700917 4 | 519.7860659210453 | 631.706279692848 | 425.40907075028326 | 648.2292528123739 | 497.9117433167677 | 549.3313634786308 | 529.270541803945 |
| SE_184_1_6 | SE_184 | SE_184_1 | GCAAACATCTGTTGACAGGA | 826.9217302081721 | 1011.3851944495029 | 1026.0404102772914 | 995.5009945182964 | 958.7774395073837 | 843.7860470118301 | 985.763121292417 | 892.4136950850386 | 1300.8984320138738 | 960.1021463370176 | 1211.3460835682627 | 997.0249899333086 |
| SE_184_1_9 | SE_184 | SE_184_1 | CTGTCAACAGATGTTTGCCG | 557.0612966277856 | 594.6385283571682 | 575.7281609159392 | 653.2568617623331 | 532.4184453910622 | 742.4277641572263 | 618.7923523396723 | 627.7147177293068 | 485.7279469703679 | 655.3772433993765 | 413.172307573671 | 363.2931569838482 |
| SE_184_2_1 | SE_184 | SE_184_2 | AGCTCCACGTCAGAGACCAG | 481.1969177096047 | 514.6871295864565 | 475.00042092721566 | 528.3312467639586 | 567.4180643102326 | 356.9197652567844 | 535.9279851557944 | 445.26149405196315 | 467.9682414138644 6 | 970.3082435645941 | 515.2915995023908 | 573.3764202876071 |
| SE_184_2_20 | SE_184 | SE_184_2 | CTCTACAACCACCAGCAGCG | 44.434850509220254 | 44.97266180852533 | 32.58838646693995 | 54.65495656178882 | 29.69664635642339 | 19.9251325269734 | 80.71204595734856 | 94.5353490556185 | 57.71904305863604 | 17.496166675845423 | 100.9455069640219 | 95.17584304369186 |
| SE_184_2_24 | SE_184 | SE_184_2 | CCTGGATGCAGGGGGAGCCA | 4288.504962560599 | 3914.6203623109714 | 3929.3693864228508 | 4280.003622183891 | 4612.52553586019 | 4892.053190426904 | 4398.268424369115 | 4403.4565590107095 | 4439.926389125849 | 3946.8435992927966 | 4648.188460203799 | 4942.179752195609 |
| SE_184_2_37 | SE_184 | SE_184_2 | CTTGCCCTCAGACTGCCCGC | 3706.516798573982 | 2919.225447615611 | 2826.301880860065 | 3589.0088142241325 | 2846.6356721657276 | 3010.42763179272 | 3840.817226957027 | 2671.568964311779 | 2658.6279218085583 | 3373.1151337140323 | 2660.970514970205 | 2702.065397630666 |
| SE_184_2_4 | SE_184 | SE_184_2 | GGAGCTGGAGCGCTTCCTGT | 464.9402650842802 | 632.6154427732563 | 631.0296652234737 | 510.112927910029 | 516.509527699221 | 541.4438186677554 | 478.8914726802681 | 780.8619831994088 | 651.7811939236747 | 901.7815907508663 | 693.706914136476 | 646.4993240894678 |
| SE_184_3_16 | SE_184 | SE_184_3 | GGAACACTGCAGGGGAGGCA | 210.25270728753 | 64.96051150120326 | 101.71526685135804 | 148.34916781056964 | 44.54496953463509 | 397.6363404295996 | 400.3317479484489 | 151.2565584889896 | 297.47506807143185 | 23.328222234460565 | 41.08247376442752 | 118.38946329825085 |
| SE_184_3_20 | SE_184 | SE_184_3 | ACAGAGTTGGGAACACTGCA | 304.541292514412 | 71.95625889364052 | 155.0417174336234 | 271.9734743193777 | 77.42339942924669 | 704.3101193230164 | 528.3948608674419 | 158.81938641343908 | 608.2699153102413 | 26.244250013768134 | 59.86303319959438 | 118.38946329825085 |
| SE_184_3_28 | SE_184 | SE_184_3 | GCCGAGTCAGAAGATATCCA | 1399.155902619594 | 1665.987271884705 | 1638.3070651107087 | 1708.6180468006837 | 1743.6173789271447 | 1894.6202102822099 | 1680.9628771383793 | 1816.9694088489875 | 2157.8042251151624 | 1518.5214660744175 | 1819.36669528179 | 2266.810017857685 |
| SE_184_3_34 | SE_184 | SE_184_3 | CTTCTCCACATAGTCTTCCA | 24.926867358830876 | 18.98845720804403 | 25.675698428498144 | 16.917010364363207 | 15.9089176909411 | 0.0 | 8.60928490211718 | 0.945353490556185 | 0.8879852778251698 | 22.599215289633673 | 0.0 | 25.534982280014887 |
| SE_184_3_37 | SE_184 | SE_184_3 | GGAGAAGGAGTTATCTCTGG | 566.8152882029804 | 891.4586092934353 | 843.3479406899906 | 641.5450853562354 | 909.990091921831 | 1087.219187884853 | 533.7756639312652 | 838.5285461233361 | 1085.1180095023574 | 843.4610351647148 | 901.4668528880095 | 1144.4314785497581 |
| SE_184_4_11 | SE_184 | SE_184_4 | GGTTAAAATTGAAATCCTAA | 22.759313675454276 | 41.97448435462364 | 53.32645058226538 | 32.53271223916001 | 19.090701229129323 | 15.593581977631358 | 15.066248578705064 | 5.67212094333711 | 39.95933750213264 | 10.206097227576498 | 1.173784964697929 | 76.60494684004466 |
| SE_184_4_12 | SE_184 | SE_184_4 | CCAGAACTCTCCCATCTTGT | 1115.2063700972597 | 1036.3700065653504 | 1056.6537430189621 | 1030.6363237365892 | 1231.3502292788412 | 969.4010129427494 | 1101.988467470999 | 1102.2821699885117 | 1116.1974942262384 | 702.0336878682976 | 870.9484438058633 | 1245.4107266570898 |
| SE_184_4_22 | SE_184 | SE_184_4 | TTAGCCAGAACTTAGGCACC | 1000.3260248782999 | 913.4447309553811 | 912.4748217014387 | 1015.0206218617924 | 1132.7149395950064 | 811.732572946699 | 881.3755418542463 | 831.9110716894428 | 886.2093072695194 | 916.3617296474041 | 934.3328318995515 | 432.93401774752516 |
| SE_184_4_24 | SE_184 | SE_184_4 | ATATCCATTTTACATTAGGG | 7.586437891818092 | 14.990887269508443 | 15.800429802152705 | 10.410467916531204 | 8.484756101835254 | 38.98395494407839 | 2.152321225529295 | 0.945353490556185 | 20.423661389978903 | 0.7290069448268927 | 10.564064682281362 | 90.53311899278006 |
| SE_184_4_5 | SE_184 | SE_184_4 | GCTCAGCCTCTGGAGTAGTT | 766.2302270736274 | 818.5024449161935 | 816.6847153987679 | 819.8243484628323 | 934.3837657146073 | 917.4224063506448 | 827.5675112600139 | 743.9931970677176 | 868.449601713016 | 755.2511948406608 | 813.4329805356648 | 1290.6772861534798 |
| SE_185_1_13 | SE_185 | SE_185_1 | CTACCACAGGGTGATAATGA | 179.90695572025763 | 272.83414830505365 | 297.24558565299776 | 230.33160265325287 | 276.81516782237514 | 607.2833870177545 | 136.67239782111022 | 262.80827037461944 | 258.40371584712443 | 92.58388199301537 | 288.75110131569056 | 696.4086076367697 |
| SE_185_1_22 | SE_185 | SE_185_1 | ACCCTTCCTACAGCTCCATG | 192.9122778205172 | 154.90585311825392 | 202.44300684008152 | 161.36225270623365 | 219.54306413498956 | 252.96255208157535 | 156.0432888508739 | 178.67180971511897 | 160.72535528635574 | 102.78997920259186 | 238.27834783336796 | 237.93960760922963 |
| SE_185_1_26 | SE_185 | SE_185_1 | TGAGCAGGCTGCCACCCCAG | 2277.015144387116 | 2653.387046702994 | 2538.931563833413 | 2458.1717367909305 | 2339.6714950810715 | 2375.422321259177 | 2157.702028593118 | 2281.1379727120743 | 2869.968417930949 | 1734.3075217431776 | 2321.7466601725037 | 2096.1899089866765 |
| SE_185_1_29 | SE_185 | SE_185_1 | TTAAGTCGCAGTCTGCCTAG | 656.7687660631092 | 704.5717016668968 | 586.5909564049192 | 746.9510730111139 | 791.2035064961374 | 549.2406096565711 | 701.6567195225501 | 681.5998666910094 | 693.5165019814576 | 875.5373407370981 | 780.5670015241228 | 763.7281063749907 |
| SE_185_1_4 | SE_185 | SE_185_1 | TGACCAAATACACCTAGCCA | 1096.7821637885586 | 1175.285561929462 | 1253.1715886823365 | 1205.0116613384869 | 1381.954650086417 | 1056.0320239295902 | 1109.5215917603516 | 1211.9431748930292 | 1070.9102450571547 | 1323.8766118056371 | 1271.209116767857 | 1558.794600093636 |
| SE_185_2_0 | SE_185 | SE_185_2 | TATTAACTAAAGTCACATTA | 56.35639576779154 | 75.95382883217611 | 70.11440724705263 | 68.96934994701923 | 80.60518296743491 | 15.593581977631358 | 89.32133085946575 | 139.91231660231537 | 56.83105778080866 | 66.33963197924723 | 254.7113373394506 | 32.49906835638259 |
| SE_185_2_12 | SE_185 | SE_185_2 | TGCAAAAGACACACAAACCT | 4772.953210795268 | 4538.241272722523 | 4636.438620069184 | 4580.60588327373 | 4767.372334718683 | 4128.833983632836 | 4989.080600776906 | 5888.6068926744765 | 4656.59479691519 | 4433.820238437161 | 6476.945435203173 | 4814.504840795535 |
| SE_185_2_2 | SE_185 | SE_185_2 | CTCCGATGCACAAAGAGCCA | 239.51468201311405 | 205.8748518345826 | 227.13117840594512 | 219.92113473672168 | 239.69435987684594 | 196.65239490401287 7 | 271.19247441669114 | 247.68261452572048 | 170.4931733424326 | 309.09894460660246 | 247.66862755126303 | 164.8167038073688 |
| SE_185_2_21 | SE_185 | SE_185_2 | ACACTTTTTTCTTAGAAACA | 35.76463577571386 | 67.95868895510495 | 79.00214901076352 | 22.12224432262881 | 86.96875004381135 | 26.85561340592067 | 23.675533480822246 | 82.2457536783881 | 64.82292528123739 | 8.019076393095819 | 127.94256115207426 | 95.17584304369186 |
| SE_185_2_23 | SE_185 | SE_185_2 | ATAATAATAGAGCAAAGTGT | 62.859056817921335 | 60.96294156266758 | 58.26408489548096 | 85.88636031388243 | 54.09032014919974 | 51.11229648223612 | 81.7882065701132 | 46.32232103725307 | 55.94307250298569 | 34.99233335180946 | 57.515463270198524 | 47.58792152184593 |
| SE_186_1_22 | SE_186 | SE_186_1 | AGCTTTGCTTTGTGGCCCAT | 118.13167574402458 | 141.91373281801324 | 151.09160998308525 | 132.73346593577284 | 171.8163110621639 | 154.20319955657675 | 135.59623720834557 | 143.69373056454012 | 124.31793889552377 | 373.25155575136904 | 100.9455069640219 | 190.3516860873837 |
| SE_186_1_24 | SE_186 | SE_186_1 | CGAGAAAGCCAACATCCGAT | 913.6238775243236 | 907.4483760476799 | 852.2356824536115 | 896.6015493112499 | 914.2324699727486 | 878.4384514065664 | 928.7266088158908 | 924.5557137639489 | 791.1948825422263 | 742.1290698377767 | 1272.3829017325552 | 745.1572101713435 |
| SE_186_1_32 | SE_186 | SE_186_1 | ATTTAAGCGCCACCTCCCCA | 126.80189047753097 | 207.87363680385042 | 181.7049427247561 | 147.04785932100324 | 191.9676068040226 | 123.88234571118245 | 192.6327496848719 | 215.5405958468102 | 179.3730261206843 | 157.46550008260883 | 191.32694924576242 | 131.15695443825828 |
| SE_186_1_33 | SE_186 | SE_186_1 | GATCCAATGGTAGCCAGCCA | 390.15966300778763 | 467.7156828086634 | 507.58880739415565 | 421.62395062196131 | 430.60137261218316 | 250.36362175970013 | 319.61970199110027 | 594.6273455598404 | 432.4488303008577 | 303.99589599258142 | 407.30338275018136 | 301.77706330926685 |
| SE_186_1_37 | SE_186 | SE_186_1 | GCAGCCCTCACAGTGCCTGC | 2498.105620091529 | 2312.5942094428356 | 2237.7358707298768 | 2480.2939811135593 | 2298.3083090846244 | 2658.7057271861463 | 2710.848583554147 | 2393.6350380882604 | 2339.8412070693225 | 2254.089473404752 | 2619.8880412057774 | 2435.1087647032377 |
| SE_186_2_14 | SE_186 | SE_186_2 | GGAGAGCCTCTTCTGCTCTG | 1095.6983869468702 | 1235.2491110074957 | 1227.4958902547382 | 1175.0815660784597 | 1309.8342232208174 | 1600.9410830368192 | 1111.6739129858809 | 1047.451667536253 | 1336.4178431268806 | 1202.8614589643728 | 1031.7569839694797 | 1330.14044058623 |
| SE_186_2_21 | SE_186 | SE_186_2 | AGTGGCATTTAGTCACAGAG | 448.68361245895574 | 543.6695116408396 | 532.2769789600193 | 553.0561080657202 | 516.509527699221 | 387.2406191711787 | 562.8320004759106 | 491.5838150892162 | 413.8011394665291 | 507.3888335995173 | 625.6273861839961 | 392.3101823020469 |
| SE_186_2_25 | SE_186 | SE_186_2 | AAGAAAATGGAGATTTTCAG | 2.167553683376598 | 16.989672238776237 | 20.738064115325425 | 3.9039254686992013 | 11.666539640023474 | 6.064170769078861 | 2.152321225529295 | 1.89070698111237 | 4.439926389125849 | 2.9160277793075706 | 10.564064682281362 | 0.0 |
| SE_186_2_41 | SE_186 | SE_186_2 | ATCTGATTCCCTTTGAGCAG | 76.94815575986922 | 86.94714616314897 | 80.97720273603261 | 87.18766880094883 | 82.72637199289373 | 83.16577054736723 | 89.32133085946575 | 113.23494886778659 | 60.38298892111544 | 69.25565975855548 | 32.86597901154201 | 71.96222278913287 |
| SE_186_2_5 | SE_186 | SE_186_2 | CTTCTTTTGGGCTCCAAGAG | 15.172875783636185 | 38.97630690072195 | 25.675698428498144 | 33.834020728726415 | 13.787728665482287 | 110.88769406315632 | 15.066248578705064 | 0.945353490556185 | 23.087617223454416 | 23.328222234460565 | 0.0 | 26.695663292742836 |
| SE_186_3_11 | SE_186 | SE_186_3 | GTTTCCTCATCTGCAAGACA | 32.51330525064897 | 20.98724217731182 | 28.638279016401 78 | 23.423552812195208 | 44.54496953463509 | 16.459892087499767 | 21.52321225529295 | 56.72120943333711 | 35.51941111300679 | 36.45034724134463 | 44.60382865853213 | 85.89039494186825 |
| SE_186_3_22 | SE_186 | SE_186_3 | ACAGATACGTACAATCACCT | 2476.430083257763 | 2095.7260402772804 | 2069.8563040820045 | 2305.9186435116617 | 2314.2172267755655 | 2031.4972076414185 | 2511.7588701926887 | 2408.7606939371594 | 2012.1746395518346 | 1736.4945425776582 | 2078.7731724800324 | 1838.518724161072 |
| SE_186_3_27 | SE_186 | SE_186_3 | CGTGCCTCCGCAATCTCGTG | 20191.8463374947 | 16112.205637267674 | 16356.407425815953 | 18764.868419547493 | 16538.91083150237 | 17513.32518109975 | 20448.127803141066 | 15319.453314462979 | 16225.266996421502 | 19083.21479473357 | 16166.540318784577 | 16268.10507439494 |
| SE_186_3_47 | SE_186 | SE_186_3 | CGCAGCAGCAGCTTTCCAGA | 1744.8807151181613 | 2176.676831532626 | 2220.9479140650897 | 1970.1810532035302 | 2490.275915888647 | 2081.7431940137863 | 1852.0724145679583 | 1964.4445533757525 | 2330.9613542910706 | 3249.9129600382876 | 2038.8644836803028 | 1570.4014102209155 |
| SE_186_3_8 | SE_186 | SE_186_3 | GGAGCCCTGAGCAGAGGCGC | 839.9270523084316 | 804.5109501302865 | 832.4851452009207 | 969.4748247269683 | 846.3544211580665 | 918.2887164605132 | 914.7365208499504 | 948.1895510278536 | 901.3050569925473 | 849.2930907233299 | 1308.7702356381908 | 1138.6280734861184 |
| SE_187_1_0 | SE_187 | SE_187_1 | TTGAATGTGGAAATAACTAG | 55.22761892610324 | 71.95625889364052 | 103.69032057662713 | 61.16149900962082 | 82.72637199289373 | 21.65775274671022 | 46.27490634887984 | 74.68292575393862 | 87.91054250469 18 | 63.42360419993593 | 75.12223774066746 | 5.803405063639747 |
| SE_187_1_5 | SE_187 | SE_187_1 | GTATTCAGTGCTCTGCAAGA | 46.60240419259685 | 48.970231747060915 | 41.47612823065085 | 66.36673296788642 | 34.999618920070425 | 40.71657516381521 | 13.990087965940416 | 26.469897735557318 | 19.535676112153734 | 53.21750697236316 | 11.73784964697929 | 9.285448101823595 |
| SE_187_1_7 | SE_187 | SE_187_1 | GGAGGCTCTGAGGTCAGCAG | 1624.58148569076 | 1949.8147375207313 | 1860.500609203481 | 1648.7578562806293 | 1913.3125009638497 | 1718.759257978923 | 1572.27065524915 | 1549.4343710215871 | 1757.322864816011 | 1299.0903756815228 | 1575.2194226246208 | 2219.2220963358395 |
| SE_187_1_8 | SE_187 | SE_187_1 | TGTCCAATATTCATGAGTAG | 132.22077468597246 | 183.8882171726369 | 196.51784566427426 | 127.52823197750725 | 123.02896347661118 | 300.60960812453785 | 131.2915947572 87 | 119.11453981007931 | 110.11017445032105 | 291.60277930575704 | 49.29896851731302 | 80.08698978822851 |
| SE_187_1_9 | SE_187 | SE_187_1 | TATTTCCATGAAAACTCCAG | 8.670214733506391 | 11.992709815606755 | 3.950107450538176 | 11.71177640609760 4 | 6.363567076376441 | 18.192512307236584 | 1.0761606127646475 | 0.945353490556185 | 4.439926389125849 | 0.7290069448268927 | 1.173784964697929 | 53.39132658548567 |
| SE_187_2_11 | SE_187 | SE_187_2 | CCTCCTTTGTCCCCTCCACG | 259.02266516350346 | 351.78615459113144 | 334.77160643311043 | 226.42767718455 37 | 278.93635684783396 | 209.64704658815492 | 243.21229848481033 | 254.30008895961376 | 292.1471564044809 | 190.99981954464587 | 158.4609702342204 | 220.52939241831038 |

| | | | | | | | | | | | | | | | |
|---|---|---|---|---|---|---|---|---|---|---|---|---|---|---|---|
| SE_187_2_16 | SE_187 | SE_187_2 | GCTTCTGAGTCATATAGCAA | 62.859056817921335 | 102.9374259172913 | 72.08946097232172 | 176.97795458103047 | 113.48361286204653 | 60.641707690788614 | 100.08293698711222 | 67.12009782948914 | 158.06137945288023 | 149.44642368951298 | 92.72901221113639 | 107.94333418369929 |
| SE_187_2_2 | SE_187 | SE_187_2 | TGAAACTCCAAAGTAGTCCA | 162.56652625324483 | 240.85358879676897 | 215.2808560543306 | 192.59365645582727 | 197.27057936766965 | 307.5400890032851 | 138.82471904663953 | 193.79746556401793 | 318.78671473923595 | 304.7249029376411 | 145.5493356225432 | 257.67118482560477 |
| SE_187_2_3 | SE_187 | SE_187_2 | ACATTTTCTTGGTAACCAGG | 76.94815575986922 | 85.94775367851507 | 72.08946097232172 | 114.51514708184324 | 115.60480188750533 | 35.51871450460476 | 60.26499431482026 | 171.10898179066947 | 102.11830694989453 | 196.831875103261 | 140.85419576375148 | 39.46315443275028 |
| SE_187_2_9 | SE_187 | SE_187_2 | CTCAGCCTTTTCCTTAACCA | 175.57184835350444 | 159.90279754142338 | 176.76730841158337 | 145.74655083143685 | 143.1802592184699 | 113.48662439276154 | 137.74855843387488 | 202.30564697902358 | 148.29354139680336 | 40.0953819654791 | 223.01914329260651 | 133.47831646371418 |
| SE_187_3_19 | SE_187 | SE_187_3 | ACCTGAAGAATGACTTTTGG | 1323.2915237014129 | 1439.1251778728106 | 1306.4980392655018 | 1355.9634461281892 | 1368.1669214209346 | 1987.3153920381296 | 1377.4855843387488 | 1816.0240553584315 | 1577.0618534175014 | 1276.491160391889 | 1900.3578578459471 | 1835.036681122888 |
| SE_187_3_4 | SE_187 | SE_187_3 | GAACAGCCTGAGACAAGCTT | 1638.670584632708 | 1522.074754097424 | 1387.4752420015343 | 1544.6531771153172 | 1532.5590708939928 | 1722.2244984183965 | 1795.035902091432 | 1180.746509704675 | 1263.6030503452166 | 1180.2622436747392 | 1450.7982163666402 | 1454.3333089481207 |
| SE_187_3_5 | SE_187 | SE_187_3 | CATTTCGATGAGTGAACTGA | 205.91759992077678 | 227.86148649652833 | 223.18107095540694 | 243.34468754891688 | 194.08879582948143 | 285.8823362565749 | 229.2222105188699 | 241.06514009182717 | 295.69909751578155 | 255.88143763423932 | 264.10161705703405 | 125.35354937461854 |
| SE_187_3_6 | SE_187 | SE_187_3 | GCCTGAGACAAGCTTCGGCA | 5304.003863222535 | 5265.799001535999 | 5267.468285292658 | 5691.923333363436 | 5797.209606578937 | 5105.165477454532 | 5085.935055925724 | 6076.732237295158 | 5136.994832218607 | 5572.529086256767 | 5976.913040241855 | 5790.63757249974 |
| SE_187_3_7 | SE_187 | SE_187_3 | CCATTTCGATGAGTGAACTG | 268.77665673869814 | 234.8572338889656 | 244.90666193336693 | 204.30543286192486 | 340.4508385861396 | 296.2780575749958 | 216.30828316569415 | 332.76442867577714 | 344.5382877961659 | 330.9691529514093 | 305.18409082146155 | 381.86405318749536 |
| SE_188_1_0 | SE_188 | SE_188_1 | TGTAGATGGAGGCTTCTGGA | 82.36703996831072 | 143.91251778728105 | 158.9918248841616 | 127.52823197750725 | 141.0590701930111 | 128.21389626052448 | 85.01668840840715 | 149.36585150787724 | 79.91867500426528 | 184.43875704120384 | 327.4860051507222 | 139.28172152735394 |
| SE_188_1_1 | SE_188 | SE_188_1 | GGTGTGTAGATGGAGGCTTC | 283.9495325223343 | 378.76975167624664 | 402.91095995489394 | 324.0258139020337 | 357.4203507898101 | 449.61494702170415 | 231.3745317443992 | 301.567763487423 | 273.49946557015227 | 295.2478126548915 | 252.36376741005475 | 508.3782835748419 |
| SE_188_1_33 | SE_188 | SE_188_1 | GTGACACGGTGTGTAGATGG | 1731.8753930179016 | 1610.0212927452067 | 1537.579325121985 | 1663.0722496658598 | 1538.9226379703691 | 1190.3100909591935 | 1647.6018981426753 | 1045.5609605551406 | 1289.3546234021464 | 1444.8917646469013 | 1071.665672769209 | 962.2045595514701 |
| SE_188_1_34 | SE_188 | SE_188_1 | CTTGTGACACGGTGTGTAGA | 690.3658481554464 | 618.6239479883817 | 606.34149365761 | 582.9862033257474 | 586.5087655393619 | 601.2192162486757 | 535.9279851567944 | 686.3266341437903 | 634.9094736449964 | 393.663750206522 | 624.4536012192982 | 703.3726937131373 |
| SE_188_1_35 | SE_188 | SE_188_1 | TCTGGGGGTCTCTTGTGACA | 429.1756293085664 | 573.6512861798564 | 530.3019252347501 | 477.58021567086894 | 512.2671496483034 | 471.2726997684143 | 391.72246304633165 | 828.129657727218 | 629.5815619780453 | 378.3546043651573 | 619.7584613605065 | 306.41978736017865 |
| SE_188_2_12 | SE_188 | SE_188_2 | GGCGCAGTCTGGAAAGGCAA | 7510.573512899911 | 7171.640469732839 | 6961.076854710901 | 8163.10815505003 | 7343.556406138412 | 7983.913972547255 | 8587.761689861887 | 7624.275901335632 | 7417.341025673643 | 8713.820011515849 | 7729.373992535862 | 9049.829856239821 |
| SE_188_2_2 | SE_188 | SE_188_2 | GCTGCCAAGATGCTGCCCCA | 968.8964964693392 | 1004.3894470570657 | 969.7513791071223 | 1016.3219303513588 | 1045.7461895511951 | 1215.4330841453775 | 939.4882149435373 | 916.9928858394994 | 1131.2932439492663 | 912.7166949232696 | 1164.3946849803456 | 916.93800005508 |
| SE_188_2_24 | SE_188 | SE_188_2 | CAGGGGCCTGATATGCACAA | 1737.2942772263432 | 1629.0097499532508 | 1661.020182951303 | 1683.893185498922 | 1396.8029732646287 | 1613.9357346848456 | 1636.8402920150288 | 2027.7832372430169 | 1418.112488686796 | 1710.979299508717 | 2475.5124905479324 | 1230.3218734916263 |
| SE_188_2_4 | SE_188 | SE_188_2 | CTTTTCTCATTGCTGCAGAA | 8.670214733506391 | 0.0 | 0.0 | 5.205233958265602 | 1.0605945127294067 | 32.919784174999535 | 0.0 | 0.0 | 0.0 | 0.0 | 0.0 | 0.0 |
| SE_188_2_45 | SE_188 | SE_188_2 | TCCTGAGTCCCGTCCCCGTG | 924.461645960119 | 942.4271130097641 | 950.0008418544313 | 956.4617398313043 | 947.1108998673602 | 1009.2512779966962 | 1097.6838250199405 | 882.9601601794768 | 901.3050569925473 | 1189.7393339574887 | 772.3505067712373 | 1044.6129114551545 |
| SE_189_1_11 | SE_189 | SE_189_1 | GGTCCTCAGCTTTTCTTGGA | 5.4188842084414945 | 8.994532361705065 | 10.862795488979984 | 2.602616979132801 | 1.0605945127294067 | 0.0 | 0.0 | 2.836060471668555 | 10.655823333902038 | 18.225173620672315 | 2.347569929395858 | 1.1606810127279494 |
| SE_189_1_14 | SE_189 | SE_189_1 | GAGTTCAGCAGTAAGCCATC | 856.1837049337562 | 926.4368332556218 | 885.811595783186 | 888.7936983738515 | 859.0815553108195 | 1155.6576865644572 | 630.6301190800834 | 816.7854158405438 | 988.3276142194139 | 686.7245420269329 | 762.9602270536539 | 999.3463519587644 |
| SE_189_1_4 | SE_189 | SE_189_1 | ACTGTCACTGATAACCATAG | 943.9696291105083 | 1006.3882320263334 | 1098.129871249613 | 939.5447294669411 | 981.0499242747012 | 750.224555146042 | 846.9384022457775 | 964.2605603673087 | 1113.533538392763 | 933.8578963232495 | 1047.0161885105526 | 833.3689671386677 |
| SE_189_1_5 | SE_189 | SE_189_1 | TCTGAGAATTTTTGCCTCTA | 176.65562519519273 | 263.8396159433486 | 240.95655448282875 | 242.04337905935049 | 233.33079280046948 | 171.52940175394494 | 164.65257375299106 | 190.01605160179318 | 354.30612585224276 | 223.07612511702916 | 322.79086529193046 | 365.61451900930405 |
| SE_189_1_6 | SE_189 | SE_189_1 | CTCAGAAAGTCTTTTCTTGA | 24.926867358830876 | 36.977521931454156 | 48.38881626909266 | 26.02616979132801 | 24.393673792776355 | 16.459892087499767 | 19.370891029763655 | 9.45353490556185 | 41.73530805778298 | 16.038152786191638 | 26.997054188052367 | 26.695663292742836 |
| SE_189_2_1 | SE_189 | SE_189_2 | CTACTGAATCAGAAAATCTA | 36.84841261740216 | 40.97509186998975 | 53.32645058226538 | 44.24448864525762 | 55.15091466192915 | 39.8502650539468 | 17.21856980423436 | 55.775855942814914 | 7.103882222601358 | 45.92743752409424 | 36.3873339056358 | 40.62383544547823 |
| SE_189_2_13 | SE_189 | SE_189_2 | TGATTGTTGATCTAAGGCAG | 285.0333093640226 | 356.78311701430096 | 323.90881094413044 | 322.7245054124673 | 388.17759165896285 | 250.36362175197013 | 311.0104170889831 | 380.9774566941426 | 380.05769890917264 | 564.9803822408418 | 627.9749561133921 | 337.7581747038333 |
| SE_189_2_16 | SE_189 | SE_189_2 | TTTCAACCTAACCCTGACTA | 36.84841261740216 | 51.968409200962604 | 59.25161175807264 | 29.93009526002721 | 27.575457330964575 | 58.90908747105179 | 44.12258512335055 | 71.84686528227006 | 84.35860139339113 | 13.122125006884067 | 68.07952795247988 | 37.14179240729438 |
| SE_189_2_4 | SE_189 | SE_189_2 | CATGGTCTCCAGGTAAGGAA | 1424.0827699784247 | 1456.1148501115867 | 1583.9930876658086 | 1453.5615828456694 | 1401.0453513155462 | 1592.2779819381353 | 1462.502272747156 | 1762.1389063967288 | 1554.8622214718723 | 2481.5396401907424 | 1626.8659610713296 | 1296.4806912171196 |
| SE_189_2_9 | SE_189 | SE_189_2 | ACTCTGCTAAAATTAGGCTA | 71.52927155142773 | 96.94107100948793 | 99.74021312608895 | 88.48897729051524 | 63.6356707637644 | 91.82887164605133 | 80.71204595734856 | 23.633837263904624 | 142.07764445202716 | 97.68693060680361 | 28.170839152750297 | 145.08512659099367 |
| SE_189_3_10 | SE_189 | SE_189_3 | TGACTCTCCTCAATTGGTGG | 80.19948628493412 | 99.93924846338962 | 104.67784743926167 | 72.87327541571842 | 100.75647870929365 | 115.21924461249836 | 82.86436718287786 | 127.62272122508497 | 24.863587779104755 | 94.77090282749604 | 213.62886357502308 | 30.177706330926686 |
| SE_189_3_12 | SE_189 | SE_189_3 | GCATTTAGCAATGATAACCA | 744.5546902398613 | 789.520062860778 | 784.096328931828 | 793.7981786355043 | 800.7488571107021 | 739.8288338276211 | 740.3985015820774 | 708.0697644265825 | 848.0259403230372 | 1262.6400284401782 | 642.0603756897672 | 697.5692886494976 |
| SE_189_3_28 | SE_189 | SE_189_3 | ACTGGAGCTTGATCCTACTG | 2012.5735950151711 | 1857.870628934413 | 1842.7251256760592 | 1911.6221711730423 | 1739.375000876227 | 1759.475833142738 | 1971.526242584834 | 1920.0129393196116 | 1735.1232328703818 | 1380.7391535021347 | 1620.99703624784 | 2234.3109495013027 |
| SE_189_3_8 | SE_189 | SE_189_3 | CATCCCATCCAAATTCTGAC | 1204.0760711157002 | 1190.2764491989703 | 1209.7204067273165 | 1162.0684811827955 | 1222.865473177006 | 979.7967342611703 | 1025.581063964709 | 1392.5056915892606 | 1331.9779167377546 | 967.3922157852866 | 1349.8527094026183 | 1122.3785393079272 |
| SE_189_3_9 | SE_189 | SE_189_3 | AGGCCAGTCAGAATTTGGAT | 3929.7748279617717 | 4266.406516902103 | 4072.5607815048597 | 4488.212980514515 | 4458.739331514426 | 4497.8820904367785 | 4338.003430054294 | 4606.707559480289 | 4085.6202632736063 | 4292.392891140744 | 4731.527192697352 | 3953.279529351396 |
| SE_19_1_1 | SE_19 | SE_19_1 | TGTCACACAGATAATAACAC | 2413.5710264398417 | 2371.558366036236 | 2334.5135032680623 | 2313.72649444906 | 2369.3681414374946 | 2105.9998770901016 | 2340.6493327631083 | 2343.531303088783 | 2807.8094484831868 | 2668.1654180664273 | 2251.319562290628 | 2974.8254356217344 |
| SE_19_1_13 | SE_19 | SE_19_1 | GTAGTACCTATCTTAAAGGG | 86.70214733506391 | 102.9374259172913 | 95.79010567555078 | 115.81645557140963 | 82.72637199289373 | 84.89839076710406 | 107.61606127646475 | 138.96696311175918 | 114.5501008394469 | 42.282402799959776 | 205.41236882213758 | 125.35354937461854 |
| SE_19_1_2 | SE_19 | SE_19_1 | TCAAATTGTGACATAACACA | 382.57322511596954 | 379.76914416088056 | 368.3475197626849 | 370.87291952642414 | 417.87423801538625 | 324.8662912006533 | 349.75219914851044 | 536.9607826359131 | 392.48949279872505 | 279.93866681352677 | 791.1310662064042 | 316.86591647473017 |
| SE_19_1_6 | SE_19 | SE_19_1 | GCATAGTAGGAGTTAACAGA | 904.9536628097296 | 1081.3426683738758 | 1078.379333996922 | 1024.1297812887572 | 885.5964181290547 | 836.8555661328828 | 735.0176985182542 | 875.3973322550273 | 827.6022789330582 | 1004.5715699714581 | 1048.1899734752506 | 1315.0515874207667 |
| SE_19_1_8 | SE_19 | SE_19_1 | CTTGTAAGATAAATTCAGAG | 100.7912462770118 | 144.91191027191493 | 150.1040831204507 | 106.70729614444484 | 135.75609762936406 | 48.51336615263089 | 53.808030638232374 | 132.3494886778659 | 94.126439449468 | 67.06863892407412 | 77.46980767006332 | 40.62383544547823 |
| SE_19_2_10 | SE_19 | SE_19_2 | TGAAATGTTCTACAGAAAGG | 118.13167574402458 | 192.88274953434197 | 197.50537252690881 | 166.56748666449926 | 149.54382629484635 | 190.58822417104992 | 72.10276105523138 | 59.557269905039654 | 64.82292528123739 | 106.43501394472632 | 53.994108376104734 | 304.0984253347228 |
| SE_19_2_11 | SE_19 | SE_19_2 | GTTCTACAGAAAGGTGGAAG | 11.921545258571289 | 22.986027146579612 | 33.5759133295745 | 15.615701874796805 | 21.211890254588134 | 17.326202197368175 | 11.837766740411123 | 8.508181415005666 | 31.96747000170611 | 31.347298627556384 | 0.0 | 9.285448101823595 |
| SE_19_2_3 | SE_19 | SE_19_2 | TCAGACTGATAACGTCTCTT | 981.9018185695988 | 989.3985597875572 | 1077.3918071342875 | 972.0774417061011 | 1223.9260676897354 | 1124.4705226091946 | 951.3259816839484 | 1297.970342533642 | 1201.4440808974548 | 1331.8956881987328 | 1541.1796586483808 | 793.9058127059175 |
| SE_19_2_7 | SE_19 | SE_19_2 | TGTAGAACATTTCATTTGGA | 213.50403781259487 | 341.7922297447925 | 333.7840795704759 | 231.63291114281927 | 355.29916176435125 | 265.9572037296015 | 199.0897133614598 | 183.3985771678999 | 210.45251084456524 | 228.9081806756443 | 272.3181118099195 | 207.76190127830296 |
| SE_19_2_8 | SE_19 | SE_19_2 | TTTCTGTAGAACATTTCATT | 0.0 | 3.9975699385355847 | 12.837849214249072 | 1.3013084895664004 | 12.727134152752882 | 0.0 | 0.0 | 0.0 | 0.0 | 2.187020834480678 | 1.173784964697929 | 0.0 |
| SE_19_3_13 | SE_19 | SE_19_3 | AACTGGGTGCCGGTCCCTGG | 82.36703996831072 | 182.888824688003 | 181.7049427247561 | 118.41907255054244 | 164.39214947305805 | 190.58822417104992 | 87.16900963393644 | 277.93392622351837 | 128.75786528464963 | 129.7632361791869 | 246.49484258656508 | 140.44240254008187 |
| SE_19_3_15 | SE_19 | SE_19_3 | GGAAGGGTATGGGTTGTCAT | 2579.3888832181515 | 2290.60757478089 | 2294.0249019000457 | 2601.3156706432346 | 2241.0362053972362 | 2332.9731258756246 | 2592.470916150036 | 2218.7446423353663 | 2197.763562617295 | 1714.6243342328514 | 2633.973460782153 | 2428.1446786268702 |
| SE_19_3_21 | SE_19 | SE_19_3 | ACTGTTGTAGAAGCAGAATT | 621.0041302873952 | 678.5874970664155 | 711.0193410968717 | 631.1346174397042 | 775.2945888051963 | 714.7058406414372 | 662.9149374630229 | 698.6162295210207 | 756.5634567070447 | 621.1139169925125 | 734.7893879009035 | 746.3178911840715 |
| SE_19_3_22 | SE_19 | SE_19_3 | CCACGTTTGAGCAAGTTATT | 1179.1492037568692 | 1176.2849544140959 | 1195.8950306504328 | 1270.0770858168069 | 1197.4112048715 | 1401.6897577670852 | 1227.8992591644628 | 981.27692319732 | 1360.3934456281602 | 1374.1780909986926 | 1032.9307689341776 | 1382.3710861589877 |
| SE_19_3_8 | SE_19 | SE_19_3 | GGATGAACCCCCTTCAGGAC | 21.675536833765978 | 34.97873696218637 | 42.463655093285396 | 26.02616979132801 | 71.05983235287025 | 21.65775274671022 | 22.599372868057596 | 62.393330376708214 | 15.095749723027886 | 8.748083337922711 | 15.259204541073078 | 95.17584304369186 |
| SE_19_4_0 | SE_19 | SE_19_4 | GCTGTGAGTACATTTATCAG | 720.7115997227188 | 830.4951547307677 | 770.2709528549443 | 709.2131268136883 | 982.1105187874307 | 888.8341727249874 | 511.17629106320754 | 874.4519787644712 | 776.0991328191984 | 621.8429239373394 | 603.3254718547355 | 667.391582318571 |
| SE_19_4_1 | SE_19 | SE_19_4 | GTGTAAAGTCTGTGGTCACA | 728.2980376145368 | 804.5109501302865 | 777.1836408933862 | 749.5536899902467 | 756.203887576067 | 426.2245740552571 | 825.4151899904846 | 657.9660294271048 | 718.3800897605623 | 756.7092087303146 | 525.8556641846722 | 493.2894304093785 |
| SE_19_4_10 | SE_19 | SE_19_4 | TATTTTTAGTGTCAGCCAAG | 22.759313675454276 | 25.984204600481302 | 19.75053725269088 | 16.917010364363207 | 25.454268305505764 | 12.994651648026132 | 43.0464245105859 | 0.0 | 31.96747000170611 | 27.702263903421922 | 0.0 | 11.606810127279495 |
| SE_19_4_2 | SE_19 | SE_19_4 | TCATCAACCCAGCTGAAAAT | 1835.9179698199785 | 2062.7460882843616 | 2116.270066625828 | 2315.0278029386263 | 2302.550687135542 | 2255.005215987468 | 2166.3113134952355 | 2346.3673635604514 | 2271.4663406767845 | 2699.5127166939833 | 2785.391721228186 | 1881.463921632006 |
| SE_19_4_7 | SE_19 | SE_19_4 | GGTCATGTATTTAGATTCTG | 250.35245042999705 | 227.86148649652833 | 293.2954782024596 | 222.52375171585447 | 252.42149402959882 | 157.66843999605038 | 259.35470767628004 | 283.6060471668555 | 158.94936473070538 | 126.84720839987932 | 250.0161974806589 | 155.53125570554522 |
| SE_19_5_0 | SE_19 | SE_19_5 | AAATGATTACAACACAGGAG | 10.837768416882989 | 19.987849692677923 | 20.738064115325425 | 9.109159426964803 | 19.090701229129323 | 26.85561340592067 | 12.913927353175769 | 10.398888396118036 | 3.551941111300679 | 96.22891671714983 | 2.347569929395858 | 52.23064557275772 |
| SE_19_5_3 | SE_19 | SE_19_5 | CCTACTCTGTACTCTCTCTG | 589.5746018784346 | 702.572916697629 | 699.1690187452572 | 648.0516278040674 | 617.2660064085147 | 801.336851628278 | 527.3187002546773 | 485.9116941458791 | 498.15974085992025 | 701.3046809234708 | 516.4653844670888 | 863.5466734695943 |
| SE_19_5_5 | SE_19 | SE_19_5 | ATGTATATCTTCCTACAGAA | 19.50798315038938 | 23.98541963121351 | 19.75053725269088 | 16.917010364363207 | 33.939024407341016 | 2.598930329605226 | 4.30464245105859 | 4.726767452780925 | 35.51941111300679 | 4.374041668961356 | 1.173784964697929 | 0.0 |
| SE_19_5_7 | SE_19 | SE_19_5 | AAGTAAAGCAACTTACAGAT | 216.75536833765977 | 273.83354078968756 | 265.64472604869235 | 193.89496494539367 | 247.11852146595177 | 443.55077625262527 | 225.99372868057597 | 224.99413075237203 | 204.23661389978906 | 225.99215289633673 | 217.15021846911688 | 161.33466076918498 |
| SE_19_5_9 | SE_19 | SE_19_5 | TGGAGCATAGAAGTCTAGGT | 302.3737388310354 | 290.8232130284638 | 293.2954782024596 | 333.1349733289985 | 326.6631099206573 | 382.90906856183665 | 331.4574687315114 | 482.13028018365435 | 380.9456841869978 | 211.41201399979886 | 170.1988198811997 | 343.561579767473 |
| SE_190_1_1 | SE_190 | SE_190_1 | TCCTCAACTCCTGTACCAGC | 205.91759992077678 | 266.83779339725027 | 256.75698428498146 | 208.20935833062407 | 214.24009157134017 | 163.73261076512924 | 230.29837113163455 | 335.60048914744567 | 264.6196127919006 | 215.7860556687602 | 215.97643350441894 | 149.7278506419055 |
| SE_190_1_18 | SE_190 | SE_190_1 | GAACCAGTGCAGAAAAGAAG | 296.9548546225939 | 337.7946598062569 | 292.307951339825 | 303.2048780689713 | 307.572408691528 | 581.2940837217022 | 287.33488360816085 | 226.8848377334844 | 262.8436422362503 | 86.75182643440023 | 428.4315121147441 | 349.3649848311128 |
| SE_190_1_26 | SE_190 | SE_190_1 | ATAGAACAGGATACATACTG | 579.82061030324 | 675.5893196125138 | 634.9797726740118 | 605.1084476483762 | 562.1150917465856 | 710.3742900920952 | 653.229491948141 | 509.54553140978373 | 609.1579005880665 | 419.90800022029015 | 415.5198775030669 | 663.9095392803871 |
| SE_190_1_32 | SE_190 | SE_190_1 | TGGAACCTGACCCAACACGG | 86.70214733506391 | 121.92588312533533 | 96.77763253818532 | 78.07850937398402 | 156.9679878839522 | 149.0053388973663 | 55.960351863761666 | 52.93979547114636 | 135.86174750725098 | 51.759493082709376 | 23.47569929395858 | 31.338387343654635 |
| SE_190_1_4 | SE_190 | SE_190_1 | TCACACATGTGCTTACAGGG | 1150.9710058729734 | 1079.3438834046078 | 1102.079978700151 | 1110.0161416001395 | 968.3227901219484 | 576.0962230624918 | 949.173660458419 | 1031.3806581967979 | 1305.3383584029996 | 569.3544239098031 | 975.415305663979 | 670.8736253567548 |
| SE_190_2_1 | SE_190 | SE_190_2 | GTTTCTGAGTGACTTTTGGA | 0.0 | 13.991494784874547 | 13.825376076883616 | 9.109159426964803 | 3.1817835381882205 | 0.0 | 0.0 | 8.508181415005666 | 0.0 | 8.019076393095819 | 19.954344399864794 | 32.49906835638259 |
| SE_190_2_16 | SE_190 | SE_190_2 | TGGATAACTTCTCCTCAGCA | 100.7912462770118 | 136.91677039484378 | 142.20386821937433 | 119.72038104010883 | 113.48361286204653 | 50.2459863723677 | 123.75847046793446 | 123.84130726286024 | 166.05324695330674 | 129.7632361791869 | 105.64064682281361 | 87.05107595459621 |
| SE_190_2_21 | SE_190 | SE_190_2 | CCTTTCTCCCAAGGTCTCCA | 134.38832836934907 | 134.91798542557598 | 139.2412876314707 | 114.51514708184324 | 195.14939034221084 | 168.93047142433971 | 148.51016456152135 | 79.40969320671954 | 171.38115862025776 | 110.0800486688608 | 83.33873249355297 | 39.46315443275028 |
| SE_190_2_5 | SE_190 | SE_190_2 | TAGTGTCTACTGCTGAAGAG | 701.2036165723293 | 853.4811818773474 | 920.375035975395 | 607.711064627509 | 744.5373479360435 | 814.3315032763043 | 467.053705939857 | 559.6492664092615 | 718.3800897605623 | 466.5644446892113 | 370.91604884454557 | 699.8906506749535 |
| SE_190_2_8 | SE_190 | SE_190_2 | TGGCTTTTCCTGCCTCCCAA | 7.586437891818092 | 4.996962423169481 | 5.925161175807264 | 13.013084895664004 | 4.242378050917627 | 0.8663101098684087 | 6.456963676587884 | 0.0 | 5.327911666951019 | 16.76715973101853 | 0.0 | 0.0 |
| SE_190_3_17 | SE_190 | SE_190_3 | GGTAACATAGAATCTTAGAA | 14.089098941947885 | 42.973876839257535 | 46.41376254382357 | 41.641871666124814 | 63.6356707637644 | 39.8502650539468 | 26.904015319116187 | 11.34424188667422 | 36.40739639083196 | 21.87020834480678 | 11.73784964697929 | 32.49906835638259 |
| SE_190_3_2 | SE_190 | SE_190_3 | TATATGATACATGATGAAGA | 52.02128840103835 | 46.97144677779312 | 42.463655093285396 | 27.32747828089441 | 50.90853661101153 | 133.41175691973496 | 49.50338818717378 | 51.994441980590175 | 106.55823333902038 | 68.5266528137279 | 48.12518355261509 | 32.49906835638259 |
| SE_190_3_6 | SE_190 | SE_190_3 | CACTCCCTTTCCTCTCACAA | 302.3737388310354 | 307.81288526724 | 297.24558565299776 | 248.5499215071825 | 302.2694361278809 | 336.1283226289426 | 263.65935012733866 | 446.2068474525193 | 273.49946557015227 | 226.72115984116363 | 433.1266519735358 | 242.58233166014142 |
| SE_190_3_7 | SE_190 | SE_190_3 | CTGAGCCCTACATAGCAAGG | 829.0892838915487 | 765.5346432295645 | 723.8571903111208 | 821.1256569163987 | 845.2938266453372 | 793.5400606394624 | 821.110547539426 | 708.0697644265825 | 720.1560603162127 | 823.0488407095618 | 583.3711274548707 | 802.0305797950131 |
| SE_190_3_8 | SE_190 | SE_190_3 | GCAGCCCTTGTGAGAGGAAA | 755.3924586567443 | 850.4830044234457 | 812.7346079482297 | 804.2086465520355 | 925.899009612772 | 1017.9143790953802 | 696.2759164587269 | 1000.1839930084437 | 988.3276142194139 | 934.5869032680764 | 1051.7113283693443 | 1002.8283949969483 |
| SE_191_1_10 | SE_191 | SE_191_1 | ACAGGCTTGCCGGCAGTGCG | 3605.7255522969704 | 3572.8281325661787 | 3507.6954160779005 | 3595.5153566719646 | 3441.629193806925 | 3801.3687621025774 | 3300.584599349174 | 3386.2562031722546 | 3550.1651407450286 | 2838.75304315592 | 2875.7731635099262 | 3262.674326778266 |
| SE_191_1_14 | SE_191 | SE_191_1 | TTTCCGCTCCTGGATCCCAC | 67.19416418467453 | 112.93135076363026 | 93.81505195028168 | 62.46280749918722 | 102.87766773475245 | 110.88769406315632 | 40.8941032850566 | 35.92343264113503 | 47.063219724734 | 37.908361130998415 | 16.432989505771005 | 64.99813671276517 |
| SE_191_1_30 | SE_191 | SE_191_1 | CCAGAGACCACACCTGGCGG | 372.8192335407748 | 431.73755336184314 | 415.74880916914304 | 460.66320530650574 | 467.7221801136684 | 425.35826394538867 | 430.464245105859 | 649.4578480120991 | 281.49133307057883 | 567.1674030753225 | 720.7039683245284 | 358.6504329329364 |
| SE_191_1_42 | SE_191 | SE_191_1 | TTTCCTTTCCGGGCAGGGGG | 5.4188842084414945 | 1.9987849692677924 | 9.87526862634544 | 6.506542447832002 | 3.1817835381882205 | 0.0 | 0.0 | 0.0 | 1.7759705556503396 | 4.374041668961356 | 0.0 | 1.1606810127279494 |
| SE_191_1_45 | SE_191 | SE_191_1 | GCTGGCTTTGTGACTCAGCC | 861.6025891421976 | 873.4690315700252 | 868.0361122557642 | 853.6583691555587 | 833.6272870053137 | 1035.2405812927484 | 903.9749147223039 | 1043.6702535740283 | 791.1948825422263 | 758.1672226199684 | 993.022080134448 | 951.7584304369185 |
| SE_191_2_1 | SE_191 | SE_191_2 | ATCCCAGTCCTGTCATGTAG | 130.05322100259588 | 137.91616287947767 | 127.39096527985618 | 132.73346593577284 | 148.48323178211695 | 86.63101098684088 | 118.37766740411122 | 89.80858160283758 | 113.66211556162173 | 128.3052222895331 | 143.20176569314734 | 44.10587848366208 |
| SE_191_2_12 | SE_191 | SE_191_2 | GTTTGCTAGACTGGGCCAGA | 110.54523785220648 | 152.90705014898612 | 118.50322351614528 | 163.96486968536647 | 144.2408537311993 | 195.78608483026036 | 124.83463108069911 | 119.11453981007931 | 95.01442472729316 | 154.54947230330123 | 95.07658214053225 | 253.02846077469297 |

| | | | | | | | | | | | | | | |
|---|---|---|---|---|---|---|---|---|---|---|---|---|---|---|
| SE_191_2_17 | SE_191 | SE_191_2 | ACGCTTCGTTCACCAAGCGG | 1316.7888626512831 | 1405.145833395258 | 1345.011586908249 | 1385.8935413882164 | 1370.2881104463936 | 1000.5881768980121 | 1348.4292477941033 | 1455.8443754565249 | 1186.348331174427 | 1976.3378274257059 | 1282.9469664148364 | 1311.5695443825828 |
| SE_191_2_3 | SE_191 | SE_191_2 | AATGCATTTTGGGATATGCA | 173.40429467012183 | 201.87728189604704 | 178.74236213685248 | 157.45832732753446 | 168.63452752397566 | 217.4438375769706 | 172.1856980403436 | 202.4606433441387 | 88.2098403240501 | 69.25331291717781 | 299.455701283811 |
| SE_191_2_5 | SE_191 | SE_191_2 | TTTGGGATATGCACGGAACA | 107.2939073271416 | 184.8876096572708 | 138.25376076883617 | 111.91253010271043 | 168.63452752397566 | 111.75400417302473 | 74.25508228076067 | 71.84686528227006 | 140.30167389637683 | 392.93474326169513 | 17.606774470086935 | 117.2287822855229 |
| SE_191_3_14 | SE_191 | SE_191_3 | GCTGCACTGCCTCCAAGTGA | 3432.3212576268425 | 3238.0316502138235 | 3374.379289622237 | 3268.8869257907977 | 3390.7206571959513 | 3496.427603428898 | 3473.846458004282 | 3146.136416570984 | 3373.45607045782 | 4030.6793979478894 | 3331.2017298127225 | 3923.101823020469 |
| SE_191_3_20 | SE_191 | SE_191_3 | GCAGCCTCTTCAGAGAGCCG | 586.3232713533697 | 592.6397433879005 | 688.3062232562772 | 720.9249032197858 | 653.3262198413146 | 747.6256248164367 | 637.0870827566713 | 749.6653180110548 | 831.1542200443589 | 597.7856947580519 | 739.4845277596953 | 705.6940557385933 |
| SE_191_3_21 | SE_191 | SE_191_3 | TCTGGGGAGACATTTATCAG | 945.0534059521966 | 1032.3724366268148 | 1062.5789041947694 | 959.0643568104372 | 1101.9576987258536 | 1073.3582261269585 | 858.7761689861887 | 1081.4843931962757 | 1352.4015781277335 | 1099.342472798954 | 1115.0957164630327 | 1205.9475722243394 |
| SE_191_3_26 | SE_191 | SE_191_3 | GTGAATGCTGAGGCTCAGCG | 675.1929723718102 | 607.6306306574089 | 573.7531071906701 | 730.0340626467506 | 618.3266009212441 | 612.4812476769626 | 761.9217138373704 | 655.1299689554362 | 431.5608450230325 | 865.3312435095215 | 870.9484438058633 | 385.3460962256792 |
| SE_191_3_9 | SE_191 | SE_191_3 | GAGATAGCTGTGGAGCTGAG | 2210.90475704413 | 2254.6294453340697 | 2413.5156522788257 | 2598.7130536641016 | 2611.1836903397993 | 2024.5667267624713 | 2551.5768128649793 | 2498.569275539997 | 2468.599072353972 | 2170.9826816944865 | 2537.7230936769224 | 2517.517116606922 |
| SE_191_4_11 | SE_191 | SE_191_4 | CGTCGCTGCCCAGGGATGAA | 10822.595541099354 | 9649.134439140267 | 9280.777455039444 | 10124.180048826596 | 10140.344136205858 | 9494.75880415776 | 11093.063596377986 | 10294.899512156855 | 8695.151840464063 | 8863.995442150188 | 9882.095617791865 | 8318.600818221214 |
| SE_191_4_13 | SE_191 | SE_191_4 | CTTCATCCCTGGGCAGCGAC | 436.7620672003845 | 427.7399834233076 | 383.1604227022031 | 356.55852614119374 | 361.6627288407277 | 280.68447559736444 | 329.3051475059821 | 376.2506892413616 | 401.36934557697674 | 313.47298627556387 | 326.31222018602426 | 188.03032406192781 |
| SE_191_4_14 | SE_191 | SE_191_4 | CAACGGCCGCTGCTGCCCA | 2047.2544539491967 | 1833.8852093031994 | 1746.9350200005083 | 1980.5915211200615 | 1806.1924551781797 | 1999.4437335762873 | 2051.162127929418 | 2119.4825258269666 | 1964.2234345492755 | 1406.9834035159029 | 2140.9837756090224 | 1605.2218406027541 |
| SE_191_4_2 | SE_191 | SE_191_4 | GCACCTTCGTGGCAGAGCTT | 644.8472208045379 | 643.6087601042292 | 676.4559009046627 | 664.9686381684306 | 678.7804881468203 | 574.363602842755 | 733.9415379054896 | 719.4140063132568 | 658.885076146276 | 607.2627850408015 | 710.1399036422471 | 641.856600038556 |
| SE_191_4_5 | SE_191 | SE_191_4 | TTCGTGGCAGAGCTTGGGGT | 347.89236618194394 | 418.7454510616025 | 417.72386289441215 | 426.82918457777936 | 470.9039636518566 | 400.2352707592048 | 304.55345341239524 | 331.8190751852209 | 323.2266411283618 | 177.14868759293492 | 380.306328562129 | 258.8318658383327 |
| SE_191_5_17 | SE_191 | SE_191_5 | ACTTAGCTGAACCTTGGCCA | 1987.64672765634 | 2020.7716039297381 | 2002.7044774228552 | 2283.796399189033 | 2402.246571332106 | 2352.898258402598 | 2116.8079253080614 | 2634.7001781800877 | 2162.2441515042883 | 2177.5437441979284 | 2967.3283907563646 | 2510.5530305305547 |
| SE_191_5_21 | SE_191 | SE_191_5 | AGGAGTTGCGGAGGGGACAG | 2971.7160999093157 | 3162.0778213816475 | 3299.3272480620117 | 2978.695132617491 | 2862.5445898566686 | 3557.9356212295547 | 3466.3133337149293 | 2722.6180528018126 | 3416.9673490712535 | 3388.424279555397 | 2767.7849467577166 | 3348.564721720134 |
| SE_191_5_28 | SE_191 | SE_191_5 | TATGCAGTGGGGGTTGGGGA | 20296.97269113846 | 23407.770775095116 | 24068.00469612911 | 21530.148959876096 | 23267.322420257726 | 23846.918394347686 | 19703.42465910793 | 21386.732016852573 | 24236.670172960185 | 21144.117427759196 | 21423.9231756666 | 20338.61338603186 |
| SE_191_5_48 | SE_191 | SE_191_5 | TTCCACCACTGGGAGACAAG | 3.2513305250648967 | 15.990279754142339 | 19.75053725269088 | 9.109159426964803 | 6.363567076376441 | 0.0 | 4.30464245105859 | 3.78141396224474 | 16.871720278678225 | 4.374041668961356 | 3.521354894093787 | 6.964086076367696 |
| SE_191_5_50 | SE_191 | SE_191_5 | TGGGACCTCTTGTCTCCCAG | 56.35639576779154 | 66.95929647047105 | 95.79010567555078 | 57.25757354092162 | 90.15053358199957 | 43.31550549342044 | 58.112673089290965 | 84.13646065950047 | 181.14899667633463 | 29.8892847379026 | 55.167893340802664 | 60.35541266185337 |
| SE_191_6_21 | SE_191 | SE_191_6 | CTCCTACCGTGCCGCGAGCT | 111.62901469389479 | 89.94532361705066 | 125.4159115545871 | 101.50206218617923 | 149.54382629484635 | 71.90373911907793 | 124.83463108069911 | 62.393330376708214 | 106.55823333902038 | 61.965590310285876 | 62.210603128990236 | 129.99627342553035 |
| SE_191_6_22 | SE_191 | SE_191_6 | CACCGAGCTCGCGGCACGGT | 1014.4151238202478 | 934.431973132693 | 943.0881538159896 | 1003.3088454556947 | 947.1108998673602 | 1366.1710432624805 | 991.1439243562403 | 1236.52236564749 | 833.8181758778344 | 583.2055558615141 | 759.4388721595601 | 1067.8265317097134 |
| SE_191_6_24 | SE_191 | SE_191_6 | GTGCCGCGAGCTCGGTGCTC | 902.786109126353 | 879.4653864778286 | 928.2752508764714 | 892.6976238425507 | 779.5369686856114 | 1004.0534173374857 | 1055.7135611221192 | 922.6650067828366 | 696.1804578149331 | 818.6747990406004 | 996.5434350285417 | 841.4937342277633 |
| SE_191_6_25 | SE_191 | SE_191_6 | TGCCGCGAGCTCGGTGCTCC | 2987.97275253464 | 2345.5741614357544 | 2275.2618915099893 | 2600.014362153668 | 2233.6120438081307 | 2547.81803312299 | 3096.1140829238907 | 2651.716541010099 | 2419.7598820735875 | 2872.287362617957 | 1961.3946760102394 | 2652.1561140833646 |
| SE_191_6_28 | SE_191 | SE_191_6 | CAGTGGAGCCCCCGCGCGCC | 3811.6431522177472 | 4406.3214647508485 | 4430.045505778565 | 4204.52772978904 | 4505.40549007452 | 4688.470314607828 | 3637.4228711445085 | 5003.756025513887 | 4242.793657448661 | 5167.930231877842 | 5395.8894827163795 | 4886.467063584667 |
| SE_192_1_13 | SE_192 | SE_192_1 | GGACAGAGAGTAGATGTGGA | 1049.0959827542733 | 1304.2071924472345 | 1311.4356735786744 | 1093.0991312357764 | 1213.3201225624414 | 887.967862615119 | 1163.329622398584 | 1624.1172967755258 | 1239.627447843937 | 1265.5560562194858 | 1210.1722986035647 | 990.0609038569409 |
| SE_192_1_15 | SE_192 | SE_192_1 | GGAGGGAAGGAAAGGCTCAG | 4028.3985205554072 | 4247.418059694059 | 3968.8704609282327 | 4019.741924270611 | 4163.8940569756505 | 3677.486416391395 | 4312.175575347943 | 4044.2222325993594 | 3541.285287966777 | 3661.072876920655 | 4456.861510958037 | 3607.3965875584668 |
| SE_192_1_20 | SE_192 | SE_192_1 | GAGCTGAGAACTCAATCCCT | 841.01082915012 | 963.4143551870759 | 1030.978044590464 | 977.2826756643667 | 820.9001528525608 | 671.3903351480168 | 985.763121292417 | 916.9928858394994 | 1014.0791872763439 | 646.6291600614538 | 971.8939507698852 | 705.6940557385933 |
| SE_192_1_22 | SE_192 | SE_192_1 | TGGTAGCAGTATGATCTCCA | 31.429528408960667 | 6.9957473924372735 | 12.837849214249072 | 18.218318853929606 | 5.3029725636470335 | 17.326202197368175 | 41.970263897821255 | 1.89070698111237 | 19.535676112153734 | 18.954180565499207 | 3.521354894093787 | 110.26469620915519 |
| SE_192_1_33 | SE_192 | SE_192_1 | CTATTGTGTGCTTTGCCCAG | 143.05854310285545 | 142.91312530264716 | 149.11655625781614 | 140.54131687317124 | 112.42301873431712 | 58.90908747105179 | 146.35784333599204 | 258.01059292183853 | 116.32607139926724 | 38.66373687582531 | 113.85714157569912 | 165.97738482009677 |
| SE_192_2_10 | SE_192 | SE_192_2 | TTACTAAATTGCTTTCCAGA | 17.340429467012783 | 31.980559508284678 | 31.60085960430541 | 24.724861301761607 | 26.51486281823517 | 50.2459863723677 | 22.599372868057596 | 2.836060471668555 | 18.647690834328564 | 12.393118062057175 | 2.347569929395858 | 135.79967848917008 |
| SE_192_2_11 | SE_192 | SE_192_2 | AGGTGGGCTGTGAGGATAAA | 7297.069475087316 | 8536.81060374274 | 8523.34435139875 | 7788.331310054907 | 8357.484760307725 | 7687.635914972259 | 7609.531692858823 | 7567.554691902261 | 8395.012816559156 | 7998.664198640667 | 7419.49476185561 | 8047.001461242873 |
| SE_192_2_12 | SE_192 | SE_192_2 | GTATTTATCACACAGCAGTT | 30.34575156727237 | 56.96537162413208 | 69.12688038441809 | 44.24448864525762 | 64.69626527649382 | 42.44919538355203 | 31.208657770174778 | 35.92343264113503 | 22.199631945629246 | 40.82438891030599 | 26.997054188052367 | 40.62383544547823 |
| SE_192_2_21 | SE_192 | SE_192_2 | AAACAAACAGAAAGTCTGCA | 1.083776841688299 | 6.9957473924372735 | 5.925161175807264 | 0.0 | 7.4241615891058474 | 51.11229648223612 | 4.30464245105859 | 0.945353490556185 | 10.655823333902038 | 0.7290069448268927 | 1.17378496497929 | 0.0 |
| SE_192_2_9 | SE_192 | SE_192_2 | TTCTGTTCTGACCCTACAAA | 0.0 | 0.9993924846338962 | 0.0 | 0.0 | 0.0 | 0.0 | 0.0 | 12.289595377230405 | 0.0 | 0.0 | 1.17378496497929 | 0.0 |
| SE_192_3_1 | SE_192 | SE_192_3 | ACCACTCTTACCCACCACAA | 1182.4005342819341 | 1258.2351381540752 | 1224.5333096668346 | 1159.4658642036627 | 1185.7446652314768 | 1017.0480689855118 | 1161.1773011730547 | 1258.2654959302822 | 1264.4910356230419 | 797.5335976406205 | 991.84829516975 | 865.8680354950503 |
| SE_192_3_12 | SE_192 | SE_192_3 | GGCTATGTAGGCAGTTGCAA | 335.9708209233727 | 457.72175796232443 | 409.8236479933358 | 391.6938553594865 | 452.8738569354567 | 326.5989114203901 | 281.9540805443376 | 457.55108942919355 | 369.40187557527065 | 433.7591321720011 | 411.9985226089731 | 453.8262759766282 |
| SE_192_3_13 | SE_192 | SE_192_3 | GAGAAGACTGAGACTCTGAG | 431.3431829194297 | 595.6379208418022 | 617.20428914659 | 595.9992882214113 | 534.539634415621 | 499.8609333940718 | 509.0239698376824 | 423.5183637691709 | 662.4370172575767 | 328.0531251721017 | 299.3151659979719 | 393.4708633147749 |
| SE_192_3_17 | SE_192 | SE_192_3 | TGTTTTTTGCCTCTGCCCAC | 105.126353643765 | 90.94471610168455 | 88.87741763710896 | 81.98243484268323 | 129.39253055298764 | 163.73261076512924 | 101.15909759987686 | 207.0324144318045 | 169.60518806460743 | 33.53431946203706 | 105.64064682281361 | 133.47831646371418 |
| SE_192_3_19 | SE_192 | SE_192_3 | TGTGAATAATGTCAGTTCAG | 1287.5268879256992 | 1387.1567686718479 | 1522.7664221824668 | 1514.72308185529 | 1318.3189793226527 | 1458.8662250184002 | 1454.9691484578034 | 1339.565896118114 | 1670.3003075891443 | 1132.8767922609911 | 1294.6848160618158 | 1333.622483624414 |
| SE_192_4_0 | SE_192 | SE_192_4 | GCAGCTAGAGAGCCAAGCAC | 3954.7016953206025 | 3102.1142723036137 | 3082.071338282412 | 3795.91686406519 | 3513.7496206725245 | 3060.673618165088 | 3996.860515807901 | 3440.1413521339573 | 2695.03531819939 | 3422.687605962261 | 3669.2517996457264 | 2440.9121697668775 |
| SE_192_4_12 | SE_192 | SE_192_4 | AATAGAACTCTGATTCCTTA | 20.59175999207768 | 33.97934477755247 | 21.725590977959968 | 23.423552812195208 | 9.545350614564661 | 52.844916701972934 | 35.513300221233365 | 32.14201867891029 | 22.199631945629246 | 13.122125006884067 | 12.911634611677218 | 1.1606810127279494 |
| SE_192_4_17 | SE_192 | SE_192_4 | TCTGAAGTTGAATAACATGG | 463.8564882425919 | 473.7120377164668 | 585.6034295422846 | 469.77236473347057 | 510.14596062284465 | 496.3956929545982 | 448.758975522858 | 547.3596710320311 | 590.5102097537379 | 688.9115628614136 | 778.219431594727 | 409.72039749296613 |
| SE_192_4_24 | SE_192 | SE_192_4 | TGAGCAGGACTGATGAATAA | 3279.5087229487926 | 3314.9848715306334 | 3209.462303562268 | 3435.454412455297 | 3520.113187748901 | 3015.6254924519308 | 3515.8167219021034 | 3254.852067984945 | 3892.927457985544 | 4014.641245161698 | 3287.771686118899 | 3166.337802721846 |
| SE_192_4_4 | SE_192 | SE_192_4 | CTTAGTATACCAGTAGGCCA | 265.5253262136332 | 286.8256430899282 | 314.033542317785 | 307.1088035376705 | 391.3593751951171 | 394.171099990126 | 270.1163138039265 | 227.8301912240406 | 336.5464202957393 | 268.2745556962965 | 247.66862755126303 | 482.843301294827 |
| SE_193_1_15 | SE_193 | SE_193_1 | GCTGGGCTCGGGGACCTCGG | 643.7634439628496 | 879.4653864778286 | 837.4227795140933 | 607.711064627509 | 806.0518296743492 | 898.3635839335399 | 552.0703943482641 | 571.9388617864919 | 739.6917364283664 | 635.6940558890504 | 617.4108914311107 | 1130.5033063970227 |
| SE_193_1_16 | SE_193 | SE_193_1 | GGCTCAGAAACAGGCCTCCG | 533.2182061106431 | 858.4781443005169 | 804.8343930471534 | 636.3398513979698 | 855.8997717726312 | 825.5935347045935 | 683.3619891055512 | 1054.0691419701463 | 735.2518100392406 | 1043.2089380427835 | 1277.0780415913468 | 950.5977494241906 |
| SE_193_1_18 | SE_193 | SE_193_1 | TGTTTCTGAGCCCCTCCCCG | 16.256652625324485 | 52.9678016855965 | 43.451181955919935 | 16.917010364363207 | 41.363185996446866 | 45.91443582302566 | 9.685455148881827 | 10.398888396118036 | 34.63142583518162 | 26.244250013768134 | 11.73784964697929 | 124.19286836189059 |
| SE_193_1_23 | SE_193 | SE_193_1 | GTCCGGGGAGCGAGTCCGCG | 474.69425665947495 | 521.6828769788938 | 523.3892371963084 | 512.7155448891617 | 563.175686259315 | 292.81281713552215 | 571.4412853780278 | 705.2337039549141 | 538.1190783620528 | 981.2433477369975 | 684.3166344188926 | 464.2724050911798 |
| SE_193_1_46 | SE_193 | SE_193_1 | GGCGGAGCCTCCCATCCTTT | 405.3325387914238 | 495.6986727841576 | 464.1376254382757 | 465.8684392647714 | 415.7530489802566 | 458.7700208651237 | 490.7292394206792 | 413.11947537305286 | 561.2066955855073 | 675.0604309097026 | 341.57142472709394 | 490.9680683839226 |
| SE_193_2_11 | SE_193 | SE_193_2 | AGCTGCTGTGCCCAGCTCCA | 0.0 | 1.9987849692677924 | 3.950107450538176 | 1.3013084895664004 | 9.545350614564661 | 6.064170769078861 | 1.0761606127646475 | 3.78141396224474 | 0.0 | 0.0 | 0.0 | 0.0 |
| SE_193_2_16 | SE_193 | SE_193_2 | TTGTTTGCTCCCAGAAGCAG | 73.69682523480432 | 83.94894870924728 | 105.66537430189621 | 84.58505182181602 | 62.5750676251035 | 111.75400417302473 | 83.94052779564251 | 64.28403735782058 | 87.02255722686664 | 73.62970142751615 | 41.08247376442752 | 220.52939241831038 |
| SE_193_2_43 | SE_193 | SE_193_2 | GCCAGAAGCGATTAGAACAC | 2490.519182199711 | 2419.529205298663 | 2432.278662668882 | 2262.9754633559705 | 2404.367760357565 | 2780.855452677592 | 2541.8913673500974 | 2639.4269456328684 | 2378.024574015805 | 2274.501667859905 | 2558.8512230414854 | 2873.846187514403 |
| SE_193_2_56 | SE_193 | SE_193_2 | AAGAGGACATCATCCCCAGC | 2501.356950616594 | 2752.326902681 75 | 2791.738440667856 | 2849.8655921500741 | 2909.2107484167627 | 2672.566688944041 | 2710.848583554147 | 2760.4321924240603 | 2140.9325048364844 | 3413.2101556795114 | 2490.7716950890053 | 3147.766906518199 |
| SE_193_3_22 | SE_193 | SE_193_3 | ATAGGCTGAGAAGAGGGGTG | 9276.04598801015 | 10465.638099086162 | 10033.272924366967 | 9524.276835136485 | 10196.555645380517 | 10575.047511163666 | 8872.944252244519 | 10318.533349420759 | 9645.296087736993 | 9697.979387032154 | 10594.583091363507 | 11542.972671579457 |
| SE_193_3_24 | SE_193 | SE_193_3 | GCTTGGATAGGCTGAGAAGA | 904.9536628097296 | 1063.3536036504656 | 959.8761104807768 | 809.413880510301 | 1051.0491621148421 | 1094.1496687638003 | 810.3489414117795 | 759.1188529166166 | 661.5490319797515 | 782.9534587440827 | 774.6980767006331 | 1181.5732709570525 |
| SE_193_3_30 | SE_193 | SE_193_3 | GGGTCTGGTATCTCTCCGGG | 10.837768416882989 | 15.990279754142339 | 33.5759133295745 | 24.724861301761607 | 38.181402458258646 | 12.128341538157722 | 7.533124289352532 | 39.70484660335977 | 4.439926389125849 | 8.019076393095819 | 98.59793703462604 | 2.321362025455899 |
| SE_193_3_35 | SE_193 | SE_193_3 | CCACCAACTGAGAGACCACC | 4176.875947866704 | 3537.8493956039924 | 3459.3065998088077 | 4005.4275308853807 | 3547.6886450798656 | 3558.801931339423 | 3961.3472155866675 | 3869.331836846465 | 3577.692684357609 | 3402.275411507108 | 3923.963136985177 | 4513.8884584989955 |
| SE_193_3_39 | SE_193 | SE_193_3 | GCAGGTGCTTGGTTATTATG | 3543.9502723207374 | 3540.8475730578944 | 3506.707889215266 | 3741.2619075034013 | 3929.502669662452 | 4153.090666709151 | 3735.3534869060913 | 3726.583459772481 | 4055.4287638275505 | 3354.1609531485333 | 3563.6111528229126 | 3310.262248300112 |
| SE_194_1_0 | SE_194 | SE_194_1 | TAATTCACTTGGGTGCAAGT | 124.63433679415438 | 95.94167852485404 | 136.27870704356707 | 145.74655083143685 | 116.66539640023474 | 99.625662634867 | 144.20552211046277 | 126.67736773452879 | 147.40555611897818 | 369.6065210272346 | 220.67157336321065 | 545.5200759821363 |
| SE_194_1_2 | SE_194 | SE_194_1 | ACAAGGCTGTTAATTCACTT | 821.5028459997305 | 781.5249229837068 | 818.659769124037 | 839.3439757703283 | 772.1128052670081 | 1094.1496687638003 | 911.5080390116564 | 768.5723878221784 | 1070.0222597793295 | 699.846667033817 | 765.3077969830497 | 661.5881772549312 |
| SE_194_1_6 | SE_194 | SE_194_1 | TGCTCACACAAAGCCTGATT | 1362.3074900021918 | 1361.1725640713667 | 1315.3857810292127 | 1355.9634461281892 | 1316.1977902971937 | 1082.8876373355108 | 1421.6081694620993 | 1650.587194511099 | 1458.0718261889288 | 1299.8193826264396 | 1381.5449034494625 | 1536.7416608518051 |
| SE_194_1_7 | SE_194 | SE_194_1 | CGTATAGAAGACACCCAATC | 15961.865324385266 | 17689.24697801996 | 17962.126104459723 | 15650.837204015099 | 17257.993911132908 | 15369.20765917544 | 14721.877182620377 | 16114.49560002073 | 16774.92988339528 | 14811.963104992805 | 15274.46374561415 | 16509.526725042353 |
| SE_194_1_8 | SE_194 | SE_194_1 | CTGATTGGGTGTCTTCATA | 99.70746943535235 | 155.9052267028878 | 166.8920397852379 | 83.28374333224963 | 193.02820131675202 | 369.0481068039421 | 59.18883370205561 | 195.6881725451303 | 138.5250342407265 | 33.53431946203706 | 142.02798072844942 | 441.0587843662076 |
| SE_194_2_12 | SE_194 | SE_194_2 | GGGAAGATCTGTTAAATGTG | 462.77271140090363 | 618.6239479883817 | 683.3685889431044 | 486.68937509783376 | 597.114710666656 | 630.6737599842015 | 362.6661265016862 | 536.9607826359131 | 706.8362811488352 | 363.77446546861944 | 460.1237061615882 | 773.0135544768143 |
| SE_194_2_28 | SE_194 | SE_194_2 | ACATTCTGCTCCAGGAGCAG | 148.47742731129696 | 148.90948021045054 | 150.1040831204507 | 152.25309327926885 | 165.45274398578746 | 146.40640856776108 | 205.54667703804768 | 97.37140952728706 | 223.77229001194277 | 99.87395144128429 | 119.72606639918877 | 90.53311899278006 |
| SE_194_2_29 | SE_194 | SE_194_2 | CATTCTGCTCCAGGAGCAGA | 13.005322100259587 | 26.983597085115196 | 25.675698428498144 | 13.013084895664004 | 22.272484767317543 | 11.262031428289314 | 19.370891029763655 | 52.93979547114636 | 10.655823333902038 | 5.832055558615141 | 5.868924823489645 | 68.48017975094902 |
| SE_194_2_3 | SE_194 | SE_194_2 | CTTGTGGACTTTACACTGGA | 557.0612966277856 | 604.6324532035072 | 597.4537518938992 | 623.3267665023058 | 660.7503814304204 | 708.6416698723583 | 574.6697672163217 | 566.2667408431548 | 539.8950489177032 | 750.1481462268725 | 536.4197288669536 | 695.2479266240417 |
| SE_194_2_6 | SE_194 | SE_194_2 | ACACTGGACGGAGAAGGTGG | 2526.283817975425 | 2747.329940258581 | 2619.908766569445 | 2315.0278029386263 | 2644.062120234411 | 2353.7645685124667 | 2149.092743691001 | 1753.6307249871232 | 2444.6234698526923 | 1191.1973478471425 | 1354.5478492614102 | 1877.981878593822 |
| SE_194_3_2 | SE_194 | SE_194_3 | CTATTTATTCCTAATCGAAT | 0.0 | 1.9987849692677924 | 2.962580587903632 | 0.0 | 0.0 | 0.0 | 0.0 | 0.0 | 0.0 | 0.0 | 0.0 | 0.0 |
| SE_194_3_3 | SE_194 | SE_194_3 | ACAAGCTACGACCTAATACT | 804.1624165327178 | 978.4052425465843 | 921.3625628380296 | 870.5753795199219 | 828.3243144416666 | 1250.0854885401138 | 736.0938591310189 | 787.4794576333021 | 850.6898961565126 | 749.4191392820456 | 1103.3578668160533 | 873.9928025841459 |
| SE_194_3_4 | SE_194 | SE_194_3 | GCGAGTGTACCCTTTCTGCA | 1301.615986867647 | 400.7563863381924 | 642.8799875750882 | 966.8722077478355 | 329.8448934588455 | 3585.657544745344 | 1355.9623720834559 | 579.5016897109414 | 2185.3317687277427 | 94.04189588266915 | 443.6907166558172 | 867.0287165077782 |
| SE_194_3_8 | SE_194 | SE_194_3 | CACGAGCGTACACCGAGCAA | 20207.019213278334 | 16062.23601303598 | 15760.928727647322 | 18328.93007554275 | 16506.032401607757 | 16365.464285524109 | 19027.59579429173 | 16294.112763226405 | 17418.71920981853 | 15045.974334282237 | 16619.621315157976 | 16750.948375689768 |
| SE_194_3_9 | SE_194 | SE_194_3 | TGGAGGAGAACAAAGGAAGA | 909.2887701764828 | 1022.3785117804758 | 968.7638522444877 | 943.4486549356403 | 965.1410065837601 | 839.454496462488 | 908.2795571733625 | 1124.9706537168601 | 1029.1749369993718 | 963.0181741163252 | 1093.96758709847 | 733.5504000440641 |
| SE_195_1_10 | SE_195 | SE_195_1 | AGGAGCCTCCACTCCACCCA | 470.3591492927217 | 568.654323756687 | 512.5264417073283 | 420.3226421299473 | 463.47980206275076 | 396.7700303197312 | 450.9112967483873 | 491.5838150892162 | 533.679151972927 | 710.7817712062204 | 463.645061055682 | 583.8225494021585 |
| SE_195_1_11 | SE_195 | SE_195_1 | GAGAAGAGAGGACCAGGCAG | 1230.0867153162192 | 1656.992739523 | 1550.4171743362342 | 1362.4699885760213 | 1530.4378818685339 | 1603.5400133664245 | 1267.7172018367546 | 1293.2435708086 1 | 1448.3039881328518 | 1689.8380981087 37 | 1008.2812846755211 | 1803.6982937792334 |
| SE_195_1_17 | SE_195 | SE_195_1 | CTGGGGCTGGAAGGACGTGA | 2769.049830513604 | 2508.4751364310796 | 2483.630059525878 | 2788.704093140796 | 2349.216845695636 | 3093.593402340088 | 2632.2888588223277 | 2264.1216098820632 | 2581.3732026377684 | 2599.638765252699 | 2218.4535832790857 | 2575.55116724332 |
| SE_195_1_33 | SE_195 | SE_195_1 | AGACACCCAGTGGCCACCAA | 742.3871365564847 | 862.4757142390524 | 998.389658123524 | 808.1125720247407 | 856.9603625603607 | 891.4331030545926 | 677.9811860417279 | 901.8627299906005 | 1026.5109811658963 | 984.1593755163051 | 893.250358135124 | 1136.3067114606624 |
| SE_195_1_40 | SE_195 | SE_195_1 | GATTGCTAAAATAGCCCAGG | 50.93751155935005 | 95.94167852485404 | 93.81505195028168 | 58.55888203048802 | 97.57469517110542 | 159.4010602157872 | 21.52321225529295 | 69.95615830115769 | 95.90241000511834 | 64.88161808959345 | 178.4153146340852 | 102.13992912005955 |
| SE_195_2_1 | SE_195 | SE_195_2 | AGCCACTCAGAGCTGAGTGG | 48.769957875973454 | 78.9520062860778 | 63.20171920861082 | 63.76411598875362 | 77.42339942924669 | 55.44384703157816 | 63.49347615311142 | 58.61191641448347 | 21.311646667804077 | 21.141201399979888 | 52.820323411406804 | 17.410215190919242 |
| SE_195_2_12 | SE_195 | SE_195_2 | GCTGCATTCCACCCACCTGC | 4752.361450803191 | 4479.277116129123 | 4292.779271872363 | 4646.972616241616 | 4395.103660750661 | 4191.208311543362 | 4690.984111041098 | 4262.598888917838 | 4416.838771902395 | 3941.0115437341815 | 3888.749588044239 | 4134.345767336956 |
| SE_195_2_24 | SE_195 | SE_195_2 | TGTGTGATCCCCAAAACCCT | 3731.443665932813 | 3949.5990992731577 | 3830.6167001593963 | 3801.1220980234557 | 4017.5320142189926 | 3928.7163482532337 | 3510.43591883828 | 4224.784749295591 | 4185.962599667851 | 3519.645529624238 | 4350.047079170525 | 3405.4380913438035 |
| SE_195_2_29 | SE_195 | SE_195_2 | GACAAACGTGTCATTGCAGT | 4465.160587755791 | 3864.6507380792764 | 3929.3693864228508 | 4138.160996821153 | 3943.2903983279343 | 4249.251088904545 | 4212.09263836083 | 4321.210805332322 | 3382.3359232360717 | 3628.267564403445 | 4364.1324987469 | 3867.3891344095277 |

| | | | | | | | | | | | | | | | |
|---|---|---|---|---|---|---|---|---|---|---|---|---|---|---|---|
| SE_195_2_34 | SE_195 | SE_195_2 | GGAGACATCACAGTGAGCAA | 3147.28794826282 | 3174.0705311972542 | 3209.462303562268 | 3030.7474722001466 | 3546.628050567136 | 1984.7164617085243 | 3281.21370831941 | 2992.0437976103253 | 3118.604295721996 | 2681.287543073311 | 2845.25475442778 | 2298.1484052013398 |
| SE_195_3_25 | SE_195 | SE_195_3 | ACAAGCTAAGAAAAGCTCTG | 438.92962088376106 | 474.7114302011007 | 483.8881629092655 | 525.7286297848258 | 462.4192075500213 | 640.2031711927541 | 578.9744096673803 | 515.2176523531208 | 353.4181405744176 | 552.5872641787846 | 624.4536012192982 | 234.4575645710458 |
| SE_195_3_3 | SE_195 | SE_195_3 | TGTAAGTCACCACTGCTCTA | 587.407048195058 | 608.6300231420428 | 584.61590267965 | 618.1215325440402 | 561.0544972338562 | 511.98927493222953 | 573.5936066035571 | 748.7199645204985 | 605.6059594767657 | 581.7475419718603 | 531.7245890081618 | 489.80738737119464 |
| SE_195_3_34 | SE_195 | SE_195_3 | AGCCACCCCTGGAAGCCAGG | 67.19416418467453 | 94.94228604022014 | 81.96472959866715 | 65.06542447832003 | 101.81707322202305 | 6.064170769078861 | 73.17892166799602 | 65.22939084837677 | 73.70277805948909 | 162.56854869639707 | 22.30191432926065 | 126.51423038734649 |
| SE_195_3_42 | SE_195 | SE_195_3 | AAGAGGTCTGCCCTGTACAG | 678.4443028968751 | 904.4501985936761 | 886.7991226458205 | 770.3746258233091 | 968.3227901219484 | 903.5614445927503 | 775.9118018033108 | 883.905513670033 | 793.8588383757018 | 937.502931047384 | 895.5979280645198 | 1812.983741881057 |
| SE_195_3_8 | SE_195 | SE_195_3 | TGACTCTACTATTGTCCCCA | 242.76601253817896 | 234.8572338889656 | 272.55741408713413 | 218.6198262471553 | 264.0880336696223 | 302.34222834407467 | 215.2321225529295 | 156.9286794323267 | 311.6828325166346 | 252.23640291010486 | 177.2415296693873 | 656.9454532040194 |
| SE_196_1_14 | SE_196 | SE_196_1 | GACTCAAGATACCACCACCG | 487.6995787597345 | 516.6859145557244 | 622.1419234597628 | 528.3312467639586 | 495.2976374446329 | 670.5240250381484 | 382.03701753144986 | 405.55664744860337 | 782.3150297639746 | 357.9424099100043 | 380.306328562129 | 769.5315114386304 |
| SE_196_1_18 | SE_196 | SE_196_1 | AGTTGGCCAGGCACAATGAT | 726.1304839311603 | 757.5395033524933 | 719.9070828605826 | 865.3701455616563 | 804.9912351616197 | 847.2512874513037 | 820.0343869266613 | 623.9333037670821 | 709.5002369823106 | 790.2435281923516 | 618.5846763958086 | 554.8055240839599 |
| SE_196_1_19 | SE_196 | SE_196_1 | GGTATCTTGAGTCTAGGCAG | 14.089098941947885 | 45.972054293159225 | 72.08946097232172 | 41.641871666124814 | 57.27210368738796 | 11.262031428289314 | 20.4470516425283 | 23.633837263904624 | 56.831057780810866 | 13.122125006884067 | 42.25625872912545 | 47.58792152184593 |
| SE_196_1_2 | SE_196 | SE_196_1 | GGTGTGCTGGATGACCAGAG | 507.2075619101239 | 586.6433884800971 | 589.5535369928228 | 575.178352388349 | 690.4470277868438 | 405.43313141841526 | 532.6995033185005 | 372.46927527913687 | 604.7179741989406 | 436.67515995130873 | 434.30043693823376 | 403.9169924293264 |
| SE_196_1_35 | SE_196 | SE_196_1 | GAGATGAGTCACGGTCCTGT | 3354.289325025285 | 3542.846358027162 | 3531.3960607811296 | 3507.0263793814493 | 3190.2682942900556 | 3227.0051592598224 | 3698.764026072093 | 3629.2120502451944 | 3470.2464657407636 | 3715.019390837845 | 3949.7864062085314 | 3016.6099520799407 |
| SE_196_2_27 | SE_196 | SE_196_2 | TCCACAATAGGAGGCCTGGG | 869.1890270340157 | 1126.3153301824009 | 1143.556106930802 | 1022.8284727991908 | 1128.4725615440889 | 1111.4758709611683 | 894.289469207422 | 1272.445798288625 | 1143.7250378388187 | 1132.1477853161643 | 1285.2945363442323 | 1122.3785393079272 |
| SE_196_2_34 | SE_196 | SE_196_2 | GGTGCCCCAGAAATAGGAAA | 33.59708209233727 | 58.96415659339988 | 55.301504307534465 | 39.03925468699201 | 44.54496953463509 | 220.04276790657582 | 64.56963676587885 | 52.93979547114636 | 36.40739639083196 | 3.6450347241344634 | 109.1620017169074 | 38.30247342002233 |
| SE_196_2_40 | SE_196 | SE_196_2 | GCAGTGACGGGTCTACAGGC | 2896.935497832823 | 2984.185959116814 | 2818.401665958989 | 2510.2240763735863 | 2778.757623351046 | 3567.4650324381073 | 2781.875183996614 | 3227.4368167588154 | 2589.3650701381953 | 2210.3490567151384 | 3514.3121843055997 | 2783.313068521623 |
| SE_196_2_5 | SE_196 | SE_196_2 | TACTGGGCCCCAGGAGACCA | 1141.2170142977789 | 1383.1591987333122 | 1394.3879300399763 | 1155.5619387349636 | 1288.6223329662291 | 1162.5881674434045 | 1028.809545803003 | 1233.6863051758214 | 1419.8884592424465 | 1242.956840929852 | 1137.3976307922933 | 1181.5732709570525 |
| SE_196_2_8 | SE_196 | SE_196_2 | ACTACCTGTATTTTACAAAA | 45.51862735090855 | 79.95139877071169 | 68.13935352178353 | 46.847105624390416 | 83.78696650562313 | 238.2352802138124 | 44.12258512335055 | 33.08737216946648 | 89.68651306034215 | 117.37011811712972 | 19.954344399864794 | 155.53125570554522 |
| SE_197_1_15 | SE_197 | SE_197_1 | TCGTGGACGAGGCAGCAGCA | 12081.944231141157 | 8951.558484865807 | 8972.669073897467 | 10692.851858767113 | 9298.23209309871 | 9741.657185470256 | 12013.18092029176 | 10109.610228007843 | 8538.866431566832 | 9126.437942287868 | 9783.497680757238 | 8590.200175199554 |
| SE_197_1_20 | SE_197 | SE_197_1 | ACGATTCTTCCCCAAGCCAA | 4068.498263697874 | 4583.213934531048 | 4519.910450278308 | 4407.531854161398 | 4754.64520056593 | 4909.379392624272 | 4130.304431790717 | 4992.411783627213 | 4813.768191090246 | 4343.4233772786265 | 5401.758407539869 | 4170.326878731522 |
| SE_197_1_29 | SE_197 | SE_197_1 | TGATGACCGCTCTCCCCCAC | 12670.435056177903 | 10597.557907057835 | 10659.364955277268 | 11309.672082821586 | 10803.215706661736 | 11192.726619499841 | 12244.555452036158 | 10933.013118282279 | 11138.88732503893 | 11134.123068341132 | 11701.462313073655 | 11193.607686748344 |
| SE_197_1_4 | SE_197 | SE_197_1 | ATCTACAGAGGAAGAAGTGG | 153.89631151973845 | 215.86877668092157 | 183.6799964500252 | 156.15701874796804 | 209.99771352042254 | 238.2352802138124 | 89.32133085946575 | 209.86847490347307 | 169.60518806460743 | 146.5303959102054 | 342.74520969179525 | 185.70896203647192 |
| SE_197_1_7 | SE_197 | SE_197_1 | AGGCGCCCTGACTCCCCCTG | 2652.0019316112675 | 2997.178061417055 | 2868.7655359533505 | 2658.573244184156 | 2895.42301975128 | 2259.33676653681 | 2550.5006522522144 | 2834.1697646874427 | 2761.634214036278 | 2599.638765252699 | 3385.195838188827 | 2004.4961089811686 |
| SE_197_2_12 | SE_197 | SE_197_2 | GGAGACAGCTGTCTACACAA | 573.3179492531101 | 651.6038999813003 | 636.9548263992809 | 722.2262117093522 | 590.7511435902795 | 619.4117285559122 | 676.9050254289633 | 650.4032015026553 | 759.2274125405202 | 776.3923962406407 | 535.2459439022556 | 710.336779789505 |
| SE_197_2_17 | SE_197 | SE_197_2 | TGTTTCCGCTGGCAGAGCAG | 67.19416418467453 | 87.94653864778286 | 94.80257881291622 | 53.35364807222242 | 85.90815553108195 | 103.09090307434064 | 50.57954879993843 | 64.28403735782058 | 28.415528890405433 | 55.404527806843845 | 42.25625872912545 | 140.44240254008187 |
| SE_197_2_22 | SE_197 | SE_197_2 | TGTCACTTCGTAATTTTGCA | 369.56790301570993 | 406.75274124599576 | 414.7612823065085 | 406.0082487447169 | 438.025533757245 | 324.8662912006533 | 270.1163138039265 | 535.0700756548007 | 441.3286830791094 | 151.63344452399366 | 717.1826134304347 | 305.2591063474507 |
| SE_197_2_32 | SE_197 | SE_197_2 | ATAGGGCTAGACAGGCCCTT | 332.71949039830776 | 394.760031430389 | 405.87354054279757 | 296.6983356211393 | 304.3906251533397 | 304.0748485638115 | 298.0964897358073 | 381.92281018469873 | 401.36934557697674 | 253.69441679975864 | 176.06774470468935 | 594.2686785167101 |
| SE_197_2_33 | SE_197 | SE_197_2 | TGACCTTTCTGCCCTCCTCT | 1018.750231187001 | 1086.3396307970452 | 1112.9427741891311 | 1176.382874568026 | 1161.3509914387005 | 1057.764644149327 | 1278.478807964401 | 920.7742998017242 | 1112.6455531149377 | 1159.8500492195863 | 846.2989595472068 | 905.3311899278006 |
| SE_197_3_1 | SE_197 | SE_197_3 | GGATCTCCTATTTAGCTCAA | 26.010644200519174 | 27.982989569749094 | 23.700644703229056 | 28.62878677046081 | 44.54496953463509 | 14.727271867762948 | 23.675533480822246 | 34.03272566002266 | 21.311646667804077 | 39.36637502065221 | 51.646538446708874 | 0.0 |
| SE_197_3_27 | SE_197 | SE_197_3 | CTACAGAAGCCCCATAGGGG | 0.0 | 1.9987849692677924 | 0.987526862634544 | 0.0 | 0.0 | 0.0 | 0.0 | 0.0 | 0.0 | 0.0 | 0.0 | 0.0 |
| SE_197_3_3 | SE_197 | SE_197_3 | AAGGCCTGCAAAGAAGGTAG | 2413.5710264398417 | 2279.614257449917 | 2279.2119989605276 | 2302.014718042962 | 2288.76295847006 | 2322.577404557204 | 2441.8084303629853 | 2481.552912709986 | 1860.3291570437307 | 2270.8566331357706 | 2361.6553489722332 | 2915.630703972609 |
| SE_197_3_31 | SE_197 | SE_197_3 | CGATGATCTTAGGATCTGCA | 503.95623138505897 | 110.93256579436247 | 202.44300684008152 | 477.58021567086894 | 90.15053358199957 | 1141.7967248065627 | 439.07353000797616 | 274.1525122612937 | 630.4695472558706 | 40.82438891030599 | 126.76877618737633 | 345.88294179292893 |
| SE_197_3_9 | SE_197 | SE_197_3 | GAGAACCCACTGCAGGAAAA | 39.01596630077876 | 9.993924846338961 | 20.738064115325425 | 36.43663770785921 | 4.242378050917627 | 47.64705604276248 | 101.15909759987686 | 11.34424188667422 | 53.27911666951019 | 0.7290069448268927 | 7.042709788187574 | 26.695663292742836 |
| SE_197_4_10 | SE_197 | SE_197_4 | TCATGGAGACAGTCAACAGA | 2210.90475704413 | 2170.6804766248224 | 2280.1995258231623 | 2072.9844238792757 | 2046.9474095677551 | 2118.128218628259 | 2181.37756207394 | 2126.1000002608603 | 2029.046359830513 | 2044.135473294607 | 1887.4462232342698 | 2303.9518102649795 |
| SE_197_4_15 | SE_197 | SE_197_4 | GAGCTTGCCTTCATGGTGGA | 859.4350354588211 | 737.5516536598154 | 794.9591244208079 | 818.5230399372658 | 706.3559454777849 | 627.2085195447279 | 935.1835724924787 | 758.1734994260604 | 642.901341145423 | 1313.6705145780606 | 893.250358135124 | 928.5448101823596 |
| SE_197_4_25 | SE_197 | SE_197_4 | TATGCACCCACCCAAAGGCT | 2540.3729169173725 | 2621.40648719471 | 2571.519950300353 | 2508.9227678840202 | 2843.4538886275395 | 1852.1710148986579 | 2555.881455316038 | 2343.531303088783 | 2564.5014823590905 | 2562.4594110665275 | 1997.7820099158753 | 2312.076577354075 |
| SE_197_4_27 | SE_197 | SE_197_4 | GCCTCATCCTAGCCTTTGGG | 612.3339155538889 | 772.5303906220017 | 738.670093250639 | 704.0078928554226 | 596.0541161539265 | 748.4919349263051 | 568.2128035397338 | 701.4522899926893 | 846.2499697673868 | 675.0604309097026 | 458.9499211968903 | 923.9020861314477 |
| SE_197_4_36 | SE_197 | SE_197_4 | TCTAGGCCCTTCTGTCCTGG | 450.8511661423323 | 439.7326932389143 | 457.2249373997939 | 421.6239506195137 | 401.96532032444514 | 691.3154676749901 | 451.98745736115194 | 455.6603824480812 | 336.5464202957393 | 364.50347241344633 | 386.17525338561865 | 486.32534433301083 |
| SE_198_1_0 | SE_198 | SE_198_1 | ACTTTATTATTTAATAACTG | 0.0 | 2.9981774539016888 | 0.987526862634544 | 0.0 | 0.0 | 6.93048087894727 | 0.0 | 0.0 | 0.8879852778251698 | 2.187020834480678 | 0.0 | 0.0 |
| SE_198_1_12 | SE_198 | SE_198_1 | GCTTGATGATAAAATTGAGA | 294.7873009392173 | 370.7746117991755 | 402.91095995489394 | 391.6938553594865 | 372.26867396802174 | 421.02671339604666 | 360.5138052761569 | 421.6276567880585 | 366.7379197417951 | 344.82028490312024 | 497.6848250319219 | 400.43494939114254 |
| SE_198_1_2 | SE_198 | SE_198_1 | ACTTCAGCCACGCATAAAGA | 523.4642145354484 | 625.619695380819 | 666.5806322783172 | 726.1301371780514 | 694.6894058377615 | 753.6897955855156 | 606.9545855992611 | 570.0481548053796 | 392.48949279872505 | 443.96522939957765 | 549.3313634786308 | 1192.019400071604 |
| SE_198_1_3 | SE_198 | SE_198_1 | AGCTGCTCCTTCTTTATGCG | 221.09047570441297 | 246.84994370457235 | 240.95655448282875 | 195.19627343496006 | 208.93711900769313 | 508.5240344927559 | 241.05997725928103 | 162.60080037566382 | 248.63587779104753 | 206.3089653860106 | 261.75404712763816 | 355.1683898947525 |
| SE_198_1_4 | SE_198 | SE_198_1 | GTACCACTACACCCTAGCCT | 1675.51899725011 | 1600.0273678988679 | 1394.3879300399763 | 1738.548142060711 | 1554.8315556613104 | 1412.0854790855062 | 1637.9164526277934 | 1421.8116497965023 | 1558.414162583173 | 1817.4143134534434 | 1233.6479978975235 | 1210.5902962752511 |
| SE_199_1_17 | SE_199 | SE_199_1 | CTTCCTGTTCTAGAGCTGAG | 53.105065242726646 | 56.96537162413208 | 83.93978332393624 | 65.06542447832003 | 78.4839939419761 | 35.51871450460476 | 43.0464245105859 | 26.46989773557318 | 59.49501361428638 | 24.057229179287457 | 93.90279717583432 | 0.0 |
| SE_199_1_22 | SE_199 | SE_199_1 | ATAGCCTTCCACTAGGCAGC | 286.1170862057109 | 294.8207829669994 | 302.1832199661705 | 305.8074950481041 | 287.42111294966924 | 239.9679004335492 | 260.43086828904467 | 313.8573588646534 | 262.8436422362503 | 228.1791737308174 | 73.94845277596953 | 395.79222534023074 |
| SE_199_1_25 | SE_199 | SE_199_1 | CTATAATTAGCAGAGTGGAG | 28.17819788389577 | 43.97326932389143 | 28.63827901640178 | 27.32747828089441 | 22.272484767317543 | 22.524062856578627 | 21.52321225529295 | 15.12565584889896 | 86.13457194904147 | 54.67552086201695 | 26.997054188052367 | 51.069964560029774 |
| SE_199_1_4 | SE_199 | SE_199_1 | GTTTTGTTTTCTGCCAGCAA | 1.083776841688299 | 3.9975699385355847 | 2.962580587903632 | 1.3013084895664004 | 3.1817835381882205 | 0.0 | 0.0 | 0.0 | 0.0 | 0.0 | 0.0 | 0.0 |
| SE_199_1_7 | SE_199 | SE_199_1 | TCAAGTGCTCCCAAGAGCAG | 630.75812186259 | 637.6124051964258 | 619.1793428718591 | 670.1738721266962 | 626.8113570230794 | 477.3368705374932 | 662.9149374630229 | 627.7147177293068 | 724.5959867053385 | 810.6557226475046 | 708.9661186775492 | 595.4293595294381 |
| SE_2_1_23 | SE_2 | SE_2_1 | CTAAAGCTGCATGGAGACCC | 550.5586355776559 | 554.6628289718124 | 577.7032146412082 | 606.4097561379426 | 623.6295734848911 | 771.8823078927521 | 673.6765435906693 | 687.2719876343465 | 546.9989311403045 | 853.6671323922913 | 733.6156029362056 | 676.6770304203945 |
| SE_2_1_38 | SE_2 | SE_2_1 | TGAAATCCACTCTCCCCGGA | 215.67159149597148 | 317.806810113579 | 311.0709617298814 | 187.38842249756166 | 380.753430069857 | 272.88768460854874 | 224.91756806781132 | 289.2781681101926 | 436.0007714121584 | 325.86610433762104 | 234.7569929395858 | 446.86218990026055 |
| SE_2_1_5 | SE_2 | SE_2_1 | GCTGGGGCTGGGGTGAGGAG | 1853.258399286991 | 2177.6762240172598 | 2145.895872504864 | 2001.4124569531239 | 1911.191311938391 | 2020.235176213129 | 2029.6389156741252 | 2311.3892844098723 | 2279.458208177211 | 2407.1809318183996 | 1855.7540291874259 | 2685.815863452475 |
| SE_2_1_7 | SE_2 | SE_2_1 | TAAATCTTAGGCCTTCACGC | 354.39502723207374 | 353.78493956039927 | 345.63440192209043 | 327.92973937073293 | 291.6634910005869 | 232.17110944473353 | 263.65935012733866 | 256.19079594072616 | 232.6521427901945 | 255.15243068941243 | 245.32105762186717 | 140.44240254008187 |
| SE_2_1_9 | SE_2 | SE_2_1 | ATACTGCAAGGAATTAGAGG | 244.93356622155557 | 296.81956793626716 | 299.22063937826687 | 234.2355281219521 | 239.69435987684594 | 297.1443676848642 | 233.5268529699285 | 377.1960427319178 | 190.02884945458632 | 293.0607918204108 | 286.4035313862947 | 248.38573672378118 |
| SE_2_2_12 | SE_2 | SE_2_2 | CCTGAACAGGAAATTTCCTG | 1160.724997448168 | 1553.0559211210746 | 1490.178035715527 | 1413.2210196691108 | 1669.3757630360863 | 2116.3955984085223 | 1163.329622398584 | 2128.936060732529 | 1778.634511483815 | 1514.1474244054561 | 2490.7716950890053 | 2266.810017857685 |
| SE_2_2_24 | SE_2 | SE_2_2 | GCCTTCCCGGTGCTTTCCTA | 639.4283365960963 | 858.4781443005169 | 909.5122404864151 | 674.0777975953954 | 1058.473323703948 | 1108.0106305216948 | 532.6995033185005 | 1342.4019565897827 | 781.4270444861494 | 818.6747990406004 | 1362.7643440142956 | 762.5674253622627 |
| SE_2_2_3 | SE_2 | SE_2_2 | TGGGTAAGAACAACAAGCAA | 201.5824925540236 | 236.8560188582334 | 230.09375899384875 | 249.85122999674888 | 144.2408537311993 | 241.70052065328605 | 204.47051642528302 | 149.36585150787724 | 255.7397600136489 | 233.28222234460566 | 206.5861537868355 | 502.5748785112021 |
| SE_2_2_30 | SE_2 | SE_2_2 | TGGAACTGTTACGAAAAAAG | 50.93751155935005 | 79.95139877071169 | 56.28903117016901 | 65.06542447832003 | 85.90815553108195 | 31.187163955262715 | 32.28481838293943 | 95.48070254617468 | 13.319779167377547 | 23.328222234460565 | 188.97937931636656 | 48.74860253457388 |
| SE_2_2_9 | SE_2 | SE_2_2 | ATTCTTATCCATGTAGCCCG | 57.440172609479845 | 11.992709815606755 | 20.738064115325425 | 18.218318853929606 | 29.69664635642339 | 20.791442636841808 | 50.57954879993843 | 144.6390840550963 | 18.647690834328564 | 54.67552086201695 | 361.52576912696213 | 18.57089620364719 |
| SE_2_3_0 | SE_2 | SE_2_3 | ATTGTGACACGTGTTCTGAG | 320.7979451397365 | 267.8371858818842 | 257.744511147616 | 374.7768449951233 | 218.48246962225778 | 153.33688944670834 | 311.0104170889831 | 188.1253446206808 | 221.99631945629244 | 324.40809044796725 | 254.7113373394506 | 165.97738482009677 |
| SE_2_3_14 | SE_2 | SE_2_3 | CCAGGGATCTACATTAACCA | 1562.806205714527 | 1670.9842343078744 | 1679.7831933413595 | 1677.3866430510902 | 1475.2869672066047 | 2069.6148524756286 | 1838.082326602018 | 1805.6251669623134 | 1831.9136281533254 | 2503.4098485355494 | 1401.4992478493273 | 2316.719301404987 |
| SE_2_3_20 | SE_2 | SE_2_3 | GAATGGCTACTCCTGCTCAG | 1024.1691153954425 | 1096.3335556433842 | 1076.404280271653 | 1037.142866184421 | 1089.2305645731008 | 1168.6523382124833 | 1054.6374005093546 | 1045.5609605551406 | 1146.3889936722942 | 1002.3845491369774 | 1082.2297374514906 | 1382.3710861589877 |
| SE_2_3_3 | SE_2 | SE_2_3 | GCAGTTCCAGTCACTAGCTT | 1946.4632076721848 | 2102.7217876697177 | 2016.529853499739 | 2117.2289125245334 | 2024.6749248004376 | 1989.914322367735 | 2158.778189205883 | 2183.766563184787 | 2131.1646667804075 | 2024.452285784281 | 2110.4653665268766 | 2131.0103393685154 |
| SE_2_3_5 | SE_2 | SE_2_3 | GGTTACATCCTAAAATCAGA | 80.19948628493412 | 89.94532361705066 | 102.70279371399258 | 57.25757354092162 | 79.5445884547055 | 129.0802063703929 | 59.18883370205561 | 172.05433528122566 | 87.02255722686664 | 70.71367364820858 | 246.49484258656508 | 408.5597164802382 |
| SE_2_4_11 | SE_2 | SE_2_4 | CTTGACACTAGCCTTGGCAA | 909.2887701764828 | 835.4921171539372 | 839.3978332393624 | 991.5970690495972 | 890.8993906927017 | 754.556105695384 | 854.4715265351301 | 974.6594487634268 | 699.7323989262338 | 677.9764586890102 | 1090.446232204376 | 805.5126228331969 |
| SE_2_4_16 | SE_2 | SE_2_4 | AGGCCTCGATGTCTCTCACA | 786.821987065705 | 913.4447309553811 | 933.2128851896441 | 813.3178059790002 | 958.7774395073837 | 891.4331030545926 | 855.5476871478947 | 796.9329925388639 | 913.7368508820997 | 1097.8844589093003 | 850.9940994059986 | 1029.524058289691 |
| SE_2_4_7 | SE_2 | SE_2_4 | GCCAGTGTGAGGGAAGAGGA | 3200.3930135055466 | 3507.8676210649755 | 3633.1113276324877 | 3065.8828014184396 | 3659.0510689164535 | 2425.6683076315444 | 2930.385348558135 | 2206.455046958136 | 3368.128158790869 | 1873.5478482051142 | 1394.4565380611398 | 1424.155602617194 |
| SE_2_4_8 | SE_2 | SE_2_4 | TTATCCTGATTTTACAGACA | 23.843090517142578 | 28.982382054382988 | 17.77548352742179 | 16.917010364363207 | 18.030106716399914 | 0.8663101098684087 | 19.370891029763655 | 2.836060471668555 | 0.0 | 62.69459725511277 | 2.347569929395858 | 0.0 |
| SE_2_4_9 | SE_2 | SE_2_4 | GAAGCTATAAACAGGACTGG | 972.1478269944041 | 1053.3596788041266 | 1097.1423443869785 | 1136.0423113914676 | 1156.0480188750535 | 1574.0854696308986 | 1051.4089186710605 | 1055.9598489512587 | 1261.8270797895664 | 1234.937764536756 | 1250.0809874032943 | 1130.5033063970227 |
| SE_2_5_0 | SE_2 | SE_2_5 | GCTCATTACAGCGTATGCGG | 935.299414377002 | 1136.30925502874 | 1092.2047100738057 | 1042.3481001426867 | 1138.0179121586534 | 932.1496782184078 | 902.8987541095393 | 1181.6918631952312 | 1364.833372017286 | 1586.3191119433184 | 1204.3033737800752 | 1198.9834861479717 |
| SE_2_5_13 | SE_2 | SE_2_5 | ATTGCTCATCCCCAAGAGCA | 96.4561389102586 | 114.93013573289807 | 125.4159115545871 | 92.39290275921444 | 116.66539640023474 | 84.03208065723565 | 102.23525821264151 | 104.93423745173654 | 103.0062922277197 | 48.114458358574915 | 161.98232512831422 | 24.37430126728694 |
| SE_2_5_24 | SE_2 | SE_2_5 | GTGAAATGCTTGGCACAAGG | 762.9788965485625 | 751.5431484446899 | 808.7845004976915 | 711.8157437928211 | 684.0834607104673 | 570.0320522934129 | 891.0609873691282 | 786.534104142746 | 618.9257386441434 | 821.590826819908 | 686.6642043482885 | 695.2479266240417 |
| SE_2_5_4 | SE_2 | SE_2_5 | TTATTCCTGAGTGTGATTGG | 54.188842084414944 | 57.964764108765976 | 57.27655803280356 | 46.847105624390416 | 94.3929116329172 | 25.12299318618385 | 36.58946083399801 | 109.66100490451745 | 16.871720278678225 | 43.01140974478667 | 24.64948425865651 | 9.285448101823595 |
| SE_2_5_6 | SE_2 | SE_2_5 | GCAGCTGCTAAATTTAGCAA | 1122.7928079890776 | 1264.2314930618786 | 1310.44814671604 | 1173.7802575888932 | 1202.7141774351473 | 1990.7806324776031 | 996.5247274200635 | 1509.7295244182274 | 1274.2588736791186 | 907.6136463094814 | 1941.4403316103746 | 2303.9518102649795 |
| SE_2_6_0 | SE_2 | SE_2_6 | TGGGCAAAGCCACAAAGCAA | 192.9122778205172 | 208.8730292884843 | 229.10623213121423 | 229.03029416368648 | 173.9375000876227 | 198.3850151598656 | 213.0798013274002 | 200.41493999791123 | 174.04511445373328 | 277.7516459790461 | 280.53460656280504 | 70.80154177640492 |
| SE_2_6_14 | SE_2 | SE_2_6 | GCTAGGAGAAAGAAGGATGG | 2812.4009041811355 | 2963.198716939502 | 2967.518222216805 | 2843.359049702585 | 2802.0907026310924 | 3186.2885840960075 | 2892.7197271113723 | 3108.322276948736 | 3230.490440727968 | 3425.6036337415685 | 3564.7849377876105 | 3429.8123926110907 |
| SE_2_6_17 | SE_2 | SE_2_6 | CGCAGGGTTAAAACTCTTGG | 2469.9274222076333 | 3195.0577733745663 | 3185.761658859039 | 2853.769517619116 | 3232.6920747992317 | 3366.4810869486364 | 2533.28208244798 | 2690.4760341229025 | 3133.700045445024 | 4137.843418837443 | 2770.1325166871125 | 3246.4247926000744 |
| SE_2_6_2 | SE_2 | SE_2_6 | TTGCCCATCCCTGACCTGCG | 761.8951197068741 | 703.5723091822629 | 671.5182665914899 | 679.283031553661 | 774.233994292467 | 751.9571753657788 | 716.7229681012552 | 659.8567364082171 | 465.304285580389 | 885.0144310198477 | 823.9970452179462 | 598.911402567622 |
| SE_2_6_33 | SE_2 | SE_2_6 | GAACAGCTCTCGAGAGCAGC | 192.9122778205172 | 202.87667438068092 | 232.06881271911786 | 214.71590077845607 | 162.27096044759924 | 296.2780575749958 | 173.26185865510826 | 76.57363273505099 | 234.42811334584482 | 329.5111390617555 | 72.7746678112716 | 362.13247597112024 |
| SE_20_1_0 | SE_20 | SE_20_1 | TCGTTATCGCTCATTTCCTA | 1329.7941847515428 | 1313.2017248089396 | 1214.6580410404893 | 1231.0378311298148 | 1338.4702750645113 | 1098.4812193131422 | 1404.389599657865 | 1216.6699423458101 | 1318.6581375703772 | 1167.869125612682 | 964.8512409816976 | 942.4729823350949 |
| SE_20_1_2 | SE_20 | SE_20_1 | CGCTCATTTCCTACGGCCCA | 1550.8846604559558 | 1842.8797416649045 | 1763.7229766652956 | 1773.6834712790037 | 1882.555260094697 | 2054.021270497997 | 1746.6086745170228 | 1553.215784983812 | 1865.6570687106816 | 2145.467438625545 | 1689.0765642003198 | 1323.1763545098624 |
| SE_20_1_3 | SE_20 | SE_20_1 | ATTTCCTACGGCCCAGGGAG | 1.083776841688299 | 8.994532361705065 | 7.900214901076352 | 7.807850937398403 | 4.242378050917627 | 0.0 | 0.0 | 0.0 | 0.0 | 0.0 | 0.0 | 1.1606810127279494 |
| SE_20_1_46 | SE_20 | SE_20_1 | GGAGAGCGACGTGAGTCCTG | 5749.4361451564255 | 5312.770448313792 | 5146.002481188609 | 5851.984277580103 | 4793.887197536918 | 4986.48099240256 | 6077.0789802819645 | 5982.196888239539 | 5138.770802774257 | 4607.323891305961 | 6530.939543579278 | 5281.09860791217 |
| SE_20_1_48 | SE_20 | SE_20_1 | GTGGAGAGCGACGTGAGTCC | 1551.968437297644 | 1275.2248103928516 | 1210.707933589951 | 1329.9372763368613 | 1200.5929884096884 | 1122.7379023894578 | 1621.7740434363238 | 1125.9160072524164 | 1236.9634920104616 | 893.0335074129435 | 1090.446232204376 | 1102.646962091552 |

| | | | | | | | | | | | | | | | |
|---|---|---|---|---|---|---|---|---|---|---|---|---|---|---|---|
| SE_20_2_11 | SE_20 | SE_20_2 | GAGCGTTTTCGGGCACACCC | 679.5280797385634 | 918.4416933785506 | 916.4249285248569 | 740.4445305632819 | 794.3852900343256 | 877.5721412966981 | 677.9811860417279 | 820.5668298027686 | 848.0259403230372 | 1099.342472798954 | 1090.446232204376 | 805.5126228331969 |
| SE_20_2_12 | SE_20 | SE_20_2 | GGGTGTGCCCGAAAACGCTC | 593.9097092451879 | 662.5972173122732 | 707.0692336463335 | 592.0953627527122 | 792.2641010088669 | 744.1603843769631 | 625.2493160162602 | 812.0586483877629 | 768.1072653187719 | 668.4993684062606 | 750.0485924419767 | 1071.3085747478974 |
| SE_20_2_2 | SE_20 | SE_20_2 | CTGTCCATAGCCCAGCACTT | 115.96412206064798 | 132.9192004563082 | 155.0417174336234 | 87.18766880094883 | 123.02896347661118 | 58.04277736118338 | 78.55972473181926 | 118.16918631952312 | 110.99815972814622 | 32.076305572383276 | 143.20176569314734 | 149.7278506419055 |
| SE_20_2_4 | SE_20 | SE_20_2 | ATAGCCCAGCACTTCGGCAG | 756.4762354984326 | 880.4647789624626 | 844.3354675525352 | 899.2041662903827 | 965.1410065837601 | 914.8234760210396 | 765.1501956756643 | 937.7906626317355 | 840.0340728226106 | 650.2741947855883 | 984.8055853815624 | 1228.0005114661706 |
| SE_20_2_8 | SE_20 | SE_20_2 | CAGGACCACATCATGCAACA | 13333.706483291142 | 11165.212838329888 | 11219.292686391054 | 12273.941673590289 | 11557.298405212345 | 12056.437799038644 | 13096.87465734576 | 10495.314452154766 | 11788.004563129129 | 11075.80251275498 | 9968.955705179511 | 11704.307332348642 |
| SE_20_3_11 | SE_20 | SE_20_3 | GAGCTTCCTTGAAATTGCTA | 43.351073667531956 | 99.93924846338962 | 82.9522564613017 | 62.46280749918722 | 88.02934455654076 | 42.44919538355203 | 71.02660044246673 | 112.49706537618601 | 69.26285167036325 | 106.43501394472632 | 227.71428315139823 | 153.20989368008932 |
| SE_20_3_13 | SE_20 | SE_20_3 | GCTTCCTTGAAATTGCTACG | 80.19948628493412 | 115.92952821753195 | 125.4159115545871 | 83.28374333224963 | 88.02934455654076 | 42.44919538355203 | 39.817942672291956 | 79.40969320671954 | 87.9105425046918 | 29.160277793075707 | 123.24742129328254 | 106.78265317097134 |
| SE_20_3_17 | SE_20 | SE_20_3 | TGAAATTGCTACGGGGAGCG | 384.7407787993461 | 514.6871295864565 | 466.11267916350477 | 391.6938553594865 | 387.1169971462335 | 631.54007009407 | 317.467380765571 | 469.84068480642395 | 458.2004033577876 | 577.373500302899 | 511.77024460829705 | 827.565562075028 |
| SE_20_3_18 | SE_20 | SE_20_3 | AATTGCTACGGGGAGCGGGG | 36.84841261740216 | 80.95079125534559 | 52.338923719630834 | 46.847105624390416 | 50.90853661101153 | 63.24063802039384 | 38.74178205952731 | 104.93423745173654 | 33.74344055735645 | 139.96933340676338 | 173.7201747752935 | 24.37430126728694 |
| SE_20_3_25 | SE_20 | SE_20_3 | ACAGAGCCCAGGACAGGAGT | 2477.5138600994514 | 2663.3809715493335 | 2725.5741408713416 | 2756.171380901636 | 2519.9725622450705 | 3099.6575731091666 | 3028.315964319718 | 2545.8369500678064 | 2652.4120248637823 | 3616.6034532862145 | 2899.2488628038845 | 2472.250557110532 |
| SE_20_4_16 | SE_20 | SE_20_4 | GTGCTTAGAGATCTTCCATT | 226.50935991285448 | 210.8718142577521 | 211.33074860379241 | 258.9603894237137 | 296.9664635642339 | 71.03742900920952 | 343.2952354719225 | 224.04877726181584 | 89.68651306034215 | 250.77838902045107 | 333.35492997421187 | 97.49720506914775 |
| SE_20_4_18 | SE_20 | SE_20_4 | AAGATCTCTAAGCACAGTGA | 358.73013459882696 | 367.7764343452738 | 427.59913152075757 | 411.21348270298256 | 349.99618920070424 | 328.3315316401269 | 434.7688875569176 | 346.9447310341199 | 455.5364475243121 | 425.0110488340784 | 287.5773163509926 | 475.87921521845925 |
| SE_20_4_2 | SE_20 | SE_20_4 | GCAGAGGGTGGAAGACAGAG | 5479.575711576039 | 6055.319064396777 | 5994.288056191683 | 5768.700534247853 | 6133.418067114159 | 6505.122615001881 | 5675.671071720751 | 5901.841841542263 | 6381.950191729496 | 6803.092809124562 | 6707.007288283967 | 7896.11292958824 |
| SE_20_4_24 | SE_20 | SE_20_4 | TCTCACCCTGAGGCCTACAG | 248.18489674662044 | 311.8104552057756 | 357.48472427370496 | 242.04337905935049 | 255.60327756778702 | 336.994632738811 | 193.70891029763655 | 145.5844375456525 | 224.66027528976795 | 325.13709739279415 | 219.49778839851274 | 451.50491395117234 |
| SE_20_4_9 | SE_20 | SE_20_4 | TGACTATCAAGGTTTCCCAC | 412.9189766832419 | 449.7266180852533 | 446.3621419108139 | 443.74619494214255 | 424.2378050917627 | 202.71656570920763 | 398.1794267229196 | 595.5726990503965 | 418.241065855655 | 387.8316946479069 | 473.0353407732654 | 215.88666836739858 |
| SE_20_5_0 | SE_20 | SE_20_5 | TCAGAATGAGCAGAACCAAC | 2006.0709339650414 | 1657.9921320076337 | 1639.2945919733431 | 1970.1810532035302 | 1787.1017539490504 | 1736.9517702861594 | 2214.7385410696447 | 2242.378479599271 | 1813.2659373189967 | 1518.5214660744175 | 2517.768749277058 | 2059.048116579382 |
| SE_20_5_21 | SE_20 | SE_20_5 | GAGTAGTCTGGATTAGAGCA | 230.84446727960767 | 275.83232575895534 | 281.445155850845 | 300.6022610898385 | 343.6326221243278 | 248.6310015322333 | 338.99059302086397 | 279.82463320463074 | 277.051406681453 | 569.3544239098031 | 140.85419576375148 | 140.44240254008187 |
| SE_20_5_24 | SE_20 | SE_20_5 | CTCATTACTTCCCTGCTCCG | 289.36841673077583 | 366.7770418606399 | 394.02321819118305 | 335.7375903081313 | 333.02667699703375 | 337.86094284867943 | 279.80175931880837 | 518.0537128247894 | 380.9456841869978 | 216.5150626135871 | 524.6818792199742 | 211.2439443164868 |
| SE_20_5_4 | SE_20 | SE_20_5 | GGCAGGTGTTGCCTTGAAGA | 1007.9124627701179 | 1020.379726811208 | 1056.6537430189621 | 934.3394955086756 | 986.3528968383483 | 693.9143980045953 | 1054.6374005093546 | 922.6650067828366 | 1043.3827014445744 | 1081.8463061231087 | 710.1399036422471 | 948.2763873987346 |
| SE_20_5_44 | SE_20 | SE_20_5 | TTTCCCTCCCTAGACCACCT | 35.76463577571386 | 62.96172653193546 | 48.38881626909266 | 61.16149900962082 | 80.60518296743491 | 22.524062856578627 | 31.208657770174778 | 129.51342820619735 | 67.4868811147129 | 4.374041668961356 | 9.390279717583432 | 113.74673924733905 |
| SE_20_6_2 | SE_20 | SE_20_6 | TGACCAAAGCCAGCAAGGCG | 27.094421042207472 | 41.97448435462364 | 34.56344019220904 | 19.519627343496005 | 54.09032014919974 | 24.256683076315444 | 5.380803063823238 | 79.40969320671954 | 39.95933750213264 | 13.122125006884067 | 24.64948425865651 | 4.642724050911798 |
| SE_20_6_3 | SE_20 | SE_20_6 | GTTACATGCAGGGGAAAAGA | 358.73013459882696 | 457.72175796232443 | 413.77375544387394 | 415.11740817168175 | 335.1478660224925 | 236.50265999407557 | 387.4178205952731 | 445.26149405196315 | 347.2022436296414 | 364.50347241344633 | 443.6907166558172 | 445.70150888753255 |
| SE_20_6_4 | SE_20 | SE_20_6 | TTTGCTTTGTGCCTAGGAGG | 54.188842084414944 | 87.94653864778286 | 55.301504307534465 | 52.05233958265602 | 86.96875004381135 | 45.048125713157255 | 63.4934761531142 | 95.48070254617468 | 102.11830694989453 | 150.17543063433988 | 95.07658214053225 | 13.928172152735392 |
| SE_20_6_5 | SE_20 | SE_20_6 | GGCGTGTGTGTTACATGCAG | 2113.364841292183 | 2064.7448732536295 | 2157.7461948564787 | 2191.4034964298185 | 2246.3391779608833 | 2779.989142567724 | 2221.1955047462325 | 2238.597065637046 | 1868.3210245441571 | 2695.138675025022 | 2413.301887418942 | 2312.076577354075 |
| SE_20_6_7 | SE_20 | SE_20_6 | CGCCCCCTACACTTACACAA | 1649.508353049591 | 1821.8924994875927 | 1810.1367392091192 | 1689.0984194571877 | 1765.8898636944623 | 1648.5881390795819 | 1381.7902267898073 | 2138.3895956380907 | 1657.868513699592 | 1716.0823481225052 | 2183.240034338148 | 1900.0348178356533 |
| SE_200_1_1 | SE_200 | SE_200_1 | CTAGCTTTCACCTCAACTGA | 295.8710777809056 | 277.83111072822317 | 340.6967676089177 | 261.5630064028465 | 312.875381255175 | 311.00532944275875 | 355.1330022123337 | 248.62796801627667 | 289.48320057100534 | 326.59511128244793 | 212.45507861032516 | 292.49161520744326 |
| SE_200_1_15 | SE_200 | SE_200_1 | CCAGACCCTGGAGAGCCAGG | 59.60772629285644 | 88.94593113241676 | 88.87741763710896 | 113.21383859227684 | 69.99923784014085 | 95.29411208552496 | 78.55972473181926 | 65.22939084837677 | 79.91867500426528 | 207.0379723308375 | 30.518409082146157 | 126.51423038734649 |
| SE_200_1_25 | SE_200 | SE_200_1 | ACAAGTAGTGGGTTAAGAGG | 725.046707089472 | 890.4587030808015 | 893.7118106842623 | 801.6060295729027 | 846.3544211580665 | 709.5079799822267 | 701.6567195225501 | 977.4955092350953 | 961.6880558846589 | 443.2362224547507 | 890.9027882057281 | 855.4219063804987 |
| SE_200_1_26 | SE_200 | SE_200_1 | TGATCTCCCTGTCCAGAGCA | 199.414938870647 | 224.86330904262664 | 232.06881271911786 | 200.40150739322567 | 195.14939034221084 | 361.2513158151264 | 188.3281072338133 | 164.4915073567762 | 237.9800544571455 | 86.02281948957334 | 191.32694924576242 | 419.0058455947897 |
| SE_200_1_31 | SE_200 | SE_200_1 | TATTGAGGGCCTTAGTACAT | 730.4655912979134 | 625.619695380819 | 660.65547110251 | 651.9555532727666 | 653.3262198413146 | 698.2459485539374 | 650.0010101098471 | 810.1679414066506 | 386.27359585394885 | 655.3772433993765 | 696.0544840658719 | 468.9151291420916 |
| SE_200_2_16 | SE_200 | SE_200_2 | AGACAATGTTGGCAAAGCCA | 628.5905681792134 | 664.596002281541 | 705.0941799210644 | 666.269946657997 | 729.6890247578318 | 821.2619841552515 | 646.7725282715531 | 756.282792444948 | 835.5941464334848 | 599.2437086477057 | 651.4506554073506 | 499.0928354730183 |
| SE_200_2_34 | SE_200 | SE_200_2 | GGTTGGCTACGCACAAAGCA | 198.3311620289587 | 206.8742443192165 | 165.9045129226034 | 199.10019890365928 | 206.8159299822343 | 221.77538812631263 | 192.6327496848719 | 235.39301914849005 | 169.60518806460743 | 121.74415978609107 | 225.36671322200237 | 120.71082532370674 |
| SE_200_2_4 | SE_200 | SE_200_2 | GGGGTAGTAACACCCACGTG | 235.17957464636086 | 281.8286806667587 | 292.307951339825 | 249.85122999674888 | 246.05792695322236 | 275.486614938154 | 295.9441685102781 | 275.09786575184984 | 311.6828325166346 | 325.13709739279415 | 408.47716771487933 | 396.95290635295873 |
| SE_200_2_5 | SE_200 | SE_200_2 | TAAACCCATGAGAGAGAAGG | 86.70214733506391 | 151.90765766435223 | 124.42838469195254 | 119.72038104010883 | 134.69550311663465 | 94.42780197565655 | 54.88419125099702 | 110.60635839507364 | 131.42182111812514 | 101.33196533093808 | 77.46980767006332 | 92.85448101823596 |
| SE_200_2_8 | SE_200 | SE_200_2 | GGTAAAATACATGAATCAAG | 239.51468201311405 | 299.81774539016885 | 258.73203801025056 | 235.53683661151848 | 264.0880336696223 | 226.10693867565467 | 228.14604990610528 | 205.14170745069214 | 258.40371584712443 | 170.5876250894929 | 251.1899824453568 | 254.18914178742094 |
| SE_200_3_12 | SE_200 | SE_200_3 | ACAACTGGCTGAGTCAAAGA | 803.0786396910295 | 944.4258979790319 | 912.4748210743187 | 861.4662200929571 | 920.596037049125 | 813.4651931664358 | 770.5309987394876 | 1225.1781237608159 | 840.9220581004358 | 921.4647782611924 | 1228.9528580387316 | 1156.0382886770376 |
| SE_200_3_19 | SE_200 | SE_200_3 | TCAGGGGCTTTGAAGTCATG | 2849.2493167985376 | 3144.0887566582373 | 3076.1461771066047 | 3051.568408033209 | 2991.9371204096565 | 2799.914275094697 | 2764.6566141923795 | 2855.912894970235 | 3233.1543965614433 | 3174.8252447211175 | 2522.4638891358495 | 3494.810529323856 |
| SE_200_3_4 | SE_200 | SE_200_3 | AGTCCATTAATTATTTTCCA | 27.094421042207472 | 36.977521931454156 | 50.36386999436174 | 50.751031093089615 | 69.99923784014085 | 136.8769973592086 | 32.28481838293943 | 116.27847933841076 | 70.15083694818841 | 14.580138896537854 | 65.73195802308402 | 55.71268861094157 |
| SE_200_3_5 | SE_200 | SE_200_3 | TGGCACAGAGGCAGCAGCAA | 273.1117641054513 | 189.88457208044028 | 224.1685978180415 | 243.34468754891688 | 240.75495438957532 | 132.54544680986655 | 308.8580958634538 | 198.52423301679886 | 117.21405667292361 | 201.93492371704926 | 142.02798072844942 | 81.24767089095646 |
| SE_200_3_6 | SE_200 | SE_200_3 | AGCAGCCTCTGACAACACCA | 1551.968437297644 | 1523.074146582058 | 1461.539756699125 | 1534.242709198786 | 1569.679878839522 | 2339.903606754572 | 1671.2774316234975 | 1820.7508228112124 | 1476.7195170232574 | 1828.3494176258469 | 1454.3195712607342 | 1814.144422893785 |
| SE_201_1_1 | SE_201 | SE_201_1 | AGTGAATCAAAGCTCCAACA | 95.37236206857031 | 116.92892070216585 | 125.4159115545871 | 132.73346593577284 | 170.75571654943448 | 90.96256153618292 | 72.10276105523138 | 106.82494443284891 | 141.189659174202 | 103.51898616541875 | 127.94256115207426 | 64.99813671276517 |
| SE_201_1_17 | SE_201 | SE_201_1 | GGTGAAGACAGTTAAATCCG | 1031.7555532872605 | 1290.21569766236 | 1314.398254166578 | 1095.7017482149092 | 1189.9870432823943 | 1035.2405812927484 | 897.517951045716 | 1169.4022678180008 | 1607.2533528635572 | 1038.8348963783221 | 1220.7363632858462 | 1089.8794709515446 |
| SE_201_1_2 | SE_201 | SE_201_1 | CTTGCTTTAGGCCAGAATTA | 260.10644200519175 | 315.8080251443112 | 312.0584885925159 | 260.2616979132801 | 359.5415398152689 | 460.01066834012505 | 239.98381664651637 | 345.05402405300754 | 280.60334779275365 | 207.7669792756644 | 377.9587586327331 | 521.1457747148493 |
| SE_201_1_3 | SE_201 | SE_201_1 | TCCGGCCCATGCTCTTGCAA | 1534.6280078306313 | 1324.1950421399124 | 1379.575027100458 | 1303.9111065455331 | 1366.0457323954759 | 1030.042720633538 | 1711.0953742957895 | 1451.117608003744 | 1434.0962236876492 | 1524.3535216330326 | 1066.9705329104174 | 1086.3974279133606 |
| SE_201_1_4 | SE_201 | SE_201_1 | CAGTGCCCTTGCAAGAGCAT | 847.5134902002497 | 732.5546912366459 | 694.2313844320845 | 901.8067832695154 | 764.6886436779023 | 971.1336331624861 | 1026.6572245774737 | 695.7801690493521 | 948.3682767172813 | 883.556417130194 | 1005.9337147461251 | 651.1420481403796 |
| SE_202_1_21 | SE_202 | SE_202_1 | TTATTTGCAGTGCACCAAGC | 108.37768416882989 | 83.94896870924728 | 103.69032057662713 | 104.10467916531204 | 91.21112809472898 | 70.17111889934111 | 115.14918556581728 | 128.56807471564116 | 87.02255722686664 | 116.64111117230283 | 90.38144228174053 | 27.856344305470785 |
| SE_202_1_22 | SE_202 | SE_202_1 | ATTGTCCCGCAACAGCAGAA | 389.0758861660993 | 483.7059625628058 | 442.4120344602757 | 467.1697477543378 | 549.3879575938327 | 278.9518553776276 | 471.3583483909156 | 625.8240107481945 | 558.5427397520318 | 541.6521600063812 | 672.5787847719133 | 356.32907090748046 |
| SE_202_1_5 | SE_202 | SE_202_1 | GTAGGCACGTATTCTTCCTG | 5503.4188020931815 | 4821.069345873915 | 5231.9173182378145 | 5179.207788474274 | 4715.403203594942 | 5161.4756345959795 | 5279.643966223361 | 4033.8233442032415 | 4497.645432184485 | 5490.151301491329 | 4366.480068676296 | 5165.030506639375 |
| SE_202_1_8 | SE_202 | SE_202_1 | GGTAAGGGTTCTTAAAGGCA | 145.22609678623206 | 211.871206742386 | 263.66967232342324 | 193.89496494539367 | 244.99733244049295 | 227.8395588953915 | 176.4903404934022 | 359.2343264113503 | 400.48136029915156 | 242.75931262735526 | 409.65095267957724 | 328.4727266020097 |
| SE_202_1_9 | SE_202 | SE_202_1 | TGGAAATTACTGCTAGAATG | 46.60240419259685 | 97.94046349412183 | 93.81505159028168 | 94.99551973834723 | 97.57469517110542 | 102.22459296447224 | 121.60614924240517 | 79.40969320671954 | 133.19779167377547 | 133.40827090332135 | 19.954344399864794 | 120.71082532370674 |
| SE_202_2_22 | SE_202 | SE_202_2 | TCTGAGAGGAAAAAACCAGC | 546.2235282109026 | 745.5467935368865 | 735.7075126627353 | 614.217607075341 | 583.3269820011737 | 560.5026410848604 | 468.12986655262165 | 598.4087595220651 | 926.1686447716521 | 683.0795073027984 | 693.706914136476 | 351.68634685656866 |
| SE_202_2_3 | SE_202 | SE_202_2 | CGTTGTCTCGGGTAAATACC | 3033.4913798855487 | 2967.196286878038 | 3065.283381617625 | 2762.677923349468 | 2973.9070136932564 | 2851.0265715769333 | 2499.921103452276 | 2524.093819785014 | 2845.104830151844 | 2675.455487514696 | 3123.441791061189 | 2839.025757132564 |
| SE_202_2_34 | SE_202 | SE_202_2 | GTTGTGGGACAATCAGAGAC | 295.8710777809056 | 368.7758268299077 | 367.3599929000504 | 331.8336648394321 | 292.72408551331625 | 159.4010602157872 | 273.34479564222045 | 330.87372169466477 | 299.2510386270822 | 468.0224585788651 | 251.1899824453568 | 255.34982280014887 |
| SE_202_2_37 | SE_202 | SE_202_2 | ATAAAGGCTTTCCTGAACTC | 79.11570944324582 | 80.95079125534559 | 87.88989077447442 | 63.76411598875362 | 84.84756101835254 | 117.81817494210358 | 95.77829453605362 | 38.75949311280358 | 57.71904305863604 | 85.29381254474644 | 38.73490383503166 | 119.5501443109788 |
| SE_202_2_7 | SE_202 | SE_202_2 | TCCCCACTAACAAGTGAAAG | 39.015966300778 76 | 35.97812944682026 | 47.40128940645811 | 48.14841411395682 | 43.48437502190568 | 4.331550549342044 | 44.12258512335055 | 82.2457536783881 | 46.17523444690883 | 85.29381254474644 | 265.27540202173196 | 19.73157721637514 |
| SE_202_3_1 | SE_202 | SE_202_3 | AGCTCCTGTTATGTGCCCCA | 114.88034521895969 | 187.88578711117248 | 199.4804262521779 | 144.44524234187045 | 220.6036586477166 | 67.57218856973589 | 250.74542277416285 | 139.91231660231537 | 213.11646667804075 | 188.8127987101652 | 125.5949912226784 | 257.67118482560477 |
| SE_202_3_10 | SE_202 | SE_202_3 | GTTCGGTAAAGTCTACAAAT | 166.90163361999802 | 178.8912547494674 | 144.17892194464343 | 184.78580551842887 | 188.7858232658344 | 427.0908841651255 | 160.34793130193248 | 136.13090264009065 | 137.6377180629013 | 336.07220156519753 | 207.75993875153344 | 454.9869569893562 |
| SE_202_3_12 | SE_202 | SE_202_3 | CCTAGGATGTCAAGATACCA | 965.6451659442744 | 901.4520211397744 | 901.6120255853388 | 857.5622946242579 | 932.2625766891485 | 1450.2031239197163 | 921.1934845265382 | 1324.440240269215 | 1048.7106131115256 | 762.5412642889297 | 1079.8821675220947 | 1319.6943114716785 |
| SE_202_3_21 | SE_202 | SE_202_3 | TCATGAACATACAAGTAAAA | 387.992109324411 | 562.6579688488836 | 512.5264417073283 | 495.7985345247986 | 645.9020582522087 | 513.7218951519664 | 387.4178205952731 | 667.4195643326666 | 595.8381214206889 | 471.66749330299956 | 963.6774560169997 | 507.2176025621139 |
| SE_202_3_24 | SE_202 | SE_202_3 | TCAACATCTGGGTCATTTAA | 348.97614302363223 | 401.75577882282624 | 430.5617121086612 | 398.20039780731855 | 418.9348325281157 | 566.5668118539393 | 351.90452037403975 | 345.9993775435637 | 396.9294191878509 | 430.11409744786664 | 331.007360044816 | 869.3500785332342 |
| SE_203_1_16 | SE_203 | SE_203_1 | GCATGTCTCAGAGGAAGGCG | 2499.1893969332173 | 2576.4338253861843 | 2347.351352482311 | 2637.7523083510937 | 2359.82279082293 | 2420.470446972334 | 2605.3848435032114 | 2393.6350380882604 | 2437.519587630091 | 2279.6047164736933 | 2466.122210830349 | 2488.5000912887235 |
| SE_203_1_26 | SE_203 | SE_203_1 | CCCGGACTCCAGACAGACGG | 311.04395356454177 | 329.79951992918575 | 288.35784388928687 | 346.1480582246625 | 395.6017532480687 | 446.1497065822305 | 360.5138052761569 | 440.5347265991822 | 268.1715539032013 | 484.7896183098836 | 265.27540202173196 | 665.070220293115 |
| SE_203_1_3 | SE_203 | SE_203_1 | ACACAGAGCTGCACGCCAGA | 4739.3561287029315 | 4511.257675637407 | 4619.650663404397 | 4731.557668063432 | 4846.916923173389 | 5495.005026895316 | 4767.391514547388 | 4434.6532241990635 | 4367.99958162201 | 5791.231169704835 | 4490.901274934276 | 4661.294947115445 |
| SE_203_1_33 | SE_203 | SE_203_1 | GGGCTGCGCAGAAAGCAGCA | 121.38300626908948 | 112.93135076363026 | 91.8399982250126 | 156.15701874796804 | 125.15015250207 | 24.256683076315444 | 149.586325174286 | 51.994441980590175 | 113.66211556162173 | 31.347298627556384 | 78.64359263476125 | 26.695663292742836 |
| SE_203_1_38 | SE_203 | SE_203_1 | GCACAGACAGATGTGCGGGA | 1930.2065550468603 | 2064.7448732536295 | 2241.685978180415 | 2125.036763461932 | 2255.8845285754483 | 2567.7431656499634 | 1944.622227265718 | 2148.7884840342085 | 2246.6027528976797 | 2014.9751955015313 | 2278.3166164786803 | 3044.466296385411 |
| SE_203_2_12 | SE_203 | SE_203_2 | GCAGGTTCCCTCAGGCGCCA | 3519.0234049619066 | 3824.6750386939207 | 3711.1259497806163 | 3669.689940577249 | 3504.20427005796 | 3506.8233247473186 | 3346.8595056980535 | 3406.1086264739347 | 3892.927457985544 | 4037.9694673961585 | 3251.3843522132634 | 3564.4513900875327 |
| SE_203_2_22 | SE_203 | SE_203_2 | GCTTCTCTCTCTTCCGAGCA | 13.005322100259587 | 50.9690167163287 | 34.56344019220904 | 18.218318853929606 | 29.69664635642339 | 0.8663101098684087 | 8.60928490211718 | 4.726767452780925 | 30.191499446055772 | 15.309145841364746 | 1.173784964697929 | 0.0 |
| SE_203_2_27 | SE_203 | SE_203_2 | TCCTGAGCCATGTTTCTAGA | 525.631768218825 | 479.7083926242702 | 498.70106563044476 | 530.9338637430914 | 372.26867396802174 | 639.3368610828857 | 499.33852432279645 | 440.5347265991822 | 428.00890391173186 | 430.84310439269353 | 613.8895365370169 | 717.3008658658728 |
| SE_203_2_30 | SE_203 | SE_203_2 | GCGATTTTAGAGGTGGGCTG | 1368.8101510523215 | 1731.9471758705422 | 1657.070075500765 | 1483.4916781056966 | 1684.2240862142978 | 1257.015969419061 | 1291.392735317577 | 1634.5161851716439 | 1446.5280175772016 | 1417.9185076883061 | 1382.7186884141604 | 1221.0364253898028 |
| SE_203_2_7 | SE_203 | SE_203_2 | GCAGTTTCCCTGGGGCGCCG | 286.1170862057109 | 340.7928372601586 | 429.5741852460267 | 296.6983356211393 | 429.5407776554097 | 362.98393603486323 | 196.93739213593048 | 325.20160075132765 | 579.8543864198359 | 304.7249029376411 | 259.4064771982423 | 443.3801468620767 |
| SE_203_3_1 | SE_203 | SE_203_3 | GACGAGGAAGAGTTGTGGCG | 14934.44487846476 | 15803.3933595158 | 15644.400557856447 | 15631.317576671601 | 16220.732477683547 | 15583.186256312936 | 15051.18233012636 | 14914.842020504931 | 15379.90501193194 | 16502.53021004637 | 15503.351813730247 | 16326.139125031337 |
| SE_203_3_2 | SE_203 | SE_203_3 | GCGAGGCAGATCCTGCCCCG | 1424.0827699784247 | 1436.127000418909 | 1452.6520149354142 | 1383.2909244090836 | 1529.3772873558046 | 1324.5881579887969 | 1549.6712823810924 | 1510.6748779087836 | 1536.2145306375437 | 1488.6321813365148 | 1665.6008649063613 | 1485.6716962917753 |
| SE_203_3_3 | SE_203 | SE_203_3 | CAGATCCTGCCCCGTGGCCG | 251.43622727168534 | 311.8104552057756 | 327.8589183946686 | 234.2355281219521 | 322.42073186973965 | 427.0908841651255 | 277.64943809327906 | 349.78079150578844 | 399.5933750213264 | 389.28970853756067 | 362.69955409166005 | 394.6315443275028 |
| SE_203_3_4 | SE_203 | SE_203_3 | TACGAGACGGCCGCGGCCAC | 9346.49148271989 | 7606.376200548584 | 7725.422646390038 | 8623.771360356535 | 8020.215705259774 | 6740.7589648860885 | 9552.00159889901 | 8242.537084159378 | 7792.070812915865 | 7960.755837509668 | 8359.69651857865 | 7103.367797895051 |
| SE_203_3_5 | SE_203 | SE_203_3 | CTACGAGACGGCCGCGGCCA | 360.89768828220355 | 355.78372452966704 | 367.3599929000504 | 294.0957186420065 | 288.4817074623986 | 422.7593336157835 | 374.5038932420973 | 241.06514009182717 | 314.3467883501101 | 393.663750206522 | 227.71428315139823 | 385.3460962256792 |
| SE_204_1_1 | SE_204 | SE_204_1 | GAGTGGTGACCACCTCAGGA | 567.8990650446686 | 622.6215179269174 | 634.9797726740118 | 540.0430231700561 | 620.447789946703 | 579.5614635019655 | 518.7094153525601 | 434.86260565584513 | 713.0521780936114 | 496.4537294271139 | 514.1178145376929 | 442.21946584934875 |
| SE_204_1_15 | SE_204 | SE_204_1 | GTGATACTGGACTTGTTCCC | 818.2515154746657 | 720.5619814210391 | 734.7199858001007 | 799.0034125937699 | 689.3864332741144 | 997.1229364585384 | 880.2993812414817 | 835.6924856516675 | 657.1091055906256 | 589.0376114201292 | 1061.1016080869279 | 873.9928025841459 |
| SE_204_1_4 | SE_204 | SE_204_1 | CTCAGGATGGAGAAGCCGAT | 366.31657249064506 | 350.7867621064976 | 366.37246603741585 | 291.4931016628737 | 321.3601373570102 | 290.2138868059169 | 382.03701753144986 | 336.5458426380019 | 420.9050216891305 | 340.4462432341589 | 235.93077790428373 | 207.76190127830296 |
| SE_204_1_5 | SE_204 | SE_204_1 | AGCATTTTCCAAGTCACAAA | 61.77527997623304 | 63.961119016569356 | 67.151826659149 | 53.35364807222242 | 84.84756101835254 | 45.91443582302566 | 59.18883370205561 | 91.69928858394995 | 55.05508722516053 | 20.412194455152996 | 16.432989505771005 | 56.873369623669525 |
| SE_204_1_9 | SE_204 | SE_204_1 | TTAGCCCCCACAATTCCCAT | 430.2594061502547 | 468.71507529329733 | 494.75095817990655 | 433.33572702561133 | 499.54001549555056 | 291.08019691578534 | 366.9707689527448 | 519.9444198059017 | 343.6503025183407 | 306.1829168272949 | 509.4226746789012 | 454.9869569893562 |
| SE_204_2_0 | SE_204 | SE_204_2 | ACTTTTGGAGTACAAAGAGA | 8.670214733506391 | 14.990887269508443 | 21.725590977959968 | 1.3013084895664004 | 6.363567076376441 | 23.390372966447035 | 0.0 | 18.9070698111237 | 18.647690834328564 | 0.7290069448268927 | 0.0 | 82.4083519036844 |

| | | | | | | | | | | | | | | | |
|---|---|---|---|---|---|---|---|---|---|---|---|---|---|---|---|
| SE_204_2_15 | SE_204 | SE_204_2 | ATATCCCTGCATGCAGCCAG | 383.6570019576578 | 275.83232575895534 | 346.62192878472496 | 313.61534598550253 | 308.63300320425736 | 426.2245740552571 | 371.2754114038034 | 327.09230773244 | 418.241065855655 | 297.4348334893722 | 268.79675691582577 | 316.86591647473017 |
| SE_204_2_28 | SE_204 | SE_204_2 | TTTCTGCCTTCACCTTCACA | 8.670214733506391 | 16.989672238776237 | 27.650752153767232 | 14.314393385230405 | 29.69664635642339 | 38.11764483420998 | 10.761606127646475 | 0.945353490556185 | 36.40739639083196 | 17.496166675845423 | 0.0 | 61.51609367458132 |
| SE_204_2_7 | SE_204 | SE_204_2 | GAGTCACCACAAGCTCTCTG | 210.25270728753 | 240.85358879676897 | 272.5574140871341 3 | 265.4669318715457 | 294.8452745387751 | 380.31013823223145 | 221.6890862295174 | 251.4640284879452 | 307.2429061275087 | 334.61418767554375 | 442.5169316911192 | 358.6504329329364 |
| SE_204_2_8 | SE_204 | SE_204_2 | TTCCTGGTCTAGTCCGCACA | 9.75399157519469 | 16.989672238776237 | 20.738064115325425 | 15.615701874796805 | 29.69664635642339 | 10.395721318420904 | 13.990087965940416 | 17.01636283001133 | 5.327911666951019 | 32.80531251721017 | 83.33873249355297 | 0.0 |
| SE_205_1_1 | SE_205 | SE_205_1 | GTTGCAGTAGAAACCATTGA | 331.63571355661946 | 324.8025575060163 | 320.9462303562268 | 330.5323563498657 | 349.99618920070424 | 339.59356306841624 | 312.0865777017478 | 444.31614056140694 | 635.7974589228215 | 306.1829168272949 | 510.59645964359913 | 450.3442329384444 |
| SE_205_1_2 | SE_205 | SE_205_1 | AGAACACTGAAAACCTTCAA | 303.4575156727237 | 361.7800794374704 | 374.2726809384922 | 412.51479119254896 | 407.2682928880922 | 512.8555850420979 | 299.172650348572 | 463.2232103725307 | 361.4100080748441 | 341.1752501789858 | 299.3151659979719 | 168.29874684555267 |
| SE_205_1_3 | SE_205 | SE_205_1 | AAATGATCAACTTCAGGATA | 69.36171786805113 | 67.95868895510495 | 64.18924607124536 | 72.87327541571842 | 64.6962652764938 2 | 15.593581977631358 | 68.87427921693744 | 51.994441980590175 | 150.0695119524537 | 56.86254169649763 | 158.4609702342204 | 112.5860582346111 |
| SE_205_1_4 | SE_205 | SE_205_1 | AAAATGATCAACTTCAGGAT | 36.84841261740216 | 48.970231747060915 | 59.25161175807264 | 36.43663770785921 | 69.99923784014085 | 29.454543735525895 | 16.142409191469714 | 13.23494886778659 | 79.03068972644012 | 26.973256958595027 | 5.868924823489645 | 9.285448101823595 |
| SE_205_1_9 | SE_205 | SE_205_1 | AATAAAGATGGCATTAAAGG | 9.75399157519469 | 28.982382054382988 | 35.55096705484358 | 13.013084895664004 | 15.9089176909411 | 6.93048087894727 | 9.685445514881827 | 0.945353490556185 | 23.975602501279585 | 82.37778476543888 | 12.911634611677218 | 16.249534178191293 |
| SE_205_2_10 | SE_205 | SE_205_2 | GCTGAATGAAAAGGGCAGGG | 1825.0802014030953 | 2213.65435346408 | 2317.725546603275 | 1997.5085314844246 | 2309.974848724648 | 2479.3795344433856 | 1918.7943725593664 | 2573.2522012939357 | 2105.4130937234777 | 2273.0436539702514 | 3035.4079187088446 | 2693.9406305415705 |
| SE_205_2_11 | SE_205 | SE_205_2 | GAGGGTTTGAGGCACACAGG | 1732.95916985959 | 1473.104522350363 | 1563.2550235504832 | 1579.78850633361 | 1434.9843757228873 | 1467.5293261170843 | 1725.0854622617298 | 1619.390529322745 | 1540.6544570266697 | 1322.4185979159834 | 1698.4668439179034 | 2249.399802666766 |
| SE_205_2_21 | SE_205 | SE_205_2 | TGGGTTGCAGAGAACGTCCA | 117.04789890233629 | 145.91130275654885 | 148.12902939518162 | 126.22692348794084 | 134.69550311663465 | 63.24063802039384 | 108.6922218892294 | 94.5353490556185 | 115.43808611727208 | 194.64485426878034 | 66.90574298778195 | 168.29874684555267 |
| SE_205_2_5 | SE_205 | SE_205_2 | GTAGGGCGAGAGGCACAGAG | 6810.453673169271 | 6785.874970664155 | 6817.885459628892 | 6695.23217881913 | 6716.745049115333 | 5836.33121018347 | 6897.113367208625 | 5798.798311071639 | 6698.072950635255 | 6536.2762673179195 | 6437.0367464034425 | 6021.613094032602 |
| SE_205_2_7 | SE_205 | SE_205_2 | AGGTGAATCTGGAGGCACAA | 1964.8874139808859 | 2294.6051447194254 | 2205.147484262937 | 2320.233036896892 | 2290.8841474955184 | 2240.277944119705 | 2404.1428089162223 | 2770.831080820178 | 2408.2160734618606 | 2195.039910873774 | 3425.1045269885567 | 2559.3016330651285 |
| SE_205_3_20 | SE_205 | SE_205_3 | GGGCATTCTCCTGTGAACAC | 266.60910305532155 | 340.7928372601586 | 319.9587034935923 | 305.8074950481041 | 389.2381861716923 | 287.6149564763117 | 269.0401531911619 | 377.1960427319178 | 209.56452556674006 | 221.61811122737538 | 411.9985226089731 | 269.2779949528843 |
| SE_205_3_3 | SE_205 | SE_205_3 | GATCTTCCTCAGTCTTTTAA | 6.5026610501297935 | 9.993924846338961 | 23.700644703229056 | 9.109159426964803 | 31.8178353818822 | 0.0 | 4.30464245105859 | 29.305958207241734 | 7.991867500426528 | 0.0 | 32.86597901154201 | 0.0 |
| SE_205_3_4 | SE_205 | SE_205_3 | GCAATGGTGAGCACACACCA | 26887.419665445006 | 22610.255572357266 | 22305.269246326447 | 25544.68565018844 | 23345.806414199702 | 23239.635007329933 | 26493.998125652855 | 24948.82396926828 | 23825.53298932713 | 22553.287852109577 | 25266.89515008762 | 23777.711226744774 |
| SE_205_3_5 | SE_205 | SE_205_3 | TTTCTTTTAACATCACAAAT | 1.083776841688299 | 2.9981774539016888 | 3.950107450538176 | 0.0 | 13.787728665482287 | 0.0 | 0.0 | 0.0 | 14.207764445202717 | 0.0 | 1.173784964697929 | 0.0 |
| SE_205_3_6 | SE_205 | SE_205_3 | CTTCTGTTAGCTGGGATCGG | 281.78197883895774 | 341.7922297447925 | 305.1458005540741 | 266.7682403611121 | 298.02705807696327 | 496.3956929545982 | 223.84140745504666 | 259.97220990295085 | 321.4506705727115 | 326.59511128244793 | 321.61708032723254 | 298.295020271083 |
| SE_206_1_17 | SE_206 | SE_206_1 | CTTTCATTAGTCTTACAAGG | 21.675536833765978 | 26.983597085115196 | 18.763010390056337 | 23.423552812195208 | 15.9089176909411 | 51.11229648223612 | 8.60928490211718 | 5.67212094333711 | 23.087617223454416 | 0.7290069448268927 | 3.521354894093787 | 62.67677468730927 |
| SE_206_1_22 | SE_206 | SE_206_1 | ATCAATGCCTTCCTGAGAGG | 225.42558307116616 | 293.8213904823655 | 288.35784388928687 | 242.04337905935049 | 216.36128059679896 | 386.3743090013103 | 212.00364071463557 | 364.90644735468743 | 129.64585056247478 | 314.20199322039076 | 388.52282331501453 | 268.1173139401563 |
| SE_206_1_32 | SE_206 | SE_206_1 | GTTTCACCGGAGCCTGTCTA | 8.670214733506391 | 15.990279754142339 | 33.5759133295745 | 24.724861301761607 | 29.69664635642339 | 4.331550549342044 | 29.056336544645482 | 36.868786131691216 | 10.655823333902038 | 37.17935418617152 | 8.216494752885502 | 0.0 |
| SE_206_2_11 | SE_206 | SE_206_2 | TTGACACTTGGAAGAAAGGA | 59.60772629285644 | 60.962941562667666 | 85.91483704920533 | 67.66804145745283 | 68.93864332741144 | 79.7005301078936 | 86.0928490211718 | 92.64464207450614 | 136.74973278507613 | 109.3510417240339 | 69.25331291717781 | 49.909283547301825 |
| SE_206_2_19 | SE_206 | SE_206_2 | GTAACTCCAGTTCTTCTGCG | 277.4468714722045 | 210.8718142577521 | 235.0313933070215 | 219.92113473672168 | 265.1486281823517 | 460.01066834012505 | 316.39122015280634 | 149.36585150787724 | 286.81924473752986 | 142.15635424124406 | 332.1811450095139 | 351.68634685656866 |
| SE_206_2_20 | SE_206 | SE_206_2 | TAGCCATGACAAATGACCAA | 1753.5509298516677 | 1777.9192301637013 | 1711.3840529456647 | 1850.4606721634213 | 1642.860900217851 | 1486.5881485341895 | 1673.4297528490267 | 1799.00769252842 | 1525.5587073036418 | 1210.8805353574687 | 1693.7717040591117 | 1651.649081111872 |
| SE_206_2_23 | SE_206 | SE_206_2 | GAAGAGGTGTCACTGACACG | 2774.468714722045 | 2190.6683263175005 | 2269.336730334182 | 2722.33736017291 | 2305.7324706737304 | 3055.4757575058775 | 2900.252851400725 | 2445.6294800688506 | 2326.5214279019447 | 2287.6237928667892 | 2402.7378227366607 | 2656.798838134276 |
| SE_206_2_5 | SE_206 | SE_206_2 | TTTGGGCAGAGTTTACCCAC | 356.5625809154503 | 308.8122777518739 | 303.170746828805 | 369.5716110368 5775 | 339.39024407341014 | 246.8983813124965 | 343.2952354719225 | 447.1522010330755 | 242.41998084627136 | 401.68282659961784 | 433.1266519735358 | 186.86964304919985 |
| SE_206_3_1 | SE_206 | SE_206_3 | GCTGCGAGTCAGCCCCTCCA | 0.0 | 4.996962423169481 | 2.962580587903632 | 2.602616979132801 | 0.0 | 0.8663101098684087 | 1.0761606127646475 | 0.945353490556185 | 1.7759705556503396 | 3.6450347241344634 | 0.0 | 0.0 |
| SE_206_3_15 | SE_206 | SE_206_3 | CAGGCCACATGATTTTACAA | 2863.3384157404857 | 3185.063848528227 | 2999.1190818211103 | 2930.5467185035336 | 3023.7549557915386 | 3376.876808267057 | 2803.398396251907 | 3102.6501560053994 | 3237.594322950569 | 3006.4246404661053 | 2882.8158732981137 | 2734.564465987049 |
| SE_206_3_16 | SE_206 | SE_206_3 | GAGAGGAGAAGCATGAGACA | 7284.064152987057 | 7289.568782919639 | 6925.525887656057 | 7256.096137822249 | 7168.5583115380605 | 7592.3418028867345 | 7553.57134099506 | 7595.915296618947 | 7063.0348998214 | 7989.9161153027435 | 8278.705356014494 | 7091.760987767771 |
| SE_206_3_20 | SE_206 | SE_206_3 | CTGGGTCTAGGAAAGACGGG | 146.30987362792035 | 176.89246978019963 | 194.5427919390051 8 | 156.15701874796804 | 191.9676068040226 | 75.36897955855156 | 122.6823098551698 | 94.5353490556185 | 157.17339417505505 | 56.13353475167074 | 24.64948425865651 | 147.40648861644956 |
| SE_206_3_26 | SE_206 | SE_206_3 | AGCTTTAACTCAGATCCCAG | 88.86970101844051 | 109.93317330972857 | 110.60300861506893 | 75.47589239485123 | 96.51410065837601 | 148.1390287874979 | 81.7882065701132 | 266.5896843368442 | 91.46248361599248 | 153.09145841364744 | 218.3240034338148 | 34.820430381838484 |
| SE_206_4_1 | SE_206 | SE_206_4 | CTATTCTTAACCCTGAACCA | 109.46146101051819 | 110.93256579436247 | 111.59053547770347 | 137.93869989403845 | 88.02934455654076 | 5.197860659210452 | 149.586325174286 | 77.51898622560716 | 108.33420389467071 | 153.09145841364744 | 77.46980767006332 | 5.803405063639747 |
| SE_206_4_22 | SE_206 | SE_206_4 | GTATCTCCCCAAGGAGTGTG | 72.61304839311603 | 80.95079125534559 | 58.264084895438096 | 58.55888203048802 | 96.51410065837601 | 2.598930329605226 | 91.47365208499504 | 70.90151179171387 | 58.607028336461205 | 71.44268059303548 | 69.25331291717781 | 37.14179240729438 |
| SE_206_4_28 | SE_206 | SE_206_4 | GCATTTTGTGGTTATGGTGG | 1126.0441385141426 | 1435.127607934275 | 1464.5023372870287 | 1228.435214150682 | 1533.619665406722 | 1279.5400322756398 | 1069.7036490880596 | 1387.7789241364796 | 1383.4810628516145 | 1162.0370700540668 | 1214.8674384623566 | 1548.3484709790846 |
| SE_206_4_32 | SE_206 | SE_206_4 | AGGAAGGCATTGATGCAGGG | 2214.156087569195 | 2585.4283577478895 | 2477.704898350071 | 2605.2195961119337 | 2734.2126538164107 | 2755.732459491408 | 2352.4870995035194 | 2684.8039131795654 | 2265.250443732008 | 2599.638765252699 | 2783.0441512987895 | 2547.694822937849 |
| SE_206_4_9 | SE_206 | SE_206_4 | GGTGCAGAAAGAAAGAAAGG | 455.18627350908554 | 583.6452110261954 | 613.2541816960519 | 514.0168533787282 | 636.356707637644 | 763.2192067940681 | 507.9478092249136 | 436.75331263695745 | 727.259942538814 | 419.90800022029015 | 422.56258729125443 | 973.8113696787495 |
| SE_207_1_16 | SE_207 | SE_207_1 | GTTCCTCTTCAAAACACTGA | 2.167553683376598 | 6.9957473924372735 | 0.0 | 0.0 | 2.121189025458813 5 | 0.8663101098684087 | 8.60928490211718 | 0.945353490556185 | 0.0 | 0.0 | 1.173784964697929 | 0.0 |
| SE_207_1_21 | SE_207 | SE_207_1 | TGTCGCCCTCTCACAGTCAA | 1778.4777972104985 | 1536.0662488822984 | 1620.5315815832867 | 1694.3036534154535 | 1735.1326228253095 | 1323.7218478789284 | 1770.284207997845 | 1800.8983995095325 | 1929.592008714094 | 1650.471723088085 | 1689.0765642003198 | 1469.422162113584 |
| SE_207_1_22 | SE_207 | SE_207_1 | TCTAGCCCTTGACTGTGAGA | 278.5306483138928 | 328.80012744455183 | 360.4473048616086 | 262.8643148924129 | 271.5121952587281 | 626.3422094348595 | 259.35470767628004 | 296.8409960346421 | 317.0107441835856 | 436.67515995130873 | 431.95286700883787 | 444.5408278748046 |
| SE_207_1_23 | SE_207 | SE_207_1 | CAGACCGAAGATCAAGAAGG | 1357.9723826354386 | 1573.0437708137526 | 1548.442120610965 | 1484.7929865952628 | 1777.5564033344856 | 1398.2245173276117 | 1170.8627466879364 | 1156.1673189502142 | 1330.2019461821044 | 1041.7509241576297 | 888.5552182763323 | 1149.0742026006699 |
| SE_207_1_37 | SE_207 | SE_207_1 | TGACCCGCAGAATCATCGGC | 1322.2077468597247 | 1645.999422192027 | 1639.2945919733431 | 1430.138030033474 | 1498.6200464866517 | 1923.2084439078674 | 1247.2701501942265 | 1809.406580924538 | 1801.7221287072696 | 1634.4335703018933 | 1881.5772984107803 | 1688.7908735191663 |
| SE_207_2_19 | SE_207 | SE_207_2 | CTAGCTCCATGACTGAGACC | 572.2341724114218 | 531.6768018252328 | 510.5513879820593 | 480.1828326500018 | 581.2057929757149 | 306.6737788934167 | 377.73237508039125 | 565.3213873525987 | 383.60964002047336 | 386.3736807582531 | 784.0883564182166 | 747.4785721967994 |
| SE_207_2_29 | SE_207 | SE_207_2 | TCAGGTTTCAGGCTGAGGGG | 842.0946059918083 | 1328.192612078448 | 1240.3337394689872 | 864.0688370720899 | 1158.1692079005122 | 1017.0480689855118 | 848.0145628585423 | 923.6103602733928 | 1027.3989664437215 | 914.9037157577503 | 883.8600784175405 | 762.5674253622627 |
| SE_207_2_30 | SE_207 | SE_207_2 | AGGCCCCCTTCTTCCCAGGA | 110.54523785220648 | 184.8876096572708 | 162.94193233469977 | 152.25309327926885 | 166.51333849851687 | 228.7058690052599 | 91.47365208499504 | 207.9777679223607 | 231.7641575123693 | 232.55321539977876 | 149.070690516637 | 298.29502027108 3 |
| SE_207_2_45 | SE_207 | SE_207_2 | ATCTCCAGCATGCAGCTCCA | 217.8391451793481 | 197.87971195751143 | 252.80687683444327 | 243.34468754891688 | 236.5125763386577 | 284.1497160368381 | 277.64943809327906 | 232.55695867682152 | 172.26914389808294 | 336.8012085100244 | 275.8394667040133 | 150.88853165463343 |
| SE_207_2_7 | SE_207 | SE_207_2 | GGGGAACCAAGAGCTCCAGG | 161.48274941155654 | 254.84508358164354 | 236.018920169656 | 170.47141213319847 | 213.17949705861076 | 182.79143318223424 | 143.1293614976981 | 230.66625169570915 | 205.12459917761421 | 163.29755564122397 | 248.84241251596094 | 422.4878886329736 |
| SE_207_3_0 | SE_207 | SE_207_3 | GTTCTTTTAAAACAAATGAA | 0.0 | 2.9981774539016888 | 0.987526862634544 | 0.0 | 1.0605945127294067 | 1.7326202197368175 | 0.0 | 0.0 | 6.215896944776189 | 0.0 | 0.0 | 5.803405063639747 |
| SE_207_3_13 | SE_207 | SE_207_3 | TGCATGATTTTCATGCCGAA | 144.14231994454374 | 185.8870021419047 | 227.1311784059451 2 | 171.77272062276487 | 220.6036586477166 | 160.2673703256556 | 147.4340039487567 | 224.04877726181584 | 218.44437834499178 | 86.02281948957334 | 104.46686185811568 | 55.71268861094157 |
| SE_207_3_22 | SE_207 | SE_207_3 | AGAGAGTTCAGCTTACACCC | 1787.1480119440048 | 1911.8378231046433 | 1878.2760927309027 | 1582.3911233127428 | 1679.9817081633803 | 1898.9517608315518 | 1855.3008964062521 | 1570.2321478138233 | 1866.5450539885069 | 1726.2884453500817 | 1403.84681777 8723 | 1662.095210226423 7 |
| SE_207_3_29 | SE_207 | SE_207_3 | TGTTCTGCGGACATCCCCAG | 292.6197472558407 | 285.8262506052943 | 327.8589183946686 | 327.92973937073293 | 326.6631099206573 | 323.13367098091646 | 256.1262258379861 | 394.21240556192913 | 285.0432741818795 | 295.9768195997184 | 262.9278320923361 | 165.97738482009677 |
| SE_207_3_7 | SE_207 | SE_207_3 | TGGATCAAACAACAAACGAG | 28.17819788389577 | 37.97691441608806 | 33.5759133295745 | 16.917010364363207 | 20.15129574185873 | 23.390372966447035 | 0.0 | 19.852423301679885 | 8.879852778251697 | 5.832055558615141 | 18.780559435166865 | 11.606810127279495 |
| SE_208_1_28 | SE_208 | SE_208_1 | GCCACGCACCCACCTCCCGG | 3373.7973081756745 | 4878.034717498047 | 4837.894100046631 | 3628.0480689111246 | 4221.166160663039 | 2532.2244511453587 | 2552.6529734777437 | 2911.6887509130497 | 3915.1270899311735 | 2725.0279597629246 | 2199.673023843919 | 1936.0159292302196 |
| SE_208_1_29 | SE_208 | SE_208_1 | GCCGCCGGGAGGTGGGTGCG | 1308.1186479177768 | 1531.069286459129 | 1549.4296474735995 | 1203.7103528489204 | 1358.62157080637 | 1892.0212799526046 | 1122.4355191135273 | 1399.1231660231538 | 1418.112488686796 | 1611.8343550122597 | 1820.540480246488 | 1990.5679368284332 |
| SE_208_1_30 | SE_208 | SE_208_1 | CGCACCCACCTCCCGGCGGC | 877.8592417675221 | 1324.1950421399124 | 1374.6373927872853 | 1314.3215744620645 | 1494.377668435734 | 1328.919708538139 | 832.9483142798372 | 1437.8826591359573 | 1272.4829031234683 | 1070.9112019507054 | 1351.0264943673162 | 1361.4788279298846 |
| SE_208_1_31 | SE_208 | SE_208_1 | ACCCACCTCCCGGCGGCTGG | 10.837768416882989 | 29.981774539016886 | 26.66322529113269 | 6.506542447832002 | 10.605945127294067 | 13.86096175789454 | 2.152321225529295 | 25.524544245016994 | 58.607028336461205 | 4.374041668961356 | 68.07952795247988 | 30.177706330926686 |
| SE_208_1_32 | SE_208 | SE_208_1 | CCCGCCAGCCGCCGGGAGGT | 149.56120415298525 | 211.871206742386 | 173.80472782367974 | 189.99103947669445 | 229.0884147495518 4 | 159.4010602157872 | 135.59623720834557 | 142.74837707398393 | 239.75602501279585 | 227.45016678599052 | 239.45213279837753 | 283.2061671056196 7 |
| SE_208_2_1 | SE_208 | SE_208_2 | TAAATGACACCACCACCTGT | 2291.104243329064 | 2137.700524631904 | 2156.758667993844 | 2247.3597614811733 | 1997.099467469473 | 2231.614843021021 | 2420.2852181076923 | 2503.296042992778 | 2393.1203237388327 | 2687.848605576753 | 2500.1619748065887 | 2879.6495925780428 |
| SE_208_2_16 | SE_208 | SE_208_2 | AGACAGTGAAGTCTGAACCG | 945.0534059521966 | 1215.2612613148178 | 1053.6911624310585 | 1000.7062284765619 | 1110.4424548276888 | 771.8823078927521 | 939.4882149435373 | 983.1676301784324 | 912.8488656042746 | 1283.0522228953312 | 1074.0132426986052 | 761.4067443495348 |
| SE_208_2_3 | SE_208 | SE_208_2 | GTTTAAGCAGATCCGACAGG | 147.39365046960864 | 208.8730292884843 | 240.95655448282875 | 183.48449702886245 | 269.3910062332693 | 389.8395494407839 | 150.66248578705066 | 252.4093819785014 | 283.26730362622914 | 151.63344452399366 | 287.5773163509926 | 286.68821014380353 |
| SE_208_2_4 | SE_208 | SE_208_2 | CTTCAGGCAGTTAAACCACA | 581.9881639866165 | 570.6531087259547 | 602.3913862070718 | 637.6411598875362 | 601.3570887175737 | 831.6577054736724 | 540.232627607853 | 686.3266341437903 | 600.2780478098148 | 667.0413545166068 | 826.344615147342 | 743.9965291586155 |
| SE_208_2_8 | SE_208 | SE_208_2 | TTAAGAAAGAAGGAAGACCA | 119.21545258571288 | 153.90644263362 | 156.02924429625796 | 156.15701874796804 | 183.48285070218736 | 173.26202197368175 | 112.99686434028798 | 178.67180971511897 | 183.81295250981015 | 59.04956253097831 | 165.503680022408 | 314.5445544492743 |
| SE_209_1_3 | SE_209 | SE_209_1 | AGCCTGCGGCGGCGGCCCGG | 4.335107366753196 | 9.993924846338961 | 7.900214901076352 | 5.205233958265602 | 16.969512203670508 | 0.0 | 2.152321225529295 | 0.945353490556185 | 24.863587779104755 | 0.7290069448268927 | 2.347569929395858 | 2.321362025455899 |
| SE_209_1_4 | SE_209 | SE_209_1 | CGGAGCCTGCGGCGGCGGCC | 74.78060207649263 | 72.95565137827442 | 110.60300861506893 | 93.69421124878083 | 68.93864332741144 | 196.65239494012877 | 48.42722757440914 | 131.40413518730972 | 153.62145306375436 | 134.86628479297514 | 46.95139858791716 | 100.9792481073316 |
| SE_209_1_5 | SE_209 | SE_209_1 | GCGCGCGGAGCCTGCGGCGG | 1198.6571869072586 | 1605.0243303220373 | 1641.269645698612 3 | 1506.9152309178917 | 1606.8006867850513 | 1586.2138111690563 | 1282.7834504154598 | 1685.5652736616778 | 1391.472930352041 | 1623.49846612949 | 1352.2002793320144 | 1902.356179861109 |
| SE_209_1_7 | SE_209 | SE_209_1 | CTGAGGGCGCGCGGAGCCTG | 3355.3731018669732 | 3026.1604434714377 | 2969.493275942074 | 3203.8215013124777 | 2961.179879540504 | 3300.641518598637 | 3582.5386798935115 | 3536.567408170688 | 2770.5140668145295 | 3573.5920435414278 | 3049.4933382852196 | 3660.7879141439525 |
| SE_209_2_17 | SE_209 | SE_209_2 | TTACAGGAAGTACAGTGTGA | 487.6995787597345 | 517.6853070403582 | 545.1148281742683 | 512.7155448891617 | 558.9333082083973 | 513.7218951519664 | 449.83513613562263 | 689.1626946154589 | 668.6529142023528 | 589.7666183649561 | 890.9027882057281 | 364.4538379965761 |
| SE_209_2_18 | SE_209 | SE_209_2 | ACTCTCTTGGAGGATGAACT | 22.759313675454276 | 56.96537162413208 | 44.43870881855448 | 33.834020728726415 | 71.05983235287025 | 2.598930329605226 | 37.665621446762664 | 58.61191641448347 | 59.49501361428638 | 12.393118062057175 | 70.42709788187574 | 2.321362025455899 |
| SE_209_2_19 | SE_209 | SE_209_2 | TCATCCTCCAAGAGAGTACA | 329.4681598732429 | 391.7618539764873 | 383.1604227022031 | 394.2964723386193 | 454.9950459609155 | 158.5347501059188 | 351.90452037403975 | 476.45815924031723 | 402.2573308548019 | 507.3888335995173 | 451.90721140870266 | 389.988820276591 |
| SE_209_2_20 | SE_209 | SE_209_2 | TCCTCCAAGAGAGTACACGG | 0.0 | 0.9993924846338962 | 0.0 | 0.0 | 0.0 | 0.0 | 0.0 | 0.0 | 0.0 | 0.0 | 0.0 | 0.0 |
| SE_209_2_21 | SE_209 | SE_209_2 | CCCTCCGTGTACTCTCTTGG | 133.30455152766078 | 130.9204154870404 | 122.45333096668347 | 137.93869989403845 | 125.15015250207 | 147.27271867762948 | 110.8445431147587 | 161.65544688510764 | 94.126439449468 | 48.114458358574915 | 246.49484258656508 | 217.04734938012655 |
| SE_209_3_11 | SE_209 | SE_209_3 | GCCACAGGAAATGAAAACAG | 1650.5921298912792 | 2174.6780465633583 | 2145.895872504864 | 1938.9496494539367 | 2458.458080506765 | 2616.2565318025945 | 1776.741171674433 | 2558.1265454450368 | 2369.144721237553 | 2109.7460983290275 | 2772.4800866165083 | 2884.2923166289543 |
| SE_209_3_18 | SE_209 | SE_209_3 | TCAGACTCTCCATTGCCTGG | 1101.1172711553118 | 1198.2715890760414 | 1275.8847065238308 | 1087.8938972775109 | 1078.6246194458067 | 1297.7325445828762 | 1011.5909759987686 | 978.4408627256515 | 1048.7106131115256 | 1122.6706950334146 | 809.911625641571 | 1500.7605494572385 |
| SE_209_3_22 | SE_209 | SE_209_3 | TCCTGCCTCAGCTTCCCAGT | 1140.1332374560905 | 980.4040274258522 | 1019.1277222388494 | 990.2957605600308 | 992.7164639147247 | 913.9571659111712 | 1063.2466854114716 | 891.4683415944825 | 960.8000706068337 | 1252.4339312126017 | 1039.9734787223651 | 450.3442329384444 |
| SE_209_3_28 | SE_209 | SE_209_3 | GTAATCCCAGCCTCCCCACT | 324.04927566480137 | 444.7296556620838 | 492.7759044546375 | 422.9252591090801 | 462.4192075500213 | 500.72724350394026 | 370.1992507910387 | 473.6220987686487 | 655.3331350349753 | 398.0377918754834 | 435.4742219029317 | 458.46900002754 |
| SE_209_3_34 | SE_209 | SE_209_3 | TGTCCTCTGGACCATCTCTG | 50.93751155935005 | 79.95139877071169 | 75.05204156022535 | 62.46280749918722 | 85.90815553108195 | 103.09090307434064 | 58.112673089290965 | 84.13646065950047 | 87.02255722686664 | 115.91210422747594 | 118.55228143449084 | 95.17584304369186 |
| SE_209_4_21 | SE_209 | SE_209_4 | CAGCAGAAGCAGCCTTTACG | 247.10111990493215 | 185.8870021419047 | 235.0313933070215 | 283.68525072547527 | 180.30106716399914 | 285.01602614670645 | 269.0401531911619 | 209.86847490347307 | 162.50130584200608 | 341.90425712381267 | 115.03092654039705 | 379.542691162039 5 |
| SE_209_4_27 | SE_209 | SE_209_4 | GACCCGGGTTTAAGAGACAA | 108.37768416882989 | 107.93438834046079 | 156.02924429625796 | 148.34916781056964 | 173.9375000876227 | 102.22459296447224 | 112.99686434028798 | 94.5353490556185 | 182.0369819541598 | 100.60295838611118 | 109.1620017169074 | 20.89225822910309 |
| SE_209_4_3 | SE_209 | SE_209_4 | CTGTGGCTCTTGCAACATAG | 376.07056406583973 | 353.78493956039927 | 356.4971974110704 | 347.4493667142289 | 403.0259148371746 | 365.5828663644685 | 541.3087882206177 | 568.1574478242671 | 335.6584350179142 | 817.9457920957735 | 564.5905680197038 | 132.31763545098624 |
| SE_209_4_5 | SE_209 | SE_209_4 | AACGGTTGCCCGTCGACAGA | 883.2781259759636 | 998.3930921492623 | 964.8137447939495 | 873.1779964990546 | 1062.7157017548654 | 1131.401003488142 | 887.8325055308342 | 1036.1074256495788 | 933.2725269942534 | 822.3198337647349 | 933.1590469348536 | 962.2045595514701 |
| SE_209_4_8 | SE_209 | SE_209_4 | GTCGACAGAGGGAAGATGAG | 1790.3993424690698 | 1933.8244577665891 | 1838.775018225521 | 1774.9847797685702 | 1912.2519064511205 | 1601.8073931466877 | 1668.0489497852036 | 1457.7350824376372 | 1756.434879538186 | 1667.9678897639303 | 1430.8438719667754 | 1530.9382557881654 |

| | | | | | | | | | | | | | | | |
|---|---|---|---|---|---|---|---|---|---|---|---|---|---|---|---|
| SE_209_5_0 | SE_209 | SE_209_5 | AAATGCATTGATCTTTAAAG | 37.93218945909046 | 51.968409200962604 | 46.41376254382357 | 19.519627343496005 | 88.02934455654076 | 61.50801780065702 | 40.8941032850566 | 86.02716764061283 | 59.49501361428638 | 24.78623612411435 | 50.47275348201095 | 31.338387343654635 |
| SE_209_5_1 | SE_209 | SE_209_5 | AACCATGTGGACCAGCCAGA | 420.50541457506 | 510.68955964792093 | 474.01289406458113 | 463.26582228563854 | 482.5705032418801 | 426.2245740552571 | 460.5967422632691 | 450.9336149953003 | 430.6725897420735 | 455.62934051680793 | 389.69660827971245 | 769.5315114386304 |
| SE_209_5_12 | SE_209 | SE_209_5 | GCTCTAACAGGGAACTCTGT | 134.38832836934907 | 226.86209401189444 | 212.31827546642697 | 174.37533760189766 | 269.3910062332693 | 90.09625142631451 | 142.05320088493346 | 221.21271679014728 | 139.41368861855165 | 471.66749330299956 | 183.11045449287693 | 92.85448101823596 |
| SE_209_5_16 | SE_209 | SE_209_5 | AGGAGACAACCTCACTGCAG | 1127.127915355831 | 308.8122777578739 | 538.2021401358265 | 926.5316445712771 | 242.87614341503414 | 2783.454383007197 | 1372.1047812749255 | 677.8184527287847 | 2160.4681809486383 | 99.87395144128429 | 372.08983380092435 | 827.565562075028 |
| SE_209_5_2 | SE_209 | SE_209_5 | TTAAGTTCAGTATAACCATG | 79.11570944324582 | 74.95443634754221 | 66.16429979651446 | 79.37981786355043 | 49.84794209828212 | 176.7272624131554 | 145.28168272322742 | 77.51898622560716 | 73.70277805948909 | 124.66018756539864 | 63.384388093688166 | 73.12290380186082 |
| SE_21_1_23 | SE_21 | SE_21_1 | GAGCTTGCTCGGGTCTCCCC | 171.23674098675122 | 285.8262506052943 | 281.445155850845 | 199.10019890365928 | 278.93635684783396 | 262.49196329012784 | 94.70213392328898 | 228.77554471459678 | 342.76231724051553 | 276.2936320893923 | 318.0957254331388 | 214.72598735467065 |
| SE_21_1_26 | SE_21 | SE_21_1 | CAAGCTCCGTGACAGCACGT | 5358.192705306458 | 4064.529235006056 | 4225.627445213114 | 4717.2432746782015 | 4366.467608906967 | 3432.3206552986353 | 5559.4457255421085 | 4619.942508348076 | 3958.6383685446067 | 4852.270224767797 | 4872.381388461104 | 3939.3513571986605 |
| SE_21_1_31 | SE_21 | SE_21_1 | AGCCCCACTCGGCGACATGG | 202.6662693957119 | 229.86027146579613 | 208.36816801588878 | 244.64599603848328 | 179.24047265126973 | 185.39036351183947 | 257.20238645075074 | 234.44766565793387 | 183.81295250981015 | 303.2668890479873 | 363.873339056358 | 169.4594278582806 |
| SE_21_1_34 | SE_21 | SE_21_1 | GGTCTGTTTCCAGCCCCACT | 0.0 | 0.0 | 0.987526862634544 | 0.0 | 0.0 | 0.8663101098684087 | 0.0 | 0.0 | 2.6639558334755096 | 0.0 | 0.0 | 0.0 |
| SE_21_1_38 | SE_21 | SE_21_1 | GCGCAAGGAGGGCTACCGCT | 7556.09214025082 | 7046.716409153602 | 7119.081152732428 | 7339.379881154498 | 7107.043829799754 | 6714.769661590036 | 7668.720526560878 | 7081.642997756382 | 6464.532822567236 | 6976.596461993363 | 6895.986667600333 | 7141.670271315073 |
| SE_21_2_1 | SE_21 | SE_21_2 | AAAGCTTGTACTTCTGGCAA | 52.02128840103835 | 81.9501837399795 | 63.20171920861082 | 68.96934994701923 | 51.96913112374093 | 48.51336615263089 | 88.2451702467011 | 32.14201867891029 | 57.71904305863604 | 97.68693060680361 | 26.997054188052367 | 29.017025318198737 |
| SE_21_2_18 | SE_21 | SE_21_2 | TCGAGGTCTTCCAAGAAGAG | 976.4829343611573 | 1098.332340612652 | 1078.379333996922 | 1013.719313372226 | 1057.4127291912184 | 783.1443393210415 | 958.8591059733009 | 988.8397511217695 | 935.9364828277289 | 1048.3119866610716 | 1097.4889419925637 | 718.4615468786008 |
| SE_21_2_20 | SE_21 | SE_21_2 | GTTCGAGGTCTTCCAAGAAG | 172.32051782843953 | 146.91069524118274 | 130.3535458677598 | 158.75963572710086 | 213.17949705861076 | 224.37431845591786 | 221.6890862295174 | 222.15807028070347 | 173.1571291759081 | 87.48083337922712 | 196.02208910455414 | 149.7278506419055 |
| SE_21_2_5 | SE_21 | SE_21_2 | GCAGAATCACTTGAATCGGG | 1684.1892119836166 | 1853.8730589958775 | 2009.6171654612972 | 1838.748895757324 | 1910.1307174256615 | 2269.732487855231 | 1654.0588618192633 | 1839.657892622336 | 2073.4456237217714 | 2431.9671679425137 | 2332.310724854785 | 2316.719301404987 |
| SE_21_2_7 | SE_21 | SE_21_2 | AATGGAGACGGAGTGTGTAG | 6012.793917686682 | 6132.272285713587 | 6258.94525537774 | 6120.053826430782 | 6302.052594638135 | 5940.288423367679 | 6530.142598255881 | 6242.16909814249 | 6477.852601734614 | 5695.002252987685 | 7259.860006656691 | 6374.460121901899 |
| SE_21_3_1 | SE_21 | SE_21_3 | GTAAAAGGTCAACTTTGTGT | 607.9988081871356 | 767.5334281988323 | 679.4184814925663 | 632.4359259292706 | 629.9931405126676 | 518.9197558111769 | 552.0703943482641 | 548.3050245225874 | 642.901341145423 | 846.3770629440223 | 605.6730417841313 | 754.4426582731671 |
| SE_21_3_15 | SE_21 | SE_21_3 | GTCTGGGAGAGTGAGAACGG | 301.2899619893471 | 370.7746117991755 | 309.0959080046123 | 301.9035695794049 | 370.147484942563 | 375.112277573021 | 307.7819352506892 | 277.93392622351837 | 429.78487446738217 | 191.72882648947277 | 358.0044142328683 | 251.86777976196504 |
| SE_21_3_17 | SE_21 | SE_21_3 | TGAGAGCCTTCTGGACCAGA | 2705.106996853994 | 2812.290451759784 | 2612.9960785310036 | 2650.765393246758 | 2749.0609769946223 | 2798.1816548749603 | 2623.6795739202107 | 2567.5800803505986 | 2537.861924024335 | 2633.1730847147364 | 2162.1119049735853 | 3201.1582331036843 |
| SE_21_3_3 | SE_21 | SE_21_3 | TGTCTGATGTAACTGCTATG | 342.47348197350243 | 322.80377253674845 | 278.48257526291414 | 329.2310478602993 | 364.8445123789159 | 338.7272529585478 | 331.4574687315114 | 239.1744331107148 | 369.40187557527065 | 261.71349131285446 | 183.11045449287693 | 279.7241240674358 |
| SE_21_3_6 | SE_21 | SE_21_3 | CTCTATCCCGACGAACGGCG | 45.51862735090855 | 62.96172653193546 | 45.42623568118903 | 63.76411598875362 | 58.33269820011737 | 48.51336615263089 | 25.827854706351538 | 64.28403735782058 | 39.07135222430747 | 48.84346530340181 | 52.820323411406804 | 19.73157721637514 |
| SE_21_4_14 | SE_21 | SE_21_4 | AGGGTCCTCTTATAAGTAGG | 58.52394945116814 | 88.94593113241676 | 95.79010567555078 | 54.65495656178882 | 120.90777445115236 | 31.187163955262715 | 77.48356411905462 | 87.9178746217252 | 81.69464555991561 | 72.17168753786237 | 120.8998513638867 | 25.534982280014887 |
| SE_21_4_15 | SE_21 | SE_21_4 | GCCACATGGCCCACTCCACA | 0.0 | 0.9993924846338962 | 0.0 | 0.0 | 0.0 | 0.0 | 0.0 | 0.0 | 0.0 | 0.0 | 0.0 | 0.0 |
| SE_21_4_16 | SE_21 | SE_21_4 | ATAAGAGGACCCTAAACTGA | 177.73940203688102 | 227.86148649652833 | 221.20601723013786 | 145.74655083143685 | 248.17911597868118 | 196.65239494012877 | 237.8314954209871 | 251.4640284879452 | 220.2203489006421 | 270.46157653077717 | 232.40942301018995 | 208.9225822910309 |
| SE_21_4_23 | SE_21 | SE_21_4 | ATGACAGCTGGCTTCCTCCA | 1847.8395150785495 | 2107.718750092887 | 2121.2077009390005 | 2135.447231378463 | 2165.7339949934485 | 2436.930339059834 | 1844.5392902786057 | 1936.083948659067 | 2078.7735353887224 | 1966.1317301981294 | 1890.9675781283636 | 2201.81188114492 |
| SE_21_4_6 | SE_21 | SE_21_4 | TAGCATTCAGAATATCCAAG | 383.6570019576578 | 504.69320474011755 | 525.3642909215774 | 501.00376848306416 | 544.0849850301856 | 835.122945913146 | 344.3713960846872 | 551.1410849942558 | 467.96824141386446 | 632.7780281097429 | 800.5213459239876 | 850.7791823295869 |
| SE_21_5_14 | SE_21 | SE_21_5 | CTCTGCAATCTCTATCCTTT | 32.51330525064897 | 29.981774539016886 | 37.52602081612674 | 42.943180155691216 | 46.66615856600939 | 18.192512307236584 | 8.60928490211718 | 60.50262339559584 | 67.48688111147129 | 19.683187510326103 | 28.170839152750297 | 83.56903291641235 |
| SE_21_5_15 | SE_21 | SE_21_5 | CAAAAGGATAGAGATTGCAG | 312.12773040623006 | 469.7144677779312 | 471.0503134766775 | 343.54544124552973 | 475.14634170277424 | 331.79677207960054 | 336.83827179533466 | 520.889773296458 | 377.39374307569716 | 417.72097938580947 | 664.3622900190278 | 357.48975192020845 |
| SE_21_5_17 | SE_21 | SE_21_5 | CCTTTCCTGCAGTTACCGAA | 321.8817219814248 | 73.95504386290831 | 158.00429802152703 | 225.12636869498726 | 66.81745430195262 | 499.8609333940718 | 435.84504816968223 | 248.62796801627767 | 549.6628897371 | 32.076305573282376 | 127.94256115207426 | 176.42351393464833 |
| SE_21_5_18 | SE_21 | SE_21_5 | TGCTGGAAAGAAACACCTTA | 687.1145176303816 | 802.5121651610186 | 800.8842855966152 | 821.1256569163987 | 908.9294974091016 | 967.6683927230125 | 636.0109221439067 | 758.1734994260604 | 737.0277805948909 | 868.2472712888291 | 787.6097113123104 | 1046.9342734806105 |
| SE_21_5_7 | SE_21 | SE_21_5 | CAGCAGGGACTGCTCCCCAG | 125.71811363584267 | 175.89307729556572 | 158.9918248841616 | 140.54131687317124 | 196.20998485494025 | 133.41175691973496 | 174.33801926787288 | 208.92312141291688 | 214.00445195586593 | 111.53806255851458 | 400.2606729619938 | 192.6730481128396 |
| SE_21_6_0 | SE_21 | SE_21_6 | TTATGTTAATGCTATATCAG | 28.17819788389577 | 39.975699385355846 | 57.27655803280356 | 59.86019052005442 | 23.33307928004695 | 9.529411208552496 | 59.18883370205561 | 18.9070698111237 | 61.27098416993672 | 134.13727848814824 | 64.5581730583861 | 38.30247342002233 |
| SE_21_6_11 | SE_21 | SE_21_6 | ATACACTTGTATGTTCTGAG | 43.351073667531956 | 60.962941562667666 | 79.98967587339807 | 24.724861301761607 | 63.6356707637644 | 58.04277736118338 | 51.655709412703075 | 100.20746999895562 | 66.59889583688773 | 75.81672226199683 | 156.11340030482455 | 33.659749369110536 |
| SE_21_6_3 | SE_21 | SE_21_6 | TTAGGATTCAAAAACAGAAT | 49.85373471766175 | 62.96172653193546 | 84.92731018657079 | 48.14841411395682 | 88.02934455654076 | 97.89304241513018 | 44.12258512335055 | 77.51898622560716 | 55.94307250298569 | 53.21750697236316 | 57.515463270198524 | 91.69380000550801 |
| SE_21_6_6 | SE_21 | SE_21_6 | TTAAGTTTGTCTCATTTACT | 46.60240419259685 | 33.97934447755247 | 35.55096705484358 | 26.02616979132801 | 22.272484767317543 | 54.57753692170975 | 40.8941032850566 | 5.67212094333711 | 23.975602501279585 | 9.477090282749604 | 68.07952795247988 | 24.37430126728694 |
| SE_21_6_8 | SE_21 | SE_21_6 | TCTCATTTACTGGGTGCAGT | 322.9654988231131 | 389.7630690072195 | 368.3475197626849 | 450.2527373899745 | 299.0876525896927 | 316.2031901019692 | 365.89460833998015 | 315.7480658457658 | 417.3530805778298 | 607.2627850408015 | 312.2268006096491 | 104.46129114551545 |
| SE_21_7_1 | SE_21 | SE_21_7 | TCTTTAAGTGAAGTAAACCA | 86.70214733506391 | 109.93317330972857 | 98.75268626345441 | 84.58505182181602 | 146.36204275665813 | 175.86095230328698 | 61.34115492575849 | 174.89039575289422 | 146.5175708411153 | 150.17543063433988 | 369.74226387984766 | 92.85448101823596 |
| SE_21_7_10 | SE_21 | SE_21_7 | GGTGCAGTCAAGAATAAAGA | 355.47880407376203 | 541.6707266715717 | 560.915257976421 | 448.95142890040813 | 554.6909301574798 | 717.3047709710424 | 432.6165663313883 | 544.5236105603626 | 404.0333014104522 | 753.0641740061801 | 590.4138372430583 | 616.3216177585412 |
| SE_21_7_13 | SE_21 | SE_21_7 | TGGTGGCTCAGCACTCTGCG | 33.59708209233727 | 17.98906472341013 | 24.688171565863602 | 27.32747828089441 | 30.757240869152795 | 6.064170769078861 | 38.74178205952731 | 22.68848377334844 | 21.311646667804077 | 3.6450347241344634 | 22.30191432926065 | 9.285448101823595 |
| SE_21_7_5 | SE_21 | SE_21_7 | GGTCAGAGTTCAGTCTCATG | 409.667646158177 | 416.7466660923347 | 444.3870881855448 | 447.65012041084174 | 418.9348325281157 | 274.62030482828555 | 486.42459696962067 | 190.01605160179318 | 502.5996672490461 | 642.2551183924925 | 193.67451917515828 | 434.0946987602531 |
| SE_21_7_8 | SE_21 | SE_21_7 | ACATAAAGTGGGAAGAATGG | 2954.3756704423026 | 3382.9435604857385 | 3428.6932670671367 | 3060.677567460174 | 3468.14405662516 | 3207.9463368427173 | 2846.444820762493 | 3386.2562031722546 | 3382.3359232360717 | 3271.0541614382673 | 3608.214981481434 | 3550.5232179347972 |
| SE_210_1_13 | SE_210 | SE_210_1 | TCAGTGATAGGACCTACTGG | 716.3764923559655 | 857.4787518158829 | 839.3978332393624 | 720.9249032197858 | 863.3239333617371 | 699.9785687736743 | 608.0307462120259 | 873.5066252739149 | 768.1072653187719 | 863.1442226750409 | 984.8055853815624 | 1244.2500456443618 |
| SE_210_1_39 | SE_210 | SE_210_1 | CCAGCCCTCCGTTATCCGTG | 1093.5308332634936 | 1149.3013573289807 | 1082.3294414474603 | 1173.7802575888932 | 1123.1695889804416 | 1155.6576865644572 | 1123.511679726292 | 1141.0416631013154 | 1024.735010610246 | 1102.9875075230887 | 777.0456466300291 | 1021.3992912005955 |
| SE_210_1_41 | SE_210 | SE_210_1 | GTTCAGCCTCTCCTTTTCAG | 60.69150313454474 | 94.94228640022014 | 92.82752508764715 | 97.59813671408003 | 101.81707322202305 | 25.989303296052263 | 71.02660044682216 | 47.26767452780925 | 68.37486639253017 | 131.22125006884067 | 55.167893340802664 | 68.48017975094902 |
| SE_210_1_42 | SE_210 | SE_210_1 | GAAATAAAGCTGTTGCCGCA | 294.7873009392173 | 321.8043800521146 | 317.9836497683232 | 344.8467497350961 | 432.7225611935979 | 429.68981449473074 | 394.95094488462564 | 486.85704763643525 | 509.7035494716475 | 831.7969204474845 | 454.25478133809855 | 470.0758101548195 |
| SE_210_1_45 | SE_210 | SE_210_1 | ACGAGCACCTCGTTCCCCGA | 13365.136011700102 | 12765.240206228757 | 12384.574384299816 | 13201.774626651133 | 12702.740478960104 | 12849.977859678107 | 13840.501640766131 | 1390.73646222466 | 13594.166618225525 | 13284.693555580465 | 13050.141237511574 | 12437.857732392706 |
| SE_210_2_14 | SE_210 | SE_210_2 | GGGCTTTGCCAGTCTCTCCG | 15.172875783636185 | 34.97873696218637 | 33.5759133295745 | 19.519627343496005 | 36.06021343279983 | 13.86096175789454 | 15.066248578705064 | 2.836060471668555 | 15.983735000853056 | 21.141201399979888 | 0.0 | 4.642724050911798 |
| SE_210_2_17 | SE_210 | SE_210_2 | TATTTAGTAGCTGAGCCCCT | 108.37768416882989 | 135.9173779102099 | 143.19139508200888 | 122.32299801924164 | 113.48361286204653 | 164.59892087499765 | 103.31141882540615 | 159.76473990399526 | 209.56452556674006 | 116.64111117230283 | 102.11929192871983 | 226.33279748195014 |
| SE_210_2_18 | SE_210 | SE_210_2 | TCAAGTCTGTGCTTAGGAGA | 1084.8606185299873 | 1171.2879919909262 | 1088.2546026232676 | 1193.2998493323892 | 1237.7167936552177 | 751.0908652559103 | 1048.1804368327666 | 884.8508671605891 | 1431.4322678541737 | 1016.9646880335152 | 1089.272447239678 | 1399.781301349907 |
| SE_210_2_31 | SE_210 | SE_210_2 | CCTCATCTCCACACAGAGGT | 836.6757217833667 | 859.4775367851507 | 988.5143894971786 | 869.2740710303555 | 1182.5628816932885 | 912.2245456914344 | 849.0907234713069 | 1130.6427747051973 | 1153.4928758948956 | 1063.6211325024365 | 1194.9130940624918 | 999.3463519587644 |
| SE_210_2_8 | SE_210 | SE_210_2 | ACAGGCCTGATGCAGTGCAG | 5302.920086380846 | 4490.27043460946 | 4615.700555953858 | 5440.77079487712 | 4642.222182216613 | 4743.914161934406 | 5671.366429269692 | 4359.970298445125 | 4977.157482100765 | 5235.7278777467743 | 4278.446196323951 | 5123.245990181169 |
| SE_210_3_15 | SE_210 | SE_210_3 | GCTGCTGACCTGTAGGAAAC | 1527.0415699388132 | 1552.0565286364408 | 1799.2739437201392 | 1531.6400922196533 | 1766.9504582071916 | 1560.224507873004 | 1265.5648806112254 | 1377.3800357403616 | 1855.0012453767797 | 967.3922157852866 | 1544.7010135424746 | 1882.624602644734 |
| SE_210_3_3 | SE_210 | SE_210_3 | GAGCGCTCGGAGCCAAGGGA | 1389.4019110443992 | 1766.9259128327265 | 1710.3965260830303 | 1426.2341064775 | 1811.4954271418267 | 1436.3421621618218 | 1454.9691484578034 | 1620.3358828133012 | 1593.9335736961798 | 1202.132452019546 | 1777.1104365526646 | 1147.913521587942 |
| SE_210_3_31 | SE_210 | SE_210_3 | GGGGTCCCTGCTAAATGCAG | 9.75399157519469 | 5.9963549078033775 | 8.887741763710896 | 5.205233958265602 | 12.727134152752882 | 0.0 | 5.380803063823238 | 0.0 | 11.543808611727208 | 3.6450347241344634 | 0.0 | 0.0 |
| SE_210_3_36 | SE_210 | SE_210_3 | GCAGCTGTCAGTCGTGGGGG | 3694.595253315411 | 3734.72971507687 | 4050.8351905268996 | 3854.475746095678 | 4075.86471241911 | 4024.0104603387585 | 3804.2277661230287 | 3714.2938643952507 | 3870.7278260399153 | 3553.9088560311015 | 3792.499220939009 | 4301.48383316978 |
| SE_210_4_1 | SE_210 | SE_210_4 | AGACGCTCCTCGCACCCCGG | 1018.750231187001 | 1111.3244429128927 | 1056.6537430189621 | 1099.6056736836083 | 1266.3498481989116 | 1356.641632053928 | 990.0677637434757 | 1318.7827893258782 | 1122.4133911710146 | 1303.4644173504842 | 1245.3858475445027 | 1147.913521587942 |
| SE_210_4_21 | SE_210 | SE_210_4 | CCAACCACTGCGTAAATCAG | 904.9536628097296 | 923.4386558017201 | 1034.9281520410022 | 942.1473464460739 | 1000.1406255038305 | 900.9625142631451 | 867.3854538883058 | 957.6430859334154 | 804.5146617096038 | 806.2816809785432 | 703.0971938540595 | 1094.5221950024563 |
| SE_210_4_25 | SE_210 | SE_210_4 | GAGCAGCCAGAGGTTGGCGG | 1626.7490393741366 | 1753.9338105324878 | 1794.3363094069666 | 1756.7664609146407 | 1622.7096044759924 | 1964.791329181551 | 1861.7578600828401 | 1762.1389063967288 | 1702.2677775908505 | 1869.1738065361528 | 1515.3563894250265 | 1925.5698001156682 |
| SE_210_4_3 | SE_210 | SE_210_4 | GATACCCTAGAGAAAAGCTA | 35.76463577571386 | 48.970231747060915 | 51.35139685699629 | 37.737946197425615 | 34.999618920070425 | 38.98395494407839 | 16.142409191469714 | 32.14201867891029 | 58.607028336461205 | 61.965590310285876 | 24.64948425865651 | 30.177706330926686 |
| SE_210_4_6 | SE_210 | SE_210_4 | TTATGAGCCAATCACAGGCA | 734.8006986646667 | 740.5498311137171 | 772.2460065802135 | 715.7196692615203 | 621.5083844594324 | 563.1015714144656 | 823.2628687649553 | 588.009871125947 | 973.231864496386 | 916.3617296474041 | 529.377019078766 | 914.6166380296241 |
| SE_210_5_28 | SE_210 | SE_210_5 | CCTGGGCCTGGCCACCGACA | 111.62901469389479 | 148.90948021045054 | 128.37849214249073 | 127.52823197750725 | 147.42263726938754 | 108.2887637335511 | 95.77829453605362 | 98.31676301784324 | 227.32423112324346 | 109.3510417240339 | 73.94845277596953 | 118.38946329285085 |
| SE_210_5_3 | SE_210 | SE_210_5 | CATCCAATACCACTTGACAA | 1268.0189047753097 | 1343.1834993479565 | 1444.751800034338 | 1383.2909244090836 | 1205.8959609733354 | 1838.3100531407633 | 1302.1543414452235 | 1309.3145844203161 | 1095.7738328362595 | 1102.2585005782616 | 1268.8615468384612 | 1710.8438127609975 |
| SE_210_5_47 | SE_210 | SE_210_5 | GGGAAGTGAGTTGGGCGTGG | 1365.5588205272566 | 1473.104522350363 | 1485.2404014023543 | 1418.4262536273766 | 1329.985518962676 | 1718.759257978923 | 1414.0750451724767 | 1420.866296305946 | 1328.425975626454 | 1442.7047438124205 | 1321.681870249868 | 981.9361367678453 |
| SE_210_5_5 | SE_210 | SE_210_5 | GAAGGGTGCACGTGACTCCG | 13905.940655702563 | 11832.80701806533 | 11652.81697908762 | 13014.38620415357 | 12005.929884096884 | 11831.197170472858 | 13950.270023268125 | 11540.875412709907 | 11736.50141701527 | 11975.397082671365 | 11136.87174505395 | 11616.095575381318 |
| SE_210_5_7 | SE_210 | SE_210_5 | TTCTGGGCTGAGGACTGGCA | 34.680858930425565 | 20.98724217731182 | 35.55096705484358 | 29.93009526002721 | 15.9089176909451 | 57.176467251314975 | 31.208657770174778 | 17.961716320567515 | 36.40739630831196 | 37.17935418127875 | 0.0 | 103.3006101327875 |
| SE_210_6_14 | SE_210 | SE_210_6 | AGAATTTTCTGGGCGTGCAG | 20.59175999207768 | 32.97995199291857 | 19.75053725269088 | 23.423552812195208 | 43.48437502190568 | 23.390372966447035 | 24.75169409358689 | 31.196665188354107 | 91.46248361599248 | 18.954180565499207 | 4.695139858791716 | 45.26655949639003 |
| SE_210_6_15 | SE_210 | SE_210_6 | GGTGCCTCTGAAGGACGCAC | 94.288585226882 | 142.91312530264716 | 149.11655625781614 | 106.70729614444484 | 128.33193604025823 | 83.16577054736723 | 174.33801926787288 | 185.28928414901227 | 131.42182111812514 | 172.04563897914667 | 129.1163461167722 | 121.87150633643469 |
| SE_210_6_21 | SE_210 | SE_210_6 | TGTCCAAGAAGGCCACAGAG | 134.38832836934907 | 163.900367479959 | 153.06666370835433 | 110.61122161314404 | 124.08955798934059 | 256.42779252104896 | 144.20552211046277 | 94.5353490556185 | 196.24474639936253 | 196.831875103261 | 82.16494752885504 | 414.3631215438779 |
| SE_210_6_37 | SE_210 | SE_210_6 | CCTCTGCTGCTCCTGCCCAG | 973.2316038360924 | 903.4508061090421 | 972.7139596950259 | 1007.2127709243939 | 984.2317078128895 | 1128.8020731585366 | 901.8225934967746 | 1161.8394398935513 | 1059.3664364454276 | 972.4952643990748 | 934.3328318995515 | 1087.5581089260886 |
| SE_210_6_6 | SE_210 | SE_210_6 | CCACCACCAACACCACAACG | 1745.9644919598495 | 1510.0820442818172 | 1438.8266388585307 | 1803.6135665390311 | 1442.4085371311993 | 1718.759257978923 | 1744.4563532919435 | 1823.5868832828808 | 1754.6589098925354 | 1739.4105703569658 | 1598.6951219185794 | 2098.5112710121325 |
| SE_210_7_2 | SE_210 | SE_210_7 | ACGAGAAGAAATCTCCCCCT | 15014.644364749693 | 16067.232975459148 | 15532.810022378742 | 15740.627489795179 | 16218.611288658089 | 16317.817229481347 | 15077.01018483271 | 16017.124190493443 | 15767.95457834154 | 15684.584417950595 | 16849.68316823877 | 16746.305651638853 |
| SE_210_7_24 | SE_210 | SE_210_7 | GCGTTGCCGCAACCCCGCGT | 1111.9550395721947 | 1035.3706140807165 | 932.2253583270095 | 1047.5533341009523 | 956.6562504819249 | 1131.401003488142 | 1262.3363987729315 | 1059.7412629134833 | 1008.7512756093929 | 1084.0333269575895 | 1213.6936534976587 | 991.2215848696688 |
| SE_210_7_25 | SE_210 | SE_210_7 | CCTTAGCTACTGGGTGAGGA | 1158.5574437647915 | 1269.2284554850482 | 1271.9345990732927 | 1085.2912802987355 | 1221.8048786642767 | 1496.9838698526103 | 1099.8361462456498 | 952.9163184806345 | 1117.0854795040636 | 954.2700907784025 | 873.2960137352592 | 1246.5714076698177 |
| SE_210_7_54 | SE_210 | SE_210_7 | GGCTCGCCTCTCCAAACCCG | 432.42695983363126 | 528.6786243713311 | 501.6636462183484 | 447.65012041084174 | 488.9340703682565 | 511.98927493222953 | 368.04692956550946 | 454.71502895752496 | 519.4713875277243 | 523.4269863857089 | 225.36671322200237 | 854.2612253677707 |
| SE_210_7_9 | SE_210 | SE_210_7 | ATTTGATTACACTTGCAACA | 48.769957875374154 | 49.96962423169481 | 54.313977444899992 | 50.751031093089615 | 56.21150917465856 | 50.24596873234677 | 41.970263897821255 | 97.37140952728706 | 48.83919028038434 | 11.66411117230282 | 46.95139858791716 | 46.42724050911798 |
| SE_211_1_13 | SE_211 | SE_211_1 | GATGGCTCCTGATAATTGAC | 909.2887701764828 | 941.4277205251302 | 1010.2399804751385 | 873.1779964990546 | 889.8387961799723 | 866.3101098684087 | 864.156972050012 | 873.5066252739149 | 878.217439769093 | 535.0910975029392 | 862.7319490529778 | 836.8510101768516 |
| SE_211_1_20 | SE_211 | SE_211_1 | TTAAAGGGCTCTCACACTTG | 140.89098941947887 | 191.88335704970808 | 201.45547997744697 | 171.77272062276487 | 183.48334570218736 | 165.46523098486605 | 109.76830250199404 | 182.45322367734371 | 111.88614500597139 | 239.11477790322208 | 157.2871852695225 | 268.1173139401563 |
| SE_211_1_31 | SE_211 | SE_211_1 | GGAGGAACTGGTTCCTCAAT | 612.3339155538889 | 738.5510461444493 | 709.0442873716026 | 766.4770035546099 | 680.9016771722792 | 1212.8341538157722 | 603.7261037609672 | 601.2448199937337 | 911.9608803264493 | 664.8543336821261 | 512.944029572995 | 1037.6488253787868 |
| SE_211_1_4 | SE_211 | SE_211_1 | CCACTTTACTGCGCTCTCCA | 215.67159149597148 | 212.87059922701988 | 207.38064115325426 | 275.8773997880769 | 263.02743915689285 | 216.5775274671022 | 189.40426784657797 | 270.37109829906893 | 263.7316275140754 | 305.453909882468 | 320.4432953625346 | 174.10215190919243 |
| SE_211_1_7 | SE_211 | SE_211_1 | TCTATGACCAGGAACTGCCT | 341.38970513181414 | 327.8007459991 8 | 394.02321819118305 | 350.0519836933617 | 349.99619920070424 | 434.02136504407275 | 243.21229848481033 | 306.29453094020397 | 446.6556947460604 | 335.3431946207064 | 137.3328408696577 | 386.50677723840715 |
| SE_211_2_19 | SE_211 | SE_211_2 | TAGCCAGCTGGTCACTCAGG | 2745.2067399964612 | 2546.4520508471674 | 2695.9483349923053 | 2752.267455432937 | 2918.7560990313273 | 3065.00516871443 | 2639.8219831116803 | 2768.940373839066 | 2860.200579874872 | 2977.2643626730296 | 2462.600855936255 | 2560.4623140778563 |
| SE_211_2_4 | SE_211 | SE_211_2 | GGTCCTGTGTGACCGCGCCC | 391.2434398494759 | 443.7302631774499 | 374.2726809384922 | 402.10432327601774 | 340.4508385861396 | 572.6309826230182 | 307.7819352506892 | 326.1469542418838 | 293.9231269601312 | 357.2134029651774 | 397.9131030325979 | 335.4368126783774 |

| | | | | | | | | | | | | | | | |
|---|---|---|---|---|---|---|---|---|---|---|---|---|---|---|---|
| SE_211_2_6 | SE_211 | SE_211_2 | GTCAAATTATGAAAGAAGAA | 35.76463577571386 | 35.97812944682026 | 52.338923719630834 | 55.95626505135522 | 51.96913112374093 | 34.65240439473635 | 39.817942672291956 | 30.25131169779792 | 55.94307250298569 | 32.076305572383276 | 19.954344399864794 | 61.51609367458132 |
| SE_211_2_8 | SE_211 | SE_211_2 | TCATTGGCTACATTTTGCTG | 2066.762437099586 | 2175.677439047992 | 2384.877373262424 | 2164.076018148924 | 2079.8258394623667 | 2578.138886968384 | 2208.2815773930565 | 2257.50413544817 | 2459.71921957572 | 2110.4751052738543 | 2571.7628576531624 | 3161.6950786709344 |
| SE_211_2_9 | SE_211 | SE_211_2 | TTTTCCGGCCACGGTCAGGG | 54.188842084414944 | 70.95686640900662 | 65.1767729338799 | 40.34056317655841 | 58.33269820011737 | 2.598930329605226 | 30.13249715741013 | 33.08737216946648 | 86.13457194904147 | 38.63736807582531 | 113.85714157569912 | 6.964086076367696 |
| SE_211_3_19 | SE_211 | SE_211_3 | GAGAGCACCAACTATCCTAA | 2389.727935922699 | 2557.44536817814 | 2459.929414822649 | 2546.660714081446 | 2895.42301975128 | 2260.2030766466783 | 2702.23929865203 | 2891.83632761137 | 2520.102218467832 | 2601.096779142353 | 3271.3386966131284 | 2590.640020408783 |
| SE_211_3_23 | SE_211 | SE_211_3 | TAGACTGTGGTTCATCCACG | 1531.3766773055663 | 1510.0820442818172 | 1383.5251345509962 | 1465.2733592517668 | 1476.3475617193342 | 1579.283330290109 | 1403.3134390451003 | 1540.9261896065816 | 1453.631899799803 | 1006.0295838611119 | 1709.0309086001846 | 1771.1992254228508 |
| SE_211_3_24 | SE_211 | SE_211_3 | GCAAAAATCTCCACAGCCCA | 1946.4632076721848 | 2441.5158399606084 | 2303.9001705263913 | 2145.8576992949943 | 2349.216845695636 | 2178.769926319048 | 2010.2680246443615 | 2430.5038242199516 | 2401.112191239259 | 2130.1582927841805 | 2689.1413541229554 | 2263.3279748195014 |
| SE_211_3_26 | SE_211 | SE_211_3 | AAAAATCTCCACAGCCCACG | 17.340429467012783 | 19.987849692677923 | 29.62580587903632 | 31.23140374959361 | 25.454268305505764 | 12.994651648026132 | 24.75169409358689 | 10.398888396118036 | 28.415528890405433 | 8.019076393095819 | 0.0 | 11.606810127279495 |
| SE_211_3_36 | SE_211 | SE_211_3 | CATTCCAAAGGAATAAGGGA | 69.36171786805113 | 112.93135076363026 | 103.69032057662713 | 52.05233958265602 | 111.36242383658771 | 71.03742900920952 | 38.74178205952731 | 35.92343264113503 | 78.14270444861494 | 22.599215289633673 | 30.518409082146157 | 124.19286836189059 |
| SE_211_4_12 | SE_211 | SE_211_4 | ATTATCATCTACTTAAGAAG | 3.2513305250648967 | 0.9993924846338962 | 7.900214901076352 | 2.602616979132801 | 8.484756101835254 | 0.0 | 5.380803063823238 | 0.945353490556185 | 39.95933750213264 | 0.0 | 0.0 | 0.0 |
| SE_211_4_16 | SE_211 | SE_211_4 | TCTGTTTCTTCCCAATAGAA | 10.837768416882989 | 16.989672238776237 | 31.60085960430541 | 32.53271223916001 | 13.787728665482287 | 17.326202197368175 | 12.913927353175769 | 8.508181415005666 | 49.72717555820951 | 14.580138896537854 | 11.73784964697929 | 2.321362025455899 |
| SE_211_4_4 | SE_211 | SE_211_4 | TGTATGGCTCAAGGCTAAAG | 526.7155450605133 | 586.6433884800971 | 569.8029997401319 | 602.5058306692434 | 566.3574697975032 | 564.8341916342025 | 498.26236371003176 | 529.3979547114636 | 575.4144600307101 | 629.8620003304353 | 515.2915995023908 | 280.88480508016374 |
| SE_211_4_5 | SE_211 | SE_211_4 | ACCCCGACTTTGCCAAACAA | 226.50935991285448 | 254.84508358164354 | 249.84429624653964 | 197.79889041409285 | 285.2999239242104 | 265.9572037296015 | 184.02346478275473 | 145.5844375456525 | 205.12459917761421 | 470.2094794133458 | 138.50662583435562 | 105.6219721582434 |
| SE_211_4_6 | SE_211 | SE_211_4 | ACCATTGTTTGGCAAAGTCG | 1343.8832836934907 | 1579.040125721556 | 1568.192657863656 | 1224.531288681983 | 1653.4668453451452 | 1570.620229191425 | 1272.0218442878133 | 1366.0357938536874 | 1674.7402339782702 | 1401.8803549021145 | 1530.6155939660994 | 1705.0404076973578 |
| SE_211_5_13 | SE_211 | SE_211_5 | CACTTCATCCACCAACAGAA | 518.0453303270069 | 515.6865220710904 | 517.4640760205011 | 560.8639590031186 | 485.7522868300683 | 604.6844566881493 | 499.33852432279645 | 423.5183637691709 | 644.6773117010732 | 567.8964100201493 | 616.2371064664128 | 562.9302911730555 |
| SE_211_5_14 | SE_211 | SE_211_5 | GAGTTCTGCGGAACGAACAG | 89.95347786012881 | 147.91008772581662 | 138.25376076883617 | 106.70729614444484 | 147.42263726938754 | 313.60425977236395 | 69.95043982970209 | 145.5844375456525 | 111.88614500597139 | 75.81672226199683 | 77.46980767006332 | 210.08326330375886 |
| SE_211_5_16 | SE_211 | SE_211_5 | AGAAAAACTCCACAAACCAA | 89.95347786012881 | 97.94046349412183 | 84.92731018657079 | 71.57196692615203 | 59.39329271284678 | 58.90908747105179 | 102.23525821264151 | 48.213028018365435 | 125.20592417334893 | 99.87395144128429 | 39.90868879972959 | 85.89039494186825 |
| SE_211_5_3 | SE_211 | SE_211_5 | GTAGCAATATGTTTTAACAA | 207.0013767624651 | 321.8043800521146 | 308.10838114197776 | 243.34468754891688 | 304.3906251533397 | 273.7539947184172 | 215.2321225529295 | 208.92312141291688 | 285.0432741818795 | 607.2627850408015 | 197.19587406925208 | 174.10215190919243 |
| SE_211_5_7 | SE_211 | SE_211_5 | GCAGGGTTGGAGAGATAGTG | 6789.861913177193 | 6803.864035387565 | 6923.550833930788 | 6885.223218295825 | 7091.134912108813 | 6785.807090599245 | 7061.765940961617 | 6850.031392570117 | 6706.952803413507 | 7291.52746215858 | 6863.120688588791 | 6642.577435842055 |
| SE_212_1_10 | SE_212 | SE_212_1 | TCACTTGGAGACGTCCTGCG | 1659.2623446247856 | 1777.9192301637013 | 1773.598245291641 | 1527.7361667509542 | 1572.8616623777102 | 1418.149649854585 | 1539.9858368662105 | 1283.7900401752993 | 1806.1620550963953 | 1466.761972991708 | 1643.2989505771006 | 1031.845420315147 |
| SE_212_1_12 | SE_212 | SE_212_1 | GCAGAGGCTGAGTGAGGTGT | 16174.285585356172 | 14450.215935321505 | 14347.77778721729 | 15407.492516466182 | 14449.539641425437 | 13096.876240990603 | 16413.601665886403 | 13943.96398570373 | 15737.763078895485 | 13447.262104276862 | 13031.360678076408 | 12015.369843759732 |
| SE_212_1_17 | SE_212 | SE_212_1 | GGGATCTCCCGCCCTCTGTG | 65.02661050129794 | 72.95565137827442 | 102.70279371399258 | 98.89944520704643 | 84.84756101835254 | 64.97325824013066 | 99.00677637434757 | 94.5353490556185 | 125.20592417334893 | 76.54572920682372 | 49.29896851731302 | 121.87150633643469 |
| SE_212_1_28 | SE_212 | SE_212_1 | GGGCTGGAGGCCCTGCTCAG | 59.60772629285644 | 67.95868895510495 | 77.02709528549444 | 74.17458390528482 | 134.69550311663465 | 46.78074593289407 | 25.827854706351538 | 121.95060028174787 | 132.3098063959503 | 80.1907639309582 | 78.64359263476125 | 124.19286836189059 |
| SE_212_1_30 | SE_212 | SE_212_1 | AGGCCCTGCTCAGGGGTAGG | 428.0918524668781 | 517.6853070403582 | 580.6657952291118 | 525.7286297848258 | 366.9657014043747 | 505.9251041631507 | 411.09335407609535 | 390.4309915997044 | 466.19227085821416 | 807.739694868197 | 373.26361877394146 | 838.0116911895794 |
| SE_212_2_0 | SE_212 | SE_212_2 | TTTTTTCCCCCCACCAACAG | 1.083776841688299 | 0.9993924846338962 | 0.0 | 2.602616979132801 | 0.0 | 0.0 | 0.0 | 0.0 | 0.0 | 0.0 | 0.0 | 0.0 |
| SE_212_2_1 | SE_212 | SE_212_2 | ATTTTTTCCCCCCACCAACA | 2.167553683376598 | 0.9993924846338962 | 0.0 | 2.602616979132801 | 1.0605945127294067 | 0.0 | 0.0 | 0.0 | 0.0 | 0.0 | 0.0 | 0.0 |
| SE_212_2_2 | SE_212 | SE_212_2 | AATTTTTTCCCCCCACCAAC | 0.0 | 2.9981774539016888 | 0.987526862634544 | 1.3013084895664004 | 1.0605945127294067 | 0.0 | 0.0 | 7.56282792444948 | 4.439926389125849 | 0.0 | 23.47569929395858 | 1.1606810127279494 |
| SE_212_2_22 | SE_212 | SE_212_2 | GAGCCCTGCAGCTACAGCCG | 70.44549470973944 | 31.980559508284678 | 77.02709528549444 | 36.43663770785921 | 29.69664635642339 | 212.24597691776015 | 76.40740350628997 | 33.08737216946648 | 171.38115862025776 | 4.374041668961356 | 66.90574298778195 | 62.67677468730927 |
| SE_212_2_23 | SE_212 | SE_212_2 | GCTGGGCTCCCTGCACCCTG | 4904.090208639553 | 4344.359130703547 | 4248.340563053808 | 4622.2477549398545 | 4466.163493103531 | 3881.0692922104713 | 4788.914726802681 | 4684.226545705897 | 4418.614742458045 | 4536.610217657753 | 4745.612612273727 | 4619.510430657238 |
| SE_212_3_10 | SE_212 | SE_212_3 | CCTGTGGTCTTTTGAGAGGT | 253.60378095506195 | 417.7460585769686 | 408.83612113070126 | 422.9252591090801 | 510.14596062284465 | 321.40105076117965 | 302.40113218686594 | 401.7752334863786 | 396.9294191878509 | 575.9154864132452 | 316.9219404684408 | 343.561579767473 |
| SE_212_3_18 | SE_212 | SE_212_3 | TCTGAGGTTTCTCTGTGAAC | 289.36841673077583 | 311.8104552057756 | 332.79655270784133 | 275.8773997880769 | 434.84375021905674 | 357.7860753756528 | 224.91756806781132 | 377.1960427319178 | 231.7641575123693 | 439.5911877306163 | 244.14727265716925 | 492.12874939665056 |
| SE_212_3_21 | SE_212 | SE_212_3 | GGTTGGCCTTGATGAGAAAT | 314.2952840896067 | 274.8329332743215 | 311.0709617298814 | 398.20039780731855 | 369.08689042983355 | 395.03741009999436 | 373.4277326293327 | 354.5075589585694 | 216.66840778934142 | 610.907819764936 | 329.83357508011807 | 286.68821014380353 |
| SE_212_3_22 | SE_212 | SE_212_3 | GGGTTGGCCTTGATGAGAAA | 156.06386520311503 | 191.88335704970808 | 174.7922546863143 | 117.11776406097604 | 167.57393301124625 | 97.89304241513018 | 134.52007659558095 | 138.96696311175918 | 81.69464555991561 | 121.74415978609107 | 82.16494752885504 | 149.7278506419055 |
| SE_212_3_27 | SE_212 | SE_212_3 | GGCCTTCGAAGCCTGAAGCG | 208.0851536041534 | 257.8432610355452 | 250.8318231091742 | 217.31851775758886 | 261.9668446441635 | 200.9839454894708 | 156.0432888508739 | 273.20715877073746 | 235.31609862367 | 253.69441679975864 | 345.09277962119114 | 234.4575645710458 |
| SE_212_4_23 | SE_212 | SE_212_4 | GCCCGGAGGCAGGGGCGCTG | 444.3485050922026 | 693.5783843359239 | 655.7178367893372 | 545.2482571283218 | 791.2035064961374 | 739.8288338276211 | 464.9013847143277 | 877.2880392361396 | 543.4469900290039 | 1123.3997019782416 | 1307.5964506734929 | 971.4900076532937 |
| SE_212_4_28 | SE_212 | SE_212_4 | CCACTGGAGTGCGGAGGCCG | 526.7155450605133 | 644.608152588863 | 628.06708463557 | 542.645640149189 | 633.1749240994558 | 631.54007009407 | 530.5471820929712 | 706.1790574454702 | 691.7405314258073 | 642.2551183924925 | 777.0456466300291 | 925.0627671441757 |
| SE_212_4_4 | SE_212 | SE_212_4 | TCAAAAGACCACAGGCACCC | 55.27261892610324 | 117.92831318679976 | 104.67784743926167 | 109.30991312357764 | 102.87766773475245 | 194.91977472039196 | 52.73187002546773 | 90.75393509339376 | 111.88614500597139 | 83.83579865509266 | 26.997054188052367 | 76.60494684004466 |
| SE_212_4_5 | SE_212 | SE_212_4 | TGAGTTATCCTGGGTGCCTG | 450.8511661423323 | 638.6117976810597 | 577.7032146412082 | 512.7155448891617 | 550.4485521065621 | 596.8876656993336 | 464.9013847143277 | 714.6872388604759 | 633.133503089346 | 977.5983130128631 | 423.7363722559524 | 955.2404734751024 |
| SE_212_4_54 | SE_212 | SE_212_4 | TTGTGCCTCTGGAGACCCCC | 264.4415493719449 | 317.806810113579 | 372.2976272132231 | 292.7944101524401 | 320.29954284428084 | 495.5293828447298 | 263.65935012733866 | 284.5514006574117 | 455.5364475243121 | 341.1752501789858 | 153.7658303754287 | 273.9207190037961 |
| SE_212_5_11 | SE_212 | SE_212_5 | GCTCTCTGCTCCCAGGACGG | 98.6236925936352 | 120.92649064070145 | 116.52816979087619 | 143.14393385230406 | 139.9984756802817 | 31.187163955262715 | 121.60614924240517 | 164.4915073567762 | 79.91867500426528 | 252.23640291010486 | 193.67451917515828 | 107.94333418369929 |
| SE_212_5_21 | SE_212 | SE_212_5 | TGAGGGCCGCTGGAGACCCA | 3171.1310387799626 | 3162.0778213816475 | 3159.0984335679063 | 3237.655522041204 | 3439.5080047814663 | 3045.946346297325 | 3194.0446986854736 | 3011.8962209120054 | 2790.0497429266834 | 4190.331918864979 | 2883.9896582628116 | 2943.48704827808 |
| SE_212_5_3 | SE_212 | SE_212_5 | GTGTCAGCTCGACCAGGCCA | 2230.4127401945193 | 2038.7606686531483 | 1957.2782417416663 | 2067.77918992101 | 2200.733613913519 | 1671.1122019361605 | 2150.1689043037654 | 1859.510315924016 | 1688.9479984234729 | 2286.1657789771352 | 1933.2238368574892 | 2084.583098859397 |
| SE_212_5_36 | SE_212 | SE_212_5 | AGGCCCTTCTCAGGCTGCCG | 305.6250693561003 | 393.7606389457551 | 342.6718213341868 | 326.62843088116654 | 398.78353678625695 | 478.2031806473616 | 309.9342564762185 | 419.73694980694614 | 280.60334779275365 | 250.04938207562418 | 370.91604884454557 | 276.24208102925195 |
| SE_212_5_43 | SE_212 | SE_212_5 | GCTAAAACAAAATACCTCAG | 903.8698859680413 | 1420.1367206647665 | 1455.6145955233178 | 1234.941756598514 | 1465.7416165920401 | 1607.0052538058983 | 1055.7135611221192 | 1640.188306114981 | 1494.4792225797607 | 1567.3649313778192 | 1396.8041079905356 | 1687.6301925064386 |
| SE_213_1_11 | SE_213 | SE_213_1 | CGCAGGCTATCCCACCGATG | 12070.022685882584 | 11058.27784247406 | 10856.870327804178 | 12306.47438582945 | 11338.815935590088 | 11559.175795974177 | 12225.184561006396 | 12374.677191380462 | 11705.421932291389 | 12080.374082726437 | 12368.172173022078 | 13408.187059033271 |
| SE_213_1_20 | SE_213 | SE_213_1 | AGGGCAATGCTGCCAGGAGA | 1860.8448371788093 | 1781.916800102237 | 1688.6709351050704 | 1989.7006805470262 | 1692.708842316133 | 1850.438394678921 | 1824.0922386360774 | 1913.3954648857184 | 1898.512523990213 | 1659.2198064260076 | 1683.2076393768302 | 1707.3617697228135 |
| SE_213_1_27 | SE_213 | SE_213_1 | CCACTCTCTGCTCGCCCCCA | 438.92962088376106 | 478.70900013963626 | 431.5492389712957 | 337.0388987976977 | 399.8441312989863 | 510.2566547124927 | 373.4277326293327 | 400.82987999582247 | 373.8418019643965 | 451.2552988478466 | 356.8306292681704 | 584.9832304148865 |
| SE_213_1_4 | SE_213 | SE_213_1 | ATGGGGTAGAGGAGCGAAGA | 11853.267317544925 | 13586.740828597818 | 13692.059950427953 | 12620.089731814951 | 12720.770585676504 | 14581.731769305055 | 12937.602886656592 | 12658.283238547318 | 12673.325885120823 | 13356.865243118327 | 12253.14124648168 | 12473.838843787273 |
| SE_213_1_40 | SE_213 | SE_213_1 | GAGCAGCATTTTCGGGCCAG | 1458.7636289124503 | 1844.8785266341724 | 1753.8477080389503 | 1750.2599184668086 | 1755.2839185671683 | 1848.7057744591843 | 1628.2310071129116 | 1841.5485996034483 | 1682.7321014786967 | 2401.3488762597844 | 2314.703950384316 | 1884.9459646701898 |
| SE_213_2_0 | SE_213 | SE_213_2 | AGGGAGTGCGGAGTTAGAGA | 4051.157834230861 | 4273.40226429454 | 4438.933247542275 | 4266.990537288227 | 4441.769819310755 | 4336.748410001254 | 4032.373816029134 | 4243.691819106714 | 4320.048376619451 | 5262.7011347053385 | 4831.298914696676 | 4964.23269143744 |
| SE_213_2_10 | SE_213 | SE_213_2 | TCCTCCTAATTCTCTCCACG | 8.670214733506391 | 8.994532361705065 | 4.93763431317272 | 5.205233958265602 | 2.1211890254588135 | 0.0 | 3.228481838293942 | 1.89070698111237 | 0.0 | 5.103048613788249 | 0.0 | 0.0 |
| SE_213_2_35 | SE_213 | SE_213_2 | GGTTGCGAAGAGGGAAAGGG | 3035.658933568925 | 3764.711489615887 | 3899.7435805438145 | 2637.7523083510937 | 3179.6623491627615 | 1957.8608483026037 | 2469.788606294866 | 2280.1926192215183 | 3036.9096501620807 | 1555.700820260589 | 1919.138417281114 | 1672.5413393409751 |
| SE_213_2_7 | SE_213 | SE_213_2 | GCCCTTTCCAAGCATTCCCA | 107.2939073271416 | 160.9021900260573 | 149.11655625781614 | 203.00412437235846 | 208.93711900769313 | 294.54543735525897 | 123.75847046793446 | 185.28928414901227 | 215.78042251151626 | 301.07986821350664 | 262.9278320923361 | 286.68821014380353 |
| SE_213_2_9 | SE_213 | SE_213_2 | TCTTATCTCACTCTGCTAGA | 87.78592417675222 | 122.92527560996923 | 103.69032057662713 | 144.44524234187045 | 123.02896347661118 | 154.20319955657675 | 111.92070372752333 | 89.80858160283758 | 118.99002722857276 | 83.83579865509266 | 34.03976397623994 | 181.06623798556012 |
| SE_213_3_25 | SE_213 | SE_213_3 | AAAAGGGCACTTTTCCTCCA | 50.93751155935005 | 92.94350107095235 | 77.02709528549444 | 63.76411598875362 | 68.93864332741144 | 155.06950966644516 | 31.208657770174778 | 82.2457536783881 | 146.517570841153 | 80.91977087578509 | 79.81737759945918 | 30.177706330926686 |
| SE_213_3_43 | SE_213 | SE_213_3 | CATCAGTGACTCCTAGCCAG | 1197.5734100655702 | 1305.2065849318685 | 1394.3879300399763 | 1111.317450089706 | 1440.2873482865343 | 1127.0694529387997 | 1349.505408406868 | 1616.5544688510763 | 1370.1612836842369 | 1764.1968064810803 | 1430.8438719667754 | 1110.7717291806475 |
| SE_213_3_44 | SE_213 | SE_213_3 | GGATGTGGGTGACCAGGCCG | 1595.319510965176 | 1849.8754890573418 | 1887.1638344946136 | 1660.469632686727 | 1729.8296502616624 | 1818.38492061379 | 1691.7244832660258 | 1458.6804359281934 | 1795.5062317624934 | 1980.7118690946672 | 1490.7069051663698 | 1946.462058344771 |
| SE_213_3_48 | SE_213 | SE_213_3 | GGTTTAACAACTCACACGTG | 314.2952840896067 | 214.86938419628768 | 233.05633958175238 | 321.4231969229009 | 258.7850611059753 | 291.08019691578534 | 263.65935012733866 | 244.84655405405192 | 174.04511445373328 | 241.30129873770147 | 323.96465025662843 | 154.37057469281729 |
| SE_213_3_5 | SE_213 | SE_213_3 | TGACTCATCTAGATTAGCCT | 466.0240419259685 | 371.7740042838094 | 449.32472249871756 | 537.4404061909233 | 508.02477159738584 | 440.95184592302 | 476.7391514547388 | 308.1852379213163 | 491.05585863731886 | 380.54162519963796 | 453.0809963734006 | 539.7166709184964 |
| SE_213_4_12 | SE_213 | SE_213_4 | GTGGGCATCTCTTCTCTGAT | 540.8046440024611 | 482.70657007817186 | 488.8257970040993 | 546.5495656178882 | 565.2968752847738 | 634.1390004236752 | 525.166379029148 | 454.71502895752496 | 527.4632550281508 | 344.09127795829335 | 571.6332778078914 | 661.5881772549312 |
| SE_213_4_15 | SE_213 | SE_213_4 | GGAGTCAAAAAGAAAGCAAG | 314.2952840896067 | 360.7806869528365 | 395.99827191645215 | 355.25721765162734 | 339.39024407341014 | 477.3368705374932 | 356.2091628250983 | 415.9555358447214 | 367.6259050196203 | 497.1827363719408 | 287.5773163509926 | 464.2724050911798 |
| SE_213_4_17 | SE_213 | SE_213_4 | TTTGGAAAGGAACAGAGCAC | 197.2473851872704 | 170.89611487239625 | 189.60515762583245 | 231.63291114281927 | 130.45312506571702 | 182.79143318223424 | 266.8878319656326 | 110.60635839507364 | 245.08393667974687 | 185.16776398603074 | 116.20471150509498 | 104.46129114551545 |
| SE_213_4_20 | SE_213 | SE_213_4 | GGAGAGCTGGAGGCACAAAG | 2282.4340285955577 | 2488.4872867384015 | 2453.0167267842075 | 2514.128001842286 | 2737.3944373545987 | 2880.481115312459 | 2386.9242391119883 | 2546.7823035583624 | 2520.990203745657 | 2850.41715427315 | 2426.2135220306195 | 3072.322640690882 |
| SE_213_4_9 | SE_213 | SE_213_4 | GTACATGAAAGTAGGCAGGA | 972.1478269944041 | 1050.361501350225 | 1125.78062340338 | 985.0905266017651 | 1167.7145585150768 | 1214.566774035509 | 910.4318783988917 | 1204.3803469685797 | 1172.140566729224 | 882.0984032405402 | 1575.2194226246208 | 1151.3955646261259 |
| SE_213_5_0 | SE_213 | SE_213_5 | GATTTCCTATGAAAGGTGAG | 0.0 | 14.990887269508443 | 3.950107450538176 | 7.807850937398403 | 1.0605945127294067 | 0.0 | 1.0761606127646475 | 2.836060471668555 | 3.551941111300679 | 18.225173620672315 | 0.0 | 15.088853165463343 |
| SE_213_5_1 | SE_213 | SE_213_5 | ACTTCCTAGGAAAAAGCTGC | 130.05322100259588 | 130.9204154870404 | 113.56558920297256 | 109.30991312357764 | 115.60480188750533 | 98.75935252499859 | 151.73864639981528 | 120.0598933006355 | 128.75786528464963 | 172.77464592397357 | 158.4609702342204 | 59.19473164912542 |
| SE_213_5_11 | SE_213 | SE_213_5 | AATTTATCCATTAATCTGCG | 5.4188842084414945 | 6.9957473924372735 | 4.93763431317272 | 3.9039254686992013 | 4.242378050917627 | 5.197860659210452 | 0.0 | 5.67212094333711 | 0.0 | 2.187020834480678 | 16.432989505771005 | 1.1606810127279494 |
| SE_213_5_2 | SE_213 | SE_213_5 | ATATCCGGCAGCTTTTTCCT | 18.42420630870108 | 26.983597085115196 | 29.62580587903632 | 44.24448864525762 | 40.30259148371746 | 10.395721318420904 | 27.980175931880833 | 22.68848377334844 | 50.615160836034676 | 45.198430579267345 | 11.73784964697929 | 147.40648861644956 |
| SE_213_5_7 | SE_213 | SE_213_5 | AGAATTCATAAGCTACAGGG | 11.921545258571289 | 35.97812944682026 | 38.51354764274722 | 32.53271223916001 | 12.727134152752882 | 16.459892087499767 | 12.913927353175769 | 7.56282792444948 | 52.391131391685015 | 26.244250013768134 | 4.695139858791716 | 35.98111139456643 |
| SE_213_6_13 | SE_213 | SE_213_6 | TTAAGGGTTTCAGGCCAAGG | 326.21682934817795 | 398.75760136892455 | 412.7862285812394 | 317.5192714542017 | 307.572408691528 | 319.66843054144283 | 291.63952605921946 | 319.5294798079905 | 349.8661994631169 | 214.32804177910643 | 367.39469395045177 | 300.6163822965389 |
| SE_213_6_19 | SE_213 | SE_213_6 | AAGAACTGAGAAGAGCCGTT | 197.2473851872704 | 266.83779339725027 | 273.5449409497687 | 294.0957186420065 | 294.8452745387751 | 181.92512307236584 | 159.27177068916782 | 283.6060471668555 | 309.0188766831591 | 244.94633346183593 | 215.97643350441894 | 254.18914178742094 |
| SE_213_6_20 | SE_213 | SE_213_6 | GTTCTTGTCACCTAAGAGAG | 14.089098941947885 | 13.991494784874547 | 20.738064115325425 | 11.711776406097604 | 42.42378050917627 | 5.197860659210452 | 4.30464245105859 | 9.45353490556185 | 66.59889583688773 | 0.7290069448268927 | 4.695139858791716 | 60.35541266185337 |
| SE_213_6_22 | SE_213 | SE_213_6 | GCGCCTTCCTAAGTACCCAA | 400.9974314246706 | 460.7199354162261 | 445.37461504817935 | 432.03441853604494 | 524.9942838010563 | 348.2566641671003 | 320.69586260386495 | 473.6220987686487 | 455.5364475243121 | 387.10268770308 | 306.35787578615947 | 385.3460962256792 |
| SE_213_7_15 | SE_213 | SE_213_7 | GCAACGCAATCTGACCGAGC | 6607.787403773558 | 7640.3555450261365 | 7710.60974345052 | 7029.668460637695 | 7566.281253811588 | 8346.03159847225 | 6484.943852519766 | 7700.849534070683 | 6607.498452297088 | 7939.614636109688 | 7597.910076489695 | 8164.230243528396 |
| SE_213_7_16 | SE_213 | SE_213_7 | GCGGCGGCGTACGACCTGCT | 14672.17088277619 | 11866.786362542884 | 11775.270310054302 | 13921.398221381352 | 12464.106713595987 | 13604.53396537349 | 14875.768150245722 | 13458.05229155785 | 12473.52919761016 | 13654.3000766077 | 13409.319436709142 | 16318.014357942242 |
| SE_213_7_18 | SE_213 | SE_213_7 | TCTCCCTTGCAGGTGACCCG | 28.17819788389577 | 28.982382054382988 | 36.53849391747813 | 28.62878677046081 | 28.63605184369398 | 8.663101098684088 | 16.142409191469714 | 9.45353490556185 | 10.655823333902038 | 23.328222234460565 | 19.954344399864794 | 4.642724050911798 |
| SE_213_7_23 | SE_213 | SE_213_7 | GGAGGGACTCCGCCTCCGAG | 997.074694353235 | 1352.1780317096616 | 1305.510512402867 | 959.0643568104372 | 1342.712653115429 | 1014.4491386559066 | 1023.4287427391797 | 956.6977324428592 | 1124.1893617266649 | 1374.1780909986926 | 954.2871762994163 | 1031.845420315147 |
| SE_213_7_31 | SE_213 | SE_213_7 | AGCCAGGCTCCTCTGCACCG | 49.85373471766175 | 69.95747392437274 | 77.02709528549444 | 71.57196692615203 | 74.24161589105847 | 99.625662634867 | 77.48356411905462 | 63.33868386726439 | 77.25471917078977 | 88.9388472688809 | 57.515463270198524 | 156.69193671827318 |
| SE_214_1_13 | SE_214 | SE_214_1 | CAGAGGCTAGAAGAACAGGA | 1717.7862940759537 | 1869.8633387500197 | 1923.7023284120917 | 1683.893185498922 | 1854.9798027637323 | 1587.0801212789247 | 1449.58834539398 | 2082.6137396952754 | 2128.500710946932 | 1529.4565702468208 | 2259.536057043513 | 2269.131379883141 |
| SE_214_1_23 | SE_214 | SE_214_1 | ACCTTCCCATTCCCACTGCA | 171.23674098675122 | 65.95990398583714 | 96.77763253818532 | 117.11776406097604 | 33.939024407341016 | 386.3743090013103 | 201.24203458698906 | 62.393330376708214 | 365.84993446396993 | 16.038152786191638 | 97.42415206992811 | 127.67491140007444 |
| SE_214_1_26 | SE_214 | SE_214_1 | AGAGCCAAAAGAAAGAGCTG | 928.7967533268721 | 1146.303179875079 | 1090.2296563485365 | 1167.2737151410613 | 1077.5640249330772 | 1105.4117001920895 | 881.3755418542463 | 1259.2108494208385 | 1136.6211556162173 | 1245.8728687091595 | 1223.083933215242 | 1535.5809798390771 |
| SE_214_1_6 | SE_214 | SE_214_1 | GCTGTGTGTGTAGATGTGGG | 3243.7440871730787 | 3224.040155428949 | 3309.2025166883573 | 3227.245054124673 | 3216.7831571082907 | 2579.871507188121 | 2970.203291230427 | 3261.469542418838 | 3157.675647946304 | 2791.367591742172 | 3503.748119623318 | 2778.670344470711 |

| | | | | | | | | | | | | | | | |
|---|---|---|---|---|---|---|---|---|---|---|---|---|---|---|---|
| SE_214_1_7 | SE_214 | SE_214_1 | GGTGGTGTGGAGAGGCCCCC | 120.29922942740117 | 152.90705014898612 | 130.3535458677598 | 117.11776406097604 | 129.39253055298764 | 97.02673230526177 | 74.25508228076067 | 108.71565141396128 | 99.45435111641902 | 110.0800486688608 | 37.56111887033373 | 18.57089620364719 |
| SE_214_2_22 | SE_214 | SE_214_2 | TGGAGGTCAGAGTCAGAAAT | 424.84052194181317 | 458.72115044695835 | 429.5741852460267 | 419.02133364038093 | 404.08650934990396 | 559.636330974992 | 340.0667536336286 | 450.9336149953003 | 440.4406978012842 | 495.724722482287 | 608.0206117135273 | 430.61265572206923 |
| SE_214_2_24 | SE_214 | SE_214_2 | GCGAAAACTCCATCTCAGGA | 1077.2741806381691 | 1274.2254179082177 | 1328.2236302434617 | 1207.6142783176197 | 1263.1680646607235 | 1567.1549887519514 | 997.6008880328282 | 1314.9867053636533 | 1396.8008420189922 | 1051.9570213852062 | 1183.1752444155125 | 1250.0534507080015 |
| SE_214_2_5 | SE_214 | SE_214_2 | TTTGGTAGCAGACATGGTCG | 312.12773040623006 | 324.8025575060163 | 349.5845093726286 | 392.9951638490529 | 314.9965702806338 | 424.4919538355203 | 375.58005385486194 | 233.5023121673777 | 532.7911666951019 | 184.43875704120384 | 342.74520969179525 | 224.01143545649424 |
| SE_214_2_8 | SE_214 | SE_214_2 | ATTGTGCTATCTCAAATCGA | 266.60910305532155 | 256.84386855091134 | 268.607306636596 | 283.68525072547527 | 217.42187510952837 | 504.1924839434139 | 260.43086828904467 | 271.3164517896251 | 431.5608450230325 | 131.95025701366757 | 279.3608215981071 | 300.6163822965389 |
| SE_214_2_9 | SE_214 | SE_214_2 | ACCAAAACCAAGATTAGGAA | 411.83519984155356 | 517.6853070403582 | 503.63869994361744 | 415.11740817168175 | 515.4489331864917 | 824.7272245947252 | 426.1596026548004 | 605.0262339559584 | 641.1253705897726 | 552.5872641787846 | 460.1237061615882 | 714.9795038404169 |
| SE_215_1_17 | SE_215 | SE_215_1 | CAGAAATGAGGTCAAATGAC | 861.6025891421976 | 804.5109501302865 | 878.8989077447442 | 867.9727625407891 | 736.0525918342083 | 602.085526358544 | 897.517951045716 | 776.1352157466279 | 915.5128214377501 | 1466.032966046881 | 770.0029368418415 | 745.1572101713435 |
| SE_215_1_2 | SE_215 | SE_215_1 | ACATCTGTGGAATACATCCC | 1667.932559358292 | 1625.012180014715 | 1607.6937323690377 | 1488.6969120639621 | 1407.4089183919227 | 1183.3796100802463 | 1435.5982574280397 | 1614.663761869964 | 1243.1793889552378 | 1350.849868764232 | 1726.6376830706536 | 1436.9230937572015 |
| SE_215_1_22 | SE_215 | SE_215_1 | AAACGATGCCTACTGCCACT | 4883.498448647475 | 4513.256460606675 | 4400.4196998995285 | 4644.369999262483 | 4385.558310136097 | 4135.764464511783 | 4794.295529866505 | 4730.54886674315 | 4851.951558036728 | 4089.728960478868 | 4470.946930534412 | 4520.852544575363 |
| SE_215_1_32 | SE_215 | SE_215_1 | GTGACTCCACAGGTGTGCCG | 596.0772629285644 | 562.6579688488836 | 574.7406340533046 | 572.5757354092162 | 529.236661851974 | 802.2031617381465 | 564.98432170144 | 653.2392619743239 | 737.0277805948909 | 622.5719308821664 | 606.8468267488294 | 644.1779620640119 |
| SE_215_1_39 | SE_215 | SE_215_1 | TTGCCAGGCTGCCTCGGGCA | 201.5824925540236 | 193.88214201897586 | 215.2808560543306 | 268.0695488506785 | 308.63300320425736 | 170.66309164407653 | 215.2321225529295 | 150.31120499843342 | 178.48504084285912 | 316.38901405487144 | 193.67451917515828 | 210.08326330375886 |
| SE_215_2_12 | SE_215 | SE_215_2 | CGAGTACTGACTAGGACACA | 5683.32575781344 | 5173.85489294968 | 5102.551299232689 | 5448.5786458145185 | 5147.0651702758105 | 5897.839227984126 | 5365.736815244532 | 5000.919965042219 | 5354.551225285773 | 5391.006356994872 | 4755.00289199131 | 5442.433268681355 |
| SE_215_2_15 | SE_215 | SE_215_2 | TACAGACAGTAAGAAAAGGG | 771.6491112820688 | 980.4040274258522 | 972.7139596950259 | 791.1955616563714 | 934.3837657146073 | 938.2138489874867 | 788.8257291564865 | 792.206225086083 | 968.7919381072602 | 1427.3955979710558 | 625.6273861839961 | 1448.529903884481 |
| SE_215_2_23 | SE_215 | SE_215_2 | ATGTCTTCCTAAAAGTTGCA | 60.69150313454474 | 68.95808143973883 | 62.21419234597627 | 72.87327541571842 | 79.5445884547055 | 97.02673230526177 | 45.19874573611519 | 128.56807471564116 | 59.49501361428638 | 83.10679171026577 | 117.3784964697929 | 95.17584304369186 |
| SE_215_2_26 | SE_215 | SE_215_2 | TGCCGCAGTAGTCAAAGCGA | 3280.592499790481 | 3189.061418466763 | 3257.851119831361 | 3267.5856173012317 | 3454.356327959678 | 3547.539899911134 | 3205.8824654258847 | 3491.1904406239914 | 2695.03531819939 | 3618.7904741206953 | 3315.9425252716496 | 3755.963757187644 |
| SE_215_2_27 | SE_215 | SE_215_2 | ATGCTGGCCTCACCCTCCCA | 5169.615534853186 | 4594.207251862021 | 4675.939694574566 | 5075.103109308961 | 4656.009910882096 | 4575.850000324935 | 5222.607453746834 | 4710.6964434414695 | 4842.1837199806505 | 4945.583113705639 | 4751.4815370972165 | 5379.756493994046 |
| SE_216_1_15 | SE_216 | SE_216_1 | AATGTCCTAATGTTTGGCCT | 239.51468201311405 | 305.81410029797223 | 253.79440369707783 | 238.13945359065127 | 238.63376536411653 | 316.2031901019692 | 243.21229848481033 | 167.32756782844476 | 147.40555611897818 | 139.96933340676338 | 117.3784964697929 | 95.17584304369186 |
| SE_216_1_16 | SE_216 | SE_216_1 | GGTGCAGCTTGCTAGAGCCA | 1722.1214014427069 | 1752.9344180478538 | 1579.0554533526358 | 1679.989260030223 | 1600.4371197086748 | 1540.2993753460307 | 1782.1219747382563 | 1423.7023567776146 | 1874.5369214889333 | 2319.7000984391725 | 1142.092770651085 | 1960.3902304975065 |
| SE_216_1_2 | SE_216 | SE_216_1 | CAGATTCTGGAAGTTTCCAT | 98.6236925936352 | 126.92284554850481 | 158.00429802152703 | 124.92561499837444 | 171.8163110621639 | 215.71121735723378 | 138.82471904663953 | 123.84130726286024 | 153.62145306375436 | 67.79764586890101 | 184.28423945757487 | 358.6504329329364 |
| SE_216_1_20 | SE_216 | SE_216_1 | AGGCCCACTTGCCTGCCAGA | 2169.7212370599746 | 2243.636128003097 | 2202.184903675033 | 2261.674154866404 | 2561.335748241517 | 2502.7699074098327 | 2524.672797545863 | 2331.2417077115524 | 2187.995724561218 | 1909.9981954464588 | 2733.7451827814766 | 2563.9443571160405 |
| SE_216_1_6 | SE_216 | SE_216_1 | TTAATGCTTTCAGCTTCCAG | 17.340429467012783 | 18.98845720804403 | 28.63827901640178 | 16.917010364363207 | 11.666539640023474 | 0.0 | 26.904015319116187 | 3.78141396222474 | 28.415528890405433 | 26.244250013768134 | 31.692194046844083 | 0.0 |
| SE_216_2_1 | SE_216 | SE_216_2 | TTTGTTGAATTATTTCCTGA | 4.335107366753196 | 17.98906472341013 | 17.77548352742179 | 2.602616979132801 | 8.484756101835254 | 0.0 | 8.60928490211718 | 0.945353490556185 | 2.6639558334755096 | 17.496166675845423 | 1.173784964697929 | 2.321362025455899 |
| SE_216_2_10 | SE_216 | SE_216_2 | GCCATAGTCCGTCGTCCCTG | 2329.0364327881543 | 2780.309892251499 | 2628.796508333156 | 2540.1541716336137 | 2563.456937266976 | 2391.0159032368083 | 2101.7416767293566 | 2911.6887509130497 | 2765.1861551475786 | 3048.7070432660653 | 2909.8129274861662 | 2754.2960432034242 |
| SE_216_2_5 | SE_216 | SE_216_2 | GGGCTTTGTGATAGGAATAAC | 13.005322100259587 | 24.984812115847404 | 26.66322529113269 | 10.410467916531204 | 22.272484767317543 | 0.0 | 3.228481838293942 | 0.0 | 9.767838056076867 | 11.664111117230282 | 16.432989505771005 | 0.0 |
| SE_216_2_6 | SE_216 | SE_216_2 | ATAGAATCAGGGATATTTGG | 32.51330525064897 | 65.95990398583714 | 79.98967587339807 | 24.724861301761607 | 40.30259148371746 | 53.71122681184134 | 11.837766740411123 | 20.79777679223607 | 38.1833669464823 | 13.851131951710961 | 3.521354894093787 | 217.04734938012655 |
| SE_216_2_9 | SE_216 | SE_216_2 | GGCTCTGCAGAGTTAAGGAG | 2.167553683376598 | 2.9981774539016888 | 4.93763431317272 | 2.602616979132801 | 1.0605945127294067 | 10.395721318420904 | 1.0761606127646475 | 0.0 | 19.535676112153734 | 0.0 | 0.0 | 0.0 |
| SE_216_3_10 | SE_216 | SE_216_3 | GATTCTGACAGTCTCCCAAG | 13.005322100259587 | 25.984204600481302 | 31.60085960430541 | 23.423552812195208 | 33.939024407341016 | 1.7326202197368175 | 13.990087965940416 | 1.89070698111237 | 35.51941111300679 | 24.78623612411435 | 1.173784964697929 | 6.964086076367696 |
| SE_216_3_16 | SE_216 | SE_216_3 | TGCTTTTTAAGATCTCCAGA | 6.502661050129793 5 | 21.986634661945715 | 25.675698428498144 | 19.519627343496005 | 12.727134152752882 | 3.465240439473635 | 18.294730416999005 | 9.45353490556185 | 46.17523444690883 | 10.206097227576498 | 4.695139858791716 | 0.0 |
| SE_216_3_2 | SE_216 | SE_216_3 | AACCTGTAGGAAGAGCACAG | 106.21013048545329 | 129.9210230024065 | 184.66752331265974 | 134.03477442533924 | 151.66501532030517 | 155.93581977631356 | 158.19561007640317 | 235.39301914849005 | 103.0062922277197 | 490.62167386849876 | 126.76877618737633 | 298.295020271083 |
| SE_216_3_21 | SE_216 | SE_216_3 | TCCCGATAACATGCTCCCAT | 405.3325387914238 | 418.7454510616025 | 430.5617121086612 | 443.74619494214255 | 396.6623477607981 | 382.90906856183665 | 394.95094488462564 | 530.3433082020198 | 483.06399113689235 | 377.6255974203304 | 546.983793549235 | 396.95290635295873 |
| SE_216_3_9 | SE_216 | SE_216_3 | TATGTCATATTGTTTTCCTG | 466.0240419259685 | 575.6500711491242 | 589.5535369928228 | 594.697979731845 | 658.6291924049616 | 726.8341821795949 | 492.8815606462085 | 714.6872388604759 | 693.5165019814576 | 819.4038059854273 | 860.384379123582 | 1001.6677139842203 |
| SE_217_1_17 | SE_217 | SE_217_1 | CATTCCGAGGGCCTGAAGAA | 240.59845885480235 | 293.8213904823655 | 281.445155850845 | 275.8773997880769 | 216.36128059679896 | 414.0962325170994 | 266.8878319656326 | 177.72645622456278 | 346.3142583518162 | 245.67534040666283 | 75.12223774066746 | 284.3668481183476 |
| SE_217_1_20 | SE_217 | SE_217_1 | GGAATGCCCAACCTGTCCTG | 442.18095140882593 | 457.72175796232443 | 417.72386289441215 | 413.81609968211535 | 408.3288874008216 | 329.1978417499953 | 397.1032661101549 | 501.037349994778 | 381.833669464823 | 440.3201946754432 | 593.9351921371521 | 490.9680683839226 |
| SE_217_1_33 | SE_217 | SE_217_1 | AGACCCCACTGACAGCCATA | 678.4443028968751 | 629.6172653193546 | 710.0318142342371 | 706.6105098345554 | 625.75076251035 | 791.8074404197256 | 816.8059050883675 | 771.408448293847 | 762.7793536518209 | 665.583340626953 | 917.8998423937805 | 690.6052025731299 |
| SE_217_1_34 | SE_217 | SE_217_1 | ATAGCTCAAGTGACCCCAGA | 223.25802938778958 | 308.8122777518739 | 297.24558565299776 | 219.92113473672168 | 326.6631099206573 | 422.7593336157835 | 224.91756806781132 | 251.4640284879452 | 265.50759806972576 | 286.4997293169688 | 282.88217649220087 | 354.0077088820246 |
| SE_217_1_7 | SE_217 | SE_217_1 | GTTGGCAGCTCCATTCTGCA | 178.8231788785693 | 53.96719417023039 | 110.60300861506893 | 89.79028578008163 | 28.63605184369398 | 588.2245646006495 | 291.63952605921946 | 75.6282792444948 | 556.7667691963815 | 12.393118062057175 | 71.60088284657367 | 106.78265317097134 |
| SE_217_2_10 | SE_217 | SE_217_2 | TCTCCTGGCTGGACCCCTGC | 311.04395356454177 | 408.75152621526354 | 375.26020780112674 | 303.2048780689713 | 435.9043447317862 | 424.4919538355203 | 302.40113218686594 | 277.93392622351837 | 365.84993446396993 | 549.671236399477 | 176.06774470468935 | 489.80738737119464 |
| SE_217_2_11 | SE_217 | SE_217_2 | GCAGGGGTCCAGCCAGGAGA | 3634.9875270225543 | 4509.258890668139 | 4480.409375772926 | 3983.305286562752 | 4230.711511277604 | 4924.972974601904 | 3761.181341612443 | 4117.014451372185 | 5006.460996378307 | 3909.6642451066255 | 3299.5095357658784 | 4780.845091426424 |
| SE_217_2_19 | SE_217 | SE_217_2 | AGATGGGGGCAACTTGGACA | 1968.1387445059509 | 2259.626407757239 | 2249.5861930814913 | 2303.3160265325287 | 2398.0041932811887 | 2320.844784337467 | 1974.7547244231282 | 2468.317963842199 | 2317.641575123693 | 1688.3800842190833 | 2590.5434170883295 | 2760.099448267064 |
| SE_217_2_3 | SE_217 | SE_217_2 | AGAGAAACAGGGGTGCACCT | 1322.2077468597247 | 1360.1731715867327 | 1219.5956753536618 | 1225.8325971715492 | 1380.8940555736876 | 1235.3582166723509 | 1272.0218442878133 | 1264.8829703641754 | 1303.5623878473493 | 1019.1517088679959 | 1259.4712671208779 | 1243.0893646316338 |
| SE_217_2_6 | SE_217 | SE_217_2 | GTGCACCTGGGGCTCCTGCA | 3.2513305250648967 | 3.9975699385355847 | 8.887741763710896 | 1.3013084895664004 | 1.0605945127294067 | 50.2459863723677 | 5.380803063823238 | 20.79777679223607 | 35.51941111300679 | 2.9160277793075706 | 0.0 | 20.89225822910309 |
| SE_217_3_11 | SE_217 | SE_217_3 | TTATTTCAGGACAATCATGC | 44.434850509220254 | 54.96658665486429 | 55.301504307534465 | 53.35364807222242 | 47.72675307282331 | 25.989303296052263 | 34.43713960846872 | 9.45353490556185 | 19.535676112153734 | 72.17168753786237 | 12.911634611677218 | 11.606810127279495 |
| SE_217_3_13 | SE_217 | SE_217_3 | GCTGTTAGTGAAGAATGGCC | 1118.4577006223244 | 1147.3025723597127 | 1180.0946008482802 | 1074.8808123818467 | 1126.35137251863 | 1156.5239966743256 | 1046.0281156072374 | 1049.3423745173654 | 1227.1956539543846 | 1030.0868130403994 | 1294.6848160618158 | 1436.9230937572015 |
| SE_217_3_23 | SE_217 | SE_217_3 | CCTTCAACACAGTTTCTCCC | 1710.1998561841356 | 1563.0498459674136 | 1726.196955885183 | 1569.3780384170789 | 1713.9207325707214 | 1652.9196896289238 | 1650.8303799809692 | 1349.019431023676 | 1840.793480931577 | 1573.9259938812613 | 1343.9837845791287 | 1663.2558912391517 |
| SE_217_3_26 | SE_217 | SE_217_3 | AGTCTGGAGGGGAGGCCAGG | 301.2899619893471 | 309.81167023650784 | 364.39741231214674 | 303.2048780689713 | 392.4199697098805 | 315.33687999210076 | 316.39122015280634 | 525.6165407492389 | 282.379318348404 | 230.3661945652981 | 611.541966607621 | 233.29688355831783 |
| SE_217_3_32 | SE_217 | SE_217_3 | GGGGTTGTCTGGAGTCTGGA | 45.51862735090855 | 107.93438834046079 | 77.02709528549444 | 46.847105624390416 | 68.93864332741144 | 24.256683076315444 | 22.599372868057596 | 45.37696754669688 | 38.1833669464823 | 67.79764586890101 | 165.503680022408 | 22.05293924183104 |
| SE_218_1_1 | SE_218 | SE_218_1 | GCTTGAGAGATTCACCCACT | 1851.0908456036145 | 2111.7163200314226 | 2114.295012900559 | 1899.9103947669446 | 1874.0705039928616 | 2041.8929289598393 | 1947.850709104012 | 1440.7187196076259 | 2237.722900119428 | 2246.7994039564833 | 1988.3917301982917 | 2176.276898864905 |
| SE_218_1_16 | SE_218 | SE_218_1 | AGAGATCCTGTGGGCTAGCG | 902.786109126353 | 1065.3523886197333 | 934.2004120522787 | 977.2826756643667 | 973.6257626855954 | 1318.523987219718 | 971.7730333264767 | 812.0586483877629 | 961.6880558846589 | 1294.7163340125614 | 600.9779019253397 | 1073.6299367733532 |
| SE_218_1_3 | SE_218 | SE_218_1 | AAGAAAGAAGAAAAGACACA | 236.26335148804915 | 310.8110627211417 | 290.3328976145559 | 257.6590809341473 | 269.3910062332693 | 378.57751801249464 | 314.23889892727703 | 408.3927079202719 | 340.9863466848652 | 389.28970853756067 | 579.849772560777 | 298.295020271083 |
| SE_218_1_4 | SE_218 | SE_218_1 | GATTATGTCAAGTTCAAAAT | 47.686181034285156 | 58.96415659339988 | 103.69032057662713 | 66.36673296788642 | 49.84794209828212 | 74.50266944868315 | 25.827854706351538 | 25.524544245016994 | 60.382998892111544 | 51.030486137882484 | 37.56111887033373 | 29.017025318198737 |
| SE_218_1_5 | SE_218 | SE_218_1 | GTTTTGAGGAATTAGCCTGG | 61.77527997623304 | 72.95565137827442 | 75.05204156022535 | 68.96934994701923 | 102.87766773475245 | 74.50266944868315 | 33.36097899570407 | 92.64464207450614 | 83.47061611556596 | 44.46942363444045 | 34.03976397623994 | 55.71268861094157 |
| SE_218_2_13 | SE_218 | SE_218_2 | TACTTACTCATTCTTCACAA | 277.4468714722045 | 346.78919216796197 | 351.5595630978977 | 266.7682403611121 | 318.178353818822 | 481.6684210868353 | 286.2587229953962 | 407.44735442971574 | 429.78487446738217 | 255.15243068941243 | 383.8276834562228 | 406.2383544547823 |
| SE_218_2_14 | SE_218 | SE_218_2 | CACAAAGGCATGCTTGTTGA | 2780.971375772175 | 2545.4526583625334 | 2434.253716394151 | 2646.8614677780583 | 2563.456937266976 | 2332.1068157657564 | 3058.448461477128 | 2499.5146290305534 | 2738.5465968128237 | 2394.0588068115153 | 2123.3770011385536 | 2417.6985495123186 |
| SE_218_2_15 | SE_218 | SE_218_2 | AATTCTCTTCAGAGATGTTG | 0.0 | 0.9993924846338962 | 0.0 | 0.0 | 0.0 | 0.0 | 0.0 | 0.0 | 0.0 | 0.7290069448268927 | 1.173784964697929 | 0.0 |
| SE_218_2_16 | SE_218 | SE_218_2 | GTGTAACCATGATGTACTAC | 403.1649851080472 | 385.76549906868394 | 419.6989166196812 | 407.3095572342833 | 405.1471038626334 | 485.9999716361773 | 383.1131781442145 | 304.4038239590916 | 565.6466219746331 | 418.44998633063636 | 246.49484258656508 | 376.06064812385563 |
| SE_218_2_17 | SE_218 | SE_218_2 | GGAGAGCCAGTAGTACATCA | 3793.2189459090464 | 3441.9077170791384 | 3325.0029464905097 | 3752.973683909499 | 3747.080413472994 | 4147.892806049941 | 3948.4332882334916 | 4183.189195711118 | 3482.678259630316 | 2915.298772362744 | 3831.2341247740405 | 3357.850169821958 |
| SE_219_1_1 | SE_219 | SE_219_1 | GTACCGTCACCTGTAAGCTT | 1805.572218252706 | 1919.8329629817147 | 1763.7229766652956 | 1810.120108986863 | 1775.4352143090268 | 1609.6041841355034 | 1830.5492023126653 | 1868.9638508295777 | 1632.116940642662 | 1663.593848094969 | 1895.6627179871555 | 1802.5376127665054 |
| SE_219_1_13 | SE_219 | SE_219_1 | GCCTGCTGTCAGCGCTTCCG | 2924.0299188750305 | 3882.6398028026865 | 3831.604227022031 | 3091.9089712097675 | 4016.4714197062635 | 3882.8019124302077 | 2782.9513446093783 | 4094.3259675988375 | 3686.0268882522796 | 3565.572967148332 | 3711.5080583748518 | 3529.6309597056943 |
| SE_219_1_2 | SE_219 | SE_219_1 | GCAGAATTCATAATGCTGGA | 472.5267029760983 | 674.58992712788 | 658.6804173772408 | 534.8377892117906 | 626.8113570230794 | 396.7700303197312 | 413.2456753016246 | 640.9496665970935 | 800.9627205983031 | 649.5451878407614 | 618.5846763958086 | 396.95290635295873 |
| SE_219_1_31 | SE_219 | SE_219_1 | GAAGATTCCAGAGGGAGGGA | 202.6662693957119 | 299.81774539016885 | 275.51999467503776 | 147.04785932100324 | 202.5735519313167 | 130.81282659012973 | 112.99686434028798 | 164.4915073567762 | 371.17784613092095 | 89.6678542137078 | 82.16494752885504 | 96.3365240564198 |
| SE_219_1_8 | SE_219 | SE_219_1 | AAGGTCTTTTGGTTTCCCAA | 94.288585226882 | 182.888824688003 | 163.9294591973343 | 108.00860463401123 | 160.14977142214042 | 157.66843999605038 | 121.60614924240517 | 113.4424188667422 | 148.29354139680336 | 138.5113195171096 | 70.42709788187574 | 354.0077088820246 |
| SE_219_2_17 | SE_219 | SE_219_2 | GCTGGTGAGAGGGAGGGCGT | 3549.369156529179 | 3150.0851115660407 | 3288.464452573032 | 3082.799811782803 | 2752.2427605328103 | 2237.6790137901 | 3302.736920574703 | 2958.0110719503027 | 2946.3351518239133 | 1925.3073412878234 | 2536.5493087122245 | 2317.879982417715 |
| SE_219_2_20 | SE_219 | SE_219_2 | TGGCCGGCTCTGCCAACAGA | 60.69150313454474 | 56.96537162413208 | 43.451181955919935 | 22.12224432262881 | 85.90815553108195 | 100.4919727447354 | 78.55972473181926 | 120.0598933006355 | 33.74344055735645 | 89.6678542137078 | 131.46391604616804 | 30.177706330926686 |
| SE_219_2_22 | SE_219 | SE_219_2 | GCATCAGATCAGCCATCTGT | 6068.066536612786 | 5440.692686346931 | 5178.590867655549 | 5708.840343727798 | 5740.998097404278 | 4979.550511523614 | 6276.168693643424 | 5640.924278148756 | 5186.722007776817 | 4772.07946083684 | 5932.309211583333 | 4690.311972433644 |
| SE_219_2_30 | SE_219 | SE_219_2 | AGTGGGAGAACAACCCGTTG | 2034.249131848937 | 2013.7758565373008 | 1895.06404939569 | 1915.5260966417413 | 1819.980183843662 | 1483.1229080947157 | 2009.1918640315969 | 2086.3951536575005 | 1955.3435817710238 | 1629.3305216881051 | 1518.8777443191202 | 1795.5735266901377 |
| SE_219_2_6 | SE_219 | SE_219_2 | GGCTAGATGCTGCCTATTAG | 377.154340907528 | 416.7466660923347 | 464.1376254382357 | 476.27890718130254 | 574.8422258993385 | 437.4866054835464 | 385.2654993697438 | 615.4251223520764 | 410.24919835522843 | 503.7437988753828 | 673.7525697366112 | 629.0891088985486 |
| SE_219_3_18 | SE_219 | SE_219_3 | GAATGCTCCACCAAGAACAC | 599.3285934536293 | 595.6379208418022 | 573.7531071906701 | 599.9032136901106 | 671.3563265577145 | 686.9839171256481 | 626.3254766290248 | 421.6276567880585 | 612.7098416993672 | 890.117479633636 | 622.1060312899024 | 1056.219721582434 |
| SE_219_3_34 | SE_219 | SE_219_3 | GTGTGTTGATGCCAAAGAGG | 370.6516798573982 | 397.7582088842907 | 395.99827191645215 | 283.68525072547527 | 410.4500764262804 | 470.40638965854595 | 297.0203291230427 | 501.98270348533424 | 461.7523444690883 | 281.39668070318055 | 346.26656458588906 | 343.561579767473 |
| SE_219_3_35 | SE_219 | SE_219_3 | TGTGTTGATGCCAAAGAGGG | 1016.5826775036244 | 967.4119251256116 | 949.0133149917968 | 988.9944520704644 | 989.5346803765365 | 989.3261454697227 | 1018.0479396753565 | 1092.8286350829499 | 1062.030392278903 | 1406.2543965710759 | 1334.5935048615454 | 934.3482152459993 |
| SE_219_3_36 | SE_219 | SE_219_3 | GAGGGGAAGTGGTTTCCCCA | 2355.0470769886733 | 2670.3767189417704 | 2508.318231091742 | 2620.8352979867304 | 2385.2770591284357 | 2253.272595767731 | 2558.033776541567 | 2312.3346379004283 | 2600.020893472097 | 3193.7794252866165 | 2977.892455438646 | 2664.923605223372 |
| SE_219_3_9 | SE_219 | SE_219_3 | GTCATGGTAGGTGTGTATGG | 3861.496886935409 | 3433.9125772020675 | 3581.7599307754913 | 3870.0914479704747 | 3544.506861541677 | 3621.1762592499485 | 3542.7207372212197 | 3653.791240999655 | 3245.5861904509957 | 3306.775501734785 | 3661.0353048928405 | 3982.2965546695946 |
| SE_219_4_15 | SE_219 | SE_219_4 | GTGAAGGCAGAGGTAGAGAA | 1000.3260248782999 | 1145.303787390445 | 1187.9948157493566 | 1165.9724066514948 | 1082.8669974967243 | 1256.1496593091927 | 1131.0448040156446 | 1137.2602491390905 | 944.8163356059806 | 950.625056054268 | 1383.8924733788583 | 793.9058127059175 |
| SE_219_4_16 | SE_219 | SE_219_4 | ACTGAATTAACAGGATTTGG | 356.5625809154503 | 504.69320474011755 | 453.2748299492557 | 432.03441853604494 | 392.4199697098805 | 333.52939229933736 | 419.7026389782125 | 325.20160075132765 | 515.9194464164236 | 449.0682780133659 | 379.1325435974311 | 318.02659748745816 |
| SE_219_4_17 | SE_219 | SE_219_4 | AGAAGTTTCCCAGCTTCACA | 62.859056817921335 | 128.9216305177726 | 107.6404280271653 | 55.95626505135522 | 101.81707322202305 | 161.133680435524 | 45.19874573611519 | 134.24019565897828 | 47.95120500255917 | 37.17935418617152 | 194.84830413985622 | 56.873369623669525 |
| SE_219_4_18 | SE_219 | SE_219_4 | TGAGGTCCGTGTGAAGGCAG | 15260.661707812937 | 13651.701340099022 | 13345.438021643227 | 13787.363446956013 | 13667.881485543865 | 12775.475190229423 | 14244.061870552874 | 13401.33108212448 | 13173.261596536393 | 11897.393339574888 | 12128.720040223701 | 10852.367469006327 |
| SE_219_4_23 | SE_219 | SE_219_4 | TATTCATAGTCCACACCATA | 153.89631151973845 | 200.87788941141312 | 192.56773821373608 | 163.96486968536647 | 160.14977142214042 | 308.4063991131535 | 193.70891029763655 | 190.96140509234937 | 308.1308914053339 | 310.55695849625624 | 218.3240034338148 | 313.38387343654637 |
| SE_219_5_12 | SE_219 | SE_219_5 | GAAAACAGAAGACTCTAGGG | 21.675536833765978 | 24.984812115847404 | 19.75053725269088 | 16.917010364363207 | 28.63605184369398 | 0.0 | 5.380803063823238 | 2.836060471668555 | 38.1833669464823 | 22.599215289633673 | 62.210603128990236 | 0.0 |
| SE_219_5_17 | SE_219 | SE_219_5 | GAGTGACTGCTCATGAGTAA | 2078.683982358157 | 2088.730292884843 | 2108.3698517247517 | 2253.8663039290054 | 2130.7343760733784 | 1796.7271678670797 | 2136.178816337825 | 2005.0947534696684 | 2117.84488761303 | 2417.387029045976 | 1834.6258998228632 | 2134.492382406699 |
| SE_219_5_2 | SE_219 | SE_219_5 | GATACAACTGCTGCTTGCTA | 1185.651864806999 | 1163.2928521138551 | 1137.6309457549946 | 1182.889417015858 | 1316.1977902971937 | 1393.0266566684013 | 1188.0813164921708 | 1127.8067142335287 | 1265.3790209008669 | 1171.5141603368165 | 1421.453592249192 | 1627.274779844585 |
| SE_219_5_29 | SE_219 | SE_219_5 | CCAAGAATTTCCAGATAGGG | 35.76463577571386 | 43.97326932389143 | 65.1767729338799 | 44.24448864525762 | 44.54496953463509 | 44.18181560328885 | 31.208657770174778 | 23.633837263904624 | 101.23032167206935 | 54.67552086201695 | 30.518409082146157 | 55.71268861094157 |
| SE_219_5_9 | SE_219 | SE_219_5 | GCCCCCTTCCCTCTAGCCCG | 41.18351998415536 | 70.95686640900662 | 66.16429979651446 | 62.46280749918722 | 97.57469517110542 | 12.128341538157722 | 55.960351863761666 | 114.38777235729839 | 59.49501361428638 | 166.94259036535843 | 105.64064682281361 | 34.820430381838484 |

| | | | | | | | | | | | | | | | |
|---|---|---|---|---|---|---|---|---|---|---|---|---|---|---|---|
| SE_219_6_10 | SE_219 | SE_219_6 | GAGGAGATCGCAGACAGCCG | 3237.2414261229487 | 3394.9362703013453 | 3532.383587643764 | 3284.502627665595 | 3620.869666458195 | 3616.8447087006066 | 3352.240308761877 | 3179.2237887404503 | 3341.488600456114 | 3622.4355088448297 | 3557.742227999423 | 4051.9374154332713 |
| SE_219_6_17 | SE_219 | SE_219_6 | GATGGAGAAGAGAGGAGTGA | 2277.015144387116 | 2245.634912972365 | 2335.5010301306966 | 2438.652109447434 | 2481.791159786812 | 2359.8287392815455 | 2544.0436885756267 | 2772.7217878012907 | 2275.90626706591 | 2843.1270848248814 | 2576.457997511954 | 2305.1124912777077 |
| SE_219_6_27 | SE_219 | SE_219_6 | TTGACCTCAGTAATACCCAG | 199.414938870647 | 184.8876096572708 | 209.35569487852334 | 214.71590077845607 | 165.45274398578746 | 170.66309164407653 | 178.6426617189315 | 133.2948421684221 | 265.50759806972576 | 88.20984032405401 | 136.15905590495976 | 618.642979783997 |
| SE_219_6_5 | SE_219 | SE_219_6 | CGTGTTAAATGTCAGTCGGG | 5086.164718043186 | 6284.179943377939 | 6681.606752585325 | 5878.010447371431 | 6230.992762285265 | 6127.411407099255 | 5142.97156840225 | 5805.415785505532 | 6205.241121442286 | 6908.798816124462 | 5740.982262337571 | 5911.348397823446 |
| SE_219_6_9 | SE_219 | SE_219_6 | CTGTCAAGAAAGAGTCACCG | 376.07056406583973 | 488.70292498597524 | 465.12515230087024 | 389.09123838035373 | 556.8121191829385 | 545.7753692170975 | 476.7391514547388 | 532.2340151831322 | 810.73055865438 | 325.13709739279415 | 735.9631728656015 | 672.0343063694827 |
| SE_22_1_14 | SE_22 | SE_22_1 | GGCACAGCCAACAAGATGAA | 505.0400082267473 | 616.625163019114 | 617.20428914659 | 501.00376848306416 | 719.0830796305378 | 486.86628174604573 | 504.7193273866197 | 529.3979547114636 | 718.3800897605623 | 731.9229726062002 | 481.2518355261509 | 432.93401774752516 |
| SE_22_1_15 | SE_22 | SE_22_1 | CAAAGCTCCGTTCATCTTGT | 1847.8395150785495 | 1653.9945620690983 | 1660.0326560886685 | 1738.548142060711 | 1624.8307935014511 | 1699.7004355618178 | 1928.4798180742482 | 1430.319831211508 | 1720.027483147354 | 2017.162216336012 | 1480.1428404840885 | 1671.3806583282471 |
| SE_22_1_16 | SE_22 | SE_22_1 | CAACAAGATGAACGGAGCTT | 2101.4432960336117 | 2103.7211801543513 | 2036.2803907524299 | 2119.8315295036664 | 2157.2492388916135 | 2024.5667267624713 | 2157.702028593118 | 2449.4108940310753 | 1880.7528184337095 | 2811.050779252498 | 2209.0633035615024 | 1887.2673266956458 |
| SE_22_1_17 | SE_22 | SE_22_1 | TAATGGCATCTGGGTCTGGT | 14673.254659617878 | 14800.003304943368 | 14640.085738557116 | 14510.890967154932 | 14765.5968062188 | 14835.560631496499 | 14047.124478416943 | 14279.564474851175 | 15101.077634694837 | 13020.064034608304 | 13510.264943673163 | 13785.408388169855 |
| SE_22_1_18 | SE_22 | SE_22_1 | ATCTTAATGGCATCTGGGTC | 242.76601253817896 | 411.7497036691652 | 349.5845093726286 | 356.55852614119374 | 349.99618920070424 | 303.20853845394305 | 279.80175931880837 | 501.98270348533424 | 553.2148280850807 | 450.5262919030197 | 428.4315121147441 | 442.21946584934875 |
| SE_22_2_15 | SE_22 | SE_22_2 | GCTGAGGTGTTTCTAATGTG | 1984.3953971312753 | 2156.688981839948 | 2069.8563040820045 | 2006.6176909113894 | 2138.158537662484 | 2151.048002803259 | 1942.4699060401888 | 2025.8925302619045 | 1865.6570687106816 | 1869.1738065361528 | 2432.082446854109 | 1758.4317342828433 |
| SE_22_2_16 | SE_22 | SE_22_2 | GTGAGGTTCTGTTAGCCTGT | 1150.9710058729734 | 1118.3201903053298 | 1117.880408502304 | 1210.2168952967525 | 992.7164639147247 | 959.8716017341968 | 1147.1872132071142 | 1182.6372166857875 | 989.2155994972392 | 1197.7584103505847 | 1482.4904104134844 | 1117.7358152570152 |
| SE_22_2_4 | SE_22 | SE_22_2 | ACCTCACTGAATCTGCTCAA | 910.3725470181711 | 1084.3408458277775 | 1063.5664310574039 | 912.2172511860467 | 844.2332321326078 | 920.8876467901184 | 773.7594805777816 | 1106.0635839507365 | 956.3601442177079 | 801.178632364755 | 1150.3092654039704 | 602.3934456058057 |
| SE_22_2_6 | SE_22 | SE_22_2 | GAATGGAGTGAGGAACATGA | 740.2195828731082 | 836.4915096385711 | 790.0214901076353 | 771.6759343128755 | 804.9912351616197 | 934.7486085480131 | 717.7991287140198 | 614.4797688615203 | 1016.7431431098194 | 852.9381254474644 | 562.242998090308 | 812.4767089095646 |
| SE_22_2_8 | SE_22 | SE_22_2 | GCTGCTGACAGAGAAGACTA | 729.3818144562251 | 843.4872570310084 | 902.5995524479732 | 904.4094002486484 | 1071.2004578567007 | 1001.4544870078805 | 877.0708994031877 | 1092.8286350829499 | 713.9401633714365 | 946.2510143853067 | 1099.8365119219595 | 670.8736253567548 |
| SE_22_3_10 | SE_22 | SE_22_3 | CTAGCCGTCTGGAAGACATG | 910.3725470181711 | 778.5267455298051 | 860.1358973546878 | 796.4007956146371 | 769.9916162415493 | 778.8127887716995 | 842.633759794719 | 855.5449089533474 | 641.1253705897726 | 799.7206184751012 | 637.3652358309755 | 709.176098776777 |
| SE_22_3_19 | SE_22 | SE_22_3 | CAGCTCTGCGGCTAATGCAG | 450.8511661423323 | 603.6330607188733 | 625.1045040476664 | 484.08675811870097 | 659.689786917691 | 617.6791083361754 | 421.8549602037418 | 628.6600712198631 | 412.91315418870397 | 1048.3119866610716 | 524.6818792199742 | 466.59376711663566 |
| SE_22_3_20 | SE_22 | SE_22_3 | TCAGCTCTGCGGCTAATGCA | 618.8365766040187 | 690.5802068820223 | 687.3186963936427 | 685.7895740014931 | 766.809832703361 | 680.0534362467008 | 549.9180731227349 | 883.905513670033 | 555.8787839185563 | 636.4230628338773 | 885.0338633822385 | 573.3764202876071 |
| SE_22_3_24 | SE_22 | SE_22_3 | GACAGATGTCCACTTCACAA | 938.5507449020669 | 1049.362108865591 | 1079.3668608595567 | 927.8329530608435 | 935.4443602273368 | 601.2192162486757 | 726.4084136161371 | 1099.446109516843 | 903.9690128260229 | 605.0757642063209 | 958.982316158208 | 582.6618683894307 |
| SE_22_3_9 | SE_22 | SE_22_3 | AAGTGGCCAGAGGGCAAGAG | 684.9469639470049 | 798.5145952224831 | 752.4954693275225 | 805.5099550416019 | 708.4771345032437 | 916.5560962407765 | 820.0343869266613 | 400.82987999582247 | 849.8019108786875 | 793.8885629164861 | 490.64211524373434 | 973.8113696787495 |
| SE_22_4_15 | SE_22 | SE_22_4 | GAGAATGCTTTGGGAATGGG | 658.9363197464858 | 744.5474010522527 | 730.7698783495625 | 711.8157437928211 | 671.3563265577145 | 694.7807081144638 | 708.113683199138 | 683.4905736721217 | 792.0828678200514 | 591.2246322546099 | 666.7098599484237 | 775.3349165022702 |
| SE_22_4_18 | SE_22 | SE_22_4 | GTGAGTCTATGAAAACCCAT | 346.80858934025565 | 399.75699385355847 | 336.74666015837954 | 322.7245054124673 | 326.6631099206573 | 444.4170863624937 | 372.351572016568 | 429.190484712508 | 333.88246446226384 | 473.85451413748024 | 544.636223619839 | 396.95290635295873 |
| SE_22_4_8 | SE_22 | SE_22_4 | AGATGCCATGAACTCAGGAG | 962.3938354192094 | 1100.3311255819197 | 1122.8180428154765 | 988.9944520704644 | 1092.412348111289 | 973.7325634920915 | 936.2597331052433 | 1007.7468209328932 | 802.7386911539535 | 1023.5257505369573 | 1086.9248773102822 | 1028.363377276963 |
| SE_22_5_12 | SE_22 | SE_22_5 | GGGAACCTGACAGGGCTGGG | 254.68755779675024 | 367.7764343452738 | 379.2103152516649 | 299.3009526002721 | 293.7846800260457 | 466.0748391092039 | 224.91756806781132 | 249.57332150683285 | 347.2022436296414 | 466.5644446892113 | 184.28423945757487 | 555.9662050966878 |
| SE_22_5_15 | SE_22 | SE_22_5 | ACTGACCCCACAGTATCTGG | 1041.5095448624552 | 1261.233315607977 | 1150.4687949692438 | 1076.1821208714132 | 1286.5011439407704 | 1127.9357630486682 | 1015.8956184498272 | 1018.1457093290112 | 1083.3420389467071 | 1293.9873270677344 | 807.5640557121752 | 1344.0686127389654 |
| SE_22_5_32 | SE_22 | SE_22_5 | ATAGGTGAAGAGCAGCCCTG | 1355.8048289520618 | 1491.093587073773 | 1322.2984690676544 | 1354.6621376386229 | 1455.135671464746 | 1522.9731731486625 | 1461.4261121343914 | 1399.1231660231538 | 1456.2958556332785 | 1090.5943894610314 | 1504.792324742745 | 1320.8549924844065 |
| SE_22_5_33 | SE_22 | SE_22_5 | GGGCTGCTCTTCACCTATCG | 175.57184835350444 | 151.90765766435223 | 175.77978154894885 | 219.92113473672168 | 233.33079280046948 | 172.39571186381335 | 198.01355274869513 | 249.57332150683285 | 238.86803973497067 | 192.45783343429966 | 115.03092654039705 | 399.2742683784146 |
| SE_22_5_6 | SE_22 | SE_22_5 | AGCTCAACACTGCCACCTGG | 1490.193157321411 | 1684.975729092749 | 1610.6563129569413 | 1553.7623365422821 | 1623.7701989887216 | 1538.5667551262939 | 1694.9529651043197 | 1677.0570922466723 | 1479.3834728567329 | 2095.1659594324897 | 1401.4992478493273 | 1839.6794051737998 |
| SE_22_6_14 | SE_22 | SE_22_6 | TCTACCTTCTTGGTAGTACC | 81.28326312662242 | 46.97144677779312 | 60.23913862070719 | 61.16149900962082 | 62.575076251035 | 2.598930329605226 | 68.87427921693744 | 19.852423301679885 | 16.871720278678225 | 45.92743752409424 | 25.823269223354437 | 55.71268861094157 |
| SE_22_6_19 | SE_22 | SE_22_6 | CCTGGAGACCTCTCATCTAG | 773.8166649654454 | 896.4550587166049 | 851.2481555909769 | 713.1170522823875 | 853.7785827471724 | 889.7004828348557 | 843.7099204074837 | 804.4958204633134 | 999.8714228311412 | 2477.894605466608 | 1058.754038157532 | 786.9417266295497 |
| SE_22_6_4 | SE_22 | SE_22_6 | CTATGTGACTGAGTAAATGT | 264.4415493719449 | 315.8080251443112 | 298.2331125156323 | 245.94730452804967 | 301.2088416151515 | 156.80212988618197 | 218.46060439122343 | 464.16856386308683 | 189.14086417676117 | 252.96540985493175 | 497.6848250319219 | 265.7959519147004 |
| SE_22_6_5 | SE_22 | SE_22_6 | CAACCAGCTAGATTGAAAAG | 209.16893044584168 | 311.8104552057756 | 302.1832199661705 | 242.04337905935049 | 308.63300320425736 | 230.43848922499672 | 209.85131948910626 | 306.29453094020397 | 262.8436422362503 | 245.67534040666283 | 241.7997027277734 | 255.34982280014887 |
| SE_22_6_9 | SE_22 | SE_22_6 | GCATCAAGTAAGGGAAGTAG | 3327.1949039830774 | 3354.9605709159896 | 3319.0777853147024 | 3293.6117870925596 | 3450.1139499087603 | 3138.6415280532447 | 2964.8224881666038 | 3204.748332985467 | 3631.8597863049445 | 3062.558175217776 | 3805.410855550686 | 3419.366263496539 |
| SE_22_8_13 | SE_22 | SE_22_8 | TCTCTGGTTGAGAACCAACA | 3.2513305250648967 | 6.9957473924372735 | 4.93763431317272 | 3.9039254686992013 | 0.0 | 0.0 | 0.0 | 0.0 | 0.0 | 0.0 | 9.390279717583432 | 0.0 |
| SE_22_8_2 | SE_22 | SE_22_8 | GCTTCCTTGTCAGTAAAACA | 98.6236925936352 | 162.90097499532507 | 164.91698605996885 | 100.20075369661284 | 132.57431409117584 | 35.51871450460476 | 76.40740350628997 | 121.95060028174787 | 96.7903952829435 | 61.23658336545898 | 91.55522724643846 | 135.79967848917008 |
| SE_22_8_4 | SE_22 | SE_22_8 | TACCCTTTCTTGTCAAACCA | 60.69150313454474 | 132.9192004563082 | 92.82752508764715 | 66.36673296788642 | 108.18064029839948 | 178.4598826328922 | 26.904015319116187 | 98.31676301784324 | 198.90870223283804 | 138.5113195171096 | 97.42415206992811 | 220.52939241831038 |
| SE_22_8_8 | SE_22 | SE_22_8 | GATGCCCAGACAATCTCAAA | 703.371170255706 | 868.4720691468558 | 760.395684228599 | 779.4837852502739 | 709.5377290159731 | 1006.6523476670909 | 874.9185781776584 | 960.479146405084 | 1070.9102450571547 | 881.3693962957133 | 944.8968965818328 | 812.4767089095646 |
| SE_22_8_9 | SE_22 | SE_22_8 | AGATGCCCAGACAATCTCAA | 1200.8247405906352 | 1439.1251778728106 | 1331.1862108313653 | 1320.8281169098964 | 1387.257622650064 | 1835.711122811158 | 1286.0119322537537 | 1660.040729416661 | 1237.8514772882868 | 1805.0211953913863 | 1282.9469664148364 | 1296.4806912171196 |
| SE_220_1_14 | SE_220 | SE_220_1 | GTATTTTGCATGTTACTCAT | 37.93218945909046 | 47.97083926242702 | 60.23913862070719 | 49.44972260352321 | 51.96913112374093 | 98.75935252499859 | 29.056336544645482 | 21.743130282792254 | 62.15896944776188 | 64.88161808959345 | 14.085419576375148 | 39.46315443275028 |
| SE_220_1_15 | SE_220 | SE_220_1 | GCCCAGCAGATGCAACCTGT | 2077.600205516469 | 1768.9246978019962 | 1801.2489974454083 | 2186.198262471553 | 2016.1901686986023 | 1911.9464124795782 | 2174.9205983973525 | 2829.4429972346616 | 2021.9424776079115 | 2590.8906819147765 | 3320.6376651304413 | 2569.74776217968 |
| SE_220_1_2 | SE_220 | SE_220_1 | GTTTGCCAATGTGGAGACCA | 120.29922942740117 | 161.90158251069118 | 147.14150253254707 | 123.62430650880805 | 156.9679878839522 | 125.61496593091927 | 77.48356411905462 | 173.94504226233803 | 120.76599778422309 | 303.9958959928142 | 167.85124995180385 | 89.37243798005211 |
| SE_220_1_3 | SE_220 | SE_220_1 | CTTCAGACTAGTCCGTCACA | 279.6144251555811 | 288.824428059196 | 205.40558742798515 | 304.5061865585377 | 295.9058690515045 | 169.79678153420812 | 293.79184728474877 | 279.82463320463074 | 251.29983362452305 | 287.9577432066226 | 208.93372371623136 | 196.15509115102344 |
| SE_220_1_6 | SE_220 | SE_220_1 | GGGCTTTCTCCGAAAATAAG | 2.167553683376598 | 3.9975699385355847 | 5.925161175807264 | 0.0 | 0.0 | 0.0 | 0.0 | 3.78141396222474 | 18.647690834328564 | 0.0 | 0.0 | 1.1606810127279494 |
| SE_220_2_11 | SE_220 | SE_220_2 | TCTGAAACGCCCTTAACGTG | 966.7289427859627 | 973.4082800334148 | 1050.7285818431549 | 851.0557521764259 | 895.1417687436193 | 769.2833775631469 | 794.2065322203098 | 828.129657727218 | 938.6004386612044 | 933.8578963232495 | 659.6671501602361 | 770.6921924513584 |
| SE_220_2_14 | SE_220 | SE_220_2 | GACACGTATAGAGAAAGAAG | 666.5227576383038 | 814.5048749766254 | 838.4103063767279 | 746.9510730111139 | 755.1432930633376 | 685.2512969059113 | 707.0375225863734 | 850.8181415005665 | 695.292472537108 | 914.1747088129234 | 717.1826134304347 | 587.3045924403424 |
| SE_220_2_31 | SE_220 | SE_220_2 | TCCTCTCCTATAAAAAAAAG | 0.0 | 0.9993924846338962 | 2.962580587903632 | 0.0 | 0.0 | 0.0 | 0.0 | 0.0 | 0.0 | 0.0 | 0.0 | 0.0 |
| SE_220_2_36 | SE_220 | SE_220_2 | GCCATACTGCTTCTTAGGCA | 1838.085523503355 | 1827.888854395396 | 1883.2137270440755 | 1673.482717582391 | 1931.3426076802498 | 1668.5132716065552 | 1826.2445598616068 | 2199.8375725242427 | 1721.8034537030042 | 1598.7122300053757 | 2364.002918901629 | 2171.6341748139935 |
| SE_220_2_6 | SE_220 | SE_220_2 | TTTTAGAGCGAGAAAGAAGA | 2.167553683376598 | 6.9957473924372735 | 1.975053725269088 | 7.807850937398403 | 2.1211890254588135 | 0.0 | 2.152321225529295 | 0.945353490556185 | 0.0 | 6.561062503442034 | 0.0 | 0.0 |
| SE_220_3_23 | SE_220 | SE_220_3 | AATTCAAGCTTCCGGCCACA | 46.60240419259685 | 52.9678016855965 | 58.264084895438096 | 40.34056317655841 | 64.69626527649382 | 51.11229648223612 | 32.28481838293943 | 81.30040018783191 | 68.37486639253807 | 33.53431946203706 | 104.46686185811568 | 3.482043038183848 |
| SE_220_3_25 | SE_220 | SE_220_3 | GAATATTCTCAGGTTCCGTG | 15.172875783636185 | 18.98845720804403 | 25.675698428498144 | 18.218318853929606 | 15.9089176909411 | 58.90908747105179 | 27.980175931880833 | 10.398888396118036 | 27.527543612580263 | 7.290069448268927 | 3.521354894093787 | 22.05293924183104 |
| SE_220_3_4 | SE_220 | SE_220_3 | TTCCAAAAGTACAGGCCTGG | 9.75399157519469 | 12.992102300240651 | 22.713117840594514 | 5.205233958265602 | 16.969512203670508 | 24.256683076315444 | 2.152321225529295 | 1.89070698111237 | 27.527543612580263 | 4.374041668961356 | 0.0 | 1.1606810127279494 |
| SE_220_3_5 | SE_220 | SE_220_3 | GCAGCTAATGGCAATACTAG | 1867.347498228939 | 1673.9824117617761 | 1905.92684488467 | 1833.5436617990583 | 1966.3422266003201 | 1813.1870599545796 | 1892.966517853015 | 2312.3346379004283 | 2027.2703892748625 | 1993.8339941015515 | 1565.8291429070373 | 2423.5019545759583 |
| SE_220_4_10 | SE_220 | SE_220_4 | TCTGTATTCCTTCTTAGGGA | 76.94815575986922 | 75.95382883217611 | 101.71526685135804 | 98.89944520704643 | 55.15091466192915 | 95.29411208552496 | 39.817942672291956 | 70.90151179171387 | 58.607028336461205 | 233.28222234460566 | 96.25036710523018 | 54.55200759821362 |
| SE_220_4_18 | SE_220 | SE_220_4 | GTATCTTCATATATAACTGG | 4.335107366753196 | 10.993317330972857 | 13.825376076883616 | 0.0 | 12.727134152752882 | 40.71657516381521 | 1.0761606127646475 | 3.78141396222474 | 13.319779167377547 | 3.6450347241344634 | 0.0 | 1.1606810127279494 |
| SE_220_4_19 | SE_220 | SE_220_4 | GGTATCTTCATATATAACTG | 3.2513305250648967 | 2.9981774539016888 | 11.850322351614528 | 1.3013084895664004 | 7.4241615891058474 | 0.0 | 0.0 | 0.0 | 12.431793889552377 | 0.7290069448268927 | 0.0 | 0.0 |
| SE_220_4_2 | SE_220 | SE_220_4 | AGTTAATTTGAGGATCACTG | 65.02661050129794 | 77.9526138014439 | 74.06451469759081 | 44.24448864525762 | 67.87804881468203 | 44.18181560328885 | 78.55972473181926 | 62.393330376708214 | 28.415528890405433 | 8.748083337922711 | 53.994108376104734 | 13.928172152735392 |
| SE_220_5_2 | SE_220 | SE_220_5 | CTAAATAGGTGGAGACACGT | 891.94834070947 | 776.5279605605373 | 740.645146975908 | 827.6321993642307 | 721.2042686559965 | 745.8930045966999 | 901.8225934967746 | 849.8727880100104 | 697.9564283705835 | 579.5605211373796 | 906.1619927468012 | 849.618501316859 |
| SE_220_5_20 | SE_220 | SE_220_5 | TTAGCCAGCAAGTAAGAAAA | 291.5359704141524 | 395.75942391502286 | 380.1978421142995 | 359.1611431203265 | 421.05602155357445 | 508.5240344927559 | 247.51694093586892 | 384.7588706563673 | 373.8418019643965 | 469.4804724685189 | 262.9278320923361 | 316.86591647473017 |
| SE_220_5_23 | SE_220 | SE_220_5 | GCAAGTAAGAAAATGGGTGA | 1687.4405425086813 | 2119.711459908494 | 2042.2055519282371 | 2014.4255418487878 | 2150.885671815237 | 2213.4223307137845 | 1869.2909843721927 | 2135.553535166422 | 2284.786119844162 | 2310.223008156423 | 2187.93517419694 | 2514.0350735687384 |
| SE_220_5_24 | SE_220 | SE_220_5 | ATGCAGGAGAAAGAAGTCAG | 1375.3128121024513 | 1472.1051298657292 | 1516.8412610066596 | 1584.9937402918758 | 1337.409680551782 | 1458.8662250184002 | 1378.5617449515134 | 1395.3417520609291 | 1231.6355803435106 | 1538.2046535847435 | 1574.045637659923 | 1578.5261773100112 |
| SE_220_5_3 | SE_220 | SE_220_5 | GCATGAACCAGGCATAAACC | 5306.171416905911 | 5159.863398164806 | 5429.4226907647235 | 5505.83621935544 | 5047.369286079247 | 4464.962306261778 | 5571.28349228258 | 5271.291063341288 | 5382.966754176179 | 5109.60967629169 | 6356.045583839285 | 5795.280296550652 |
| SE_221_1_1 | SE_221 | SE_221_1 | AAAAGATCACTCCAGCTGCA | 40.09974314246706 | 8.994532361705065 | 20.738064115325425 | 42.943180155691216 | 8.484756101835254 | 207.0481162585497 | 72.10276105523138 | 11.34424188667422 | 133.19779167377547 | 1.4580138896537853 | 9.390279717583432 | 61.51609367458132 |
| SE_221_1_13 | SE_221 | SE_221_1 | GGCACTGTGACAATAAATGC | 1044.7608753875202 | 1174.286169444828 | 1051.7161087057893 | 1033.238940715722 | 1154.987424362324 | 933.0159883282762 | 1111.6739129858809 | 1014.3642953667866 | 1062.030392278903 | 759.6252365096221 | 1253.6023422973883 | 1037.6488253787868 |
| SE_221_1_14 | SE_221 | SE_221_1 | GCAACGCTGCAGGGGAGTCA | 1809.9073256194592 | 486.7041400167074 | 875.9363271568405 | 1568.0767299275126 | 424.2378050917627 | 3348.2885746414 | 2243.79487761429 | 919.828946311168 | 3078.644958219864 | 128.3052222895331 | 489.4683302790364 | 747.4785721967994 |
| SE_221_1_17 | SE_221 | SE_221_1 | GCCTCCGATGTAGCAGGCAT | 1439.255645762061 | 1465.1093824732918 | 1381.550080825727 | 1501.709996959626 | 1433.9237812101578 | 1749.9464219341855 | 1569.042173410856 | 1549.4343710215871 | 1458.0718261889288 | 1528.727563301994 | 2011.8674294922503 | 1698.07632162099 |
| SE_221_1_3 | SE_221 | SE_221_1 | TTTGCTTTAATAAAGAGAAG | 9.75399157519469 | 7.9951398770711695 | 3.950107450538176 | 6.506542447832002 | 10.605945127294067 | 0.0 | 2.152321225529295 | 0.0 | 0.0 | 0.0 | 2.347569929395858 | 0.0 |
| SE_222_1_0 | SE_222 | SE_222_1 | ATCTTGTTCTGACAAACCAT | 74.78060207649263 | 95.94167852485404 | 81.96472959866715 | 93.69421124878083 | 106.05945127294068 | 180.19250285262902 | 120.52998862964051 | 89.80858160283758 | 73.70277805948909 | 251.50739596527796 | 188.97937931636656 | 109.10401519642724 |
| SE_222_1_1 | SE_222 | SE_222_1 | AGTGGGTTCTCCTAAGACAT | 6.5026610501297935 | 0.9993924846338962 | 0.0 | 0.0 | 14.848323178211695 | 0.0 | 5.380803063823238 | 4.726767452780925 | 0.0 | 0.0 | 8.216494752885502 | 0.0 |
| SE_222_1_12 | SE_222 | SE_222_1 | GCTTGACTGAAGGCTCTAGA | 542.9721976858377 | 750.543755960056 | 734.7199858001007 | 645.4490108249346 | 610.9024393321383 | 877.5721412966981 | 517.6332547397955 | 607.8622944276269 | 792.9708530978767 | 649.5451878407614 | 773.5242917359352 | 812.4767089095646 |
| SE_222_1_3 | SE_222 | SE_222_1 | GATTAAGGCCTATAACCCTG | 141.97476626116716 | 196.88031947287755 | 182.69246958739063 | 283.68525072547527 | 184.54344521491677 | 251.22993186183854 | 112.99686434028798 | 90.75393509339376 | 286.81924473752986 | 147.2594028550323 | 123.24742129328254 | 205.44053925284706 |
| SE_222_1_36 | SE_222 | SE_222_1 | TGTCCCCAGAAGCAGTGCAC | 1415.4125552449184 | 1173.2867769601942 | 1115.9053547770347 | 1353.3608291490564 | 1146.5026682604887 | 1262.2138300782715 | 1409.7704027216882 | 813.0040018783191 | 1131.2932439492663 | 1211.6095423022955 | 993.022080134448 | 1367.2822329935245 |
| SE_222_2_1 | SE_222 | SE_222_2 | CGATATCCTTTTCTGTACAG | 1006.8286859284296 | 1146.303179875079 | 1143.556106930802 | 1037.142866184421 | 1077.5640249330772 | 919.1550265703817 | 939.4882149435373 | 1235.5770121569337 | 1242.2914036774125 | 1183.9072783988736 | 1108.053006674845 | 1470.582843126312 |
| SE_222_2_11 | SE_222 | SE_222_2 | GTATGAGTGAATAACAAGGC | 201.5824925540236 | 310.8110627211417 | 279.470102125576 | 219.92113473672168 | 321.3601373570102 | 258.1604127407858 | 196.93739213593048 | 203.25100046957976 | 223.77229001194277 | 277.0226390342192 | 186.63180938697073 | 162.4953417819129 |
| SE_222_2_2 | SE_222 | SE_222_2 | AAAGTTGTACCATCTCAGGC | 87.78592417675222 | 147.91008772581662 | 117.51569665351074 | 102.80337067574564 | 138.9378811675523 | 102.22459296447224 | 64.56963676587885 | 96.42605603673087 | 221.1083341784673 | 53.94651391719005 | 124.42120625798047 | 45.26655949639003 |
| SE_222_2_26 | SE_222 | SE_222_2 | TAAGGAGGCATCATGTGCGG | 8982.34246391262 | 9542.19944328444 | 9542.472073637598 | 9443.595708783369 | 9694.894440859507 | 9227.93529031829 | 8653.40748724053 | 9165.202090942214 | 9639.968176070042 | 7386.298364986076 | 9400.843782265714 | 8809.568886605137 |
| SE_223_1_1 | SE_223 | SE_223_1 | AATAGTATAGGAGAAGCCAT | 29.26197472558407 | 39.975699385355846 | 32.58838646693995 | 35.13532921829281 | 46.6661585600939 | 12.994651648026132 | 30.13249715741013 | 27.415251226129364 | 5.327911666951019 | 26.244250013768134 | 21.128129364562724 | 58.034050636397474 |
| SE_223_1_26 | SE_223 | SE_223_1 | CTTAGGCTAAAATACCAGGG | 99.7074694353235 | 112.93135076363026 | 103.69032057662713 | 124.92561499837444 | 181.36166167672854 | 164.59892087499765 | 119.45382801687587 | 247.68261452572048 | 159.83735000853056 | 57.59154864132452 | 185.45802442227279 | 59.19473164912542 |
| SE_223_1_39 | SE_223 | SE_223_1 | AATGGTTTGCTCACTGAGCA | 75.86437891818092 | 129.9210230024065 | 111.59053547770347 | 139.24000838360485 | 150.60442080757576 | 76.23528966841997 | 94.70213392328898 | 178.67180971511897 | 122.54196833987343 | 125.38919451022554 | 100.9455069640219 | 68.48017975094902 |
| SE_223_1_6 | SE_223 | SE_223_1 | ACTGTACTGGAATTAGCCCG | 1096.7821637885586 | 1366.169526494536 | 1280.8223408370036 | 1100.9069821731748 | 1122.1089944677124 | 843.7860470118301 | 941.6405361690665 | 1458.6804359281934 | 1537.102515915369 | 812.8427434819853 | 1605.7378317067669 | 959.8831975260142 |
| SE_223_1_9 | SE_223 | SE_223_1 | CTGTGCTTGAAGCAAAGCCA | 45.51862735090855 | 46.97144677779312 | 44.43870881855448 | 36.43663770785921 | 27.575457330964575 | 24.256683076315444 | 51.655709412703075 | 10.398888396118036 | 22.199631945629246 | 24.78623612411435 | 15.259204541073078 | 15.088853165463343 |
| SE_223_2_12 | SE_223 | SE_223_2 | TGTTATCTCACAGTTTGTGT | 250.35245042999705 | 249.84812115847404 | 237.00644703229057 | 232.9342196323857 | 242.87614341503414 | 257.2941026309174 | 311.0104170889831 | 334.6551356568895 | 181.14899667633463 | 139.2403264619365 | 300.48895096266983 | 246.06437469832528 |

| | | | | | | | | | | | | | | | |
|---|---|---|---|---|---|---|---|---|---|---|---|---|---|---|---|
| SE_223_2_15 | SE_223 | SE_223_2 | TTATCTCTGCTGGTAGAAAA | 8.670214733506391 | 24.984812115847404 | 13.825376076883616 | 16.917010364363207 | 29.69664635642339 | 1.7326202197368175 | 1.0761606127646475 | 10.398888396118036 | 13.319779167377547 | 1.4580138896537853 | 11.73784964697929 | 67.31949873822107 |
| SE_223_2_2 | SE_223 | SE_223_2 | AGTAGTTCTTACATTTTGGG | 18.42420630870108 | 40.97509186998975 | 23.700644703229056 | 41.641871666124814 | 36.06021343279983 | 42.44919538355203 | 35.513300221233365 | 38.75949311280358 | 24.863587779104755 | 46.65644446892113 | 26.997054188052367 | 35.98111139456643 |
| SE_223_2_4 | SE_223 | SE_223_2 | ACACATTTAGCTTTAAACCT | 514.7939998019419 | 728.5571212981104 | 666.5806322783172 | 640.243776866669 | 753.0221040378788 | 692.1817777848586 | 718.8752893267845 | 872.5612717833587 | 540.7830341955284 | 1113.922611695492 | 1140.918985686387 | 735.87176206952 |
| SE_223_2_6 | SE_223 | SE_223_2 | AAAGATGTGAGAGAAATCCA | 224.34180622947787 | 289.82382054382987 | 324.89633780676496 | 222.52375171585447 | 253.4820885423282 | 312.73794966249557 | 293.79184728474877 | 126.67736773452879 | 267.2835686253761 | 204.12194455152994 | 187.80559435166865 | 320.34795951291403 |
| SE_223_3_13 | SE_223 | SE_223_3 | GCAGACAAGGACAGCCCAAT | 2015.8249255402359 | 2037.7612761685143 | 2072.818884669908 | 2236.949293564642 | 1940.8879582948143 | 1800.1924083065533 | 2296.5267476397576 | 2154.4606049775457 | 2120.5088434465056 | 2319.7000984391725 | 2178.544894479356 | 2146.0991925339786 |
| SE_223_3_17 | SE_223 | SE_223_3 | TGTTCAGGGCAACAGCAAGG | 955.8911743690796 | 879.4653864778286 | 886.7991226458205 | 860.1649116033907 | 953.4744669437366 | 889.7004828348557 | 909.3557177861271 | 1008.6921744234494 | 873.7775133799671 | 648.8161808959345 | 1428.4963020373796 | 1164.1630557661333 |
| SE_223_3_19 | SE_223 | SE_223_3 | TTCCTATCCCAGACACTTGG | 5.4188842084414945 | 1.9987849692677924 | 4.93763431317272 | 1.3013084895664004 | 5.3029725636470335 | 0.0 | 1.0761606127646475 | 16.071009339455145 | 0.8879852778251698 | 0.0 | 2.347569929395858 | 0.0 |
| SE_223_3_20 | SE_223 | SE_223_3 | TCTAATGTGGGACAGATGTG | 1154.2223363980384 | 1043.3657539577875 | 942.100626953355 | 1129.5357689436355 | 1087.109375547642 | 994.5240061289333 | 988.991603130711 | 967.0966208389773 | 866.6736311573657 | 1081.8463061231087 | 1271.209116767857 | 984.2574987933011 |
| SE_223_3_34 | SE_223 | SE_223_3 | AATCCTTCCCAGTCTGTAGG | 8.670214733506391 | 9.993924846338961 | 11.850322351614528 | 18.218318853929606 | 18.030106716399914 | 59.7753975809202 | 4.30464245105859 | 0.945353490556185 | 19.535676112153734 | 18.225173620672315 | 0.0 | 78.92630886550056 |
| SE_224_1_1 | SE_224 | SE_224_1 | GTGTGCAAAGAGGTGCCAGA | 1084.8606185299873 | 1110.3250504282587 | 1214.6580410404893 | 1322.129425399463 | 1331.0461134754055 | 1307.2619557914288 | 1171.9389073007012 | 1265.8283238547317 | 1140.173096727518 | 1145.2699103230484 | 1028.2356290753858 | 1297.6413722298475 |
| SE_224_1_24 | SE_224 | SE_224_1 | GCTTGGCTAATGGTGGAAAC | 1633.2517004242663 | 1494.0917645276747 | 1628.431796484363 | 1614.923835551903 | 1339.5308695772408 | 1366.1710432624805 | 1684.1913589766732 | 1588.1938641343909 | 1736.011218148207 | 1904.8951468326704 | 1752.460952294008 | 1169.966460829773 |
| SE_224_1_28 | SE_224 | SE_224_1 | ACTAGAACAGAGGTGCCCTG | 1089.1957258967404 | 1371.1664889177055 | 1351.9242749466907 | 1224.531288681983 | 1222.865473177006 | 1352.310081504586 | 1012.6671366115332 | 1214.7792353646978 | 1282.2507411795452 | 1423.0215563020945 | 1471.926345731203 | 1234.9645975425383 |
| SE_224_1_30 | SE_224 | SE_224_1 | CCTGTCATCCACTAGAACAG | 938.5507449020669 | 972.408887548781 | 992.4644969477167 | 1000.7062284765619 | 856.9603662853607 | 1101.0801496427475 | 895.3656298201867 | 785.5887506521898 | 927.0566300494772 | 668.4993684062606 | 717.1826134304347 | 759.085382324079 |
| SE_224_1_41 | SE_224 | SE_224_1 | GGTTTTGCTTCTGCTCAAAA | 2.167553683376598 | 7.9951398770711695 | 6.912688038441808 | 1.3013084895664004 | 4.242378050917627 | 0.0 | 1.0761606127646475 | 0.0 | 13.319779167377547 | 0.0 | 0.0 | 0.0 |
| SE_224_2_11 | SE_224 | SE_224_2 | GCACCATCTGGGCGTAGGAG | 1726.4565088094603 | 1673.9824117617761 | 1727.1844827478176 | 1819.2292684138279 | 1705.435976468886 | 1563.6897483124778 | 1646.5257375299107 | 1788.608804132302 | 1958.0075376044995 | 2001.1240635498204 | 1934.3976218221871 | 1731.7360709901006 |
| SE_224_2_20 | SE_224 | SE_224_2 | TCCTGGCCACGTGGGCGCAG | 342.47348197350243 | 386.7648915533178 | 395.01074505381763 | 460.66320530650574 | 416.8136435026569 | 304.94115867367987 | 407.86487223780136 | 370.57856829802455 | 324.11462640618697 | 314.20199322039076 | 281.70839152750295 | 271.59935697834015 |
| SE_224_2_3 | SE_224 | SE_224_2 | AAGGTGGGCGGCAGCCCTGA | 288.2846398890875 | 309.81167023650784 | 284.40773643874866 | 309.7114205168033 | 375.45045750621 | 461.74328855986187 | 285.1825623826316 | 307.2398844307601 | 369.40187557527065 | 301.80887515833354 | 385.00146842092073 | 272.76003799106815 |
| SE_224_2_33 | SE_224 | SE_224_2 | CACTCAGCTCGTGCTGGCCG | 1146.6358985062202 | 1126.3153301824009 | 1077.3918071342875 | 990.2957605600308 | 1009.6859761183953 | 856.7806986598563 | 1095.531503794411 | 1427.4837707398394 | 804.5146617096038 | 1300.5483895711766 | 1351.0264943673162 | 748.6392532095274 |
| SE_224_2_6 | SE_224 | SE_224_2 | TGGCTGAAGCGGATGAAGGT | 112.71279153558308 | 137.91616287947767 | 180.71741586212156 | 105.40598765487843 | 147.42263726938754 | 189.7219140611815 | 86.0928490211718 | 214.595242356254 | 164.2772763976564 | 64.88161808959345 | 200.71722896334586 | 135.79967848917008 |
| SE_225_1_16 | SE_225 | SE_225_1 | GGAGGAGGAATGAGTCACAC | 9518.812000548329 | 8798.651434716821 | 8487.793384343906 | 8694.042018793121 | 8222.78925719109 | 8430.929989239354 | 9496.04124703525 | 8440.11596368562 | 8055.80244042994 | 7309.752635779253 | 8748.219341893666 | 7651.209235902643 |
| SE_225_1_22 | SE_225 | SE_225_1 | TCTTTTCCTTCTGGAGAGAG | 11.921545258571289 | 6.9957473924372735 | 10.862795488979984 | 13.013084895664004 | 0.0 | 27.72192351578908 | 9.685445514881827 | 4.726767452780925 | 8.879852778251697 | 2.187020834480678 | 7.042709788187574 | 11.606810127279495 |
| SE_225_1_24 | SE_225 | SE_225_1 | ACATAAATGGCATCACACTG | 8039.456611643801 | 7147.655050101625 | 7241.534483699112 | 7603.545504536478 | 7431.585750694953 | 6948.673391254506 | 7892.561934015925 | 7882.357404257471 | 6489.39641034634 | 7053.142191200186 | 7343.198739150244 | 7690.672390335393 |
| SE_225_1_26 | SE_225 | SE_225_1 | GCTCCTTCTTGCTTTTGCCA | 50.93751155935005 | 85.94775367851507 | 84.92731018657079 | 33.834020728726415 | 103.93826224748186 | 69.3048087894727 | 53.808030638232374 | 107.7702979234051 | 63.04695472558706 | 18.225173620672315 | 120.8998513638867 | 24.37430126728694 |
| SE_225_1_27 | SE_225 | SE_225_1 | GTCTGAATCAGACTTCAGGG | 33.59708209233727 | 52.9678016855965 | 71.10193410968716 | 24.724861301761607 | 30.757240869152795 | 2.598930329605226 | 20.4470516425283 | 49.15838150892162 | 26.639558334755094 | 16.76715973101853 | 73.94845277596953 | 1.1606810127279494 |
| SE_225_2_10 | SE_225 | SE_225_2 | TTTACCTGAAAACAAAACAG | 16.256652625324485 | 19.987849692677923 | 12.837849214249072 | 10.410467916531204 | 12.727134152752882 | 122.14972549144564 | 4.30464245105859 | 105.87959094229272 | 25.751573056929924 | 8.748083337922711 | 219.49778839851274 | 47.58792152184593 |
| SE_225_2_16 | SE_225 | SE_225_2 | GTTCCAATCCCTGTTAGCGG | 27.094421042207472 | 38.97630690072195 | 32.58838646693995 | 27.32747828089441 | 50.90853661101153 | 65.83956834999907 | 27.980175931880833 | 88.86322811228139 | 27.527543612580263 | 9.477090282749604 | 38.73490383503166 | 134.63899747644214 |
| SE_225_2_4 | SE_225 | SE_225_2 | TGATCATCTGGAGATCACAG | 133.30455152766078 | 167.89793741849456 | 197.50537252690881 | 126.22692348794084 | 125.15015250207 | 143.80747823815585 | 163.5764131402264 | 76.57363273505099 | 131.42182111812514 | 204.85095149635683 | 115.03092654039705 | 210.08326330375886 |
| SE_225_2_42 | SE_225 | SE_225_2 | GCTGGCAGGGAGAACCAACA | 3569.9609165212564 | 3265.015247298939 | 3254.888539243457 | 3736.0566735451357 | 3219.9649406464787 | 3443.5826867269247 | 3851.578833084673 | 3484.5729661900978 | 3587.460522413686 | 2958.31018210753 | 3297.1619658364825 | 3609.7179495839227 |
| SE_225_2_6 | SE_225 | SE_225_2 | CTGATTGTGCCAAGGACCCA | 381.48944827428124 | 342.79162222942637 | 426.61160465812304 | 356.55852614119374 | 464.54039657548014 | 396.7700303197312 | 373.4277326293327 | 577.6109827298291 | 316.12275890576046 | 225.99215289633673 | 569.2857078784956 | 443.3801468620767 |
| SE_225_3_10 | SE_225 | SE_225_3 | GAACACTTCGTTGTCCTCTG | 74.78060207649263 | 108.93378082509469 | 104.67784743926167 | 74.1745839052848 | 79.5445884547055 | 110.02138395328791 | 78.55972473181926 | 88.86322811228139 | 121.65398306204825 | 59.04956253097831 | 134.98527094026184 | 150.88853165463343 |
| SE_225_3_11 | SE_225 | SE_225_3 | ACTTTCACGGACAACCTCAG | 216.75536833765977 | 235.8566263735995 | 266.6322529113269 | 200.40150739322567 | 200.45236290585788 | 303.20853845394305 | 228.14604990610528 | 166.38221433788857 | 219.33236362281693 | 236.92725706874012 | 269.9705418805237 | 279.7241240674358 |
| SE_225_3_12 | SE_225 | SE_225_3 | TTGTCCGTGAAAGTTCACCC | 200.4987157123353 | 243.85176625067066 | 300.2081662409014 | 223.82506020542087 | 279.9969513605634 | 261.62565318025946 | 117.30150679134657 | 259.97220990295085 | 215.78042251151626 | 207.7669792756644 | 214.80264853972102 | 143.92444557826573 |
| SE_225_3_14 | SE_225 | SE_225_3 | CAGAATAGTGAAACTAACAG | 166.90163361999802 | 234.8572338889656 | 260.7070917355196 | 242.04337905935049 | 253.4820885423282 | 305.8074687835483 | 242.13613787204568 | 185.28928414901227 | 318.78671473923595 | 191.72882648947277 | 242.9734876924713 | 282.04548609289174 |
| SE_225_3_9 | SE_225 | SE_225_3 | TCAGAGGACAACGAAGTGTT | 3343.451556608402 | 3307.9891241381965 | 3338.8283225673936 | 3159.57701266722 | 3193.4500778282436 | 3003.497150913773 | 3105.7995284387725 | 3562.0919524157052 | 3321.95292434396 | 3080.0543418936213 | 3124.615576025887 | 2901.7025318198735 |
| SE_226_1_10 | SE_226 | SE_226_1 | ATATCCTTACTCTTAGGAAG | 9.75399157519469 | 30.98116702365078 | 14.81290293951816 | 0.0 | 22.272484767317543 | 6.064170769078861 | 1.0761606127646475 | 21.743130282792254 | 5.327911666951019 | 0.7290069448268927 | 1.173784964697929 | 0.0 |
| SE_226_1_16 | SE_226 | SE_226_1 | CAGTGTGCTTAAAGGCAAGG | 2127.4539402341306 | 2111.7163200314226 | 1960.24082232957 | 2147.159007784561 | 1929.2214186547908 | 1539.4330652361623 | 1970.4500819720695 | 2162.9687863925515 | 2030.8223303861632 | 2357.608459570171 | 1912.0957074929263 | 1689.9515545318943 |
| SE_226_1_17 | SE_226 | SE_226_1 | ACGTTATCAAAAGCAACAGA | 7266.723723520045 | 8491.837941934216 | 8515.444136497674 | 7948.392254271574 | 8088.093754074456 | 9370.876458446577 | 7514.829558935533 | 9638.824189710862 | 9809.57336413465 | 8107.2862334198735 | 9772.933616074957 | 10427.558218347898 |
| SE_226_1_24 | SE_226 | SE_226_1 | TAGAACTACCTAACTACCCA | 335.9708209233727 | 403.7545637920941 | 379.2103152516649 | 426.82918457777936 | 305.45121966606916 | 343.92511361775826 | 278.7255987060437 | 392.32169858081676 | 343.6503025183407 | 261.71349319285446 | 325.13843522132635 | 582.6618683894307 |
| SE_226_1_7 | SE_226 | SE_226_1 | GCATAGTTATTTAGACCTGC | 912.5401007015477 | 1040.3675765038859 | 1009.252453612504 | 918.7237936338787 | 1040.443216987548 | 719.9037013006476 | 1061.0943641859424 | 1037.052779140135 | 986.5516436637637 | 871.8923060129637 | 908.5095626761971 | 796.2271747313733 |
| SE_226_2_1 | SE_226 | SE_226_2 | TTACAGGGTTTCTATTTTGG | 41.18351998415536 | 49.96962423169481 | 48.38881626909266 | 31.23140374959361 | 18.030106716399914 | 50.2459863723677 | 23.675533480822246 | 51.04908849003399 | 55.05508722516053 | 4.374041668961356 | 55.167893340802664 | 41.78451645820618 |
| SE_226_2_3 | SE_226 | SE_226_2 | TTAAACTCTCAAATGCTAAG | 7.586437891818092 | 9.993924846338961 | 8.887741763710896 | 7.807850937398403 | 8.484756101835254 | 6.93048087894727 | 9.685445514881827 | 23.633837263904624 | 3.551941111300679 | 30.61829168272949 | 84.5125174582509 | 1.1606810127279494 |
| SE_226_2_5 | SE_226 | SE_226_2 | TCTCAAATGCTAAGAGGAGG | 359.81391144051526 | 508.69077467865316 | 463.15009857560113 | 353.95590916206095 | 407.2682928880922 | 349.1229742769687 | 314.23889892727703 | 401.7752334863786 | 606.493944754591 | 403.1408404892716 | 431.95286700883787 | 600.0720835803498 |
| SE_226_2_8 | SE_226 | SE_226_2 | TTTTGCTGGGACAAAGGCAA | 24.926867358830876 | 36.977521931454156 | 54.31397744489992 | 27.32747828089441 | 22.272484767317543 | 40.71657516381521 | 10.761606127646475 | 63.33868386726439 | 29.303514168230603 | 8.748083337922711 | 91.5552272464 3846 | 41.78451645820618 |
| SE_226_2_9 | SE_226 | SE_226_2 | ACTATACGATCACAAGGAGA | 469.27537245103343 | 502.6944197708498 | 435.49934642183393 | 343.54544124552973 | 461.35861303729195 | 399.36896064933643 | 392.79862365909634 | 458.4964429197497 | 444.88062419041006 | 702.7626948131245 | 454.25478133809855 | 450.3442329384444 |
| SE_227_1_0 | SE_227 | SE_227_1 | GCTCAGCATTAATTATGCCA | 938.5507449020669 | 1090.3372007355808 | 1154.418902419782 | 966.8722077478355 | 1091.3517535985595 | 655.7967531703854 | 953.4783029094776 | 1107.9542909318488 | 1146.3889936722942 | 1043.9379449921103 | 1176.132534627325 | 755.6033392858951 |
| SE_227_1_18 | SE_227 | SE_227_1 | CTAACTGACCAGTTTCAAGA | 252.52000411337366 | 259.842046004813 | 305.1458005540741 | 221.22244322628808 | 272.5727897714575 | 522.3849962506505 | 265.8116713528679 | 172.99968877178185 | 336.5464202957393 | 513.9498961029593 | 127.94256115207426 | 205.44053925284706 |
| SE_227_1_22 | SE_227 | SE_227_1 | GTTTCAAGAGGGCAAAGGGG | 94.288585226882 | 110.93256579436247 | 117.51569665351074 | 92.39290275921444 | 159.089176909411 | 42.44919538355203 | 78.55972473181926 | 126.67736773452879 | 86.13457194904147 | 56.13353475167074 | 136.15905590495976 | 34.820430381838484 |
| SE_227_1_36 | SE_227 | SE_227_1 | TCTGCTGGTGGAAAGAGCAG | 693.6171786805113 | 797.5152027378491 | 734.7199858001007 | 631.1346174397042 | 920.596037049125 | 596.0213555894652 | 598.345300697144 | 996.402579046219 | 683.7486639253807 | 712.968792040701 | 881.5125084881447 | 567.5730152239672 |
| SE_227_1_9 | SE_227 | SE_227_1 | GATCTCCCTGACTTACTCCC | 138.72343573610226 | 187.88578711117248 | 201.45547997744697 | 148.34916781056964 | 136.81669214209347 | 97.89304241513018 | 114.07302495305264 | 102.09817698006798 | 198.02071695501286 | 142.88536118607095 | 85.68630242294881 | 85.89039494186825 |
| SE_227_2_14 | SE_227 | SE_227_2 | CTACAATAGCCACTACTAGG | 15.172875783636185 | 35.97812944682026 | 55.301504307534465 | 32.53271223916001 | 57.27210368738796 | 3.465240439473635 | 19.370891029763655 | 6.617474433893295 | 27.527543612580263 | 21.141201399979888 | 7.042709788187574 | 4.642724050911798 |
| SE_227_2_19 | SE_227 | SE_227_2 | TGAGCTTTTCTAATAGCCTG | 138.72343573610226 | 249.84812115847404 | 268.607306636596 | 173.07402911233126 | 201.5129574185873 | 189.7219140611815 | 138.82471904663953 | 128.56807471564116 | 313.4588030722849 | 333.88518073071685 | 245.32105762186717 | 191.51236710011165 |
| SE_227_2_3 | SE_227 | SE_227_2 | TGTCCAACAGAAATATAACA | 176.65562519519273 | 207.87363680385042 | 213.30580232906152 | 186.08711400799527 | 187.725228753105 | 178.4598826328922 | 137.74855843387488 | 154.09261896065814 | 210.45251084456524 | 44.46942363444045 | 170.1988198811997 | 427.1306126838854 |
| SE_227_2_5 | SE_227 | SE_227_2 | CTAAGGAGAGGAGGAAGGAA | 1574.7277509730984 | 1851.8742740266096 | 1757.7978154894884 | 1751.561226956375 | 1761.6474856435445 | 2031.4972076414185 | 1720.7808198106713 | 1628.8440642283067 | 1930.4799939919192 | 1738.681563412139 | 1576.3932075893188 | 2090.386503923037 |
| SE_227_2_6 | SE_227 | SE_227_2 | TAACCTTTATATATAAGAAA | 2.167553683376598 | 0.9993924846338962 | 1.975053725269088 | 0.0 | 0.0 | 0.0 | 0.0 | 0.0 | 0.8879852778251698 | 0.0 | 0.0 | 0.0 |
| SE_228_1_0 | SE_228 | SE_228_1 | CCTGCAGGCCGCGAGCCCGG | 3.2513305250648967 | 4.996962423169481 | 3.950107450538176 | 14.314393385230405 | 1.0605945127294067 | 6.93048087894727 | 8.60928490211718 | 5.67212094333711 | 0.8879852778251698 | 0.7290069448268927 | 11.73784964697929 | 1.1606810127279494 |
| SE_228_1_29 | SE_228 | SE_228_1 | GTAGTCATCGGCGTGGCCCA | 1036.0906606540138 | 975.4070650026827 | 891.7367569589933 | 969.4748247269683 | 953.4744669437366 | 771.0159977828838 | 1049.2565974455313 | 949.1349045184097 | 1093.109877002784 | 713.6977989855279 | 833.3873249355296 | 840.3330532150354 |
| SE_228_1_31 | SE_228 | SE_228_1 | GTTTGCAGTAGTCATCGGCG | 707.7062776224592 | 885.461741385632 | 927.2877240138369 | 835.4400503016291 | 1020.2919212456893 | 797.8716111888044 | 757.6170713863119 | 763.8456203693975 | 890.6492336586452 | 814.3007573716391 | 982.4580154521666 | 942.4729823350949 |
| SE_228_1_34 | SE_228 | SE_228_1 | GTTTGAAGTCGAGCTCGTCG | 186.4096167703874 | 183.8882171726369 | 165.9045129226034 | 169.17010364363205 | 154.84679885849337 | 181.92512307236584 | 204.47051642528302 | 104.93423745173654 | 113.66211556162173 | 110.0800486688608 | 104.46686185811568 | 157.85261773100112 |
| SE_228_1_35 | SE_228 | SE_228_1 | CTCGACTTCAAACTCGTCTT | 641.595890279473 | 678.5874970664155 | 609.3040742455137 | 684.4882655119267 | 589.6905490775501 | 563.9678815243341 | 542.3849488333823 | 600.2994665031774 | 595.8381214206889 | 549.671236399477 | 561.0692131256101 | 869.3500785332342 |
| SE_228_2_14 | SE_228 | SE_228_2 | GTCACTCAGATCCACCAAAG | 164.73407993662144 | 247.84933618920624 | 252.80687683444327 | 252.45384697588167 | 259.84565561870465 | 423.62564372565186 | 150.66248578705066 | 284.5514006574117 | 379.1697136313475 | 153.09145841364744 | 142.02798072844942 | 329.6334076147376 |
| SE_228_2_15 | SE_228 | SE_228_2 | TGTCACTCAGATCCACCAAA | 407.5000924748004 | 446.7284406313516 | 461.1750448503321 | 433.33572702561133 | 526.0548783137857 | 712.9732204217004 | 448.758975522858 | 464.16856386308683 | 488.3919028038434 | 397.3087849306565 | 527.0294491493702 | 508.3782835748419 |
| SE_228_2_18 | SE_228 | SE_228_2 | TTTGGTGGATCTGAGTGACA | 158.23141888649164 | 162.90097499532507 | 157.0167711588925 | 222.52375171585447 | 147.42263726938754 | 166.33154109473446 | 216.30828316569415 | 162.60080037566382 | 206.90056973326455 | 158.9235139722626 | 373.26361877394146 | 69.64086076367697 |
| SE_228_2_19 | SE_228 | SE_228_2 | GTCAAGTTCTCTTTGAAAAC | 112.71279153558308 | 96.94107100948793 | 101.71526685135804 | 96.29682822791364 | 88.02934455654076 | 69.3048087894727 | 85.01668840840715 | 103.98888396118035 | 63.93494000341222 | 72.90069448268926 | 120.8998513638867 | 39.46315443275028 |
| SE_228_2_8 | SE_228 | SE_228_2 | GGCCAGGATCCACCCCTGGG | 197.2473851872704 | 264.8390084279825 | 277.49504840030687 | 149.65047630013606 | 268.3304117205399 | 468.67376943880913 | 171.10953742957895 | 190.01605160179318 | 256.62774529147407 | 129.03422923436 | 119.72606639918877 | 571.0550582621511 |
| SE_228_3_12 | SE_228 | SE_228_3 | ATAGAAGGCACCAAGGCCAA | 897.3672249179115 | 869.4714616314897 | 867.0485853931297 | 914.8198681651795 | 860.1421498235488 | 790.9411303098572 | 921.1934845265382 | 1288.5168076280802 | 917.2887919934004 | 688.9115628614136 | 1125.659781145314 | 312.2231924238184 |
| SE_228_3_14 | SE_228 | SE_228_3 | TCTCCTCATAGAAGGCACCA | 76.94815575986922 | 72.95565137827442 | 74.06451469759081 | 91.09159426964803 | 119.84717993842297 | 229.5721791151283 | 97.93061576158291 | 59.557269905039654 | 165.1652616754816 | 71.44268059303548 | 73.94845277596953 | 177.58419494737626 |
| SE_228_3_16 | SE_228 | SE_228_3 | CTTCTATGAGGAGACCACAC | 69.36171786805113 | 86.94714616314897 | 79.00214901076352 | 85.88636031138243 | 42.42378050917627 | 30.320853845394307 | 78.55972473181926 | 51.04908849003399 | 144.74160028550267 | 276.2936320893923 | 7.042709788187574 | 199.6371341892073 |
| SE_228_3_26 | SE_228 | SE_228_3 | CAAGACTTCAGAGCAGACAG | 1359.0561594771268 | 1398.1500860028207 | 1449.6894343475105 | 1380.6883074299508 | 1403.1665403410052 | 1215.4330841453775 | 1254.803274483579 | 1609.936994417183 | 1259.1631239560907 | 1122.6706950334146 | 2534.2017387828287 | 1693.4335975700783 |
| SE_228_3_31 | SE_228 | SE_228_3 | AGATGGCTGTCAAGTTTGAC | 2749.5418473632144 | 2419.529205298663 | 2488.567693839051 | 2521.935852779684 | 2395.8830042557297 | 2915.1335197071953 | 2649.507428626562 | 2358.656958937682 | 2842.4408743183685 | 2074.7537649773367 | 2386.30483323089 | 2266.810017857685 |
| SE_228_4_11 | SE_228 | SE_228_4 | TCTTTCTATCTCCAAGGAAG | 15.172875783636185 | 14.990887269508443 | 8.887741763710896 | 6.506542447832002 | 4.242378050917627 | 0.0 | 3.228481838293942 | 13.23494886778659 | 5.327911666951019 | 0.0 | 9.390279717583432 | 9.285448101823595 |
| SE_228_4_13 | SE_228 | SE_228_4 | TCTATCTCCAAGGAAGCGGG | 65.02661050129794 | 153.90644263362 | 105.66537430189621 | 98.8994452070464 3 | 68.93864332741144 | 52.844916701972934 | 55.960351863761666 | 105.87959094229272 | 46.17523444690883 | 130.49224312401378 | 24.64948425865651 | 51.069964560029774 |
| SE_228_4_14 | SE_228 | SE_228_4 | TCACAGCTGACAGGATGCTG | 3809.475598534371 | 3512.864583488145 | 3472.144449023057 | 3595.5153566719646 | 3326.0243919194195 | 4184.277830664414 | 3935.5193608803156 | 3602.742152509621 | 3390.327790736498 | 3127.4397933073697 | 3056.536048073407 | 4150.595301515147 |
| SE_228_4_15 | SE_228 | SE_228_4 | TTGATGACTTCACAGCTGAC | 319.71416829804815 | 301.8165303594366 | 287.3703170266523 | 255.0564639550145 | 351.0567837134336 | 224.37431845591786 | 320.69586260386495 | 350.72614499634466 | 346.3142583518162 | 220.8891042825485 | 360.3519841622642 | 360.97179495839225 |
| SE_228_4_16 | SE_228 | SE_228_4 | AATGGAGAGGGTATCAGCAA | 602.5799239786942 | 629.6172653193546 | 573.7531071906701 | 579.0822778570482 | 656.5080033795027 | 418.4277830664414 | 770.5309987394876 | 472.6767452780925 | 727.259942538814 | 500.82777109607525 | 588.0662673136625 | 360.97179495839225 |
| SE_228_5_10 | SE_228 | SE_228_5 | ATCTCTGCCTAGCAGAGCAG | 16.256652625324485 | 12.992102300240651 | 7.900214901076352 | 15.615701874796805 | 14.848323178211695 | 15.593581977631358 | 18.294730416999005 | 4.726767452780925 | 0.8879852778251698 | 0.0 | 0.0 | 3.482043038183848 |
| SE_228_5_11 | SE_228 | SE_228_5 | TACAGGGTGAGGGAACAGGA | 2035.3329086906253 | 1965.8050172748738 | 1991.8416819338754 | 1966.277127734831 | 1801.9500771272621 | 2036.695068300629 | 1773.512689836139 | 1698.8002225294645 | 2239.4988706750783 | 1636.620591136374 | 1692.5979190944136 | 1988.2465748029774 |
| SE_228_5_25 | SE_228 | SE_228_5 | AAAACCCTAGGCACTTGCAA | 28.17819788389577 | 51.968409200962604 | 43.451181955919935 | 50.751031093089615 | 39.24199697098805 | 103.09090307434064 | 19.370891029763655 | 42.54090707502832 | 5.327911666951019 | 37.17935418617152 | 99.77172199932397 | 15.088853165463343 |
| SE_228_5_7 | SE_228 | SE_228_5 | ACAAGTCCATATTTGAGCAG | 179.90695572025763 | 215.86877668092157 | 211.33074860379241 | 212.11328379932328 | 199.39176839312847 | 251.22993186183854 | 174.33801926787288 | 320.47483329854674 | 253.07580418017338 | 192.45783343429966 | 353.30927437407667 | 208.9225822910309 |
| SE_228_5_9 | SE_228 | SE_228_5 | CTTTGGGAGGATAAAACAGG | 87.78592417675222 | 71.95625889364052 | 80.97720273603261 | 71.57196692615203 | 124.08955798934059 | 63.24063802039384 | 91.47365208499504 | 90.75393509339376 | 90.57449833816732 | 12.393118062057175 | 143.20176569314734 | 37.14179240729438 |
| SE_229_1_11 | SE_229 | SE_229_1 | GGCTACACTTAAAGCTTAAA | 62.859056817921335 | 95.94167852485404 | 86.90236391183987 | 58.55888203048802 | 30.757240869152795 | 207.9144263684181 | 44.12258512335055 | 34.03272566002266 | 59.49501361428638 | 53.21750697236316 | 7.042709788187574 | 71.96222278913287 |
| SE_229_1_12 | SE_229 | SE_229_1 | GGGAAGCTGGGCCAACCAGG | 793.3246481158347 | 945.4252904636658 | 941.1131000907204 | 922.6277191025779 | 900.4447413072663 | 579.5614635019655 | 778.0641230288401 | 861.2170298966845 | 816.058470321331 | 960.1021463370176 | 1147.9616954745745 | 721.9435899167845 |

| | | | | | | | | | | | | | | | |
|---|---|---|---|---|---|---|---|---|---|---|---|---|---|---|---|
| SE_229_1_23 | SE_229 | SE_229_1 | GTTCTGTTTGACAAGAGCAA | 6.5026610501297935 | 3.9975699385355847 | 9.87526862634544 | 5.205233958265602 | 5.3029725636470335 | 25.12299318618385 | 7.533124289352532 | 8.508181415005666 | 8.879852778251697 | 9.477090282749604 | 0.0 | 3.482043038183848 |
| SE_229_1_44 | SE_229 | SE_229_1 | CAGCTTTTCCAGGGAGGCAA | 464.9402650842802 | 394.7600314303389 | 399.9483793609719 | 379.9820789358865 | 496.3582319573623 | 692.1817777848588 | 401.4079085612145 | 503.8734104664466 | 567.4225925302835 | 387.8316946479069 | 571.6332778078914 | 557.1268861094158 |
| SE_23_1_11 | SE_23 | SE_23_1 | GCTATGCTATGATTCTCATG | 674.109195530122 | 461.7193279008600 4 | 527.3393446468465 | 616.8202240544738 | 583.3269820011737 | 894.0320333841978 | 699.5043982970209 | 556.813205937593 | 492.8318291929692 | 498.6407502615946 | 664.3622900190278 | 677.8377114331224 |
| SE_23_1_18 | SE_23 | SE_23_1 | TCCCCCTTCACAGCATGAAA | 29.26197472558407 | 31.980559508284678 | 32.58838646693995 | 32.53271223916001 | 9.545350614564661 | 4.331550549342044 | 21.52321225529295 | 67.12009782948914 | 25.75157305692924 | 32.80531251721017 | 71.60088284657367 | 1.1606810127279494 |
| SE_23_1_22 | SE_23 | SE_23_1 | GTATACTCCACATTAAGGGA | 31.429528408960667 | 38.97630690072195 | 46.41376254382357 | 18.218318853929606 | 54.09032014919974 | 56.31015714144657 | 5.380803063823238 | 87.9178746217252 | 31.96747000170611 | 17.496166675845423 | 56.341678305500594 | 85.89039494186825 |
| SE_23_1_27 | SE_23 | SE_23_1 | TCAGGAGGAAGGCCTCATCA | 4733.937244494489 | 4212.439322731872 | 4425.107871465392 | 4600.125510617226 | 4810.856709740589 | 3528.481077494029 | 4590.901174053986 | 4417.636861369053 | 4807.552294145469 | 4183.770856361537 | 4841.862979378957 | 4029.8844761914406 |
| SE_23_1_9 | SE_23 | SE_23_1 | GTATATATGAAAGACTCAGT | 43.351073667531956 | 73.95504386290831 | 60.23913862070719 | 41.641871666124814 | 81.66577748016432 | 47.64705604276248 | 15.066248578705064 | 46.32232103725307 | 68.37486693253807 | 35.72134029651774 | 109.1620017169074 | 20.89225822910309 |
| SE_23_2_17 | SE_23 | SE_23_2 | TCGAGCCTAAAACAGCTTGA | 2737.620302104643 | 2499.4806040693743 | 2592.258014415678 | 2734.049136579007 | 2207.0971809898956 | 2577.272576858516 | 2677.487604558443 | 2535.438061671688 | 2543.189835691286 | 2644.1081888871395 | 2638.6686006409445 | 3125.713967276368 |
| SE_23_2_2 | SE_23 | SE_23_2 | CGAACCTCAGTTGAATCATC | 4061.9956026477444 | 3782.700554339297 | 3667.6747678246966 | 3695.716110368577 | 3911.472562946052 | 3546.6735898012653 | 4159.360768335362 | 3184.895909683787 | 3650.507477139273 | 3879.0459534238958 | 3048.3195533205217 | 3524.9882356547823 |
| SE_23_2_3 | SE_23 | SE_23_2 | AAAACCAGATGATTCAACTG | 20.59175999207768 | 18.98845720804403 | 27.650752153767232 | 23.423552812195208 | 9.545350614564661 | 52.844916701972934 | 12.913927353175769 | 25.52454424501699 4 | 36.40739639083196 | 14.580138896537854 | 52.820323411406804 | 17.410215190919242 |
| SE_23_2_4 | SE_23 | SE_23_2 | TTGGGAATTTCGATCGTTTT | 108.37768416882989 | 156.9046200875217 | 179.729888999487 | 135.33608291490566 | 96.51410065837601 | 50.2459863723677 | 75.33124289352533 | 61.44797688615203 | 112.77413028379657 | 145.07238020055163 | 56.341678305500594 | 56.873369623669525 |
| SE_23_2_6 | SE_23 | SE_23_2 | TGTATGTAGCACTCACCAAT | 613.4176923955772 | 620.6227329576495 | 592.5161175807265 | 687.0908824910595 | 491.05525939371535 | 453.0801874611778 | 600.4976219226733 | 467.9499778253116 | 571.8625189194094 | 1402.6093618469415 | 462.471276090984 | 355.1683898947525 |
| SE_23_3_11 | SE_23 | SE_23_3 | TATTCCCCCTGGCCTATGGG | 98.6236925936352 | 106.9349958558269 | 90.85247136237805 | 63.76411598875362 | 97.57469517110542 | 266.8235138394699 | 71.02660044246673 | 104.93423745173654 | 97.67838056076867 | 69.98466670338169 | 93.90279717583432 | 67.31949873822107 |
| SE_23_3_17 | SE_23 | SE_23_3 | ACTTCCTTAACTCCTCCCAT | 282.86575568064603 | 280.82928818212486 | 335.75913329574496 | 291.4931016628737 | 324.5419208951985 | 129.94651648026132 | 304.55345341239524 | 378.141396222474 | 455.5364475243121 | 306.1829168272949 | 385.00146842092073 | 215.88666836739858 |
| SE_23_3_2 | SE_23 | SE_23_3 | TTATTCAAAGAAGTTGAGTG | 20.59175999207768 | 27.982989569749094 | 32.58838646693995 | 18.218318853929606 | 19.090701229129323 | 19.9251325269734 | 8.60928490211718 | 99.26211650839943 | 24.863587779104755 | 14.580138896537854 | 75.12223774066746 | 19.73157721637514 |
| SE_23_3_6 | SE_23 | SE_23_3 | ACTCATCTATGATTAAGAAA | 35.76463577571386 | 67.95868895510495 | 54.31397744489992 | 35.13532921829281 | 43.48437502190568 | 94.42780197565655 | 24.75169409358689 | 67.12009782948914 | 79.03068972644012 | 13.122125006884067 | 30.518409082146157 | 64.99813671276517 |
| SE_23_3_7 | SE_23 | SE_23_3 | GGGAGTTTAAGAGAAATGAC | 97.53991575194691 | 173.89429232629794 | 153.06666370835433 | 152.25309327926885 | 183.48285070218736 | 144.67378834802426 | 107.61606127646475 | 92.64464207450614 | 159.83735000853056 | 179.3357084274156 | 178.4153146340852 | 42.945197470934126 |
| SE_23_4_0 | SE_23 | SE_23_4 | TTTTAAGACATAATTACAAA | 2.167553683376598 | 4.996962423169481 | 7.900214901076352 | 1.3013084895664004 | 0.0 | 0.0 | 0.0 | 0.0 | 3.551941111300679 | 0.0 | 0.0 | 0.0 |
| SE_23_4_12 | SE_23 | SE_23_4 | TGTAGTTCCAGCACTTTGCA | 562.4801808362271 | 671.591749673978 3 | 569.8029997401319 | 599.9032136901106 | 644.8414637394793 | 625.4758993249911 | 619.868512952437 | 718.4686528227006 | 450.2085358573611 | 731.1939656613733 | 723.0515382539243 | 754.4426582731671 |
| SE_23_4_2 | SE_23 | SE_23_4 | ATGCACGTGCCACCATGCCC | 8153.253180021073 | 5012.952702923623 | 4915.90872219476 | 7364.10474245626 | 5732.513341302443 | 6977.261624880164 | 9412.100719239606 | 8586.645754721829 | 5973.4769639299175 | 11087.466623872211 | 11631.03521519178 | 10889.509261413621 |
| SE_23_4_4 | SE_23 | SE_23_4 | CTATGAGCCTTTTATTTGTA | 17.340429467012783 | 17.98906472341013 | 15.800429802152705 | 15.615701874796805 | 26.51486281823517 | 19.9251325269734 | 11.837766740411123 | 35.92343264113503 | 0.8879852778251698 | 5.103048613788249 | 34.03976397623994 | 10.446129114551544 |
| SE_23_4_5 | SE_23 | SE_23_4 | TGATAATCCCTACAAATAAA | 147.39365046960864 | 231.8590564350639 | 228.11870526857967 | 214.71590077845607 | 234.3913873131989 | 149.0053388973663 | 205.54667703804768 | 113.4424188667422 | 157.17339417505505 | 110.80905561368769 | 79.81737759945918 | 226.33279748195014 |
| SE_23_5_11 | SE_23 | SE_23_5 | GATCCTGTCTTCTTCCCAAG | 88.86970101844051 | 91.94410858631845 | 87.88989077447442 | 85.88636031138243 | 136.81669214209347 | 85.76470087697247 | 81.7882065701132 | 141.80302358342774 | 161.6133205641809 | 44.46942363444045 | 343.9189946564932 | 110.26469620915519 |
| SE_23_5_4 | SE_23 | SE_23_5 | ACTCCAGCCTGCATAACAGA | 372.8192335407748 | 450.7260105698872 | 450.3122493613521 | 399.5017062968849 5 | 505.903582571927 | 389.8395494407839 | 397.1032661101549 | 495.36522905144096 | 362.29799335266927 | 360.129430744485 | 511.77024460829705 | 360.97179495839225 |
| SE_23_5_5 | SE_23 | SE_23_5 | ATTGATTTGCTGAAGAAGCT | 36.84841261740216 | 57.964764108765976 | 41.47612823065085 | 32.53271223916001 | 68.93864332741144 | 86.63101098684088 | 34.43713960846872 | 17.01636283001133 | 5.327911666951019 | 30.61829168272949 | 38.73490383503166 | 39.46315443275028 |
| SE_23_5_6 | SE_23 | SE_23_5 | TAGGGGAGAGTATATCAGAG | 1013.3313469785595 | 1455.1154576269528 | 1520.791368457198 | 1102.208290662741 | 1364.9851782827466 | 994.5240061289333 | 1060.0182035731777 | 1387.7789241364796 | 1581.5017998066274 | 1099.342472798954 | 1694.9454890238096 | 1192.019400071604 |
| SE_23_5_7 | SE_23 | SE_23_5 | CTATGAGGAAGAAACACAAT | 668.6903113216804 | 764.5352507449306 | 735.7075126627353 | 693.5974249388914 | 734.9919973214788 | 1172.117578651957 | 757.6170713863119 | 658.911382917661 | 777.8751033748488 | 1582.674077219184 | 616.2371064664128 | 1008.631800060588 |
| SE_23_6_0 | SE_23 | SE_23_6 | GTATGAAATCACCTTACCTT | 340.30592829012585 | 313.8092401750434 | 291.3204244771905 | 301.9035695794049 | 340.4508385861396 | 265.0908936197331 | 291.63952605921946 | 251.4640284879452 | 378.28172835352234 | 315.6600711004455 | 164.32989505771008 | 441.05878483662076 |
| SE_23_6_2 | SE_23 | SE_23_6 | TCCTTCTCTGAAACATTGGA | 54.188842084414944 | 68.95808143973883 | 47.40128940645811 | 61.16149900962082 | 81.66577748016432 | 70.17111889934111 | 55.960351863761666 | 46.32232103725307 | 31.96747000170611 | 7.290069448268927 | 18.780559435166865 | 3.482043038183848 |
| SE_23_6_3 | SE_23 | SE_23_6 | GCCATCCAATGTTTCAGAGA | 547.307305052591 | 602.6336682342394 | 620.1668697344936 | 588.191437284013 | 537.7214179538092 | 601.2192162486757 | 454.13977858668125 | 517.1083593342332 | 589.6222244759127 | 320.7630557238328 | 571.6332778078914 | 651.1420481403796 |
| SE_23_6_4 | SE_23 | SE_23_6 | CAGCAGAAAGTGCACTCACT | 3957.9530258456675 | 3601.8105146205617 | 3702.2382080169054 | 3971.5935101566542 | 3582.6882639999358 | 3960.7698223183647 | 3832.20794205491 | 3818.282748356431 | 4436.374448014548 | 3545.160772693179 | 4024.9086439491984 | 3289.3699900710085 |
| SE_230_1_14 | SE_230 | SE_230_1 | ACAGTCTGAATTTAAACACG | 1123.876584830766 | 1370.1670964330717 | 1420.0636284684742 | 1240.1469905567797 | 1310.8948177335467 | 1594.8769122677404 | 1131.0448040156446 | 1284.7353936658553 | 1363.0574014616357 | 1444.1627577020743 | 1809.9764155642065 | 1806.0196558046894 |
| SE_230_1_4 | SE_230 | SE_230_1 | GGTAAGAGAAGAATGCTGAA | 385.8245556410344 | 470.7138602625651 | 450.3122493613521 | 473.67629020216975 | 379.69283555712764 | 298.87698790460104 | 378.80853569315593 | 280.76998669518696 | 680.1967224180801 | 354.29737518586984 | 321.61708032723254 | 395.79222534023074 |
| SE_230_1_7 | SE_230 | SE_230_1 | TCATCTGGAGGTATGTTTGG | 714.2089386725889 | 767.5334281988323 | 801.8718124592498 | 782.0864022294066 | 811.3548022379962 | 626.3422094348595 | 633.8586009183773 | 993.5665185745504 | 858.6817636569392 | 1026.4417783162648 | 1108.053006674845 | 511.8603266130257 |
| SE_230_1_8 | SE_230 | SE_230_1 | TTTTTTGGTCCAGAAGATAG | 1.083776841688299 | 5.9963549078033775 | 0.987526862634544 | 0.0 | 3.1817835381882205 | 0.0 | 0.0 | 0.0 | 0.0 | 0.0 | 0.0 | 0.0 |
| SE_230_1_9 | SE_230 | SE_230_1 | GATGGATGCAGAGCTCAGGA | 925.5454228018073 | 1172.2873844755602 | 1078.379333996922 | 1020.225855820058 | 1191.0476377951238 | 1391.2940364486644 | 955.630624135007 | 1060.6866164040396 | 1016.7431431098194 | 1581.2160633295302 | 858.0368091941862 | 1021.3992912005955 |
| SE_230_2_10 | SE_230 | SE_230_2 | AAGCAGATGTTTCTGTCCAC | 501.7886777016824 | 564.6567538181513 | 573.7531071906701 | 575.178352388349 | 594.9935216411972 | 435.75398526380957 | 506.871648612149 | 420.6823032975023 | 359.6340375191938 | 481.14458358574916 | 328.65979011542015 | 516.5030506639375 |
| SE_230_2_14 | SE_230 | SE_230_2 | AATGGTCATGACTGGAACAC | 907.1212164931062 | 755.5407183832256 | 717.9320291353135 | 787.2916361876722 | 911.0506864345604 | 512.8555850420979 | 830.7959930543078 | 645.6764340498744 | 629.5815619780453 | 452.71331273750036 | 599.8041169606417 | 329.6334076147376 |
| SE_230_2_5 | SE_230 | SE_230_2 | ATTTGTTTCTCCCACAGACA | 5.4188842084414945 | 14.990887269508443 | 6.912688038441808 | 1.3013084895664004 | 10.605945127294067 | 4.331550549342044 | 2.152321225529295 | 17.961716320567515 | 6.215896944776189 | 0.7290069448268927 | 37.56111887033373 | 27.856344305470785 |
| SE_230_2_7 | SE_230 | SE_230_2 | ACAGGAACCTCCCTGTCTGT | 710.9576081475241 | 761.5370732910 29 | 721.8821365858516 | 750.854998479813 | 744.5373479360435 | 847.2512874513037 | 644.6202070460239 | 764.7909738599536 | 583.4063275311365 | 789.5145212475247 | 866.2533039470716 | 624.4463848476368 |
| SE_230_3_17 | SE_230 | SE_230_3 | CTTTGCATCCTGTTAGTTCG | 339.22215144843756 | 382.76732161478225 | 414.7612823065085 | 404.7069402551505 | 362.7233233534571 | 683.5186766861744 | 374.5038932420973 | 354.5075589585694 | 665.1009730910522 | 393.663750206522 | 262.9278320923361 | 763.7281063749907 |
| SE_230_3_2 | SE_230 | SE_230_3 | TGGTGCCATTATAATCCCAA | 23.843090517142578 | 23.98541963121351 | 13.825376076883616 | 13.013084895664004 | 47.72675307282331 | 8.663101098684088 | 18.294730416999005 | 10.398888396118036 | 34.63142583518162 | 5.103048613788249 | 10.564064682281362 | 1.1606810127279494 |
| SE_230_3_20 | SE_230 | SE_230_3 | GCTTTCCGCTGTGCTCAGTG | 225.42558307166616 | 239.8541963121351 | 231.0812858564833 | 203.0041243723584 6 | 306.51181417879855 | 277.2192351578908 | 273.34479564222045 | 196.63352603565 865 | 229.98818695671898 | 210.68300705491 97 | 219.49778839851274 | 141.60308355280984 |
| SE_230_3_27 | SE_230 | SE_230_3 | TGAAAGCCAAGTTCTGGGGA | 316.4628377729833 | 448.72722560061936 | 416.73633603177757 | 372.17422801599054 | 450.75266790999785 | 460.01066834012505 | 338.99059302086397 | 438.6440196180698 | 320.5626852948863 | 326.59511128244793 | 394.39174813850417 | 391.14950128931895 |
| SE_230_3_5 | SE_230 | SE_230_3 | AGACTTGTAGACTTAATTTT | 67.19416418467453 | 110.93256579436247 | 123.440857829318 | 119.72038104010883 | 145.30144824392872 | 132.54544680986655 | 91.47365208499504 | 156.9286794323267 | 53.27911666951019 | 62.69459725511277 | 165.503680022408 | 114.907420260067 |
| SE_230_4_11 | SE_230 | SE_230_4 | GTGGCTACTTCTAGGAAGGA | 17.340429467012783 | 11.992709815606755 | 18.763010390056337 | 18.218318853929606 | 22.272484767317543 | 48.51336615263089 | 15.066248578705064 | 2.836060471668555 | 23.97560250127958 5 | 0.0 | 11.73784964697929 | 0.0 |
| SE_230_4_7 | SE_230 | SE_230_4 | GGGAAGAGCCTAGTACCTAG | 524.5479913771367 | 660.5984323430054 | 717.9320291353135 | 503.60638546219695 | 669.2351375322556 | 585.6256342710443 | 523.0140578036187 | 494.41987556088475 | 724.5959867053385 | 538.7361322270737 | 602.1516868900376 | 425.96993167115744 |
| SE_230_4_8 | SE_230 | SE_230_4 | AAGAGCCTAGTACCTAGGGG | 287.2008630473992 | 391.7618539764873 | 363.4098854495122 | 333.1349733289985 | 371.2080794552923 6 | 610.7486274572282 | 345.4475566941585 3 | 393.267052071373 | 445.76860946823524 | 306.1829168272949 | 291.0986712450864 | 399.2742683784146 |
| SE_230_5_0 | SE_230 | SE_230_5 | TTTATACAGATACTGCGAAG | 32.51330525064897 | 16.989672238776237 | 8.887741763710896 | 39.03925468699201 | 18.030106716399914 | 178.4598826328922 | 58.112673089290965 | 35.92343264113503 | 1.7759705556503396 | 83.83579865509266 | 42.25625872912545 | 9.285448101823595 |
| SE_230_5_1 | SE_230 | SE_230_5 | ACACATAGTACAAACAGTAT | 1446.842083653879 | 1375.1640588562411 | 1382.5376076883617 | 1410.618402689978 | 1415.8936744937758 | 1354.0427017243228 | 1545.3666399300337 | 1188.3093376291245 | 1405.6806947972439 | 1769.2998155089685 | 963.6774560169997 | 1426.4769646426498 |
| SE_230_5_10 | SE_230 | SE_230_5 | GAGCAAAATTCTCTGCAACG | 160.39897256986825 | 259.842046004813 | 276.50752153767235 | 176.97795458103047 | 294.8452745387751 | 343.92511361775826 | 158.19561007640317 | 334.6551356568895 | 293.9231269601312 | 292.33178487558393 | 277.01325166871123 | 97.49720506914775 |
| SE_230_5_2 | SE_230 | SE_230_5 | GTACAAACAGTATAGGTCCT | 834.5081680999901 | 732.5546913266459 | 825.5724571624788 | 874.4793049886212 | 679.8410826595497 | 851.5828380006458 | 727.4845742289017 | 613.5344153709641 | 828.4902642108834 | 1434.6856674193248 | 291.0986712450864 | 691.7658835858579 |
| SE_230_5_3 | SE_230 | SE_230_5 | CATTTCTTGTTAAAATTCAG | 8.670214733506391 | 26.983597085115196 | 12.837849214249072 | 10.410467916531204 | 5.3029725636470335 | 1.7326202197368175 | 9.685445514881827 | 1.89070698111237 | 11.543808611727208 | 35.72134029651774 | 21.128129364562724 | 0.0 |
| SE_231_1_10 | SE_231 | SE_231_1 | AATCACGTATGTCATCCCGG | 193.9960546622055 | 207.87363680385042 | 234.04386644438694 | 223.82506020542087 | 154.84679885849337 | 153.33688944670834 | 96.85445514881827 | 101.15282348951179 | 136.74973278507613 | 319.305041834179 | 90.38144228174053 | 221.69007343103834 |
| SE_231_1_14 | SE_231 | SE_231_1 | GACAATTTACGAAGATGTCA | 117.04789890233629 | 264.8390084279825 | 203.43053370271608 | 139.24000838360485 | 205.7553354695049 | 234.77003977433876 | 76.40740350628997 | 139.91231660231537 | 219.33236321281693 | 103.51898616541875 | 165.503680022408 | 299.455701283811 |
| SE_231_1_15 | SE_231 | SE_231_1 | GTAAATTGTCAAAAATTCCT | 48.769957875973454 | 84.94836119388117 | 106.65290116453076 | 96.29682822791364 | 101.81707322202305 | 123.01603560131404 | 46.27490634887984 | 86.97252113116902 | 70.15083694818841 | 87.48083337922712 | 32.86597901154201 | 26.695663292742836 |
| SE_231_1_16 | SE_231 | SE_231_1 | TAAATTGTCAAAAATTCCTT | 10.837768416882989 | 16.989672238776237 | 22.713117840594514 | 18.218318853929606 | 18.030106716399914 | 64.10694813026224 | 26.904015319116187 | 61.44797688615203 | 8.879852778251697 | 1.4580138896537853 | 26.997054188052367 | 2.321362025455899 |
| SE_231_1_17 | SE_231 | SE_231_1 | TGTCAAAAATTCCTTGGGAC | 138.72343573610226 | 234.8572338889656 | 215.2808560543306 | 199.10019890365928 | 223.78544218590483 | 130.81282659012973 | 178.6426617189315 | 352.61685197745703 | 213.11646667804075 | 250.77838902045107 | 495.33725510252606 | 82.4083519036844 |
| SE_231_2_15 | SE_231 | SE_231_2 | TCCAACCTGGACCTGCCAGT | 15.172875783636185 | 25.984204600481302 | 16.78795666478725 | 36.43663770785921 | 10.605945127294067 | 47.64705604276248 | 9.685445514881827 | 0.0 | 3.551941111300679 | 10.206097227576498 | 3.521354894093787 | 143.92444557826573 |
| SE_231_2_17 | SE_231 | SE_231_2 | GAAACAGCTCAGAAAGCCA | 347.89236618194394 | 442.7308706926816 | 435.4993464218339 3 | 373.47553650555693 | 369.0868904298335 | 333.52939229933736 | 399.2555875356842 | 487.80240112699147 | 516.8074316942489 | 567.8964100201493 | 562.242998090308 | 393.4708633147749 |
| SE_231_2_18 | SE_231 | SE_231_2 | AGATAGATAAAGTATCCTAT | 95.37236206857031 | 92.94350107095235 | 113.56558920297256 | 96.29682822791364 | 103.93826224748186 | 82.29946043749882 | 65.6457973786435 | 190.01605160179318 | 84.35860139339113 | 85.29381254476644 | 146.72312058724114 | 76.60494684004466 |
| SE_231_2_19 | SE_231 | SE_231_2 | CTGAGCAGTAGCAACAGCAG | 1383.9830268359576 | 1550.0767436671729 | 1535.604271396716 | 1568.0767299275126 | 1543.1650160212869 | 1734.3528399565544 | 1528.1480701257994 | 1437.8826591359573 | 1357.7294897946847 | 1811.5822578948282 | 1618.6494663184442 | 1564.5980051572758 |
| SE_231_2_25 | SE_231 | SE_231_2 | GCAGGGGTGCAAAGCAGATA | 11106.545073621688 | 11170.209800753059 | 10912.171832111711 | 11100.161461001396 | 11162.757246477006 | 11830.33086036299 | 11618.229975407134 | 11238.362295731928 | 9674.599601905225 | 9889.708213521626 | 10917.373956655438 | 10046.85484617313 |
| SE_232_1_0 | SE_232 | SE_232_1 | GGAATGTGCACACACCTAAC | 1472.8527278543982 | 1397.150693518187 | 1345.9991137708835 | 1421.0288706065094 | 1263.1680646607235 | 1166.9191791297467 | 1598.0985099555014 | 1311.2052914014287 | 1136.6211556162173 | 1855.3226745844418 | 1430.8438719667754 | 1361.4788279298846 |
| SE_232_1_10 | SE_232 | SE_232_1 | GCGCCCAGAGTTTCTCACAA | 273.1117641054513 | 435.73512330037875 | 453.2748299492557 | 314.9166544750689 | 431.66196668086855 | 361.2513158151264 | 263.65935012733866 | 527.5072477303512 | 541.6710194733536 | 543.110173896035 | 665.5360749837257 | 482.843301294827 |
| SE_232_1_5 | SE_232 | SE_232_1 | ATATGTTTCAGACCACCATA | 48.769957875973454 | 122.92527560996923 | 93.81505195028168 | 68.96934994701923 | 94.3929116329172 | 42.44919538355203 | 26.904015319116187 | 55.775855942814914 | 133.19779167377547 | 44.46942363444045 | 51.646538446708874 | 9.285448101823595 |
| SE_232_1_8 | SE_232 | SE_232_1 | GGGGATGCTTTCCTGCAGAT | 78.03193260155751 | 21.986634661945715 | 37.526020780112674 | 65.06542447832003 | 15.9089176909411 | 294.54543735525897 | 134.52007659558095 | 72.79221877282625 | 177.59705556503397 | 5.103048613788249 | 34.03976397623994 | 48.74860253457388 |
| SE_232_1_9 | SE_232 | SE_232_1 | CGCCCAGAGTTTCTCACAAA | 489.86713244311113 | 682.585067004951 | 804.8343930471534 | 576.4796608779154 | 804.9912351616197 | 996.25662634867 | 471.3583483909156 | 590.8459315976156 | 990.1035847750643 | 630.5910072752622 | 806.3902707474773 | 697.5692886494976 |
| SE_232_2_1 | SE_232 | SE_232_2 | GTTCTGCTGCTATATTACAC | 72.61304839311603 | 75.95382883217611 | 81.96472959866715 | 109.30991312357764 | 77.42339942924669 | 130.81282659012973 | 60.26499431482026 | 75.6282792444948 | 57.71904305863604 | 33.53431946203706 | 26.997054188052367 | 11.606810127279495 |
| SE_232_2_13 | SE_232 | SE_232_2 | TCTCCTGGTGTGGTGGCACA | 202.6662693957119 | 218.86695413482326 | 219.23096350486878 | 195.19627343496006 | 189.84641777856382 | 299.7432980144694 | 157.11944946363852 | 178.67180971511897 | 181.14899667633463 | 309.09894466066246 | 131.46391604616804 | 289.0095721692594 |
| SE_232_2_15 | SE_232 | SE_232_2 | GGTACAAACATACTGTTAGA | 199.414938870647 | 244.85115873530455 | 246.881715658636 | 281.0826337463425 | 300.1482471024221 | 421.02671339604666 | 231.3745317443992 | 312.91200537409725 | 260.17968640277473 | 483.33160442022984 | 476.5566956673592 | 401.59563040387053 |
| SE_232_2_25 | SE_232 | SE_232_2 | TATCACTCTGTAGAGTGCAG | 109.46146101051819 | 123.92466809460312 | 124.42838469195254 | 115.81645557140963 | 93.3323171201878 | 107.42245362368269 | 144.20552211046277 | 79.40969320671954 | 50.615160836034676 | 298.1638404341991 | 65.73195802308402 | 91.69380000550801 |
| SE_232_2_9 | SE_232 | SE_232_2 | TTTCAAAGTAGCTAGAAGGG | 15.172875783636185 | 36.977521931454156 | 25.675698428498144 | 9.109159426964803 | 42.42378050917627 | 60.641706907886614 | 4.30464245105859 | 68.06545132004533 | 39.07135222430747 | 3.6450347241344634 | 83.33873249355297 | 112.5860582346111 |
| SE_233_1_10 | SE_233 | SE_233_1 | GTTTTAAAACCATATGACAT | 1.083776841688299 | 8.994532361705065 | 12.837849214249072 | 3.9039254686992013 | 3.1817835381882205 | 8.663101098684088 | 9.685445514881827 | 5.67212094333711 | 0.0 | 4.374041668961356 | 0.0 | 1.1606810127279494 |
| SE_233_1_13 | SE_233 | SE_233_1 | GTAAGCTGGAAAGAAACAAG | 328.3843830315546 | 447.7278331159855 | 465.12515230087024 | 353.95590916206095 | 361.6627288407277 | 372.51334724341575 | 301.3249715741013 | 446.2068475425193 | 388.0495664095992 | 610.907819764936 | 566.9381379490998 | 331.95476964019355 |
| SE_233_1_14 | SE_233 | SE_233_1 | AGTAAGCTGGAAAGAAACAA | 83.45081680999901 | 154.90583511825392 | 141.21634135673 98 | 105.40598765487843 | 114.54420737457792 | 248.6310015322333 | 40.89410328505 66 | 178.67180971511897 | 209.56452556674006 | 38.63736807582531 | 188.97937931636656 | 408.5597164802382 |
| SE_233_1_15 | SE_233 | SE_233_1 | GAGTAAGCTGGAAAGAAACA | 340.30592829012585 | 468.71507529329733 | 458.21246426242845 | 429.43180155691215 | 425.2983996044921 | 431.42243471446756 | 338.99059302086397 | 421.62765678 80585 | 498.15974085992025 | 508.1178405443442 | 509.4226746789012 | 403.9169924293264 |
| SE_233_1_4 | SE_233 | SE_233_1 | GAGCAGAGATGGACTTTTGG | 911.4563238598594 | 1140.3068249672756 | 1143.556106930802 | 1069.675578423581 | 1068.0186743185127 | 973.7325634920915 | 964.2399090371241 | 962.3698533861964 | 1009.639260887218 | 1019.8807158128228 | 1022.3667042518962 | 1255.8568557716412 |
| SE_234_1_13 | SE_234 | SE_234_1 | TTTCTAGGCCCAGAGCAGCG | 33.59708209233727 | 55.96597913949819 | 64.18924607124536 | 57.25757354092162 | 89.08939906927017 | 57.17646725131495 | 61.3411549725845 | 86.02716764061283 | 53.27911666951019 | 139.2403264619365 | 124.42120625798047 | 54.55200759821362 |
| SE_234_1_2 | SE_234 | SE_234_1 | AGACTCTTACTTAAGTTGTG | 52.02128840103835 | 81.9501837399795 | 73.07698783495626 | 29.93009526002721 | 46.6661585600939 | 100.4919727447354 | 47.35106696164449 | 52.93979547114636 | 155.39742361940472 | 84.56480559991955 | 52.820323411406804 | 117.2287822855229 |
| SE_234_1_21 | SE_234 | SE_234_1 | TGCCCCCTCCACACACAAAA | 163.65030309493312 | 210.8718142577521 | 191.58021135110155 | 197.79889041409285 | 166.51333849851687 | 181.92512307236584 | 186.175786008284 | 243.90120056349573 | 181.14899667633463 | 160.3815278619164 | 213.62886357502308 | 71.96222278913287 |

| | | | | | | | | | | | | | | | |
|---|---|---|---|---|---|---|---|---|---|---|---|---|---|---|---|
| SE_234_1_34 | SE_234 | SE_234_1 | CTTTTCCTAGAGGCAAAGGG | 5.4188842084414945 | 11.992709815606755 | 11.850322351614528 | 6.506542447832002 | 8.484756101835254 | 25.12299318618385 | 10.761606127646475 | 11.34424188667422 | 4.439926389125849 | 7.290069448268927 | 0.0 | 0.0 |
| SE_234_1_6 | SE_234 | SE_234_1 | GTTGATCTCCCTAGGTTGGT | 162.56652625324483 | 216.86816916555549 | 211.33074860379241 | 165.26617817493286 | 219.5430641349872 | 97.02673230526177 | 152.81480701257993 | 180.56251669623134 | 168.71720278678225 | 180.0647153722425 | 179.58909959878315 | 85.89039494186825 |
| SE_235_1_10 | SE_235 | SE_235_1 | GAGATCTCAGGAGGTCTGAG | 163.65030309493312 | 303.81531532870446 | 312.0584885925159 | 295.3970271315729 | 272.5727897714575 | 207.9144263684181 | 191.55658907210724 | 260.91756339350707 | 231.7641575123693 | 196.831875103261 | 239.45213279837753 | 341.24021774201714 |
| SE_235_1_12 | SE_235 | SE_235_1 | CTAAACTTGTCCTTGATAAG | 199.414938870647 | 281.8286806667587 | 260.7070917355196 | 209.51066682019047 | 177.11928362581094 | 172.39571186381335 | 216.30828316569415 | 169.21827480955713 | 248.63587779104753 | 298.89284737902597 | 184.28423945757487 | 224.01143545649424 |
| SE_235_1_32 | SE_235 | SE_235_1 | TCACAGTTCCAACAAGGGTC | 6.5026610501297935 | 10.993317330972857 | 8.887741763710896 | 14.314393385230405 | 2.1211890254588135 | 0.0 | 1.0761606127646475 | 0.0 | 10.655823333902038 | 0.0 | 3.521354894093787 | 1.1606810127279494 |
| SE_235_1_6 | SE_235 | SE_235_1 | TCGGTATGTATACATCTACA | 1322.2077468597247 | 1430.1306455111055 | 1368.712231611478 | 1353.3608291490564 | 1345.894436653617 | 1467.5293261170843 | 1053.5612398965898 | 1210.052467911917 | 1411.89659174202 | 1168.598132557509 | 1139.7452007216891 | 1035.3274633533308 |
| SE_235_1_7 | SE_235 | SE_235_1 | TCTGGAAACGAGATCTCAGG | 487.6995787597345 | 485.70474753207355 | 536.2270864105574 | 503.60638546219695 | 476.2069362155036 | 547.5079894368344 | 498.26236371003176 | 440.5347265991822 | 468.85622669168964 | 492.07968775815255 | 448.3858565146089 | 776.4955975149982 |
| SE_235_2_30 | SE_235 | SE_235_2 | GCAAGTTGCAGCTCTGTCCC | 5824.216747232918 | 5203.8366674886975 | 5208.216673534585 | 5628.159217374682 | 5228.730947755976 | 5595.496999640052 | 5896.283997337504 | 5249.547933058495 | 5350.999284174473 | 5140.22796797442 | 4696.313643756414 | 5275.295202848531 |
| SE_235_2_32 | SE_235 | SE_235_2 | TTGGCTGCAGGGAAATCGCC | 149.56120415298525 | 54.96658665486429 | 39.50107450538176 | 118.41907255054244 | 23.33307928004695 | 405.43313141841526 | 202.31819519975372 | 41.59555358447214 | 174.93309973155846 | 6.561062503442034 | 14.085419576375148 | 59.19473164912542 |
| SE_235_2_36 | SE_235 | SE_235_2 | AGGCCTCCCTTCTCAGGCGT | 430.2594061502547 | 554.6628289718124 | 505.61375366888655 | 369.57161103685775 | 467.7221801136684 | 507.6577243828875 | 417.5503177526832 | 303.4584704685354 | 404.0333014104522 | 467.2934516340382 | 319.2695103978367 | 423.64856964570157 |
| SE_235_2_44 | SE_235 | SE_235_2 | TGGAGAAGCATCATTGTAGA | 593.9097092451879 | 642.6093676195952 | 688.3062232562772 | 615.5189155649074 | 641.659680201291 | 711.2406002019636 | 693.047434620433 | 596.5180525409528 | 639.3494000341223 | 773.4763684613331 | 570.4594928431935 | 1037.6488253787868 |
| SE_235_2_9 | SE_235 | SE_235_2 | ATTAAAAAAAAAATGCTCCA | 0.0 | 0.0 | 0.987526862634544 | 2.602616979132801 | 0.0 | 0.0 | 0.0 | 6.617474433893295 | 0.0 | 0.0 | 4.695139858791716 | 0.0 |
| SE_235_3_10 | SE_235 | SE_235_3 | GTAATAGTCGTGTATCAGTC | 306.7088461977886 | 423.742413484772 | 421.6739703449503 | 230.33160265325287 | 384.99580812077465 | 168.93047142433971 | 276.5732774805144 | 350.72614499634466 | 352.5301552965924 | 323.67908350314036 | 387.34903835031656 | 168.29874684555267 |
| SE_235_3_11 | SE_235 | SE_235_3 | TCAGGATTCTCCAAAGAAAT | 117.04789890233629 | 173.89429232629794 | 193.55526507637063 | 106.70729614444484 | 173.9375000876227 | 255.56148241118058 | 80.71204595734856 | 154.09261896065814 | 184.70093778763533 | 187.3547848205114 | 63.384388093688166 | 595.4293595294381 |
| SE_235_3_24 | SE_235 | SE_235_3 | GCCTGGACCGACAGGAAGCG | 2071.097544466339 | 2143.6968795397074 | 2245.636085630953 | 2044.355637108815 | 2159.370427917072 | 2322.577404557204 | 1792.8835808659028 | 1653.4232549827675 | 2150.7003428925614 | 1595.796202226068 | 1624.5183911419338 | 2385.199481155936 |
| SE_235_3_5 | SE_235 | SE_235_3 | GCCAGGCCTTTGAATCAGCT | 1141.2170142977789 | 1108.326265458991 | 1056.6537430189621 | 1145.1514708184325 | 1169.8357475405357 | 1033.5079610730115 | 1147.1872132071142 | 907.5393509339376 | 923.5046889381766 | 1494.46423689513 | 914.3784874996867 | 1431.1196886935616 |
| SE_235_3_9 | SE_235 | SE_235_3 | GTAGTAGGATCACTGGAGGC | 561.3964039945388 | 733.5540837212798 | 661.6429979651446 | 511.4142363995954 | 677.7198936340909 | 494.6630727348614 | 493.9577212589732 | 905.6486439528252 | 737.9157658727161 | 632.7780281097429 | 981.2842304874687 | 785.7810456168218 |
| SE_236_1_10 | SE_236 | SE_236_1 | ACTACAGGCATGTATCACCA | 2252.088277028285 | 2008.7788941141314 | 2105.407271136848 | 2333.246121792556 | 2279.2176078554953 | 1898.0854507216836 | 2139.407298176119 | 2616.73846185952 | 1872.760950933283 | 2321.8871192736533 | 2224.3225081025757 | 1706.2010887100857 |
| SE_236_1_2 | SE_236 | SE_236_1 | CATCTCAAGGATGGAAACAT | 431.34318299194297 | 369.7752193145416 | 376.24773466376126 | 435.9383440047441 | 344.6932166370572 | 827.3261549243304 | 429.38808449309437 | 355.45291244912556 | 321.4506705727115 | 258.06845846872 | 370.91604884454557 | 284.3668481183476 |
| SE_236_1_25 | SE_236 | SE_236_1 | ACGAAAGCATTCTAAAGTGG | 6379.110490177327 | 7430.483123253018 | 7568.405875231146 | 6867.004899441895 | 7512.190933662388 | 6343.122624456489 | 6108.287638052139 | 5731.6782132421495 | 6508.932086458494 | 5261.972127760511 | 4225.625872912545 | 5412.255562350429 |
| SE_236_1_5 | SE_236 | SE_236_1 | ACTGTAACTTAACAACCCAT | 1399.155902619594 | 1310.203547355038 | 1336.123845144538 | 1283.0901707124708 | 1323.6219518862997 | 1398.2245173276117 | 1331.2106779898688 | 1476.642152248761 | 1376.377180629013 | 1246.6018756539866 | 1099.8365119219595 | 1182.7339519697805 |
| SE_236_1_9 | SE_236 | SE_236_1 | ACTGCCACTAGATATCCATG | 1737.2942772263432 | 1803.9034347641825 | 1730.1470633357212 | 1678.6879515406565 | 1811.4954277418267 | 1885.9571091835257 | 1728.313944100024 | 1773.4831482834031 | 1928.7040234362687 | 1887.398980156825 | 1680.8600694474344 | 1366.1215519807965 |
| SE_236_2_0 | SE_236 | SE_236_2 | TCAAAATTTACATTTATTTC | 49.85373471766175 | 65.95990398583714 | 58.264084895438096 | 41.641871666124814 | 79.5445884547055 | 34.65240439473635 | 30.13249715741013 | 138.021609621203 | 47.063219724734 | 30.61829168272949 | 203.06479889274172 | 66.15881772549312 |
| SE_236_2_1 | SE_236 | SE_236_2 | TTTATTTCAGGTCACTTCAG | 1.083776841688299 | 5.9963549078033775 | 5.925161175807264 | 6.506542447832002 | 2.1211890254588135 | 0.0 | 1.0761606127646475 | 0.0 | 39.07135222430747 | 0.7290069448268927 | 0.0 | 0.0 |
| SE_236_2_12 | SE_236 | SE_236_2 | TCCATGCTAGAATAACGCAG | 847.5134902002497 | 829.4957622461338 | 917.4124553874914 | 761.2654663963442 | 722.264863168726 | 1263.08014018814 | 825.4151899904846 | 811.1132948972067 | 1244.955359510888 | 679.434472578664 | 1093.96758709847 | 1332.461802611686 |
| SE_236_2_2 | SE_236 | SE_236_2 | TAATTCTTGAAGCAATCAGG | 31.429528408960667 | 47.97083926242702 | 37.526020780112674 | 76.77720088441762 | 26.51486281823517 | 66.70587845986748 | 64.56963676587885 | 45.37696754669688 | 56.831057780810866 | 174.23265981362735 | 10.564064682281362 | 172.94147089646447 |
| SE_236_2_3 | SE_236 | SE_236_2 | TGATAATTCTTGAAGCAATC | 123.55055995246607 | 177.89186226483352 | 174.7922546863143 | 187.38842249756166 | 134.69550311663465 | 156.80212988618197 | 124.83463108069911 | 55.775855942814914 | 195.35676112153735 | 69.98466670338169 | 38.73490383503166 | 123.03218734916264 |
| SE_236_3_10 | SE_236 | SE_236_3 | CAATTTACGAGCTGTTCCAT | 134.38832836934907 | 168.89732990312845 | 150.1040831204507 | 128.82954046707366 | 100.75647870929365 | 161.133680435524 | 107.61606127646475 | 133.2948421684221 | 150.95749723027888 | 242.75931262735526 | 204.23858385743966 | 254.18914178742094 |
| SE_236_3_11 | SE_236 | SE_236_3 | GTCACATTTGAAGAAACCGA | 285.0333093640226 | 350.7867621064976 | 295.27053192772865 | 296.6983356211393 | 253.4820885423282 | 414.9625426269678 | 300.24881096133663 | 341.2726100907828 | 317.0107441835856 | 424.2820418892515 | 239.45213279837753 | 518.8244126893934 |
| SE_236_3_2 | SE_236 | SE_236_3 | CTTCCCCTTGATTCAGTGGT | 55.27261892610324 | 81.9501837399795 | 65.1767729338799 | 39.03925468699201 | 98.63528968383483 | 25.12299318618385 | 62.417315540349556 | 77.51898622560716 | 65.71091055906257 | 110.80905561368769 | 43.43004369382337 | 134.63899747644214 |
| SE_236_3_3 | SE_236 | SE_236_3 | GAACCGACCACTGAATCAAG | 1031.7555532872605 | 1083.3414533431435 | 1032.953098315733 | 1044.9507171218195 | 1105.1394822640418 | 1072.49191601709 | 1004.0578517094161 | 1505.9481104560027 | 1055.8144953341268 | 984.888382461132 | 1367.4594838730873 | 992.3822658823967 |
| SE_236_3_9 | SE_236 | SE_236_3 | CGATGGAACAGCTCGTAAAT | 6194.868427090317 | 6242.205459023316 | 6213.519019696551 | 6401.136460177124 | 6445.232853856604 | 7905.946062659098 | 6058.784249864965 | 6504.0320150265525 | 6469.8607342341875 | 6109.078197649361 | 6528.591973649881 | 7076.672134602308 |
| SE_236_4_1 | SE_236 | SE_236_4 | ACGCACTCTTGTGAGACCTG | 8890.221432369115 | 7626.364050241262 | 7713.572324038423 | 8812.461091343665 | 7888.701985681328 | 8873.61445538211 | 9818.889430864643 | 8545.050201137356 | 8188.112246825891 | 8955.12131025355 | 9348.023458854306 | 7538.6231776680315 |
| SE_236_4_11 | SE_236 | SE_236_4 | ACTTGTTCCTGGGTGTTCAG | 187.49339361207572 | 219.86634661945715 | 228.11870526857967 | 174.37533760189766 | 117.72599091296415 | 250.36362175197013 | 200.16587397422444 | 333.7097821663333 | 159.83735000853056 | 110.0800486688608 | 328.65979011542015 | 175.26283292192036 |
| SE_236_4_12 | SE_236 | SE_236_4 | ACTCTCAGACATCTCAAGGA | 79.11570944324582 | 120.92649064070145 | 112.57806234033802 | 108.00860463401123 | 133.63490860390525 | 153.33688944670834 | 69.95043982970209 | 70.90151179171387 | 92.35046889381766 | 59.7785694758052 | 28.170839152750297 | 141.60308355280984 |
| SE_236_4_18 | SE_236 | SE_236_4 | TCTGTTTCCTCCTTCCAGAA | 72.61304839311603 | 67.95868895510495 | 73.07698783495626 | 100.20075369661284 | 49.84794209828212 | 80.566840217762 | 86.0928490211718 | 80.35504669727572 | 67.4868811147129 | 36.45034724134463 | 28.170839152750297 | 81.24767089095646 |
| SE_236_4_4 | SE_236 | SE_236_4 | AAGGCTGCGGAAGGAGTCAG | 3222.068550339313 | 3274.0097796606437 | 3242.050690029208 | 3179.0966400107163 | 3417.2355200141487 | 3628.1067401288956 | 3111.180331502596 | 3779.523255243628 | 3203.8508823932125 | 3925.702397892817 | 3727.9410478806226 | 3795.4269116203945 |
| SE_237_1_2 | SE_237 | SE_237_1 | GTATGGAGCTTCTAGCCCGC | 604.7474776620708 | 599.6354907803377 | 593.5036444433609 | 537.4404061909233 | 526.0548783137857 | 495.5293828447298 | 595.11681885885 | 664.583503860998 | 523.023328639025 | 249.3203751307973 | 591.5876222077562 | 581.5011873767027 |
| SE_237_1_24 | SE_237 | SE_237_1 | GCCATCAGAATAGCAAGAGG | 1231.1704921579076 | 1399.1494784874546 | 1464.5023372870287 | 1315.6228829516308 | 1292.8647110171469 | 1488.3207687539261 | 1281.7072898026952 | 1268.6643843264003 | 1656.9805284217668 | 1813.040271784482 | 1230.1266430034295 | 1666.7379342773354 |
| SE_237_1_5 | SE_237 | SE_237_1 | AGTTTGTTTCTAAAGATAGA | 3.2513305250648967 | 5.9963549078033775 | 17.77548352742179 | 3.9039254686992013 | 16.969512203670508 | 1.7326202197368175 | 0.0 | 0.945353490556185 | 7.991867500426528 | 0.0 | 1.173784964697929 | 6.964086076367696 |
| SE_237_1_7 | SE_237 | SE_237_1 | CTAAGATCCTTCCTTCTACA | 238.43090517142576 | 261.8408309740808 | 276.50752153767235 | 312.3140374959361 | 260.90625013143404 | 367.3154865842053 | 207.69899826357695 | 257.1361494312823 | 430.67285974520735 | 78.00374309647752 | 489.4683302790364 | 637.2138759876442 |
| SE_237_1_9 | SE_237 | SE_237_1 | GTTTGGTCAAAACCACAAAA | 6.5026610501297935 | 6.9957473924372735 | 29.62580587903632 | 6.506542447832002 | 11.666539640023474 | 10.395721318420904 | 9.685445514881827 | 17.961716320567515 | 19.535676112153734 | 5.103048613788249 | 8.216494752885502 | 112.5860582346111 |
| SE_237_2_1 | SE_237 | SE_237_2 | GGACAGATAATAGAGCTGTG | 834.5081680999901 | 923.4386558017201 | 943.0881538159896 | 877.0819219677539 | 954.5350614564661 | 784.0106494309099 | 882.4517024670109 | 972.7687417823144 | 1248.5073006221887 | 830.3389101578307 | 1257.123697191482 | 1094.5221950024563 |
| SE_237_2_13 | SE_237 | SE_237_2 | TGGCGTGAGACACCATGCCG | 221.09047570441297 | 193.88214201897586 | 190.592684488467 | 184.78580551842887 | 230.14900926228125 | 97.89304241513018 | 216.30828316569415 | 327.09230773244 | 115.43808611727208 | 153.09145841364744 | 420.2150173618586 | 52.23064557275772 |
| SE_237_2_26 | SE_237 | SE_237_2 | TGTGTGGCCTTCATCCCACA | 1874.9339361207572 | 1544.0613887593695 | 1535.604271396716 | 1574.5832723753444 | 1527.2560983303458 | 1477.058737325637 | 1807.9498294446078 | 1386.8335706459234 | 1539.7664717488444 | 1414.2734729641718 | 1351.0264943673162 | 1246.5714076698177 |
| SE_237_2_35 | SE_237 | SE_237_2 | GCTCCTTCCCAGGAACCTGA | 35.76463577571386 | 65.95990398583714 | 79.98967587339807 | 29.93009526002721 | 82.72637199289373 | 58.04277736118338 | 16.142409191469714 | 6.617474433893295 | 110.99815972814622 | 2.187020834480678 | 0.0 | 10.446129114551544 |
| SE_237_2_6 | SE_237 | SE_237_2 | TGTTGTCCTAGACACTTAGG | 251.43622727168534 | 280.82928818212486 | 269.5948334992305 | 239.44076208021767 | 252.42149402959882 | 330.0641518598637 | 308.8580958634538 | 369.63321480746833 | 277.051406681453 | 300.35086126867975 | 246.49484258656508 | 246.06437469832528 |
| SE_237_3_12 | SE_237 | SE_237_3 | CTAGGTTAGCCCCATGGCAG | 107.2939073271416 | 130.9204154870404 | 171.82967409841066 | 137.93869989403845 | 134.69550311663465 | 277.2192351578908 | 95.77829453605362 | 85.08181415005664 | 133.19779167377547 | 158.9235139722626 | 130.29013108147012 | 774.1742354895423 |
| SE_237_3_17 | SE_237 | SE_237_3 | GTTTCTTGCAGTCAGAGCTA | 7.586437891818092 | 20.98724217731182 | 5.925161175807264 | 39.03925468699201 | 1.0605945127294067 | 10.395721318420904 | 24.75169409358689 | 3.78141396222474 | 7.103882222601358 | 10.206097227576498 | 21.128129364562724 | 0.0 |
| SE_237_3_2 | SE_237 | SE_237_3 | TTTGCAGGTGTAGGGAACAA | 490.9509092847994 | 603.6330607188733 | 567.8279460148628 | 562.165267492685 | 463.47980206275076 | 404.5668213085469 | 588.6598551822622 | 529.3979547114636 | 651.7811939236747 | 480.41557664092227 | 558.7216431962142 | 444.5408278748046 |
| SE_237_3_22 | SE_237 | SE_237_3 | GTTGTGATGTCAGAAAGAGA | 249.26867358830876 | 208.8730292884843 | 235.0313933070215 | 271.9734743193777 | 294.8452745387751 | 278.0855452677592 | 287.33488360816085 | 330.87372169466477 | 276.1634214036278 | 272.64859736525784 | 431.95286700883787 | 255.34982280014887 |
| SE_237_3_4 | SE_237 | SE_237_3 | AATCCAAGCAGGTCAGGCAT | 212.42026097090658 | 291.8226055130977 | 310.0834348672468 | 184.78580551842887 | 190.90701229129323 | 199.251325269734 | 222.76524684228204 | 107.7702979234051 | 277.93939195927817 | 190.99981954464587 | 59.86303319959438 | 185.70896203647192 |
| SE_237_4_1 | SE_237 | SE_237_4 | ATACTGTAAGGTCTTCCCAG | 39.01596630077876 | 90.94471610168455 | 97.76515940081985 | 37.737946197425615 | 93.3323171201878 | 90.96256153618292 | 32.28481838293943 | 25.524544245016994 | 126.09390945117411 | 34.26332640686395 | 31.692194046844083 | 44.10587848366208 |
| SE_237_4_10 | SE_237 | SE_237_4 | ATGTGCCGCTAGGCATTCCG | 1521.6226857303716 | 1502.086904404746 | 1548.442120610965 | 1519.9283158135556 | 1322.5613573735702 | 1283.0052727151133 | 1397.9326359812771 | 1740.3957761139366 | 1428.7683120206982 | 1614.0213758467403 | 1199.6082339212835 | 1391.6565342608114 |
| SE_237_4_14 | SE_237 | SE_237_4 | GGGATTTCAAAGCCAGTGGA | 43.351073667531956 | 96.94107100948793 | 71.10193410968716 | 71.57196692615203 | 42.42378050917627 | 82.29946043749882 | 57.03651247652632 | 33.08737216946648 | 111.88614500597139 | 34.26332640686395 | 37.56111887033373 | 30.177706330926686 |
| SE_237_4_5 | SE_237 | SE_237_4 | GAATTTTCTGCATTACCGTG | 5.4188842084414945 | 14.990887269508443 | 22.713117840594514 | 7.807850937398403 | 26.51486281823517 | 0.0 | 1.0761606127646475 | 6.617474433893295 | 2.6639558334755096 | 1.4580138896537853 | 1.173784964697929 | 0.0 |
| SE_237_4_7 | SE_237 | SE_237_4 | GTATACATCTGCCACTCGTC | 600.4123702953176 | 552.6640440025446 | 603.3789130697064 | 504.90769395176335 | 563.175686259315 | 397.6363404295996 | 606.9545855992611 | 509.54553140978373 | 731.6998689279399 | 516.13691693744 | 440.1693617617234 | 626.7677468730927 |
| SE_238_1_13 | SE_238 | SE_238_1 | AGAGATATGGAGGTGTGAGG | 461.68893455921534 | 525.6804469174294 | 612.2666548334173 | 459.36189681693935 | 536.6608234410799 | 534.5133377888081 | 544.5372700589116 | 447.1522010330755 | 747.683603928793 | 565.7093891856687 | 358.0044142328683 | 655.7847721912914 |
| SE_238_1_27 | SE_238 | SE_238_1 | ACTAGCAAGGGATAGAGCTG | 1075.1066269547925 | 1146.303179875079 | 1125.78062340338 | 1022.8284727991908 | 876.05106751449 | 833.3903256934092 | 1071.8559703135888 | 747.7746110299423 | 1219.2037864539582 | 728.2779378820658 | 893.250358135124 | 1115.4144532315595 |
| SE_238_1_32 | SE_238 | SE_238_1 | ACTTCCTCATCTCCAGCACG | 309.9601767228535 | 330.79891241381966 | 350.5720362352631 | 303.2048780689713 | 351.0567837134336 | 314.4705698822324 | 302.40113218686594 | 399.88452650526625 | 385.38561057612367 | 463.64841690990374 | 825.1708301826441 | 313.38387343654637 |
| SE_238_1_35 | SE_238 | SE_238_1 | AGGAAGTGAGGAAATACCCA | 1673.3514435667335 | 1758.9307729556572 | 1786.4360945058902 | 1759.3690778937735 | 1993.9176839312847 | 1598.3421527072142 | 1912.3374088827786 | 1768.7563808306222 | 1685.3960573121722 | 2370.7305845770547 | 2353.438854219348 | 1573.8834532590995 |
| SE_238_1_8 | SE_238 | SE_238_1 | GGAAGCTTTGAGGGAGAGAG | 365.2327956489567 | 426.7405909386737 | 395.99827191645215 | 456.75927983780656 | 405.1471038626334 | 258.1604127407858 | 403.5602297867428 | 341.2726100907828 | 609.1579005880665 | 378.3546043651573 | 414.34609253836896 | 322.66932153836996 |
| SE_238_2_23 | SE_238 | SE_238_2 | TAGACTCACATCCGACACGG | 2303.0257885876354 | 2235.6409881260256 | 2237.7358707298768 | 2197.9100388776505 | 2256.9451230881778 | 2177.9036162091797 | 2438.579948524691 | 2382.290796201586 | 2151.5883281703864 | 2464.043473514897 | 2562.372577935579 | 2175.116217852177 |
| SE_238_2_5 | SE_238 | SE_238_2 | CATATCTCCCTCAACAACCA | 881.110572292587 | 815.5042674612592 | 841.3728869646316 | 799.0034125937699 | 748.7797259869611 | 830.791395363804 | 846.9384022457775 | 640.9496665970935 | 763.667338929646 | 735.5680073303347 | 573.9808477372873 | 1147.913521587942 |
| SE_238_2_6 | SE_238 | SE_238_2 | GTAGTCCCAGCTACCCAGGT | 15.172875783636185 | 27.982989569749094 | 16.78795666478725 | 16.917010364363207 | 11.666539640023474 | 10.395721318420904 | 18.294730416999005 | 0.0 | 0.8879852778251698 | 3.6450347241344634 | 16.432989505771005 | 1.1606810127279494 |
| SE_238_2_7 | SE_238 | SE_238_2 | ATATTCATATGCAAGACTGT | 45.51862735090855 | 32.97995199291857 | 44.43870881855448 | 42.943180155691216 | 28.63605184369398 | 17.326202197368175 | 57.03651247652632 | 69.95615830115769 | 47.063219724734 | 26.244250013768134 | 80.99116256415711 | 121.87150633643469 |
| SE_238_2_8 | SE_238 | SE_238_2 | TTAGCTACAAGGAGAGACCT | 360.89768828220355 | 317.806810113579 | 342.6718213341868 | 368.27030254729135 | 302.2694361278809 | 279.818165487496 | 407.86487223780136 | 259.0268564123947 | 504.37563780469645 | 290.14476404110326 | 211.28129364562722 | 247.22505571105322 |
| SE_238_3_12 | SE_238 | SE_238_3 | GGCAGGTATTTCCTGAGCCA | 649.182328171291 | 594.6385283571682 | 617.20428914659 | 668.8725636371298 | 743.4767534233141 | 681.7860564664377 | 549.9180731227349 | 554.9224989564806 | 675.7567964249542 | 500.82777109607525 | 692.5331291717781 | 306.41978736017865 |
| SE_238_3_22 | SE_238 | SE_238_3 | ACTAATTCTAGGATCAGCAA | 120.29922942740117 | 154.90583511825392 | 160.96687860943067 | 180.88188004972966 | 101.81707322202305 | 33.78609428486794 | 156.0432888508739 | 216.48594933736638 | 148.29354139680336 | 237.656264013567 | 181.936669528179 | 134.63899747644214 |
| SE_238_3_3 | SE_238 | SE_238_3 | GGCTGTGTGTGGAGAGCAGA | 766.2302270736274 | 1016.3821568726725 | 1036.9032057662712 | 836.7413587911955 | 981.0499242747012 | 702.5774991032795 | 843.7099204074837 | 1041.779546592916 | 903.9690128260229 | 823.0488407095618 | 1435.5390118255673 | 792.7451316931895 |
| SE_238_3_34 | SE_238 | SE_238_3 | CTGGAATGCTCAATGTGAGG | 451.93494298402067 | 434.73573081574483 | 510.5513879820593 | 530.9338637430914 | 476.2069362155036 | 245.16576109275968 | 443.3781724590348 | 390.4309915997044 | 766.3312947631215 | 377.6255974203304 | 204.23858385743966 | 625.6070658603647 |
| SE_238_3_35 | SE_238 | SE_238_3 | GCTCAATGTGAGGAGGTAGA | 1146.6358985062202 | 1175.285561929462 | 1216.6330947657582 | 1128.2344604540692 | 1292.8647110171469 | 1200.7058122776145 | 1236.50854406658 | 1349.9647845142322 | 1169.4766108957485 | 803.3656531992357 | 978.9366605580728 | 998.1856709460366 |
| SE_238_4_1 | SE_238 | SE_238_4 | GAACAGTTCTAGATTTTGCA | 114.88034521895969 | 125.92345306387092 | 152.07913684571977 | 143.14393385230406 | 201.5129574185873 | 170.66309164407653 | 160.34793130193248 | 187.17999113012462 | 184.70093778763533 | 89.6678542137078 | 204.23858385743966 | 84.7297139291403 |
| SE_238_4_14 | SE_238 | SE_238_4 | GCTGGGGTCAGGGTGAAGGT | 3087.6802219699634 | 3038.1531532870445 | 2988.2562863321305 | 3089.3063542306345 | 3057.6939801988797 | 2600.662949824963 | 2747.438044388145 | 3146.136416570984 | 3298.865307120506 | 2501.951834645896 | 3069.4476826850846 | 2487.3394102759958 |
| SE_238_4_21 | SE_238 | SE_238_4 | CTTGGGAAGATGCTCCAGAG | 406.4163156331121 | 518.6846995249921 | 496.72601190517565 | 495.7985345247986 | 521.8125002628681 | 673.989265477622 | 436.92120878244685 | 582.33775018261 | 421.79300696695566 | 191.72882648947277 | 414.34609253836896 | 393.4708633147749 |
| SE_238_4_24 | SE_238 | SE_238_4 | AGGTCCCTCCCTGAAGTCAG | 214.5878146542832 | 210.8718142577521 | 232.06881271911786 | 264.1656233819793 | 182.42225618945795 | 173.26202197368175 | 294.8680078975134 | 247.68261452572048 | 193.580790565887 | 220.1600973377216 | 273.4918967746175 | 269.2779949528843 |
| SE_238_4_36 | SE_238 | SE_238_4 | GTTGAAAAGTTAGAGCCTCA | 148.47742731129696 | 154.90583511825392 | 156.02924429625796 | 170.47141213319847 | 152.72560983303458 | 124.74865582105086 | 163.5764131402264 | 134.24019565897828 | 154.50943834157954 | 172.04563897914667 | 104.46686185811568 | 167.1380658328247 |
| SE_239_1_19 | SE_239 | SE_239_1 | GCCAGCCGCCTCCTTTTACA | 587.407048195058 | 689.5808143973884 | 757.4331036406953 | 704.0078928554226 | 845.2938266453372 | 925.2191973394605 | 609.1069068247905 | 695.7801690493521 | 545.2229605846543 | 842.003021275061 | 1014.1502094990107 | 779.977640553182 |
| SE_239_1_2 | SE_239 | SE_239_1 | GGCGGAAGATGAGAATGGGA | 1450.093414178944 | 1697.9678313929896 | 1694.5960962808776 | 1469.177284720466 | 1721.3448941598272 | 1597.4758425973457 | 1546.4428005427985 | 1762.1389063967288 | 1587.7176767514036 | 1653.3877508673925 | 1579.9145624834125 | 1828.0725950465203 |
| SE_239_1_5 | SE_239 | SE_239_1 | TCTGGGGCGGAAGATGAGAA | 1008.9962396118062 | 1259.2345306387092 | 1226.5083633921038 | 1270.0770858168069 | 1219.6836896388177 | 1214.566774035509 | 1032.0380276412968 | 1124.9706537618601 | 1349.737622294258 | 1444.1627577020743 | 1207.824728674169 | 1452.0119469226647 |
| SE_239_1_7 | SE_239 | SE_239_1 | CATGGCGTGTAGAGGAGACG | 15149.032693119043 | 13067.056736588193 | 13033.379533050713 | 14064.542155233656 | 12761.073177160222 | 13177.443081208365 | 14995.221978262598 | 13191.462607221005 | 13476.064576274777 | 15093.359785695986 | 12851.771578477625 | 13877.102188175364 |

| | | | | | | | | | | | | | | | |
|---|---|---|---|---|---|---|---|---|---|---|---|---|---|---|---|
| SE_239_1_9 | SE_239 | SE_239_1 | CGATGGGCTTTAGAACAAAG | 3422.567266051648 | 3973.584518904371 | 4097.2489530707235 | 3751.6723754199325 | 3970.865855658899 | 3595.1869559538964 | 3486.760385357458 | 4497.0465545757725 | 4269.433215783416 | 4083.896904920253 | 4374.696563429182 | 4946.822476246521 |
| SE_239_2_15 | SE_239 | SE_239_2 | GACTAAACTCTAATGATAGG | 174.48807151181612 | 199.87849692677923 | 207.38064115325426 | 192.59365645582727 | 219.5430641349872 | 238.2352802138124 | 214.15596194016484 | 200.41493999791123 | 156.28540889722987 | 65.61062503442034 | 228.88806811609615 | 175.26283292192036 |
| SE_239_2_17 | SE_239 | SE_239_2 | TCTTAGAAGAAACAAATGTG | 246.01734306324386 | 242.85237376603678 | 281.445155850845 | 340.94282426639694 | 248.17911597868118 | 448.7486369118357 | 315.3150595400417 | 272.2618052801813 | 233.54012806801964 | 492.80869470297944 | 255.88512230414852 | 205.44053925284706 |
| SE_239_2_2 | SE_239 | SE_239_2 | GACAATGGCTTGGACAAGGG | 1777.3940203688103 | 1562.0504534827796 | 1692.6210425556085 | 1673.482717582391 | 1796.6471045636151 | 1698.8341254519496 | 1777.8173322871976 | 1972.952734790758 | 1876.3128920445838 | 1663.593848094969 | 1983.6965903395 | 1604.0611595900261 |
| SE_239_2_4 | SE_239 | SE_239_2 | CTTTTCCTACAGAAATGGAG | 7.586437891818092 | 12.992102300240651 | 12.837849214249072 | 11.711776406097604 | 24.393673792776355 | 0.0 | 8.60928490211718 | 2.836060471668555 | 13.319779167377547 | 8.748083337922711 | 16.432989505771005 | 4.642724050911798 |
| SE_239_2_9 | SE_239 | SE_239_2 | ACATAGTAAGAATAGCAATG | 1072.939073271416 | 1191.2758416836043 | 1198.8576112383364 | 1156.86324722453 | 1387.257622650064 | 1186.84485051972 | 1173.0150679134658 | 1523.9098267765703 | 1136.6211556162173 | 1565.1779105433386 | 2094.0323770211053 | 834.5296481513957 |
| SE_239_3_0 | SE_239 | SE_239_3 | TGGTGGTTGGAGGAAGCAGA | 3.2513305250648967 | 4.996962423169481 | 9.87526862634544 | 5.205233958265602 | 5.3029725636470335 | 11.262031428289314 | 5.380803063823238 | 1.89070698111237 | 0.0 | 5.832055558615141 | 1.173784964697929 | 16.249534178191293 |
| SE_239_3_12 | SE_239 | SE_239_3 | CAAGGGACCTGAGAGAAACG | 3222.068550339313 | 3489.8785563415654 | 3394.129826874928 | 3184.301873968982 | 3542.3856725162186 | 3028.620144099957 | 2949.7562395878986 | 3407.0539799644907 | 3310.4091157322327 | 3102.653557183255 | 3070.6214676497825 | 3556.326622998437 |
| SE_239_3_15 | SE_239 | SE_239_3 | GTGTTTCCTCAGTTGCCAGT | 40.09974314246706 | 60.962941562667666 | 89.8649444997435 | 61.16149900962082 | 81.66577748016432 | 9.529411208552496 | 40.8941032850566 | 61.44797688615203 | 72.81479278166393 | 40.0953819654791 | 34.03976397623994 | 26.695663292742836 |
| SE_239_3_2 | SE_239 | SE_239_3 | ATACTGCTGAAGAGTGGTGG | 596.0772629285644 | 655.601469919836 | 715.9569754100444 | 584.2875118153138 | 663.9321649686086 | 720.7700114105161 | 516.5570941270308 | 585.1738106542786 | 475.960108914291 | 607.9917919856284 | 755.9175172654662 | 1265.142303873465 |
| SE_239_3_22 | SE_239 | SE_239_3 | TTGTTAATATCACTTGCACA | 114.88034521895969 | 139.91494784874547 | 108.62795488979984 | 139.24000838360485 | 134.69550311663465 | 82.29946043749882 | 123.75847046793446 | 352.61685197745703 | 105.67024806119521 | 261.71349319285446 | 370.91604884454557 | 142.76376456553777 |
| SE_24_1_14 | SE_24 | SE_24_1 | CACACACCAGTACTCCCCTA | 6302.162334417458 | 5849.4442125621945 | 6005.1508516806625 | 6081.014571743789 | 6329.628051969099 | 6592.61993609859 | 6111.516119890433 | 5975.579413805645 | 5590.755309187269 | 6771.745510497006 | 5321.94102994041 | 6917.658835858579 |
| SE_24_1_15 | SE_24 | SE_24_1 | GGAGCCCTGCAGGGAGGAGG | 117.04789890233629 | 21.986634661945715 | 45.42623568118903 | 96.29682822791364 | 14.848323178211695 | 253.82886219144376 | 171.10953742957895 | 69.95615830115769 | 134.08577695160065 | 11.664111117230282 | 38.73490383503166 | 13.928172152735392 |
| SE_24_1_17 | SE_24 | SE_24_1 | GCATGTGTGATGTCGCCCTA | 6065.898982929409 | 5799.4745883304995 | 5632.853224467439 | 6116.149900962082 | 5835.3910090371955 | 6265.154714568332 | 5899.512479175797 | 6454.873633517631 | 5978.804875596868 | 5965.463829518462 | 6640.101545296185 | 5086.104197773874 |
| SE_24_1_28 | SE_24 | SE_24_1 | TGTAGGTATGTGGATCTTTG | 721.7953765644071 | 698.5753467590935 | 781.1337483439244 | 772.9772428024419 | 765.7492381906317 | 921.7539568999869 | 794.2065322203098 | 839.4738996138923 | 860.4577342125896 | 647.3581670062807 | 860.384379123582 | 869.3500785332342 |
| SE_24_1_7 | SE_24 | SE_24_1 | CGAGGTTCAGTGCTCTCTAG | 4365.453118320468 | 4518.253423029844 | 4314.504862850323 | 4329.453344787415 | 4503.2843010490615 | 4355.8072324183595 | 4160.436928948127 | 4236.128991182265 | 3987.0538974350125 | 3559.01190464489 | 4242.058862418316 | 4271.306126838854 |
| SE_24_10_0 | SE_24 | SE_24_10 | GGACTACTTTGGACAGAAGT | 405.3325387914238 | 487.70353250134133 | 533.2645058226537 | 400.80301478645134 | 512.2671496483034 | 538.8448883381502 | 419.7026389782125 | 561.5399733903739 | 419.1290511334801 | 406.0568682685792 | 626.8011711486942 | 409.72039749296613 |
| SE_24_10_15 | SE_24 | SE_24_10 | TACATATTTCAGCACTAAGG | 263.3577725302566 | 266.83779339725027 | 309.0959080046123 | 287.5891761941745 | 515.4489331864917 | 195.78608483026036 | 188.3281072338133 | 329.92836820410855 | 233.54012806801964 | 245.67534040666283 | 288.75110131569056 | 276.24208102925195 |
| SE_24_10_4 | SE_24 | SE_24_10 | GGTAGCTTTAGTTTGTACGT | 107.2939073271416 | 171.89550735703014 | 173.80472782367974 | 176.97795458103047 | 173.9375000876227 | 203.58287581907607 | 121.60614924240517 | 178.67180971511897 | 148.29354139680336 | 174.96166675845424 | 92.72901221113639 | 128.83559241280238 |
| SE_24_10_5 | SE_24 | SE_24_10 | CTGGACAGCAGATGAATGAG | 896.2834480762232 | 973.4082800334148 | 970.7389059697567 | 943.4486549356403 | 1044.6855950384656 | 894.0320333841978 | 941.6405361690665 | 1044.6156070645845 | 965.2399969959596 | 909.8006671439621 | 1223.083933215242 | 839.1723722023074 |
| SE_24_10_7 | SE_24 | SE_24_10 | TCTGCTGTCCAGGAAAGCTT | 1130.3792458808957 | 1147.3025723597127 | 1180.0946008482802 | 1016.3219303513588 | 1276.9557933262058 | 956.4063612947232 | 1015.8956184498272 | 1024.7631837629046 | 1242.2914036774125 | 1226.1896811988333 | 1113.9219314983347 | 797.3878557441012 |
| SE_24_2_18 | SE_24 | SE_24_2 | CACCCTCCAGTCGGATCAGA | 363.0652419655801 | 432.73694584647706 | 389.08558387801037 | 429.43180155691215 | 289.54230197512805 | 460.01066834012505 | 419.7026389782125 | 397.9938195241539 | 393.37747807655023 | 655.3772433993765 | 338.0500698330036 | 283.20616710561967 |
| SE_24_2_22 | SE_24 | SE_24_2 | TCCTCACCTCCCTATCTCAT | 579.82061030324 | 544.6689041254734 | 560.915257976421 | 537.4404061909233 | 582.2663874884443 | 967.6683927230125 | 700.5805589097855 | 484.0209871647667 | 602.0540183654651 | 731.1939656613733 | 609.1943966782252 | 569.8943772494232 |
| SE_24_2_29 | SE_24 | SE_24_2 | CCCAGGAATGGGACACCACA | 314.2952840896067 | 311.8104552057756 | 316.99612290568865 | 277.1787082776433 | 368.02629591710416 | 300.60960812433785 | 299.172650348572 | 207.9777679223607 | 221.99631945629244 | 462.91940996507685 | 186.63180938697073 | 217.04734938012655 |
| SE_24_2_35 | SE_24 | SE_24_2 | ATATCCCTGCGGACACACCT | 184.24206308701082 | 228.86087898116222 | 223.18107095540694 | 180.88188004972966 | 187.725228753105 | 306.6737788934167 | 263.65935012733866 | 148.42049801732105 | 258.40371584712443 | 220.1600973377216 | 220.67157336321065 | 233.29688355831783 |
| SE_24_2_4 | SE_24 | SE_24_2 | GGATAAACAGAAGGGCTCTG | 112.71279153558308 | 202.87667438068092 | 209.35569487852334 | 161.36225270623365 | 229.08841474955184 | 161.99999054539242 | 133.4439159828163 | 270.37109829906893 | 254.85177473582374 | 211.41201399799886 | 196.02208910455414 | 264.6352709019725 |
| SE_24_3_30 | SE_24 | SE_24_3 | ACCTGTTCACACCTGTACGG | 459.52138087583876 | 530.6774093405988 | 510.5513879820593 | 481.4841411395682 | 587.5693600520914 | 553.5721602059132 | 469.2060271653863 | 422.5730102786147 | 445.76860946823524 | 342.63326406863956 | 336.8762848683056 | 745.1572101713435 |
| SE_24_3_34 | SE_24 | SE_24_3 | CCCTGCAAAGAATCAGCATG | 565.7315113612921 | 540.6713341869379 | 654.7303099267027 | 511.4142363995954 | 732.8708082960201 | 604.6844566881493 | 545.6134306716763 | 698.6162295210207 | 655.3331350349753 | 670.6863892407413 | 691.3593442070802 | 704.5333747258653 |
| SE_24_3_37 | SE_24 | SE_24_3 | TGTGGTGCCAGCCTAGGCAG | 3181.968807196846 | 3085.1246000648375 | 3077.1337039692394 | 2887.6035383478425 | 2910.271342929492 | 2676.898239493383 | 3027.2398037069534 | 2696.1481550662397 | 2875.2963295979 | 3365.0960573209363 | 2718.4859782404037 | 2555.8195900269448 |
| SE_24_3_38 | SE_24 | SE_24_3 | GCTGATTCTTTGCAGGGCAA | 348.97614302363223 | 344.7904071986942 | 383.1604227022031 | 411.21348270298256 | 323.48132638246904 | 614.2138678967018 | 394.95094488462564 | 311.96665188354103 | 414.6891247443543 | 267.5455487514696 | 436.6480068676296 | 523.4671367403052 |
| SE_24_3_8 | SE_24 | SE_24_3 | AGAAACTCAGAACAGGCAGT | 290.4521935724641 | 359.78129446820265 | 363.4098854495122 | 291.4931016628737 | 424.2378050917627 | 171.52940175394494 | 238.90765603375175 | 305.34917744964775 | 369.40187557527065 | 277.0226390342192 | 275.8394667040133 | 389.988820276591 |
| SE_24_4_19 | SE_24 | SE_24_4 | ACAGGCCTCTGAGTCAGGAT | 2182.726559160234 | 2346.5735539203883 | 2317.725546603275 | 2403.5167802291417 | 2361.9439798483886 | 2312.181683238783 | 2559.109937154332 | 2215.9085818636977 | 2491.6866895774265 | 3104.111571072909 | 2568.2415027590687 | 1966.1936355611463 |
| SE_24_4_20 | SE_24 | SE_24_4 | TCTGCGGGAGAGAGGCCAGG | 2125.2863865507543 | 2157.6883743245817 | 2156.758667993844 | 2115.9276040349673 | 2331.186738979236 | 1895.4865203920783 | 2031.7912368996545 | 2620.519875821745 | 1832.8016134311504 | 1826.162396791366 | 3033.0603487794488 | 2256.363888743134 |
| SE_24_4_24 | SE_24 | SE_24_4 | CCACAGCCCTGAGCCATCCG | 298.0386314642822 | 349.78736962186366 | 347.6094556473595 | 300.6022610898385 | 461.35861303729195 | 194.91977472039196 | 258.2785470635154 | 223.10342377125966 | 311.6828325166346 | 288.6867501514495 | 456.6023512674944 | 213.5653063419427 |
| SE_24_4_32 | SE_24 | SE_24_4 | TCTGGCCTCTCTCAACACCT | 1921.536340313354 | 1908.8396456507417 | 1867.4132972419227 | 1730.7402911233125 | 2044.8262205422961 | 2467.251192905228 | 1825.1683992488422 | 2152.569897996433 | 1973.1032873275274 | 1654.8457647570463 | 2010.6936445275524 | 1703.8797266846298 |
| SE_24_4_6 | SE_24 | SE_24_4 | AGTTGGAGGCCAATACCAAG | 351.14369670700887 | 421.7436285155042 | 437.474400147103 | 472.37498171260336 | 446.5102898590802 | 250.36362175197013 | 488.57691819515 | 399.88452650526625 | 216.66840778934142 | 575.1864794684183 | 400.2606729619938 | 159.01329874372908 |
| SE_24_5_14 | SE_24 | SE_24_5 | GCCCGCCCTCAGAAAACCCA | 152.81253467805016 | 182.888824688003 | 177.75483527421792 | 200.40150739322567 | 200.45236290585788 | 122.14972549144564 | 173.26185865510826 | 329.92836820410855 | 159.83735000853056 | 152.36245146882055 | 383.8276834562228 | 204.2798582401191 |
| SE_24_5_23 | SE_24 | SE_24_5 | AGGTGATGTCTACGGCTCAG | 3480.0074386611277 | 3437.910147140603 | 3404.0050955012734 | 3575.9957293284683 | 3278.2976388465963 | 3171.5613122282443 | 3878.4828484037894 | 3023.24046279868 | 3313.9610568435337 | 4262.503606402842 | 3453.275366141307 | 3520.345511603871 |
| SE_24_5_3 | SE_24 | SE_24_5 | GGTCACTCGAGGTAGCAGCG | 235.17957464636086 | 272.83414830505365 | 298.2331125156323 | 213.41459228888968 | 304.3906251533397 | 234.77003977433876 | 221.6890862295174 | 340.3272566002266 | 331.21850862878836 | 186.62577787568452 | 296.96759606857603 | 160.17397975645702 |
| SE_24_5_34 | SE_24 | SE_24_5 | GGAACCCAAACAGGAAGCTG | 52.02128840103835 | 86.94714616314897 | 79.00214901076352 | 70.27065843658562 | 103.93826224748186 | 30.320853845394307 | 55.960351863761666 | 22.68848377334844 | 134.08577695160065 | 94.77090282749604 | 52.820323411406804 | 31.338387343654635 |
| SE_24_5_9 | SE_24 | SE_24_5 | TCCCACATCAGCTTCTCCAG | 126.80189047753097 | 209.8724217731182 | 246.881715658636 | 208.20935833062407 | 171.8163110621639 | 198.3850151598656 | 91.47365208499504 | 194.7428190545741 | 294.81111223795637 | 150.90443757916677 | 159.63475519891836 | 313.38387343654637 |
| SE_24_6_0 | SE_24 | SE_24_6 | CCACATGATTTCTCTGGGGC | 108.37768416882989 | 132.9192004563082 | 164.91698605996885 | 156.15701874796804 | 114.54420737477592 | 128.21389626052448 | 147.4340039487567 | 154.09261896065814 | 175.8210850093836 | 363.04545852379255 | 56.341678305500594 | 378.3820101493115 |
| SE_24_6_12 | SE_24 | SE_24_6 | GGTGCAAATGATATCTGTGA | 792.2408712741465 | 824.4987998229643 | 862.110951079957 | 753.4576154589458 | 854.8391772599018 | 757.1550360249893 | 829.7198324415432 | 912.2661183867185 | 679.3087375362549 | 755.2511948406608 | 821.6494752885503 | 482.843301294827 |
| SE_24_6_40 | SE_24 | SE_24_6 | CTGGCTCCCTGGGCCATGAG | 164.73407993662144 | 161.90158251069118 | 192.56773821373608 | 176.97795458103047 | 153.786204345764 | 246.8983813124965 | 145.28168272322742 | 107.7702979234051 | 110.11017445032105 | 72.17168753786237 | 86.86008738764674 | 29.017025318198737 |
| SE_24_6_49 | SE_24 | SE_24_6 | CAGAGTCCAGGGTAAGGATC | 1133.6305764059607 | 1133.3110775748382 | 1153.4313755571475 | 1123.0292264958036 | 1018.1707322202304 | 738.9625237177527 | 1154.7203374964668 | 1157.1126724407704 | 984.7756731081133 | 648.8161808959345 | 1004.7599297814272 | 1163.0023747534053 |
| SE_24_6_50 | SE_24 | SE_24_6 | AGCATTCCTGATCCTTACCC | 402.0812082663589 | 418.7454510616025 | 409.8236479933358 | 469.77236473347057 | 408.3288874008216 | 298.0106777947326 | 456.29209981221055 | 691.9987550871274 | 388.0495664095992 | 502.28578498572904 | 845.1251745825089 | 452.6655949639003 |
| SE_24_7_18 | SE_24 | SE_24_7 | GATAAGAGCACTGATAAACG | 386.9083324827227 | 464.7175053547617 | 417.72386289441215 | 292.7944101524401 | 384.99580812077465 | 450.48125713157253 | 349.75219914851044 | 414.064828863609 | 374.7297872422217 | 469.4804724685189 | 335.7024999036077 | 487.48602534573877 |
| SE_24_7_19 | SE_24 | SE_24_7 | CTAGACAGCTGTGGCAAGAA | 572.2341724114218 | 780.5255304990729 | 665.5931054156827 | 684.4882655119267 | 806.0518296743492 | 611.6149375670966 | 583.2790521184389 | 613.5344153709641 | 730.8118836501147 | 396.5797779858296 | 639.7128057603713 | 950.5977494241906 |
| SE_24_7_3 | SE_24 | SE_24_7 | CAGATGGAGACATTACGCCG | 2498.105620091529 | 2573.435647932283 | 2454.004253646842 | 2436.0494924683017 | 2518.911967732341 | 2228.1496025815472 | 2485.9310154863356 | 2339.7498891265577 | 2497.0146012443774 | 2160.047577522083 | 2497.814404877193 | 2582.5152533196874 |
| SE_24_7_40 | SE_24 | SE_24_7 | ATACCCACAGCACAGACCCA | 865.9376965089508 | 766.5340357141984 | 868.0361122557642 | 769.0733173337427 | 716.9618906050789 | 505.9251041631507 | 780.2164442543694 | 1075.8122722529386 | 985.6636583859384 | 851.4801115578106 | 762.9602270536539 | 543.1987139566603 |
| SE_24_7_45 | SE_24 | SE_24_7 | TGGCTGGCTGGGGAGGAACA | 36.84841261740216 | 93.94289355558624 | 99.74021312608895 | 52.05233958265602 | 103.93826224748186 | 85.76470087697247 | 31.208657770174778 | 24.57919075446081 | 48.83919028038434 | 23.328222234460565 | 16.432989505771005 | 22.05293924183104 |
| SE_24_8_2 | SE_24 | SE_24_8 | TTTCTGGATCCCGCAGTGGC | 5.4188842084414945 | 7.9951398770711695 | 10.862795488979984 | 10.410467916531204 | 0.0 | 19.9251325269734 | 2.152321225529295 | 0.0 | 3.551941111300679 | 8.748083337922711 | 0.0 | 1.1606810127279494 |
| SE_24_8_46 | SE_24 | SE_24_8 | TGCAAGGTAGGCCAGGGAGC | 4196.383931017093 | 3784.699339308565 | 3860.2425060384326 | 3497.9172199544846 | 3630.4150170727594 | 3959.9035122084965 | 3779.476072029442 | 3434.46923119062 | 3498.661994631169 | 2972.890321004068 | 3810.1059954094776 | 3977.6538306186826 |
| SE_24_8_48 | SE_24 | SE_24_8 | TTCTTGTGCAAGGTAGGCCA | 8.670214733506391 | 12.992102300240651 | 13.825376076883616 | 9.109159426964803 | 0.0 | 67.57218856973589 | 0.0 | 16.071009339455145 | 15.095749723027886 | 0.0 | 1.173784964697929 | 0.0 |
| SE_24_8_5 | SE_24 | SE_24_8 | GCCTGCCGCTCCGGCCACTG | 855.0999280920678 | 929.4350107095235 | 908.5247136237805 | 818.5230399372658 | 974.6863571983248 | 803.0694718480149 | 944.8690180073605 | 1219.5060028174787 | 920.840733104701 | 934.5869032680764 | 1034.1045538988756 | 601.2327645930778 |
| SE_24_8_9 | SE_24 | SE_24_8 | GGAGCGGCAGGCCGGGAGGT | 2987.97275253464 | 3108.110627211417 | 3029.7324145627813 | 3115.332524021963 | 3187.086510751867 | 2732.342086524961 | 3387.75360898311 | 3147.08177006154 | 2623.9964959733766 | 3139.8329113694267 | 3325.332804989233 | 3020.0919951181245 |
| SE_24_9_14 | SE_24 | SE_24_9 | TCCTTGCCAGAGGTACACCA | 954.8073975273913 | 892.4574887780693 | 924.3251434259332 | 812.0164974894338 | 823.0213418780196 | 942.5453995368287 | 783.4449260926633 | 1120.2438863090792 | 844.4739992117364 | 632.7780281097429 | 852.1678843706965 | 1021.3992912005955 |
| SE_24_9_27 | SE_24 | SE_24_9 | CCAGGTCACTAGATAGGAAG | 723.9629302477837 | 663.596609796907 | 740.645146975908 | 752.1563069693794 | 631.053735073997 | 596.0213555894652 | 751.1601077097239 | 564.3760338620425 | 803.6266764317786 | 584.6635697511679 | 657.3195802308403 | 913.4559570168961 |
| SE_24_9_28 | SE_24 | SE_24_9 | TTCTGTCTCCAAAGAGAGGA | 6.5026610501297935 | 0.0 | 0.0 | 3.9039254686992013 | 0.0 | 1.7326202197368175 | 0.0 | 0.0 | 0.0 | 8.748083337922711 | 0.0 | 0.0 |
| SE_24_9_31 | SE_24 | SE_24_9 | TTTCGGAGCATAGCGTCCAG | 200.4987157123353 | 184.8876096572708 | 207.38064115325426 | 184.78580551842887 | 198.33117388039906 | 168.93047142433971 | 190.4804284593426 | 275.09786575184984 | 370.2898608530958 | 145.07238202055163 | 287.5773163509926 | 272.76003799106815 |
| SE_24_9_37 | SE_24 | SE_24_9 | GCTGGTGCTATGGCCCGGGA | 995.9909175115467 | 1175.285561929462 | 1113.9303010517656 | 1181.5881085262915 | 1094.5335371367478 | 1035.2405812927484 | 1305.3828232835174 | 1091.8832815923936 | 1251.1712564556642 | 2417.387029045976 | 1108.053006674845 | 1232.6432355170823 |
| SE_240_1_10 | SE_240 | SE_240_1 | CTTTGCAGATAGAGAAAGGG | 45.51862735090855 | 61.96233404730156 | 75.05204156022535 | 58.55888203048802 | 64.69626527649382 | 121.28341538157723 | 35.513300221233365 | 106.82494443284891 | 55.94307250298569 | 56.13353475167074 | 142.02798072844942 | 88.21175696732416 |
| SE_240_1_12 | SE_240 | SE_240_1 | GATTAGTTCAGCAAAGAGAC | 57.440172609479845 | 69.95747392437274 | 80.97720273603261 | 79.37981786355043 | 81.66577748016432 | 8.663101098684088 | 69.95043982970209 | 13.23494886778659 | 43.51127861343332 | 58.320555586151414 | 18.780559435166865 | 49.909283547301825 |
| SE_240_1_2 | SE_240 | SE_240_1 | ATACTTATATGAACTTATAA | 0.0 | 2.9981774539016888 | 3.950107450538176 | 2.602616979132801 | 13.787728665482287 | 0.8663101098684087 | 0.0 | 7.56282792444948 | 4.439926389125849 | 0.0 | 0.0 | 0.0 |
| SE_240_1_21 | SE_240 | SE_240_1 | GCTTTCACATGAGTCTCACT | 365.2327956489567 | 363.7788644067382 | 375.26020780112674 | 386.48862140122094 | 435.9043447317862 | 176.7272624131554 | 292.7156866719841 | 490.63846159866 | 245.08393667974687 | 355.7553890755236 | 390.87039324441037 | 233.29688355831783 |
| SE_240_1_4 | SE_240 | SE_240_1 | TCAGTCAGCAAACATGAGAG | 2373.4712832973746 | 2569.438077993747 | 2605.095863629927 | 2180.9930285132873 | 2325.883766415589 | 2260.2030766466783 | 2189.9868469760577 | 2126.1000002608603 | 2301.6578401228403 | 2882.4934598455334 | 2029.4742039627192 | 1956.9081874593228 |
| SE_240_2_10 | SE_240 | SE_240_2 | GCAGCCCTCAAGTGTCTGTG | 1581.2304120232282 | 1430.1306455111055 | 1480.3027670891815 | 1510.819156386591 | 1499.6806409993812 | 1246.62024810064 | 1535.681194415152 | 1806.5705204528695 | 1225.4196833987344 | 1880.837917653383 | 1530.6155939660994 | 1393.9778962862672 |
| SE_240_2_6 | SE_240 | SE_240_2 | CCCTCAAGTGTCTGTGTGGG | 236.26335148804915 | 224.86330904262664 | 228.11870526857967 | 235.53683661151848 | 286.3605184369398 | 259.89303296052265 | 245.3646197103396 | 530.3433082020198 | 253.96378945799856 | 225.26314595150984 | 517.6391694317867 | 451.50491395117234 |
| SE_240_2_7 | SE_240 | SE_240_2 | GCCCTCAAGTGTCTGTGTGG | 122.46678311077778 | 178.8912547494674 | 221.20601723013786 | 119.72038104010883 | 153.786204345764 | 200.9839454894708 | 107.61606127646475 | 174.89039575289422 | 158.94936473070538 | 73.62970142751615 | 57.515463270198524 | 153.20989368008932 |
| SE_240_2_8 | SE_240 | SE_240_2 | AGCCCTCAAGTGTCTGTGTG | 145.22609678623206 | 143.91251778728105 | 171.82967409841066 | 196.49758192452646 | 226.96722572409305 | 310.1390193328903 | 157.11944946363852 | 120.0598933006355 | 127.86988000682445 | 146.5303959102054 | 171.37260484589763 | 138.12104051462597 |
| SE_240_2_9 | SE_240 | SE_240_2 | CAGCCCTCAAGTGTCTGTGT | 671.9416418467454 | 627.6184803500868 | 600.4163324818028 | 707.9118183241219 | 760.4462656269847 | 428.8235043848623 | 661.8387768502582 | 608.8076479181832 | 665.9889583688773 | 553.3162711236115 | 592.7614071724541 | 551.323481045776 |
| SE_241_1_2 | SE_241 | SE_241_1 | GGGTCCCGCGGCCGGGCCCC | 0.0 | 0.9993924846338962 | 0.0 | 0.0 | 0.0 | 0.0 | 0.0 | 0.0 | 0.0 | 0.0 | 0.0 | 0.0 |
| SE_241_1_4 | SE_241 | SE_241_1 | GGGAGCCTGGGGCCCGGCCG | 52.02128840103835 | 88.94593113241676 | 95.79010567555078 | 40.34056317655841 | 66.81745430195262 | 26.85561340592067 | 35.513300221233365 | 72.79221877282625 | 125.20592417334893 | 52.48850002753627 | 144.37555065784528 | 117.2287822855229 |
| SE_241_1_5 | SE_241 | SE_241_1 | GCCGGGCCCCAGGCTCCCGC | 37.93218945909046 | 99.93924846338962 | 77.02709528549444 | 40.34056317655841 | 43.48437502190568 | 149.0053388973663 | 26.904015319116187 | 20.79777679223607 | 54.167101947335354 | 16.038152786191638 | 2.347569929395858 | 85.89039494186825 |
| SE_241_1_6 | SE_241 | SE_241_1 | CCCGGCGGGAGCCTGGGGCC | 0.0 | 0.0 | 2.962580587903632 | 0.0 | 0.0 | 0.0 | 0.0 | 0.0 | 0.0 | 0.0 | 0.0 | 0.0 |
| SE_241_2_10 | SE_241 | SE_241_2 | TTAAAATTAGGAGGACACTG | 31.429528408960667 | 12.992102300240651 | 20.738064115325425 | 15.615701874796805 | 11.666539640023474 | 5.197860659210452 | 36.58946083399801 | 11.34424188667422 | 21.311646667804077 | 1.4580138896537853 | 5.868924823489645 | 1.1606810127279494 |
| SE_241_2_13 | SE_241 | SE_241_2 | CCCAGCCTCTTAAAATTAGG | 44.434850509220254 | 35.97812944682026 | 49.376343131727204 | 42.943180155691216 | 21.211890254588134 | 14.727271867762948 | 48.42722757440914 | 14.180302358342775 | 26.639558334755094 | 18.225173620672315 | 2.347569929395858 | 2.321362025455899 |
| SE_241_2_14 | SE_241 | SE_241_2 | GCAGCTCTTTGCATCCGCCA | 1231.1704921579076 | 1265.2308855465126 | 1308.4730929907707 | 1123.0292264958036 | 1091.3517535985595 | 937.3475388776183 | 1149.3395344326434 | 1052.1784349890338 | 1267.1549914565173 | 603.6177503166671 | 734.7893879009035 | 853.1005443550429 |
| SE_241_2_16 | SE_241 | SE_241_2 | GCTGGGTCAGATCATTTCTA | 744.5546902398613 | 839.4896870924728 | 806.8094467724225 | 713.1170522823875 | 747.7191314742317 | 502.4598637236771 | 629.5539584673188 | 782.7526901805212 | 808.0666028209044 | 599.9727155925326 | 811.085410606269 | 341.24021774201714 |
| SE_241_2_5 | SE_241 | SE_241_2 | ATGTATGTAACAATCCCTAA | 197.2473851872704 | 248.84872867384016 | 264.6571991860578 | 160.06094421666725 | 250.30030500414 | 168.93047142433971 | 187.25194662104866 | 216.48594933736638 | 308.1308914053339 | 444.69423634440454 | 196.02208910455414 | 164.8167038073688 |
| SE_242_1_13 | SE_242 | SE_242_1 | TTATTTTTATTGCCACCCAG | 2.167553683376598 | 0.0 | 3.950107450538176 | 1.3013084895664004 | 11.666539640023474 | 4.331550549342044 | 2.152321225529295 | 8.508181415005666 | 8.879852778251697 | 0.0 | 8.216494752885502 | 2.321362025455899 |

| | | | | | | | | | | | | | | | |
|---|---|---|---|---|---|---|---|---|---|---|---|---|---|---|---|
| SE_242_1_16 | SE_242 | SE_242_1 | AGTTGGTCTGACAAACCAGT | 218.92292202103638 | 202.87667438068092 | 191.58021135110155 | 226.4276771845537 | 234.3913873131989 | 147.27271867762948 | 148.51016456152135 | 364.90644735468743 | 98.56636583859384 | 261.71349319285446 | 192.50073421046037 | 189.19100507465575 |
| SE_242_1_21 | SE_242 | SE_242_1 | GCCAAGATGGAACCAAGAGA | 1108.7037090471297 | 1235.2491110074957 | 1191.9449231998947 | 1123.0292264958036 | 1440.2873482865343 | 1337.5828096368232 | 1104.1407886965283 | 1249.7573145152767 | 1105.5416708923365 | 1395.3192923986726 | 1441.4079366490569 | 1067.8265317097134 |
| SE_242_1_33 | SE_242 | SE_242_1 | TTCAAAAGTACAGAGTTGCA | 44.434850509220254 | 37.97691441608806 | 35.55096705484358 | 41.641871666124814 | 28.63605184369398 | 56.31015714144657 | 36.58946083399801 | 25.524544245016994 | 72.81479278166393 | 61.965590310285876 | 82.16494752885504 | 13.928172152735392 |
| SE_242_1_6 | SE_242 | SE_242_1 | AAGGTTGTCTTCCCAAACCT | 371.7354566990865 | 417.7460585769686 | 405.87354054279757 | 389.09123838035373 | 434.84375021905674 | 369.91441691381056 | 427.23576326756506 | 578.5563362203852 | 315.2347736273563 | 360.129430744485 | 1021.1929192871983 | 198.47645317647937 |
| SE_242_2_13 | SE_242 | SE_242_2 | ATCACTTTCTAGGTGACGGA | 29.26197472558407 | 30.98116702365078 | 31.60085960430541 | 20.820935833062407 | 57.27210368738796 | 1.7326202197368175 | 8.60928490211718 | 48.213028018365435 | 21.311646667804077 | 1.4580138896537853 | 4.695139858791716 | 8.124767089095647 |
| SE_242_2_23 | SE_242 | SE_242_2 | GTATAGAAAAGGGATATACA | 45.51862735090855 | 84.94836119388117 | 55.301504307534465 | 49.44972260352321 | 50.90853661101153 | 21.65775274671022 | 46.27490634887984 | 13.23494886778659 | 111.88614500597139 | 45.198430579267345 | 48.12518355261509 | 117.2287822855229 |
| SE_242_2_25 | SE_242 | SE_242_2 | TGTGCTTAGTGTATAGAAAA | 329.4681598732429 | 389.7630690072195 | 441.4245075976164 | 490.59330056653295 | 454.9950459609155 | 291.08019691578534 | 359.43764466339223 | 501.98270348533424 | 493.7198144707944 | 508.1178405443462 | 543.4624386551411 | 409.72039749296613 |
| SE_242_2_3 | SE_242 | SE_242_2 | GCTCTGCATGCCAAGTACTG | 381.48944827428124 | 364.7782568913721 | 331.8090258452068 | 372.17422801599054 | 351.0567837134336 | 307.5400890032851 | 371.2754114038034 | 279.82463320463074 | 391.6015075208999 | 296.7058265445453 | 473.0353407732654 | 320.34795951291403 |
| SE_242_2_6 | SE_242 | SE_242_2 | GGTGCAATCTTGGTTCACTG | 1376.3965889441397 | 1350.1792467403936 | 1305.510512402867 | 1469.177284720466 | 1324.682546399029 | 1233.625596452614 | 1534.6050338023872 | 1438.8280126265136 | 1358.6174750725097 | 1691.2961119983909 | 1631.5611009301213 | 1145.5921595624861 |
| SE_242_3_17 | SE_242 | SE_242_3 | CTTAGGAATACAGTAACCCA | 358.73013459882696 | 372.77339676844326 | 361.4348317242431 | 448.95142890040813 | 282.1181403860222 | 271.1550643888119 | 309.9342564762185 | 300.62240999686685 | 385.38561057612367 | 697.6596461993363 | 349.78791947998286 | 355.1683898947525 |
| SE_242_3_2 | SE_242 | SE_242_3 | TTAATTTCCCTGCTGTAAGC | 5.4188842084414945 | 10.993317330972857 | 7.900214901076352 | 2.602616979132801 | 3.1817835381882205 | 0.8663101098684087 | 20.4470516425283 | 0.945353490556185 | 27.527543612580263 | 40.82438891030599 | 1.173784964697929 | 1.1606810127279494 |
| SE_242_3_27 | SE_242 | SE_242_3 | GAGGGGAGATTGAGGCACGT | 12045.095818523754 | 11232.17213480036 | 11270.644083248051 | 12177.644845362376 | 11645.327749768887 | 10873.924499068266 | 12328.495979831801 | 11276.176435354175 | 11360.883644495223 | 11390.733512920198 | 11643.946849803457 | 11214.499944977448 |
| SE_242_3_4 | SE_242 | SE_242_3 | AATCAGAGCCAGACTGGGCG | 222.1742525461013 | 229.86027146579613 | 237.00644703229057 | 268.0695488506785 | 260.90625013143404 | 268.5561340592067 | 235.6791741954578 | 259.0268564123947 | 211.34049612239042 | 252.23640291010486 | 197.19587406925208 | 218.20803039285448 |
| SE_242_3_6 | SE_242 | SE_242_3 | TAGTGTTTCTGACAGGAGGA | 340.30592829012585 | 351.78615459113144 | 399.9483793669903 | 325.3271223916001 | 345.7538111497866 | 460.87697844999343 | 399.2555873356842 | 483.07563367421056 | 250.41184834669787 | 212.14102094462575 | 271.1443268452216 | 480.52193926937105 |
| SE_242_4_10 | SE_242 | SE_242_4 | CAGAAAGTAACATCGTGTGG | 2397.3143738145172 | 2202.6610361331072 | 2330.563395817524 | 2563.577724445809 | 2300.4294981100834 | 2838.898230038775 | 2442.8845909757497 | 2313.279991390985 | 2607.1247756946987 | 2410.825966542534 | 2619.8880412057774 | 2943.48704827808 |
| SE_242_4_14 | SE_242 | SE_242_4 | AGTACTGTAGGAGGCCAAGG | 456.27005035077383 | 614.6263780498462 | 579.6782683664774 | 556.9600335344194 | 505.903582571927 | 476.4705604276248 | 454.13977858668125 | 432.97189867473276 | 516.8074316942489 | 790.9725351371785 | 453.0809963734006 | 669.7129443440268 |
| SE_242_4_20 | SE_242 | SE_242_4 | TGAGGCCTCCAAGTGCAGGG | 2484.016521149581 | 2575.4344329015503 | 2670.2726365638073 | 2471.1848216865947 | 2469.064025634059 | 2715.8821944374613 | 2545.119849188391 | 2744.3611830846053 | 2647.084113196831 | 3344.6838628657833 | 2751.351957251946 | 2706.708121681578 |
| SE_242_4_8 | SE_242 | SE_242_4 | GTGGCATTCACAATACGAAA | 88.86970101844051 | 96.94107100948793 | 89.8649444997435 | 68.96934994701923 | 111.36242383658771 | 28.588233625657487 | 82.86436718287786 | 62.393330376708214 | 61.27098416993672 | 12.393118062057175 | 92.72901221113639 | 112.5860582346111 |
| SE_242_5_11 | SE_242 | SE_242_5 | AATGGTAGATTCTTGAGCTA | 88.86970101844051 | 153.90644263362 | 131.34107273039436 | 97.59813671748003 | 134.69550311663465 | 353.4545248263108 | 125.91079169346375 | 111.55171188562983 | 186.47690834328566 | 212.14102094462575 | 38.73490383503166 | 117.2287822855229 |
| SE_242_5_2 | SE_242 | SE_242_5 | CCCACACTCAAAATTTAGCA | 163.65030309493312 | 245.85055121993847 | 228.11870526857967 | 212.11328379932328 | 224.84603669863424 | 297.1443676848642 | 195.86123152316583 | 212.70453537514163 | 340.9863466848652 | 230.3661945652981 | 280.53460656280504 | 406.2383544547823 |
| SE_242_5_4 | SE_242 | SE_242_5 | ATGAGGGCGGAGCAAAAGTG | 5288.830987438899 | 6212.223684484298 | 6412.999445948729 | 6101.835507576851 | 6059.176451223101 | 5697.721592604525 | 5399.097794240236 | 5517.0829708858955 | 5239.113139168502 | 5678.964100201494 | 5023.7996489071365 | 4959.589967386528 |
| SE_242_5_7 | SE_242 | SE_242_5 | ATGTTCTCCCAGTAAAAGCA | 14.089098941947885 | 3.9975699385355847 | 1.975053725269088 | 11.711776406097604 | 4.242378050917627 | 0.0 | 5.380803063823238 | 4.726767452780925 | 15.983735000853056 | 0.0 | 2.347569929395858 | 1.1606810127279494 |
| SE_242_5_8 | SE_242 | SE_242_5 | TCTCCCAGTAAAAGCAGGGA | 31.429528408960667 | 41.97448435462364 | 44.43870881855448 | 31.23140374959361 | 7.4241615891058474 | 0.8663101098684087 | 10.761606127646475 | 0.0 | 53.27911666951019 | 18.954180565499207 | 1.173784964697929 | 1.1606810127279494 |
| SE_242_6_1 | SE_242 | SE_242_6 | CACTTGAGAAAAACACAGAA | 140.89098941947887 | 181.8894322033691 | 220.2184903675033 | 189.99103947669445 | 238.63376536411653 | 172.39571186381335 | 185.09962539551935 | 328.9830147135524 | 224.66027528976795 | 266.8165418066427 | 393.21796317380625 | 87.05107595459621 |
| SE_242_6_33 | SE_242 | SE_242_6 | CATTGCAACCTCAAACTCGA | 971.0640501527158 | 886.4611338702659 | 854.2107361788806 | 998.1036114974291 | 926.9596041255015 | 768.4170674532785 | 995.4485668072989 | 936.8453091411793 | 665.1009730910522 | 885.0144310198477 | 1213.6936534976587 | 1006.3104380351322 |
| SE_242_6_4 | SE_242 | SE_242_6 | CACGATGCTGAGCTAACCCG | 13719.531038932177 | 12299.52330838936 | 12218.669871377213 | 12971.44302399788 | 11922.142917591262 | 11569.5715172926 | 13106.560102860642 | 11174.078258374107 | 11989.577221195443 | 12074.542027167823 | 9802.278240192405 | 10006.231010727652 |
| SE_242_6_6 | SE_242 | SE_242_6 | GTAGTTCATCATCAGCCCTC | 511.54266927687706 | 474.7114302011007 | 527.3393446468465 | 541.3443316596226 | 526.0548783137857 | 541.4438186677554 | 487.5007575823853 | 362.07038688301884 | 441.3286830791094 | 457.0873544064617 | 468.3402009144737 | 237.93960760922963 |
| SE_242_6_7 | SE_242 | SE_242_6 | ATTGAGGGAAAAGAGACAAA | 212.42026097090658 | 287.8250355745621 | 292.307951339825 | 249.85122999674888 | 277.8757623351046 | 469.5400795486775 | 249.66926216139822 | 236.33837263904624 | 229.1002016788938 | 160.3815278619164 | 238.2783478336796 | 266.95663292742836 |
| SE_242_7_10 | SE_242 | SE_242_7 | AACACCAGAGAAAGCAGTGG | 634.0094523876548 | 709.5686640900662 | 721.8821365858516 | 648.0516278040674 | 639.5384911758323 | 803.0694718480149 | 703.8090407480795 | 481.1849266930982 | 638.461414756297 | 1152.5599797713173 | 454.25478133809855 | 694.0872456113137 |
| SE_242_7_13 | SE_242 | SE_242_7 | AATGTAAATGGAAATCAGAA | 87.78592417675222 | 153.90644263362 | 162.94193233469977 | 121.02168952967524 | 132.57431409117584 | 83.16577054736723 | 116.22534617858193 | 137.07625613064684 | 232.6521427901945 | 335.34319462037064 | 149.070690516637 | 44.10587848366208 |
| SE_242_7_17 | SE_242 | SE_242_7 | TTCACAACAGAGAGAATGCA | 29.26197472558407 | 39.975699385355846 | 44.43870881855448 | 13.013084895664004 | 66.81745430195262 | 1.7326202197368175 | 21.52321225529295 | 56.72120943331 | 19.535676112153734 | 21.87020834480678 | 110.33576868160533 | 1.1606810127279494 |
| SE_242_7_19 | SE_242 | SE_242_7 | CACAACAGAGAGAATGCAGG | 1001.4098017199882 | 1000.39187711853 | 1056.6537430189621 | 1054.0598765487844 | 1089.2305645731008 | 1053.433093599985 | 1082.6175764412353 | 1133.478835176866 | 966.1279822737847 | 1092.7814102955122 | 922.5949822525722 | 677.8377114331224 |
| SE_242_7_2 | SE_242 | SE_242_7 | TTTTACGCTGTGCTTCTCAA | 31.429528408960667 | 16.989672238776237 | 22.713117840594514 | 16.917010364363207 | 77.42339942924669 | 12.128341538157722 | 16.142409191469714 | 29.305958207241734 | 0.0 | 14.580138896537854 | 24.64948425865651 | 5.803405063639747 |
| SE_243_1_0 | SE_243 | SE_243_1 | TTATTCAGAAAACTCAGTAG | 7.586437891818092 | 10.993317330972857 | 16.78795666478725 | 10.410467916531204 | 9.545350614564661 | 28.588233625657487 | 6.456963676587884 | 0.0 | 22.199631945629246 | 18.954180565499207 | 0.0 | 33.659749369110536 |
| SE_243_1_12 | SE_243 | SE_243_1 | GCTGCTCTTTCAGAAGCCTA | 118.13167574402458 | 102.9374259172913 | 119.49075037877982 | 113.21383859227684 | 110.3018293238583 | 168.93047142433971 | 128.06311291899306 | 103.98888396118035 | 208.6765402889149 | 293.7897987652377 | 96.25036710523018 | 66.15881772549312 |
| SE_243_1_26 | SE_243 | SE_243_1 | GCTGTTAAGAAGGAAAAGAA | 148.47742731129696 | 236.8560188532334 | 239.9690276201942 | 180.88188004972966 | 224.84603669863424 | 277.2192351578908 | 160.34793130193248 | 210.81382839402926 | 231.7641575123693 | 142.15635424124406 | 173.7201747752935 | 215.88666836739858 |
| SE_243_1_3 | SE_243 | SE_243_1 | GTTCTTGTTTACATAACGCA | 7.586437891818092 | 15.990279754142339 | 15.800429802152705 | 6.506542447832002 | 11.666539640023474 | 1.7326202197368175 | 0.0 | 18.9070698111237 | 3.551941111300679 | 0.0 | 19.954344399864794 | 35.98111139456643 |
| SE_243_1_33 | SE_243 | SE_243_1 | GAAGAACCTAAACTTAGCAA | 533.2182061106431 | 643.6087601042292 | 714.9694485474099 | 633.737234418837 | 641.659680201291 | 767.5507573434102 | 637.0870827566713 | 612.5890618804079 | 623.3656650332692 | 477.4995488616147 | 640.8865907250693 | 930.8661722078155 |
| SE_243_2_14 | SE_243 | SE_243_2 | GCTGGGCTGCCAAGTCACAT | 6813.705003694336 | 5861.436922377801 | 5774.069565824179 | 6205.940186742164 | 5581.908920494868 | 6249.5611325907 | 6626.997053404699 | 5493.449133621991 | 6376.622280062545 | 5492.33832232581 | 6256.273861839962 | 6160.8948155599555 |
| SE_243_2_5 | SE_243 | SE_243_2 | GCCCGCACACACTATAACTT | 4245.153888893067 | 4117.497036691652 | 4116.999490323414 | 4295.619324058688 | 4266.771724710404 | 4386.994396373621 | 4099.095774020542 | 4600.090085046396 | 3909.7991782642225 | 4419.9691064854505 | 4611.801126298164 | 4457.015088875326 |
| SE_243_2_6 | SE_243 | SE_243_2 | AGTTCCACCAGCAGCCACCA | 1072.939073271416 | 1263.2321005772449 | 1107.0176130133239 | 1139.946236860167 | 1308.773628708088 | 1056.0320239295902 | 1060.0182035731777 | 1115.5171188562983 | 1411.0086064641948 | 1137.2508339299525 | 994.1958650991459 | 1173.4485038679568 |
| SE_243_2_8 | SE_243 | SE_243_2 | GGCAATATCTGCATGCTGCT | 1145.552121664532 | 1217.2600462840855 | 1190.9573963372602 | 974.6800586852339 | 1147.5632627732182 | 1143.5293450262996 | 964.2399090371241 | 1268.6643843264003 | 1333.7538872934051 | 976.8693060680362 | 1265.3401919443675 | 1605.2218406027541 |
| SE_243_2_9 | SE_243 | SE_243_2 | ATTCTTAGACGTGTCCCTGG | 14.089098941947885 | 39.975699385355846 | 36.53849391747813 | 16.917010364363207 | 26.51486281823517 | 5.197860659210452 | 8.60928490211718 | 69.95615830115769 | 49.72717555820951 | 32.80531251721017 | 22.30191432926065 | 58.034050636397474 |
| SE_243_3_10 | SE_243 | SE_243_3 | AGGTAATTAACAAACTTTAA | 119.21545258571288 | 169.89672238776234 | 214.29332919169605 | 124.92561499837444 | 161.21036593486983 | 275.486614938154 | 105.46374005093546 | 82.2457536783881 | 145.62958556332785 | 97.68693060680361 | 111.50957164630326 | 128.83559241280238 |
| SE_243_3_15 | SE_243 | SE_243_3 | AATGTGATACATACACACAT | 288.2846398890875 | 205.8748518345826 | 253.7944036970783 | 274.5760912985105 | 237.57317085138712 | 187.12298373157628 | 321.7720232166296 | 258.08150292183853 | 283.26730362622914 | 250.04938207562418 | 254.7113373394506 | 255.34982280014887 |
| SE_243_3_18 | SE_243 | SE_243_3 | GGATGCACAGGAAGCCACAA | 2151.2970307512733 | 2353.5693013128257 | 2227.860602103531 | 2179.691720023721 | 2494.5182939395645 | 2046.2244795091815 | 2092.056231214475 | 2518.421698841677 | 2324.7454573462946 | 2094.4369524876624 | 2259.536057043513 | 2171.6341748139935 |
| SE_243_3_2 | SE_243 | SE_243_3 | GACCCAGTAAGTCCATTACT | 150.64498099467355 | 212.87059922701988 | 206.3931142906197 | 226.4276771845537 | 218.48246962225778 | 238.2352802138124 | 188.3281072338133 | 60.50262339559584 | 340.9863466848652 | 549.671236399477 | 72.77466781126716 | 262.31390887651656 |
| SE_243_3_7 | SE_243 | SE_243_3 | AGTACAGTTGAACTCTGCCA | 613.4176923955772 | 639.6111901656935 | 690.2812769815463 | 676.6804145745282 | 588.6299545648208 | 758.0213461348576 | 676.9050254289633 | 544.5236105603626 | 832.0422053221 | 602.1597364270133 | 671.4049998072154 | 598.91140256762 |
| SE_244_1_10 | SE_244 | SE_244_1 | GTTCTGGGAGTGTTTCAACG | 47.686181034285156 | 45.972054293159225 | 49.376343131727204 | 57.25757354092162 | 19.090701229129323 | 16.459892087499767 | 30.13249715741013 | 30.25131169779792 | 39.95933750213264 | 47.38545141374802 | 2.347569929395858 | 51.069964560029774 |
| SE_244_1_14 | SE_244 | SE_244_1 | AGTGACTGTGTGACACACGA | 2430.9114559068544 | 2370.558973551602 | 2328.5883420922546 | 2404.8180887187087 | 2281.3387969886154 | 2126.7913197269436 | 2426.74218178428 | 2473.990084785536 | 1931.3679792697442 | 2778.245466735288 | 2310.0088105255246 | 2601.086149523335 |
| SE_244_1_19 | SE_244 | SE_244_1 | GACTTCTTATGCTTGTCAGG | 22.759313675454276 | 46.97144677779312 | 46.41376254382357 | 23.423552812195208 | 22.272484767317543 | 94.42780197565655 | 30.13249715741013 | 53.88514896170255 | 31.07948472388094 | 154.54947230330123 | 35.21354894093787 | 189.19100507465575 |
| SE_244_1_25 | SE_244 | SE_244_1 | AGTGTTGTACAAGAGATAAG | 204.8338230790885 | 215.86877668092157 | 234.04386644438694 | 201.70281588279207 | 284.239329411481 | 125.61496591692927 | 204.47051642528302 | 243.90120056349573 | 174.93309973155846 | 144.34337507572474 | 269.9705418805237 | 334.2761316656494 |
| SE_244_1_6 | SE_244 | SE_244_1 | GCACACGTTGTGGCAAACAT | 5403.711332657858 | 4239.422919816988 | 4018.2468040599597 | 5010.037684830641 | 4027.0773648335576 | 4122.769812863758 | 5247.359147840421 | 3926.0530462798365 | 4242.793657448661 | 5186.1554054985145 | 4225.625872912545 | 4764.595557248233 |
| SE_244_2_10 | SE_244 | SE_244_2 | GACTGTGTGACACACGAGGG | 1028.5042227621957 | 921.4398708324522 | 855.1982630415151 | 1111.317450089706 | 942.8685218164426 | 846.3849773414353 | 1027.7333851902383 | 1130.6427747051973 | 665.9889583688773 | 1627.8725077984514 | 977.7628755933749 | 1221.0364253898028 |
| SE_244_2_19 | SE_244 | SE_244_2 | ACTGGAGCTTACATTCCAGC | 1741.6293845930963 | 1482.099054712068 | 1459.5647029738561 | 1501.7099969359626 | 1516.6501532030516 | 1360.9731826032 | 1581.9561007640318 | 1424.6477102681708 | 1163.2607139509723 | 1478.4260841089383 | 1471.926345731203 | 1026.0420152515073 |
| SE_244_2_3 | SE_244 | SE_244_2 | CTGCCATTTACCCTGTACAG | 157.14764204480335 | 189.88457208044028 | 205.40558742798515 | 149.65047630013606 | 230.14900926228125 | 115.21924461249836 | 102.23525821264151 | 294.0049355629735 | 178.48504084285912 | 261.71349319285446 | 204.23858385743966 | 111.42537722188314 |
| SE_244_2_8 | SE_244 | SE_244_2 | GCACATGGTAGGCACTCAGT | 9669.456981543002 | 7764.280213120739 | 7967.366727735502 | 9357.709348471986 | 8636.42111715556 | 8849.357772305795 | 9663.922302626534 | 8890.104225190364 | 7675.744741520768 | 8832.64814352263 | 7736.416702324051 | 8449.757772659472 |
| SE_244_3_13 | SE_244 | SE_244_3 | CCAGGCAGCACTGTGTTTGG | 6756.264831084855 | 5629.577865942737 | 5700.992577989223 | 6358.193280021433 | 5770.694743760702 | 5323.475625141372 | 6888.504082306508 | 6010.557492956224 | 5502.844766682577 | 5918.078378104715 | 6952.328345905834 | 4956.107924348344 |
| SE_244_3_27 | SE_244 | SE_244_3 | CTGGCAGGCCAAGAAGGGAA | 187.49339361207572 | 184.8876096572708 | 223.18107095540694 | 192.59365645582727 | 135.75609762936406 | 177.5935725230238 | 168.95721620404964 | 123.84130726286024 | 120.76599778422309 | 286.4997293169688 | 295.7938111038781 | 52.23064557275772 |
| SE_244_3_50 | SE_244 | SE_244_3 | TCCCCTGCCAGTATCAGAGA | 97.53991575194691 | 104.9362108865591 | 114.55311606560711 | 119.72030810410883 | 102.87766774375245 | 76.23526896684917 | 110.84454431147587 | 103.98888396118035 | 126.98189472899928 | 125.38919451022554 | 73.94845277596953 | 153.20989368008932 |
| SE_244_3_54 | SE_244 | SE_244_3 | ATACTGGCAGGGGAGGACAT | 873.524134400769 | 712.5668415439679 | 764.345791679137 | 795.0994871250707 | 813.475991263455 | 931.2833681085394 | 961.0114271988302 | 924.5557137639489 | 854.2418372678134 | 555.5032919580922 | 875.643583664655 | 814.7980709350205 |
| SE_244_3_6 | SE_244 | SE_244_3 | AGACAGAGTGCTAGTAACCT | 1364.4755436855684 | 1598.0285829296 | 1519.8038415945632 | 1402.8105517525796 | 1458.3174550029344 | 1239.6897672216928 | 1434.522096815275 | 1272.445798288265 | 1164.1486992287976 | 1673.7999453225455 | 1623.3446061772358 | 1129.3426253842947 |
| SE_245_1_1 | SE_245 | SE_245_1 | GAGTCTTTTTCTGTTGGCCG | 1.083776841688299 | 21.986634661945715 | 19.75053725269088 | 35.13532921829281 | 15.9089176909411 | 34.65240439473635 | 15.066248578705064 | 78.46433971616335 | 50.615160830346767 | 10.206097227576498 | 73.94845277596953 | 0.0 |
| SE_245_1_16 | SE_245 | SE_245_1 | ACAAAGGCCGAGGGGTTCCG | 544.055974527526 | 798.5145952224831 | 738.670093250639 | 529.632555253525 | 704.2347564523261 | 473.8716300980196 | 471.3583483909156 | 417.84624282583377 | 576.3024453085352 | 672.873410075222 | 754.7437323007683 | 421.32720762024564 |
| SE_245_1_35 | SE_245 | SE_245_1 | GGATCCTGCAAGCAGCCAGC | 207.0013767624651 | 191.88335704970808 | 220.2184903675033 | 265.4669318715457 | 205.7553354695049 | 112.62031428289313 | 238.90765603375175 | 181.50780718678753 | 115.43808611727208 | 272.64859736525784 | 125.5949912226784 | 287.84889115653147 |
| SE_245_1_51 | SE_245 | SE_245_1 | GGTCCGGGGCAGCTTCCGCG | 242.76601253817896 | 410.7503111845313 | 386.12300329010674 | 239.44076208021767 | 357.4203507898101 | 304.94115867367987 | 219.53676500398808 | 259.97220990295085 | 371.17784613092095 | 391.47672937204135 | 368.56847891514974 | 331.95476964019355 |
| SE_245_1_9 | SE_245 | SE_245_1 | ACTGTCAGTCAGGGACGCAA | 2705.106996853994 | 2452.5091572915812 | 2455.979307372111 | 2482.8965980926992 | 2365.125763386577 | 2461.187022136149 | 2710.848583554147 | 2550.563717520587 | 2585.8131290268943 | 1871.3608273706334 | 2598.759911841215 | 2498.946220403275 |
| SE_245_2_17 | SE_245 | SE_245_2 | ATAGTCCCCACCCTGCAGGC | 8.670214732506391 | 17.98906472341013 | 20.738064115325425 | 2.602616979132801 | 1.0605945127294067 | 98.75935252499859 | 4.30464245105859 | 22.68848377334844 | 53.27911666951019 | 1.4580138896537853 | 16.432989505771005 | 4.642724050911798 |
| SE_245_2_2 | SE_245 | SE_245_2 | CTAGGAAATCAACCCCAGGG | 70.44549470973944 | 105.935603371193 | 117.51569665351074 | 87.18766880094883 | 84.84756101835254 | 85.76470087697247 | 48.42722757440914 | 62.393330376708214 | 63.04695472558706 | 88.93884712688809 | 29.344624117448227 | 174.10215190919243 |
| SE_245_2_26 | SE_245 | SE_245_2 | CTACGCGGCAACCTCACAAC | 930.9643070102487 | 746.5461860215205 | 763.3582648165026 | 880.9858474364531 | 765.7492381906317 | 629.8074498743332 | 966.3922302626535 | 875.3973322550273 | 803.6266764317786 | 658.2932711786841 | 909.683334764089 | 713.8188228276889 |
| SE_245_2_48 | SE_245 | SE_245_2 | GCAGACTTGAAGAACGCTGG | 2030.9978013238722 | 2270.619725088212 | 2311.8003854274675 | 2043.0543286192487 | 2236.7938273463187 | 2617.1228419124627 | 1998.4302579039504 | 2356.7662519565692 | 2318.529560401518 | 2103.185035855585 | 2126.8983560326474 | 2033.5131342993675 |
| SE_245_2_9 | SE_245 | SE_245_2 | GCAGCCATGTCCTCCCAGGG | 2260.7584917671914 | 2608.414384894469 | 2729.52424832188 | 2286.3990161681654 | 2615.426068390717 | 2919.4650702565373 | 2308.3645148021763 | 2778.393908744628 | 2671.9477009759357 | 2399.1618554253037 | 2637.4948156762466 | 2266.810017857685 |
| SE_246_1_11 | SE_246 | SE_246_1 | TATTTGCTGCCTTTTGTGCA | 338.13837460674927 | 337.7946598062569 | 358.4722551363395 | 356.55852614119374 | 363.7839178661865 | 186.25667362170788 | 456.29209981221055 | 335.60048914744567 | 439.5521252345904 | 283.58370153766112 | 408.47716771487933 | 256.51050381287683 |
| SE_246_1_14 | SE_246 | SE_246_1 | GCAGGTCCTTCGGATCCACA | 1355.8048289520618 | 1313.2017248089396 | 1262.0593304469473 | 1363.7712970655878 | 1359.6825163190993 | 1561.9571280927941 | 1395.7803147557477 | 1264.8829703641754 | 1145.501008394469 | 1408.4414174055566 | 1562.3077880129435 | 1472.9042051517679 |
| SE_246_1_15 | SE_246 | SE_246_1 | TGCAAAACCATAGAGACCGG | 575.4855029364867 | 714.5656265132358 | 798.9092318713462 | 579.0822778570482 | 784.839939419761 | 678.320816026964 | 464.9013847143277 | 727.9221877282624 | 880.8813956025684 | 547.4842155649964 | 496.511040067224 | 549.00211902032 |
| SE_246_1_29 | SE_246 | SE_246_1 | GCTGGCTGCGGAAACCCAGA | 1517.2875783636184 | 1698.9672238776236 | 1741.9973856873357 | 1592.8015912292742 | 1826.3437509200385 | 1998.5774234664188 | 1698.1814469426138 | 2017.3843488468988 | 1638.3328375874382 | 2117.036167777296 | 2015.3887843863442 | 2040.4772203757352 |
| SE_246_1_31 | SE_246 | SE_246_1 | GGAAGCAAACCACATCCCCT | 3695.679030157099 | 3611.804439466901 | 3529.42100705586 | 3694.414801879011 | 3330.266769970337 | 3168.9623818986393 | 4080.8010436035433 | 3900.5285020348819 | 3662.9392710288253 | 4199.080002202902 | 3477.924850399964 | 3349.725402732862 |
| SE_246_2_20 | SE_246 | SE_246_2 | ACAGGTCTGGGAAGGGCGCG | 1899.860803479588 | 1871.8621237192876 | 1901.9767374341318 | 1850.4606721634213 | 2045.8868150550256 | 1587.9464313887931 | 1857.4532176317816 | 1691.237394605015 | 1664.9723959221933 | 1381.4681604469615 | 1586.9572722716 | 1818.7871469446968 |
| SE_246_2_28 | SE_246 | SE_246_2 | GAGGGGACTGTGGAAGCACA | 3771.54340907528 | 3619.799579343972 | 3632.123800769583 | 3983.305286562752 | 3608.142532305442 | 3798.7698317729723 | 4229.311208165044 | 4015.861627882674 | 3946.2065746550547 | 4615.342967699057 | 3643.4285304223718 | 3870.8711774477115 |
| SE_246_2_30 | SE_246 | SE_246_2 | GGTGCAGCTCAGGGAAGAAG | 2164.3023528515328 | 2218.6513158872494 | 2274.274364647355 | 2325.4382708551575 | 2183.7641017098485 | 1691.037334463134 | 2339.5731721503435 | 2329.35100073044 | 2129.388696224757 | 1741.5975911914466 | 2572.9366426178603 | 1949.944101382955 |
| SE_246_2_6 | SE_246 | SE_246_2 | TATTAGATAGAAACAGAGGT | 18.42420630870108 | 51.968409200962604 | 36.53849391747813 | 49.44972260352321 | 54.09032014919974 | 46.78074593289407 | 29.056336544645482 | 60.50262339559584 | 108.33420389467071 | 131.95025701366757 | 71.60088284657367 | 12.767491140007444 |

| | | | | | | | | | | | | | | | |
|---|---|---|---|---|---|---|---|---|---|---|---|---|---|---|---|
| SE_246_2_7 | SE_246 | SE_246_2 | ATCTTAGCAGAAGGCCTTGA | 305.6250693561003 | 255.84447606627742 | 297.24558565299776 | 372.17422801599054 | 233.33079280046948 | 243.43314087302286 | 311.0104170889831 | 379.0867497130302 | 185.58892306546048 | 195.37386121360723 | 342.74520969179525 | 164.8167038073688 |
| SE_246_3_10 | SE_246 | SE_246_3 | TCATCTGCGTGTTATCAGCC | 2114.448618133871 | 1879.8572635963587 | 1943.4528656647826 | 2052.1634880462134 | 1968.463415625779 | 1972.5881201703667 | 1932.784460525307 | 1926.630413753505 | 1461.6237673002295 | 1539.6626674743973 | 1684.3814243415281 | 2098.5112710121325 |
| SE_246_3_2 | SE_246 | SE_246_3 | AGAAAGGAGCAGTCTCCCCA | 444.3485050922026 | 621.6221254422834 | 635.9672995366464 | 498.4011515039314 | 527.1154728265152 | 510.2566547124927 | 420.77879959097714 | 424.46371725972705 | 506.15160836034676 | 620.3849100476856 | 606.8468267488294 | 501.41419749847415 |
| SE_246_3_21 | SE_246 | SE_246_3 | TGAGGAGGAAAGTCTCCTGT | 5651.896229404479 | 6169.249807645041 | 5882.6975207139785 | 5548.7793995111315 | 5939.329271284678 | 5854.523722490706 | 5306.547981542477 | 5134.214807210641 | 5585.427397520318 | 6441.505364490424 | 4733.874762626748 | 5730.282159837886 |
| SE_246_3_22 | SE_246 | SE_246_3 | ATAAGATGGCCATGCCCGCG | 516.9615534853186 | 604.6324532035072 | 560.915257976421 | 612.9162985857746 | 642.7202747140204 | 758.0213461348576 | 606.9545855992611 | 570.0481548053796 | 707.7242664266603 | 645.1711461718 | 604.4992568194334 | 740.5144861204317 |
| SE_246_3_8 | SE_246 | SE_246_3 | CCAAATACACACAATCCACG | 3318.524689249571 | 3194.0583808899323 | 3106.7595098482757 | 3102.3194391262987 | 3191.328888802785 | 2949.785924101932 | 3170.3691652046514 | 3094.1419745903936 | 3040.461591273381 | 3288.5503281141127 | 3211.4756634135338 | 2921.4341090362486 |
| SE_246_4_10 | SE_246 | SE_246_4 | AAAACCGCTAGAAGGACAGG | 121.38300626908948 | 160.9021900260573 | 147.14150253254707 | 126.22692348794084 | 79.5445884547055 | 170.66309164407653 | 120.52998862964051 | 59.557269905039654 | 121.65398306204825 | 139.2403264619365 | 129.1163461167722 | 129.99627342553035 |
| SE_246_4_11 | SE_246 | SE_246_4 | GGAACCGCCTGTCCTTCTAG | 713.1251618309007 | 745.5467935368865 | 810.7595542229607 | 814.6191144685666 | 738.1737808596671 | 1118.4063518401156 | 795.2826928330745 | 737.3757226338244 | 751.2355450400936 | 1021.3387297024766 | 630.3225260427879 | 571.0550582621511 |
| SE_246_4_3 | SE_246 | SE_246_4 | CAGCACGTGAGGTTGCTTAA | 2143.710592859455 | 2162.6853367477515 | 2153.7960874059404 | 2174.486486065455 | 2239.975610884507 | 2059.2191311572074 | 1919.870533172131 | 2133.6628281853095 | 2439.2955581857414 | 3335.9357795278606 | 2045.9071934684903 | 1942.9800153065873 |
| SE_246_4_5 | SE_246 | SE_246_4 | GCTAGGATTCTAGTTACCAA | 392.3272166911642 | 505.69259722475147 | 438.46192700973756 | 415.11740817168175 | 419.9954270408451 | 721.6363215203845 | 319.61970199110027 | 426.3544242408394 | 556.7667691963815 | 540.9231530615543 | 462.471276090984 | 588.4652734530704 |
| SE_246_4_6 | SE_246 | SE_246_4 | TTTTGTTTATGCCATCAGGA | 3.2513305250648967 | 0.9993924846338962 | 8.887741763710896 | 1.3013084895664004 | 3.1817835381882205 | 0.8663101098684087 | 0.0 | 0.0 | 3.551941111300679 | 0.7290069448268927 | 12.911634611677218 | 1.1606810127279494 |
| SE_247_1_11 | SE_247 | SE_247_1 | GGAGTCCAGACACCTTTCCA | 262.27399568856833 | 359.78129446820265 | 385.13547642747216 | 348.7506752037953 | 355.29916176435125 | 265.9572037296015 | 338.99059302086397 | 266.5896843368442 | 287.70723001535504 | 562.7933614063611 | 331.007360044816 | 233.29688355831783 |
| SE_247_1_14 | SE_247 | SE_247_1 | TGGTGGGGAGGCCTAATGGA | 21.675536833765978 | 31.980559508284678 | 30.613332741670867 | 50.751031093089615 | 71.05983235287025 | 4.331550549342044 | 23.675533480822246 | 25.524544245016994 | 39.95933750213264 | 42.282402799959776 | 26.997054188052367 | 6.964086076367696 |
| SE_247_1_4 | SE_247 | SE_247_1 | TCAATGAACGGAGAGTAGGA | 851.8485975670029 | 731.555298752012 | 810.7595542229607 | 866.6714540512227 | 870.748094950843 | 1134.8662439276154 | 727.4845742289017 | 684.4359271626779 | 791.1948825422263 | 450.5262919030197 | 936.6804018289474 | 373.7392860983997 |
| SE_247_1_5 | SE_247 | SE_247_1 | TCGCTTTTAGGGGAGTACAG | 138.72343573610226 | 154.90583511825392 | 150.1040831204507 | 188.68973098712806 | 142.1196647057405 | 69.3048087894727 | 176.4903404934022 | 124.78666075341643 | 177.59705556503397 | 141.42734729641717 | 112.68335661100119 | 226.33279748195014 |
| SE_247_1_7 | SE_247 | SE_247_1 | ACTCTCCGTTCATTGACCAC | 518.0453303270069 | 457.72175796232443 | 450.3122493613521 | 419.02133364038093 | 489.9946648809859 | 349.1229742769687 | 424.00728142927113 | 484.9663406553229 | 302.8029797383829 | 289.41575709627637 | 422.56258729125443 | 312.2231924238184 |
| SE_247_2_10 | SE_247 | SE_247_2 | GATATTGCAAGAGGACCCAC | 291.5359704141524 | 373.7727892530772 | 389.08558387801037 | 338.3402072872641 | 355.29916176435125 | 491.19783229538774 | 275.49711686774975 | 365.8518008452436 | 404.0333014104522 | 371.79354186171525 | 484.7731904202447 | 513.0210076257537 |
| SE_247_2_13 | SE_247 | SE_247_2 | ACTCAAATCTGACTAACTAC | 842.0946059918083 | 964.4137476717099 | 1015.1776147883113 | 789.894253166805 | 965.1410065837601 | 1081.1550171157742 | 772.6833199650168 | 894.304402066151 | 1228.0836392322099 | 1176.6172089506047 | 816.9543354297587 | 1147.913521587942 |
| SE_247_2_17 | SE_247 | SE_247_2 | AGATATTGAGAGGATGGCAT | 423.7567451001249 | 437.7339082696465 | 437.474400147103 | 434.63703551517773 | 449.6920733972685 | 501.59355361380864 | 415.3979965271539 | 547.3596710320311 | 416.46509530000463 | 450.5262919030197 | 519.9867393611826 | 328.4727266020097 |
| SE_247_2_20 | SE_247 | SE_247_2 | TCAATCCACTATATAAACAA | 215.67159149597148 | 313.8092401750434 | 368.3475197626849 | 239.44076208021767 | 270.45160074599875 | 368.1817966940737 | 146.35784333599204 | 369.63321480746833 | 225.54826056759313 | 207.03797237308375 | 347.440349550587 | 685.9624785222181 |
| SE_247_2_28 | SE_247 | SE_247_2 | ATGCTCTTATCAAAATAAAA | 16.256652625324485 | 21.986634661945715 | 23.700644703229056 | 13.013084895664004 | 31.8178353818822 | 1.7326202197368175 | 6.456963676587884 | 38.75949311280358 | 12.431793889552377 | 11.664111117230282 | 15.259204541073078 | 0.0 |
| SE_248_1_13 | SE_248 | SE_248_1 | GAGGCCCAAGAGAAAATAGG | 447.59983561726744 | 634.6142277425241 | 602.3913862070718 | 595.9992882214113 | 548.3273630811033 | 842.9197369019616 | 474.5868302292095 | 477.40351273087344 | 792.9708530978767 | 585.3925766959948 | 462.471276090984 | 417.8451645820618 |
| SE_248_1_18 | SE_248 | SE_248_1 | ATTTGCCTCAAGGATTCCAA | 37.93218945909046 | 48.970231747060915 | 46.413762542382357 | 42.943180155691216 | 32.87842989461161 | 5.197860659210452 | 43.0464245105859 | 54.83050245225873 | 46.17523444690883 | 78.00374309647752 | 142.02798072844942 | 17.410215190919242 |
| SE_248_1_19 | SE_248 | SE_248_1 | CTTTTCCTGGAGAGGCAACA | 11.921545258571289 | 17.98906472341013 | 28.63827901640178 | 5.205233958265602 | 15.9089176909411 | 22.524062856578627 | 2.152321225529295 | 9.45353490556185 | 7.103882222601358 | 4.374041668961356 | 8.216494752885502 | 2.321362025455899 |
| SE_248_1_20 | SE_248 | SE_248_1 | CAAGACTATGAGCTTCTTAA | 71.52927155142773 | 118.92770567143364 | 114.55311606560711 | 96.29682822791364 | 138.9378811675523 | 55.44384703157816 | 86.0928490211718 | 212.70453537514163 | 79.91867500426528 | 180.79372231706938 | 224.19292825730446 | 34.820430381838484 |
| SE_248_1_7 | SE_248 | SE_248_1 | TCTACTATGGTAATAAACAC | 80.19948628493412 | 99.93924846338962 | 97.76515940081985 | 84.58505182181602 | 69.99923784014085 | 223.50800834640945 | 89.32133085946575 | 70.90151179171387 | 93.23845417164283 | 216.51506261355871 | 44.6038286585213 | 98.6578860818757 |
| SE_248_2_11 | SE_248 | SE_248_2 | TTTGTACTGACAAGGAGAGG | 128.96944416090756 | 125.92345306387092 | 78.01462214812898 | 127.52823197750725 | 133.63490860390525 | 273.7539947184172 | 78.55972473181926 | 247.68261452572048 | 109.22218917249589 | 132.67926395849446 | 180.76288456348107 | 156.69193671827318 |
| SE_248_2_13 | SE_248 | SE_248_2 | GCTGTTGATTCAGAACTGGC | 574.4017260947984 | 766.5340357141984 | 735.7075126627353 | 598.6019052005443 | 683.0228667179779 | 640.2031711927541 | 465.97754532709234 | 555.8678524470367 | 656.2211203128005 | 368.8775140248077 | 419.0412323971607 | 820.6014759986602 |
| SE_248_2_14 | SE_248 | SE_248_2 | GTGTGGTCGCTGCAGATGAA | 644.8472208045379 | 147.91008772581662 | 278.4825752629414 | 556.9600335344194 | 127.2713415275288 | 1206.7699830466934 | 793.1303716075452 | 363.9610938641312 | 1008.7512756093929 | 40.82438891030599 | 125.5949912226784 | 322.66932153836996 |
| SE_248_2_3 | SE_248 | SE_248_2 | GCTATGTTAGAGGAATGGGG | 383.6570019576578 | 471.713252747199 | 458.21246426242845 | 473.67629020216975 | 473.0251526773154 | 563.9678815243341 | 430.464245105859 | 415.01018235416524 | 379.1697136313475 | 479.6865696960954 | 484.7731904202447 | 265.7959519147004 |
| SE_248_2_4 | SE_248 | SE_248_2 | TTTGCCCTAAATCAGTAAAA | 20.591759992077 68 | 15.990279754142339 | 31.60085960430541 | 35.13532921829281 | 20.15129574185873 | 0.0 | 26.904015319116187 | 13.23494886778659 | 42.62329333560815 | 96.22891671714983 | 16.432989505771005 | 2.321362025455899 |
| SE_248_3_2 | SE_248 | SE_248_3 | ATATTCCTCCAACTGCAGGA | 1.083776841688299 | 0.9993924846338962 | 5.925161175807264 | 0.0 | 6.363567076376441 | 0.0 | 0.0 | 0.0 | 0.0 | 0.0 | 0.0 | 0.0 |
| SE_248_3_34 | SE_248 | SE_248_3 | AAGTATTGCTATGAAATTGG | 104.0425768020767 | 97.94046349412183 | 122.45333096668347 | 93.69421124878083 | 118.78658542569356 | 50.2459863723677 | 95.77829453605362 | 179.61716320567515 | 79.03068972644012 | 104.24799311024564 | 181.936669528179 | 112.5860582346111 |
| SE_248_3_40 | SE_248 | SE_248_3 | TCCAGGTAGCTTCAGCCGAT | 1708.032302500759 | 1426.1330755725699 | 1453.6395417980489 | 1642.2513138327975 | 1309.8342232208174 | 966.8020826131442 | 1734.7709077766117 | 1493.6585150787723 | 1363.0574014616357 | 1479.884097998592 | 1795.8909959878315 | 1326.6583975480462 |
| SE_248_3_6 | SE_248 | SE_248_3 | TGCAGTTGGAGGAATATGGG | 1573.64397413141 | 1773.9216602251656 | 1730.1470633357212 | 1488.6969120639621 | 1657.7092233960627 | 1498.7164900723471 | 1388.2471904663953 | 1880.308092716252 | 1692.4999395347736 | 2118.49418166695 | 1449.6244314019423 | 2189.0443900049127 |
| SE_248_3_8 | SE_248 | SE_248_3 | TGGTGCAGTCTGTTTCTGTG | 211.3364841292183 | 209.8724217731182 | 235.0313933070215 | 205.60674135149128 | 166.51333849851687 | 247.7646914223649 | 152.81480701257993 | 150.31120499843342 | 247.74789251322238 | 288.6867501514495 | 142.02798072844942 | 182.22691899828806 |
| SE_248_4_11 | SE_248 | SE_248_4 | AAGGTATTTATATAACAAGA | 17.340429467012783 | 33.97934447755247 | 30.613332741670867 | 15.615701874796805 | 32.87842989461161 | 17.326202197368175 | 10.761606127646475 | 54.83050245225873 | 39.07135222430747 | 40.0953819654791 | 70.42709788187574 | 17.410215190919242 |
| SE_248_4_12 | SE_248 | SE_248_4 | GCCAAGGGTTATAATAACTG | 602.5799239786942 | 784.5231004376085 | 799.8967587339806 | 622.0254580127394 | 935.4443602273368 | 570.0320522934129 | 698.4282376842563 | 697.6708760304646 | 579.8543864198359 | 1153.288986716144 | 987.1531553109584 | 590.7866354785263 |
| SE_248_4_24 | SE_248 | SE_248_4 | GGATGTGGTGTAACTAATCA | 751.0573512899912 | 715.5650189978696 | 750.5204156022535 | 869.2740710303555 | 777.4157778306552 | 860.2459309093299 | 856.6238477606594 | 1152.3859049879895 | 678.4207522584297 | 909.8006671439621 | 1068.1443178751153 | 379.5426911620395 |
| SE_248_4_7 | SE_248 | SE_248_4 | GTTTGCTCCACTCACATGGG | 67.19416418467453 | 69.95747392437274 | 111.59053547770347 | 81.98243484268323 | 75.30221040378788 | 23.390372966447035 | 85.01668840840715 | 184.3439306584561 | 103.0062922277197 | 51.030486137882484 | 281.70839152750295 | 88.21175696732416 |
| SE_249_1_10 | SE_249 | SE_249_1 | CACGCGCCACATCAAGGCAG | 3685.9250385819046 | 3109.110019696051 | 2919.1294059477123 | 3103.620747615865 | 2601.638339725235 | 2645.7110755381204 | 3202.653983587591 | 2643.2083595950935 | 2647.084113196831 | 1299.0903756815228 | 2361.6553489722332 | 2163.5094077248978 |
| SE_249_1_24 | SE_249 | SE_249_1 | CGCAGCCGCCATCTTGGCTC | 179.90695572025763 | 200.87788941141312 | 256.75698428498146 | 222.52375171585447 | 195.14939034221084 | 184.52405340197106 | 235.6791741954578 | 204.19635396013595 | 247.74789251322238 | 307.6409307169487 | 407.30338275018136 | 102.13992912005955 |
| SE_249_1_27 | SE_249 | SE_249_1 | GGCTGCGAAGCCAACCCTCA | 406.4163156331121 | 484.7053554074396 4 | 501.6636462183484 | 472.37498171260336 | 510.14596062284465 | 385.5079988914419 | 437.99736939525115 4 | 486.8570476364325 5 | 570.086548363159 | 468.751465523692 | 511.77024460829705 | 159.01329874372908 |
| SE_249_1_28 | SE_249 | SE_249_1 | GAGAGCGAGTCCGTGAGGGT | 4485.752347747869 | 4648.174446032252 | 4648.288942420799 | 4352.876897599609 | 4546.768676070967 | 4619.165505818355 | 4364.90744537341 | 4850.608760043785 | 4245.457613282137 | 3657.4278421965205 | 4997.9763796837815 | 4903.877278775586 |
| SE_249_1_31 | SE_249 | SE_249_1 | GTTCTGCCTCGCGCAGCTCG | 0.0 | 2.9981774539016886 | 3.950107450538176 | 2.602616979132801 | 2.1211890254588135 | 0.0 | 0.0 | 0.0 | 0.0 | 0.0 | 0.0 | 0.0 |
| SE_249_2_30 | SE_249 | SE_249_2 | GCCCCTGACGGAGGCTTTGC | 19.50798315038938 | 29.981774539016886 | 30.613332741670867 | 29.930095260027 21 | 10.605945127294067 | 62.37432791052543 | 15.066248578705064 | 0.945353490556185 | 38.1833669464823 | 4.374041668961356 | 0.0 | 0.0 |
| SE_249_2_33 | SE_249 | SE_249_2 | CAGCCAGCAAAGCCTCCGTC | 1631.0841467408898 | 1860.8688063883146 | 1705.4588917698575 | 1764.574311852039 | 1489.074695872087 | 1837.443743030895 | 1748.760995742552 | 1509.7295244182274 | 1648.9886609213404 | 1669.425903653584 | 1272.3829017325552 | 1457.8153519863044 |
| SE_249_2_39 | SE_249 | SE_249_2 | GGGGATTGAGTTGGGGGCAA | 186.4096167703874 | 246.84994370457235 | 252.80687683444327 | 235.53683661151848 | 249.2397104914106 | 191.45453428091832 | 209.85131948190626 | 269.4257448085127 | 239.7560250127958 5 | 240.57229179287458 | 381.4801135268269 | 125.35354937461854 |
| SE_249_2_41 | SE_249 | SE_249_2 | GGGACGTCAACAGTCTAGGC | 2786.3902599806165 | 2421.5279902679304 | 2394.752641888769 | 2434.748183978735 | 2523.1543457832586 | 1905.882241710499 | 2685.0207288477955 | 2438.066652144401 | 1702.2677775908505 | 1975.608820480879 | 2199.673023843919 | 2444.3942128050617 |
| SE_249_2_52 | SE_249 | SE_249_2 | TTCACAGCCAATTCATCCTT | 138.72343573610226 | 163.900367479959 | 161.95440547206522 | 134.03477442533924 | 200.45236290585788 | 123.01603560131404 | 163.5764131402264 | 180.56251669623134 | 258.40371584712443 | 174.23265981562736 | 302.83652089206566 | 244.90369368559732 |
| SE_249_3_25 | SE_249 | SE_249_3 | GGGTCCTTCCAGGAGCGCGC | 20.59175999207768 | 44.97266180852533 | 63.20171920861082 | 20.820935833062407 | 38.18140245825864 6 | 48.51336615263089 | 20.4470516425283 | 0.945353490556185 | 55.05508722516053 | 41.553395855132884 | 3.521354894093787 | 56.87336962369525 |
| SE_249_3_26 | SE_249 | SE_249_3 | CTTGCCGGCGCGCTCCTGGA | 1462.0149594375152 | 1739.9423157476133 | 1840.75007195079 | 1503.0113054491926 | 1738.3144063634977 | 2167.5078948907585 | 1371.0286206621608 | 1871.7999113012463 | 1964.2234543492755 | 1425.208577136575 | 2388.6524031602858 | 2198.329838106736 |
| SE_249_3_30 | SE_249 | SE_249_3 | CAGGAAGCAGGGCTGGCAGT | 14844.491400604631 | 12377.475922190804 | 12453.701264684234 | 13813.389616747341 | 12440.773634315941 | 12614.3415097939 | 14093.399384765824 | 12965.523122978077 | 12481.521065110586 | 12634.419360794876 | 13935.175100893814 | 13178.372218513137 |
| SE_249_3_31 | SE_249 | SE_249_3 | AGAGCTTCAGGAAGCAGGGC | 277.4468714722045 | 310.8110627211417 | 360.4473048616086 | 342.24413275596334 | 328.7842989461941 | 326.5989114203901 | 305.62961402515987 | 346.9447310341199 | 509.7035494716475 | 527.8010280546703 | 281.70839152750295 | 425.96993167115744 |
| SE_249_3_32 | SE_249 | SE_249_3 | CCGAAGAGCTTCAGGAAGCA | 971.0640501527158 | 886.4611338702659 | 942.100626953355 | 795.0994871250707 | 1009.6859761183953 | 641.0694813026224 | 964.2399090371241 | 1054.0691419701463 | 978.5597761633371 | 847.8350768336761 | 1723.1163281765598 | 745.1572101713435 |
| SE_249_4_0 | SE_249 | SE_249_4 | GGAGGTATAGGAGTAGACTC | 1284.2755574006342 | 1453.116672657685 | 1472.4025521881051 | 1329.9372763368613 | 1555.8921501740397 | 1216.299394255246 | 1426.9889725259225 | 1458.6804359281934 | 1621.46111730876 | 1882.2959315430369 | 1250.0809874032943 | 985.4181798060291 |
| SE_249_4_10 | SE_249 | SE_249_4 | GCATGTATCAGAGCTCCAGC | 1777.3940203688103 | 1964.8056247902398 | 1947.402973115321 | 1892.1025438295462 | 2235.733232833589 | 1831.379572261816 | 1661.5919861086156 | 2158.2420189397703 | 1712.9236009247525 | 2028.0973205084154 | 2814.7363453456337 | 2222.7041393740233 |
| SE_249_4_13 | SE_249 | SE_249_4 | GAGAAAACCGAGACACAGCT | 870.272803875704 | 1018.3809418419402 | 1042.8283669420784 | 1038.4441746739876 | 921.6566315618545 | 1425.9464408434008 | 1070.7798097008242 | 1058.7959094229273 | 1058.4784511676023 | 1958.8416607498605 | 946.0706815465307 | 993.5429468951247 |
| SE_249_4_16 | SE_249 | SE_249_4 | ATAGGGTCTCTCCCTCACCC | 912.5401007015477 | 957.4180002792725 | 941.1131000907204 | 1056.6624935279171 | 854.8391772599018 | 729.4331125092001 | 1013.743297224298 | 850.8181415005665 | 1075.3501714462807 | 650.2741947855883 | 1159.699545121554 | 850.7791823295869 |
| SE_249_4_29 | SE_249 | SE_249_4 | CAGCATTGTTCAGGAAAGGG | 2700.771889487241 | 2466.500652076456 | 2494.9817805094765 | 2652.066270136324 | 2618.6078519289053 | 2359.8287392815455 | 3050.9153371877755 | 2845.514006574117 | 2345.1691187362735 | 2690.7646333560606 | 2842.907184498384 | 2097.3505899994047 |
| SE_25_1_1 | SE_25 | SE_25_1 | GTGAGAGAGTGGGAGTACAA | 1907.447241371406 | 2143.6968795397074 | 2033.3178101645262 | 2057.368722004479 | 2239.975610884507 | 2167.5078948907585 | 2101.7416767293566 | 2040.0728326202473 | 2059.2378592765685 | 2858.436230666246 | 1804.107490740717 | 1663.2558912391517 |
| SE_25_1_15 | SE_25 | SE_25_1 | CCCACCCCCTCCCCAAACAA | 198.3311620289587 | 241.8529812814029 | 219.23096350486878 | 206.90804984105768 | 212.1189025458813 5 | 401.9676909789416 | 192.6327496848719 | 117.22383282896693 | 146.5175708413 | 387.10268770808 | 132.63770101086598 | 409.72039749296613 |
| SE_25_1_2 | SE_25 | SE_25_1 | AGTAACGGTCTGTTTAGGGA | 251.43622727168534 | 277.83111072822317 | 301.1956931035359 | 282.3839422359089 | 290.60289648785744 | 226.97324878552308 | 263.65935012733866 | 256.19079594072616 | 254.85177473582374 | 651.7322086752421 | 319.2695103978367 | 200.79781520193526 |
| SE_25_1_34 | SE_25 | SE_25_1 | GCTCATTGTTGAACACTGGG | 645.9309976462262 | 744.5474010522527 | 745.5827812890808 | 746.9510730111139 | 734.9919973214788 | 727.7004922894633 | 654.3056525609056 | 845.1460205572293 | 878.217439769093 | 661.2092989579917 | 863.9057340176757 | 773.0135544768143 |
| SE_25_1_7 | SE_25 | SE_25_1 | TGAAAAACAGACGCCCTGGG | 403.1649851080472 | 575.6500711491242 | 538.2021401358265 | 480.18283265000 18 | 623.6295734848911 | 654.930443060517 | 401.4079085612135 | 880.1240997078082 | 642.901341145423 | 215.7860556876602 | 1023.5404892165941 | 854.2612253677707 |
| SE_25_2_10 | SE_25 | SE_25_2 | AAGCACTAAGTAGTGCCCAG | 264.4415493719449 | 286.8256430899282 | 300.2081662409014 | 312.3140374959361 | 308.63300320425736 | 335.2620125190742 | 305.62961402515987 | 303.4584704685354 | 143.85361500767752 | 337.5302154548513 | 245.32105762186717 | 496.77147344756236 |
| SE_25_2_22 | SE_25 | SE_25_2 | TAGCTGCACCACTTACCTGC | 10746.731162181171 | 9470.2431843908 | 9546.422181088137 | 9442.294400293802 | 9479.593754775438 | 9402.06362240184 | 9219.467965954735 | 10118.118409422848 | 9658.615866904373 | 9217.563810391232 | 8732.960137352591 | 11313.157831059323 |
| SE_25_2_29 | SE_25 | SE_25_2 | ATCTGCATCATCAACACAAA | 384.7407787993461 | 482.70657007817186 | 411.7987017186049 | 369.57161103685775 | 374.38986299348056 | 394.171099990126 | 351.90452037403975 | 255.24544245016995 | 463.5283150247386 | 334.61418767554375 | 268.79675691582577 | 622.1250228221809 |
| SE_25_2_38 | SE_25 | SE_25_2 | AGAGGGATGGGGGCTGAAAA | 825.8379533664838 | 1002.3906620877979 | 959.8761104807768 | 1017.6232388409252 | 961.9592230455719 | 1123.6042124993262 | 970.6968727137 12 | 1237.4677191380463 | 1215.6518453426575 | 1296.1743479022152 | 924.942552181968 | 1039.9701874042428 |
| SE_25_2_4 | SE_25 | SE_25_2 | GCCATGGCAGCCTATAAGAA | 784.6544333823284 | 921.4398708324522 | 875.9363271568405 | 912.2172551186046 7 | 1105.1394822640418 | 973.7325634920915 | 817.8820657011131 | 1195.872165553574 | 935.9364828277289 | 1543.3077021985318 | 1116.2695014277306 | 793.9058127059175 |
| SE_25_3_18 | SE_25 | SE_25_3 | TTGCCTACTAAGAAGTCTAG | 23.843090517142578 | 48.970231747060915 | 53.32645058226538 | 42.943180155691216 | 34.99961892007042 5 | 33.78609428486794 | 38.74178205952731 | 15.12565584889896 | 11.543808611727208 | 104.977000055072 54 | 22.30191432926065 | 0.0 |
| SE_25_3_22 | SE_25 | SE_25_3 | GGAAGAATGTGCTCTCTCAG | 717.4602691976539 | 843.4872570310084 | 855.1982631455751 | 727.4314456716178 | 764.6886436779023 | 736.3635938814474 | 712.4183256501966 | 767.6270343316222 | 713.0521780316114 | 756.7092087303146 | 1002.4123598520314 | 626.7677468730927 |
| SE_25_3_35 | SE_25 | SE_25_3 | GCTTTCTGGCGGGGAAAACG | 215.67159149597148 | 305.81410029797223 | 253.79440369707783 | 208.20935833062407 | 231.20960377501066 | 440.08553581315164 | 167.88105559128502 | 394.21240556192913 | 174.04511445373328 | 309.82795155142935 | 289.9248862803885 | 221.69007343103834 |
| SE_25_3_37 | SE_25 | SE_25_3 | TTAGGACAGGGGAACCCCTG | 325.13305250648966 | 340.7928372601586 | 358.47225113633395 | 376.07815344864897 | 266.20922266950811 | 522.38499625065605 | 344.37139068486872 | 311.96665188354103 | 515.0314611385985 | 271.91195904204309 5 | 241.7997027277734 | 439.8981038238928 |
| SE_25_3_8 | SE_25 | SE_25_3 | GTCAGGACTCCGGGGAGCAG | 407.5000924748004 | 504.69320474011755 | 536.2270864105574 | 473.67629020216975 | 597.114710666656 | 560.5026410848604 | 391.72246304633165 | 434.86260565584513 | 451.09652113518626 | 881.3693962957133 | 423.7363722559524 | 807.8339848586528 |
| SE_25_4_11 | SE_25 | SE_25_4 | GCTTTCAAAGAGCACCAAGT | 133.30455152766078 | 152.90705014898612 | 174.79225468631 43 | 114.51514708184324 | 165.45274398578746 | 97.02673230526177 | 128.06311291899306 | 233.50231216737777 | 85.2465866712163 | 123.93118062015175 | 219.49778839851274 | 205.44053925284706 |
| SE_25_4_30 | SE_25 | SE_25_4 | CGACAGGACGTGGTTCACAC | 10134.397246627283 | 7793.262595175122 | 7719.49748521423 | 8418.164619005045 | 8378.696650562313 | 8344.298978252513 | 9497.117407648015 | 7450.330859073294 | 7347.190188725454 | 7457.741045579111 | 6886.59638788275 | 7747.545759959063 |
| SE_25_4_32 | SE_25 | SE_25_4 | GAACTTTCCAGGGTTCCTCA | 5.418884208441 4945 | 15.990279754142339 | 11.850322351614528 | 14.314393385230405 | 21.211890254588134 | 4.331550549342044 | 9.685445514881827 | 17.961716320567515 | 15.095749723027886 | 38.637368075825 31 | 19.954344399864794 | 27.856344305470785 |
| SE_25_4_4 | SE_25 | SE_25_4 | TTGAGAAATGCGCTTCCCAT | 1010.0800164534946 | 1019.3803343265741 | 1002.3397655740622 | 1034.5402492052883 | 903.6265248454546 | 829.9250852593555 | 1058.9420429604131 | 876.3426857455835 | 978.5597761633371 | 890.8464865784629 | 1282.9469664148364 | 994.7036279078527 |
| SE_25_4_5 | SE_25 | SE_25_4 | ACACAACTCCTCTTAGGCTG | 2398.3981506562054 | 2484.489716799866 | 2497.455435602762 | 2410.0233226769737 | 2658.9104434126225 | 2974.042607178247 | 2356.791741954578 | 2538.2741221433566 | 2281.234178732861 | 3084.428383562583 | 2840.5596145689883 | 3130.35669132728 |
| SE_25_5_12 | SE_25 | SE_25_5 | GCAAGCACAAACTGGAGCCG | 2759.295838938409 | 3293.997629353322 | 3243.0382168918427 | 2999.516068450553 | 3449.053355396031 | 3790.9730407841566 | 2632.2888588223277 | 3303.0650960033104 | 3199.4109560040865 | 2595.264723583738 | 3800.7157156918943 | 4389.695590137105 |

| | | | | | | | | | | | | | | | |
|---|---|---|---|---|---|---|---|---|---|---|---|---|---|---|---|
| SE_25_5_14 | SE_25 | SE_25_5 | GCCCATCTTCTAAGACACAG | 275.27931778882794 | 347.7885846525959 | 391.0606376032794 | 338.3402072872641 | 357.4203507898101 | 240.83421054341764 | 361.58996588892154 | 475.51280574976107 | 404.0333014104522 | 855.125146281945 | 474.20912573796335 | 266.95663292742836 |
| SE_25_5_22 | SE_25 | SE_25_5 | GGTCAGAGATAATAAGATCA | 164.73407993662144 | 232.85844891969782 | 209.35569487852334 | 166.56746866449926 | 244.99733244049295 | 299.7432980144694 | 213.0798013274002 | 166.38221433788857 | 406.69725724392777 | 250.77838920045107 | 196.02208910455414 | 148.56716962917753 |
| SE_25_5_24 | SE_25 | SE_25_5 | TGAATGGCTCAGATGTCACG | 2383.225274872569 | 2225.647063279687 | 2182.4343664223425 | 2469.883513197028 | 2111.643674844249 | 2723.678985426277 | 2307.2883537674043 | 2149.7338375247646 | 2400.224205961434 | 2033.2003691222037 | 2108.1177965974807 | 2590.640020408783 |
| SE_25_5_33 | SE_25 | SE_25_5 | TAGTCCTGCTGCATTGCGGG | 1264.767574250245 | 1324.1950421399124 | 1385.5001882762654 | 1219.326054723717 | 1418.0148635192168 | 1342.7806702960336 | 1217.1376530368163 | 1372.6532682875807 | 1308.8902995143003 | 1075.2852436196667 | 1169.0898248391372 | 743.9965291586155 |
| SE_250_1_10 | SE_250 | SE_250_1 | GACCCATTGGCAGAGCTGAG | 172.32051782843953 | 265.8384009126164 | 288.35784388928687 | 214.71590077845607 | 284.239329411481 | 164.59892087499765 | 134.52007659558095 | 162.60080037566382 | 196.24474639936253 | 86.75182643440023 | 50.47275348201095 | 52.23064557275772 |
| SE_250_1_22 | SE_250 | SE_250_1 | GAAGAGTTCAGCTTTGAAGG | 382.57322511596954 | 448.72722560061936 | 465.12515230087024 | 415.11740817168175 | 381.81402458258646 | 400.2352707592048 | 358.3614840506276 | 190.96140509234937 | 268.1715539032013 | 204.85095149635683 | 199.54344399864794 | 324.9906835638258 |
| SE_250_1_36 | SE_250 | SE_250_1 | GGAGCCTGTGACTTTATCAA | 115.96412206064798 | 187.88578711117248 | 191.58021135110155 | 109.30991312357764 | 124.08955798934059 | 90.96256153658292 | 73.73782169957492 | 73.73757226338243 | 198.02071651001286 | 37.93083611309846 | 55.167893340802664 | 56.873369623669525 |
| SE_250_1_43 | SE_250 | SE_250_1 | CAGAGGCACTGACTCTCCCG | 4724.183252919295 | 4862.044437743905 | 4896.158184942069 | 3989.8118290105836 | 4377.073554034262 | 4002.3527075920483 | 3878.4828484037894 | 3193.404091098793 | 4631.731209136085 | 2555.8983485630856 | 2953.2429711799896 | 2377.0747140668404 |
| SE_250_1_9 | SE_250 | SE_250_1 | GCCTGTTTCTGATCTCCGAA | 490.9509092847994 | 616.625163019114 | 537.214613273192 | 571.2744269196498 | 618.3266009212441 | 481.6684210868353 | 366.9707689527448 | 730.758248199931 | 841.810043378261 | 406.0568682685792 | 515.2915995023908 | 752.1212962477113 |
| SE_250_2_10 | SE_250 | SE_250_2 | GGACGCCAGTCCCTCCTGCG | 256.8551114801268 | 307.81288526724 | 287.3703170266523 | 311.0127290063697 | 260.90625013143404 | 212.24597691776015 | 294.8680078975134 | 185.28928414901227 | 270.8355097366768 | 301.07986821350664 | 190.1531642810645 | 175.26283292192036 |
| SE_250_2_14 | SE_250 | SE_250_2 | TCAGTCTTGACAGCTGCCGG | 575.4855029364867 | 600.6348833264716 | 650.7802024761645 | 694.8987334284578 | 627.8719515358088 | 291.9465070256537 | 683.3619891055147 | 720.3593598038129 | 720.1560603162127 | 796.0755837509668 | 610.3681816429231 | 453.8262759766282 |
| SE_250_2_31 | SE_250 | SE_250_2 | AGAGATCCTTGGGTCCTAAA | 99.7074694353235 | 171.89550735703014 | 168.86709351050703 | 130.13084895664005 | 111.36242383658771 | 158.5347501059188 | 86.0928490211718 | 171.10898179066947 | 129.64585056247478 | 68.5266528137279 | 153.7658303754287 | 25.534982280014887 |
| SE_250_2_36 | SE_250 | SE_250_2 | GGTCTTGTGGGGAATCAGCC | 254.68755779675024 | 308.8122777518739 | 269.5948334992305 | 245.94730452804967 | 311.81478674244556 | 277.2192351578908 | 233.5268529699285 | 383.8135171658111 | 177.59705556503397 | 236.19825012391323 | 453.0809963734006 | 258.8318658383327 |
| SE_251_1_10 | SE_251 | SE_251_1 | ACCTGCTCCCTCCAAGCCAG | 1998.4844960732232 | 2072.740013130701 | 2061.956089180928 | 1942.8535749226357 | 2002.40244003312 | 1930.1389247868146 | 1951.0791909423058 | 1773.4831482834031 | 1651.6526167548159 | 2072.566744142856 | 1858.1015991168217 | 1922.0877570774842 |
| SE_251_1_16 | SE_251 | SE_251_1 | ACAACCCCACACCTAGGGCA | 2995.5591904264584 | 3044.149508194848 | 2939.8674700630377 | 2973.489898659225 | 2825.4237819111395 | 2983.5720183867998 | 2915.31909997943 | 2802.9730994990887 | 2934.7913432121863 | 3324.2716684106304 | 3096.4447368731367 | 2685.815863452475 |
| SE_251_1_19 | SE_251 | SE_251_1 | ATTCCAGAAGAATAAGAACA | 7.586437891818092 | 19.987849692677923 | 12.837849214249072 | 10.410467916531204 | 22.272484767317543 | 7.796790988815679 | 9.685445514881827 | 36.868786131691216 | 46.17523444690883 | 199.0188959377417 | 4.695139858791716 | 0.0 |
| SE_251_1_34 | SE_251 | SE_251_1 | CCGACTTTCGGGCTACAGAG | 47.686181034285156 | 70.95686640900662 | 73.07698783495626 | 71.57196692615203 | 41.363185996446866 | 4.331550549342044 | 69.95043982970209 | 49.15838150892162 | 53.27911666951019 | 41.553395855132884 | 34.03976397623994 | 55.71268861094157 |
| SE_251_1_8 | SE_251 | SE_251_1 | CTTTTCTCAGCCAGGCAGGA | 8.670214733506391 | 23.98541963121351 | 27.650752153767232 | 29.93009526002721 | 25.454268305505764 | 6.064170769078861 | 30.132497157741013 | 1.89070698111237 | 29.303514168230603 | 82.37778476543888 | 2.347569929395858 | 16.249534178191293 |
| SE_251_2_14 | SE_251 | SE_251_2 | GGTCAGGGACTGCTCACACA | 939.6345217437552 | 1023.3779042651097 | 988.5143894971786 | 893.9989323321171 | 996.9588419656424 | 868.0427300881455 | 843.7099204074837 | 947.2441975372974 | 1013.1912019985187 | 916.3617296474013 | 1353.3740642967123 | 1251.2141317207295 |
| SE_251_2_15 | SE_251 | SE_251_2 | GCTGGCTCGGGGCAGCCCCA | 2.167553683376598 | 6.9957473924372735 | 3.950107450538176 | 0.0 | 7.4241615891058474 | 17.326202197368175 | 5.380803063823238 | 0.945353490556185 | 13.319779167377547 | 3.6450347241344634 | 0.0 | 8.124767089095647 |
| SE_251_2_16 | SE_251 | SE_251_2 | GTGTCCTCAGGGCTGAGCAG | 179.90695572025763 | 243.85176625067066 | 237.00644703229057 | 226.4276771845537 | 195.14939034221084 | 325.73260131052166 | 186.175786008284 | 76.57363273505099 | 166.94123223113192 | 199.74790288256858 | 89.2076573170426 | 247.22505571105322 |
| SE_251_2_17 | SE_251 | SE_251_2 | TCCCAGGGCACAGATCCATG | 1221.4165005827128 | 1231.25154106896 | 1209.7204067273165 | 1354.6621376386229 | 1072.2610523694302 | 1339.3154298565598 | 1283.8596110282244 | 1378.3253892309178 | 1181.0204195074757 | 1298.3613687366958 | 1584.6097023422042 | 912.2952760041683 |
| SE_251_2_8 | SE_251 | SE_251_2 | TTTGAGCCAGACAGCCTGCG | 916.8752080683008 | 918.4416933785506 | 1003.3272924366968 | 920.0251021234451 | 1092.412348111289 | 989.3261454697227 | 848.0145628585423 | 1113.626411875186 | 1310.6662700699505 | 1033.7318477645338 | 1051.7113283693443 | 1122.3785393079272 |
| SE_251_3_11 | SE_251 | SE_251_3 | AAGGGAAGTTAAAGACAAAG | 1121.7090311473894 | 1412.1415807876954 | 1419.0761016058398 | 1293.500638629002 | 1450.8932934138284 | 1917.1442731387885 | 1153.6441768837021 | 1505.0027569654465 | 1533.5505748040682 | 961.5601602266714 | 1429.6700870020775 | 1587.811625411835 |
| SE_251_3_23 | SE_251 | SE_251_3 | CTGTGTCCCCTGAAGCCAGG | 307.7926230394769 | 323.80316502138237 | 264.6571991860578 | 279.7813252567761 | 340.4508385861396 | 325.73260131052166 | 315.3150595400417 | 194.7428190545741 | 171.38115862025776 | 303.2668890479873 | 159.63475519891836 | 95.17584304369186 |
| SE_251_3_25 | SE_251 | SE_251_3 | ACCCCTTCTCCAGGGATGCG | 35.76463577571386 | 69.95747392437274 | 63.20171920861082 | 65.06542447832003 | 57.27210368738796 | 21.65775274671022 | 83.94052779564251 | 70.90151179171387 | 8.879852778251697 | 48.11445835574915 | 46.95139858791716 | 10.446129114551544 |
| SE_251_3_39 | SE_251 | SE_251_3 | ATAGAAGTCCCAGGAAGACA | 280.6982019972694 | 326.80134247528406 | 318.9711766309577 | 359.1611431203265 | 299.0876525896927 | 151.60426922697152 | 331.4574687315114 | 297.78634952519826 | 178.48504084285912 | 702.0336878682976 | 471.86155580856746 | 141.60308355280984 |
| SE_251_3_9 | SE_251 | SE_251_3 | CTGTATTTCATTCTCCCAGA | 9.75399157519469 | 31.980559508284678 | 34.56344019220904 | 29.93009526002721 | 20.15129574185873 | 2.598930329605226 | 18.29473041699005 | 0.0 | 12.431793895523377 | 17.496166675845423 | 0.0 | 29.017025318198737 |
| SE_252_1_13 | SE_252 | SE_252_1 | GTCTGGGCACCAATTTCCCT | 972.1478269944041 | 867.4726766622219 | 900.6244987227042 | 1013.719313372226 | 1175.1387201041828 | 1244.021317771035 | 1302.1543414452235 | 1036.1074256495788 | 945.7043208838058 | 1255.3499589919093 | 1259.4712671208779 | 1606.3825216154821 |
| SE_252_1_14 | SE_252 | SE_252_1 | GCTTTACAGCTTGGTCTGGA | 262.27399568856833 | 373.7727892530772 | 332.79655270784133 | 331.8336648394321 | 281.0575458732928 | 596.8876656993336 | 221.6890862295174 | 236.33837263904624 | 384.49762529829854 | 98.4159375516305 | 211.28129364562722 | 334.2761316656494 |
| SE_252_1_22 | SE_252 | SE_252_1 | GGAAATTCGTCTGTGTAACT | 84.53459365168732 | 117.92831318679976 | 126.40343841722164 | 93.69421124878083 | 104.99885676021127 | 115.21924461249836 | 45.19874573611519 | 152.2019119795458 | 158.06137945288023 | 91.85487504818848 | 77.46980767006332 | 81.24767089095646 |
| SE_252_1_5 | SE_252 | SE_252_1 | TGAGCGGCCCAGTACCTACA | 8629.031213522236 | 7766.278998090007 | 8219.18607770731 | 8302.348163433635 | 8318.242763336737 | 7365.368554101211 | 8596.370974764004 | 8580.028280287936 | 8181.008364603289 | 9100.922699218929 | 8752.914481752458 | 8606.449709377745 |
| SE_252_1_6 | SE_252 | SE_252_1 | AGACAGGCCTTGTAGGTACT | 1105.452378522065 | 1129.3135076363026 | 1175.1569665351074 | 1147.7540877975653 | 1160.290396925971 | 944.2780197565655 | 1096.6076440717557 | 1294.1889285714174 | 829.3782494887085 | 1105.1745283559762 | 1103.3578668160533 | 1481.0289722408634 |
| SE_252_2_13 | SE_252 | SE_252_2 | CAAAGGATGCTTTTAGGGCG | 592.8259324034995 | 799.5139877071169 | 828.5350377503825 | 698.802658897157 | 774.233994292467 | 618.5454184460439 | 446.6066542973287 | 772.3538017844031 | 746.7956186509678 | 548.2132225098233 | 478.90426559675507 | 951.7584304369185 |
| SE_252_2_17 | SE_252 | SE_252_2 | GCGGGGATAACAAGAGTTGG | 4997.295017024746 | 5210.832414881134 | 5350.42054175396 | 5198.72741581777 | 5032.520962901035 | 4963.090619436113 | 5396.945473014707 | 5382.842775226917 | 5255.98485944718 | 5446.410884801715 | 6164.718634593523 | 6033.2199041598815 |
| SE_252_2_19 | SE_252 | SE_252_2 | GACAGAGTCTCACTCCGACG | 1436.004315236996 | 1410.1427958184274 | 1398.3380374905144 | 1244.0509160254787 | 1217.5625006613359 | 920.8876467901184 | 1329.0583567643396 | 1470.0246778148676 | 1831.0256428755001 | 954.2700907784025 | 1540.005873683683 | 1207.1082532370674 |
| SE_252_2_20 | SE_252 | SE_252_2 | AGACAGAGTCTCACTCCGAC | 312.12773040623006 | 358.78190198356873 | 333.7840795704759 | 231.63291114281927 | 303.33003064061035 | 265.0908936197331 | 300.24881096133663 | 432.02654518417654 | 315.2347736279353 | 744.3160906682574 | 339.2238547977015 | 280.88480508016374 |
| SE_252_2_5 | SE_252 | SE_252_2 | ACTCAAATGTTAAAGCTACT | 269.8604335803864 | 351.78615459113144 | 355.50967054843585 | 308.4101120272369 | 342.5720276115984 | 394.171099990126 | 219.53676500398808 | 335.60048914744567 | 385.38561057612367 | 282.85469459283433 | 266.4491869864299 | 359.8111139456643 |
| SE_252_3_12 | SE_252 | SE_252_3 | GCTTTCACTCACCCAGTACC | 588.4908250367463 | 550.6652590332768 | 525.3642909215774 | 592.0953627527122 | 574.8422258993385 | 528.4491670197293 | 583.2790521184389 | 610.6983548992955 | 411.1371836330536 | 554.7742850132653 | 647.9293005132569 | 596.5900405421166 |
| SE_252_3_13 | SE_252 | SE_252_3 | TCTCTTGTAGCTGTCACCAC | 966.7289427859627 | 1002.3906620877979 | 1005.3023461619658 | 921.3264106130115 | 996.9588419656424 | 797.005301078936 | 799.587335284133 | 1471.9153847959801 | 1057.5904658977773 | 548.2132225098233 | 1016.4977794284065 | 1355.6754228662448 |
| SE_252_3_15 | SE_252 | SE_252_3 | GTGACAGCTACAAGAGAAGG | 1341.7157300101142 | 1678.9793741849455 | 1547.4545937483306 | 1467.8759762308996 | 1391.5000007009817 | 1244.8876278809034 | 1234.3562228410506 | 1310.2599379108724 | 1478.4954875789076 | 1152.5599797713173 | 1147.9616954745745 | 1225.6791494407146 |
| SE_252_3_24 | SE_252 | SE_252_3 | TCACTAACCACCTCAGCCCA | 4809.801623412671 | 4570.221832230807 | 4459.671311657601 | 4653.479158689448 | 4939.188645780847 | 3834.288546277577 | 4935.2725701386735 | 4591.581903631391 | 5065.068024714768 | 5998.9981489804995 | 5289.075050928868 | 4758.792152184593 |
| SE_252_3_5 | SE_252 | SE_252_3 | AGACCCTCTTGGCATCTGGT | 280.6982019972694 | 280.82928818124446 | 235.0313933070215 | 273.2747828089441 | 327.7237044333867 | 227.8395588593915 | 283.0302411571023 | 308.1852379213163 | 144.74160028550267 | 358.6714168548312 | 591.5876222077562 | 292.49161520744326 |
| SE_253_1_14 | SE_253 | SE_253_1 | TGACCACTGTACTTCAGCCT | 1991.9818350230935 | 1747.9374556246844 | 1727.1844827478176 | 1845.2554382051558 | 2009.8266016222258 | 1695.3688850124759 | 1983.3640093252452 | 1680.838506208897 | 2037.9262126087647 | 1428.1246049158826 | 1977.8276655160105 | 1862.893025428359 |
| SE_253_1_17 | SE_253 | SE_253_1 | CTTCTTGCCTCATAAAAACA | 22.759313675454276 | 40.97509186998975 | 50.36389699436174 | 40.34056317655841 | 96.51410058367601 | 44.1818156032885 | 33.36097899570407 | 57.66656292392729 | 48.83919028038434 | 145.07238202055163 | 76.29602270536539 | 15.088853165463343 |
| SE_253_1_18 | SE_253 | SE_253_1 | CTTGCCTCATAAAAACAGGG | 260.10644200519175 | 336.795267321623 | 313.04601545515044 | 279.7813252567761 | 349.99618920070424 | 220.04276790657582 | 294.8680078975134 | 371.5239217885807 | 346.3142583518162 | 292.33178487558393 | 383.8276834562228 | 113.74673924733905 |
| SE_253_1_2 | SE_253 | SE_253_1 | AATGCAAAACAGTAATCACA | 658.9363197464858 | 825.4981923075983 | 809.7720273603261 | 676.6804145745282 | 813.475991263455 | 1098.4812193131422 | 690.8951133949037 | 739.2664296149367 | 876.4414692134426 | 678.7054656338371 | 922.5949822525722 | 1349.8720178026051 |
| SE_253_1_8 | SE_253 | SE_253_1 | GTCATTGTGGAATTAGCCAA | 480.1131408679164 | 636.6130127117918 | 594.4911713059955 | 495.7985345247986 | 654.3868143554044 | 615.0801780065782 | 473.5106969164494 | 647.5671410309867 | 412.0251689108788 | 578.8315141925527 | 750.0485924419767 | 733.5504000440641 |
| SE_254_1_24 | SE_254 | SE_254_1 | GCTGTGAGATTCTAGAAGGT | 428.0918524668781 | 629.6172653193546 | 598.4412787565337 | 430.7331004647854 | 659.689786917691 | 712.9732204217004 | 390.64630243356703 | 508.6001779192275 | 447.54458002388554 | 512.4918822133055 | 563.4167830550059 | 625.6070658603647 |
| SE_254_1_3 | SE_254 | SE_254_1 | CTCCTGATAAGCATCCAGCA | 34.680858934025565 | 50.9690167163287 | 46.413762543822357 | 29.93009526002721 | 54.09032014919974 | 64.10694813026224 | 45.19874573611519 | 68.06545132004533 | 70.15083694818841 | 43.74041668961356 | 32.86597901154201 | 39.46315443275028 |
| SE_254_1_31 | SE_254 | SE_254_1 | GTGCCATGTGGGGTAGGCTG | 1160.724997448168 | 1242.244858399933 | 1174.169439672473 | 1133.4396944123348 | 1222.865473177006 | 920.0213366802501 | 1202.0714044581111 | 1203.4349934780234 | 1233.4115508991608 | 991.449444964574 | 1017.6715643931045 | 1116.5751342442873 |
| SE_254_1_5 | SE_254 | SE_254_1 | GCACATTCCTCTCTTAGAGA | 429.1756293085664 | 456.7223654776906 | 448.337195636083 | 442.44488645257616 | 459.2374240118331 | 546.6416793269659 | 357.285323437983 | 392.3216985808176 | 461.7523444690883 | 223.8051320618506 | 460.1237061615882 | 448.0228709129885 |
| SE_254_1_9 | SE_254 | SE_254_1 | AGAGGCCACCCTAAATCCCT | 2706.1907736956823 | 3111.108804665319 | 3030.7199414254155 | 2681.9967969963513 | 3155.268675369985 | 2830.2351289400913 | 2884.1104422092553 | 3107.37692345818 | 3036.9096501620807 | 3643.5767102448094 | 2819.4314852044254 | 3580.700924265724 |
| SE_254_2_1 | SE_254 | SE_254_2 | ACAGTTCTTCAGTCAGTAGA | 259.02266516350346 | 342.79162222942637 | 330.8214989825723 | 359.1611431203265 | 314.9965702806338 | 190.58822417104992 | 350.82835976127507 | 255.24544245016995 | 221.99631945629244 | 213.59903483427954 | 382.65389849152484 | 125.35354937461854 |
| SE_254_2_16 | SE_254 | SE_254_2 | ACATAGGATGAGGAAGATGT | 2851.4168704819144 | 2509.474528915713 | 2589.2954338277746 | 2601.3156706432346 | 2468.0034311213294 | 2731.4757764150927 | 2610.7656465670348 | 2495.7332150683283 | 2115.1809317795546 | 2202.3299803220425 | 2861.687743933551 | 2803.044645737998 |
| SE_254_2_25 | SE_254 | SE_254_2 | AGGTCTGAAGGGGAAAAGGG | 2441.7492243237375 | 3535.8506106347245 | 3393.142300012293 | 2597.411745174535 | 2830.7267544747865 | 2607.59340370391 | 2165.2351528824706 | 2680.0771457267847 | 3261.5699254518486 | 2378.020654025324 | 2263.057411937607 | 2413.055825461407 |
| SE_254_2_26 | SE_254 | SE_254_2 | CCAAGGTCTGAAGGGGAAAA | 85.61837049337561 | 97.94046349412183 | 92.82752508764715 | 59.86019052005442 | 72.12042686559965 | 63.24063802039384 | 78.55972473181926 | 38.75949311280358 | 100.34233639424419 | 34.26332640686395 | 57.515463270198524 | 124.19286836189059 |
| SE_254_2_9 | SE_254 | SE_254_2 | AAGAAAGTCTTGACTTCAAG | 100.7912462770118 | 112.93135076363026 | 117.51569665351074 | 127.52823197750725 | 84.84756101835254 | 45.048125713157255 | 119.45382801687587 | 72.79221877282625 | 213.11646667804075 | 213.59903483427954 | 273.4918967746175 | 39.46315443275028 |
| SE_255_1_15 | SE_255 | SE_255_1 | GCTCTCATCATGGAAGCCCA | 66.11038734298623 | 71.95625889364052 | 110.60300861506893 | 66.36673296788642 | 108.18064029839948 | 75.36897955855156 | 95.77829453605362 | 58.61191641448347 | 12.431793889552377 | 147.2594028550323 | 32.86597901154201 | 5.803405063639747 |
| SE_255_1_2 | SE_255 | SE_255_1 | AAATGCCTACGTGACCACAG | 404.24876194973547 | 475.71082268573457 | 409.8236479933358 | 425.52787608821296 | 369.08689042893355 | 185.39036351183947 | 424.00728142927113 | 258.08150292183853 | 412.9131541870397 | 156.73649313778193 | 210.1075068609293 | 163.65602279464088 |
| SE_255_1_23 | SE_255 | SE_255_1 | TGTCCTCTTCAGCTCACTAA | 109.46146101051819 | 178.8912547494674 | 165.9045129226034 | 117.11776406097604 | 219.5430641349872 | 72.77004922894633 | 44.12258512335055 | 153.14726547010198 | 201.57265806631355 | 75.81672226199683 | 140.85419576375148 | 251.86777976196504 |
| SE_255_1_26 | SE_255 | SE_255_1 | GATCTCCTCTGGGAATGAGG | 24.926867358830876 | 52.9678016855965 | 38.51354764274722 | 11.711776406097604 | 30.757240869152795 | 15.5938519776313558 | 2.152321225529295 | 28.36060471668555 | 38.1833669464823 | 0.0 | 1.173784964697929 | 31.338387343654635 |
| SE_255_1_5 | SE_255 | SE_255_1 | TCCGAACAGGCAGAGTTAGA | 1044.7608753875202 | 1116.321405336062 | 1082.3294414474603 | 1181.5881085262915 | 1234.5320128170295 | 566.5668118539393 | 1225.7469379389336 | 1133.478835176866 | 1168.5886256179235 | 443.23622245475076 | 943.7231116171349 | 745.1572101713435 |
| SE_256_1_1 | SE_256 | SE_256_1 | GGAAGATACTGATCAGAGGG | 547.307305052591 | 830.4951547307677 | 775.2085871681171 | 664.9686381684306 | 866.5057168999253 | 609.0160072374913 | 564.98432170144 | 859.3263229155722 | 779.651073930499 | 620.3849100476856 | 996.5434350285417 | 479.3612582566431 |
| SE_256_1_11 | SE_256 | SE_256_1 | TGACAGGACCCAATACCAAA | 2240.166731769714 | 2803.2959193980787 | 2715.698872244996 | 2587.001277258004 | 2863.605184369398 | 2923.7966208058797 | 2590.3185949245067 | 3119.6665188354104 | 3070.653090719437 | 2162.2345983565638 | 2731.397612852081 | 2999.1997368890216 |
| SE_256_1_12 | SE_256 | SE_256_1 | ATATCATTCTTGGCAATCCA | 39.01596630077876 | 72.95565137827442 | 79.98967587339807 | 52.05233958265602 | 89.08993906927017 | 140.34223779868222 | 66.72195799140815 | 51.04908849003399 | 45.28724916908366 | 227.45016678599052 | 56.341678305500594 | 60.35541266185337 |
| SE_256_1_20 | SE_256 | SE_256_1 | GCTGAACCCAACAATGAAAC | 3.2513305250648967 | 4.996962423169481 | 6.912688038441808 | 1.3013084895664004 | 2.1211890254588135 | 0.8663101098684087 | 3.228481838293942 | 0.0 | 0.0 | 18.225173620672315 | 0.0 | 0.0 |
| SE_256_1_5 | SE_256 | SE_256_1 | ATATAAGAAAACAAGCCTGG | 24.926867358830876 | 29.981774539016886 | 45.42623568118903 | 16.917010364363207 | 53.02972563647034 | 2.598930329605226 | 9.685445514881827 | 32.14201867891029 | 21.311646667804077 | 60.50757642063209 | 44.6038286585213 | 0.0 |
| SE_257_1_3 | SE_257 | SE_257_1 | AGGTGGCTTTGAACATGCAA | 311.04395356454177 | 333.7970898677213 | 337.73418702101407 | 313.61534598550253 | 321.36013735701020 | 547.5079894368344 | 334.68595056980536 | 328.03766122299962 | 509.7035494716475 | 215.05704872393332 | 226.5404981867003 | 199.6371341892073 |
| SE_257_1_4 | SE_257 | SE_257_1 | GGTGGCTTTGAACATGCAAG | 204.8338230790885 | 197.87971195751143 | 269.5948334992305 | 157.45832723753446 | 225.90663121136365 | 242.56683076315446 | 166.80489497852037 | 256.19079594072616 | 154.50943834157954 | 204.12194455152994 | 140.85419576375148 | 228.65415950740604 |
| SE_257_1_47 | SE_257 | SE_257_1 | GGAGAGCCGAGCTCCGAGAA | 544.055974527526 | 749.5443634754222 | 811.7470810855951 | 775.5798597815747 | 754.0826985506081 | 751.9571753657788 | 689.818952782139 | 755.3374389543918 | 1016.7431431098194 | 775.6633892958138 | 1004.7599297814272 | 725.4256329549684 |
| SE_257_1_49 | SE_257 | SE_257_1 | GAGCTCGGCTCTCCCAGTGT | 290.4521935724641 | 357.7825094989348 | 387.11053015274126 | 343.54544124552973 | 449.6207339726285 | 269.4224441690751 | 248.59310154863357 | 332.76442867577714 | 396.04143390102457 | 454.90033571981504 | 362.69955409166005 | 502.5748785112021 |
| SE_257_1_7 | SE_257 | SE_257_1 | ACATGCAGGGGGTTGTAGGT | 2240.166731769714 | 2257.6276227879716 | 2334.5135032680623 | 2194.006113408951 | 2146.6432937643194 | 2348.566707853256 | 2124.341049597414 | 2195.1108050714615 | 2286.5620903998124 | 1769.2998550948685 | 2415.649457348338 | 2710.190164719762 |
| SE_257_2_0 | SE_257 | SE_257_2 | GTTGCCAAAGCTTAGCCACA | 649.182328171291 | 686.5826369434867 | 691.2688038441809 | 671.4751806162326 | 658.6291924049616 | 584.7593241611759 | 722.1037711650785 | 600.2994665031774 | 749.4595744844434 | 470.2094794133458 | 627.9749561133921 | 937.8302528841832 |
| SE_257_2_1 | SE_257 | SE_257_2 | CATTAAGCTTCTGCTCCAGT | 1238.7569300497257 | 1125.3159376977671 | 1135.6558920297257 | 1142.5488538392997 | 995.898247452913 | 999.7218667881436 | 1208.5283681346991 | 1222.3420632891473 | 1121.5254058931894 | 1027.1707852610919 | 762.9602270536539 | 782.2990025786379 |
| SE_257_2_2 | SE_257 | SE_257_2 | ATAGCCGATCTGACAGGAGG | 795.4922017992114 | 872.4696390853913 | 811.7470810855951 | 793.7981786355043 | 749.8403204996906 | 639.3368610828857 | 744.7031440333136 | 700.5069365021133 | 824.9383230995827 | 897.4075490810949 | 1028.2356290753858 | 683.6411164967623 |
| SE_257_2_25 | SE_257 | SE_257_2 | GCTGCAGTAAGAATCACATG | 859.4350354588211 | 227.86148649652833 | 428.5866583833921 | 694.8987334284578 | 191.9676068040226 | 2080.8768839039176 | 1322.6013930877518 | 437.69866612751366 | 1618.7971614752846 | 80.91977087578509 | 332.1811450095139 | 517.6637316766654 |
| SE_257_2_6 | SE_257 | SE_257_2 | TTAGGAACTGGATCAGCAGG | 921.2103154350541 | 1015.3827643880385 | 998.389658123524 | 940.8460379565075 | 1001.20122001656 | 1213.7004639256406 | 888.9086661435988 | 934.954602160067 | 1017.6311283876446 | 843.4610351647148 | 860.384379123582 | 1715.4865368119092 |
| SE_257_3_14 | SE_257 | SE_257_3 | GCTAAATCTGGGGATCGGGG | 6017.129025053436 | 8849.620451433151 | 8955.88111723268 | 6215.0493461765 | 8417.938647533301 | 9538.0743065118 | 5477.657518972055 | 7574.172166336154 | 8823.0217204070886 | 6954.7262536488556 | 6743.394622189602 | 8404.491213163083 |
| SE_257_3_2 | SE_257 | SE_257_3 | GTTCAATGTATGGTAGTCGG | 161.48274941155654 | 208.8730292884843 | 159.97935174679614 | 134.03477442533924 | 155.90739337122278 | 239.9679004335492 | 121.60614924240517 | 136.13090264009065 | 220.2203489006421 | 295.2478126548915 | 90.38144228174053 | 188.03032406192781 |
| SE_257_3_24 | SE_257 | SE_257_3 | GACCCGTTCATCTAGTGCAT | 487.6995787597345 | 347.7885846525959 | 382.1728958395685 | 420.3226421299473 | 363.7839178661865 | 424.4919538355203 | 541.3087882206177 | 341.2726100907828 | 285.0432741818795 | 240.57229179287458 | 446.03828658521303 | 305.2591063474507 |

| | | | | | | | | | | | | | | | |
|---|---|---|---|---|---|---|---|---|---|---|---|---|---|---|---|
| SE_257_3_3 | SE_257 | SE_257_3 | AGAGCTCTATGACCCCAAAA | 85.61837049337561 | 114.93013573289807 | 127.39096527985618 | 144.44524234187045 | 114.54420737477592 | 116.08555472236677 | 115.14918556581728 | 49.15838150892162 | 147.40555611897818 | 132.67926395849446 | 72.7746678112716 | 167.1380658328247 |
| SE_257_3_31 | SE_257 | SE_257_3 | GAAAGGTTACAGGAGTGAAA | 356.5625809154503 | 494.6992798937786 | 469.0752597514084 | 387.78992989078733 | 379.69283555712764 | 210.51335669802333 | 470.28218777815096 | 536.9607826359131 | 618.9257386441434 | 554.7742850132653 | 491.81590020843225 | 379.5426911620395 |
| SE_257_4_10 | SE_257 | SE_257_4 | AGGGTTGGAGAGGTGAGCAG | 2102.5270728753 | 2337.579021558683 | 2316.73801974064 | 2296.8094840846966 | 2243.1573944226952 | 1889.4223496229995 | 2155.549707367589 | 2076.9416187519387 | 2252.8186498424557 | 2348.860376232248 | 1849.885104363936 | 1996.3713418920731 |
| SE_257_4_14 | SE_257 | SE_257_4 | GGAGAGAGTGACAAAAGGTA | 2165.3861296932214 | 2551.4490132703368 | 2478.6924252127055 | 2428.241641530903 | 2423.4584615866943 | 2423.0693773019393 | 2083.4469463123573 | 2447.520187049963 | 2413.5439851288115 | 2014.2461885567045 | 3020.1487141677712 | 2267.9706988704133 |
| SE_257_4_5 | SE_257 | SE_257_4 | GAGATGATACATCACATGGG | 745.6384670815496 | 754.5413258985916 | 864.086004805226 | 719.6235947302195 | 748.7797259869611 | 901.8288243730135 | 873.8424175648937 | 651.3485549932115 | 836.48213171131 | 1395.3192923986726 | 537.5935138316515 | 791.5844506804615 |
| SE_257_4_7 | SE_257 | SE_257_4 | TGTTCTAGTTCAAACCACCA | 61.77527997623304 | 93.94289355558624 | 72.08946097232172 | 78.07850937398402 | 28.63605184369398 | 171.52940175394494 | 34.43713960846872 | 89.80858160283758 | 100.34233639424419 | 77.27473615165061 | 16.432989505771005 | 55.71268861094157 |
| SE_257_4_9 | SE_257 | SE_257_4 | GTAGACTAGTCAAGAGACCG | 415.0865303666185 | 514.6871295864565 | 566.8404191522283 | 412.51479119254896 | 483.63109780460945 | 429.38808449309437 | 584.2284571637223 | 482.1760058590672 | 510.30486137882485 | 550.5051484433287 | 756.764020298623 | |
| SE_258_1_14 | SE_258 | SE_258_1 | CTGGCTTTCTCTTTGCGCAA | 110.54523785220648 | 121.92588312533533 | 160.96687860943067 | 106.70729614444484 | 82.72637199289373 | 111.75400417302473 | 89.32133085946575 | 93.58999556506231 | 102.11830694989453 | 101.33196533093808 | 76.29602270536539 | 126.51423038734649 |
| SE_258_1_27 | SE_258 | SE_258_1 | AATGGAACCAGAGCTTGGCA | 482.2806945512930 4 | 627.6184803500868 | 659.6679442398754 | 689.6934994701923 | 692.5682168123026 | 478.2031806473616 | 736.0938591310189 | 611.6437083898517 | 526.5752697503257 | 718.0718406544893 | 591.5876222077562 | 438.7374228111649 |
| SE_258_1_32 | SE_258 | SE_258_1 | ACTGTCCCAGACCTTTCTTA | 358.73013459882696 | 436.7345157850126 | 418.7113897570467 | 391.6938553594865 | 559.9939027211268 | 582.1603938315707 | 347.59987792298114 | 542.6329035792502 | 428.00890391173186 | 238.3852709583939 | 477.7304806320571 | 347.04362280565687 |
| SE_258_1_35 | SE_258 | SE_258_1 | GCCGTGGTAGAGACAGGGAG | 3097.4342135451584 | 3111.108804665319 | 3113.6727897886717 | 3255.873840895134 | 3383.2964956068076 | 3339.6254735427156 | 3602.9857315360396 | 3529.4930989348254 | 3298.865307120506 | 3264.4930989348254 | 3048.3195533205217 | 2922.594790048977 |
| SE_258_1_8 | SE_258 | SE_258_1 | AGAGGAGTTTCAGTTGGGCA | 1605.0735025403708 | 1939.8208126743925 | 1917.7771672362844 | 2059.971338983612 | 2162.5522114552605 | 2442.9945098289127 | 1859.6055388573109 | 2001.3133395074437 | 1863.8810981550314 | 2227.116216446157 | 1851.058889328634 | 2538.4093748360256 |
| SE_258_2_10 | SE_258 | SE_258_2 | TGCAGGACGTCCCCATTCCA | 4912.760423373059 | 4205.4435753394355 | 4400.4196998995285 | 4847.374123634841 | 4654.949316369366 | 4476.224337690068 | 4946.03417626632 | 3830.5723437336615 | 3991.493823824138 | 4627.736085761115 | 4393.4771228643485 | 3761.767162251284 |
| SE_258_2_15 | SE_258 | SE_258_2 | CTGGGCCTCCGAGGGCCGGG | 24.926867358830876 | 41.97448435462364 | 43.451181955919935 | 24.724861301761607 | 72.12042686559965 | 32.05347406513112 | 22.599372868057596 | 32.14201867891029 | 43.51127861343332 | 49.5724722482287 | 19.954344399864794 | 41.78451645820618 |
| SE_258_2_23 | SE_258 | SE_258_2 | GCAGCGGTGCCAATGCCCGC | 1377.4803657858279 | 1073.3475284968044 | 1129.7307308539184 | 1280.487553733338 | 1278.016387838935 | 1316.7913669999812 | 1416.227366398276 | 1141.9870165918715 | 1237.8514772882868 | 1164.2240908885476 | 800.5213459239876 | 1861.732344415631 |
| SE_258_2_5 | SE_258 | SE_258_2 | TTGACCCTAAGGACACAAGC | 327.30060618986624 | 290.8232130284638 | 353.53461682316674 | 314.9166544750689 | 278.93635684783396 | 251.22993186183854 | 244.288459097575 | 220.2673632995911 | 292.1471564044809 | 317.11802099969833 | 226.5404981867003 | 314.5445544492743 |
| SE_258_2_9 | SE_258 | SE_258_2 | GTTCCAGGTAGACCCCTCAG | 42.26729682584366 | 32.97995199291857 | 31.60085960430541 | 39.03925468699201 | 48.78734758555271 | 3.465240439473635 | 25.827854706351538 | 13.23494886778659 | 79.91867500426528 | 6.561062503442034 | 52.820323411406804 | 5.803405063639747 |
| SE_258_3_1 | SE_258 | SE_258_3 | GCTTGTGTGGTTTGACTGAA | 1519.455132046995 | 1555.0547060903425 | 1531.6541639461777 | 1499.1073799804933 | 1600.4371197086748 | 1168.6523382124833 | 1618.5455615980297 | 1770.6470878117345 | 1547.7583392492709 | 1616.9374036260479 | 1766.5463718703832 | 1258.1782177970972 |
| SE_258_3_11 | SE_258 | SE_258_3 | GCACTCACTGAGCATTTCCT | 1479.355388904528 | 1582.0383031754577 | 1550.4171743362342 | 1471.779901699599 | 1585.588796530463 | 1619.133595344056 | 1604.5554736208994 | 1716.761938850032 | 1793.730261206843 | 1448.5367993710356 | 1878.0559435166865 | 2136.813744432155 |
| SE_258_3_17 | SE_258 | SE_258_3 | GTATATACCACGCTTCTCTA | 28.17819788389577 | 47.97083926242702 | 52.338923719630834 | 40.34056317655841 | 55.15091466192915 | 34.65240439473635 | 19.370891029763655 | 26.46989773557318 | 96.7903952829435 | 8.748083337922711 | 18.780559435166865 | 12.767491140007444 |
| SE_258_3_2 | SE_258 | SE_258_3 | CATCATGGCTTGCCTGGCCA | 5405.878886341235 | 4693.147107840777 | 4636.438620069184 | 5270.299382743921 | 5160.8528989412935 | 4190.342001433493 | 5446.448861201881 | 5093.564607116725 | 5390.07063639878 | 4975.472398443542 | 5242.123652340951 | 5487.699828177745 |
| SE_258_3_9 | SE_258 | SE_258_3 | ACTCACTGAGCATTTCCTTG | 613.4176923955772 | 669.5929647047104 | 638.92988012455 | 707.9118183241219 | 617.2660064085147 | 751.0908652559103 | 652.1533313353764 | 588.9552246165033 | 754.7874861513943 | 948.4380352197874 | 487.12076034964053 | 617.4822987712691 |
| SE_259_1_11 | SE_259 | SE_259_1 | GAAGGGTCCGGATGGCCACA | 3328.278680824766 | 3252.023144998698 | 3282.5392913972246 | 3436.7557209448637 | 3296.3277455629964 | 3593.4543357341595 | 3682.6216168806236 | 3533.7313476990194 | 3061.7732379411855 | 4283.644807802822 | 3629.3431108459968 | 3281.2452229819132 |
| SE_259_1_13 | SE_259 | SE_259_1 | ATTTTGAGCAGGACCCCACT | 32.51330525064897 | 39.975699385355846 | 53.32645058226538 | 20.820935833062407 | 65.75685978922321 | 69.3048087894727 | 40.8941032850566 | 50.10373499947781 | 9.767838056076867 | 16.76715973101853 | 69.25331291771781 | 4.642724050911798 |
| SE_259_1_17 | SE_259 | SE_259_1 | CTTGGTTTCTCCCCTGTTAG | 113.79656837727138 | 149.90887269508443 | 146.1539756699125 | 106.70729614444484 | 156.9679878839522 | 153.33688944670834 | 123.75847046793446 | 139.91231660231537 | 141.189659174202 | 69.2556597585548 | 254.7113373394506 | 103.3006101327875 |
| SE_259_1_26 | SE_259 | SE_259_1 | AGTACTAACCTCCACAGCCA | 438.92962088376106 | 449.7266180852533 | 442.4120344602757 | 447.65012041084174 | 346.814405662516 | 229.5721791151283 | 451.98745736115194 | 370.57856829802455 | 269.9475244588516 | 288.6867501514495 | 207.75993875153344 | 372.57860508567177 |
| SE_259_1_42 | SE_259 | SE_259_1 | AGTCCCTCCCTGAAGAGCTG | 45.51862735090855 | 41.97448435462364 | 61.22666548334173 | 31.23140374959361 | 44.54496953463509 | 38.98395494407839 | 39.817942672291956 | 55.775855942814914 | 16.871720278678225 | 169.1296111998391 | 53.994108376104734 | 13.928172152735392 |
| SE_259_2_17 | SE_259 | SE_259_2 | AGACTCGGCTGAAGAAAAGT | 300.2061851476588 | 429.7387689325753 7 | 434.51181955268935 | 353.95590916206095 | 428.48011824268035 | 348.2566641671003 | 361.58996588892154 | 408.39277079202719 | 445.76860946823524 | 290.14476404110326 | 428.4315121147441 | 380.7033721747674 |
| SE_259_2_19 | SE_259 | SE_259_2 | TAGACTCCAGGGAATAGCCA | 590.6583787201229 | 686.5826369434867 | 593.5036444433609 | 517.9207788474274 | 618.3266009122441 | 509.39034460262434 | 594.0406582460854 | 630.5507782009754 | 585.1822980867869 | 841.2740143302341 | 718.3563983951326 | 630.2497899112766 |
| SE_259_2_39 | SE_259 | SE_259_2 | GCAAGTGCACTGAGGCCCTG | 4345.945135170078 | 4164.468483469445 | 4101.199060521261 | 4427.051481504895 | 4093.8948191355103 | 3952.973031329549 | 4738.335178002742 | 4653.029880517543 | 4157.547070777445 | 4320.824161988993 | 3662.2090898575384 | 3952.1188483386677 |
| SE_259_2_42 | SE_259 | SE_259_2 | AGCCTCGATCCTGAGCCGCA | 696.8685092055762 | 744.5474010522527 | 787.0589095197316 | 818.5230399372658 | 852.717988234443 | 782.2780292111731 | 810.3489414117795 | 1452.0629614943002 | 668.6529142023528 | 665.583340626953 | 1334.5935048615454 | 877.4748456223298 |
| SE_259_2_6 | SE_259 | SE_259_2 | GCGCAGGTAAGCCACTTGCC | 20174.505908027684 | 15355.665526399815 | 15645.388084719081 | 18487.68971126985 | 16352.246197261993 | 18366.640639320132 | 21363.940484603783 | 17931.465008869716 | 16115.15682197118 | 15776.439292998784 | 19672.63600833729 | 18144.926271976034 |
| SE_259_3_10 | SE_259 | SE_259_3 | GTATAGGCTGACTAATGCAC | 1249.5946984666086 | 1044.3651464424215 | 969.7513791071223 | 982.4879096226323 | 783.7793449070316 | 809.9999527269622 | 962.0875871159 49 | 745.88390404883 | 939.4884239390296 | 998.010507468016 | 645.5817305838609 | 1266.302984886193 |
| SE_259_3_13 | SE_259 | SE_259_3 | TGATTGAACCACACTCTAAG | 343.5572588151908 | 417.7460585769686 | 396.9857987790867 | 360.4624516098929 | 419.9954270408451 | 207.9144263684181 | 376.6562144676266 | 462.27785688197446 | 396.9294191878509 | 267.5455487514696 | 556.3740732668184 | 401.59563040387053 |
| SE_259_3_14 | SE_259 | SE_259_3 | AGGAGGGAAAGGCAGTTGAA | 3503.8505291782703 | 3273.01038717601 | 3369.441655309064 | 3432.851795476164 | 3256.0251540792788 | 3616.8447087006066 | 3700.9163472976225 | 3427.851756756727 | 3709.1145054757344 | 3648.679758858598 | 3626.995540916601 | 3021.2526761308523 |
| SE_259_3_3 | SE_259 | SE_259_3 | ACAGAGACCTCAAAATTTAG | 591.7421555165812 | 659.5990398583715 | 718.919555997948 | 748.2523815006803 | 696.8105948632202 | 587.3582544907811 | 775.9118018033108 | 597.463406031509 | 600.2780478098148 | 860.2281948957333 | 852.1678843706965 | 481.68262028209904 |
| SE_259_3_4 | SE_259 | SE_259_3 | AAGTTCCCAGGCCAACCAGT | 454.10249666739725 | 535.6743717637684 | 489.81332386673387 | 448.95142890040813 | 539.842606979268 | 465.2085289993355 | 458.4444210377398 | 674.9823922571161 | 410.24919835522843 | 531.4460627788047 | 539.9410837610474 | 328.4727266020097 |
| SE_259_4_0 | SE_259 | SE_259_4 | CAGATTGGAGCTCTACACCT | 1303.7835405510236 | 1429.1312530264715 | 1407.2257792542252 | 1372.8804564925524 | 1568.6192843267925 | 1694.5025749026074 | 1443.1313817173923 | 1272.445798288625 | 1093.997862280609 | 1938.4294662947075 | 1157.351975192158 | 1267.4636658989207 |
| SE_259_4_21 | SE_259 | SE_259_4 | AGTCCAGCTCTGCCCACGGC | 289.36841673077583 | 390.7624614918534 | 450.3122493613521 | 324.0258139020337 | 413.6318599644686 | 448.7486369118357 | 209.85131948910626 | 356.3982659396817 | 399.5933750213264 | 349.1943265720816 | 450.73342644400475 | 974.9720506914775 |
| SE_259_4_5 | SE_259 | SE_259_4 | TCACAGACAGGAGATGTGAT | 1240.9244837331023 | 1275.2248103928516 | 1271.9345990732927 | 1292.1993301394357 | 1125.2907780059006 | 803.0694718480149 | 1210.6806893602284 | 1196.8175190441302 | 1704.931733424326 | 1331.8956881987328 | 1656.210585188778 | 994.7036279078527 |
| SE_259_4_6 | SE_259 | SE_259_4 | CAGATGCAGTCACAACTCCC | 11347.14353247649 | 9752.071865057558 | 9933.532711240878 | 10977.838417982153 | 9748.984761008707 | 10918.972624781423 | 11627.915420922016 | 10097.320632630612 | 8793.718206302656 | 11378.34039485814 | 10618.058790657466 | 9179.826129665353 |
| SE_259_4_8 | SE_259 | SE_259_4 | CAGCCCTTAAGAAAGGCTGG | 319.71416829804815 | 390.7624614918534 | 365.38493917478127 | 347.4493667142289 | 274.6939787969163 | 567.4331219638077 | 396.02710549739027 | 432.02654518417654 | 435.1127861343332 | 664.1253267372992 | 429.60529707944204 | 490.9680683839226 |
| SE_26_1_10 | SE_26 | SE_26_1 | TCTGGCTTGGACGGTTACCC | 1712.3674098675124 | 1549.0583511825391 | 1519.8038415945632 | 1603.2120591458054 | 1434.9843757228873 | 1369.6362837019542 | 1689.5721620404966 | 1465.2979103620867 | 1396.8008420189922 | 1221.815639529872 | 1205.477158744773 | 944.7943443605509 |
| SE_26_1_30 | SE_26 | SE_26_1 | GCATGGCAATGTGAAGAAGG | 8609.523230371846 | 8315.94486463865 | 8134.25876752074 | 8349.195269058026 | 7936.428738754151 | 6454.876628629513 | 7954.979249556274 | 7920.171543879718 | 7487.491862621831 | 6878.909531386559 | 6145.938075158357 | 6264.195425692743 |
| SE_26_1_5 | SE_26 | SE_26_1 | GTTGTTCAGACAGACCGTGG | 238.43090517142576 | 239.8541963121351 | 267.61977977396145 | 242.04337905995049 | 248.17911597868118 | 179.3261927427606 | 248.59310154863357 | 177.72645622456278 | 182.03698195441598 | 191.72882648947277 | 173.7201747752935 | 277.40276204197994 |
| SE_26_1_8 | SE_26 | SE_26_1 | AGCAGGTTATAAGAATGCTG | 281.78197883895774 | 424.7418059694059 | 398.9608525043558 | 350.0519836933617 | 368.02629591710416 | 393.30478988025754 | 279.80175931880837 | 355.45291244912556 | 504.37563780469645 | 368.1485071375808 | 491.81590020843225 | 345.88294179292893 |
| SE_26_1_9 | SE_26 | SE_26_1 | GGACAGACACGTGCAATAGG | 4368.704448845533 | 3839.6659259634293 | 3907.6437954448907 | 4308.632408954352 | 3939.048020277017 | 3763.2511172683676 | 4139.989877305598 | 4103.779502504399 | 4191.2905113348015 | 3561.1989254793707 | 4080.0765372900014 | 3162.855759683662 |
| SE_26_2_10 | SE_26 | SE_26_2 | GTGTCTGTGAGAGATTTCTG | 20.59175999207768 | 20.98724217731182 | 35.55096705484538 | 11.711776406097604 | 18.370810617639914 | 8.663101098684088 | 11.837766740411123 | 32.14201867891029 | 9.767838056076867 | 83.83579865509266 | 90.38144228174053 | 2.32136202545589 9 |
| SE_26_2_2 | SE_26 | SE_26_2 | ATTCTGATGCAGGTACCCGA | 11.921545258571289 | 35.97812944682026 | 22.713117840594514 | 20.820935833062407 | 40.30259148371746 | 2.598930329605226 | 22.599372868057596 | 84.13646065950047 | 14.207764445202717 | 8.748083337922711 | 221.8453583279086 | 8.124767089095647 |
| SE_26_2_24 | SE_26 | SE_26_2 | AATGGAGTTTGAATGCCTGT | 619.920353445707 | 668.5935722200766 | 546.1023550369028 | 642.8463938458018 | 566.3574697975032 | 447.0160166920989 | 684.4381497183158 | 481.1849266930982 | 812.5065292100304 | 422.8240279955977 | 599.8041169606417 | 557.1268861094158 |
| SE_26_2_6 | SE_26 | SE_26_2 | GGACAGTAGCAAAAGCCTGT | 1781.7291277355635 | 2023.7697813836398 | 2041.2180250656024 | 2035.2464776818504 | 1997.099467469473 | 1531.6362742473466 | 1810.102150670137 | 1835.8764786601114 | 1633.8929111983125 | 1812.3112648396552 | 2274.7952615845866 | 1780.4846735246745 |
| SE_26_2_9 | SE_26 | SE_26_2 | GATTCCTGAAATTCATTAAA | 0.0 | 0.9993924846338962 | 1.975053725269088 | 0.0 | 6.363567076376441 | 0.0 | 0.0 | 11.34424188667422 | 0.0 | 0.0 | 0.0 | 0.0 |
| SE_260_1_12 | SE_260 | SE_260_1 | TCCTAGACCCCTAAACCAGA | 109.46146101051819 | 116.92892070216585 | 130.3535458677598 | 106.70729614444484 | 113.48361286204653 | 84.03208065723565 | 128.06311291899306 | 71.84686528227006 | 153.62145306375436 | 134.86628479297514 | 72.7746678112716 | 42.945197470934126 |
| SE_260_1_14 | SE_260 | SE_260_1 | CTAGACCCCTAAACCAGAGG | 215.67159149597148 | 247.84933618920624 | 271.5698877244996 | 284.9865592150417 | 225.90663121136365 | 232.17110944473353 | 179.71882233169612 | 293.05958207241736 | 186.47690834328566 | 138.5113195171096 | 492.98968517313017 | 94.01516203096391 |
| SE_260_1_21 | SE_260 | SE_260_1 | GGTGGCCATTCGCAGAAAGG | 189.6609472954523 | 189.88457208044028 | 226.1436515433106 | 192.59365645582727 | 221.664253160446 | 215.71121735723378 | 192.63274968487 19 | 485.9116941458791 | 251.29983362452305 | 103.51988616541875 | 437.8217918323275 | 242.58233166014142 |
| SE_260_1_22 | SE_260 | SE_260_1 | TCTTTGGATGTAAGGATGGG | 187.49339361207572 | 206.8742443192165 | 221.20601723013786 | 222.52375171585447 | 244.99733244049295 | 134.27806702960336 | 172.1856980423436 | 189.070698111237 | 285.9312594597047 | 176.41968064810803 | 165.503680022408 | 78.92630886550056 |
| SE_260_1_7 | SE_260 | SE_260_1 | ATAGTAAGTAAGATGAGCCA | 549.4748587359676 | 484.70535504743964 | 517.4640760205011 | 521.8247043161264 | 617.2660064085147 | 554.4384703517816 | 642.4678858204945 | 510.4908849003399 | 504.37563780469645 | 438.1331738409625 | 562.242998090308 | 764.8887873877187 |
| SE_260_2_11 | SE_260 | SE_260_2 | AAATCGTCTCCAAAGCCCAA | 79.11570944324582 | 150.9082651797183 | 111.59053547770347 | 145.74655083143685 | 110.3018293238583 | 24.256683076315444 | 101.15909759987686 | 77.51898622560716 | 99.45435111641902 | 140.69834035159028 | 36.3873339056358 | 69.64086076367697 |
| SE_260_2_12 | SE_260 | SE_260_2 | ATATAGCTATCTCCATGCCA | 86.70214733506391 | 143.91251778728105 | 150.1040831204507 | 94.99551973834723 | 115.60480188750533 | 133.41175691973496 | 64.56963676587885 | 120.0598933006355 | 241.53199556844618 | 92.58388199301537 | 64.5581730583861 | 237.93960760922963 |
| SE_260_2_20 | SE_260 | SE_260_2 | TTTGGGGGTTCAGCTTCAGG | 102.95879996038839 | 143.91251778728105 | 142.20386821937433 | 117.11776406097604 | 170.75571654943448 | 220.09078014644423 | 107.61606127646475 | 130.45878169675353 | 156.28540889722987 | 147.2594028550323 | 223.01914329260651 | 157.85261773100112 |
| SE_260_2_31 | SE_260 | SE_260_2 | TCCCTGCTCCTGCAACCCCA | 742.3871365564847 | 709.5686640900662 | 677.4434277672972 | 769.0733173337427 | 783.7793449070316 | 557.0374006453868 | 881.3755418542463 | 933.0638951789546 | 661.5490319797515 | 909.8006671439621 | 850.9940994059986 | 359.8111139456643 |
| SE_260_2_9 | SE_260 | SE_260_2 | GCTTGGACTGCTAGGTTGGG | 29591.442885457313 | 30403.51816753239 | 30135.369740155747 | 30133.099384399567 | 29504.678749619365 | 30243.75224561002 | 30780.345846294447 | 29097.03508582882 | 30432.143456346395 | 28346.706042648893 | 28652.090988276446 | 28672.303057418536 |
| SE_261_1_12 | SE_261 | SE_261_1 | AGAATTAGAAAATATTCCAA | 0.0 | 9.993924846338962 | 7.900214901076352 | 2.602616979132801 | 0.0 | 0.0 | 2.152321225529295 | 0.8879852778251698 | 0.0 | 0.0 | 0.0 | 0.0 |
| SE_261_1_14 | SE_261 | SE_261_1 | TGTGGGAGTTAACCAGGCAA | 2763.630946305162 | 2976.1908192397427 | 2957.6429535904595 | 3176.4940230315833 | 2970.7252301550684 | 2946.320683662458 | 3199.425501749297 | 3474.17407779398 | 2992.510386270822 | 3558.282897700063 | 3941.5699114556455 | 3114.1071571490884 |
| SE_261_1_2 | SE_261 | SE_261_1 | GTGCCCTAAGACAGGAACAT | 209.16893004584168 | 217.86756165018937 | 216.26838291696515 | 193.89496494539367 | 195.14939034221084 | 80.566840217762 | 199.0897133614598 | 431.0811916936204 | 207.78855501108973 | 156.73649313778193 | 161.98232512831422 | 324.9906835638258 |
| SE_261_1_6 | SE_261 | SE_261_1 | GTAGGCACTATACACACAGG | 3063.837131452821 | 3250.02436002943 | 3127.497573963601 | 3071.088035376705 | 2992.997714922386 | 3201.8821660736385 | 2990.650342872955 | 3175.4423747782253 | 2682.603524309838 | 3037.771939093662 | 3461.4918608941925 | 2984.110883723558 |
| SE_261_1_7 | SE_261 | SE_261_1 | ACTCAATCATGGATAACTA | 348.97614302363223 | 397.7582088842907 | 407.8485942680667 | 348.7506752037953 | 360.6021345298673 | 276.3529250480 2237 | 333.60978995704073 | 332.7644286757714 | 334.770449740089 | 336.8012085100244 | 473.0353407732654 | 487.48602534573877 |
| SE_261_2_0 | SE_261 | SE_261_2 | GACAGTCTCAATTTTCTAAG | 143.05854310285545 | 235.8566263735995 | 241.94408134546327 | 171.77272062276487 | 215.30068608406958 | 193.18715450065514 | 112.99686434028798 | 150.31120499843342 | 341.87433196269035 | 189.5418056549921 | 65.73195802308402 | 254.18914178742094 |
| SE_261_2_1 | SE_261 | SE_261_2 | TCTGCAGTCTTCCTGTTGCC | 438.92962088376106 | 107.93438834046079 | 230.09375899384875 | 201.70281588279207 | 102.87667773475245 | 989.3261454697227 | 430.4642451058 59 | 182.45322367734371 | 968.7919381072602 | 29.160277793075707 | 104.46686185811568 | 336.59749369110534 |
| SE_261_2_2 | SE_261 | SE_261_2 | ATCTTCTCAAAACATCACAG | 23.843090517142578 | 56.96537162413208 | 64.18924607124536 | 50.75103109389165 | 46.6661585600939 | 34.65240439473635 | 20.4470516425283 | 62.393330376708214 | 45.28724916908366 | 53.94651391719005 | 56.341678305500594 | 18.57089620364719 |
| SE_261_2_6 | SE_261 | SE_261_2 | TGTTCATGTAGGAAAATCAA | 176.65562519519273 | 183.8882171726369 | 218.24343664223423 | 206.90804984105768 | 208.93711900769313 | 239.1015903236808 | 177.56650110616684 | 216.48594933736638 | 207.78855501108973 | 96.95792366197672 | 96.25036710523018 | 253.02846077469297 |
| SE_261_2_8 | SE_261 | SE_261_2 | GCTTTTCCTCACTTGACAGA | 41.18351998415536 | 71.95625889364052 | 94.80257881291622 | 54.65495656178882 | 64.69626527649382 | 112.62031428293813 | 45.19874575611519 | 146.52979103620868 | 83.47061611556596 | 32.076350572383276 | 232.40942301018995 | 111.42537722188314 |
| SE_261_3_1 | SE_261 | SE_261_3 | GCTGAGGAGCCCAGAACTAA | 54.188842084414944 | 46.97144677779312 | 37.526020780112674 | 49.44972260352321 | 54.09032014919974 | 47.64705604276248 | 35.513300221233365 | 73.73757226338243 | 68.37486639253807 | 33.53431946203706 | 151.41826044603283 | 107.94333418369929 |
| SE_261_3_12 | SE_261 | SE_261_3 | AGTTCAGTGGAGAGAAGTGA | 215.67159149597148 | 331.7983084982435 | 323.90881094413044 | 192.59365645582727 | 293.29878064424217 | 310.1390193328903 | 151.73866439981528 | 366.79715437357998 | 433.3368155786828 3 | 461.46139607542307 | 342.74520969179525 | 540.87735191312244 |
| SE_261_3_25 | SE_261 | SE_261_3 | AGCGGACTAAGAAGCTGAG | 2000.6520497565998 | 2122.7096373623954 | 2217.985333477186 | 2085.99750877494 | 2055.43216566959 | 2342.502537084177 | 2315.8976386695213 | 2313.279991390985 | 2159.580195670813 | 2402.077883204611 | 2481.381415371422 | 2495.4641773650915 |
| SE_261_3_5 | SE_261 | SE_261_3 | AAGAAAACTGATGCTCACAG | 384.7407787993461 | 472.7126452318979 | 491.7883775920029 | 441.14357796300976 | 519.6913112374093 | 389.8395494407839 | 427.2357632675650 6 | 650.4032015026553 | 451.9845064130114 5 | 340.4462423251589 | 424.9101572206503 | 459.629681040268 |
| SE_261_3_6 | SE_261 | SE_261_3 | AGAAAACTGATGCTCACAGA | 161.48274941155654 | 215.86877668092157 | 171.82967409841066 | 156.15701874796804 | 203.6341464440461 | 168.93047142433971 | 147.4340039487567 | 193.79746556401793 | 198.90870223283804 | 98.4159375516305 | 142.02798072844942 | 47.58792152184593 |
| SE_262_1_24 | SE_262 | SE_262_1 | CTATTTGTTTTATTTTGGAG | 1.083776841688299 | 2.9981774539016888 | 4.93763431317272 | 7.807850937398403 | 7.4241615891058474 | 0.8663101098684087 | 0.0 | 0.0 | 0.0 | 27.702263903421922 | 0.0 | 0.0 |
| SE_262_1_25 | SE_262 | SE_262_1 | ACTCAACTCTGTCTTCTCAA | 367.40034933233335 | 390.7624614918534 | 494.75095817990655 | 379.98208795533889 | 391.3593751971511 | 531.04809734933345 | 295.9441685102781 | 295.8956425440859 | 440.4406978012842 | 548.2132225096333 | 228.88806811609615 | 586.1439114276145 |
| SE_262_1_30 | SE_262 | SE_262_1 | CCTGCCAGTGCTTTGCCGGG | 1344.967060535179 | 1768.9246978019962 | 1791.3737288193063 | 1465.2733592517668 | 1978.0087662403437 | 1581.0159505098459 | 1440.979060491863 | 1857.6196089429036 | 2083.2134617778484 | 1641.7236397501622 | 1629.2135310007254 | 2449.036936855973 |
| SE_262_1_7 | SE_262 | SE_262_1 | TGACCAAGACAATACCACCG | 2152.3808075929614 | 2103.7211801543513 | 2074.7939383954177 | 2182.2943370028534 | 2054.3715711568607 | 1822.716471163132 | 2129.7218526612373 | 2383.2361496921426 | 2068.1177120548205 | 2446.5473068390515 | 2483.7289853008180 | 1868.6964304919986 |

| | | | | | | | | | | | | | | | |
|---|---|---|---|---|---|---|---|---|---|---|---|---|---|---|---|
| SE_262_1_9 | SE_262 | SE_262_1 | TTGGTGGAAACAGCAAGCAA | 291.5359704141524 | 289.82382054382987 | 287.3703170266523 | 251.15253848631528 | 282.1181403860222 | 635.871620643412 | 257.20238645075074 | 366.7971543357998 | 383.60964002047336 | 222.34711817220227 | 350.9617044446808 | 488.6467063584667 |
| SE_262_2_12 | SE_262 | SE_262_2 | AGGAATGTGGAAGCAAGGGA | 1326.5428542264779 | 1579.040125721556 | 1542.5169594351578 | 1284.3914792020373 | 1337.409680551782 | 1467.5293261170843 | 1298.9258596069294 | 1499.3306360221095 | 1492.7032520241105 | 1322.4185979159834 | 1381.5449034494625 | 681.3197544713063 |
| SE_262_2_13 | SE_262 | SE_262_2 | TTTTGCTGCCCTCTGCTCAG | 210.25270728753 | 209.8724217731182 | 226.1436515433106 | 175.67664609146405 | 146.36204275665813 | 86.63101098684088 | 154.96712823810924 | 260.91756339350707 | 319.6747000170611 | 170.5876250894929 | 289.9248862803885 | 80.08698987822851 |
| SE_262_2_21 | SE_262 | SE_262_2 | GGATTTAGGTTGTTACCAGG | 31.429528408960667 | 55.96597913949819 | 45.42623568118903 | 32.53271223916001 | 36.06021343279983 | 5.197860659210452 | 53.808030638232374 | 23.633837263904624 | 56.831057780810866 | 27.702263903421922 | 34.03976397623994 | 19.73157721637514 |
| SE_262_2_4 | SE_262 | SE_262_2 | AGAAGAGCTGAAGCTCAAAG | 151.72875783636184 | 163.900367479959 | 166.89203978523796 | 238.13945359065127 | 171.8163110621639 | 203.58287581907607 | 222.76524684228204 | 121.00524679119168 | 190.02884945458632 | 121.01515284126418 | 194.84830413985622 | 85.89039494186825 |
| SE_262_2_6 | SE_262 | SE_262_2 | GGAGGCCTGGACAATGGAAG | 1721.0376246010187 | 1740.941708232247 | 1703.4838380445885 | 1747.6573014876758 | 1895.28239424745 | 1623.465145893398 | 1942.4699060401888 | 1781.0459762078526 | 1844.3454220428775 | 1584.8610980536646 | 1363.9381289789935 | 1576.2048152845553 |
| SE_262_3_1 | SE_262 | SE_262_3 | CTCAAGCACTTTTCACCAGA | 193.9960546622055 | 190.88396456507417 | 231.0812858564833 | 176.97795458103047 | 176.05868911308153 | 174.12833208355016 | 182.94730416999008 | 247.68261452572048 | 217.5563930671666 | 126.84720839987932 | 289.9248862803885 | 364.4538379965761 |
| SE_262_3_14 | SE_262 | SE_262_3 | TGCCTGGCTCACCCACTCCG | 1407.8261173531002 | 1271.227240454316 | 1329.2111571060964 | 1372.8804564925524 | 1314.076601271735 | 1884.2244889637889 | 1423.7604906876286 | 1662.8767898883293 | 1586.8296914735784 | 1442.7047438124205 | 1611.6067565302565 | 2283.0595520358765 |
| SE_262_3_2 | SE_262 | SE_262_3 | GAAACAGGCAGAAGTCTGCA | 59.60772629285644 | 25.984204600481302 | 50.36386999436174 | 26.02616979132801 | 7.4241615891058474 | 114.35293450262995 | 132.36775537005164 | 49.15838150892162 | 192.69280528806183 | 5.103048613788249 | 52.820323411406804 | 27.856344305470785 |
| SE_262_3_22 | SE_262 | SE_262_3 | GTGGAGATGCCTGCAAGCGA | 469.27537245103343 | 447.7278331159855 | 464.1376254382357 | 467.1697477543378 | 489.9946648809859 | 362.11762592499485 | 415.3979965271539 | 643.785727068762 | 475.07212363646585 | 402.41183354444473 | 473.0353407732654 | 459.629681040268 |
| SE_262_3_6 | SE_262 | SE_262_3 | GTACAGGGCAGCAGACCTTG | 5153.358882227862 | 4650.173231001519 | 4576.199481448477 | 5036.06385462197 | 4975.248859213647 | 4908.513082514404 | 5607.872953116578 | 5006.592085985556 | 4550.9245488539955 | 4878.514474781566 | 5000.323949613177 | 4851.6466332028285 |
| SE_263_1_10 | SE_263 | SE_263_1 | GGGAGGCTGGAGCCCCGGGA | 72.61304839311603 | 131.9198079716743 | 125.4159115545871 | 110.61122161314404 | 101.81707322202305 | 168.0641613144713 | 87.16900963393644 | 131.40413518730972 | 110.99815972814622 | 55.404527806843845 | 99.77172199932397 | 38.30247342002233 |
| SE_263_1_25 | SE_263 | SE_263_1 | GGAACTGAAAGGGGTCAGCG | 320.7979451397365 | 422.74302100013807 | 419.6989166196812 | 314.9166544750689 | 355.29916176435125 | 530.1817872394662 | 311.0104170889831 | 416.9008893352776 | 282.379318348404 | 281.39668070318055 | 286.4035313862947 | 595.4293595294381 |
| SE_263_1_34 | SE_263 | SE_263_1 | GCAGAGCCTGGGGATGGTGG | 755.3924586567443 | 967.4119251256116 | 1044.8034206673476 | 572.5757354092162 | 802.8700461361609 | 440.08553581315164 | 549.9180731227349 | 367.74250782635596 | 880.8813956025684 | 401.68282659961784 | 266.4491869864299 | 257.67118482560477 |
| SE_263_1_36 | SE_263 | SE_263_1 | GCCTGCCCGAGAGGTCAGGG | 127.88566731921927 | 181.8894322033691 | 202.44300684008152 | 124.92561499837444 | 141.0590701930111 | 29.454543735525895 | 78.55972473181926 | 183.3985771678999 | 114.5501008394469 | 44.46942363444045 | 119.72606639918877 | 55.71268861094157 |
| SE_263_1_5 | SE_263 | SE_263_1 | GCCTTGGCAGAGACCTTTGA | 997.074694353235 | 1354.1768166789293 | 1319.3358884797508 | 1042.3481001426867 | 1154.987424362324 | 1341.0480500762967 | 970.696872713712 | 1149.549844516321 | 1685.3960573121722 | 1312.9415076332336 | 1021.1929192871983 | 1250.0534507080015 |
| SE_263_2_0 | SE_263 | SE_263_2 | GATGTCTGAGGGTTTACAGT | 331.63571355661946 | 393.7606389457551 | 370.322573487954 | 273.2747828089441 | 384.99580812077465 | 477.3368705374932 | 207.69899826357695 | 302.5131169779792 | 409.36121307740325 | 358.6714168548312 | 240.62591776307545 | 586.1439114276145 |
| SE_263_2_1 | SE_263 | SE_263_2 | CCACTTCCTGGGCAGCTCAG | 187.49339361207572 | 208.8730292884843 | 238.98150075755964 | 201.70281588279207 | 293.7846800260457 | 293.67912724539053 | 172.1856980423436 | 284.5514006574117 | 218.44437834499178 | 282.85469459283433 | 264.10161705703405 | 197.3157721637514 |
| SE_263_2_10 | SE_263 | SE_263_2 | CTGGGCTGTAACTCTGACCA | 1367.7263742106331 | 1277.2235953621193 | 1290.697609463349 | 1432.7406470126068 | 1438.1661592610756 | 1350.5774612848493 | 1560.4328885087389 | 1712.9805248878072 | 1030.950907555022 | 1340.6437715366555 | 2203.194378738013 | 1471.74352413904 |
| SE_263_2_24 | SE_263 | SE_263_2 | TGTGGCTGTGATCTCAGTAA | 2472.0949758910097 | 2420.5285977832964 | 2457.95436109738 | 2298.110792574263 | 2658.9104434126225 | 2130.256560166417 | 2251.3280019036424 | 2202.6736329959112 | 2566.2774529147405 | 2305.1199595426347 | 2351.091284289952 | 1806.0196558046894 |
| SE_263_2_3 | SE_263 | SE_263_2 | GCTGCTACAGCAGAGGAAGT | 592.8259324034995 | 553.6634364871785 | 583.6283758170156 | 628.5320004605715 | 656.5080033795027 | 752.8234854756472 | 628.4777978545541 | 701.4522899926893 | 522.1353433611998 | 497.1827363719408 | 636.1914508662776 | 481.68262028209904 |
| SE_264_1_1 | SE_264 | SE_264_1 | GAGCTGGTCACCATACATGT | 2246.669392819844 | 2089.729685369477 | 1870.3758778298263 | 2119.8315295036664 | 2211.339559040813 | 2255.005215987468 | 2293.298265801464 | 1783.8820366795212 | 1749.3309973155845 | 1766.383827315561 | 1902.705427775343 | 1698.07632162099 |
| SE_264_1_20 | SE_264 | SE_264_1 | GGATGTGGCCGGCATCGTGG | 2756.044508413344 | 2825.2825540600247 | 2814.4515585084505 | 2851.1669006399834 | 2899.665397802198 | 2403.144244774966 | 2871.1965148560794 | 3120.6118723259665 | 2614.2286579172996 | 3354.88996009336 | 2785.391721228186 | 2394.4849292577596 |
| SE_264_1_25 | SE_264 | SE_264_1 | TGATGCCCTTGACAAGTGTG | 1990.898058181405 | 1555.0547060903425 | 1585.9681413910778 | 1984.4954465887606 | 1581.3464184795455 | 1751.6790421539224 | 2093.132391827239 | 1371.7079147970244 | 1631.2289553648368 | 1752.53269536385 | 1824.0618351405817 | 1073.6299367733532 |
| SE_264_1_26 | SE_264 | SE_264_1 | TGTGGAACTGAAAGCAAGGC | 1424.0827699784247 | 1678.9793741849455 | 1712.3715798082994 | 1488.6969120639621 | 1693.7694368288626 | 1206.7699830466934 | 1332.2868386026337 | 1314.0413518730973 | 1710.259645091277 | 1639.5366189156816 | 1412.0633125316085 | 1556.47323806818 |
| SE_264_1_9 | SE_264 | SE_264_1 | GCAGTGGGCCTCATTCCCTA | 8491.391554627822 | 7318.551164974022 | 7301.773622319819 | 7967.91188161507 | 7487.7972598696115 | 7871.2936582643615 | 8347.777873215371 | 8574.356159344597 | 7215.768367607329 | 7367.344184420577 | 8313.918904955432 | 7125.420737136881 |
| SE_264_2_10 | SE_264 | SE_264_2 | TGAAAAGCCGGACTGACGGG | 433.51073667531955 | 774.5291755912696 | 778.1711677560207 | 564.7678844718178 | 750.90091501242 | 909.6256153618292 | 354.056841599569 | 694.834815558796 | 857.793778379114 | 610.1788128201091 | 705.4447637834553 | 1212.9116583007071 |
| SE_264_2_20 | SE_264 | SE_264_2 | ATAAATTCAGATTAGACCAA | 15.172875783636185 | 11.992709815606755 | 9.87526862634544 | 9.109159426964803 | 19.090701229129323 | 0.8663101098684087 | 8.60928490211718 | 14.180302358342775 | 25.751573056929924 | 0.0 | 3.521354894093787 | 1.1606810127279494 |
| SE_264_2_26 | SE_264 | SE_264_2 | CAGAACCTGGGTGAACCCAG | 1.083776841688299 | 3.9975699385355847 | 2.962580587903632 | 9.109159426964803 | 21.211890254588134 | 0.0 | 8.60928490211718 | 32.14201867891029 | 10.655823333902038 | 50.30147919305559 | 37.56111887033373 | 0.0 |
| SE_264_2_27 | SE_264 | SE_264_2 | ACTCACTCCACCAATCACGT | 4186.629939441898 | 3821.676861240019 | 3793.0906793792838 | 3742.563215992968 | 3824.503812902241 | 3599.5185065032383 | 4171.198535075773 | 3772.9057808097346 | 3554.6050671341545 | 3838.9505714584166 | 3246.6892123544717 | 3599.271820469371 |
| SE_264_2_33 | SE_264 | SE_264_2 | AATGTTCTGTGGGTTCACCC | 149.56120415298525 | 119.92709815606754 | 139.2412876314707 | 113.21383859227684 | 110.3018293238583 | 77.10159977828837 | 135.59623720834557 | 70.90151179171387 | 86.13457194904147 | 37.908361130998415 | 92.72901221113639 | 148.56716962917753 |
| SE_265_1_10 | SE_265 | SE_265_1 | GGCCCGCTCTACTTCCACCG | 186.4096167703874 | 213.8699917116538 | 262.6821454607887 | 227.72898567412008 | 255.60327756778702 | 300.60960812433785 | 216.30828316569415 | 280.76998669518696 | 215.78042251151626 | 185.89677093085763 | 244.14727265716925 | 240.26096963468552 |
| SE_265_1_12 | SE_265 | SE_265_1 | CTAAAGAAACCCTAAGCCCA | 573.3179492531101 | 936.4307581019607 | 871.9862197063023 | 735.2392966050162 | 897.2629577690781 | 1106.278010301958 | 623.0969947907308 | 868.779857821134 | 877.3294544912677 | 662.6673128476455 | 766.4815819477477 | 1057.380402595162 |
| SE_265_1_16 | SE_265 | SE_265_1 | GTAGTCAGCCAGCCCACCAT | 3737.946326982943 | 3244.028005121627 | 3354.628752369546 | 3382.100764383075 | 3059.8151692243387 | 3069.336719263772 | 3554.558503961631 | 3265.250956381063 | 3575.0287285241334 | 2792.825605631826 | 4100.030881689866 | 3375.260385012877 |
| SE_265_1_18 | SE_265 | SE_265_1 | CTCAAGAGTGAAGTGCCAAT | 532.1344292689548 | 413.748488638433 | 410.8111748559703 | 446.34881192127534 | 460.2980185245625 | 345.6577338374951 | 420.77879959097714 | 509.54553140978373 | 357.8580669635434 | 279.93866681352677 | 385.00146842092073 | 249.54641773650914 |
| SE_265_1_2 | SE_265 | SE_265_1 | AGACATGAGACACACAGTAA | 1367.7263742106331 | 1246.2424283384685 | 1214.6580410404893 | 1233.6404481089476 | 1311.9554122462762 | 951.2085006355128 | 1326.9060355388103 | 1453.0083149848563 | 1440.3121206324254 | 797.5335976406205 | 1250.0809874032943 | 1152.5562456388539 |
| SE_265_2_11 | SE_265 | SE_265_2 | TTGAGAGAAGAATCTGGGAG | 118.13167574402458 | 148.90948021045054 | 168.86709351050703 | 134.03477442533924 | 181.36166167672854 | 218.310147686839 | 77.48356411905462 | 83.19110716894428 | 247.74789251322238 | 112.26706950334147 | 76.29602270536539 | 211.2439443164868 |
| SE_265_2_15 | SE_265 | SE_265_2 | GTACATCATGATACTCCATA | 767.3140039153157 | 599.6354907803377 | 559.9277311137864 | 692.296116449325 | 462.4192075500213 | 583.8930140513074 | 738.2461803565482 | 419.73694980694614 | 546.9989311403045 | 747.232118447565 | 500.0323949613178 | 1027.2026962642353 |
| SE_265_2_7 | SE_265 | SE_265_2 | CAGTCGCCTATAAAGTACAA | 37.93218945909046 | 50.9690167163287 | 53.32645058226538 | 62.46280749918722 | 61.51448173830559 | 49.379676262499295 | 40.8941032850566 | 198.52423301679886 | 40.84732277995781 | 21.141201399979888 | 346.26656458588906 | 16.249534178191293 |
| SE_265_2_8 | SE_265 | SE_265_2 | TAGGAAAGTTCACATCGGCG | 2395.1468201311404 | 3092.120347457275 | 3039.6076831891264 | 2570.0842668936407 | 3188.1471052645966 | 2935.0586522341687 | 2658.116713528679 | 3297.392975059973 | 3070.653090719437 | 2830.004959817997 | 2997.846799838511 | 2948.1297723289917 |
| SE_265_2_9 | SE_265 | SE_265_2 | TGTCTCCTAAGGTGGCCGTG | 92.12103154350541 | 97.94046349412183 | 108.62795488979984 | 93.69421124878083 | 74.24161589105847 | 6.064170769078861 | 78.55972473181926 | 76.57363273505099 | 63.93494000341222 | 111.53806255851458 | 57.515463270198524 | 42.945197470934126 |
| SE_265_3_12 | SE_265 | SE_265_3 | ACCGGCTTCGGGGCGCGCGG | 707.7062776224592 | 1069.349958558269 | 1127.7556771286493 | 684.4882655119267 | 1049.9885676021127 | 825.5935347045935 | 521.9378971908541 | 845.1460205572293 | 1076.2381567241057 | 407.514882158233 | 694.880699101174 | 521.1457747148493 |
| SE_265_3_2 | SE_265 | SE_265_3 | CGTTTTGCCGTGCCGCTCGG | 2658.504592661397 | 2447.512194868412 | 2474.742317762167 | 2597.411745174535 | 2340.7320895938005 | 2637.047974439436 | 2851.8256238263157 | 2887.109560158589 | 2493.462660133077 | 2338.6542790046715 | 2682.098644334768 | 2771.7062583943434 |
| SE_265_3_4 | SE_265 | SE_265_3 | TTTTGCCGTGCCGCTCGGAG | 140.89098941947887 | 103.93681840192521 | 139.2412876314707 | 195.19627343496006 | 178.17987813854035 | 73.63635933881474 | 148.51016456152135 | 143.69373056454012 | 104.78226278337003 | 131.22125006884067 | 131.46391604616804 | 76.60494684004466 |
| SE_265_3_42 | SE_265 | SE_265_3 | GGCTACTTTGGGGAGCTGGG | 37.93218945909046 | 69.95747392437274 | 67.151826659149 | 40.34056317655841 | 42.42378050917627 | 26.85561340592067 | 19.370891029763655 | 43.48626056558451 | 15.095749723027886 | 22.599215289633673 | 37.56111887033373 | 27.856344305470785 |
| SE_265_4_13 | SE_265 | SE_265_4 | ACCACATTGAGCGCTGTAAG | 1414.32877840323 | 1413.1409732723291 | 1573.1302921768286 | 1441.8498064395717 | 1547.4073940722044 | 1376.5667645809015 | 1303.2305020579881 | 1170.347621308557 | 1533.5505748040682 | 1551.3267785916275 | 1051.7113283693443 | 1508.8853165463343 |
| SE_265_4_15 | SE_265 | SE_265_4 | AATGGATATCTGGCTCAGGC | 67.19416418467453 | 88.94593113241676 | 119.49075037877982 | 76.77720088441762 | 130.45312506571702 | 109.1550738434195 | 41.970263897821255 | 80.35504669727572 | 152.7334677859292 | 95.49990977232294 | 143.20176569314734 | 55.71268861094157 |
| SE_265_4_25 | SE_265 | SE_265_4 | GCCTGGAACCAGATTGTGAA | 1892.2743655877698 | 2146.695056993609 | 2102.4446905489444 | 1879.0894589338823 | 2083.0076230005548 | 2039.2939986302342 | 2059.771412831535 | 2167.695553845332 | 2160.4681809486383 | 2216.9101192185803 | 2462.600855936255 | 3097.857622970897 |
| SE_265_4_3 | SE_265 | SE_265_4 | TGGAAAGAGAAGGCAAACAG | 332.71949039830776 | 310.8110627211417 | 321.93375721886133 | 239.44076208021767 | 335.1478660224925 | 395.9037202098628 | 259.35470767628004 | 387.59493112803585 | 315.2347736279353 | 282.85469459283433 | 207.75993875153344 | 365.61451900930405 |
| SE_265_4_4 | SE_265 | SE_265_4 | CTGCACTTACACTACAGCCT | 126.80189047753097 | 137.91616287947767 | 151.09160998308525 | 141.84262536273764 | 114.54420737477592 | 84.89839076710406 | 120.52998862964051 | 117.22383282896693 | 253.96378945799856 | 761.0832503992759 | 41.08247376442752 | 178.74487596010422 |
| SE_265_5_1 | SE_265 | SE_265_5 | GGAAAATTCAAGCCACAGAA | 74.78060207649263 | 127.92223803313871 | 127.39096527985618 | 110.61122161314404 | 120.90777445115236 | 81.43315032763041 | 157.11944946363852 | 124.78666075341643 | 123.4299536176986 | 384.1866599237724 | 102.11929192871983 | 192.6730481128396 |
| SE_265_5_14 | SE_265 | SE_265_5 | CTCAGCACTGAGCACAGGCG | 21.675536833765978 | 19.987849692677923 | 14.81290293951816 | 13.013084895664004 | 15.9089176909411 | 65.83956834999907 | 15.066248578705064 | 2.836060471668555 | 53.27911666951019 | 0.7290069448268927 | 1.173784964697929 | 26.695663292742836 |
| SE_265_5_16 | SE_265 | SE_265_5 | TGGAGACTGCAGGGGCCACA | 1.083776841688299 | 6.9957473924372735 | 4.93763431317272 | 0.0 | 0.0 | 17.326202197368175 | 1.0761606127646475 | 1.89070698111237 | 9.767838056076867 | 0.0 | 0.0 | 4.642724050911798 |
| SE_265_5_22 | SE_265 | SE_265_5 | CACACAGCCAGGGAACAATA | 1777.3940203688103 | 1942.8189901282942 | 1906.9143717473046 | 1698.2075788841526 | 1708.6177600070744 | 1585.347501059188 | 1546.4428005427985 | 1700.6909295105768 | 2024.606433441387 | 2285.4367720323085 | 1604.564046742069 | 1837.358043148344 |
| SE_265_5_9 | SE_265 | SE_265_5 | TTTGGCTCTAAAGTTTCCAT | 40.09974314246706 | 44.97266180852533 | 44.43870881855448 | 33.834020728726415 | 55.15091466192915 | 25.12299318618385 | 31.208657770174778 | 19.852423301679885 | 117.21405667292241 | 50.30147919305559 | 32.86597901154201 | 11.606810127279495 |
| SE_265_6_10 | SE_265 | SE_265_6 | TATTCTCTGAAACACCCACT | 57.440172609479845 | 51.968409200962604 | 69.12688038441809 | 59.86019052005442 | 39.24199697098805 | 40.71657516381521 | 32.28481838293943 | 67.12009782948914 | 34.63142583518162 | 26.244250013768134 | 88.03387235234467 | 17.410215190919242 |
| SE_265_6_15 | SE_265 | SE_265_6 | GCTTGGTTCTACACATTTTA | 437.84584404207277 | 444.7296556620838 | 468.08773288877387 | 465.8684392647714 | 519.6913112374093 | 431.42243471446756 | 530.5471820929712 | 489.69310810810384 | 543.4469900290039 | 412.6179307720212 | 527.0294491493702 | 387.6674582511351 |
| SE_265_6_22 | SE_265 | SE_265_6 | CTAGCTGGAATTTACCACCA | 725.046707089472 | 662.5972173122732 | 748.5453618769844 | 681.8856485327939 | 765.7492381906317 | 711.2406002019636 | 654.3056525609056 | 737.3757226338244 | 675.7567964249542 | 892.3045004681167 | 758.2650871948622 | 634.8925139621883 |
| SE_265_6_3 | SE_265 | SE_265_6 | TCTCGACCTTGGCTGCACAT | 1048.012205912585 | 777.5273530451713 | 674.4808471793935 | 990.2957605600308 | 761.506860139714 | 785.7432696506468 | 963.1637484243595 | 880.1240997078082 | 738.8037511505413 | 732.6519795510271 | 1034.1045538988756 | 749.7999342222554 |
| SE_265_6_4 | SE_265 | SE_265_6 | GATAAGAAGCCAAGTTTCAG | 85.61837049337561 | 195.88092698824366 | 232.06881271911786 | 131.43215744620645 | 182.42225618945795 | 207.9144263684181 | 128.06311291899306 | 175.8357492434504 | 227.32423112324346 | 62.69459725511277 | 139.68041079905356 | 107.94333418369929 |
| SE_265_7_12 | SE_265 | SE_265_7 | TGTACTAAGTTTAAATCCAG | 123.55055995246607 | 157.9040125721556 | 172.81720096104522 | 109.30991312357764 | 131.51371957844643 | 95.29411208552496 | 111.92070372752333 | 80.35504669727572 | 123.4299536176986 | 183.70975009637695 | 127.94256115207426 | 201.9584962146632 |
| SE_265_7_13 | SE_265 | SE_265_7 | GTACTAAGTTTAAATCCAGC | 172.32051782843953 | 194.88153450360974 | 204.41806056535063 | 162.66356119580004 | 180.30106716399914 | 130.81282659012973 | 148.51016456152135 | 175.8357492434504 | 218.44437834499178 | 309.82795155142935 | 311.0530156449512 | 164.8167038073688 |
| SE_265_7_15 | SE_265 | SE_265_7 | AATTCAGGCTGCTTGACGTG | 423.7567451001249 | 307.81288526724 | 259.7195648728851 | 446.34881192127534 | 258.7850611059753 | 99.625662634867 | 414.3218359143893 | 374.35998226024924 | 280.60334779275365 | 422.8240279995977 | 973.0677357345832 | 241.4216506474135 |
| SE_265_7_28 | SE_265 | SE_265_7 | GCACACAGCAAAAGCCCACG | 7886.644076965751 | 7814.249837352434 | 7976.254469499212 | 7829.973181721031 | 8432.786970711513 | 8858.887183514347 | 8003.406477130683 | 7594.024589637834 | 7935.924427923543 | 9642.57485922531 | 7984.085329875314 | 8831.621825846967 |
| SE_265_7_7 | SE_265 | SE_265_7 | GTTCTGGAGCCATCTCAAGA | 32.51330525064897 | 63.961119016569356 | 51.35139685699629 | 36.43663770785921 | 92.27172260745839 | 110.02138395328791 | 8.60928490211718 | 69.95615830115769 | 30.191499446055772 | 16.76715973101853 | 70.42709788187574 | 176.42351393464833 |
| SE_266_1_14 | SE_266 | SE_266_1 | ATTCATTCTACAAATCTACT | 0.0 | 11.992709815606755 | 4.93763431317272 | 0.0 | 4.242378050917627 | 0.0 | 0.0 | 0.0 | 0.8879852778251698 | 0.0 | 0.0 | 0.0 |
| SE_266_1_15 | SE_266 | SE_266_1 | ATAATAGTATCATCTTCATA | 11.921545258571289 | 12.992102300240651 | 26.66322529113269 | 33.834020728726415 | 42.42378050917627 | 0.8663101098684087 | 12.913927353175769 | 43.48626056558451 | 1.7759705556503396 | 15.309145841364746 | 45.77761362321923 | 0.0 |
| SE_266_1_3 | SE_266 | SE_266_1 | ACTCTAGAGGGAGAATGCCT | 260.10644200519175 | 287.8250355745621 | 254.78193055971235 | 303.2048780689713 | 299.0876525896927 | 285.01602614670645 | 234.60301358269314 | 308.1852379213163 | 285.9312594597047 | 103.51898616541875 | 417.8674474324627 | 215.88666836739858 |
| SE_266_1_6 | SE_266 | SE_266_1 | AGTGAATTCGATAATGAAGG | 212.42026097090658 | 271.8347558204198 | 277.49504840030687 | 244.64599603848328 | 224.84603669863424 | 246.8983813124965 | 230.29837113163455 | 120.0598933006355 | 194.4687758437122 | 298.89284737902597 | 59.86303319959438 | 81.24767089095646 |
| SE_266_1_7 | SE_266 | SE_266_1 | ACAAAGTGAAGAGAGAGAGG | 307.7926230394769 | 442.730870692816 | 424.63655093285394 | 206.90804984105768 | 327.7237044333867 | 213.11228702762855 | 258.2785470635154 | 239.1744331107148 | 451.98450641301145 | 96.22891671714983 | 137.3328408696577 | 163.65602279464088 |
| SE_267_1_24 | SE_267 | SE_267_1 | GCTGGAGGAGAGGAATACCC | 4058.7442721226794 | 3859.653775656107 | 3688.412831940022 | 3711.331812243374 | 3961.3205050443344 | 3441.850066507188 | 3890.3206151442005 | 4312.702623917316 | 3808.568856592153 | 3042.87498770745 | 3999.0853747258443 | 3143.124182467287 |
| SE_267_1_27 | SE_267 | SE_267_1 | ATAGCCTTCAGGGAGAGGAA | 254.68755779675024 | 340.7928372601586 | 350.5720362352631 | 312.3140374959361 | 358.48094530253945 | 485.1336615263089 | 345.44755669745183 | 395.15775905248535 | 482.1760058590672 | 472.39650024782645 | 258.2326922335444 | 451.50491395117234 |
| SE_267_1_29 | SE_267 | SE_267_1 | GGTTGGAGCAGATCCACCCA | 485.5320250763579 | 461.71932790086004 | 478.95052837775387 | 532.2351722326578 | 384.99580812077465 | 272.02137449868036 | 397.1032661101549 | 639.058959615981 | 450.2085358573611 | 569.3544239098031 | 508.24888971420324 | 500.2535164857462 |
| SE_267_1_6 | SE_267 | SE_267_1 | GTAACTCAACCACAGTATGG | 237.34712832973747 | 325.80194999065014 | 305.1458005540741 | 303.2048780689713 | 314.9965702806338 | 228.7058690052599 | 231.3745317443992 | 190.96140509234937 | 315.2347736279353 | 707.1367364820859 | 147.89690555193906 | 509.5389645875698 |
| SE_267_1_8 | SE_267 | SE_267_1 | CCTCACCTCTAAGTCACCAT | 2184.8941128436104 | 1845.8779191188062 | 1910.8644791978427 | 2221.3335916898454 | 2006.6448180840375 | 1715.2940175394492 | 2353.563260116284 | 2067.4880838463764 | 2090.31734400045 | 2751.2722097766928 | 1952.0043962926559 | 2032.3524532866395 |
| SE_267_2_11 | SE_267 | SE_267_2 | GCTTCCTTGGCCCCACCATG | 139.80721257779055 | 132.9192004563082 | 156.02924429625796 | 152.25309327926885 | 130.45312506571702 | 124.74865582105086 | 159.27177068916782 | 107.7702979234051 | 64.82292528123739 | 83.10679171026577 | 82.16494752885504 | 80.08698987822851 |
| SE_267_2_12 | SE_267 | SE_267_2 | CATAATTCCTCAGGAGACAG | 475.77803350116324 | 435.73512330037875 | 439.4494538723721 | 468.4710562439042 | 539.842606979268 | 626.3422094348595 | 492.8815606462085 | 639.058959615981 | 640.2373853119474 | 348.4653196272547 | 615.0633215017148 | 949.4370684114626 |
| SE_267_2_27 | SE_267 | SE_267_2 | GCTGCTTCGAGGAAGAAGGT | 835.5919449416784 | 972.408887548781 | 948.0257881291623 | 818.5230399372658 | 870.748094950843 | 895.7646536039347 | 843.7099204074837 | 899.9765230094881 | 775.2111475413732 | 952.8120768887487 | 822.8232602532482 | 1024.8813342387793 |
| SE_267_2_35 | SE_267 | SE_267_2 | TTACTGGCAGGGTTCATGCA | 453.01871982570896 | 449.7266180852533 | 500.6761193557138 | 378.6807704638225 | 491.05525939371535 | 375.9785876828894 | 334.68595056980536 | 484.0209871647667 | 414.6891247443543 | 417.72097938580947 | 518.8129543964847 | 367.93588103476 |
| SE_267_2_4 | SE_267 | SE_267_2 | CATTATCCCCAGAAACCAGA | 395.5785472162291 | 467.7156828086634 | 511.5389148446938 | 564.7678844718178 | 537.7214179538092 | 727.7004922894633 | 570.3651247652632 | 534.1247221642445 | 559.430725029857 | 852.2091185026375 | 698.4020539952678 | 591.9473164912542 |
| SE_267_3_11 | SE_267 | SE_267_3 | GCCTATCGATGGTGGCCCCA | 83.45081680999901 | 108.93378082509469 | 129.36601900512528 | 63.76411598875362 | 119.84717993842297 | 122.14972549144564 | 54.88419125099702 | 136.13090264009065 | 225.54826056759313 | 70.71367364820858 | 37.56111887033373 | 170.62010887100857 |

| | | | | | | | | | | | | | | | |
|---|---|---|---|---|---|---|---|---|---|---|---|---|---|---|---|
| SE_267_3_13 | SE_267 | SE_267_3 | CTTCACTGAGGGCAGCCTTA | 113.79656837727138 | 191.88335704970808 | 139.2412876314707 | 101.50206218617923 | 127.2713415275288 | 168.93047142433971 | 124.83463108069911 | 86.02716764061283 | 99.45435111641902 | 169.858618144666 | 190.1531642810645 | 40.62383544547823 |
| SE_267_3_2 | SE_267 | SE_267_3 | CATACAGCTGGTGAGCAGCA | 4446.73638144709 | 4048.5389552519136 | 4018.2468040599597 | 4653.479158689448 | 4072.6829288809217 | 4401.721668241385 | 4553.235552607223 | 4292.850200615636 | 4343.135993842905 | 4722.5069885886105 | 3912.2252873381976 | 4281.752255953405 |
| SE_267_3_20 | SE_267 | SE_267_3 | GGAAAACTTGGCCAAGAAAA | 69.36171786805113 | 123.92466809460312 | 123.440857829318 | 84.58505182181602 | 144.2408537311993 | 157.66843999605038 | 62.417315540349556 | 117.22383282896693 | 113.66211556162173 | 29.160277793075707 | 105.64064682281361 | 210.08326330375886 |
| SE_267_3_21 | SE_267 | SE_267_3 | CGGTGTGGATCTCACCCACA | 4058.7442721226794 | 3602.8099071051956 | 3497.820147451555 | 3831.0521932834827 | 4129.955032568309 | 3001.7645306940362 | 4296.033166156472 | 3011.8962209120054 | 3567.036861023707 | 3939.553529844528 | 3258.427062001451 | 3117.589200187272 |
| SE_267_4_17 | SE_267 | SE_267_4 | CCACCCTCGTCCACACCACA | 1582.3141888649163 | 1455.1154576269528 | 1355.874382397229 | 1469.177284720466 | 1511.3471806394045 | 1349.7111511749808 | 1462.502272747156 | 1330.1123612125523 | 1651.6526167548159 | 2137.448362232449 | 1670.296004765153 | 1373.0856380571643 |
| SE_267_4_24 | SE_267 | SE_267_4 | ACAAAGCCCAGGGGTCTGAG | 227.59313675454277 | 301.8165303594366 | 318.9711766309577 | 231.63291114281927 | 317.11775930609264 | 229.5721791151283 | 186.175786008284 | 286.4421076385241 | 361.4100080748441 | 145.07238202055163 | 281.70839152750295 | 243.74301267286938 |
| SE_267_4_45 | SE_267 | SE_267_4 | TGGAGTGGCGGTCAGCCAAG | 767.3140039153157 | 646.6069375581309 | 605.3539667949755 | 672.776489105829 | 642.7202747140204 | 498.99462328420344 | 634.934761531142 | 554.9224989564806 | 713.0521780936114 | 581.0185350270334 | 728.920463077414 | 843.8150962532193 |
| SE_267_4_5 | SE_267 | SE_267_4 | GGGGCCTGAGAAACTCTTAG | 26.010644200519174 | 41.97448435462364 | 46.41376254382357 | 35.13532921829281 | 18.030106716399914 | 28.588233625657487 | 31.208657770174778 | 30.25131169779792 | 72.81479278166393 | 1.4580138896537853 | 32.86597901154201 | 24.37430126728694 |
| SE_267_4_6 | SE_267 | SE_267_4 | CTTGGAACTGACATACAGGA | 1937.7929929386785 | 1924.829925404884 | 1945.4279193900518 | 2044.355637108815 | 2070.280488847802 | 2053.1549603881285 | 1784.2742959637856 | 1873.6906182823586 | 1892.2966270454367 | 2294.184855370231 | 1885.098653304874 | 1828.0725950465203 |
| SE_268_1_11 | SE_268 | SE_268_1 | TGAGAGTCAGTGACATATAG | 1331.9617384349194 | 1357.174994132831 | 1343.0365331829798 | 1397.605317794314 | 1354.3791927554523 | 1209.3689133762987 | 1281.7072898026952 | 1501.2213430032218 | 1210.3239336757065 | 1580.4870563847032 | 1468.4049908371092 | 976.1327317042055 |
| SE_268_1_13 | SE_268 | SE_268_1 | CGGTTTCCTGATCTGTAAAA | 5.4188842084414945 | 4.996962423169481 | 4.93763431317272 | 3.9039254686992013 | 6.363567076376441 | 4.331550549342044 | 2.152321225529295 | 0.0 | 0.0 | 2.187020834480678 | 34.03976397623994 | 3.482043038183848 |
| SE_268_1_26 | SE_268 | SE_268_1 | TCACATACACACTTCCTGAA | 2809.149573656071 | 2991.1817065092514 | 3076.1461771066047 | 2929.2454100139676 | 2999.3612819987625 | 3162.0319010196918 | 3020.7828400303656 | 2832.27905770633 | 2990.734415715172 | 3625.351536624137 | 2253.6671322200236 | 2921.4341090362486 |
| SE_268_1_28 | SE_268 | SE_268_1 | GTATGTGAGACCTAAAGCAA | 387.992109324411 | 408.75152621526354 | 433.5242926965648 | 369.57161103685775 | 415.75304898992744 | 415.8288527368362 | 371.2754114038034 | 380.9774566941426 | 430.67285974520735 | 216.5150626135871 | 295.7938111038781 | 326.1513645765538 |
| SE_268_1_9 | SE_268 | SE_268_1 | GGAGGTTTCTGAGGTCAGAG | 203.7500462374002 | 195.88092698824366 | 226.1436515433106 | 175.67664609146405 | 186.6646342403756 | 210.51335669802333 | 244.288459097575 | 151.2565584889896 | 221.99631945629244 | 166.94259036535843 | 181.936669528179 | 192.6730481128396 |
| SE_268_2_0 | SE_268 | SE_268_2 | CTCTGTCTCCTTCAAAGCAC | 133.30455152766078 | 100.93864094802352 | 89.8649444997435 | 102.80337067574564 | 61.51448173830559 | 124.74865582105086 | 94.70213392328898 | 159.76473990399526 | 54.167101947335354 | 54.67552086201695 | 192.50073421046037 | 67.31949873822107 |
| SE_268_2_11 | SE_268 | SE_268_2 | CTGTTTCCTTAGAGCCTGGA | 23.843090517142578 | 31.980559508284678 | 26.66322529113269 | 26.02616979132801 | 39.24199697098805 | 0.8663101098684087 | 8.60928490211718 | 77.51898622560716 | 36.40739639083196 | 23.328222234460565 | 46.95139858791716 | 31.338387343654635 |
| SE_268_2_4 | SE_268 | SE_268_2 | GTTAATGTGGAAGTAGGCAG | 259.02266516350346 | 361.7800794374704 | 387.11053015274126 | 249.85122999674888 | 286.3605184369398 | 403.70051119867844 | 246.44078032310426 | 244.84655405405192 | 404.0333014104522 | 372.52254880654215 | 237.10456286898167 | 557.1268861094158 |
| SE_268_2_6 | SE_268 | SE_268_2 | TTATATACTCCTCCTCTTAA | 10.837768416882989 | 17.98906472341013 | 10.862795488979984 | 7.807850937398403 | 3.1817835381882205 | 18.192512307236584 | 1.0761606127646475 | 0.0 | 19.535676112153734 | 0.0 | 5.868924823489645 | 37.14179240729438 |
| SE_268_2_8 | SE_268 | SE_268_2 | CAGTTTGGAGACTTGTAGAC | 360.89768828220355 | 375.77157422234495 | 395.01074505381763 | 390.3925468699201 | 378.6322410443982 | 366.44917647433687 | 370.1992507910387 | 521.8351267870141 | 379.1697136313475 | 643.7131322821463 | 467.16641594977574 | 342.40089875474507 |
| SE_268_3_0 | SE_268 | SE_268_3 | GGTACCATAGGGTAGAACAC | 331.63571355661946 | 329.79951992918575 | 331.8090258452068 | 318.8205799437681 | 250.30030500414 | 319.66843054144283 | 295.9441685102781 | 413.11947537305286 | 374.7297872422217 | 341.90425712381267 | 561.0692131256101 | 236.7789265965017 |
| SE_268_3_1 | SE_268 | SE_268_3 | GCTTCCAGACTCTTTCTCTG | 48.769957875973454 | 90.94471610168455 | 75.05204156022535 | 61.16149900962082 | 97.57469517110542 | 74.50266944868315 | 44.12258512335055 | 96.42605603673087 | 182.92496723198497 | 39.36637502065221 | 171.37260484589763 | 33.659749369110536 |
| SE_268_3_10 | SE_268 | SE_268_3 | TGTATTGTCATGGCGCTGGG | 496.3697934932409 | 439.7326932389143 | 539.189666998461 | 477.58021567086894 | 467.7221801136684 | 356.05345515591597 | 378.80853569315593 | 468.89533131586774 | 282.379318348404 | 427.19806966855907 | 450.73342644400475 | 364.4538379965761 |
| SE_268_3_11 | SE_268 | SE_268_3 | CTATAGGCAGAGCAGCAGTG | 528.8830987438898 | 487.70353250134133 | 538.2021401358265 | 498.4011515039314 | 426.3589941172215 | 602.9518364684125 | 589.7360157950268 | 531.2886616925759 | 557.6547544742066 | 568.6254169649762 | 592.7614071724541 | 305.2591063474507 |
| SE_268_3_25 | SE_268 | SE_268_3 | GTAGACCCCACCTATTTATG | 95.37236206857031 | 100.93864094802352 | 99.74021312608895 | 109.30991312357764 | 124.08955798934059 | 69.3048087894727 | 77.48356411905462 | 118.16918631952312 | 84.35860139339113 | 40.0953819654791 | 137.3328408696577 | 188.03032406192781 |
| SE_269_1_18 | SE_269 | SE_269_1 | TATTTTACTACAGTTTTGAA | 4.335107366753196 | 7.9951398770711695 | 5.925161175807264 | 6.506542447832002 | 6.363567076376441 | 24.256683076315444 | 27.980175931880833 | 43.48626056558451 | 3.551941111300679 | 81.64877782061198 | 31.692194046844083 | 0.0 |
| SE_269_1_3 | SE_269 | SE_269_1 | AAACTTTTAGAAGACAACAT | 1.083776841688299 | 5.9963549078033775 | 7.900214901076352 | 1.3013084895664004 | 6.363567076376441 | 0.0 | 1.0761606127646475 | 0.0 | 3.551941111300679 | 0.7290069448268927 | 0.0 | 0.0 |
| SE_269_1_5 | SE_269 | SE_269_1 | AGTTGGTGTGTCGATAAAAA | 68.27794102636283 | 164.89975996459287 | 179.729888999487 | 97.59813671748003 | 149.54382629484635 | 193.18715450065514 | 89.32133085946575 | 132.3494886778659 | 66.59889583688773 | 86.75182643440023 | 59.86303319959438 | 186.86964304919985 |
| SE_269_1_8 | SE_269 | SE_269_1 | GTTCCTGTATGTTTGTTGCT | 44.434850509220254 | 42.973876839257535 | 55.301504307534465 | 58.55888203048802 | 47.72675307282331 | 116.08555472236677 | 63.4934761531142 | 53.88514896170255 | 101.23032167206935 | 192.45783343429966 | 79.81737759945918 | 25.534982280014887 |
| SE_269_1_9 | SE_269 | SE_269_1 | GCAGTTGCCAGGTCACATGA | 2412.4872495981535 | 2119.711459908494 | 2196.259742499226 | 2626.040531944996 | 2366.1863578993066 | 2676.0319293835146 | 2830.3024115710227 | 2527.875233747239 | 2428.6397348518394 | 2311.6810220460766 | 2773.653871581206 | 3022.4133571435805 |
| SE_269_2_3 | SE_269 | SE_269_2 | TCAGGGCTGCTCAGTCGCAT | 24327.538765377245 | 20659.4414423519 | 21326.630125455613 | 23260.889250999408 | 20499.170742033974 | 20489.100408497736 | 23138.529335052684 | 20184.242376865106 | 20317.99114191771 | 20973.529802669702 | 20443.81273014383 | 20224.866646784518 |
| SE_269_2_4 | SE_269 | SE_269_2 | GCAGAGGTTGACTTGAATGA | 5046.06497490072 | 5516.646515179107 | 5347.457961166056 | 5688.019407894736 | 5003.884911057341 | 4908.513082514404 | 5244.130666002127 | 5269.4003563601755 | 5764.800423641002 | 5912.2463225461 | 5328.983739728598 | 6345.443096583699 |
| SE_269_2_6 | SE_269 | SE_269_2 | AATATACTGCAGCCTTCCAG | 0.0 | 0.9993924846338962 | 3.950107450538176 | 0.0 | 0.0 | 23.390372966447035 | 0.0 | 0.0 | 0.0 | 0.0 | 0.0 | 0.0 |
| SE_269_2_8 | SE_269 | SE_269_2 | TGTGGGCTCTGAACTTAGTT | 1796.9020035191995 | 1786.9137625254064 | 1990.8541550712407 | 1910.3208626834758 | 2034.220275415002 | 1875.561387865105 | 1904.804284593426 | 2130.826767713641 | 1864.7690834328566 | 2024.452285784281 | 2859.340174004155 | 1908.159584924749 |
| SE_269_2_9 | SE_269 | SE_269_2 | ACCTTACTACAAGATCGGAG | 542.9721976858377 | 658.5996473737376 | 619.1793428718591 | 543.9469486387554 | 654.386814354044 | 853.3154582203827 | 457.3682604249752 | 700.506936502133 | 849.8019108786875 | 591.2246322546099 | 666.7098599484237 | 810.1553468841087 |
| SE_27_1_17 | SE_27 | SE_27_1 | CAGATGTCTTTATCTAGTGG | 346.80858934025565 | 406.75274124599576 | 377.23526152639585 | 329.2310478602993 | 415.75304898992744 | 514.5882052618348 | 334.68595056980536 | 385.7042241469235 | 301.91499446055775 | 231.09520151012498 | 401.4344579266917 | 188.03032406192781 |
| SE_27_1_18 | SE_27 | SE_27_1 | ATATCACCTCAAATATAACA | 84.53459365168732 | 73.95504386290831 | 108.62795488979984 | 97.59813671748003 | 114.54420737477592 | 36.38502461447317 | 81.7882065701132 | 49.15838150892162 | 144.74160028550267 | 67.79764586890101 | 83.33873249355297 | 23.21362025455899 |
| SE_27_1_19 | SE_27 | SE_27_1 | TCATTGTGAAGCTAAGTGGA | 1381.815473152581 | 1360.1731715867327 | 1267.9844916227546 | 1310.4176489933652 | 1353.318598242723 | 1395.6255869980064 | 1505.5486972577419 | 1311.2052914014287 | 1322.2100786816777 | 1378.552132667654 | 1143.2665556157829 | 1258.1782177970972 |
| SE_27_1_20 | SE_27 | SE_27_1 | GGAGGTCCTAAACTGACAGG | 1511.868694155177 | 1383.1591987333122 | 1364.76212416094 | 1440.5484979500052 | 1362.8639488572876 | 1354.9090118341912 | 1598.0985099555014 | 1413.3034683814965 | 1395.0248714633417 | 1410.6284382400372 | 1278.2518265560448 | 1323.1763545098624 |
| SE_27_1_6 | SE_27 | SE_27_1 | GCAAATTTAATCCACTGAAG | 126.80189047753097 | 213.8699917116538 | 220.2184903675033 | 113.21383859227684 | 149.54382629484635 | 139.4759276888138 | 139.90087965940418 | 124.78666075341643 | 233.54012806801964 | 206.3089653860106 | 120.8998513638867 | 318.02659748745816 |
| SE_27_2_12 | SE_27 | SE_27_2 | TCTCCTTGTACCAAGTCCTT | 74.78060207649263 | 98.93985597875572 | 88.87741763710896 | 68.96934994701923 | 125.15015250207 | 39.8502650539468 | 45.19874573611519 | 64.28403735782058 | 36.40739639083196 | 112.26706950334147 | 35.21354894093787 | 45.26655949639003 |
| SE_27_2_5 | SE_27 | SE_27_2 | AAACGAAAAACTGCAAATAC | 2268.3449296536096 | 2768.3171824358924 | 2757.175000475647 | 2308.5212604907942 | 2539.0632634741996 | 2756.5987696012767 | 2241.6425563887606 | 2620.519875821745 | 2430.41570540749 | 2053.6125635773565 | 2369.8718437251187 | 2488.5000912887235 |
| SE_27_2_6 | SE_27 | SE_27_2 | AAAACTGCAAATACAGGAAG | 75.86437891818092 | 129.9210230024065 | 141.2163413567398 | 105.40598765487843 | 96.51410065837601 | 122.14972549144564 | 92.54981269775968 | 85.08181415005664 | 124.31793889552377 | 48.114458358574915 | 92.72901221113639 | 94.01516203096391 |
| SE_27_2_8 | SE_27 | SE_27_2 | ACGAATAGAAAAATTGAAAA | 596.0772629285644 | 1172.2873844755602 | 1175.1569665351074 | 939.5447294669411 | 1077.5640249330772 | 1423.3475105137954 | 634.934761531142 | 1216.6699423458101 | 1439.4241353546001 | 1414.2734729641718 | 1157.351975192158 | 1474.0648861644959 |
| SE_27_3_13 | SE_27 | SE_27_3 | AGCTGGGTGGAAAGTCCAGG | 423.7567451001249 | 602.6336682342394 | 648.8051487508955 | 573.8770438987826 | 623.6295734848911 | 555.30478042565 | 441.22585123350547 | 614.4797688615203 | 670.4288847580032 | 506.6598266546904 | 573.9808477372873 | 449.1835519257164 |
| SE_27_3_16 | SE_27 | SE_27_3 | GCCAGGCCATCTCATCCACC | 4951.776389673838 | 3949.5990992731577 | 4046.8850830763613 | 4933.260483946224 | 4073.7435233936512 | 4777.700255924274 | 5177.408708010719 | 4030.987283731573 | 3948.87053048853 | 3556.824883810409 | 3587.0868521168713 | 5064.051258532043 |
| SE_27_3_7 | SE_27 | SE_27_3 | GCAAGCAGGGAAAGAGTGAG | 4013.225644771771 | 3872.6458779563477 | 3992.5711056314617 | 3914.3359366157324 | 3601.7789652290653 | 4518.67353307362 | 4199.178711007655 | 4780.652601742628 | 3829.880503259957 | 3978.91990486518 | 4003.780514584636 | 4061.222863535095 3 |
| SE_27_3_8 | SE_27 | SE_27_3 | CCAATTCCTTTATCGTTTGG | 145.22609678623206 | 139.91494784874547 | 139.2412876314707 | 126.22692348794084 | 165.45274398578746 | 34.65240439473635 | 111.92070372752333 | 110.60635839507364 | 105.67024806119521 | 139.96933340676338 | 104.46686185811568 | 170.62010887100857 |
| SE_27_3_9 | SE_27 | SE_27_3 | AAGATACTCTGCGAGCACAA | 193.9960546622055 | 211.871206742386 | 236.018920169656 | 239.44076208021767 | 265.1486281823517 | 264.22458350986466 | 205.54667703804768 | 217.43130282792256 | 190.02884945458632 | 83.83579865509266 | 225.36671322200237 | 359.8111139456643 |
| SE_27_4_1 | SE_27 | SE_27_4 | CATCCTTGATGGTGGAGCGG | 1394.8207952528408 | 1295.2126600855295 | 1303.5354586775982 | 1456.1641998248022 | 1440.2873482865343 | 1666.7806513868184 | 1491.5586092918013 | 1000.1839930084437 | 1381.7050922959643 | 1330.437674309079 | 721.8777532892263 | 1376.567681095348 |
| SE_27_4_11 | SE_27 | SE_27_4 | ATGCCCTGCTTCCTCCCAGG | 366.31657249064506 | 366.7770418606399 | 395.99827191645215 | 392.9951638490529 | 403.0259148371746 | 389.8395494407839 | 318.54354137833565 | 424.46371725972705 | 272.61148029232714 | 509.57585443399796 | 230.0618530807941 | 701.0513316876815 |
| SE_27_4_12 | SE_27 | SE_27_4 | CATCCTCCTGGGAGGAAGCA | 121.38300626908948 | 118.92770567143364 | 118.50322351614528 | 85.88636031138243 | 86.96875004381135 | 123.88234571118245 | 143.1293614976981 | 57.66656292392729 | 131.42182111812514 | 90.39686115853469 | 29.344624117448227 | 175.26283292192036 |
| SE_27_4_2 | SE_27 | SE_27_4 | CCATCCTTGATGGTGGAGCG | 783.5706565406401 | 619.6233404730157 | 630.0421383608391 | 629.8333089501378 | 621.5083844594324 | 479.9358008670984 | 671.5242223651401 | 720.3593598038129 | 544.334975306829 | 758.1672226199684 | 1011.8026395696148 | 663.9095392803871 |
| SE_27_4_5 | SE_27 | SE_27_4 | TCCACCATCAAGGATGGCTA | 122.46678311077778 | 81.9501837399795 | 96.77763253818532 | 135.33608291490566 | 104.99885676021127 | 104.82352329407746 | 92.54981269775968 | 65.22939084837677 | 80.80666028209045 | 47.38545141374802 | 52.820323411406804 | 38.30247342002233 |
| SE_27_5_12 | SE_27 | SE_27_5 | CTTCCAGAGGATTTGGGAAA | 45.51862735090855 | 85.94775367851507 | 65.1767729338799 | 61.16149900962082 | 40.30259148371746 | 45.91443582302566 | 34.43713960846872 | 53.88514896170255 | 14.207764445202717 | 96.22891671714983 | 26.997054188052367 | 24.37430126728694 |
| SE_27_5_17 | SE_27 | SE_27_5 | AATGAGATTCCTTCTTCCAG | 23.843090517142578 | 57.964764108765976 | 45.42623568118903 | 29.93009526002721 | 27.575457330964575 | 73.63635933881474 | 13.990087965940416 | 17.01636283001133 | 39.95933750213264 | 6.561062503442034 | 10.564064682281362 | 61.51609367458132 |
| SE_27_5_20 | SE_27 | SE_27_5 | TCCGTATCCAATATTGTGGG | 188.577170453764 | 159.90279754142338 | 159.97935174679614 | 192.59365645582727 | 154.84679885849337 | 147.27271867762948 | 194.7850709104012 | 147.47514452676486 | 190.02884945458632 | 205.57995844118372 | 178.4153146340852 | 19.73157721637514 |
| SE_27_5_21 | SE_27 | SE_27_5 | TTCCGTATCCAATATTGTGG | 3.2513305250648967 | 3.9975699385355847 | 4.93763431317272 | 3.9039254686992013 | 2.1211890254588135 | 0.0 | 0.0 | 0.0 | 0.0 | 0.0 | 0.0 | 1.1606810127279494 |
| SE_27_5_22 | SE_27 | SE_27_5 | TTTCCGTATCCAATATTGTG | 11.921545258571289 | 6.9957473924372735 | 3.950107450538176 | 10.410467916531204 | 15.9089176909411 | 0.0 | 30.13249715741013 | 29.305958207241734 | 9.767838056076867 | 0.0 | 2.347569929395858 | 1.1606810127279494 |
| SE_270_1_10 | SE_270 | SE_270_1 | GCAGAAAAGCCTGTAGGCCA | 811.7488544245359 | 1002.3906620877979 | 1061.591377332135 | 802.9073380624691 | 1132.7149395950064 | 1179.9143696407727 | 703.8090407480795 | 1243.1398400813832 | 1044.2706867223997 | 636.4230628338773 | 900.2930679233116 | 1291.8379671662078 |
| SE_270_1_11 | SE_270 | SE_270_1 | TCAAAAGCAGAAAAGCCTGT | 1473.9365046960866 | 1755.9325955017555 | 1813.0993197970229 | 1611.0199100832037 | 1673.6181410870038 | 1445.871573370374 | 1627.154846500147 | 1796.1716320567516 | 1560.1901331388233 | 2165.1506261358713 | 1901.531642810645 | 1221.0364253898028 |
| SE_270_1_12 | SE_270 | SE_270_1 | GAGTCCATATTTTAAAATAG | 2.167553683376598 | 13.991494784874547 | 16.78795666478725 | 13.013084895664004 | 11.666539640023474 | 0.8663101098684087 | 5.380803063823238 | 0.945353490556185 | 79.91867500426528 | 8.019076393095819 | 1.173784964697929 | 1.1606810127279494 |
| SE_270_1_2 | SE_270 | SE_270_1 | GCAGCTTGTGAGCAGAGGCC | 1917.2012329466008 | 1906.840860681474 | 1701.5087843193194 | 1924.6352560687062 | 1669.3757630360863 | 2046.2244795091815 | 2024.2581126103019 | 2234.8156516748213 | 1640.1088081430885 | 2274.501667859905 | 2150.374055326606 | 2173.9555368394495 |
| SE_270_1_3 | SE_270 | SE_270_1 | TTGAAGGACAAAGAACTTGA | 188.577170453764 | 206.8742443192165 | 275.51999467503776 | 169.17010364363205 | 202.5735519313167 | 259.89303296052265 | 126.9869523062284 | 202.30564697902358 | 281.49133307057883 | 90.39686115853469 | 163.15611009301213 | 188.03032406192781 |
| SE_270_2_13 | SE_270 | SE_270_2 | TGTGCTACAAAACTGCCTCA | 2502.4407274582823 | 2541.455088423998 | 2506.343177366473 | 2642.9575423093593 | 2623.9108244925524 | 2105.1335669802334 | 2500.997264065041 | 2493.842508087216 | 2259.922532065057 | 2727.2149805974054 | 2376.914553513306 | 2100.8326330375885 |
| SE_270_2_14 | SE_270 | SE_270_2 | ATTTGCATAAGCCATACAAA | 27.094421042207472 | 35.97812944682026 | 42.463655093285396 | 36.43663770785921 | 36.06021343279983 | 25.989303296052263 | 3.228481838293942 | 24.57919075446081 | 23.975602501279585 | 2.9160277793075706 | 70.42709788187574 | 190.3516860873837 |
| SE_270_2_2 | SE_270 | SE_270_2 | GTTTATTTAGAAATTCCACT | 1.083776841688299 | 0.0 | 0.0 | 2.602616979132801 | 0.0 | 0.0 | 0.0 | 0.0 | 0.0 | 0.0 | 0.0 | 0.0 |
| SE_270_2_4 | SE_270 | SE_270_2 | GTTTCTAAGAAAAGCCCCAG | 0.0 | 1.9987849692677924 | 0.0 | 0.0 | 0.0 | 0.0 | 0.0 | 0.0 | 0.0 | 0.0 | 0.0 | 0.0 |
| SE_270_2_6 | SE_270 | SE_270_2 | AAAGCCAGTTAAAACCAAAA | 69.36171786805113 | 105.935603371193 | 99.74021312608895 | 68.96934994701923 | 139.9984756802817 | 20.791442636841808 | 85.01668840840715 | 79.40969320671954 | 103.89427750554486 | 170.5876250894929 | 130.29013108147012 | 56.873369623669525 |
| SE_270_3_1 | SE_270 | SE_270_3 | GGTAGGAAATTGCTTCAACC | 414.0027535249302 | 385.76549906868394 | 439.4494538723721 | 463.26582228563854 | 470.9039636518566 | 388.97323933091553 | 462.7490634887984 | 567.212094333711 | 509.7035494716475 | 338.2592223996782 | 671.4049998072154 | 393.4708633147749 |
| SE_270_3_11 | SE_270 | SE_270_3 | AACAAAGTACCAAGTCTGGG | 48.769957875973454 | 71.95625889364052 | 54.31397744489992 | 63.76411598875362 | 123.02896347661118 | 70.17111889934111 | 65.6457973786435 | 48.213028018365435 | 5.327911666951019 | 75.08771531716994 | 61.03681816429231 | 34.820430381838484 |
| SE_270_3_14 | SE_270 | SE_270_3 | GTACTTTGTTATGGCACCAC | 198.3311620289587 | 284.8268581206604 | 242.93160820809783 | 192.59365645582727 | 178.17987813854035 | 226.10693867565467 | 208.7751588763416 | 176.7811027340066 | 280.60334779275365 | 391.47672937204135 | 111.50957164630326 | 96.3365240564198 |
| SE_270_3_2 | SE_270 | SE_270_3 | GTAGGAAATTGCTTCAACCG | 658.9363197464858 | 1055.3584637733943 | 1008.2649267498695 | 787.2916361876722 | 860.1421498235488 | 647.9999621815697 | 640.3155645949653 | 732.6489551810433 | 1169.4766108957485 | 809.9267157026777 | 705.4447637834553 | 829.8869241004838 |
| SE_270_3_7 | SE_270 | SE_270_3 | CCTCCTCTTTCGATCTTGGT | 171.23674098675122 | 136.91677039484378 | 143.19139508200888 | 139.24000838360485 | 130.45312506571702 | 110.88769406315632 | 225.99372868057597 | 130.45878169675353 | 207.78855501108973 | 243.48831957218215 | 157.2871852695225 | 91.69380000550801 |
| SE_271_1_0 | SE_271 | SE_271_1 | TTGTAGGAGTCTCAACAGGC | 48.769957875973454 | 85.94775367851507 | 54.31397744489992 | 45.545797134824014 | 92.27172260745839 | 62.37432791052543 | 39.817942672291956 | 55.775855942814914 | 63.93494000341222 | 64.15261114476655 | 21.128129364562724 | 176.42351393464833 |
| SE_271_1_1 | SE_271 | SE_271_1 | GGCAGGTGCGTAGCTTTCAA | 876.7754649258338 | 994.3955222107267 | 1000.3647118487931 | 1017.6232388409252 | 979.9893297619718 | 987.593525249986 | 993.2962455817697 | 1195.872165553574 | 904.856998103848 | 1263.369035385005 | 982.4580154521666 | 1031.845420315147 |
| SE_271_1_10 | SE_271 | SE_271_1 | CGTGTGTGTCATCCTCCACC | 40186.445289802126 | 32948.970825894925 | 32748.36581868675 | 36089.18834114498 | 32581.463431047374 | 32447.64516512125 | 39533.836270522086 | 33123.29560210761 | 30613.29245302273 | 33554.00264954739 | 34261.60933456785 | 32106.758174080536 |
| SE_271_1_2 | SE_271 | SE_271_1 | GCAGGTGCGTAGCTTTCAAC | 5998.704818744734 | 5606.591838796157 | 5399.796884885687 | 5596.927813625089 | 5431.304499687292 | 5469.015723599265 | 5475.505197746526 | 5182.427835229007 | 5808.311702254436 | 4893.82362062293 | 5223.343092905784 | 5284.580650950354 |
| SE_271_1_4 | SE_271 | SE_271_1 | GTCTGTCAAGACGTGTGAGG | 182.0745094036342 | 277.83111072822317 | 228.11870526857967 | 196.49758192452646 | 242.87614341503414 | 122.14972549144564 | 246.44078032310426 | 236.33837263904624 | 125.20592417334893 | 158.19450702743572 | 280.53460656280504 | 226.33279748195014 |
| SE_271_2_1 | SE_271 | SE_271_2 | CTCATCCGCAGCCGCAAAGT | 459.52138087583876 | 503.6938122554837 | 475.00042092721566 | 411.21348270298256 | 554.6909301574798 | 356.9197652657844 | 449.83513613562263 | 372.46927527913687 | 443.1046536347597 | 562.7933614063611 | 241.7997027277734 | 581.5011873767027 |
| SE_271_2_2 | SE_271 | SE_271_2 | TCATCCGCAGCCGCAAAGTG | 4614.721791908777 | 4482.275293583024 | 4684.827436338277 | 4377.601758901371 | 4502.2237065363315 | 4721.390098782827 | 4466.066542973287 | 4653.975234008099 | 4669.026590804743 | 4726.152023312745 | 5017.930724083647 | 4210.950714177001 |
| SE_271_2_4 | SE_271 | SE_271_2 | GGTTCATGAGCCCCACTTTG | 15.172875783636185 | 32.97995199291857 | 27.650752153767232 | 24.724861301761607 | 13.787728665482287 | 25.989303296052263 | 15.066248578705064 | 9.45353490556185 | 7.103882222601358 | 231.82420845495187 | 0.0 | 2.321362025455899 |

| | | | | | | | | | | | | | | | |
|---|---|---|---|---|---|---|---|---|---|---|---|---|---|---|---|
| SE_271_2_52 | SE_271 | SE_271_2 | GCGGCGTGTCGTACTCGAAC | 12059.184917465702 | 8949.559699896541 | 8438.417041212178 | 10778.738219078496 | 9493.38148344092 | 9743.389805689992 | 11686.028094011306 | 9345.764607638444 | 9663.055793293497 | 7880.565073578709 | 8609.71271605931 | 11474.492491828509 |
| SE_271_2_6 | SE_271 | SE_271_2 | CGCCGTGCAACTGCTCATCC | 24095.61052125595 | 19335.24640021193 | 19262.698982549417 | 21861.98262471553 | 18880.7035516589 | 18056.501619987244 | 24041.428089162226 | 20047.16612073446 | 19125.426913798507 | 18683.71898896843 | 19556.431296832197 | 18837.85283657462 |
| SE_271_3_0 | SE_271 | SE_271_3 | GAATCCCTCCCTCACCACTG | 252.52000411337366 | 255.84447606627742 | 228.11870526857967 | 230.33160265325287 | 270.45160074599875 | 194.91977472039196 | 336.83827179533466 | 263.7536238651756 | 160.72533528635574 | 363.77446546861944 | 215.97643350441894 | 109.10401519642724 |
| SE_271_3_14 | SE_271 | SE_271_3 | GTAGCACCAAGAGGGTGAGG | 963.4776122608978 | 950.4222528868353 | 967.7763253818531 | 961.66697378967 | 843.1726376198784 | 763.2192067940681 | 859.8523295989534 | 838.5285461233361 | 816.058470321331 | 815.029764351466 | 502.3799648907136 | 523.4671367403052 |
| SE_271_3_16 | SE_271 | SE_271_3 | CAGGCTTTGTAGCACCAAGA | 1257.1811363584268 | 1308.2047623857702 | 1373.6498659246508 | 1454.8628913352356 | 1499.6806409993812 | 1393.8929667782697 | 1350.5815690196325 | 1754.5760784722793 | 1236.9634920104616 | 1573.1969869364343 | 1834.6258998228632 | 1551.8305140172683 |
| SE_271_3_20 | SE_271 | SE_271_3 | AGATCACTATGATCGCTGTG | 644.8472208045379 | 557.661006425714 | 573.7531071906701 | 546.5495656178882 | 626.8113570230794 | 749.3582450361736 | 612.3353886630844 | 675.9277457476722 | 452.87249169083657 | 468.0224585788651 | 734.7893879009035 | 1089.8794709515446 |
| SE_271_3_27 | SE_271 | SE_271_3 | TGTTTCTGTGTGGCCCTGCA | 24.926867358830876 | 0.9993924846338962 | 9.87526862634544 | 24.742861301761607 | 4.242378050917627 | 34.65240439473635 | 24.75169409358689 | 3.78141396224274 | 26.639558334755094 | 0.0 | 12.911634611677218 | 1.1606810127279494 |
| SE_271_4_14 | SE_271 | SE_271_4 | CTTAGGTCAGAGCCCTAAAA | 26.010644200519174 | 43.97326932389143 | 42.463655093285396 | 26.02616979132801 | 46.6661585600939 | 30.320853845394307 | 19.370891029763655 | 48.213028018365435 | 99.45435111641902 | 26.244250013768134 | 17.606774470468935 | 44.10587848366208 |
| SE_271_4_15 | SE_271 | SE_271_4 | GTGCGGGCCGAGGCTCCTGA | 348.97614302363223 | 683.584459489585 | 608.3165473828791 | 471.07367322303696 | 634.2355186121853 | 473.8716300980196 | 371.2754114038034 | 618.261182823745 | 650.8932086458494 | 602.1597364270133 | 630.3225260427879 | 564.0909721857835 |
| SE_271_4_22 | SE_271 | SE_271_4 | CTGCTGCCCCATGTTCTCCA | 154.98008836142674 | 164.89975996459287 | 175.77978154894885 | 154.85571025840164 | 196.20998485494025 | 181.92512307236584 | 122.6823098551698 | 302.5131169779792 | 193.580790565887 | 163.29755564122397 | 280.53460656280504 | 293.6522962201712 |
| SE_271_4_29 | SE_271 | SE_271_4 | GAGCGCACACGCATGAGCAG | 7197.362005651993 | 6192.235834791621 | 6122.666548334173 | 6869.607516421028 | 6628.715704558792 | 6443.614597201225 | 7495.458667905769 | 6765.894931910616 | 6059.611535878958 | 6274.562774125665 | 6438.21053136814 | 5861.439114276144 |
| SE_271_4_32 | SE_271 | SE_271_4 | GCAAGGACCATCCCTGGGCG | 6278.319243900316 | 6266.190878654529 | 6385.348693794962 | 6509.145064811135 | 6475.990094725757 | 7206.833803995292 | 6462.344479651708 | 7427.642375299945 | 6606.6104670192635 | 6315.3871630353715 | 8103.811396274502 | 6918.819516871306 |
| SE_271_5_16 | SE_271 | SE_271_5 | CTCACCGAGAGCTCAGACCG | 592.8259324034995 | 576.649463633758 | 552.0275162127101 | 588.191437284013 | 550.4485521065621 | 565.700501744071 | 619.868512952437 | 682.5452201815656 | 457.3124180799624 | 748.6901323372188 | 484.7731904202447 | 401.59563040387053 |
| SE_271_5_20 | SE_271 | SE_271_5 | CGAGGTCTCCATGAGCCCCG | 675.1929723718102 | 645.6075450734969 | 649.7926756135299 | 782.0864022294066 | 609.8418448194088 | 572.6309826230182 | 702.7328801353148 | 612.5890618804079 | 778.7630886526739 | 809.9267157026777 | 523.5080942552763 | 685.9624785222181 |
| SE_271_5_33 | SE_271 | SE_271_5 | GCTCTTCTGCACTCTAGCTT | 99.7074694353235 | 154.90583511825392 | 123.440857829318 | 83.28374333224963 | 102.87766773475245 | 57.176467251314975 | 102.23525821264151 | 69.0108048106015 | 119.87801250639792 | 83.83579865509266 | 51.646538446708874 | 51.069964560029774 |
| SE_271_5_4 | SE_271 | SE_271_5 | ACATTCTTCGTGGTTGTAAT | 285.0333093640226 | 270.83536333578587 | 288.35784388928687 | 265.4669318715457 | 302.2694361278809 | 487.7325918559141 | 256.1262258379861 | 305.34917744964775 | 348.09022890746655 | 191.72882648947277 | 399.0868879972959 | 637.2138759876442 |
| SE_271_5_5 | SE_271 | SE_271_5 | CTAGCATCTTAGAGCTGGAG | 274.19554094713965 | 330.79891241381966 | 367.3599929000504 | 258.9603894237137 | 281.0575458732928 | 345.6577338374951 | 223.84140745504666 | 312.91200537409725 | 459.97637391343795 | 205.57995844118372 | 271.1443268452216 | 630.2497899112766 |
| SE_271_6_1 | SE_271 | SE_271_6 | TCGTGCAGACTGGAGTACAG | 17687.23805635304 | 16251.121192631786 | 16014.7231313444 | 18272.973810491396 | 17179.50991719093 | 27179.613387011457 | 20922.714633370277 | 26857.492666701215 | 18572.212085713425 | 36206.12991482762 | 38872.23667590132 | 33482.16517416316 |
| SE_271_6_10 | SE_271 | SE_271_6 | GTGAAGCTTGGGATGGACAA | 794.4084249575232 | 775.5285680759034 | 778.1711677560207 | 799.0034125937699 | 799.6882625979727 | 859.3796289894615 | 747.93162587143 | 607.8622944276269 | 982.1117172746377 | 723.1748892682755 | 912.0309175702909 | 955.2404734751424 |
| SE_271_6_17 | SE_271 | SE_271_6 | GCAGATTTTGTCCAAATGGA | 364.1490188072684 | 392.7612464611212 | 352.5470899605322 | 361.7637600994593 | 389.2381861716923 | 280.68447559736444 | 410.01719346333067 | 468.89533131586774 | 289.48320057100534 | 371.06453491688836 | 534.0721589375577 | 221.69007343103834 |
| SE_271_6_22 | SE_271 | SE_271_6 | GGCTATTGCAAGAAGCAGGG | 83.45081680999901 | 100.93864094802352 | 100.72773998872348 | 92.39290275921444 | 68.93864332741144 | 80.566840217762 | 100.08293698711222 | 138.96696311175918 | 99.45435111641902 | 54.67552086201695 | 146.72312058724114 | 104.46129114551545 |
| SE_271_6_24 | SE_271 | SE_271_6 | TGGGCAATAGGAAATTGAGA | 149.56120415298525 | 192.88274953434197 | 161.95440547206522 | 114.51514708184324 | 193.02820131675202 | 213.97859713749696 | 182.94730416999008 | 204.19635396013595 | 222.88430473411762 | 68.5266528137279 | 258.2326922335444 | 193.83372912556754 |
| SE_272_1_10 | SE_272 | SE_272_1 | CTGCCGGGCTGCTAGCAAAG | 324.04927566480137 | 385.76549906868394 | 405.87354054279757 | 316.2179629646353 | 352.11737822616305 | 267.67898239493383 | 411.09335407609535 | 265.64433084628797 | 395.15344863220054 | 596.3276808683981 | 292.2724562097843 | 441.05878483662078 |
| SE_272_1_13 | SE_272 | SE_272_1 | GTACTTCAGTGGAAGCCTGC | 243.84978937986725 | 371.7740042838094 | 345.63440192209043 | 208.20935833062407 | 290.60289648785744 | 257.2941026309174 | 256.1262258379861 | 220.2673632995911 | 285.9312594597047 | 371.06453491688836 | 199.54344399864794 | 347.04362280565687 |
| SE_272_1_16 | SE_272 | SE_272_1 | ACTTTTCTATTGCATTGACA | 24.926867358830876 | 24.984812115847404 | 31.60085960430541 | 19.519627343496005 | 7.4241615891058474 | 56.31015714144657 | 22.599372868057596 | 0.945353490556185 | 11.543808611727208 | 2.187020834480678 | 11.73784964697929 | 61.51609367458132 |
| SE_272_1_8 | SE_272 | SE_272_1 | AGAGGAGGACTGCAAGACAT | 6697.740881633687 | 6789.872540602691 | 6914.663092167078 | 6736.874050485255 | 7288.405491476483 | 6205.379316987412 | 6827.162927378924 | 6711.064429458357 | 7085.23453176703 | 8172.896858454294 | 7690.639088700831 | 6626.3279016638635 |
| SE_272_1_9 | SE_272 | SE_272_1 | CCGGGCTGCTAGCAAAGAGG | 509.3751155935005 | 540.6713341869379 | 503.63869994361744 | 480.1828326500018 | 478.32812524096244 | 330.0641518598637 | 447.6828149100933 | 504.8187639570028 | 531.0151961394515 | 373.25155575136904 | 629.14874107809 | 945.9550253732788 |
| SE_272_2_1 | SE_272 | SE_272_2 | GCTTCCTCCCCATTGCCCAG | 379.3218949509504 | 408.75152621526354 | 367.3599929000504 | 334.4362818185649 | 433.78315570632736 | 220.04276790657582 | 351.90452037403975 | 397.9938195241539 | 426.2329333560815 | 351.38134740656227 | 597.4565470312459 | 111.42537722188314 |
| SE_272_2_14 | SE_272 | SE_272_2 | GCTTCTCAGCCTTTAGAGAA | 2.167553683376598 | 7.9951398770711695 | 2.962580587903632 | 0.0 | 5.3029725636470335 | 11.262031428289314 | 0.0 | 0.0 | 5.327911666951019 | 0.0 | 0.0 | 0.0 |
| SE_272_2_20 | SE_272 | SE_272_2 | CCATGGCCCAAGTTACCGCC | 292.6197472558407 | 328.80012744455183 | 344.64687505945585 | 353.95590916206095 | 313.9359757679044 | 328.3315316401269 | 266.8878319656326 | 315.7480658457658 | 339.21037612921486 | 223.80513206185606 | 313.4005855743471 | 511.8603266130257 |
| SE_272_2_4 | SE_272 | SE_272_2 | AGCTGGACTTCATCTCCAGA | 167.98541046168634 | 225.86270152250163 | 298.2331126156393 | 266.7682403611121 | 247.11852146557177 | 226.10693867565467 | 246.44078032310426 | 93.58999556506231 | 292.1471564044809 | 347.7363126824278 | 107.98821675220947 | 167.1380658328247 |
| SE_272_2_8 | SE_272 | SE_272_2 | GGCATTTGCACAAAGACCGG | 121.38300626908948 | 230.85966395043002 | 214.29332919169605 | 196.49758192452646 | 242.87614341503414 | 200.9839454894708 | 129.1392735317577 | 206.08706094124832 | 142.96562972985234 | 143.61436813089784 | 159.63475519891836 | 395.79222534023074 |
| SE_272_3_0 | SE_272 | SE_272_3 | TCTGGGTTTTTAGGAGGTGT | 2508.9433885084118 | 3375.947813093301 | 3390.1797194243895 | 2860.276060066948 | 3540.2644834907594 | 3356.0853656302156 | 2608.6133253415055 | 3381.529435719474 | 3450.7107896286097 | 2739.6080986594625 | 3793.673005903707 | 2944.6477292908075 |
| SE_272_3_2 | SE_272 | SE_272_3 | TCTTCTTCTGGGTTTTTAGG | 7.586437891818092 | 18.98845720804403 | 14.81290293951816 | 16.917010364363207 | 19.090701229129323 | 56.31015714144657 | 3.228481838293942 | 27.415251226129364 | 55.05508722516053 | 37.908361130998415 | 8.216494752885502 | 146.24580760372163 |
| SE_272_3_21 | SE_272 | SE_272_3 | CTAGAAGGAAGCTCATACCG | 645.9309976462262 | 678.5874970664155 | 736.6950395253698 | 624.6280749918722 | 775.2945888051963 | 776.2138584420942 | 797.4350140586038 | 605.0262339559584 | 824.9383230995827 | 1144.5409033782214 | 670.2312148425175 | 831.0476051132118 |
| SE_272_3_24 | SE_272 | SE_272_3 | GGGAGCTTGAAAGACCCTGC | 303.4575156727237 | 405.75334876136185 | 333.7840795704759 | 361.7637600994593 | 363.7839178661865 | 305.8074687835483 | 347.59987792298114 | 312.91200537409725 | 406.69725724392777 | 437.40416686910356 | 277.01325166871123 | 319.1872785001861 |
| SE_272_3_25 | SE_272 | SE_272_3 | CCTCATTCAGGCAGCCTGCA | 179.906955572025763 | 64.96051150120326 | 108.62795488979984 | 166.56748666449926 | 57.27210368738796 | 556.1710905355184 | 259.35470767628004 | 80.35504669727572 | 302.8029797383829 | 10.206097227576498 | 69.25331291717781 | 141.60308355280984 |
| SE_272_4_10 | SE_272 | SE_272_4 | AATATGAGCAGATAGTAAGG | 99.7074694353235 | 105.935603371193 | 110.60300861506893 | 111.91253010271043 | 97.57469517110542 | 115.21924461249836 | 95.77829453605362 | 46.32232103725307 | 72.81479278166393 | 199.74790288256858 | 32.86597901154201 | 136.96035950189804 |
| SE_272_4_16 | SE_272 | SE_272_4 | AGTGGCTTCCCATTGCATTG | 387.992109324411 | 445.7290481467177 | 502.6511730809829 | 527.0299382743922 | 439.0861282699744 | 376.8448977927578 | 379.88469630592056 | 427.29977773139564 | 399.5933750213264 | 801.178632364755 | 359.1781991975663 | 429.4519747093413 |
| SE_272_4_18 | SE_272 | SE_272_4 | TTAAGCTGAGACCTGAATGA | 213.50403781259487 | 283.8274656360265 | 280.4526789882105 | 231.63291114281927 | 301.2088416151515 | 740.6951439374894 | 248.59310154863357 | 264.6989773557318 | 365.84993446396993 | 160.3815278619164 | 217.15021846911688 | 521.1457747148493 |
| SE_272_4_6 | SE_272 | SE_272_4 | ACAGGGCTAGGGAATCAGAG | 3835.48624273489 | 3525.8566857883857 | 3607.4356292039893 | 3773.7946197425613 | 3630.4150170727594 | 4180.81259022494 | 3953.814091297315 | 3462.8298359073056 | 3379.6719674025962 | 3734.7025783481713 | 3044.798198426428 | 2929.5588761253443 |
| SE_272_4_7 | SE_272 | SE_272_4 | GCATCACCTATGTCACACCC | 8528.239967245225 | 6833.845809926582 | 6945.276424908749 | 7771.414299690543 | 6984.014866323143 | 7593.208112996603 | 8213.25779661979 | 7114.730369925848 | 7558.530684847845 | 7012.31780228988 | 6873.684753271073 | 6339.639691520059 |
| SE_272_5_13 | SE_272 | SE_272_5 | GCATAGTAAGTGGGAGGCAG | 6525.420363805248 | 6128.274175712753 | 6087.1155812793295 | 5991.224285693707 | 6138.721039677806 | 5888.309816775574 | 6701.2521356854595 | 6956.856337002965 | 5310.151961394516 | 6613.55100346957 | 7808.0175851706235 | 5700.1044535069595 |
| SE_272_5_23 | SE_272 | SE_272_5 | AATTAAAAGTACAAAAGCAA | 1.083776841688299 | 5.9963549078033775 | 10.862795488979984 | 1.3013084895664004 | 6.363567076376441 | 0.8663101098684087 | 0.0 | 0.0 | 0.8879852778251698 | 0.0 | 1.173784964697929 | 0.0 |
| SE_272_5_26 | SE_272 | SE_272_5 | CTGTCTGAGTCAGAGACAGA | 59.60772629285644 | 90.94471610168455 | 84.92731018657079 | 67.66804145745283 | 84.84756101835254 | 142.07485801841904 | 102.23525821264151 | 114.38777235729839 | 84.35860139339113 | 205.57995844118372 | 106.81443178751154 | 184.54828102374395 |
| SE_272_5_3 | SE_272 | SE_272_5 | TGAGTTGGTAAAGGATGCAT | 1638.670584632708 | 1600.0273678988679 | 1631.3943770722667 | 1280.487553733338 | 1722.4054886725567 | 1554.1603371039253 | 1517.386463998153 | 1537.1447756443567 | 1423.4404003537472 | 1572.4679799916075 | 1901.531642810645 | 1788.6094406137702 |
| SE_272_5_5 | SE_272 | SE_272_5 | GGCTCCTGTTAGGATGGCAT | 35.76463577571386 | 63.961119016569356 | 56.28903117016901 | 33.834020728726415 | 90.15053358199957 | 32.05347406513112 | 39.817942672291956 | 12.289595377230405 | 43.51127861343332 | 18.954180565499207 | 25.823269223354437 | 22.05293924183104 |
| SE_272_6_18 | SE_272 | SE_272_6 | TGTGTTAAGAGTAATTGCAA | 235.17957464636086 | 325.80194999065014 | 302.1832199661705 | 277.1787082776433 | 354.2385672516219 | 330.0641518598637 | 237.8314954209871 | 225.93948424292822 | 342.76231724051553 | 260.98448626482047 | 169.0250349165018 | 153.20989368008932 |
| SE_272_6_19 | SE_272 | SE_272_6 | CCTGGTTGCATATTCAGCTG | 1541.130668880761 | 1424.1342906033021 | 1465.4898641496634 | 1575.884580864911 | 1429.6814031592403 | 1231.026666123009 | 1722.9331410362006 | 1555.1064919649243 | 1443.864061743726 | 891.5754935232898 | 1460.1884960842237 | 842.6544152404913 |
| SE_272_6_4 | SE_272 | SE_272_6 | CTAACAGGAGCCAAAGCGTG | 311.04395356454177 | 274.8329332743215 | 252.80687683444327 | 327.92973970373293 | 237.57317085138712 | 286.74864636644327 | 357.2853231437863 | 276.98857273296225 | 175.82108500938036 | 232.55321539977876 | 295.79381110381781 | 284.3668481183476 |
| SE_272_6_5 | SE_272 | SE_272_6 | GCCAAAGGTGAAAATTTCCT | 606.9150313454473 | 809.5079125534559 | 835.4477257888243 | 697.5013504075906 | 745.597942448773 | 900.0962041532766 | 519.7855759653247 | 828.129657727218 | 1109.9815972814622 | 874.8083337922712 | 727.746678112716 | 630.2497899112766 |
| SE_272_6_7 | SE_272 | SE_272_6 | CACCAAGATGGCTAACCAGA | 391.2434398494759 | 381.76792913014833 | 423.6490240702194 | 406.0082487447169 | 369.08689042983355 | 447.88232680196734 | 333.60978995704073 | 513.3269453720085 | 232.6521427901945 | 725.3619101027582 | 482.4256204908488 | 434.0946987602531 |
| SE_273_1_15 | SE_273 | SE_273_1 | CACCAGCTGCCTGCAGGGGG | 8.670214733506391 | 7.9951398770711695 | 15.800429802152705 | 9.109159426964803 | 6.363567076376441 | 60.641707690788614 | 19.370891029763655 | 21.743130282792254 | 47.95120500255917 | 0.7290069448268927 | 8.216494752885502 | 0.0 |
| SE_273_1_19 | SE_273 | SE_273_1 | TGGGGACACCAGCTGCCTGC | 203.7500462374002 | 260.8414384894469 | 196.51784566427426 | 222.52375171585447 | 179.24047265126973 | 162.86630065526083 | 181.87114355722542 | 300.62240999686685 | 194.4687758437122 | 220.8891042825485 | 204.23858385743966 | 318.02659748745816 |
| SE_273_1_20 | SE_273 | SE_273_1 | CAGCTGGTGTCCCCACAACC | 2353.963300146985 | 2131.7041697241007 | 2189.347054460784 | 2266.8793888246696 | 2363.004574361118 | 2597.1977093854894 | 2542.967527962862 | 2605.394219972846 | 2994.2863568264725 | 2238.0513206185606 | 2712.617053416914 | 3137.3207774036473 |
| SE_273_1_28 | SE_273 | SE_273_1 | GGCCCTGCCCCAGGCTTCCC | 0.0 | 0.0 | 0.987526862634544 | 0.0 | 0.0 | 0.0 | 0.0 | 0.0 | 4.439926389125849 | 0.0 | 0.0 | 0.0 |
| SE_273_1_7 | SE_273 | SE_273_1 | AAAATAGAAGCTGAGATCAA | 23.843090517142578 | 37.97691441608806 | 37.526020780112674 | 28.62878677046081 | 39.24199697098805 | 17.326202197368175 | 50.57954879993843 | 37.8141396224274 | 22.199631945629246 | 29.160277793075707 | 43.43004369382337 | 11.606810127279495 |
| SE_273_2_12 | SE_273 | SE_273_2 | ATTGGCTCACACATCTAGGG | 229.76069043791938 | 312.8098476904095 | 284.40773643874866 | 330.5323563498657 | 273.63338428418695 | 269.4224441690751 | 325.00050505492356 | 232.55695867682152 | 302.8029797383829 | 359.4004237996581 | 137.3328408696577 | 386.50677723840715 |
| SE_273_2_13 | SE_273 | SE_273_2 | CTTGGTTCCCGAATGCTGAG | 228.67691359623106 | 213.8699917116538 | 272.55741408713413 | 270.6721658298113 | 183.48285070218736 | 510.2566547124927 | 309.9342564762185 | 266.5896843368442 | 185.58892306546048 | 375.4385765858497 | 188.97937931636656 | 280.88480508016374 |
| SE_273_2_26 | SE_273 | SE_273_2 | TGTGTTGCAGACTCCCCACG | 1613.743717273877 | 1774.9210527097794 | 1952.3406074284935 | 1707.3167383111174 | 1893.161205221991 | 1715.2940175394492 | 1623.926364661853 | 1859.510315924016 | 1634.7808964761375 | 2130.1582927841805 | 1429.6700870020775 | 1493.7964633808708 |
| SE_273_2_27 | SE_273 | SE_273_2 | CCAGGAACACTGGAGACAGG | 1281.0242268755694 | 1370.1670964330717 | 1334.148791419269 | 1055.3611850383509 | 1197.4112048715 | 1198.9731920578777 | 1281.7072898026952 | 1209.1071144213606 | 1346.1856811829573 | 895.2205282474242 | 1430.8438719667754 | 1089.8794709515446 |
| SE_273_2_35 | SE_273 | SE_273_2 | GCACCTGGAGTGAGTGGGAA | 709.8738313058358 | 887.4605263548998 | 878.8989077447442 | 844.5492097285699 | 854.8391777208918 | 1069.8929856874847 | 784.5210867025438 | 866.8891508400217 | 1028.2869517215465 | 974.6822852335555 | 535.2459439022556 | 1322.0156734971345 |
| SE_273_3_0 | SE_273 | SE_273_3 | AGATCTGGCCTTGGTCCCAG | 1625.6652625324484 | 1657.9921320076337 | 1616.5814741327486 | 1600.6094421666726 | 1630.1337660650981 | 1574.951779740767 | 1625.0025252746177 | 1632.6254781905316 | 1925.152082324968 | 1758.3647509224652 | 1379.1973335200666 | 1214.0723393134351 |
| SE_273_3_1 | SE_273 | SE_273_3 | GTGCTACACAGGAAAAGTAG | 476.8618103428515 | 454.72358050842274 | 491.7883775920029 | 415.11740817168175 | 398.78353678625695 | 524.1176164703872 | 382.03701753144986 | 347.89008452467607 | 277.93939195927817 | 217.244069558414 | 328.65979011542015 | 525.7884987657611 |
| SE_273_3_2 | SE_273 | SE_273_3 | ACATTGCAGAGTTTTTTGCA | 596.0772629285644 | 1085.3402383124112 | 1009.252453612504 | 812.0164974894338 | 922.7172260745839 | 784.8769595407783 | 836.176796118131 | 1015.3096488573427 | 967.01596755161 | 1378.552132667654 | 977.7628755933749 | 831.0476051132118 |
| SE_273_3_3 | SE_273 | SE_273_3 | AATTTAGTGATAATTCCACT | 0.0 | 0.0 | 3.950107450538176 | 2.602616979132801 | 0.0 | 0.0 | 1.0761606127646475 | 0.0 | 0.0 | 5.103048613788249 | 0.0 | 0.0 |
| SE_273_3_7 | SE_273 | SE_273_3 | ATCATGATAGCCTATCCAAA | 207.0013767624651 | 212.87059922701988 | 235.0313933070215 | 270.6721658298113 | 189.84641777856382 | 256.42779252104896 | 173.26185865510826 | 232.55695867682152 | 283.26730362622914 | 220.8891042825485 | 267.6229719511278 | 345.88294179292893 |
| SE_274_1_10 | SE_274 | SE_274_1 | TCTTGGACATCCACTGCCTA | 1753.5509298516677 | 1420.1367206647665 | 1315.3857810292127 | 1525.1335497781212 | 1455.135671464746 | 1816.652300394053 | 1849.920093342429 | 1502.1666964937779 | 1703.1557628686755 | 1448.5367993710356 | 1652.689230294684 | 1397.4599393244512 |
| SE_274_1_11 | SE_274 | SE_274_1 | GGTCTGAGATGTGTTGTAGA | 74.78060207649263 | 132.9192004563082 | 149.11655625781614 | 88.48897729051524 | 147.42263726958714 | 128.21389626052448 | 76.40740350628997 | 72.79221877282625 | 82.5826308377408 | 46.65644446892113 | 95.07658214053225 | 110.26469620915519 |
| SE_274_1_2 | SE_274 | SE_274_1 | GAGGCCTCATTATCGCTGGA | 1037.174437495702 | 1171.2879919909262 | 1241.321266331622 | 1000.7062284765619 | 999.0800309911011 | 1267.4116907374819 | 938.4120543307725 | 1091.8832815923936 | 1057.5904658897773 | 1190.4683409023157 | 1477.7952705546927 | 969.1686456278378 |
| SE_274_1_28 | SE_274 | SE_274_1 | CTTCCCTCATGAATGTATTA | 44.434850509220254 | 70.95686640900662 | 53.32645058226538 | 53.35364807222242 | 58.33269820011737 | 6.064170769078861 | 76.40740350628997 | 106.82494443284891 | 100.34233639424419 | 134.13727784814824 | 76.29602270536539 | 17.410215190919242 |
| SE_274_1_7 | SE_274 | SE_274_1 | GTATCAAAGGAAGGCAAAGG | 296.9548546225939 | 433.73633833111097 | 430.5617121086612 | 372.17422801599054 | 357.4203507898101 | 265.0908936197331 | 259.35470767628004 | 410.2834149013493 | 364.9619491861448 | 300.35086126867975 | 293.4462411744823 | 196.15509115102344 |
| SE_274_2_2 | SE_274 | SE_274_2 | TTATCTTTAGTGTCAGCTAG | 11.921545258571289 | 18.98845720804403 | 6.912688038441808 | 9.109159426964803 | 4.242378050917627 | 0.8663101098684087 | 15.066248578705064 | 13.23494886778659 | 7.991867500426528 | 6.561062503442034 | 0.0 | 0.0 |
| SE_274_2_34 | SE_274 | SE_274_2 | CCAGAGCCTAAGACCCAGGA | 204.8338230790885 | 228.86087898116222 | 228.11870526857967 | 223.82506020542087 | 226.90621503143404 | 236.50265994077557 | 165.7287343657557 | 319.5294798079905 | 289.48320057100534 | 104.24799311024564 | 224.19292825730446 | 169.4594278582806 |
| SE_274_2_35 | SE_274 | SE_274_2 | GCAGCCAGAGCCTAAGACCC | 5251.982574821496 | 5162.861575618707 | 5015.648935320849 | 5214.343117692567 | 5517.212655218374 | 5173.603976134137 | 5489.495285712467 | 5676.847710789891 | 5180.506110832041 | 6889.844635558962 | 6197.584613605065 | 5024.588104099293 |
| SE_274_2_40 | SE_274 | SE_274_2 | GCAGAGGACCTGGAGCCTGC | 8073.0536937361385 | 7689.3257767731975 | 7715.547377763693 | 7858.601968491493 | 7869.611284452198 | 8076.609154303174 | 8182.049138869615 | 8365.433037931682 | 7336.5343653915525 | 8377.018803005823 | 9272.90122111364 | 7983.164005542836 |
| SE_274_2_44 | SE_274 | SE_274_2 | CAGCAGTTAGGTCCTTGCAG | 1481.5229425879047 | 1454.116065142319 | 1568.192657863656 | 1604.5133676353717 | 1442.4085371311993 | 1576.684399960504 | 1472.1877182620378 | 1647.7511340394306 | 1683.620086756522 | 2089.3339038738745 | 1302.9013108147012 | 1400.941982362635 |
| SE_275_1_13 | SE_275 | SE_275_1 | CTACACAGCCTTGCCAAGAG | 158.23141888649164 | 193.88214201897586 | 219.23096350486878 | 183.48449702886245 | 240.75495438957532 | 219.1764577967074 | 139.90087965940418 | 283.6060471668555 | 269.9475244588516 | 201.93492371704926 | 167.85124995180385 | 169.4594278582806 |
| SE_275_1_19 | SE_275 | SE_275_1 | CTGCCAGGACACTAATCAGG | 650.2661050129793 | 669.5929647047104 | 697.7883775920029 | 664.9686381684306 | 669.2353755322556 | 526.7165467999926 | 623.0969947907308 | 704.2883504643578 | 548.7749016595549 | 691.0985836958943 | 771.1767218065394 | 718.4615468786008 |
| SE_275_1_3 | SE_275 | SE_275_1 | TTTTGGGTTGATGATTACAG | 19.50798315038938 | 22.986027146579612 | 16.78795666478725 | 33.834020728726415 | 26.51486281823517 | 0.0 | 30.13249715741013 | 5.67212094333711 | 5.327911666951019 | 16.038152786191638 | 12.911634611677218 | 0.0 |
| SE_275_1_8 | SE_275 | SE_275_1 | GGCAGGTATCAGATGCCAGT | 1788.2317887856932 | 2029.7661362914432 | 1868.4008241045574 | 2026.1373182548855 | 1937.706174756626 | 1375.700454471033 | 1771.3603686106098 | 1960.6631394135277 | 1894.9605828789124 | 1403.3383687917683 | 2022.4314941745317 | 1332.461802611686 |

| | | | | | | | | | | | | | | | |
|---|---|---|---|---|---|---|---|---|---|---|---|---|---|---|---|
| SE_275_1_9 | SE_275 | SE_275_1 | GCATGGTAAAGAGTCTGGC | 2709.4421042207473 | 2951.2060071238952 | 2903.3289761455594 | 2917.53363360787 | 2837.090321551163 | 2954.983784761142 | 2943.299275911311 | 2806.7545134613133 | 3207.4028235045134 | 3469.344050431182 | 2851.1236792512696 | 2951.6118153671755 |
| SE_276_1_1 | SE_276 | SE_276_1 | GCAGGATTCCTTCGACTGTG | 5020.054330700201 | 4591.209074408119 | 4528.798192042019 | 4981.408898060181 | 4899.946648809859 | 4478.823268019673 | 5547.607958801757 | 4125.5226327871915 | 4574.900151355275 | 4691.888696905881 | 4440.428521452265 | 4201.665266075177 |
| SE_276_1_16 | SE_276 | SE_276_1 | AGCGCGGGCTCACTGAGGAG | 849.6810438836263 | 1294.2132676008955 | 1244.2838469195256 | 651.9555532727666 | 896.2023632563487 | 891.4331030545926 | 552.0703943482641 | 562.4853268809301 | 1030.950907555022 | 195.37386121360723 | 163.15611009301213 | 185.70896203647192 |
| SE_276_1_26 | SE_276 | SE_276_1 | ACACACTGCTCCACCAGCAG | 3076.8424535530808 | 3103.1136647882477 | 3164.036067881079 | 3309.2274889673563 | 3205.116617468267 | 3296.309968049295 | 3325.3362934427605 | 3043.0928861003595 | 2740.3225673684738 | 2663.7913763974657 | 2489.5979101243074 | 3014.2885900544848 |
| SE_276_1_32 | SE_276 | SE_276_1 | AGGAAACAGTCCACCACGAG | 3246.9954176981437 | 4024.5535356207 | 4007.38400857098 | 3415.9347851118014 | 3625.1120445091124 | 3622.9088794696854 | 3030.4682855452475 | 3926.0530462798365 | 4542.932681353569 | 2891.9705501282833 | 3841.798189456322 | 4032.205838216896 |
| SE_276_1_9 | SE_276 | SE_276_1 | AGGCCAGGCCAACCTGGTGA | 1954.049645564003 | 2152.6914119014123 | 1975.053725269088 | 1846.5567466947223 | 2075.583461411449 | 1885.0907990736573 | 1942.4699060401888 | 2144.0617165814274 | 1877.2008773224088 | 1734.3075217431776 | 2229.0176479613674 | 1671.3806583282471 |
| SE_276_2_17 | SE_276 | SE_276_2 | TCTTCTTGAATCTCCCAGGA | 27.094421042207472 | 44.97266180852533 | 43.451181955919935 | 48.14841411395682 | 39.24199697098805 | 6.93048087894727 | 58.112673089290965 | 132.3494886778659 | 61.27098416993672 | 85.29381254474644 | 107.98821675220947 | 27.856344305470785 |
| SE_276_2_18 | SE_276 | SE_276_2 | GGTTCCTCTGGAATCATGGG | 1.083776841688299 | 0.9993924846338962 | 1.975053725269088 | 3.9039254686992013 | 16.969512203670508 | 0.0 | 1.0761606127646475 | 0.0 | 1.7759705556503396 | 0.0 | 0.0 | 16.249534178191293 |
| SE_276_2_24 | SE_276 | SE_276_2 | ATATAGAGCAAGCAGTGCTG | 300.2061851476588 | 303.81531532870446 | 347.6094556473595 | 390.3925468699201 | 307.572408691528 | 357.7860753756528 | 368.04692956550946 | 396.1031125430415 | 324.11462640618697 | 594.1406600339175 | 582.1973424901728 | 135.79967848917008 |
| SE_276_2_4 | SE_276 | SE_276_2 | TAGTCGGGATGATTTACAGG | 854.0161512503795 | 1132.3116850902043 | 1173.1819128098384 | 968.1735162374019 | 1006.504192580207 | 920.8876467901184 | 841.5575991819543 | 1148.6044910257647 | 1008.7512756093929 | 1102.9875075230887 | 946.0706815465307 | 969.1686456278378 |
| SE_276_2_5 | SE_276 | SE_276_2 | GCTCCCTTCCTCTTCTCTCA | 88.86970101844051 | 107.93438834046079 | 113.56558920297256 | 100.20075369661284 | 134.69550311663465 | 116.95186483223517 | 109.76838250199404 | 110.60635839507364 | 62.15896944776188 | 56.13353475167074 | 131.46391604616804 | 156.69193671827318 |
| SE_276_3_11 | SE_276 | SE_276_3 | GATAGTCCAGGGATGTCGGG | 751.0573512899912 | 911.4459459861133 | 1029.9905177278295 | 719.6235947302195 | 934.3837657146073 | 917.4224063506448 | 721.0276105523138 | 758.1734994260604 | 1138.3971261718677 | 833.2549379371383 | 759.4388721595601 | 1187.3766760206922 |
| SE_276_3_13 | SE_276 | SE_276_3 | GAGGATAAAGAACCCCAGGA | 151.72875783636184 | 357.7825094989348 | 333.7840795704759 | 238.13945359065127 | 303.33003064061035 | 239.9679004335492 | 149.586325174286 | 158.81938641343908 | 341.87433196269035 | 72.90069448268926 | 491.81590020843225 | 146.24580760372163 |
| SE_276_3_37 | SE_276 | SE_276_3 | TGAGTGTGCGAGATAGGGAG | 2051.58956131595 | 2325.5863117430763 | 2424.3784477678055 | 2225.237517158545 | 2407.5495438957532 | 1918.8768933585254 | 2080.2184644740637 | 2066.5427303558204 | 1956.231567048849 | 2243.154369232349 | 2110.4653665268766 | 1456.6546709735765 |
| SE_276_3_8 | SE_276 | SE_276_3 | CATCTCTCCGCCAAGAACTG | 332.71949039830776 | 413.748488638433 | 389.08558387801037 | 443.74619494214255 | 366.9657014043747 | 468.67376943880913 | 418.6264783654479 | 374.35998226024924 | 304.57895029403323 | 468.0224585788651 | 516.4653844670888 | 237.93960760922963 |
| SE_276_3_9 | SE_276 | SE_276_3 | ACAGTGGAGCCCAACCCCAG | 1905.2796876880295 | 2382.5516833672086 | 2415.4907060040946 | 2085.99750877494 | 2298.3083090846244 | 2459.4544019164123 | 2080.2184644740637 | 2363.3837263904625 | 2452.615337353119 | 2926.962883479974 | 2347.5699293958583 | 2511.7137115432824 |
| SE_276_4_10 | SE_276 | SE_276_4 | CAAACATGTGAAATAAAGGC | 477.9455871845398 | 506.6919897093854 | 510.5513879820593 | 426.82918457777936 | 458.1768294991037 | 349.98928438683714 | 464.9013847143277 | 428.2451312219518 | 500.82369669339573 | 439.5911877306163 | 450.73342644400475 | 345.88294179292893 |
| SE_276_4_19 | SE_276 | SE_276_4 | AAGCCAGGCACTCACCACAG | 2272.6800370203628 | 2340.577199012585 | 2441.166404432593 | 2692.4072649128825 | 2649.365092798058 | 2163.1763443414166 | 2604.308682890447 | 2764.213606386285 | 2635.540304585104 | 4021.20230766514 | 2920.3769921684475 | 2914.470022959881 |
| SE_276_4_31 | SE_276 | SE_276_4 | CCTGCTCTTTAACATGGACA | 808.4975238994709 | 867.4726766622219 | 909.5122404864151 | 813.3178059790002 | 955.5956559691955 | 783.1443393210415 | 850.1668840840715 | 997.3479325367751 | 824.0503378217576 | 1312.2125006884069 | 936.6804018289474 | 1305.766139318943 |
| SE_276_4_32 | SE_276 | SE_276_4 | AGGGCTGGAGAGGTTAACCT | 2907.773266249706 | 3153.0832890199426 | 3110.7096172988136 | 2881.0969959000104 | 3174.3593765991145 | 3215.743127831533 | 3012.173555128248 | 2979.754202233095 | 3031.5817384951297 | 3532.038647686295 | 3115.2252963083038 | 2873.846187514403 |
| SE_276_4_6 | SE_276 | SE_276_4 | CCTGGGGCACAAAATGTAAG | 707.7062776224592 | 558.660398910348 | 563.8778385643246 | 697.5013504075906 | 656.5080033795027 | 588.2245646006495 | 762.9978744501351 | 951.0256114995221 | 905.7449833816731 | 537.2781183374199 | 796.9999910298939 | 680.1590734585784 |
| SE_277_1_34 | SE_277 | SE_277_1 | TGAGTGGCAGAGGAGTCAAA | 2520.864933766983 | 2518.4690612774184 | 2476.7173714874366 | 2924.040176055702 | 2690.728278794505 | 2666.502518174962 | 2887.338924047549 | 2788.7927971407457 | 2642.6441868077054 | 2235.135292839253 | 2822.952840098519 | 3212.7650432309642 |
| SE_277_1_35 | SE_277 | SE_277_1 | GTGGCAGAGGAGTCAAAGGG | 308.8763998811652 | 360.7806869528365 | 331.8090258452068 | 338.3402072872641 | 302.2694361278809 | 276.35292504802237 | 298.0964897358073 | 350.72614499634466 | 344.5382877961659 | 271.19058347560406 | 376.7849736680352 | 181.06623798556012 |
| SE_277_1_39 | SE_277 | SE_277_1 | GTGGGCTTTTCCATTTAAAA | 19.50798315038938 | 29.981774539016886 | 43.451181955919935 | 11.711776406097604 | 51.96913112374093 | 8.663101098684088 | 11.837766740411123 | 55.775855942814914 | 54.167101947335354 | 5.832055558615141 | 78.64359263476125 | 11.606810127279495 |
| SE_277_1_41 | SE_277 | SE_277_1 | TCAGCAGCACAACTGAGGGG | 4040.3200658139785 | 4075.5225523370286 | 4278.953895795479 | 4056.17856197847 | 4245.559834455815 | 3858.5452293538924 | 3736.429647518856 | 4144.429702598315 | 4407.958919124143 | 4227.51127305115 | 3622.300401057809 | 2962.0579444817267 |
| SE_277_1_43 | SE_277 | SE_277_1 | ATAGTCCTGGCACTGTGCAA | 579.82061030324 | 584.6446035108293 | 549.0649356248065 | 540.0430231700561 | 509.0853661101152 | 714.7058406414372 | 582.2028915056743 | 617.3158293331888 | 572.7505041972345 | 312.0149723859101 | 463.645061055682 | 580.3405063639747 |
| SE_277_2_13 | SE_277 | SE_277_2 | GCAAGGCCTTTTGTTCCCCA | 1321.1239700180363 | 1299.2102300240651 | 1417.1010478805706 | 1452.2602743561029 | 1353.318598242723 | 1386.9624858993225 | 1491.5586092918013 | 1577.7949757382728 | 1411.0086064641948 | 1467.4909799365348 | 1291.1634611677218 | 1641.2029519973205 |
| SE_277_2_15 | SE_277 | SE_277_2 | ATTTGCTCAACAGTCCCAAG | 46.60240419259685 | 112.93135076363026 | 62.21419234597627 | 31.23140374959361 | 43.48437502190568 | 173.26202197368175 | 59.18883370205568 | 63.33868386726439 | 5.327911666951019 | 41.553395855132884 | 56.341678305500594 | 95.17584304369186 |
| SE_277_2_19 | SE_277 | SE_277_2 | TTATATTCTGCTTCACAAAG | 0.0 | 7.9951398770711695 | 5.925161175807264 | 10.410467916531204 | 4.242378050917627 | 12.128341538157722 | 2.152321225529295 | 0.945353490556185 | 0.0 | 0.0 | 0.0 | 3.482043038183848 |
| SE_277_2_25 | SE_277 | SE_277_2 | GAGCAGGAGAGGGTGCCAAG | 2379.9739443475046 | 2444.51401741451 | 2526.0937146191636 | 2417.831173614372 | 2372.549924975683 | 2070.481162585497 | 2495.6164610012174 | 2714.1098713868073 | 2139.156534280834 | 2539.131188832067 | 2378.088338478004 | 2240.1143545649425 |
| SE_277_2_27 | SE_277 | SE_277_2 | AGGCACTTCCCTATATGGTG | 1341.7157300101142 | 1517.0777916742545 | 1567.2051310010213 | 1514.72308185529 | 1585.588796530463 | 1091.550738434195 | 1357.0385326962205 | 1549.4343710215871 | 1299.1224614582234 | 1216.7125909160839 | 1516.5301743897244 | 1048.0949544933383 |
| SE_277_3_0 | SE_277 | SE_277_3 | CCTGGGTTGCAGCCGAGTAG | 1443.5907531288142 | 1329.192004563082 | 1321.31094220502 | 1423.6314875856422 | 1308.773628708088 | 1284.7378929348502 | 1350.5815690196325 | 1174.1290352707817 | 1275.1468589569438 | 1595.796202226068 | 780.5670015241228 | 1833.87600011016 |
| SE_277_3_1 | SE_277 | SE_277_3 | GCCTTCAGCGGAGGCCCAGA | 1.083776841688299 | 1.9987849692677924 | 1.975053725269088 | 1.3013084895664004 | 0.0 | 7.796790988815679 | 8.60928490211718 | 3.78141396222474 | 0.0 | 3.6450347241344634 | 18.780559435166865 | 3.482043038183848 |
| SE_277_3_17 | SE_277 | SE_277_3 | CTAAAAGGTAAAGGAGGCAG | 761.8951197068741 | 861.4763217544186 | 864.086004805226 | 815.920422958133 | 941.8079273037132 | 1241.4223874414297 | 1014.8194578370626 | 1091.8832815923936 | 930.608571160778 | 1176.6172089506047 | 1395.6303230258377 | 1520.4921266736137 |
| SE_277_3_34 | SE_277 | SE_277_3 | AGATCCACCACAACCTCACG | 1903.112134004653 | 1915.835393043179 | 1997.7668431096827 | 1885.5960013817141 | 1986.493522342179 | 1901.5506911611571 | 1987.668651776304 | 1758.3574924345041 | 1919.824170658017 | 2117.036167777296 | 1835.799684787561 | 2319.040663430443 |
| SE_277_3_35 | SE_277 | SE_277_3 | TGAGCACTGCACAGTCCCCA | 5147.939998019419 | 5046.932047401176 | 5174.64076020501 | 4922.850016029693 | 5124.792685508493 | 3932.181588692707 | 4327.241823926647 | 4154.828590994433 | 4713.425854696001 | 3340.309821196822 | 2178.544894479356 | 2684.655182439747 |
| SE_278_1_0 | SE_278 | SE_278_1 | AGATTGACAAGGCGCTTCCA | 41.18351998415536 | 48.970231747060915 | 61.22666548334173 | 35.13532921829281 | 38.181402458258646 | 55.44384703157816 | 35.513300221233365 | 45.37696754669688 | 42.62329333560815 | 72.90069448268926 | 31.692194046844083 | 71.96222278913287 |
| SE_278_1_12 | SE_278 | SE_278_1 | AAAATAAGTGAATGACCAGA | 42.26729682584366 | 67.95868895510495 | 59.25161175807264 | 45.545797134824014 | 61.51448173830559 | 63.24063802039384 | 52.73187002546773 | 79.40969320671954 | 26.639558334755094 | 35.72134029651774 | 164.32989505771008 | 78.92630886550056 |
| SE_278_1_18 | SE_278 | SE_278_1 | TGTGCAGGAAGTAGAGGCCT | 9116.730792281971 | 8447.864672610325 | 8788.989077447442 | 8494.941819889462 | 8633.239333617372 | 7916.341783977519 | 9194.716275461147 | 9169.928858394995 | 8411.884536837833 | 7338.912913572329 | 8690.703878623466 | 7726.653501729959 |
| SE_278_1_23 | SE_278 | SE_278_1 | TTAGCCAGAAGACAGCTCTG | 604.7474776620708 | 641.6099751349614 | 663.6180516904136 | 588.191437284013 | 566.3574697975032 | 490.33152218551936 | 581.1267308929097 | 761.0095598977289 | 638.461414756297 | 1233.4797506471023 | 491.81590020843225 | 817.1194329604764 |
| SE_278_1_26 | SE_278 | SE_278_1 | CAAAGGGTCTCTCTAAGCTG | 911.4563238598594 | 1150.3007498136144 | 1233.4210514305455 | 999.4049199869955 | 1117.8666164167946 | 1067.2940553578796 | 1013.743297224298 | 884.8508671605891 | 1317.770152292552 | 1593.6091813915873 | 861.5581640882799 | 1403.263344388091 |
| SE_278_2_10 | SE_278 | SE_278_2 | GTTACTGGAAGAGCACAGGA | 244.93356622155557 | 272.83414830505365 | 261.6946185981542 | 253.7551554654481 | 302.2694361278809 | 311.00532944275875 | 220.61292561675273 | 207.9777679223607 | 561.2066955855073 | 408.97289604788676 | 354.4830593387746 | 270.4386759656122 |
| SE_278_2_11 | SE_278 | SE_278_2 | GTAGTCCGTTTTACACATGA | 406.4163156331121 | 447.7278331159855 | 429.5741852460267 | 442.44488645257616 | 465.6009910882096 | 440.95184592302 | 404.63639039950743 | 426.3544242408394 | 387.161581131774 | 294.5188057100646 | 523.5080942552763 | 459.629681040268 |
| SE_278_2_14 | SE_278 | SE_278_2 | ATATTCAGAAGAAAGAAACG | 7.586437891818092 | 41.97448435462364 | 35.55096705484358 | 18.218318853929606 | 14.848323178211695 | 74.50266944868315 | 15.066248578705064 | 17.961716320567515 | 49.72717555820951 | 29.160277793075707 | 7.042709788187574 | 66.15881772549312 |
| SE_278_2_16 | SE_278 | SE_278_2 | ATGACCGGCCAAGTTCACCA | 3491.928983919699 | 3792.694479185636 | 3648.91175743464 | 3642.362462296355 | 3755.5651695748293 | 3377.743118376926 | 3552.4061827361015 | 3340.8792356255576 | 3506.6538621315954 | 3959.236717354854 | 3369.936633647754 | 4002.0281318859697 |
| SE_278_2_2 | SE_278 | SE_278_2 | CATTTTGTAGGGGAGCACAG | 210.25270728753 | 249.84812115847404 | 326.87139153203407 | 226.4276771845537 | 219.5430641349872 | 295.41174746512735 | 248.59310154863357 | 274.1525122612937 | 178.48504084285912 | 468.751465523692 | 226.5404981867003 | 309.9018303983625 |
| SE_278_3_19 | SE_278 | SE_278_3 | GCTCTGACAGTGCCACGCAT | 928.7967533268721 | 866.473284177588 | 847.2980481404388 | 841.946592749461 | 849.5362046962548 | 609.8823173473597 | 822.1867081521907 | 769.5177413127346 | 781.4270444861494 | 587.5795975304754 | 537.5935138316515 | 379.5426911620395 |
| SE_278_3_2 | SE_278 | SE_278_3 | TCTTCGTGCGTCCTGTCTGA | 901.7023322846646 | 965.4131401563437 | 974.6890134202949 | 925.2303360817107 | 849.5362046962548 | 643.6684116322277 | 773.7594805777816 | 681.5998666910094 | 831.1542200443589 | 772.0183545716793 | 764.1340120183518 | 1042.2915494296985 |
| SE_278_3_38 | SE_278 | SE_278_3 | ACCCTTCCTCGACAGAGCAG | 289.36841673077583 | 225.86270152726053 | 219.23096350486878 | 243.34468754891688 | 205.7553354695049 | 286.74864636644327 | 212.00364071463557 | 241.06514009182717 | 407.58524252175295 | 153.09145841364744 | 188.97937931636656 | 239.1002886219576 |
| SE_278_3_6 | SE_278 | SE_278_3 | GCACGTAGAGTACTGAGCCA | 19074.472413714062 | 17880.130942585038 | 17572.052993719077 | 18227.42801335657 | 17642.98971925368 | 17023.8599690241 | 18223.70381655654 | 17724.432594437912 | 16590.228945607647 | 15734.885897143651 | 17211.208937365733 | 18371.259069457985 |
| SE_278_3_9 | SE_278 | SE_278_3 | CAAGCTGCACACATAACCAA | 5258.485235871626 | 4648.174446032252 | 4802.343132991788 | 5206.535266755168 | 4997.521343980965 | 5171.005045804532 | 5125.752998598016 | 5170.138239851776 | 4169.978864666998 | 3995.6870645961985 | 4892.335732860968 | 5530.645025648679 |
| SE_279_1_11 | SE_279 | SE_279_1 | CACGATGGAGGACATTGGAA | 5578.199404169674 | 5745.5073941602695 | 5505.462259187583 | 5846.779043621837 | 5672.059454076867 | 5501.935507774264 | 5326.995033185005 | 5438.618631169732 | 5522.380442794731 | 4925.899926195314 | 5853.665618948572 | 5175.4766357539265 |
| SE_279_1_21 | SE_279 | SE_279_1 | TGGGAAGGTCGCTTGCGCCG | 559.2288503111622 | 639.6111901656935 | 639.9174069871846 | 541.3443316596226 | 617.2660064085147 | 611.6149375670966 | 534.8518245440298 | 577.6109827298291 | 602.0540183654651 | 435.2171460616549 | 519.9867393611826 | 422.4878886329736 |
| SE_279_1_26 | SE_279 | SE_279_1 | CTTCTGTCACGCAGGCTGGA | 209.16893044584168 | 203.8760668653148 | 140.22881449410525 | 187.38842249756166 | 214.24009157134017 | 181.92512307236584 | 253.9739046124568 | 269.4257448085127 | 234.42811334584482 | 628.4039864407815 | 140.85419576375148 | 290.17025318198733 |
| SE_279_1_4 | SE_279 | SE_279_1 | GCAAAAGGAATAAAGGCAAG | 1496.6958183715408 | 1835.8839942724674 | 1678.7956664787248 | 1365.072605555154 | 1713.9207325707214 | 2056.6202008276023 | 1383.9425480153366 | 1312.1506448919847 | 2106.3010790013027 | 1332.6246951435598 | 1408.5419576375148 | 2189.0443900049127 |
| SE_279_1_5 | SE_279 | SE_279_1 | TGTTAAGTAATACCTTAACC | 164.73407993662144 | 154.90583511825392 | 185.6550501752943 | 149.65047630013606 | 152.72560983303458 | 95.29411208552496 | 154.96712823810924 | 169.21827480955713 | 146.517570841153 | 69.98466670338169 | 331.007360044816 | 406.2383544547823 |
| SE_279_2_1 | SE_279 | SE_279_2 | TGGAGAGCGGAAGTGAGCTG | 1334.129292118296 | 1335.1883594708852 | 1256.13416927114 | 1365.072605555154 | 1198.4717993842296 | 1192.9090212887988 | 1320.4490718622224 | 1097.5554025357308 | 1258.2751386782656 | 874.8083337922712 | 1025.88805914599 | 918.098681067808 |
| SE_279_2_10 | SE_279 | SE_279_2 | ATTGTCCGCACAGAACCAGT | 1012.2475701368712 | 1009.3864094802351 | 950.0008418544313 | 1124.33053498537 | 1060.5945127294067 | 811.732572946699 | 1061.0943641859424 | 947.2441975372974 | 1072.6862156128052 | 1266.2850631643125 | 829.8659700414358 | 849.618501316859 |
| SE_279_2_3 | SE_279 | SE_279_2 | AAAGTACCAGAGGAGAGTAC | 199.414938870647 | 122.92527560996923 | 168.86709351050703 | 154.85571025840164 | 163.33155496032865 | 118.68448505197199 | 217.3844437784588 | 121.95060028174787 | 134.08577695160065 | 53.21750697236316 | 198.36965903395 | 81.24767089095646 |
| SE_279_2_5 | SE_279 | SE_279_2 | GCAGTCATGTTTTCAAACGT | 2762.547169463474 | 2531.461163577659 | 2616.9461859815415 | 2652.066701736324 | 2804.2118916565514 | 3212.2778873920597 | 2496.6926216139823 | 3025.131169779792 | 2749.2024201467257 | 2695.138675025022 | 3111.70394141421 | 2404.9310583723113 |
| SE_279_2_6 | SE_279 | SE_279_2 | TGTTCTACATTCAGCTTGAA | 74.78060207649263 | 66.95929647047105 | 78.01462214812898 | 98.89944520704643 | 104.99885676021127 | 49.379676262499295 | 109.76838250199404 | 61.44797688615203 | 79.03068972644012 | 83.83579865509266 | 104.46686185811568 | 38.30247342002233 |
| SE_28_1_38 | SE_28 | SE_28_1 | TTGCTCCCTGCGCTTCCGAG | 58.52394945116814 | 151.90765766435223 | 128.37849214249073 | 54.65495656178882 | 95.45350614564661 | 35.51871450460476 | 20.4470516425283 | 120.0598933006355 | 118.10204195074758 | 11.664111117230282 | 197.19587406925208 | 41.78451645820618 |
| SE_28_1_40 | SE_28 | SE_28_1 | CAGTCGCCACTCGGAAGCGC | 380.4056714325929 | 380.7685366455145 | 349.5845093726286 | 394.2964723386193 | 358.48094530253945 | 557.9037107552552 | 284.1064017698669 | 288.33281461963645 | 558.5427397520318 | 399.49580576513716 | 221.8453583279086 | 719.6222278913286 |
| SE_28_1_41 | SE_28 | SE_28_1 | CTTCCGAGTGGCGACTGCAG | 122.46678311077778 | 33.97934447755247 | 82.9522564613017 | 59.86019052005442 | 25.454268305505764 | 446.1497065822305 | 142.05320088493346 | 102.09817698006798 | 204.23661389978906 | 10.93510417240339 | 55.167893340802664 | 132.31763545098624 |
| SE_28_1_47 | SE_28 | SE_28_1 | AGCGGGTCCGGTGCAGGTGA | 2528.4513716588012 | 2550.4496207857032 | 2560.6571548113725 | 2689.8046479337495 | 2746.9397879691633 | 2799.914275094697 | 2838.91169647314 | 2768.940373839066 | 2657.7399365307333 | 3464.970008762221 | 2825.3004100279154 | 3313.7442913382956 |
| SE_28_1_57 | SE_28 | SE_28_1 | CACCTGTGCGAGGCCGCGTT | 1038.2582143373904 | 1040.3675765038859 | 978.6391208708332 | 1116.5226840479716 | 1000.1406255038305 | 1046.5026127210378 | 908.2795571733625 | 1070.1401513096014 | 1151.7169053392452 | 757.4382156751415 | 1305.248880744097 | 973.8113696787495 |
| SE_28_2_16 | SE_28 | SE_28_2 | AGTAGAGTGAGAAACCACAG | 151.72875783636184 | 171.89550735703014 | 202.44300684008152 | 223.82506020542087 | 255.60327756778702 | 310.1390193328903 | 227.06988929334062 | 231.61160518626534 | 226.43624584541828 | 262.44250013768135 | 193.67451917515828 | 125.35354937461854 |
| SE_28_2_17 | SE_28 | SE_28_2 | GCTTAGTGCACTGAATGAGC | 2291.104243329064 | 2226.6464557643208 | 2249.5861930814913 | 2287.700324657732 | 2255.8845285754483 | 2434.3314087302283 | 2667.802159043561 | 2340.695242617114 | 2620.444554862076 | 2646.2952097216203 | 2489.5979101243074 | 2374.7533520413845 |
| SE_28_2_26 | SE_28 | SE_28_2 | CATAGGGCAGAAGGAATCGT | 5971.610397702527 | 5323.763765644765 | 5184.516028831356 | 5491.52182597021 | 5071.762959879623 | 4793.293837901905 | 5782.2109723844505 | 5073.712183815045 | 5179.618125544215 | 5613.353475167073 | 4967.457970601636 | 4636.920645848158 |
| SE_28_2_7 | SE_28 | SE_28_2 | AGCTTTGGCATGCACTTACG | 558.1450734694739 | 662.5972173122732 | 580.6657952291118 | 593.3966712422786 | 558.9333082083973 | 803.0694718480149 | 661.8387768502582 | 731.7036016904872 | 534.5671372507522 | 858.7701810060795 | 854.5154543000923 | 428.29129369661337 |
| SE_28_2_8 | SE_28 | SE_28_2 | GCTTTGGCATGCACTTACGT | 1155.3061132397265 | 1189.2770567143364 | 1187.9948157493566 | 1280.487553733338 | 1078.6246194458067 | 1234.4919065624824 | 1074.008291539118 | 1306.4785239486478 | 840.0340728226106 | 1546.9527369226662 | 1536.4845187895892 | 1136.3067114606624 |
| SE_28_3_14 | SE_28 | SE_28_3 | GTAAAAATGTTAACTCACAT | 143.05854310285545 | 175.89307729556572 | 156.02924429625796 | 165.26617817493286 | 164.39214947305805 | 168.0641613144713 | 159.27177068916782 | 222.15807028070347 | 201.57265806631355 | 143.61436813089784 | 465.9926309850778 | 200.79781520193526 |
| SE_28_3_22 | SE_28 | SE_28_3 | GCTGTGGCCCAAGGAGAAAA | 6.5026610501297935 | 2.9981774539016888 | 9.87526862634544 | 6.506542447832002 | 5.3029725636470335 | 0.0 | 6.456963676587884 | 22.68848377334844 | 5.327911666951019 | 16.038152786191638 | 14.085419576375148 | 0.0 |
| SE_28_3_23 | SE_28 | SE_28_3 | TAGCTCTTAGTGGTGCCAAT | 548.3910818942793 | 602.6336682342394 | 639.9174069871846 | 573.8770438987826 | 571.6604423611502 | 712.9732204217004 | 491.8054000334439 | 804.4958204633134 | 634.9094736449964 | 645.1711461718 | 561.0692131256101 | 695.2479266240417 |
| SE_28_3_25 | SE_28 | SE_28_3 | AGGGAAGTCCTGAAGCTAAG | 2935.951464133602 | 3224.040155428949 | 3327.9655270784133 | 3138.7560768341577 | 3318.600230330314 | 3560.53455155916 | 3188.6638956216507 | 3610.3049804340703 | 3638.963668527546 | 2991.8445015695675 | 3461.4918608941925 | 3335.797230580127 |
| SE_28_3_9 | SE_28 | SE_28_3 | TTTGTCATGACCCAGCTCCA | 89.95347786012881 | 162.90097499532507 | 151.09160998308525 | 102.80337067574564 | 136.81669214209347 | 58.04277736118338 | 72.10276105523138 | 43.48626056558451 | 174.93309973155846 | 97.68693060680361 | 15.259204541073078 | 9.285448101823595 |
| SE_28_4_13 | SE_28 | SE_28_4 | GAATCAGGAAAATTGGTGCA | 145.22609678623206 | 234.8572338889656 | 229.10623213121423 | 191.29234796626088 | 225.90663121136365 | 309.27270922302193 | 171.10953742957895 | 240.119786601271 | 159.83735000853056 | 247.8623612411435 | 207.75993875153344 | 116.06810127279495 |
| SE_28_4_16 | SE_28 | SE_28_4 | GCAGGGATTGGCACACTGCA | 3056.250693561003 | 753.5419334139577 | 1296.6227706391562 | 2544.058097102313 | 658.6291924049616 | 6261.689474128858 | 4280.966917577768 | 1610.8823479077394 | 4911.446571651014 | 217.244069558414 | 881.5125084881447 | 1716.6472178246372 |
| SE_28_4_19 | SE_28 | SE_28_4 | AGACATACAAAGTCACCTAA | 361.98146512389184 | 409.75091869989745 | 461.1750448503321 | 365.6676855681585 | 303.33003064061035 | 512.8555850420979 | 413.2456753016246 | 349.78079150578844 | 532.7911666951019 | 398.0377918754834 | 433.1266519735358 | 542.0380329439524 |
| SE_28_4_3 | SE_28 | SE_28_4 | GATCTTAACAGTTTTAGCCT | 107.2939073271416 | 164.89975996459287 | 128.37849214249073 | 123.62430650880805 | 146.36204275665813 | 71.03742900920952 | 112.99686434028798 | 51.994441980590175 | 143.85361500767752 | 51.030486137882484 | 66.90574298778195 | 84.7297139291403 |
| SE_28_4_4 | SE_28 | SE_28_4 | TAAAGTTAAGGGAACTCGGA | 93.2048083851937 | 165.89915244922676 | 146.1539756699125 | 109.30991312357764 | 128.33193604025823 | 191.45453428091832 | 120.52998862964051 | 119.11453981007931 | 162.50130584200608 | 293.7897987652377 | 51.646538446708874 | 270.4386759656122 |

| | | | | | | | | | | | | | | | |
|---|---|---|---|---|---|---|---|---|---|---|---|---|---|---|---|
| SE_28_5_10 | SE_28 | SE_28_5 | AGTCTGTTTCCACCACATGG | 93.2048083851937 | 111.93195827899638 | 118.50322351614528 | 92.39290275921444 | 99.69588419656424 | 195.78608483026036 | 130.21543414452233 | 119.11453981007931 | 106.55823333902038 | 139.96933340676338 | 134.98527094026184 | 123.03218734916264 |
| SE_28_5_22 | SE_28 | SE_28_5 | CTCTGTGCCTGATCTCCTGG | 158.23141888649164 | 166.89854493386065 | 215.2808560543306 | 157.45832723753446 | 254.5426830550576 | 207.9144263684181 | 122.6823098551698 | 208.92312141291688 | 158.06137945288023 | 179.3357084274156 | 345.09277962119114 | 255.34982280014887 |
| SE_28_5_26 | SE_28 | SE_28_5 | GGATGCACCAGAAGAGCCAG | 1698.2783109255645 | 1790.911332463942 | 1926.6649089999953 | 1941.5522664330695 | 1826.3437509200385 | 2294.855481041415 | 1835.9300053764887 | 1776.3192087550717 | 2012.1746395518346 | 1818.1433203982704 | 1849.885104363936 | 2200.651200132192 |
| SE_28_5_28 | SE_28 | SE_28_5 | GCTGAGGCCTTGGCCCCAGA | 299.1224083059705 | 269.83597085115196 | 276.50752153767235 | 223.82506020542087 | 286.3605184369398 | 275.486614938154 | 288.41104422092553 | 306.29453094020397 | 287.70723001535504 | 248.5913681859704 | 258.2326922335444 | 423.64856964570157 |
| SE_28_5_38 | SE_28 | SE_28_5 | GCCACAGTCTTCCTACACCT | 3177.6336998300926 | 3110.109412180685 | 3159.0984335679063 | 3196.0136503750796 | 3259.206937617467 | 3608.181607601922 | 3225.2533564556484 | 3291.720854116636 | 3009.3821065495003 | 3007.153647410932 | 4962.762830742844 | 3543.5591318584297 |
| SE_280_1_0 | SE_280 | SE_280_1 | TGTACTGTTGCATCTAGCAG | 688.1982944720698 | 701.5735242129952 | 703.1191261957954 | 688.3921909806259 | 782.7187503943022 | 474.73794020788796 | 769.454838126723 | 546.4143175414749 | 748.5715892066181 | 856.5831601715988 | 766.4815819477477 | 615.1609367458132 |
| SE_280_1_10 | SE_280 | SE_280_1 | GAAGGGCAATGATTAAGCAG | 5175.034419061627 | 5801.473373299767 | 5668.404191522282 | 5585.216037218991 | 5506.60671009108 | 5055.785801192033 | 5385.107706274296 | 5434.837217207508 | 5361.655107508375 | 7147.913094027683 | 5216.3003831175965 | 4824.950969910085 |
| SE_280_1_5 | SE_280 | SE_280_1 | ACTGTGGTATATTCATCCCA | 554.893742944409 | 624.6203028961851 | 660.65547110251 | 629.8333089501378 | 672.4169210704439 | 503.32617383354545 | 558.527358024852 | 657.9660294271048 | 670.4288847580032 | 752.3351670613532 | 358.0044142328683 | 398.11358736568667 |
| SE_280_1_8 | SE_280 | SE_280_1 | ATGATTAAGCAGAGGCACGG | 793.3246481158347 | 808.508520068822 | 775.2085871681171 | 731.335371136317 | 789.0823174706786 | 862.8448694289351 | 878.1470600159523 | 851.7634949911227 | 927.0566300494772 | 869.7052851784829 | 1031.7569839694797 | 692.9265645985859 |
| SE_280_1_9 | SE_280 | SE_280_1 | GCAATGATTAAGCAGAGGCA | 1630.0003698992016 | 1812.8979671258876 | 1734.0971707862593 | 1605.8146761249382 | 1891.0400161965322 | 1921.4758236881305 | 1418.3796876238055 | 1587.2485106438346 | 1807.0500403742205 | 1840.0135287430771 | 1755.982307188102 | 1887.2673266956458 |
| SE_281_1_13 | SE_281 | SE_281_1 | CCTTGCTCTCCCTATCAGCG | 717.4602691976539 | 838.4902946078389 | 789.0339632450007 | 705.309201344989 | 758.3250766015258 | 531.914407459203 | 609.1069068247905 | 813.9493553688753 | 883.5453514360439 | 716.6138267648355 | 839.2562497590193 | 699.8906506749535 |
| SE_281_1_24 | SE_281 | SE_281_1 | GCTTGGGCCCCACTTCAGAC | 44.434850509220254 | 43.97326932389143 | 43.451181955919935 | 29.93009526002721 | 58.33269820011737 | 22.524062856578627 | 45.19874573611519 | 49.15838150892162 | 79.91867500426528 | 24.057229179287457 | 37.56111887033373 | 22.05293924183104 |
| SE_281_1_26 | SE_281 | SE_281_1 | AAAGTCCCTGGAAAATTCCA | 7.586437891818092 | 11.992709815606755 | 16.78795666478725 | 11.711776406097604 | 13.787728665482287 | 0.0 | 17.21856980423436 | 24.57919075446081 | 8.879852778251697 | 1.4580138896537853 | 9.390279717583432 | 0.0 |
| SE_281_1_6 | SE_281 | SE_281_1 | GCTTCTTCTCACGTATGCAT | 172.32051782843953 | 196.88031947287755 | 175.77978154894885 | 225.12636869498726 | 189.84641777856382 | 370.78072702367893 | 216.30828316569415 | 122.89595377230405 | 125.20592417334893 | 164.75556953087775 | 143.20176569314734 | 78.92630886550056 |
| SE_281_1_9 | SE_281 | SE_281_1 | TTACATTTTGAACTGGCGGG | 17.340429467012783 | 22.986027146579612 | 40.488601368016305 | 16.917010364363207 | 24.393673792776355 | 23.390372966447035 | 16.142409191469714 | 16.071009339455145 | 47.95120500255917 | 42.282402799959776 | 12.911634611677218 | 18.57089620364719 |
| SE_281_2_20 | SE_281 | SE_281_2 | ACAGAGCCACACAAGCCTCG | 3162.4608240464563 | 3187.062633497495 | 3092.9341337713918 | 3320.939265373454 | 2967.54344661688 | 3162.8982111295604 | 3356.5449512129353 | 3241.6171191171584 | 2924.135519878284 | 3648.679758858598 | 3263.1222018602425 | 3845.3361951676966 |
| SE_281_2_26 | SE_281 | SE_281_2 | GGGCTGGAAGGAATTCACAG | 339.22215144843756 | 369.7752193145416 | 355.50967054843585 | 379.9820789533889 | 399.8441312989863 | 397.6363404295996 | 342.2190748591579 | 481.1849266930982 | 336.5464202957393 | 347.7363126824278 | 420.2150173618586 | 437.57674179843696 |
| SE_281_2_29 | SE_281 | SE_281_2 | GGAAACCCACCCCAAAAGTG | 9.75399157519469 | 6.9957473924372735 | 5.925161175807264 | 10.410467916531204 | 12.727134152752882 | 0.0 | 3.228481838293942 | 0.0 | 0.0 | 6.561062503442034 | 0.0 | 0.0 |
| SE_281_2_3 | SE_281 | SE_281_2 | CTCTCGAGAACAGCACCAAG | 58.52394945116814 | 52.9678016855965 | 41.47612823065085 | 42.943180155691216 | 29.69664635642339 | 10.395721318420904 | 60.26499431482026 | 49.15838150892162 | 18.647690834328564 | 37.908361130998415 | 37.56111887033373 | 1.1606810127279494 |
| SE_281_2_4 | SE_281 | SE_281_2 | GGATCTCTGTTTTACAGAGG | 1.083776841688299 | 12.992102300240651 | 15.800429802152705 | 5.205233958265602 | 3.1817835381882205 | 12.994651648026132 | 9.685445514881827 | 3.78141396222474 | 5.327911666951019 | 25.515243068941242 | 1.173784964697929 | 0.0 |
| SE_281_3_0 | SE_281 | SE_281_3 | GTGCTTCCTCAATGCCCACG | 70.44549470973944 | 102.9374259172913 | 66.16429979651446 | 70.27065843658562 | 99.69588419656424 | 187.12298373157628 | 82.86436718287786 | 34.03272566002266 | 120.76599778422309 | 53.21750697236316 | 44.6038286585213 | 109.10401519642724 |
| SE_281_3_15 | SE_281 | SE_281_3 | TAGAGGTCAGCTCTTCAGAG | 832.3406144166136 | 1042.3663614731538 | 1128.7432039912837 | 1054.0598765487844 | 1201.6535829224179 | 1191.176401069062 | 965.3160696498887 | 1195.872165553574 | 1212.987889509182 | 1577.5710286053957 | 1109.2267916395429 | 1642.3636330100485 |
| SE_281_3_2 | SE_281 | SE_281_3 | TTTGCTAGCTGATTCCACGT | 423.7567451001249 | 346.78919216796197 | 397.97332564172126 | 379.9820789533889 | 406.2076983753628 | 371.64703713354737 | 385.2654993697438 | 397.0484660335977 | 490.16787335949374 | 301.80887515833354 | 458.9499211968903 | 504.896240536658 |
| SE_281_3_23 | SE_281 | SE_281_3 | TCACGAAGAATGAAATCAAG | 3093.0991061784052 | 3435.911362171335 | 3419.805525303426 | 3176.4940230315833 | 3302.6913126393724 | 3143.8393887124553 | 3032.6206067707767 | 2843.6232995930045 | 3954.198442155481 | 3391.3403073347044 | 2741.961677534362 | 3444.901245776554 |
| SE_281_3_4 | SE_281 | SE_281_3 | GTACAGCAACTTCCAGGCAG | 1751.383376168291 | 2003.7819316909618 | 1925.6773821373608 | 1841.3515127364567 | 1952.5544979348379 | 1985.5827718183928 | 1802.5690263807844 | 1902.9965764896003 | 1902.0644651015136 | 1433.227653529671 | 2435.603801748203 | 2115.9214862030517 |
| SE_282_1_12 | SE_282 | SE_282_1 | CATTCCTGTTTTCCTCCCAG | 365.2327956489567 | 389.7630690072195 | 392.048164465914 | 404.7069402551505 | 361.6627288407277 | 194.91977472039196 | 399.2555873356842 | 300.62240999686685 | 356.97008168571824 | 187.3547848205114 | 356.8306292681704 | 138.12104051462597 |
| SE_282_1_17 | SE_282 | SE_282_1 | TTTAACATATCATTAGGCCC | 34.680858934025565 | 19.987849692677923 | 18.763010390056337 | 32.53271223916001 | 15.9089176909411 | 1.7326202197368175 | 59.18883370205561 | 28.36060471668555 | 19.535676112153734 | 71.44268059303548 | 86.86008738764674 | 32.49906835638259 |
| SE_282_1_2 | SE_282 | SE_282_1 | ATACTCACATGAGCGAGGCA | 368.48412617402164 | 409.75091869989745 | 445.37461504817935 | 385.18731291165454 | 457.1162349863743 | 304.94115867367987 | 383.1131781442145 | 405.55664744860337 | 280.60334779275365 | 589.0376114201292 | 401.4344579266917 | 342.40089875474507 |
| SE_282_1_23 | SE_282 | SE_282_1 | CTGTGCTTACAGTTTCTCAG | 70.44549470973944 | 90.94471610168455 | 93.81505195028168 | 89.79028578008163 | 125.15015250207 | 57.176467251314975 | 112.99686434028798 | 125.7320142439726 | 90.57449833816732 | 99.87395144128429 | 53.994108376104734 | 164.8167038073688 |
| SE_282_1_8 | SE_282 | SE_282_1 | TCCTTAGAATGGCCTCCCAG | 210.25270728753 | 296.81956793626716 | 253.79440369707783 | 277.1787082776433 | 179.24047265126973 | 237.36897010394398 | 217.3844437784588 | 324.25624726077143 | 319.6747000170611 | 546.0262016753426 | 274.6656817393154 | 291.3309341947153 |
| SE_282_2_1 | SE_282 | SE_282_2 | GCAGTTTCTTCTAGCTGTGA | 302.3737388310354 | 372.77339676844326 | 330.8214989825723 | 268.0695488506785 | 373.3292684807512 | 320.5347406513112 | 304.55345341239524 | 490.63846159866 | 245.08393667974687 | 504.4728058202097 | 644.407945619163 | 239.1002886219576 |
| SE_282_2_14 | SE_282 | SE_282_2 | TCTCCCAGTTGTGTGACCAT | 96.4561389102586 | 111.93195827899638 | 122.45333096668347 | 80.68112635311682 | 115.60480188750533 | 213.11228702762855 | 89.32133085946575 | 87.9178746217252 | 107.44621861684554 | 64.88161808959345 | 69.25331291717781 | 242.58233166014142 |
| SE_282_2_15 | SE_282 | SE_282_2 | GCTTTTCAAACGAAGCACAG | 53.105065242726646 | 92.94350107095235 | 84.92731018657079 | 62.46280749918722 | 72.12042686559965 | 24.256683076315444 | 89.32133085946575 | 55.775855942814914 | 199.7966875106632 | 99.1449444964574 | 63.384388093688166 | 41.78451645820618 |
| SE_282_2_26 | SE_282 | SE_282_2 | TCACAGTCACTAATAACTGG | 1207.327401640765 | 1320.197472201377 | 1421.051155331109 | 1238.8456820672131 | 1381.954650086417 | 1573.2191595210302 | 1021.2764215136505 | 1417.0848823437213 | 1348.849637016433 | 909.0716601991352 | 1494.2282600604638 | 1518.170764648158 |
| SE_282_2_7 | SE_282 | SE_282_2 | GGAGTTCAAGTCAATCCGAA | 296.9548546225939 | 334.7964823523552 | 324.89633780676496 | 347.4493667142289 | 285.2999239242104 | 336.1283226289426 | 276.5732774805144 | 283.6060471668555 | 296.5870827936067 | 334.61418767554375 | 164.32989505771008 | 429.4519747093413 |
| SE_282_3_0 | SE_282 | SE_282_3 | CAGAAAGAATCCATAATCCG | 676.2767492134985 | 759.5382883217611 | 815.6971885361334 | 785.9903276981058 | 872.8692839763017 | 672.2566452578852 | 711.342165037432 | 782.7526901805212 | 857.793778379114 | 1597.2542161157219 | 820.4756903238524 | 851.9398633423149 |
| SE_282_3_11 | SE_282 | SE_282_3 | GCTTACTTATGTTTTAGCAA | 152.81253467805016 | 229.86027146579613 | 237.00644703229057 | 201.70281588279207 | 203.6341464440461 | 279.818165487496 | 163.5764131402264 | 225.93948424292822 | 184.70093778763533 | 191.72882648947277 | 338.0500698330036 | 379.5426911620395 |
| SE_282_3_14 | SE_282 | SE_282_3 | ATAGTTATGGAAAAAACACA | 17.340429467012783 | 51.968409200962604 | 36.53849391747813 | 54.65495656178882 | 56.21150917465856 | 43.31550549342044 | 49.50338818717378 | 50.10373499947781 | 86.13457194904147 | 53.21750697236316 | 65.73195802308402 | 1.1606810127279494 |
| SE_282_3_25 | SE_282 | SE_282_3 | TTAGGTGAATAGAATCCAAA | 140.89098941947887 | 192.88274953434197 | 149.11655625781614 | 148.34916781056964 | 208.93711900769313 | 141.20854790855063 | 90.39749147223039 | 155.03797245121433 | 243.3079661240965 | 189.5418056549921 | 98.59793703462604 | 103.3006101327875 |
| SE_282_3_7 | SE_282 | SE_282_3 | CAGAGACAAGGTAAAGCCGA | 2002.8196034399764 | 1839.8815642110028 | 1835.8124376376174 | 1808.8188004972967 | 1928.1608241420615 | 2382.352802138124 | 1857.4532176317816 | 1880.308092716252 | 1992.638963439681 | 1629.3305216881051 | 1635.0824558242152 | 1641.2029519973205 |
| SE_283_1_1 | SE_283 | SE_283_1 | AGCAGACAAACGTCAAACAT | 2370.2199527723096 | 2095.7260402772804 | 2024.4300684008153 | 2312.425185959494 | 2532.699696397823 | 1879.0266283045785 | 2510.6827095799226 | 2207.400400448692 | 1991.7509781618558 | 2706.0737791974257 | 2278.3166164786803 | 1951.104782395683 |
| SE_283_1_12 | SE_283 | SE_283_1 | TCAGTGAAAGGACGAACAAG | 2015.8249255402359 | 1929.8268878280535 | 2120.220174076366 | 2154.966858721959 | 1895.28239424745 | 1943.9998865447092 | 1963.9931182954817 | 2340.695242617114 | 1862.1051275993811 | 2786.993550073211 | 2055.2974731860736 | 1345.2292937516934 |
| SE_283_1_17 | SE_283 | SE_283_1 | ACTTCCTCTTTGCGTCAGAG | 97.53991575194691 | 59.96354907803377 | 64.18924607124536 | 66.36673296788642 | 115.60480188750533 | 133.41175691973496 | 55.960351863761666 | 121.00524679119168 | 167.8292175089571 | 23.328222234460565 | 190.1531642810645 | 233.29688355831783 |
| SE_283_1_18 | SE_283 | SE_283_1 | CACTTCCTCTTTGCGTCAGA | 274.19554094713965 | 274.8329332743215 | 242.93160820809783 | 266.7682403611121 | 275.75457330964576 | 454.8128076809146 | 247.51694093586892 | 173.94504226233803 | 311.6828325166346 | 307.6409307169487 | 179.58909959878315 | 409.72039749296613 |
| SE_283_1_20 | SE_283 | SE_283_1 | GGTTTCTGGCAACTAAAACC | 26.010644200519174 | 7.9951398770711695 | 5.925161175807264 | 9.109159426964803 | 14.848323178211695 | 0.0 | 7.533124289352532 | 5.67212094333711 | 0.8879852778251698 | 1.4580138896537853 | 0.0 | 16.249534178191293 |
| SE_284_1_18 | SE_284 | SE_284_1 | AGGTGGGGCAGAAGAGGCCG | 701.2036165723293 | 873.4690315700252 | 803.8468661845188 | 745.6497645215475 | 810.2942077252668 | 586.4919443809127 | 743.6269834203714 | 789.3701646144145 | 792.9708530978767 | 912.7166949232696 | 997.7172199932397 | 436.41606078570896 |
| SE_284_1_23 | SE_284 | SE_284_1 | TTTCGGTGTTCACCCAACCT | 62.859056817921335 | 74.95443634754221 | 79.98967587339807 | 106.70729614444484 | 63.6356707637644 | 119.5507951618404 | 81.7882065701132 | 81.30040018783191 | 50.615160836034676 | 79.4617569861313 | 146.72312058724114 | 138.12104051462597 |
| SE_284_1_3 | SE_284 | SE_284_1 | AGACAAGCGTGGCTATCTAA | 418.33786089168336 | 499.6962423169481 | 509.5638611194247 | 407.3095572342833 | 480.44931426642125 | 453.0801874611778 | 399.2555873356842 | 384.7588706563673 | 353.4181405744176 | 331.6981598962362 | 441.3431467264213 | 318.02659748745816 |
| SE_284_1_5 | SE_284 | SE_284_1 | GTGTGCGTGTGAATGTCAAG | 2670.4261379199684 | 2585.4283577478895 | 2609.0459710804653 | 2637.7523083510937 | 2653.6074708489755 | 2871.818014213775 | 2489.1594973246297 | 2302.8811029948665 | 2559.1735706921395 | 2496.1197790872807 | 2258.3622720788153 | 2714.832888770674 |
| SE_284_1_7 | SE_284 | SE_284_1 | GTCTGCGTGAGAACAATGTG | 410.75142299986527 | 402.75517130746016 | 408.83612113070126 | 399.50170629688495 | 363.7839178661865 | 637.6042408631488 | 391.72246304633165 | 400.82987999582247 | 389.8255369652495 | 518.3239377719207 | 347.440349550587 | 513.0210076257537 |
| SE_285_1_16 | SE_285 | SE_285_1 | TGGACAATACAGGCCAGGCA | 673.0254186884337 | 576.649463633758 | 606.34149365761 | 607.711064627509 | 592.8723326157384 | 545.7753692170975 | 802.815817122427 | 789.3701646144145 | 691.7405314258073 | 681.6214934131447 | 982.4580154521666 | 444.5408278748046 |
| SE_285_1_18 | SE_285 | SE_285_1 | GCTCTTAATAAGGATTATGG | 44.434850509220254 | 63.961119016569356 | 68.13935352178353 | 49.44972260352321 | 80.60518296743491 | 86.63101098684088 | 36.58946083399801 | 27.415251226129364 | 86.13457194904147 | 150.17543063433988 | 3.521354894093787 | 10.446129114551544 |
| SE_285_1_2 | SE_285 | SE_285_1 | ATTCATAGTGTGTATTTAAG | 2.167553683376598 | 10.993317330972857 | 18.763010390056337 | 11.711776406097604 | 20.15129574185873 | 2.598930329605226 | 7.533124289352532 | 17.01636283001133 | 19.535676112153734 | 18.225173620672315 | 73.94845277596953 | 13.928172152735392 |
| SE_285_1_5 | SE_285 | SE_285_1 | ATTAGCAGTTGGATAGGAAA | 80.19948628493412 | 79.95139877071169 | 108.62795488979984 | 70.27065843658562 | 50.90853661101153 | 94.42780197565655 | 72.10276105523138 | 120.0598933006355 | 100.34233639424419 | 75.08771531716994 | 174.89395973999143 | 284.3668481183476 |
| SE_285_1_7 | SE_285 | SE_285_1 | ATGTGATCTGTAGGACTCAG | 745.6384670815496 | 786.5218854068763 | 736.6950395253698 | 780.7850937398402 | 734.9919973214788 | 771.8823078927521 | 735.0176985182542 | 834.7471321611114 | 716.604119204912 | 731.9229726062002 | 1058.754038157532 | 562.9302911730555 |
| SE_285_2_14 | SE_285 | SE_285_2 | GCCTCTGCCAAAACACAGGA | 106.21013048545329 | 194.88153450360974 | 181.7049427247561 | 149.65047630013606 | 138.9378811675523 | 207.9144263684181 | 128.06311291899306 | 169.21827480955713 | 245.08393667974687 | 119.5571389516104 | 190.1531642810645 | 73.12290380186082 |
| SE_285_2_25 | SE_285 | SE_285_2 | AATAAGGGAGACACACACCT | 383.6570019576578 | 333.7970898677213 | 301.1956931035359 | 333.1349733289985 | 393.4805642226099 | 281.5507857072328 | 383.1131781442145 | 404.61129395804716 | 414.6891247443543 | 271.19058347560406 | 326.31222018602426 | 249.54641773650914 |
| SE_285_2_3 | SE_285 | SE_285_2 | GTATCTTGAAGGATAAGTAA | 13.005322100259587 | 32.97995199291857 | 40.488601368016305 | 10.410467916531204 | 37.120807945529236 | 84.03208065723565 | 1.0761606127646475 | 29.305958207241734 | 52.391131391685015 | 5.832055558615141 | 1.173784964697929 | 12.767491140007444 |
| SE_285_2_4 | SE_285 | SE_285_2 | GCCAACTGAAAAGGCACGTA | 2111.197287608806 | 2131.7041697241007 | 2097.5070562357714 | 2177.089103044588 | 2162.5522114552605 | 1456.2672946887951 | 2163.0828316569414 | 2092.067274600837 | 1946.463728992772 | 2184.1048067013703 | 2434.430016783505 | 1757.2710532701155 |
| SE_285_2_8 | SE_285 | SE_285_2 | GAATGTTGAGAGGGTGACAG | 403.1649851080472 | 388.7636765225856 | 405.87354054279757 | 498.4011515039314 | 412.57126545173924 | 371.64703713354737 | 357.285323437863 | 452.8243219764126 | 472.4081678029903 | 481.14458358574916 | 534.0721589375577 | 242.58233166014142 |
| SE_286_1_0 | SE_286 | SE_286_1 | ATCAGATGTGGGTTTCAGAA | 79.11570944324582 | 164.89975996459287 | 148.12902939518162 | 132.73346593577284 | 257.72446659324584 | 104.82352329407746 | 101.15909759987686 | 243.90120056349573 | 89.68651306034215 | 255.88143763423932 | 136.15905590495976 | 78.92630886550056 |
| SE_286_1_24 | SE_286 | SE_286_1 | AAGCTCTGACCAGACCACGG | 1331.9617384349194 | 1620.0152175915457 | 1621.5191084459213 | 1794.5044071120662 | 1655.588034370604 | 1540.2993753460307 | 1569.042173410856 | 1888.8162741312576 | 1673.852248700445 | 2066.005681639414 | 1746.5920274705184 | 1802.5376127665054 |
| SE_286_1_26 | SE_286 | SE_286_1 | GCTGTGAAACAAGGCAAGGA | 1839.1693003450432 | 1837.882779241735 | 1873.33845841773 | 1862.172448569519 | 1766.9504582071916 | 1434.609541942085 | 1878.9764298870746 | 1707.3084039444702 | 1743.1151003708082 | 1761.2807787017728 | 1781.8055764114563 | 1908.159584924749 |
| SE_286_1_27 | SE_286 | SE_286_1 | GGCTGTGAAACAAGGCAAGG | 599.3285934536293 | 575.6500711491242 | 575.7281609159392 | 618.1215325440402 | 538.7820124665386 | 529.3154771295978 | 590.8121764077914 | 544.5236105603626 | 367.6259050196203 | 644.4421392269732 | 609.1943966782252 | 670.8736253567548 |
| SE_286_1_33 | SE_286 | SE_286_1 | TGAAGGATGTGGCACACAAG | 4511.762991948388 | 4162.469698500177 | 4088.361211307012 | 4384.108301349203 | 4023.895581295369 | 5043.657459653876 | 4666.232416947511 | 4391.166963633479 | 3874.2797671512158 | 5137.311940195113 | 4212.714238300868 | 5932.24065605255 |
| SE_287_1_27 | SE_287 | SE_287_1 | CTGTGTCTAAGGTAACTGTA | 130.05322100259588 | 145.91130275654885 | 162.94193233469977 | 148.34916781056964 | 169.69512203670507 | 86.63101098684088 | 96.85445514881827 | 160.71009339455145 | 142.96562972985234 | 94.04189588266915 | 95.07658214053225 | 32.49906835638259 |
| SE_287_1_28 | SE_287 | SE_287_1 | TACCTTAGACACAGCCCTAG | 783.5706565406401 | 826.4975847922321 | 801.8718124592498 | 702.7065843658562 | 813.475991263455 | 811.732572946699 | 708.113683199138 | 714.6872388604759 | 888.873263102995 | 1018.422701923169 | 747.7010225125808 | 925.0627671441757 |
| SE_287_1_30 | SE_287 | SE_287_1 | GAAGTAGGAGAAGACCACTA | 1033.9231069706373 | 1104.3286955204553 | 1170.2193322219348 | 1054.0598765487844 | 1065.8974852930537 | 1313.3261265605076 | 1041.7234731561787 | 1275.2818587602935 | 1210.3239336757065 | 982.7013616266513 | 1260.6450520855758 | 1129.3426253842947 |
| SE_287_1_31 | SE_287 | SE_287_1 | GGAAGTAGGAGAAGACCACT | 610.1663618705123 | 624.6203028961851 | 667.5681591409518 | 586.8901287944466 | 686.2046497359262 | 467.8074593289407 | 608.0307462120259 | 750.6106715016109 | 678.4207522584297 | 753.793180951007 | 598.6303319959438 | 727.7469949804243 |
| SE_287_1_33 | SE_287 | SE_287_1 | GGTCTTCTCCTACTTCCTCA | 33.59708209233727 | 19.987849692677923 | 17.77548352742179 | 18.218318853929606 | 31.8178353818822 | 54.57753692170975 | 43.0464245105859 | 84.13646065950047 | 14.207764445202717 | 44.46942363444045 | 36.3873339056358 | 12.767491140007444 |
| SE_288_1_17 | SE_288 | SE_288_1 | CGAGGGGGATGGGAGTGCAG | 31670.12686781547 | 27558.247763779687 | 26841.96765326954 | 30225.492287158784 | 28239.389495933185 | 29710.105217937078 | 32902.53457466633 | 30654.977638265413 | 27700.700741756173 | 32330.728996127862 | 32274.391389334258 | 29056.488472631485 |
| SE_288_1_18 | SE_288 | SE_288_1 | GCTACATCAGAGGCCAGCTA | 2081.935312883222 | 2277.6154724806493 | 2169.5965172080932 | 2043.0543286192487 | 2558.153964703329 | 2314.780613568388 | 2122.1887283718847 | 2479.6622057288732 | 2338.953221791497 | 1664.322855039796 | 2745.483032428456 | 2486.1787292632675 |
| SE_288_1_3 | SE_288 | SE_288_1 | TCTTGCTCTGGTCCTGAGCG | 653.5174355380442 | 646.6069375581309 | 693.2438575694499 | 546.5495656178882 | 730.7496192705612 | 808.2673325072253 | 626.3254766290248 | 712.7965318793634 | 466.19227085821416 | 743.5870837234305 | 507.07510474950533 | 682.4804354840343 |
| SE_288_1_33 | SE_288 | SE_288_1 | AGCAGCAAAGAGCTGAGTGG | 834.5081680999901 | 895.455666231971 | 975.6765402829295 | 901.8067832695154 | 795.445884547055 | 825.5935347045935 | 749.0077864841946 | 808.2772344255382 | 1149.940934783595 | 756.7092087303146 | 638.5390207956734 | 771.8528734640864 |
| SE_288_1_34 | SE_288 | SE_288_1 | ACTGTCACTTGGTCCTCTGG | 1181.3167574402457 | 1189.2770567143364 | 1204.7827724141437 | 1223.2299801924164 | 1244.0773634315942 | 1109.7432507414317 | 1277.4026473516365 | 1412.3581148909404 | 1178.3564636740002 | 1517.0634521847637 | 1286.4683213089302 | 1164.1630557661333 |
| SE_288_2_15 | SE_288 | SE_288_2 | TCCCGCCCCCACCAGCCGGC | 113.79656837727138 | 190.88396456507417 | 232.06881271911786 | 186.08711400799527 | 250.30030500414 | 198.3850151598656 | 178.6426617189315 | 199.46958650735505 | 380.9456841869978 | 290.87377098593015 | 199.54344399864794 | 225.1721164692222 |
| SE_288_2_18 | SE_288 | SE_288_2 | GCCCTCCCGCCCCCACCAGC | 20.59175999207768 | 26.983597085115196 | 24.688171565863602 | 15.615701874796805 | 31.8178353818822 | 19.05882241710499 | 18.294730416999005 | 34.03272566002266 | 39.95933750213264 | 65.61062503442034 | 24.64948425865651 | 17.410215190919242 |
| SE_288_2_26 | SE_288 | SE_288_2 | GCGGCCATCGGTCTCTGGGG | 738.0520291897316 | 1038.3687915346181 | 1163.3066441834928 | 750.854998479813 | 974.6863571983248 | 654.0641329506486 | 627.4016372417894 | 883.905513670033 | 837.3701169891351 | 1033.7318477645338 | 672.5787847719133 | 526.949179778489 |
| SE_288_2_27 | SE_288 | SE_288_2 | TGTGCCCCCCCAGAGACCGA | 1386.1505805193342 | 1493.092372043041 | 1426.9763165069162 | 1353.3608291490564 | 1544.2256105340161 | 1673.7111322657656 | 1334.439159828163 | 1749.8493110194984 | 1415.4485328533206 | 1185.3652922885274 | 1535.3107338248913 | 2034.6738153120953 |
| SE_288_2_7 | SE_288 | SE_288_2 | GCCTACCGAGTGATTCCGGC | 669.7740881633687 | 934.431973132693 | 953.9509493049695 | 606.4097561379426 | 903.6265248454546 | 913.0908558013028 | 594.0406582460854 | 847.0367275383418 | 815.1704850435059 | 776.3923962406407 | 559.8954281609122 | 1221.0364253898028 |
| SE_289_1_17 | SE_289 | SE_289_1 | TGTGCTCCATTGTTAACTGG | 1209.4949553241415 | 1158.2958896906857 | 1283.7849214249072 | 1223.2299801924164 | 1346.9550311663465 | 790.0748201999887 | 1197.7667620070526 | 1568.341440832711 | 1372.8252395177126 | 1162.7660769988938 | 1129.1811360394076 | 1276.7491140007444 |

| | | | | | | | | | | | | | | | |
|---|---|---|---|---|---|---|---|---|---|---|---|---|---|---|---|
| SE_289_1_19 | SE_289 | SE_289_1 | TCATCTCCACCATCCAAGGA | 473.6104798177866 | 572.6518936952225 | 569.8029997401319 | 491.89460905609934 | 622.5689789721617 | 602.085526358544 | 419.7026389782125 | 695.7801690493521 | 597.6140919763393 | 384.1866599237724 | 537.5935138316515 | 531.5919038294008 |
| SE_289_1_20 | SE_289 | SE_289_1 | TTGTTAACTGGGGGTGAGCA | 311.04395356454177 | 332.79769738308744 | 344.64687505945585 | 262.8643148924129 | 356.3597562770807 | 310.1390193328903 | 170.0333768168143 | 364.90644735468743 | 343.6503025183407 | 102.06097227576497 | 194.84830413985622 | 120.71082532370674 |
| SE_289_1_21 | SE_289 | SE_289_1 | ACTCAGGTCCCACCCCTAAT | 654.6012123797325 | 713.5662340286019 | 691.2688038441809 | 748.2523815006803 | 658.6291924049616 | 695.6470182243322 | 758.6932319990765 | 690.1080481060151 | 776.0991328191984 | 839.0869934957534 | 854.5154543000923 | 846.1364582786751 |
| SE_289_1_4 | SE_289 | SE_289_1 | GCTGTATCTCCGGTAAAGAA | 101.8750231187001 | 198.87910444214535 | 224.1685978180415 | 150.95178478970246 | 110.3018293238583 | 67.57218856973589 | 102.23525821264151 | 247.68261452572048 | 229.1002016788938 | 159.6525209170895 | 258.2326922335444 | 185.70896203647192 |
| SE_289_2_14 | SE_289 | SE_289_2 | CTTAAACAAGTAAGAGGGGG | 1310.2862016011534 | 1723.9520359934709 | 1771.623191566372 | 1535.5440176883526 | 1841.19207409825 | 1872.9624575354997 | 1212.8330105857576 | 1647.7511340394306 | 1928.7040234362687 | 1835.6394870741158 | 1650.3416603652881 | 1628.435460857313 |
| SE_289_2_15 | SE_289 | SE_289_2 | CTCCCCCTCTTACTTGTTTA | 4.335107366753196 | 11.992709815606755 | 15.800429802152705 | 11.711776406097604 | 1.0605945127294067 | 0.8663101098684087 | 1.0761606127646475 | 0.945353490556185 | 11.543808611727208 | 0.7290069448268927 | 0.0 | 0.0 |
| SE_289_2_16 | SE_289 | SE_289_2 | TGCCTTAAACAAGTAAGAGG | 263.3577725302566 | 344.7904071986942 | 322.9212840814959 | 268.0695488506785 | 275.75457330964576 | 147.27271867762948 | 146.35784333599204 | 361.1250333924627 | 191.80482001023668 | 265.3585279169889 | 273.4918967746175 | 177.58419494737626 |
| SE_289_2_17 | SE_289 | SE_289_2 | CTGCCTTAAACAAGTAAGAG | 52.02128840103835 | 97.94046349412183 | 79.98967587339807 | 94.99551973834723 | 110.3018293238583 | 40.71657516381521 | 32.28481838293943 | 93.58999556506231 | 70.15083694818841 | 77.27473615165061 | 31.692194046844083 | 49.909283547301825 |
| SE_289_2_18 | SE_289 | SE_289_2 | CCTGCCTTAAACAAGTAAGA | 406.4163156331121 | 524.6810544327955 | 517.4640760205011 | 451.554045879541 | 605.5994667684912 | 524.9839265802557 | 406.78871162503674 | 698.6162295210207 | 517.695416972074 | 293.7897987652377 | 829.8659700414358 | 395.79222534023074 |
| SE_289_3_1 | SE_289 | SE_289_3 | TCAGCAGGATAGCAGTAAGA | 2940.286571500355 | 3080.127637641668 | 3073.183596518701 | 3276.6947767281963 | 3043.906251533397 | 2688.1602709216722 | 3013.249715741013 | 3301.174389022198 | 3262.4579107296736 | 3776.984981148131 | 3576.5227874345896 | 3165.177121709118 |
| SE_289_3_2 | SE_289 | SE_289_3 | GAGGGATAAAAGCATAAGAG | 512.6264461185654 | 575.6500711491242 | 659.6679442398754 | 564.7678844718178 | 583.3269820011737 | 673.989265477622 | 510.1001304504429 | 496.3105825419971 | 802.7386911539535 | 723.9038962131044 | 640.8865907250693 | 786.9417266295497 |
| SE_289_3_3 | SE_289 | SE_289_3 | AGAGAGGCTACATAAAAGCC | 1079.4417343215457 | 1242.244858399933 | 1243.2963200568909 | 1083.9899718088116 | 1235.592607329759 | 1113.2084911809052 | 1173.0150679134658 | 1141.0416631013154 | 935.9364828277289 | 1188.281320067835 | 1510.6612495662346 | 1539.063022877261 |
| SE_289_3_6 | SE_289 | SE_289_3 | GCATATGATTTATAAAGGAA | 61.77527997623304 | 144.91191027191493 | 102.70279371399258 | 100.20075369661284 | 173.9375000876227 | 77.10159977828837 | 58.112673089290965 | 106.82494443284891 | 70.15083694818841 | 132.67926395849446 | 104.46686185811568 | 182.22691899828806 |
| SE_289_3_7 | SE_289 | SE_289_3 | TCAGATAGTGTTCAGAAACA | 882.1943491342753 | 958.4173927639065 | 896.674391272166 | 877.0819219677539 | 824.0819363907491 | 1172.9838887618255 | 883.5278630797756 | 988.8397511217695 | 1078.9021125575812 | 761.8122573441028 | 936.6804018289474 | 1182.7339519697805 |
| SE_29_1_1 | SE_29 | SE_29_1 | GACAGATAAAGGCTGCAGCA | 193.9960546622055 | 60.962941562667666 | 91.8399982250126 | 217.31851775758886 | 67.87804881468203 | 356.9197652657844 | 259.35470767628004 | 138.96696311175918 | 351.6421700187672 | 13.122125006884067 | 68.07952795247988 | 211.2439443164868 |
| SE_29_1_10 | SE_29 | SE_29_1 | TCTCAGCATCTGTTTCTGGA | 404.24876194973547 | 454.72358050842274 | 478.95052837775387 | 446.34881192127534 | 318.178353818822 | 697.379638444069 | 335.76211118257004 | 310.07594490242866 | 471.5201825251652 | 497.9117433167677 | 244.14727265716925 | 545.5200759821363 |
| SE_29_1_12 | SE_29 | SE_29_1 | GGACACATCCCAAAATAGGA | 307.7926230394769 | 377.7703591916128 | 353.53461682316674 | 330.5323563498657 | 362.7233233534571 | 521.518686140782 | 366.9707689527448 | 444.31614056140694 | 310.79484723880944 | 224.53413900668295 | 559.8954281609122 | 537.3953088930406 |
| SE_29_1_19 | SE_29 | SE_29_1 | GGAGCAATAAGAAGAGGAGG | 425.92429878350146 | 681.5856745203172 | 637.9423532619154 | 543.9469486387554 | 550.4485521065621 | 360.38500570525804 | 349.75219914851044 | 347.89008452467607 | 873.7775133799671 | 344.09127795829335 | 254.7113373394506 | 212.40462532921475 |
| SE_29_1_29 | SE_29 | SE_29_1 | AGACTTCCTGGGGGTCTCAG | 11.921545258571289 | 28.982382054382988 | 15.800429802152705 | 13.013084895664004 | 18.030106716399914 | 52.844916701972934 | 2.152321225529295 | 16.071009339455145 | 45.28724916908366 | 1.4580138896537853 | 16.432989505771005 | 40.62383544547823 |
| SE_29_2_11 | SE_29 | SE_29_2 | TATAGATTAACAATATAAAA | 0.0 | 4.996962423169481 | 0.987526862634544 | 3.9039254686992013 | 4.242378050917627 | 0.0 | 0.0 | 0.0 | 0.0 | 0.0 | 0.0 | 0.0 |
| SE_29_2_12 | SE_29 | SE_29_2 | ATAGGAATAGAAAAGTCTGT | 180.99073256194592 | 159.90279754142338 | 154.05419057098888 | 139.24000838360485 | 199.39176839312847 | 173.26202197368175 | 166.80489497852037 | 183.3985771678999 | 215.78042251151626 | 117.37011811712972 | 163.15611009301213 | 169.4594278582806 |
| SE_29_2_13 | SE_29 | SE_29_2 | TGTGGTCAAGACAATGAAGA | 751.0573512899912 | 844.4866495156423 | 829.522564613017 | 731.335371136317 | 781.6581558815727 | 662.7272340493327 | 570.3651247652632 | 695.7801690493521 | 791.1948825422263 | 1156.9340214402787 | 612.7157515723189 | 875.1534835968739 |
| SE_29_2_6 | SE_29 | SE_29_2 | AGAGTTCAGGAACCTGTGGG | 117.04789890233629 | 217.86756165018937 | 191.58021135110155 | 111.91253010271043 | 164.39214947305805 | 241.70052065328605 | 88.2451702467011 | 295.8956425440859 | 158.94936473070538 | 100.60295838611118 | 261.75404712763816 | 335.4368126783774 |
| SE_29_2_7 | SE_29 | SE_29_2 | GATCCCATCTCTAAAAGATG | 8.670214733506391 | 32.97995199291857 | 18.763010390056337 | 7.807850937398403 | 30.757240869152795 | 25.989303296052263 | 5.380803063823238 | 5.67212094333711 | 13.319779167377547 | 2.9160277793075706 | 0.0 | 51.069964560029774 |
| SE_29_3_11 | SE_29 | SE_29_3 | AGAAGGAAACCAGAATACTG | 104.0425768020767 | 155.9052276028878 | 128.37849214249073 | 117.11776406097604 | 136.81669214209347 | 65.83956834999907 | 102.23525821264151 | 86.97252113116902 | 134.9737622294258 | 185.16776398603074 | 46.95139858791716 | 13.928172152735392 |
| SE_29_3_13 | SE_29 | SE_29_3 | AAAAATGAGATAAATAACAA | 2.167553683376598 | 8.994532361705065 | 5.925161175807264 | 9.109159426964803 | 3.1817835381882205 | 0.0 | 1.0761606127646475 | 34.97807915057884 | 0.0 | 0.7290069448268927 | 31.69219404684083 | 0.0 |
| SE_29_3_17 | SE_29 | SE_29_3 | TGGAGTATAGTAGGGACACA | 291.5359704141524 | 266.83779339725027 | 241.94408134546327 | 234.2355281219521 | 299.0876525896927 | 187.12298373157628 | 279.80175931880837 | 342.21796358133895 | 244.1959514019217 | 88.20984032405401 | 356.8306292681704 | 340.0795367292892 |
| SE_29_3_33 | SE_29 | SE_29_3 | GCCCGAGGTGGTCTACTCCA | 59.60772629285644 | 111.93195827899638 | 120.47827724141437 | 110.61122161314404 | 126.21074701479941 | 23.390372966447035 | 69.95043982970209 | 105.87959094229272 | 154.50943834157954 | 144.34337507572474 | 113.85714157569912 | 61.51609367458132 |
| SE_29_3_8 | SE_29 | SE_29_3 | CTATGCCCTTGGACAATGCA | 117.04789890233629 | 140.91434033337936 | 139.2412876314707 | 94.99551973834723 | 127.2713415275288 | 110.02138395328791 | 85.01668840840715 | 46.32232103725307 | 96.7903952829435 | 32.076305572383276 | 83.33873249355297 | 91.69380000550801 |
| SE_29_4_10 | SE_29 | SE_29_4 | GGTGTGAGACAATGAACGCA | 1023.0853385537541 | 1121.3183677592315 | 996.4146043982549 | 1115.2213755584053 | 1037.2614334493599 | 1081.1550171157742 | 1220.3661348751102 | 1311.2052914014287 | 943.9283503281555 | 1434.6856674193248 | 1343.9837845791287 | 699.8906506749535 |
| SE_29_4_34 | SE_29 | SE_29_4 | CATGTATCTGGCAAAGAGAA | 1019.8340080286893 | 1237.2478959767634 | 1203.7952455515092 | 996.8023030078627 | 1193.1688268205826 | 1699.7004355618178 | 1029.8857064157676 | 1187.3639841385684 | 1361.2814309059852 | 1171.5141603368165 | 1335.7672898262433 | 1259.3388988098252 |
| SE_29_4_5 | SE_29 | SE_29_4 | GCTCCCTCTAACTATACAGT | 66.11038734298623 | 106.9349958558269 | 117.51569665351074 | 87.18766880094883 | 112.42301834931712 | 51.978606592104526 | 67.79811860417279 | 98.31676301784324 | 133.19779167377547 | 58.320555586151414 | 126.76877618737633 | 109.10401519642724 |
| SE_29_4_6 | SE_29 | SE_29_4 | AGAAAGCTAGAAGCTTGCAC | 68.27794102636283 | 114.93013573289807 | 100.72773998872348 | 55.95626505135522 | 146.36204275665813 | 105.68983340394587 | 37.665621446762664 | 99.26211650839943 | 182.0369819541598 | 51.030486137882484 | 92.72901221113639 | 163.65602279464088 |
| SE_29_4_9 | SE_29 | SE_29_4 | TTTGCATAGGGCCCACAGAT | 353.31125039038545 | 468.71507529329733 | 406.86106740543215 | 318.8205799437681 | 395.6017532480687 | 560.5026410848604 | 289.48720483369016 | 380.03210320358636 | 453.76047696866175 | 381.99963908929175 | 419.0412323971607 | 965.686602589654 |
| SE_29_5_1 | SE_29 | SE_29_5 | ATTTCAGCAGGGTTGAAAAA | 11.921545258571289 | 15.990279754142339 | 18.763010390056337 | 15.615701874796805 | 12.727134152752882 | 1.7326202197368175 | 47.35106696164449 | 26.46989773557318 | 6.215896944776189 | 63.42360419993966 | 83.33873249355297 | 1.1606810127279494 |
| SE_29_5_16 | SE_29 | SE_29_5 | TCTGAGTGACAGAAACAGGA | 361.98146512389184 | 502.6944197708498 | 475.98794778985024 | 398.20039780731855 | 426.3589941172215 | 556.1710905355184 | 269.0401531911619 | 473.6220987686487 | 320.5626852948863 | 249.3203751307973 | 400.2606729619938 | 405.07767344205433 |
| SE_29_5_19 | SE_29 | SE_29_5 | CCCAAGTACGAGGTTCCAGA | 62.859056817921335 | 126.92284554850481 | 111.59053547770347 | 61.16149900962082 | 76.36280491651729 | 59.7753975809202 | 47.35106696164449 | 71.84686528227006 | 36.40739639083196 | 57.59154864132452 | 133.8114859755639 | 146.24580760372163 |
| SE_29_5_2 | SE_29 | SE_29_5 | TCTGGAATCTTGAACCACAC | 882.1943491342753 | 773.5297831066356 | 803.8468661845188 | 966.8722077478355 | 685.1440552231968 | 606.4170769078861 | 969.6207121009473 | 534.1247221642445 | 805.402646987429 | 1030.0868130403994 | 573.9808477372873 | 360.97179495839225 |
| SE_29_5_3 | SE_29 | SE_29_5 | TGTGTGGTTCAAGATTCCAG | 214.5878146542832 | 264.8390084279825 | 314.033542317785 | 264.1656233819793 | 343.6326221243278 | 369.91441691381056 | 235.6791741954578 | 418.79159631639 | 253.07580418017338 | 565.7093891856687 | 238.2783478336796 | 391.14950128931895 |
| SE_29_6_12 | SE_29 | SE_29_6 | TAATCTCCAAAGGTCCACGA | 193.9960546622055 | 285.8262506052943 | 271.5698872244996 | 201.70281588279207 | 292.72408551331625 | 454.8128076809146 | 230.29837113163455 | 218.37665631847872 | 191.80482001023668 | 110.80905561368769 | 244.14727265716925 | 129.99627342553035 |
| SE_29_6_13 | SE_29 | SE_29_6 | AATCTCCAAAGGTCCACGAT | 29.26197472558407 | 51.968409200962604 | 41.47612823065085 | 28.62878677046081 | 42.42378050917627 | 3.465240439473635 | 29.056336544645482 | 30.25131169779792 | 76.3667338929646 | 52.48850002753627 | 18.780559435166865 | 26.695663292742836 |
| SE_29_6_7 | SE_29 | SE_29_6 | TCTTTGCAGTCGGATACTTC | 372.8192335407748 | 387.7642840379517 | 404.88601368016305 | 337.0388987976977 | 463.47980206275076 | 221.77538812631263 | 321.7720232166296 | 354.5075589585694 | 337.4344055735645 | 340.4462432341589 | 433.1266519735358 | 403.9169924293264 |
| SE_29_6_8 | SE_29 | SE_29_6 | ATCACACTCTTAAAAATCTG | 143.05854310285545 | 151.90765766435223 | 175.77978154894885 | 153.55440176883525 | 216.36128059679896 | 239.1015903236808 | 221.6890862295174 | 290.2235216007488 | 206.90056973326455 | 243.48831957218215 | 311.0530156449512 | 201.9584962146632 |
| SE_29_6_9 | SE_29 | SE_29_6 | ACACTCTTAAAAATCTGTGG | 48.769957875973454 | 56.96537162413208 | 93.81505195028168 | 59.86019052005442 | 92.27172260745839 | 36.38502461447317 | 68.87427921693744 | 66.17474433893295 | 65.71091055906257 | 33.53431946203706 | 51.646538446708874 | 69.64086076367697 |
| SE_290_1_4 | SE_290 | SE_290_1 | TTATGGCCAATTTTAAACAA | 295.8710777809056 | 468.71507529329733 | 415.74880916914304 | 411.21348270298256 | 504.84298805919764 | 408.0320617480205 | 309.9342564762185 | 330.87372169466477 | 447.54458002388554 | 771.2893476268524 | 318.0957254331388 | 392.3101823020469 |
| SE_290_1_41 | SE_290 | SE_290_1 | GAGTTGCAAGGGTAAGTCGG | 1349.302167901932 | 1543.0619962747357 | 1432.9014776827235 | 1452.2602743561029 | 1535.7408544321809 | 1012.7165184361698 | 1407.618081496159 | 1382.1068031931425 | 1451.8559292441525 | 963.7471810611521 | 990.6745102050521 | 778.816959540454 |
| SE_290_1_42 | SE_290 | SE_290_1 | GAGGAGTTGCAAGGGTAAGT | 6754.097277401479 | 6791.871325571959 | 7146.7319048861955 | 6976.3148125654725 | 6817.5015278246265 | 7244.951448829503 | 7032.709604416971 | 6823.561494834544 | 7093.226399267456 | 7255.806121862062 | 6868.989613412281 | 7260.059734613324 |
| SE_290_1_43 | SE_290 | SE_290_1 | ACCCTTGCAACTCCTCTCCA | 630.75812186259 | 711.5674490593341 | 774.2210603054825 | 736.5406050945827 | 672.4169210704439 | 668.7914048184116 | 625.2493160162602 | 929.2824812167298 | 784.9789855974501 | 729.0069448268927 | 779.3932165594249 | 667.391582318571 |
| SE_290_1_44 | SE_290 | SE_290_1 | ACCTTGGAGAGGAGTTGCAA | 1663.5974519915387 | 1930.8262803126875 | 1958.2657686043008 | 1906.4169372147767 | 1861.3433698401088 | 1717.8929478690545 | 2056.5429309932415 | 1754.5760784722793 | 1678.292175089571 | 2386.03973041842 | 1724.2901131412577 | 1615.6679697173056 |
| SE_290_2_20 | SE_290 | SE_290_2 | GAAGACCGCTCACAGCACCT | 3483.2587691861927 | 3771.7072370083242 | 3838.5169150604725 | 3622.842834952859 | 3875.412349513252 | 3829.0906856183665 | 3807.456247961323 | 4317.429391370097 | 3630.083815749294 | 3854.9887242446084 | 5188.129543964847 | 3570.2547951511724 |
| SE_290_3_10 | SE_290 | SE_290_3 | AGTAAAACCAAAAACTGTGG | 82.36703996831072 | 96.94107100948793 | 95.79010567555078 | 94.99551973834723 | 108.18064029839948 | 71.03742900920952 | 115.14918556581728 | 202.30564697902358 | 60.382998892111544 | 156.73649313778193 | 179.58909959878315 | 32.49906835638259 |
| SE_290_3_12 | SE_290 | SE_290_3 | AGTTCACAGAGATAGCATAT | 141.97476626116716 | 220.86573910409106 | 190.592684488467 | 216.01720926802247 | 177.11928362581094 | 170.66309164407653 | 144.20552211046277 | 115.33312584785457 | 196.24474639936253 | 516.13691693744 | 53.994108376104734 | 83.56903291641235 |
| SE_290_3_15 | SE_290 | SE_290_3 | TTAGAAAATAACATTAGAAG | 8.670214733506391 | 17.98906472341013 | 28.63827901640178 | 13.013084895664004 | 32.87842989461161 | 0.8663101098684087 | 19.370891029763655 | 12.289595377230405 | 0.0 | 17.496166675845423 | 24.64948425865651 | 2.321362025455899 |
| SE_290_3_2 | SE_290 | SE_290_3 | GTTGTCAAAGTGAAACTGTG | 217.8391451793481 | 237.85541134286728 | 255.7694574223469 | 204.30543286192486 | 219.5430641349872 | 249.49731164210172 | 253.9739046124568 | 178.67180971511897 | 227.32423112324346 | 387.8316946479069 | 136.15905590495976 | 289.0095721692594 |
| SE_291_1_1 | SE_291 | SE_291_1 | AGTGACAGTGAAGGGACAAA | 980.8180417279106 | 1115.322012851428 | 1194.9075037877983 | 1044.9507171218195 | 963.0198175583013 | 1216.299394255246 | 948.0974998456544 | 743.0478435771614 | 1200.5560956196296 | 1230.5637228677947 | 578.675987596079 | 920.4200430932639 |
| SE_291_1_10 | SE_291 | SE_291_1 | TGTGTCCCAAGGCAAGCCCA | 1131.4630227225841 | 1112.3238353975264 | 1149.4812681066091 | 981.1866011330659 | 1013.9283541693128 | 991.0587656894596 | 1025.581063964709 | 1247.8666075341641 | 1092.2218917249588 | 944.063993550826 | 1039.9734787223651 | 935.5088962587272 |
| SE_291_1_22 | SE_291 | SE_291_1 | GCTTGGCTGTGCCCTGATTG | 2546.8755779675025 | 2235.6409881260256 | 2270.3242571968167 | 2709.3242752772458 | 2369.3681414374946 | 2256.737836207205 | 2752.8188474519684 | 2536.3834151622445 | 2709.243082644593 | 1935.5134385154 | 2145.6789154678145 | 2852.9539292852996 |
| SE_291_1_29 | SE_291 | SE_291_1 | AGGACTTGAACCCAGATAGT | 1818.5775403529656 | 2139.6993096011715 | 2181.4468395597078 | 1962.3732022661318 | 2198.61242488806 | 1699.7004355618178 | 1772.4365292233745 | 2153.5152514869897 | 2122.2848140021556 | 2113.391133053162 | 2037.6906987156049 | 1822.2691899828806 |
| SE_291_1_4 | SE_291 | SE_291_1 | GCACCCACGTAAGGTGTGCA | 1595.319510965176 | 1377.1628438255088 | 1349.9492212214218 | 1440.5484979500052 | 1339.5308695772408 | 1534.235204576952 | 1281.7072898026952 | 1424.6477102681708 | 1596.5975295296553 | 1396.7773062883264 | 1239.516922721013 | 1568.0800481954598 |
| SE_291_2_12 | SE_291 | SE_291_2 | TGTCCCCTGCCATTTCAGTG | 161.48274941155654 | 190.88396456507417 | 146.1539756699125 | 192.59365645582727 | 126.21074701479941 | 303.20853845394305 | 151.73864639981528 | 113.4424188667422 | 258.40371584712443 | 92.58388199301537 | 241.7997027277734 | 147.40648861644956 |
| SE_291_2_13 | SE_291 | SE_291_2 | AGGCCCCACTGAAATGGCAG | 424.84052194181317 | 467.7156828086634 | 503.63869994361744 | 434.63703551517773 | 539.842606979268 | 617.6791083361754 | 422.93112081650645 | 388.540284618592 | 517.695416972074 | 965.9342018956328 | 568.1119229137977 | 517.6637316766654 |
| SE_291_2_14 | SE_291 | SE_291_2 | CAGGCCCCACTGAAATGGCA | 1495.6120415298526 | 1457.1142425962207 | 1439.8141657211652 | 1431.4393385230405 | 1432.8631866974285 | 1079.4223968960373 | 1566.8898521853266 | 1632.6254781905316 | 1664.084410644368 | 1701.5022092259674 | 1631.5611009301213 | 1081.7547038624489 |
| SE_291_2_24 | SE_291 | SE_291_2 | CACAGGCCCTGCTCACAGGT | 784.6544333823284 | 725.5589438442087 | 741.6326738385426 | 827.6321993642307 | 716.9618906050789 | 933.8822984381446 | 830.7959930543078 | 551.1410849942558 | 705.0603105931848 | 664.8543336821261 | 522.3343092905784 | 1150.2348836133979 |
| SE_291_2_28 | SE_291 | SE_291_2 | CTCACAGGTGGGCAGCTGGT | 739.1358060314199 | 618.6239479883817 | 705.0941799210644 | 701.4052758762898 | 836.8090705435019 | 584.7593241611759 | 703.8090407480795 | 886.7415741417016 | 725.4839719831638 | 992.9074588542278 | 554.0265033374225 | 807.8339848586528 |
| SE_292_1_1 | SE_292 | SE_292_1 | AAGGAGATAAAGATGTACAT | 1056.6824206460915 | 1267.2296705157803 | 1209.7204067273165 | 1191.9985764428227 | 1364.9851378827466 | 1516.9090023795836 | 1050.332758058296 | 1183.5825701763436 | 1303.5623878473493 | 1046.124965826591 | 1248.9072024385964 | 1696.915640608262 |
| SE_292_1_11 | SE_292 | SE_292_1 | ATACTTTTTAGTGACTCCCG | 3.2513305250648967 | 9.993924846338961 | 17.77548352742179 | 11.711776406097604 | 9.545350614564661 | 0.0 | 6.456963676587884 | 10.398888396118036 | 31.07948472388094 | 9.477090282749604 | 3.521354894093787 | 5.803405063639747 |
| SE_292_1_16 | SE_292 | SE_292_1 | CTAAGGAATAAAGAGTACAG | 67.19416418467453 | 127.92223803313871 | 135.29118018093254 | 124.92561499837444 | 147.42263726938754 | 63.24063802039384 | 107.61606127646475 | 192.85211207346174 | 109.22218917249589 | 160.3815278619164 | 165.503680022408 | 113.74673924733905 |
| SE_292_1_2 | SE_292 | SE_292_1 | GTAAAGGAACAGACCGACAA | 1353.6372752686852 | 1449.1191027191494 | 1399.3255643531488 | 1346.8542867012245 | 1385.1364336246052 | 1342.7806702960336 | 1338.7438022792214 | 1271.5004447980689 | 1992.638963439681 | 1310.754486798753 | 1471.926345731203 | 1505.4032735081505 |
| SE_292_1_7 | SE_292 | SE_292_1 | GGAGAAATACAAGGCCTCTG | 144.14231994454374 | 243.85176625067066 | 277.49504840030687 | 219.92113473672168 | 239.69435987684594 | 206.1818061486813 | 198.01355274869513 | 245.7919075446081 | 260.17968640277473 | 291.60277793075704 | 332.1811450095139 | 83.56903291641235 |
| SE_293_1_28 | SE_293 | SE_293_1 | CACAGGGCGATGTCACCAGT | 7060.806123599267 | 6264.192093685261 | 6167.105257152727 | 6432.367863926717 | 6365.688265401899 | 5679.529080297288 | 6982.130055617033 | 6931.331792757948 | 7110.0981195461345 | 5836.429600284103 | 6671.793739343028 | 6360.531949749163 |
| SE_293_1_32 | SE_293 | SE_293_1 | GGGGAAACTGAGGCACTGGT | 872.4403575590807 | 1096.3335556433842 | 1022.090302826753 | 904.4094002486484 | 1090.29115908583 | 934.7486085480131 | 1067.5513278625303 | 905.6486439528252 | 943.0403650503304 | 1193.3843686816233 | 970.7201658051873 | 940.151620309639 |
| SE_293_1_35 | SE_293 | SE_293_1 | TCTTTGATGTGGGGAAACTG | 87.78592417675222 | 132.9192004563082 | 140.22881449410525 | 121.02168952967524 | 136.81669214209347 | 341.32618328815306 | 122.6823098551698 | 156.9286794323267 | 189.14086417676117 | 123.93118062057175 | 134.98527094026184 | 257.67118482560477 |
| SE_293_1_37 | SE_293 | SE_293_1 | GGAGCTTCTTCCTCCCCAGC | 33.59708209233727 | 52.9678016855965 | 29.62580587903632 | 41.641871666124814 | 53.02972563647034 | 116.08555472236677 | 8.60928490211718 | 13.23494886778659 | 63.93494000341222 | 29.160277793075707 | 49.29896851731302 | 62.67677468730927 |
| SE_293_1_43 | SE_293 | SE_293_1 | AGACCTCGCTCCCTTATGTG | 249.26867358830876 | 277.83111072822317 | 284.40773643874866 | 230.33160265325287 | 303.33003064061035 | 452.21387735130935 | 134.52007659558095 | 210.81382839402926 | 452.87249169083657 | 214.32804177910643 | 200.71722896334586 | 277.40276204197994 |
| SE_293_2_4 | SE_293 | SE_293_2 | GCCATCGCCCTCCACTCCTC | 502.8724545433707 | 371.7740042838094 | 354.5221436858013 | 430.73311004647854 | 299.0876525896927 | 414.0962325170994 | 637.0870827566713 | 354.5075589585694 | 221.99631945629244 | 209.95400011014507 | 470.68777084386954 | 178.74487596010422 |
| SE_293_2_45 | SE_293 | SE_293_2 | GGAGGAAGAAGTCCCCTGGG | 456.27005035077383 | 502.6944197708498 | 624.1169771850318 | 494.4972260352322 | 655.4474088667733 | 418.4277830664414 | 431.5404057186236 | 673.0916852760038 | 439.55271252345904 | 653.9192295097228 | 782.9145714535186 | 533.9132658548567 |
| SE_293_2_47 | SE_293 | SE_293_2 | AGGAAGAAGTCCCCTGGGAG | 455.18627350908554 | 811.5066975227237 | 770.2709528549443 | 612.9162985857746 | 773.1733997797376 | 1240.5560773315613 | 532.6995033185005 | 690.1080481060151 | 809.8425733765548 | 860.9572018405602 | 829.8659700414358 | 1013.2745241114999 |
| SE_293_2_51 | SE_293 | SE_293_2 | GTCCCCTGGGAGGGGGCCTG | 0.0 | 0.9993924846338962 | 4.93763431317272 | 0.0 | 1.0605945127294067 | 0.0 | 0.0 | 0.0 | 3.551941111300679 | 0.0 | 0.0 | 0.0 |
| SE_293_2_6 | SE_293 | SE_293_2 | TCGCCCTCCACTCCTCTGGG | 442.18095140882593 | 420.7442360308703 | 440.4369807350066 | 497.099843014365 | 504.84298805919764 | 833.3903256934092 | 387.4178205952731 | 388.540284618592 | 625.1416355889195 | 363.04545852379255 | 480.078050561453 | 506.05692154938595 |
| SE_293_3_1 | SE_293 | SE_293_3 | GGAGGAGGAAGAAGTCCCCT | 4522.6007603652715 | 4624.189026401038 | 4707.540554178871 | 4546.771862545003 | 4815.0990877915065 | 5183.9996974525575 | 4440.238688266935 | 4832.647043723217 | 4765.816986087686 | 4939.022051202198 | 4486.206135075485 | 4490.674838244437 |
| SE_293_3_23 | SE_293 | SE_293_3 | AGAAGGCAGCCCGATGGGAG | 1091.363279580117 | 1405.145833395258 | 1398.3380374905144 | 1185.4920339949908 | 1296.0464945553351 | 1313.3261265605076 | 1053.5612398965898 | 1035.1620721590225 | 1272.4829031234683 | 1073.827229730013 | 1348.6789244379204 | 1461.2973950244884 |

| | | | | | | | | | | | | | | | |
|---|---|---|---|---|---|---|---|---|---|---|---|---|---|---|---|
| SE_293_3_25 | SE_293 | SE_293_3 | CTGCGTTTCTCCTCTCCCAT | 455.18627350908554 | 398.75760136892455 | 406.86106740543215 | 445.04750343170895 | 415.75304898992744 | 399.36896064933643 | 446.6066542973287 | 415.01018235416524 | 516.8074316942489 | 623.3009378269933 | 521.1605243258805 | 623.2857038349089 |
| SE_293_3_34 | SE_293 | SE_293_3 | GCTCCCTCCTCCCAGAGGAG | 7.586437891818092 | 6.9957473924372735 | 8.887741763710896 | 1.3013084895664004 | 16.969512203670508 | 0.0 | 1.0761606127646475 | 5.67212094333711 | 11.543808611727208 | 2.187020834480678 | 3.521354894093787 | 12.767491140007444 |
| SE_293_4_38 | SE_293 | SE_293_4 | GGCTGTGCAGGCCTGGAGCG | 1044.7608753875202 | 1228.2533636150583 | 1204.7827724141437 | 1173.7802575888932 | 1192.108232307853 | 970.2673230526178 | 954.5544635222423 | 1525.8005337576826 | 1128.6292881157908 | 935.3159102129033 | 1356.895419190806 | 995.8643089205806 |
| SE_293_4_41 | SE_293 | SE_293_4 | GCTCCAGGCCTGCACAGCCC | 538.6370903190846 | 539.6719417023039 | 594.4911713059955 | 477.58021567086894 | 555.7515246702092 | 421.02671339604666 | 375.58005385486194 | 381.92281018469873 | 406.69725724392777 | 532.9040766684585 | 234.7569929395858 | 604.7148076312617 |
| SE_293_4_44 | SE_293 | SE_293_4 | CTCCCCAGGCCCCCTCCCGG | 17.340429467012783 | 21.986634661945715 | 29.62580587903632 | 27.32747828089441 | 44.54496953463509 | 45.91443582302566 | 17.21856980423436 | 26.46989773557318 | 50.615160836034676 | 36.45034724134463 | 23.47569929395858 | 8.124767089095647 |
| SE_293_4_48 | SE_293 | SE_293_4 | GGAAGAAGTCCCCCGGGAGG | 23.843090517142578 | 55.96597913949819 | 39.50107450538176 | 32.53271223916001 | 34.999618920070425 | 37.251334724341575 | 24.75169409358689 | 18.9070698111237 | 32.855455279531284 | 18.954180565499207 | 0.0 | 33.659749369110536 |
| SE_293_4_52 | SE_293 | SE_293_4 | GGAGGAGGAAGAAGTCCCCC | 2972.799876751004 | 3112.1081971499525 | 2876.6657508544267 | 2957.874196784428 | 3027.997333842456 | 3238.267190688112 | 2972.3556124559564 | 3478.900845246761 | 2764.2981698697536 | 3398.6303767829736 | 3487.315130117547 | 2872.685506501675 |
| SE_294_1_18 | SE_294 | SE_294_1 | CTGAGTGGACGGGAAGGCGG | 1850.0070687619263 | 1694.969653939088 | 1662.9952366765722 | 1774.9847797685702 | 1900.585366811097 | 1706.6309164407653 | 1661.5919861086156 | 1565.5053803610424 | 1642.7727639765642 | 1894.689049605094 | 1588.131057236298 | 2228.507544437663 |
| SE_294_1_2 | SE_294 | SE_294_1 | GCTCGCTGTCCCGGTGCCAG | 2214.156087569195 | 2474.495791953527 | 2437.2162969820547 | 2354.0670576256184 | 2428.7614341503413 | 2649.1763159775937 | 2194.291489427116 | 2463.591196389418 | 2719.01092070067 | 3023.191800197124 | 1957.8733211161457 | 2442.0728507796057 |
| SE_294_1_26 | SE_294 | SE_294_1 | CCGCAGAGTGCGAACTGCAG | 91.0372547018171 | 40.97509186998975 | 83.93978332393624 | 96.29682822791364 | 51.96913112374093 | 370.78072702367893 | 157.11944946363852 | 47.26767452780925 | 231.7641575123693 | 3.6450347241344634 | 93.90279717583432 | 94.01516203096391 |
| SE_294_1_30 | SE_294 | SE_294_1 | TCCCCCAAAGGCGGACGCAG | 2.167553683376598 | 3.9975699385355847 | 4.93763431317272 | 7.807850937398403 | 0.0 | 0.0 | 6.456963676587884 | 0.0 | 13.319779167377547 | 17.496166675845423 | 0.0 | 0.0 |
| SE_294_1_8 | SE_294 | SE_294_1 | CAGCGGGAGCTGGAAAGCTT | 2511.1109421917886 | 2474.495791953527 | 2545.8442518718543 | 2818.6341884008234 | 2697.0918458708816 | 2076.5453333545756 | 2786.1798264476724 | 2984.480969685876 | 2316.753589845868 | 3510.168439341488 | 3103.487446661324 | 3104.821709047265 |
| SE_294_2_11 | SE_294 | SE_294_2 | GCATTGAAGTCCGGCGGTGG | 831.2568375749253 | 1186.2788792604347 | 1037.8907326289059 | 892.6976238425507 | 1033.019055398442 | 908.7593052519608 | 731.7892166799603 | 793.1515785766392 | 1145.501008394469 | 489.89266692367187 | 422.56258729125443 | 402.75631141659846 |
| SE_294_2_14 | SE_294 | SE_294_2 | CTCCTCGCTCCCGCCACCGC | 120.29922942740117 | 117.92831318679976 | 124.42838469195254 | 105.40598765487843 | 171.8163110621639 | 137.743307469077 | 154.96712823810924 | 55.775855942814914 | 105.67024806119521 | 152.36245146882055 | 11.73784964697929 | 91.69380000550801 |
| SE_294_2_15 | SE_294 | SE_294_2 | CGGCGGTGGCGGGAGCGAGG | 836.6757217833667 | 847.484826969544 | 928.2752508764714 | 789.894253166805 | 891.959985205431 | 969.4010129427494 | 745.7793046459007 | 552.086438484812 | 868.449601713016 | 688.9115628614136 | 659.6671501602361 | 875.1534835968739 |
| SE_294_2_5 | SE_294 | SE_294_2 | GCACTCACGCTGGCGAGCTG | 2883.9301757325634 | 2645.3919068259233 | 2722.611560283438 | 2809.5250289738588 | 2921.9378825695158 | 2455.1228513670703 | 2784.027505222143 | 3324.8082262861026 | 2071.669653166121 | 2963.4132307213185 | 3449.7540112472134 | 3014.2885900544848 |
| SE_294_2_6 | SE_294 | SE_294_2 | CACGCTGGCGAGCTGGGGAC | 7665.553601261338 | 7213.614954087463 | 7246.472118012284 | 7278.218382144878 | 7423.100994593118 | 7256.213480257791 | 7693.472220654465 | 8343.689907648888 | 7341.862277058503 | 7350.577024689559 | 7702.37693834781 | 6120.270980114477 |
| SE_294_3_1 | SE_294 | SE_294_3 | TCAACCCCAGAGATAAAGCC | 642.6796671211613 | 685.5832444588527 | 754.4705230527917 | 737.8419135841491 | 686.2046497359262 | 759.7539663545945 | 592.9644976333208 | 520.889773296458 | 771.6592064300726 | 676.5184447993564 | 489.4683302790364 | 632.5711519367325 |
| SE_294_3_2 | SE_294 | SE_294_3 | ACCCCAGAGATAAAGCCGGG | 49.85373471766175 | 64.96051150120326 | 69.12688038441809 | 45.545797134824014 | 118.78658542569356 | 39.8502650539468 | 58.112673089290965 | 69.95615830115769 | 47.95120500255917 | 20.412194455152996 | 126.76877618737633 | 92.85448101823596 |
| SE_294_3_3 | SE_294 | SE_294_3 | CAGCGCTGACTGTTTAGAAG | 1270.1864584586863 | 1293.2138751162618 | 1346.9866406335182 | 1267.474468837674 | 1446.6509153629108 | 1516.0426922697152 | 1207.4522075219345 | 1659.0953759261047 | 1449.191973410677 | 1474.7810493848037 | 1297.0323859912116 | 1377.728362108076 |
| SE_294_3_30 | SE_294 | SE_294_3 | TCACCAGAGGGGCCAGCAGA | 471.44292613441 | 651.6038999813003 | 654.7303099267027 | 636.3398513979698 | 688.325838761385 | 579.5614635019655 | 652.1533313353764 | 803.5504669727572 | 828.4902642108834 | 1328.2506534745985 | 626.8011711486942 | 724.2649519422405 |
| SE_294_3_33 | SE_294 | SE_294_3 | TGAAGTGAAAGCTCACCAGA | 275.27931778882794 | 344.7904071986942 | 300.2081662409014 | 299.3009526002721 | 415.75304898992744 | 347.3903540572319 | 245.3646197103396 | 268.48039131795656 | 371.17784613092095 | 279.2096598686999 | 334.5287149389098 | 389.988820276591 |
| SE_295_1_11 | SE_295 | SE_295_1 | GCCCGAGGAACAAAAAGCAG | 315.379060931295 | 351.78615459113144 | 324.89633780676496 | 369.57161103685775 | 312.875381255175 | 189.7219140611815 | 293.79184728474877 | 399.88452650526625 | 276.1634214036278 | 357.9424099100043 | 323.96465025662843 | 235.61824558377373 |
| SE_295_1_13 | SE_295 | SE_295_1 | GAAGGAAACCTTTTTCACCG | 1109.787485888818 | 1338.1865369247869 | 1337.1113720071726 | 1175.0815660784597 | 1499.6806409993812 | 1231.8929762328771 | 1306.458983896282 | 1591.0299246060595 | 1343.5217253494818 | 1368.3460354400775 | 1463.7098509783175 | 1086.3974279133606 |
| SE_295_1_17 | SE_295 | SE_295_1 | TGAGGAAGTGGTTAAAGGCG | 5212.966608520718 | 5768.493421306849 | 5742.468706219874 | 5448.5786458145185 | 5903.269057851878 | 5088.705585367033 | 5277.491644997831 | 5780.836594751071 | 6008.108389765099 | 4514.740009312946 | 5370.066213493025 | 5227.707281326684 |
| SE_295_1_22 | SE_295 | SE_295_1 | TTAGATAAAGCCGGAGCCCA | 126.80189047753097 | 197.87971195751143 | 186.64257703792882 | 156.15701874796804 | 142.1196647057405 | 356.9197652657844 | 87.16900963393644 | 227.8301912240406 | 261.9556569584251 | 98.4159375516305 | 278.1870366334092 | 685.9624785222181 |
| SE_295_1_4 | SE_295 | SE_295_1 | ATTAGAGACAAAGCACTCGG | 37.93218945909046 | 72.95565137827442 | 66.16429979651446 | 41.641871666124814 | 50.90853661101153 | 87.49732109670929 | 12.913927353175769 | 61.44797688615203 | 102.11830694989453 | 53.94651391719005 | 28.170839152750297 | 23.21362025455899 |
| SE_296_1_12 | SE_296 | SE_296_1 | TTACCCAATCCATTTTCCAG | 4.335107366753196 | 27.982989569749094 | 13.825376076883616 | 16.917010364363207 | 9.545350614564661 | 10.395721318420904 | 2.152321225529295 | 0.945353490556185 | 8.879852778251697 | 13.122125006884067 | 8.216494752885502 | 41.78451645820618 |
| SE_296_1_13 | SE_296 | SE_296_1 | TACCCAATCCATTTTCCAGT | 238.43090517142576 | 294.8207829669994 | 256.75698428498146 | 269.37085734024487 | 257.72446659324584 | 155.93581977631356 | 249.66926216139822 | 250.518674997389 | 277.051406681453 | 242.03030568252836 | 147.89690555193906 | 155.53125570554522 |
| SE_296_1_18 | SE_296 | SE_296_1 | AGTCTCTCCTCTAATCCCTT | 6.5026610501297935 | 16.989672238776237 | 27.650752153767232 | 10.410467916531204 | 9.545350614564661 | 7.796790988815679 | 10.761606127646475 | 3.78141396222474 | 0.8879852778251698 | 31.347298627556384 | 1.173784964697929 | 1.1606810127279494 |
| SE_296_1_26 | SE_296 | SE_296_1 | TTTAGCTCCTTTCAGAGCAT | 13.005322100259587 | 6.9957473924372735 | 6.912688038441808 | 2.602616979132801 | 16.969512203670508 | 12.128341538157722 | 16.142409191469714 | 2.836060471668555 | 11.543808611727208 | 0.7290069448268927 | 0.0 | 19.73157721637514 |
| SE_296_1_5 | SE_296 | SE_296_1 | ACTGTCCCTGATAACAGCAT | 1431.6692078702429 | 1247.2418208231024 | 1266.99696476012 | 1250.5574584733108 | 1245.1379579443235 | 1425.0801307335323 | 1307.5351445090466 | 1330.1123612125523 | 1359.505460350335 | 1043.2089380472835 | 1097.4889419925637 | 1434.6017317317455 |
| SE_297_1_14 | SE_297 | SE_297_1 | CTGGCCCGAGTGAGGCGATA | 727.2142607728485 | 704.5717016668968 | 778.1711677560207 | 806.8112635311683 | 860.1421498235488 | 966.8020826131442 | 724.2560923906077 | 830.9657181988866 | 815.1704850435059 | 766.9153059578911 | 949.5920364406246 | 836.8510101768516 |
| SE_297_1_26 | SE_297 | SE_297_1 | GGAGTAGTAAGTGGAAACTG | 858.3512586171328 | 1004.3894470570657 | 966.7887985192186 | 835.4400503016291 | 1019.2313267329599 | 1049.9678531605114 | 872.7662569521291 | 929.2824812167298 | 864.8976606017154 | 879.9113824060595 | 1316.9867303910764 | 927.3841291696316 |
| SE_297_1_27 | SE_297 | SE_297_1 | TTTCCACTTACTACTCCCAA | 20.59175999207768 | 12.992102300240651 | 19.75053725269088 | 2.602616979132801 | 23.33307928004695 | 18.192512307236584 | 21.52321225529295 | 33.08737216946648 | 19.535676112153734 | 4.374041668961356 | 19.954344399864794 | 2.321362025455899 |
| SE_297_1_6 | SE_297 | SE_297_1 | ATAACCACTTACAGCCCGAC | 355.47880407376203 | 362.77947192210434 | 409.8236479933358 | 433.33572702561133 | 373.3292684807512 | 466.0748391092039 | 399.2555873356842 | 241.06514009182717 | 481.288020581242 | 439.5911877306163 | 342.74520969179525 | 511.8603266130257 |
| SE_297_1_7 | SE_297 | SE_297_1 | ATGCTCTTGTATTTAAGTGA | 151.72875783636184 | 182.888824688003 | 161.95440547206522 | 135.33608291490566 | 164.39214947305805 | 181.92512307236584 | 134.52007659558095 | 114.38777235729839 | 272.61148029232714 | 255.15243068941243 | 169.0250349165018 | 196.15509115102344 |
| SE_297_2_1 | SE_297 | SE_297_2 | CCTTCAGTCAGGCAGACAGT | 1185.651864806999 | 1047.3633238963232 | 1066.5290116453075 | 1253.1600754524436 | 1118.927210929524 | 1058.6309542591955 | 1104.1407886965283 | 1156.1673189502142 | 911.9608803264493 | 1113.1936047506651 | 1113.9219314983347 | 863.5466734695943 |
| SE_297_2_22 | SE_297 | SE_297_2 | AATGTCCCAGGGAGCCTCCA | 81.28326312662242 | 144.91191027191493 | 137.26623390620162 | 78.07850937398402 | 119.84717993842297 | 205.31549603881288 | 47.35106696164449 | 86.02716764061283 | 136.74973278507613 | 147.9884097998592 | 98.59793703462604 | 67.31949873822107 |
| SE_297_2_5 | SE_297 | SE_297_2 | ATATGCTAATATTTTATGGA | 10.837768416882989 | 9.993924846338961 | 23.700644703229056 | 16.917010364363207 | 21.211890254588134 | 0.8663101098684087 | 22.599372868057596 | 10.398888396118036 | 17.759705556503395 | 8.748083337922711 | 36.3873339056358 | 11.606810127279495 |
| SE_297_2_6 | SE_297 | SE_297_2 | TTGTTCCCTAAAAAATGCAG | 32.51330525064897 | 18.98845720804403 | 15.800429802152705 | 16.917010364363207 | 20.15129574185873 | 4.331550549342044 | 37.665621446762664 | 1.89070698111237 | 85.2465866712163 | 1.4580138896537853 | 8.216494752885502 | 164.8167038073688 |
| SE_297_2_7 | SE_297 | SE_297_2 | TCTGGCCGCTGCATTTTTTA | 1011.1637932951829 | 1225.2551861611566 | 1207.7453530020473 | 1178.9854915471587 | 1173.0175310787238 | 1482.2565979848473 | 1042.7996337689433 | 1139.1509561202029 | 1053.1505395006513 | 1287.4262645642925 | 1024.714274181292 | 1850.1255342883514 |
| SE_297_3_20 | SE_297 | SE_297_3 | GCTTCCTCAGAAGAGGCTAT | 152.81253467805016 | 147.91008772581662 | 143.19139508200888 | 184.78580551842887 | 166.51333849851687 | 359.5186955953896 | 205.54667703804768 | 190.01605160179318 | 139.41368861855165 | 177.8776945377618 | 346.26656458588906 | 120.71082532370674 |
| SE_297_3_3 | SE_297 | SE_297_3 | CTGATCGTCATTGCTCCAGA | 484.4482482346696 | 562.6579688488836 | 530.3019252347501 | 432.03441853604494 | 486.8128813427977 | 535.3796478986766 | 385.2654993697438 | 575.7202757487166 | 475.07212363646585 | 320.7630557238328 | 498.85860999661986 | 374.8999671111277 |
| SE_297_3_4 | SE_297 | SE_297_3 | GGTGTCTGTGAGATTCAGGC | 99.7074694353235 | 126.92284554850481 | 138.25376076883617 | 111.91253010271043 | 139.9984756802817 | 65.83956834999907 | 142.05320088493346 | 63.33868386726439 | 87.02255722686664 | 113.72508339299526 | 98.59793703462604 | 146.24580760372163 |
| SE_297_3_5 | SE_297 | SE_297_3 | GAGCAGTGTGATACACAATG | 1333.0455152766076 | 1292.2144826316278 | 1347.9741674961526 | 1371.5791480029861 | 1403.1665403410052 | 1483.1229080947157 | 1258.031756321873 | 1082.4297466868318 | 1385.2570334072648 | 1250.975917322948 | 984.8055853815624 | 1758.4317342828433 |
| SE_297_3_6 | SE_297 | SE_297_3 | GCCCTAGCCAGTTTGGCCCA | 230.84446727960767 | 225.86270152726053 | 226.1436515433106 | 279.7813252567761 | 230.14900926228125 | 200.1176353796024 | 247.51694093586892 | 366.7971543357998 | 119.87801250639792 | 181.52272926189627 | 376.7849736680352 | 285.52752913107554 |
| SE_297_4_10 | SE_297 | SE_297_4 | GCTCTAAGCCTCTCATAACA | 84.53459365168732 | 121.92588312533533 | 101.71526685135804 | 87.18766880094883 | 64.69626527649382 | 163.73261076512924 | 52.73187002546773 | 32.14201867891029 | 53.27911666951019 | 113.72508339299526 | 29.344624117448227 | 59.19473164912542 |
| SE_297_4_21 | SE_297 | SE_297_4 | GTTCTGTGTAAACAGCTCTG | 22.759313675454276 | 17.98906472341013 | 21.725590977959968 | 15.615701874796805 | 25.454268305505764 | 0.8663101098684087 | 7.533124289352532 | 84.13646065950047 | 36.40739639083196 | 2.187020834480678 | 76.29602270536539 | 5.803405063639747 |
| SE_297_4_25 | SE_297 | SE_297_4 | GGTGGCCGCCTGTAATGTGA | 867.0214733506391 | 812.5060900073576 | 838.4103063767279 | 763.8680833754771 | 831.5060979798549 | 878.4384514065664 | 967.4683908754181 | 890.5229881039263 | 723.7080014275134 | 1309.2964729090993 | 618.5846763958086 | 579.1798253512468 |
| SE_297_4_6 | SE_297 | SE_297_4 | ATAAAGGTTTCCCAGGACAG | 91.0372547018171 | 103.93681840192521 | 119.49075037877982 | 78.07850937398402 | 88.02934455654076 | 183.65774329210265 | 92.54981269775968 | 78.46433971616335 | 124.31793889552377 | 35.72134029651774 | 174.89395973999143 | 139.28172152735394 |
| SE_297_4_8 | SE_297 | SE_297_4 | AGAGGCTTAGAGCACTGGAG | 906.0374396514179 | 1156.2971047214178 | 1047.7660012552512 | 676.6804145745282 | 812.4153967507256 | 796.1389909690677 | 563.9081610886752 | 473.6220987686487 | 667.7649289245277 | 225.99215289633673 | 458.9499211968903 | 107.94333418369929 |
| SE_298_1_11 | SE_298 | SE_298_1 | GAATACTATAGCTCTTAAGG | 0.0 | 6.9957473924372735 | 7.900214901076352 | 1.3013084895664004 | 0.0 | 5.197860659210452 | 0.0 | 0.0 | 61.27098416993672 | 11.664111117230282 | 0.0 | 13.928172152735392 |
| SE_298_1_12 | SE_298 | SE_298_1 | GTAGGGGGTGGTGTTCATCA | 3860.4131100937207 | 3831.670786086358 | 3986.6459444556544 | 3871.3927564600413 | 3829.806785465888 | 3699.144169138105 | 3876.33052717826 | 3533.7313476990194 | 4050.1008521605995 | 3878.316946479069 | 4117.637656160335 | 3359.0108508346857 |
| SE_298_1_2 | SE_298 | SE_298_1 | GCTTGTAGTGGTAACCCGTT | 2267.2611528119214 | 2210.6561760101786 | 2349.3264062075805 | 2117.2289125245334 | 2233.6120438081307 | 2157.978483682206 | 2005.9633821933028 | 2227.252823750372 | 2545.8537915247616 | 1580.4870563847032 | 2345.2223594664624 | 2553.4982280014888 |
| SE_298_1_22 | SE_298 | SE_298_1 | GAGCAACCTTGCAAGAACCT | 862.6863659838859 | 856.479359331249 | 802.8593393218842 | 802.9073380624691 | 727.567835732373 | 1301.19778502235 | 821.110547539426 | 876.3426857455835 | 912.8488656042746 | 1235.666771481583 | 717.1826134304347 | 1228.0005114661706 |
| SE_298_1_4 | SE_298 | SE_298_1 | CTAAACCTTGAGCTAGATGC | 428.0918524668781 | 372.77339676844326 | 360.4473048616086 | 346.1480582246625 | 371.20807945529236 | 423.62564372565186 | 427.23576326756506 | 338.43654961911426 | 279.7153625149285 | 307.6409307169487 | 329.83357508011807 | 405.07767344205433 |
| SE_298_2_1 | SE_298 | SE_298_2 | CTGACATGCCATATTCCAAG | 428.0918524668781 | 447.7278331159855 | 428.5866583833921 | 412.51479119254896 | 407.2682928880922 | 151.60426922697152 | 408.94103285056605 | 215.5405958468102 | 207.78855501108973 | 636.4230628338773 | 145.5493356225432 | 184.54828102374395 |
| SE_298_2_12 | SE_298 | SE_298_2 | ACAGCCACTGGGTTGTAATG | 3356.456878708662 | 3301.992769230393 | 3425.730686479233 | 3527.8473152145116 | 3510.5678371343365 | 3505.95701463745 | 3595.4526072466874 | 3250.125300532164 | 3315.737027399184 | 3856.446738134262 | 3611.7363363755276 | 2927.2375140998884 |
| SE_298_2_18 | SE_298 | SE_298_2 | TTTCAGCCCGAAGTGTGTGT | 193.9960546622055 | 215.86877668092157 | 196.51784566427426 | 189.99103947669445 | 137.87728665482288 | 298.0106777947326 | 206.6228376508123 | 150.31120499843342 | 381.833669464823 | 193.18684037912655 | 132.63770101086598 | 289.0095721692594 |
| SE_298_2_19 | SE_298 | SE_298_2 | TTATGTCCTGTCCTTGAAGG | 208.0851536041534 | 180.8900397187352 | 187.63010390056337 | 176.97795458103047 | 198.33117388039906 | 181.05881296249743 | 217.3844437784588 | 149.36585150787724 | 114.5501008394469 | 69.2556597585548 | 106.81443178751154 | 54.55200759821362 |
| SE_298_2_26 | SE_298 | SE_298_2 | TCCTGTAACCACTCAGTGCA | 579.82061030324 | 663.596609796907 | 638.92988012455 | 610.3136816066418 | 557.8727136956679 | 625.4758993249911 | 605.8784249864965 | 758.1734994260604 | 504.37563780469645 | 586.1215836408217 | 720.7039683245284 | 329.6334076147376 |
| SE_298_3_10 | SE_298 | SE_298_3 | GCGTATGATGAGCAGTGAAG | 4146.530196299432 | 3792.694479185636 | 4001.4588473951726 | 4459.584193744055 | 3782.0800323930644 | 4489.218989338094 | 4644.709204692219 | 3957.2497114681905 | 4282.752994950793 | 3889.9810575962993 | 3706.81291851606 | 5025.748785112021 |
| SE_298_3_17 | SE_298 | SE_298_3 | ATAGCCAACCAGCAACCGCT | 339.22215144843756 | 522.6822694635277 | 487.83827014146476 | 426.82918457777936 | 468.7827746263978 | 397.6363404295996 | 264.7355107401033 | 410.2834149013843 | 522.1353433611998 | 395.8507710410027 | 490.64211524373434 | 678.9983924458504 |
| SE_298_3_21 | SE_298 | SE_298_3 | ATAGGGTAACTTCCAGACAT | 442.18095140882593 | 570.6531087259547 | 546.1023550369028 | 533.5364807222242 | 493.17644841917416 | 417.561472956573 | 467.053705939857 | 397.0484660335977 | 602.9420036432903 | 733.380986495854 | 284.05596145689884 | 806.6733038459248 |
| SE_298_3_34 | SE_298 | SE_298_3 | CCTGCATCCTGTTTTATCAG | 1830.4990856115369 | 1833.8852093031994 | 1820.0120078354646 | 1980.5915211200615 | 1919.6760680402263 | 2156.2458634624695 | 1991.9732942273624 | 2306.6625169570916 | 2092.9812998339253 | 1942.074501018842 | 2024.7790641039276 | 2842.507800170748 |
| SE_298_3_8 | SE_298 | SE_298_3 | GCACTTCTAGAGACAGACCT | 443.2647282505143 | 526.6798394020633 | 496.72601190517565 | 484.08675811870097 | 534.539634415621 | 508.5240344927559 | 554.2227155737935 | 563.4306803714862 | 469.7442119695148 | 480.41557664092227 | 635.0176659015796 | 435.25537977298103 |
| SE_299_1_13 | SE_299 | SE_299_1 | CAGAGCACCAGACTCCGACG | 3281.676276632169 | 3633.7910741288465 | 3501.7702549020933 | 3573.3931123493358 | 3739.656251883888 | 2735.8073269644347 | 3644.955995433861 | 3639.6109386413123 | 3806.7928860365028 | 3744.9086755757476 | 3405.150182588692 | 3001.521098914477 |
| SE_299_1_17 | SE_299 | SE_299_1 | AGGACACCGGCCCAGTGCAG | 5154.442659069549 | 6217.220646907468 | 5976.512572664261 | 5574.80556930246 | 6173.7206585978765 | 6261.689474128858 | 5508.86617674223 | 6101.311428049618 | 6324.2311486708595 | 5845.177683622025 | 7433.580181431985 | 7236.846114358765 |
| SE_299_1_18 | SE_299 | SE_299_1 | GAGGCAGCGTGATTGCACCA | 23441.00930887622 | 19200.328414786414 | 18512.17856694716 | 21280.297729879345 | 18686.614719779416 | 18290.40534965171 | 23050.284164805984 | 19385.41867734513 | 19382.942644367806 | 16879.426800521873 | 19623.337039819977 | 18731.070183403648 |
| SE_299_1_22 | SE_299 | SE_299_1 | GCAATGTCAACTGGGCAACA | 5763.525244098373 | 5140.874940956762 | 5228.95473764991 | 5936.569329401918 | 5258.427594112399 | 5013.336605808481 | 5839.247484860977 | 5322.340151831321 | 5752.36862975145 | 5519.311579284405 | 5125.918940835856 | 5072.176025621139 |
| SE_299_1_37 | SE_299 | SE_299_1 | CTTCCTGAGTTCCTGCTGAA | 96.4561389102586 | 137.91616287947767 | 107.6404280271653 | 122.32299801924164 | 125.15015250207 | 103.09090307434064 | 105.46374005093546 | 74.68292575393862 | 118.99002722857276 | 61.23658336545898 | 62.210603128990236 | 291.3309341947153 |
| SE_299_2_2 | SE_299 | SE_299_2 | TCTCCTCCCACCCAGCAATA | 139.80721257779055 | 138.91555536411158 | 147.14150253254707 | 141.84262536273764 | 116.66539640023474 | 141.20854790855063 | 123.75847046793446 | 306.29453094020397 | 128.75786528464963 | 163.29755564122397 | 355.6568443034725 | 171.7807898837365 |
| SE_299_2_42 | SE_299 | SE_299_2 | GGGCCGGGTTCACGTGGCCA | 1098.9497174719352 | 1229.2527560996923 | 1219.5956753536618 | 1437.9458809708724 | 1299.2282780935232 | 1196.3742617282724 | 1392.5518329174538 | 1380.2160962120302 | 1528.2226631371173 | 1054.1440422196868 | 1555.265078224756 | 1338.2652076753257 |
| SE_299_2_5 | SE_299 | SE_299_2 | CTGGTGTCCGTATTGCTGGG | 479.0293640262281 | 436.7345157850126 | 469.0752597514084 | 395.5977808281857 | 391.3593751971511 | 460.01066834012505 | 393.87478427186096 | 280.76998669518696 | 599.3900625319897 | 195.37386121360723 | 304.01030585676364 | 315.70523546200224 |
| SE_299_2_53 | SE_299 | SE_299_2 | GCCGGCTTCAGAGACCACAG | 562.4801808362271 | 861.4763217544186 | 791.9965438329043 | 680.5843400432275 | 776.3551833179257 | 890.5667929447242 | 636.0109221439067 | 738.3210761243805 | 902.1930422703725 | 1292.5293131780807 | 838.0824647943214 | 1091.0401519642726 |
| SE_299_2_54 | SE_299 | SE_299_2 | GGCCGGCTTCAGAGACCACA | 335.9708209233727 | 400.7563863381924 | 432.5367658339303 | 378.6807704638225 | 428.48018314268035 | 397.6363404295996 | 408.94103285056605 | 587.0645176353909 | 469.7442119695148 | 588.3086044753023 | 700.7496239246636 | 425.96993167115744 |
| SE_299_3_15 | SE_299 | SE_299_3 | GAAGCAGTCATCCCTCCACG | 3344.5353334500905 | 3369.951458185498 | 3714.08853036852 | 3589.0088142241325 | 3594.3548036399593 | 2903.0051781690377 | 3408.2006606256386 | 3225.5461097777034 | 3380.5599526804212 | 3841.866599237724 | 3619.952831128413 | 3849.978919218608 |
| SE_299_3_2 | SE_299 | SE_299_3 | AGAGCCAGCCTTGTGTGGCG | 1959.4685297724445 | 2155.689589355314 | 2160.7087754443824 | 2059.971338983612 | 2001.3418455203905 | 2278.3955889539147 | 1901.5758027551321 | 2307.6078704476477 | 2077.8855501108974 | 1843.6585634672115 | 2239.5817126436486 | 2306.2731722904355 |
| SE_299_3_20 | SE_299 | SE_299_3 | GAGGTCCTCATGGCTCCCGG | 438.92962088376106 | 618.6239479883817 | 600.4163324818028 | 528.3312467639586 | 648.0232472776676 | 366.44917647433687 | 394.95094488462564 | 656.0753224459924 | 581.6303569754862 | 721.7168753786237 | 545.810008584537 | 549.00211902032 |
| SE_299_3_27 | SE_299 | SE_299_3 | GGAGAGAAGAGAAGGCTCCG | 683.8631871053166 | 947.4240754329336 | 952.963422442335 | 761.2654663963442 | 836.8090705435019 | 1049.101543050643 | 608.0307462120259 | 934.954602160067 | 882.6573661582188 | 969.5792366197672 | 1078.7083825573968 | 1138.6280734861184 |

| | | | | | | | | | | | | | | | |
|---|---|---|---|---|---|---|---|---|---|---|---|---|---|---|---|
| SE_299_3_33 | SE_299 | SE_299_3 | GGGATTGCCAGACCCCCAGG | 20.59175999207768 | 38.97630690072195 | 39.50107450538176 | 11.711776406097604 | 42.42378050917627 | 10.395721318420904 | 18.294730416999005 | 96.42605603673087 | 63.93494000341222 | 2.187020834480678 | 96.25036710523018 | 2.321362025455899 |
| SE_299_4_17 | SE_299 | SE_299_4 | GCCTGCATCCCAGCTGCCAG | 1930.2065550468603 | 2091.7284703387445 | 2184.4094201476114 | 1964.9758192452646 | 2335.4291170301535 | 2286.1923799427304 | 2052.2382885421825 | 2624.3012897839694 | 2030.8223303861632 | 2851.1461612179774 | 2457.9057160774632 | 2589.4793393960554 |
| SE_299_4_21 | SE_299 | SE_299_4 | GAGGGTTAGGAAGAGAACAG | 1168.3114353399862 | 1397.150693518187 | 1361.7995435730363 | 1236.2430650880804 | 1129.5331560568181 | 1285.6042030447186 | 1248.3463108069911 | 1244.0851935719395 | 1411.89659174202 | 1168.598132557509 | 1030.5831990047816 | 1175.7698658934128 |
| SE_299_4_27 | SE_299 | SE_299_4 | CGTCCTGGTGAGAACATGGC | 10758.652707439744 | 10093.864094802351 | 10001.672064762663 | 9469.621878574697 | 10092.617383133034 | 10092.51277996696 | 9768.309882064705 | 9568.868031409706 | 9568.929353844029 | 10354.814644321183 | 9900.876177227032 | 9741.59573982568 |
| SE_299_4_39 | SE_299 | SE_299_4 | CTAGAGGTGCCCGTATCCCT | 1683.1054351419282 | 1536.0662488822984 | 1508.9410461055834 | 1660.46963268672 7 | 1476.3475617193342 | 1583.6148808394512 | 1630.3833283384408 | 2007.930813941337 | 1436.7601795211247 | 1826.891403736193 | 1779.4580064820605 | 1101.486281078824 |
| SE_299_4_5 | SE_299 | SE_299_4 | GCCCGCTCCACAGAACCCCA | 348.97614302363223 | 408.75152621526354 | 479.9380552403884 | 387.78992989078733 | 504.84298805919764 | 439.2192257032832 | 321.7720232166296 | 470.7860382969801 | 400.48136029915156 | 411.15991688236744 | 692.5331291717781 | 558.2875671221436 |
| SE_3_1_12 | SE_3 | SE_3_1 | TACAGGAGACTGGAGATGGA | 928.7967533268721 | 1058.356641227296 | 1174.169439672473 | 957.7630483208707 | 1157.1086133877827 | 1186.84485051972 | 1006.2101729349454 | 1561.7239663988175 | 1100.2137592253853 | 1996.021014936032 | 1582.2621324128083 | 1866.3750684665426 |
| SE_3_1_16 | SE_3 | SE_3_1 | GGTGCTCAGCAGATACCCAG | 402.0812082663589 | 512.6883446171887 | 507.58880739415565 | 426.82918457777936 | 465.6009910882096 | 556.1710905355184 | 389.5701418208024 | 358.2889729207941 | 491.05585863731886 | 209.95400011014507 | 152.59204541073078 | 290.17025318198733 |
| SE_3_1_17 | SE_3 | SE_3_1 | AATTGCTGTTCCAGTCAACA | 121.38300626908948 | 54.96658665486429 | 78.01462214812898 | 58.55888203048802 | 47.72675307282331 | 36.38502461447317 | 74.25508228076067 | 9.45353490556185 | 91.46248361599248 | 59.7785694758052 | 31.692194046844083 | 49.909283547301825 |
| SE_3_1_20 | SE_3 | SE_3_1 | GAGACAGCCTAGAAATGGAA | 919.0427617516775 | 1017.3815493573063 | 1203.7952455515092 | 966.8722077478355 | 1009.6859761183953 | 809.1336426170938 | 933.0312512669493 | 680.6545132004532 | 1108.205626725812 | 865.3312435095215 | 496.511040067224 | 796.2271747313733 |
| SE_3_1_9 | SE_3 | SE_3_1 | TCTGAGCGAACCTTTCCATG | 2294.3555738541286 | 2515.4708838235165 | 2553.744466772931 | 2122.434146482799 | 2366.1863578993066 | 1671.9785120460288 | 1878.9764298870746 | 1942.70142309296 | 2567.165438192566 | 1587.0481188881454 | 1671.4697897298508 | 1413.7094735026424 |
| SE_3_2_12 | SE_3 | SE_3_2 | TGAGGTTAGCAGTTACCCAG | 1935.625439255302 | 2768.3171824358924 | 2549.7943593223927 | 1441.8498064395717 | 1844.3738576364383 | 1449.3368138098479 | 1245.117828968697 | 1018.1457093290112 | 2162.2441515042883 | 411.15991688236744 | 536.4197288669536 | 555.9662050966878 |
| SE_3_2_20 | SE_3 | SE_3_2 | TGGATTCTATCACCCTAAGA | 0.0 | 0.9993924846338962 | 1.975053725269088 | 1.3013084895664004 | 0.0 | 1.7326202197368175 | 0.0 | 0.0 | 0.0 | 0.0 | 0.0 | 0.0 |
| SE_3_2_24 | SE_3 | SE_3_2 | GTGCCAGGTACAAGATACAG | 254.68755779675024 | 304.8147078133383 | 342.6718213341868 | 212.11328379932328 | 317.11775930609264 | 272.02137449868036 | 134.52007659558095 | 117.22383282896693 | 309.90686196098426 | 364.50347241344633 | 131.46391604616804 | 85.89039494186825 |
| SE_3_2_28 | SE_3 | SE_3_2 | TCTCCCACCACAGCAGAACA | 249.26867358830876 | 307.81288526724 | 308.10838114197776 | 282.3839422359089 | 335.1478660224925 | 171.52940175394494 | 233.5268529699285 | 206.08706094124832 | 261.9556569584251 | 153.09145841364744 | 130.29013108147012 | 155.53125570554522 |
| SE_3_2_9 | SE_3 | SE_3_2 | ATACTGATGTGACATCCACT | 373.90301038246315 | 399.75699385355847 | 403.8984868175285 | 425.52787608821296 | 309.69359771698674 | 331.79677207960054 | 436.92120878244685 | 280.76998669518696 | 354.30612585224276 | 204.12194455152994 | 221.8453583279086 | 289.0095721692594 |
| SE_3_3_15 | SE_3 | SE_3_3 | GGTTGGGTCATTGTGCCTTG | 521.2966608520718 | 593.6391358725343 | 559.9277311137864 | 573.8770438987826 | 626.8113570230794 | 603.8181465782809 | 475.6629908419742 | 382.86816367525495 | 617.149768088493 | 618.1978892132049 | 497.6848250319219 | 586.1439114276145 |
| SE_3_3_18 | SE_3 | SE_3_3 | GTGACTGATCTTTGACCGTA | 360.89768828220355 | 495.6986723784125 | 457.2249373997939 | 403.40563176558413 | 445.44969534635084 | 384.64168878157346 | 358.3614840506276 | 426.3544242408394 | 453.76047696866175 | 519.7819516615745 | 779.3932165594249 | 285.52752913107554 |
| SE_3_3_20 | SE_3 | SE_3_3 | GGACTAGACTCTCAGAGAAG | 148.47742731129696 | 247.84933618920624 | 299.22063937826687 | 226.4276771845537 | 266.2092226950811 | 218.310147686839 | 156.0432888508739 | 182.45322367734371 | 353.4181405744176 | 260.98448624802757 | 261.75404712763816 | 249.54641773650914 |
| SE_3_3_6 | SE_3 | SE_3_3 | ATAGGGCTAGGAAAAGGGGG | 558.1450734694739 | 754.5413258985916 | 822.6098765745752 | 530.9338637430914 | 639.5384911758323 | 665.3261643789378 | 464.9013847143277 | 587.0645176353909 | 757.4514419848698 | 452.71331273750036 | 507.07510474950533 | 602.3934456058057 |
| SE_30_1_25 | SE_30 | SE_30_1 | GAGACCTAAGGATAAAAAGG | 63.94283365960963 | 124.92406057923702 | 104.67784743926167 | 76.77720088441762 | 82.72637199289373 | 130.81282659012973 | 95.77829453605362 | 87.9178746217252 | 93.23845417164283 | 92.58388199301537 | 125.5949912226784 | 90.53311899278006 |
| SE_30_1_27 | SE_30 | SE_30_1 | TCCCCAGTACTCCTCCGCAG | 97.53991575194691 | 150.9082651797183 | 122.45333096668347 | 79.37981786355043 | 143.1802592184699 | 59.7753975809202 | 66.72195799140815 | 88.86322811228139 | 123.4299536176986 | 72.17168753786237 | 130.29013108147012 | 123.03218734916264 |
| SE_30_1_37 | SE_30 | SE_30_1 | GGTGTCTGGTAGGTGTGTAG | 223.25802938778958 | 342.79162222942637 | 353.53461682316674 | 244.64599603848328 | 266.2092226950811 | 225.24062856578627 | 173.26185865510826 | 192.85211207346174 | 336.5464202957393 | 187.3547848205114 | 304.01030585676364 | 262.31390887651656 |
| SE_30_1_52 | SE_30 | SE_30_1 | GCAGGGGGACTGGGGCCCAG | 5995.453488219669 | 6958.769870505819 | 6820.848040216795 | 6053.687093462895 | 6804.774393671873 | 6109.218894792019 | 5870.456142631152 | 5276.963184284625 | 6249.640385333545 | 5373.510190319026 | 4724.484482909164 | 5495.82459526684 |
| SE_30_1_58 | SE_30 | SE_30_1 | GCAGACAGAACTAAACAGGC | 1880.3528203291985 | 2687.366391180547 | 2821.3642465468924 | 2140.6524653367287 | 2530.5785073723646 | 1587.0801212789247 | 1650.8303799809692 | 2125.1546467703038 | 2399.3362206836086 | 2534.757147163106 | 2454.3843611833695 | 1743.34288111738 |
| SE_30_2_12 | SE_30 | SE_30_2 | GTATAAATGACCCCTCCTCT | 128.96944416090756 | 129.9210230024065 | 146.1539756699125 | 163.96486968536647 | 135.75609762936406 | 170.66309164407653 | 139.90087965940418 | 105.87959094229272 | 126.09390945117411 | 85.29381254474644 | 173.7201747752935 | 110.26469620915519 |
| SE_30_2_19 | SE_30 | SE_30_2 | AGGATTAGTGGACAAATGGA | 927.7129764851838 | 1024.3772967497437 | 1071.4666459584803 | 1017.6232388409252 | 972.565168172866 | 1176.449129201299 | 949.173660458419 | 1253.5387284775013 | 1123.3013764488398 | 1016.2356810886884 | 1279.4256115207427 | 1103.8076431042798 |
| SE_30_2_27 | SE_30 | SE_30_2 | GACAGGCCGACAGAACTTGG | 7009.868612039917 | 6043.326354581171 | 5948.861820510493 | 6678.315168454767 | 6337.052213558205 | 7458.063735857131 | 7360.938591310189 | 6848.1406855890045 | 6036.5239186555045 | 5768.631954415202 | 6840.81877425953 | 7174.169339671455 |
| SE_30_2_3 | SE_30 | SE_30_2 | CTAGAGCAAAGAACTCTCAG | 445.43228193389086 | 491.70110243987693 | 544.1273013116338 | 497.099843014365 | 605.5994667684912 | 419.29409317630984 | 413.2456753016246 | 465.11391735364305 | 409.36121307740325 | 595.5986739235713 | 767.6553669124456 | 327.31204558928175 |
| SE_30_2_45 | SE_30 | SE_30_2 | TTTAGATTCGACTTTCCCCT | 13.005322100259587 | 18.98845720804403 | 19.75053725269088 | 15.615701874796805 | 24.393673792776355 | 19.9251325269734 | 5.380803063823238 | 0.945353490556185 | 16.871720278678225 | 16.038152786191638 | 0.0 | 39.46315443275028 |
| SE_30_3_12 | SE_30 | SE_30_3 | AGACTTCATCCTTTTCCAAG | 53.105065242726646 | 86.94714616314897 | 69.12688038441809 | 48.1484141395682 | 97.57469517110542 | 94.42780197565655 | 94.70213392328898 | 35.92343264113503 | 108.33420389467071 | 78.00374309647752 | 38.73490383503166 | 66.15881772549312 |
| SE_30_3_16 | SE_30 | SE_30_3 | TGTGACCAATGGGGATTGCG | 5288.830987438899 | 4668.162295724929 | 4772.717327112751 | 4529.85485218064 | 4072.6829288809217 | 3566.5987223282386 | 4930.967927687615 | 3529.0045802462387 | 4368.887566899835 | 3499.9623421139117 | 2381.609693372098 | 3565.6120711002604 |
| SE_30_3_2 | SE_30 | SE_30_3 | CCTGGGGTGTTCCCATTCTA | 602.5799239786942 | 782.5243154683407 | 731.7574052121971 | 651.9555532727666 | 636.356707637644 | 613.3475577868334 | 541.3087882206177 | 494.41987556088475 | 837.3701169891351 | 607.9917919856284 | 463.645061055682 | 513.0210076257537 |
| SE_30_3_38 | SE_30 | SE_30_3 | GGGTCCCTGGAACCCCACCG | 0.0 | 1.9987849692677924 | 2.962580587903632 | 0.0 | 0.0 | 1.7326202197368175 | 0.0 | 0.0 | 2.6639558334755096 | 2.187020834480678 | 0.0 | 0.0 |
| SE_30_3_5 | SE_30 | SE_30_3 | GATGGTCTACATGGAATGAG | 1183.4843111236223 | 1201.2697665299431 | 1194.9075037877983 | 1306.5137235246661 | 1116.8060219040653 | 975.4651837118282 | 1369.9524600493962 | 1237.4677191380463 | 1170.3645961735738 | 1819.6013342879241 | 1347.5051394732225 | 921.5807241059919 |
| SE_30_4_15 | SE_30 | SE_30_4 | TTGTTATTTGATCTTCACAA | 7.586437891818092 | 27.982989569749094 | 23.700644703229056 | 11.711776406097604 | 18.030106716399914 | 6.93048087894727 | 15.066248578705064 | 6.617474433893295 | 19.535676112153734 | 19.683187510326103 | 0.0 | 1.1606810127279494 |
| SE_30_4_19 | SE_30 | SE_30_4 | CTGACTACACCAGAACCCAG | 499.6211240183058 | 720.5619814210391 | 753.4829961901571 | 559.5626505135522 | 893.0205797181604 | 693.048087894727 | 459.5205816505045 | 838.5285461233361 | 1063.8063628345535 | 919.2777574267117 | 773.5242917359352 | 602.3934456058057 |
| SE_30_4_27 | SE_30 | SE_30_4 | GATGGGGCTCGGCCACCAGA | 1010.0800164534946 | 1202.269159014577 | 1228.4834171173727 | 1063.1690359757492 | 1420.1360525446755 | 1147.8608955756415 | 1066.4751672497657 | 1328.22165423144 | 1220.9797570096084 | 1339.1857576470018 | 855.6892392647902 | 925.0627671441757 |
| SE_30_4_8 | SE_30 | SE_30_4 | ACTAAGAAGGTCCCTCACCA | 363.0652419655801 | 486.7041400167074 | 488.8257970040993 | 368.27030254729135 | 461.35861303729195 | 529.3154771295978 | 312.0865777017478 | 214.595242356254 | 411.1371836330536 | 197.5608820480879 | 279.3608215981071 | 696.4086076367697 |
| SE_30_4_9 | SE_30 | SE_30_4 | TTAGCCTCGGGTCACAGCCA | 643.7634439628496 | 683.584459489585 | 790.0214901076353 | 556.9600335344194 | 821.9607473652902 | 809.1336426170938 | 510.1001304504429 | 546.4143175414749 | 841.810043378261 | 436.67515995130873 | 536.4197288669536 | 968.0079646151098 |
| SE_300_1_1 | SE_300 | SE_300_1 | GGAAGGATGGCTTCTCTTGG | 704.4549470973943 | 929.4350107095235 | 921.3625628380296 | 813.3178059790002 | 760.4462656269847 | 1190.3100909591935 | 663.9910980757875 | 781.807336689965 | 1023.8470253324208 | 872.6213129577906 | 1271.209116767857 | 1456.6546709735765 |
| SE_300_1_13 | SE_300 | SE_300_1 | GAGGCTGCACCTCAAGCGAG | 3944.947703745408 | 4016.558395743629 | 3865.180140351605 | 4056.17856197847 | 3906.169590382405 | 3562.2671717788967 | 4286.347720641591 | 4060.2932419388144 | 4280.089039117318 | 4377.68670368549 | 3838.276834562228 | 4604.421577491776 |
| SE_300_1_15 | SE_300 | SE_300_1 | AAAGTTCTAGAAACTCAACC | 11.921545258571289 | 9.993924846338961 | 17.77548352742179 | 20.820935833062407 | 8.484756101835254 | 18.192512307236584 | 26.904015319116187 | 8.508181415005666 | 31.96747000170611 | 3.6450347241344634 | 34.03976397623994 | 20.89225822910309 |
| SE_300_1_16 | SE_300 | SE_300_1 | CCACCTGCCAAGAGGTTAAG | 108.37768416882989 | 134.91798542557598 | 128.37849214249073 | 81.98243484268323 | 109.24123481112889 | 65.83956834999907 | 80.71204595734856 | 69.95615830115769 | 80.80666028209045 | 34.992333351690846 | 61.03681816429231 | 55.71268861094157 |
| SE_300_1_25 | SE_300 | SE_300_1 | ATAAAATCTTGGGTAAGCCT | 3.2513305250648967 | 15.990279754142339 | 18.763010390056337 | 15.615701874796805 | 30.757240869152795 | 5.197860659210452 | 6.456963676587884 | 17.01636283001133 | 17.759705556503395 | 19.683187510326103 | 32.86597901154201 | 5.803405063639747 |
| SE_300_2_0 | SE_300 | SE_300_2 | GTAAATGGCATGTAAATCAT | 899.534778601288 | 902.4514136244082 | 860.1358973546878 | 1116.5226840479716 | 823.0213418780196 | 1060.3635744789324 | 1049.2565974455313 | 1037.052779140135 | 851.5778814343378 | 1057.0600699989943 | 942.549326652437 | 978.4540937296614 |
| SE_300_2_14 | SE_300 | SE_300_2 | TGAACCACCACACCTGACCA | 2090.6055276167285 | 1795.9082948871114 | 1882.2262001814408 | 2037.8490946609832 | 1790.2835374872386 | 2150.1816926933907 | 2074.8376614102403 | 1827.3682972451056 | 1830.137657597675 | 1649.7427161432581 | 1797.0647809525294 | 2859.9180153616676 |
| SE_300_2_20 | SE_300 | SE_300_2 | CTGTGGCTCGTCCTGCTACA | 1355.8048289520618 | 1259.2345306387092 | 1234.40857829318 | 1314.3215744620645 | 1285.4405494280409 | 1468.3956362269528 | 1348.4292477941033 | 998.2932860273314 | 1418.112488686796 | 1140.895868654087 | 806.3902707474773 | 1275.5884329880164 |
| SE_300_2_7 | SE_300 | SE_300_2 | GCATAAGATAATATAAGGGG | 144.14231994454374 | 252.84629861237573 | 281.445155850845 | 186.08711400799527 | 233.33079280046948 | 121.28341538157723 | 99.00677637434757 | 211.75918188458544 | 196.24474639936253 | 220.8891042825485 | 105.64064682281361 | 152.0492126673614 |
| SE_300_2_9 | SE_300 | SE_300_2 | TGAAATATCCAACAATAAGT | 82.36703996831072 | 81.9501837399795 | 95.79010567555078 | 83.28374333224963 | 67.87804881468203 | 41.582885273683615 | 71.02660044246673 | 89.80858160283758 | 40.84732277995781 | 139.96933340676338 | 92.72901221113639 | 23.21362025455899 |
| SE_301_1_13 | SE_301 | SE_301_1 | CTAATTTATCTTCTGACCCG | 20.59175999207768 | 73.95504386290831 | 59.25161175807264 | 23.42355281219520 8 | 78.4839939419761 | 55.44384703157816 | 24.75169409358689 | 24.57919075446081 | 129.64585056247478 | 24.78623612411435 | 55.167893340802664 | 23.21362025455899 |
| SE_301_1_14 | SE_301 | SE_301_1 | ATAAATAGCCTTCCCAGGAA | 14.089098941947885 | 43.97326932389143 | 47.40128940645811 | 19.519627343496005 | 39.24199697098805 | 93.56149186578814 | 6.456963676587884 | 7.56282792444948 | 37.29538166865713 | 19.683187510326103 | 19.954344399864794 | 95.17584304369186 |
| SE_301_1_15 | SE_301 | SE_301_1 | AATTTATCTTCTGACCCGAG | 6.5026610501297935 | 0.9993924846338962 | 3.950107450538176 | 7.807850937398403 | 0.0 | 0.0 | 5.380803063823238 | 7.56282792444948 | 0.8879852778251698 | 0.0 | 1.173784964697929 | 1.1606810127279494 |
| SE_301_1_7 | SE_301 | SE_301_1 | TATGGCACTCTAGCTCTCAA | 2099.275742350235 | 1939.8208126743925 | 1989.8666282086062 | 2072.9844238792757 | 2199.6730194007896 | 2009.8394548947083 | 2248.0995200653488 | 2162.023432901995 | 2042.3661389978904 | 1898.3340843292285 | 2462.600855936255 | 1792.091483651954 |
| SE_301_1_8 | SE_301 | SE_301_1 | TTTGGTTGCTTGTAACACAG | 178.8231788785693 | 185.8870021419047 | 201.45547997744697 | 166.56748666449926 | 145.30144824392872 | 63.24063802039384 | 168.95721620404964 | 135.18554914953447 | 143.85361500767752 | 158.19450702743572 | 157.2871852695225 | 156.69193671827318 |
| SE_302_1_0 | SE_302 | SE_302_1 | ATAAAAAATAAGTTTCCTTG | 0.0 | 0.0 | 1.975053725269088 | 2.602616979132801 | 3.1817835381882205 | 0.0 | 4.30464245105859 | 0.0 | 0.0 | 4.374041668961356 | 0.0 | 0.0 |
| SE_302_1_1 | SE_302 | SE_302_1 | AGCACACTGAATTCTTCCGG | 361.98146512389184 | 565.6561463027853 | 494.75095817990655 | 411.2134827029825 6 | 599.2358996921148 | 516.3208254815715 | 332.53362934427605 | 532.2340151831322 | 465.304285580389 | 659.7512850683379 | 635.0176659015796 | 539.7166709184964 |
| SE_302_1_12 | SE_302 | SE_302_1 | TGTTATTTATCCCTCCCCCC | 404.24876194973547 | 551.6646515179107 | 489.81332386673387 | 409.91217421341617 | 466.66158560093896 | 593.42242525986 | 362.6661265016862 | 448.0975545236317 | 637.5734294784719 | 847.1060698888492 | 367.39469395045177 | 442.21946584934875 |
| SE_302_1_24 | SE_302 | SE_302_1 | AAACTCTGTCTCAAAAGGAG | 27.094421042207472 | 24.984812115847404 | 18.763010390056337 | 18.218318853929606 | 9.545350614564661 | 32.05347406513112 | 18.294730416999005 | 0.945353490556185 | 26.639558334755094 | 27.702263903421922 | 2.347569929395858 | 30.177706330926686 |
| SE_302_1_28 | SE_302 | SE_302_1 | GGACTGAAAATAAATCCATG | 228.67691359623106 | 348.78797713722975 | 366.37246603741585 | 287.5891761941745 | 374.38986299348056 | 195.78608483026036 | 253.9739046124568 | 296.8409960346421 | 420.0170364113053 | 317.8470279445252 | 345.09277962119114 | 507.2176025621139 |
| SE_302_2_18 | SE_302 | SE_302_2 | GCAGCCAGAATTGAGTCTGT | 4711.177930819035 | 4442.299594197669 | 4429.05797891593 | 4758.885146344326 | 4601.919590732896 | 4498.748400546647 | 4612.424386309279 | 4471.522010330755 | 4509.189240796212 | 3683.6720922102886 | 4915.8114321549265 | 3533.113002743878 |
| SE_302_2_20 | SE_302 | SE_302_2 | GCCTCTGTGCCCCAGAGTGG | 21.675536833765978 | 42.973876839257535 | 39.50107450538176 | 27.32747828089441 | 18.030106716399914 | 9.529411208552496 | 33.36097899570407 | 19.852423301679885 | 67.4868811147129 | 266.0875348618158 | 35.21354894093787 | 1.1606810127279494 |
| SE_302_2_22 | SE_302 | SE_302_2 | TCTGGCTGCCAGGCCATCAA | 722.8791534060954 | 763.5358582602967 | 712.9943948221407 | 687.0908824910595 | 715.9012960923495 | 647.1336520717014 | 662.9149374630229 | 541.687550088694 | 445.76860946823524 | 586.1215836408217 | 748.8748074772788 | 717.3008658658728 |
| SE_302_2_27 | SE_302 | SE_302_2 | GTGTGTGGCAGGGTGACAAG | 1919.3687866299774 | 1626.011572499349 | 1724.221902159914 | 1868.678991017351 | 1658.7698179087922 | 1483.9892182045842 | 1944.622227265718 | 1805.6251669623134 | 1657.868513699592 | 1353.0368895987128 | 1723.1163281765598 | 1428.7983266681058 |
| SE_302_2_4 | SE_302 | SE_302_2 | TGTCTTTAGGAGTTTAATGA | 18.42420630870108 | 31.980559508284678 | 33.5759133295745 | 13.013084895664004 | 38.181402458258646 | 1.7326202197368175 | 9.685445514881827 | 6.617474433893295 | 47.063219724734 | 18.954180565499207 | 10.564064682281362 | 1.1606810127279494 |
| SE_302_3_0 | SE_302 | SE_302_3 | TCTTACATCCTTGACCTAGT | 800.9110860076529 | 551.6646515179107 | 563.8778385643246 | 756.0602324380786 | 553.6303356447503 | 417.561472956573 | 735.0176985182542 | 670.2556248043352 | 592.2861803093882 | 788.7855143026978 | 555.2002883021204 | 506.05692154938595 |
| SE_302_3_1 | SE_302 | SE_302_3 | TCAAGCACGAAAAAACATAT | 2285.6853591206223 | 2408.5358879676896 | 2370.064470322906 | 2264.2767718455366 | 2302.550687135542 | 2799.914275094697 | 2186.7583651377636 | 2409.7060474277155 | 2881.5122265426758 | 2058.715612191145 | 2396.868897913171 | 2284.2202330486043 |
| SE_302_3_2 | SE_302 | SE_302_3 | GGAGAACACACATTTCCTTA | 1790.3993424690698 | 1889.8511884426978 | 1910.8644791978427 | 1789.2991731538007 | 1790.2835374872386 | 2734.9410168545664 | 1858.5293782445463 | 1624.1172967755258 | 2375.3606181823293 | 1745.971632860408 | 1844.0161795404465 | 2105.4753570885005 |
| SE_302_3_3 | SE_302 | SE_302_3 | GTTGGCTGTGGTGTGCCAGG | 956.9749512107679 | 913.4447309553811 | 1000.3647118487931 | 965.5708992582692 | 823.0213418780196 | 627.2085195447279 | 1182.7005134283477 | 661.7474433893295 | 1003.4233639424418 | 870.4342921233098 | 528.2032341140681 | 1012.1138430987719 |
| SE_302_3_6 | SE_302 | SE_302_3 | GTTAAAGTACTAATTTAAGT | 33.59708209233727 | 46.97144677779312 | 38.51354764274722 | 36.43663770785921 | 36.06021343279983 | 24.256683076315444 | 12.913927353175769 | 29.305958207241734 | 67.4868811147129 | 32.076305572383276 | 38.73490383503166 | 19.73157721637514 |
| SE_302_4_1 | SE_302 | SE_302_4 | GCTCTCATGCAACAAAGCGA | 303.4575156727237 | 281.8286806667587 | 356.4971974110704 | 255.0564639550145 | 328.7842989461161 | 388.1069292210471 | 194.7850709104012 | 234.44766565793387 | 483.06399113689235 | 279.93866681352677 | 275.8394667040133 | 449.1835519257164 |
| SE_302_4_10 | SE_302 | SE_302_4 | TTTCCCTGCTTCCCACAGAA | 17.340429467012783 | 21.986634661945715 | 30.613332741670867 | 18.218318853929606 | 14.848323178211695 | 38.98395494407839 | 11.837766740411123 | 7.56282792444948 | 13.319779167377547 | 4.374041668961356 | 29.344624117448227 | 218.20803039285448 |
| SE_302_4_13 | SE_302 | SE_302_4 | GCTGCTCCTTCAGACAACCA | 953.723620685703 | 1054.3590712887606 | 999.3771849861586 | 978.5839841539331 | 1022.4131102711481 | 1465.7967058973475 | 997.6008880328282 | 1064.4680303662644 | 959.9120853290085 | 1560.8038688743773 | 1338.1148597556391 | 1901.1954988483812 |
| SE_302_4_26 | SE_302 | SE_302_4 | GTTAACCTGAAAGCACAAGG | 138.72343573610226 | 161.90158251069118 | 150.1040831204507 | 178.27926307059687 | 142.1196647057405 | 79.7005301078936 | 139.90087965940418 | 140.85767009287156 | 205.12459917761421 | 253.69441679975864 | 126.76877618737633 | 67.31949873822107 |
| SE_302_4_9 | SE_302 | SE_302_4 | CCTTGTTTACCGATGAGACC | 1081.6092880049223 | 1026.3760817190114 | 1107.0176130133239 | 1182.889417015858 | 1136.957317645924 | 1303.796715351955 | 1224.6707773261687 | 907.5393509339376 | 943.9283503281555 | 1199.9454311850652 | 1159.699545121554 | 1332.461802611686 |
| SE_303_1_19 | SE_303 | SE_303_1 | TCTCCCAGTCATGGCAACCG | 378.2381177492163 | 433.73633833111097 | 437.474400147103 | 433.33572702561133 | 414.69245447719805 | 338.7272529585478 | 386.3416599825084 | 535.0700756548007 | 338.3223908513897 | 316.38901405487144 | 498.85860999661986 | 456.14763800208414 |
| SE_303_1_23 | SE_303 | SE_303_1 | TGACTGGGAGAAGAGAGGAT | 1468.517620487645 | 1686.9745140620169 | 1600.781044330596 | 1395.0027008151812 | 1386.1970281373347 | 1126.2031428289313 | 1177.3197103645243 | 967.0966208389773 | 1735.1232328703818 | 968.8502296749404 | 583.3711274548707 | 1064.3444886715297 |
| SE_303_1_36 | SE_303 | SE_303_1 | GCCAGTGGCCATGAGAACAA | 163.65030309493312 | 157.9040125721556 | 147.14150253254707 | 161.36225270623365 | 167.57393301124625 | 101.35828285460381 | 165.7287343657557 | 84.13646065950047 | 189.14086417676117 | 88.9388472688809 | 45.77761362321923 | 169.4594278582806 |
| SE_303_1_5 | SE_303 | SE_303_1 | TTGAGACCATCGCAAACAAT | 913.623877543236 | 802.5121651610186 | 823.5974034372098 | 942.1473464460739 | 756.203887576067 | 861.1122492091982 | 794.2065322203098 | 934.0092486695108 | 998.0954522754909 | 1236.3957784264098 | 970.7201658051873 | 393.4708633147749 |
| SE_303_1_9 | SE_303 | SE_303_1 | TGTGGTGAAGATCAGAGGAA | 1251.7622521499852 | 1343.1834993479565 | 1431.9139508200888 | 1298.7058725872675 | 1334.2278970135937 | 1591.411671828267 | 1269.869523062284 | 1634.5161851716439 | 1468.727649522831 | 1558.6168480398965 | 1508.3136796368387 | 1492.635782368143 |
| SE_304_1_20 | SE_304 | SE_304_1 | CCTCGTCTGTGACATGAGGG | 800.9110860076529 | 791.5188478300457 | 828.5350377503825 | 865.3701455616563 | 721.2042686559965 | 764.9518270138049 | 863.0808114372472 | 540.7421965981379 | 900.4170717147222 | 809.9267157026777 | 681.9690644894968 | 727.7469949804243 |

| | | | | | | | | | | | | | | | |
|---|---|---|---|---|---|---|---|---|---|---|---|---|---|---|---|
| SE_304_1_3 | SE_304 | SE_304_1 | GCACACATCCTAAGTGCCCA | 201.5824925540236 | 226.86209401189444 | 231.0812858564833 | 217.31851775758886 | 217.42187510952837 | 159.4010602157872 | 148.51016456152135 | 180.56251669623134 | 235.31609862367 | 185.89677093085763 | 139.68041079905356 | 105.6219721582434 |
| SE_304_1_31 | SE_304 | SE_304_1 | GGCTGCTTTTGCCATCTGAG | 137.63965889441397 | 182.888824688003 | 160.96687860943067 | 130.13084895664005 | 184.54344521491677 | 239.1015903236808 | 102.23525821264151 | 236.33837263904624 | 118.10204195074758 | 136.32429868262892 | 237.10456286898167 | 143.92444557826573 |
| SE_304_1_33 | SE_304 | SE_304_1 | TGGCTTCCTGTGTACCCCAT | 22.759313675454276 | 5.9963549078033775 | 10.862795488979984 | 28.62878677046081 | 12.727134152752882 | 17.326202197368175 | 8.60928490211718 | 0.945353490556185 | 1.7759705556503396 | 0.0 | 0.0 | 22.05293924183104 |
| SE_304_1_7 | SE_304 | SE_304_1 | TACCACATCAAAGTCCCCCA | 2163.2185760098446 | 2739.3348003815095 | 2562.632208536642 | 2308.5212604907942 | 2563.456937266976 | 3034.684314869036 | 1912.3374088827786 | 1854.783548471235 | 2624.884481251202 | 2380.2076748598047 | 1710.2046935648825 | 2280.7381900104206 |
| SE_304_2_23 | SE_304 | SE_304_2 | GTGGCTGGAGGGGAGAGTAA | 105.126353643765 | 103.93681840192521 | 116.52816979087619 | 117.11776406097604 | 113.48361286204653 | 178.4598826328922 | 110.8445431147587 | 69.0108048106015 | 103.0062922277197 | 160.3815278619164 | 59.86303319959438 | 190.3516860873837 |
| SE_304_2_24 | SE_304 | SE_304_2 | TTTCAAACTGTACGATTCGG | 68.27794102636283 | 61.96233404730156 | 76.03956842285989 | 42.943180155691216 | 65.75685978922321 | 63.24063802039384 | 76.40740350628997 | 63.33868386726439 | 16.871720278678225 | 17.496166675845423 | 59.86303319959438 | 20.89225822910309 |
| SE_304_2_25 | SE_304 | SE_304_2 | TTGGGTTTCAGCTTTCCTGT | 44.434850509220254 | 88.94593113241676 | 60.23913862070719 | 66.36673296788642 | 50.90853661101153 | 91.82887164605133 | 35.513300221233365 | 17.961716320567515 | 39.07135222430747 | 40.82438891030599 | 37.56111887033373 | 49.909283547301825 |
| SE_304_2_28 | SE_304 | SE_304_2 | GAGAGGACAGAACCAGCCGT | 7977.681331667568 | 7772.275352997811 | 7721.4725389395 | 8012.156370260327 | 7457.040019000458 | 8066.213432984754 | 8278.903593998433 | 7747.171855107936 | 7905.732928477487 | 6530.444211759304 | 7894.877672558271 | 8730.642577739636 |
| SE_304_2_30 | SE_304 | SE_304_2 | CCAGGCTGCTCTGTGACCTA | 3105.0206514369766 | 2806.2940968519806 | 2714.7113453823617 | 3129.646917407193 | 2664.21341597627 | 2931.593411794695 | 3367.306557340582 | 2714.1098713868073 | 2838.0009479292426 | 4554.835391278425 | 2288.8806811609616 | 2969.0220305580947 |
| SE_305_1_12 | SE_305 | SE_305_1 | TCGTGCCTCCCACATCTCTA | 198.3311620289587 | 181.8894322033691 | 183.6799964500252 | 179.58057156016326 | 230.14900926228125 | 213.97859713749696 | 236.75533480822244 | 231.61160518626534 | 274.38745084797745 | 122.47316673091797 | 158.4609702342204 | 311.06251141109044 |
| SE_305_1_14 | SE_305 | SE_305_1 | GGGAGGCACGATGAGTCCAT | 9365.99946587028 | 7744.292363428061 | 7551.617918566359 | 8353.099194526725 | 7206.739713996319 | 7683.304364422917 | 9590.743380958538 | 6996.561183606325 | 7492.819774288782 | 7804.019344371886 | 7065.011702516835 | 7372.645792847935 |
| SE_305_1_20 | SE_305 | SE_305_1 | CTTGCTCTCCAGGAAGCAGT | 535.3857597940197 | 607.6306306574089 | 553.0150430753447 | 566.0691929613841 | 524.9942838010563 | 622.0106588855175 | 595.11681885885 | 547.3596710320311 | 457.3124180799624 | 569.3544239098031 | 510.59645964359913 | 882.1175696732415 |
| SE_305_1_30 | SE_305 | SE_305_1 | GCTCCCCTAAAAGAAAAAAG | 5.4188842084414945 | 13.991494784874547 | 18.763010390056337 | 7.807850937398403 | 8.484756101835254 | 44.18181560328885 | 5.380803063823238 | 59.557269905039654 | 9.767838056076867 | 0.7290069448268927 | 80.99116256415711 | 18.57089620364719 |
| SE_305_1_8 | SE_305 | SE_305_1 | ACAGAAAAGGAGAAGACATA | 1158.5574437647915 | 1322.1962571706447 | 1347.9741674961526 | 1378.085690450818 | 1279.0769823516646 | 1299.4651648026131 | 1369.9524600493962 | 1366.0357938536874 | 1288.4666381243214 | 1124.8577158678954 | 1488.359335236974 | 1052.7376785442502 |
| SE_305_2_0 | SE_305 | SE_305_2 | GGCCCTGATGCACAAAGTTC | 177.73940203688102 | 196.88031947287755 | 227.13117840594512 | 197.79889041409285 | 127.2713415275288 | 168.93047142433971 | 188.3281072338133 | 201.36029348846742 | 198.90870223283804 | 494.2667085926332 | 238.2783478336796 | 129.99627342553035 |
| SE_305_2_1 | SE_305 | SE_305_2 | TACCGGAACTTTGTGCATCA | 1856.509729812056 | 1774.9210527097996 | 1742.9849125499702 | 1945.4561919017688 | 1798.768293589074 | 2204.7592296151 | 1857.4532176317816 | 1871.7999113012463 | 1828.3616870420246 | 2113.391133053162 | 1727.8114680353515 | 2584.8366153451434 |
| SE_305_2_2 | SE_305 | SE_305_2 | TTACCGGAACTTTGTGCATC | 496.3697934932409 | 404.75395627672793 | 414.7612823065085 | 372.17422801599054 | 398.78353678625695 | 500.72724350394026 | 440.14969062074084 | 412.17412188249665 | 394.26546335437536 | 453.44231968232725 | 184.28423945757487 | 445.70150888753255 |
| SE_305_2_27 | SE_305 | SE_305_2 | TTAGCATCCGTTCAGGGCGT | 1708.032302500759 | 1608.022507775939 | 1495.1156700286997 | 1703.4128128424181 | 1388.3182171627934 | 1444.1389531506375 | 1598.0985099555014 | 1307.4238774392038 | 1527.334677859292 | 828.8808962681769 | 1110.4005766042408 | 1211.7509772879791 |
| SE_305_2_28 | SE_305 | SE_305_2 | CATGGTGCCGACGCCCTGAA | 5038.478537008901 | 4811.075421027576 | 4958.372377288046 | 4957.985345247986 | 5098.277822690258 | 5179.6681469032155 | 4938.501051976968 | 5623.907915318745 | 5136.994832218607 | 4984.220481781465 | 5002.671519542573 | 4517.370501537179 |
| SE_305_3_13 | SE_305 | SE_305_3 | CTCGTACTTTCCGGCTGGTG | 732.6331449812901 | 973.4082800334148 | 951.9758955797005 | 752.1563069693794 | 1055.2915401657597 | 860.2459390993299 | 620.9446735652016 | 927.3917742356175 | 876.4414692134426 | 460.7323891305962 | 931.9852619701556 | 915.7773190423521 |
| SE_305_3_16 | SE_305 | SE_305_3 | CTGAGCCGACTCGTACTTTC | 1126.0441385141426 | 1193.274626652872 | 1078.379333996922 | 1011.1166963930931 | 1076.5034304203477 | 1063.828814918406 | 1136.4256070794677 | 1313.095998382541 | 1010.5272461650433 | 1029.3578060955724 | 934.3328318995515 | 1037.6488253787868 |
| SE_305_3_17 | SE_305 | SE_305_3 | AAGTACGAGTCGGCTCAGCC | 1587.733073073358 | 1473.104522350363 | 1356.8619092598635 | 1544.6531771153172 | 1546.3467995594751 | 931.2833681085394 | 1472.1877182620378 | 1586.3031571532783 | 1412.784577019845 | 1485.7161535572072 | 1448.4506464372444 | 1458.9760329990324 |
| SE_305_3_5 | SE_305 | SE_305_3 | ACTGCCACGCCCCCGCATGC | 9114.563238598594 | 7263.584578319157 | 7109.2058841060825 | 8409.05545957808 | 7579.008387964341 | 8022.0316173814645 | 9412.100719239606 | 7630.893375769525 | 7014.195709541016 | 7650.927885958238 | 8288.095635732077 | 9233.217456250837 |
| SE_305_3_7 | SE_305 | SE_305_3 | ATGGGAGTTTCTGGCCCTGC | 2550.1269084925675 | 2576.4338253861843 | 2720.636506558169 | 2639.05361684066 | 2855.120428267563 | 3106.5880539881136 | 2463.3316426182782 | 3122.502579307079 | 3035.13367960643 | 3012.2566960247204 | 4048.384343243157 | 3126.8746482890956 |
| SE_306_1_1 | SE_306 | SE_306_1 | TCCTGATAGCTGACACGTGG | 1217.0813932159597 | 1182.2813093218992 | 1180.0946008482802 | 1315.6228829516308 | 1187.8658542569356 | 898.3635839335399 | 1039.5711519306494 | 1190.200044610237 | 921.7287183825263 | 1034.4608547093608 | 903.8144228174053 | 704.5333747258653 |
| SE_306_1_18 | SE_306 | SE_306_1 | AATACAGATGAGGAGATCAT | 23.843090517142578 | 48.970231747060915 | 40.488601368016305 | 55.95626505135522 | 62.575076251035 | 6.93048087894727 | 19.370891029763655 | 34.03272566002266 | 71.03882222601358 | 94.04189588266915 | 23.47569929395858 | 24.37430126728694 |
| SE_306_1_19 | SE_306 | SE_306_1 | TGTTCCTCAATATCCAGGGA | 106.21013048545329 | 103.93681840192521 | 79.00214901076352 | 140.54131687317124 | 102.87766773475245 | 107.42245362368269 | 153.8909676253446 | 90.75393509339376 | 206.90056973326455 | 368.1485071375808 | 76.29602270536539 | 414.3631215438779 |
| SE_306_1_2 | SE_306 | SE_306_1 | TGAGACGTTCCCAGAACTGA | 1311.3699784428418 | 1404.146440910624 | 1512.8911535561215 | 1346.8542867012245 | 1468.9234001302284 | 1130.5346933782735 | 1453.8929878450388 | 1469.0793243243115 | 1562.8540889722988 | 1138.7088478196063 | 1763.0250169762894 | 1530.9382557881654 |
| SE_306_1_20 | SE_306 | SE_306_1 | TTATATAGTCTAAAAACGGG | 1.083776841688299 | 3.9975699385355847 | 6.912688038441808 | 0.0 | 16.969512203670508 | 0.0 | 1.0761606127646475 | 21.743130282792254 | 0.0 | 2.187020834480678 | 14.085419576375148 | 0.0 |
| SE_306_2_2 | SE_306 | SE_306_2 | TCACACACATACAAAAAAAG | 967.812719627651 | 1305.2065849318685 | 1311.4356735786744 | 1076.1821208714132 | 1396.8029732646287 | 1303.796715351955 | 942.7166967818312 | 1207.2164074402483 | 1624.1250731422356 | 1229.834715922968 | 1191.3917391683979 | 1487.993058317231 |
| SE_306_2_29 | SE_306 | SE_306_2 | GTACCTCTCGATGGGAAGAG | 150.64498099467355 | 107.93438834046079 | 119.49075037877982 | 122.32299801924164 | 90.15053358199957 | 79.7005301078936 | 191.55658907210724 | 111.55171188562983 | 23.975602501279585 | 78.73275004130441 | 42.25625872912545 | 99.81856709460365 |
| SE_306_2_5 | SE_306 | SE_306_2 | AGAACCCTCACGAAGTTCCA | 84.53459365168732 | 137.91616287947767 | 142.20386821937433 | 115.81645557140963 | 169.69512203670507 | 155.06950966644516 | 67.79811860417279 | 142.74837707398393 | 189.14086417676117 | 80.1907639309582 | 136.15905590495976 | 164.8167038073688 |
| SE_306_2_6 | SE_306 | SE_306_2 | TTAGGATGCACTGAGCTGGA | 1080.5255111632341 | 1073.3475284968044 | 1100.1049249748821 | 1156.86324722453 | 922.7172260745839 | 954.6737410749864 | 1118.1308766624688 | 1064.4680303662644 | 1112.6455531149377 | 1292.5293131780807 | 943.7231116171349 | 911.1345949914403 |
| SE_306_2_8 | SE_306 | SE_306_2 | AGAAGCTGTAGGCTTCTTAA | 3.2513305250648967 | 3.9975699385355847 | 8.887741763710896 | 1.3013084895664004 | 14.848323178211695 | 1.7326202197368175 | 2.152321225529295 | 16.071009339455145 | 0.8879852778251698 | 0.0 | 19.954344399864794 | 4.642724050911798 |
| SE_307_1_1 | SE_307 | SE_307_1 | GATTCAAAGCCTGAGAAGAA | 100.7912462770118 | 93.94289355558624 | 96.77763253818532 | 88.48897729051524 | 74.24161589105847 | 97.89304241513018 | 69.95043982970209 | 47.26767452780925 | 103.0062922277197 | 134.13727784814824 | 92.72901221113639 | 55.71268861094157 |
| SE_307_1_11 | SE_307 | SE_307_1 | GACGAGCCATTTCATCAGAG | 8609.523230371846 | 10056.886572870897 | 9654.062609115303 | 9469.621878574697 | 9791.408541517883 | 9208.876467901186 | 9280.809124482319 | 9350.491375091226 | 10689.566774459394 | 10618.71515834852 | 10351.609603671037 | 9634.813086654709 |
| SE_307_1_27 | SE_307 | SE_307_1 | TGGATGATAAGCCTTGAGGA | 47.686181034285156 | 60.962941562667666 | 72.08946097232172 | 46.847105624390416 | 74.24161589105847 | 127.34758615065608 | 19.370891029763655 | 2.836060471668555 | 34.63142583518162 | 17.496166675845423 | 38.73490383503166 | 51.069964560029774 |
| SE_307_1_3 | SE_307 | SE_307_1 | GTGAGCTCATACGGATCAGG | 1592.068180440111 | 1583.0376956600915 | 1659.045129226034 | 1657.8670157075942 | 1612.1036593486983 | 1425.9464408434008 | 1423.7604906876286 | 1476.642152248761 | 1514.0148986919144 | 1184.6362853437006 | 1088.0986622749801 | 1441.5658178081133 |
| SE_307_1_32 | SE_307 | SE_307_1 | GCACAGAGCGAGCCTGTGGA | 2730.033864212825 | 2698.35970851152 | 2694.9608081296706 | 2715.830817725078 | 2909.2107484167627 | 2764.3955605900924 | 2685.0207288477955 | 2706.5470434623576 | 2859.312594597047 | 2389.684765142554 | 3089.402027084949 | 2949.2904533417195 |
| SE_307_2_2 | SE_307 | SE_307_2 | TATGTAGAACACACAACACA | 2248.83694650322 | 2369.559581066968 | 2344.3887718944075 | 2420.433790593505 | 2273.914635291848 | 1973.454430280235 | 2431.0468242353386 | 1935.1385951685106 | 2502.3425129113284 | 2702.428744473291 | 2083.468312338824 | 2059.048116579382 |
| SE_307_2_3 | SE_307 | SE_307_2 | GCCTGCCTAGATGCAGACAC | 3179.801253513469 | 2481.491539345964 | 2547.8193055971237 | 2705.4203498085467 | 2566.638720805164 | 2356.3634988420717 | 2872.272675468844 | 2657.388661953436 | 2387.7924120718817 | 2036.1163969015113 | 3030.712778850053 | 2897.059807768962 |
| SE_307_2_4 | SE_307 | SE_307_2 | ACCTGTGTCTGCATCTAGGC | 732.6331449812901 | 628.6178728347207 | 633.0047189487427 | 649.3529362936338 | 721.2042686559965 | 641.9357914124909 | 521.9378971908541 | 523.7258337681264 | 615.3737975328427 | 303.9958959928142 | 507.07510474950533 | 392.3101823020469 |
| SE_307_2_5 | SE_307 | SE_307_2 | AGAAACCTGTGTCTGCATCT | 240.59845885480235 | 235.8566263735995 | 261.6946185981542 | 219.92113473672168 | 303.33003064061035 | 381.17644834209983 | 295.9441685102781 | 260.91756339350707 | 177.59705556503397 | 269.0035626411234 | 232.40942301018995 | 371.41792407294383 |
| SE_307_2_7 | SE_307 | SE_307_2 | ACATATTTAATGCTGCTACT | 320.7979451397365 | 404.75395627672793 | 367.3599929000504 | 370.87291952642414 | 381.81402458258646 | 473.00531998815114 | 349.75219914851044 | 476.45815924031723 | 371.17784613092095 | 112.26706950334147 | 494.16347013782814 | 461.95104306572387 |
| SE_308_1_1 | SE_308 | SE_308_1 | TTAGCAGGACAGTTGGAGAG | 1140.1332374560905 | 1115.322012851428 | 1127.7556771286493 | 990.2957605600308 | 1052.1097566275714 | 1343.646980405902 | 1089.0745401178233 | 1159.948732912439 | 967.9039528294351 | 1160.579056164413 | 1284.1207513795343 | 1266.302984886193 |
| SE_308_1_22 | SE_308 | SE_308_1 | CTGGTGTGTGGGATGCACAA | 671.9416418467454 | 551.6646515179107 | 531.2894520973847 | 629.8333089501378 | 694.6894058377615 | 589.9571848203864 | 648.9248494970824 | 763.8456203693975 | 487.5039175260182 | 317.11802099969833 | 1041.147263687063 | 784.6203646040938 |
| SE_308_1_28 | SE_308 | SE_308_1 | ACAGTACTGGGAAAGCAGGT | 1871.6826055956922 | 1978.7971195751145 | 1935.5526507637062 | 2178.3904115341543 | 1893.161205221991 | 2106.86618719997 | 2144.7881012399425 | 2102.4661629969555 | 1817.7058637081225 | 2639.7341472181784 | 2596.412341911819 | 2510.5530305305547 |
| SE_308_1_3 | SE_308 | SE_308_1 | GATTCTGTCCTCTTGGCCAG | 46.60240419259685 | 47.97083926242702 | 66.16429979651446 | 33.834020728726415 | 106.05945127294068 | 57.176467251314975 | 75.33124289352533 | 57.66656292392729 | 27.527543612580263 | 42.282402799959776 | 123.24742129328254 | 10.446129114551544 |
| SE_308_1_7 | SE_308 | SE_308_1 | GCTATCTTATACGTATGTTG | 123.55055995246607 | 240.85358879676897 | 208.36816801588878 | 162.66356119580004 | 159.089176909411 | 258.1604127407858 | 217.3844437784588 | 213.64988886569782 | 198.02071695501286 | 227.45016678599052 | 151.41826044603283 | 280.88480508016374 |
| SE_309_1_17 | SE_309 | SE_309_1 | CTTCATTATAAACTGAGCGA | 20.59175999207768 | 30.98116702365078 | 42.463655093285396 | 16.917010364363207 | 60.45388722557618 | 32.919784174999535 | 3.228481838293942 | 18.9070698111237 | 23.087617223454416 | 16.76715973101853 | 10.564064682281362 | 27.856344305470785 |
| SE_309_1_21 | SE_309 | SE_309_1 | GTGCATGCTCTGTTTCCTGG | 1287.5268879256992 | 1346.1816768018582 | 1423.0262090563779 | 1341.649052742959 | 1305.5918451698997 | 1166.9197179927467 | 1228.9754197772274 | 1332.0030681936646 | 1509.5749723027886 | 1223.2736534195258 | 1704.335768741393 | 865.8680354950503 |
| SE_309_1_31 | SE_309 | SE_309_1 | TTAAAATTCTGAGAGAGGGA | 0.0 | 2.9981774539016888 | 1.975053725269088 | 1.3013084895664004 | 7.4241615891058474 | 0.8663101098684087 | 0.0 | 0.0 | 3.551941111300679 | 0.0 | 0.0 | 0.0 |
| SE_309_1_6 | SE_309 | SE_309_1 | CCTGGACGCGCTTCTCTCCA | 4700.340162402153 | 4234.425957393818 | 4455.721204207062 | 4619.6451379607215 | 4685.706557238519 | 4998.609333940718 | 4487.58975522858 | 3910.9273904309375 | 4426.606609958471 | 5194.17448189161 | 4536.678888557496 | 5301.990866141273 |
| SE_309_1_9 | SE_309 | SE_309_1 | ACACTAGGCATGACAGCAAG | 4391.463762520987 | 3865.6501305639104 | 3896.780999955911 | 3837.5587357313148 | 3652.687501840077 | 3473.903540572319 | 4166.893892624715 | 3249.1799470416076 | 3698.458682141832 | 3816.3513561687832 | 3531.9189587760684 | 3109.4644330981764 |
| SE_309_2_12 | SE_309 | SE_309_2 | GCAGGAGAACAGGGTCTGGA | 11002.50249681961 | 11647.919408408061 | 11589.615259879009 | 10835.995792619417 | 11292.149777029994 | 11262.897738399182 | 10671.208636174244 | 11699.694799123346 | 11095.376046425496 | 10515.925179127926 | 11899.831972107604 | 11363.067114606625 |
| SE_309_2_13 | SE_309 | SE_309_2 | GTGGCGTGCATCCTACCTAA | 771.6491112820688 | 633.6148352578901 | 661.6429979651446 | 644.1477023353682 | 622.5689789721617 | 791.8074404197256 | 639.2394039822005 | 725.086127256594 | 630.4695472558706 | 497.1827363719408 | 689.0117742776844 | 844.9757772659472 |
| SE_309_2_19 | SE_309 | SE_309_2 | TATCAGACAAGGAAACAGAG | 371.7354566990865 | 436.7345157850126 | 440.4369807350066 | 351.3532921829281 | 543.0243905174563 | 396.7700303197312 | 399.2555873356842 | 575.7202757487166 | 545.2229605846543 | 394.3927571513489 | 409.65095267957724 | 748.6392532095274 |
| SE_309_2_20 | SE_309 | SE_309_2 | TGGATTTCCTTTGTATTAGG | 1.083776841688299 | 3.9975699385355847 | 3.950107450538176 | 3.9039254686992013 | 5.3029725636470335 | 0.0 | 1.0761606127646475 | 0.0 | 0.0 | 0.0 | 0.0 | 1.1606810127279494 |
| SE_309_2_6 | SE_309 | SE_309_2 | GGGTGCATCGACCTGGCCAA | 3507.1018597033353 | 3081.127030126302 | 3160.085960430541 | 3507.0263793814493 | 3171.177593060926 | 3299.775208488769 | 3607.2903739870985 | 3023.24046279868 | 3218.0586468384154 | 3183.5733280590403 | 3931.0058467733643 | 3154.7309925945665 |
| SE_309_3_13 | SE_309 | SE_309_3 | AAGCTGTGTGTAATTCTGAA | 239.51468201311405 | 407.7521337306296 | 410.8111748559703 | 256.35777244458086 | 392.4199697098805 | 313.60425977236395 | 178.6426617189315 | 162.60080037566382 | 324.11462640618697 | 201.93492371704926 | 224.19292825730446 | 364.4538379965761 |
| SE_309_3_14 | SE_309 | SE_309_3 | GAGGTCACATATGTAGACAG | 1952.9658687223146 | 1812.8979671258876 | 1914.8145866483808 | 1808.8188004972967 | 1798.768293589074 | 1911.0801023697097 | 1994.1256154528917 | 1740.3957761139366 | 1990.8629928840307 | 2068.9217094187215 | 2009.5198595628544 | 1909.3202659374767 |
| SE_309_3_16 | SE_309 | SE_309_3 | GGATGGCATTACAAAACAGA | 2187.061666526987 | 2652.3876542183607 | 2613.983605393638 | 2259.071537887271 | 2512.5484006559645 | 2542.6201724637795 | 2327.7354054099324 | 2785.011383178521 | 2408.2160734618606 | 2863.5392792800344 | 2623.409396099871 | 2378.235395079568 |
| SE_309_3_21 | SE_309 | SE_309_3 | GGATGTTTCCGGAAACTGAG | 92.12103154350541 | 137.91616287947767 | 128.37849214249073 | 102.80337067574564 | 119.84717993842297 | 102.22459296447224 | 154.96712823810924 | 94.5353490556185 | 138.5257033407265 | 185.16776398603074 | 113.85714157569912 | 201.9584962146632 |
| SE_309_3_22 | SE_309 | SE_309_3 | GCTTTGTGAGGAACCTACGA | 366.31657249064506 | 438.73330075428044 | 460.1875179876975 | 342.24413275596334 | 398.78353678625695 | 539.7111984480186 | 335.76211118257004 | 627.7147177293068 | 396.0414339100257 | 272.64859736525784 | 450.73342644400475 | 552.4841620585039 |
| SE_31_1_15 | SE_31 | SE_31_1 | GTAGCCTTACCTGAGGGCGG | 399.9136545829823 | 444.7296556620838 | 499.6885924930793 | 376.0781534846897 | 461.35861303729195 | 321.40105076117965 | 393.87478427186096 | 567.212094333711 | 330.3305233509632 | 427.92707661338596 | 530.5508040434639 | 300.6163822965389 |
| SE_31_1_16 | SE_31 | SE_31_1 | TGGGTAGTTATGCCACCCGA | 153.89631151973845 | 173.89429232629794 | 175.77978154894885 | 158.75963572710086 | 138.9378811675523 | 89.2299413164461 | 227.06988929334062 | 155.98332594177052 | 100.34233639424419 | 169.858618144666 | 177.2415296693873 | 67.31949873822107 |
| SE_31_1_34 | SE_31 | SE_31_1 | GAATCTGTCCCAGCTCTAAG | 4.335107366753196 | 5.9963549078033775 | 11.850322351614528 | 13.013084895664004 | 16.969512203670508 | 0.0 | 15.066248578705064 | 17.961716320567515 | 0.0 | 2.9160277793075706 | 4.695139858791716 | 0.0 |
| SE_31_1_43 | SE_31 | SE_31_1 | ATGAGGCGAGAGGGGCCATG | 6508.079934338235 | 5308.772878375256 | 5317.832155287019 | 6389.424683771026 | 5646.605185771362 | 5883.978266226232 | 7018.719516451031 | 6041.7541581445785 | 5432.693929734389 | 5249.579009698454 | 5924.092716830448 | 6048.308757325344 |
| SE_31_1_44 | SE_31 | SE_31_1 | GTACACAGTGAGGTTGGGCA | 2604.3157505769823 | 2256.6282303033377 | 2275.2618915099893 | 2481.595289603126 | 2478.6093762486234 | 2359.8287392815455 | 2752.8188474519684 | 2583.6510896900536 | 1875.4249067667586 | 2514.344952707953 | 2729.050042922685 | 2227.346863424935 |
| SE_31_10_25 | SE_31 | SE_31_10 | GCCCTCAGAGGAGCTAAGGA | 177.73940203688102 | 286.8256430899282 | 321.93375721886133 | 261.5630064028465 | 264.0880336696223 | 277.2192351578908 | 190.4804284593426 | 215.5405958468102 | 419.1290511334801 | 387.10268770308 | 164.32989505771008 | 269.2779949528843 |
| SE_31_10_38 | SE_31 | SE_31_10 | GTAGGTTGTAAATTCTTGAG | 176.65562519519273 | 262.8402234587147 | 262.6821454607887 | 231.63291114281927 | 197.27057936766965 | 240.83421054341764 | 223.84140745504666 | 259.97220990295085 | 295.69909751578155 | 312.743979330737 | 180.76288456348107 | 301.77706330926685 |
| SE_31_10_4 | SE_31 | SE_31_10 | GGTCCAAAGGTCACACCGAT | 755.3924586567443 | 784.5231004376085 | 756.4455767780607 | 717.0209777510867 | 779.536966856114 | 546.6416793269659 | 526.2425396419126 | 609.7530014087394 | 436.0007714121584 | 788.0565073578709 | 451.90721140870266 | 862.3859924568665 |
| SE_31_10_44 | SE_31 | SE_31_10 | TTATCACCCCTCTCTGTAGG | 59.60772629285644 | 42.973876839257535 | 46.41376254382357 | 58.55888203048802 | 49.84794209828212 | 6.93048087894727 | 35.513300221233365 | 19.852423301679885 | 45.28724916908366 | 60.50757642063209 | 5.868924823489645 | 18.57089620364719 |
| SE_31_10_8 | SE_31 | SE_31_10 | TCCTTTTCTGAGTTTCCTGT | 7.586437891818092 | 25.984204600481302 | 27.650752153767232 | 11.711776406097604 | 13.787728665482287 | 17.326202197368175 | 8.60928490211718 | 12.289595377230405 | 9.767838056076867 | 1.4580138896537853 | 65.73195802308402 | 5.803405063639747 |
| SE_31_11_12 | SE_31 | SE_31_11 | CCTAACATCAAAATGTACAG | 867.0214733506391 | 1068.350566073635 | 1023.0778296893876 | 1029.3350152470227 | 891.959985205431 | 1264.8127604078768 | 881.3755418542463 | 983.1676301784324 | 820.4983967104569 | 1277.9491742815428 | 1156.1781902274602 | 1571.5620912336435 |
| SE_31_11_13 | SE_31 | SE_31_11 | TTTTCTAGTCCACAGCTCTG | 2.167553683376598 | 2.9981774539016888 | 2.962580587903632 | 0.0 | 8.484756101835254 | 0.0 | 0.0 | 21.743130282792254 | 0.0 | 0.7290069448268927 | 11.73784964697929 | 0.0 |
| SE_31_11_14 | SE_31 | SE_31_11 | GCAACAGGATACTTACTGAG | 1870.598828754004 | 2164.684121717019 | 2222.9229677903586 | 1919.4300221104406 | 2342.8532786192595 | 2216.0212610433896 | 1604.5554736320894 | 2492.8971545966597 | 2152.4763134482114 | 2351.7764040115558 | 2683.272429299466 | 2115.9214862030517 |
| SE_31_11_35 | SE_31 | SE_31_11 | CATCTCCCTCACCAGCACAC | 1205.1598479573884 | 1236.2485034921297 | 1320.3234153423853 | 1000.7062284765619 | 1227.1078512279237 | 1377.43307469077 | 1107.3692705348224 | 1017.200355838455 | 1198.7801250639793 | 1253.1629381574285 | 1024.714274181292 | 1484.5110152790473 |
| SE_31_11_39 | SE_31 | SE_31_11 | TCTCCATCTGACATAGCGAC | 139.80721257779055 | 185.8870021419047 | 160.96687860943067 | 131.43215744620645 | 61.51448173830559 | 203.58287581907607 | 119.45382801687587 | 76.57363273505099 | 95.01442472729316 | 116.64111117230283 | 143.20176569314734 | 293.6522962201712 |
| SE_31_12_14 | SE_31 | SE_31_12 | GCTAGCTCCTGGGACATCCA | 923.3778691184307 | 1199.2709815606754 | 1062.5789041947694 | 1077.4834293609795 | 1118.927210929524 | 961.6042219539337 | 1005.1340123221808 | 1016.2550023478989 | 1165.0366845066228 | 1376.3651118331734 | 872.1222287705613 | 1248.8927696952735 |
| SE_31_12_20 | SE_31 | SE_31_12 | CTTGGGATCTGGGCACAGAG | 577.6530566198633 | 651.6038999813003 | 650.7802024761645 | 575.178352388349 | 625.75076251035 | 672.2566452578852 | 624.1731554034956 | 820.5668298027686 | 610.0458858658916 | 656.1062503442034 | 615.0633215017148 | 913.4559570168961 |

| | | | | | | | | | | | | | | | |
|---|---|---|---|---|---|---|---|---|---|---|---|---|---|---|---|
| SE_31_12_24 | SE_31 | SE_31_12 | GTGAGCCATAGGCTGCAAAA | 231.92824412129596 | 289.82382054382987 | 268.607306636596 | 229.03029416368648 | 279.9969513605634 | 139.4759276888138 | 221.6890862295174 | 352.61685197745703 | 186.47690834328566 | 270.46157653077717 | 370.91604884454557 | 95.17584304369186 |
| SE_31_12_3 | SE_31 | SE_31_12 | ACAGAGGCTCTCTTTGACTA | 854.0161512503795 | 1027.3754742036454 | 1153.4313755571475 | 969.4748247269683 | 1251.5015250207 | 1153.0587562348156 | 841.5575991819543 | 1332.0030681936646 | 1113.5335383927563 | 1177.3462158954317 | 1460.1884960842237 | 1295.3200102043916 |
| SE_31_12_8 | SE_31 | SE_31_12 | GGATTCCCTTTTGTCCCCCA | 21.675536833765978 | 37.97691441608806 | 24.688171565863602 | 28.62878677046081 | 27.575457330964575 | 6.064170769078861 | 7.533124289352532 | 17.961716320567515 | 29.303514168230603 | 36.45034724134463 | 15.259204541073078 | 3.482043038183848 |
| SE_31_13_26 | SE_31 | SE_31_13 | CTTGCAGCGAACTCCGGCCC | 2916.4434809832123 | 2895.2400279843973 | 2896.4162881071175 | 2978.695132671491 | 3104.3601387589733 | 3370.8126374979769 | 2921.776063656126 | 2698.9842155379083 | 3002.278224326899 | 3013.7147099143745 | 3108.1825865201163 | 2880.8102735907705 |
| SE_31_13_29 | SE_31 | SE_31_13 | GGCTGGAGCGGAGCGCACCG | 740.21958287310832 | 872.4696390853913 | 909.5122404864151 | 888.7936983738515 | 919.5354425363956 | 849.8502177809089 | 751.1601077097239 | 978.4408627256515 | 1079.7900978354064 | 903.9686115853469 | 1536.4845187895892 | 1421.834240591738 |
| SE_31_13_32 | SE_31 | SE_31_13 | CGCTCTGGGCACCACCGCCG | 123.55055995246607 | 170.89611487239625 | 174.7922546863143 | 163.96486968536647 | 178.17987813854035 | 145.54009845789267 | 186.175786008284 | 218.37665631847872 | 253.07580418017338 | 196.10286815843412 | 107.98821675220947 | 261.15322786378863 |
| SE_31_13_33 | SE_31 | SE_31_13 | CGCGGGTTCCCCCGAAGTTC | 834.5081689999901 | 926.4368332556218 | 979.6266477334677 | 927.8329530608435 | 1019.2313267329599 | 1372.2352140315595 | 675.8288648161986 | 846.9071740477856 | 701.5083694818841 | 610.90781697436 | 671.4049998072154 | 644.1779620640119 |
| SE_31_13_42 | SE_31 | SE_31_13 | CCACCGCCGCCGTCGCCGCG | 22.759313675454276 | 24.984812115847404 | 27.650752153767232 | 14.314393385230405 | 59.39329271284678 | 14.727271867762948 | 10.761606127646475 | 44.431614056140695 | 29.303514168230603 | 10.93510417240339 | 38.73490383503166 | 10.446129114551544 |
| SE_31_2_16 | SE_31 | SE_31_2 | AGAGCTTGTGGGAGATAGAC | 652.4336586963559 | 660.5984323430054 | 643.8675144377227 | 642.8463938458018 | 622.5689789721617 | 874.9732109670928 | 599.4214613099086 | 904.703290462269 | 546.1109458624794 | 441.0492016202701 | 1213.6936534976587 | 531.5919038294008 |
| SE_31_2_18 | SE_31 | SE_31_2 | ATTTAAGTGAGGCTGGTCCA | 24.926867358830876 | 40.97509186998975 | 45.42623568118903 | 31.23140374959361 | 32.87842989461161 | 45.91443582302566 | 8.60928490211718 | 12.289595377230405 | 50.615160836034676 | 93.31288893784226 | 2.347569929395858 | 76.60494684004466 |
| SE_31_2_37 | SE_31 | SE_31_2 | GATAGGCCCAGTGGACCCTG | 720.7115997227188 | 807.5091275841881 | 821.6223497119406 | 730.0340626467506 | 750.90091501242 | 548.3742995467027 | 691.9712740076683 | 851.7634949911227 | 903.0810275481977 | 820.1328129302542 | 705.4447637834553 | 698.7299696622256 |
| SE_31_2_42 | SE_31 | SE_31_2 | GACGTGAGTCACCTCTAGGA | 1982.2278434478987 | 2501.4793890386422 | 2496.4679087401273 | 2052.1634880462134 | 2658.9104434126225 | 2473.315363674307 | 1833.7776841509592 | 2474.9354382760926 | 2156.9162398373373 | 2609.1158555354486 | 1903.8792127400409 | 2072.9762887321176 |
| SE_31_2_6 | SE_31 | SE_31_2 | GTCACCCCCTACCCTTGGCA | 410.75142299986527 | 408.75152621526354 | 399.9483793669903 | 452.8553543691074 | 438.025533757245 | 480.80211097696684 | 449.83513613562263 | 349.78079150578844 | 457.3124180799624 | 447.6102641237121 | 404.9558128207855 | 343.561579767473 |
| SE_31_3_19 | SE_31 | SE_31_3 | GCGCCACGCGGCGTGAGGGG | 716.3764923559655 | 773.5297831066356 | 792.9840706955389 | 683.1869570223603 | 757.2644820887964 | 381.17644834209983 | 626.3254766290248 | 411.2287683919405 | 567.4225925302835 | 293.7897987652377 | 318.0957254331388 | 203.11917727739116 |
| SE_31_3_20 | SE_31 | SE_31_3 | TTTTGCTGATGAGGAAGCTG | 83.45081680999901 | 91.94410858631845 | 108.62795448979984 | 71.57196692615203 | 103.93826224748186 | 103.95721318420905 | 80.71204595734856 | 114.38777235729839 | 41.73530805778298 | 99.87395144128429 | 57.515463270198524 | 19.73157721637514 |
| SE_31_3_23 | SE_31 | SE_31_3 | GTCCCGCGTGTTAGAGCCCG | 307.7926230394769 | 291.8226055130977 | 313.04601545515044 | 291.4931016628737 | 328.7842989461161 | 430.5561246045991 | 356.2091628250983 | 420.6823032975023 | 434.224800856508 | 146.5303959102054 | 403.7820278560876 | 437.57674179843696 |
| SE_31_3_42 | SE_31 | SE_31_3 | TTAGGGACTGGGCCAGCCTG | 1520.5389088886834 | 1310.203547355038 | 1267.9844916227546 | 1518.6270073239893 | 1594.0735526322983 | 1675.4437524855025 | 1811.1783112829016 | 1398.1778125325977 | 1284.0267117351955 | 2162.2345983565638 | 1574.045637659923 | 1556.47323806818 |
| SE_31_3_9 | SE_31 | SE_31_3 | AGCTGAGTCCACGCGGGCCG | 1837.0017466616666 | 1794.9089024024775 | 1793.348782544332 | 1942.8535749226357 | 1963.160443062132 | 1821.8501610532635 | 1951.0791909423058 | 2550.563717520587 | 2038.81419788659 | 1694.2121397776984 | 2448.51543635988 | 1548.3484709790846 |
| SE_31_4_16 | SE_31 | SE_31_4 | ATATACAGATAAGGAAACTG | 36.84841261740216 | 60.96294156266766 | 62.21419234597627 | 41.641871666124814 | 42.42378050917627 | 77.96790998815578 | 35.513300221233365 | 6.617474433893295 | 65.71091055906257 | 18.225173620672315 | 10.564064682281362 | 83.56903291641235 |
| SE_31_4_28 | SE_31 | SE_31_4 | TTTCAAGTCAGGATTCTCGT | 20.59175999207768 | 44.97266180852533 | 45.42623568118903 | 16.917010364363207 | 63.6356707637644 | 24.256683076315444 | 3.228481838293942 | 33.08737216946648 | 32.855455279531284 | 11.664111117230282 | 14.085419576375148 | 5.803405063639747 |
| SE_31_4_29 | SE_31 | SE_31_4 | CACCGGCTAGAGCTCCAGAG | 544.055974527526 | 857.4787518158829 | 783.1088020691934 | 654.5581702518995 | 701.0529729141379 | 546.6416793269659 | 514.4047729015015 | 467.9499778253116 | 656.2211203128005 | 559.8773336270535 | 394.39174813850417 | 320.34795951291403 |
| SE_31_4_41 | SE_31 | SE_31_4 | GAATCCTCCACAGGGCCAGT | 50.93751155935005 | 104.9362108865591 | 86.90236391183987 | 87.18766880094883 | 96.51410065837601 | 64.10694813026224 | 50.57954879993843 | 66.17474433893295 | 71.03882226601358 | 136.32429868262892 | 38.73490383503166 | 177.58419494737626 |
| SE_31_4_44 | SE_31 | SE_31_4 | GCTTCCCAGAATCCTCCACA | 126.80189047753097 | 210.8718142577521 | 213.30580232906152 | 163.96486968536647 | 155.90739337122278 | 76.23528966841997 | 129.1392735317577 | 151.2565588489896 | 206.90056973326455 | 228.1791737308174 | 213.62886357502308 | 40.62383544547823 |
| SE_31_5_14 | SE_31 | SE_31_5 | TGTGAGACTGACTTCAGGGA | 997.074694353235 | 1197.2721965914077 | 1055.6662161563277 | 957.7630483208707 | 879.2328510526781 | 933.0159883282762 | 939.4882149435373 | 1013.4189418762303 | 1259.1631239560907 | 947.7090282749605 | 907.3357777114992 | 1223.3577874152586 |
| SE_31_5_17 | SE_31 | SE_31_5 | AGATGGCAAAACGCTCCCTC | 8671.298510348079 | 8782.66115496268 | 8769.23854019475 | 8432.479012390275 | 8765.813647708546 | 8916.929960875532 | 8337.016267087723 | 8278.460516800513 | 8030.05086737301 | 8682.472712888291 | 8589.758371659445 | 8596.003580263194 |
| SE_31_5_2 | SE_31 | SE_31_5 | GGGCAGAGCATGCCAGGCAG | 1600.7383951736174 | 1682.9769441234812 | 1649.1698605996885 | 1679.9892600300223 | 1658.7698179087922 | 1632.128246992082 | 1618.5455615980297 | 1940.8107161118478 | 1875.4249067667586 | 1753.2617023086768 | 2320.5728752078057 | 1728.2540279519167 |
| SE_31_5_21 | SE_31 | SE_31_5 | TGATCACTTCAGAGAGCATG | 546.2235282109026 | 628.6178728347207 | 580.6657952291118 | 564.7678844718178 | 620.447789946703 | 461.74328855986187 | 503.643166773855 | 557.7585594281492 | 586.070283364612 | 494.9957155374601 | 600.9779019253397 | 646.4993240894678 |
| SE_31_5_22 | SE_31 | SE_31_5 | CGCTCCCTCCGGGAGAGGGA | 0.0 | 0.9993924846338962 | 1.975053725269788 | 1.3013084895664004 | 1.0605945127294067 | 0.0 | 9.685445514881827 | 0.0 | 0.0 | 17.496166675845423 | 0.0 | 0.0 |
| SE_31_6_11 | SE_31 | SE_31_6 | AATGCAAAACTGAATCCTGA | 1322.2077468597247 | 1607.023115291305 | 1521.7788953198324 | 1262.2692348794085 | 1565.4375007886044 | 1432.0106116124796 | 1278.478807964401 | 1856.6742554523473 | 1403.0167389637681 | 1421.5635424124407 | 1703.161983776695 | 1651.649081111872 |
| SE_31_6_24 | SE_31 | SE_31_6 | GGAACAGCCTCCCTGACAGA | 1360.1399363188152 | 1538.065033851566 | 1498.0782506166033 | 1374.181764982119 | 1525.1349093048868 | 1851.3047047887894 | 1388.2471904663953 | 1516.3469988521208 | 1553.974236194047 | 1383.6551812814423 | 2082.294527374126 | 1376.567681095348 |
| SE_31_6_36 | SE_31 | SE_31_6 | CTCCTGTCTCCGGTTTCCTA | 8.670214733506391 | 24.984812115847404 | 27.650752153767232 | 14.314393385230405 | 12.727134152752882 | 38.11764483420998 | 5.380803063823238 | 37.8141396224274 | 49.72717555820951 | 21.870208344806678 | 19.954344399864794 | 112.5860582346111 |
| SE_31_6_39 | SE_31 | SE_31_6 | GGAGAGAAAGGGGCTGAGAG | 9464.623158463914 | 13194.978974621332 | 12374.699115673471 | 9778.031990601932 | 10417.159304028233 | 8768.790932088034 | 8188.506102526203 | 7365.249044923237 | 10596.328320287752 | 6913.172857793423 | 6458.164875768006 | 6978.014248520432 |
| SE_31_7_17 | SE_31 | SE_31_7 | CAAAAGTTAACCAGGTGTGG | 190.74472431714062 | 246.84994370457235 | 273.54494049497687 | 219.92113473672168 | 186.6646342403756 | 187.12298373157628 | 188.3281073132183 | 221.21271679014728 | 177.59705556503397 | 314.93100016521765 | 248.84241251596094 | 190.3516860873837 |
| SE_31_7_18 | SE_31 | SE_31_7 | ATACAAAAGTTAACCAGGTG | 17.340429467012783 | 46.97144677779312 | 40.48860136801630 | 18.218318853929606 | 14.848323178211695 | 4.331550549342044 | 24.75169409358689 | 27.415251226129364 | 23.975602501279585 | 0.7290069448268927 | 7.042709788187574 | 12.767491140007444 |
| SE_31_7_20 | SE_31 | SE_31_7 | GAGTTTCACCATGTTGTCCA | 342.47348197350243 | 372.77339676844326 | 357.48472427370496 | 366.9689940577249 | 389.2381861716923 | 356.05345515591597 | 385.2654993697438 | 467.0046243347554 | 321.4506705727115 | 282.12568764800744 | 534.0721589375577 | 445.70150888753255 |
| SE_31_7_23 | SE_31 | SE_31_7 | CAGGAGTTTTGAGACCACCC | 4888.917332855916 | 5359.741895091585 | 5259.568070391581 | 5030.858620663704 | 5296.608996570657 | 5582.5023479920255 | 5271.034681321244 | 5542.607515130912 | 5997.452566431197 | 7050.226163420879 | 4787.868871002853 | 6531.152058620171 |
| SE_31_7_3 | SE_31 | SE_31_7 | TCTTCTCCCCGCCTGCTACA | 299.1224083059705 | 407.7521337306296 | 382.1728958395685 | 321.4231969229009 | 345.7538111497866 | 407.16575163815213 | 357.285323437863 | 236.33837263904624 | 460.86435919126313 | 823.7778476543887 | 291.0986712450864 | 501.41419749847415 |
| SE_31_8_1 | SE_31 | SE_31_8 | AGTAGGGAGTTACATACTTT | 330.5519367149312 | 445.7290481467177 | 420.6864434823158 | 343.5454412455297 | 403.0259148371746 | 704.3101193230164 | 336.83827179533466 | 333.7097821663333 | 396.9294191878509 | 371.7935418617152 | 362.69955409166005 | 302.9377443219948 |
| SE_31_8_11 | SE_31 | SE_31_8 | CCTGAGCGAGAAGGGGACAG | 9143.825213324179 | 9785.051817050478 | 9657.025189703207 | 7751.894672347047 | 7100.680262723378 | 6032.11729501373 | 7872.1148823733965 | 5210.788439945692 | 7575.402405126523 | 4238.446377223554 | 3497.8791947998284 | 4166.844835693339 |
| SE_31_8_20 | SE_31 | SE_31_8 | CAGTTTCCAGACTCCCCGGT | 162.56652625324483 | 200.87788941141312 | 186.64257703792882 | 184.78580551842887 | 283.17873489875516 | 225.24062856578627 | 122.6823098551698 | 276.043219242406 | 219.33236362281693 | 137.7823125722827 | 483.5994054555468 | 449.1835519257164 |
| SE_31_8_36 | SE_31 | SE_31_8 | GGGCAGATGTCACGCAGCCC | 3930.8586048034604 | 3198.0559508284678 | 3117.6223053372555 | 3914.3359366157324 | 3193.4500778282436 | 2710.684333778251 | 4452.076455007346 | 3411.780747417272 | 3064.437193774661 | 3947.572606237624 | 3844.1457593857176 | 2862.239377387123 |
| SE_31_8_41 | SE_31 | SE_31_8 | GCGGAGCCGCCTCCTCCCCT | 230.84446727960767 | 309.81167023650784 | 192.56773821373608 | 287.5891761941745 | 312.875381255175 | 170.66309164407653 | 264.7355107401033 | 226.8848377334844 | 223.77229001194277 | 179.3357084274156 | 362.69955409166005 | 77.76562785277261 |
| SE_31_9_23 | SE_31 | SE_31_9 | GCTCACTTCCCAGTATTTCG | 720.7115997227188 | 768.5328206834662 | 816.6847153987679 | 762.5667748859106 | 835.7484760307725 | 578.695153392097 | 679.0573466549926 | 674.0370387665599 | 954.5841736620575 | 751.6061601165263 | 591.5876222077562 | 631.4104709240045 |
| SE_31_9_24 | SE_31 | SE_31_9 | CATTTACACACCCAAATCAG | 835.5919449416784 | 927.4362257402556 | 894.6993375468969 | 970.7761332165347 | 1025.5948938093363 | 1114.0748012907736 | 854.4715265351301 | 1083.375100177388 | 951.0322325507568 | 1046.124965826591 | 1035.2783388635735 | 1331.301121598958 |
| SE_31_9_25 | SE_31 | SE_31_9 | GGTTGGGGCAGGCACAAAGA | 658.9363197464858 | 643.6087601042292 | 679.4184814925663 | 722.2262117093522 | 680.9016771722792 | 580.4277736118338 | 848.0145628585423 | 691.9987550871274 | 578.0784158641856 | 1008.9456116404194 | 739.4845277596953 | 771.8528734640864 |
| SE_31_9_26 | SE_31 | SE_31_9 | GAAGGAAGACTTCAGCTAAA | 1752.4671530099793 | 2437.518270022073 | 2262.4240422957405 | 2105.5171361184357 | 2369.3681414374946 | 3028.620144099957 | 2081.294625086828 | 2202.6736329959112 | 2528.0940859682582 | 2773.8714250663265 | 2061.166398009563 | 3117.589200187272 |
| SE_31_9_3 | SE_31 | SE_31_9 | GCAGCTGTACCCCAAGGCCC | 3054.0831398776263 | 2891.2424580458614 | 2714.7113453823617 | 3015.13177032535 | 2950.5739344132094 | 2708.9517135585143 | 2862.5872299539624 | 2877.656025253027 | 2838.0009479292426 | 2776.787452845634 | 2732.5713978167787 | 2661.441562185188 |
| SE_310_1_19 | SE_310 | SE_310_1 | TAGGAGGGTGACAAGCCACA | 14160.628213499314 | 12539.377504701495 | 12465.551587035849 | 12945.416854020655 | 13463.186744587088 | 11122.5555006005 | 14037.439032902063 | 13885.352069289245 | 13511.583987387783 | 12010.389416023056 | 13957.477015223074 | 12856.863577987497 |
| SE_310_1_21 | SE_310 | SE_310_1 | AGACATTTGAAAGAGAAACA | 113.79656837727138 | 156.9046200875217 | 160.96687860943067 | 156.15701874796804 | 215.30068608406958 | 138.6096175789454 | 158.19561007640317 | 236.33837263904624 | 138.5270333407265 | 177.14868759293492 | 178.4153146340852 | 82.4083519036844 |
| SE_310_1_24 | SE_310 | SE_310_1 | TCTTGGCTAAAACTGTCGCA | 751.0573512899912 | 771.5309981373679 | 783.1088020691934 | 838.0426672807619 | 925.899009612757 | 524.1176164703872 | 743.6269834203714 | 1109.8449979129612 | 768.1072653187719 | 844.1900421095417 | 904.9882077821032 | 582.6618683894307 |
| SE_310_1_33 | SE_310 | SE_310_1 | TATTAAGTCCTGGAGTCCAG | 115.96412206064798 | 80.95079125534559 | 112.57806234033802 | 89.79028578008163 | 109.24123481112889 | 63.24063802039384 | 95.77829453605362 | 130.45878169675353 | 125.20592417334893 | 256.6104445790662 | 230.0618530807941 | 105.6219721582434 |
| SE_310_1_7 | SE_310 | SE_310_1 | TCAGGACTAGTGTATGAAAT | 1616.9950477989419 | 1797.9070798563791 | 1721.2593215720103 | 1751.561226956375 | 1569.679878839522 | 2021.967796432866 | 1790.7312596403733 | 1631.6801246999753 | 1865.6570687106816 | 1985.8149177084556 | 2223.1487231378774 | 1901.1954988483812 |
| SE_310_2_0 | SE_310 | SE_310_2 | CAAGGAGCTGCTGCTGCAGA | 409.667646158177 | 98.93985597875572 | 224.1685978180415 | 301.9035695794049 | 98.63528968383483 | 1224.9624953539299 | 487.5007575823853 | 194.7428190545741 | 914.6248361599248 | 34.992333351690846 | 120.8998513638867 | 226.33279748195014 |
| SE_310_2_2 | SE_310 | SE_310_2 | TGCAGCAGCAGCTCCTTGAA | 1949.7145381972498 | 2108.718142577521 | 1981.9664133075298 | 2004.0150739322567 | 2175.279345608013 | 2306.1175124697043 | 2254.5564837419365 | 2525.03917327557 | 2420.647867351413 | 2843.1270848248814 | 2900.4226477685825 | 2445.5548938177894 |
| SE_310_2_3 | SE_310 | SE_310_2 | GCAGCAGCTCCTTGAAGGGT | 4337.274920436572 | 4628.186596339573 | 4571.261847135304 | 4528.553543691073 | 4718.58498713313 | 5028.930187786113 | 4156.132286497069 | 4956.488350986078 | 4266.7692599499405 | 4148.049516065019 | 4864.164893708218 | 4528.977311664458 |
| SE_310_2_4 | SE_310 | SE_310_2 | CAGCAGCTCCTTGAAGGGTC | 764.0626733902508 | 958.4173927639065 | 873.9612734315715 | 771.6759343128755 | 967.2621956092189 | 908.7593052519608 | 726.4084136161371 | 1072.97621178127 | 893.3131894921208 | 943.3349866059991 | 961.3298860876039 | 1027.2026962642353 |
| SE_310_2_5 | SE_310 | SE_310_2 | TTTCAGAGCCCGACCCTTCA | 26.010644200519174 | 47.97083962242702 | 52.338923719630834 | 71.57196692615203 | 67.87804881468203 | 90.96256153618292 | 41.970263897821255 | 2.836060471668555 | 166.05324695330674 | 53.94651391719005 | 3.521354894093787 | 127.67491140007444 |
| SE_310_3_11 | SE_310 | SE_310_3 | ATGAAGGATCCCTCTCCCAC | 2140.45926233439 | 2163.684729232385 | 2028.3801758513534 | 2223.9362086689785 | 2028.917302851355 | 2198.6950588460213 | 2312.6691568312276 | 2246.1598935614957 | 2180.0038570607917 | 2603.2837999768335 | 2428.5610919600153 | 2103.1539950630445 |
| SE_310_3_15 | SE_310 | SE_310_3 | ATAGTGGCTGGGCAGAGGGC | 1169.3952121816744 | 1226.2545786457906 | 1293.6601900512526 | 1388.4961583673494 | 1313.0160067590054 | 1185.9785404098516 | 1349.505408406868 | 1468.1339708337553 | 1167.7006403400983 | 1724.830431460428 | 1582.2621324128083 | 1261.660260835281 |
| SE_310_3_20 | SE_310 | SE_310_3 | CGTGGCTCTGGTTTTTATAG | 3305.5193671493116 | 4109.501896814581 | 4276.97884207021 | 3457.576656777926 | 4227.529727739416 | 4006.6842581413903 | 3310.2700448640558 | 3862.714362412572 | 4742.729368864232 | 4289.476863361437 | 3662.2090898575384 | 4396.659676213472 |
| SE_310_3_23 | SE_310 | SE_310_3 | CCACACTCAAGCAAAAGAAG | 496.3697934932409 | 537.6731567330362 | 523.3892371963084 | 560.8639590031186 | 565.2968752847738 | 1023.1122397545907 | 542.3849488333823 | 658.911382917661 | 535.4551225285774 | 338.2592223996782 | 436.6480068676296 | 538.5559899057686 |
| SE_310_3_24 | SE_310 | SE_310_3 | CCCACACTCAAGCAAAAGAA | 279.6144251555811 | 243.85176625067066 | 278.4825752629414 | 321.4231969229009 | 225.90663121136365 | 317.0695002118376 | 323.9243444421589 | 256.19079594072616 | 128.75786528464963 | 299.62185432385286 | 227.71428315139823 | 600.0720835803498 |
| SE_311_1_0 | SE_311 | SE_311_1 | GCAGTGTCTGGTAGTCTGGG | 2529.53514850049 | 2863.2594684761125 | 2855.9276867391013 | 2544.058097102313 | 2622.850229979823 | 3121.315325855877 | 2556.957615928802 | 2980.6995557236514 | 2981.85456293692 | 3101.1955432936015 | 3200.9115987312525 | 3349.725402732862 |
| SE_311_1_1 | SE_311 | SE_311_1 | GTCATCATTCTGGTTTTGGC | 739.1358060314199 | 685.5832444588527 | 702.1315993331608 | 672.7764891025929 | 696.8105948632202 | 760.6202764644629 | 821.1105475339426 | 559.6492664092615 | 613.5978269771923 | 621.8429239373394 | 663.1885050543299 | 695.2479266240417 |
| SE_311_1_10 | SE_311 | SE_311_1 | CATGAAGCCAGGAATCCATG | 2094.9406349834817 | 2092.7278628233785 | 1941.4778119395135 | 2275.9885482516343 | 1851.798019225544 | 2138.9196612651012 | 2216.890862295174 | 1972.0073813002018 | 2297.2179137337143 | 1865.5287718120182 | 2032.995558856813 | 2028.8704102484555 |
| SE_311_1_11 | SE_311 | SE_311_1 | GTGACAGTTCACAAGTGCCA | 1376.3965889441397 | 1349.17985425576 | 1445.7393268969724 | 1486.0942950848294 | 1603.618903246863 | 1397.3582072177433 | 1443.1313817173923 | 1714.8712318689195 | 1258.2751386782656 | 1457.2848827089583 | 1687.902779235622 | 1310.408863369855 |
| SE_311_1_17 | SE_311 | SE_311_1 | ATGTGTGTGGTGCTGTCACA | 3288.178937682299 | 2664.380364033967 | 2654.4722067616544 | 3176.4940230315833 | 2508.306022605047 | 2782.588072897329 | 3303.8130811874629 | 2165.8048466422 | 2400.2242065961434 | 2515.0739596527796 | 1987.2179452335938 | 2498.946220403275 |
| SE_311_2_14 | SE_311 | SE_311_2 | GCCCCGCGTCCTTCCACGCGG | 241.68223569649066 | 187.88578711117248 | 175.77978154894885 | 205.60674135149128 | 127.2713415275288 | 116.95186483223517 | 227.06988929334062 | 121.00524679119168 | 118.10204195074758 | 215.05704877393332 | 183.11045449287693 | 78.92630886550056 |
| SE_311_2_25 | SE_311 | SE_311_2 | AGTTTGCTGACTGACTCACC | 351.14369670700887 | 354.7843320450331 | 334.77160643311043 | 346.1480582246625 | 302.2694361278809 | 299.7432980144694 | 394.95094488462564 | 330.87372169466477 | 321.4506705727115 | 234.01122928943255 | 289.9248862803885 | 321.50864052564197 |
| SE_311_2_4 | SE_311 | SE_311_2 | TCCCATCCCTTGTGGGGCGT | 266.60910305532155 | 195.88092698824366 | 191.58021135110155 | 213.41459228888968 | 193.02820131675202 | 125.61496593091927 | 272.2686350294558 | 131.40413518730972 | 156.28540889722987 | 260.2554793032007 | 115.03092654039705 | 119.5501443109788 |
| SE_311_2_6 | SE_311 | SE_311_2 | CCCCCACGCCCCACAAGGGA | 19.50798315038938 | 9.993924846338961 | 15.800429802152705 | 15.615701874796805 | 32.87842989461161 | 3.465240439473635 | 2.152321225529295 | 12.289595377230405 | 81.69464555991561 | 3.6450347241344634 | 12.911634611677218 | 0.0 |
| SE_311_2_64 | SE_311 | SE_311_2 | GCGCCGCGTCCTCACCGCCG | 4753.445227644879 | 4428.308099412794 | 4462.633892245504 | 4659.98570113728 | 4294.347182041368 | 4276.106702310465 | 4646.861525917748 | 4186.9706096733435 | 4415.950786624569 | 4549.732342664637 | 4077.7289673606756 | 3885.9600306131747 |
| SE_311_3_18 | SE_311 | SE_311_3 | AGCTGAGACCACAGTCAGCT | 1106.536155363753 | 967.4119251256116 | 990.4894432224477 | 1047.5533341009523 | 901.5053358199957 | 1127.9357630486682 | 1002.9816910966514 | 709.0151179171388 | 895.0891600477711 | 1353.7658965435396 | 1090.446232204376 | 990.0609038569409 |
| SE_311_3_19 | SE_311 | SE_311_3 | AAGCTGAGACCACAGTCAGC | 966.7289427859627 | 1002.3906620877979 | 1126.7681502660148 | 938.2434209773747 | 955.5956559691955 | 885.3689322855137 | 929.8027694284576 | 975.6048022539829 | 1020.2950842211201 | 1062.5161186127827 | 1092.793802133772 | 669.7129443440268 |
| SE_311_3_20 | SE_311 | SE_311_3 | TCTCAGCTTTTCTAGCCTCC | 238.43090517142576 | 246.84994370457235 | 238.98150075755964 | 265.4669318715457 | 303.33003064061035 | 169.79678153420812 | 212.00364071463557 | 205.14170745069214 | 341.87433196269035 | 301.80887515833354 | 91.55522724643846 | 138.12104051462597 |
| SE_311_3_21 | SE_311 | SE_311_3 | CTAGCCTCCAGGCTTTCTCC | 119.21545258571288 | 119.92709815606754 | 104.67784743926167 | 122.32299801924164 | 173.93750008762227 | 153.33688944670834 | 119.45382801687587 | 168.27292131900094 | 69.26285167036325 | 28.431270848248815 | 109.1620017169074 | 212.40462532921475 |
| SE_311_3_29 | SE_311 | SE_311_3 | ACTGGAAATGCCCCATGACT | 1590.984403598423 | 1824.8906769414943 | 1809.1492123464848 | 1518.6270073239893 | 1952.5544979348379 | 2246.3421148887837 | 1576.5752977002085 | 2118.5371723364105 | 1900.2884945458634 | 1293.9873270677344 | 2298.270960878545 | 2084.583098859397 |
| SE_312_1_11 | SE_312 | SE_312_1 | GGGATTGGATGGGTGACACG | 471.44292613441 | 439.7326932389143 | 540.1771938610956 | 512.7155448891617 | 524.9942838010563 | 632.4063802039384 | 693.047434620433 | 380.03210320358636 | 539.8950489177032 | 734.8390003855078 | 332.1811450095139 | 550.162800033048 |
| SE_312_1_12 | SE_312 | SE_312_1 | TAGGCATTCATGGGAATGGA | 3051.91558619425 | 2897.238812953665 | 3003.0691892716486 | 3138.7560768341577 | 2834.969132525704 | 3148.1709392617972 | 2860.434908728433 | 2644.1537130856495 | 2964.9828426582417 | 2983.0961182316447 | 2602.2812667353087 | 2985.271564736286 |
| SE_312_1_21 | SE_312 | SE_312_1 | TCAAGGGTTCTAAGAAGTGA | 576.569279778175 | 616.625163019114 | 642.8799875750882 | 582.9862033257474 | 803.9306406488903 | 589.0908747105179 | 610.1830674375551 | 640.9496665970935 | 737.9157658727161 | 839.0869934957534 | 670.2312148425175 | 389.988820276591 |
| SE_312_1_4 | SE_312 | SE_312_1 | AAGAGTGAAGAGCACAGCAA | 939.6345217437552 | 920.4404783478184 | 1012.2150342004077 | 1028.0337067574565 | 1038.3220279620891 | 1127.0694529387997 | 1079.3890946029414 | 991.6758115934381 | 1165.0366845066228 | 1094.9684311299927 | 739.4845277596953 | 1368.4429140062523 |

| | | | | | | | | | | | | | | | |
|---|---|---|---|---|---|---|---|---|---|---|---|---|---|---|---|
| SE_312_1_8 | SE_312 | SE_312_1 | CATGTCTGCTAGGCATTCAT | 1869.5150519123156 | 1479.1008772581663 | 1673.8580321655522 | 1804.9148750285974 | 1666.193979497898 | 1020.5133094249854 | 1843.463129665841 | 1729.0515342272624 | 1676.5162045339205 | 1415.0024799089986 | 1883.924868340176 | 1333.622483624414 |
| SE_312_2_23 | SE_312 | SE_312_2 | AGCTGTTAGGAGTCAGGCTA | 320.7979451397365 | 366.7770418606399 | 368.3475197626849 | 318.8205799437681 | 377.5716465316688 | 303.20853845394305 | 405.7125510122721 | 458.4964429197497 | 260.17968640277473 | 266.8165418066427 | 301.66273592736775 | 443.3801468620767 |
| SE_312_2_3 | SE_312 | SE_312_2 | GATGGATATGATGACCTGGG | 1113.038816413883 | 1172.2873844755602 | 1339.0864257324417 | 1167.2737151410613 | 1052.1097566275714 | 736.3635933881474 | 1110.5977523731162 | 1330.1123612125523 | 948.3682767172813 | 790.2435281923516 | 1373.328408696577 | 1292.9986481789356 |
| SE_312_2_4 | SE_312 | SE_312_2 | CTAATTCTCAACCTTCATGC | 286.1170862057109 | 226.86209401189444 | 234.04386644438694 | 343.54544124552973 | 307.572408691528 | 120.41710527170882 | 326.0766656676882 | 372.46927527913687 | 225.54826056759313 | 395.8507710410027 | 367.39469395045177 | 210.08326330375886 |
| SE_312_2_5 | SE_312 | SE_312_2 | AAGATCTCGTCCCTAAGAAG | 406.4163156331121 | 567.654931272053 | 587.5784832675537 | 510.112927910029 | 520.7519057501387 | 440.08553581315164 | 329.3051475059821 | 232.55695867682152 | 661.5490319797515 | 269.0035626411234 | 129.1163461167722 | 211.2439443164868 |
| SE_312_2_8 | SE_312 | SE_312_2 | TTTCTCATCCGTAATCACAT | 13.005322100259587 | 10.993317330972857 | 14.81290293951816 | 15.615701874796805 | 4.242378050917627 | 0.8663101098684087 | 4.30464245105859 | 19.852423301679885 | 38.1833669464823 | 1.4580138896537853 | 9.390279717583432 | 0.0 |
| SE_313_1_11 | SE_313 | SE_313_1 | GGCCTTTTACAAAGAGGAGG | 3.2513305250648967 | 12.992102300240651 | 5.925161175807264 | 6.506542447832002 | 7.4241615891058474 | 0.0 | 2.152321225529295 | 4.726767452780925 | 0.8879852778251698 | 0.0 | 16.432989505771005 | 0.0 |
| SE_313_1_27 | SE_313 | SE_313_1 | GGCATCTGTGGAGATAAGAG | 123.55055995246607 | 331.7983048984535 | 289.3453707519214 | 184.78580551842887 | 284.239329411481 | 106.55614351381428 | 87.16900963393644 | 104.93423745173654 | 249.5238630688727 | 114.45409033782215 | 36.3873339056358 | 59.19473164912542 |
| SE_313_1_32 | SE_313 | SE_313_1 | AGAGCGGCAGGGGATCCACT | 5981.364389277722 | 6055.319064396777 | 6206.606331658109 | 5651.5827701868775 | 6217.205033619783 | 5893.5076774347845 | 5695.041962750514 | 6431.239796253727 | 6249.640385333545 | 6241.757461607855 | 5522.658258903756 | 6995.424463711352 |
| SE_313_1_6 | SE_313 | SE_313_1 | GGATGTTCCCAGGGAGACTG | 503.95623138505897 | 631.6160502886224 | 633.9922458113773 | 534.8377892117906 | 601.3570887175737 | 416.6951628467046 | 514.4047729015015 | 634.3321921632001 | 643.7893264232481 | 653.1902225648959 | 608.0206117135273 | 507.2176025621139 |
| SE_313_1_7 | SE_313 | SE_313_1 | CGTGTGCTCTTAAAGTTTGG | 5151.191328544484 | 5343.751615337443 | 5333.632585089173 | 5543.574165552865 | 5182.0647891958815 | 5097.368686465717 | 5296.862536027595 | 5286.416719190187 | 5413.158253622235 | 6183.436906021703 | 5488.618494927516 | 6315.265390252773 |
| SE_313_2_0 | SE_313 | SE_313_2 | GCATTGCTATACAGAAAACA | 888.6970101844051 | 1000.39187711853 | 1033.9406251783676 | 1021.5271643096244 | 1035.140244423901 | 1196.3742617282724 | 933.0312512669493 | 860.2716764061283 | 1167.7006403400983 | 803.3656531992357 | 785.2621413829145 | 1085.2367469006326 |
| SE_313_2_12 | SE_313 | SE_313_2 | CTCTCTGCGATATATTTGCA | 4.335107366753196 | 10.993317330972857 | 16.78795666478725 | 6.506542447832002 | 1.0605945127294067 | 6.064170769078861 | 2.152321225529295 | 12.289595377230405 | 0.0 | 11.664111117230282 | 0.0 | 0.0 |
| SE_313_2_15 | SE_313 | SE_313_2 | ACTTGCAGTCACACACATGT | 926.6291996434956 | 1012.3845869341368 | 985.551808909275 | 845.8505182181602 | 967.2621956092189 | 930.417057998671 | 955.630624135007 | 789.3701646144145 | 707.7242664266603 | 1507.586361902014 | 674.9263547013092 | 950.5977494241906 |
| SE_313_2_35 | SE_313 | SE_313_2 | GTACAGACCTTCCTTGTCCT | 1072.939073271416 | 868.4720691468558 | 913.4623479369533 | 957.7630483208707 | 772.1128052670081 | 759.7539663545945 | 1136.4256070794677 | 761.0095598977289 | 904.856998103848 | 1156.2050144954517 | 480.078050561453 | 568.7336962366952 |
| SE_313_2_5 | SE_313 | SE_313_2 | TGCTGGATGGTGACCCCAAG | 1551.968437297644 | 1612.0200777144746 | 1546.4670668856959 | 1591.5002827397077 | 1628.0125770396394 | 1487.4544586440577 | 1414.0750451727467 | 1269.6097378169566 | 1971.327316771877 | 1154.0179936609711 | 1101.0102968866574 | 1111.9324101933755 |
| SE_313_3_1 | SE_313 | SE_313_3 | ATAGGATGTGAGCTAATTAA | 233.01202096298425 | 282.82807315139263 | 304.15827369143955 | 236.83814510108488 | 328.7842989461161 | 258.1604127407858 | 179.71882233169612 | 103.98888396118035 | 178.48504084285912 | 479.6865696960954 | 123.24742129328254 | 316.86591647473017 |
| SE_313_3_19 | SE_313 | SE_313_3 | GGTTTCGAGCACCTACTCCA | 50.93751155935005 | 66.95929647047105 | 65.1767729338799 | 32.53271223916001 | 34.999618920070425 | 33.78609428486794 | 29.056336544645482 | 41.59555358447214 | 116.32607139509724 | 110.0800486688608 | 32.86597901154201 | 77.76562785277261 |
| SE_313_3_5 | SE_313 | SE_313_3 | GAGGGTCTGAACCCAAGTGA | 978.6504880445339 | 1087.3390232816791 | 1076.404280271653 | 1038.4441746739876 | 1088.1699700603713 | 1010.1175881065645 | 1067.5513278625303 | 1277.1725657414058 | 1238.7394625661118 | 1022.7967435921304 | 1354.5478492614102 | 1219.8757443770749 |
| SE_314_1_10 | SE_314 | SE_314_1 | CAGAACTTAAGAGAGGGTGA | 932.048083851937 | 1223.256401191889 | 1132.693311441822 | 1059.26511050705 | 1084.988186522183 | 1058.6309542591955 | 936.2597331052433 | 745.88390404883 | 1130.405258671441 | 1405.525389626249 | 969.5463808404894 | 864.7073544823223 |
| SE_314_1_15 | SE_314 | SE_314_1 | GGAGGAACAGAGAAACGATG | 4312.3480530777415 | 4483.274686067658 | 4465.596472833408 | 4411.435779630097 | 4562.677593761908 | 4598.3740631815135 | 4176.579338139597 | 4257.872121465058 | 4316.49643550815 | 3778.442995037785 | 4537.852673522194 | 4714.6862737009305 |
| SE_314_1_2 | SE_314 | SE_314_1 | TCACAGTCGCCTGTATGCGG | 313.2115072479184 | 463.7181128701278 | 447.34966877344846 | 433.33572702561133 | 461.35861303729195 | 523.2513063605189 | 312.0865777017478 | 466.0592708441992 | 586.070283364612 | 384.1866599237724 | 556.3740732668184 | 573.3764202876071 |
| SE_314_1_24 | SE_314 | SE_314_1 | CAAGTACCAAAGGCTGCGAG | 1984.3953971312753 | 2353.5693013128257 | 2271.3117840594514 | 2002.7137654426904 | 2209.218370015354 | 1436.3421621618218 | 1844.5392902786057 | 2162.023432901995 | 2121.3968287243306 | 1995.2920079912053 | 2386.30483323089 | 2006.8174710066246 |
| SE_314_1_7 | SE_314 | SE_314_1 | ATGCAGTCAAACAAAGCAGA | 754.308681815056 | 751.5431484446899 | 790.0214901076353 | 801.6060295729027 | 797.5670735725139 | 732.8983529486737 | 835.1006355053664 | 761.9549133882851 | 753.8995008735692 | 867.5182643440022 | 551.6789334080266 | 530.4312228166729 |
| SE_314_2_1 | SE_314 | SE_314_2 | GCCCTCAAATTCTTTCAGAT | 26.010644200519174 | 55.96597913949819 | 58.264084895438096 | 26.02616979132801 | 51.96913112374093 | 56.31015714144657 | 20.4470516425283 | 51.994441980590175 | 79.03068972644012 | 11.664111117230282 | 41.08247376442752 | 10.446129114551544 |
| SE_314_2_10 | SE_314 | SE_314_2 | GATGGTATTTCACCCTGTGA | 487.6995787597345 | 515.6865220710904 | 587.5784832675537 | 486.68937509783376 | 645.9020582522087 | 427.9571942749939 | 435.84504816968223 | 700.506936502133 | 632.2455178115209 | 492.07968775815255 | 773.5242917359352 | 465.4330861039077 |
| SE_314_2_2 | SE_314 | SE_314_2 | TCTCAGTCCCTTGAGTAGCT | 331.63571355661946 | 396.7588163996568 | 382.1728958395685 | 403.40563176558413 | 351.0567837134336 | 304.0748485638115 | 297.0203291230427 | 315.7480658457658 | 350.75418474094204 | 260.98448624802757 | 183.11045449287693 | 330.7940886274656 |
| SE_314_2_3 | SE_314 | SE_314_2 | AGAGGCCAGAACTTAAGAGA | 1088.111949055052 | 1492.092979558407 | 1472.4025521881051 | 1385.8935413882164 | 1411.6512964428405 | 1341.0480500762967 | 1330.1345173771042 | 1405.740640457047 | 1464.287723133705 | 1817.4143134534434 | 1421.453592249192 | 1315.0515874207667 |
| SE_314_2_31 | SE_314 | SE_314_2 | TGGCTTTTAAAACCTTCCGC | 7.586437891818092 | 1.9987849692677924 | 3.950107450538176 | 5.205233958265602 | 16.969512203670508 | 1.7326202197368175 | 1.0761606127646475 | 7.56282792444948 | 0.0 | 0.0 | 10.564064682281362 | 0.0 |
| SE_315_1_13 | SE_315 | SE_315_1 | GTAATTATTTTCAATCAGCT | 3.2513305250648967 | 13.991494784874547 | 13.825376076883616 | 7.807850937398403 | 6.363567076376441 | 0.8663101098684087 | 1.0761606127646475 | 3.78141396222474 | 10.655823333902038 | 0.0 | 0.0 | 1.1606810127279494 |
| SE_315_1_16 | SE_315 | SE_315_1 | GGGTTTGAGGAGAAGCCGTT | 39.01596630077876 | 51.968409200962604 | 63.20171920861082 | 24.724861301761607 | 83.78696650562313 | 65.83956834999907 | 27.980175931880833 | 63.33868386726439 | 44.39926389125849 | 2.187020834480678 | 250.0161974806589 | 0.0 |
| SE_315_1_2 | SE_315 | SE_315_1 | AGAGAAATAAAAATTATCCA | 13.005322100259587 | 27.982989569749094 | 45.42623568118903 | 24.724861301761607 | 36.06021343279983 | 47.64705604276248 | 10.761606127646475 | 59.557269905039654 | 15.095749723027886 | 3.6450347241344634 | 32.86597901154201 | 56.873369623669525 |
| SE_315_1_21 | SE_315 | SE_315_1 | TTGGGGCTAAGAGTGAGGGG | 6169.9415597314855 | 7307.557847643049 | 7405.463942896446 | 6182.5166339299685 | 6661.594134453404 | 6605.614587746617 | 4972.938191585436 | 4892.204313628257 | 6280.719870057426 | 3537.1416963000834 | 3958.002900961417 | 3542.3984508457015 |
| SE_315_1_28 | SE_315 | SE_315_1 | ATTGTTTCTGGGCTACCCTT | 10.837768416882989 | 13.991494784874547 | 17.77548352742179 | 7.807850937398403 | 11.666539640023474 | 14.727271867762948 | 15.066248578705064 | 81.30040018783191 | 1.7759705556503396 | 1.4580138896537853 | 68.07952795247988 | 0.0 |
| SE_315_2_1 | SE_315 | SE_315_2 | TTAACAAAACACCTGCACCC | 444.3485050922026 | 427.7399834233076 | 393.0356913285485 | 468.4710562439042 | 383.9352136080452 | 344.7914237276267 | 435.84504816968223 | 397.0484660335977 | 289.48320057100534 | 522.697979440882 | 244.14727265716925 | 204.2798582401191 |
| SE_315_2_2 | SE_315 | SE_315_2 | CATAAAACCTTTTTGAACAG | 320.7979451397365 | 399.75699385355847 | 390.0731107406449 | 433.33572702561133 | 399.8441312989863 | 371.64703713354737 | 364.81844772721547 | 397.0484660335977 | 507.03959363817194 | 646.6291600614538 | 453.0809963734006 | 412.04175951842205 |
| SE_315_2_21 | SE_315 | SE_315_2 | CTATTTTTCTGATAGCCACG | 0.0 | 0.9993924846338962 | 0.0 | 0.0 | 0.0 | 0.0 | 0.0 | 0.0 | 0.0 | 0.0 | 0.0 | 0.0 |
| SE_315_2_3 | SE_315 | SE_315_2 | TTGCCATCGAAAAGCCCGAT | 105.126353643765 | 155.9052276028878 | 133.31612645566344 | 108.00860463401123 | 156.9679878839522 | 98.75935252499859 | 64.56963676587885 | 75.6282792444948 | 232.6521427901945 | 74.35870837234305 | 86.86008738764674 | 49.909283547301825 |
| SE_315_2_9 | SE_315 | SE_315_2 | ATTGTCACAGAAGATAGTGC | 328.3843830315546 | 320.8049875674807 | 297.24558565299776 | 307.1088035376705 | 290.60289648785744 | 715.5721507513056 | 332.53362934427605 | 188.1253446206808 | 312.57081779445974 | 158.19450702743572 | 190.1531642810645 | 398.11358736568667 |
| SE_316_1_12 | SE_316 | SE_316_1 | AGGGGTCCCGGGCAGCTAAG | 904.9536628097296 | 1223.256401191889 | 1174.169439672473 | 1184.1907255054243 | 1244.0773634315942 | 1322.8555377690602 | 1169.7865860751717 | 1275.2818587602935 | 1014.0791872763439 | 1526.5405424675132 | 1085.7510923455843 | 1136.3067114606624 |
| SE_316_1_2 | SE_316 | SE_316_1 | ACATGTAAACTTATTTTACG | 798.7435323242763 | 938.4295430712285 | 1017.1526685135804 | 895.3002408216835 | 847.415015670796 | 869.7753503078824 | 856.6238477606594 | 1024.7631837629046 | 903.9690128260229 | 835.4419587716189 | 1071.665672769209 | 766.0494684004466 |
| SE_316_1_7 | SE_316 | SE_316_1 | TGGATATTTCCCTTAGCCAG | 17.340429467012783 | 22.986027146579612 | 28.63827901640178 | 36.43663770785921 | 67.87804881468203 | 48.51336615263089 | 16.142409191469714 | 55.775855942814914 | 70.15083694818841 | 5.832055558615141 | 35.21354894093787 | 8.124767089095647 |
| SE_316_1_8 | SE_316 | SE_316_1 | TGTTTCCTCTCACATTATGG | 16.256652625324485 | 5.9963549078033775 | 11.850322351614528 | 20.820935833062407 | 13.787728665482287 | 1.7326202197368175 | 13.990087965940416 | 3.78141396222474 | 34.63142583518162 | 18.225173620672315 | 0.0 | 0.0 |
| SE_316_1_9 | SE_316 | SE_316_1 | TTGCTTCCTGACAGATGCAG | 162.56652625324483 | 103.93681840192521 | 133.31612645566344 | 175.67664609146405 | 99.69588419656424 | 115.21924461249836 | 179.71882233169612 | 105.87959094229272 | 56.831057780810866 | 60.50757642063209 | 88.03387235234467 | 63.83745570003722 |
| SE_316_2_10 | SE_316 | SE_316_2 | TCAGGCATCGGGACATGCAG | 8118.572321087047 | 6458.074235704237 | 6750.733632969743 | 7338.078572664932 | 6841.895201617403 | 5917.7643605111 | 7798.9359607054 | 6709.173722477245 | 7474.172083454454 | 5369.865155594891 | 6898.334237529729 | 6888.64181054038 |
| SE_316_2_11 | SE_316 | SE_316_2 | CTGCATGTCCCGATGCCTGA | 2482.932744307893 | 2525.4648086698558 | 2534.9814563828745 | 2373.5866849691142 | 2386.337653641165 | 1832.2458823716845 | 2351.4109388907546 | 2491.9518011061036 | 2391.344353183182 | 2871.5583556731303 | 2361.6553489722332 | 1810.662379855601 |
| SE_316_2_14 | SE_316 | SE_316_2 | CCTCCATCTCCATCAGGCAT | 863.7701428255742 | 726.5583363288425 | 782.1212752065588 | 815.920422958133 | 697.8711893759496 | 667.0587845986747 | 768.3786775139583 | 860.2716764061283 | 756.5634567070447 | 739.2130420544692 | 920.2474123231764 | 711.497460802233 |
| SE_316_2_15 | SE_316 | SE_316_2 | AGGATTCCTCCATCTCCATC | 237.34712832973747 | 243.85176625067066 | 282.4326827134796 | 239.44076208021767 | 198.33117388039906 | 181.05881296249743 | 266.8878319656326 | 157.8740329228829 | 221.1083341784673 | 146.5303959102054 | 150.24447548133492 | 298.295020271083 |
| SE_316_2_8 | SE_316 | SE_316_2 | ATTGAGATATGTATTTCTGC | 58.52394945116814 | 73.95504386290831 | 62.21419234597627 | 76.77720088441762 | 73.18102137832906 | 27.72192351578908 | 68.87427921693744 | 46.32232103725307 | 113.66211556162173 | 8.748083337922711 | 76.29602270536539 | 98.6578860818757 |
| SE_317_1_14 | SE_317 | SE_317_1 | AGATGGTCATGTGAAAAGGG | 1428.4178773451779 | 1685.9751215773829 | 1697.5586768687813 | 1630.5395374266998 | 1546.3467995594751 | 1400.823447657217 | 1530.3003913513287 | 1742.2864830950489 | 1412.784577019845 | 1322.4185979159834 | 1313.4653754969827 | 1614.5072887045776 |
| SE_317_1_15 | SE_317 | SE_317_1 | GTGGAAGGAGAGAAGACGGT | 859.4350354588211 | 901.4520211397744 | 912.4748210743187 | 845.8505182181602 | 845.2938266453372 | 581.2940837217022 | 848.0145628585423 | 619.2065363143012 | 729.0359130944644 | 1000.9265352473236 | 642.0603756897672 | 728.9076759931522 |
| SE_317_1_22 | SE_317 | SE_317_1 | ATAGTTCCCTTCTAGACGTG | 131.13699784428417 | 172.89489984166403 | 182.69246958739063 | 137.93869989403845 | 155.90739337122278 | 202.71656570920763 | 118.37766740411122 | 170.1636283001133 | 184.70093778763533 | 88.20984032405401 | 163.15611009301213 | 183.38760001101602 |
| SE_317_1_5 | SE_317 | SE_317_1 | CGACTGACGTGACATGAACG | 4220.227021534236 | 3453.900426894745 | 3617.310897830335 | 3849.2705121374124 | 3653.7480963528064 | 3630.705670458501 | 3966.7280186504904 | 3546.0209430762498 | 3329.9447918443866 | 3540.7867310242177 | 3624.6479709872046 | 4352.55379772981 |
| SE_317_1_8 | SE_317 | SE_317_1 | TGACTTGCATCCATATCAGA | 757.5600123401209 | 801.5127726763848 | 866.0610585304951 | 743.0471475424147 | 850.5967992089842 | 925.2191973394605 | 671.5242223651401 | 723.1954202754815 | 616.2617828106678 | 704.9497156476052 | 957.8085311935101 | 775.3349165022702 |
| SE_317_2_16 | SE_317 | SE_317_2 | TTTTCCCCACAGCACACAGT | 41.18351998415536 | 43.97326932389143 | 55.301504307534465 | 22.12224432262881 | 40.30259148371746 | 203.58287581907607 | 19.370891029763655 | 90.75393509339376 | 42.62329333560815 | 29.160277793075707 | 34.03976397623994 | 61.51609367458132 |
| SE_317_2_19 | SE_317 | SE_317_2 | AAGTGTGTCATGGTGACAAA | 528.8830987438898 | 618.6239479883817 | 694.2313844320845 | 685.7895740014931 | 684.0834607104673 | 730.2994226190685 | 738.2461803565482 | 721.3047132943692 | 825.8263083774079 | 847.1060698888492 | 492.98968517313017 | 735.87176206952 |
| SE_317_2_2 | SE_317 | SE_317_2 | TGGTAGAAAAGGCTGACCGG | 20.59175999207768 | 8.994532361705065 | 6.912688038441808 | 9.109159426964803 | 6.363567076376441 | 18.192512307236584 | 22.599372868057596 | 68.06545132004533 | 2.6639558334755096 | 0.7290069448268927 | 183.11045449287693 | 1.1606810127279494 |
| SE_317_2_23 | SE_317 | SE_317_2 | GGATGGTTCAGGGTTTCGAT | 335.9708209233727 | 366.7770418606399 | 475.98794778985024 | 353.95590916206095 | 340.4508385861396 | 603.8181465782809 | 309.9342564762185 | 301.567763487423 | 438.66472724563386 | 324.40809044796725 | 333.35492997421187 | 169.4594278582806 |
| SE_317_2_6 | SE_317 | SE_317_2 | ACATGACGACAGATTTGGAA | 3672.919716481645 | 4136.485493899696 | 4137.73755443874 | 3807.628640471288 | 4321.922639372333 | 3456.5773383749506 | 3720.287238327386 | 4321.210805332322 | 4171.754835222648 | 3620.248488010349 | 3778.4138013626334 | 4576.565233186305 |
| SE_317_3_1 | SE_317 | SE_317_3 | CTCTTTGTAGTCCAAGAACA | 27.094421042207472 | 15.990279754142339 | 21.725590977959968 | 7.807850937398403 | 44.54496953463509 | 3.465240439473635 | 12.913927353175769 | 25.524544245016994 | 68.37486639253807 | 2.9160277793075706 | 3.521354894093787 | 1.1606810127279494 |
| SE_317_3_18 | SE_317 | SE_317_3 | GGAAAACTACTGAGAAAGCA | 163.65030309493312 | 293.8213904823655 | 260.7070917355196 | 229.03029416368648 | 253.4820885423282 | 452.21387735130935 | 270.1163138039265 | 254.30008895961376 | 366.7379197417951 | 316.38901405487144 | 123.24742129328254 | 277.40276204197994 |
| SE_317_3_19 | SE_317 | SE_317_3 | TTAATGAGAAGATGGAGCAG | 136.55588205272565 | 183.8882171726369 | 160.96687860943067 | 175.67664609146405 | 184.54344521491677 | 99.625662634867 | 131.291594757287 | 184.3439306584561 | 146.517570841153 | 196.831875103261 | 78.64359263476125 | 206.601220265575 |
| SE_317_3_2 | SE_317 | SE_317_3 | GATCCTGTAAGTACAGGATG | 26.010644200519174 | 46.97144677779312 | 37.526020780112674 | 31.23140374959361 | 37.120807945529236 | 8.663101098684088 | 53.808030638232374 | 52.93979547114636 | 37.29538166865713 | 0.7290069448268927 | 68.07952795247988 | 35.98111139456643 |
| SE_317_3_26 | SE_317 | SE_317_3 | TCTGTGACCCAAACACAAGA | 159.31519572817993 | 130.9204154870404 | 123.440857829318 | 149.65047630013606 | 88.02934455654076 | 157.66843999605038 | 136.67239782111022 | 215.5405958468102 | 156.28540889722987 | 103.51898616541875 | 146.72312058724114 | 116.06810127279495 |
| SE_317_4_16 | SE_317 | SE_317_4 | GCTTCGGAGTATGATCAGGT | 882.1943491342753 | 1115.322012851428 | 1126.7681502660148 | 1086.5925887879444 | 1138.0179121586534 | 1124.4705226091946 | 1068.627488475295 | 1176.019742251894 | 1357.7294897946847 | 2299.2879039840195 | 1022.3667042518962 | 1399.781301349907 |
| SE_317_4_21 | SE_317 | SE_317_4 | GCCGTTATATCCAGGAGCTT | 1301.615986867647 | 1370.1670964330717 | 1327.2361033808272 | 1233.6404481089476 | 1338.4702750645113 | 1858.2351856677367 | 1182.7005134283477 | 1283.7900401752993 | 1462.5117525780547 | 1165.6821047782014 | 1354.5478492614102 | 1926.7304811283961 |
| SE_317_4_23 | SE_317 | SE_317_4 | CTGGATATAACGGCCAACCC | 131.13699784428417 | 133.9185929409421 | 157.0167711588925 | 117.11776406097604 | 124.08955798934059 | 271.1550643888119 | 131.291594757287 | 160.71009339455145 | 66.59889583688773 | 66.33963197924723 | 104.46686185811568 | 33.657949369110536 |
| SE_317_4_7 | SE_317 | SE_317_4 | TAAACCAAGGATGATCTCAG | 172.32051782843953 | 171.89550735703014 | 159.97935174679614 | 175.67664609146405 | 137.87728665482288 | 201.85025559933922 | 194.7850709104012 | 208.92312141291688 | 161.6133205641809 | 153.09145841364744 | 286.4035313862947 | 215.88666836739858 |
| SE_317_4_8 | SE_317 | SE_317_4 | AGGGCCCCTGAGATCATCCT | 2003.9033802816648 | 2124.7084223316633 | 2068.86877721937 | 2161.473401169791 | 2171.0369675570955 | 1778.534655559843 | 1919.870533172131 | 2467.372610351643 | 2402.000176517084 | 2657.9593208388505 | 2449.689221324578 | 2603.407511548791 |
| SE_318_1_14 | SE_318 | SE_318_1 | GCTCCCCAAAGAGATCTAGC | 70.44549470973944 | 94.94228604022014 | 114.55311606560711 | 91.09159426964803 | 108.18064029839948 | 113.48662439276154 | 67.79811860417279 | 55.775855942814914 | 71.92680750383876 | 109.3510417240339 | 43.43004369382337 | 83.56903291641235 |
| SE_318_1_24 | SE_318 | SE_318_1 | CCAACTGAGGAAATCCCGCA | 427.00807562518975 | 433.73633833111097 | 420.6864434823158 | 353.95590916206095 | 379.69283555712764 | 480.80211097696684 | 330.3813081187468 | 363.9610938641312 | 410.24919835522843 | 220.8891042825485 | 311.0530156449512 | 631.4104709240045 |
| SE_318_1_6 | SE_318 | SE_318_1 | TATTAGCTCATAATTCTGCC | 343.5572588151908 | 450.7260105698872 | 469.0752597514084 | 538.7417146804897 | 561.0544972338562 | 722.5026316302528 | 456.29209981221055 | 463.2232103725307 | 508.8155641938223 | 576.644493358072 | 660.840935124934 | 810.1553468841087 |
| SE_318_1_8 | SE_318 | SE_318_1 | TGATCCCTAGATGACCCCTC | 547.307305052591 | 541.6707266715717 | 548.0774087621719 | 628.5320004605715 | 589.6905490775501 | 649.7325824013066 | 577.8982490546157 | 691.9987550871274 | 721.0440455940378 | 621.8429239373394 | 731.2680330068098 | 392.3101823020469 |
| SE_318_1_9 | SE_318 | SE_318_1 | GGAGCCAGAGGGGTCATCTA | 348.97614302363223 | 528.6786243713311 | 515.489022295232 | 517.9207788474274 | 551.5091466192915 | 386.3743090013103 | 518.7094153525601 | 543.5782570698063 | 471.5201825251652 | 584.6635697511679 | 846.2989595472068 | 629.0891088985486 |
| SE_318_2_12 | SE_318 | SE_318_2 | AGGCTGCTACAACAAAATAT | 1291.8619952924523 | 1551.0571361518068 | 1692.6210425556085 | 1414.5223281586773 | 1710.7389490325331 | 1758.6095230328697 | 1195.6144407815234 | 1909.6140509234938 | 1564.6300595279492 | 1694.2121397776984 | 1889.7937931636657 | 1898.8741368229253 |
| SE_318_2_13 | SE_318 | SE_318_2 | AATCGCATACCTACAGGGGA | 346.80858934025565 | 396.7588163996568 | 452.2873030866212 | 430.73311004647854 | 359.5415398152689 | 361.2513158151264 | 405.7125510122721 | 440.5347265991822 | 302.8029797383829 | 433.7591321720011 | 349.78791947998286 | 371.41792407294383 |
| SE_318_2_18 | SE_318 | SE_318_2 | TAGACAGTATCCCCAACCCA | 3453.9967944606087 | 3960.5924166041304 | 3842.467022511011 | 3547.3669425580074 | 3998.4413129898635 | 3825.6254451788927 | 3483.5319035191637 | 3929.834460242061 | 3779.2653424239224 | 3717.9354186171527 | 4512.029404298839 | 3690.9656204748794 |
| SE_318_2_22 | SE_318 | SE_318_2 | GGAGATAGTCAACGAGGAAA | 758.6437891818092 | 953.420430340737 | 879.8864346073788 | 830.2348163433635 | 767.8704272160904 | 799.6042314085413 | 788.8257291564865 | 939.6813696128479 | 861.3457194904147 | 887.9304587991553 | 840.4300347237172 | 708.0154177640492 |
| SE_318_2_35 | SE_318 | SE_318_2 | ATAAGAGACATCTTCCAAAG | 41.18351998415536 | 79.95139877071169 | 69.12688038441809 | 42.943180155691216 | 50.90853661101153 | 34.65240439473635 | 63.4934761531142 | 24.57919075446081 | 60.382998892111544 | 5.103048613788249 | 57.515463270198524 | 46.42724050911798 |
| SE_319_1_0 | SE_319 | SE_319_1 | ACTGGGGTGGAGGGAGAAAT | 14573.547190182555 | 17267.50334950446 | 17508.851274510467 | 15916.304135886643 | 16831.634917015686 | 17746.362600654353 | 15869.064395827492 | 16967.20444850241 | 18473.645719874832 | 18623.2114125478 | 16435.3370757004 | 18595.27050491448 |
| SE_319_1_4 | SE_319 | SE_319_1 | CTAATCATGGTGGTTCTACC | 1.083776841688299 | 1.9987849692677924 | 1.975053725269088 | 1.3013084895664004 | 0.0 | 0.0 | 0.0 | 0.0 | 4.439926389125849 | 0.0 | 0.0 | 0.0 |

| | | | | | | | | | | | | | | | |
|---|---|---|---|---|---|---|---|---|---|---|---|---|---|---|---|
| SE_319_1_5 | SE_319 | SE_319_1 | GTGGCTGGCATAACTAGCAT | 331.63571355661946 | 255.84447606627742 | 254.78193055971235 | 303.2048780689713 | 268.3304117205399 | 201.85025559933922 | 298.0964897358073 | 258.08150292183853 | 167.8292175089571 | 248.5913681859704 | 253.53755237475266 | 123.03218734916264 |
| SE_319_1_6 | SE_319 | SE_319_1 | AGGAAGGGAGGCTAAGGGAA | 2461.257207474127 | 2541.455088423998 | 2495.4803818774926 | 2428.241641530903 | 2414.9737054848592 | 1851.3047047887894 | 2655.96439230315 | 2431.4491777105077 | 2338.065236513672 | 2883.9514737351874 | 2421.5183821718274 | 1696.915640608262 |
| SE_319_1_7 | SE_319 | SE_319_1 | GGAGGAGGAAGGGAGGCTAA | 6180.779328148368 | 6218.220039392102 | 6112.791279707828 | 5927.460169974954 | 5939.329271284678 | 6031.2509849038615 | 5856.466054665211 | 5905.623255504488 | 5798.543864198358 | 5023.586856802117 | 5657.643529844018 | 5972.864491498028 |
| SE_32_1_15 | SE_32 | SE_32_1 | TCTGTGTGAACCAACCATCA | 594.9934860868761 | 527.6792318866972 | 597.4537518938992 | 559.5626505135522 | 606.6600612812207 | 403.70051119867844 | 414.3218359143893 | 576.6656292392729 | 500.82369669339573 | 349.9233335169085 | 390.87039324441037 | 559.4482481348716 |
| SE_32_1_2 | SE_32 | SE_32_1 | AGTGGTGCCTCTTTTCAAAA | 241.68223569649066 | 269.83597085115196 | 278.4825752629414 | 219.92113473672168 | 329.8448934588455 | 421.89302350591504 | 244.288459097575 | 433.9172521652889 | 171.38115862025776 | 167.67159731018532 | 414.34609253836896 | 179.90555697283216 |
| SE_32_1_20 | SE_32 | SE_32_1 | AGGGCAGTTTAGGTACTGCA | 1103.2848248386883 | 291.8226055130977 | 471.0503134766775 | 942.1473464460739 | 211.05830803315195 | 2173.5720656598373 | 1190.2336377177 | 637.1682526348687 | 1427.880326742873 | 86.02281948957334 | 191.32694924576242 | 566.4123342112393 |
| SE_32_1_4 | SE_32 | SE_32_1 | AAAATGGTATGACAGTCCCT | 54.188842084414944 | 110.93256579436247 | 94.80257881291622 | 54.6549565617882 | 74.24161589105847 | 110.02138395328791 | 26.904015319116187 | 70.90151179171387 | 140.30167389637683 | 91.85487504818848 | 49.29896851731302 | 168.29874684555267 |
| SE_32_1_6 | SE_32 | SE_32_1 | AACAGGAGCCAGAGAACCTA | 870.272803875704 | 1010.3858019648691 | 1020.115249101484 | 1044.9507171218195 | 1021.3525157584187 | 1049.9678531605114 | 835.1006355053664 | 856.4902624439036 | 1093.109877002784 | 852.9381254474644 | 1128.0073510747097 | 1583.168901360923 |
| SE_32_2_0 | SE_32 | SE_32_2 | GAAGGAACCAACAGGATACG | 2619.4886263606186 | 2792.302602067106 | 2749.2747855745706 | 2529.7437037170826 | 2606.941312288882 | 2817.2404772920654 | 2321.2784417333446 | 3079.0163187414946 | 2259.034546787232 | 2696.5966889146757 | 2875.7731635099262 | 2567.426400154224 |
| SE_32_2_20 | SE_32 | SE_32_2 | GGGGTGGTGAAGGAGTCGGA | 277.4468714722045 | 366.7770418606399 | 335.75913329574496 | 287.5891761941745 | 356.3597562770807 | 113.48662439276154 | 209.85131948910626 | 310.07594490242866 | 344.5382877961659 | 172.04563897914667 | 220.67157336321065 | 178.74487596010422 |
| SE_32_2_29 | SE_32 | SE_32_2 | CTGTTGACAGATCCTCCCCA | 300.2061851476588 | 328.80012744455183 | 364.39741231214674 | 348.7506752037953 | 355.29916176435125 | 273.7539947184172 | 357.285323437863 | 315.7480658457658 | 358.7460522413686 | 290.14476404110326 | 292.2724562097843 | 321.50864052564197 |
| SE_32_2_3 | SE_32 | SE_32_2 | GCACTTTGCAGACTATCTGT | 1186.7356416486873 | 1381.1604137640445 | 1438.8266388585307 | 1450.9589658665366 | 1445.5903208501813 | 1540.2993753460307 | 1464.6545939726852 | 1705.4176969633577 | 1215.6518453426575 | 1981.4408760394942 | 1741.8968876117267 | 1989.4072558157054 |
| SE_32_2_6 | SE_32 | SE_32_2 | GCTATCTTAACAACAGTAAA | 100.7912462770118 | 211.871206742386 | 182.69246958739063 | 173.07402911233126 | 152.72560983303458 | 189.7219140611815 | 131.291594757287 | 224.04877726181584 | 259.29170112494955 | 113.72508339299526 | 225.36671322200237 | 142.76376456553777 |
| SE_32_3_10 | SE_32 | SE_32_3 | ATTCTCCCCGACCCCCCAGA | 4.335107366753196 | 7.9951398770711695 | 7.900214901076352 | 3.9039254686992013 | 2.1211890254588135 | 0.8663101098684087 | 8.60928490211718 | 0.0 | 1.7759705556503396 | 2.187020834480678 | 0.0 | 20.89225822910309 |
| SE_32_3_26 | SE_32 | SE_32_3 | GGGGCCAGTCGGCCTCAGCA | 41.18351998415536 | 34.97873696218637 | 32.58838646693995 | 53.35364807222242 | 25.454268305505764 | 24.256683076315444 | 39.817942672291956 | 16.071009339455145 | 42.62329333560815 | 67.06863892407412 | 12.911634611677218 | 64.99813671276517 |
| SE_32_3_3 | SE_32 | SE_32_3 | CACTGCTGTGAATTCACCCA | 1212.7462858492065 | 1222.257008707255 | 1154.418902419782 | 1250.5574584733108 | 1309.8342232208174 | 1046.5026127210378 | 1293.5450565431063 | 1641.133659605537 | 1037.1668044997982 | 1245.8728687091595 | 1226.6052881093358 | 950.5977494241906 |
| SE_32_3_31 | SE_32 | SE_32_3 | ACCATCTGCCTTAAGAGGAA | 878.9430186092104 | 990.3979522721911 | 976.6640671455641 | 890.0950068634179 | 1019.2313267329599 | 1393.0266566684013 | 817.8820657011321 | 696.7255225399083 | 1106.4296561701615 | 1309.2964729090993 | 856.8630242294882 | 1159.5203317152216 |
| SE_32_3_4 | SE_32 | SE_32_3 | TCATTGGAAGTGAAGCAGGT | 1588.8168499150463 | 1744.9392781707827 | 1579.0554533526358 | 1600.6094421666726 | 1871.9493149674029 | 1717.026637759186 | 1584.108421989561 | 1734.7236551705994 | 1735.1232328703818 | 1823.2463690120585 | 1326.3770101086598 | 1739.8608380791961 |
| SE_32_4_10 | SE_32 | SE_32_4 | CCCTCAGGACTGCTGAGAAA | 166.90163361999802 | 225.86270152726053 | 224.1685978180415 | 170.47141213319847 | 164.39214947305805 | 77.96790988815678 | 131.291594757287 | 180.56251669623134 | 270.8355097366768 | 268.2745556962965 | 172.54638981059557 | 314.5445544492743 |
| SE_32_4_16 | SE_32 | SE_32_4 | CCCCAGTCATTCATTCCCTG | 57.440172609479845 | 76.95322131681 | 72.08946097232172 | 89.79028578008163 | 74.24161589105847 | 6.93048087894727 | 55.960351863761666 | 112.49706537618601 | 106.55823333902038 | 77.27473615165061 | 41.08247376442752 | 56.873369623669525 |
| SE_32_4_7 | SE_32 | SE_32_4 | ACACGGACCCCATCTTCCAC | 16119.01296643007 | 14217.357486401806 | 14160.147683316727 | 14954.637162097073 | 14207.724092523133 | 14521.090061614266 | 15764.676816389321 | 14827.869499373763 | 14679.284627727882 | 13949.54788926259 | 14389.429882231912 | 15228.134886990696 |
| SE_32_4_8 | SE_32 | SE_32_4 | CAGGACTGCTGAGAAAAGGC | 2490.519182199711 | 2543.453873393266 | 2451.041673058938 | 2671.58632907982 | 2595.2747726488583 | 2221.2191217026 | 2827.073929732729 | 2514.6402848794523 | 2347.833074569749 | 3567.030981037986 | 2032.995558856813 | 2979.4681596726464 |
| SE_32_4_9 | SE_32 | SE_32_4 | GCCTTTTCTCAGCAGTCCTG | 223.25802938778958 | 205.8748518345826 | 225.15612468067604 | 277.1787082776433 | 248.17911597868118 | 260.759343070391 | 297.0203291230427 | 213.64988886569782 | 189.14086417676117 | 284.3127084824881 | 275.8394667040133 | 475.87921521845925 |
| SE_32_5_22 | SE_32 | SE_32_5 | GTATATGGTAAGTGTGCCTC | 588.4908250367463 | 414.7478811230669 | 395.01074505381763 | 556.9600335344194 | 448.63147888453904 | 304.94115867367987 | 652.1533313353764 | 646.6217875404305 | 469.7442119695148 | 685.2665281372791 | 841.6038196884151 | 355.1683898947525 |
| SE_32_5_23 | SE_32 | SE_32_5 | GATCTGAGTCATGATGTATA | 164.73407993662144 | 148.90948021045054 | 163.9294591973343 | 140.54131687317124 | 182.42225618945795 | 145.54009845789267 | 136.67239782111022 | 212.70453537514163 | 166.94123223113192 | 50.30147919305559 | 274.6656817393154 | 249.54641773650914 |
| SE_32_5_24 | SE_32 | SE_32_5 | TCAGATCCCATTTGCTGACT | 1318.9564163346597 | 1242.244858399933 | 1291.6851363259836 | 1244.0509160254787 | 1358.62157080637 | 1599.2084628170826 | 1273.098004900578 | 1363.1997333820188 | 1240.5154331217623 | 1167.869125612682 | 1145.6141255451787 | 973.8113696787495 |
| SE_32_5_25 | SE_32 | SE_32_5 | TCAGTTCCAAGTCAGCAAAT | 917.9589849099892 | 1084.3408458277775 | 1075.4167534090184 | 1016.3219303513588 | 1093.4729426240183 | 1111.4758709611683 | 896.4417904329514 | 947.2441975372974 | 1109.9815972814622 | 1323.8766118056371 | 1126.8335661100118 | 1156.0382886770376 |
| SE_32_5_26 | SE_32 | SE_32_5 | TTCAGTTCCAAGTCAGCAAA | 10.837768416882989 | 21.986634661945715 | 11.850322351614528 | 19.519627343496005 | 22.27248476731754 3 | 32.919784174999535 | 22.599372868057596 | 2.836060471668555 | 6.215896944776189 | 53.94651391719005 | 15.259204541073078 | 1.1606810127279494 |
| SE_32_6_12 | SE_32 | SE_32_6 | GGCTCTCTGAGAGCTCTGAG | 1.083776841688299 | 4.996962423169481 | 4.93763431317272 | 0.0 | 1.0605945127294067 | 0.0 | 1.0761606127646475 | 13.23494886778659 | 0.0 | 0.0 | 0.0 | 0.0 |
| SE_32_6_13 | SE_32 | SE_32_6 | CAGTTCACACACTGACTGCA | 290.4521935724641 | 64.96051150120326 | 81.96472959866715 | 264.1656233819793 | 88.02934455654076 | 411.49730218749414 | 350.82835976127507 | 74.68292575393862 | 323.2266411283618 | 26.973256958595027 | 44.6038286585213 | 141.60308355280984 |
| SE_32_6_33 | SE_32 | SE_32_6 | GCATAGTGACTACCATTTCA | 3349.954217658532 | 3306.9897316535626 | 3468.1943415725186 | 3270.188234280364 | 3489.355946879748 | 2857.9570524558803 | 3154.226756013182 | 3440.1413521339573 | 3689.5788293635806 | 2896.3445917972444 | 3580.0441423286834 | 3038.6628913217714 |
| SE_32_6_4 | SE_32 | SE_32_6 | ATGAAGCTCTTCAGAACTAG | 337.054597765061 | 371.7740042838094 | 377.23526152639585 | 339.6415157768305 | 348.9355946879748 | 355.1871450460476 | 362.6661265016862 | 385.7042241469235 | 391.6015075208999 | 477.4995488616147 | 369.7422638798 4766 | 344.722260780201 |
| SE_32_6_9 | SE_32 | SE_32_6 | TTATAGTGAGGACTAAGTGA | 85.61837049337561 | 88.94593113241676 | 68.13935352178353 | 54.65495656178882 | 98.63528968383483 | 92.69518175591973 | 48.42722757440914 | 78.46433971616335 | 113.66211556162173 | 34.992333351690846 | 38.73490383503166 | 25.534982280014887 |
| SE_32_7_14 | SE_32 | SE_32_7 | CAACACTTCACAAAGGCCCA | 1322.2077468597247 | 1496.0905494969427 | 1465.4898641496634 | 1473.0812101891654 | 1384.075839111876 | 1235.3582166723509 | 1362.4193357600436 | 1520.1284128143454 | 1418.112488686796 | 1509.0443757916678 | 1397.9778929552335 | 1414.8701545153704 |
| SE_32_7_19 | SE_32 | SE_32_7 | CTGTTAACGATACATTTGGG | 208.0851536041534 | 269.83597085115196 | 261.6946185981542 | 214.71590077845607 | 203.6341464440461 | 399.36896064933643 | 170.0333768168143 | 273.20715877073746 | 218.44437834499178 | 162.56854869639707 | 250.0161974806589 | 366.77520002203204 |
| SE_32_7_32 | SE_32 | SE_32_7 | TCTCCTCCCGCTAGAAGGGA | 23.843090517142578 | 32.97995199291857 | 47.40128940645811 | 42.943180155691216 | 9.545350614564661 | 48.51336615263089 | 35.513300221233365 | 29.305958207241734 | 138.5257033407265 | 48.84346530340181 | 4.695139858791716 | 18.57089620364719 |
| SE_32_7_39 | SE_32 | SE_32_7 | CAGGGGGCACAGTTGAACCT | 21564.991595913772 | 18768.59086142457 | 19097.781996489448 | 21223.040156338426 | 19617.816701955835 | 18390.03101228658 | 21917.08703956481 | 22407.713786653254 | 18124.66750568954 | 19030.726294706034 | 21823.010063663896 | 19422.836066989505 |
| SE_32_7_9 | SE_32 | SE_32_7 | GGTTAGGGCCAGGGATCCCA | 398.829877741294 | 484.70535504743964 | 473.0253672019466 | 385.18731291165454 | 519.6913112374093 | 572.6309826230182 | 340.0667536336286 | 330.87372169466477 | 464.4163003025638 | 444.69423634440454 | 281.70839152750295 | 295.9736582456271 |
| SE_32_8_1 | SE_32 | SE_32_8 | ATAATACGTGTCTATGGCAG | 366.31657249064506 | 418.7454510616025 | 383.1604227022031 | 398.20039780731855 | 518.6307167246799 | 440.08553581315164 | 392.79862365909634 | 490.63846159866 | 446.6565947460604 | 521.9689724960551 | 659.6671501602361 | 427.1306126838854 |
| SE_32_8_13 | SE_32 | SE_32_8 | TGCTAGGTGGCATTCCCAGG | 574.4017260947984 | 580.6470335722937 | 572.7655803280355 | 503.60638546219695 | 561.0544972338562 | 537.1122681184135 | 546.6895912844409 | 674.0370387665599 | 626.9176061445698 | 587.5795975304754 | 642.0603756897672 | 500.2535164857462 |
| SE_32_8_14 | SE_32 | SE_32_8 | CTATATGGTCTAAAAGAAGG | 11.921545258571289 | 18.98845720804403 | 34.56344019220904 | 10.410467916531204 | 31.8178353818822 | 21.65775274671022 | 11.837766740411123 | 0.945353490556185 | 4.439926389125849 | 2.187020834480678 | 0.0 | 0.0 |
| SE_32_8_3 | SE_32 | SE_32_8 | TATGAGGAGTCTTCCTAGAG | 282.86575568064603 | 455.72297299305666 | 445.37461504817935 | 291.4931016628737 | 434.84375021905674 | 332.6630821894689 | 201.24203458698906 | 432.97189867473276 | 424.45696280043114 | 454.90033357198104 | 186.63180938697073 | 287.8488915653147 |
| SE_32_8_32 | SE_32 | SE_32_8 | GACCTGGAAGACAGGATCCT | 338.13837460674927 | 366.7770418606399 | 347.6094556473595 | 321.4231969229009 | 308.63300320425736 | 558.7700208651237 | 417.5503177526832 | 341.2726100907828 | 268.1715539032013 | 420.63700716511704 | 376.7849736680352 | 324.9906835638258 |
| SE_320_1_2 | SE_320 | SE_320_1 | GCTTGGGGCTTATTATTACA | 656.7687660631092 | 888.4599188395337 | 882.8490151952824 | 800.3047210833363 | 940.7473327909838 | 867.1764199782772 | 643.5440464332592 | 918.8835928206119 | 851.5778814343378 | 772.7473615165062 | 1036.4521238282714 | 864.7073544823223 |
| SE_320_1_26 | SE_320 | SE_320_1 | GGGACTTGGGAGGGAAACTA | 170.15296414506292 | 168.89732990312845 | 210.3432217411579 | 204.30543286192486 | 184.54344521491677 | 427.0908841651255 | 125.91079169346375 | 191.90675858290555 | 214.00445195586593 | 116.64111117230283 | 102.11929192871983 | 207.76190127830296 |
| SE_320_1_35 | SE_320 | SE_320_1 | GCTCTGGGAGGCATTGGGGA | 301.2899619893471 | 410.7503111845313 | 415.74880916914304 | 313.61534598550253 | 322.42073186973965 | 314.4705698822324 | 213.0798013274002 | 404.61129395804716 | 369.40187557527065 | 429.38509050303975 | 315.7481555037429 | 473.5578531930034 |
| SE_320_1_44 | SE_320 | SE_320_1 | ACTTGCCCAAGACGTCCTTG | 1290.778218450764 | 1332.1901820169835 | 1370.6872853367472 | 1193.2998849323892 | 1502.8624245375693 | 1227.5614256835352 | 1200.9952438453465 | 1576.8496222477165 | 1098.437788669735 | 1450.7238202055164 | 1726.6376830706536 | 1518.170764648158 |
| SE_320_1_5 | SE_320 | SE_320_1 | GCCACTTACTGAACTTGCGT | 1187.8194184903757 | 1131.3122926055705 | 1184.0447082988183 | 1134.741002901901 | 1304.5312506571704 | 1803.657648746027 | 1374.2571025004547 | 1146.7137840446524 | 959.0241000511834 | 1737.952556467312 | 1164.3946849803456 | 1621.4713747809453 |
| SE_320_2_0 | SE_320 | SE_320_2 | ATTTGTGTCTCTCTTCCCCA | 30.34575156727237 | 47.97083926242702 | 53.32645058226538 | 39.03925468699201 | 15.9089176909411 | 29.454543735525895 | 31.208657770174778 | 13.23494886778659 | 24.863587779104755 | 16.038152786191638 | 15.259204541073078 | 2.321362025455899 |
| SE_320_2_18 | SE_320 | SE_320_2 | GTGCAGTGTGATCCTTCCCA | 2559.880900067762 | 2217.6519234026155 | 2278.224472097893 | 2201.8139643463496 | 2272.854040779119 | 2485.4437052124645 | 2577.4046675713307 | 2093.0126280913937 | 1898.512523990213 | 2287.6237928667892 | 2068.209107797751 | 2263.3279748195014 |
| SE_320_2_22 | SE_320 | SE_320_2 | CTACTGCTGATGCTGTGCGA | 273.1117641054513 | 309.81167023650784 | 315.02106918041954 | 297.9996441107057 | 317.11775930609264 | 352.58821471644234 | 370.1992507910387 | 306.29453094020397 | 315.2347736279353 | 422.0950210547708 | 352.1354894093787 | 269.2779949528843 |
| SE_320_2_23 | SE_320 | SE_320_2 | GCGAGCACTCCAGAAACGGT | 9523.147107915083 | 9404.283280404963 | 9255.101756610946 | 9499.551973834723 | 9217.626910131274 | 9016.555623510398 | 9904.982279885815 | 10163.495376969546 | 8739.551104355322 | 8032.198518102703 | 10072.24878207293 | 8185.1225017575 |
| SE_320_2_31 | SE_320 | SE_320_2 | GCTGGCCTAGCTGAGTTGGG | 2296.5231275375054 | 2216.652530917982 | 2137.0081307411533 | 2383.9971528856454 | 2299.368903597354 | 2441.261889609176 | 2445.036912201279 | 2379.4547357299175 | 2266.138429009833 | 2640.463154163005 | 2413.301887418942 | 2153.063278610346 |
| SE_321_1_1 | SE_321 | SE_321_1 | GGCTGAGCTGAACATCACAG | 1920.4525634716656 | 1744.9392781707827 | 1871.363404692461 | 1850.4606721634213 | 1810.4348332290974 | 2073.9464030249706 | 1875.7479480487805 | 1818.8601158300999 | 2121.3968287243306 | 2208.891042825485 | 1767.720156835081 | 3022.4133571435805 |
| SE_321_1_2 | SE_321 | SE_321_1 | GCTGAGCTGAACATCACAGA | 5307.2551937476 | 4796.084533758068 | 4776.667434563289 | 5619.0500579477175 | 5178.8830056576935 | 5442.160110193344 | 5811.2673089290965 | 5704.2629620160205 | 5390.958621676606 | 5883.086044753024 | 5345.416729234369 | 6030.898542134425 |
| SE_321_1_5 | SE_321 | SE_321_1 | AAAGCTGCTGGCTCTGCCCT | 382.57322511596954 | 430.7381608772093 | 457.2249373997939 | 426.82918457777936 | 450.75266790999785 | 237.36897010394398 | 426.1596026548004 | 542.6329035792502 | 689.9645608701569 | 272.64859736525784 | 492.98968517313017 | 300.6163822965389 |
| SE_321_1_6 | SE_321 | SE_321_1 | TTTATGGTCTGGAGGGACAG | 744.5546902398613 | 886.4611338702659 | 804.8343930471534 | 930.4355700399764 | 820.9001528525608 | 874.9732109670928 | 864.156972050012 | 1037.998132630691 | 725.4839719831638 | 1102.9875075230887 | 870.9484438058633 | 1275.5884329880164 |
| SE_321_1_8 | SE_321 | SE_321_1 | CTCAAGGGAACACTGCAAAG | 477.9455871845398 | 436.7345157850126 | 416.73633603177757 | 489.29199207696655 | 494.23704293190355 | 647.1336520717014 | 432.6165663313883 | 764.7909738599536 | 495.49578502644476 | 256.6104445790662 | 697.2282690305699 | 500.2535164857462 |
| SE_321_2_22 | SE_321 | SE_321_2 | AAACAAGAAAATGGACTCCC | 65.02661050129794 | 81.9501837399795 | 88.87741763710896 | 87.18766880094883 | 117.72599091296415 | 51.11229648223612 | 97.93061576158291 | 56.7212094333711 | 83.47061611556596 | 161.83954175157018 | 66.90574298778195 | 125.35354937461854 |
| SE_321_2_24 | SE_321 | SE_321_2 | ATTCAACTCTCCAGCAGCCA | 44.434850509220254 | 59.96354907803377 | 62.21419234597627 | 45.545797134824014 | 60.45388722557618 | 35.51871450460476 | 29.056336544645482 | 14.180302358342775 | 63.93494000341222 | 147.2594028550323 | 8.216494752885502 | 13.928172152735392 |
| SE_321_2_25 | SE_321 | SE_321_2 | CATTCAACTCTCCAGCAGCC | 795.4922017992114 | 1024.3772967497437 | 980.6141745961022 | 909.6146342069139 | 940.7473327909838 | 1349.7111511749808 | 754.3885895480179 | 1233.6863051758214 | 1212.0999042313567 | 1017.6936949783421 | 1328.7245800380556 | 1104.9683241170078 |
| SE_321_2_26 | SE_321 | SE_321_2 | TGAATGCCAGCCACCAGCCA | 2900.186828357888 | 3319.9818339538033 | 3078.1212308318736 | 2546.660714081446 | 3004.6642545624095 | 1476.1924272157685 | 2117.8840859208262 | 1539.0354826254693 | 2342.505162902798 | 885.7434379646746 | 825.1708301826441 | 954.0797924623745 |
| SE_321_2_27 | SE_321 | SE_321_2 | GAATGCCAGCCACCAGCCAT | 3161.377047204768 | 3015.167126140465 | 2960.605534178363 | 3306.6248719882237 | 3035.421495431562 | 2766.1281808098292 | 3367.306557340582 | 3167.879546853776 | 2756.306302369327 | 2998.4055640730094 | 2995.499229909115 | 3047.9483394235954 |
| SE_321_3_19 | SE_321 | SE_321_3 | GATAATGCAAGCCAGTGCAG | 803.0786396910295 | 881.4641714470964 | 785.0838557944625 | 827.6321993642307 | 882.4146345908664 | 1240.5560773315613 | 754.3885895480179 | 813.9493553688753 | 878.217439769093 | 718.8008475993162 | 827.51840011204 | 869.3500785332342 |
| SE_321_3_23 | SE_321 | SE_321_3 | GAGAATTGCATGGATCCGGG | 223.25802938778958 | 329.79951992918575 | 345.63440192209043 | 270.6721658298113 | 442.26791180816264 | 294.54543735525897 | 315.3150595400417 | 452.8243219764126 | 240.644010290621 | 548.9422294546501 | 681.9690644894968 | 206.601220265575 |
| SE_321_3_4 | SE_321 | SE_321_3 | CCTATAAACAGTTCAGCCCA | 1558.471098347774 | 1659.9909169769016 | 1733.1096439236248 | 1755.4651524250742 | 1702.2541929306979 | 1540.2993753460307 | 1574.4229764746792 | 1452.0629614943002 | 1911.8323031575906 | 2031.74235523255 | 1478.9690555193906 | 1864.053706441087 |
| SE_321_3_5 | SE_321 | SE_321_3 | CTGGAATGTAAGTGGCACCA | 912.5401007015477 | 1000.39187711853 | 950.0008418544313 | 947.3525804043395 | 987.4134913510777 | 689.5828474552534 | 1068.627488475295 | 997.3479325367751 | 851.5778814343378 | 1140.16686170926 | 867.4270889117696 | 999.3463519587644 |
| SE_321_3_7 | SE_321 | SE_321_3 | GGTGCCACTTACATTCCAGT | 638.344559754408 | 743.5480085676188 | 734.7199858001007 | 774.2785512920083 | 689.3864332741144 | 793.5400606394624 | 768.3786775139583 | 706.1790574454702 | 895.9771453255963 | 828.15188932335 | 453.0809963734006 | 652.3027291531076 |
| SE_322_1_11 | SE_322 | SE_322_1 | TTAGGACATCAGATGGATGG | 1119.5414774640128 | 1059.35603371193 | 951.9758955797005 | 1169.876332120194 | 1021.3525157584187 | 1345.3796006256387 | 1262.3363987729315 | 929.2824812167298 | 808.9545880987297 | 1029.3578060955724 | 916.7260574290826 | 1388.1744912226275 |
| SE_322_1_2 | SE_322 | SE_322_1 | TCATCTCCTGAAATAGTGAC | 381.48944827428124 | 290.8232130284638 | 340.6967676089177 | 291.4931016628737 | 309.69359771698674 | 210.51335669802333 | 366.9707689527448 | 534.1247221642445 | 358.7460522413686 | 363.77446546861944 | 568.1119229137977 | 84.7297139291403 |
| SE_322_1_20 | SE_322 | SE_322_1 | TTAGGTCTTGACACTTGGGA | 262.27399568856833 | 245.85055121993847 | 234.04386644438694 | 242.04337905935049 | 166.51333849851687 | 289.3475766960485 | 326.0766656676882 | 164.4915073567762 | 256.62774529147407 | 305.453909882468 | 301.66273592736775 | 247.22505571105322 |
| SE_322_1_31 | SE_322 | SE_322_1 | CAGCGGCCTAGGTTAGAGGA | 803.0786396910295 | 737.5516536598154 | 738.670093250639 | 879.6845389468867 | 810.2942077252668 | 891.4331030545926 | 964.2399090371241 | 771.408448293847 | 853.3538519899881 | 1161.30806310924 | 777.0456466300291 | 1228.0005114661706 |
| SE_322_1_7 | SE_322 | SE_322_1 | AGGTGGCCAAAGCCCATCAA | 358.73013459882696 | 421.7436285155042 | 415.74880916914304 | 337.0388987976977 | 348.9355946879748 | 233.90372966447035 | 352.9806809868044 | 328.9830147135524 | 465.304285580389 | 424.2820418892515 | 316.9219404684408 | 286.68821014380353 |
| SE_322_2_0 | SE_322 | SE_322_2 | ACATCTCTGTCTGACTTCAA | 434.59451351700784 | 496.6980648630464 | 462.1625717129666 | 469.77236473347057 | 469.84336913912716 | 337.86094284867943 | 475.6629908419742 | 464.16856386308683 | 410.24919835522843 | 750.1481462268725 | 417.8674474324627 | 422.4878886329736 |
| SE_322_2_11 | SE_322 | SE_322_2 | GCAGTTTCTCAGCCAGAAGT | 576.569279778175 | 555.6622214564463 | 573.7531071906701 | 572.5757354092162 | 622.5689789721617 | 662.7272340493327 | 586.5075339567329 | 625.8240107481945 | 557.6547544742066 | 586.1215836408217 | 327.4860051507222 | 356.32907090748046 |
| SE_322_2_3 | SE_322 | SE_322_2 | GCTCATTCCTGACACAATGA | 503.95623138505897 | 517.6853070403582 | 540.1771938610956 | 575.178352388349 | 502.7217990337388 | 421.89302350591504 | 534.8518245440298 | 419.73694980694614 | 453.76047696866175 | 376.1675835306766 | 349.78791947998286 | 537.3953088930406 |
| SE_322_2_5 | SE_322 | SE_322_2 | TGTCAGGAATGAGCTCTGCA | 348.97614302363223 | 58.96415659339988 | 161.95440547206522 | 395.5977808281857 | 104.99885676021127 | 679.1871261368325 | 549.9180731227349 | 179.61716320567515 | 623.3656650332692 | 36.45034724134463 | 117.3784964697929 | 247.22505571105322 |
| SE_322_2_7 | SE_322 | SE_322_2 | GGAAAGCCAGAGTTTGCCCT | 771.6491112820688 | 904.4501985936761 | 924.3251434259332 | 822.4269654059651 | 1023.4737047838776 | 1095.8822889835371 | 724.2560923906077 | 773.2991552749594 | 958.1361147733581 | 914.9037157577503 | 717.1826134304347 | 1091.0401519642726 |
| SE_322_3_1 | SE_322 | SE_322_3 | AATAAATCTGAGTAAAGAGA | 1.083776841688299 | 1.9987849692677924 | 3.950107450538176 | 0.0 | 0.0 | 0.0 | 0.0 | 0.0 | 0.0 | 0.0 | 1.173784964697929 | 0.0 |
| SE_322_3_22 | SE_322 | SE_322_3 | TTGTATCTTCTGTAGAGCCA | 61.77527997623304 | 38.97630690072195 | 28.63827901640178 | 50.751031093089615 | 51.96913112374093 | 0.8663101098684087 | 80.71204595734856 | 25.524544245016994 | 94.126439449468 | 11.664111117230282 | 11.73784964697929 | 23.21362025455899 |
| SE_322_3_32 | SE_322 | SE_322_3 | AGTGGTTCCAGCTACCAGGG | 15.172875783636185 | 12.992102300240651 | 11.850322351614528 | 20.820935833062407 | 14.848323178211695 | 16.459892087499767 | 7.533124289352532 | 26.46989773557318 | 23.975602501279585 | 7.290069448268927 | 2.347569929395858 | 10.446129114551544 |

| | | | | | | | | | | | | | | | |
|---|---|---|---|---|---|---|---|---|---|---|---|---|---|---|---|
| SE_322_3_4 | SE_322 | SE_322_3 | GTAACAATGAGAATTTTACT | 73.69682523480432 | 141.91373281801324 | 119.49075037877982 | 96.29682822791364 | 116.66539640023474 | 133.41175691973496 | 47.35106696164449 | 138.021609621203 | 28.415528890405433 | 56.13353475167074 | 61.03681816429231 | 420.1665266075177 |
| SE_322_3_5 | SE_322 | SE_322_3 | CTGTATGTGGAACAGAAACA | 411.83519984155356 | 455.72297299520666 | 420.6864434823158 | 507.5103109308962 | 358.48094530253945 | 570.0320522934129 | 601.5737825354379 | 253.35473546905757 | 478.6240647745665 | 429.38509050303975 | 352.1354894093787 | 627.9284278858206 |
| SE_323_1_20 | SE_323 | SE_323_1 | CGTGCAGGCAGTCCTCACAA | 82150.28459997306 | 75387.1732933887 | 74493.1013559742 | 81582.93313638635 | 80266.85331787424 | 76222.29501677194 | 81030.5894987269 | 78620.32304210513 | 75405.93382235777 | 72626.58787143436 | 75942.71343099131 | 73476.91151074284 |
| SE_323_1_25 | SE_323 | SE_323_1 | GCAGGGCCGTGTGAACGGGG | 4858.571581288644 | 5428.699976531324 | 5297.094091171694 | 5086.814885715059 | 5506.60671009108 | 5225.5825827262415 | 5090.239698376782 | 4627.505336272526 | 5737.272880028422 | 4474.644627347467 | 4294.879185829723 | 3810.515764785858 |
| SE_323_1_42 | SE_323 | SE_323_1 | AAAGCTTTTCAGAAAGGCCA | 42.26729682584366 | 38.97630690072195 | 48.38881626909266 | 67.66804145745283 | 40.30259148371746 | 89.2299413164461 | 64.56963676587885 | 4.726767452780925 | 59.49501361428638 | 44.46942363444045 | 10.564064682281362 | 67.31949873822107 |
| SE_323_1_46 | SE_323 | SE_323_1 | GGAGGGGGACGCCAGAGTCA | 5542.43476839396 | 5953.381030964119 | 5794.807629939504 | 5928.76147846452 | 5976.450079230207 | 5688.192181395972 | 5653.071698852693 | 5786.508715694408 | 5922.861803093882 | 6368.604670007734 | 5605.996991397309 | 6109.824850999926 |
| SE_323_1_50 | SE_323 | SE_323_1 | TGAAATCTACCAGCAACAAT | 419.42163773337165 | 526.6798394020633 | 530.3019252347501 | 452.8553543691074 | 417.87423801538625 | 327.4652128249665 | 517.6332547397955 | 412.17412188249665 | 533.679151972027 | 433.758213217200011 | 409.65095267957724 | 287.84889115653147 |
| SE_323_2_1 | SE_323 | SE_323_2 | TCATATCAGCCCCGGCCGAG | 1084.8606185299873 | 1403.1470484259903 | 1524.741475907736 | 1215.422129255018 | 1481.6505342829812 | 767.5507573434102 | 1051.4089186710605 | 1210.997821402473 | 1361.2814309059852 | 901.0525838060394 | 1116.2695014277306 | 557.1268861094158 |
| SE_323_2_19 | SE_323 | SE_323_2 | TCATTCCAAAGGGCCACCAG | 1015.4989006619361 | 1103.3293030358213 | 981.6017014587368 | 931.7368785295428 | 1105.1394822640418 | 756.2887259151208 | 967.4683908754181 | 874.4519787644712 | 860.4577342125896 | 591.2246322546099 | 740.6583127243932 | 677.8377114331224 |
| SE_323_2_2 | SE_323 | SE_323_2 | ACGTGCTTAACCACGTGACA | 26694.50738762449 | 22403.38132803805 | 21970.497639893336 | 24687.12335556418 | 22004.154355597002 | 22511.0682049306 | 26002.192725619414 | 22303.724902692073 | 22600.113305928397 | 22013.82271293768 | 20605.795055272145 | 21975.173613978266 |
| SE_323_2_3 | SE_323 | SE_323_2 | ATGCAGACTGAGACTCTGAG | 1053.4310901210265 | 990.3979522721911 | 972.7139596950259 | 998.1036114974291 | 1024.5342992966068 | 958.13898151446 | 1007.28633354771 | 1129.697421214641 | 1036.2788192219732 | 579.5605211373796 | 746.5272375478829 | 473.5578531930034 |
| SE_323_2_30 | SE_323 | SE_323_2 | GTTCCAGTCATTCTGATTAT | 34.680858934025565 | 39.975699385355846 | 44.43870881855448 | 50.751031093089615 | 26.51486281823517 | 11.262031428289314 | 50.57954879993843 | 34.97807915057884 | 21.311646667804077 | 61.965590310285876 | 16.432989505771005 | 4.642724050911798 |
| SE_323_3_12 | SE_323 | SE_323_3 | GCTGACTCCCCAAGGTCACA | 592.8259324034995 | 558.6603989103048 | 496.72601190517565 | 605.1084476483762 | 592.8723326157384 | 890.5667929447242 | 504.7193273866197 | 389.4856381091482 | 579.8543864198359 | 352.11035435138916 | 444.8645016205151 | 815.9587519477484 |
| SE_323_3_24 | SE_323 | SE_323_3 | ACTCCACAGTTGCCTCCAGA | 161.48274941155654 | 281.8286806667587 | 287.3703170266523 | 210.81197530975686 | 275.75457330964576 | 155.93581977631356 | 140.9770402721688 | 223.10342377125966 | 221.99631945629244 | 185.89677093085763 | 176.06774470468935 | 210.08326330375886 |
| SE_323_3_3 | SE_323 | SE_323_3 | TATTGCTGCTACAAAAAAAT | 288.2846398890875 | 419.7448435462364 | 388.0980570153758 | 356.55852614119374 | 397.7229422735275 | 563.1015714144656 | 318.54354137833565 | 516.1630058436771 | 277.051406681453 | 411.88892382719433 | 592.7614071724541 | 307.5804683729066 |
| SE_323_3_31 | SE_323 | SE_323_3 | AGTAGGCCAGGCAGCAAGAA | 1076.1904037964807 | 993.3961297260928 | 1035.9156789036367 | 992.8983775391636 | 991.6558694019953 | 1000.5881768980121 | 1185.9289952666416 | 1088.101867630169 | 990.1035847750643 | 853.6671323922913 | 1443.7555065784527 | 764.8887873877187 |
| SE_323_3_9 | SE_323 | SE_323_3 | GAGAGCTTCATCCCTAGCCC | 1213.8300626908947 | 1158.2958896906857 | 1101.0924518375166 | 1337.7451272742596 | 1038.3220279620891 | 1354.9090118341912 | 1330.1345173771042 | 1288.5168076280802 | 1179.2444489518255 | 1374.9070979435196 | 1062.2753930516258 | 1427.6376456553778 |
| SE_324_1_23 | SE_324 | SE_324_1 | GGAAATGCCTTTAACCGGCG | 56.35639576779154 | 150.9082651797183 | 118.50322351614528 | 67.66804145745283 | 88.02934455654076 | 110.88769406315632 | 34.43713960846872 | 103.98888396118035 | 150.0695119524537 | 58.320555586151414 | 124.42120625798047 | 132.31763545098624 |
| SE_324_1_27 | SE_324 | SE_324_1 | CCCGGCCCCACGCCGGTTAA | 3.2513305250648967 | 12.992102300240651 | 11.850322351614528 | 2.602616979132801 | 14.848323178211695 | 4.3315505493420444 | 1.0761606127646475 | 0.0 | 24.863587779104755 | 1.4580138896537853 | 1.173784964697929 | 0.0 |
| SE_324_1_28 | SE_324 | SE_324_1 | GCAGCTGCCCGGCCCCACGC | 169.06918730337463 | 229.86027146579613 | 187.63010390056337 | 144.44524234187045 | 215.30068608406958 | 185.39036351183947 | 151.73864639981528 | 207.0324144318045 | 144.74160028550267 | 125.38919451022554 | 262.9278320923361 | 271.59935697834015 |
| SE_324_1_30 | SE_324 | SE_324_1 | TGGGGCCGGGCAGCTGCAGG | 26.010644200519174 | 8.994532361705065 | 26.66322529113269 | 6.506542447832002 | 7.4241615891058474 | 56.31015714144657 | 7.533124289352532 | 17.961716320567515 | 76.3667338299646 | 2.9160277793075706 | 10.564064682281362 | 20.89225822910309 |
| SE_324_1_31 | SE_324 | SE_324_1 | ATCTGCCTCCTGCAGCTGCC | 18.42420630870108 | 15.990279754142339 | 15.800429802152705 | 16.917010364363207 | 3.1817835381882205 | 51.978606592104526 | 32.28481838293943 | 26.46989777557318 | 75.47874861513944 | 8.019076393095819 | 30.518409082146157 | 20.89225822910309 |
| SE_324_2_2 | SE_324 | SE_324_2 | GGTTTTCCCCAAAGACAGAG | 0.0 | 4.996962423169481 | 0.987526862634544 | 1.3013084895664004 | 0.0 | 45.048125713157255 | 2.152321225529295 | 0.0 | 0.0 | 0.0 | 0.0 | 52.23064557275772 |
| SE_324_2_20 | SE_324 | SE_324_2 | GTGATTCCACCAATTCCCAG | 69.36171786805113 | 42.973876839257535 | 53.32645058226538 | 65.06542447832003 | 51.96913112374093 | 151.60426922697152 | 62.417315540349556 | 40.650200093915956 | 10.655823333902038 | 108.622034779207 | 28.170839152750297 | 16.249534178191293 |
| SE_324_2_4 | SE_324 | SE_324_2 | ATGGTTTTCCCCAAAGACAG | 144.14231994454374 | 155.9052276028878 | 184.66752331265974 | 156.15701874796804 | 169.69512203670507 | 178.4598826328922 | 132.36775537005164 | 175.8357492434504 | 158.94936473070538 | 118.09912506195661 | 230.0618530807941 | 126.51423038734649 |
| SE_324_2_44 | SE_324 | SE_324_2 | GGAATGGCCAAGCAGGAGCG | 1171.5627658650512 | 1298.2108375394312 | 1474.3776059133743 | 1212.8195122758852 | 1426.4996196210152 | 1367.037353372349 | 1305.3828232835174 | 1523.9098267765703 | 1217.4278158983077 | 1248.7888964884671 | 1647.9940904358923 | 1546.0271089536286 |
| SE_324_2_9 | SE_324 | SE_324_2 | CAGGCCCTGAGGCTCCCTGG | 992.7395869864818 | 1183.280701806533 | 1114.9178279144003 | 877.0819219677539 | 1123.1695889804416 | 518.9197558111769 | 663.9910980757875 | 673.0916852760038 | 1200.5560956196296 | 486.9766391443643 | 507.07510474950533 | 419.0058455947897 |
| SE_324_3_14 | SE_324 | SE_324_3 | CTGTGGTCTGAATACTGCTG | 1053.4310901210265 | 1045.3645389270555 | 1114.9178279144003 | 1041.0467916531204 | 1057.4127291912184 | 1432.876921722348 | 1027.7333851902383 | 1109.8449979129612 | 1178.3564636740002 | 1409.1704243503834 | 926.116337146666 | 855.4219063804987 |
| SE_324_3_16 | SE_324 | SE_324_3 | ATCGGTCCAGGACTTACTGA | 1339.5481763267373 | 1455.1154576269528 | 1433.889004545358 | 1309.116340503799 | 1459.3780495156636 | 1549.8287865545833 | 1355.9623720834559 | 1728.1061807367062 | 1423.4404003537472 | 1386.5712090607499 | 2583.5007073001416 | 1158.3596507024936 |
| SE_324_3_21 | SE_324 | SE_324_3 | GTATATTGAGGAAAGCACTT | 68.27794102636283 | 90.94471610168455 | 87.88989077447442 | 75.47589239485123 | 103.93826224748186 | 10.395721318420904 | 87.16900963393644 | 41.59555358447214 | 130.53383584029996 | 167.67159731018532 | 107.98821675220947 | 60.35541266185337 |
| SE_324_3_24 | SE_324 | SE_324_3 | TTAAAGATAACGACCAGGCA | 85.61837049337561 | 75.95382883217611 | 69.12688038441809 | 66.36673296788642 | 82.72637199289373 | 226.97324875532508 | 65.6457973786435 | 140.85767009287156 | 107.44621861684554 | 93.31288893784226 | 171.37260484589763 | 207.76190127830296 |
| SE_324_3_25 | SE_324 | SE_324_3 | AAGATAACGACCAGGCAGCG | 896.2834480762232 | 991.397344756825 | 949.0133149917968 | 1026.73239826789 | 1002.2618145292894 | 986.7272151401176 | 855.5476871478947 | 1030.4353047062416 | 1067.358303945854 | 757.4382156751415 | 1207.824728674169 | 779.977640553182 |
| SE_325_1_0 | SE_325 | SE_325_1 | GCTTATTTCATTTTTTGAGA | 28.17819788389577 | 43.97326932389143 | 28.63827901640178 | 46.847105624390416 | 46.6661585600939 | 37.251334724341575 | 44.12258512335055 | 49.15838150892162 | 24.863587779104755 | 44.46942363444045 | 104.46686185811568 | 61.51609367458132 |
| SE_325_1_1 | SE_325 | SE_325_1 | TTATTAGACATGAGAGCAGA | 19.50798315038938 | 30.98116702365078 | 39.50107450538176 | 19.519627343496005 | 50.90853661101153 | 21.65775274671022 | 15.066248578705064 | 28.36060471668555 | 63.93494000341222 | 5.103048613788249 | 3.521354894093787 | 29.017025318198737 |
| SE_325_1_2 | SE_325 | SE_325_1 | ATTACTTTACGGCAACAGGT | 69.36171786805113 | 39.975699385355846 | 53.32645058226538 | 79.37981786355043 | 40.30259148371746 | 188.8556039513131 | 94.70213392328898 | 51.04908849003399 | 127.86988000682445 | 123.93118062057175 | 15.259204541073078 | 24.37430126728694 |
| SE_325_1_3 | SE_325 | SE_325_1 | GAAGAGTCAAGGATACCCTG | 1137.9656837727139 | 1207.2661214377465 | 1076.404280271653 | 1121.7279180062371 | 1076.5034304203477 | 1619.9999054539244 | 1148.2633738198788 | 924.5557137639489 | 1182.7963900631262 | 898.8655629715587 | 865.0795189823737 | 1330.14044058623 |
| SE_325_1_7 | SE_325 | SE_325_1 | GCTTTATTCTATAGGCAACA | 32.51330525064897 | 62.96172653193546 | 67.151826659149 | 52.05233958265602 | 67.87804881468203 | 72.77004922894633 | 31.208657770174778 | 6.617474433893295 | 64.82292528123739 | 66.33963197924723 | 3.521354894093787 | 3.482043038183848 |
| SE_325_2_1 | SE_325 | SE_325_2 | GTACTTCCTAGCAGACTGAA | 246.01734306324386 | 248.84872867384016 | 211.33074860379241 | 173.07402911233126 | 244.99733244049295 | 158.5347501059188 | 172.1856980423436 | 250.518674997389 | 318.78671473923595 | 160.3815278619164 | 375.6111887033373 | 178.74487596010422 |
| SE_325_2_15 | SE_325 | SE_325_2 | AGTGCAGGTGCCTTTCCATA | 1172.6465427067394 | 1379.1616287947768 | 1248.2339543700637 | 1305.2124150350996 | 1186.805259744206 | 1134.8662439276154 | 1062.170524798707 | 1304.5878169675352 | 1139.2851114496927 | 1469.6780007710156 | 847.4727445119048 | 1169.966460829773 |
| SE_325_2_19 | SE_325 | SE_325_2 | TTGTCGATGTCTTCTCACAG | 357.6463577571386 | 341.7922297447925 | 404.88601368016305 | 324.0258139020337 | 411.5106709390098 | 320.5347406513112 | 267.9639925783972 | 336.5458426380019 | 445.76860946823524 | 273.37760431008473 | 543.4624386551411 | 241.4216506474135 |
| SE_325_2_2 | SE_325 | SE_325_2 | AGTGACTTCTCATGTCACAC | 136.55588205272565 | 205.8748518345826 | 178.74236213685248 | 200.40150739322567 | 161.21036593486983 | 200.1176353796024 | 165.7287343657557 | 152.2019119795458 | 258.40371584712443 | 384.9156668685993 | 106.81443178751154 | 298.295020271083 |
| SE_325_2_4 | SE_325 | SE_325_2 | ACTGAAAGGTCATCTCCTAA | 602.5799239786942 | 598.6360982957038 | 601.4038593444374 | 533.5364807222242 | 544.0849850301856 | 702.5774991032795 | 553.1465549610288 | 586.1191641448347 | 838.2581022669602 | 435.94615300648184 | 835.7348948649254 | 591.9473164912542 |
| SE_326_1_10 | SE_326 | SE_326_1 | TTAGCTAGATTCTTTGAACT | 113.79656837727138 | 113.93074324826416 | 138.25376076883617 | 105.40598765487843 | 145.30144829392872 | 53.71122681184134 | 116.22534617858193 | 126.67736773452879 | 51.50314611385985 | 69.98466670338169 | 144.37555065784527 | 83.56903291641235 |
| SE_326_1_34 | SE_326 | SE_326_1 | CTATGACCTTGCTTCACATT | 211.3364841292183 | 306.81349278260615 | 327.8589183946686 | 236.83814510108488 | 314.9965702806338 | 278.9518553776276 | 279.80175931880837 | 315.7480658457658 | 340.9863466848652 | 139.2403264619365 | 448.3858565146089 | 237.93960760922963 |
| SE_326_1_35 | SE_326 | SE_326_1 | CCTATGACCTTGCTTCACAT | 876.7754649258338 | 735.5528686905476 | 707.0692336463335 | 878.3832304573203 | 831.5060979798549 | 910.4919254716975 | 759.7693926118411 | 1000.1839930084437 | 667.7649289245277 | 809.9267157026777 | 995.3696500638438 | 651.1420481403796 |
| SE_326_1_5 | SE_326 | SE_326_1 | GGGAGGGCCAGCGGCTACAA | 1033.9231069706373 | 1413.1409732723291 | 1397.3505106278799 | 1379.3869989403845 | 1433.9237812101578 | 1495.2512496328734 | 1214.985331811287 | 1492.713161588216 | 1490.0392961906348 | 1673.0709383777187 | 1743.0706725764246 | 1332.461802611686 |
| SE_326_1_6 | SE_326 | SE_326_1 | GGAGGGCCAGCGGCTACAAG | 1884.6879276959519 | 2137.700524631904 | 2082.6941532962533 | 1929.8404900269718 | 2251.6421505245307 | 2163.1763443414166 | 1915.5658907210725 | 2445.6294800688506 | 2210.1953565068475 | 2392.6007929218617 | 2321.7466601725037 | 2223.864820386751 |
| SE_326_2_17 | SE_326 | SE_326_2 | TTCCCTGATCTAGACAGCAA | 0.0 | 3.9975699385355847 | 2.962580587903632 | 5.205233958265602 | 15.9089176909411 | 2.598930329605226 | 30.13249715741013 | 18.90706981111237 | 1.7759705556503396 | 109.3510417240339 | 22.30191432926065 | 0.0 |
| SE_326_2_19 | SE_326 | SE_326_2 | AAACAACTCTAGCTGAAGGG | 34.680858934025565 | 64.96051150120326 | 64.18924607124536 | 26.02616979132801 | 73.18102137832906 | 136.01068724934018 | 5.380803063823238 | 65.22939084837677 | 112.77413028379657 | 39.36637502065221 | 50.47275348201095 | 163.65602279464088 |
| SE_326_2_23 | SE_326 | SE_326_2 | TGGCAATTCACAGCCACAAA | 33.59708209233727 | 44.97266180852533 | 47.40128940645811 | 33.834020278726415 | 32.87842989461161 | 59.7753975809202 | 20.44705516425283 | 29.305958207241734 | 44.39926389125849 | 53.21750697236316 | 23.47569929395858 | 26.695663292742836 |
| SE_326_2_6 | SE_326 | SE_326_2 | CATGCTACCCAGGAAAGGAA | 2272.6800370203628 | 2214.653745948714 | 2182.4343664223425 | 2184.8969539819864 | 2433.0038122012593 | 2334.7057460953615 | 2231.9571108738787 | 2525.03917327557 | 2422.423837907063 | 1900.521105163709 | 2187.93517419694 | 2193.6871140558246 |
| SE_326_2_9 | SE_326 | SE_326_2 | GGCAGCCGTGGACACCCTAG | 1689.608096192058 | 1441.1239628433705 | 1425.9887896442815 | 1604.5133653537717 | 1419.0754580319463 | 1495.2512496328734 | 1524.9195882875056 | 1681.7838596994532 | 1610.805293974858 | 1216.7125909160839 | 1525.9204541073077 | 1923.2484380902122 |
| SE_327_1_1 | SE_327 | SE_327_1 | GGAGAGAAAAGGTCATCCCA | 2410.3196959147767 | 2709.3530258424926 | 2746.3122049866067 | 2510.2240763735863 | 2713.0007635618226 | 3193.2190649749546 | 2269.6227323206417 | 2635.645531670644 | 2203.091474284246 | 2437.799223501129 | 2662.144299934903 | 2030.0310912611835 |
| SE_327_1_11 | SE_327 | SE_327_1 | GGCAAGAACAGAATATCGGA | 585.2394945116814 | 758.5388958371273 | 730.7698783495625 | 576.4796608779154 | 834.6878815180431 | 719.0373911907792 | 599.4214613099086 | 691.9987550871274 | 601.1660330876399 | 650.2741947855883 | 469.5139858791716 | 674.3556683949387 |
| SE_327_1_19 | SE_327 | SE_327_1 | GGCTGTGAGCTAGAGAAGAG | 287.2008630473992 | 410.7503111845313 | 417.72386289441215 | 360.46245166098929 | 397.7229422735275 | 314.4706988222324 | 390.64630243356703 | 319.5294798079905 | 395.15344863220054 | 212.87002788945974 | 217.15021846911688 | 196.15509115102344 |
| SE_327_1_21 | SE_327 | SE_327_1 | AGCTCACCTCTAACCTTACA | 482.28069455129304 | 512.6883446171887 | 509.5638611194247 | 605.1084476483762 | 593.9329271284678 | 661.8609239394642 | 584.3552127312036 | 544.5236105603626 | 523.023328639025 | 583.934562806341 | 478.90426559675507 | 356.32907090748046 |
| SE_327_1_6 | SE_327 | SE_327_1 | TGAGAAGGCAAAGGAAAACA | 8313.652152590941 | 8841.62531155608 | 8662.58563903022 | 8415.562002025912 | 8382.939028613231 | 8028.095788150544 | 8211.10547539426 | 7655.472566523986 | 8593.033533514168 | 8476.892754447108 | 7214.082393033472 | 7589.693142228061 |
| SE_327_2_17 | SE_327 | SE_327_2 | GCAAAAGTAGGGAAAGAGAC | 1094.614610105182 | 1146.303179875079 | 1223.5457828042001 | 1043.6494086322532 | 1152.8662353368652 | 1086.3528777749846 | 881.3755418542463 | 814.8947088594315 | 953.6961883842323 | 964.476188005979 | 739.4845277596953 | 876.3141646096018 |
| SE_327_2_24 | SE_327 | SE_327_2 | CCTTGGCATGAACCAATCAC | 3716.270790149177 | 3382.9435604857385 | 3375.3668164848714 | 3591.611431203265 | 3317.5396358175844 | 3377.743118376926 | 3964.575697424961 | 3154.6445979859895 | 3682.474947140979 | 3537.1416963000834 | 3304.20467562467 | 2962.0579444817267 |
| SE_327_2_3 | SE_327 | SE_327_2 | ATAAGCAGACAATTTCAGGG | 80.19946826493412 | 108.93378082509469 | 115.54064292824165 | 100.20075369661284 | 123.02896347661118 | 115.21924461249836 | 78.55972471825459 | 95.47800254617468 | 217.5563930761666 | 168.4006042550122 | 136.15905590495976 | 210.08326330375886 |
| SE_327_2_5 | SE_327 | SE_327_2 | CTGATTTCCCAGCTACTGAG | 40.09974314246706 | 52.9678016855965 | 67.151826659149 | 48.14841411395682 | 46.6661585600939 | 25.989303296052263 | 47.35106696164449 | 44.431614056140695 | 31.07948472388094 | 50.30147919305559 | 84.5125174582509 | 37.14179240729438 |
| SE_327_2_7 | SE_327 | SE_327_2 | ATACCAAACTTCTTGCACGC | 488.7833556014228 | 570.6531087259547 | 559.9277311137864 | 573.8770438987826 | 456.0556404736449 | 348.2566641671003 | 425.08344204203576 | 496.3105825419971 | 412.91315418870397 | 614.5528544900705 | 569.2857078784956 | 287.84889115653147 |
| SE_327_3_0 | SE_327 | SE_327_3 | TGATTTTTCTTATAAACACA | 3.2513305250648967 | 0.9993924846338962 | 0.0 | 2.602616979132801 | 3.1817835381882205 | 0.8663101098684087 | 2.152321225529295 | 6.617474433893295 | 0.0 | 0.7290069448268927 | 29.344624117448227 | 2.321362025455899 |
| SE_327_3_15 | SE_327 | SE_327_3 | TACTCATGTGAAAAATTACA | 157.14764204480335 | 195.88092698824366 | 238.98150075755964 | 200.40150739322567 | 234.3913873131989 | 170.66309164467653 | 118.37766740411122 | 165.43686084733238 | 306.3549208496836 | 105.70600699899943 | 251.1899824453568 | 175.26283292192036 |
| SE_327_3_21 | SE_327 | SE_327_3 | GAGTGTGCAGGAACAGGCAG | 4046.822726864108 | 3861.652560625375 | 4011.3341160215177 | 3859.6809800539436 | 3790.5647884949 | 4377.464985165069 | 3776.247590191148 | 3431.6331707189515 | 3402.7595846260506 | 2905.821682079994 | 3169.2194046844083 | 3443.740564763826 |
| SE_327_3_22 | SE_327 | SE_327_3 | AGTGTGCAGGAACAGGCAGA | 3348.8704408168437 | 3095.1185249111763 | 3246.9883243423806 | 3190.808416416814 | 3392.841846221372 | 2693.358131580883 | 3245.700408098177 | 2852.1314810080103 | 3379.6719674025962 | 2149.8414802945063 | 2696.184063911143 | 2452.5189798941574 |
| SE_327_3_3 | SE_327 | SE_327_3 | GCTGATAAAGATTATCCATA | 275.27931778827984 | 382.76732161478425 | 350.5720362352631 | 296.6983556121193 | 362.72332335534571 | 314.4706839272366 | 222.76524684228204 | 247.68261452572048 | 254.85177473582374 | 276.2936320893923 | 210.1075086809293 | 372.57860508567177 |
| SE_328_1_17 | SE_328 | SE_328_1 | GCCAGTGTCCGGGAGGCAGA | 948.3047364772616 | 1023.3779042651097 | 1100.1049249748821 | 988.9944520704644 | 1121.0483999549829 | 784.0106494309099 | 1033.1141882540617 | 1124.9706537618601 | 919.0647625490507 | 1366.8880215504237 | 1008.2812846755211 | 807.8339848586528 |
| SE_328_1_18 | SE_328 | SE_328_1 | CAGAGCAGCCAGTGTCCGGG | 1382.8992499942694 | 1870.8627312346537 | 1950.3655537032246 | 1821.8318853929607 | 2233.6120438081307 | 2554.7485140019376 | 1546.4428005427985 | 2348.2580705415635 | 1918.0482001023668 | 2259.19252018540314 | 2725.528688028591 | 2854.114610298028 |
| SE_328_1_22 | SE_328 | SE_328_1 | GGCTGCTCTGGATGATGACG | 570.0666187280452 | 547.6670815793751 | 544.1273013116338 | 523.126012805693 | 592.8723326157384 | 507.6577243828875 | 669.3719011396107 | 784.6433971616335 | 661.5490319797515 | 742.1290698337767 | 727.746678112716 | 646.4993240894678 |
| SE_328_1_24 | SE_328 | SE_328_1 | GATGACGAGGAGAGAAGGCC | 1942.1281003054316 | 1738.9429232629793 | 1736.0722245115285 | 1838.748895757324 | 1915.4336899893085 | 1659.850170507871 | 2123.2648889846496 | 1951.2096045079659 | 1529.1106484144923 | 1573.9259938812613 | 2139.8099906443244 | 1703.8797266846298 |
| SE_328_1_27 | SE_328 | SE_328_1 | GCACTCAGAGAAGCTGCCCT | 563.5639576779155 | 562.6579688488836 | 561.9021748390556 | 547.8508741074546 | 635.2961131249147 | 514.5882052618348 | 444.4543330717994 | 649.4578480120991 | 691.7405314258073 | 415.5339855315288 | 573.9808477372873 | 617.4822987712691 |
| SE_328_2_10 | SE_328 | SE_328_2 | TTCACAGGTTGAAGACACCG | 78.03193260155751 | 86.94714616314897 | 70.11440724705263 | 98.89944520704643 | 79.5445884547055 | 74.50266944868315 | 64.56963676587885 | 34.97807915057884 | 87.9105425046918 | 61.23658336545898 | 112.68335661100119 | 175.26283292192036 |
| SE_328_2_13 | SE_328 | SE_328_2 | AGACACCGAGGCTCAGAGAG | 836.6757217833667 | 894.4562737447337 | 920.375035975395 | 757.361540927645 | 879.2328510526781 | 619.4117285559122 | 705.9613619736087 | 963.3152068767525 | 946.592306161631 | 807.0106879233701 | 1070.4918878045112 | 393.4708633147749 |
| SE_328_2_14 | SE_328 | SE_328_2 | CATTTCCCCTCTCTGAGCCT | 381.48944827428124 | 394.760031430389 | 372.2976272132231 | 454.15666285867377 | 371.20807945529236 | 255.56148241118058 | 462.7490634887984 | 580.4470432014976 | 514.1434758607733 | 651.7322086752421 | 551.6789334080266 | 558.2875671221436 |
| SE_328_2_31 | SE_328 | SE_328_2 | CAGAGCTATTTCCTTCCACA | 613.4176923955772 | 643.6087601042492 | 596.4662250312646 | 645.4490108249346 | 636.356707637644 | 824.7272245492752 | 679.0573466544926 | 535.0700756548007 | 411.1371836330536 | 613.0948450941667 | 458.9499211968903 | 749.7999342222554 |
| SE_328_2_6 | SE_328 | SE_328_2 | TCACTGCCTTACCTCCCTCA | 2429.827679065166 | 1857.870628934413 | 1959.2532954669355 | 2026.1373182548855 | 2000.281251007661 | 2068.74854236576 | 2351.4109388907546 | 1740.3957761139366 | 1835.4655692646259 | 2456.7534040666283 | 2253.6671322200236 | 1602.9004785772981 |
| SE_328_3_13 | SE_328 | SE_328_3 | TCTAAGCCTACCATGCTGGA | 860.5188123005094 | 747.5455785061544 | 717.9320291353135 | 827.6321993642307 | 774.233994292467 | 504.1924839434139 | 931.9550906541847 | 1068.249444328489 | 710.3882222601359 | 972.4952643990748 | 1029.4094140400837 | 452.6655949639003 |
| SE_328_3_18 | SE_328 | SE_328_3 | GGCAGAGGATACATAGGAAA | 722.8791534060954 | 960.4161777331742 | 900.6244987227042 | 853.6583691565587 | 849.5362046962548 | 1147.8608955756415 | 745.7793046469907 | 746.8292575393862 | 866.6733611573657 | 839.0869934957534 | 895.5979280645198 | 985.4181798060291 |
| SE_328_3_19 | SE_328 | SE_328_3 | CAGGATTGCTTGAGACCAGA | 1317.8726394929715 | 1351.1786392250276 | 1383.5251345509962 | 1397.605317794314 | 1496.498857461193 | 1591.411671828267 | 1345.2007659558094 | 1685.5652736616778 | 1402.1287536859431 | 1596.5252091708949 | 1520.0515292838181 | 1962.7115925229625 |
| SE_328_3_20 | SE_328 | SE_328_3 | TTATTGACCAAACCCAGAGA | 19.50798315038938 | 27.982989569749094 | 49.376343131727204 | 14.314393385230405 | 36.06021343279983 | 28.588233625657487 | 9.685445514881827 | 34.97807915057884 | 42.62329333560815 | 2.9160277793075706 | 53.994108376104734 | 33.659749369110536 |

| | | | | | | | | | | | | | | | |
|---|---|---|---|---|---|---|---|---|---|---|---|---|---|---|---|
| SE_328_3_22 | SE_328 | SE_328_3 | GTAGTCCCTGCTCAAGCAGA | 346.80858934025565 | 237.85541134286728 | 243.91913507073238 | 268.0695488506785 | 214.24009157134017 | 438.3529155934148 | 285.1825623826316 | 270.37109829906893 | 150.95749723027888 | 387.8316946479069 | 184.28423945757487 | 175.26283292192036 |
| SE_329_1_36 | SE_329 | SE_329_1 | ATGGCACCGCAAGCCGTCCC | 4791.3774171039695 | 3994.5717610816832 | 4097.2489530707235 | 4311.235025933484 | 3839.3521360804525 | 3340.491783652584 | 4745.868302292095 | 3991.282437128213 | 3774.825416034797 | 4037.2404604513317 | 3764.3283817862584 | 3718.82196478035 |
| SE_329_1_37 | SE_329 | SE_329_1 | GCGTGGCCGGGGACGGCTTG | 6698.824658475375 | 6395.1125091723015 | 6460.400735355187 | 7006.2449078255 | 7423.100994593118 | 6997.1867574071375 | 7652.578117369409 | 7750.007915579605 | 6635.913981187494 | 7804.019344371886 | 8195.36662352094 | 7097.564392831411 |
| SE_329_1_39 | SE_329 | SE_329_1 | CAAGCCGTCCCCGGCCACGC | 800.9110860076529 | 751.5431484446899 | 725.8322440363899 | 808.1125720207447 | 681.9622716850085 | 1101.0801496427475 | 1038.4949913178848 | 720.3593598038129 | 654.4451497571501 | 726.819923992412 | 724.2253232186222 | 1072.4692557606252 |
| SE_329_1_40 | SE_329 | SE_329_1 | CCGTCCCGCGTGGCCGGGGA | 584.1557176699931 | 551.6646515179107 | 595.4786981686301 | 611.6149900962082 | 648.0232472776676 | 507.6577243828875 | 672.6003829779047 | 789.3701646144145 | 388.0495664095992 | 416.2629654961557 | 612.7157515723189 | 387.6674582511351 |
| SE_329_1_42 | SE_329 | SE_329_1 | TTACCCGTCCCGCGTGGCCG | 27.094421042207472 | 37.97691441608806 | 41.47612823065085 | 37.737946197425615 | 26.51486281823517 | 113.48662439276154 | 36.58946083399801 | 9.45353490556185 | 49.72717555820951 | 64.88161808959345 | 7.042709788187574 | 145.08512659099367 |
| SE_329_2_12 | SE_329 | SE_329_2 | GAAATGATAGAACCAAACTG | 173.40429467012783 | 217.86756165018937 | 218.24343664223423 | 191.29234796626088 | 168.63452752397566 | 254.69517230131217 | 165.7287343657557 | 155.03797245121433 | 187.36489362111084 | 143.61436813089784 | 191.32694924576242 | 377.22132913658356 |
| SE_329_2_13 | SE_329 | SE_329_2 | GCTCCCCAAACACAAAGACA | 355.47880407376203 | 397.7582088842907 | 348.59698250999406 | 325.3271223916001 | 316.0571647933632 | 381.17644834209983 | 350.82835976127507 | 449.98826150474406 | 408.4732277995781 | 462.91940996507685 | 386.17525338561865 | 465.4330861039077 |
| SE_329_2_14 | SE_329 | SE_329_2 | TCCCCAAACACAAAGACACG | 217.8391451793481 | 231.8590564350639 | 183.6799964500252 | 296.6983356211393 | 204.6947409567755 | 134.27806702960336 | 232.45069235716386 | 253.35473546905757 | 142.96562972985234 | 281.39668070318055 | 440.1693617617234 | 76.60494684004466 |
| SE_329_2_4 | SE_329 | SE_329_2 | ACTCCATCCCAGGATTTGGG | 86.70214733506391 | 144.91191027191493 | 149.11655625781614 | 98.89944520704643 | 177.11928362581094 | 299.7432980144694 | 107.61606127646475 | 145.5844375456525 | 89.68651306034215 | 209.95400011014507 | 156.11340030482455 | 305.2591063474507 |
| SE_329_2_5 | SE_329 | SE_329_2 | CTGTGGTAACCACCTTACCA | 332.71949039830776 | 375.77157422234495 | 357.48472427370496 | 357.85983463076013 | 363.7839178661865 | 451.34756724144097 | 405.7125510122721 | 265.64433084628797 | 450.2085358573611 | 487.7056460891912 | 153.7658303754287 | 337.7581747038333 |
| SE_329_3_1 | SE_329 | SE_329_3 | ACTAGCAGACCCATCTACAA | 778.1517723321986 | 982.4028123951199 | 998.389658123524 | 977.2826756643667 | 920.596037049125 | 1052.5667834901167 | 1029.8857064157676 | 959.5337929145278 | 1181.908404785301 | 1832.7234592948082 | 1116.2695014277306 | 1283.713200077112 |
| SE_329_3_21 | SE_329 | SE_329_3 | TTGTCCCCATTTGACTAACG | 122.46678311077778 | 105.935603371193 | 132.3285995930289 | 128.82954046707366 | 142.1196647057405 | 71.03742900920952 | 167.88105559128502 | 87.9178746217252 | 71.92680750383876 | 298.89284737902597 | 77.46980767006332 | 97.49720506914775 |
| SE_329_3_5 | SE_329 | SE_329_3 | TGTGGTGATGAGTCTGAGAG | 1556.3035446643971 | 1576.0419482676543 | 1778.5358796048138 | 1571.9806553962117 | 1583.4676075050043 | 1425.9464408434008 | 1531.3765519640933 | 1630.734771209419 | 1719.1394978695287 | 1452.910841039997 | 1414.4108824610046 | 1305.766139318943 |
| SE_329_3_6 | SE_329 | SE_329_3 | GGAAATGCAGAGTGAGACAA | 716.3764923559655 | 917.4423008939167 | 816.6847153987679 | 727.4314456676178 | 984.2317078128895 | 938.2138489874867 | 848.0145628585423 | 806.3865274444258 | 844.4739992117364 | 944.7930004956529 | 860.384379123582 | 1317.3729494462227 |
| SE_329_3_8 | SE_329 | SE_329_3 | GATGATGCATGAGCTCAGTA | 1544.381999405826 | 1601.0267603835016 | 1625.4692158964594 | 1642.2513138327975 | 1573.9222568904397 | 1726.5560489677387 | 1302.1543414452235 | 1641.133659605537 | 1742.227115092983 | 1390.2162437848842 | 1848.7113193992382 | 1835.036681122888 |
| SE_329_4_14 | SE_329 | SE_329_4 | ATGCATCTGGAGTCAGCAGG | 980.8180417279106 | 1133.3110775748382 | 1062.5789041947694 | 925.2303360817107 | 881.354040078137 | 1164.3207876631413 | 996.5247274200635 | 984.1129836689886 | 900.4170717147222 | 814.3007573716391 | 996.5434350285417 | 1510.0459975590622 |
| SE_329_4_2 | SE_329 | SE_329_4 | CTCAGCGCCCATAAACACTA | 87.78592417675222 | 98.93985597875572 | 100.72773998872348 | 110.61122161314404 | 97.57469517110542 | 94.42780197565655 | 92.54981269775968 | 120.0598933006355 | 193.580790565887 | 301.07986821350664 | 62.210603128990236 | 74.28358481458876 |
| SE_329_4_25 | SE_329 | SE_329_4 | AGAACATGCTTGGTTAGCGT | 1612.6599404321887 | 1480.1002697428003 | 1420.0636284684742 | 1674.7840260719574 | 1532.5590708939928 | 1337.5828096368232 | 1777.8173322871976 | 1609.936994417183 | 1546.8703539714459 | 1733.5785147983506 | 1725.4638981059556 | 1713.1651747864535 |
| SE_329_4_6 | SE_329 | SE_329_4 | AAGGTTCATCGTCTTACCCG | 729.3818144562251 | 672.5911421586121 | 656.7053636519718 | 694.8987334284578 | 562.1150917465856 | 633.2726903138068 | 683.3619891055512 | 640.0043131065372 | 702.3963547597093 | 622.5719308821664 | 639.7128057603713 | 384.1854152129513 |
| SE_329_4_7 | SE_329 | SE_329_4 | AGGTTCATCGTCTTACCCGA | 345.72481249856736 | 392.7612464611212 | 368.3475197626849 | 343.54544124552973 | 354.2385672516219 | 417.561472956573 | 383.1131781442145 | 470.7860382969801 | 370.2898608530958 | 481.8735905305765 | 296.96759606857603 | 311.06251141109044 |
| SE_329_5_0 | SE_329 | SE_329_5 | CTTGTTGAATGAGTGAGCAA | 509.3751155935005 | 512.6883446171887 | 502.6511730809829 | 609.0123731170754 | 389.2381861716923 | 484.2673514164405 | 588.6598551822622 | 333.7097821663333 | 488.3919028038434 | 718.0718406544893 | 142.02798072844942 | 743.9965291586155 |
| SE_329_5_19 | SE_329 | SE_329_5 | GAAGGCCTCAGGGGAAATGA | 1644.0894688411495 | 1723.9520359934709 | 1736.0722245115285 | 1616.2251440414693 | 1730.8902447743917 | 2034.962448080892 | 1706.790731844731 | 1767.811027340066 | 1681.8441162008717 | 1579.7580494398765 | 1871.0132337284988 | 1364.9608709680685 |
| SE_329_5_20 | SE_329 | SE_329_5 | CCCTGACTAGAAACAATGAT | 382.57322511596954 | 370.7746117991755 | 314.033542317785 | 399.50170629688495 | 418.9348325281157 | 330.9304619697321 | 267.9639925783972 | 396.1031125430415 | 396.9294191878509 | 212.14102094462575 | 616.2371064664128 | 575.6977823130629 |
| SE_329_5_22 | SE_329 | SE_329_5 | AGAACAGCGTGAGAAAAGAG | 559.2288503111622 | 713.5662340286019 | 694.2313844320845 | 585.5888203048802 | 619.3871954339735 | 358.6523854855212 | 526.2425396419126 | 532.2340151831322 | 838.2581022669602 | 535.8201044477661 | 539.9410837610474 | 709.176098776777 |
| SE_329_5_5 | SE_329 | SE_329_5 | GATGTGGCTAGAGCAGACTA | 1534.6280078306313 | 1473.104522350363 | 1503.0158849297761 | 1582.3911233127428 | 1655.5880343370604 | 1614.8020447947138 | 1518.4626246109176 | 1792.3902180945267 | 1349.737622294258 | 1417.1895007434794 | 1772.4152966938727 | 1458.9760329990324 |
| SE_33_1_2 | SE_33 | SE_33_1 | GTAAAATTGCACTTCTTAAA | 95.37236206857031 | 127.92223803313871 | 149.11655625781614 | 141.84262536273764 | 162.27096044759924 | 176.7272624131554 | 101.15909759987686 | 182.45322367734371 | 142.96562972985234 | 53.21750697236316 | 125.5949912226784 | 176.42351393464833 |
| SE_33_1_4 | SE_33 | SE_33_1 | ACACACATACCAAAAAGCAC | 6502.661050129794 | 6277.184195985502 | 6100.940957356213 | 6294.4291640326679 | 6118.5697435935947 | 5897.839227984126 | 6691.566690170578 | 6383.9721217259175 | 5926.413744205183 | 5444.952870912061 | 5850.144264054478 | 6228.214314298177 |
| SE_33_1_6 | SE_33 | SE_33_1 | ATAGTATCACTTACTTCACA | 183.15828624532253 | 150.9082651797183 | 196.51784566427426 | 127.52823197750725 | 131.51371957844643 | 91.82887164605133 | 194.7850709104012 | 122.89595377230405 | 82.5826308377408 | 158.9235139722626 | 99.77172199932397 | 55.71268861094157 |
| SE_33_1_7 | SE_33 | SE_33_1 | AATACAACTTTACTATAAGA | 7.586437891818092 | 22.986027146579612 | 11.850322351614528 | 14.314393385230405 | 44.54496953463509 | 0.0 | 6.456963676587884 | 5.67212094333711 | 8.879852778251697 | 10.206097227576498 | 5.868924823489645 | 1.1606810127279494 |
| SE_33_1_8 | SE_33 | SE_33_1 | CTTGGGTTGCATTATCTCAG | 762.9788965485625 | 636.6130127117918 | 654.7303099267027 | 706.6105098345554 | 639.5384911758323 | 637.6042408631488 | 728.5607348416663 | 767.6270343316222 | 689.9645608701569 | 303.2668890479873 | 923.7687672172701 | 813.6373899222925 |
| SE_33_2_1 | SE_33 | SE_33_2 | GGTTTGAGCTTGGTGCATGA | 119.21545258571288 | 165.89915244922676 | 158.9918248841616 | 205.60674135149128 | 169.69512203670507 | 113.48662439276154 | 115.14918556581728 | 163.54615386622 | 161.6133205641809 | 213.59903483427954 | 62.210603128990236 | 126.51423038734649 |
| SE_33_2_19 | SE_33 | SE_33_2 | GTAACTCTACCTATGTACTG | 188.577170453764 | 260.8414384894469 | 209.35569487852334 | 127.52823197750725 | 298.02705807696327 | 109.1550738434195 | 176.4903404934022 | 120.0598933006355 | 311.6828325166346 | 122.47316768097 | 184.28423945757487 | 159.01329874372908 |
| SE_33_2_22 | SE_33 | SE_33_2 | TTACTTTTCTGAAATGCCTG | 17.340429467012783 | 14.990887269508443 | 22.713117840594514 | 10.410467916531204 | 6.363567076376441 | 0.8663101098684087 | 8.60928490211718 | 0.0 | 16.871720278678225 | 20.412194455152996 | 1.173784964697929 | 2.321362025455899 |
| SE_33_2_4 | SE_33 | SE_33_2 | AGTGGAAAGGAAATTCAGAG | 198.3311620289587 | 307.81288526724 | 313.04601545515044 | 294.0957186420065 | 282.1181403860222 | 198.3850151598656 | 167.88105559128502 | 129.51342820619735 | 203.34862862196388 | 138.5113195171096 | 166.67746498710594 | 171.7807898837365 |
| SE_33_2_8 | SE_33 | SE_33_2 | AGGTCATCACCGTTAAAGGT | 1506.4498099467355 | 1660.9903094615354 | 1592.8808294295195 | 1549.8584110735828 | 1657.7092233960627 | 1621.732525673661 | 1359.1908539217497 | 1897.3244555462634 | 1661.4204548108926 | 1885.2119593223445 | 2129.245925962043 | 1917.4450330265724 |
| SE_33_3_12 | SE_33 | SE_33_3 | TCACGCTTCCTTCTGCCACA | 8465.380910427302 | 6898.806321427785 | 7025.266100782146 | 7513.755218756396 | 7115.52858590159 | 6747.689445765036 | 8369.301085470663 | 6733.752913231706 | 7491.043803733132 | 6995.550642558862 | 7427.711256608495 | 7074.350772576852 |
| SE_33_3_15 | SE_33 | SE_33_3 | GGTAAAGGTGACAATAGAGG | 203.7500462374002 | 342.79162222942637 | 312.0584885925159 | 253.7551554654481 | 311.81478674244556 | 432.28874482433594 | 209.85131948910626 | 174.89039575289422 | 292.1471564044809 | 254.42342374458553 | 164.32989505771008 | 333.1154506529215 |
| SE_33_3_16 | SE_33 | SE_33_3 | GAAAGGGTAGCCACTGTCAG | 1378.5641426275163 | 1738.9429232629793 | 1630.4068502096322 | 1595.404208208407 | 1724.5266776980154 | 1645.9892087499766 | 1572.27065524915 | 1363.1997333820188 | 1751.1069678712347 | 1933.3264176809193 | 1515.3563894250265 | 1947.622739357499 |
| SE_33_3_23 | SE_33 | SE_33_3 | AAGAGTTTCCTGGGGACATG | 59.60772629285644 | 79.95139877071169 | 82.9522564613017 | 91.09159426964803 | 69.99923784014085 | 114.35293450262995 | 72.10276105523138 | 79.40969320671954 | 138.5257033407265 | 67.06863892407412 | 89.2076573170426 | 99.81856709040365 |
| SE_33_3_5 | SE_33 | SE_33_3 | AATGTTTCTTATGCACTCCG | 70.44549470973944 | 62.96172653193546 | 43.451181955919935 | 59.86019052005442 | 71.05983235287025 | 68.4384986796043 | 73.17892166799602 | 82.2457536783881 | 24.863587779104755 | 35.72134029651774 | 50.47275348201095 | 104.46129114551545 |
| SE_33_4_11 | SE_33 | SE_33_4 | AGTCAGAACCCCTTTCACAG | 150.64498099467355 | 193.88214201897586 | 230.09375899384875 | 163.96486968536647 | 242.87614341503414 | 167.1978512046029 | 143.1293614976981 | 277.93392622351837 | 401.36934557697674 | 174.23265981362735 | 133.8114859755639 | 473.5578531930034 |
| SE_33_4_12 | SE_33 | SE_33_4 | TTTTAAGCAAAGACATGCCA | 41.18351998451536 | 30.98116702365078 | 29.62580587903632 | 26.02616971921502 | 63.6356707637644 | 19.05882241710499 | 32.28481838293943 | 19.852433010679885 | 25.751573056929924 | 242.03030568252836 | 57.515463270198524 | 5.803405063639747 |
| SE_33_4_3 | SE_33 | SE_33_4 | CAAAGACTAGGAGACAGAGG | 1207.327401640765 | 1235.2491110074957 | 1260.0842767216782 | 1160.7671726932292 | 1408.4695129046522 | 1138.3314843670892 | 1292.4688959303417 | 1783.8820366795212 | 1219.2037864539582 | 1651.200730032912 | 1709.0309086001846 | 1636.5602279464088 |
| SE_33_4_6 | SE_33 | SE_33_4 | TGTAAGTTGACCTTATTCAA | 305.6250693561003 | 298.81835290553494 | 305.1458005540741 | 311.0127290063697 | 277.8757623351046 | 256.42779252104896 | 362.6661265016862 | 324.25624726077143 | 320.5626852948863 | 329.5111390617555 | 556.3740732668184 | 324.9906835638258 |
| SE_33_4_9 | SE_33 | SE_33_4 | TGGCTCTTATGAAAGCAAAT | 0.0 | 0.0 | 0.0 | 0.0 | 1.0605945127294067 | 0.0 | 0.0 | 0.0 | 0.0 | 0.0 | 0.0 | 0.0 |
| SE_330_1_12 | SE_330 | SE_330_1 | GGAAGGCCGAATGAACCACG | 7036.963033082125 | 6848.83669719609 | 6561.12847534391 | 6339.974961167503 | 6564.019439282299 | 6467.004970167671 | 6723.851508553517 | 7013.577546436337 | 6645.68181924357 | 5583.464190429171 | 7008.670024211334 | 6938.551094087682 |
| SE_330_1_15 | SE_330 | SE_330_1 | AAGAGCATCCGGTCACCTCG | 4287.42118571891 | 4866.04200768294 | 4690.752597514084 | 4415.339705698797 | 5160.8528989412935 | 5257.6360567913725 | 4185.188623041714 | 4421.418275331277 | 4749.83325108683 | 5678.964100201494 | 4723.3106979444665 | 4555.672974957201 |
| SE_330_1_18 | SE_330 | SE_330_1 | CTGGAGCTGTTGAGTCACCA | 2060.2597760494564 | 1831.8864243339317 | 1833.8373839123483 | 2009.2203078905222 | 1856.0403977264618 | 1729.1549792973437 | 1926.327496848719 | 1983.3516231868762 | 1650.7646314769906 | 2274.501667859905 | 2328.789369960691 | 1673.7020203537031 |
| SE_330_1_5 | SE_330 | SE_330_1 | TTCTTTATTGTGTAGCCCAG | 0.0 | 0.0 | 0.0 | 0.0 | 0.0 | 13.86096175789454 | 0.0 | 0.0 | 0.0 | 0.0 | 0.0 | 0.0 |
| SE_330_1_6 | SE_330 | SE_330_1 | GTAGGATGAATAAGACAAGC | 671.9416418467454 | 756.5401108678594 | 775.2085871681171 | 637.6411598875362 | 704.2347564523261 | 496.3956929545982 | 564.98432170144 | 790.3155181049707 | 879.1054250469181 | 470.2094794133458 | 690.1855592423823 | 716.1401848531448 |
| SE_330_2_2 | SE_330 | SE_330_2 | TCTGACTGCATAGTACGGAG | 841.01082915012 | 919.4410858631845 | 927.2877240138369 | 778.1824767207075 | 815.5971802889138 | 1038.7058217322221 | 941.6403561690665 | 1019.0910628195675 | 679.3087375362549 | 868.2472712888291 | 863.9057340176757 | 495.6107924348344 |
| SE_330_2_21 | SE_330 | SE_330_2 | AACTACCTACCAGATGGCAA | 173.40429467012783 | 172.89489984166403 | 181.7049427247561 | 140.54131687317124 | 273.63338428418695 | 481.6684210868353 | 218.46060439122343 | 228.77554471459678 | 255.7397600136489 | 207.0379723308375 | 327.4860051507222 | 150.88853165463343 |
| SE_330_2_27 | SE_330 | SE_330_2 | ACCTCCTAAGGCACAAAGCA | 397.7461008996057 | 455.72297299305666 | 466.11267916350477 | 363.0650685890257 | 371.20807945529236 | 356.05345515591597 | 307.7819352506892 | 504.8187639570028 | 420.0170364113053 | 328.0531251721017 | 508.24888971420324 | 337.7581747038333 |
| SE_330_2_5 | SE_330 | SE_330_2 | GCAACGAGCAACCATCTAA | 1956.2171992473795 | 2108.718142577521 | 2189.347054460784 | 1912.9234796626085 | 2319.5201993392125 | 1639.0587278710293 | 1677.7343953000855 | 2156.351311958658 | 2164.908107337764 | 1963.9447093636488 | 2409.7805325248482 | 1790.930802639226 |
| SE_330_2_7 | SE_330 | SE_330_2 | GCAACCATCTAAAGGAAGTG | 1083.7768416882989 | 1255.2369607001735 | 1163.3066441834928 | 1047.5533341009523 | 1174.0781255914533 | 1284.7378929348502 | 1151.4918556581728 | 1327.2763007408837 | 1157.0448170061961 | 1343.5597993159631 | 1449.6244314019423 | 1374.246319069892 |
| SE_331_1_16 | SE_331 | SE_331_1 | AGGACTTGGGACTTTGCGAA | 863.7701428252761 | 1326.1938271091803 | 1227.4958902547382 | 938.2434209773747 | 1268.4710372243705 | 1068.1603654677479 | 842.633759794719 | 910.3754114056062 | 1212.987889509182 | 1002.3845491369774 | 946.0706815465307 | 1161.8416937406773 |
| SE_331_1_18 | SE_331 | SE_331_1 | GGACTTTGCGAAAGGATCGC | 252.52000411337366 | 290.8232130284638 | 274.53246781240324 | 231.63291114281927 | 243.93673792776355 | 396.7700303197312 | 307.7819352506892 | 240.119786601271 | 285.9312594597047 | 292.33178487558393 | 250.0161974806589 | 214.72598735467065 |
| SE_331_2_11 | SE_331 | SE_331_2 | GATGTGGCTGTGCTCCAAGG | 1922.6201171550422 | 2173.6786540787243 | 2123.18275466427 | 1955.8666598183 | 1913.3125009638497 | 1918.010583248657 | 1915.5658907210725 | 2132.7174746947535 | 2230.6190178968263 | 2009.1431399429162 | 2190.2827441263357 | 1736.3787950410124 |
| SE_331_2_24 | SE_331 | SE_331_2 | CTGTGACTCATGGCAGGGGA | 640.5121543377847 | 761.537073291029 | 721.8821365858516 | 685.7895740014931 | 651.2050308158557 | 1064.6951250282743 | 809.2727807990149 | 817.7307693311001 | 628.6935767002249 | 519.7819516615745 | 676.1001396660072 | 509.5389645875698 |
| SE_331_2_3 | SE_331 | SE_331_2 | GGCCTCATAGACCGCAGGAA | 117.04789890233629 | 130.9204154870404 | 153.06666370835433 | 143.14393385230406 | 149.54382629484635 | 181.92512307236584 | 110.8445431147587 | 96.42605603673087 | 165.1652616754816 | 88.9388472688809 | 103.29307689341775 | 113.74673924733905 |
| SE_331_2_4 | SE_331 | SE_331_2 | TTTCCTCTAGAGCAAAGTGC | 11.921545258571289 | 19.987849692677923 | 5.925161175807264 | 15.615701874796805 | 16.969512203670508 | 17.326202197368175 | 11.837766740411123 | 55.775855942814914 | 0.8879852778251698 | 1.4580138896537853 | 70.42709788187574 | 2.321362025455899 |
| SE_331_3_5 | SE_331 | SE_331_3 | CCACAGCCTGACTCAAAGCA | 466.0240419259685 | 382.76732161478225 | 373.28515407585763 | 361.7637600994593 | 510.14596062284465 | 460.87697844999343 | 486.42459696962067 | 536.0154291453568 | 445.76860946823524 | 156.00748619295504 | 600.9779019253397 | 219.36871140558245 |
| SE_331_3_6 | SE_331 | SE_331_3 | ACCTCCTTGGAGATGAGATA | 74.78060207649263 | 85.94775367851507 | 101.71526685135804 | 67.66804145745283 | 123.02896347661108 | 100.4919727447354 | 58.112657089290965 | 90.75393509339376 | 128.75786528464963 | 48.84346530340181 | 123.24742129328254 | 147.40648861644956 |
| SE_331_3_7 | SE_331 | SE_331_3 | GCCTTCCTATTGTACTTCTA | 72.61304839311603 | 112.93135076363026 | 123.440857829318 | 79.37981786355043 | 116.66539640023474 | 28.588233625657487 | 89.32133085946575 | 148.42049801732105 | 93.23845417164283 | 54.67552086201695 | 131.46391604616804 | 26.695663292742836 |
| SE_331_3_8 | SE_331 | SE_331_3 | CCTTCCCTATCTCATCTCCA | 494.2022398098643 | 552.6640440025446 | 576.7156877785737 | 541.3443316596226 | 662.8715704558792 | 540.5775085578871 | 583.2790521184389 | 540.7421965981379 | 552.3268428072556 | 932.3998824335957 | 738.3107427949974 | 571.0550582621511 |
| SE_331_3_9 | SE_331 | SE_331_3 | TCCCTATCTCATCTCCAAGG | 42.26729682584366 | 29.981774530916886 | 57.276558032805 | 18.218318853929606 | 72.12042686559965 | 54.57753692170975 | 17.21856980423436 | 107.7702979234051 | 154.5094383415795 | 8.748083337922711 | 235.93077790428373 | 1.1606810127279494 |
| SE_331_4_20 | SE_331 | SE_331_4 | AGCGCTGCAAAACCAACGCC | 1881.436597170887 | 2101.7223951850838 | 2017.5173803623734 | 1920.7313306000071 | 2040.5838424913786 | 2085.20843445326 | 2096.3608736655333 | 2192.274744599793 | 2219.0752092850994 | 2607.657841645795 | 2236.060357749555 | 2077.6190127830296 |
| SE_331_4_37 | SE_331 | SE_331_4 | TCCTGCCTTGACCAGGACTT | 363.0652419655801 | 291.8226055130977 | 306.13332741670865 | 321.42319692209009 | 353.17797277889244 | 180.19250285262902 | 366.9707689526427 | 293.0595820724173 | 343.6503025183407 | 204.12194455152994 | 306.35787578615947 | 169.45942785828806 |
| SE_331_4_38 | SE_331 | SE_331_4 | TCCCAAGTCCTGGTCAAGGC | 288.2846398890875 | 376.77096670697887 | 315.02106918041954 | 197.79889041409285 | 348.93559467897448 | 166.33154109473446 | 204.47051642528302 | 377.1960427319178 | 221.1083341784673 | 215.7860556687602 | 355.6568443034725 | 194.9944101382955 |
| SE_331_4_42 | SE_331 | SE_331_4 | GGGACTTTGCGAAAGGATCG | 339.22215144843756 | 456.7223654776906 | 441.4245075976412 | 335.7375903081313 | 425.2983996044921 | 290.2138868059169 | 352.9806809868044 | 285.49675414796786 | 351.6421700187672 | 936.7739241025571 | 250.0161974806589 | 596.590040542166 |
| SE_331_4_46 | SE_331 | SE_331_4 | AAGGATCGCGGGGCCCGGAG | 1023.0853385537541 | 1746.9380631400504 | 1590.9057757042503 | 887.4923898824851 | 1275.8951988134763 | 1240.5560773315613 | 829.7198324415432 | 861.2170298966845 | 1345.2976959051323 | 1094.8352420079427 | 760.612657124258 | 472.39717218027545 |
| SE_332_1_1 | SE_332 | SE_332_1 | AGCAACCTCAGCATCTCAG | 253.60378095506195 | 332.79769738308744 | 360.4473048616086 | 274.5760912985105 | 380.753430069857 | 175.86095230328698 | 241.05997725928103 | 177.72645622456278 | 350.75418474094204 | 443.23622245475076 | 161.98232512831422 | 328.4727266020097 |
| SE_332_1_10 | SE_332 | SE_332_1 | GGAGGTGAGAAGGAGACAAG | 2589.142874793346 | 2680.3706437881096 | 2581.395218926916 | 2571.3855753835207 | 2573.0022878051867 | 2276.6629687341783 | 2707.6201017158533 | 2711.2738109151387 | 2837.1129626514175 | 2743.2531333835973 | 2856.992604074759 | 2755.456724216152 |
| SE_332_1_11 | SE_332 | SE_332_1 | TGGCCACCCAGATTATCCAG | 27.094421042207472 | 15.990279754142339 | 22.713117840594514 | 29.930952560022721 | 42.42378050917627 | 39.8502650539468 | 38.74178205952731 | 50.10373499947781 | 85.2465866712163 | 4.374041668961356 | 50.47275348201095 | 96.33652405640198 |
| SE_332_1_12 | SE_332 | SE_332_1 | TAGCCTTTCTATCCAAACAG | 100.7912462770118 | 164.89975996459287 | 129.36601900512528 | 62.46280749918722 | 103.93860224748186 | 109.1550738434195 | 80.71204595734856 | 80.35504669727572 | 29.303514168230603 | 128.30352223411806 | 52.820323411406804 | 124.19286836189059 |
| SE_332_1_14 | SE_332 | SE_332_1 | TTAGTTTCAAAACTTCCCGT | 9.75399157519469 | 25.984204600481302 | 20.738064115325425 | 24.724861301761607 | 25.454268305505764 | 0.0 | 34.43713960846872 | 14.180302358342775 | 20.423661389978903 | 48.114458358574915 | 8.216494752885502 | 0.0 |
| SE_333_1_20 | SE_333 | SE_333_1 | TGACAAATAGGGCATCCCTC | 2594.561759001788 | 2578.432610355452 | 2565.5947891245455 | 2413.927248145673 | 2339.6714950810715 | 2373.68970103944 | 2474.0932487459245 | 2149.7338375247646 | 2276.7942523437355 | 1681.8190217156414 | 2148.0264853972103 | 2117.08216721578 |
| SE_333_1_21 | SE_333 | SE_333_1 | AGGCCCCAGAGGCAAGGTGA | 1315.7050858095595 | 1416.1391507262308 | 1425.0012267811647 | 1355.9634461281892 | 1257.8650920970765 | 1023.1122397545907 | 1223.5946167134 | 1358.472965929238 | 1532.6625896526243 | 927.2968338198075 | 996.5434350285417 | 1093.3615139897283 |
| SE_333_1_23 | SE_333 | SE_333_1 | GGAGTGGTGAGGAGACCTAA | 883.2781259759636 | 1009.3864094802351 | 1012.2150342004077 | 901.8067832695154 | 1001.20122001656 | 1005.7860375572226 | 889.9848267563634 | 1040.8341931023597 | 1082.454053668882 | 1397.5063132331532 | 1009.455069640219 | 1217.5543823516189 |
| SE_333_1_40 | SE_333 | SE_333_1 | ACCAACCTGTCAGGAGCGCA | 2209.8209802024417 | 2085.732115430941 | 2018.504907225008 | 2145.8576992949943 | 2004.5236290585788 | 2363.293979721019 | 2226.576307810056 | 1981.4609162057639 | 2167.5720631712393 | 1757.6357439776382 | 1949.65682636326 | 2086.904460884853 |

| | | | | | | | | | | | | | | | |
|---|---|---|---|---|---|---|---|---|---|---|---|---|---|---|---|
| SE_333_1_9 | SE_333 | SE_333_1 | TTTGACAAGCACGCCCACAG | 366.31657249064506 | 393.7606389457551 | 463.15009857560113 | 370.87291952642414 | 366.9657014043747 | 424.4919538355203 | 297.0203291230427 | 427.29977773139564 | 280.60334779275365 | 341.90425712381267 | 397.9131030325979 | 198.47645317647937 |
| SE_333_2_13 | SE_333 | SE_333_2 | CTTCAACCCAGATCCCCCGA | 358.73013459882696 | 487.70353250134133 | 515.489022295232 | 411.21348270298256 | 442.26791180816264 | 814.3315032763043 | 313.1627383145124 | 574.7749222581605 | 487.5039175260182 | 607.9917919856284 | 484.7731904202447 | 662.7488582676591 |
| SE_333_2_14 | SE_333 | SE_333_2 | GCAGTGCCCCATCTGCCATA | 234.09579780467257 | 288.824428059196 | 228.11870526857967 | 227.72898567412008 | 353.17797273889244 | 113.48662439276154 | 271.19247441669114 | 211.75918188458544 | 428.896889189557 | 201.20591677222237 | 180.76288456348107 | 201.9584962146632 |
| SE_333_2_16 | SE_333 | SE_333_2 | AGAGGGTGACCAATGAGAGG | 4853.152697080202 | 5557.621607049096 | 5499.537098011776 | 4835.662347228744 | 5211.761435552305 | 5738.43816776834 | 4475.7519884881685 | 5105.854202493955 | 5221.353433611998 | 5715.4144474428385 | 4643.493320345007 | 5277.616564873986 |
| SE_333_2_26 | SE_333 | SE_333_2 | GGTGGCTGGGAGCCTGTGCG | 230.84446727960767 | 314.8086326596773 | 297.24558565299776 | 271.9734743193777 | 266.2092226950811 | 248.6310015322333 | 215.2321225529295 | 246.7372610351643 | 278.8273772371033 | 172.77464592397357 | 143.20176569314734 | 240.26096963468552 |
| SE_333_2_34 | SE_333 | SE_333_2 | ATCCACTTAACCCAGAGCGT | 254.68755779675024 | 248.84872867384016 | 290.3328976145559 | 287.5891761941745 | 295.9058690515045 | 226.97324878552308 | 139.90087965940418 | 381.92281018469873 | 302.8029797383829 | 150.90443757916677 | 354.4830593387746 | 153.20989368008932 |
| SE_333_3_16 | SE_333 | SE_333_3 | ATGGAAGGTGAGTAGCCAAA | 1986.562950814652 | 2071.740620646067 | 1950.3655537032246 | 1980.5915211200615 | 2232.551449295401 | 2112.0640478591804 | 2180.301401461176 | 2210.2364609203605 | 2162.2441515042883 | 2107.5590774945467 | 2332.310724854785 | 2031.1917722739115 |
| SE_333_3_18 | SE_333 | SE_333_3 | AGAAGTGCAAGGAAAGACCG | 1425.1665468201131 | 1615.0182551683763 | 1531.6541639461777 | 1487.3956035743956 | 1372.4092994718524 | 1104.545390082221 | 1388.2471904663953 | 1119.298532818523 | 1687.1720278678226 | 1485.7161535572072 | 1235.9955678269193 | 1582.008220348195 |
| SE_333_3_4 | SE_333 | SE_333_3 | GCCTTGATCAGATTACTGTG | 1372.0614815773865 | 1620.0152175915457 | 1510.9160998308523 | 1281.7888622229045 | 1780.7381868726739 | 1796.7271678670797 | 1351.6577296323971 | 1507.838817437115 | 1607.2533528635572 | 1132.1477853161643 | 1892.1413630930615 | 1953.4261444211388 |
| SE_333_3_6 | SE_333 | SE_333_3 | TTAATACAACAGCCAATGCA | 94.288585226882 | 110.93256579436247 | 146.1539756699125 | 97.59813671748003 | 150.60442080757576 | 136.01068724934018 | 83.94052779564251 | 129.51342820619735 | 167.8292175089571 | 118.8281320067835 | 59.86303319959438 | 170.62010887100857 |
| SE_333_3_7 | SE_333 | SE_333_3 | ATAGATTATTGATTATAACG | 3.2513305250648967 | 17.98906472341013 | 11.850322351614528 | 2.602616979132801 | 13.787728665482287 | 19.05882241710499 | 17.21856980423436 | 4.726767452780925 | 19.535676112153734 | 12.393118062057175 | 0.0 | 12.767491140007444 |
| SE_333_4_13 | SE_333 | SE_333_4 | GGTGGTCTGGGACTTGGCCT | 113.79656837727138 | 90.94471610168455 | 88.87741763710896 | 102.80337067574564 | 83.78696650562313 | 77.96790988815678 | 146.35784333599204 | 114.38777235729839 | 82.5826308377408 | 244.94633346183593 | 34.03976397623994 | 110.26469620915519 |
| SE_333_4_16 | SE_333 | SE_333_4 | GCTGTAAAGCTAATGACTGT | 1618.0788246406303 | 1549.0583511825391 | 1498.0782506166033 | 1539.4479431570517 | 1510.2865861266753 | 1710.9624669901073 | 1663.744307334145 | 1292.2982215903048 | 1200.5560956196296 | 1361.0559659918085 | 1671.4697897298508 | 1149.0742026006699 |
| SE_333_4_20 | SE_333 | SE_333_4 | GCAATTCAACGCAGTGGCCA | 969.9802733110275 | 1170.2885995062925 | 1248.2339543700637 | 1133.4396944123348 | 1122.1089944677124 | 863.7111795388035 | 1004.0578517094161 | 1018.1457093290112 | 1178.3564636740002 | 1313.6705145780606 | 1034.1045538988756 | 1209.4296152625234 |
| SE_333_4_21 | SE_333 | SE_333_4 | GTACCTCCTGTGTAGGAGCA | 137.63965889441397 | 135.9173779102099 | 106.65290116453076 | 96.29682822791364 | 144.2408537311993 | 110.02138395328791 | 96.85445514881827 | 107.7702979234051 | 140.30167389637683 | 37.908361130998415 | 64.5581730583861 | 63.83745570003722 |
| SE_333_4_26 | SE_333 | SE_333_4 | GGCCAGATCAATGCTCACCT | 1734.0429467012782 | 1455.1154576269528 | 1443.7642731717033 | 1735.9455250815781 | 1711.7995435452624 | 1626.0640762230032 | 1595.9461887299722 | 1615.6091153605203 | 1315.1061964590765 | 1285.2392437298117 | 1905.0529977047388 | 1055.059040569706 |
| SE_334_1_1 | SE_334 | SE_334_1 | TAGGGTCTATCTATAAAATG | 274.19554094713965 | 420.7442360308703 | 423.6490240702194 | 297.9996441107057 | 357.4203507898101 | 330.9304619697321 | 247.51694093586892 | 285.49675414796786 | 321.4506705727115 | 521.9689724960551 | 262.9278320923361 | 376.06064812385563 |
| SE_334_1_15 | SE_334 | SE_334_1 | TCGTCTCTCAGTATCTGTCG | 848.597267041938 | 951.4216453714691 | 1056.6537430189621 | 896.6015493112499 | 983.1711133001601 | 1121.8715922795893 | 731.7892166799603 | 699.5615830115769 | 1002.5353786646167 | 839.8160004405803 | 882.6862934528426 | 860.0646304314105 |
| SE_334_1_21 | SE_334 | SE_334_1 | GCATCTGCTTTGAACACCTA | 851.8485975670029 | 960.4161777331742 | 892.7242838216278 | 828.9335078537971 | 944.9897108419015 | 983.2619747006439 | 754.3885895480179 | 700.506936502133 | 902.1930422703725 | 1136.5218269851257 | 793.4786361358 | 682.4804354840343 |
| SE_334_1_3 | SE_334 | SE_334_1 | AGATTAAAAGGGAAATACAA | 49.85373471766175 | 98.93985597875572 | 117.51569665351074 | 98.89944520704643 | 80.60518296743491 | 169.79678153420812 | 35.513300221233365 | 124.78666075341643 | 203.34862862196388 | 46.65644446892113 | 43.43004369382337 | 34.820430381838484 |
| SE_334_1_4 | SE_334 | SE_334_1 | TCCCTCAATGAACATACAGA | 139.80721257779055 | 158.9034050567895 | 142.20386821937433 | 93.69421124878083 | 134.69550311663465 | 122.14972549144564 | 105.46374005093546 | 99.26211650839943 | 99.45435111641902 | 126.11820145505243 | 100.9455069640219 | 113.74673924733905 |
| SE_335_1_10 | SE_335 | SE_335_1 | TGGTGTTCCAGGAACCACTG | 11.921545258571289 | 8.994532361705065 | 7.900214901076352 | 16.917010364363207 | 20.15129574185873 | 0.8663101098684087 | 19.370891029763655 | 25.524544245016994 | 11.543808611727208 | 35.72134029651774 | 53.994108376104734 | 15.088853165463343 |
| SE_335_1_25 | SE_335 | SE_335_1 | ATCTGATGAGCAAGACCACT | 266.60910305532155 | 313.8092401750434 | 317.9836497683232 | 283.68525072547527 | 328.7842989461161 | 411.49730218749414 | 287.33488360816085 | 294.0049355629735 | 281.49133307057883 | 207.7669792756644 | 379.1325435974311 | 547.8414380075922 |
| SE_335_1_27 | SE_335 | SE_335_1 | CTTGAGGGACTGTGCCACGA | 2746.2905168381494 | 2581.4307878093537 | 2431.2911358062474 | 2541.45548012318 | 2416.0342999975887 | 2097.3367759914177 | 2600.0040404393885 | 2506.1321034644466 | 2306.097766511966 | 2587.974654135469 | 2405.0853926660566 | 1974.318402650242 |
| SE_335_1_36 | SE_335 | SE_335_1 | CTATAAGCCCCTGACTTTGG | 21.675536833765978 | 16.989672238776237 | 25.675698428498144 | 20.820935833062407 | 19.090701229129323 | 1.7326202197368175 | 7.533124289352532 | 5.67212094333711 | 1.7759705556503396 | 8.019076393095819 | 7.042709788187574 | 0.0 |
| SE_335_1_7 | SE_335 | SE_335_1 | ACAGTGGGTACAGCCCACCG | 1664.6812288332271 | 1719.9544660549354 | 1784.461040780621 | 1890.80123533998 | 1936.6455802438968 | 1783.7325162190537 | 1788.578938414844 | 1677.0570922466723 | 1576.1738681396764 | 1794.0860912189828 | 1653.863015259382 | 1960.3902304975065 |
| SE_335_2_11 | SE_335 | SE_335_2 | AAGCCAATGTGACAAGGTCA | 1240.9244837331023 | 1277.2235953621193 | 1267.9844916227546 | 1050.155951080085 | 1228.168445740653 | 1167.7860281026149 | 1103.0646280837636 | 1085.2658071585004 | 1377.2651659068383 | 1084.0333269575895 | 1220.7363632858462 | 1125.860582346111 |
| SE_335_2_12 | SE_335 | SE_335_2 | AATCCTTGACCTTGTCACAT | 47.686181034285156 | 87.94653864778286 | 66.16429979651446 | 27.32747828089441 | 64.69626527649382 | 89.2299413164461 | 31.208657770174778 | 6.617474433893295 | 95.01442472729316 | 43.74041668961356 | 19.954344399864794 | 51.069964560029774 |
| SE_335_2_13 | SE_335 | SE_335_2 | TTTCTGAACACAGCTCAGAT | 19.50798315038938 | 37.97691441608806 | 19.75053725269088 | 15.615701874796805 | 27.575457330964575 | 8.663101098684088 | 32.28481838293943 | 74.68292575393862 | 5.327911666951019 | 13.122125006884067 | 111.50957164630326 | 4.642724050911798 |
| SE_335_2_14 | SE_335 | SE_335_2 | AACACAGCTCAGATAGGATA | 1237.6731532080373 | 1186.2788792604347 | 1140.5935263428983 | 1104.8109076418739 | 1294.9859000426056 | 925.2191973394605 | 1056.7897217348839 | 1199.6535795157988 | 1349.737622294258 | 1370.5330562745582 | 1448.4506464372444 | 950.5977494241906 |
| SE_335_2_9 | SE_335 | SE_335_2 | ACCTTGTCACATTGGCTTAA | 716.3764923559655 | 772.5303906220017 | 782.1212752065588 | 717.0209777510867 | 860.1421498235488 | 697.379638444069 | 909.3557177861271 | 890.5229881039263 | 585.1822980867869 | 734.8390003855078 | 868.6008738764675 | 1123.539220320655 |
| SE_335_3_2 | SE_335 | SE_335_3 | GAGCCTTTCATACATCCCTT | 88.86970101844051 | 159.90279754142338 | 159.97935174679614 | 110.61122161314404 | 102.87766773475245 | 127.34758615065608 | 106.5399006637001 | 126.67736773452879 | 181.14899667633463 | 121.74415978609107 | 57.515463270198524 | 33.659749369110536 |
| SE_335_3_3 | SE_335 | SE_335_3 | TATGCATTTGAGTTTCTAGG | 211.3364841292183 | 313.8092401750434 | 323.90881094413044 | 236.83814510108488 | 335.1478660224925 | 120.41710527170882 | 250.74542277416285 | 251.4640284879452 | 212.22848140021557 | 367.4195001927539 | 259.4064771982423 | 177.58419494737626 |
| SE_335_3_5 | SE_335 | SE_335_3 | TTTTGCAGGTGTTATCACAG | 65.02661050129794 | 85.94775367851507 | 98.75268626345441 | 79.37981786355043 | 131.51371957844643 | 43.31550549342044 | 80.71204595734856 | 139.91231660231537 | 112.77413028379657 | 244.94633346183593 | 125.5949912226784 | 121.87150633643469 |
| SE_335_3_6 | SE_335 | SE_335_3 | GTCTCCAGAGCAAATGAAAG | 35.76463577571386 | 58.96415659339988 | 66.16429979651446 | 55.95626505135522 | 66.81745430195262 | 28.588233625657487 | 22.599372868057596 | 44.431614056140695 | 57.71904305863604 | 48.84346530340181 | 43.43004369382337 | 2.321362025455899 |
| SE_335_3_7 | SE_335 | SE_335_3 | GCATAAAATAGGGCTGAAAG | 1272.3540121420629 | 1501.087511920112 | 1533.629217671447 | 1279.1862452437717 | 1368.1669214209346 | 1405.154998206559 | 1478.6446819386256 | 1538.090129134913 | 1443.864061743726 | 1442.7047438124205 | 1556.438863189454 | 1775.8419494737627 |
| SE_336_1_11 | SE_336 | SE_336_1 | AGTAGTTCCTTTCAGGGCAG | 49.85373471766175 | 79.95139877071169 | 56.28903117016901 | 45.545797134824014 | 32.87842989461161 | 116.95186483223517 | 10.761606127646475 | 31.196665188354107 | 80.80666028209045 | 2.9160277793075706 | 41.08247376442752 | 1.1606810127279494 |
| SE_336_1_15 | SE_336 | SE_336_1 | GTACCCACACATTTCCTGGT | 163.65030309493312 | 151.90765766435223 | 195.5303188016397 | 184.78580551842887 | 168.63452752397566 | 266.8235138394699 | 160.34793130193248 | 114.38777235729839 | 183.81295250981015 | 210.68300705497197 | 171.37260484589763 | 341.24021774201714 |
| SE_336_1_28 | SE_336 | SE_336_1 | TATAGGGTTCTTCTTCTAAG | 17.340429467012783 | 34.97873696218637 | 34.56344019220904 | 42.943180155691216 | 20.15129574185873 | 6.93048087894727 | 39.817942672291956 | 0.945353490556185 | 17.759705556503395 | 26.244250013768134 | 10.564064682281362 | 11.606810127279495 |
| SE_336_1_4 | SE_336 | SE_336_1 | TTAGAAATGATGCAACCTTT | 170.15296414506292 | 155.9052276028878 | 170.8421472357761 | 161.36225270623365 | 151.66501532030517 | 261.62565318025946 | 90.39749147223039 | 170.1636283001133 | 257.51573056929925 | 163.29755564122397 | 281.70839152750295 | 75.44426582731671 |
| SE_336_1_5 | SE_336 | SE_336_1 | TCTTCATTCCCCCTGCATGT | 624.2554608124601 | 733.5540837212798 | 592.5161175807265 | 605.1084476483762 | 579.0846039502561 | 796.1389909690677 | 620.9446735652016 | 502.9280569758904 | 562.9826661411577 | 460.0033821857693 | 393.21796317380625 | 750.9606152349833 |
| SE_336_2_11 | SE_336 | SE_336_2 | GATGAGAGCCTAAGTTTAGA | 783.5706565406401 | 877.4666015085609 | 823.5974034372098 | 900.505474779949 | 1030.8978663729833 | 916.5560962407765 | 841.5575991819543 | 1072.0308582907137 | 795.6348089313522 | 931.6708754887688 | 1347.5051394732225 | 1168.805779817045 |
| SE_336_2_25 | SE_336 | SE_336_2 | TGAGCTTCAGACTTAGGTTT | 1077.2741806381691 | 1197.2721965914077 | 1295.6352437765217 | 1060.5664189966164 | 1217.562500613359 | 1247.4865582105085 | 1075.0844521518827 | 1156.1673189502142 | 1035.390833944148 | 1030.0868130403994 | 1129.1811360394076 | 1136.3067114606624 |
| SE_336_2_5 | SE_336 | SE_336_2 | TTCAGGACTTGGGTACCAAT | 49.85373471766175 | 47.97083926242702 | 75.05204156022535 | 59.86019052005442 | 57.27210368738796 | 11.262031428289314 | 76.40740350628997 | 49.15838150892162 | 72.81479278166393 | 34.26332640686395 | 69.25331291717781 | 3.482040308183848 |
| SE_336_2_6 | SE_336 | SE_336_2 | GAGTGGGGATACATGCCGGA | 734.8006986646667 | 1000.39187711853 | 988.5143894971786 | 778.1824767607075 | 852.717988234443 | 737.2299034980158 | 601.5737825354379 | 815.8400623499876 | 848.9139256008623 | 739.942048999296 | 584.5449124195686 | 726.5863139676964 |
| SE_336_2_8 | SE_336 | SE_336_2 | GAAGAGGGTCAAACTCAGAG | 716.3764923559655 | 1174.286169444828 | 1160.3440635955892 | 931.7368785295428 | 1080.7458084712655 | 1050.8341632703798 | 797.4350140586038 | 787.4794576333021 | 1314.2182111812513 | 809.1977087578508 | 713.6612585363408 | 1322.0156734971345 |
| SE_336_3_1 | SE_336 | SE_336_3 | TTCTATGCATAGCTACCTAG | 4.335107366753196 | 4.996962423169481 | 2.962580587903632 | 5.205233958265602 | 0.0 | 0.0 | 2.152321225529295 | 0.0 | 0.0 | 0.0 | 1.173784964697929 | 0.0 |
| SE_336_3_17 | SE_336 | SE_336_3 | TTTGCTGCAGAAGTAGACAG | 55.27261892610324 | 19.987849692677923 | 35.55096705484358 | 102.80337067574564 | 11.666539640023474 | 116.08555472236677 | 72.10276105523138 | 42.54090707502832 | 79.03068972644012 | 2.9160277793075706 | 15.259204541073078 | 95.17584304369186 |
| SE_336_3_18 | SE_336 | SE_336_3 | CTTAATGAGAAGAAACAGAA | 82.36703996831072 | 197.87971195751143 | 157.0167711588925 | 127.52823197750725 | 131.51371957844643 | 94.42780197565655 | 128.06311291899306 | 69.0108048106015 | 90.57449833816732 | 155.27847924812812 | 80.99116256415711 | 40.62383544547823 |
| SE_336_3_2 | SE_336 | SE_336_3 | TGAATCTGACAGCTATAAGA | 57.440172609479845 | 96.94107100948793 | 80.97720273603261 | 85.88636031138243 | 78.4839939419761 | 102.22459296447224 | 38.74178205952731 | 52.93979547114636 | 112.77413028379657 | 51.030486137882484 | 37.56111887033373 | 26.695663292742836 |
| SE_336_3_9 | SE_336 | SE_336_3 | CTATCCTCTGGCCTGTCTTG | 128.96944416090756 | 189.88457208044028 | 181.7049427247561 | 136.63739140447206 | 171.8163110621639 | 150.73795911710312 | 124.83463108069911 | 128.56807471564116 | 238.86803973497067 | 145.07238202055163 | 113.85714157569912 | 211.2439443164868 |
| SE_337_1_15 | SE_337 | SE_337_1 | GCCAACAGCTCAGAGCTGTA | 676.2767492134985 | 687.5820294281206 | 763.3582648165026 | 700.1039673867234 | 818.778963827102 | 986.7272151401176 | 776.9879624160754 | 810.1679414066506 | 921.7287183825263 | 908.3426532543083 | 635.0176659015796 | 1033.006101327875 |
| SE_337_1_16 | SE_337 | SE_337_1 | CCTCCCGGAAGACATCCAGA | 707.7062776224592 | 791.5188478300457 | 763.3582648165026 | 812.0164974894338 | 738.1737808596671 | 1146.1282753559049 | 768.3786775139583 | 965.2059138578649 | 1167.7006403400983 | 846.3770629440223 | 809.911625641571 | 788.1024076422776 |
| SE_337_1_20 | SE_337 | SE_337_1 | TGGCCTGGAGAGCTTGGTGA | 35.76463577571386 | 54.96658665486429 | 49.376343131727204 | 31.23140374959361 | 55.15091466192915 | 31.187163955262715 | 33.36097899570407 | 35.92343264113503 | 64.82292528123739 | 33.53431946203706 | 31.692194046844083 | 47.58792152184593 |
| SE_337_1_29 | SE_337 | SE_337_1 | AGATCTTCCTGTGAACCACA | 213.50403781259487 | 241.8529812814029 | 248.8567693839051 | 219.92113473672168 | 236.5125763386577 | 365.5828663644685 | 229.2222105188699 | 191.90675858290555 | 396.9294191878509 | 436.67515995130873 | 143.20176569314734 | 575.6977823130629 |
| SE_337_1_9 | SE_337 | SE_337_1 | ACATCATTAATCTCCTGGAA | 270.9442104220747 | 468.71507529329733 | 471.0503134766775 | 320.1218884333345 | 502.7217990337388 | 297.1443676848642 | 307.7819352506892 | 469.84068480642395 | 561.2066955855073 | 390.01871548238756 | 523.5080942552763 | 327.31204558928175 |
| SE_337_2_19 | SE_337 | SE_337_2 | TTTTCTGCTGGGCTCCCCCC | 3.2513305250648967 | 16.989672238776237 | 12.837849214249072 | 9.109159426964803 | 5.3029725636470335 | 0.0 | 8.60928490211718 | 0.945353490556185 | 29.303514168230603 | 25.515243068941242 | 0.0 | 1.1606810127279494 |
| SE_337_2_2 | SE_337 | SE_337_2 | TTAAAACAAAAGCAGCAGTG | 81.28326312662242 | 86.94714616314897 | 86.90236391183987 | 55.95626505135522 | 94.3929116329172 | 52.844916701972934 | 87.16900963393644 | 86.02716764061283 | 63.93494000341222 | 62.69459725511277 | 119.72606639918877 | 100.9792481073316 |
| SE_337_2_33 | SE_337 | SE_337_2 | GGGGTTGGCACAGACCCAAG | 189.6609472954523 | 164.89975996459287 | 172.81720096104522 | 197.79889041409285 | 151.66501532030517 | 95.29411208552496 | 207.69899826357695 | 154.09261896065814 | 119.87801250639792 | 251.50739596527796 | 219.49778839851274 | 236.7789265965017 |
| SE_337_2_36 | SE_337 | SE_337_2 | CAATGCCCATGTCACCAGCG | 1510.7849173134887 | 1231.25154106896 | 1180.0946008482802 | 1495.203454511794 | 1269.5316317370998 | 1427.6790610631376 | 1488.3301274535074 | 1470.9700313054238 | 1024.735010610246 | 1500.296292453745 | 1662.0795100122675 | 1319.6943114716785 |
| SE_337_2_47 | SE_337 | SE_337_2 | AGACAAGCACCTACAAAGGG | 417.2540840499951 | 567.654931272053 | 567.8279460148628 | 451.554045879541 | 638.4778966631029 | 398.502650539468 | 421.8549602037418 | 444.31614056140694 | 469.7442119695148 | 413.3469377168481 | 551.6789334080266 | 639.5352380131002 |
| SE_338_1_11 | SE_338 | SE_338_1 | TTAAATACTCCCTTCATGGG | 4.335107366753196 | 10.993317330972857 | 7.900214901076352 | 7.807850937398403 | 3.1817835381882205 | 0.0 | 4.30464245105859 | 0.0 | 3.551941111300679 | 0.0 | 1.173784964697929 | 0.0 |
| SE_338_1_2 | SE_338 | SE_338_1 | GAAAGAGTGCAGCAAGCGCG | 1802.320887727641 | 1801.9046497949148 | 1680.770720203994 | 1812.7227259659958 | 1929.2214186547908 | 2302.6522720302305 | 1925.2513362359543 | 1761.1935529061727 | 1778.634511483815 | 1945.7195357429764 | 1489.5331202016719 | 1846.6434912501675 |
| SE_338_1_28 | SE_338 | SE_338_1 | GTATTGATAATGGTTATTCA | 18.42420630870108 | 43.97326932389143 | 29.62580587903632 | 15.615701874796805 | 45.60556404736449 | 25.989303296052263 | 9.685445514881827 | 22.68848377334844 | 21.311646667804077 | 0.7290069448268927 | 29.344624117448227 | 125.35354937461854 |
| SE_338_1_34 | SE_338 | SE_338_1 | TCTCTGCTGAAGCAAAGGGA | 115.96412206064798 | 125.92345306387092 | 139.2412876314707 | 109.30991312357764 | 176.05868911308153 | 112.62031428289313 | 130.21543414452233 | 137.07625613064684 | 125.20592417334893 | 167.67159731018532 | 178.4153146340852 | 189.19100507465575 |
| SE_338_1_6 | SE_338 | SE_338_1 | TGTAATACCAGCTACTCGCG | 286.1170862057109 | 363.7788644067382 | 343.6593481968213 | 277.1787082776433 | 382.87461909531584 | 202.71656570920763 | 221.6890862295174 | 202.30564697902358 | 340.09836140704004 | 492.80869470297944 | 251.1899824453568 | 297.13433925835506 |
| SE_338_2_0 | SE_338 | SE_338_2 | TGAAATCACCATAAACCATG | 174.48807151181612 | 183.8882171726369 | 193.55526507637063 | 179.58057156016326 | 201.5129574185873 | 123.01603560131404 | 187.25194662104866 | 138.021609621203 | 182.0369819541598 | 231.82420845495187 | 73.94845277596953 | 148.56716962917753 |
| SE_338_2_11 | SE_338 | SE_338_2 | GTTTTAGCCTCATCATTTAG | 7.586437891818092 | 12.992102300240651 | 9.87526862634544 | 2.602616979132801 | 9.545350614564661 | 0.0 | 6.456963676587884 | 10.398888396118036 | 34.63142583518162 | 15.309145841364746 | 3.521354894093787 | 0.0 |
| SE_338_2_14 | SE_338 | SE_338_2 | CTAGGAAATTCAGTAGGGGG | 55.27261892610324 | 81.9501837399795 | 78.01462214812898 | 65.06542447832003 | 81.66577748016432 | 181.92512307236584 | 61.3411549275849 | 49.15838150892162 | 87.9105425046918 | 101.33196533093808 | 62.210603128990236 | 157.85261773100112 |
| SE_338_2_2 | SE_338 | SE_338_2 | GTTCTTATTAAGATACCACA | 1.083776841688299 | 6.9957473924372735 | 2.962580587903632 | 1.3013084895664004 | 3.1817835381882205 | 2.598930329605226 | 0.0 | 0.0 | 0.0 | 0.0 | 0.0 | 0.0 |
| SE_338_2_9 | SE_338 | SE_338_2 | TGAACTTTGGATGGAGGGAC | 449.767389300644 | 640.6105826503275 | 579.6782683664774 | 506.20900244132974 | 481.50990877915063 | 675.7218856973589 | 520.8617365780893 | 537.9061361264693 | 459.0883886356128 | 605.8047711511477 | 545.810008584537 | 739.3538051077038 |
| SE_339_1_13 | SE_339 | SE_339_1 | GATATCGCGTGAACCCCGGG | 376.07056406583973 | 394.760031430389 | 456.23741053715935 | 353.95590916206095 | 366.9657014043747 | 318.8021204315744 | 303.47729279963056 | 440.5347265991822 | 356.08209640789306 | 578.1025072477258 | 370.91604884454557 | 362.13247597112024 |
| SE_339_1_14 | SE_339 | SE_339_1 | AGATATCGCGTGAACCCCGG | 654.6012123797325 | 740.5498311137171 | 728.7948246242935 | 714.4183607719539 | 845.2938266453372 | 886.2352423953821 | 691.9712740076683 | 709.0151179171388 | 860.4577342125896 | 972.4952643990748 | 605.6730417841313 | 833.3689671386677 |
| SE_339_1_20 | SE_339 | SE_339_1 | AGTAGATTTCTCATACAGCA | 42.26729682584366 | 67.95868895510495 | 67.151826659149 | 37.737946197425615 | 38.181402458258646 | 108.2887637335511 | 35.513300221233365 | 92.64464207450614 | 122.54196833987343 | 15.309145841364746 | 341.57142472709734 | 315.70523546200224 |
| SE_339_1_21 | SE_339 | SE_339_1 | GCCCCTTCAGGCTCCTACAG | 2.167553683376598 | 7.9951398770711695 | 6.912688038441808 | 0.0 | 5.3029725636470335 | 0.0 | 0.0 | 0.0 | 0.0 | 0.0 | 0.0 | 0.0 |
| SE_339_1_9 | SE_339 | SE_339_1 | TGAAAGCTCAAGAAAGACCA | 891.94834070947 | 971.4094950641471 | 902.5995524479732 | 948.6538888939059 | 1000.1406255038305 | 1133.999933817747 | 758.6932319990765 | 1206.271053949692 | 989.2155994972392 | 885.7434379646746 | 1254.7761272620862 | 1154.8776076643096 |
| SE_339_2_10 | SE_339 | SE_339_2 | AATAGATCTTCATTTCTGGG | 8.670214733506391 | 7.9951398770711695 | 8.887741763710896 | 31.23140374959361 | 11.666539640023474 | 0.8663101098684087 | 19.370891029763655 | 2.836060471668555 | 3.551941111300679 | 28.431270848248815 | 0.0 | 4.642724050911798 |
| SE_339_2_11 | SE_339 | SE_339_2 | GGGTGTGGTGGTGCTATTAA | 249.26867358830876 | 300.81713787480277 | 284.40773643874866 | 264.1656233819793 | 326.6631099206573 | 255.56148241118058 | 259.35470767628004 | 241.06514009182717 | 278.8273772371033 | 200.47690982739547 | 136.15905590495976 | 575.6977823130629 |
| SE_339_2_20 | SE_339 | SE_339_2 | CCCTCACCTCCAATCAAGCA | 429.1756293085664 | 517.6853070403582 | 554.0025699379792 | 455.45797134824016 | 482.5705032918801 | 397.6363404295996 | 511.17629106320754 | 339.3819031096704 | 618.0377533663182 | 832.5259309923114 | 727.746678112716 | 410.8810785056941 |
| SE_339_2_21 | SE_339 | SE_339_2 | ATTTTAGCAGCCGAGCTGAG | 35.76463577571386 | 23.98541963121351 | 27.650752153767232 | 36.43663770785921 | 21.211890254588134 | 32.05347406513112 | 19.370891029763655 | 25.524544245016994 | 39.95933750213264 | 8.019076393095819 | 21.128129364562724 | 2.321362025455899 |
| SE_339_2_3 | SE_339 | SE_339_2 | TCCCTACGTGCAGATAACTG | 356.5625809154503 | 402.75517130746016 | 385.13547642747216 | 326.62843088116654 | 357.4203507898101 | 298.0106777947326 | 319.61970199110027 | 476.45815924031723 | 575.4144600307101 | 449.0682780133659 | 458.9499211968903 | 228.65415950740604 |

| | | | | | | | | | | | | | | |
|---|---|---|---|---|---|---|---|---|---|---|---|---|---|---|
| SE_339_3_10 | SE_339 | SE_339_3 | GTAGTTCTATAATAAGCAGG | 13.005322100259587 | 23.98541963121351 | 27.650752153767232 | 35.13532921829281 | 29.69664635642339 | 1.7326202197368175 | 30.13249715741013 | 20.79777679223607 | 40.84732277995781 | 31.347298627556384 | 18.780559435166865 | 1.1606810127279494 |
| SE_339_3_14 | SE_339 | SE_339_3 | AGAAGGTTGCTGTGCTGCAA | 1039.3419911790786 | 995.3949146953606 | 953.9509493049695 | 1104.8109076418739 | 908.9294974091016 | 992.7913859091964 | 999.7532092583575 | 1219.5060028174787 | 770.7712211522473 | 891.5754935232898 | 1103.3578668160533 | 1053.898359556978 |
| SE_339_3_15 | SE_339 | SE_339_3 | CTGTTAGACTGAACTCGCCA | 273.1117641054513 | 311.8104552057756 | 298.2331125156323 | 327.92973937073293 | 331.9660824843043 | 330.0641518598637 | 236.75533480822244 | 616.3704758426326 | 340.9863466848652 | 139.96933340676338 | 490.64211524373434 | 453.8262759766282 |
| SE_339_3_6 | SE_339 | SE_339_3 | TCACTTCCAGAGCTTGACCC | 1231.1704921579076 | 1146.303179875079 | 1037.8907326289059 | 1202.409044359354 | 1040.443216987548 | 1439.8074026012953 | 1232.2039016155213 | 1143.8777235729838 | 787.6429414309256 | 1061.4341116679557 | 1309.9440206028887 | 1053.898359556978 |
| SE_339_3_9 | SE_339 | SE_339_3 | GGAGGGGTGGAGGTTGCAGT | 6986.025521522774 | 6986.752860075569 | 7029.216208232684 | 7299.03931797794 | 6899.167305304791 | 6092.759002704519 | 7623.521780824763 | 7418.188840394384 | 6813.511036752528 | 8135.717504268122 | 7192.954263668909 | 6492.849585200149 |
| SE_34_1_0 | SE_34 | SE_34_1 | TCCCCATTGCAAGTACCAAT | 161.48274941155654 | 122.92527560996923 | 141.2163413567398 | 144.44524234187045 | 126.21074701479941 | 183.65774329210265 | 114.07302495305264 | 182.45322367734371 | 141.189659174202 | 192.45783343429966 | 136.15905590495976 | 42.945197470934126 |
| SE_34_1_18 | SE_34 | SE_34_1 | GCCAGCATAAGGAACTATAT | 778.1517723321986 | 851.4823969080795 | 919.3875091127605 | 818.5230399372658 | 903.6265248454546 | 945.144329866434 | 823.2628687649553 | 1006.8014674423371 | 779.651073930499 | 747.9611253923919 | 1334.5935048615454 | 838.0116911895794 |
| SE_34_1_2 | SE_34 | SE_34_1 | TAAACACAAATACTGCTCAA | 699.0360628889528 | 821.5006223690626 | 833.4726720635551 | 692.296116449325 | 760.4462656269847 | 963.3368421736706 | 681.2096678800218 | 673.0916852760038 | 819.6104114326317 | 1241.4988270401982 | 697.2282690305699 | 879.7962076477856 |
| SE_34_1_20 | SE_34 | SE_34_1 | GTTTGTTTGAGCCTGGTTGG | 47.686181034285156 | 58.96415659339988 | 62.21419234597627 | 52.05233958265602 | 63.6356707637644 | 57.176467251314975 | 64.56963676587885 | 128.56807471564116 | 31.07948472388094 | 83.83579865509266 | 66.90574298778195 | 9.285448101823595 |
| SE_34_1_4 | SE_34 | SE_34_1 | GGTGCGAGTGGATCTGAAAT | 1233.3380458412842 | 1272.22663293895 | 1281.809867699638 | 1415.8236366482438 | 1395.7423787518992 | 1483.9892182045842 | 1479.7208425513902 | 1288.5168076280802 | 1061.1424070010778 | 1598.7122300053757 | 1615.1281114243504 | 1144.4314785497581 |
| SE_34_2_13 | SE_34 | SE_34_2 | GGACTCTGAAGATCTGCAGA | 4.335107366753196 | 3.9975699385355847 | 3.950107450538176 | 10.410467916531204 | 0.0 | 15.593581977631358 | 29.056336544645482 | 10.398888396118036 | 6.215896944776189 | 0.0 | 4.695139858791716 | 2.321362025455899 |
| SE_34_2_14 | SE_34 | SE_34_2 | GCTGGGGTAGGGAGGCTGAG | 5877.321812475645 | 5816.464260569276 | 6054.527194812389 | 5699.731184300834 | 5728.270963251526 | 6198.448836108464 | 6228.81762668178 | 5282.635305227962 | 6049.843697822882 | 6698.11580906949 | 4603.584631545277 | 5582.875671221437 |
| SE_34_2_15 | SE_34 | SE_34_2 | GGAGAGTTCTGTATAGACAG | 203.7500462374002 | 400.7563863381924 | 378.22278838903037 | 238.13945359065127 | 312.875381255175 | 674.8555755874904 | 223.84140745504666 | 227.8301912240406 | 381.833669464823 | 450.5262919030197 | 246.49484258656508 | 362.13247597112024 |
| SE_34_2_4 | SE_34 | SE_34_2 | TTGTGTCTGGAGCTGCACCT | 573.3179492531101 | 596.637313326436 | 582.640848954381 | 455.45797134824016 | 615.144817383056 | 591.6898050401231 | 473.5106696164449 | 364.90644735468743 | 686.4126197588563 | 504.4728058202097 | 388.52282331501453 | 514.1816886384815 |
| SE_34_2_7 | SE_34 | SE_34_2 | CATGATTAAGAGATTTTGGG | 1233.3380458412842 | 1876.859086142457 | 1721.2593215720103 | 1354.6621376386229 | 1823.1619673818502 | 1210.235223486167 | 1171.9389073007012 | 1600.4834595116213 | 1641.884778698739 | 1380.0101465573077 | 1610.4329715655585 | 1602.9004785772981 |
| SE_340_1_1 | SE_340 | SE_340_1 | TATTGTTACAGCTAGAACTA | 93.2048083851937 | 73.95504386290831 | 91.8399982250126 | 94.99551973834723 | 86.96875004381135 | 47.64705604276248 | 69.95043982970209 | 91.69928858394995 | 188.252878898936 | 143.61436813089784 | 71.60088284657367 | 6.964086076367696 |
| SE_340_1_10 | SE_340 | SE_340_1 | TCTGATTCAGTGAGTCTGAG | 358.73013459882696 | 313.8092401750434 | 335.75913329574496 | 274.5760912985105 | 371.20807945529236 | 298.0106777947326 | 263.65935012733866 | 396.1031125430415 | 366.7379197417951 | 268.2745556962965 | 414.34609253836896 | 481.68262028209904 |
| SE_340_1_14 | SE_340 | SE_340_1 | TGTAGTTTCTTTGGAGACAG | 82.36703996831072 | 137.91616287947767 | 137.26623390620162 | 144.44524234187045 | 112.42301834931712 | 183.65774329210265 | 102.23525821264151 | 89.80858160283758 | 98.56636583859384 | 181.52272926189627 | 163.15611009301213 | 120.71082532370674 |
| SE_340_1_27 | SE_340 | SE_340_1 | AGAGATGCCAGGATTAGTGG | 1677.6865509334866 | 1873.8609086885554 | 1939.5027582142445 | 1845.2554382051558 | 2160.4310224298015 | 2105.9998770901016 | 1600.250831181031 | 1840.6032461128923 | 2072.5576384439464 | 1740.1395773017928 | 1829.9307599640713 | 1949.944101382955 |
| SE_340_1_4 | SE_340 | SE_340_1 | GATGATTCCCAAACACACAT | 301.2899619893471 | 293.8213904823655 | 297.24558565299776 | 282.3839422359089 | 282.1181403860222 | 168.93047142433971 | 350.82835976127507 | 292.11422858186114 | 465.304285580389 | 365.9614863031001 | 339.2238547977015 | 119.5501443109788 |
| SE_340_2_0 | SE_340 | SE_340_2 | GCAGTGATGTTGATCATAGA | 2124.2026097090657 | 2059.74791083046 | 2037.2679176150643 | 2145.8576992949943 | 1981.1905497785317 | 1911.0801023697097 | 1872.5194662104866 | 2235.7610051653774 | 2237.722900119428 | 1864.0707579223645 | 2403.9116077013587 | 1588.9723064245627 |
| SE_340_2_13 | SE_340 | SE_340_2 | CTCTAATTAGAGAAGCAGGC | 4.335107366753196 | 9.993924846338961 | 8.887741763710896 | 19.519627343496005 | 7.4241615891058474 | 0.0 | 17.21856980423436 | 0.945353490556185 | 24.863587779104755 | 59.7785694758052 | 3.521354894093787 | 0.0 |
| SE_340_2_25 | SE_340 | SE_340_2 | TGGTAGCCCCAGATAAGCAG | 1.083776841688299 | 3.9975699385355847 | 1.975053725269088 | 0.0 | 0.0 | 0.8663101098684087 | 3.228481838293942 | 0.0 | 0.0 | 6.561062503442034 | 0.0 | 0.0 |
| SE_340_2_3 | SE_340 | SE_340_2 | AATTCTCATGAGGAGAAGAG | 0.0 | 1.9987849692677924 | 11.850322351614528 | 1.3013084895664004 | 1.0605945127294067 | 0.8663101098684087 | 10.761606127646475 | 0.0 | 0.0 | 1.4580138896537853 | 0.0 | 0.0 |
| SE_340_2_30 | SE_340 | SE_340_2 | AAGCTCCTGGTTGAGCCCAG | 73.69682523480432 | 80.95079125534559 | 76.03956842285989 | 61.16149900962082 | 107.12004578567009 | 106.55614351381428 | 86.0928490211718 | 91.69928858394995 | 59.49501361428638 | 312.0149723859101 | 5.868924823489645 | 47.58792152184593 |
| SE_340_3_16 | SE_340 | SE_340_3 | GAAAAAGGATGCAAGTGAGG | 399.9136545829823 | 451.72540305452105 | 438.46192700973756 | 374.7768449951233 | 393.4805642226099 | 210.51335669802333 | 422.93112081650645 | 432.97189867473276 | 365.84993446396993 | 450.5262919030197 | 664.3622900190278 | 203.11917722739116 |
| SE_340_3_28 | SE_340 | SE_340_3 | GAAGTTTTAGTGAAGGCCAA | 101.8750231187001 | 80.95079125534559 | 98.75268626345441 | 98.89944520704643 | 106.05945127294068 | 224.37431845591786 | 79.63588534458391 | 32.14201867891029 | 27.527543612580263 | 109.3510417240339 | 68.07952795247988 | 211.2439443164868 |
| SE_340_3_32 | SE_340 | SE_340_3 | CCTACTCATCCTCACTAACA | 1409.9936710364768 | 1321.1968646860107 | 1354.8868555345944 | 1398.9066262838805 | 1148.6238572859475 | 1618.2672852341875 | 1349.505408406868 | 1069.1947978190453 | 1355.9535192390342 | 1372.7200771090388 | 1234.8217828622214 | 1601.7397975645702 |
| SE_340_3_6 | SE_340 | SE_340_3 | TCACAAGTGTGAATTCAGGG | 298.0386314642822 | 413.748488638433 | 360.4473048616086 | 344.8467497350961 | 452.8738569354567 | 300.60960812433785 | 246.44078032310426 | 418.79159631639 | 517.695416972074 | 269.7325695859503 | 428.4315121147441 | 508.3782835748419 |
| SE_340_3_8 | SE_340 | SE_340_3 | AGAGTTGCCCTCTTTTACCC | 361.98146512389184 | 412.74909615379914 | 429.5741852460267 | 351.3532921829281 | 452.8738569354567 | 392.43847977038916 | 420.77879959097714 | 435.8079591464013 | 258.40371584712443 | 409.70190299271366 | 498.85860999661986 | 365.61451900930405 |
| SE_341_1_12 | SE_341 | SE_341_1 | GCTAACAGCAGGCCTGACTA | 3848.4915648351493 | 3490.8779488261994 | 3608.423156066624 | 3725.6462056286045 | 3880.715322076899 | 3988.491745834154 | 3615.8996588892155 | 3717.1299248669193 | 3889.3755168742437 | 3337.3937934175146 | 4233.84236766543 | 4483.710752168069 |
| SE_341_1_13 | SE_341 | SE_341_1 | ATAGAATCAGCTTTGTTGAG | 151.72875783636184 | 182.888824688003 | 178.74236213685248 | 170.47141213319847 | 161.21036593486983 | 264.22458350986466 | 130.21543414452233 | 188.1253446206808 | 190.02884945458632 | 357.9424099100043 | 134.98527094026184 | 168.29874684555267 |
| SE_341_1_14 | SE_341 | SE_341_1 | GCAGAACTGTCATCCCAGGT | 5466.57038947578 | 5828.456970384883 | 5854.059241697577 | 5806.438480445278 | 6211.902061056136 | 6674.919396536089 | 6009.280861677791 | 6962.528457946302 | 5683.993763358912 | 6721.44403130395 | 6574.3695872731005 | 5871.885243390696 |
| SE_341_1_16 | SE_341 | SE_341_1 | CCTTCCGGAAACAAACTGCC | 3385.718853434246 | 3611.804439466901 | 3491.8949862757477 | 3395.1138492787386 | 3246.479803464714 | 4903.315221855193 | 3262.918977902411 | 3559.2558919440366 | 3663.8272563066507 | 3542.2447449138713 | 3527.2238189172767 | 4060.062182522367 |
| SE_341_1_9 | SE_341 | SE_341_1 | GATGATTCAGAAGCCCCTTC | 66.11038734298623 | 114.93013573289807 | 108.62795488979984 | 53.35364807222242 | 115.60480188750533 | 208.7807364782865 | 74.25508228076067 | 89.80858160283758 | 114.5501008394469 | 184.43875704120384 | 77.46980767006332 | 268.1173139401563 |
| SE_341_2_11 | SE_341 | SE_341_2 | GGACCCAGCGAGTGGTCACA | 294.7873009392173 | 352.78554707576535 | 404.88601368016305 | 343.54544124552973 | 407.2682928880922 | 446.1497065822305 | 315.3150595400417 | 459.4417964103059 | 344.5382877961659 | 260.2554793032007 | 812.2591955709669 | 607.0361696567176 |
| SE_341_2_2 | SE_341 | SE_341_2 | ACCATGCCGACTCCACCAGG | 3705.433021732294 | 4301.38525386429 | 4247.3530361911735 | 3906.5280856783343 | 4151.166922822898 | 4561.122728457172 | 3932.290879042022 | 3785.195376186965 | 4079.40436632883 | 5036.708981809002 | 4001.43294465524 | 4345.5897116534425 |
| SE_341_2_31 | SE_341 | SE_341_2 | GCGGCCCTGCTCCATACCCA | 2360.465961197115 | 1902.8432907429383 | 2026.4051221260843 | 2203.115272835916 | 2056.4927601823197 | 2027.1656570920763 | 2438.579948524691 | 1925.6850602629488 | 2203.091474284246 | 2310.95201510125 | 1963.7422459396353 | 2440.9121697668775 |
| SE_341_2_43 | SE_341 | SE_341_2 | CCACTCAACAGCTGACCCAG | 847.5134902002497 | 943.426505494398 | 915.4374016622223 | 939.5447294669411 | 959.8380340201131 | 719.0373911907792 | 913.6603602371857 | 1100.3914630073994 | 935.0484975499038 | 1266.2850631643125 | 1419.1060223197962 | 834.5296481513957 |
| SE_341_2_5 | SE_341 | SE_341_2 | GTTTATGTTGGGCCTCCTGG | 14.089098941947885 | 25.984204600481302 | 20.738064115325425 | 6.506542447832002 | 21.211890254588134 | 9.529411208552496 | 34.43713960846872 | 8.508181415005666 | 0.0 | 21.141201399979888 | 25.823269223354437 | 11.606810127279495 |
| SE_341_3_17 | SE_341 | SE_341_3 | GCGAAGGCGATGGAGTTGGG | 10545.148669627148 | 9842.01718867461 | 9607.64884657148 | 10206.162483669279 | 9974.89139222007 | 9853.41118964328 | 10538.840880804193 | 9657.731259521986 | 10027.129757201817 | 10655.894512534689 | 8235.27531232067 | 12078.046618447042 |
| SE_341_3_26 | SE_341 | SE_341_3 | GCCCGGAATCATCAGATACA | 458.4376040341504 | 529.678016855965 | 525.3642909215774 | 538.7417146804897 | 550.4485521065621 | 351.72190460657396 | 439.07353000797616 | 570.9935082959357 | 502.5996672490461 | 422.8240279995977 | 503.5537498554116 | 291.3309341947153 |
| SE_341_3_30 | SE_341 | SE_341_3 | ACAGGGGGTGCACTCTCAGG | 4871.576903388904 | 4962.983078691928 | 5723.705695829817 | 5041.269088580235 | 5512.9702771674565 | 5217.785791737426 | 5003.070688742846 | 5075.602890796157 | 5632.490617245052 | 6054.402676787344 | 6304.399045392577 | 5535.287749699591 |
| SE_341_3_7 | SE_341 | SE_341_3 | ACACTTGCTGGTCAAAGACA | 1105.452378522065 | 1057.3572487426623 | 1005.3023461619658 | 1169.876332120194 | 1070.1398633439715 | 892.299413164461 | 1292.4688959303417 | 1068.249444328489 | 1273.3708884012935 | 1022.0677366473035 | 956.6347462288122 | 760.2460633368069 |
| SE_341_3_8 | SE_341 | SE_341_3 | TTTGACCAGCAAGTGTCCAG | 114.88034521895969 | 133.9185929409421 | 124.42838469195254 | 98.89944520704643 | 168.63452752397566 | 142.07485801841904 | 91.47365208499504 | 178.67180971511897 | 113.66211556162173 | 41.553395855132884 | 200.71722896334586 | 120.71082532370674 |
| SE_342_1_12 | SE_342 | SE_342_1 | TCTCTGCTAGGTGATAAGGG | 141.97476626116716 | 115.92952821753195 | 137.26623390620162 | 124.92561499837444 | 110.3018293238583 | 54.57753692170975 | 120.52998862964051 | 141.80302358342774 | 152.7334677859292 | 72.17168753786237 | 313.4005855743471 | 75.44426582731671 |
| SE_342_1_15 | SE_342 | SE_342_1 | ACTGGTAGAAAACTTCACTT | 629.6743450209017 | 842.4878645463745 | 714.9694485474099 | 740.4445305632819 | 844.2332321326078 | 1212.8341538157722 | 596.1929794716146 | 787.4794576333021 | 737.0277805948909 | 717.3428337096624 | 609.1943966782252 | 868.1893975205062 |
| SE_342_1_19 | SE_342 | SE_342_1 | ATAAGTTCGATGGTAGAAAG | 76.94815575986922 | 88.94593113241676 | 58.264084895438096 | 106.70729614444484 | 45.60556404736449 | 162.86630065526083 | 71.02660044246673 | 67.12009782948914 | 49.72717555820951 | 53.21750697236316 | 32.86597901154201 | 88.21175696732416 |
| SE_342_1_30 | SE_342 | SE_342_1 | GCTGATCATCCTGAATAGAG | 1145.552121664532 | 1224.255793676523 | 1238.3586857437183 | 1293.500638629002 | 1407.4089183919227 | 1180.7806797506412 | 1385.0187086281012 | 1329.167007721996 | 1418.112488686796 | 1423.7505632469213 | 1426.1487321079837 | 1306.926820331671 |
| SE_342_1_5 | SE_342 | SE_342_1 | GGAAGGCTCTGGATTTACCC | 778.1517723321986 | 878.4659939931947 | 905.5621330358769 | 792.4968701459379 | 978.9287352492424 | 760.6202764644629 | 892.1371479818928 | 813.9493553688753 | 1235.187521454811 | 606.5337780959746 | 1076.360812628001 | 978.4540937296614 |
| SE_342_2_17 | SE_342 | SE_342_2 | GCCTACCTGTGCTCACCCTG | 4154.116634191249 | 3577.825094989348 | 3700.2631542916365 | 3820.6417253669515 | 3692.990093323794 | 3519.817976395345 | 4200.254871620419 | 4208.713739956135 | 3620.315977693217 | 4380.602731464798 | 3750.2429622098834 | 3952.1188483386677 |
| SE_342_2_18 | SE_342 | SE_342_2 | CTAAAGTTATCCAACAGGCA | 30.34575156727237 | 47.97083926242702 | 39.50107450538176 | 41.641871666124814 | 36.06021343279983 | 4.331550549342044 | 50.57954879993843 | 35.92343264113503 | 32.855455279531284 | 9.477090282749604 | 18.780559435166865 | 2.321362025455899 |
| SE_342_2_19 | SE_342 | SE_342_2 | GTTTCTACATCAACCAAATG | 5.4188842084414945 | 0.9993924846338962 | 2.962580587903632 | 1.3013084895664004 | 20.15129574185873 | 0.0 | 1.0761606127646475 | 0.0 | 6.215896944776189 | 0.0 | 4.695139858791716 | 0.0 |
| SE_342_2_2 | SE_342 | SE_342_2 | AGATTAGTAGCAATACCAAA | 246.01734306324386 | 412.74909615379914 | 323.90881094413044 | 404.7069402551505 | 355.29916176435125 | 304.0748485638115 | 336.83827179533466 | 246.7372610351643 | 555.8787839185563 | 401.68282659961784 | 269.9705418805237 | 232.1362025455899 |
| SE_342_2_27 | SE_342 | SE_342_2 | TGTAGTACTCCTTTTCTATG | 204.8338230790885 | 254.84508358164354 | 227.13117840594512 | 239.44076208021767 | 272.5727897714575 | 232.17110944473353 | 281.9540805443376 | 313.8573588646534 | 329.442538073138 | 142.15635424124406 | 322.79086529193046 | 284.3668481183476 |
| SE_342_3_13 | SE_342 | SE_342_3 | TTAGGGAAAGCACCCAATGA | 930.9643070102487 | 1022.3785117804758 | 1052.703635568424 | 1059.26511050705 | 1258.9256866098058 | 967.6683927230125 | 1060.0182035731777 | 1176.019742251894 | 903.0810275481977 | 1028.6287991507456 | 1473.100130695901 | 900.6884658768888 |
| SE_342_3_29 | SE_342 | SE_342_3 | TTTCCAGGTGGCTCTCTCAA | 0.0 | 0.0 | 0.0 | 0.0 | 1.0605945127294067 | 0.0 | 0.0 | 0.0 | 0.0 | 0.0 | 2.347569929395858 | 0.0 |
| SE_342_3_32 | SE_342 | SE_342_3 | GTGAGGTGGCCCTTTCCAGG | 292.6197472558407 | 428.73937590794145 | 408.83612113070126 | 317.5192714542017 | 394.5411587353393 | 529.3154771295978 | 305.62961402515987 | 499.1466430136657 | 458.2004033577876 | 322.95007655831347 | 639.7128057603713 | 293.6522962201712 |
| SE_342_3_5 | SE_342 | SE_342_3 | GCAAAACGTCAAAATATGAA | 772.7328881237571 | 933.432580648059 | 951.9758955797005 | 918.7237936338787 | 975.7469517110542 | 1260.4812098585346 | 818.9582263138967 | 856.4902624439036 | 1204.1080367309303 | 1027.1707852610919 | 1221.9101482505441 | 926.2234481569036 |
| SE_342_3_6 | SE_342 | SE_342_3 | AGACCTTCCCATCTGAACCC | 908.2049933347945 | 998.3930921492623 | 978.6391208708332 | 939.5447294669411 | 1035.140244423901 | 923.4865771197237 | 1000.8293698711221 | 949.1349045184097 | 964.3520117181343 | 1269.930097888447 | 1218.3887933564504 | 673.1949873822107 |
| SE_343_1_10 | SE_343 | SE_343_1 | GTTCTCTGAGTAACATACAC | 15.172875783636185 | 2.9981774539016888 | 3.950107450538176 | 11.711776406097604 | 15.9089176909411 | 0.0 | 15.066248578705064 | 18.9070698111237 | 0.0 | 11.664111117230282 | 0.0 | 8.124767089095647 |
| SE_343_1_21 | SE_343 | SE_343_1 | GATCTAGCCCCTCAATTCCC | 480.1131408679164 | 441.7314782081821 | 426.61160465812304 | 494.4972260352322 | 453.9344514481861 | 523.2513063605189 | 425.08344204203576 | 368.6878613169122 | 275.2754361258026 | 386.3736807582531 | 476.5566956673592 | 539.7166709184964 |
| SE_343_1_28 | SE_343 | SE_343_1 | GGGCTACTCAGAGAAGACTT | 68.27794102636283 | 117.92831318679976 | 118.50322351614528 | 102.80337067574564 | 84.84756101835254 | 213.97859717349696 | 81.7882065701132 | 160.71009339455145 | 105.67024806119521 | 43.74041668961356 | 140.85419576375148 | 48.74860253457388 |
| SE_343_1_8 | SE_343 | SE_343_1 | TGTTACTCAGAGAACACAGG | 193.9960546622055 | 260.8414384894469 | 250.8318231091742 | 217.31851775758886 | 243.93673792776355 | 269.4224441690751 | 143.1293614976981 | 223.10342377125966 | 132.3098063959503 | 303.2668890479873 | 174.89395973999143 | 200.79781520193526 |
| SE_343_1_9 | SE_343 | SE_343_1 | GTATGTTACTCAGAGAACAC | 95.37236206857031 | 149.90887269508443 | 141.2163413567398 | 75.47589239485123 | 186.6646342403756 | 85.76470087697247 | 73.17892166799602 | 133.2948421684221 | 142.96562972985234 | 75.08771531716994 | 93.90279717583432 | 52.23064557275772 |
| SE_343_2_15 | SE_343 | SE_343_2 | AAAGAAACCAGTCCTCTGGG | 29.26197472558407 | 43.97326932389143 | 52.338923719630834 | 39.03925468699201 | 50.90853661101153 | 8.663101098684088 | 38.74178205952731 | 54.83050245225873 | 39.07135222430747 | 33.53431946203706 | 36.3873339056358 | 97.49720506914775 |
| SE_343_2_2 | SE_343 | SE_343_2 | TTAGGGACTCTATTTCACTA | 166.90163361999802 | 230.85966395043002 | 177.75483527421792 | 174.37533760189766 | 161.21036593486983 | 589.9571848203864 | 162.50025252746178 | 169.21827480955713 | 188.252878898936 | 188.0837917653383 | 355.6568443034725 | 263.4745898892445 |
| SE_343_2_3 | SE_343 | SE_343_2 | ACTCAAACTTTACATGGCAG | 482.28069455129304 | 593.6391358725343 | 605.3539667949755 | 569.9731184300834 | 646.9626527649381 | 572.6309826230182 | 483.19611513132674 | 508.6001779192275 | 648.2292528123739 | 996.5524935783623 | 433.1266519735358 | 525.7884987657611 |
| SE_343_2_4 | SE_343 | SE_343_2 | TCAGTGGTTGGCAGATAAGA | 1057.7661974877797 | 1164.292244598489 | 1239.3462126063528 | 1181.5881085262915 | 1046.8067840639244 | 1216.299394255246 | 1222.5184561006395 | 899.031169518932 | 1395.912856741167 | 1362.5139798814623 | 676.1001396660072 | 1462.4580760372162 |
| SE_343_2_6 | SE_343 | SE_343_2 | TTCTGTCTCAGCCTCCCTAG | 4.335107366753196 | 6.9957473924372735 | 5.925161175807264 | 15.615701874796805 | 9.545350614564661 | 0.8663101098684087 | 27.980175931880833 | 2.836060471668555 | 0.0 | 142.15635424124406 | 1.173784964697929 | 0.0 |
| SE_343_3_12 | SE_343 | SE_343_3 | CTTAGATGCCCATATTTCTA | 83.45081680999901 | 106.9349958558269 | 102.70279371399258 | 83.28374333224963 | 146.36204275665813 | 219.1764577967074 | 104.3875794381708 | 103.04353047062416 | 114.5501008394469 | 231.09520151012498 | 88.03387235234467 | 49.909283547301825 |
| SE_343_3_2 | SE_343 | SE_343_3 | ATATCTGTATTTGATAATAA | 2.167553683376598 | 5.9963549078033775 | 8.887741763710896 | 2.602616979132801 | 8.484756101835254 | 0.0 | 1.0761606127646475 | 0.0 | 17.759705556503395 | 1.4580138896537853 | 2.347569929395858 | 0.0 |
| SE_343_3_21 | SE_343 | SE_343_3 | ATACCAGTCTGTATCAGCCA | 252.52000411337366 | 254.84508358164354 | 229.10623213121423 | 258.9603894237137 | 265.1486281823517 | 171.52940175394494 | 229.2222105188699 | 229.72089820515296 | 139.41368861855165 | 258.7974654135469 | 207.75993875153344 | 141.60308355280984 |
| SE_343_3_23 | SE_343 | SE_343_3 | TCAGAATACCTCTCAGGGAG | 138.72343573610226 | 166.89854493386065 | 202.44300684008152 | 187.38842249756166 | 237.57317085138712 | 103.09090307434064 | 163.5764131402264 | 343.16331707189516 | 229.1002016788938 | 314.93100016521765 | 410.82473764427516 | 104.46129114551545 |
| SE_343_3_8 | SE_343 | SE_343_3 | GCATGTGCTTCAGATAGGGA | 2003.9033802816648 | 1867.864553780752 | 1941.4778119395135 | 1981.8928296096278 | 2087.2500010514723 | 1737.8180803960279 | 2050.0859673166533 | 1799.9530460189762 | 2061.901815110044 | 1955.925632970553 | 1619.823251283142 | 1669.0592963027914 |
| SE_344_1_10 | SE_344 | SE_344_1 | TCTCATTTAATATATGAGCA | 4.335107366753196 | 14.990887269508443 | 9.87526862634544 | 5.205233958265602 | 13.787728665482287 | 0.0 | 7.533124289352532 | 24.57919075446081 | 0.8879852778251698 | 65.61062503442034 | 1.173784964697929 | 1.1606810127279494 |
| SE_344_1_13 | SE_344 | SE_344_1 | ACAGTCATAACATCAAGTGA | 1851.0908456036145 | 1908.8396456507417 | 1877.2885658682683 | 1920.7313306000071 | 1804.071266152721 | 1886.8234192933942 | 1793.9597414786674 | 1580.6310362099414 | 1963.3354492714504 | 1612.5633619570865 | 1764.1988019409873 | 1783.9667165628582 |
| SE_344_1_15 | SE_344 | SE_344_1 | ATAAACTCTGGAGCCCCAGA | 0.0 | 0.9993924846338962 | 0.0 | 0.0 | 0.0 | 0.0 | 0.0 | 0.0 | 0.0 | 0.0 | 0.0 | 4.642724050911798 |
| SE_344_1_17 | SE_344 | SE_344_1 | GACTGTCTATAAGGAAAAAG | 60.69150313454474 | 116.92892070216585 | 106.65290116453076 | 123.62430650880805 | 163.33155496032865 | 233.90372966447035 | 63.4934761531142 | 51.04908849003399 | 117.21405667292241 | 252.23640291010486 | 23.47569929395858 | 179.90555697283216 |
| SE_344_1_9 | SE_344 | SE_344_1 | TTTTGACTTTCGAAACAGAG | 2.167553683376598 | 3.9975699385355847 | 0.987526862634544 | 0.0 | 5.3029725636470335 | 0.0 | 0.0 | 11.34424188667422 | 16.871720278678225 | 0.0 | 1.173784964697929 | 0.0 |
| SE_345_1_11 | SE_345 | SE_345_1 | GCCAGAATATGCCCAAACCC | 248.18489674662044 | 213.8699917116538 | 222.1935440927724 | 258.9603894237137 | 220.6036586477166 | 169.79678153420812 | 253.9739046124568 | 217.43130282792256 | 266.39558334755094 | 199.0188959377417 | 292.2724562097843 | 140.44240254008187 |

| | | | | | | | | | | | | | | | |
|---|---|---|---|---|---|---|---|---|---|---|---|---|---|---|---|
| SE_345_1_12 | SE_345 | SE_345_1 | GCACAGTCTGAGGCCCTGAG | 1176.9816500734926 | 1383.1591987333122 | 1363.7745972983053 | 1211.5182037863187 | 1375.5910830100406 | 857.6470087697246 | 1014.8194578370626 | 1098.500756026287 | 1135.7331703383923 | 909.8006671439621 | 1003.5861448167293 | 892.5636987877931 |
| SE_345_1_2 | SE_345 | SE_345_1 | ACTACACTCCAGCCTGGGGA | 448.68361245895574 | 559.6597913949819 | 486.85074327883024 | 524.4273212952594 | 471.964558164586 | 401.9678909789416 | 510.1001304504429 | 443.3707870708508 | 494.6077997486196 | 507.3888335995173 | 347.440349550587 | 608.1968506694456 |
| SE_345_1_25 | SE_345 | SE_345_1 | CTAGTGCTCCTCCCGCCCCC | 2287.852912803999 | 1976.7983346058465 | 2012.5797460492008 | 2089.901434243639 | 2148.764482789778 | 2420.470446972334 | 2108.1986404059444 | 2220.6353493164784 | 2337.177251235847 | 1664.322855039796 | 2131.593495891439 | 2328.3261115322666 |
| SE_345_1_7 | SE_345 | SE_345_1 | TAGGGACAAGGGAGAAGCCG | 2193.564327577117 | 2563.4417230859435 | 2539.919090696047 | 2039.1504031505494 | 2412.8525164594002 | 1694.5025749026074 | 1800.4167051552552 | 1556.9971989460366 | 2635.540304585104 | 1091.3233964058584 | 983.6318004168645 | 1211.7509772879791 |
| SE_345_2_10 | SE_345 | SE_345_2 | GACCTTCTGATCTCAAGCCA | 152.81253467805016 | 203.8760668653148 | 276.50752153767235 | 240.7420705697841 | 173.9375000876227 | 181.92512307236584 | 137.74855843387488 | 131.40413518730972 | 255.7397600136489 | 284.3127084824881 | 137.3328408696577 | 141.60308355280984 |
| SE_345_2_18 | SE_345 | SE_345_2 | TCTGGAATAGTAGTATCGGT | 433.51073667531955 | 470.7138602625651 | 511.5389148446938 | 490.59330056653295 | 553.6303356447503 | 456.5454279006514 | 414.3218359143893 | 716.5779458415882 | 419.1290511334801 | 344.09127795829335 | 419.0412323971607 | 221.69007343103834 |
| SE_345_2_2 | SE_345 | SE_345_2 | AGTTCCTCCAAGAGACCCTG | 1.083776841688299 | 4.996962423169481 | 1.975053725269088 | 1.3013084895664004 | 0.0 | 41.582885273683615 | 3.228481838293942 | 13.23494886778659 | 18.647690834328564 | 43.01140974478667 | 0.0 | 26.695663292742836 |
| SE_345_2_20 | SE_345 | SE_345_2 | CTATTCCAGATAAGAAGCCC | 5.4188842084414945 | 23.98541963121351 | 33.5759133295745 | 11.711776406097604 | 14.848323178211695 | 4.331550549342044 | 2.152321225529295 | 6.617474433893295 | 13.319779167377547 | 18.954180565499207 | 0.0 | 4.642724050911798 |
| SE_345_2_32 | SE_345 | SE_345_2 | TCAGGAACCTGGGGGGACGA | 870.272803875704 | 1089.3378082509469 | 1062.5789041947694 | 826.3308908746643 | 830.4455034671255 | 645.4010318519645 | 770.5309987394876 | 831.9110716894428 | 967.01596755161 | 966.6632088404597 | 524.6818792199742 | 1050.4163165187942 |
| SE_346_1_14 | SE_346 | SE_346_1 | CTTGAGGGAGAACTTCCCGT | 2193.564327577117 | 2459.5049046840186 | 2579.420165201429 | 2226.538825648111 | 2198.61242488806 | 2842.363470478249 | 2311.5929962184628 | 2230.0888842220406 | 2053.9099476096176 | 2022.9942718946272 | 2332.310724854785 | 1764.235139346483 |
| SE_346_1_24 | SE_346 | SE_346_1 | TGTGCCTTAGAGCAGTGTGG | 1637.5868077910197 | 1442.1233553267123 | 1458.5771761112214 | 1564.1728044588133 | 1547.4073940722044 | 1827.048021712474 | 1555.0520854449155 | 1454.8990219659688 | 1547.7583392492709 | 1759.093757867292 | 1602.2164768126731 | 1417.1915165408263 |
| SE_346_1_34 | SE_346 | SE_346_1 | GTGTGGCCCCACATGCTGGG | 534.3019829523314 | 453.7241880237889 | 441.4245075976412 | 549.152182597021 | 463.47980206275076 | 403.70051119867844 | 676.9050254289633 | 343.16331707189516 | 348.09022890746655 | 364.50347241344633 | 590.4138372430583 | 369.0965620474879 |
| SE_346_1_42 | SE_346 | SE_346_1 | AGCCTGGGCCACTAAACCCG | 655.6849892214209 | 1001.391269603164 | 911.4872942116841 | 664.9686381684306 | 920.596037049125 | 915.689786130908 | 582.2028915056743 | 989.7851046123257 | 747.683603928793 | 561.3353475167073 | 961.3298860876039 | 544.3593949694083 |
| SE_346_2_14 | SE_346 | SE_346_2 | TTAGCTACACCTAACTACAA | 153.89631151973845 | 141.91373281801324 | 132.3285995930289 | 110.61122161314404 | 159.089176909411 | 292.81281713552215 | 95.77829453605362 | 125.7320142439726 | 194.4687758437122 | 26.244250013768134 | 100.9455069640219 | 124.19286836189059 |
| SE_346_2_22 | SE_346 | SE_346_2 | CAGGGAAGACACTAGCTCCA | 14303.686756602168 | 12916.148471408475 | 13408.639740851839 | 13874.551115756962 | 12913.798786993257 | 14385.945684474795 | 14247.290352391168 | 13842.811162214217 | 13543.551457389489 | 12853.85045118777 | 13552.52120240229 | 13542.826056509713 |
| SE_346_2_33 | SE_346 | SE_346_2 | GGTAACATCACAGAGACGGC | 986.236925936352 | 1071.3487435275367 | 1095.1672906617093 | 882.2871559260195 | 960.8986285328425 | 816.064123496041 | 959.9352665860655 | 1070.1401513096014 | 1145.501008394469 | 1008.2166046955925 | 1173.7849646979291 | 1210.5902962752511 |
| SE_346_2_7 | SE_346 | SE_346_2 | AGTTCAAAATGACCAGGCCA | 280.6982019972694 | 311.8104552057756 | 306.13332741670865 | 320.1218884333345 | 299.0876525896927 | 245.16576109275968 | 326.0766656676882 | 230.66625169570915 | 408.4732277995781 | 402.41183354444473 | 200.71722896334586 | 367.93588103476 |
| SE_346_2_9 | SE_346 | SE_346_2 | AAAAGGATGTCAGTTAACAA | 110.54523785220648 | 157.9040125721556 | 150.1040831204507 | 109.30991312357764 | 207.87652449496372 | 54.57753692170975 | 128.06311291899306 | 233.5023121673777 | 174.93309973155846 | 33.53431946203706 | 298.141381033274 | 44.10587848366208 |
| SE_346_3_14 | SE_346 | SE_346_3 | CAGAGGAGAAGGGAAATGGG | 5758.106359889932 | 6023.338504888492 | 5810.608059741657 | 5773.905768206118 | 6192.811359827006 | 5579.903417662421 | 5733.783744810042 | 5674.011650318223 | 5406.9423566774585 | 5948.696669787444 | 4743.265042344331 | 4668.259033191813 |
| SE_346_3_16 | SE_346 | SE_346_3 | CATCCACCTAAAAAGCCCAG | 470.3591492927217 | 540.6713341869379 | 570.7905266027665 | 467.1697477543378 | 610.9024393321383 | 504.1924839434139 | 460.5967422632691 | 485.9116941458791 | 764.5553242074711 | 534.3620905581123 | 346.26656458588906 | 784.6203646040938 |
| SE_346_3_17 | SE_346 | SE_346_3 | TCCAGTCTCAGAAAGCAGCG | 317.54661461467157 | 231.8590564350639 | 272.55741408713413 | 312.3140374959361 | 283.1787348987516 | 146.40640856776108 | 319.61970199110027 | 349.78079150578844 | 333.88246446226384 | 371.79354186171525 | 431.95286700883787 | 146.24580760372163 |
| SE_346_3_20 | SE_346 | SE_346_3 | GAAAGGGGTGATGGTGTCAG | 6486.404397504469 | 7187.630749486982 | 6933.426102557134 | 6853.991814546231 | 6741.138722908109 | 6075.43280050715 | 6903.5703308852135 | 6919.987550871274 | 7158.937309826519 | 7822.244517992558 | 6812.64793510678 | 6827.1257168657985 |
| SE_346_3_54 | SE_346 | SE_346_3 | AGCAGAGACTGGCCATCAAG | 1091.363279580117 | 1106.327480489723 | 1083.3169683100948 | 1136.0423113914676 | 1080.7458084712655 | 888.8341727249874 | 1103.0646280837636 | 1070.1401513096014 | 895.9771453255963 | 797.5335976406205 | 1071.665672769209 | 818.2801139732044 |
| SE_347_1_21 | SE_347 | SE_347_1 | ACGGGGAAAGTGAGTCAGAA | 8799.184177667299 | 10771.452199384134 | 10630.726676260867 | 9860.014425444617 | 10105.344517285788 | 10120.23470348275 | 9698.359442235003 | 10041.544776687797 | 10940.866608083918 | 8855.247358812265 | 9641.469700028789 | 9085.810967634388 |
| SE_347_1_31 | SE_347 | SE_347_1 | TTTGGCCCTGTTGCCTAGCG | 332.71949039830776 | 353.78493956039927 | 290.3328976145559 | 291.4931016628737 | 360.6021343279983 | 439.2192257032832 | 304.55345341239524 | 312.91200537409725 | 118.10204195074758 | 314.93100016521765 | 233.58320797488787 | 220.52939241831038 |
| SE_347_1_37 | SE_347 | SE_347_1 | CTAACCTGGTTAGTGCCGGA | 261.19021884688004 | 311.8104552057756 | 321.93375721886133 | 269.37085734024487 | 317.11775930609264 | 267.6898239493383 | 229.2222105188699 | 455.6603824480812 | 271.72349501450196 | 295.9768195997184 | 420.2150173618586 | 174.10215190919243 |
| SE_347_1_47 | SE_347 | SE_347_1 | GCACACAGAGGACTCAGTTG | 6753.01350055979 | 6632.967920515169 | 6733.945676304956 | 6723.860965589591 | 7204.61852497086 | 5583.368658101895 | 6480.639210068707 | 6948.34815558796 | 6208.793062553587 | 5733.639621063511 | 6574.3695872731005 | 5618.856782616003 |
| SE_347_1_6 | SE_347 | SE_347_1 | CTAACCAGTAGGACAGGCAT | 1943.21187714712 | 1675.9811967310438 | 1624.481689033825 | 1961.0718937765655 | 1613.1642538614276 | 1405.154998206559 | 1832.7015235381946 | 1433.1558916831764 | 1490.92728146846 | 1275.0331465022352 | 1363.9381289789935 | 1599.4184355391144 |
| SE_347_2_17 | SE_347 | SE_347_2 | GAGCAAAGCCTGACCCCGCG | 1786.0642351023166 | 2134.702347178002 | 2121.2077009390005 | 1970.1810532035302 | 1909.0701229129322 | 1756.0105927032646 | 1690.6483226532612 | 1885.034860169033 | 2088.5413734447993 | 1718.269368956986 | 1647.9940904358923 | 1563.4373241445478 |
| SE_347_2_18 | SE_347 | SE_347_2 | GGAGCAAAGCCTGACCCCGC | 2475.3463064160746 | 2452.5091572915812 | 2402.652856789846 | 2233.045368095943 | 2369.3681414374946 | 2144.1175219243114 | 2440.7322697502204 | 2035.3460651674663 | 2367.368750681903 | 2237.3223136737333 | 2122.2032161738557 | 2219.2220963358395 |
| SE_347_2_21 | SE_347 | SE_347_2 | CTTTCCAACGTACAGCCCCA | 223.25802938778958 | 209.8724217731182 | 151.09160998308525 | 144.44524234187045 | 191.9676068040226 | 105.68983340394587 | 146.35784333599204 | 172.99968877178185 | 161.6133205641809 | 25.515243068941242 | 91.55522724643846 | 89.37243798005211 |
| SE_347_2_32 | SE_347 | SE_347_2 | TGAGGCCAAAACAGGCACCA | 11696.119675500122 | 10924.35924953312 | 10194.239802976397 | 10463.821564603426 | 10413.977520490045 | 9845.614398654465 | 11077.99734779928 | 9450.69884509018 | 10069.753050537425 | 8527.92324058499 | 8289.269420696775 | 8570.46859798318 |
| SE_347_2_33 | SE_347 | SE_347_2 | GAAAACCTTGGTGCCTGTTT | 78.03193260155751 | 151.90765766435223 | 115.54064292824165 | 115.81645557140963 | 177.11928362581094 | 116.08555472236677 | 107.61606127646475 | 172.99968877178185 | 220.2203489006421 | 58.320555586151414 | 98.59793703462604 | 134.63899747644214 |
| SE_347_3_22 | SE_347 | SE_347_3 | GATCTGGAGGTGCGGGTGGG | 1879.2690434875103 | 2147.694449478243 | 2240.6984513177804 | 1903.814320235644 | 2038.4626534659199 | 2234.213773350626 | 1806.8736688318431 | 2287.7554471459675 | 1847.0093778763533 | 1753.9907092535038 | 2209.0633035615024 | 1905.838222899293 |
| SE_347_3_26 | SE_347 | SE_347_3 | CGGTTGAGCCCACTTTCCCG | 394.4947703745408 | 549.665866548643 | 544.1273013116338 | 420.3226421299473 | 497.41882647009174 | 351.72190460657396 | 321.7720232166296 | 365.8518008452436 | 626.0296208667447 | 594.8696669787444 | 253.53755237475266 | 354.0077088820246 |
| SE_347_3_36 | SE_347 | SE_347_3 | GGCTTCACGTTTTCACCAAG | 42.26729682584366 | 97.94046349412183 | 83.93978332393624 | 55.95626505135522 | 119.84717993842297 | 94.42780197565655 | 36.58946083399801 | 83.19110716894428 | 140.30167389637683 | 117.37011811712972 | 125.5949912226784 | 97.49720506914775 |
| SE_347_3_38 | SE_347 | SE_347_3 | TGGCGCTCAAAGCTGCCGAC | 88.86970101844051 | 98.93985597875572 | 83.93978332393624 | 115.81645557140963 | 69.99923784014085 | 117.81817494210358 | 87.16900963393644 | 56.7212094333711 | 62.15896944776188 | 147.2594028550323 | 59.86303319959438 | 81.24767089095646 |
| SE_347_3_39 | SE_347 | SE_347_3 | CTGGCGCTCAAAGCTGCCGA | 24.926867358830876 | 52.9678016855965 | 36.53849391747813 | 13.013084895664004 | 46.6661585600939 | 3.465240439473635 | 10.761606127646475 | 55.775855942814914 | 23.087617223454416 | 16.038152786191638 | 96.25036710523018 | 70.80154177640492 |
| SE_348_1_0 | SE_348 | SE_348_1 | ATACAAACACTACACAACAT | 883.2781259759636 | 871.4702466007575 | 851.2481555909769 | 946.0512719147731 | 772.1128052670081 | 655.7967531703854 | 794.2065322203098 | 705.2337039549141 | 978.5597761633371 | 626.2169656063008 | 718.3563983951326 | 855.4219063804987 |
| SE_348_1_1 | SE_348 | SE_348_1 | TTTCCTTTACAAAGTGTAAG | 1.083776841688299 | 0.0 | 1.975053725269088 | 0.0 | 0.0 | 0.0 | 0.0 | 0.0 | 0.0 | 0.0 | 0.0 | 0.0 |
| SE_348_1_4 | SE_348 | SE_348_1 | AAGTGGGATGAGCCTCACTA | 778.1517723321986 | 870.4708541161235 | 856.1857899041497 | 789.894253166805 | 864.3845278744665 | 855.0480784401194 | 1048.1804368327666 | 880.1240997078082 | 918.1767772712255 | 984.1593755163051 | 920.2474123231764 | 605.8754886439896 |
| SE_348_1_5 | SE_348 | SE_348_1 | AACATGTTGGAAACAGTCAG | 236.26335148804915 | 271.8347558204198 | 260.7070971355196 | 301.9035695794049 | 296.9664635642339 | 254.69517230131217 | 246.44078032310426 | 231.61160518626534 | 393.37747807655023 | 402.41183354444473 | 159.63475519891836 | 337.7581747038333 |
| SE_348_1_8 | SE_348 | SE_348_1 | TGTGGTGGTTGGAAATGTCA | 1452.2609678623205 | 1297.2114450547972 | 1343.0365331829798 | 1410.618402689978 | 1517.710747715781 | 1274.3421716164291 | 1482.9493243896843 | 1406.6859939476033 | 1394.1368861855165 | 1868.4447995913258 | 1305.248880744097 | 1221.0364253898028 |
| SE_348_2_17 | SE_348 | SE_348_2 | CCATGGTCCACTTATTCGGG | 596.0772629285644 | 586.6433884800971 | 681.3935352178354 | 651.9555532727666 | 727.567835732373 | 668.7914048184116 | 577.8982490546157 | 837.58319263278 | 840.9220581004358 | 980.5143407921706 | 1003.5861448167293 | 773.0135544768143 |
| SE_348_2_25 | SE_348 | SE_348_2 | GGTCTGGGCCCCCTTCACGG | 1.083776841688299 | 1.9987849692677924 | 0.987526862634544 | 0.0 | 0.0 | 0.0 | 1.0761606127646475 | 0.0 | 8.879852778251697 | 0.0 | 0.0 | 0.0 |
| SE_348_2_3 | SE_348 | SE_348_2 | GCTGTTGGAGATATAAAAGG | 140.89098941947887 | 186.8863946265386 | 192.56773821373608 | 179.58057156016326 | 241.81554890230473 | 194.05346461052355 | 181.87114355722542 | 246.7372610351643 | 174.04511445373328 | 238.3852709583939 | 265.27540202173196 | 320.34795951291403 |
| SE_348_2_30 | SE_348 | SE_348_2 | CTTGGGACTAGAGTAGGTGG | 8884.802548160675 | 8395.896263409362 | 8773.18864764529 | 9638.791982218328 | 10055.496575187506 | 11631.079535093255 | 10520.546150387194 | 12843.57252269633 | 9087.641333262787 | 16390.992147487854 | 15085.484366297784 | 16643.00504150607 |
| SE_348_2_4 | SE_348 | SE_348_2 | TTAATGAGAGTGACACTGAT | 176.65562519519273 | 137.91616287947767 | 147.14150253254707 | 162.66356119580004 | 151.66501532030517 | 116.08555472236677 | 235.6791741954578 | 277.93392622351837 | 106.55823333902038 | 396.5797779858296 | 316.9219404684408 | 152.0492126673614 |
| SE_348_3_12 | SE_348 | SE_348_3 | AGGGTTTTAGGAGATCTGGT | 162.56652625324483 | 226.86209401189444 | 241.94408134546327 | 227.72898567412008 | 163.33155496032865 | 220.04276790657582 | 138.82471904663953 | 183.3985771678999 | 250.41184834669787 | 263.17150708250824 | 213.62886357502308 | 120.71082532370674 |
| SE_348_3_17 | SE_348 | SE_348_3 | CTAGGTCACAAAGTTCTGGA | 325.13305250648966 | 447.7278331159855 | 412.7862285812394 | 360.4624516098929 | 390.2987806844217 | 272.02137449868036 | 261.50702890180935 | 352.61685197745703 | 348.97821418529173 | 298.1638404341991 | 190.1531642810645 | 170.62010887100857 |
| SE_348_3_3 | SE_348 | SE_348_3 | TCTAGGACTGAGGTGAGAGA | 255.77133463843853 | 330.79891241381966 | 323.90881094413044 | 300.6022610898385 | 238.63376536411653 | 165.46523098486605 | 171.10953742957895 | 252.4093819785014 | 159.83735000853056 | 111.53806255851458 | 82.16494752885504 | 22.05293924183104 |
| SE_348_3_4 | SE_348 | SE_348_3 | TAGTTAGGTGGCACACCAAG | 1408.9098941947886 | 1471.1057373810952 | 1361.7995435730363 | 1529.0374752405205 | 1475.2869672066047 | 1123.6042124993262 | 1536.7573550279167 | 1246.921254043608 | 1315.9941817369015 | 2204.5170011565233 | 1034.1045538988756 | 1425.3162836299218 |
| SE_348_3_7 | SE_348 | SE_348_3 | ACAACCAAACACAATCCCAA | 2909.9408199330824 | 3107.111234726783 | 2929.992201436692 | 3041.1579401166778 | 3355.721038275843 | 2699.4223023499617 | 2749.5903656136743 | 3183.005202702675 | 2986.294489326046 | 2633.902091659563 | 2964.9808208269687 | 3037.5022103090437 |
| SE_349_1_1 | SE_349 | SE_349_1 | GTTCTTCAATGAGAACACAT | 1.083776841688299 | 1.9987849692677924 | 5.925161175807264 | 14.314393385230405 | 9.545350614564661 | 0.0 | 0.0 | 3.78141396222474 | 0.0 | 0.0 | 0.0 | 0.0 |
| SE_349_1_12 | SE_349 | SE_349_1 | GCCCTATCAGAGTTTTCCAA | 22.759313675454276 | 43.97326932389143 | 47.40128940645811 | 28.62878677046081 | 49.84794209828212 | 89.2299413164461 | 18.294730416999005 | 63.33868386726439 | 32.855455279531284 | 49.5724722482287 | 38.73490383503166 | 9.285448101823595 |
| SE_349_1_23 | SE_349 | SE_349_1 | TTAGCACACTATATAAAGAG | 53.105065242726646 | 94.94228604022014 | 76.03956842285989 | 49.44972260352321 | 62.575076251035 | 35.51871450460476 | 71.02660044246673 | 28.36060471668555 | 139.41368861855165 | 18.225173620672315 | 29.344624117448227 | 104.46129114551545 |
| SE_349_1_6 | SE_349 | SE_349_1 | CCACTGCCTGCAAGTATGCA | 257.9388883218151 | 206.8742443192165 | 268.607306636596 | 253.7551554654481 | 217.42187510952837 | 58.90908747105179 | 271.19247441669114 | 280.76998669518696 | 129.64585056247478 | 173.50365286880046 | 280.53460656280504 | 171.7807898837365 |
| SE_349_1_7 | SE_349 | SE_349_1 | TTGGCTTGAATGGTTAGCCA | 329.4681598732429 | 238.8548038275012 | 289.3453707519214 | 303.2048780689713 | 299.0876525896927 | 336.1283226289426 | 369.1230901782741 | 199.46958650735505 | 243.3079661240965 | 417.72097938580947 | 201.8910139280438 | 482.843301294827 |
| SE_35_1_10 | SE_35 | SE_35_1 | TTCTGTGTCATGAACCCCTG | 15.172875783636185 | 15.990279754142339 | 11.850322351614528 | 18.218318853929606 | 10.605945127294067 | 0.0 | 32.28481838293943 | 10.398888396118036 | 0.0 | 2.187020834480678 | 24.64948425865651 | 0.0 |
| SE_35_1_13 | SE_35 | SE_35_1 | GCAGATGTTGGCAGTTACCA | 4800.047631837476 | 5396.71941702304 | 5172.665706479742 | 4864.291133999205 | 5406.910825894515 | 4958.759068886771 | 4864.245969696207 | 4499.882615047441 | 5456.669532235668 | 3736.8895991826516 | 4216.235593194961 | 4595.136129389952 |
| SE_35_1_15 | SE_35 | SE_35_1 | GTTTCTAGTGTCCATGCCAG | 17.340429467012783 | 39.975699385355846 | 51.35139685699629 | 31.23140374959361 | 32.87842989461161 | 130.81282659012973 | 16.142409191469714 | 38.75949311280358 | 57.71904305863604 | 13.851131951710961 | 22.30191432926065 | 306.41978736017865 |
| SE_35_1_18 | SE_35 | SE_35_1 | TCTTGGACCAGGAGGCAGCA | 762.9788965485625 | 665.5953947661749 | 693.2438575694499 | 720.9249032197858 | 705.2953509650555 | 939.9464692072235 | 711.342165037432 | 474.56745225920486 | 597.6140919763393 | 325.13709739279415 | 315.7481555037429 | 525.7884987657611 |
| SE_35_1_9 | SE_35 | SE_35_1 | AGGCCTCCTGCCAGGCCACA | 59.60772629285644 | 68.95808143973883 | 86.90236391183987 | 75.47589239485123 | 43.48437502190568 | 50.2459863723677 | 65.6457973786435 | 60.50262339559584 | 61.27098416993672 | 46.65644446892113 | 55.167893340802664 | 52.23064557275772 |
| SE_35_2_20 | SE_35 | SE_35_2 | GGTTAGACATGAATTTGCGG | 87.78592417675222 | 149.90887269508443 | 170.8421472357761 | 115.81645557140963 | 217.42187510952837 | 309.27270922302193 | 51.655709412703075 | 116.27847933841076 | 178.48504084285912 | 48.114458358574915 | 96.25036710523018 | 152.0492126673614 |
| SE_35_2_22 | SE_35 | SE_35_2 | GTGTACTCACCTCCTCTGCC | 617.7527997623304 | 616.625163019114 | 620.1668697344936 | 584.2875118153138 | 622.5689789721617 | 510.2566547124927 | 619.868512952437 | 692.9441085776837 | 545.2229605846543 | 742.8580767786036 | 665.5360749837257 | 796.2271747313733 |
| SE_35_2_23 | SE_35 | SE_35_2 | CCAGAGAGGCCTGGCAGAGG | 34.680858934025565 | 36.977521931454156 | 22.713117840594514 | 36.43663770785921 | 14.848323178211695 | 20.791442636841808 | 35.513300221233365 | 58.61191641448347 | 25.751573056929924 | 1.4580138896537853 | 51.646538446708874 | 1.1606810127279494 |
| SE_35_2_24 | SE_35 | SE_35_2 | TTGCCAGAGAGGCCTGGCAG | 39.01596630077876 | 49.96962423169481 | 80.97720273603261 | 19.519627343496005 | 48.78734758555271 | 49.379676262499295 | 16.142409191469714 | 5.67212094333711 | 21.311646667804077 | 18.225173620672315 | 15.259204541073078 | 8.124767089095647 |
| SE_35_2_25 | SE_35 | SE_35_2 | GCTTGTTTGCCAGAGAGGCC | 360.89768828220355 | 531.6768018252328 | 540.1771938610956 | 442.44488645257616 | 494.23704293190355 | 474.73794020788796 | 363.74228711445085 | 649.4578480120991 | 468.85622669168964 | 508.84684748917107 | 550.5051484433287 | 761.4067443495348 |
| SE_35_3_16 | SE_35 | SE_35_3 | GCAATGTGTGGCCAAATCCC | 4489.003678272934 | 4018.5571807128968 | 4075.5233620927634 | 4371.095216453539 | 4260.408157634027 | 4670.277802300591 | 4402.573066820173 | 3732.2555807158183 | 4138.899379943116 | 3872.484890920454 | 3376.9793434359417 | 4721.650359777298 |
| SE_35_3_20 | SE_35 | SE_35_3 | TCTCCATCCAGAAGCCCCTG | 65.02661050129794 | 76.95322131681 | 105.66537430189621 | 91.09159426964803 | 98.63528968383483 | 115.21924461249836 | 49.50338818717378 | 70.90151179171387 | 144.74160028550267 | 243.48831957218215 | 84.5125174582509 | 210.08326330375886 |
| SE_35_3_32 | SE_35 | SE_35_3 | TGCTCAATGCCTCTGAGCAC | 361.98146512389184 | 327.800734959918 | 292.307951339825 | 409.91217421341617 | 375.45045750621 | 351.72190460657396 | 459.5205816505045 | 325.20160075132765 | 359.6340375191938 | 266.0875348618158 | 233.58320797488787 | 395.79222534023074 |
| SE_35_3_5 | SE_35 | SE_35_3 | TAAGTGTTGTATTTGTAGGA | 275.27931778882794 | 432.73694584647706 | 411.7987017186049 | 350.0519836933617 | 404.08650934990396 | 397.6363404295996 | 265.8116713528679 | 419.73694980694614 | 432.4488303008577 | 401.68282659961784 | 308.70544571555536 | 501.41419749847415 |
| SE_35_3_6 | SE_35 | SE_35_3 | GCTCTGAGTGAATTCCAGGG | 52.02128840103835 | 94.94228604022014 | 135.29118018093254 | 74.17458390528482 | 97.57469517110542 | 12.994651648026132 | 27.980175931880833 | 37.8141396222474 | 79.91867500426528 | 62.69459725511277 | 22.30191432926065 | 53.39132658548567 |
| SE_35_4_16 | SE_35 | SE_35_4 | AGAGAGTCCTGGCTAGCAAG | 1382.8992499942694 | 1343.1834993479565 | 1241.321266331622 | 1367.6752225342868 | 1308.773628708088 | 1717.8929478690545 | 1319.3729112494577 | 1518.2377058332331 | 1043.3827014445744 | 1340.6437715366555 | 1320.5080852851702 | 1344.0686127389654 |
| SE_35_4_20 | SE_35 | SE_35_4 | TTAAGAAGTGATGGGGGCAT | 666.5227576383038 | 644.608152588863 | 659.6679442398754 | 581.684894836181 | 711.6589180414319 | 466.0748391092039 | 637.0870827566713 | 621.0972432954136 | 645.5652969788985 | 704.9497156476052 | 532.8983739728598 | 481.68262028209904 |
| SE_35_4_23 | SE_35 | SE_35_4 | GGGCCTTTAAGAAGTGATGG | 15.172875783636185 | 27.982989569749094 | 30.613332741670867 | 28.62878677046081 | 49.84794209828212 | 40.71657516381521 | 5.380803063823238 | 17.01636283001133 | 53.27911666951019 | 8.748083337922711 | 11.73784964697929 | 45.26655949639003 |
| SE_35_4_3 | SE_35 | SE_35_4 | CATTCACTACATATTATCCA | 458.4376040341504 | 609.6294156266766 | 543.1397744489992 | 540.0430231700561 | 615.144817383056 | 517.18713559144 | 390.64630243356703 | 630.5507782009754 | 540.7830341955284 | 499.36975720642147 | 530.5508040434639 | 649.9813671276517 |
| SE_35_4_4 | SE_35 | SE_35_4 | GAAAGTTTATTAAGGCACAG | 47.686181034285156 | 52.9678016855965 | 66.16429979651446 | 91.09159426964803 | 80.60518296743491 | 200.9839454894708 | 68.87427921693744 | 86.97252113116902 | 89.68651306034215 | 80.91977087578509 | 43.43004369382337 | 105.6219721582434 |
| SE_350_1_16 | SE_350 | SE_350_1 | GAGTCTGTCCTTAAACTCAC | 282.86575568064603 | 327.800734959918 | 326.87139153203407 | 297.9996441107057 | 346.814405662516 | 248.6310015322333 | 289.48720483369016 | 357.34361943023794 | 366.7379197417951 | 362.31645157896565 | 343.9189946564932 | 286.68821014380353 |
| SE_350_1_2 | SE_350 | SE_350_1 | CTACTCCAGCCAGGCGCCAA | 34.680858934025565 | 56.96537162413208 | 45.42623568118903 | 28.62878677046081 | 40.30259148371746 | 12.128341538157722 | 10.761606127646475 | 15.12565584889896 | 17.759705556503395 | 61.965590310285876 | 39.90868879972959 | 0.0 |
| SE_350_1_20 | SE_350 | SE_350_1 | TCCAACCTGGAACAGCCAAC | 1359.0561594771268 | 1238.2472884613974 | 1276.8722333864655 | 1260.967926389842 | 1200.5929884096884 | 908.7593052519608 | 1309.6874657345759 | 1262.046909892507 | 1207.659977842231 | 1795.5441051086366 | 1052.8851133340424 | 828.7262430877558 |

| | | | | | | | | | | | | | | | |
|---|---|---|---|---|---|---|---|---|---|---|---|---|---|---|---|
| SE_350_1_4 | SE_350 | SE_350_1 | GCTTCATGATAACAGTTTGT | 435.6782903586962 | 479.7083926242702 | 521.4141834710392 | 528.3312467639586 | 478.32812524096244 | 732.8983529486737 | 400.3317479484489 | 348.8354380152323 | 395.15344863220054 | 532.9040766684585 | 528.2032341140681 | 876.3141646096018 |
| SE_350_1_6 | SE_350 | SE_350_1 | TGCTTAAGCCGAGGAGGCAG | 693.6171786805113 | 677.5881045817816 | 675.4683740420281 | 672.776489105829 | 714.8407015796201 | 626.3422094348595 | 600.4976219226733 | 645.6764340498744 | 522.1353433611998 | 370.33552797206147 | 862.7319490529778 | 1380.049724133532 |
| SE_351_1_17 | SE_351 | SE_351_1 | CAAGGAGTGCCAGGAAGCCA | 3644.7415185977493 | 3548.842712934965 | 3484.9822982373057 | 3618.9389094841595 | 3484.052974316101 | 3502.4917741979766 | 3522.273685578691 | 3577.217608264604 | 3254.466043229247 | 3099.7375294039475 | 3574.175217505194 | 3499.4532533747674 |
| SE_351_1_24 | SE_351 | SE_351_1 | GTGTCTACTAAAAATAGCTG | 19.50798315038938 | 46.97144677779312 | 53.32645058226538 | 32.53271223916001 | 37.120807945529236 | 26.85561340592067 | 19.370891029763655 | 40.650200093915956 | 96.7903952829435 | 57.59154864132452 | 28.170839152750297 | 3.482043038183848 |
| SE_351_1_3 | SE_351 | SE_351_1 | ATCCTAGCCATTCTCCCAGG | 125.71811363584267 | 167.89793741849456 | 169.85462037314159 | 171.77272062276487 | 141.0590701930111 | 172.39571186381335 | 187.25194662104866 | 207.0324144318045 | 126.09390945117411 | 114.45409033782215 | 240.62591776307545 | 51.069964560029774 |
| SE_351_1_33 | SE_351 | SE_351_1 | AGTCTTAGCCCAGCTGCTGA | 203.7500462374002 | 321.8043800521146 | 332.79655270784133 | 342.24413275596334 | 339.39024407341014 | 334.3957024092058 | 287.33488360816085 | 290.2235216007488 | 314.3467883501101 | 231.82420845495187 | 377.9587586327331 | 192.6730481128396 |
| SE_351_1_36 | SE_351 | SE_351_1 | CTCCACTTTATCAATCCCGG | 32.51330525064897 | 41.97448435462364 | 49.376343131727204 | 31.23140374959361 | 19.090701229129323 | 22.524062856578627 | 9.685445514881827 | 56.7212094333711 | 44.39926389125849 | 2.187020834480678 | 123.24742129328254 | 23.21362025455899 |
| SE_351_2_16 | SE_351 | SE_351_2 | GCTGCATGTTGGGATCACTT | 3384.6350765925577 | 3125.1002994501932 | 3165.0235947437136 | 3280.5987021968954 | 3176.480565624573 | 3355.219055520347 | 3188.6638956216507 | 3173.551667797113 | 3067.1011496081364 | 3052.3520779901996 | 2790.0868610869775 | 3425.1696685601787 |
| SE_351_2_17 | SE_351 | SE_351_2 | CAGACGAGATCACATGAGCA | 6495.074612237975 | 5805.470943238303 | 5642.728493093785 | 5758.290066331322 | 5754.785826069761 | 5669.999669088736 | 5839.247484860977 | 5830.940329750549 | 5575.659559464241 | 5716.143454387665 | 6286.7922709221075 | 5624.660187679643 |
| SE_351_2_4 | SE_351 | SE_351_2 | GTGCAGGCTGTGGCTTGCCG | 2425.492571698413 | 2458.5055121993846 | 2521.156080305991 | 2266.8793888246696 | 2550.729803114223 | 2158.8447937920746 | 2160.930510431412 | 2273.575144787625 | 2655.963965975083 | 1998.9370427153397 | 2440.2989416069945 | 2199.4905191194644 |
| SE_351_2_5 | SE_351 | SE_351_2 | GCAAGCCACAGCCTGCACCG | 3308.7706976743766 | 3075.1306752184987 | 3092.9341337713918 | 3365.1837540187116 | 3342.99390412309 | 3038.149555308509 | 3513.664400676574 | 3260.524188928282 | 2972.9747101586686 | 3029.023855755739 | 3688.032359080893 | 3095.536260945441 |
| SE_351_2_7 | SE_351 | SE_351_2 | AGACTGGCTTCATCTTACTG | 913.623877543236 | 1004.3894470570657 | 969.7513791071223 | 1022.8284727991908 | 897.2629577690781 | 869.7753503078824 | 1004.0578517094161 | 749.6653180110548 | 905.7449833816731 | 929.4838546542882 | 780.5670015241228 | 739.3538051077038 |
| SE_352_1_10 | SE_352 | SE_352_1 | TGAATTGTAGGATACCCCGT | 365.2327956489567 | 355.78372452966704 | 403.8984868175285 | 355.25721765162734 | 373.3292684807512 | 144.67378834802426 | 297.0203291230427 | 432.97189867473276 | 486.615932248193 | 452.71331273750036 | 307.5316607508574 | 359.8111139456643 |
| SE_352_1_12 | SE_352 | SE_352_1 | TGTACCCAAGAAGTGTGTGT | 1106.536155363753 | 1031.3730441421808 | 1060.6038504695002 | 943.4486549356403 | 1068.0186743185127 | 946.0106399763023 | 1014.8194578370626 | 979.3862162162077 | 1094.8858475584343 | 948.4380352197874 | 879.1649385587489 | 761.4067443495348 |
| SE_352_1_22 | SE_352 | SE_352_1 | CTCTCTCTATTGAGGAGGAG | 0.0 | 1.9987849692677924 | 0.987526862634544 | 0.0 | 7.4241615891058474 | 0.0 | 0.0 | 0.0 | 0.0 | 0.0 | 0.0 | 0.0 |
| SE_352_1_5 | SE_352 | SE_352_1 | GAATTTATATTTCAAATGGG | 1.083776841688299 | 0.0 | 1.975053725269088 | 1.3013084895664004 | 0.0 | 0.0 | 0.0 | 11.34424188667422 | 0.0 | 0.0 | 7.042709788187574 | 0.0 |
| SE_352_1_8 | SE_352 | SE_352_1 | CAACTTTCCGGACACCAACG | 387.992109324411 | 439.7326932389143 | 399.9483793669903 | 273.2747828089441 | 404.08650934990396 | 252.96255208157535 | 361.58996588892154 | 352.61685197745703 | 411.1371836330536 | 201.93492371704926 | 251.1899824453568 | 468.9151291420916 |
| SE_353_1_0 | SE_353 | SE_353_1 | GGACCGGCGGAAAGGAAAGT | 2207.653426519065 | 2541.455088423998 | 2511.2808116796455 | 2354.0670576256184 | 2386.337653641165 | 2081.7431940137863 | 2261.0134474185243 | 2043.854246582472 | 2777.617949037131 | 2405.7229179287456 | 2048.254763397886 | 2451.358298881429 |
| SE_353_1_13 | SE_353 | SE_353_1 | TCCGAGATTGTCTGAAAACT | 714.2089386725889 | 834.4927246693034 | 803.8468661845188 | 819.8243484268323 | 800.7488571107021 | 622.8769689953858 | 849.0907234713069 | 892.4136950850386 | 936.8244681055542 | 591.9536391994368 | 732.4418179715077 | 656.9454532040194 |
| SE_353_1_14 | SE_353 | SE_353_1 | ACTCCTAGATGACATAACGG | 61.77527997623304 | 117.92831318679976 | 103.69032057662713 | 72.87327541571842 | 77.42339942924669 | 63.24063802039384 | 35.513300221233365 | 81.30040018783191 | 57.71904305863604 | 67.06863892407412 | 83.33873249355297 | 88.21175696732416 |
| SE_353_1_22 | SE_353 | SE_353_1 | TGGTCAGCCCCAAGCACGGG | 2.167553683376598 | 2.9981774539016888 | 4.93763431317272 | 0.0 | 0.0 | 0.0 | 0.0 | 0.0 | 0.0 | 0.0 | 0.0 | 0.0 |
| SE_353_1_5 | SE_353 | SE_353_1 | TAAAGCCTGCTTGACCACCG | 1024.1691153954425 | 915.4435159246489 | 913.4623479369533 | 1113.9200670688388 | 974.6863571983248 | 1127.9357630486682 | 1087.9983795050587 | 1068.249444328489 | 1078.9021125575812 | 704.2207087027783 | 1147.9616954745745 | 1172.287822855229 |
| SE_353_2_18 | SE_353 | SE_353_2 | ATGCTAACGCTGATTCAGGG | 125.71811363584267 | 144.91191027191493 | 146.1539756699125 | 139.24000838360485 | 99.69588419656424 | 230.43848922499672 | 111.92070372752333 | 113.4424188667422 | 135.86174750725098 | 92.58388199301537 | 134.98527094026184 | 129.99627342553035 |
| SE_353_2_21 | SE_353 | SE_353_2 | GAATCAGCGTTAGCATGTGG | 535.3857597940197 | 491.70110243987693 | 496.72601190517565 | 537.4404061909233 | 500.60061000828 | 575.2299129526234 | 737.1700197437835 | 657.9660294271048 | 429.78487446738217 | 779.3084240199482 | 623.2798162546003 | 922.7414051187199 |
| SE_353_2_22 | SE_353 | SE_353_2 | GGTGATTCAGATTCAGACAC | 106.21013048545329 | 166.89854493386065 | 168.86709351050703 | 136.63739140447206 | 93.3323171201878 | 52.844916701972934 | 139.90087965940418 | 91.69928858394995 | 126.98189472899928 | 271.91959042043095 | 51.646538446708874 | 110.26469620915519 |
| SE_353_2_24 | SE_353 | SE_353_2 | AGACACAGGAATTATGGTGA | 1366.642597368945 | 1601.0267603835016 | 1564.2425504131177 | 1517.3256988344228 | 1597.2553361704865 | 1718.759257978923 | 1504.4725366449773 | 1562.6693198893738 | 1221.8677422874337 | 1746.700639805235 | 1694.9454890238096 | 1095.6828760151843 |
| SE_353_2_25 | SE_353 | SE_353_2 | AATTATGGTGACGGTAGAAA | 17.340429467012783 | 43.97326932389143 | 43.451181955919935 | 22.12224432262881 | 44.54496953463509 | 11.262031428289314 | 10.761606127646475 | 21.743130282792254 | 47.95120500255917 | 2.187020834480678 | 1.173784964697929 | 25.534982280014887 |
| SE_353_3_15 | SE_353 | SE_353_3 | ATCTGAGAAGAACAAGCAGG | 145.22609678623206 | 213.8699917116538 | 236.018920169656 | 152.25309327926885 | 206.8159299822343 | 183.65774329210265 | 111.92070372752333 | 157.8740329228829 | 198.02071695501286 | 134.86628479297514 | 261.75404712763816 | 212.40462532921475 |
| SE_353_3_19 | SE_353 | SE_353_3 | AAGAAGTGAAGACTGAGCAG | 391.2434398494759 | 484.70535504743964 | 494.75095817990655 | 433.33572702561133 | 487.8734758555271 | 397.6363404295996 | 399.2555873356842 | 441.4800800897384 | 456.4244328021373 | 701.3046809234708 | 306.35787578615947 | 458.46900002754 |
| SE_353_3_20 | SE_353 | SE_353_3 | AGAAGTGAAGACTGAGCAGG | 318.63039145635986 | 547.6670815793751 | 456.23741053715935 | 446.34881192127534 | 434.84375021905674 | 612.4812476769649 | 439.07353000797616 | 427.29977773139564 | 638.461414756297 | 713.6977989855279 | 761.7864420889559 | 631.4104709240045 |
| SE_353_3_22 | SE_353 | SE_353_3 | GGCTGGAGTGTTCTAATAGC | 436.7620672003845 | 617.6245555037478 | 599.4288056191682 | 394.2964723386193 | 618.3266009212441 | 373.3796573532842 | 291.63952605921946 | 433.9172521652889 | 654.4451497571501 | 376.8965904755035 | 285.22974642159676 | 318.02659748745816 |
| SE_353_3_23 | SE_353 | SE_353_3 | GGGATAATGAGTGCTAGGGA | 164.73407993662144 | 240.85358879676897 | 266.6322529113269 | 249.85122999674888 | 182.42225618945795 | 218.310147686839 | 116.22534617858193 | 238.22907962015861 | 179.3730261206843 | 284.3127084824881 | 413.172307573671 | 132.31763545098624 |
| SE_353_4_0 | SE_353 | SE_353_4 | AATGGTAAAACTTTATTATC | 53.105065242726646 | 79.95139877071169 | 94.80257881291622 | 49.44972260352321 | 84.84756101835254 | 129.0802063703929 | 51.655709412703075 | 34.03272566002266 | 134.08577695160065 | 120.28614589643729 | 9.390279717583432 | 96.3365240564198 |
| SE_353_4_2 | SE_353 | SE_353_4 | CCTGTCCAGAGCGCAGGGAG | 2135.040378125949 | 1989.7904369060873 | 1910.8644791978427 | 2143.2550823158617 | 2000.281251007661 | 2218.6201913729947 | 2216.890862295174 | 2473.04473129498 | 1937.5838762145204 | 2768.0393695077114 | 2085.81588226822 | 2453.679660906885 |
| SE_353_4_3 | SE_353 | SE_353_4 | AAATATGGAGTTGGGCATGG | 96.4561389102586 | 80.95079125534559 | 118.50322351614528 | 111.91253010271043 | 102.87766773475245 | 58.04277736118338 | 99.00677637434757 | 43.48626056558451 | 67.4868811147129 | 112.26706950334147 | 32.86597901154201 | 49.909283547301825 |
| SE_353_4_4 | SE_353 | SE_353_4 | ATGGAGTTGGGCATGGTGGT | 5398.292448449417 | 5029.9423751624 | 5009.723774145042 | 5517.547995761538 | 4837.3715725588245 | 4960.491689106509 | 5375.422260759414 | 5598.383371073727 | 5200.041786944194 | 4819.464912250587 | 5044.927778271699 | 4431.480106595311 |
| SE_353_4_6 | SE_353 | SE_353_4 | AGTTGGGCATGGTGGTTGGG | 592.8259324034995 | 631.6160502886224 | 652.7552562014336 | 620.724149523173 | 691.5076222995732 | 512.8555850420979 | 670.4480617523753 | 588.009871125947 | 668.6529142023528 | 450.5262919030197 | 815.7805504650606 | 759.085382324079 |
| SE_353_5_16 | SE_353 | SE_353_5 | GCAGGATATAGCCCGGAGGA | 1094.614610105182 | 1347.181069286492 | 1279.8348139743691 | 1259.6666179002757 | 1338.4702750645113 | 1975.187050499972 | 1006.2101729349454 | 1306.4785239486478 | 1248.5073006221887 | 854.3961393371181 | 1348.6789244379204 | 1614.5072887045776 |
| SE_353_5_19 | SE_353 | SE_353_5 | AAGCTCTGCAGGATATAGCC | 32.51330525064897 | 12.992102300240651 | 23.700644703229056 | 27.32747828089441 | 16.969512203670508 | 181.92512307236584 | 71.02660044246673 | 37.8141396222474 | 74.59076333731426 | 6.561062503442034 | 9.390279717583432 | 16.249534178191293 |
| SE_353_5_21 | SE_353 | SE_353_5 | ATATCCTGCAGAGCTTCAGC | 10.837768416882989 | 1.9987849692677924 | 11.850322351614528 | 20.820935833062407 | 0.0 | 18.192512307236584 | 22.599372868057596 | 2.836060471668555 | 48.83919028038434 | 0.7290069448268927 | 0.0 | 0.0 |
| SE_353_5_26 | SE_353 | SE_353_5 | TCTCCGTGATGAGGTGGTCA | 252.52000411337366 | 267.8371858818842 | 274.53246781240324 | 252.45384697588167 | 270.45160074599875 | 423.62564372565186 | 262.583189514574 | 271.3164517896251 | 380.05769890917264 | 341.90425712381267 | 181.936669528179 | 304.0984253347228 |
| SE_353_5_27 | SE_353 | SE_353_5 | TCACCCTGACCACCTCATCA | 521.2966608520718 | 439.7326932389143 | 537.214613273192 | 571.2744269196498 | 482.5705032918801 | 549.2406096565711 | 681.2096678800218 | 335.60048914744567 | 449.3205505795359 | 681.6214934131447 | 331.007360044816 | 495.6107924348344 |
| SE_354_1_0 | SE_354 | SE_354_1 | TGAGAAACGTGATATTTGGT | 2022.3275865903659 | 2102.7217876697177 | 2203.172430537668 | 1763.2730033624725 | 2173.1581565825545 | 1877.2940080848416 | 1855.3008964062521 | 2011.7122279035616 | 2023.718448163562 | 2349.589383177075 | 1967.263600833729 | 2302.7911292522517 |
| SE_354_1_19 | SE_354 | SE_354_1 | GTCTCACTCTTCCTAAGGCC | 125.71811363584267 | 153.90644263362 | 147.14150253254707 | 122.32299801924164 | 110.3018293238583 | 37.251334724341575 | 83.94052779564251 | 64.28403735782058 | 113.66211556162173 | 185.89677093085763 | 151.41826044603283 | 157.85261773100112 |
| SE_354_1_3 | SE_354 | SE_354_1 | CTATTCCTAATATCAGCATT | 16.256652625324485 | 19.987849692677923 | 16.78795666478725 | 9.109159426964803 | 18.030106716399914 | 26.85561340592067 | 8.60928490211718 | 70.90151179171387 | 68.37486639253807 | 2.187020834480678 | 68.07952795247988 | 63.83745570003722 |
| SE_354_1_5 | SE_354 | SE_354_1 | AACACATGTGAGGAAACCAG | 1449.0096373372558 | 1532.068678943763 | 1573.1302921768286 | 1466.5746677413333 | 1625.8913880141806 | 1354.9090118341912 | 1364.571656985573 | 1363.1997333820188 | 1624.1250731422356 | 1496.6512577296107 | 1455.493356225432 | 1658.6131671882397 |
| SE_354_1_7 | SE_354 | SE_354_1 | GGATGTAGTGCAAACCACAC | 3067.088461977886 | 2704.356063419323 | 2951.717792414652 | 2916.2323251183034 | 2832.8479435002455 | 2211.6897104940476 | 3159.607559077005 | 2498.569275539997 | 2676.3876273650617 | 3514.5424810104496 | 2615.1929013469858 | 3372.939022987421 |
| SE_355_1_14 | SE_355 | SE_355_1 | GCATTTGTCTGGTTTTCCAT | 275.27931778882794 | 384.76610658405 | 340.6967676089177 | 379.9820789533889 | 403.0259148371746 | 534.5133377888081 | 311.0104170889831 | 352.61685197745703 | 351.6421700187672 | 402.41183354444473 | 221.8453583279086 | 294.81297723289913 |
| SE_355_1_19 | SE_355 | SE_355_1 | TCACTGATAAGGGATGGTGA | 1763.3049214268624 | 1753.9338105324878 | 1703.4838380445885 | 1564.1728044588133 | 1534.6802599194516 | 1718.759257978923 | 1481.8731637769197 | 1335.7844821558895 | 1526.4466925814668 | 1779.505952322445 | 1189.044169239002 | 2179.758941903089 |
| SE_355_1_2 | SE_355 | SE_355_1 | TTGCGAAACATTCATTTCCA | 597.1610397702527 | 536.6737642484022 | 576.7156877785737 | 510.112927910029 | 609.8418448194088 | 595.1550454795968 | 475.6629908419742 | 544.5236105603626 | 553.2148280850807 | 524.1559933305358 | 442.5169316911192 | 444.5408278748046 |
| SE_355_1_3 | SE_355 | SE_355_1 | AAGGCACCCAGAACTCAACA | 6040.972115570578 | 6256.19695380819 | 6121.679021471538 | 5898.831383204493 | 5985.995429844772 | 5963.678796334126 | 5761.763920741922 | 5571.913473338154 | 5841.167157533967 | 5751.864794684183 | 5128.266510765252 | 6122.592342139933 |
| SE_355_1_7 | SE_355 | SE_355_1 | GGCTGTTCTTCACCTCCAGG | 138.72343573610226 | 120.92649064070145 | 113.56558920297256 | 85.88636031138243 | 111.36242383658771 | 348.2566641671003 | 116.22534617858193 | 101.15282348951179 | 118.10204195074758 | 73.62970142751615 | 145.5493356225432 | 159.01329874372908 |
| SE_356_1_15 | SE_356 | SE_356_1 | ATATCCTGTCCTGGAATTAG | 82.36703996831072 | 125.92345306387092 | 104.67784743926167 | 88.48897729051524 | 81.66577748016432 | 90.96256153618292 | 64.56963676587885 | 64.28403735782058 | 166.94123223113192 | 64.88161808959345 | 179.58909959878315 | 91.69380000550801 |
| SE_356_1_38 | SE_356 | SE_356_1 | TCTGGCTCACCATCGCCCCC | 3920.0208363865772 | 3443.9065020484063 | 3400.054988050735 | 4036.6589346349742 | 3468.14405662516 | 3191.4864447552177 | 4186.264783654478 | 3809.7745669414257 | 3313.0730715657087 | 4049.633578513389 | 4047.2105582784593 | 3830.247342002233 |
| SE_356_1_6 | SE_356 | SE_356_1 | GCCTGGGCCAGACAGAGATG | 1304.8673173927118 | 1281.221165300655 | 1261.0718035843126 | 1099.6056736836083 | 1299.2282780935232 | 1205.903672936825 | 1353.8100508579266 | 1346.1833705520075 | 1268.0429767343423 | 1529.4565702468208 | 1035.2783388635735 | 981.9361367678453 |
| SE_356_1_9 | SE_356 | SE_356_1 | GGACAGGATATCCTCATTGG | 969.9802733110275 | 1066.3517811043673 | 1153.4313755571475 | 992.8983775391636 | 1016.0495431947717 | 1346.245910735507 | 1065.399006637001 | 1079.5936862151632 | 1006.9753050537425 | 857.3121671164257 | 876.817368629353 | 1157.1989696897656 |
| SE_356_2_12 | SE_356 | SE_356_2 | ACTCGGCCCCTGAGTCTGAG | 664.3552039549272 | 736.5522611751815 | 708.0567605089681 | 641.5450853562354 | 660.7503814304204 | 453.94649757104617 | 569.2889641524985 | 723.1954202754815 | 712.1641928157861 | 790.9725351371785 | 703.0971938540595 | 901.8491468896167 |
| SE_356_2_17 | SE_356 | SE_356_2 | GGGATCAGTGTATCAGGCCA | 555.9775197860973 | 714.5656265132358 | 744.5952544264462 | 671.4751806162626 | 778.4763723433846 | 784.8769595407783 | 626.3254766290248 | 661.7474433893295 | 791.1948825422263 | 720.25886148897 | 528.2032341140681 | 667.391582318571 |
| SE_356_2_2 | SE_356 | SE_356_2 | TCAAGGCAGGAAGTGGTGAG | 6995.779513097969 | 5914.404724063397 | 5909.360746005112 | 6548.184319498127 | 5795.088417553478 | 6704.3739402716155 | 7077.908350153087 | 6688.375945685009 | 6193.697312830559 | 6846.104218869349 | 6941.764281223553 | 5630.463592743283 |
| SE_356_2_5 | SE_356 | SE_356_2 | ACAGGCAATCAAAGGAACAG | 4171.457063658262 | 4487.272256006194 | 4497.197332437713 | 4486.9116720249485 | 4503.2843010490615 | 4629.561227136776 | 4102.324255858836 | 4452.614940519631 | 4916.774483317965 | 3694.607196382692 | 4747.960182203123 | 5072.176025621139 |
| SE_356_2_7 | SE_356 | SE_356_2 | ACCAGAGCACAGGCAATCAA | 3278.424946107104 | 3416.922904963291 | 3302.2898286499153 | 3464.083199225758 | 3490.4165413924775 | 2905.6041084986427 | 3418.962266753285 | 3147.08177006154 | 3186.9791621145346 | 3779.9010089274384 | 3264.2959868249404 | 3222.0504913327877 |
| SE_356_3_13 | SE_356 | SE_356_3 | TCTGTGGCAGGAAGGAGAGA | 1097.8659406302468 | 1106.327480489723 | 1080.3543877221912 | 1159.4658642036627 | 1156.0480188750535 | 1300.3314749124816 | 1125.6640009518212 | 1885.034860169033 | 1130.405258671441 | 1058.518083888648 | 2082.294527374126 | 1493.7964633808708 |
| SE_356_3_34 | SE_356 | SE_356_3 | GCATGCCCAGATGTAGCCTT | 2310.612226479453 | 2064.7448732536295 | 2082.6941532962533 | 2450.363885853532 | 2074.5228668987197 | 2029.7645874216817 | 2480.5502124225122 | 2307.6078704476477 | 2514.774306800881 | 2448.0053207287056 | 2166.807044832377 | 1998.692703917529 |
| SE_356_3_45 | SE_356 | SE_356_3 | CACTTGAGAGAGGAAGGGCG | 849.6810438836263 | 843.4872570310084 | 855.1982630415151 | 687.0908824910595 | 1022.4131102711481 | 891.4331030545926 | 820.0343869266613 | 726.0314807471501 | 942.1523797725051 | 497.9117433167677 | 933.1590469348536 | 797.3878557441012 |
| SE_356_3_5 | SE_356 | SE_356_3 | GGTCCCACAGCGCCACCTAG | 440.01339772544935 | 511.68895213255485 | 539.189666998461 | 530.9338637430914 | 479.3887197536919 | 405.43313141841526 | 503.643166773855 | 603.135526974846 | 475.960108914291 | 416.2629654961557 | 588.0662673136625 | 629.0891088985486 |
| SE_356_3_9 | SE_356 | SE_356_3 | ACGCTGCTAGACATGTCATG | 17370.775218580056 | 16090.219002605729 | 16424.546779337736 | 17017.21111805982 | 16547.395587604206 | 16399.25037980898 | 17553.255754804166 | 15044.355448711129 | 15494.455112771388 | 15548.989126212793 | 15837.880528669157 | 14763.862481899518 |
| SE_356_4_1 | SE_356 | SE_356_4 | GCAACCTCATCACCAATCCT | 3831.1511353681367 | 3384.9423454550065 | 3464.2442341219803 | 3562.9826444328046 | 3625.1120445091124 | 3290.2457972802163 | 3620.204301340274 | 3006.2240999686683 | 3567.9248463015324 | 3319.1686197968424 | 3705.6391335513617 | 4006.6708559368813 |
| SE_356_4_15 | SE_356 | SE_356_4 | CATGCCAGACACACACAAGA | 5294.24987164734 | 5095.902279148237 | 5205.254092946681 | 5341.871349670074 | 5321.002670363434 | 5186.598627782163 | 4957.871943006731 | 5534.099333715907 | 5671.56196946936 | 4936.835030367717 | 5399.410837610473 | 5369.310364879494 |
| SE_356_4_2 | SE_356 | SE_356_4 | AGACAGCTCAGACAGGCCTG | 2177.3076749517927 | 2194.665896256036 | 2317.725546603275 | 2356.669674604751 | 2260.126906626366 | 2369.358150490098 | 2326.659244797168 | 2543.946243086694 | 2329.18538373542 | 1700.7732022811406 | 2290.0544661256595 | 2994.5570128381096 |
| SE_356_4_23 | SE_356 | SE_356_4 | GCCACTTTGTACATGTCCCA | 276.36309463051623 | 363.7788644067382 | 370.322573487954 | 439.84226947344337 | 321.3601373570102 | 292.81281713552215 | 286.2587229953962 | 344.1086705624513 | 346.3142583518162 | 608.7207989304553 | 268.79675691582577 | 490.9680683839226 |
| SE_356_4_9 | SE_356 | SE_356_4 | GGTTCAGAGTTGATGTTCAA | 34.680858934025565 | 58.96415659339988 | 58.264084895438096 | 53.35364807222242 | 38.181402458258646 | 103.95721318420905 | 53.808030638232374 | 34.03272566002266 | 34.63142583518162 | 20.412194455152996 | 8.216494752885502 | 73.12290380186082 |
| SE_356_5_1 | SE_356 | SE_356_5 | GACACTTCAAAGAAAATACA | 196.1636083455821 | 269.83597085115196 | 280.4576289882105 | 214.71590077845607 | 226.96722572409305 | 198.3850151598656 | 171.10953742957895 | 224.99413075237203 | 287.70723001535504 | 290.14476404110326 | 205.41236882213758 | 259.9925468510607 |
| SE_356_5_12 | SE_356 | SE_356_5 | GGACAGGTCTGCGAATGGGG | 828.0055070498604 | 1104.3286955204553 | 1087.267075760633 | 946.0512719147731 | 1018.1707322202304 | 1109.7432507414317 | 982.5346394541232 | 933.0638951789546 | 1228.0836392322099 | 914.9037157577503 | 789.9572812417063 | 789.2630886550056 |
| SE_356_5_19 | SE_356 | SE_356_5 | TGTTAGGCTCTGCCAAGAGG | 170.15296414506292 | 220.86573910409106 | 232.06881271911786 | 169.17010364363205 | 200.45236290585788 | 285.01602614670645 | 187.25194662104866 | 116.27847933841076 | 160.72533528635574 | 252.23640291010486 | 137.3328408696577 | 89.37243798005211 |
| SE_356_5_2 | SE_356 | SE_356_5 | CAGCGAGTATAGCTAGTAAG | 1465.2662899625802 | 1610.0212927452067 | 1614.6064204074794 | 1664.373558155426 | 1644.98208924331 | 2185.7004071979954 | 1461.4261121343914 | 1440.7187196076259 | 1592.1576031405293 | 1753.2617023086768 | 1467.2312058724112 | 2413.055825461407 |
| SE_356_5_27 | SE_356 | SE_356_5 | GCTCTGGGCTCTGCACCCAT | 477.9455871845398 | 501.69502728621586 | 568.8154728774973 | 482.7854496291346 | 562.1150917465856 | 845.518667231567 | 528.3948608674419 | 713.7418853699197 | 775.2111475413732 | 634.9650489442236 | 819.3019053591545 | 438.7374228111649 |
| SE_357_1_14 | SE_357 | SE_357_1 | TAGACGATCAGGGCAATGCG | 5737.514599897854 | 5000.9599931080165 | 5054.162482963597 | 5576.106877792025 | 5055.854042181082 | 5148.480982947953 | 5744.545350937688 | 4510.281503443559 | 4339.584052731605 | 4672.934516340382 | 4946.329841237073 | 4483.710752168069 |
| SE_357_1_15 | SE_357 | SE_357_1 | GGAGTCATGAACCTGAGAAG | 309.9601767228535 | 394.760031430389 | 349.5845093726286 | 391.6938553594865 | 389.2381861716923 | 336.1283226289426 | 365.89460833998015 | 397.9938195241539 | 396.0414339100257 | 626.9459725511277 | 517.6391694317867 | 297.13433925835506 |
| SE_357_1_16 | SE_357 | SE_357_1 | TGATGGGGTGGCCAATTACA | 3442.0752492020374 | 3505.8688360957076 | 3397.0924074628315 | 3444.563571882262 | 3124.5114345008324 | 3170.695002118376 | 3349.011826923583 | 3083.7430861942753 | 3570.588802135008 | 3228.0427516934806 | 3123.441791061189 | 3239.460706523707 |
| SE_357_1_23 | SE_357 | SE_357_1 | GACCAGTGTGAAGAAACACA | 709.8738313058358 | 786.5218854068763 | 730.7698783495625 | 814.6191144685666 | 776.3551833179257 | 774.4812382223574 | 747.93162587143 | 646.6217875404305 | 872.8895281021419 | 868.976278233656 | 428.4315121147441 | 740.5144861204317 |
| SE_357_1_29 | SE_357 | SE_357_1 | TTACCAGAGTAAGAAATGCA | 131.13699784428417 | 148.90948021045054 | 144.17892194464343 | 184.78580551842887 | 126.21074701479941 | 149.8716490072347 | 156.0432888508739 | 107.7702979234051 | 111.88614500597139 | 298.89284737902597 | 151.41826044603283 | 53.39132658548567 |

| | | | | | | | | | | | | | | | |
|---|---|---|---|---|---|---|---|---|---|---|---|---|---|---|---|
| SE_357_2_0 | SE_357 | SE_357_2 | GGATGCTCATAAATAAGCAG | 192.9122778205172 | 245.85055121993847 | 247.86924252127056 | 248.5499215071825 | 275.75457330964576 | 274.62030482828555 | 202.31819519975372 | 240.119786601271 | 220.2203489006421 | 336.8012085100244 | 379.1325435974311 | 304.0984253347228 |
| SE_357_2_20 | SE_357 | SE_357_2 | CAGTATGTAGCCACCTGCCA | 408.5838693164887 | 323.80316502138237 | 327.8589183946686 | 305.8074950481041 | 410.4500764262804 | 236.50265999407557 | 352.9806809868044 | 401.7752334863786 | 245.97192195757202 | 195.37386121360723 | 238.2783478336796 | 420.1665266075177 |
| SE_357_2_21 | SE_357 | SE_357_2 | TTAAAATTCCATTCTCCTGA | 14.089098941947885 | 17.98906472341013 | 13.825376076883616 | 13.013084895664004 | 11.666539640023474 | 20.791442636841808 | 16.142409191469714 | 35.92343264113503 | 19.535676112153734 | 4.374041668961356 | 14.085419576375148 | 77.76562785277261 |
| SE_357_2_3 | SE_357 | SE_357_2 | TTCAAGATAACAAGAAAAAA | 5.4188842084414945 | 0.9993924846338962 | 0.0 | 1.3013084895664004 | 7.4241615891058474 | 0.0 | 5.380803063823238 | 32.14201867891029 | 1.7759705556503396 | 0.0 | 2.347569929395858 | 0.0 |
| SE_357_2_7 | SE_357 | SE_357_2 | GTGGGAGAAGAAACAAGGGA | 463.8564882425919 | 616.625163019114 | 645.8425681629918 | 474.97759869173615 | 638.4778966631029 | 529.3154771295978 | 507.9478092249136 | 579.5016897109414 | 477.73607946994133 | 493.53770164780633 | 570.4594928431935 | 391.14950128931895 |
| SE_358_1_1 | SE_358 | SE_358_1 | AGTTTCACAGGAAAATTGAG | 24.926867358830876 | 36.977521931454156 | 29.62580587903632 | 28.62878677046081 | 23.33307928004695 | 2.598930329605226 | 30.13249715741013 | 7.56282792444948 | 55.05508722516053 | 26.973256958595027 | 1.173784964697929 | 1.1606810127279494 |
| SE_358_1_12 | SE_358 | SE_358_1 | CTTGAATCTGTAAGAACAAC | 386.9083324827227 | 448.72722560061936 | 532.2769789600193 | 386.48862140122094 | 512.2671496483034 | 458.27804812038823 | 455.21593919944587 | 379.0867497130302 | 396.9294191878509 | 474.58352108230713 | 626.8011711486942 | 538.5559899057686 |
| SE_358_1_3 | SE_358 | SE_358_1 | TCTATGGAGGACAGACCAAG | 368.48412617402164 | 275.83232575895534 | 324.89633780676496 | 363.0650685890257 | 347.8750001752454 | 388.1069292210471 | 340.0667536336286 | 432.97189867473276 | 346.3142583518162 | 396.5797779858296 | 630.3225260427879 | 493.2894304093785 |
| SE_358_1_7 | SE_358 | SE_358_1 | AGTTCATCCAAGCAGTCAAT | 189.6609472954523 | 199.87849692677923 | 174.7922546863143 | 182.18318853929605 | 145.30144824392872 | 232.17110944473353 | 191.55658907210724 | 130.45878169675353 | 85.2465866712163 | 111.53806255851458 | 218.3240034338148 | 128.83559241280238 |
| SE_358_1_9 | SE_358 | SE_358_1 | AGTCAATCGGTCATTTCACA | 579.82061030324 | 463.7181128701278 | 591.5285907180919 | 582.9862033257474 | 583.3269820011737 | 668.7914048184116 | 652.1533313353764 | 687.2719876343465 | 423.56897752260596 | 295.9768195997184 | 839.2562497590193 | 497.9321544602903 |
| SE_358_2_13 | SE_358 | SE_358_2 | GGCCATAAAGAGGAAGAGAG | 391.2434398494759 | 452.72479553915497 | 475.00042092721566 | 378.6807704638225 | 453.9344514481861 | 436.620295373678 | 345.44755669745183 | 541.687550088694 | 589.6222244759127 | 390.01871548238756 | 395.5655331032021 | 403.9169924293264 |
| SE_358_2_20 | SE_358 | SE_358_2 | GGTGACATCAGGGGAGAGAG | 657.8525429047975 | 806.5097350995542 | 736.6950395253698 | 642.8463938458018 | 527.1154728265152 | 533.6470276789398 | 588.6598551822622 | 344.1086705624513 | 723.7080014275134 | 650.2741947855883 | 299.3151659979719 | 239.1002886219576 |
| SE_358_2_30 | SE_358 | SE_358_2 | CTAAGTGCTAGAGCTGGGAA | 277.4468714722045 | 340.7928372601586 | 325.88386466939954 | 265.4669318715457 | 418.9348325281157 | 318.8021204315744 | 185.09962539551935 | 357.34361943023794 | 436.88875668998355 | 344.09127795829335 | 350.9617044446808 | 598.911402567622 |
| SE_358_2_36 | SE_358 | SE_358_2 | GAGCGGGATGACCTCAGCAA | 6315.167656517718 | 6061.31541930458 | 6209.568912246013 | 6221.55588861696 | 5757.967609607949 | 6008.726922047283 | 6168.552632366959 | 6069.169409370707 | 5776.34423225273 | 6418.9061492007895 | 6536.808468402767 | 6093.575316821734 |
| SE_358_2_6 | SE_358 | SE_358_2 | AGAAGGAATGAGATTATGCG | 929.8805301685604 | 1042.3663614731538 | 1022.090302826753 | 948.6538888939059 | 914.2324699727486 | 918.2887164605132 | 931.9550906541847 | 1015.3096488573427 | 1039.830760333274 | 531.4460627788047 | 951.9396063700204 | 776.4955975149982 |
| SE_358_3_12 | SE_358 | SE_358_3 | TAATTCTGAATGATGCCATG | 146.30987362792035 | 147.91008772581662 | 197.50537252690881 | 161.36225270623365 | 169.69512203670507 | 214.84490724736537 | 135.59623720834557 | 187.17999113012462 | 207.78855501108973 | 219.4310903928947 | 134.98527094026184 | 227.4934784946781 |
| SE_358_3_23 | SE_358 | SE_358_3 | GCATCTCACCCGGCAGAGGA | 1350.3859447436205 | 1514.0796142203528 | 1477.340186501278 | 1540.7492516466182 | 1546.3467995594751 | 1577.5507100703724 | 1450.664506006745 | 1391.5603380987043 | 1347.9616517386078 | 1383.6551812814423 | 1084.5773073808864 | 1166.4844177915893 |
| SE_358_3_28 | SE_358 | SE_358_3 | CCTGATTAGGGAGACAGCTG | 986.236925936352 | 1009.3864094802351 | 995.4270775356204 | 962.9682822791364 | 1060.5945127294067 | 907.0266850322239 | 1099.8361462454698 | 941.5720765939602 | 951.920217828582 | 1545.4947230330124 | 1288.815891238326 | 1063.1838076588017 |
| SE_358_3_5 | SE_358 | SE_358_3 | GCAGAGGCCAGAGGACTGAG | 3752.0354259248907 | 3958.593631634863 | 3802.965948005629 | 4009.33145635408 | 3579.5064804617477 | 3304.9730691479795 | 3892.4729363697297 | 3454.3216544923 | 3627.4198599158185 | 3708.4583283344027 | 3568.3062926817042 | 2960.897263468999 |
| SE_358_3_6 | SE_358 | SE_358_3 | TCCCCCACACAGCATCCTGC | 143.05854310285545 | 159.90279754142338 | 166.89203978523796 | 135.33608291490566 | 146.36204275665813 | 103.09090307434064 | 117.30150679134657 | 254.30008895961376 | 134.08577695160065 | 81.64877782061198 | 159.63475519891836 | 152.0492126673614 |
| SE_358_4_13 | SE_358 | SE_358_4 | TGGACGATCAGGGACTACCA | 276.36309463051623 | 346.78919216796197 | 321.93375721886133 | 324.0258139020337 | 283.1787348987516 | 431.42243471446756 | 309.9342564762185 | 399.88452650526625 | 223.77229001194277 | 247.1333542963166 | 328.65979011542015 | 316.86591647473017 |
| SE_358_4_14 | SE_358 | SE_358_4 | GAGTGCAAATAGTTTCAGAG | 510.4588924351888 | 760.537680806395 | 730.7698783495625 | 551.7547995761538 | 846.3544211580665 | 805.6684021776201 | 404.63639039950743 | 754.3920854638357 | 659.7730614241011 | 917.8197435370579 | 645.5817305838609 | 976.1327317042055 |
| SE_358_4_16 | SE_358 | SE_358_4 | CTGGGCCTCCAGGGACACAG | 291.5359704141524 | 315.8080251443112 | 304.15827369143955 | 378.6807704638225 | 324.5419208951985 | 229.5721791151283 | 344.3713960846872 | 232.55695867682152 | 346.3142583518162 | 204.12194455152994 | 89.2076573170426 | 255.34982280014887 |
| SE_358_4_26 | SE_358 | SE_358_4 | GCCTCTCTAAAGGTCTGAGG | 48.769957875973454 | 108.93378082509469 | 92.82752508764715 | 49.44972260352321 | 65.75685978922321 | 183.65774329210265 | 32.28481838293943 | 135.18554914953447 | 174.93309973155846 | 41.553395855132884 | 7.042709788187574 | 174.10215190919243 |
| SE_358_4_27 | SE_358 | SE_358_4 | CAGGTGTCATGCCCTCTGAA | 2221.742525461013 | 2129.7053847548327 | 2096.519529373137 | 2213.5257407524473 | 2163.61280596799 | 2144.98383203418 | 2138.3311375633543 | 1986.1876836585448 | 2575.1573056929924 | 2230.0322442254646 | 1599.8689068832773 | 1946.462058344771 |
| SE_359_1_19 | SE_359 | SE_359_1 | CCTGCATTCGCGGCGTGTGC | 424.84052194181317 | 334.7964823523552 | 351.5595630978977 | 394.2964723386193 | 364.8445123789159 | 388.97323933091553 | 509.02396983767824 | 380.03210320358636 | 340.9863466848652 | 294.5188057100646 | 313.4005855743471 | 349.3649848311128 |
| SE_359_1_20 | SE_359 | SE_359_1 | CTGCACACGCCGCGAATGCA | 622.0879071290835 | 630.6166578039885 | 592.5161175807265 | 495.7985345247986 | 522.8730947755976 | 470.40638965854595 | 682.2858284927865 | 612.5890618804079 | 687.3006050366814 | 618.1978892132049 | 467.16641594977574 | 465.4330861039077 |
| SE_359_1_23 | SE_359 | SE_359_1 | CTCCAGCCAGATGCGGTAAG | 54.188842084414944 | 67.95868895510495 | 84.92731018657079 | 48.1484411395682 | 45.60556404736449 | 116.08555472236677 | 33.36097899570407 | 80.35504669727572 | 74.59076333731426 | 17.496166675845423 | 62.210603128990236 | 58.034050636397474 |
| SE_359_1_24 | SE_359 | SE_359_1 | GACCGCTTACCGCATCTGGC | 4090.17380053164 | 3866.6495230485443 | 3967.882934065598 | 3790.7116301069245 | 3756.6257640875588 | 3619.4436390302117 | 3938.7478427186097 | 3771.015073828622 | 3493.334082964218 | 3620.9774949551756 | 3858.2311789620926 | 3670.073362245776 |
| SE_359_1_36 | SE_359 | SE_359_1 | TGGAGAAGGCGGCCTGGCGC | 1066.4364122212862 | 938.4295430712285 | 929.2627777391059 | 1020.225855820058 | 891.959985205431 | 1078.5560867861689 | 1163.329622398584 | 945.353490556185 | 976.7838056076868 | 763.9992781785835 | 963.6774560169997 | 1085.2367469006326 |
| SE_359_2_1 | SE_359 | SE_359_2 | AGACCAGCCTTGCCAACGTG | 2967.3809925425626 | 2428.5237376603677 | 2431.2911358062474 | 2791.306710119929 | 2546.4874250633056 | 2649.1763159775937 | 3022.935161255895 | 2677.241085255116 | 2669.2837451424602 | 2478.623612411435 | 2469.643565724443 | 2677.6910963633795 |
| SE_359_2_18 | SE_359 | SE_359_2 | GAGTTTTCCGAGAAAGGGAT | 26.010644200519174 | 39.975699385355846 | 36.53849391747813 | 54.6549565617882 | 31.8178353818822 | 48.51336615263089 | 40.8941032850566 | 47.26767452780925 | 77.25471917078977 | 30.61829168272949 | 32.86597901154201 | 63.83745570003722 |
| SE_359_2_2 | SE_359 | SE_359_2 | ACGATAGTTTACAAATGCCG | 3334.7813418748956 | 4310.379786225994 | 4415.232602839046 | 3924.7464045322636 | 3998.4413129898635 | 3260.7912535446903 | 3404.9721787873445 | 3769.1243668475095 | 4875.927160538007 | 4551.190356554291 | 4551.938093098569 | 3707.2151546530704 |
| SE_359_2_21 | SE_359 | SE_359_2 | ACTATCTCAGAACCTTAGAG | 35.76463577571386 | 59.96354907803377 | 59.25161175807264 | 46.847105624390416 | 75.30221040378788 | 219.1764577967074 | 30.13249715741013 | 53.88514896170255 | 6.215896944776189 | 10.206097227576498 | 62.210603128990236 | 110.26469620915519 |
| SE_359_2_3 | SE_359 | SE_359_2 | TGACTGAGAATGCCTAACGT | 2693.185451595423 | 2834.2770864217296 | 2748.287258711936 | 2752.267455432937 | 2896.4836142640097 | 2427.4009278512813 | 2579.55698879686 | 2546.7823035583624 | 2229.7310326190013 | 3315.5235850727076 | 2271.273906690493 | 2385.199481155936 |
| SE_359_3_14 | SE_359 | SE_359_3 | ACAACCGCAAACTCTCCAAG | 1428.4178773451779 | 1970.8019796980432 | 1798.2864168575047 | 1695.6049619050198 | 1867.7069369164853 | 2149.315382583522 | 1533.5288731896226 | 1930.4118277157297 | 2237.722900119428 | 1915.101244060247 | 1862.7967389756134 | 2439.7514887541497 |
| SE_359_3_19 | SE_359 | SE_359_3 | TCTGTATTCATGAAACTGAG | 135.47210521103736 | 167.89793741849456 | 156.02924429625796 | 121.02168952967524 | 180.30106716399914 | 254.69517230131217 | 103.31141882540615 | 145.5844375456525 | 157.17339417505505 | 84.56480559991955 | 52.820323411406804 | 60.35541266185337 |
| SE_359_3_23 | SE_359 | SE_359_3 | TGCCCTCACACAGAACCACA | 2092.7730813001053 | 1987.7916519368196 | 2000.7294236975863 | 2037.8490946609832 | 1971.6451991639672 | 1933.6041652262884 | 1898.347320916838 | 2153.5152514869897 | 2255.482605675931 | 1965.4027232533026 | 2486.0765552302137 | 2025.3883672102718 |
| SE_359_3_24 | SE_359 | SE_359_3 | GTCCTTGTGGTTCTGTGTGA | 49.85373471766175 | 87.94653864778286 | 65.1767729338799 | 89.79028578008163 | 106.05945127294068 | 28.588233625657487 | 75.33124289352533 | 99.26211650839943 | 17.759705556503395 | 68.5266528137279 | 109.1620017169074 | 31.338387343654635 |
| SE_359_3_6 | SE_359 | SE_359_3 | TGCACTGCTCAGTCTCTGTA | 2394.0630432894523 | 2242.636735518463 | 2157.7461948564787 | 2408.7220141874072 | 2399.064787793918 | 2098.203086101286 | 2324.5069235716387 | 2543.0008895961378 | 2576.933276248643 | 2081.3148274807786 | 2441.4727265716924 | 2785.634430547079 |
| SE_36_1_14 | SE_36 | SE_36_1 | GTCTCCTGAGGCCATGCACG | 89.95347786012881 | 101.9380334326574 | 93.81505195028168 | 96.29682822791364 | 117.72599091296415 | 84.89839076710406 | 83.94052779564251 | 118.16918631952312 | 83.47061611556596 | 231.09520151012498 | 190.1531642810645 | 169.4594278582806 |
| SE_36_1_16 | SE_36 | SE_36_1 | CGGTGGCTACTGGTGTTGAA | 721.7953765644071 | 790.5194553454119 | 812.7346079482297 | 717.0209777510867 | 709.5377290159731 | 699.1122586638058 | 627.4016372417894 | 743.9931970677176 | 895.0891600477711 | 649.5451878407614 | 842.777604653113 | 854.2612253677707 |
| SE_36_1_19 | SE_36 | SE_36_1 | AGGCAGGCAGCCTAGCTGGT | 8528.239967245225 | 7382.512283990591 | 7234.62179566067 | 8457.203873692037 | 7423.100994593118 | 8554.812334950537 | 8588.83785047465 | 7159.161983981989 | 7612.697786795181 | 6806.008836903869 | 6655.360749837258 | 7680.2262612208415 |
| SE_36_1_21 | SE_36 | SE_36_1 | TAGGCTGCCTGCCTCTTCCG | 5414.549101074741 | 4935.000089122179 | 4882.332808865186 | 5513.6440702928385 | 5028.278584850117 | 4837.475653505195 | 5473.352876520997 | 5158.793997965102 | 5010.900922767433 | 4637.213176043864 | 5446.362236198391 | 5681.533557303313 |
| SE_36_1_22 | SE_36 | SE_36_1 | AAGCTCACCTCGGAAGAGGC | 407.5000924748004 | 379.76914416088056 | 379.2103152516649 | 382.58469593252175 | 467.7221801136684 | 295.41174746512735 | 350.82835976127507 | 344.1086705624513 | 225.54826056759313 | 436.67515995130873 | 397.9131030325979 | 275.081400016524 |
| SE_36_2_2 | SE_36 | SE_36_2 | GCTAGACTTTCATCTGGCAG | 900.6185554429763 | 1140.3068249672756 | 1063.5664310574039 | 981.1866011330659 | 1109.3818603149596 | 1230.1603560131405 | 829.7198324415432 | 1235.5770121569337 | 1426.9923414650477 | 1189.010327012662 | 1226.6052881093358 | 1620.310693768 2173 |
| SE_36_2_23 | SE_36 | SE_36_2 | TATAGCAAGCCACCCCCCAG | 1411.0774478781652 | 1662.9890944308033 | 1652.1324411875921 | 1739.8494505502774 | 1798.768293589074 | 2221.2191217026 | 1368.8762994366316 | 2102.4661629969555 | 1686.2840425899974 | 1970.5057718670907 | 2128.0721409973453 | 2247.07844064131 |
| SE_36_2_30 | SE_36 | SE_36_2 | CAAGCCATCCTGGCAACGCA | 699.0360628889528 | 685.5832444588527 | 703.1191261957954 | 862.7675285825235 | 549.3879575938327 | 835.9892560230144 | 783.4449260926633 | 566.2667408431548 | 552.3268428072556 | 1152.5599797713173 | 543.4624386551411 | 735.87176206952 |
| SE_36_2_36 | SE_36 | SE_36_2 | CACCTGGGATGAAATACCTG | 275.27931778882794 | 321.8043800521146 | 280.4576289882105 | 243.34468754891688 | 285.2999239242104 | 595.1550454795968 | 361.58996588892154 | 183.3985771678999 | 247.74789251322238 | 330.2401460065824 | 419.0412323971607 | 870.5107595459621 |
| SE_36_3_11 | SE_36 | SE_36_3 | GCTGATGTAAACCTTTCTGT | 1682.02165830024 | 1934.8238502512231 | 1874.3259852803646 | 1715.1245892485158 | 2007.705412596767 | 2105.1335669802334 | 1918.7943725593664 | 2324.6242332776587 | 1744.0030856486335 | 3120.1497238591005 | 2628.1045359586633 | 2488.5000912887235 |
| SE_36_3_17 | SE_36 | SE_36_3 | AGCAGGGCATTGAAGCCACA | 2726.78253368776 | 2471.4976144996253 | 2305.8752242516603 | 2573.98819236234 | 2513.608995168694 | 2870.9517041039066 | 2820.616966056141 | 2618.6291688406322 | 2335.4012806801966 | 3026.8368349212583 | 2785.391721228186 | 3065.3585546145146 |
| SE_36_3_21 | SE_36 | SE_36_3 | GGGAATGTATCTCAAGGTGG | 177.73940203688102 | 218.86695413482326 | 206.3931142906197 | 183.4844970288 6245 | 189.84641777856382 | 340.4598731782846 | 178.6426617189315 | 317.63877282687815 | 216.66840778934142 | 430.84310439269353 | 531.7245890081618 | 164.8167038073688 |
| SE_36_3_30 | SE_36 | SE_36_3 | CTGCTTAATTAATAAAGGGG | 15.172875783636185 | 14.990887269508443 | 20.738064115325425 | 16.917010364363207 | 18.030106716399914 | 40.71657516381521 | 18.294730416999005 | 2.836060471668555 | 7.991867500426528 | 34.26332640686395 | 11.73784964697929 | 15.088853165463343 |
| SE_36_3_5 | SE_36 | SE_36_3 | ATTTCCACTCCCAGACAACG | 37.93218945909046 | 42.973876839257535 | 25.675698428498144 | 19.519627343496005 | 53.02972563647034 | 90.09625142631451 | 8.60928490211718 | 41.59555358447214 | 51.50314611385985 | 9.477090282749604 | 48.12518355261509 | 38.30247342002233 |
| SE_36_4_12 | SE_36 | SE_36_4 | ATATAGCAAGCCACCCCCCA | 371.7354566990865 | 519.684092009626 | 455.2498836745248 | 426.82918457777936 | 520.7519057501387 | 823.8609144848567 | 313.1627383145124 | 408.3927079202719 | 511.4795200272978 | 399.49580576513716 | 279.3608215981071 | 405.07767344205433 |
| SE_36_4_30 | SE_36 | SE_36_4 | GGTCTCAGCAGGTTCAAACC | 253.60378095506195 | 289.82382054382987 | 275.51999467503776 | 283.68525072547527 | 260.90625013143404 | 356.9197652657844 | 236.75533480822244 | 233.5023121673777 | 254.85177473582374 | 348.4653196272547 | 190.1531642810645 | 196.15509115102344 |
| SE_36_4_35 | SE_36 | SE_36_4 | CCATCAGTCTCACCCAACGT | 18101.240809877967 | 15330.680714283968 | 15920.90807939412 | 17073.167383111173 | 16166.642157534347 | 15835.282498284643 | 17214.2651617833 | 16021.850957946224 | 15550.398185274373 | 17064.594564507905 | 17428.35915583485 | 16378.369770604095 |
| SE_36_4_44 | SE_36 | SE_36_4 | AATTATCCAGAGTGTCTGGA | 7.586437891818092 | 18.98845720804403 | 12.837849214249072 | 13.013084895664004 | 9.545350614564661 | 0.0 | 8.60928490211718 | 87.9178746217252 | 5.327911666951019 | 0.7290069448268927 | 34.03976397623994 | 10.446129114551544 |
| SE_36_4_7 | SE_36 | SE_36_4 | TCTTTGGGAGGCAAACAGAG | 176.65562519519273 | 176.89246978019963 | 203.43053370271608 | 186.08711400799527 | 220.6036586477166 | 136.8769973592086 | 255.05006522522146 | 244.84655405405192 | 134.08577695160065 | 451.2552988478466 | 178.4153146340852 | 378.3820101493115 |
| SE_360_1_12 | SE_360 | SE_360_1 | GGTGACAGAAAGAAAACAAG | 352.22747354869716 | 477.7096076550024 | 477.9630015151193 | 357.85983463076013 | 509.0853661101152 | 255.56148241118058 | 281.9540805443376 | 474.5674522592 0486 | 436.88875668998355 | 554.7742850132653 | 504.7275348201095 | 477.03989623118724 |
| SE_360_1_25 | SE_360 | SE_360_1 | TACACAGCACGTGTGATGGG | 6360.686283868627 | 5522.64287008691 | 5624.953009566363 | 5920.953627527122 | 5971.14710666656 | 4786.363357022959 | 6058.784249864965 | 5403.640552019154 | 5563.227765574688 | 5773.005996084163 | 5204.562533470617 | 3883.6386685877187 |
| SE_360_1_4 | SE_360 | SE_360_1 | TTTGGAAACAGGAAGCAGAA | 80.19948628493412 | 72.95565137827442 | 76.03956842285989 | 92.39290275921444 | 129.39253055298764 | 18.192512307236584 | 60.26499431482026 | 40.650200093915956 | 136.74973278507613 | 12.393118062057175 | 7.042709788187574 | 42.945197470934126 |
| SE_360_1_5 | SE_360 | SE_360_1 | AGACAGTGCTGACAGATGGG | 2179.475228635169 | 2137.700524631904 | 2056.0309280051206 | 2015.7268503383543 | 2253.7633395499893 | 1600.074772926951 | 2361.0963844056364 | 2022.1111162996797 | 1847.0093778763533 | 2027.3683135635886 | 1963.7422459396353 | 2563.9443571160405 |
| SE_360_1_51 | SE_360 | SE_360_1 | GAGGCTTCCTCTTACAGGAA | 107.2939073271416 | 178.8912547494674 | 179.729888999487 | 145.74655083143685 | 123.02896347661118 | 81.43315032763041 | 125.91079169346375 | 150.31120499843342 | 94.126439449468 | 80.1907639309582 | 130.29013108147012 | 102.13992912005955 |
| SE_360_2_18 | SE_360 | SE_360_2 | ATGCACTGCTCCCTCTGCAA | 1721.0376246010187 | 1762.9283428941928 | 1769.6481378411029 | 1752.8625354459414 | 1769.0716472326505 | 2002.0426639058926 | 1741.2278714531997 | 1502.1666964937779 | 1874.5369214889333 | 1618.3954175157016 | 1443.7555065784527 | 1901.1954988483812 |
| SE_360_2_27 | SE_360 | SE_360_2 | AGGGACCCGCAGCCTCAGCC | 4548.6114045657905 | 5248.809329297223 | 5334.620111951807 | 4706.83280676167 | 5144.943981250352 | 4820.149451307826 | 4484.361273390286 | 4712.587150422582 | 5053.524216103041 | 4855.915259491932 | 5094.226746789012 | 5026.909466124749 |
| SE_360_2_35 | SE_360 | SE_360_2 | GCCAGGATCAGCTGGCTGGA | 1107.6199322054415 | 1149.3013573289807 | 1150.4687949692438 | 1162.0684811827955 | 1334.2278970135937 | 1149.5935157953784 | 1091.2268613433525 | 1395.3417520609291 | 1411.89659174202 | 1111.7355908610114 | 1617.4756813537463 | 1164.1630557661333 |
| SE_360_2_41 | SE_360 | SE_360_2 | GGATCTGGCAGCCAGATTTG | 138.72343573610226 | 249.84812115847404 | 222.1935440927724 | 256.35777244458086 | 238.63376536411653 | 389.8395494407839 | 166.80489497852037 | 180.56251669623134 | 324.11462640618697 | 548.9422294546501 | 116.20471150509498 | 351.68634685656866 |
| SE_360_2_9 | SE_360 | SE_360_2 | GGGACCGTAATTCCTCTGTG | 482.28069455129304 | 543.6695116408396 | 580.6657952291118 | 458.06058832737295 | 543.0243905174563 | 513.7218951519664 | 491.8054000334439 | 493.4745220703286 | 599.3900625319897 | 338.9882293445051 | 424.9101572206503 | 424.8092506584295 |
| SE_360_3_12 | SE_360 | SE_360_3 | TCATGTGAAGTCCACGTTCC | 5477.4081578926625 | 4779.094861519292 | 4672.9771139866625 | 4935.863100925357 | 4823.5838438933415 | 5329.53979591045 | 4958.948103619496 | 5580.4216547531605 | 5004.685025822657 | 3949.759627072104 | 5581.347507138653 | 5806.887106677931 |
| SE_360_3_13 | SE_360 | SE_360_3 | GAACGTGGACTTCACATGAT | 4184.462385758522 | 3881.640410318053 | 3917.5190640712362 | 4339.863812703946 | 3554.052212156242 | 4693.668175267038 | 4118.466665050306 | 3578.1629617551603 | 3662.0512857510003 | 4331.030259216569 | 3921.615567055781 | 4249.253187597023 |
| SE_360_3_14 | SE_360 | SE_360_3 | TGGACTTCACATGATAGGTC | 82.36703996831072 | 74.95443634754221 | 87.88989077447442 | 104.10467916531204 | 72.12042686559965 | 73.63635933881474 | 92.54981269775968 | 24.57919075446081 | 94.126439449468 | 35.72134029651774 | 44.6038286585213 | 83.56903291641235 |
| SE_360_3_16 | SE_360 | SE_360_3 | GGTCTGGATCTGTTTCTGTT | 60.69150313454474 | 94.94228604022014 | 119.49075037877982 | 100.20075369661284 | 113.48361286204653 | 68.4384986796043 | 60.26499431482026 | 47.26767452780925 | 134.9737622294258 | 129.7632361791869 | 41.08247376442752 | 64.99813671276517 |
| SE_360_3_6 | SE_360 | SE_360_3 | GCAGACACTGAGGAACACAG | 6285.905681792134 | 5577.609456741775 | 5627.915590154266 | 5954.787648255849 | 6040.085749993971 | 5654.406087111104 | 6431.135821881533 | 6564.534638422148 | 5477.093193625 6475 | 6039.8225378908055 | 6987.541894846771 | 5707.068539583327 |
| SE_361_1_11 | SE_361 | SE_361_1 | TGACTGGCTCCTGAATTTCA | 607.9988081871356 | 685.5832444588527 | 797.9217050087116 | 662.3660211892978 | 706.3559454777849 | 540.5775085578871 | 633.8586009183773 | 719.4140063132568 | 1038.0547897776235 | 440.3201946754432 | 862.7319490529778 | 608.1968506694456 |
| SE_361_1_14 | SE_361 | SE_361_1 | AGAAAATGAAGTTCCTTCAT | 35.76463577571386 | 30.98116702365078 | 42.463655093285396 | 32.53271223916001 | 59.39329271284678 | 77.10159977828837 | 20.4470516425283 | 28.36060471668555 | 40.84732277995781 | 47.38545141374802 | 68.07952795247988 | 67.31949873822107 |
| SE_361_1_15 | SE_361 | SE_361_1 | CATTTTCTGCAGCCTTTGTC | 10.837768416882989 | 8.994532361705065 | 19.75053725269088 | 2.602616979132801 | 6.363567076376441 | 24.256683076315444 | 3.228481838293942 | 18.9070698111237 | 3.551941111300679 | 1.4580138896537853 | 17.606774470468935 | 6.964086076367696 |
| SE_361_1_8 | SE_361 | SE_361_1 | AAAAGTTTAGAGCACTCAAG | 5.4188842084414945 | 10.993317330972857 | 15.800429802152705 | 14.314393385230405 | 26.51486281823517 | 28.588233625657487 | 24.75169409358689 | 31.196665188354107 | 1.7759705556503396 | 24.78623612411435 | 5.868924823489645 | 9.285448101823595 |
| SE_361_1_9 | SE_361 | SE_361_1 | GAGCACTCAAGAGGTTTGAC | 368.48412617402164 | 353.78493956039927 | 359.459777998974 | 279.7813252567761 | 397.7229422735275 | 423.62564372565186 | 301.3249715741013 | 334.6551356568895 | 478.6240647477665 | 230.3661945652981 | 260.58026216294024 | 261.15322786378863 |
| SE_362_1_10 | SE_362 | SE_362_1 | TTACAAGAAGTTTAATGATA | 16.256652625324485 | 14.990887269508443 | 13.825376076883616 | 24.724861301761607 | 21.211890254588134 | 2.598930329605226 | 26.904015319116187 | 0.945353490556185 | 26.639558334755094 | 195.37386121360723 | 4.695139858791716 | 23.21362025455899 |
| SE_362_1_12 | SE_362 | SE_362_1 | ATGACTTTATATCTCGTGTA | 156.06386520311503 | 210.8718142577521 | 211.33074860379241 | 222.52375171585447 | 224.84603669863424 | 312.73794966249557 | 200.16587397422444 | 324.25624726077143 | 238.86803973497067 | 212.87002788945264 | 275.8394667040133 | 313.38387343654637 |

| | | | | | | | | | | | | | | | |
|---|---|---|---|---|---|---|---|---|---|---|---|---|---|---|---|
| SE_362_1_16 | SE_362 | SE_362_1 | CTGGGATGAGCCACTGAGCG | 395.5785472162291 | 517.6853070403582 | 512.5264417073283 | 439.84226947344337 | 534.539634415621 | 489.4652120756509 | 436.92120878244685 | 575.7202757487166 | 470.63219724734 | 486.9766391443643 | 460.1237061615882 | 538.5559899057686 |
| SE_362_1_17 | SE_362 | SE_362_1 | GTACCTGAGCCCAGGAGTCG | 314.2952840896067 | 435.73512330037875 | 462.1625717129666 | 420.3226421299473 | 383.9352136080452 | 355.1871450460476 | 312.0865777017478 | 408.3927079202719 | 539.0070636398781 | 631.3200142200891 | 505.9013197848074 | 538.5559899057686 |
| SE_362_1_3 | SE_362 | SE_362_1 | AAGGGATTGATAGTTAAACG | 536.469536635708 | 719.5625889364053 | 679.4184814925663 | 575.178352388349 | 637.4173021503734 | 828.1924650341988 | 567.1366429269692 | 544.5236105603626 | 585.1822980867869 | 879.1823754612326 | 448.3858565146089 | 503.7355595239301 |
| SE_363_1_12 | SE_363 | SE_363_1 | ACCGGCAAGCCTCGAACCCG | 2137.2079318093256 | 2088.730292884843 | 2167.6214634828243 | 2283.796399189033 | 2081.9470284878253 | 2125.925009617075 | 2251.3280019036424 | 1529.5819477199072 | 2798.9295957049353 | 1695.6701536673522 | 1946.1354714691663 | 2626.6211318033497 |
| SE_363_1_37 | SE_363 | SE_363_1 | GACCACTCACCACAGCCACC | 5270.4067811301975 | 4870.039577620976 | 4800.368079266518 | 4432.25671546316 | 4707.979042005836 | 4724.855339222301 | 4992.309082615199 | 4683.2811922153405 | 4391.087198845465 | 4242.820418892516 | 4826.603774837884 | 4970.03609650108 |
| SE_363_1_38 | SE_363 | SE_363_1 | ATGGGAATATAGAGGGCGGA | 435.6782903586962 | 552.6640440025446 | 668.5556860035863 | 482.7854496291346 | 578.0240094375267 | 909.6256153618292 | 359.43764466339223 | 594.6273455598404 | 505.26362308252163 | 556.961305847746 | 1044.6686185811568 | 928.5448101823596 |
| SE_363_1_40 | SE_363 | SE_363_1 | GCAATGGATGGGAATATAGA | 869.1890270340157 | 1014.3833719034046 | 1073.4416996837494 | 951.2565058730387 | 1011.807165143854 | 939.080159097355 | 793.1303716075452 | 1174.1290352707817 | 927.0566300494772 | 759.6252365096221 | 1294.6848160618158 | 754.4426582731671 |
| SE_363_1_41 | SE_363 | SE_363_1 | GGACACGCTGAGCTGTGGCC | 3727.10855856606 | 2963.198716939502 | 3087.996499458219 | 3546.0656340684413 | 3409.8113584250427 | 2876.149564763117 | 3852.654993697438 | 2972.1913743086457 | 2643.5321720855304 | 2817.61184175594 | 2967.3283907563646 | 2040.4772203757352 |
| SE_364_1_12 | SE_364 | SE_364_1 | GGAAGAGCAGTAAGAATTAA | 1235.5055995246607 | 1559.052276028878 | 1499.0657774792378 | 1314.3215744620645 | 1564.376906275875 | 1860.834115997342 | 1261.2602381601669 | 1653.4232549827675 | 2051.245991776142 | 1404.0673757365953 | 1868.66566379910 3 | 1142.1101165243022 |
| SE_364_1_15 | SE_364 | SE_364_1 | CACCGAGAAAGACAAGATTG | 3205.811897713988 | 3211.0480531287085 | 3013.9319847606284 | 3199.9175758437786 | 3060.8757637370677 | 3199.2832357440334 | 3064.905425153716 | 2908.852690441381 | 3316.625012677009 | 2761.4783070042695 | 2626.9307509939654 | 3435.6157976747304 |
| SE_364_1_18 | SE_364 | SE_364_1 | GCTCGAGAAACAGAGACGAA | 958.0587280524562 | 1214.2618688301839 | 1140.5935263428983 | 921.3264106130115 | 1205.8959609733354 | 1243.1550076611666 | 899.6702722712453 | 822.457536783881 | 1166.812655062273 | 1071.6402088955322 | 711.313688606945 | 1102.646962091552 |
| SE_364_1_7 | SE_364 | SE_364_1 | AGCTCAGCATCCTGAATCTT | 822.5866228414188 | 884.4623489009981 | 845.3229944151697 | 763.8680833754771 | 853.7785827471724 | 965.0694623934073 | 725.3322530033724 | 580.4470432014976 | 991.8795553307147 | 715.8848198200086 | 532.8983739728598 | 672.0343063694827 |
| SE_364_1_8 | SE_364 | SE_364_1 | CAAGAATTGTAGTTCGGGCG | 278.5306483138928 | 545.6682966101073 | 529.3143983721156 | 417.72002515081454 | 388.17759165896285 | 497.2620030644666 | 283.0302411571023 | 432.97189867473276 | 454.64846224648693 | 511.03386832365175 | 328.65979011542015 | 503.7355595239301 |
| SE_364_2_24 | SE_364 | SE_364_2 | ATGACAAGCTGCCTAAGAAG | 848.597267041938 | 1105.3280880050893 | 994.4395506729859 | 914.8198681651795 | 888.7782016672428 | 993.6576960190648 | 840.4814385691897 | 829.0750112177742 | 977.6717908855119 | 803.3656531992357 | 1092.793802133772 | 795.0664937186453 |
| SE_364_2_5 | SE_364 | SE_364_2 | AATCTTTACTAAACTCGCGG | 1.083776841688299 | 7.9951398770711695 | 10.862795488979984 | 2.602616979132801 | 15.9089176909411 | 0.8663101098684087 | 0.0 | 4.726767452780925 | 6.215896944776189 | 1.4580138896537853 | 12.911634611677218 | 1.1606810127279494 |
| SE_364_2_6 | SE_364 | SE_364_2 | GAATCTTTACTAAACTCGCG | 13.005322100259587 | 19.987849692677923 | 23.700644703229056 | 22.12224432262881 | 18.030106716399914 | 19.05882241710499 | 7.533124289352532 | 7.56282792444948 | 11.543808611727208 | 10.206097227576498 | 17.606774470468935 | 81.24767089095646 |
| SE_364_2_8 | SE_364 | SE_364_2 | GCTCAAGGACCCACAGTTTG | 766.2302270736274 | 687.5820294281206 | 747.5578350143498 | 870.5753795199219 | 610.9024393321383 | 868.0427300881455 | 790.9780503820159 | 956.6977324428592 | 650.8932086458494 | 719.5298545441431 | 680.7952795247988 | 649.9813671276517 |
| SE_364_2_9 | SE_364 | SE_364_2 | CTACTGTCTAAATCACAAGT | 69.36171786805113 | 99.93924846338962 | 118.50322351614528 | 114.51514708184324 | 121.96836896388177 | 146.40640856776108 | 114.07302495305264 | 27.415251226129364 | 79.03068972644012 | 330.9691529514093 | 29.344624117448227 | 46.42724050911798 |
| SE_364_3_10 | SE_364 | SE_364_3 | GATGTTGTCGTATCGCTCGG | 1525.957793097125 | 1491.093587073773 | 1323.2859959302891 | 1311.7189574829317 | 1271.6528207625586 | 1411.2191689756378 | 1570.1183340236207 | 1153.3312584785458 | 1415.4485328533206 | 1147.456931157529 | 1251.2547723679922 | 835.6903291641236 |
| SE_364_3_13 | SE_364 | SE_364_3 | TGAAGGAAAACAGGACGCTG | 2369.1361759306214 | 2572.436255447649 | 2659.409841074827 | 2255.167612418572 | 2605.8807177761523 | 2149.315382583522 | 2227.65246842282 | 2865.3664298757967 | 2583.149173193419 | 2462.5854596252434 | 2706.7481285934246 | 2589.4793393960554 |
| SE_364_3_2 | SE_364 | SE_364_3 | ACAAGCTCCATCACTCAAAG | 884.3619028176519 | 902.4514136244082 | 983.5767551840058 | 880.9858474364531 | 954.5350614564661 | 732.8983529486737 | 880.2993812414817 | 687.2719876343465 | 848.0259403230372 | 1000.1975283024967 | 703.0971938540595 | 1064.3444886715297 |
| SE_364_3_3 | SE_364 | SE_364_3 | CACTACACAGAAAGATTGTG | 1917.2012329466008 | 1795.9082948871114 | 1794.3363094069666 | 2013.1242333592215 | 2025.7355193131668 | 1912.8127225894464 | 1867.1386631466635 | 2087.3405071480565 | 2046.8060653870164 | 1488.6321813365148 | 2398.042682877869 | 1623.7927368064013 |
| SE_364_3_4 | SE_364 | SE_364_3 | AACTGCCCTATTCAGCTGGG | 316.4628377729833 | 312.8098476904095 | 309.0959080046123 | 286.2878677046081 | 337.2690550479513 | 149.0053388973663 | 276.5732774805144 | 398.9391730147101 | 260.17968640277473 | 318.5760348893521 | 407.30338275018136 | 383.0247342002233 |
| SE_364_4_18 | SE_364 | SE_364_4 | GATTCACCGAGTTCTCTAGG | 18.42420630870108 | 32.97995199291857 | 22.713117840594514 | 33.834020728726415 | 34.999618920070425 | 90.96256153618292 | 18.294730416999005 | 40.650200093915956 | 38.1833669464823 | 1.4580138896537853 | 14.085419576375148 | 3.482043038183848 |
| SE_364_4_19 | SE_364 | SE_364_4 | CTACCACATTCAATGCCCGG | 286.1170862057109 | 381.76792913014833 | 402.91095995489394 | 337.0388987976977 | 475.14634170277424 | 357.7860753756528 | 405.7125510122721 | 338.43654961911426 | 402.2573308548019 | 657.5642642338572 | 433.1266519735358 | 191.51236710011165 |
| SE_364_4_2 | SE_364 | SE_364_4 | TCTCCTCCCACTTTTAGGGA | 15.172875783636185 | 20.98724217731182 | 27.650752153767232 | 15.615701874796805 | 36.06021343279983 | 1.7326202197368175 | 18.294730416999005 | 62.393330376708214 | 10.655823333902038 | 28.431270848248815 | 11.7378496469792 9 | 6.964086076367696 |
| SE_364_4_22 | SE_364 | SE_364_4 | TCAGACTATGGCATGTTCTG | 1638.670584632708 | 1395.151908548919 | 1393.4004031773416 | 1508.2165394074582 | 1618.4672264250746 | 1331.5186388677441 | 1542.1381580917398 | 1429.3744777209517 | 1510.4629575806139 | 1776.5899245431374 | 1828.7569749993734 | 1233.8039165298103 |
| SE_364_4_5 | SE_364 | SE_364_4 | CCAGGAGCCGTCCCTAAAAG | 254.68755779675024 | 307.81288526724 | 298.2331125156323 | 300.6022610898385 | 303.33003064061035 | 560.5026410848604 | 285.1825623826316 | 535.0700756548007 | 256.62774529147407 | 340.4462432341589 | 624.4536012192982 | 506.05692154938595 |
| SE_365_1_1 | SE_365 | SE_365_1 | TTAGTGGTTACCATATCGGA | 159.31519572817993 | 159.90279754142338 | 152.07913684571977 | 154.85571025840164 | 149.54382629484635 | 173.26202197368175 | 159.27177068916782 | 116.27847933841076 | 111.88614500597139 | 245.67534040666283 | 103.29307689341775 | 133.47831646371418 |
| SE_365_1_10 | SE_365 | SE_365_1 | GCCAGCCTAGTAATGACTGC | 2198.9832117855585 | 2027.7673513221753 | 1970.1160909559153 | 1867.3776825277846 | 1857.1009917891913 | 1889.4223496229995 | 2139.407298176119 | 1887.8709206407013 | 1803.4980992629198 | 1982.169882984321 | 2358.1339940781395 | 1706.2010887100857 |
| SE_365_1_14 | SE_365 | SE_365_1 | AAGGTGTGAGAATGATTAAG | 695.7847323638879 | 761.537073291029 | 674.4808471793935 | 750.854998479813 | 720.1436741432672 | 841.1871166822249 | 794.2065322203098 | 727.9221877282624 | 852.465866712163 | 980.5143407921706 | 613.8895365370169 | 1197.822805135244 |
| SE_365_1_21 | SE_365 | SE_365_1 | ATCATTCCCTATCTTCCTAG | 53.105065242726646 | 102.9374259172913 | 114.55311606560711 | 92.39290275921444 | 69.99923784014085 | 101.35828285460381 | 86.0928490211718 | 103.04353047062416 | 202.4606433441387 | 13.851131951710961 | 71.60088284657367 | 94.01516203096391 |
| SE_365_1_6 | SE_365 | SE_365_1 | GAAAGAACAGACACAAATGG | 2286.769135962311 | 2523.466023700588 | 2426.353501493075 | 2290.302941636865 | 2530.5785073723646 | 2645.7110755381204 | 2321.2784417333446 | 2236.706358655934 | 2424.1998084627135 | 2344.4863345632866 | 2156.2429801500957 | 3196.515509052773 |
| SE_366_1_10 | SE_366 | SE_366_1 | GCAGGAAACAGCTTTGAGAG | 3038.91026409399 | 3377.946598062569 | 3488.932405687844 | 3197.314958864646 | 3527.537349338007 | 3650.6308029854745 | 3206.9586260386495 | 2854.0221879891224 | 3524.4135676880987 | 3078.5963280039678 | 2317.051520313712 | 3740.874904022181 |
| SE_366_1_14 | SE_366 | SE_366_1 | ATAGATGAGGGAATTCGAAG | 136.55588205272565 | 180.8900397187352 | 205.4055874279851 5 | 154.85571025840164 | 119.84717993842297 | 240.83421054341764 | 158.19561007640317 | 62.393330376708214 | 198.90870223283804 | 157.46550008260883 | 77.46980767006332 | 80.08698987822851 |
| SE_366_1_21 | SE_366 | SE_366_1 | TGAGGTCCCAAAAATTCGAG | 4322.1020446529365 | 4883.031679921217 | 4920.846356507933 | 4428.352789994461 | 4621.010291962025 | 5510.598608872948 | 4529.560019126401 | 5353.536817019675 | 5208.921639722446 | 4511.823981533638 | 5135.309220553439 | 5900.902268708895 |
| SE_366_1_24 | SE_366 | SE_366_1 | CATGCTGTCTACTTACCCTT | 7394.609390839263 | 6643.961237846142 | 6800.10997610147 | 6960.699110690676 | 6630.836893584251 | 7487.518279592657 | 7235.027799616725 | 6271.475056349731 | 6248.75240005572 | 6464.104579780057 | 6523.896833791089 | 8338.332395437588 |
| SE_366_1_5 | SE_366 | SE_366_1 | GTAGTGAGATTGCAGCCAGC | 1298.364656342582 | 1255.2369607001735 | 1294.6477169138873 | 1405.4131687317124 | 1429.6814031592403 | 1244.021317771035 | 1369.9524600493962 | 1505.0027569654465 | 1334.6418725712301 | 1493.7352299503032 | 1383.8924733788583 | 958.7225165132862 |
| SE_366_2_22 | SE_366 | SE_366_2 | AATGGCTCCAGATGTCTGGG | 122.46678311077778 | 171.89550735703014 | 172.81720096104522 | 186.08711400799527 | 191.9676068040226 | 204.44918592894447 | 123.75847046793446 | 218.37665631847872 | 295.69909751578155 | 194.64485426878034 | 375.6111887033373 | 159.01329874372908 |
| SE_366_2_27 | SE_366 | SE_366_2 | TGTCTGGGAGGCAGGAAGGG | 398.829877741294 | 539.6719417023039 | 545.1148281742683 | 359.1611431203265 | 497.41882647009174 | 460.87697844999343 | 325.00050505492356 | 494.41987556088475 | 603.8299889211155 | 559.8773336270535 | 360.3519841622642 | 431.77333673479717 |
| SE_366_2_31 | SE_366 | SE_366_2 | ACACAGAGAGGAGCTGGCTG | 3061.6695777694445 | 3396.9350552706132 | 3251.9259586555536 | 3019.035695794049 | 3135.1173796281264 | 3483.4329517808715 | 2905.6336544645483 | 2811.481280914094 | 3085.748840442465 | 3540.7867310242177 | 2466.122210830349 | 3630.6102078130257 |
| SE_366_2_46 | SE_366 | SE_366_2 | GGAACAATGACTCTGAGAAC | 497.4535703349292 | 411.7497036691652 | 409.8236479933358 | 437.2396524943106 | 487.8734758555271 | 448.7486369118357 | 437.99736939521154 | 604.0808804654022 | 348.97821418529173 | 312.0149723859101 | 575.1546327019852 | 365.61451900930405 |
| SE_366_2_6 | SE_366 | SE_366_2 | ATTGTTGCTGAGTTGGGTGG | 460.60515771752705 | 465.71689783939564 | 534.2520326852883 | 494.4972260352322 | 494.23704293190355 | 399.36896064933643 | 445.5304936845641 | 554.9224989564806 | 523.023328639025 | 502.28578498572904 | 558.7216431962142 | 631.4104709240045 |
| SE_366_3_1 | SE_366 | SE_366_3 | TATGCTGGCAGCTTAAGAAG | 1438.1718689203726 | 1814.8967520951555 | 1670.8954515776486 | 1799.7096410703318 | 1920.7366625529555 | 1762.9410735822119 | 1566.8898521853266 | 1959.7177859229714 | 1512.2389281362641 | 1914.3722371154201 | 2000.129579845271 | 1958.0688684720508 |
| SE_366_3_2 | SE_366 | SE_366_3 | GGTCAGTTCCAGGAACTGGA | 73.69682523480432 | 111.93195827899638 | 140.22881449410525 | 70.27065843658562 | 132.57431409117584 | 127.34758615065608 | 66.72195799140815 | 98.31676301784324 | 81.69464555991561 | 90.39686115853469 | 99.77172199932397 | 127.67491140007444 |
| SE_366_3_3 | SE_366 | SE_366_3 | TTAGCTGTGTGAAGAGAGGA | 462.77271140090363 | 454.72358050842274 | 453.2748299492557 | 543.9469486387554 | 459.2374240118331 | 236.50265999407557 | 481.04379390579743 | 551.1410849942558 | 483.06399113689235 | 476.0415349719609 | 664.3622900190278 | 409.72039749296613 |
| SE_366_3_4 | SE_366 | SE_366_3 | TAGAGCCTAGTTGTGTACAT | 2616.2372958355536 | 2199.6628586792053 | 2207.1225379882057 | 2387.901078354345 | 2232.551449295401 | 1948.3314370940511 | 2462.2554820055134 | 2098.684749034731 | 2154.252284003862 | 2020.8072510601464 | 1840.4948246463528 | 1871.0177925174544 |
| SE_366_3_7 | SE_366 | SE_366_3 | AGTTGTGTACATGGGATGGA | 379.3218945909046 | 435.73512330037875 | 479.9380552403884 | 416.41871666124814 | 380.753430069857 | 271.1550643888119 | 368.04692956550946 | 371.5239217885807 | 602.0540183654651 | 288.6867501514495 | 252.36376741005475 | 450.3442329384444 |
| SE_367_1_0 | SE_367 | SE_367_1 | GCTGTGCTCTCCAGAACCAT | 1528.1253467805016 | 1561.0510609981459 | 1514.8662072813906 | 1644.8539308119302 | 1619.527820937804 | 1121.0052821697209 | 1503.3963760322126 | 1396.2871055514852 | 1507.7990017471384 | 1065.808153336917 | 1394.4565380611398 | 754.4426582731671 |
| SE_367_1_13 | SE_367 | SE_367_1 | AATGAGAGTGACCAGCCTCA | 871.3565807173924 | 948.4234679175675 | 880.8739614700132 | 864.0688370720899 | 958.7774395073837 | 729.4331125092001 | 779.1402836416048 | 897.1404625378195 | 887.0972925473446 | 387.8316946479069 | 545.810008584537 | 557.1268861094158 |
| SE_367_1_19 | SE_367 | SE_367_1 | AGAATGATCTGAAAACGAAA | 79.11570944324582 | 105.935603371193 | 140.22881449410525 | 143.14393385230406 | 88.02934455654076 | 197.51870504999718 | 88.2451702467011 | 140.85767009287156 | 179.3730261206843 | 125.38919451022554 | 106.81443178751154 | 100.9792481073316 |
| SE_367_1_20 | SE_367 | SE_367_1 | GAAATGTTCTACCAGCTACA | 82.36703996831072 | 84.94836119388117 | 92.82752508764715 | 87.18766880094883 | 108.18064029839948 | 79.7005301078936 | 64.56963676587885 | 59.557269905039654 | 103.89427750554486 | 64.15261114476655 | 30.518409082146157 | 153.20989368008932 |
| SE_367_1_3 | SE_367 | SE_367_1 | ACAGCTGGATAGCAGCATCA | 2418.989910648283 | 2161.6859442631176 | 2313.7754391527365 | 2309.8225689803608 | 1922.8578515784145 | 1826.1817116026057 | 2047.933646091124 | 1652.4779014922115 | 1854.1132600989545 | 738.4840351096423 | 1136.2238458275954 | 1041.1308684169705 |
| SE_367_2_16 | SE_367 | SE_367_2 | GTTACGGAACAGGTAACTAA | 373.90301038246315 | 462.71872038549395 | 438.46192700973756 | 447.65012041084174 | 445.44969534635084 | 251.22993186183854 | 430.464245105859 | 569.1028013148234 | 531.9031814172768 | 348.4653196272547 | 570.4594928431935 | 359.8111139456643 |
| SE_367_2_17 | SE_367 | SE_367_2 | GAGGGCTAAGGTATCCCAAG | 1842.4206308701082 | 2072.740013130701 | 1995.7917893844135 | 1821.8318853929607 | 1857.1009917891913 | 1643.3902784203713 | 1832.7015235381946 | 1718.6526458311444 | 1964.2234345492755 | 2323.345133163307 | 1925.0073421046036 | 1383.5317671717157 |
| SE_367_2_18 | SE_367 | SE_367_2 | TTATTCCCTCATCGTGTGCG | 107.2939073271416 | 95.94167852485404 | 106.65290116453076 | 45.545797134824014 | 89.08993906927017 | 157.66843999605038 | 96.85445514881827 | 63.33868386726439 | 148.29354139680336 | 173.50365286880046 | 119.72606639918877 | 71.96222278913287 |
| SE_367_2_22 | SE_367 | SE_367_2 | TGGCAATAGTCCCACCCGCT | 131.13699784428417 | 159.90279754142338 | 147.14150253254707 | 102.80337067574564 | 89.08993906927017 | 146.40640856776108 | 142.05320088493346 | 119.11453981007931 | 67.4868811147129 | 190.99981954464587 | 115.03092654039705 | 106.78265317097134 |
| SE_367_2_3 | SE_367 | SE_367_2 | GCTCCTTGAAGACGGAGGGT | 85.61837049337561 | 168.89732990312845 | 152.07913684571977 | 75.47589239485123 | 156.9679878839522 | 96.16042219539337 | 51.655709412703075 | 83.19110716894428 | 157.17339417505505 | 139.2403264619365 | 99.77172199932397 | 39.46315443275028 |
| SE_368_1_16 | SE_368 | SE_368_1 | AGATCTAGAGATGTAATCGG | 94.288585226882 | 179.89064723410132 | 144.17892194464343 | 178.27926307059687 | 103.93826224748186 | 319.66843054144283 | 157.11944946363852 | 98.31676301784324 | 77.25471917078977 | 324.40809044796725 | 26.997054188052367 | 49.909283547301825 |
| SE_368_1_17 | SE_368 | SE_368_1 | GAGCAACCAGAGACCCAGGG | 288.2846398890875 | 451.72540305452105 | 466.1126791635047 7 | 270.6721658298113 | 259.84565561870465 | 222.64169823618104 | 232.45069235716386 | 190.96140509234937 | 518.5834022498991 | 334.61418767554375 | 154.93961534012664 | 113.74673924733905 |
| SE_368_1_27 | SE_368 | SE_368_1 | GGTGTTGCGACTCAAAAGAA | 145.22609678623206 | 227.86148649652833 | 220.2184903675033 | 214.71590077845607 | 160.14977142214042 | 161.133680435524 | 236.75533480822244 | 247.68261452572048 | 340.9863466848652 | 340.4462432341589 | 129.1163461167722 | 89.37243798005211 |
| SE_368_1_8 | SE_368 | SE_368_1 | TATTGCTGAAAATACTCCAG | 135.47210521103736 | 205.8748518345826 | 238.98150075755964 | 169.17010364363205 | 173.9375000876227 | 295.41174746512735 | 139.90087965940418 | 113.4424188667422 | 222.88430473411762 | 82.37778476543888 | 95.07658214053225 | 248.38573672378118 |
| SE_369_1_1 | SE_369 | SE_369_1 | ACAAAATGCCTTAGACAGCG | 1244.175814258167 | 1343.1834993479565 | 1285.7599751501764 | 1242.7496075359124 | 1258.9256866098058 | 931.2833681085394 | 1188.0813164921708 | 1146.7137840446524 | 1163.2607139509723 | 1747.4296467500617 | 1266.5139769090654 | 1381.2104051462597 |
| SE_369_1_10 | SE_369 | SE_369_1 | CTGCAGCAATGTGTGAGCAG | 159.31519572817993 | 66.95929647047105 | 86.90236391183987 | 102.80337067574564 | 57.27210368738796 | 297.1443676848642 | 213.0798013274002 | 119.11453981007931 | 285.0432741818795 | 14.580138896537854 | 46.95139858791716 | 106.78265317097134 |
| SE_369_1_11 | SE_369 | SE_369_1 | GAGCTGGTATCACAAGATAC | 576.569279778175 | 502.6944197708498 | 482.900635828292 | 501.00376848306416 | 405.1471038626334 | 476.4705604276248 | 488.57691819515 | 397.0484660335977 | 504.37563780469645 | 338.2592223996782 | 440.1693617617234 | 762.5674253622627 |
| SE_369_1_24 | SE_369 | SE_369_1 | TCAAAAAGCCAGCATGAGGT | 988.4044796197286 | 1157.2964972060518 | 1222.5582559415654 | 1042.3481001426867 | 1253.6227140461588 | 1060.3635744789324 | 933.0312512669493 | 1075.8122722529386 | 1166.812655062273 | 1003.1135560818043 | 983.6318004168645 | 1104.9683241170078 |
| SE_369_1_25 | SE_369 | SE_369_1 | ACAGATCTCCTACCTCATGC | 2526.283817975425 | 2298.6027146579613 | 2250.573719944126 | 2099.010593670604 | 1967.4028211130494 | 1820.1175408335268 | 2293.298265801464 | 1985.2423301679885 | 2147.1484017812604 | 2043.4064663497802 | 1827.5831900346755 | 1976.639764675698 |
| SE_369_2_35 | SE_369 | SE_369_2 | ATAATCTCCTTCCCTCCACC | 130.05322100259588 | 109.93317330972857 | 96.77763253818532 | 130.13084895664005 | 134.69550311663465 | 237.36897010394398 | 151.73864639981528 | 58.61191641448347 | 87.9105425046918 | 195.37386121360723 | 69.25331291717781 | 99.81856709460365 |
| SE_369_2_37 | SE_369 | SE_369_2 | CTGCCACCAATGATATGAGG | 176.65562519519273 | 182.888824688003 | 202.44300684008152 | 199.10019890365928 | 309.69359771698674 | 185.39036351183947 | 203.39435581251837 | 313.8573588646534 | 249.5238630688727 | 198.2898889929148 | 233.5832079748878 7 | 350.5256658438407 |
| SE_369_2_49 | SE_369 | SE_369_2 | GGAGCCGGGCCTTTCCCCCA | 170.15296414506292 | 244.85115873530455 | 283.42020957611413 | 275.8773997880769 | 258.7850611059753 | 388.97323933091553 | 174.33801926787288 | 306.29453094020397 | 316.12275890576046 | 217.97307650324092 | 288.75110131569056 | 193.83372912556754 |
| SE_369_2_6 | SE_369 | SE_369_2 | GGGAAAAGAGCCGGACTACG | 139.80721257779055 | 202.87667438068092 | 223.18107095540694 | 148.34916781056964 | 200.45236290585788 | 155.93581977631356 | 140.9770402721688 | 181.50787018678753 | 227.32423112324346 | 201.93492371704926 | 178.4153146340852 | 181.06623798556012 |
| SE_369_2_7 | SE_369 | SE_369_2 | GCAATGGTGGGATGGCCTGG | 5551.104983127467 | 5048.930832370444 | 5221.054522748835 | 5650.281461697311 | 5362.36585635988 | 5274.96225898874 | 5969.462919005499 | 5465.088528905306 | 4693.890178583847 | 5540.452780684384 | 5352.459439022557 | 5310.115633230369 |
| SE_369_3_16 | SE_369 | SE_369_3 | GATAAGGAATCAGAGACCGA | 200.4987157123353 | 250.84751364310793 | 306.13332741670865 | 221.22244322628808 | 307.572408691528 | 206.1818061486813 | 202.31819519975372 | 147.47514452676486 | 190.9168347324115 | 310.55695849625624 | 273.4918967746175 | 73.12290380186082 |
| SE_369_3_2 | SE_369 | SE_369_3 | ATATAGGAGAAGCCGTAAGA | 9.75399157519469 | 10.993317330972857 | 24.688171565863602 | 32.53271223916001 | 28.63605184369398 | 18.192512307236584 | 19.370891029763655 | 14.180302358342775 | 21.311646667804077 | 40.0953819654791 | 3.521354894093787 | 0.0 |
| SE_369_3_3 | SE_369 | SE_369_3 | AATATAGGAGAAGCCGTAAG | 10.837768416882989 | 19.987849692677923 | 45.42623568118903 | 31.23140374959361 | 26.51486281823517 | 12.994651648026132 | 19.370891029763655 | 41.59555358447214 | 39.07135222430747 | 39.36637502065221 | 41.08247376442752 | 69.64086076367697 |
| SE_369_3_7 | SE_369 | SE_369_3 | TGAGAATTCCTCAGAGAGCA | 326.21682934817795 | 441.7314782081821 | 405.87354054279757 | 338.3402072872641 | 453.9344514481861 | 283.28340592696964 | 356.2091628250983 | 315.7480658457658 | 293.9231269601312 | 276.2936320893923 | 342.74520969179525 | 284.3668481183476 |
| SE_369_3_9 | SE_369 | SE_369_3 | TTACCATACCATGCTCTCTG | 229.76069043791938 | 254.84508358164354 | 277.49504840030687 | 225.12636869498726 | 177.11928362581094 | 254.69517230131217 | 242.13613787204568 | 214.595242356254 | 159.83735000853056 | 182.98074315155006 | 98.59793703462604 | 273.9207190037961 |
| SE_369_4_14 | SE_369 | SE_369_4 | TTATAGACTCCAAAATTAGG | 7.586437891818092 | 10.993317330972857 | 10.862795488979984 | 3.9039254686992013 | 5.3029725636470335 | 0.0 | 34.43713960846872 | 21.743130282792254 | 46.17523444690883 | 2.9160277793075706 | 0.0 | 0.0 |
| SE_369_4_15 | SE_369 | SE_369_4 | TTTCCTCCTAGACCACAGAG | 10.837768416882989 | 15.990279754142339 | 6.912688038441808 | 6.506542447832002 | 21.211890254588134 | 2.598930329605226 | 7.533124289352532 | 23.633837263904624 | 24.863587779104755 | 0.0 | 9.390279717583432 | 0.0 |
| SE_369_4_20 | SE_369 | SE_369_4 | GAAACTAGAAAAGCACCAGG | 66.11038734298623 | 99.93924846338962 | 149.11655625781614 | 96.29682822791364 | 143.1802592184699 | 65.83956834999907 | 99.00677637434757 | 241.06514009182717 | 121.65398306204825 | 156.00748619295504 | 171.37260484589763 | 148.56716962917753 |
| SE_369_4_26 | SE_369 | SE_369_4 | CTTTCCTCTCTGTCAACCTG | 37.93218945909046 | 35.97812944682026 | 29.62580587903632 | 37.737946197425615 | 12.727134152752882 | 0.8663101098684087 | 35.513300221233365 | 19.852423301679885 | 55.05508722516053 | 67.06863892407412 | 18.780559435166865 | 9.285448101823595 |

| | | | | | | | | | | | | | | | |
|---|---|---|---|---|---|---|---|---|---|---|---|---|---|---|---|
| SE_369_4_34 | SE_369 | SE_369_4 | TGATATGCCCCAAAAGGGAG | 476.8618103428515 | 603.6330607188733 | 582.640848954381 | 485.38806660826737 | 474.0857471900448 | 446.1497065822305 | 365.89460833998015 | 545.4689640509188 | 696.1804578149331 | 435.94615300648184 | 620.9322463252045 | 531.5919038294008 |
| SE_37_1_26 | SE_37 | SE_37_1 | TATCTGGTAGGTGCAGGGCA | 1623.4977088490718 | 1326.1938271091803 | 1391.4253494520726 | 1581.0898148231765 | 1501.80183002484 | 1315.0587467802445 | 1642.221095078852 | 1159.0033794218828 | 1491.8152667462853 | 1135.063813095472 | 943.7231116171349 | 1085.2367469006326 |
| SE_37_1_28 | SE_37 | SE_37_1 | TGGATTATCTGGTAGGTGCA | 85.61837049337561 | 97.94046349412183 | 108.62795488979984 | 100.20075369661284 | 128.33193604025823 | 120.41710527170882 | 92.54981269775968 | 136.13090264009065 | 109.22218917249589 | 72.17168753786237 | 116.20471150509498 | 149.7278506419055 |
| SE_37_1_30 | SE_37 | SE_37_1 | CTGTACATGGATTATCTGGT | 857.2674817754445 | 849.4836119388118 | 913.4623479369533 | 848.4531351972931 | 988.4740858638071 | 606.4170769078861 | 1064.3228460242362 | 1321.6041797975467 | 758.339427262695 | 1094.239424185166 | 1235.9955678269193 | 675.5163494076666 |
| SE_37_1_34 | SE_37 | SE_37_1 | GTCTGATCTCCGAATAACTG | 328.3843830315546 | 358.78190198356873 | 332.79655270784133 | 299.3009526002721 | 377.5716465316688 | 631.54007009407 | 484.27227574409136 | 446.2068475425193 | 371.17784613092095 | 383.45765297894553 | 390.87039324441037 | 265.7959519147004 |
| SE_37_1_35 | SE_37 | SE_37_1 | TTATTCGGAGATCAGACATC | 160.39897256986825 | 184.8876096572708 | 209.35569487852334 | 171.77272062276487 | 121.96836896388177 | 101.35828285460381 | 111.92070372752333 | 98.31676301784324 | 222.88430473411762 | 96.95792366197672 | 71.60088284657367 | 170.62010887100857 |
| SE_37_2_1 | SE_37 | SE_37_2 | AATCCTGCCCCCGTTCCTAG | 95.37236206857031 | 86.94714616314897 | 98.75268626345441 | 93.69421124878083 | 99.69588419656424 | 217.4438375769706 | 116.22534617858193 | 73.73757226338243 | 40.84732277995781 | 40.0953819654791 | 69.25331291717781 | 76.60494684004466 |
| SE_37_2_11 | SE_37 | SE_37_2 | GATGTGAGTTGGAACTTGCG | 1457.679852070762 | 1423.1348981186682 | 1377.599973375189 | 1540.7492516466182 | 1563.3163117631454 | 1561.957128092741 | 1550.747442993857 | 1614.663761869964 | 1530.8866189705927 | 1364.701000715943 | 1863.9705239403113 | 1865.2143874538147 |
| SE_37_2_18 | SE_37 | SE_37_2 | TCAAACCCTGGAAAAGAGGT | 343.5572588151908 | 422.74302100013807 | 381.185368976934 | 351.3532921829281 | 431.66196668086855 | 198.3850151598656 | 444.4543330717994 | 376.2506892413616 | 598.5020772541644 | 188.0837917653383 | 480.078050561453 | 193.83372912556754 |
| SE_37_2_19 | SE_37 | SE_37_2 | GTTAGACAGTAGAAAATGAA | 133.30455152766078 | 115.92952821753195 | 141.2163413567398 | 122.32299801924164 | 94.3929116329172 | 100.4919727447354 | 148.51016456152135 | 158.81938641343908 | 149.18152667462851 | 113.72508339299526 | 143.20176569314734 | 55.71268861094157 |
| SE_37_2_4 | SE_37 | SE_37_2 | GTGATAAGCATGACTGGAGG | 770.5653344403805 | 893.4568812627032 | 971.7264328323913 | 771.6759343128755 | 767.8704272160904 | 866.3101098684087 | 675.8288648161986 | 694.834815558796 | 908.4089392151487 | 890.8464865784629 | 643.2341606544651 | 623.2857038349089 |
| SE_37_3_0 | SE_37 | SE_37_3 | ATATGGAGTTCAAAATCTAC | 165.81785677830973 | 157.9040125721556 | 114.55311606560711 | 126.22692348794084 | 111.36242383658771 | 135.14437713947177 | 145.28168272322742 | 85.08181415005664 | 118.10204195074758 | 100.60295838611118 | 71.60088284657367 | 53.39132658548567 |
| SE_37_3_10 | SE_37 | SE_37_3 | CCCCATATACTAATTCTTAG | 17.340429467012783 | 15.990279754142339 | 21.725590977959968 | 10.410467916531204 | 27.575457330964575 | 22.524062856578627 | 0.0 | 20.79777679223607 | 39.95933750213264 | 0.0 | 75.12223774066746 | 2.3213620254558999 |
| SE_37_3_12 | SE_37 | SE_37_3 | TAGTATATGGGGCCAAAGCA | 456.27005035077383 | 493.6998874091447 | 535.2395595479229 | 437.2396524943106 | 532.4184453901622 | 406.2994415282837 | 550.9942337354995 | 536.9607826359131 | 570.086548363759 | 605.8047711511477 | 524.6818792199742 | 431.77333673479717 |
| SE_37_3_5 | SE_37 | SE_37_3 | TTAGCCTCTTTCCTCTGTAG | 156.06386520311503 | 110.93256579436247 | 100.72773998872348 | 148.34916781056964 | 144.2408537311993 | 110.02138395328791 | 225.99372868057597 | 194.7428190545741 | 106.55823333902038 | 82.37778476543888 | 109.1620017169074 | 176.42351393464833 |
| SE_37_3_6 | SE_37 | SE_37_3 | GTTAGCCTCTTTCCTCTGTA | 171.23674098675122 | 185.8870021419047 | 194.54279193900518 | 156.15701874796804 | 214.24009157134017 | 148.1390287874979 | 152.81480701257993 | 206.08706094124832 | 144.74160028550267 | 156.73649313778193 | 183.11045449287693 | 106.78265317097134 |
| SE_370_1_16 | SE_370 | SE_370_1 | TAGACCCCTCAGCAGCGCTG | 666.5227576383038 | 736.5522611751815 | 710.0318142342371 | 668.8725636371298 | 773.1733997797376 | 656.6630632802538 | 657.5341343991996 | 698.6162295210207 | 760.1153978183453 | 798.2626045854474 | 689.0117742776844 | 788.1024076422776 |
| SE_370_1_17 | SE_370 | SE_370_1 | GTAACTCCTCCTGGAAACTT | 136.55588205272565 | 197.87971195751143 | 166.89203978523796 | 157.45832723753446 | 161.21036593486983 | 270.28875427894354 | 151.73864639981528 | 174.89039575289422 | 209.56452556674006 | 256.6104445790662 | 158.4609702342204 | 141.60308355280984 |
| SE_370_1_2 | SE_370 | SE_370_1 | TTCTTGCTCTCCGGTTTCCA | 0.0 | 0.0 | 4.93763431317272 | 0.0 | 5.3029725636470335 | 0.0 | 0.0 | 0.0 | 0.0 | 0.0 | 0.0 | 0.0 |
| SE_370_1_22 | SE_370 | SE_370_1 | GTTTAACCGCCTCATCCGTG | 224.34180622947787 | 232.85844891969782 | 312.0584885925159 | 219.92113473672168 | 263.02743915689285 | 252.96255208157535 | 213.0798013274002 | 201.36029348846742 | 240.644010290621 | 190.99981954464587 | 260.58026216294024 | 327.31204558928175 |
| SE_370_1_25 | SE_370 | SE_370_1 | AGTGCTCCTATGACACCAGG | 325.13305250648966 | 338.7940522908908 | 340.6967676089177 | 356.55852614119374 | 342.5720276115984 | 358.6523854855212 | 419.7026389782125 | 357.34361943023794 | 452.87249169083657 | 240.57229179287458 | 286.4035313862947 | 338.9188557165612 |
| SE_370_2_12 | SE_370 | SE_370_2 | CAGTTCCAAACATATCTAGG | 201.5824925540236 | 236.8560188582334 | 248.8567693839051 | 247.2486130176161 | 264.0880336696223 | 169.79678153420812 | 207.69899826357695 | 270.37109829906893 | 321.4506705727115 | 196.10286815843412 | 410.82473764427516 | 521.1457747148493 |
| SE_370_2_13 | SE_370 | SE_370_2 | CCTCACCTCCTCAGAAGGTG | 344.64103565687907 | 324.8025575060163 | 246.881715658636 | 277.1787082776433 | 316.0571647933632 | 323.13367098091646 | 266.8878319656326 | 338.43654961911426 | 255.7397600136489 | 367.4195001927539 | 254.7113373394506 | 356.32907090748046 |
| SE_370_2_17 | SE_370 | SE_370_2 | TGCCCCACGATAATAAGGCA | 449.767389300644 | 476.7102151703685 | 429.5741852460267 | 458.06058832737295 | 471.964558164586 | 416.6951628467046 | 441.22585123350547 | 312.91200537409725 | 462.64032974691344 | 529.988048889151 | 208.93372371623136 | 425.96993167115744 |
| SE_370_2_2 | SE_370 | SE_370_2 | ACAAAGATGAAAGGAAAGAG | 286.1170862057109 | 385.76549906868394 | 372.2976272132231 | 400.80301478645134 | 347.8750001752454 | 277.2192351578908 | 294.8680078975134 | 470.7860382969801 | 352.5301552965924 | 207.7669792756644 | 343.9189946564932 | 524.6278177530331 |
| SE_370_2_7 | SE_370 | SE_370_2 | GTATTCGAACAGTCCAACCT | 363.0652419655801 | 327.800734959918 | 322.9212840814959 | 307.1088035376705 | 303.33003064061035 | 346.5240439473635 | 442.3020118462701 | 357.34361943023794 | 336.5464202957393 | 269.0035626411234 | 327.4860051507222 | 386.50677723840715 |
| SE_370_3_1 | SE_370 | SE_370_3 | AGCAGCCTGGATTTTTAGGA | 282.86575568064603 | 360.7806869528365 | 366.37246603741585 | 283.68525072547527 | 412.57126545173924 | 267.6898239493383 | 274.42095625498513 | 388.540284618592 | 332.99447918443866 | 457.8163613512886 | 241.7997027277734 | 381.86405318749536 |
| SE_370_3_3 | SE_370 | SE_370_3 | CCACATGGCAAGAAGCAGCC | 1189.9869721737523 | 1042.3663614731538 | 1064.5539579200386 | 1154.2606302453971 | 1174.0781255914533 | 974.5988736019598 | 1129.96864340288 | 1505.0027569654465 | 975.0078350520364 | 1530.9145841364746 | 1913.2694924576244 | 1301.1234152680313 |
| SE_370_3_4 | SE_370 | SE_370_3 | AGGAGCTCATTGTCTCCACA | 2844.9142094317845 | 2949.2072221546277 | 2750.2623124372053 | 2570.0842668936407 | 2740.576220892787 | 2762.6629403703555 | 2576.328506958566 | 2968.409960346421 | 2616.892613750775 | 2223.4711817220227 | 3291.293041012993 | 2671.88769129974 |
| SE_370_3_6 | SE_370 | SE_370_3 | ACAATGAGCTCCTTAACCAA | 1170.4789890233628 | 1286.2181277238244 | 1313.4107273039435 | 1361.168680086455 | 1183.623476206018 | 1040.4384419519588 | 1108.445431147587 | 1154.2766119691019 | 1209.4359483978812 | 1689.1090911639103 | 1255.9499122267841 | 1252.3748127334575 |
| SE_370_4_14 | SE_370 | SE_370_4 | TTACTGATCCTGGTGGCCAA | 159.31519572817993 | 204.87545934994873 | 190.592684488467 | 184.78580551842887 | 199.39176839312847 | 141.20854790855063 | 151.73864639981528 | 196.6335260356865 | 180.26101139850945 | 402.4118335444473 | 132.63770101086598 | 228.65415950740604 |
| SE_370_4_2 | SE_370 | SE_370_4 | GCAGCTACTCCCCTAGAGAG | 743.470913398173 | 846.4854344849101 | 820.6348228493061 | 961.66697378957 | 867.5663114126547 | 1069.0266755776163 | 842.633759794719 | 791.2608715955269 | 990.1035847750643 | 1257.5369798263898 | 679.6214945601009 | 785.7810456168218 |
| SE_370_4_20 | SE_370 | SE_370_4 | GGTGGTCCTGAAGGGCAAGG | 13.005322100259587 | 14.990887269508443 | 31.60085960430541 | 10.410467916531204 | 26.51486281823517 | 1.7326202197368175 | 11.837766740411123 | 43.48626056558451 | 2.6639558334755096 | 7.290069448268927 | 52.820323411406804 | 8.124767089095647 |
| SE_370_4_3 | SE_370 | SE_370_4 | CCAAGTGACAAAGTTCAGCA | 1515.1200246802418 | 1447.1203177498817 | 1365.7496510235744 | 1431.4393385230405 | 1468.9234001302284 | 1656.3849300683976 | 1364.571656985573 | 1414.2488218720528 | 1568.1820006392497 | 1129.2317575368568 | 1034.1045538988756 | 2035.8344963248232 |
| SE_370_4_4 | SE_370 | SE_370_4 | TTTTTGATACGATAAAATGG | 0.0 | 5.9963549078033775 | 6.912688038441808 | 5.205233958265602 | 5.3029725636470335 | 0.0 | 2.152321225529295 | 9.45353490556185 | 0.0 | 45.92743752409424 | 0.0 | 1.1606810127279494 |
| SE_371_1_20 | SE_371 | SE_371_1 | GCCCACCTAGGACAGTGCCA | 765.1464502319391 | 656.6008624044698 | 698.1814918826226 | 770.3746258233091 | 790.142911983408 | 1014.4491386559066 | 780.2164442543694 | 750.6106715016109 | 735.2518100392406 | 852.9381254474644 | 629.14874107809 | 704.5333747258653 |
| SE_371_1_25 | SE_371 | SE_371_1 | GCTGTGGAGGTGCAAAACAA | 2109.02973392543 | 2102.7217876697177 | 2097.5070562357714 | 2326.739579344724 | 2223.0060986808367 | 2557.3474443315426 | 2315.8976386695213 | 1833.9857716789988 | 2259.922532065057 | 2720.6539180939635 | 2339.3534346429724 | 1944.1406963193153 |
| SE_371_1_28 | SE_371 | SE_371_1 | TTTGCACCTCCACAGCCTGG | 501.7886777016824 | 487.70353250134133 | 412.7862285812394 | 439.84226947344337 | 395.6017532480687 | 450.48125713157253 | 479.96763329303275 | 734.5396621621558 | 618.9257386441434 | 444.69423634440454 | 633.8438809368816 | 818.2801139732044 |
| SE_371_1_29 | SE_371 | SE_371_1 | TCATCAGCGTGGCAGACTAG | 3655.5792870146324 | 3010.1701637172955 | 3096.88424122193 | 3547.3669425580074 | 3097.9965716825973 | 3416.727073321004 | 3669.7076895274477 | 3220.819342324922 | 3452.48676018426 | 3830.931495065321 | 3098.7923068025325 | 3610.8786305966505 |
| SE_371_1_33 | SE_371 | SE_371_1 | GGTTTGCTATAGAGTGAGAT | 35.76463577571386 | 67.95868895510495 | 65.1767729338799 | 71.57196692615203 | 16.969512203670508 | 3.465240439473635 | 48.42722757440914 | 42.54090707502832 | 55.94307250298569 | 38.63736807582531 | 62.210603128990236 | 51.069964560029774 |
| SE_371_2_15 | SE_371 | SE_371_2 | GGTTTTCCTTGTGCTCCCCA | 13.005322100259587 | 11.992709815606755 | 20.738064115325425 | 2.602616979132801 | 11.666539640023474 | 12.994651648026132 | 2.152321225529295 | 19.852423301679885 | 18.647690834328564 | 0.7290069448268927 | 7.042709788187574 | 1.1606810127279494 |
| SE_371_2_20 | SE_371 | SE_371_2 | GCTTCTGACCCAGAGGCCCG | 32.51330525064897 | 47.97083926242702 | 44.43870881855448 | 66.36673296788642 | 41.363185996446866 | 4.331550549342044 | 25.827854706351538 | 45.37696754669688 | 56.831057780810866 | 122.47316673091797 | 56.341678305500594 | 212.40462532921475 |
| SE_371_2_4 | SE_371 | SE_371_2 | CCCTCTCCCTTCTTTAACCA | 67.19416418467453 | 93.94289355558624 | 79.98967587339807 | 85.88636031138243 | 69.99923784014085 | 42.44919538355203 | 34.43713960846872 | 50.10373499947781 | 49.72717555820951 | 51.759493082709376 | 53.994108376104734 | 219.36871140558245 |
| SE_371_2_5 | SE_371 | SE_371_2 | CAAGGATTCTGAAACCCAGA | 518.0453303270069 | 642.6093676195952 | 719.9070828605826 | 572.5757354092162 | 725.4466467069142 | 522.3849962506505 | 540.232627607853 | 857.4356159344597 | 753.011515595744 | 840.5450073854072 | 922.5949822525722 | 662.7488582676591 |
| SE_371_2_8 | SE_371 | SE_371_2 | CTAACGCAACTGGAAGACAC | 58.52394945116814 | 72.95565137827442 | 70.11440724705263 | 53.35364807222242 | 43.48437502190568 | 32.919784174999535 | 23.675533480822246 | 38.75949311280358 | 109.22218917249589 | 79.4617569861313 | 17.606774470468935 | 52.23064557275772 |
| SE_372_1_27 | SE_372 | SE_372_1 | GCGGCCCTGTGGAGAAGCCA | 641.595890279473 | 712.5668415439679 | 575.7281609159392 | 694.8987334284578 | 796.5064790597845 | 878.4384514065664 | 723.1799317778431 | 945.353490556185 | 572.7505041972345 | 932.3998824335957 | 892.0765731704261 | 406.2383544547823 |
| SE_372_1_30 | SE_372 | SE_372_1 | TGCCATGGTGAGCGGCCCTG | 2102.5270728753 | 2040.759453622416 | 2101.4571636863097 | 2134.1459228888966 | 2130.7343760733784 | 2542.6201724637795 | 2169.539795333529 | 2125.1546467703038 | 1996.1909045509817 | 2838.75304315592 | 2110.4653665268766 | 2384.038800143208 |
| SE_372_1_31 | SE_372 | SE_372_1 | GGGCCGCTCACCATGGCAGC | 855.0999280920678 | 1005.3888395416996 | 994.4395506729859 | 970.7761332165347 | 917.4142535109369 | 959.0052916243285 | 900.7464328840099 | 866.8891508400217 | 846.2499697673868 | 1052.686028330033 | 773.5242917359352 | 908.8132329659844 |
| SE_372_1_32 | SE_372 | SE_372_1 | GAGCCGGCTGCCATGGTGAG | 1760.0535909017974 | 2159.6871592938496 | 2137.995657603788 | 2023.5347012757527 | 2265.429879190013 | 1672.8448221558972 | 1973.6785638103634 | 2410.651400918272 | 1649.8766461991654 | 2053.6125635773565 | 2022.4314941745317 | 2025.3883672102718 |
| SE_372_1_34 | SE_372 | SE_372_1 | CTGGCTGGTGGTGATGGAGC | 908.2049933347945 | 905.4495910783099 | 934.2004120522787 | 864.0688370720899 | 908.9294974091016 | 1017.0480689855118 | 957.7829453605362 | 840.4192531044484 | 670.4288847580032 | 906.8846393646545 | 677.2739246307051 | 913.4559570168961 |
| SE_372_2_3 | SE_372 | SE_372_2 | GGGTCAGGATGGAGGCCGAC | 494.2022398098643 | 557.661006425714 | 543.1397744489992 | 514.0168533787282 | 589.6905490775501 | 443.55077625262527 | 362.6661265016862 | 558.7039129187053 | 432.4488303008577 | 438.8621807857894 | 579.849772560777 | 389.988820276591 |
| SE_372_2_32 | SE_372 | SE_372_2 | CCAAGGGAGGGATCATGGCA | 1863.0123908621858 | 1698.9672238776236 | 1626.456742759094 | 1769.7795458103046 | 1782.8593758981328 | 1749.9464219341855 | 1842.3869690530764 | 1695.9641620577959 | 1381.7050922959643 | 1778.7769455776182 | 1651.515445329986 | 1374.246319069892 |
| SE_372_2_34 | SE_372 | SE_372_2 | GCACGGTCGTATCCAAGGGA | 7858.465879081856 | 8309.948509730846 | 7994.029953026634 | 6730.367508037423 | 7339.314028087495 | 5891.775057215048 | 6421.450376366652 | 5372.4438868308 | 6750.464082026941 | 4472.457606512987 | 3779.5875863273313 | 4791.291220540976 |
| SE_372_2_36 | SE_372 | SE_372_2 | CCAGGCACGGTCGTATCCAA | 2673.6774684450334 | 2473.496399468893 | 2486.5926401137817 | 2279.892473720334 | 2488.1547268631884 | 2358.0961190618086 | 2418.132896882163 | 2767.0496668579535 | 2391.344353183182 | 2219.097140053061 | 2369.8718437251187 | 2812.3300938398215 |
| SE_372_2_41 | SE_372 | SE_372_2 | TTATTAGGATCACAGCCCCG | 144.14231994454374 | 213.8699917116538 | 192.56773821373608 | 166.56748666449926 | 135.75609762936406 | 434.8876751539412 | 119.45382801687587 | 263.7536238651756 | 244.1959514019217 | 75.08771531716994 | 220.67157336321065 | 542.0380329439524 |
| SE_372_3_24 | SE_372 | SE_372_3 | TCTGTCTTGGTGAGTCCTGA | 333.80326723999605 | 327.800734959918 | 355.50967054843585 | 350.0519836933617 | 318.178353818822 | 235.63634988420716 | 390.64630243356703 | 491.5838150892162 | 396.0414339100257 | 416.2629654961557 | 666.7098599484237 | 213.5653063419427 |
| SE_372_3_26 | SE_372 | SE_372_3 | AGGACTCACCAAGACAGAAG | 2681.2639063368515 | 3065.1367503721594 | 3160.085960430541 | 2976.092515638358 | 3090.5724100934913 | 3268.588044533506 | 2681.7922470095014 | 2879.5467322341397 | 3321.064939066135 | 3281.989265610671 | 2913.33428238026 | 3686.3228964239674 |
| SE_372_3_27 | SE_372 | SE_372_3 | GGACTCACCAAGACAGAAGA | 211.3364841292183 | 266.83779339725027 | 317.9836497683232 | 242.04337905935049 | 275.75457330964576 | 262.49196329012784 | 170.0333768168143 | 273.20715877073746 | 226.43624584541828 | 244.94633346183593 | 287.5773163509926 | 224.01143545649424 |
| SE_372_3_28 | SE_372 | SE_372_3 | CTCACCAAGACAGAAGAGGG | 227.59313675454277 | 174.89368481093183 | 205.40558742798515 | 212.11328379932328 | 243.93673792776355 | 135.14437713947177 | 194.7850709104012 | 202.30564697902358 | 158.94936473070538 | 115.18309728264904 | 151.41826044603283 | 257.67118482560477 |
| SE_372_3_31 | SE_372 | SE_372_3 | AAGACAGAAGAGGGCGGTCG | 794.4084249575232 | 995.3949146953606 | 1029.9905177278295 | 878.3832304573203 | 1033.019055398442 | 842.0534267920933 | 670.4480617523753 | 986.9490441406572 | 1060.2544217232528 | 1176.6172089506047 | 883.8600784175405 | 1043.4522304424265 |
| SE_372_4_0 | SE_372 | SE_372_4 | CTGTTCACTTCGGGCTGAGC | 365.2327956489567 | 343.7910147140603 | 304.15827369143955 | 335.7375903081313 | 281.0575458732928 | 572.6309826230182 | 256.1262258379861 | 418.79159631639 | 270.8355097366768 | 167.67159731018532 | 476.5566956673592 | 711.497460802233 |
| SE_372_4_1 | SE_372 | SE_372_4 | CTGCTCAGCCCGAAGTGAAC | 200.4987157123353 | 173.89429232629794 | 191.58021135110155 | 218.6198262471553 | 193.02820131675202 | 88.3636312065777 | 184.02346478275473 | 117.22383282896693 | 209.56452556674006 | 112.99607644816837 | 123.24742129328254 | 47.58792152184593 |
| SE_372_4_17 | SE_372 | SE_372_4 | CTGCCTCCGACTGCAAATCC | 585.2394945116814 | 625.619695380819 | 608.3165473828791 | 585.5888203048802 | 593.9329271284678 | 390.70585955065235 | 583.2790521184389 | 558.7039129187053 | 652.6691792014998 | 521.2399655512282 | 502.3799648907136 | 516.5030506639375 |
| SE_372_4_18 | SE_372 | SE_372_4 | CAGGATTTGCAGTCGGAGGC | 259.02266516350346 | 509.6901671632871 | 427.59913152075757 | 294.0957186420065 | 476.2069362155036 | 413.22992240723096 | 252.89774399969215 | 322.3655402796591 | 475.07212363646585 | 361.58744463413876 | 227.71428315139823 | 875.1534835968739 |
| SE_372_4_20 | SE_372 | SE_372_4 | CTATGGAAAGTGATTGACCA | 563.5639576779155 | 576.649463633758 | 568.8154728774973 | 512.7155448891617 | 543.0243905174563 | 747.6256248164367 | 540.232627607853 | 474.56745225920486 | 759.2274125405202 | 607.2627850408015 | 500.0323949613178 | 575.6977823130629 |
| SE_373_1_1 | SE_373 | SE_373_1 | TTAAGGACACCGGAACTCCA | 563.5639576779155 | 574.6506786644903 | 519.4391297457702 | 554.3574165552866 | 522.8730947755976 | 589.0908747105179 | 457.3682604249752 | 764.7909738599536 | 956.3601442177079 | 669.2283753510875 | 650.2768704426527 | 721.9435899167845 |
| SE_373_1_15 | SE_373 | SE_373_1 | AGTTCTGGAGAGGATTCAAT | 24.926867358830876 | 51.968409200962604 | 35.55096705484358 | 32.53271223916001 | 33.939024407341016 | 30.320853845394307 | 20.4470516425283 | 21.743130282792254 | 27.527543612580263 | 23.328222234460565 | 0.0 | 13.928172152735392 |
| SE_373_1_20 | SE_373 | SE_373_1 | ATGGCCAGCCCAGACCCAAA | 3061.6695777694445 | 3567.8311701430093 | 3522.5083190174187 | 3248.0659899577354 | 3534.9615109271126 | 3555.3366908999496 | 3282.2898689321746 | 4089.5992001460563 | 3521.7496118546233 | 3202.527508624539 | 3400.4550427299005 | 3292.8520331091927 |
| SE_373_1_24 | SE_373 | SE_373_1 | GCTTCCTGCTCAGAGCCAGA | 365.2327956489567 | 344.7904071986942 | 322.9212840814959 | 426.8291845777936 | 369.08689042983355 | 369.0481068039421 | 361.58996588892154 | 359.2343264113503 | 403.1453161326271 | 365.2324793582732 | 145.5493356225432 | 345.8829417929893 |
| SE_373_1_27 | SE_373 | SE_373_1 | TCTTCAATTTGCTTTCTGCA | 13.005322100259587 | 9.993924846338961 | 7.900214901076352 | 16.917010364363207 | 2.1211890254588135 | 79.7005301078936 | 25.827854706351538 | 3.78141396222474 | 26.639558334755094 | 1.4580138896537853 | 2.347569929395858 | 16.249534178191293 |
| SE_373_2_17 | SE_373 | SE_373_2 | TACAGGATGGAAGGAAACAG | 1514.0362478385537 | 1636.0054973456881 | 1592.8808294295195 | 1297.4045640977013 | 1528.316692843075 | 1518.6416225993205 | 1207.4522075219345 | 1601.4288130021773 | 1660.5324695330676 | 777.8504101302944 | 1557.6126481541519 | 1234.9645975425383 |
| SE_373_2_28 | SE_373 | SE_373_2 | TTAAATACCCGTCCTCAGCA | 29.26197472558407 | 57.964764108765976 | 50.36386999436174 | 53.35364807222242 | 23.33307928004695 | 101.35828285460381 | 9.685445514881827 | 26.46989773557318 | 11.543808611727208 | 21.141201399979888 | 25.823269223354437 | 33.659749369110536 |
| SE_373_2_29 | SE_373 | SE_373_2 | GCTGACATTTGGGCACTGAT | 4680.832179251763 | 4008.5632558665575 | 4015.284223472056 | 4934.56179243579 | 4174.500002102945 | 4743.914161639406 | 5216.150490070247 | 4736.220987686487 | 4280.977024395143 | 4205.641064706344 | 4522.59346898112 | 4012.474261000521 |
| SE_373_2_32 | SE_373 | SE_373_2 | TGAAAAGCACTCAGAGAAGA | 370.6516798573982 | 404.75395627672793 | 426.61160465812304 | 430.73311004647854 | 440.1467227827038 | 828.1924650341988 | 432.6165663313883 | 481.1849266930982 | 472.4081678029903 | 406.7858752134061 | 322.79086529193046 | 374.8999671111277 |
| SE_374_1_2 | SE_374 | SE_374_1 | CCACTTCCTGAGGCTGGCCT | 266.60910305532155 | 224.86330904262664 | 196.51784566427426 | 287.5891761941745 | 266.2092226950811 | 506.7914142730191 | 304.55345341239524 | 261.8629168840632 | 280.60334779275365 | 157.46550008260883 | 279.3608215981071 | 406.2383544547823 |
| SE_374_1_20 | SE_374 | SE_374_1 | CTATAGGTTCAACTTAAAAT | 19.50798315038938 | 26.983597085115196 | 30.613332741670867 | 16.917010364363207 | 15.9089176909411 | 78.83421999802519 | 20.4470516425283 | 0.945353490556185 | 21.311646667804077 | 1.4580138896537853 | 1.173784964697929 | 60.35541266185337 |
| SE_374_1_24 | SE_374 | SE_374_1 | AAGTTGAACCTATAGCAGGG | 27.094421042207472 | 72.95565137827442 | 66.16429979651446 | 118.41907255054244 | 75.30221040378788 | 320.5347406513112 | 55.960351863761666 | 83.19110716894428 | 87.9105425046918 | 106.43501394472632 | 92.72901221113639 | 113.74673924733905 |
| SE_374_1_25 | SE_374 | SE_374_1 | GTTGGACAAATGAAGCAAAG | 217.8391451793481 | 310.8110627211417 | 317.9836497683232 | 261.5630064028465 | 330.9054879715749 | 474.73794020788796 | 267.9639925783972 | 429.190484712508 | 369.40187557527065 | 132.67926395849446 | 289.9248862803885 | 552.4841620585039 |
| SE_374_1_38 | SE_374 | SE_374_1 | ATTGCTTCCTCAAGTGCCTA | 39.01596630077876 | 78.9520062860778 | 79.98967587339807 | 79.37981786355043 | 92.27172260745839 | 77.10159977828837 | 44.12258512335055 | 53.88514896170255 | 23.087617223454416 | 152.36245146882055 | 23.47569929395858 | 85.89039494186825 |
| SE_374_2_10 | SE_374 | SE_374_2 | TGAGCCCCTGGCCCAACCAG | 198.3311620289587 | 223.86391655799275 | 229.10623213121423 | 221.22244322628808 | 184.54344521491677 | 188.8556039513131 | 258.2785470635154 | 170.1636283001133 | 269.9475244588516 | 238.3852709583939 | 149.070690516637 | 507.2176025621139 |
| SE_374_2_22 | SE_374 | SE_374_2 | GGAGAGAGTCTAGTGGCGGT | 1662.5136751498505 | 2241.637343033829 | 2190.3345813234187 | 1764.574311852039 | 2038.4626534659199 | 2812.042616632855 | 1632.5356495639703 | 2248.0506005426078 | 2028.1583745526877 | 1829.8074315155006 | 1948.4830413985621 | 2315.558620392259 |

| | | | | | | | | | | | | | | |
|---|---|---|---|---|---|---|---|---|---|---|---|---|---|---|
| SE_374_2_3 | SE_374 | SE_374_2 | CTTGTTATCATCCTTCCACA | 419.42163773337165 | 363.7788644067382 | 393.0356913285485 | 381.2833874429553 | 441.2073172954332 | 427.0908841651255 | 408.94103285056605 | 400.82987999582247 | 358.7460522413686 | 343.36227101346645 | 446.03828658521303 | 471.23649116754746 |
| SE_374_2_39 | SE_374 | SE_374_2 | ACCTCAATAGGGAAAGCACC | 1251.7622521499852 | 1104.3286955204553 | 1267.9844916227546 | 1251.8587669628773 | 1073.3216468821597 | 971.9999432724536 | 1531.3765519640933 | 1117.4078258374107 | 1069.1342745015045 | 1234.937766354784 | 1235.9955678269193 | 906.4918709405285 |
| SE_374_2_4 | SE_374 | SE_374_2 | CACTAACTGAGGGAGCAAGT | 1316.7888626512831 | 1411.1421883030614 | 1293.6601900512526 | 1277.8849367542052 | 1313.0160067590054 | 1468.3956362269528 | 1223.594616713404 | 1309.3145844203161 | 1287.5786528464962 | 911.9876879784428 | 1362.7643440142956 | 1548.3484709790846 |
| SE_374_3_3 | SE_374 | SE_374_3 | TGAACACGAGTCACATCCAA | 1828.3315319281603 | 1723.9520359934709 | 1596.8309368800576 | 1678.6879515406565 | 1649.2244672942275 | 1721.358188308528 | 1591.6415462789137 | 1388.7242776270357 | 1953.5676111253735 | 1214.525570081603 | 1427.3225170726816 | 2098.5112710121325 |
| SE_374_3_30 | SE_374 | SE_374_3 | GTACTCACCATTCCACCTCA | 526.7155450605133 | 557.661006425714 | 596.4662250312646 | 593.3966712422786 | 611.9630338448677 | 354.32083493617915 | 490.7292394206792 | 466.0592708441992 | 416.46509530000463 | 796.0755837509668 | 746.5272375478829 | 543.1987139566803 |
| SE_374_3_32 | SE_374 | SE_374_3 | GCACACTGAGAAGTGATGGG | 2721.3636494793186 | 2639.39555191812 | 2640.6468306847705 | 2881.0969959000104 | 2786.1817849401514 | 2718.4811247670664 | 3057.3723008643633 | 3104.5408629865115 | 2782.945860704082 | 3351.9739323140525 | 2854.6450341453633 | 3025.895400181764 |
| SE_374_3_7 | SE_374 | SE_374_3 | GACCAGGGTGAAGATTAAAA | 166.90163619999802 | 233.8578414043317 | 270.5823603618651 | 223.82506020542087 | 281.0575458732928 | 211.37966680789174 | 146.3578433599204 | 211.75918188458544 | 302.8029797383829 | 190.99981954464587 | 185.45802442227279 | 493.2894304093785 |
| SE_374_3_8 | SE_374 | SE_374_3 | GCTCTGAGACAGCTGAGTGG | 390.15966300778763 | 367.7764343452738 | 368.3475197626849 | 357.85983463076013 | 512.2671496483034 | 307.5400890032851 | 392.79862365909634 | 258.08150292183853 | 309.90686196098426 | 274.8356181997385 | 265.27540202173196 | 155.53125570554522 |
| SE_375_1_10 | SE_375 | SE_375_1 | TATCCTGATGGGAGAGACGG | 370.6516798573982 | 381.76792913014833 | 402.91095995489394 | 403.40563176558413 | 384.99580812077465 | 493.796762624993 | 352.9806809868044 | 538.8514896170254 | 457.3124180799624 | 295.9768195997184 | 787.6097113123104 | 436.41606078570896 |
| SE_375_1_12 | SE_375 | SE_375_1 | AGTGTGCTCGGTATCCTGAT | 1049.0959827542733 | 1078.344490919974 | 977.6515940081986 | 1047.5533341009523 | 1122.1089944677124 | 1150.4598259052468 | 1136.4256070794677 | 1119.298532818523 | 1003.4233639424418 | 1177.3462158954317 | 990.6745102050521 | 833.3689671386677 |
| SE_375_1_20 | SE_375 | SE_375_1 | ACCTCCCACACACTAAGCCC | 2721.3636494793186 | 2704.356063419323 | 2546.831778734489 | 2803.0184865260267 | 2492.397104914106 | 2126.7913197269436 | 2528.9774399969215 | 2555.290484973368 | 2286.5620903998124 | 2848.23013343867 | 2301.7923157726386 | 2221.543458361295 |
| SE_375_1_22 | SE_375 | SE_375_1 | GGATCACTGTAGCACAGCCA | 743.470913398173 | 916.4429084092828 | 951.9758955797005 | 770.3746258233091 | 969.3833846346778 | 653.1978228407802 | 757.6170713863119 | 790.3155181049707 | 954.5841736620575 | 626.9459725511277 | 819.3019053591545 | 790.4237696677336 |
| SE_375_1_5 | SE_375 | SE_375_1 | GTTTAGATATCGCTTCCTGC | 43.351073667531956 | 38.97630690072195 | 52.338923719630834 | 39.03925468699201 | 48.78734758555271 | 25.989303296052263 | 48.42722757440914 | 4.726767452780925 | 2.6639558334755096 | 27.702263903421922 | 21.128129364562724 | 54.55200759821362 |
| SE_375_2_15 | SE_375 | SE_375_2 | ATAGAACTGAGAAAGGTAGA | 89.95347786012881 | 143.91251778728105 | 164.91698605996885 | 145.74655083143685 | 181.36166167672854 | 288.4812665861801 | 99.00677637434757 | 274.1525122612937 | 141.189659174202 | 57.59154864132452 | 281.70839152750295 | 227.4934784946781 |
| SE_375_2_16 | SE_375 | SE_375_2 | CATAGAACTGAGAAAGGTAG | 443.2647282505143 | 535.6743717637684 | 524.3767640589429 | 473.67629020216975 | 436.96493924451556 | 660.9946138295959 | 463.82522410156304 | 400.82987999582247 | 537.2310930842277 | 314.93100016521765 | 273.4918967746175 | 413.20244053115 |
| SE_375_2_22 | SE_375 | SE_375_2 | ACAGCAGCCCTCGGAAGAAG | 3711.9356827824236 | 3937.606389457551 | 3890.8558387801036 | 4149.872773227251 | 4054.652822164522 | 4199.005102532177 | 3995.784355195136 | 3809.7745669414257 | 3709.1145054757344 | 4301.869981423493 | 3236.1251476721905 | 4158.720068604242 |
| SE_375_2_23 | SE_375 | SE_375_2 | ACAAAAATAATTGCCAGGAG | 49.85373471766175 | 70.95686640900662 | 73.07698783495626 | 105.40598765487843 | 44.54496953463509 | 113.48662439276154 | 79.63588534458391 | 44.431614056140695 | 149.18152667462851 | 143.61436813089784 | 183.11045449287693 | 10.446129114551544 |
| SE_375_2_24 | SE_375 | SE_375_2 | GACTGTCACCTCTTCTTCCG | 1127.127915355831 | 1252.2387832462719 | 1385.5001882762654 | 1142.5488538392997 | 1468.9234001302284 | 1568.0212988618198 | 1164.4057830113486 | 923.6103602733928 | 1330.2019461821044 | 1084.7623339024162 | 1216.0412234270545 | 1128.181944371567 |
| SE_375_3_10 | SE_375 | SE_375_3 | TGAGAGGCAGAGAGAGCACA | 9875.37458146378 | 9086.476470291384 | 9110.922834666304 | 9606.259269979168 | 9111.567458858333 | 9714.801572064336 | 9587.514899120244 | 9162.366030470545 | 9417.083871335926 | 8451.377511187276 | 9052.229647750428 | 9434.015271452772 |
| SE_375_3_16 | SE_375 | SE_375_3 | TCAGGGTTAGCCACACACTG | 6020.380355578501 | 5249.808721781857 | 5322.769789600193 | 5915.748393568856 | 5103.580795253905 | 5495.005026895316 | 5761.763920741922 | 5293.03419362408 | 5446.901694179592 | 4988.594523450426 | 5541.438818338923 | 5799.923020601564 |
| SE_375_3_19 | SE_375 | SE_375_3 | ATAGCTGTCCTGTGTCCCAG | 605.831254503759 | 603.6330607188733 | 642.8799875750882 | 568.6718099405169 | 631.053735073997 | 421.89302350591504 | 562.8320004759106 | 671.2009782948913 | 609.1579005880665 | 521.9689724960551 | 733.6156029362056 | 254.18914178742094 |
| SE_375_3_22 | SE_375 | SE_375_3 | TTTGGAGCCAGAGAGTACGT | 463.8564882425919 | 427.7399834233076 | 507.58880739415565 | 469.77236473347057 | 403.0259148371746 | 300.60960812433785 | 408.94103285056605 | 394.21240556192913 | 261.0676716805999 | 394.3927571513489 | 501.2061799260157 | 275.081400016524 |
| SE_375_3_26 | SE_375 | SE_375_3 | GACTTTCCAACAAATAGCCC | 66.11038734298623 | 97.94046349412183 | 94.80257881291622 | 50.751031093089615 | 82.72637199289373 | 155.93581977631356 | 69.95043982970209 | 93.58999556506231 | 88.79852778251698 | 282.12568764800744 | 130.29013108147012 | 112.5860582346111 |
| SE_375_4_16 | SE_375 | SE_375_4 | CCTCCTTTCCCCAGTGCAGA | 139.80721257779055 | 255.84447606627742 | 258.73203801025056 | 196.49758192452646 | 171.8163110621639 | 161.99999054539242 | 163.5764131402264 | 153.14726547010198 | 245.08393667974687 | 432.3011182823473 | 133.8114859755639 | 148.56716962917753 |
| SE_375_4_17 | SE_375 | SE_375_4 | CGTGAGTAATGGAGTCACCA | 18376.520127666798 | 16492.97417391319 | 16513.424196974844 | 17529.92666294898 | 17707.685984530173 | 15956.56591366622 | 18053.670439739726 | 17877.579859908015 | 16492.550565046877 | 17676.231391217665 | 17274.59332545942 | 16856.57034784801 |
| SE_375_4_20 | SE_375 | SE_375_4 | AGTGGGCTCCAGACTGAACT | 598.244816611941 | 584.6446035108293 | 557.9526773885174 | 519.2220873369938 | 584.3875765139031 | 537.1122681184135 | 456.29209981221055 | 480.239573202542 | 386.27359585394885 | 779.3084240199482 | 403.7820278560876 | 817.1194329604764 |
| SE_375_4_22 | SE_375 | SE_375_4 | CAAACACACTTTCTTGCGGG | 753.2249049733678 | 1037.3693990499842 | 1027.0279371399258 | 921.3264106130115 | 1092.412348111289 | 846.3849773414353 | 862.0046508244826 | 1045.5609605551406 | 1078.9021125575812 | 722.4458823234506 | 1322.855655214566 | 1182.7339519697805 |
| SE_375_4_26 | SE_375 | SE_375_4 | GTTTCCTCACGTAAGATGAG | 16.256652625324485 | 20.98742217731182 | 26.66322529113269 | 2.602616979132801 | 14.848323178211695 | 0.0 | 6.456963676587668 | 5.67212094333711 | 31.07948472388094 | 11.66411117230282 | 18.780559435166865 | 0.0 |
| SE_375_5_1 | SE_375 | SE_375_5 | GAAACTATAAGAGTACAGAG | 13.005322100259587 | 22.986027146579612 | 21.725590977959968 | 22.12224432262881 | 39.24199697098805 | 25.12299318618385 | 20.44705164125283 | 54.83050245225873 | 15.983735000853056 | 28.431270848248815 | 19.954344399864794 | 17.410215190919242 |
| SE_375_5_10 | SE_375 | SE_375_5 | ACATGTTTAAAAGGTGGTGA | 773.8166649654454 | 916.4429084092828 | 914.4498747995877 | 874.4793049886212 | 1008.6253816056658 | 902.6951344828819 | 898.5941116584806 | 681.5998666910094 | 1221.8677422874337 | 1309.2964729090993 | 694.8806991011174 | 1029.524058289691 |
| SE_375_5_13 | SE_375 | SE_375_5 | GCATTGTGTGAAATGGGCAG | 3374.8810850173627 | 3475.887061556691 | 3469.1818684351533 | 3467.987124964457 | 3193.4500778824036 | 3235.6682603585507 | 3287.670671995998 | 2839.84188563078 | 4556.254605520946 | 3299.4854322865162 | 2709.0956985228204 | 3347.4040407074062 |
| SE_375_5_2 | SE_375 | SE_375_5 | GTCCAATGTCCACAGCACAA | 979.7342648862223 | 851.4823969080765 | 908.5247136237805 | 766.4707003546099 | 930.1413876636897 | 1069.8929856878447 | 710.2660044246674 | 717.5232993321445 | 787.6429414309256 | 663.3963197924724 | 716.0088284657367 | 1135.1460304479344 |
| SE_375_5_5 | SE_375 | SE_375_5 | ATAAGAGTACAGAGTGGGTG | 418.33786089168336 | 522.6822694635277 | 477.9630015151193 | 435.9383440047441 | 521.8125002628681 | 881.90369184604 | 415.3979965271539 | 405.55664744860337 | 654.4451497571501 | 391.47672937204135 | 482.4256204908488 | 597.750721554894 |
| SE_376_1_0 | SE_376 | SE_376_1 | CAGTGTTGGAGGCTGAAAAG | 1377.4803657858279 | 1549.0583511825391 | 1497.0907237539689 | 1426.234104564775 | 1578.1646349413572 | 1094.1496687638003 | 1408.6942421089236 | 1779.15526922674 | 1331.9779167377546 | 1936.242445460227 | 1671.4697897298508 | 1598.2577545263864 |
| SE_376_1_1 | SE_376 | SE_376_1 | CCAGTGTTGGAGGCTGAAAA | 355.47880407376203 | 480.7077851089041 | 417.72386289441215 | 385.18731291165454 | 363.7839178661865 | 281.5507857072328 | 287.33488360816085 | 304.4038239590916 | 317.0107441835856 | 188.8127987101652 | 211.28129364562722 | 243.74301267286938 |
| SE_376_1_10 | SE_376 | SE_376_1 | ACTTTATTGTAAAACTTCAG | 2.167553683376598 | 7.9951398770711695 | 13.825376076883616 | 22.12224432262881 | 8.484756101835254 | 62.37432791052543 | 9.685445514881827 | 10.398888396118036 | 62.15896944776188 | 18.954180565499207 | 5.868924823489645 | 170.62010887100857 |
| SE_376_1_11 | SE_376 | SE_376_1 | AAATAGATCTTGAATCATAG | 18.42420630870108 | 12.992102300240651 | 18.763010390056337 | 7.807850937398403 | 15.9089176909411 | 20.791442636841808 | 17.21856980423436 | 29.305958207241734 | 17.759705556503395 | 0.0 | 59.86303319959438 | 0.0 |
| SE_376_1_5 | SE_376 | SE_376_1 | TGGCCCAACTCCAGTGTTGG | 10.837768416882989 | 18.98845720804403 | 28.63827901640178 | 5.205233958265602 | 20.15129574185873 | 0.0 | 27.980175931880833 | 21.743130282792254 | 0.8879852778251698 | 0.7290069448268927 | 9.390279717583432 | 9.285448101823595 |
| SE_377_1_15 | SE_377 | SE_377_1 | TCCAGCACAGACCTGAAGGA | 701.2036165723293 | 769.5322131681 | 832.4851452009207 | 767.7720088441763 | 681.9622716850085 | 325.73260131052166 | 785.5972473181927 | 853.6542019722351 | 681.0847080919052 | 978.32731995769 | 838.0824647943214 | 493.2894304093785 |
| SE_377_1_21 | SE_377 | SE_377_1 | CTGCTGCCAGGACCGCCGGT | 345.72481249856736 | 415.74727360770083 | 408.83612113070126 | 408.6108657238497 | 298.02705807696327 | 343.92511361775826 | 334.68595056980536 | 277.93392622351837 | 376.505757797872 | 304.7249029376411 | 606.8468267488294 | 474.7185342057313 |
| SE_377_1_26 | SE_377 | SE_377_1 | GCTCACGAGCAGCTGCAGGG | 518.0453303270069 | 123.92466809460312 | 222.1935440927724 | 390.3925468699201 | 94.3929116329172 | 830.791395363804 | 558.527358024852 | 233.5023121673777 | 812.5065292100304 | 43.74041668961356 | 172.54638981059557 | 409.72039749296613 |
| SE_377_1_27 | SE_377 | SE_377_1 | GATGCTCACGAGCAGCTGCA | 221.09047570441297 | 61.96233404730156 | 100.72773998872348 | 189.99103947669445 | 48.78734758555271 | 618.5454184460439 | 294.8680078975134 | 125.7320142439726 | 417.3530805778298 | 20.412194455152996 | 149.070690516637 | 129.99627342553035 |
| SE_377_1_9 | SE_377 | SE_377_1 | TGTGCTGGAGAGGAGGTGGG | 39048.47960602941 | 38181.789875438 | 39094.21343797633 | 38629.342512778596 | 39159.27059899516 | 40166.468244048774 | 39078.62033132264 | 41311.00218381473 | 39313.77220515374 | 45692.69728785998 | 43629.587137822025 | 41642.91337465337 |
| SE_377_2_18 | SE_377 | SE_377_2 | CTCGGGCAGGTAACACAATC | 2288.9366896456872 | 1909.8390381353756 | 2012.5797460492008 | 2298.110792574263 | 2274.9752298045773 | 2302.6522720302305 | 2576.328506958566 | 2543.0008895961378 | 2148.924372336911 | 2857.7072237214193 | 3395.759902871109 | 1969.6756785993302 |
| SE_377_2_19 | SE_377 | SE_377_2 | CTTTCGGGGTCTGCTCGGGC | 23.843090517142578 | 71.95625889364052 | 56.28903117016901 | 20.820935833062407 | 46.6661585600939 | 90.96256153618292 | 18.294730416999005 | 28.36060471668555 | 48.83919028038434 | 11.664111117230282 | 140.85419576375148 | 60.35541266185337 |
| SE_377_2_24 | SE_377 | SE_377_2 | GACACGGCTGGACCTGGCGG | 4710.094153977347 | 5294.781383590382 | 5131.189578249091 | 5332.762190243109 | 5324.184453901622 | 5562.577215465052 | 5218.302811295775 | 5637.142864186531 | 5070.395936381719 | 5945.05163506331 | 6022.690653865074 | 4723.9717218027545 |
| SE_377_2_25 | SE_377 | SE_377_2 | TGAGACACGGCTGGACCTGG | 4569.203164557868 | 4291.39132901795 | 4121.937124636587 | 4680.806636970342 | 4472.527060179908 | 4597.507753071645 | 5093.468180215076 | 4941.362695137179 | 4268.5452305055915 | 4241.362405002861 | 4959.24147584875 | 5155.745058537552 |
| SE_377_2_28 | SE_377 | SE_377_2 | ACAGAATGGCTACTGAGACA | 2103.610849716988 | 1976.7983346058465 | 1898.0266299835937 | 2244.757144502041 | 2156.188644378884 | 1989.0480122578665 | 2272.8512141589354 | 2492.8971545966597 | 2350.4970304032245 | 2012.0591677222237 | 2717.312193275706 | 1906.998903912021 |
| SE_378_1_0 | SE_378 | SE_378_1 | ACTGGTGTGAGGGCAGGACA | 2424.408794856725 | 2138.699917116538 | 2028.3801758513534 | 2344.9578981986538 | 2140.279726687943 | 1435.4758520519533 | 2778.64670215832 | 2387.9629171449233 | 1967.7753756605762 | 3189.4053836176554 | 2428.5610919600153 | 2341.093602672274 |
| SE_378_1_12 | SE_378 | SE_378_1 | GCTACCACGTGGCTTTTGGG | 586.3232713533697 | 646.6093755581309 | 786.071382657097 | 709.2131268168383 | 714.8407015796201 | 806.5347122874886 | 571.4412857802576 | 653.2392619743239 | 727.259942528814 | 890.117479633636 | 352.1354894093787 | 974.9720506914775 |
| SE_378_1_13 | SE_378 | SE_378_1 | TGTGAGCTGAGGAAAACATG | 2971.7160999093157 | 2702.357278450055 | 2863.8279016401775 | 2964.38073923226 | 2585.7294220342937 | 2516.6308691677273 | 2821.6931266689057 | 2794.464918084083 | 2745.650479035425 | 2587.245647190642 | 2557.6774380767874 | 2996.8783748635656 |
| SE_378_1_18 | SE_378 | SE_378_1 | GCACATCTCTTTAGCAGCAG | 2628.158841094125 | 2857.263113568309 | 3010.969404172725 | 2727.5425941311755 | 2932.54382769681 | 3233.0693300289013 | 2798.0175931880835 | 2620.519875821745 | 2695.03531819939 | 3117.233696077993 | 2835.8644747101966 | 3216.247086269148 |
| SE_378_1_5 | SE_378 | SE_378_1 | GGTACAGAGGAGGAACTCTG | 450.8511661423323 | 568.654323756687 | 503.63869994361744 | 520.5233958265602 | 499.54001549555056 | 687.8502272355165 | 437.99736939521154 | 562.4853268809301 | 539.0070636398781 | 491.35068081332565 | 584.5449124195686 | 457.30831901481207 |
| SE_378_2_10 | SE_378 | SE_378_2 | CATTCTCACTTTGGTAGCCG | 225.42558307116616 | 267.8371858818842 | 289.3453707519214 | 191.29234796626088 | 307.572408691528 | 386.3743090013103 | 143.1293614976981 | 260.91756339350707 | 293.035141682306 | 287.2287362617957 | 226.5404981867003 | 516.5030506639375 |
| SE_378_2_14 | SE_378 | SE_378_2 | GGGAAGTGTATGTTTGGCAG | 10622.096825387018 | 13105.03365100428 | 12742.059108573521 | 12027.994369062239 | 12905.31403089142 | 12995.517958135999 | 11727.998357909128 | 13047.768876656466 | 12958.369159302703 | 13455.281180669957 | 13780.235485553687 | 13277.030104595013 |
| SE_378_2_20 | SE_378 | SE_378_2 | GGGTACTTAGAGCACTTTAA | 23.843090517142578 | 36.977521931454156 | 44.43870881855448 | 22.12224432262881 | 32.87842989461161 | 10.395721318420904 | 0.0 | 41.59555358447214 | 21.311646667804077 | 2.9160277793075706 | 25.823269223354437 | 24.37430126728694 |
| SE_378_2_23 | SE_378 | SE_378_2 | ATTACTACATCCCTAACAAA | 21.675536833765978 | 12.992102300240651 | 19.75053725269088 | 18.218318853929606 | 11.666539640023474 | 8.663101098684088 | 32.28481838293943 | 6.617474433893295 | 1.7759705556503396 | 25.515243068941242 | 8.216494752885502 | 68.48017975094902 |
| SE_378_2_6 | SE_378 | SE_378_2 | GTGAGAATGAGGACTCAAGG | 897.3672249179115 | 892.4574887780693 | 912.4748210743187 | 754.7589239485122 | 940.7473327909838 | 1102.8127698624844 | 836.176796118131 | 1163.7301468746637 | 754.7874861513943 | 1046.8539727714178 | 894.4241430998219 | 1263.981622860737 |
| SE_378_3_0 | SE_378 | SE_378_3 | GTATTTTGCAGAAAAATAAC | 6.5026610501297935 | 12.992102300240651 | 6.912688038441808 | 15.615701874796805 | 44.54496953463509 | 0.8663101098684087 | 15.066248578705064 | 19.852423301679885 | 17.759705556503395 | 2.9160277793075706 | 48.12518355261509 | 1.1606810127279494 |
| SE_378_3_11 | SE_378 | SE_378_3 | GTAAGGTCTAATGCAGCAGA | 1344.967060535179 | 1391.1543386103835 | 1464.5023372870287 | 1456.1641998248022 | 1461.4992385411224 | 1613.9357346848456 | 1510.929500321565 | 1439.7733661170698 | 1391.472930352041 | 1489.3611882813416 | 1207.824728674169 | 1582.008220348195 |
| SE_378_3_13 | SE_378 | SE_378_3 | GCAGGACTTGAGGAAAAGCA | 5301.836309539158 | 5978.365843079967 | 5806.657952291119 | 4814.8414113956815 | 4850.0987067115775 | 2922.930310696011 | 4158.284607722598 | 3555.474477981812 | 4401.743022179367 | 3031.9398835350466 | 2806.5198505927483 | 2377.0747140668404 |
| SE_378_3_3 | SE_378 | SE_378_3 | GCTGAAGTGAGAAGAAAAT | 455.18627350908554 | 636.6130127117918 | 615.2292354613209 | 549.152182597021 | 575.9028204120679 | 479.9358008670984 | 542.3849485338923 | 536.9607826359131 | 846.2499697678368 | 397.3087849306565 | 572.8070627725893 | 437.57674179843696 |
| SE_378_3_6 | SE_378 | SE_378_3 | ATATAGCACTCAAGAGGCAG | 244.93356622155557 | 240.85358879676897 | 227.13117840594512 | 219.92113473672168 | 292.72408551331625 | 343.92511361775826 | 152.81480701257993 | 205.14170745069214 | 155.39742361940472 | 163.29755564122397 | 300.48895096266983 | 222.85075444376628 |
| SE_379_1_22 | SE_379 | SE_379_1 | CCAGGAATGTGGTATCAAGG | 440.01339772544935 | 400.7563863381924 | 403.8984868175285 | 359.1611431203265 | 392.4199697098805 | 521.518686140782 | 436.92120878244685 | 555.8678524470367 | 412.91315418870397 | 489.163659978845 | 415.5198775030669 | 284.3668481183476 |
| SE_379_1_29 | SE_379 | SE_379_1 | AGCTCGTTCTCACCAGGCCA | 626.4230144958368 | 596.637313326436 | 619.1793428718591 | 577.7809693674818 | 566.3574697975032 | 738.0962136078842 | 556.3750367993227 | 700.506936502133 | 677.5327669806045 | 454.90033357198104 | 542.2886536904432 | 422.4878886329736 |
| SE_379_1_42 | SE_379 | SE_379_1 | TGTCATCCTCTTCCTCTGCC | 1455.5122983873855 | 1079.3438834046078 | 1244.2838469195256 | 1284.3914792020373 | 1321.5007628608407 | 1684.1068535841866 | 1246.1939895814617 | 1297.0249890430857 | 1026.5109811658963 | 913.4457018680965 | 1232.4742129328256 | 863.5466734695943 |
| SE_379_1_43 | SE_379 | SE_379_1 | TGAGCCAGATAAAGGCACAG | 2120.951279184001 | 2066.7436582228975 | 2192.3096350486876 | 2255.167614288512 | 2268.611662728201 | 3346.555954421663 | 2073.7615007974755 | 2612.9570478972955 | 2378.024574015805 | 2507.783890204511 | 2550.6347282886 | 3212.7650432309642 |
| SE_379_2_17 | SE_379 | SE_379_2 | TCAGGCACCTCCTTTGGGTG | 2472.0949758910097 | 2534.4593410315606 | 2450.054146196304 | 2426.940333041337 | 2596.335367161588 | 2461.187022136149 | 2663.4975165925025 | 2662.115429406217 | 2509.44639513393 | 2137.448362232449 | 2608.1501915587983 | 2667.244967248828 |
| SE_379_2_18 | SE_379 | SE_379_2 | GCACCTCCTTTGGGTGTGGA | 273.1117641054513 | 313.8092401750434 | 361.4348317242431 | 213.41459228888968 | 284.239329411481 | 329.1978417499953 | 189.40426784657797 | 228.77554471459678 | 375.6177725200468 | 125.38919451022554 | 433.1266519735558 | 92.85448101823596 |
| SE_379_2_19 | SE_379 | SE_379_2 | ATCGAGTCCCAACCCCCATG | 3490.845207078011 | 3639.78742903665 | 3650.8868111599095 | 3549.9695595371404 | 3571.0217243599127 | 4044.8019029756006 | 3318.8793297661728 | 3579.1083152457163 | 3596.3403751919377 | 3563.3859463138515 | 2813.562560380936 | 3707.2151546530704 |
| SE_379_2_2 | SE_379 | SE_379_2 | CAGGTGGCCTGGTCCGTGGA | 1179.1492037568692 | 1057.3572487426623 | 1024.0653565520222 | 1258.3653094107092 | 1107.2606712895006 | 1175.5828190914306 | 1169.7865860751717 | 897.1404625378195 | 1206.7719925644058 | 960.8311532818445 | 1044.6686185811568 | 1295.3200102043916 |
| SE_379_2_24 | SE_379 | SE_379_2 | GGACACCCGGACCAGTGGCT | 807.4137470577826 | 1014.3833719034046 | 974.6890134202949 | 858.8636031138243 | 985.2923023256188 | 862.8448694289351 | 889.9848267563634 | 938.7360161222917 | 913.7368508820997 | 809.9267157026777 | 832.2135399708317 | 653.4634101658355 |
| SE_38_1_2 | SE_38 | SE_38_1 | ATCTGTTTCGATCTGCGCCG | 56.35639576779154 | 64.96051150120326 | 66.16429979651446 | 85.88636031138243 | 46.6661585600939 | 128.21389626052448 | 83.94052779564251 | 129.51342820619735 | 68.37486639253807 | 42.282402799959776 | 172.54638981059557 | 11.606810127279495 |
| SE_38_1_31 | SE_38 | SE_38_1 | ACCCACCCCGGCTTCGCGTG | 279.6144251555811 | 453.7241880237889 | 461.1750448503321 | 325.3271223916001 | 368.02629591710416 | 230.43848922499672 | 270.1163138039265 | 273.20715877073746 | 293.035141682306 | 279.2096598566899 | 296.96759606857603 | 543.1987139566803 |
| SE_38_1_32 | SE_38 | SE_38_1 | CCCGCACGCGAAGCCGGGGT | 693.6171786805113 | 903.4508061090421 | 837.4227795140933 | 593.3966712422786 | 683.0228661797379 | 261.62565318025946 | 373.4277326293327 | 233.5023121673777 | 630.4695472558706 | 242.03030568252836 | 138.50662583435562 | 55.71268861094157 |
| SE_38_1_33 | SE_38 | SE_38_1 | GCCCGCACGCGAAGCCGGGG | 536.469536635708 | 722.560766390307 | 676.4559009046627 | 589.4927457735794 | 624.6901679976206 | 506.7914142730191 | 411.09335407609535 | 325.20160075132765 | 463.5283150247386 | 246.40434735148972 | 348.61413451528495 | 319.1872785001861 |
| SE_38_1_36 | SE_38 | SE_38_1 | GAAAATGCCCGCAGCACCGC | 861.6025891421976 | 942.4271130097641 | 824.5849302998442 | 873.1779964990546 | 1046.8067840639244 | 1096.7485990934053 | 971.7730333264767 | 897.1404625378195 | 985.6636583859384 | 888.6594657439822 | 1058.754038157532 | 723.1042709295125 |
| SE_38_2_14 | SE_38 | SE_38_2 | ACCCCGGCTTCGCGTGCGGG | 1.083776841688299 | 4.996962423169481 | 4.93763431317272 | 6.506542447832002 | 0.0 | 0.0 | 0.0 | 0.945353490556185 | 0.0 | 0.0 | 0.0 | 0.0 |
| SE_38_2_15 | SE_38 | SE_38_2 | AAAATGCCCGCAGCACCGCG | 93.2048083851937 | 140.91434033337936 | 121.46580410404891 | 100.20075369661284 | 167.57393301124625 | 109.1550738434195 | 66.72195799140815 | 265.64433084628797 | 102.11830694989453 | 114.45409033782215 | 176.06774470468935 | 162.4953417819129 |
| SE_38_3_12 | SE_38 | SE_38_3 | AATATTGCATCAACTCTGCA | 0.0 | 0.9993924846338962 | 3.950107450538176 | 0.0 | 0.0 | 0.0 | 0.0 | 2.836060471668555 | 0.0 | 0.0 | 0.0 | 1.1606810127279494 |
| SE_38_3_21 | SE_38 | SE_38_3 | TTTTTAAAGAAGTAGCAAAA | 2.167553683376598 | 0.0 | 0.0 | 0.0 | 0.0 | 0.0 | 0.0 | 0.0 | 0.0 | 0.0 | 0.0 | 0.0 |

| | | | | | | | | | | | | | | | |
|---|---|---|---|---|---|---|---|---|---|---|---|---|---|---|---|
| SE_38_3_24 | SE_38 | SE_38_3 | GCTACTTCTTTAAAAAAGGG | 13.005322100259587 | 39.975699385355846 | 37.526020780112674 | 14.314393385230405 | 26.51486281823517 | 34.65240439473635 | 2.152321225529295 | 3.78141396222474 | 42.62329333560815 | 2.9160277793075706 | 10.564064682281362 | 31.338387343654635 |
| SE_38_3_4 | SE_38 | SE_38_3 | TATACTACTTTTTTTCCCCG | 16.256652625324485 | 40.97509186998975 | 29.62580587903632 | 33.834020728726415 | 32.87842989461161 | 79.7005301078936 | 36.58946083399801 | 50.10373499947781 | 29.303514168230603 | 34.26332640686395 | 12.911634611677218 | 149.7278506419055 |
| SE_38_3_9 | SE_38 | SE_38_3 | TAATGCAATGAGATGCAAAA | 618.8365766040187 | 695.5771693051918 | 719.9070828605826 | 640.243776866669 | 750.90091501242 | 720.7700114105161 | 570.3651247652632 | 681.5998666910094 | 645.5652969788985 | 430.84310439269353 | 1065.7967479457195 | 1016.7565671496837 |
| SE_380_1_1 | SE_380 | SE_380_1 | CATGGTCCAGCTCTACACTA | 4472.74702564761 | 3988.57540617388 | 3908.6313223075254 | 4429.654098484027 | 4046.1680660626866 | 3738.1281240821836 | 4233.615850616123 | 3995.0638510904378 | 4209.050216891304 | 4158.255613292596 | 3637.559605598882 | 3892.924116689542 |
| SE_380_1_17 | SE_380 | SE_380_1 | CCAGAGGAGCTGCAGATGCA | 414.0027535249302 | 126.92284554850481 | 281.445155850845 | 450.2527373899745 | 72.12042686559965 | 1170.3849584322202 | 574.6697672163217 | 203.25100046957976 | 855.1298225456385 | 29.8892847379026 | 159.63475519891836 | 276.24208102925195 |
| SE_380_1_31 | SE_380 | SE_380_1 | GGATGCAGAGGAGTGTTGGG | 1829.4153087698485 | 1995.7867918138907 | 2016.529853499739 | 2023.5347012757527 | 2134.976754124296 | 1493.5186294131368 | 2002.734900355009 | 2228.198177240928 | 2294.553957900239 | 1837.0975009637696 | 2189.108959161638 | 1821.1085089701526 |
| SE_380_1_32 | SE_380 | SE_380_1 | GCTAAGCCCTTGGGACACAC | 135.47210521103736 | 161.90158251069118 | 158.9918248841616 | 139.24000838360485 | 107.12004578567009 | 127.34758615065608 | 166.80489497852037 | 95.48070254617468 | 69.26285167036325 | 153.82046535847434 | 61.03681816429231 | 225.1721164692222 |
| SE_380_1_4 | SE_380 | SE_380_1 | GAGAATGCCATCCCTGACCA | 2880.6788452074984 | 2628.402234587147 | 2696.93586185494 | 2757.4726893912025 | 2682.2435226926696 | 2580.7378172979897 | 2932.5376697836646 | 3355.0595379839006 | 2706.5791268111175 | 2851.875168162804 | 3824.1914149858526 | 3144.284863480015 |
| SE_381_1_13 | SE_381 | SE_381_1 | GCTTGCCAAAAGGCTCCACT | 825.8379533664838 | 1007.3876245109674 | 1075.4167534090184 | 896.6015493112499 | 988.4740858638071 | 954.6737410749864 | 779.1402836416048 | 859.3263229155722 | 1006.9753050537425 | 867.5182643440022 | 1012.9764245343127 | 669.7129443440268 |
| SE_381_1_15 | SE_381 | SE_381_1 | CCTTCGCTCCCCAAGGCTGA | 111.62901469389479 | 87.94653864778286 | 105.66537430189621 | 91.09159426964803 | 98.63528968383483 | 19.05882241710499 | 91.47365208499504 | 88.86322811228139 | 90.57449833816732 | 37.908361130998415 | 90.38144228174053 | 53.39132658548567 |
| SE_381_1_19 | SE_381 | SE_381_1 | GCACCTCTAAGATCATCAGA | 358.73013459882696 | 452.72479553915497 | 495.73848504254113 | 491.89460905609934 | 480.44931426642125 | 870.6416604177508 | 346.5237173102165 | 552.086438484812 | 507.03959363817194 | 349.9233335169085 | 559.8954281609122 | 798.5485367568292 |
| SE_381_1_8 | SE_381 | SE_381_1 | CAGCACTCCAGATCAAAGGA | 399.9136545829823 | 472.7126452318329 | 499.6885924930793 | 379.9820789533889 | 484.6916923173389 | 356.9197652657844 | 417.5503177526832 | 560.5946198998178 | 394.26546335437536 | 360.85843768931187 | 477.7304806320571 | 287.84889115653147 |
| SE_381_1_9 | SE_381 | SE_381_1 | ACTCTGACACTGAAAAGCAA | 252.52000411337366 | 282.82807315139263 | 238.98150075755964 | 236.83814510108488 | 277.8757623351046 | 129.94651648026132 | 193.70891029763655 | 276.043219242406 | 236.20408390149515 | 136.32429868262892 | 227.71428315139823 | 364.4538379965761 |
| SE_381_2_1 | SE_381 | SE_381_2 | TCCATATGCCAGAAATGTGA | 601.4961471370059 | 675.5893196125138 | 721.8821365858516 | 575.178352388349 | 746.6585369615024 | 613.3475577868334 | 681.2096678800218 | 815.8400623499876 | 645.5652969788985 | 815.7587712612928 | 569.2857078784956 | 912.2952760041683 |
| SE_381_2_3 | SE_381 | SE_381_2 | ATATGCCAGAAATGTGAGGG | 276.36309463051623 | 270.83536333578587 | 266.6322529113269 | 274.5760912985105 | 351.0567837134336 | 203.58287581907607 | 275.49711686774975 | 391.3763450902606 | 349.8661994631169 | 467.2934516340382 | 437.8217918323275 | 226.33279748195014 |
| SE_381_2_5 | SE_381 | SE_381_2 | CCTACAGTAAGAGTAAAAGA | 472.5267029760983 | 576.649463633758 | 596.4662250312646 | 605.1084476483762 | 588.6299545648208 | 698.2459485539374 | 485.348436356856 | 469.84068480642395 | 676.6447817027794 | 790.9725351371785 | 500.0323949613178 | 853.1005443550429 |
| SE_381_2_8 | SE_381 | SE_381_2 | GGTCCCATCAGGTTAGCAGA | 147.39365046960864 | 183.8882171726369 | 168.86709351050703 | 160.06094421666725 | 176.05868911308153 | 126.48127604078768 | 134.52007659558095 | 259.97220990295085 | 143.85361500767752 | 84.56480559991955 | 196.02208910455414 | 111.42537722188314 |
| SE_381_2_9 | SE_381 | SE_381_2 | GGGTACTGTATAATAGTCAC | 58.52394945116814 | 94.94228604022014 | 82.9522564613017 | 72.87327541571842 | 80.60518296743491 | 46.78074593289407 | 35.513300221233365 | 8.508181415005666 | 65.71091055906257 | 26.973256958595027 | 23.47569929395858 | 55.71268861094157 |
| SE_381_3_1 | SE_381 | SE_381_3 | CTCTGTACGCAGTTAGGCAG | 204.8338230790885 | 275.83232575895534 | 239.9690276201942 | 213.41459228888968 | 194.08879582948143 | 406.2994415282837 | 202.31819519975372 | 212.70453537514163 | 247.74789251322238 | 89.6678542137078 | 97.42415206992811 | 200.79781520193526 |
| SE_381_3_10 | SE_381 | SE_381_3 | AGGCATTCTAAGATGAAGGC | 433.51073667531955 | 547.6670815793751 | 578.6907415038428 | 461.96451379607214 | 491.05525939371535 | 415.8288527368362 | 330.3813081187468 | 362.07038688301884 | 524.7992991946753 | 320.7630557238328 | 350.9617044446808 | 515.3423696512095 |
| SE_381_3_12 | SE_381 | SE_381_3 | TGTACTTGAATATAACCAAT | 174.48807151181612 | 188.8851795958064 | 196.51784566427426 | 163.96486968536647 | 201.5129574185873 | 172.39571186381335 | 205.54667703804768 | 224.99413075237203 | 98.56636583859384 | 250.04938207562418 | 171.37260484589763 | 212.40462532921475 |
| SE_381_3_15 | SE_381 | SE_381_3 | TACCACGAGATACTTATGGA | 641.595890279473 | 704.5717016668968 | 708.0567605089681 | 638.9424683771026 | 668.1745430195263 | 910.4919254716975 | 575.7459278290864 | 471.7313917875363 | 711.276207537961 | 887.2014518543284 | 429.60529707944204 | 702.2120127004094 |
| SE_381_3_19 | SE_381 | SE_381_3 | TCAGCAGGTCCCTTCTGTGA | 2526.283817975425 | 2982.187174147546 | 2875.678223991792 | 2883.6996128791434 | 3069.360519838903 | 2600.662949824963 | 2587.0901130862126 | 3087.5245001565004 | 3147.907809890227 | 2915.298772362744 | 2837.0382596748946 | 3263.8350077909936 |
| SE_382_2_0 | SE_382 | SE_382_2 | CCAGCTCATCTTTATGACAA | 1050.1797595959617 | 1081.3426683738758 | 1110.967720463862 | 1221.9286717028501 | 1072.2610523694302 | 641.9357914124909 | 1353.8100508579266 | 1250.7026680058327 | 1060.2544217232528 | 1019.1517088679959 | 1139.7452007216891 | 970.3293266405657 |
| SE_382_2_2 | SE_382 | SE_382_2 | GCTGGTTTGGCTAATCCCAG | 1103.2848248386883 | 1296.2120525701635 | 1361.7995435730363 | 1149.0553962871315 | 1302.4100616317114 | 1311.5935063407708 | 1042.7996337689433 | 1194.926812063018 | 1244.955359510888 | 1152.5599797713173 | 1130.3549210041056 | 1142.1101165243022 |
| SE_382_2_3 | SE_382 | SE_382_2 | TCACACTTACTAGTACCTCT | 2314.9473338462067 | 2072.740013130701 | 2095.5320025105025 | 2406.119397208274 | 2111.643674844249 | 2449.0586805979915 | 2095.2847130527684 | 1870.85455781069 | 2513.886321523056 | 2166.608640025525 | 2190.2827441263357 | 2380.556757105024 |
| SE_382_2_5 | SE_382 | SE_382_2 | ATTGTTCTGTTGAGCACAAG | 189.6609472954523 | 126.92284554850481 | 157.0167711588925 | 197.79889041409285 | 186.6646342403756 | 107.42245362368269 | 214.15596194016484 | 232.55695867682152 | 180.26101139850945 | 136.32429868262892 | 181.936669528179 | 131.15695443825828 |
| SE_382_2_6 | SE_382 | SE_382_2 | TCTTGTGCTCAACAGAACAA | 1789.3155656273816 | 1847.8767040880741 | 1861.4881360661154 | 1958.4692767974327 | 2017.2507632113316 | 1689.304714243397 | 1859.6055388573109 | 2121.373232808079 | 2453.5033226309442 | 1998.9370427153397 | 1665.6008649063613 | 1534.4202988263492 |
| SE_383_1_13 | SE_383 | SE_383_1 | GTATCTATCTCACAGGGTGT | 108.37768416882989 | 120.92649064070145 | 96.77763253818532 | 143.14393385230406 | 109.24123481112889 | 115.21924461249836 | 123.75847046793446 | 84.13646065950047 | 114.5501008394469 | 121.74415978609107 | 142.02798072844942 | 24.37430126728694 |
| SE_383_1_2 | SE_383 | SE_383_1 | GGAGTGCTCTGCCCAGTACT | 835.5919449416784 | 805.5103426149203 | 881.8614883326478 | 823.7282738955315 | 718.0224851178084 | 959.0052916243285 | 848.0145628585423 | 813.0040018783191 | 812.5065292100304 | 581.7475419178603 | 483.5994054555468 | 1219.8757443770749 |
| SE_383_1_26 | SE_383 | SE_383_1 | GGAGTGCTGGGGGCATGTGG | 1100.0334943136234 | 1621.0146100761797 | 1489.1905088528924 | 1124.33053498537 | 1477.4081562320637 | 1346.245910735507 | 1010.514815386004 | 1401.013873004266 | 1590.381632584879 | 1599.4412369502024 | 1115.0957164630327 | 1486.8323773045033 |
| SE_383_1_27 | SE_383 | SE_383_1 | CTGAGGAGCCTAGTTTTGGA | 470.3591492927217 | 530.6774093405988 | 520.4266566084048 | 486.68937509783376 | 479.3887197536919 | 701.711188993411 | 628.4777978545541 | 596.5180525409528 | 807.1786175430793 | 472.39650024782645 | 985.9793703462603 | 507.2176025621139 |
| SE_383_1_7 | SE_383 | SE_383_1 | TCTTGCAGCCCACTTACTAA | 332.71949039830776 | 391.7618539764873 | 374.2726809384922 | 309.7114205168033 | 408.3288874008216 | 298.0106777947326 | 286.2587229953962 | 437.69866612751366 | 399.5933750213264 | 250.04938207562418 | 434.30043693823376 | 289.0095721692594 |
| SE_383_2_12 | SE_383 | SE_383_2 | ACTCCTGGAGAGCAGCACGG | 198.3311620289587 | 163.900367479959 | 229.10623213121423 | 186.08711400799527 | 201.5129574185873 | 154.20319955657675 | 215.2321225529295 | 307.2398844307601 | 221.1083341784673 | 118.09912506195661 | 246.49484258656508 | 251.86777976196504 |
| SE_383_2_16 | SE_383 | SE_383_2 | AGGGGCTTCACTCTACAGGG | 411.83519984155356 | 602.6336682342394 | 668.5556860035863 | 523.126012805693 | 567.4180643102326 | 331.79677207960054 | 378.80853569315593 | 602.1901734842899 | 420.9050216891305 | 545.2971947305157 | 657.3195802308403 | 385.3460962256792 |
| SE_383_2_17 | SE_383 | SE_383_2 | CGAAGGGGCTTCACTCTACA | 6974.103976264203 | 7058.709118969209 | 7123.031260182966 | 6809.747325900974 | 6530.080414874958 | 7234.555727511081 | 6889.580242919273 | 7637.510850203419 | 6125.322446438021 | 7932.324566661419 | 7852.621413829145 | 8583.236089123186 |
| SE_383_2_22 | SE_383 | SE_383_2 | GATTGCTTCTGACCACAGGA | 106.21013048545329 | 217.86756165018937 | 215.2808560543306 | 137.93869989403845 | 193.02820131675202 | 115.21924461249836 | 111.92070372752333 | 190.96140509234937 | 79.03068972644012 | 231.09520151012498 | 272.3181118099195 | 85.89039494186825 |
| SE_383_2_9 | SE_383 | SE_383_2 | GTCACTGAAGGATAAACAGG | 354.39502723207374 | 489.7023174706091 | 524.3767640589429 | 333.1349733289985 | 468.7827746263978 | 578.695153392097 | 379.88469630592056 | 465.11391735364305 | 557.6547544742066 | 512.4918822133055 | 519.9867393611826 | 754.4426582731671 |
| SE_384_1_0 | SE_384 | SE_384_1 | AGTATCTGTTGCGAACTACA | 34.680858934025565 | 81.9501837399795 | 72.08946097232172 | 54.65495656178882 | 69.99923784014085 | 31.187163955262715 | 45.19874573611519 | 202.30564697902358 | 58.607028336461205 | 45.92743752409424 | 134.98527094026184 | 20.89225822910309 |
| SE_384_1_2 | SE_384 | SE_384_1 | TGACTCATATTTCTCTACAT | 248.18489674662044 | 241.8529812814029 | 272.55741408713413 | 235.53683661151848 | 221.664253160446 | 190.58822417104992 | 295.9441685102781 | 263.7536238651756 | 237.09206917932033 | 322.95007655831347 | 156.11340030482455 | 313.38387343654637 |
| SE_384_1_21 | SE_384 | SE_384_1 | AGAGACGTCTAGTAACTGAA | 848.597267041938 | 980.4040274258522 | 908.5247136237805 | 936.9421124878083 | 990.5952748892659 | 993.6576960190648 | 963.1637484243595 | 872.5612717833587 | 1038.9427750554487 | 972.4952643990748 | 1157.351975192158 | 986.578860818757 |
| SE_384_1_4 | SE_384 | SE_384_1 | TTTCCTGGCCTGTGTTACAG | 52.02128840103835 | 31.980559508284678 | 37.526020780112674 | 62.46280749918722 | 42.42378050917627 | 0.0 | 49.50338818717378 | 26.46989773557318 | 80.80666028209045 | 43.74041668961356 | 0.0 | 1.1606810127279494 |
| SE_384_1_8 | SE_384 | SE_384_1 | AAGCTAGTAGATAATCCCTA | 179.90695572025763 | 223.86391655799275 | 225.15612468067604 | 166.56748666449926 | 240.75495438957532 | 221.77538812631263 | 184.02346478275473 | 263.7536238651756 | 277.051406681453 | 314.93100016521765 | 213.62886357502308 | 117.2287822855229 |
| SE_385_1_12 | SE_385 | SE_385_1 | TCTTTATTATACAGCACATC | 42.26729682584366 | 47.97083926242702 | 35.55096705484358 | 31.23140374959361 | 56.21150917465856 | 47.64705604276248 | 68.87427921693744 | 10.398888396118036 | 15.095749723027886 | 59.04956253097831 | 65.73195802308402 | 22.05293924183104 |
| SE_385_1_13 | SE_385 | SE_385_1 | GAGTATTCTGTATACTTGTT | 10.837768416882989 | 38.97630690072195 | 23.700644703229056 | 24.724861301761607 | 21.211890254588134 | 28.588233625657487 | 7.533124289352532 | 1.89070698111237 | 71.92680750383876 | 40.82438891030599 | 2.347569929395858 | 87.05107595459621 |
| SE_385_1_14 | SE_385 | SE_385_1 | AGAGTATTCTGTATACTTGT | 78.03193260155751 | 79.95139877071169 | 72.08946097232172 | 98.89944520704643 | 85.90815553108195 | 178.4598826328922 | 66.72195799140815 | 67.12009782948914 | 134.9737622294258 | 196.831875103261 | 63.384388093688166 | 74.28358481458876 |
| SE_385_1_15 | SE_385 | SE_385_1 | TTCATTAGTGTAATATTGAA | 1.083776841688299 | 5.9963549078033775 | 10.862795488979984 | 0.0 | 5.3029725636470335 | 1.7326202197368175 | 0.0 | 0.0 | 2.6639558334755096 | 2.9160277793075706 | 7.042709788187574 | 0.0 |
| SE_385_1_17 | SE_385 | SE_385_1 | TTTTGAAGAGATGCGACTCT | 23.843090517142578 | 22.986027146579612 | 20.738064115325425 | 16.917010364363207 | 13.787728665482287 | 0.0 | 19.370891029763655 | 56.7212094333711 | 33.74344055735645 | 12.393118062057175 | 34.03976397623994 | 0.0 |
| SE_385_2_15 | SE_385 | SE_385_2 | TTTTCAGTTTGATATCACGA | 7.586437891818092 | 2.9981774539016888 | 20.738064115325425 | 5.205233958265602 | 2.1211890254588135 | 0.8663101098684087 | 12.913927353175769 | 6.617474433893295 | 25.751573056929924 | 2.9160277793075706 | 10.564064682281362 | 2.321362025455899 |
| SE_385_2_16 | SE_385 | SE_385_2 | TATCACGATGGACAAGTCCC | 1198.6571869072586 | 1149.3013573289807 | 1140.5935263428983 | 1159.4658642036627 | 1084.988186522183 | 1320.256607439455 | 1288.164253479283 | 1167.5115608368885 | 1135.7331703383923 | 749.4191392820456 | 1392.1089681317437 | 1547.1877899663566 |
| SE_385_2_2 | SE_385 | SE_385_2 | GCAGATAGCACTAGATGTGG | 4966.949265457474 | 5067.919289578487 | 5088.7259231558055 | 5147.97638472468 | 5326.305642927081 | 5078.309864048612 | 5040.736310189609 | 5151.231170040652 | 5316.367858339291 | 4944.125099815986 | 5098.921886647804 | 6097.0573598599185 |
| SE_385_2_3 | SE_385 | SE_385_2 | TTTGCAGATAGCACTAGATG | 261.19021884688004 | 271.8347558204198 | 244.90666193336693 | 301.9035695794049 | 254.5426830550576 | 274.62030482828555 | 303.47729279963056 | 310.07594490242866 | 341.87433196269035 | 184.43875704120384 | 258.2326922335444 | 443.3801468620767 |
| SE_385_2_5 | SE_385 | SE_385_2 | TATCTGCAAACGTGTCTCCA | 794.4084249575232 | 880.4647789624626 | 956.9135298928732 | 748.2523815006803 | 872.8692839763017 | 789.2085100901204 | 697.3520770714915 | 724.1407737660377 | 841.810043378261 | 1330.437674309079 | 622.1060312899024 | 798.5485367568292 |
| SE_386_1_10 | SE_386 | SE_386_1 | ACTCAGTGGGCTGATCCCAA | 351.14369670700887 | 403.7545637920941 | 430.5617121086612 | 404.7069402551505 | 370.147484942563 | 547.5079894368344 | 317.467380765571 | 323.3108937702153 | 388.0495664095992 | 432.3011182823473 | 262.9278320923361 | 344.722260780201 |
| SE_386_1_11 | SE_386 | SE_386_1 | ATAGCTTATCAGAACTCAGT | 134.38832836934907 | 223.86391655799275 | 217.25590977959968 | 173.07402911233126 | 185.60403972764618 | 119.5507951618404 | 203.39435581251837 | 257.1361494312823 | 165.1652616754816 | 341.1752501789858 | 201.8910139280438 | 81.24767089095646 |
| SE_386_1_12 | SE_386 | SE_386_1 | AATAGCTTATCAGAACTCAG | 48.769957875973454 | 36.977521931454156 | 68.13935352178353 | 28.62878677046081 | 72.12042686559965 | 19.05882241710499 | 54.88419125099702 | 79.40969320671954 | 18.647690834328564 | 19.683187510326103 | 36.3873339056358 | 3.482043038183848 |
| SE_386_1_7 | SE_386 | SE_386_1 | TGATGATGCTCTCTGGAATG | 2309.528449637765 | 2326.5857042277103 | 2378.952212086617 | 2515.429310331852 | 2391.640626204812 | 2140.652281484838 | 2320.2022811205798 | 2049.5263675258093 | 2199.5395331729455 | 2276.6886886943857 | 1889.7937931636657 | 2154.2239596230743 |
| SE_386_1_9 | SE_386 | SE_386_1 | GAATGTGGATGCAATCCATT | 725.046707089472 | 748.5449709907882 | 752.4954693275225 | 805.5099550416019 | 813.475991263455 | 991.9250757993279 | 794.2065322203098 | 679.709159709897 | 814.2824997656807 | 647.3581670062807 | 625.6273861839961 | 624.4463848476368 |
| SE_387_1_10 | SE_387 | SE_387_1 | GCCTTATCTTTCTCTTCCAG | 280.6982019972694 | 399.75699385355847 | 385.13547642747216 | 288.8904846837409 | 354.2385672516219 | 608.149697127623 | 276.5732774805144 | 300.62240999686685 | 384.49762529829854 | 221.61811122737538 | 285.22974642159676 | 714.9795038404169 |
| SE_387_1_19 | SE_387 | SE_387_1 | ATGGAGCATTAAGGCCACCT | 5471.989273684221 | 4724.128274864427 | 4588.0498038000915 | 5552.683324979831 | 4711.160825544025 | 4699.732346036118 | 5682.128035397339 | 5133.269453720084 | 4327.152258842052 | 4193.976953589114 | 4399.346047687838 | 4965.393372450168 |
| SE_387_1_23 | SE_387 | SE_387_1 | CATACACAGCAGTCTCCAGA | 2025.5789171154306 | 2755.3250801356517 | 2749.2747855745706 | 2283.796399189033 | 2903.9077758531157 | 3215.743127831533 | 1629.3071677256762 | 2248.0506005426078 | 2705.6911415332925 | 2452.3793623976667 | 1923.8335571399057 | 2012.6208760702643 |
| SE_387_1_34 | SE_387 | SE_387_1 | GATGGCCTTCCGCCATAACG | 524.5479913771367 | 368.7758268299077 | 344.64687505945585 | 573.8770438987826 | 514.3883386737623 | 511.98927493222953 | 746.8554652586654 | 397.9938195241539 | 466.19227085821416 | 634.9650489442236 | 436.6480068676296 | 235.61824558377373 |
| SE_387_1_6 | SE_387 | SE_387_1 | CTTTCCAGCAGACATTGCTC | 113.79656837727138 | 153.90644263362 | 157.0167711588925 | 148.34916781056964 | 160.14977142214042 | 118.68448505197199 | 108.6922218892294 | 149.36585150787724 | 101.23032167206935 | 106.43501394472632 | 115.03092654039705 | 2.321362025455899 |
| SE_387_2_12 | SE_387 | SE_387_2 | GTGCACAGACAGGTCCCAGG | 1080.5255111632341 | 1322.1962571706447 | 1314.398254166578 | 1201.1077358697876 | 1359.6821653190993 | 1236.2245267822193 | 1022.3525821264151 | 1217.6152958363664 | 1367.4973278507614 | 1472.5940285503232 | 1259.4712671208779 | 1391.6565342608114 |
| SE_387_2_15 | SE_387 | SE_387_2 | TCTTCCGTGTGTGTATGTGA | 841.01082915012 | 927.4362257402556 | 900.6244987227042 | 870.5753795199219 | 818.778963827102 | 609.8823173473597 | 823.2628687649553 | 821.5121832933248 | 924.3926742160018 | 625.487958661474 | 703.0971938540595 | 530.4312228166729 |
| SE_387_2_3 | SE_387 | SE_387_2 | TATCCCTTTGTTCTTTAAAA | 0.0 | 7.9951398770711695 | 14.81290293951816 | 18.218318853929606 | 24.393673792776355 | 13.86096175789454 | 8.60928490211718 | 12.289595377230405 | 18.647690834328564 | 0.0 | 0.0 | 0.0 |
| SE_387_2_4 | SE_387 | SE_387_2 | GGAGTACTCAAGGCCCAGAA | 343.5572588151908 | 483.7059625628058 | 462.1625717129666 | 458.06058832737295 | 585.4481710266325 | 430.5561246045991 | 338.99059302086397 | 592.736638578728 | 497.27175558209507 | 699.846667033817 | 521.1605243258805 | 354.0077088820246 |
| SE_387_2_5 | SE_387 | SE_387_2 | CTTGCTGCTTCGACCCAGGA | 1324.3753005431013 | 1425.1336830879359 | 1437.839111995896 | 1341.649052742959 | 1387.257622650064 | 1629.5293166624767 | 1273.098004900578 | 1613.7184083794077 | 1387.9209892407405 | 1238.5827992608906 | 1462.5360660136196 | 1463.6187570499442 |
| SE_387_3_18 | SE_387 | SE_387_3 | GGCCACGTGACAAGGCCACA | 2495.9380664081523 | 2006.7801091448634 | 1951.353080565859 | 2286.3990161681654 | 1806.1924551781797 | 1830.5132621519476 | 2681.7922470095014 | 2094.903335072506 | 2111.6289906682537 | 1958.1126538050337 | 1987.2179452335938 | 2430.4660406523262 |
| SE_387_3_2 | SE_387 | SE_387_3 | GCCTATGTAGAGAGACCACA | 726.1304839311603 | 715.5650189978696 | 742.6202007011772 | 769.0733173337427 | 657.5685978922322 | 647.1336520717014 | 884.6040236925402 | 978.4408627256515 | 808.0666028209044 | 755.2511948406608 | 930.8114770054577 | 673.1949873822107 |
| SE_387_3_24 | SE_387 | SE_387_3 | TTACAGTGACAGGCAAGCGG | 654.6012123797325 | 761.537073291029 | 687.3186963936427 | 727.4314456676178 | 761.506860139714 | 912.2245456914344 | 531.6233427057358 | 696.7255225399083 | 728.1479278166393 | 461.46139607542307 | 1150.3092654039704 | 899.5277848641608 |
| SE_387_3_31 | SE_387 | SE_387_3 | ACATGTCAAGCTGCAGGCAA | 945.0534059521966 | 182.888824688003 | 328.8464452573032 | 830.2348163433635 | 189.84641777856382 | 1286.470513154587 | 1163.329622398584 | 465.11391735364305 | 1509.5749723027886 | 65.61062503442034 | 381.4801135268269 | 480.52193926937105 |
| SE_387_3_8 | SE_387 | SE_387_3 | GGGTGCTCACGGCTTCTCCA | 44.434850509220254 | 68.95808143973883 | 56.28903117016901 | 46.847105624390416 | 32.87842989461161 | 39.8502650539468 | 3.228481838293942 | 51.04908849003399 | 34.63142583518162 | 34.26332640686395 | 23.47569929395858 | 18.57089620364719 |
| SE_388_1_1 | SE_388 | SE_388_1 | CTCAGGATACCAGTAGGCAT | 384.7407787993461 | 472.7126452318329 | 489.81332386673387 | 455.45797134824016 | 379.69283555712764 | 674.8555755874904 | 411.09335407609535 | 380.9774566941426 | 312.57081779445974 | 403.8698474340985 | 410.82473764427516 | 516.5030506639375 |
| SE_388_1_10 | SE_388 | SE_388_1 | TCTAGGTTGTTATTAGTGGG | 159.31519572817993 | 220.86573910409106 | 205.40558742798515 | 199.10019890365928 | 167.57393301124625 | 148.1390287874979 | 172.1856980423436 | 191.90675858290555 | 259.29170112494955 | 275.5646251445654 | 124.42120625798047 | 121.87150633643469 |
| SE_388_1_14 | SE_388 | SE_388_1 | GATGCAGTAGAATTAGGAAC | 1067.5201890629744 | 1431.1300379957393 | 1376.6124465125545 | 1158.1645557140964 | 1269.5316317370998 | 1540.2993753460307 | 884.6040236925402 | 1341.4566030992264 | 1430.5442825763485 | 1170.7851533919895 | 1262.9926220149716 | 1688.7908735191663 |
| SE_388_1_18 | SE_388 | SE_388_1 | TTAGAAGAGGAATAACTGCC | 447.59983561726744 | 513.6877371018227 | 534.2520326852883 | 485.38806660826737 | 487.8734758555271 | 340.4598731782846 | 358.3614840506276 | 445.26149405196315 | 449.3205505795359 | 530.7170558339778 | 515.2915995023908 | 183.38760001101602 |
| SE_388_1_4 | SE_388 | SE_388_1 | TGAGACCCTACTACAGCCAG | 1981.1440666062103 | 1609.021900260573 | 1808.16168548385 | 1763.2730033624725 | 1618.4672264250746 | 1653.7859997387923 | 1683.1151983639086 | 1679.8931527183408 | 1565.5180448057743 | 1815.2272926189628 | 1183.1752444155125 | 1593.6150304754747 |
| SE_389_1_21 | SE_389 | SE_389_1 | CAGGCAGGCAAGTTACCCCA | 29298.82313820147 | 25982.205815512032 | 26033.18315277185 | 28148.603937810807 | 26979.40321481065 | 26062.939655391077 | 29113.373057122008 | 27624.17434754228 | 25645.014823590904 | 25045.762596472723 | 25319.715473499025 | 26077.02031295884 |
| SE_389_1_27 | SE_389 | SE_389_1 | TCACACCCTGTTTACAGAGG | 550.5586355776559 | 555.6622214564463 | 532.2769789600193 | 525.7286297848258 | 459.2374240118331 | 583.8930140513074 | 431.5404057186236 | 614.4797688615203 | 741.4677069840168 | 664.8543336821261 | 835.7348948649254 | 529.270541803945 |
| SE_389_1_29 | SE_389 | SE_389_1 | CGTCACACCCTGTTTACAGA | 11554.144909238954 | 11998.706170514557 | 11991.538692971268 | 11651.91621557755 | 12487.439792876035 | 12312.865591559694 | 11062.931099220576 | 12389.80284722936 | 12000.233044529345 | 11343.348061506449 | 12910.460826712522 | 14694.221621135839 |

| | | | | | | | | | | | | | | | |
|---|---|---|---|---|---|---|---|---|---|---|---|---|---|---|---|
| SE_389_1_39 | SE_389 | SE_389_1 | GCCATGTCGGAGGCTCCGAG | 341.38970513181414 | 580.6470335722937 | 609.3040742455137 | 498.4011515039314 | 679.8410826595497 | 472.13900987828276 | 327.1528262804528 | 734.5396621621558 | 525.6872844725006 | 688.9115628614136 | 531.7245890081618 | 555.9662050966878 |
| SE_389_1_50 | SE_389 | SE_389_1 | GAGGACAGCCATCGGCTCAG | 3959.0368026873557 | 3865.6501305639104 | 3949.1199236755415 | 4122.545294946357 | 3726.929117731135 | 4361.00509307757 | 4334.774948216 | 4853.444820515454 | 4379.543390233737 | 5055.663162374501 | 5847.7966941250825 | 4472.103942040789 |
| SE_389_2_13 | SE_389 | SE_389_2 | GTGCCGAGCCTCAGAGTTTG | 852.9323744086912 | 750.543755960056 | 804.8343930471534 | 871.8766880094882 | 933.3231712018779 | 936.4812287677498 | 872.7662569521291 | 1040.8341931023597 | 927.9446153273025 | 933.8578963232495 | 888.5552182763323 | 840.3330532150354 |
| SE_389_2_17 | SE_389 | SE_389_2 | AGAATGTTCTCCCCTCTCCC | 295.8710777809056 | 344.7904071986942 | 296.25805879036324 | 311.0127290063697 | 322.42073186973965 | 484.2673514164405 | 344.3713960846872 | 321.4201867891029 | 369.40187557527065 | 183.70975009637695 | 482.4256204908488 | 183.38760001101602 |
| SE_389_2_20 | SE_389 | SE_389_2 | CCGGCAGGCAGGATTCAGCC | 1309.202424759465 | 1058.356641227296 | 1085.292022035364 | 1211.5182037863187 | 1087.109375547642 | 1099.3475294230107 | 1307.5351445090466 | 822.457536783881 | 1150.8289200614201 | 1053.41503527486 | 955.4609612641142 | 958.7225165132862 |
| SE_389_2_23 | SE_389 | SE_389_2 | TGCCTGCCGGTGTTACCTCA | 1135.7981300893373 | 989.3985597875572 | 992.4644969477167 | 1016.3219303513588 | 1087.109375547642 | 1074.224536236827 | 956.7067847477716 | 889.57763461337 | 1058.4784511676023 | 776.3923962406407 | 1018.8453493578024 | 961.0438785387421 |
| SE_389_2_31 | SE_389 | SE_389_2 | GGGTCTTGAGAAATAGCTGA | 31.429528408960667 | 71.95625889364052 | 69.12688038441809 | 65.06542447832003 | 89.08993906927017 | 71.03742900920952 | 49.50338818717378 | 25.524544245016994 | 23.087617223454416 | 23.328222234460565 | 115.03092654039705 | 55.71268861094157 |
| SE_389_3_1 | SE_389 | SE_389_3 | TAAGAAAAGTGAATTAAACA | 287.2008630473992 | 405.75334876136185 | 459.199991125063 | 320.1218884333345 | 386.05640263350404 | 549.2406096565711 | 347.59987792298114 | 504.8187639570028 | 374.7297872422217 | 357.2134029651774 | 572.8070627725893 | 341.24021774201714 |
| SE_389_3_20 | SE_389 | SE_389_3 | GCGCCTCTCCCCCTTTCAGG | 49.85373471766175 | 105.935603371193 | 95.79010567555078 | 72.87327541571842 | 78.4839939419761 | 92.69518175591973 | 45.19874573611519 | 76.57363273505099 | 154.50943834157954 | 109.3510417240339 | 14.085419576375148 | 75.44426582731671 |
| SE_389_3_21 | SE_389 | SE_389_3 | TCAAACGATCACAAACGTGG | 2049.4220076325732 | 2154.69019687068 | 2119.2326472137315 | 2158.8707841906585 | 2145.58269925159 | 1956.9945381927353 | 1943.5460666529534 | 1970.1166743190895 | 2424.1998084627135 | 2045.5934871842608 | 1679.6862844827365 | 2233.1502684885745 |
| SE_389_3_26 | SE_389 | SE_389_3 | CTACTTCCCCACGCACCCCA | 1492.3607110047876 | 1493.092372043041 | 1450.6769612101452 | 1483.4916781056966 | 1345.894436653617 | 1313.3261265605076 | 1383.9425480153366 | 1973.8980882813144 | 1188.1243017300771 | 1848.0326051361728 | 1605.7378317067669 | 1050.4163165187942 |
| SE_389_3_8 | SE_389 | SE_389_3 | GAGCTGCTGTACTTGACGGA | 2253.1720538699733 | 2363.5632261591645 | 2457.95436109738 | 2492.005757519657 | 2401.1859768193767 | 1936.2030955558935 | 2223.3478259717617 | 2829.4429972346616 | 2096.5332409452258 | 1947.1775496326302 | 2901.5964327332804 | 1961.5509115102345 |
| SE_389_4_10 | SE_389 | SE_389_4 | CCTACCCCCCCCCCAGTCCA | 133.30455152766078 | 192.88274953434197 | 176.76730841158337 | 186.08711400799527 | 176.05868911308153 | 195.78608483026036 | 152.81480701257993 | 121.00524679119168 | 191.80482001023668 | 161.1105348067433 | 205.41236882213758 | 197.3157721637514 |
| SE_389_4_12 | SE_389 | SE_389_4 | GGGAGGTAAGGCATGAGCAC | 2287.852912803999 | 1914.836000558545 | 1896.0515762583245 | 2033.9451691922839 | 1866.6463424037559 | 2342.502537084177 | 2151.2450649165303 | 1807.5158739434257 | 2053.9099476096176 | 2831.462973707651 | 1758.3298771174977 | 1474.0648861644959 |
| SE_389_4_36 | SE_389 | SE_389_4 | GCCCCTTTTGATCTCCCCAT | 19.50798315038938 | 20.98724217731182 | 21.725590977959968 | 11.711776406097604 | 15.9089176909411 | 25.989303296052263 | 6.456963676587884 | 5.67212094333711 | 29.303514168230603 | 20.412194455152996 | 17.606774470468935 | 22.05293924183104 |
| SE_389_4_41 | SE_389 | SE_389_4 | GCCGCAGCCCTGGGACCCTG | 58.52394945116814 | 101.9380334326574 | 63.20171920861082 | 81.98243484268323 | 112.42301834931712 | 25.12299318618385 | 97.93061576158291 | 88.86322811228139 | 106.55823333902038 | 53.94651391719005 | 117.3784964697929 | 121.87150633643469 |
| SE_389_4_6 | SE_389 | SE_389_4 | GGATGACTGAGAAGCCAGTG | 671.9416418467454 | 783.5237079529746 | 813.7221348108643 | 710.5144353032547 | 747.7191314742317 | 1136.5988641473523 | 779.1402836416048 | 803.5504669727572 | 703.2843400375344 | 1143.8118964333946 | 994.1958650991459 | 1058.54108360789 |
| SE_389_5_1 | SE_389 | SE_389_5 | CGCGGCAAGGAACGGGCCCT | 23961.222192886602 | 21895.68994584403 | 22287.493762799026 | 23893.32517692868 | 23423.229813628946 | 22486.811521854284 | 24446.064479561734 | 24978.129927475522 | 23091.169164565716 | 23115.35220657111 | 25475.82887380385 | 23713.873771044735 |
| SE_389_5_16 | SE_389 | SE_389_5 | TCCCTATTCGAGGCCCCAGC | 37.93218945909046 | 34.97873696218637 | 48.38881626909266 | 18.218318853929606 | 48.78734758555271 | 51.11229648223612 | 6.456963676587884 | 74.68292575393862 | 12.431793889552377 | 36.45034724134463 | 21.128129364562724 | 15.088853165463343 |
| SE_389_5_2 | SE_389 | SE_389_5 | TCTGGGAGAGAGGACCTCTG | 792.2408712741465 | 875.467816539293 | 915.4374016622223 | 838.0426672807619 | 872.8692839763017 | 547.5079894368344 | 739.3223409693128 | 699.5615830115769 | 943.0403650503304 | 736.2970142751616 | 821.6494752885503 | 723.1042709295125 |
| SE_389_5_20 | SE_389 | SE_389_5 | TTCTGAGTCACACAGCCAGA | 9.75399157519469 | 20.98724217731182 | 15.800429802152705 | 9.109159426964803 | 11.666539640023474 | 24.256683076315444 | 12.913927353175769 | 13.23494886778659 | 9.767838056076867 | 49.5724722482287 | 1.173784964697929 | 25.534982280014887 |
| SE_389_5_4 | SE_389 | SE_389_5 | CAGAGGTCCTCTCTCCCAGA | 643.7634439628496 | 735.5528686905476 | 769.2834259923098 | 732.6366796258834 | 947.1108998673602 | 741.5614540473579 | 780.2164442543694 | 965.2059138578649 | 816.058470321331 | 925.1098129853268 | 915.5522724643846 | 623.2857038349089 |
| SE_389_6_0 | SE_389 | SE_389_6 | GGCGGGGAGAGGGGATGGGG | 3.2513305250648967 | 3.9975699385355847 | 3.950107450538176 | 1.3013084895664004 | 1.0605945127294067 | 11.262031428289314 | 1.0761606127646475 | 0.945353490556185 | 4.439926389125849 | 2.9160277793075706 | 0.0 | 11.606810127279495 |
| SE_389_6_13 | SE_389 | SE_389_6 | AGACTCACACACCTGCCACC | 3641.4901880726843 | 3341.968468615749 | 3252.913485518188 | 3577.297037818035 | 3142.5415412172324 | 3402.8661115631094 | 3556.71082518716 | 3508.2068034540025 | 3199.4109560040865 | 3099.7375294039475 | 3658.6877349634447 | 3587.6650103420916 |
| SE_389_6_21 | SE_389 | SE_389_6 | GAATAGGGACCGGGGCCGCA | 537.5533134773963 | 707.5698791207985 | 695.218911294719 | 658.4620957205987 | 724.3860521941848 | 634.1390004236752 | 622.0208341779662 | 650.4032015026553 | 701.5083694818841 | 891.5754935232898 | 821.6494752885503 | 530.4312228166729 |
| SE_389_6_41 | SE_389 | SE_389_6 | AGAGCCCTGGACTGGGGGGG | 144.14231994454374 | 164.89975996459287 | 166.89203978523796 | 160.06094421666725 | 186.6646342403756 | 195.78608483026036 | 126.9869523062284 | 202.30564697902358 | 249.5238630688727 | 167.67159731018532 | 308.70544571555536 | 269.2779949528843 |
| SE_39_1_10 | SE_39 | SE_39_1 | TCCAGCAACTTCAGACGCTG | 924.461645960119 | 1003.3900545724317 | 1059.6163236068658 | 848.4531351972931 | 1033.019055398442 | 1522.9731731486625 | 944.8690180073605 | 1103.227523479068 | 1171.252581451399 | 1223.2736534195258 | 1135.0500608628975 | 901.8491468896167 |
| SE_39_1_13 | SE_39 | SE_39_1 | CGGTAGTGTCTGTCATTTCG | 9131.903668065606 | 7865.218854068763 | 8577.658328843649 | 8924.373621446373 | 9302.474471149626 | 8551.347094511062 | 8681.38766317241 | 7985.400934728094 | 8200.544040715444 | 9023.647963067277 | 9356.239953607192 | 9004.563296743432 |
| SE_39_1_4 | SE_39 | SE_39_1 | GTAACCGAACTGCAGCGCCC | 317.54661461467157 | 96.94107100948793 | 177.75483527421792 | 191.29234796626088 | 83.78696650562313 | 756.2887259151208 | 444.4543330717994 | 138.021609621203 | 650.8932086458494 | 21.87020834480678 | 75.12223774066746 | 218.20803039285448 |
| SE_39_1_6 | SE_39 | SE_39_1 | GCTGTATATACTGCTGACCG | 2224.9938559860775 | 2497.4818191001063 | 2561.644681674007 | 2302.014718042962 | 2368.307546924765 | 2141.5185915947063 | 2118.960246533591 | 2231.9795912031527 | 2535.1979681908597 | 1692.7541258880447 | 1934.3976218221871 | 1795.5735266901377 |
| SE_39_1_9 | SE_39 | SE_39_1 | CGGTCAGCAGTATATACAGC | 3568.8771396795682 | 3736.728500046138 | 3907.6437954448907 | 3422.441327559633 | 3217.84375162102 | 3052.0105170664037 | 3129.4750619195947 | 3122.502579307079 | 3224.2745437831913 | 3341.7678350864758 | 2578.80556744135 | 3451.865331852922 |
| SE_39_10_1 | SE_39 | SE_39_10 | TCCTACATAGATCCACAGAC | 347.89236618194394 | 396.7588163996568 | 460.1875179876975 | 356.55852614119374 | 479.3887197536919 | 268.5561340592067 | 321.7720232166296 | 428.2451312219518 | 420.0170364113053 | 504.4728058202097 | 278.1870366334092 | 439.8981038238928 |
| SE_39_10_17 | SE_39 | SE_39_10 | TGAACAGCGTTATCTTTCCA | 1476.1040583794631 | 1510.0820442818172 | 1527.7040564956396 | 1405.4131687317124 | 1399.984756802817 | 1520.3742428190574 | 1274.1741655133426 | 1538.090129134913 | 1266.267006178692 | 1406.2543965710759 | 1392.1089681317437 | 1209.4296152625234 |
| SE_39_10_20 | SE_39 | SE_39_10 | AGGAAGTGAATCTAATTGAA | 1291.8619952924523 | 1838.882171726369 | 1767.6730841158337 | 1314.3215744620645 | 1705.435976468886 | 1591.411671828267 | 1266.64104122399 | 1557.942552436593 | 2051.245991776142 | 1799.1891398327712 | 1914.4432774223224 | 2185.562346966729 |
| SE_39_10_6 | SE_39 | SE_39_10 | TATATTACAGAAGACCAGAG | 20.59175999207768 | 42.973876839257535 | 40.488601368016305 | 26.02616979132801 | 28.63605184369398 | 118.68448505197199 | 9.685445514881827 | 37.8141396222474 | 27.527543612580263 | 5.103048613788249 | 34.03976397623994 | 146.24580760372163 |
| SE_39_10_9 | SE_39 | SE_39_10 | CTTTTCCCAGACACAACACA | 176.65562519519273 | 232.85844891969782 | 232.06881271911786 | 197.79889041409285 | 215.30068608406958 | 132.54544680986655 | 173.26185865510826 | 340.3272566002266 | 169.60518806460743 | 123.20217367574486 | 244.14727265716925 | 450.3442329384444 |
| SE_39_11_22 | SE_39 | SE_39_11 | TCAGCTGAGACATGGAAGTG | 5783.033227248763 | 5461.679928524242 | 5516.325054676563 | 5693.224641853002 | 5388.880719178116 | 4958.759068886771 | 5685.356517235633 | 5913.186083428937 | 5499.292825571277 | 5529.51767651198 | 5345.416729234369 | 6352.4071826600675 |
| SE_39_11_23 | SE_39 | SE_39_11 | TTCAGCTGAGACATGGAAGT | 184.24206308701082 | 178.8912547494674 | 131.34107273039436 | 156.15701874796804 | 163.33155496032865 | 213.11228702762855 | 217.3844437784588 | 252.4093819785014 | 171.38115862025776 | 212.14102094462575 | 123.24742129328254 | 105.6219721582434 |
| SE_39_11_24 | SE_39 | SE_39_11 | CTTCAGCTGAGACATGGAAG | 299.1224083059705 | 268.8365783665181 | 303.170746828805 | 327.92973937073293 | 337.2690550479513 | 239.9679004335492 | 316.39122015280634 | 127.62272122508497 | 238.86803973497067 | 252.23640291010486 | 86.86008738764674 | 314.5445544492743 |
| SE_39_11_25 | SE_39 | SE_39_11 | CACATTCTTCAGCTGAGACA | 1537.879338355696 | 1467.1081674425595 | 1580.0429802152705 | 1513.4217733657238 | 1652.4062508324157 | 1451.935744139453 | 1447.4360241684508 | 1888.8162741312576 | 1788.402349539892 | 1382.1971673917885 | 2132.767280856137 | 1532.0989368008932 |
| SE_39_11_26 | SE_39 | SE_39_11 | CATGTCTCAGCTGAAGAATG | 5282.328326388769 | 4771.09972164222 | 4722.35345711839 | 4995.723291445412 | 4876.613569529813 | 5184.866007562427 | 5541.15099512517 | 4816.576034383763 | 4728.521604419029 | 4745.835210823071 | 4062.469762819532 | 4169.166197718794 |
| SE_39_2_1 | SE_39 | SE_39_2 | GGGGTTTTCCCTGAGCAGGG | 9.75399157519469 | 14.990887269508443 | 13.825376076883616 | 15.615701874796805 | 23.33307928004695 | 52.844916701972934 | 7.533124289352532 | 5.67212094333711 | 9.767838056076867 | 4.374041668961356 | 28.170839152750297 | 0.0 |
| SE_39_2_2 | SE_39 | SE_39_2 | GCTGGGGTTTTCCCTGAGCA | 1215.9976163742713 | 1718.9550735703015 | 1600.781044330596 | 1480.8890611265638 | 1801.9500771272621 | 2090.40629511247 | 1494.7870911300954 | 1838.7125391317797 | 1511.3509428584389 | 1396.7773062883264 | 1932.050051892791 | 1847.8041722628955 |
| SE_39_2_26 | SE_39 | SE_39_2 | CATGATGGTGGTGTACCGCA | 8794.849070300546 | 7439.4776556147235 | 7642.470389928736 | 7835.178415679297 | 7689.310217288199 | 6620.341859614379 | 8287.51287890055 | 6672.304936345554 | 6735.368332303913 | 7547.408899792819 | 6218.712742969628 | 7161.401848531448 |
| SE_39_2_3 | SE_39 | SE_39_2 | TTTACAGAATAGACACATGG | 44.434850509220254 | 59.96354907803377 | 55.301504307534465 | 52.05233958265602 | 36.06021343279983 | 18.192512307236584 | 37.665621446762664 | 41.59555358447214 | 63.04695472558706 | 33.53431946203706 | 5.868924823489645 | 44.10587848366208 |
| SE_39_2_5 | SE_39 | SE_39_2 | GTGACTGAGTTTGTTCTGCT | 1649.508353049591 | 1775.9204451944336 | 1766.6855572531992 | 1752.8625354459414 | 1939.8273637820848 | 1222.3635650243248 | 1966.145439521011 | 1822.6415297923247 | 1596.5975295296553 | 1945.7195357429764 | 1939.0927616809788 | 1634.2388659209528 |
| SE_39_3_12 | SE_39 | SE_39_3 | GGAAGCATGCCAGTGGTGCG | 3346.702887133467 | 3392.9374853320774 | 3400.054988050735 | 3534.3538576623437 | 3270.8734772574903 | 3694.812618588763 | 3585.7671617318056 | 3301.174389022198 | 2734.994655701523 | 3442.370793472587 | 3245.515427389774 | 3353.207445771046 |
| SE_39_3_16 | SE_39 | SE_39_3 | GCTATTTTTCTTTTCTCACA | 19.50798315038938 | 16.989672238776237 | 15.800429802152705 | 40.34056317655841 | 9.545350614564661 | 39.8502650539468 | 33.36097899570407 | 56.7212094333711 | 23.087617223454416 | 90.39686115853469 | 5.868924823489645 | 123.03218734916264 |
| SE_39_3_22 | SE_39 | SE_39_3 | TAAGAAGGAACTCACCACAA | 1556.3035446643971 | 1653.9945620690983 | 1756.810288626854 | 1708.6180468006837 | 1613.1642538614276 | 1759.475833142738 | 1600.250831181031 | 1209.1071144213606 | 1618.7971614752846 | 1392.403264619365 | 1332.2459349321496 | 1212.9116583007071 |
| SE_39_3_23 | SE_39 | SE_39_3 | CTTCTTGACATTTAGTGCCA | 46.60240419259685 | 59.96354907803377 | 62.21419234597627 | 44.24448864525762 | 74.24161589105847 | 104.82352329407746 | 30.13249715741013 | 60.50262339559584 | 85.2465866712163 | 7.290069448268927 | 42.25625872912545 | 85.89039494186825 |
| SE_39_3_6 | SE_39 | SE_39_3 | TGGTCTACCGTGTTCCAGGA | 2.167553683376598 | 0.9993924846338962 | 0.987526862634544 | 0.0 | 0.0 | 0.0 | 0.0 | 0.0 | 0.0 | 0.0 | 0.0 | 0.0 |
| SE_39_4_3 | SE_39 | SE_39_4 | TGTCCATCAAGGCGTCCATT | 679.5280797385634 | 767.5334281988323 | 785.0838557944625 | 758.6628494172114 | 686.2046497359262 | 751.9571753657788 | 656.457973786435 | 417.84624282583377 | 659.7730614241011 | 665.583340626953 | 281.70839152750295 | 687.123159534946 |
| SE_39_4_8 | SE_39 | SE_39_4 | ATGGACACGTCCGGGACCCA | 5475.240604209286 | 5809.468513176838 | 5741.481179357239 | 5671.102397530373 | 5774.937121811619 | 5108.630717894006 | 5504.5615342911715 | 5823.377501826099 | 5393.622577510081 | 6039.8225378908055 | 5204.562533470617 | 5255.563625632155 |
| SE_39_5_13 | SE_39 | SE_39_5 | GTTGGAGGTTATAGAGGCCA | 360.89768828220355 | 440.7320857235482 | 492.7759044546375 | 478.88152416043533 | 445.44969534635084 | 437.4866054835464 | 382.03701753144986 | 374.35998226024924 | 380.9456841869978 | 1005.300576916285 | 347.440349550587 | 160.17397975645702 |
| SE_39_5_16 | SE_39 | SE_39_5 | CTCTATAACCTCCAACACCA | 48.769957875973454 | 45.972054293159225 | 44.43870881855448 | 61.16149900962082 | 90.15053358199957 | 53.71122681184134 | 76.40740350628997 | 106.82494443284891 | 18.647690834328564 | 32.80531251721017 | 140.85419576375148 | 292.49161520744326 |
| SE_39_5_20 | SE_39 | SE_39_5 | GTGTCTTTAAGTTCAAGACA | 32.51330525064897 | 33.97934447755247 | 35.55096705484358 | 41.641871666124814 | 28.63605184369398 | 1.7326202197368175 | 29.056336544645482 | 7.56282792444948 | 43.51127861343332 | 199.0188959377417 | 3.521354894093787 | 4.642724050911798 |
| SE_39_5_5 | SE_39 | SE_39_5 | CATAGAGGTGTTGAGCAAGG | 2814.5684578645123 | 2714.349988265662 | 2732.4868289097835 | 2750.9661469433704 | 2875.2717240094216 | 3408.06397222232 | 2964.8224881666038 | 2786.902090159633 | 2956.10298987999 | 3725.954495010248 | 2570.5890726884645 | 3067.6799166399705 |
| SE_39_5_9 | SE_39 | SE_39_5 | GTACTACTGCATGGAGAGTG | 2736.536525262955 | 2926.221195008048 | 2677.185324602249 | 2570.0842668936407 | 2899.665397802198 | 2287.058690052599 | 2544.0436885756267 | 2917.360871856387 | 2711.9070384780684 | 2325.5321539977876 | 2332.310724854785 | 2162.34872671217 |
| SE_39_6_12 | SE_39 | SE_39_6 | CTATCCTGCAGTAACCCTAG | 23.843090517142578 | 9.993924846338961 | 30.613332741670867 | 10.410467916531204 | 10.605945127294067 | 99.625662634867 | 33.36097899570407 | 10.398888396118036 | 24.863587779104755 | 5.832055558615141 | 24.64948425865651 | 13.928172152735392 |
| SE_39_6_17 | SE_39 | SE_39_6 | AACCAGATATGCTGTCCAAG | 101.8750231187001 | 160.9021900260573 | 118.50322351614528 | 128.82954046707366 | 82.72637199289373 | 58.90908747105179 | 123.75847046793446 | 106.82494443284891 | 78.14270444861494 | 105.70600699989943 | 71.60088284657367 | 47.58792152184593 |
| SE_39_6_19 | SE_39 | SE_39_6 | GCGCCCAGCACGGAAGCCTG | 4623.392006642283 | 5492.661095547894 | 5502.499678599679 | 5038.666471601103 | 5823.7244693971725 | 5364.192200305187 | 4272.35763267565 | 5822.432148335543 | 5033.100554713063 | 5411.418551450024 | 5339.5478044108795 | 6368.656716838259 |
| SE_39_6_3 | SE_39 | SE_39_6 | TCAACCTCATGAGGAAGAGG | 447.59983561726744 | 503.6938122554837 | 490.8008507293684 | 394.2964723386193 | 513.3277441610329 | 375.112277573021 | 388.4939812080377 | 354.5075589585694 | 553.2148280850807 | 355.7553890755236 | 380.306328562129 | 555.9662050966878 |
| SE_39_6_31 | SE_39 | SE_39_6 | GAAGGGGGCATACTCTAGAA | 1140.1332374560905 | 1636.0054973456881 | 1624.481689033825 | 1417.12494513781 | 1473.165778181146 | 1502.1817305118207 | 1118.1308766624688 | 1373.5986217781367 | 1625.0130584200608 | 1536.7466396950897 | 1392.1089681317437 | 1608.7038836409379 |
| SE_39_7_10 | SE_39 | SE_39_7 | GACCCTCGAGACTTATGCTC | 266.60910305532155 | 338.7940522908908 | 267.61977977396145 | 251.15253848631528 | 292.72408551331625 | 381.17644834209983 | 190.4804284593426 | 347.89008452467607 | 386.27359585394885 | 444.69423634440454 | 321.61708032723254 | 285.52752913107554 |
| SE_39_7_11 | SE_39 | SE_39_7 | GGCCTGAGCATAAGTCTCGA | 1110.8712627305065 | 1127.3147226670349 | 1216.6330947657582 | 1112.6187585792725 | 1252.5621195334293 | 1128.8020731585366 | 1071.8559703135888 | 1103.227523479068 | 1445.6400322993763 | 891.5754935232898 | 1211.3460835682627 | 984.2574987933011 |
| SE_39_7_49 | SE_39 | SE_39_7 | GATGCTTGTCCATTAGGTGA | 2683.4314600202283 | 2883.2473181687906 | 2942.8300506509413 | 2771.7870827764327 | 2598.4565561870463 | 3321.432961235479 | 2720.5340290690287 | 2367.165140352687 | 2702.1392004219915 | 2792.0965986869987 | 2275.9690465492845 | 2884.2923166289543 |
| SE_39_7_50 | SE_39 | SE_39_7 | CCATTAGGGCCATCACCTAA | 22.759313675454276 | 61.96233404730156 | 57.27655803280356 | 39.03925468699201 | 47.72675307282331 | 19.05882241710499 | 23.675533480822246 | 24.57919075446081 | 73.70277805948909 | 31.347298627556384 | 16.432989505771005 | 20.89225822910309 |
| SE_39_7_9 | SE_39 | SE_39_7 | TTAAGCGCTTGCTTAGTACA | 515.8777766436302 | 583.6452110261954 | 582.640848954381 | 455.45797134824016 | 580.1451984629855 | 427.0908841651255 | 439.07353000797616 | 463.2232103725307 | 713.9401633714365 | 306.1829168272949 | 523.5080942552763 | 561.7696101603275 |
| SE_39_8_19 | SE_39 | SE_39_8 | AAACAAGCCTTGCCAGGTCC | 135.47210521103736 | 111.93195827899638 | 94.80257881291622 | 113.21383859227684 | 144.2408537311993 | 117.81817494210358 | 130.21543414452233 | 74.68292575393862 | 149.18152667462851 | 35.72134029651774 | 119.72606639918877 | 77.76562785277261 |
| SE_39_8_20 | SE_39 | SE_39_8 | CATAGTAAACAAGCCTTGCC | 5356.025151623573 | 5025.944805223864 | 4829.006358282921 | 4865.592442488772 | 4464.042304078073 | 3989.358055944022 | 5132.209962274604 | 4923.400978816611 | 4857.279469703679 | 6012.120273987383 | 4610.627341333466 | 4330.50085848798 |
| SE_39_8_27 | SE_39 | SE_39_8 | CATGATCCTGGGACTGTCTG | 4918.1793075815 | 3978.5814813275406 | 3853.3298179999906 | 4649.5752332207485 | 4125.712654517392 | 3903.5933550670497 | 4723.268929424038 | 4057.4571814671463 | 4038.557043548872 | 4461.522502340583 | 3212.6494483782317 | 4103.007379993302 |
| SE_39_8_29 | SE_39 | SE_39_8 | CTGGAAAAGGAAGAGCGGGA | 324.04927566480137 | 368.7758268299077 | 396.9857987790867 | 348.7506752037953 | 362.7233233534571 | 701.711188993411 | 465.97754532709234 | 328.9830147135524 | 330.3305233509632 | 1076.7432575093205 | 309.8792306802533 | 952.9191114496465 |
| SE_39_8_31 | SE_39 | SE_39_8 | CGACCTGGAAAAGGAAGAGC | 2379.9739443475046 | 2529.462378608391 | 2426.353501493075 | 2482.896598092692 | 2391.640626204812 | 2417.8715166427287 | 2364.3248662439305 | 2595.940685067284 | 2399.3362206836086 | 2997.6765571281826 | 2260.709842008211 | 2216.9007343103835 |
| SE_39_9_12 | SE_39 | SE_39_9 | ACATCAAGTTCCCCTTCCCG | 3765.0407480251506 | 4076.5219448216626 | 4469.546580283946 | 4394.518769265735 | 4825.7050329188005 | 5241.176164703873 | 4556.464034445517 | 5044.406225607803 | 4945.190012208371 | 6550.127399269631 | 7210.561038139378 | 6211.964780119985 |
| SE_39_9_2 | SE_39 | SE_39_9 | GCTGGAACAAGGGGGCTACA | 790.0733175907699 | 1043.3657539577875 | 1066.5290116453075 | 827.6321993642307 | 1251.5015250207 | 1160.8555472236676 | 778.0641230288401 | 1210.997821402473 | 1038.0547897776235 | 966.6632088404597 | 1133.8762758981995 | 1003.9890760096763 |
| SE_39_9_27 | SE_39 | SE_39_9 | TCACCCTTCCTTCTGACCCA | 1321.1239700180363 | 1195.2734116221397 | 1209.7204067273165 | 1203.7103528489204 | 1081.806402983995 | 912.2245456914344 | 1178.395870977289 | 1228.9595377230405 | 1443.864061743726 | 893.0335074129435 | 1510.6612495662346 | 1552.9911950299963 |
| SE_39_9_3 | SE_39 | SE_39_9 | AAGAAGCTCTCAAACTAAGG | 81.28326312662242 | 143.91251778728105 | 120.47827724141437 | 87.18766880094883 | 126.21074701479941 | 82.29946043749882 | 135.59623720834557 | 206.08706094124832 | 152.7334677859292 | 207.7669792756644 | 313.4005855743471 | 51.069964560029774 |
| SE_39_9_6 | SE_39 | SE_39_9 | CATGAAACATACCATCCACT | 6864.642515253685 | 5732.515291860029 | 5627.915590154266 | 6126.560368878613 | 5974.328890204748 | 5759.229610405181 | 6704.480617523754 | 6176.939707294113 | 6452.101028677684 | 5585.651211263652 | 5739.808477372873 | 6272.320192781839 |
| SE_390_1_10 | SE_390 | SE_390_1 | ATATTACTGAACCTTCACCA | 81.28326312662242 | 133.9185929409421 | 106.65290116453076 | 80.68112635311682 | 153.786204345764 | 82.29946043749882 | 59.18883370205561 | 168.27292131900094 | 60.382998892111544 | 71.44268059303548 | 144.37555065784528 | 42.945197470934126 |
| SE_390_1_11 | SE_390 | SE_390_1 | ACAGGATGAGATAGGAAGCT | 1613.743717273877 | 1732.946568355176 | 1638.3070651107087 | 1630.5395374266998 | 1582.4070129922748 | 1568.0212988618198 | 1428.065133138687 | 1292.2982215903048 | 1444.7520470215513 | 1392.403264619365 | 1160.8733300862518 | 2504.749625466915 |
| SE_390_1_12 | SE_390 | SE_390_1 | TTAGGCAATTCCCTGAACTG | 1383.9830268359576 | 1558.0528835442442 | 1489.1905088528924 | 1454.8628913352356 | 1491.195884897546 | 1205.903672936825 | 1289.2404140920478 | 1539.9808361160253 | 1745.7790562042837 | 2009.8721468877432 | 1547.0485834718704 | 1107.2896861424638 |

| | | | | | | | | | | | | | | | |
|---|---|---|---|---|---|---|---|---|---|---|---|---|---|---|---|
| SE_390_1_19 | SE_390 | SE_390_1 | GGAATCAGACTGCTTGGGAT | 549.4748587359676 | 629.6172653193546 | 641.8924607124536 | 598.6019052005443 | 660.7503814304204 | 503.32617383354545 | 588.6598551822622 | 610.6983548992955 | 592.2861803093882 | 347.7363126824278 | 285.22974642159676 | 771.8528734640864 |
| SE_390_1_6 | SE_390 | SE_390_1 | CCAATCCAGACCTCAAAAGA | 228.67691359623106 | 282.82807315139263 | 314.033542317785 | 294.09571862420065 | 288.4817074623986 | 186.25667362170788 | 205.54667703804768 | 195.6881725451303 | 222.88430473411762 | 244.94633346183593 | 184.28423945757487 | 154.37057469281729 |
| SE_391_1_12 | SE_391 | SE_391_1 | AATCGTGGACAGGATAGCCT | 500.7049008599941 | 645.6075450734969 | 557.9526777885174 | 532.2351722326578 | 505.903582571927 | 337.86094284867943 | 500.41468493556107 | 473.6220987686487 | 657.1091055906256 | 349.9233335169085 | 379.1325435974311 | 587.3045924403424 |
| SE_391_1_15 | SE_391 | SE_391_1 | AGAGCCATCTGAGTCTACAG | 642.6796671211613 | 810.5073050380898 | 891.7367569589933 | 754.7589239485122 | 769.9916162415493 | 835.9892560230144 | 760.8455532246057 | 1012.4735883856741 | 469.7442119695148 | 770.5603406820255 | 1003.5861448167293 | 633.7318329494603 |
| SE_391_1_18 | SE_391 | SE_391_1 | GTGGAGAAACACGACAGACG | 520.2128840103835 | 639.6111901656935 | 642.8799875750882 | 624.6280749918722 | 610.9024393321383 | 829.0587751440671 | 598.345300697144 | 572.8842152770482 | 618.0377533663182 | 686.7245420269329 | 562.242998090308 | 927.3841291696316 |
| SE_391_1_36 | SE_391 | SE_391_1 | CATCGGGGAAGTGAGCCTGG | 7316.577458237706 | 7275.577288134764 | 7204.995989781633 | 7141.5809907404055 | 7314.920354294718 | 6827.389975872929 | 7358.786270084659 | 7213.047132943691 | 7322.3266009463505 | 7089.592538441531 | 6647.144255084372 | 7607.1033574189805 |
| SE_391_1_38 | SE_391 | SE_391_1 | GAGAGAACCTGAGAGCTGAA | 1109.787485888818 | 1366.169526494536 | 1344.0240600456145 | 1125.6318434749364 | 1283.319360402582 | 1657.2512401782658 | 1231.1277410027567 | 1195.8721655553574 | 1304.922431240138 | 1166.7422549097414 | 1889.5886887211018 |
| SE_391_2_13 | SE_391 | SE_391_2 | TGACGCTGTGTTCCAGCCGG | 6266.397698641745 | 6550.018344290555 | 6449.537939866207 | 6176.0100914821137 | 6402.809073347428 | 6171.593222702544 | 5858.618375890741 | 5356.372877491344 | 6191.0333569970835 | 6643.440288207473 | 4805.4756454733215 | 6525.348653556532 |
| SE_391_2_14 | SE_391 | SE_391_2 | GGATGACGCTGTGTTCCAGC | 267.69287989700985 | 253.84569109700962 | 294.28300506509413 | 284.9865592150417 | 256.66387208051646 | 398.502650539468 | 343.2952354719225 | 348.8354380152323 | 282.379318348404 | 137.7823125722827 | 322.79086529193046 | 157.85261773100112 |
| SE_391_2_2 | SE_391 | SE_391_2 | GGAGGTCACCCTCCTTACCG | 1175.8978732318044 | 1219.2588312533533 | 1183.0571814361838 | 1268.7757773272403 | 982.1105187874307 | 1013.5828285460382 | 1288.164253479283 | 1187.3639841385684 | 1062.030392278903 | 920.0067643715386 | 1549.3961534012662 | 1270.9457089371047 |
| SE_391_2_5 | SE_391 | SE_391_2 | CCCCATTTCCTCGGTAAGGA | 28.17819788389577 | 58.96415659339988 | 56.28903117016901 | 29.93009526002721 | 43.48437502190568 | 15.593581977631358 | 16.142409191469714 | 60.50262339559584 | 63.04695472558706 | 32.076305572383276 | 23.47569929395858 | 58.034050636397474 |
| SE_391_2_8 | SE_391 | SE_391_2 | GAGAGGGTCCCCCATTTCCT | 902.786109126353 | 990.3979522721911 | 1020.115249101484 | 969.4748247269683 | 917.4142535109369 | 1097.6149092032738 | 899.6702722712453 | 1241.249133100271 | 1117.9734647818887 | 670.6863892407413 | 1369.807053802483 | 1066.6658506969854 |
| SE_391_3_24 | SE_391 | SE_391_3 | TGGACATCACCTGTGGACGA | 333.80326723999605 | 358.78190198356873 | 341.6842944715522 | 356.55852614119374 | 377.5716465316688 | 220.90907801644423 | 344.3713960846872 | 273.20715877073746 | 259.29170112494955 | 243.48831957218215 | 316.9219404684408 | 253.02846077469297 |
| SE_391_3_26 | SE_391 | SE_391_3 | TCTTCAATGGACATCACCTG | 509.3751155935005 | 525.6804469174294 | 565.8528922895937 | 439.84226947344337 | 538.7820124665386 | 449.61494702170415 | 394.95094488462564 | 564.3760338620425 | 435.1127861343332 | 331.6981598962362 | 422.56258729125443 | 256.51050381287683 |
| SE_391_3_30 | SE_391 | SE_391_3 | GATCCTCTCCACCTCTTCAA | 70.44549470973944 | 73.95504386290831 | 132.3285995930289 | 101.50206218617923 | 112.42301834931712 | 302.34222834407467 | 91.47365208499504 | 117.22383282896693 | 183.81295250981015 | 69.98466670338169 | 83.33873249355297 | 98.6578860818757 |
| SE_391_3_35 | SE_391 | SE_391_3 | GGACGAGACGCAGGAGGCCG | 5858.897606166944 | 5580.607634195676 | 5682.229567599166 | 5676.307631488638 | 5875.693600520914 | 6254.758993249911 | 6427.907340043239 | 6183.557181728006 | 5147.650655552509 | 5339.975870856989 | 6437.0367464034425 | 5638.588359832378 |
| SE_391_3_37 | SE_391 | SE_391_3 | GGAGGCCGTGGAGTACCAGC | 2974.9674304343807 | 3265.015247298939 | 3115.6472516119866 | 3121.8390664697945 | 3182.8441327009496 | 2805.1121357539073 | 2971.2794518431915 | 2961.792485912528 | 3329.0568065665616 | 3107.7566057970434 | 2815.9101303103316 | 2634.7458988924454 |
| SE_392_1_23 | SE_392 | SE_392_1 | AATGAGAAGAAAGCATAAAA | 84.53459365168732 | 108.93378082509469 | 77.02709528549444 | 104.10467916531204 | 85.90815553108195 | 50.2459863723677 | 78.55972473181926 | 124.78666075341643 | 71.92680750383876 | 85.29381254474644 | 124.42120625798047 | 58.034050636397474 |
| SE_392_1_4 | SE_392 | SE_392_1 | AATCCATAGAAATATATCAG | 3.2513305205648967 | 4.996962423169481 | 0.987526862634544 | 3.9039254686992013 | 6.363567076376441 | 0.0 | 0.0 | 2.836060471668555 | 0.0 | 0.0 | 18.780559435166865 | 0.0 |
| SE_392_1_7 | SE_392 | SE_392_1 | CACAAAATGATTCTAGGCAG | 844.2621596751849 | 1136.30925502874 | 1077.3918071342875 | 879.6845389468867 | 974.6863571983248 | 1297.7325445828762 | 754.3885895480179 | 1033.2713651779102 | 1187.236316452252 | 1234.937764536756 | 1343.9837845791287 | 1118.8964962697432 |
| SE_392_1_8 | SE_392 | SE_392_1 | GTTTAATCTCTGCTACAGCT | 5.4188842084414945 | 13.991494784874547 | 14.81290293951816 | 6.506542447832002 | 39.24199697098805 | 0.0 | 8.60928490211718 | 0.0 | 5.327911666951019 | 20.41219455152996 | 1.173784964697929 | 0.0 |
| SE_392_1_9 | SE_392 | SE_392_1 | ACAACTTGCCTGGAAACAGA | 943.9696291105083 | 964.4137476717099 | 974.6890134202949 | 847.1518267077267 | 1033.019055398442 | 822.1282942651198 | 910.4318783988917 | 1009.6375279140055 | 1112.6455531149377 | 1115.3806255851457 | 765.3077969830497 | 857.7432684059546 |
| SE_392_2_13 | SE_392 | SE_392_2 | CTGTAGCCAGGTAGACAGCA | 136.55588205272565 | 123.92466809460312 | 128.37849214249073 | 134.03477442533924 | 158.0285823966816 | 150.73795911710312 | 148.51016456152135 | 195.6881725451303 | 248.63587779104753 | 47.38545141374802 | 72.7746678112716 | 38.30247342002233 |
| SE_392_2_29 | SE_392 | SE_392_2 | GTTGAGCTCCAGATTCCACA | 816.0839617912891 | 892.4574887780693 | 792.9840706955389 | 739.1432220737155 | 851.6573937217136 | 701.711188993411 | 691.9712740076683 | 1008.6921744234494 | 762.7793536518209 | 618.9268961580318 | 997.7172199392397 | 963.3652405641981 |
| SE_392_2_3 | SE_392 | SE_392_2 | GGGTTGCCGTAAGAATTAAA | 39.01596630077876 | 70.95686640900662 | 72.08946097232172 | 36.43663770785921 | 30.757240869152795 | 113.48662439276154 | 43.0464245105859 | 9.45353490556185 | 58.607028336461205 | 87.48083337922712 | 3.521354894093787 | 113.74673924733905 |
| SE_392_2_4 | SE_392 | SE_392_2 | AGAGAAAATGACCTACCCCT | 2774.468714722045 | 3266.014639783573 | 3299.3272480620117 | 3016.4330788149164 | 3148.9051082936085 | 3641.101391776922 | 3082.1239949579503 | 3167.879546853776 | 2941.8952254347873 | 2955.3941543282226 | 3412.1928923768796 | 3846.4968761804244 |
| SE_392_2_5 | SE_392 | SE_392_2 | GCATTTTCTGATCTGCTTAT | 97.53991575194691 | 146.91069524118274 | 143.19139508200888 | 106.70729614444484 | 130.45312506571702 | 143.80747823815585 | 106.5399006637001 | 103.98888396118035 | 187.36489362111084 | 308.36993766177557 | 110.33578668160533 | 139.28172152735394 |
| SE_392_3_13 | SE_392 | SE_392_3 | GTTTAAGGTGAAGATTGTCA | 70.44549470973944 | 94.94228604022014 | 98.75268626345441 | 67.66804145745283 | 94.3929116329172 | 36.38502461447317 | 59.18883370205561 | 179.61716320567715 | 130.5338358402996 | 16.038152786191638 | 257.05890726884644 | 75.44426582731671 |
| SE_392_3_14 | SE_392 | SE_392_3 | TGAGCATGTACAAAGGAAGG | 2271.5962601786746 | 2370.558973551602 | 2339.451137581235 | 2290.302941636865 | 2341.79268410653 | 1845.2405340197106 | 2050.0859673166533 | 2216.8539353542537 | 2057.4618887209185 | 2250.4444386806176 | 1811.1502005289044 | 1708.5224507355415 |
| SE_392_3_5 | SE_392 | SE_392_3 | TTCTAGGCAGAGGCTGATAA | 7.586437891818092 | 11.992709815606755 | 7.900214901076352 | 14.314393385230405 | 12.727134152752882 | 38.11764483420998 | 2.152321225529295 | 0.945353490556185 | 4.439926389125849 | 0.0 | 0.0 | 27.856344305470785 |
| SE_392_3_9 | SE_392 | SE_392_3 | CATGAAGACAGGTTTCTAGA | 838.8432754667433 | 1191.2758416836043 | 1245.27137378216 | 931.7368785295428 | 1164.5327749288855 | 1409.486548755901 | 862.0046508244826 | 1423.7023567776146 | 1367.4973287507614 | 1009.6746185852463 | 1162.0471150509497 | 1108.4503671551918 |
| SE_393_1_12 | SE_393 | SE_393_1 | ACTGCATCTGCAGAGCCCAT | 244.93356622155557 | 58.96415659339988 | 103.69032057662713 | 196.49758192452646 | 27.575457330964575 | 722.5026316302528 | 326.0766656676882 | 117.22383282896693 | 357.8580669635434 | 16.76715973101853 | 84.5125174582509 | 186.86964304919985 |
| SE_393_1_14 | SE_393 | SE_393_1 | TGGTGCTTCTCTTTGACAGC | 0.0 | 0.9993924846338962 | 5.925161175807264 | 0.0 | 0.0 | 0.0 | 0.0 | 0.0 | 0.0 | 0.0 | 0.0 | 0.0 |
| SE_393_1_26 | SE_393 | SE_393_1 | CGCCGTAAGACACGTCCGCG | 3589.468899671646 | 3194.0583808899323 | 3180.8240245458665 | 3513.5329218292813 | 3422.5384925777958 | 3246.0639816769276 | 3703.068668523152 | 3865.5504228842406 | 3120.3802662776466 | 3436.538737913972 | 3643.4285304223718 | 2863.4000583998513 |
| SE_393_1_30 | SE_393 | SE_393_1 | TTACCCCGGCTATAAAATGG | 35.76463577571386 | 53.96719417023039 | 49.376343131727204 | 33.834020728726415 | 49.84794209828212 | 19.9251325269734 | 36.58946083399801 | 41.59555358447214 | 56.831057780810866 | 21.141201399979888 | 99.77172199932397 | 10.446129114551544 |
| SE_393_1_8 | SE_393 | SE_393_1 | AGTCAAGTGGCCAGAGCACG | 1134.714353247649 | 1377.1628438255088 | 1286.7475020128109 | 1184.1907255054243 | 1255.7439030716175 | 1036.9732015124853 | 1044.9515949944728 | 1137.2602491390905 | 1458.0718261889288 | 1240.0408131505444 | 1471.9263457131203 | 1000.5070329714924 |
| SE_393_2_12 | SE_393 | SE_393_2 | TTTGGGTGATGAGTAAACAA | 80.19948628493412 | 99.93924846338962 | 93.81505195028168 | 101.50206218617923 | 138.9378811675523 | 171.52940175394494 | 77.48356411905462 | 68.06545132004533 | 79.91867500426528 | 45.92743752409424 | 140.85419576375148 | 38.30247342002233 |
| SE_393_2_34 | SE_393 | SE_393_2 | GCGCGGCCACACTTATCTGG | 3227.487434547754 | 3143.0893641736034 | 3019.8571459364357 | 3362.5811370395786 | 3205.116617468267 | 3735.5291937525785 | 3406.0483394001094 | 3535.622054680132 | 3433.8390693499314 | 3845.511633961859 | 4054.253268066647 | 4087.918526827838 |
| SE_393_2_39 | SE_393 | SE_393_2 | CCAGGGTCCTGAGGAGAAAT | 466.0240419259685 | 652.6032924659343 | 658.6804173772408 | 501.00376848306416 | 613.0236283575971 | 579.5614635019655 | 560.6796792503814 | 632.4414851820877 | 525.6872844725006 | 461.46139607542307 | 699.5758389599657 | 580.3405063639747 |
| SE_393_2_41 | SE_393 | SE_393_2 | CATTTCTCCTCAGGACCCTG | 48.769957875973454 | 60.962941562667666 | 73.07698783495626 | 74.17458390528482 | 90.15053358199957 | 49.379676262499295 | 43.0464245105859 | 86.97252113116902 | 36.40739639083196 | 23.328222234460565 | 134.98527094026184 | 91.69380000550801 |
| SE_393_2_7 | SE_393 | SE_393_2 | AAAATGGCCATAGAACAAAG | 60.69150313454474 | 85.94775367851507 | 76.03956842285989 | 78.07850937398402 | 125.15015250207 | 254.69517230131217 | 65.6457973786435 | 116.27847933841076 | 78.14270444861494 | 65.61062503442034 | 125.5949912226784 | 78.92630886550056 |
| SE_393_3_11 | SE_393 | SE_393_3 | TGATTCTTCACCTCTTCCCA | 27.094421042207472 | 23.98541963121351 | 23.700644703229056 | 22.12224432262881 | 31.8178353818822 | 4.331550549342044 | 19.370891029763655 | 26.46989773557318 | 38.1833669464823 | 43.01140974478667 | 53.994108376104734 | 15.088853165463343 |
| SE_393_3_14 | SE_393 | SE_393_3 | AGAAGCCGGCTAGGTCCCCG | 57.440172609479845 | 109.93317330972857 | 105.66534730189621 | 84.58505182181602 | 154.84679885849337 | 212.24597679170615 | 58.112673409205569 | 101.15282348951779 | 97.67838506076867 | 84.56480555991955 | 117.3784964697929 | 373.7392860983997 |
| SE_393_3_24 | SE_393 | SE_393_3 | ATAACCACAGGACAGACAGG | 1014.4151238202478 | 1188.2776642297026 | 1059.6163236068658 | 1018.9245473304916 | 1239.8349853806765 | 1103.6790799723526 | 1082.6175764412353 | 1195.8721655553574 | 1238.7394625661118 | 1162.0370700540668 | 1210.1722986035647 | 855.4219063804987 |
| SE_393_3_3 | SE_393 | SE_393_3 | CCGCAGACGTGACAGCCGCG | 3898.345299552811 | 3244.028005121627 | 3267.7263884577064 | 3441.9609549031293 | 3134.056785115397 | 3403.732421672978 | 3539.4922553829256 | 3165.9888398726634 | 2802.4815368616236 | 3591.8172171621 | 2984.9351652268338 | 3473.9182710947525 |
| SE_393_3_35 | SE_393 | SE_393_3 | TGGCATCCACAGAATTGTAG | 21.675536833765978 | 34.97873696218637 | 28.63827901640178 | 45.545797134824014 | 26.51486281823517 | 58.04277736118338 | 50.57954879993843 | 23.633837263904624 | 22.199631945629246 | 37.17935418617152 | 30.518409082146157 | 59.19473164912542 |
| SE_394_1_11 | SE_394 | SE_394_1 | TTTAAGGCCCATCTGCTTGT | 172.32051782843953 | 175.89307729556572 | 187.63010390056337 | 205.60674135149128 | 197.27057936766965 | 252.09624197170695 | 159.27177068916782 | 158.81938641343908 | 159.83735000853056 | 161.83954175157018 | 107.98821675220947 | 290.17025318198733 |
| SE_394_1_25 | SE_394 | SE_394_1 | TCTGAAAACACGAATCCAAC | 5538.099661027208 | 4908.016490247854 | 5012.686354732946 | 4982.710206549747 | 5077.06593243567 | 5470.748343819001 | 5147.276210853309 | 5828.104269278881 | 5260.424785836306 | 4185.957877196018 | 6086.075041958762 | 5065.211939544772 |
| SE_394_1_27 | SE_394 | SE_394_1 | ACACGAATCCAACAGGACCA | 2028.8302476404956 | 1763.9277353788268 | 1914.8145866483808 | 2032.6438607027176 | 2022.5537357749786 | 1943.9998865447092 | 1817.6352749594896 | 2038.182125639135 | 1806.1620550963953 | 1585.5901049984916 | 2045.9071934684903 | 1861.732344415631 |
| SE_394_1_34 | SE_394 | SE_394_1 | TTAATTACTTATTTAATGGA | 0.0 | 5.9963549708033775 | 1.975053725269088 | 3.9039254686992013 | 8.484756101835254 | 0.8663101098684087 | 17.21856980423436 | 1.890706981181237 | 0.8879852778251698 | 5.103048613788249 | 0.0 | 0.0 |
| SE_394_1_8 | SE_394 | SE_394_1 | AATACTGGTATGAGACAGG | 86.70214733506391 | 74.95443634754221 | 69.12688038441809 | 92.39290275921444 | 92.27172260745839 | 42.44919538355203 | 108.6922218892294 | 34.97807915057884 | 83.47061611556596 | 30.61829168272949 | 15.259204541073078 | 110.26469620915519 |
| SE_395_1_12 | SE_395 | SE_395_1 | GTTGTGAGAGAGAAGGCTGA | 340.30592829012585 | 373.7727892530772 | 352.5470899605322 | 337.0388987976977 | 340.4508385861396 | 278.0855426577592 | 290.56336544645484 | 498.2012895231095 | 315.2347736279353 | 320.7630557238328 | 240.62591776307545 | 220.52939241831038 |
| SE_395_1_14 | SE_395 | SE_395_1 | GCAGAGGGAGAAGCAAGCAT | 22059.193835723636 | 19667.044705110442 | 19865.090368756486 | 21808.628976643304 | 19740.845665432447 | 20710.00948651418 | 23732.56999329877 | 21752.583817697818 | 19781.648034111306 | 21607.7658446691 | 21015.446007951723 | 21346.084505079718 |
| SE_395_1_17 | SE_395 | SE_395_1 | ATTTATGATTGTGTGCCCGT | 24.926867358830876 | 37.97691441608806 | 35.55096705484358 | 32.53271223916001 | 34.999618920070425 | 5.197860659210452 | 22.599372868075596 | 53.88514896170255 | 20.423661389978903 | 15.309145841364746 | 24.64948425865651 | 19.73157721637514 |
| SE_395_1_18 | SE_395 | SE_395_1 | GGATAATCCCGGGCAAACCA | 93.2048083851937 | 179.89064723410132 | 188.61763076319792 | 121.02168952967524 | 181.36166167672854 | 29.454543735525895 | 89.32133085946575 | 301.567763487423 | 159.83735000853056 | 60.50757642603209 | 204.23858385743966 | 53.39132658548567 |
| SE_395_1_23 | SE_395 | SE_395_1 | AGAGCCTGAACTTTTCCAGA | 163.65030309493312 | 333.7970898677213 | 329.8339721199377 | 199.10019890365928 | 343.6326221243278 | 362.98393603486323 | 199.0897133614598 | 447.1522010330755 | 203.34862862196388 | 358.6714168548312 | 503.5537498554116 | 192.6730481128396 |
| SE_395_2_1 | SE_395 | SE_395_2 | AGAACATCAGGCTTTGTAGG | 396.6623240579174 | 530.6774093405988 | 531.2894520973847 | 439.84226947344337 | 565.2968752847738 | 467.7272340493327 | 417.5503177526832 | 467.0046243347554 | 598.5020772541644 | 514.6789030477862 | 394.39174813850417 | 695.2479266240417 |
| SE_395_2_11 | SE_395 | SE_395_2 | CTTTCCTCATCTTCTTTATA | 8.670214733506391 | 7.9951398770711695 | 24.688171565863602 | 0.0 | 4.242378050917627 | 0.0 | 1.0761606127646475 | 3.78141396222474 | 9.767838056076867 | 0.7290069448268927 | 0.0 | 8.124767089095647 |
| SE_395_2_18 | SE_395 | SE_395_2 | GAGGTCATATAGCTGGTAAG | 1347.1346142185555 | 1504.0856893740138 | 1491.1655625781616 | 1392.4000838360485 | 1528.316692843075 | 1231.026666123009 | 1307.5351445090466 | 1715.8165853594758 | 1353.2895634055587 | 1432.498646584844 | 1591.6524121303917 | 1590.1329874372907 |
| SE_395_2_19 | SE_395 | SE_395_2 | GCAGTAAGTGCTGTGATGGT | 8583.512586171328 | 7536.418726624212 | 7367.937922116333 | 8294.540314266236 | 7279.920735374648 | 7173.914019820293 | 8504.879232267901 | 7706.52165501402 | 6778.879610917346 | 7623.954628999643 | 7674.20609919506 | 6517.223886467436 |
| SE_395_2_32 | SE_395 | SE_395_2 | ACAAGACTAAGCCAAGGGTT | 1376.3965889441397 | 1567.0474159059493 | 1506.9659923803142 | 1531.6400922196533 | 1930.2820131675203 | 1332.3849489776126 | 1426.9889725259225 | 1682.7292131900092 | 1465.1757084115302 | 2046.3224941290878 | 1517.7039593544223 | 1456.6546709735765 |
| SE_395_3_1 | SE_395 | SE_395_3 | GGCTTAGTCTGACCAGTAGA | 390.15966300778763 | 410.7503111845313 | 377.2446731260336 | 472.37498171260336 | 362.72332335354571 | 437.48660548354644 | 449.83513613562263 | 365.8518008452436 | 439.55271252345904 | 425.0110488340784 | 338.0500698330036 | 434.0946987602531 |
| SE_395_3_14 | SE_395 | SE_395_3 | GAACCGCTAGCACTAGGTGG | 828.0055070498604 | 978.4052424565843 | 974.6890134202949 | 938.2434209773747 | 944.9897108419015 | 615.9464881164386 | 856.6238477606594 | 1072.0308582907137 | 1038.0547897776235 | 1044.6669519369373 | 787.6097113123104 | 766.0494684004466 |
| SE_395_3_16 | SE_395 | SE_395_3 | GGACTGGTAGCTTTGTCGTG | 1283.191780558946 | 1297.2114450547972 | 1261.0718035843126 | 1294.8019471185685 | 1369.227515933664 | 1305.529335571692 | 1335.5153204409276 | 1292.2982215903048 | 1363.945386739461 | 1446.349778536555 | 1477.7952705546927 | 1527.4562127499814 |
| SE_395_3_25 | SE_395 | SE_395_3 | TTTAAATTCTAGCAGAGACA | 0.0 | 2.9981774539016888 | 4.93763431317272 | 2.602616979132801 | 7.4241615891058474 | 0.0 | 0.0 | 15.12565584889896 | 14.207764445202717 | 0.0 | 0.0 | 1.1606810127279494 |
| SE_395_3_26 | SE_395 | SE_395_3 | CTTCTCTCCAGAGTCACACA | 44.434850509220254 | 48.970231747060915 | 48.38881626909266 | 40.34056317655841 | 71.05983235287025 | 8.663101098684088 | 71.02660044246673 | 106.82494443284891 | 10.655823333902038 | 46.65644446892113 | 29.344624117448227 | 114.907420260067 |
| SE_396_1_1 | SE_396 | SE_396_1 | CCTGTGCCTGGAGGACACAG | 280.6982019972694 | 205.8748518345826 | 304.15827369142955 | 305.8074950481041 | 209.99771352402254 | 129.9465164826132 | 290.56336544645484 | 317.63877282687815 | 306.3549208496836 | 87.48083337922712 | 68.07952795247988 | 358.6504329329364 |
| SE_396_1_10 | SE_396 | SE_396_1 | GCTGGAACTCAAATGCTGTG | 3103.936874595288 | 3104.1130572728816 | 3015.9070384858974 | 3146.5639277715563 | 3094.814788144409 | 2786.919623446671 | 3027.2398037069534 | 2895.6177415735947 | 3220.722602671891 | 3669.8209602585775 | 3158.655340002127 | 3424.008987547451 |
| SE_396_1_12 | SE_396 | SE_396_1 | GTACGGACTGTACCACCAAG | 161.48274941155654 | 176.89246978019963 | 140.22881449410525 | 174.37533760189766 | 177.11928362581094 | 53.71122681184134 | 161.42409191469713 | 116.27847933841076 | 96.7903952829435 | 248.5913681859704 | 133.8114859755639 | 53.39132658548567 |
| SE_396_1_28 | SE_396 | SE_396_1 | CTCTTGCTCAACCAAAAGCA | 35.76463577571386 | 55.96597913949819 | 52.338923719630834 | 36.43663770785921 | 42.42378050917627 | 103.95721318420905 | 26.904015319116187 | 119.11453981007931 | 116.32607139509724 | 8.019076393095819 | 149.070690516637 | 73.12290380186082 |
| SE_396_1_31 | SE_396 | SE_396_1 | GGAGAGTTCTCCATAACCAA | 146.30987362792035 | 157.9040125721556 | 170.8421472357761 | 227.72898567412008 | 195.14939034221084 | 153.33688944670834 | 212.00364071463557 | 323.3108937702153 | 224.66027528976795 | 181.52272926189627 | 227.71428315139823 | 54.55200759821362 |
| SE_397_1_10 | SE_397 | SE_397_1 | AAGGAGGGTGAGGCTCCGAG | 2816.7360115478887 | 3218.043800521146 | 3127.497573963601 | 2743.1582960050723 | 2874.211129496692 | 2750.534598832198 | 2590.3185949245667 | 2876.710671762471 | 3080.420928775514 | 2892.699557071331 | 2785.391721228186 | 2319.040663430443 |
| SE_397_1_24 | SE_397 | SE_397_1 | GGAAGCTTCTTAATAACGTG | 233.01202096298425 | 297.8189604209011 | 294.28300506509413 | 251.15253848631528 | 216.36128059679896 | 248.6310015322333 | 236.75533480822244 | 433.9172521652889 | 216.66840778934142 | 328.0531251721017 | 678.447709595403 | 156.69193671827318 |
| SE_397_1_44 | SE_397 | SE_397_1 | GCTCCTGTGTCCCATCCGCC | 79.11570944324582 | 90.94471610168455 | 100.7277998872348 | 87.18766808094883 | 85.90815553108195 | 252.96255208157535 | 80.71204595734856 | 161.65544685010764 | 76.3667338929646 | 70.71367364820858 | 109.1620017169074 | 61.51609367458132 |
| SE_397_1_7 | SE_397 | SE_397_1 | GTAGTGGGCCCTGCAAATTA | 1113.038816413883 | 1394.1525160642852 | 1438.8266388585307 | 1254.4613839420101 | 1521.9531257666986 | 1874.6950777552365 | 1235.4323834538152 | 1703.5269899822454 | 1651.6526167548159 | 1428.8536118607096 | 1778.2842215173625 | 1543.7057469281729 |
| SE_397_1_8 | SE_397 | SE_397_1 | CTGCGTTTCAGGGAGCAGAA | 277.4468714722045 | 301.8165303594366 | 321.93375721886133 | 277.1787082776433 | 276.85156782237514 | 282.41709581710126 | 214.15596194046484 | 193.79746556401793 | 288.59521529318016 | 182.25173620672317 | 176.06774470468935 | 178.74487596010422 |
| SE_398_1_10 | SE_398 | SE_398_1 | GGAACGGGATATCCATGTGG | 1501.030925738294 | 1744.9392781707827 | 1742.9849125499702 | 1591.5002827397077 | 1711.7995435452624 | 1655.5186199585291 | 1283.8596110282244 | 1930.4118277157297 | 1517.566839803215 | 1525.0825285778594 | 1881.5772984107803 | 1254.6961747589132 |
| SE_398_1_2 | SE_398 | SE_398_1 | GCTCCCAGCTGTGAGGAAGT | 484.4482482346696 | 544.6689041254734 | 507.58880739415565 | 503.60638546219695 | 603.4782777430324 | 451.34756724144097 | 499.33852432279645 | 603.135526976846 | 455.5364475243121 | 371.06453491688836 | 657.3195802308403 | 446.86218990026055 |
| SE_398_1_28 | SE_398 | SE_398_1 | AGGTCTCCTAAGGACCAGCA | 192.912278205172 | 226.86209401189444 | 214.29332919169605 | 201.70281588279207 | 309.69315957192674 | 401.10158086907325 | 187.25194662104866 | 212.70453575145163 | 175.82108500093836 | 227.45016678599052 | 136.15905590495976 | 235.61824558377373 |
| SE_398_1_8 | SE_398 | SE_398_1 | CAGGGTTTTGTTTTATTGCG | 592.8259324034995 | 831.4945472154017 | 820.6348228493061 | 723.5275201989186 | 715.9012960923495 | 782.2780292111731 | 525.166379029148 | 822.457536783881 | 827.6022789330582 | 696.9306392545094 | 860.384379123582 | 604.7148076312617 |
| SE_398_1_9 | SE_398 | SE_398_1 | CTGGGAACGGGATATCCATG | 496.3697934932409 | 530.6774093405988 | 555.9776236632483 | 517.9207788474274 | 603.4782777430324 | 546.6416793269659 | 432.6165663313883 | 584.2284571637223 | 465.304285580389 | 674.3314239648757 | 552.8527183727246 | 318.02659748745816 |

| | | | | | | | | | | | | | | | |
|---|---|---|---|---|---|---|---|---|---|---|---|---|---|---|---|
| SE_399_1_12 | SE_399 | SE_399_1 | GACCTTATCTTCAGGAGCAG | 173.40429467012783 | 230.85966395043002 | 245.89418879600146 | 161.36225270623365 | 259.84565561870465 | 153.33688944670834 | 151.73864639981528 | 253.35473546905757 | 281.49133307057883 | 172.04563897914667 | 241.7997027277734 | 304.0984253347228 |
| SE_399_1_16 | SE_399 | SE_399_1 | GAGAACACCACATAGCACAT | 3915.685729019824 | 3307.9891241381965 | 3438.5685356934823 | 3862.2835970330766 | 3771.4740872657703 | 3839.4864069367873 | 4005.469800710018 | 4094.3259675988375 | 3908.023207708572 | 3436.538737913972 | 3213.8232333429296 | 3709.5365166785264 |
| SE_399_1_19 | SE_399 | SE_399_1 | TCCTGCTTATCTCTCAACAG | 269.8604335803864 | 446.7284406313516 | 469.0752597514084 | 396.89908931775216 | 427.4195886299509 | 271.1550643888119 | 317.467380765571 | 513.3269453720085 | 514.1434758607733 | 785.1404795785634 | 453.0809963734006 | 480.52193926937105 |
| SE_399_1_7 | SE_399 | SE_399_1 | AATGTGCTATACCTATACAA | 136.55588205272565 | 166.89854493386065 | 212.31827546642697 | 161.36225270623365 | 133.63490860390525 | 141.20854790855063 | 103.31141882540615 | 198.52423301679886 | 205.12459917761421 | 64.15261114476655 | 143.20176569314734 | 170.62010887100857 |
| SE_399_1_9 | SE_399 | SE_399_1 | TTATCTTCAGGAGCAGAGGA | 33.59708209233727 | 43.97326932389143 | 62.21419234597627 | 61.16149900962082 | 54.09032014919974 | 51.11229648223612 | 49.50338818717378 | 87.9178746217252 | 23.087617223454416 | 13.122125006884067 | 113.8571415756912 | 80.08698987822851 |
| SE_4_1_11 | SE_4 | SE_4_1 | GCCACAGCCTCTACCCCAAT | 1320.040193176348 | 1498.0893344662104 | 1421.051155331109 | 1004.6101539452611 | 1250.4409305079705 | 852.4491481105142 | 993.2962455817697 | 591.7912850881718 | 1236.0755067326363 | 161.1105348067433 | 171.37260484589763 | 484.0039823075549 |
| SE_4_1_2 | SE_4 | SE_4_1 | ACTAGGATATGAGTGTACCA | 502.8724545433707 | 647.6063300427647 | 622.1419234597628 | 342.24413275596334 | 260.90625013143404 | 178.4598826328922 | 307.7819352506892 | 117.22383282896693 | 404.9212866882774 | 29.160277793075707 | 38.73490383503166 | 64.99813671276517 |
| SE_4_1_29 | SE_4 | SE_4_1 | ATGCCCCAGTGCACAAGTAA | 445.43228193389086 | 601.6342757496055 | 622.1419234597628 | 353.95590916206095 | 429.5407776554097 | 251.22993186183854 | 246.44078032310426 | 184.3439306584561 | 666.8769436467026 | 36.45034724134463 | 58.68924823489645 | 78.92630886550056 |
| SE_4_1_3 | SE_4 | SE_4_1 | GTATGCTTGCTAGTTGGCTT | 1341.7157300101142 | 1553.0559211210746 | 1464.5023372870287 | 1043.6494086322532 | 1051.0491621148421 | 696.5133283342007 | 815.7297444756027 | 490.63846159866 | 842.6980286560861 | 221.61811122737538 | 139.68041079905356 | 200.79781520193526 |
| SE_4_1_7 | SE_4 | SE_4_1 | TTACTCTGTGTCCCTCCACT | 130.05322100259588 | 183.8882171726369 | 203.43053370271608 | 83.28374333224963 | 194.08879582948143 | 138.6096175789454 | 49.50338818717378 | 49.15838150892162 | 107.44621861684554 | 10.93510417240339 | 18.780559435166865 | 45.26655949639003 |
| SE_4_2_0 | SE_4 | SE_4_2 | AAATTACAGTTAAGATTCCG | 8.670214733506391 | 14.990887269508443 | 22.713117840594514 | 16.917010364363207 | 27.575457330964575 | 7.796790988815679 | 24.75169409358689 | 1.89070698111237 | 10.655823333902038 | 0.0 | 7.042709788187574 | 1.1606810127279494 |
| SE_4_2_14 | SE_4 | SE_4_2 | AGGCAAAGTGATGTGAAGAA | 4600.6326929668285 | 5814.465475600008 | 6017.988700894912 | 4384.108301349203 | 5382.517152101739 | 2926.3955511354848 | 3203.7301442003554 | 2876.710671762471 | 4724.969663307728 | 1331.166681253906 | 1301.7275258500033 | 1231.4825545043543 |
| SE_4_2_20 | SE_4 | SE_4_2 | GATTTGGTCCATCCAAGGAC | 53.105065242726646 | 56.96537162413208 | 57.27655803280356 | 37.737946197425615 | 26.51486281823517 | 59.7753975809202 | 33.36097899570407 | 16.071009339455145 | 37.29538166865713 | 1.4580138896537853 | 9.390279717583432 | 19.73157721637514 |
| SE_4_2_27 | SE_4 | SE_4_2 | GTTTAGCCAGGGTAGAGGGT | 110.54523785220648 | 168.89732990312845 | 168.86709351050703 | 68.96934994701923 | 155.90739337122278 | 193.18715450065514 | 39.817942672291956 | 25.524544245016994 | 166.94123223113192 | 9.477090282749604 | 4.695139858791716 | 11.606810127279495 |
| SE_4_2_45 | SE_4 | SE_4_2 | GGGAGGGGACATCAAAAACA | 2518.6973800836067 | 3098.1167023650783 | 3075.15865024397 | 2766.581848818167 | 2960.1192850277744 | 3046.8126564071936 | 2886.2627634347846 | 2745.3065365751613 | 2869.0804326531234 | 2575.581536073412 | 2488.4241251596095 | 2014.9422380957203 |
| SE_4_3_20 | SE_4 | SE_4_3 | AGCAGGCAAGTCTACCCAGT | 2436.330340115296 | 2690.3645686344485 | 2601.145756179389 | 2177.089103044588 | 2472.245809172247 | 1821.8501610532635 | 1716.4761773596126 | 1388.7242776270357 | 2342.505162902798 | 716.6138267648355 | 626.8011711486942 | 738.1931240949758 |
| SE_4_3_34 | SE_4 | SE_4_3 | GGGATTCGCAGTTTGTCCTG | 244.93356622155557 | 281.8286806667587 | 255.7694574223469 | 300.6022610898385 | 225.90663121136365 | 246.8983813124965 | 203.39435581251837 | 288.33281461963645 | 200.68467278848837 | 164.75556953087775 | 266.4491869864299 | 240.26096963468552 |
| SE_4_3_37 | SE_4 | SE_4_3 | CGTATGACAGACATGAAAGG | 3271.9222850569745 | 3908.624007403168 | 4026.147018961036 | 2816.0315714216904 | 3301.6307181266434 | 2475.9142940039123 | 2238.414074550467 | 1647.7511340394306 | 3346.816512123065 | 758.8962295647952 | 498.85860999661986 | 763.7281063749907 |
| SE_4_3_4 | SE_4 | SE_4_3 | AGGCCCACACGCCTAAAGGT | 2326.8688791047775 | 2408.5358879676896 | 2442.1539312952273 | 2170.582560596756 | 2268.611662728201 | 2151.914312913127 | 1881.1287511126038 | 1387.7789241364796 | 2357.600912625826 | 755.9802017854877 | 968.3725958757915 | 836.8510101768516 |
| SE_4_3_7 | SE_4 | SE_4_3 | ATAAGTTTCTCGGGACACAG | 23.843090517142578 | 28.982382054382988 | 29.62580587903632 | 52.05233958265602 | 80.60518296743491 | 7.796790988815679 | 21.52321225529295 | 10.398888396118036 | 28.415528890405433 | 37.17935418617152 | 2.347569929395858 | 0.0 |
| SE_4_4_1 | SE_4 | SE_4_4 | GCTATTTTGGCAGAGCTGGG | 102.95879996038839 | 136.91677039484378 | 150.1040831204507 | 59.86019052005442 | 138.9378811675523 | 136.01068724934018 | 39.817942672291956 | 69.95615830115769 | 143.85361500767752 | 21.87020834480678 | 30.518409082146157 | 37.14179240729438 |
| SE_4_4_10 | SE_4 | SE_4_4 | ATAAGAAAAGAGATTCATGC | 110.54523785220648 | 166.89854493386065 | 167.87956664787248 | 105.40598765487843 | 135.75609762936406 | 122.14972549144564 | 109.76838250199404 | 100.20746999895562 | 173.1571291759081 | 80.91977087578509 | 66.90574298778195 | 38.30247342002233 |
| SE_4_4_19 | SE_4 | SE_4_4 | CTCAGGCAGAGGAAAATCAA | 537.5533134773963 | 626.6190878654529 | 564.8653654269592 | 618.1215325440402 | 594.9935216411972 | 303.20853845394305 | 485.348436356856 | 422.5730102786147 | 766.3312947631215 | 489.89266692367187 | 448.3858565146089 | 324.9906835638258 |
| SE_4_4_2 | SE_4 | SE_4_4 | GGAGCCCGTTTAGCATGACA | 639.4283365960963 | 814.5048749766254 | 739.6576201132735 | 584.2875118153138 | 483.63109780460945 | 543.1764388874923 | 547.7657518972055 | 362.07038688301884 | 906.6329686594984 | 591.2246322546099 | 251.1899824453568 | 331.95476964019355 |
| SE_4_4_8 | SE_4 | SE_4_4 | ATTCCATCGTTGGCAAGTGA | 20.59175999207768 | 40.97509186998975 | 27.650752153767232 | 11.711776406097604 | 34.999618920070425 | 0.8663101098684087 | 13.990087965940416 | 38.75949311280358 | 67.4868811147129 | 25.515243068941242 | 14.085419576375148 | 40.62383544547823 |
| SE_4_5_0 | SE_4 | SE_4_5 | TATGAAGTCAGAGATCCTAG | 947.2209596355732 | 1305.2065849318685 | 1246.2589006447945 | 888.7936983738515 | 1171.9569365659945 | 820.3956740453831 | 713.4944862629612 | 692.9441085776837 | 1297.3464909025731 | 552.5872641787846 | 334.5287149389098 | 654.6240911785635 |
| SE_4_5_1 | SE_4 | SE_4_5 | GCAATATGAAAAGAAAGATG | 508.2913387518122 | 636.6130127117918 | 708.0567605089681 | 553.0561080657202 | 715.9012960923495 | 509.39034460262434 | 433.6927269441529 | 532.2340151831322 | 569.1985630859339 | 258.06845846872 | 288.75110131569056 | 333.1154506529215 |
| SE_4_5_11 | SE_4 | SE_4_5 | GTGTACAAAGGACATCTCCA | 292.6197472558407 | 354.7843320450331 | 320.9462303562268 | 264.1656233819793 | 328.7842989461161 | 320.5347406513112 | 267.9639925783972 | 163.54615386622 | 515.0314611385985 | 94.77090282749604 | 57.515463270198524 | 218.20803039285448 |
| SE_4_5_13 | SE_4 | SE_4_5 | ACAGCTATAAAACAGCCAAG | 568.9828418863569 | 805.5103426149203 | 906.5496598985114 | 592.0953627527122 | 733.9314028087495 | 718.1710810809109 | 459.5205816505045 | 503.8734104664466 | 570.9745336415841 | 327.3241182272748 | 406.12959778548344 | 183.38760001101602 |
| SE_4_5_22 | SE_4 | SE_4_5 | TCAGAGTTAGAATTTGCCAG | 438.92962088376106 | 668.5935722200766 | 594.4911713059955 | 547.8508741074546 | 557.8727136956679 | 314.4705698822324 | 513.3286122887368 | 406.50200093915953 | 472.4081678029903 | 674.3314239648757 | 235.93077790428373 | 294.81297723289913 |
| SE_4_6_18 | SE_4 | SE_4_6 | ATTACTGCTGGAAGGCAGCA | 26.010644200519174 | 33.97934447755247 | 31.60085960430541 | 31.23140374959361 | 20.15129574185873 | 0.8663101098684087 | 3.228481838293942 | 0.0 | 5.327911666951019 | 0.7290069448268927 | 0.0 | 0.0 |
| SE_4_6_22 | SE_4 | SE_4_6 | TCTGCATGATCTATTTGTGG | 1019.8340080286893 | 1116.321405336062 | 1179.1070739856457 | 895.3002408216835 | 1042.5644060130069 | 740.6951439374894 | 807.1204595734856 | 504.8187639570028 | 923.5046889381766 | 288.6867501514495 | 385.00146842092073 | 275.081400016524 |
| SE_4_6_28 | SE_4 | SE_4_6 | GCAGAAATCAGGTGGCAATG | 2665.007253711527 | 3090.121562488007 | 2871.728116541254 | 2700.215115850281 | 2832.8479435002455 | 2168.374205000627 | 2383.695757273694 | 1732.832948189487 | 2050.358006498317 | 1106.632542247223 | 1500.0971848839533 | 1754.9496912446596 |
| SE_4_6_5 | SE_4 | SE_4_6 | ATTCTGGCAGCTCCTACTGA | 14.089098941947885 | 56.96537162413208 | 51.35139685699629 | 24.724861301761607 | 55.15091466192915 | 2.598930329605226 | 4.30464245105859 | 12.289595377230405 | 54.167101947335354 | 7.290069448268927 | 1.173784964697929 | 4.642724050911798 |
| SE_4_6_7 | SE_4 | SE_4_6 | TGGCTCGGCAATAATATAGG | 5.4188842084414945 | 5.9963549078033775 | 4.93763431317272 | 3.9039254686992013 | 6.363567076376441 | 0.0 | 0.0 | 0.0 | 0.0 | 0.7290069448268927 | 0.0 | 13.928172152735392 |
| SE_4_7_0 | SE_4 | SE_4_7 | ACACAATTATCTCATCAAAG | 255.77133463843853 | 297.8189604209011 | 328.8464452573032 | 300.6022610898385 | 293.7846800260457 | 278.0855452677592 | 230.29837113163455 | 182.45322367734371 | 317.89872946141077 | 263.17150708250824 | 120.8998513638867 | 199.6371341892073 |
| SE_4_7_10 | SE_4 | SE_4_7 | TTACTAAGAGTAACACAATA | 9.75399157519469 | 13.991494784874547 | 18.763010390056337 | 2.602616979132801 | 3.1817835381882205 | 45.048125713157255 | 0.0 | 7.56282792444948 | 1.7759705556503396 | 0.7290069448268927 | 1.173784964697929 | 6.964086076367696 |
| SE_4_7_12 | SE_4 | SE_4_7 | TTACTGAGCTTTAGTAAAGT | 39.01596630077876 | 34.97873696218637 | 55.301504307534465 | 32.53271223916001 | 33.939024407341016 | 34.65240439473635 | 50.57954879993843 | 22.68848377334844 | 53.27911666951019 | 6.561062503442034 | 46.95139858791716 | 89.37243798005211 |
| SE_4_7_14 | SE_4 | SE_4_7 | AGAGGAATTCACCATCAAGG | 301.2899619893471 | 422.74302100013807 | 453.2748299492557 | 294.0957186420065 | 426.3589941172215 | 354.32083493617915 | 215.2321225529295 | 409.3380614108281 | 286.81924473752986 | 143.61436813089784 | 171.37260484589763 | 219.36871140558245 |
| SE_4_7_5 | SE_4 | SE_4_7 | ATTGTGTGTTTTAGACTGGG | 119.21545258571288 | 191.88335704970808 | 178.74236213685248 | 147.04785932100324 | 142.1196647057405 | 28.588233625657487 | 86.0928490211718 | 64.28403735782058 | 143.85361500767752 | 29.8892847379026 | 57.515463270198524 | 304.0984253347228 |
| SE_40_10_13 | SE_40 | SE_40_10 | GTTACCTCATAATTTATCAG | 9.75399157519469 | 16.989672238776237 | 24.688171565863602 | 13.013084895664004 | 27.575457330964575 | 41.582885273683615 | 22.599372868057596 | 45.37696754669688 | 61.27098416993672 | 25.515243068941242 | 7.042709788187574 | 15.088853165463343 |
| SE_40_10_14 | SE_40 | SE_40_10 | CACATGAGAAAAATGAACCT | 1544.381999405826 | 1735.9447458090776 | 1728.172009610452 | 1595.404208208407 | 1806.1924551781797 | 1904.1496214907625 | 1558.2805672832096 | 1775.3738552645154 | 1656.9805284217668 | 1944.9905287981496 | 2436.7775867129008 | 1815.3051039065128 |
| SE_40_10_18 | SE_40 | SE_40_10 | GGAAAGATAACGCTGTTCAG | 343.5572588151908 | 452.72479553915497 | 437.474400147103 | 353.95590916206095 | 496.35823195736236 | 423.62564372565186 | 399.2555873356842 | 453.7696754669688 | 587.8462539202624 | 351.38134740656227 | 409.65095267957724 | 269.2779949528843 |
| SE_40_2_7 | SE_40 | SE_40_2 | CATGGGTAACTGGACTGCAG | 1132.5467995642723 | 283.8274656360265 | 585.6034295422846 | 815.920422958133 | 263.02743915689285 | 2727.1442258657507 | 1330.1345173771042 | 706.1790574454702 | 2259.922532065057 | 96.95792366197672 | 373.26361877394146 | 1048.0949544933383 |
| SE_40_2_8 | SE_40 | SE_40_2 | CGCTGCAGTCCAGTTACCCA | 1716.7025172342655 | 514.6871295864565 | 1029.002990865195 | 1363.7712970655878 | 431.66196668086855 | 4174.748419455862 | 2217.9670229079384 | 1016.2550023478989 | 3246.4741757288207 | 155.27847924812812 | 591.5876222077562 | 1301.1234152680313 |
| SE_40_2_9 | SE_40 | SE_40_2 | GCTGCAGTCCAGTTACCCAT | 1171.5627658650512 | 273.83354078968756 | 484.87568955356113 | 1029.3350152470227 | 218.48246962225778 | 2559.9463746611477 | 1451.7406666195095 | 583.2831036731661 | 2074.3336089995964 | 102.78997922059186 | 247.66862755126303 | 798.5485367568292 |
| SE_40_3_17 | SE_40 | SE_40_3 | ATGTAAGCAGATATTCTGAG | 621.0041302873952 | 741.549223598351 | 791.0090169702697 | 688.3921909806259 | 769.9916162415493 | 716.438460861174 | 713.4944862629612 | 589.9005781070595 | 955.4721589398827 | 605.0757642063209 | 744.179667618487 | 731.2290380186081 |
| SE_40_3_20 | SE_40 | SE_40_3 | TGTGGTGAGTTCCTTCTTAG | 902.786109126353 | 967.4119251256116 | 1040.8533132168095 | 831.5361248329299 | 1105.1394822640418 | 894.8983434940662 | 845.8622416330129 | 963.3152068767525 | 987.4396289415888 | 682.3505003579716 | 929.6376920407598 | 1072.4692557606252 |
| SE_40_3_21 | SE_40 | SE_40_3 | ACCACCACTGAAGGAGCCAG | 1908.5310182130943 | 1837.882779241735 | 1674.8455590281867 | 1836.1462787781911 | 1976.9481717276142 | 2130.256560166417 | 1680.9628771383793 | 2362.4383728999064 | 1950.0156701040728 | 1656.3037786467 | 1971.9587406925207 | 2230.828906463119 |
| SE_40_3_23 | SE_40 | SE_40_3 | TTACATGTGAGTTTTTAAAG | 22.759313675454276 | 69.95747392437274 | 38.51354764274722 | 22.12224432262881 | 46.6661585600939 | 0.8663101098684087 | 18.294730416999005 | 15.12565584889896 | 79.91867500426528 | 26.244250013768134 | 14.085419576375148 | 15.088853165463343 |
| SE_40_3_4 | SE_40 | SE_40_3 | AAAAATCAGGATTCTCCCAG | 2.167553683376598 | 4.996962423169481 | 2.962580587903632 | 0.0 | 7.4241615891058474 | 5.197860659210452 | 4.30464245105859 | 0.0 | 5.327911666951019 | 0.0 | 0.0 | 0.0 |
| SE_40_5_2 | SE_40 | SE_40_5 | CCTTGCTCAACACCTCTATG | 851.8485975670029 | 917.4423008939167 | 900.6244987227042 | 844.5492097285938 | 1007.5647870929364 | 819.5293639355147 | 885.6801843053049 | 834.7471321611114 | 928.8326006051276 | 1022.7967435921304 | 1068.1443178751153 | 1186.2159950079642 |
| SE_40_5_21 | SE_40 | SE_40_5 | TTGCCCAGACTGAAGCGCAG | 238.43090517142576 | 275.83232575895534 | 272.55741408713413 | 265.4669318715457 | 333.02667699703375 | 333.52939229933736 | 263.65935012733866 | 320.47483329854674 | 401.36934557697674 | 177.14868759293492 | 326.31222018602426 | 269.2779949528843 |
| SE_40_5_8 | SE_40 | SE_40_5 | AGGGGGAGCAGGAAATGCTA | 6589.363197464857 | 6499.0493275742265 | 6192.7809555812255 | 6613.2497439764475 | 6825.9862839264615 | 6421.090534344646 | 7145.706468757259 | 6783.8566482311835 | 6776.215655083871 | 6483.787767290383 | 7526.309193643121 | 6069.201015554448 |
| SE_40_6_10 | SE_40 | SE_40_6 | CCAGTATTGCAGAATAGGAG | 220.00669886272468 | 253.84569109700962 | 285.39526330138324 | 245.94730452804967 | 294.8452745387751 | 194.05346461052355 | 230.29837113163455 | 205.14170745069214 | 259.29170112494955 | 164.02656258605086 | 266.4491869864299 | 127.67491140007444 |
| SE_40_6_21 | SE_40 | SE_40_6 | CTACAAAGTGTGAAGACAAA | 202.6662693957119 | 277.83111072822317 | 275.51999467503776 | 290.1917931733073 | 217.42187510952837 | 229.5721791151283 | 213.0798013274002 | 251.4640284879452 | 325.89059696183733 | 441.0492016202701 | 312.2268006096491 | 159.01329874372908 |
| SE_40_6_22 | SE_40 | SE_40_6 | AGACAGGAGCTGAGTCTGGG | 1064.2688585379096 | 1313.2017248089396 | 1291.6851363259836 | 1339.0464357638261 | 1292.8647110171469 | 1982.1175313789192 | 1183.7766740411123 | 1418.9755893248337 | 1222.755727565259 | 2358.3374665149977 | 1102.1840818513554 | 1362.6395089426126 |
| SE_40_6_24 | SE_40 | SE_40_6 | GCTCTCTGAGATAGGCCTGG | 60.69150313454474 | 81.9501837399795 | 119.49075037877982 | 76.77720088441762 | 91.21112809472898 | 64.97325824013066 | 27.980175931880833 | 70.90151179171387 | 107.44621861684554 | 26.244250013768134 | 97.42415206992811 | 75.44426582731671 |
| SE_40_8_12 | SE_40 | SE_40_8 | CTGGTGCTGGCGTCAGGCGC | 1993.0656118647817 | 2056.749733376558 | 2010.6046923239317 | 1889.4999268504134 | 1951.4939034221084 | 1837.443743030895 | 1856.377057019017 | 1585.3578036627223 | 2195.0996067838196 | 1953.7386121360723 | 1186.6965993096062 | 2237.7929925394865 |
| SE_40_8_21 | SE_40 | SE_40_8 | CTGGCAAGGCTTGTTTACTA | 448.68361245895574 | 455.72297299305666 | 538.2021401358265 | 590.7940542631458 | 588.6299545648208 | 760.6202764644629 | 418.6264783654479 | 666.4742108421104 | 413.8011394665291 | 987.8044102404396 | 877.991153594051 | 834.5296481513957 |
| SE_40_8_23 | SE_40 | SE_40_8 | TCACTGCTGCTCCAGATCAT | 3560.206924946062 | 3018.1653035943664 | 3063.3083278923555 | 3421.140019070067 | 3469.2046511378894 | 3481.7003315611346 | 3541.644576608455 | 3130.0654072315288 | 3134.5880307228495 | 2889.7835292938025 | 2918.0294222390517 | 3130.35669132728 |
| SE_40_8_6 | SE_40 | SE_40_8 | CAGCTGACCCGGGAGGCCGC | 924.461645960119 | 1044.3651464424215 | 1029.9905177278295 | 1074.8808123818467 | 919.5354425363956 | 1070.7592957973532 | 1209.6045287474637 | 1143.8777235729838 | 1112.6455531149377 | 1721.9144036811203 | 1118.6170713571264 | 1240.768002606178 |
| SE_40_9_13 | SE_40 | SE_40_9 | GGACCTAATTCAATATAAGG | 23.843090517142578 | 35.97812944682026 | 55.301504307534465 | 76.77720088441762 | 57.27210368738796 | 93.56149186578814 | 18.294730416999005 | 31.196665188354107 | 19.535676112153734 | 19.683187510326103 | 36.3873339056358 | 87.05107595459621 |
| SE_40_9_17 | SE_40 | SE_40_9 | ATGTCGAGTTAGGAACAAAA | 453.01871982570896 | 632.6154427732563 | 658.6804173772408 | 503.60638546219695 | 514.3883386737623 | 481.6684210868353 | 356.2091628250983 | 412.17412188249665 | 502.5996672490461 | 363.04545852379255 | 545.810008584537 | 792.7451316931895 |
| SE_40_9_9 | SE_40 | SE_40_9 | TGGGCTTTGCTGCCACCTAG | 52.02128840103835 | 35.97812944682026 | 51.35139685699629 | 57.25757354092162 | 34.999618920070425 | 24.256683076315444 | 58.112673089290965 | 23.633837263904624 | 51.50314611385985 | 8.019076393095819 | 11.73784964697929 | 95.17584304369186 |
| SE_400_1_45 | SE_400 | SE_400_1 | GCCCCCAGCATGGCTTGGCA | 209.16893044584168 | 210.8718142577521 | 184.66752331265974 | 213.41459228888968 | 277.8757623351046 | 104.82352329407746 | 178.6426617189315 | 210.81382839402926 | 168.71720278678225 | 188.8127987101652 | 186.63180938697073 | 150.88853165463343 |
| SE_400_1_46 | SE_400 | SE_400_1 | GCCCTGCCAAGCCATGCTGG | 373.90301038246315 | 451.72540305452105 | 413.77375544387394 | 429.43180155691215 | 393.4805642226099 | 389.8395494407839 | 364.81844772721547 | 397.9938195241539 | 353.4181405744176 | 472.39650024782645 | 427.25772715004615 | 432.93401774752516 |
| SE_400_1_50 | SE_400 | SE_400_1 | GGCTTGGCAGGGCTGGCCCG | 589.5746018784346 | 633.6148352578901 | 657.6928905146063 | 654.5581702518995 | 670.295732044985 | 625.4758993249911 | 637.0870827566713 | 741.1571365960491 | 673.9808258693039 | 709.3237573165666 | 771.1767218065394 | 792.7451316931895 |
| SE_400_1_51 | SE_400 | SE_400_1 | GGCAGGGCTGGCCCGCGGCG | 825.8379533664838 | 1045.3645389270555 | 1102.079978700151 | 1013.719313372226 | 1149.684451798677 | 1062.9625048085375 | 1028.809545803003 | 1196.8175190441302 | 1244.0673742330628 | 1273.5751326125815 | 1158.525760156856 | 843.8150962532193 |
| SE_400_1_55 | SE_400 | SE_400_1 | GGCGACCCACTGCCACGCCG | 1998.4844960732232 | 2300.601499627229 | 2322.6631809164473 | 2334.5474302821226 | 2294.065931033707 | 2099.0693962111545 | 2315.8976386695213 | 2283.028679693187 | 2219.0752092850994 | 2149.1124733496795 | 2310.0088105255245 | 2476.893281161444 |
| SE_400_2_10 | SE_400 | SE_400_2 | TTAATCTTCACACAGTCCTT | 58.52394945116814 | 57.964764108765976 | 52.338923719630834 | 66.36673296788642 | 64.69626527649382 | 75.36897955855156 | 104.3875794381708 | 18.9070698111237 | 121.65398306204825 | 130.49224312401378 | 88.03387235234467 | 77.76562785277261 |
| SE_400_2_11 | SE_400 | SE_400_2 | GCAAGACCGAGCCAGCGGCA | 5595.539833636687 | 4987.967890807776 | 5151.927642364416 | 5417.347242064925 | 5318.881481337975 | 5392.780433930844 | 5882.293909371563 | 5199.444198059017 | 5111.243259161677 | 4703.5528080231115 | 5412.322472222151 | 5632.784954768738 |
| SE_400_2_12 | SE_400 | SE_400_2 | AAACTGCAAGACCGAGCCAG | 330.5519367149312 | 370.7746117991755 | 432.5367658339303 | 303.2048780689713 | 416.8136435026569 | 311.00532944275875 | 396.02710549739027 | 447.1522010330755 | 357.8580669635434 | 239.8432848480477 | 501.2061799260157 | 438.7374228111649 |
| SE_400_2_13 | SE_400 | SE_400_2 | GAGGTACCGAGTCATCCAGG | 1638.670584632708 | 2019.7722114451042 | 2086.6442607467916 | 1681.2905685197893 | 1740.4355953889565 | 1974.3207403901035 | 1668.0489497852036 | 1423.7023567776146 | 2247.4907381755047 | 1384.384188226269 | 1496.5758299898596 | 2005.6567899938966 |
| SE_400_2_5 | SE_400 | SE_400_2 | ACTGTGCATGATGGACACAG | 4584.376040341504 | 4095.5104020297067 | 4085.3986307191085 | 4322.946802339582 | 4333.589179012356 | 4053.4650040742845 | 4590.901174053986 | 3722.8020458102565 | 3612.3241101927906 | 3820.7253978377444 | 3987.347525078865 | 3974.171787580499 |
| SE_400_3_1 | SE_400 | SE_400_3 | CATATGACTCAGGGCTGCGA | 1061.0175280128447 | 1086.3396307970452 | 1107.0176130133239 | 1070.9768869131476 | 1210.1383390242531 | 1539.4330652361623 | 1134.2732858539384 | 1551.3250780026997 | 1035.390833944148 | 1086.949354736897 | 1813.4977704583005 | 886.7602937241534 |
| SE_400_3_10 | SE_400 | SE_400_3 | GAGGTAGGAGACTGTCAGGT | 2856.8357546903558 | 3428.9156147788976 | 3355.6162792321807 | 2922.7388675661355 | 3204.0560229555376 | 3578.7270638663963 | 2894.8720483369016 | 3183.005202702675 | 2993.3983715486474 | 3970.171821527257 | 3058.883618002803 | 3410.0808153947155 |
| SE_400_3_2 | SE_400 | SE_400_3 | CTGCTGTGTCATATGACTCA | 477.9455871845398 | 427.7399834233076 | 471.0503134766775 | 386.48862140122094 | 411.5106709390098 | 328.3315316401269 | 455.21593919944587 | 498.2012895231095 | 513.2554905829481 | 351.38134740656227 | 379.1325435974311 | 337.7581747038333 |
| SE_400_3_6 | SE_400 | SE_400_3 | AGTCTCCTACCTCACGGGCA | 55.27261892610324 | 89.94532361705066 | 87.88989077447442 | 57.25757354092162 | 81.66577748016432 | 161.133680435524 | 59.18883370205561 | 134.24019565897828 | 48.83919028038434 | 149.44642368951298 | 89.2076573170426 | 100.9792481073316 |
| SE_400_3_9 | SE_400 | SE_400_3 | CCTGACAGTCTCCTACCTCA | 3009.648289368406 | 2730.340268019804 | 2934.9298357498647 | 2866.78260251478 | 3173.298782086385 | 2592.8661588361474 | 2988.498021647426 | 3493.0811476051035 | 2548.517747358237 | 3696.7942172171724 | 3750.2429622098834 | 2798.401921687086 |
| SE_401_1_25 | SE_401 | SE_401_1 | GAGAAGCAGGTGAAGTGGAA | 2026.662693957119 | 2476.4945769227947 | 2374.0145777734438 | 2433.4468754891686 | 2428.7614341503413 | 2537.4223118045693 | 2445.036912201279 | 2664.006136387329 | 2633.7643340294535 | 2840.9400639904006 | 2842.907184498384 | 2957.415220430815 |

| | | | | | | | | | | | | | | | |
|---|---|---|---|---|---|---|---|---|---|---|---|---|---|---|---|
| SE_401_1_26 | SE_401 | SE_401_1 | GGAGAAGCAGGTGAAGTGGA | 2438.4978937986725 | 2788.3050321285705 | 2574.4825308882564 | 2606.5209046015 | 2724.667303201846 | 3025.1549036604833 | 2638.7458224989155 | 2600.667452520065 | 2348.721059847574 | 3413.939522624338 | 2508.3784695594745 | 3309.1015672873837 |
| SE_401_1_27 | SE_401 | SE_401_1 | GCGAGGAGAAGCAGGTGAAG | 15498.008836142675 | 15325.683751860797 | 14964.98207636388 | 15840.828243491793 | 15466.649779132938 | 18062.56579075632 | 16743.98297400515 | 16962.477681049626 | 15379.90501193194 | 18577.283975023707 | 17184.211883177682 | 17833.863760564942 |
| SE_401_1_28 | SE_401 | SE_401_1 | CTTCACCTGCTTCTCCTCGC | 184.24206308701082 | 259.842046004813 | 226.1436515433106 | 210.81197530975686 | 176.05868911308153 | 420.1604032861782 | 124.83463108069911 | 90.75393509339376 | 199.7966875106632 | 166.21358342053153 | 149.070690516637 | 219.36871140558245 |
| SE_401_1_32 | SE_401 | SE_401_1 | AGAAGAAGAGAACCCTGCGG | 30.34575156727237 | 48.970231747060915 | 49.376343131727204 | 40.3405631765841 | 44.54496953463509 | 71.90373911907793 | 38.74178205952731 | 37.8141396222474 | 17.759705556503395 | 13.122125006884067 | 45.7761362321923 | 84.7297139291403 |
| SE_401_2_11 | SE_401 | SE_401_2 | ATACGCTGCGGAAGGCCGCA | 108.37768416882989 | 120.92649064070145 | 130.3535458677598 | 123.62430650880805 | 134.69550311663465 | 171.52940175394494 | 75.33124289352533 | 116.27847933841076 | 182.0369819541598 | 104.24799311024564 | 45.7761362321923 | 241.4216506474135 |
| SE_401_2_20 | SE_401 | SE_401_2 | TCCCACATCCAGCCCTAAGG | 255.77133463843853 | 242.85237376603678 | 247.86924252127056 | 292.7944101524401 | 296.9664635642339 | 134.27806702960336 | 178.6426617189315 | 568.1574478242671 | 163.38929111983123 | 219.4310903928947 | 388.52282331501453 | 89.37243798005211 |
| SE_401_2_21 | SE_401 | SE_401_2 | AAGTGGCGCTCAGCATACAG | 1902.0283571629645 | 1803.9034347641825 | 1714.3466335335684 | 1754.1638439355079 | 1784.9805649235916 | 1909.3474821499728 | 1941.393745427424 | 2145.95242356254 | 1935.8079056588701 | 1880.108910708556 | 1707.8571236354867 | 1948.783420370227 |
| SE_401_2_5 | SE_401 | SE_401_2 | ACAAGCCATGATGATGATGG | 1483.6904962712813 | 1414.140365756963 | 1375.6249196499198 | 1262.2692348794085 | 1359.6821653190993 | 1421.6148902940588 | 1405.4657602706295 | 1347.1287240425636 | 1121.5254058931894 | 1155.476007550625 | 1348.6789244379204 | 1422.994921604466 |
| SE_401_2_6 | SE_401 | SE_401_2 | TACCCAGAGACAATACGCTG | 690.3658481554464 | 718.5631964517713 | 765.3333185417716 | 685.7895740014931 | 783.7793449070316 | 459.1443582302566 | 586.5075339567329 | 632.4414851820877 | 709.5002369823106 | 572.9994586339376 | 686.6642043482885 | 536.2346278803126 |
| SE_401_3_15 | SE_401 | SE_401_3 | GCCAGCACAGGACCCGGGTG | 851.8485975670029 | 815.5042674612592 | 925.3126702885678 | 884.8897729051523 | 934.3837657146073 | 894.0320333841978 | 871.6900963393645 | 1190.200044610237 | 785.8669708752752 | 662.6673128476455 | 1086.9248773102822 | 576.8584633257909 |
| SE_401_3_21 | SE_401 | SE_401_3 | GACCCGGGTGGGGCAGGGAG | 165.81785677830973 | 210.8718142577521 | 217.25590977959968 | 189.99103947669445 | 207.87652449496372 | 133.41175691973496 | 167.88105559128502 | 196.6335260356865 | 220.2203489006421 | 236.92725706874012 | 165.503680022408 | 59.19473164912542 |
| SE_401_3_28 | SE_401 | SE_401_3 | AAGTTGAATGATGCTGCCCG | 390.15966300778763 | 321.8043800521146 | 348.59698250999406 | 381.2833874429553 | 337.2690550479513 | 446.1497065822305 | 418.6264783654479 | 408.3927079202719 | 465.304285580389 | 326.59511128244793 | 240.62591776307545 | 414.3631215438779 |
| SE_401_3_29 | SE_401 | SE_401_3 | CAAGTTGAATGATGCTGCCC | 1741.6293845930963 | 1445.121532780614 | 1583.9930876658086 | 1484.7929865952628 | 1368.1669214209346 | 1526.4384135881362 | 1510.929500321565 | 1443.5547800792945 | 1147.2769789501194 | 1520.708486908898 | 1020.0191343225003 | 1228.0005114661706 |
| SE_402_1_1 | SE_402 | SE_402_1 | CTAACATCCTTCCCACCGAG | 1198.6571869072586 | 1084.3408458277775 | 1001.3522387114276 | 1156.86324722453 | 1001.20122001656 | 1155.6576865644572 | 1280.6311291899306 | 1057.850555932371 | 1141.0610820053432 | 1333.3537020883866 | 1184.3490293802104 | 788.1024076422776 |
| SE_402_1_15 | SE_402 | SE_402_1 | CAGGGCATAGAAGCCTAGCA | 4312.3480530777415 | 3989.5747986585134 | 4096.261426208089 | 4398.422694734433 | 4242.378050917627 | 4310.759106705202 | 4313.251735960707 | 4568.893419858042 | 3952.4224715998307 | 4375.499682851009 | 4148.156065242481 | 4372.285374946186 |
| SE_402_1_17 | SE_402 | SE_402_1 | ATAGAAGCCTAGCACGGCCA | 533.2182061106431 | 592.6397433879005 | 541.1647207237302 | 637.6411598875362 | 613.0236283575971 | 647.9999621815697 | 517.6332547397955 | 642.8403735782058 | 366.7379197417951 | 660.4802920131648 | 542.2886536904432 | 387.6674582511351 |
| SE_402_1_26 | SE_402 | SE_402_1 | TTCCGTCACAAGCATCCGGG | 19.50798315038938 | 62.96172653193546 | 52.338923719630834 | 14.314393385230405 | 40.30259148371746 | 68.4384986796043 | 8.60928490211718 | 63.33868386726439 | 42.62329333560815 | 6.561062503442034 | 90.38144228174053 | 3.482043038183848 |
| SE_402_1_7 | SE_402 | SE_402_1 | GAGCCACATCCTTTGCCCGG | 1436.004315236996 | 1818.894322033691 | 1779.5234064674482 | 1706.015429821551 | 1924.9790406038733 | 2034.962448080892 | 1694.9529651043197 | 2392.6896845977044 | 2166.6840778934143 | 1745.971632860408 | 2435.603801748203 | 2314.397939379531 |
| SE_403_1_20 | SE_403 | SE_403_1 | TTTTGTGTCTTCTCTTCACA | 17.340429467012783 | 19.987849692677923 | 41.47612823065085 | 22.12224432262881 | 15.9089176909411 | 108.2887637335511 | 4.30464245105859 | 12.289595377230405 | 56.831057780810866 | 2.187020834480678 | 11.73784964697929 | 112.5860582346111 |
| SE_403_1_23 | SE_403 | SE_403_1 | TCTCCCCTGCTTTTAAGTAG | 24.926867358830876 | 33.97934447755247 | 31.60085960430541 | 23.423552812195208 | 48.78734758555271 | 26.85561340592067 | 31.208657770174778 | 5.67212094333711 | 31.07948472388094 | 78.73275004130441 | 1.173784964697929 | 56.873369623669525 |
| SE_403_1_24 | SE_403 | SE_403_1 | CTCCCCTGCTTTTAAGTAGG | 7.586437891818092 | 10.993317330972857 | 7.900214901076352 | 2.602616979132801 | 10.605945127294067 | 1.7326202197368175 | 11.837766740411123 | 16.071009339455145 | 7.991867500426528 | 33.53431946203706 | 30.518409082146157 | 4.642724050911798 |
| SE_403_1_26 | SE_403 | SE_403_1 | GTTCCCCCTACTTAAAAGCA | 2.167553683376598 | 0.0 | 2.962580587903632 | 0.0 | 9.545350614564661 | 0.0 | 0.0 | 0.0 | 0.0 | 2.9160277793075706 | 0.0 | 0.0 |
| SE_403_2_15 | SE_403 | SE_403_2 | GGGTTGGGCCTGCCACTTAA | 320.7979451397365 | 328.80012744455183 | 354.5221436858013 | 301.9035695794049 | 329.8448934588455 | 526.7165467999926 | 274.42095625498513 | 469.84068480642395 | 512.367505305123 | 395.8507710410027 | 573.9808477372873 | 239.1002886219576 |
| SE_403_2_18 | SE_403 | SE_403_2 | AGTAAGTGAGACACACGAAG | 208.0851536041534 | 295.82017545163325 | 295.27053192772865 | 234.2355281219521 | 257.72446659324584 | 347.3903540572319 | 187.25194662104866 | 216.48594933736638 | 212.22848140021557 | 220.1600973377216 | 131.46391604616804 | 243.74301267286938 |
| SE_403_2_3 | SE_403 | SE_403_2 | AGTGGTAAATGACAAGGCCA | 664.3552039549272 | 759.5382883217611 | 785.0838557944625 | 664.9686381684306 | 702.1135674268672 | 794.4063707493308 | 496.11004248450246 | 730.758248199931 | 556.7667691963815 | 776.3923962406407 | 488.2945453143385 | 890.2423367623372 |
| SE_403_2_32 | SE_403 | SE_403_2 | GGATGGATGAGGCTATGCAG | 1158.5574437647915 | 1223.256401191889 | 1214.6580410404893 | 1076.1821208714132 | 1296.0464945553351 | 1568.8876089716882 | 1259.1079169346376 | 1738.5050691328242 | 1228.0836392322099 | 1437.6016951986323 | 2124.5507861032515 | 1846.6434912501675 |
| SE_403_2_4 | SE_403 | SE_403_2 | GCTACACTAGCAGGAACACG | 8948.745381820285 | 7999.137447009705 | 7711.597270313155 | 8455.90256520247 | 7664.916543495423 | 8500.234798028827 | 8444.632328364189 | 7380.374700772137 | 7546.986876236118 | 7116.565795400126 | 6856.077978800604 | 7472.464359942538 |
| SE_403_3_13 | SE_403 | SE_403_3 | TATGTTACGGTACAGCAGGA | 1082.6930648466107 | 1062.3542111658317 | 980.6141745961022 | 1137.3436198810339 | 1083.9275920094537 | 1363.5721129328754 | 1107.3692705348224 | 1015.3096488573427 | 946.592306161631 | 1234.2087575919293 | 897.9454979939158 | 875.1534835968739 |
| SE_403_3_18 | SE_403 | SE_403_3 | GTTGGGAAATCCAAGATCCA | 136.55588205272565 | 186.8863946265386 | 140.22881449410525 | 139.24000838360485 | 181.36166167672854 | 207.9144263684181 | 176.4903404934022 | 239.1744331107148 | 190.9168347324115 | 128.3052222895331 | 203.06479889274172 | 95.17584304369186 |
| SE_403_3_20 | SE_403 | SE_403_3 | ATGGCGGGATGGGAAAAGGA | 2657.420815819709 | 3329.975758800142 | 3185.761658859039 | 2352.765749136052 | 2404.367760357565 | 1801.0587184164217 | 1824.0922386360774 | 1212.8885283835853 | 3045.7895029403326 | 878.4533685164057 | 1194.9130940624918 | 1145.5921595624861 |
| SE_403_3_28 | SE_403 | SE_403_3 | CCTTCCCTGTACATTCCAGG | 500.7049008599941 | 556.6616139410802 | 572.7655803280355 | 568.6718099405169 | 506.9641770846564 | 404.5668213085469 | 515.4809335142661 | 512.3815918814523 | 700.620384204059 | 736.2970142751616 | 652.6244403720485 | 514.1816886384815 |
| SE_403_3_9 | SE_403 | SE_403_3 | TGATGCCTCCCGTTATGTTA | 2078.683982358157 | 1749.936240593952 | 1907.901898609939 | 1966.277127734831 | 1884.676449120156 | 1950.9303674236564 | 2083.4469463123573 | 1846.2753670562292 | 2132.940637336058 | 1772.944889819003 | 2071.730462691845 | 1380.049724133532 |
| SE_403_4_0 | SE_403 | SE_403_4 | TGGAGATGCCTGTAACCCAC | 1055.598643804403 | 909.4471610168455 | 869.0236391183987 | 1013.719313372226 | 909.990091921831 | 925.2191973394605 | 992.2200849690049 | 763.8456203693975 | 951.920217828582 | 1076.7432575093205 | 652.6244403720485 | 1079.433341836993 |
| SE_403_4_1 | SE_403 | SE_403_4 | TCTGATTTCCGGTGGGTTAC | 144.14231994454374 | 224.86330904262664 | 177.75483527421792 | 150.95178478970246 | 162.27096044759924 | 70.17111889934111 | 94.70213392328898 | 160.71009339455145 | 389.8255369652495 | 147.2594028550323 | 136.15905590495976 | 164.8167038073688 |
| SE_403_4_2 | SE_403 | SE_403_4 | AGATGCGTCTGATTTCCGGT | 2012.5735950151711 | 2098.724217731182 | 2095.5320025105025 | 1928.5391815374055 | 1839.0708850727913 | 1846.9731542394475 | 2027.486594448596 | 2035.3460651674663 | 1997.078889828807 | 2351.047397066729 | 2103.4226567386886 | 1888.4280077083738 |
| SE_403_4_3 | SE_403 | SE_403_4 | CAGATGCGTCTGATTTCCGG | 1318.9564163346597 | 1428.1318605418376 | 1587.9431951163467 | 1462.670742272634 | 1399.984756802817 | 1473.5934968861632 | 1376.4094237259842 | 1447.3361940415193 | 1715.587556758228 | 1358.139938212501 | 1302.9013108147012 | 1428.7983266681058 |
| SE_403_4_4 | SE_403 | SE_403_4 | TGTCAGATGCGTCTGATTTC | 1070.7715195880394 | 1130.3129001209365 | 1020.115249101484 | 1133.4396944123348 | 1079.685213958536 | 1126.2031428289313 | 956.7067847477716 | 935.8999556506232 | 1125.9653322823153 | 898.1365560267318 | 725.3991081833201 | 2059.048116579382 |
| SE_404_1_10 | SE_404 | SE_404_1 | TAGCTTTAGACACAAAACAG | 346.80858934025565 | 395.75942391502286 | 399.9483793669903 | 379.9820789533889 | 431.66196668086855 | 344.7914237276267 | 288.41104422092553 | 667.4195643326666 | 485.7279469703679 | 443.23622245475076 | 714.8350435010387 | 391.14950128931895 |
| SE_404_1_2 | SE_404 | SE_404_1 | GCACATACAGAGATGACACG | 5745.101037789673 | 5138.876155987494 | 5288.206349407983 | 5499.329676907609 | 5153.428737352187 | 5261.9676073407145 | 5852.161412214153 | 4809.95855994987 | 5303.048079171914 | 5638.139711291188 | 4868.86003356701 | 5153.423696512095 |
| SE_404_1_23 | SE_404 | SE_404_1 | AGTGCTGAGGAAAGGTGGAG | 912.5401007015477 | 1041.3669689885198 | 1040.8533132168095 | 991.5970690495972 | 954.5350614564661 | 1045.6363026111694 | 896.4417904329514 | 958.5884394239716 | 1293.7945497912724 | 969.5792366197672 | 836.9086798296235 | 1057.380402595162 |
| SE_404_1_3 | SE_404 | SE_404_1 | TTATCTCCACGTGTGACAGG | 79.11570944324582 | 67.95868895510495 | 64.18924607124536 | 71.57196692615203 | 107.12004578567009 | 71.90373911907793 | 96.85445514881827 | 57.66656292392729 | 55.94307250298569 | 37.17935418617152 | 26.997054188052367 | 63.83745570003722 |
| SE_404_1_41 | SE_404 | SE_404_1 | GAGGAGGAAAAGGAGCTGGA | 3846.324011151773 | 4654.1708009400545 | 4400.4196998995285 | 4113.436135519391 | 4373.891770496073 | 5033.2617383354545 | 4060.353991961015 | 5130.433393248416 | 4406.182948568492 | 4495.785828747447 | 4979.195820248615 | 5751.174418066989 |
| SE_404_2_1 | SE_404 | SE_404_2 | GAATTTGGAGAAGTCTGTGG | 189.6609472954523 | 340.7928372601586 | 353.53461682316674 | 242.04337905935049 | 268.3304117205399 | 164.59892087499765 | 215.2321225529295 | 392.32169858081676 | 291.2591711266557 | 385.6446738134262 | 798.1737759945918 | 246.06437469832528 |
| SE_404_2_13 | SE_404 | SE_404_2 | TATAAAGTGCAAACTTGGCA | 319.71416829804815 | 406.75274124599576 | 400.9359062296249 | 284.9865592150417 | 398.78353678625695 | 224.37431845591786 | 263.65935012733866 | 418.79159631639 | 285.9312594597047 | 220.8891042825485 | 361.52576912696213 | 554.8055240839599 |
| SE_404_2_17 | SE_404 | SE_404_2 | ACTTTATAAGCGTTGTCACA | 140.89098941947887 | 177.89186226483352 | 173.80472782367974 | 174.37533760189766 | 143.1802592184699 | 149.8716490072347 | 160.34793130193248 | 136.13090264009065 | 163.38929111983123 | 191.72882648947277 | 88.03387235234467 | 185.70896203647192 |
| SE_404_2_18 | SE_404 | SE_404_2 | CGTTGTCACAGGGAAGACCA | 9976.165827740791 | 8779.662977508779 | 8285.350377503824 | 9746.80058685234 | 9321.565172378756 | 9227.93529031829 | 10869.222188922939 | 9510.256114995222 | 9412.6439449468 | 10558.207581927887 | 9983.041124755886 | 10359.078038596948 |
| SE_404_2_4 | SE_404 | SE_404_2 | AAACTCTCTCCCAGTAGCAA | 13.005322100259587 | 25.984204600481302 | 23.700644703229056 | 9.109159426964803 | 22.272484767317543 | 6.064170769078861 | 15.066248578705064 | 13.23494886778659 | 19.535676112153734 | 7.290069448268927 | 1.173784964697929 | 24.37430126728694 |
| SE_404_3_10 | SE_404 | SE_404_3 | AAATCAGAAAGACAGCGCAG | 44.434850509220254 | 87.94653864778286 | 63.20171920861082 | 59.86019052005442 | 110.3018293238583 | 22.524062856578627 | 60.26499431482026 | 10.398888396118036 | 78.14270444861494 | 54.67552086201695 | 30.518409082146157 | 22.05293924183104 |
| SE_404_3_17 | SE_404 | SE_404_3 | TTACTACTGCTATTGATGAA | 122.46678311077778 | 94.94228604022014 | 96.77763253818532 | 97.59813671748003 | 106.05945127294068 | 99.625662634867 | 110.8445431147587 | 138.021609621203 | 72.81479278166393 | 110.0800486688608 | 185.45802442227279 | 16.249534178191293 |
| SE_404_3_3 | SE_404 | SE_404_3 | AGGAAACTGCTTTACTGACG | 2324.701325421401 | 2543.453873393266 | 2582.3827457893326 | 2476.3900556448602 | 2318.459604826483 | 2500.1709770802277 | 2301.907550703581 | 2538.2741221433566 | 2345.1691187362735 | 2813.237800086979 | 2623.409396099871 | 2497.785539390547 |
| SE_404_3_5 | SE_404 | SE_404_3 | AGCAGTTTCCTTCTACCGCA | 256.8551114801268 | 239.8541963121351 | 212.31827546642697 | 240.7420705697841 | 320.29954284428084 | 282.41709581710126 | 312.0865777017478 | 328.0376612229962 | 237.9800544571455 | 613.8238475442436 | 176.06774470468935 | 320.34795951291403 |
| SE_404_3_7 | SE_404 | SE_404_3 | AGAGCTGAAGAGCGACCCTG | 977.5667112028456 | 1111.3244429128927 | 1142.5685800681674 | 1128.2344604540692 | 995.898247452913 | 1250.0854885401138 | 1029.8857064157676 | 1292.2982215903048 | 976.7838056076868 | 1363.2429868262893 | 924.942552181968 | 846.1364582786751 |
| SE_405_1_11 | SE_405 | SE_405_1 | GGATGGATAACTAGAAGTTG | 710.9576081475241 | 823.4994073383305 | 913.4623479369533 | 771.6759343128755 | 855.8997717726312 | 925.2191973394605 | 742.5508228076068 | 979.3862162162077 | 1082.454053668882 | 1011.1326324749001 | 758.2650871948622 | 1015.5958861369558 |
| SE_405_1_13 | SE_405 | SE_405_1 | TCCTTAGGAGTAGATCTGGA | 240.59845885480235 | 269.83597085115196 | 269.5948334992305 | 234.2355281219521 | 302.2694361278809 | 188.8556039513131 | 298.0964897358073 | 274.1525122612937 | 357.8580669635434 | 253.69441679975864 | 201.8910139280438 | 127.67491140007444 |
| SE_405_1_18 | SE_405 | SE_405_1 | CTAAGGAAGATGACCCAAGG | 445.43228193389086 | 600.6348832649716 | 508.5763342567902 | 528.3312467639586 | 537.7214179538092 | 504.1924839434139 | 589.7360157950268 | 441.4800800897384 | 587.8462539202624 | 770.5603406820255 | 591.5876222077562 | 348.20430381838486 |
| SE_405_1_20 | SE_405 | SE_405_1 | CTAGATTGGATACAACACCT | 572.2341724114218 | 541.6707266715717 | 554.9900968006137 | 638.9424683771026 | 468.7827746263978 | 812.5988830565674 | 511.17629106320754 | 748.7199645204985 | 643.7893264232481 | 487.7056460891912 | 616.2371064664128 | 712.6581418149609 |
| SE_405_1_7 | SE_405 | SE_405_1 | GCAGCCTTCCTGTTCGACCC | 5674.6555430799335 | 4494.268003398631 | 4698.652812415161 | 4915.042165092294 | 4808.73552071513 | 5582.5023479920255 | 5550.8364406400515 | 4668.155536366442 | 4415.950786624569 | 6133.864433773475 | 3862.9263188208843 | 6090.093273783551 |
| SE_405_2_10 | SE_405 | SE_405_2 | GCTAGTATGTAGTACAACAA | 621.0041302873952 | 576.649463633758 | 590.5410638554573 | 592.0953627527122 | 632.1143295867264 | 641.9357914124909 | 636.0109221439067 | 567.212094333711 | 542.5590047511787 | 763.2702712337566 | 341.57142472709734 | 391.14950128931895 |
| SE_405_2_12 | SE_405 | SE_405_2 | GATACTATCAGACTAGGCTC | 272.027987263763 | 320.8049875674807 | 316.99612290568865 | 314.9166544750689 | 269.3910062332693 | 153.33688944670834 | 388.4939812080377 | 402.72058697693484 | 242.41998084627136 | 233.28222234460566 | 273.4918967746175 | 230.97552153286193 |
| SE_405_2_14 | SE_405 | SE_405_2 | CAGGGTTTACTATTGAACCA | 651.3498818546676 | 744.5474010522527 | 801.8718124592498 | 817.2217314476994 | 655.4474088667733 | 865.4437997585403 | 724.2560923906077 | 917.9382393300557 | 798.2987647648276 | 790.9725351371785 | 599.8041169606417 | 604.7148076312617 |
| SE_405_2_15 | SE_405 | SE_405_2 | ATAGTATCTAAAAATGAGAT | 19.50798315038938 | 22.986027146579612 | 14.81290293951816 | 16.917010364363207 | 14.848323178211695 | 10.395721318420904 | 39.817942672291956 | 20.79777679223607 | 5.327911666951019 | 45.198430579267345 | 1.173784964697929 | 2.321362025455899 |
| SE_405_2_7 | SE_405 | SE_405_2 | CAGCTCCCTTTGGTATACAT | 265.5253262136332 | 278.830503212857 | 328.8464452573032 | 321.4231969229009 | 325.60251540792785 | 487.7325918559141 | 284.1064017698669 | 352.61685197745703 | 367.6259050196203 | 199.0188959377417 | 214.80264853972102 | 384.1854152129513 |
| SE_406_1_16 | SE_406 | SE_406_1 | TGTGGGAGTTTCTCGTCGGG | 357.6463577571386 | 618.6239479883817 | 700.1565456078918 | 361.7637600994593 | 591.811738103009 | 557.9037107552552 | 297.0203291230427 | 584.2284571637223 | 751.2355450400936 | 629.8620003304353 | 654.9720103014444 | 470.0758101548195 |
| SE_406_1_18 | SE_406 | SE_406_1 | GGAGGCCTAGGAGAGGAGAA | 764.0626733902508 | 1146.303179875079 | 1034.9281520410022 | 595.9992882214113 | 877.1116620272194 | 421.02671339604666 | 456.29209981221055 | 461.3325033914183 | 896.8651306034214 | 314.20199322039076 | 322.79086529193046 | 313.38387343654637 |
| SE_406_1_27 | SE_406 | SE_406_1 | TATGTCCCAAGGCTGCACAG | 2025.5789171154306 | 1858.870021419047 | 1821.9870615607338 | 2010.5216163800887 | 1901.6459613238262 | 1885.9571091835257 | 1806.8736688318431 | 1857.6196089429036 | 2511.2223656895803 | 1245.1438617643328 | 2340.5272196076703 | 2085.7437798721253 |
| SE_406_1_3 | SE_406 | SE_406_1 | ATACCATGAGGAAACCAGCC | 486.6158019180462 | 428.73937590794145 | 433.5242926965648 | 451.554045879541 | 426.3589941172215 | 567.4331219638077 | 428.3119238803297 | 282.66069367629933 | 598.5020772541644 | 454.17132662715414 | 429.60529707944204 | 286.68821014380353 |
| SE_406_1_36 | SE_406 | SE_406_1 | GCTTCAGCCATGGCTCCAAG | 381.48944827428124 | 404.75395627672793 | 369.3350466253195 | 339.6415157768305 | 394.5411587353393 | 432.28874482433594 | 305.62961402515987 | 481.1849266930982 | 360.5220227970189 | 294.5188057100646 | 488.2945453143385 | 219.36871140558245 |
| SE_407_1_29 | SE_407 | SE_407_1 | GCTCCTAGAGAACAAGGACC | 54.188842084414944 | 76.95322131681 | 62.21419234597627 | 88.48897729051524 | 86.96875004381135 | 78.83421999802519 | 34.43713960846872 | 49.15838150892162 | 117.21405667292241 | 74.35870837234305 | 116.20471150509498 | 80.08698987822851 |
| SE_407_1_37 | SE_407 | SE_407_1 | TCCAGCCCAGGCCAAGCACA | 2169.7212370599746 | 1856.8712364497792 | 1860.500609203481 | 1824.4345023720934 | 1957.8574704984849 | 2119.860838847996 | 1998.4302579039504 | 1780.1006227172963 | 2085.877417611324 | 1852.4066468051342 | 1856.9278141521238 | 2276.095465959509 |
| SE_407_1_4 | SE_407 | SE_407_1 | CTGACCTGCTGTCATCGTAG | 441.09717456713764 | 422.74302100013807 | 395.01074505381763 | 420.3226421299473 | 445.44969534635084 | 333.52939229933736 | 351.90452037403975 | 605.0262339559584 | 418.241065855655 | 486.2476321995374 | 725.3991081833201 | 871.67144055869 |
| SE_407_1_5 | SE_407 | SE_407_1 | TCTACCTCTACGATGACAGC | 450.8511661423323 | 424.7418059694059 | 423.6490240702194 | 347.4493667142289 | 388.17759165896285 | 308.4063991131535 | 331.4574687315114 | 314.8027123552096 | 378.28172835352234 | 822.3198337647349 | 413.172307573671 | 311.06251141109044 |
| SE_407_1_8 | SE_407 | SE_407_1 | GGTAGAAAGCAAACAATCTG | 626.4230144958368 | 790.5194553454119 | 686.3311695310081 | 646.750319314501 | 715.9012960923495 | 1190.3100909591935 | 670.4480617523753 | 934.954602160067 | 585.1822980867869 | 642.2551183924925 | 995.3696500638438 | 778.816959540454 |
| SE_407_2_21 | SE_407 | SE_407_2 | CTAGCTGTTAGCCATCTCCA | 317.54661461467157 | 299.81774539016885 | 272.55741408713413 | 346.1480582246625 | 397.7229422735275 | 394.171099990126 | 328.2289868932175 | 375.30533575080545 | 287.70723001535504 | 201.93492371704926 | 616.2371064664128 | 157.85261773100112 |
| SE_407_2_23 | SE_407 | SE_407_2 | CAGCTAGAGCAGGCTGTCCT | 2182.726559160234 | 1919.8329629817147 | 1901.9767374341318 | 2143.2550823158617 | 1841.19207409825 | 2112.0640478591804 | 2230.8809502611143 | 1793.335571585083 | 1961.5594787158 | 1611.1053480674327 | 1410.8895275669106 | 2208.775967221288 |
| SE_407_2_24 | SE_407 | SE_407_2 | CTAGAGCAGGCTGTCCTCGG | 1316.7888626512831 | 1450.1184952037834 | 1495.1156700286997 | 1359.8673715968885 | 1506.0442080757575 | 1514.3100720499785 | 1296.7735383814002 | 1815.0787018678752 | 1470.5036200784812 | 1053.41503527486 | 1583.4359173775063 | 1340.5865697007816 |
| SE_407_2_25 | SE_407 | SE_407_2 | TAGAGCAGGCTGTCCTCGGA | 5359.276482148638 | 5597.597306434453 | 5616.065267802652 | 5563.093792896362 | 5713.422640073314 | 6158.598571054517 | 5234.445220487245 | 6285.655358708074 | 5385.630710009655 | 6446.608413104212 | 6718.745137930946 | 5165.030506639375 |
| SE_407_2_26 | SE_407 | SE_407_2 | AGAGCAGGCTGTCCTCGGAG | 2631.41017161919 | 3008.1713787480276 | 3078.1212308318736 | 2866.78260251478 | 3178.601754650032 | 2805.1121357539073 | 2804.4745568646713 | 3006.2240999686683 | 3162.1155743354298 | 2794.2836195214795 | 3016.6273592736775 | 3310.262248300112 |
| SE_408_1_12 | SE_408 | SE_408_1 | TGTGTTTCAGATATGCAAGG | 217.8391451793481 | 327.800734959918 | 290.3328976145559 | 262.8643148924129 | 261.9668446441635 | 236.50265999407557 | 232.45069235716386 | 173.94504226233803 | 362.29799335266927 | 394.3927571513489 | 138.50662583435562 | 496.77147344756236 |
| SE_408_1_4 | SE_408 | SE_408_1 | GACTAGCACACAAGTCCGGT | 1160.724997448168 | 1087.3390232816791 | 1062.5789041947694 | 1056.6624935279171 | 1199.532393896959 | 869.7753503078824 | 1112.7500735986455 | 966.1512673484211 | 848.0259403230372 | 1105.1745283575692 | 1061.1016080869279 | 1022.5599722133235 |
| SE_408_1_5 | SE_408 | SE_408_1 | TGAGCCAGCAGAAAGTACTG | 2885.0139525742516 | 3027.1598359560717 | 2996.1565012332067 | 2968.284664700959 | 2783.0000014019633 | 2997.4329801446943 | 2847.520981375257 | 2583.6510896900536 | 2827.3451245953406 | 2709.71881392156 | 2321.7466601725037 | 3233.657301460067 |
| SE_408_1_8 | SE_408 | SE_408_1 | ATATCCTGTATCCGAACAGA | 21.675536833765978 | 31.980559508284678 | 20.738064115325425 | 3.9039254686992013 | 32.87842989461161 | 28.588233625657487 | 20.4470516425283 | 4.726767452780925 | 1.7759705556503396 | 13.851131951710961 | 1.173784964697929 | 3.482043038183848 |

| | | | | | | | | | | | | | | |
|---|---|---|---|---|---|---|---|---|---|---|---|---|---|---|
| SE_408_1_9 | SE_408 | SE_408_1 | ACAGAAGGAGAGTCAGCTGG | 3052.999363035938 | 3055.1428255258206 | 3052.4455324033756 | 3068.485418397572 | 2994.0583094351155 | 3784.908870015078 | 3200.501662362062 | 3724.692752791369 | 3147.907809890227 | 3680.0270574861543 | 3443.885086423724 | 3444.901245776554 |
| SE_408_2_1 | SE_408 | SE_408_2 | ATCTCTGGACCCTTTCCCAA | 44.434850509220254 | 54.96658665486429 | 58.264084895438096 | 33.834020728726415 | 55.15091466192915 | 50.2459863723677 | 21.52321225529295 | 105.87959094229272 | 31.07948472388094 | 61.23658336545898 | 96.25036710523018 | 3.482043038183848 |
| SE_408_2_10 | SE_408 | SE_408_2 | ACTTCCTGTGGACTTCCTGT | 292.6197472558407 | 362.77947192210434 | 374.2726809384922 | 261.5630064028465 | 374.38986299348056 | 388.97323933091553 | 267.9639925783972 | 306.29453094020397 | 377.39374307569716 | 280.66767375835366 | 300.48895096266983 | 156.69193671827318 |
| SE_408_2_2 | SE_408 | SE_408_2 | AATCGGGGATTTCTTGCCCA | 72.61304839311603 | 125.92345306387092 | 124.42838469195254 | 134.03477442533924 | 98.63528968383483 | 168.0641613144713 | 112.99686434028798 | 75.6282792444948 | 147.40555611897818 | 80.91977087578509 | 46.95139858791716 | 104.46129114551545 |
| SE_408_2_22 | SE_408 | SE_408_2 | AATGGAGCTACAATCAACAA | 745.6384670815496 | 749.5443634754222 | 791.0090169702697 | 754.7589239485122 | 815.5971802889138 | 662.7272340493327 | 609.1069068247905 | 778.9712762182965 | 741.4677069840168 | 435.94615300648184 | 754.7437323007683 | 697.5692886494976 |
| SE_408_2_24 | SE_408 | SE_408_2 | CATTTGAATTAGCTGTAGAA | 27.094421042207472 | 47.97083926242702 | 52.338923719630834 | 53.35364807222242 | 51.96913112374093 | 72.77004922894633 | 32.28481838293943 | 47.26767452780925 | 51.50314611385985 | 53.21750697236316 | 42.25625872912545 | 241.4216506474135 |
| SE_409_1_0 | SE_409 | SE_409_1 | CATACTTCTAACCTTTACCA | 340.30592829012585 | 336.795267321623 | 345.63440192209043 | 394.2964723386193 | 446.5102898590802 | 630.6737599842015 | 317.467380765571 | 349.78079150578844 | 223.77229001194277 | 380.54162519963796 | 308.70544571555536 | 553.6448430712319 |
| SE_409_1_10 | SE_409 | SE_409_1 | GTCACTTTGGGGCTGCACAG | 341.38970513181414 | 436.7345157850126 | 462.1625717129666 | 350.0519836933617 | 421.05602155357445 | 427.9571942749939 | 360.5138052761569 | 302.5131169779792 | 536.3431078064026 | 420.63700716511704 | 388.52282331501453 | 432.93401774752516 |
| SE_409_1_11 | SE_409 | SE_409_1 | GTGCAGCCCCAAAGTGACAG | 635.0932292293431 | 706.5704866361646 | 674.4808471793935 | 632.4359259292706 | 572.7210368738796 | 421.02671339604666 | 590.8121764077914 | 515.2176523531208 | 618.0377533663182 | 820.8618198750811 | 339.2238547977015 | 688.283840547674 |
| SE_409_1_21 | SE_409 | SE_409_1 | CTTTTCACCACACAACCCCG | 151.72875783636184 | 112.93135076363026 | 124.42838469195254 | 160.06094421666725 | 180.30106716399914 | 226.10693867565467 | 166.80489497852037 | 120.0598933006355 | 131.42182111812514 | 41.553395855132884 | 142.02798072844942 | 157.85261773100112 |
| SE_409_1_23 | SE_409 | SE_409_1 | TCTTAGATGGCTCGACCACG | 729.3818144562251 | 755.5407183832256 | 776.1961140307516 | 705.309201344989 | 884.5358236163253 | 540.5775085578871 | 655.3818131736704 | 890.5229881039263 | 682.8606786475556 | 721.7168753786237 | 772.3505067712373 | 555.9662050966878 |
| SE_41_1_0 | SE_41 | SE_41_1 | GCTTTAAGAGAAATGAGAGT | 60.69150313454474 | 128.9216305177726 | 122.45333096668347 | 89.79028578008163 | 179.24047265126973 | 241.70052065328605 | 89.32133085946575 | 122.89595377230405 | 65.71091055906257 | 325.13709739279415 | 136.15905590495976 | 44.10587848366208 |
| SE_41_1_2 | SE_41 | SE_41_1 | AATACCTGGAGAAATCACAG | 29.26197472558407 | 26.983597085115196 | 25.675698428498144 | 33.834020728726415 | 42.42378050917627 | 8.663101098684088 | 16.142409191469714 | 22.68848377334844 | 39.07135222430747 | 100.60295838611118 | 8.216494752885502 | 2.321362025455899 |
| SE_41_1_3 | SE_41 | SE_41_1 | TTAGAATTGCAGGAAATACC | 226.50935991285448 | 246.84994370457235 | 316.99612290568865 | 222.52375171585447 | 236.5125763386577 | 193.18715450065514 | 212.00364071463557 | 120.0598933006355 | 381.833669464823 | 547.4842155649964 | 127.94256115207426 | 228.65415950740604 |
| SE_41_1_5 | SE_41 | SE_41_1 | ATATTCTCTAATATCAAGGG | 5.4188842084414945 | 4.996962423169481 | 1.975053725269088 | 15.615701874796805 | 6.363567076376441 | 0.0 | 10.761606127646475 | 0.0 | 0.0 | 0.7290069448268927 | 0.0 | 0.0 |
| SE_41_1_6 | SE_41 | SE_41_1 | TTAATATTCTCTAATATCAA | 3.2513305250648967 | 0.0 | 0.987526862634544 | 0.0 | 2.1211890254588135 | 0.0 | 0.0 | 0.0 | 0.8879852778251698 | 0.0 | 0.0 | 0.0 |
| SE_41_2_20 | SE_41 | SE_41_2 | TGAAGGTGCCTTCTGAACGA | 1656.0110140997208 | 1756.9319879863895 | 1727.1844827478176 | 1719.0285147172149 | 1620.5884154505336 | 1862.5667362170789 | 1400.0849572068064 | 1527.691240738795 | 1939.3598467701709 | 1111.0065839161844 | 1197.2606639918877 | 1478.7076102154076 |
| SE_41_2_25 | SE_41 | SE_41_2 | TTATAAAACACTCTCCCGGC | 8.670214733506391 | 14.990887269508443 | 14.81290293951816 | 5.205233958265602 | 3.1817835381882205 | 35.51871450460476 | 0.0 | 0.0 | 9.767838056076867 | 0.0 | 2.347569929395858 | 38.30247342002233 |
| SE_41_2_41 | SE_41 | SE_41_2 | TCTCCCACACCAGGTCCCAG | 201.5824925540236 | 306.81349278260615 | 285.39526330138324 | 231.63291114281927 | 344.6932166370572 | 262.49196329012784 | 159.27177068916782 | 415.01018235416524 | 322.3386558505366 | 199.74790288256858 | 416.6936624677648 | 364.4538379965761 |
| SE_41_2_48 | SE_41 | SE_41_2 | GGACCTTGCATTTGCTTCCA | 625.3392376541485 | 768.5328206834662 | 682.38106208047 | 637.6411598875362 | 706.3559454777849 | 593.42242525986 | 613.4115492758491 | 748.7199645204985 | 849.8019108786875 | 862.415215730214 | 590.4138372430583 | 583.8225494021585 |
| SE_41_2_5 | SE_41 | SE_41_2 | CTTCCTGCCTTGAGTCTGGA | 102.95879996038839 | 112.93135076363026 | 108.62795488979984 | 145.74655083143685 | 113.48361286204653 | 83.16577054736723 | 109.76838250199404 | 85.08181415005664 | 33.74344055735645 | 113.72508339299526 | 42.25625872912545 | 113.74673924733905 |
| SE_410_1_10 | SE_410 | SE_410_1 | TATATCTTAGTGTTTCTGTG | 28.17819788389577 | 48.970231747060915 | 30.613332741670867 | 16.917010364363207 | 37.120807945529236 | 58.90908747105179 | 24.75169409358689 | 80.35504669727572 | 48.83919028038434 | 9.477090282749604 | 113.85714157569912 | 12.767491140007444 |
| SE_410_1_12 | SE_410 | SE_410_1 | CTATTTCATTCTGGGTGGCG | 7.586437891818092 | 21.986634661945715 | 18.763010390056337 | 14.314393385230405 | 28.63605184369398 | 22.524062856578627 | 3.228481838293942 | 11.34424188667422 | 28.415528890405433 | 1.4580138896537853 | 2.347569929395858 | 6.964086076367696 |
| SE_410_1_13 | SE_410 | SE_410_1 | GACCAGAGTAATGGTAGTGG | 1743.796938276473 | 2016.7740339912025 | 1969.1285640932808 | 1791.9017901329335 | 2032.0990863895433 | 1998.5774234664188 | 1698.1814469426138 | 2723.563406292369 | 1612.5812645305084 | 1854.593667639615 | 2731.397612852081 | 1605.2218406027541 |
| SE_410_1_18 | SE_410 | SE_410_1 | ACAAGGGTTAGAACTATGGG | 419.42163773337165 | 457.72175796232443 | 494.75095817990655 | 407.3095572342833 | 572.7210368738796 | 785.7432696506468 | 450.9112967483873 | 559.6492664092615 | 493.7198144707944 | 511.03386832365175 | 487.12076034964053 | 714.9795038404169 |
| SE_410_1_19 | SE_410 | SE_410_1 | AGAAGGTGGGACATTATACG | 356.5625809154503 | 367.7764343452738 | 373.28515407585763 | 244.64599603848328 | 370.147484942563 | 336.994632738811 | 305.62961402515987 | 435.8079591464013 | 253.07580418017338 | 234.01122928943255 | 482.4256204908488 | 510.69964560029774 |
| SE_411_1_27 | SE_411 | SE_411_1 | CCTTTGCACCCCTCTCTGGG | 68.27794102636283 | 135.9173779102099 | 115.54064292824165 | 109.30991312357764 | 150.60442080757576 | 105.68983340394587 | 90.39749147223039 | 146.52979103620868 | 165.1652616754816 | 215.7860556687602 | 367.39469395045177 | 106.78265317097134 |
| SE_411_1_28 | SE_411 | SE_411_1 | CTTCCCCATGTTTACAAGGA | 52.02128840103835 | 69.95747392437274 | 92.82752508764715 | 55.95626505135522 | 62.575076251035 | 53.71122681184134 | 69.95043982970209 | 34.03272566002266 | 83.47061611556596 | 51.759493082709376 | 16.432989505771005 | 100.9792481073316 |
| SE_411_1_37 | SE_411 | SE_411_1 | CCGCTTCTCCTGCATGGCCT | 48.769957875973454 | 32.97995199291857 | 46.41376254382357 | 50.751031093089615 | 36.06021343279983 | 15.593581977631358 | 87.16900963393644 | 50.10373499947781 | 9.767838056076867 | 34.992333351690846 | 36.3873339056358 | 2.321362025455899 |
| SE_411_1_40 | SE_411 | SE_411_1 | AGGCCATGCAGGAGAAGCGG | 5050.4000822674725 | 5691.540199990039 | 5666.429137797014 | 4488.212980514515 | 4984.794209828212 | 3775.3794588065252 | 4078.648722378014 | 3357.895598455569 | 4939.86210054142 | 2996.218543238529 | 3015.4535743089796 | 2626.6211318033497 |
| SE_411_1_41 | SE_411 | SE_411_1 | GCACTCCCTTCCAGCAACTG | 1081.6092880049223 | 1066.3517811043673 | 953.9509493049695 | 1094.4004397253427 | 989.5346803765365 | 1407.7539285361643 | 1147.1872132071142 | 1001.1293464989999 | 1081.566068391057 | 1669.425903653584 | 1190.2179542037 | 918.098681067808 |
| SE_411_2_15 | SE_411 | SE_411_2 | TGTCAGTAGAAACTTCCTGC | 610.1663618705123 | 639.6111901656935 | 656.7053636519718 | 594.697979731845 | 572.7210368738796 | 516.3208254815715 | 495.03388187173783 | 553.9771454659244 | 546.1109458624794 | 401.68282659961784 | 713.6612585363408 | 654.6240911785635 |
| SE_411_2_16 | SE_411 | SE_411_2 | GTCAGTAGAAACTTCCTGCA | 66.11038734298623 | 29.981774539016886 | 40.488601368016305 | 49.44972260352321 | 53.02972563647034 | 173.26202197368175 | 120.52998862964051 | 51.04908849003399 | 145.62958556332785 | 8.748083337922711 | 11.73784964697929 | 39.46315443275028 |
| SE_411_2_18 | SE_411 | SE_411_2 | TGCTGGATGAGCAGAACAAG | 1279.940450033881 | 1073.3475284968044 | 1089.242129485902 | 1331.2385848264275 | 1140.1391011841122 | 1412.0854790855062 | 1262.3363987729315 | 1365.0904403631312 | 840.0340728226106 | 1057.7890769438213 | 1227.7790730740337 | 1333.622483624414 |
| SE_411_2_19 | SE_411 | SE_411_2 | GTTCTGCTCATCCAGCAAGA | 59.60772629285644 | 49.96962423169481 | 59.25161175807264 | 58.55888203048802 | 39.24199697098805 | 91.82887164605133 | 63.4934761531142 | 41.59555358447214 | 110.11017445032105 | 8.748083337922711 | 30.518409082146157 | 106.78265317097134 |
| SE_411_2_20 | SE_411 | SE_411_2 | AGATGCAGCTGCCATCTTGC | 7984.183992717698 | 6772.882868363914 | 6736.90825689286 | 7366.707359435393 | 7218.406253636343 | 6448.812457860435 | 8148.68815985391 | 7092.9872396430565 | 6571.091055906257 | 7273.302288537908 | 6444.07945619163 | 7344.789448542464 |
| SE_412_1_1 | SE_412 | SE_412_1 | AAAGAGAGTACACATCGACG | 484.4482482346696 | 640.6105826503275 | 661.6429979651446 | 433.33572702561133 | 509.0853661101152 | 275.486614938154 | 309.9342564762185 | 266.5896843368442 | 489.27988808166856 | 353.56836824104295 | 320.4432953625346 | 217.04734938012655 |
| SE_412_1_10 | SE_412 | SE_412_1 | TGTATCTCCAGAACTCATGT | 69.36171786805113 | 106.9349958558269 | 109.61548175243439 | 87.18766880094883 | 100.75647870929365 | 108.2887637335511 | 100.08293698711222 | 34.03272566002266 | 106.55823333902038 | 172.04563897914667 | 31.692194046844083 | 48.74860253457388 |
| SE_412_1_11 | SE_412 | SE_412_1 | AAGAAGGCAGATTCCGTCAG | 753.2249049733678 | 971.4094950641471 | 971.7264328323913 | 681.8856485327939 | 755.1432930633376 | 528.4491670197293 | 644.6202070460239 | 690.1080481060151 | 621.5896944776189 | 425.7400557789053 | 572.8070627725893 | 673.1949873822107 |
| SE_412_1_7 | SE_412 | SE_412_1 | GGTGGCAGCAGTGAGAGGAA | 810.6650775828476 | 998.3930921492623 | 1017.1526685135804 | 822.4269654059651 | 719.0830796305378 | 748.4919349263051 | 611.2592280503197 | 792.206225086083 | 932.3845417164283 | 344.82028490312024 | 416.6936624677648 | 279.7241240674358 |
| SE_412_1_8 | SE_412 | SE_412_1 | GCATCTTTTGATTTCACCAG | 65.02661050129794 | 104.9362108865591 | 119.49075037877982 | 102.80337067574564 | 113.48361286204653 | 166.33154109473446 | 45.19874573611519 | 156.9286794323267 | 47.95120500255917 | 91.85487504818848 | 134.98527094026184 | 78.92630886550056 |
| SE_413_1_24 | SE_413 | SE_413_1 | TCCAGCTGTCACCAAGAAGA | 1265.851351091933 | 1234.2497185228617 | 1352.9118018093254 | 1311.7189574829317 | 1374.5304884973111 | 1194.6416415085357 | 1392.5518329174538 | 1219.5060028174787 | 1348.849637016433 | 1252.4339312126017 | 1383.8924733788583 | 1450.8512659099367 |
| SE_413_1_25 | SE_413 | SE_413_1 | GTGACAGCTGGAAGTAGCAG | 1347.1346142185555 | 1389.1555536411156 | 1429.9388970948198 | 1441.8498064395717 | 1619.527820937804 | 1235.3582166723509 | 1369.9524600493962 | 1497.439929040997 | 1514.0148986919144 | 1509.0443757916678 | 1257.123697191482 | 1903.516860873837 |
| SE_413_1_26 | SE_413 | SE_413_1 | CTGGGGGCTGTAGAAATGTT | 867.0214733506391 | 995.3949146953606 | 944.075680678624 | 1030.6363237365892 | 998.0194364783717 | 1031.7753408532749 | 1016.9717790625918 | 926.4464207450613 | 779.651073930499 | 574.4574725235914 | 815.7805504650606 | 844.9757772659472 |
| SE_413_1_27 | SE_413 | SE_413_1 | CAAACATTTCTACAGCCCCC | 393.4109935328525 | 462.71872038549395 | 491.7883775920029 | 396.89908931775216 | 428.48018314268035 | 444.4170863624937 | 377.73237508039125 | 508.6001779192275 | 539.8950489177032 | 459.2743752409424 | 662.014720089632 | 486.32534433301083 |
| SE_413_1_28 | SE_413 | SE_413_1 | GGGTGACAGGAATGTCCTGG | 531.0506524272664 | 758.5388958371273 | 663.6180516904136 | 562.165267492685 | 558.9333082083973 | 584.7593241611759 | 510.1001304504429 | 502.9280569758904 | 560.3187103076822 | 581.0185350270334 | 517.6391694317867 | 637.2138759876442 |
| SE_413_2_22 | SE_413 | SE_413_2 | CAATGCTTCTGGGTACACAG | 592.8259324034995 | 672.5911421586121 | 665.5931054156827 | 633.737234418837 | 564.2362807720444 | 676.5881958072272 | 574.6697672163217 | 545.4689640509188 | 816.058470321331 | 457.0873544064617 | 610.3681816429231 | 925.0627671441757 |
| SE_413_2_31 | SE_413 | SE_413_2 | TGATGGGGTGACTTTCTGGG | 808.4975238994709 | 1296.2120525701635 | 1285.7599751501764 | 947.3525804043395 | 1319.379573835382 | 1385.2298656795856 | 826.4913506032492 | 1073.9215652718262 | 1182.7963900631262 | 1171.5141603368165 | 829.8659700414358 | 1292.9986481789356 |
| SE_413_2_35 | SE_413 | SE_413_2 | GGAGGGGTCACAGATGAAAA | 1145.552121664532 | 1422.1355056340342 | 1377.599973375189 | 1100.9069821731748 | 1311.9554122462762 | 1503.9143507315575 | 1153.6441768837021 | 1390.6149846081482 | 1581.5017798066274 | 1179.5332367299122 | 1544.7010135424746 | 1678.3447444046149 |
| SE_413_2_44 | SE_413 | SE_413_2 | GCCCAGGCTAAAAGGCCCTG | 79.11570944324582 | 95.94167852485404 | 122.45333096668347 | 78.07850937398402 | 96.51410065837601 | 43.31550549342044 | 69.95043982970209 | 112.49706537618601 | 131.42182111812514 | 110.80905561368769 | 35.21354894093787 | 63.83745570003722 |
| SE_413_2_7 | SE_413 | SE_413_2 | GGCCCAAAGGACATGAACAA | 225.42558307116616 | 256.84386855091134 | 259.7195648728851 | 189.99103947669445 | 322.42073186973965 | 291.08019691578534 | 217.3844437784588 | 250.518674997389 | 229.1002016788938 | 187.3547848205114 | 212.45507861032516 | 132.31763545098624 |
| SE_413_3_1 | SE_413 | SE_413_3 | GAAGAAACTTAACAATGTAA | 106.21013048545329 | 129.9210230024065 | 145.16644880727796 | 137.93869989403845 | 120.90777445115236 | 123.01603560131404 | 90.39749147223039 | 227.8301912240406 | 126.09390945117411 | 47.38545141374802 | 269.9705418805237 | 129.99627342553035 |
| SE_413_3_25 | SE_413 | SE_413_3 | TTAGGTTGCTGACACACCAG | 651.3498818546676 | 621.6221254422834 | 646.8300950256263 | 650.6542447832002 | 681.9622716850085 | 675.7218856973589 | 708.113683199138 | 469.84068480642395 | 1075.3501714462807 | 559.1483266822266 | 915.5522724643846 | 868.1893975205062 |
| SE_413_3_26 | SE_413 | SE_413_3 | GTTAGGTTGCTGACACACCA | 1246.3433679415436 | 1257.2357456694415 | 1247.2464275074292 | 1218.0247462341508 | 1238.774390867947 | 1302.0640951322184 | 1270.9456836750487 | 1136.3148956485343 | 1197.892139786154 | 908.3426532543083 | 1321.681870249868 | 1525.1348507245254 |
| SE_413_3_3 | SE_413 | SE_413_3 | CTCTCAGGAGAAATGGCCAG | 80.19948628493412 | 68.95808143973883 | 92.82752508764715 | 76.77720088441762 | 82.72637199289373 | 151.60426922697152 | 86.0928490211718 | 91.69928858394995 | 92.35046889381766 | 31.347298627556384 | 161.98232512831422 | 146.24580760372163 |
| SE_413_3_4 | SE_413 | SE_413_3 | CTAGGAATAGAAAGAAACAC | 418.33786089168336 | 510.68955964792093 | 459.199991125063 | 469.77236473347057 | 415.75304898992744 | 417.561472956573 | 487.5007575823853 | 342.21796358133895 | 469.7442119695148 | 365.9614863031001 | 404.9558128207855 | 694.0872456113137 |
| SE_413_4_19 | SE_413 | SE_413_4 | TGTGCAGTGGAACATTGCCT | 9584.922387891316 | 8180.027486728441 | 7979.217050087116 | 9116.967277902202 | 8184.607854732832 | 8438.72678022817 | 9532.630707869248 | 8877.814629813134 | 7572.738449293048 | 7747.885809620215 | 8453.599315754485 | 7657.012640966283 |
| SE_413_4_2 | SE_413 | SE_413_4 | TCACAGAGTCCTTTGTGCCA | 997.074694353235 | 1196.2728041067737 | 1056.6537430189621 | 929.1342615504099 | 1210.1383390242531 | 1112.3421810710367 | 952.402142296713 | 1191.145398100793 | 1012.3032167206935 | 769.1023267923717 | 1495.4020450251617 | 1563.4373241445478 |
| SE_413_4_26 | SE_413 | SE_413_4 | CTGCCCCTGGGAATTCACAT | 128.96944416090756 | 189.88457208044028 | 205.40558742798515 | 134.03477442533924 | 172.8769055748933 | 499.8609333940718 | 130.21543414452233 | 175.8357492434504 | 162.50130584200608 | 177.8776945377618 | 83.33873249355297 | 160.17397975645702 |
| SE_413_4_30 | SE_413 | SE_413_4 | ATAGGGAGCTGCTCCCTGAG | 464.9402650842802 | 597.6367058110699 | 711.0193410968717 | 520.5233958265602 | 649.083841790397 | 415.8288527368362 | 431.5404057186236 | 448.0975545236317 | 639.3494000341223 | 518.3239377719207 | 180.76288456348107 | 348.20430381838486 |
| SE_413_4_37 | SE_413 | SE_413_4 | GCAGGAGGAGAAGAGAGCCT | 10005.427802466376 | 9780.054854627308 | 9254.114229748313 | 9879.534052788113 | 10080.95084349301 | 9891.52883447749 | 10776.67237622518 | 10429.139707815833 | 10588.336452787324 | 11470.924276851156 | 11141.566884912743 | 10041.05144110949 |
| SE_413_5_13 | SE_413 | SE_413_5 | AGATCACCTTTCCAGCTGGG | 525.631768218825 | 474.7114302011007 | 554.0025699379792 | 546.5495656178882 | 530.2972563647033 | 1043.9036823914325 | 577.8982490546157 | 784.6433971616335 | 551.4388575294304 | 476.0415349719609 | 1034.1045538988756 | 723.1042709295125 |
| SE_413_5_19 | SE_413 | SE_413_5 | CATTCTTTCTACCTTTCTAG | 15.172875783636185 | 43.97326932389143 | 24.688171565863602 | 32.53271223916001 | 28.63605184369398 | 5.197860659210452 | 7.533124289352532 | 37.8141396222474 | 3.551941111300679 | 54.67552086201695 | 17.606774470468935 | 42.945197470934126 |
| SE_413_5_21 | SE_413 | SE_413_5 | TGAACCACAAACTTAAGAGG | 242.76601253817896 | 305.81410029797223 | 331.8090258452068 | 329.2310478602993 | 277.8757623351046 | 421.89302350591504 | 269.0401531911619 | 373.4146287696931 | 276.1634214036278 | 363.04545852379255 | 372.0898338092435 | 163.65602279464088 |
| SE_413_5_3 | SE_413 | SE_413_5 | TTAACACCTTTGAGGTAAAG | 49.85373471766175 | 22.986027146579612 | 18.763010390056337 | 48.14841411395682 | 38.181402458258646 | 48.51336615263089 | 91.47365208499504 | 1.89070698111237 | 2.6639558334755096 | 1.4580138896537853 | 35.21354894093787 | 5.803405063639747 |
| SE_413_5_9 | SE_413 | SE_413_5 | CCTCCATTCAGCTGTTTGCA | 567.8990650446686 | 672.5911421586121 | 615.2292354213209 | 675.3791060849618 | 715.9012960923495 | 920.0213366802501 | 626.3254766290248 | 622.9879502765259 | 562.0946808633324 | 535.0910975029392 | 842.777604653113 | 821.7621570113882 |
| SE_414_1_10 | SE_414 | SE_414_1 | CATCTGATTAAACCATCCAA | 254.68755779675024 | 346.78919216796197 | 325.88386466939954 | 262.8643148924129 | 374.38986299348056 | 323.99998109078484 | 246.44078032310426 | 519.9444198059017 | 123.4299536176986 | 180.0647153722425 | 727.746678112716 | 330.7940886274656 |
| SE_414_1_16 | SE_414 | SE_414_1 | GACAGGAGTTCAGGAAGCTG | 1669.0163361999803 | 1852.8736665112435 | 1844.7001794013283 | 1810.120108986863 | 1760.5868911308153 | 1667.6469614966868 | 1616.3932403725005 | 2053.307781488034 | 1560.1901331388233 | 1632.2465494674127 | 2308.8350255608266 | 1579.6868583227392 |
| SE_414_1_25 | SE_414 | SE_414_1 | CATAGACAAGGGGGACACAG | 1022.0015617120658 | 1073.3475284968044 | 1097.1423443869785 | 1057.9638020174837 | 1055.2915401657597 | 1075.0908463466953 | 1072.9321309263535 | 1001.1293464989999 | 943.9283503281555 | 834.712951826792 | 963.6774560169997 | 775.3349165022702 |
| SE_414_1_3 | SE_414 | SE_414_1 | GATGATTTGGTGTGATACCA | 930.9643070102487 | 1013.3839794187708 | 993.4520238103513 | 934.3394955086756 | 1023.4737047838776 | 1290.802063703929 | 973.925354552006 | 1176.019742251894 | 1297.3464909025731 | 1024.2547574817843 | 1444.9292915431506 | 1716.6472178246372 |
| SE_414_1_6 | SE_414 | SE_414_1 | TGCCAGGGAAAAAAATGGCG | 443.2647282505143 | 667.5941797354426 | 632.0171920861081 | 400.80301478645134 | 641.659680201291 | 554.4384703157816 | 455.21593919944587 | 501.037349994778 | 794.7468236535269 | 294.5188057100646 | 636.1914508662776 | 415.5238025566059 |
| SE_415_2_0 | SE_415 | SE_415_2 | GAGTAGCCAAAACCTGTTTT | 941.8020754271317 | 1255.2369607001735 | 1224.5333096668346 | 1185.4920339949908 | 1256.804497584347 | 1523.839483258531 | 1175.167389138995 | 1539.0354826254693 | 1386.14501868509 | 1447.0787854813818 | 1267.6877618737633 | 1561.115962119092 |
| SE_415_2_2 | SE_415 | SE_415_2 | AGAGGTGATTAACCTAAAAC | 634.0094523876548 | 904.4501985936761 | 929.2627777391059 | 722.2262117093522 | 881.354040078137 | 847.2512874513037 | 891.0609873691282 | 694.834815558796 | 753.011515595744 | 1147.456931157529 | 747.7010225125808 | 726.5863139676964 |
| SE_415_2_4 | SE_415 | SE_415_2 | TTTATCATTTATTTTGCCAG | 4.335107366753196 | 5.9963549078033775 | 3.950107450538176 | 3.9039254686992013 | 8.484756101835254 | 1.7326202197368175 | 3.228481838293942 | 8.508181415005666 | 0.0 | 22.599215289633673 | 0.0 | 0.0 |
| SE_415_2_7 | SE_415 | SE_415_2 | AGGAAATTTATGATAATGCT | 374.98678722415144 | 525.6804469174294 | 539.189666998461 | 459.36189681693935 | 506.9641770846564 | 557.9037107552552 | 489.6530788079146 | 654.18461546488 | 490.16787335949374 | 564.9803822408418 | 753.5699473360704 | 482.843301294827 |
| SE_415_2_8 | SE_415 | SE_415_2 | TGCTGTAGAAAATTAACAGT | 1610.4923867488121 | 2083.7333304616736 | 2069.8563040820045 | 2041.7530201296822 | 2021.4931412622493 | 1926.6736843473411 | 1713.2476955213187 | 1975.7887952624267 | 2233.282973730302 | 2439.9862443356096 | 1402.6730328140252 | 2274.934784946781 |
| SE_416_1_10 | SE_416 | SE_416_1 | ATCAAACTCAAACTATTAGG | 131.13699784428417 | 158.9034050567895 | 162.94193233469977 | 91.09159426964803 | 151.66501532030517 | 71.03742900920952 | 132.36775537005164 | 100.20746999895562 | 174.93309973155846 | 43.01140974478667 | 109.1620017169074 | 64.99813671276517 |
| SE_416_1_14 | SE_416 | SE_416_1 | TCCATGCCTAAAGCAGCCGG | 1076.1904037964807 | 1089.3378082509469 | 1078.379333996922 | 1115.2213755584053 | 1057.4127291912184 | 944.2780197565655 | 923.3458057520676 | 966.1512673484211 | 1030.950907555022 | 1202.8614589643728 | 812.2591955709669 | 1077.1119798115371 |
| SE_416_1_18 | SE_416 | SE_416_1 | TCCTTAATACAGAGACTAGA | 99.7074694353235 | 126.92284554850481 | 137.26623390620162 | 88.48897729051524 | 141.0590701930111 | 233.03741955460194 | 91.47365208499504 | 195.6881725451303 | 77.25471917078977 | 156.00748619295504 | 102.11929192871983 | 129.99627342553035 |
| SE_416_1_21 | SE_416 | SE_416_1 | AACATGTGAACACATTATGA | 352.22747354869716 | 359.78129446820265 | 366.37246603741585 | 391.6938553594865 | 398.78353678625695 | 282.41709581710126 | 488.57691819515 | 339.3819031096704 | 409.36121307740325 | 492.80869470297944 | 332.1811450095139 | 387.6674582511351 |
| SE_416_1_7 | SE_416 | SE_416_1 | ATGAAAAGCAGGGTAACAGG | 721.7953765644071 | 884.4623489009981 | 915.4374016622223 | 727.4314456676178 | 804.9912351616197 | 732.8983529486737 | 719.9514499395492 | 1031.3806581967979 | 805.402646987429 | 871.8923060129637 | 681.9690644894968 | 702.2120127004094 |

| | | | | | | | | | | | | | | | |
|---|---|---|---|---|---|---|---|---|---|---|---|---|---|---|---|
| SE_416_2_15 | SE_416 | SE_416_2 | AAGTGACACTCCCTGCACAG | 499.6211240183058 | 589.6415659339988 | 599.4288056191682 | 633.737234418837 | 639.5384911758323 | 740.6951439374894 | 513.3286122887368 | 599.3541130126213 | 595.8381214206889 | 761.8122573441028 | 509.4226746789012 | 718.4615468786008 |
| SE_416_2_23 | SE_416 | SE_416_2 | GCATCACAAGGTCAGAGGAG | 2511.1109421917886 | 2587.427142717157 | 2658.4223142121928 | 2520.6345442901174 | 2699.21303489634 | 1858.2351856677367 | 2310.516835605698 | 2136.498888656978 | 2664.8438187533347 | 1808.6662301155206 | 1493.0544750957658 | 1865.2143874538147 |
| SE_416_2_33 | SE_416 | SE_416_2 | TCCTTATCAGTCAGTGCCAG | 852.9323744086912 | 742.5486160829848 | 744.5952544264462 | 838.0426672807619 | 827.2637199289372 | 602.085526358544 | 770.5309987394876 | 817.7307693311001 | 934.1605122720786 | 500.09876415124836 | 973.0677357345832 | 1055.059040569706 |
| SE_416_2_4 | SE_416 | SE_416_2 | TACTGCAGCCTCAACCACCA | 2534.954032708931 | 582.6458185415614 | 989.5019163598131 | 2167.9799436781623 | 485.7522868300683 | 5107.764407784148 | 3219.8725533391825 | 1257.3201424397262 | 4146.891247443543 | 222.34711817220227 | 687.8379893129865 | 1471.74352413904 |
| SE_416_2_6 | SE_416 | SE_416_2 | GGTGGCCTAAAGTGACTGTG | 472.5267029760983 | 437.7339082696465 | 420.6864434823158 | 392.9951638490529 | 357.4203507898101 | 373.37965735532842 | 472.43450900368026 | 397.9938195241539 | 386.27359585394885 | 295.9768195997184 | 413.172307573671 | 694.0872456113137 |
| SE_416_3_24 | SE_416 | SE_416_3 | CCCATGGTCGAGGACAGTAC | 1495.6120415298526 | 1139.3074324826416 | 1153.4313755571475 | 1345.552978211658 | 1077.5640249330772 | 1089.818118214458 | 1177.3197103645243 | 1055.0144954607024 | 1090.4459211693086 | 991.449444964574 | 1408.5419576375148 | 1095.6828760151843 |
| SE_416_4_2 | SE_416 | SE_416_4 | GGCATGGAGTGTTCTGGGAG | 704.4549470973943 | 942.4271100097641 | 989.3771849861586 | 796.4007956146371 | 993.7770584274541 | 1077.6897766763004 | 718.8752893267845 | 864.9984438589092 | 1056.702480611952 | 839.8160004405803 | 984.8055853815624 | 1132.8246684224787 |
| SE_416_4_32 | SE_416 | SE_416_4 | ATTGACACATGAATGAGAGC | 476.8618103428515 | 408.75152621526354 | 484.87568955356113 | 331.8336648394321 | 510.14596062284465 | 623.7432791052543 | 574.6697672163217 | 430.13583820306417 | 508.8155641938223 | 780.0374309647751 | 419.0412323971607 | 461.95104306572387 |
| SE_416_4_33 | SE_416 | SE_416_4 | CATTGACACATGAATGAGAG | 2019.0762560653009 | 2108.718142577521 | 2158.7337217191134 | 2205.7178898150487 | 2031.0384918768139 | 2189.1656476374687 | 2210.4338986185858 | 1888.8162741312576 | 2378.024574015805 | 2366.3565429080936 | 1561.1340030482456 | 1992.8892988538892 |
| SE_416_4_8 | SE_416 | SE_416_4 | GCTCCTGAGTTCCAGTGGCA | 244.93356622155557 | 268.8365783665181 | 339.70924074628317 | 230.33160265325287 | 313.9359757679044 | 292.81281713552215 | 332.53362934427605 | 346.9447310341199 | 349.8661994631169 | 266.0875348618158 | 518.8129543964847 | 240.26096963468552 |
| SE_416_4_9 | SE_416 | SE_416_4 | CCTGCGCTCCTGAGTTCCAG | 4942.022398098643 | 4355.3524480345195 | 4242.415401878001 | 4356.780823068309 | 4423.739712594355 | 4121.9035027538885 | 5098.8489832789 | 4223.839395805035 | 4513.629167185338 | 4346.339405057934 | 4386.434413076161 | 4237.646377469743 |
| SE_417_1_11 | SE_417 | SE_417_1 | GCCCTATTTGAACCCCAAAT | 112.71279153558308 | 110.93256579436247 | 87.88989077447442 | 127.52823197750725 | 107.12004578567009 | 217.4438375769706 | 102.23525821264151 | 59.557269905039654 | 39.07135222430747 | 34.99233335169084 6 | 160.80854016361627 | 268.1173139401563 |
| SE_417_1_19 | SE_417 | SE_417_1 | TTGCTGCTCTTGGTGTCAGA | 423.7567451001249 | 534.6749792791345 | 482.900635828292 | 409.91217421341617 | 495.2976374446329 | 328.3315316401269 | 468.12986655262165 | 471.7313917875363 | 571.8625189194094 | 603.6177503166671 | 272.3181118099195 | 687.123159534946 |
| SE_417_1_20 | SE_417 | SE_417_1 | CAAAGCATATGCCTATTTGA | 285.0333093640226 | 284.8268581206604 | 284.40773643874866 | 312.3140374959361 | 338.32964956068076 | 292.81281713552215 | 286.2587229953962 | 254.30008895961376 | 267.2835686253761 | 324.40809044796725 | 370.91604884454557 | 289.0095721692594 |
| SE_417_1_46 | SE_417 | SE_417_1 | CCAAGGTTACCTAGATTAGG | 113.79656837727138 | 101.9380334326574 | 103.69032057662713 | 115.81645557140963 | 124.08955798934059 | 142.07485801841904 | 111.92070372752333 | 74.68292575393862 | 103.0062922277197 | 355.7553890755236 | 45.77761362321923 | 68.48017975094902 |
| SE_417_1_7 | SE_417 | SE_417_1 | TAGACCCCTCAAGTGCACAG | 1196.489633223882 | 955.4192153100048 | 1024.0653565520222 | 1193.2998849323892 | 1013.9283541693128 | 1135.7325540374839 | 1019.1241002881212 | 1359.418319419794 | 909.2969244929739 | 1097.1554519644735 | 1270.0353318031591 | 1022.5599722133235 |
| SE_417_2_13 | SE_417 | SE_417_2 | ATATCCCCCTACCTTTCATG | 190.74472413714062 | 214.86938419628768 | 186.64257703792882 | 148.34916781056964 | 204.6947409567755 | 343.92511361775826 | 189.40426784657797 | 164.4915073567762 | 265.50759806972576 | 101.33196533093808 | 159.63475519891836 | 142.76376456553777 |
| SE_417_2_15 | SE_417 | SE_417_2 | CGTATCCCCATGAAAGGTAG | 2862.2546388987976 | 2795.300779521008 | 2661.3848948000964 | 2816.0315714216904 | 2541.1844524996586 | 2397.9463841157553 | 2645.2027861755037 | 2465.4819033705303 | 2736.7706262571733 | 2392.6007929218617 | 2569.4152877237666 | 2538.4093748360256 |
| SE_417_2_24 | SE_417 | SE_417_2 | TTGTTCACTTTGGAACCCAC | 108.37768416882989 | 150.9082651797183 | 136.27870704356707 | 70.27065843496522 | 201.5129574185873 | 103.09090307434464 | 68.87427921693744 | 191.90675858290555 | 110.99815978214622 | 232.55312359977876 | 191.32694924576242 | 210.08326330375886 |
| SE_417_2_27 | SE_417 | SE_417_2 | CTAAGACTTGGGGAGAAGGA | 380.4056714325929 | 422.74302100013807 | 356.4971974110704 | 355.25721765162734 | 344.6932166370572 | 408.89837185788895 | 437.99736939521154 | 420.6823032975023 | 364.0739639083196 | 395.1217640961758 | 338.0500698330036 | 293.6522962201712 |
| SE_417_2_9 | SE_417 | SE_417_2 | GGTCTCTGCTTCATCAGGTG | 32.51330525064897 | 36.977521931454156 | 41.47612823065085 | 37.73794619742561 5 | 23.33307928004695 | 26.85561340592067 | 25.827854706351538 | 12.289595377230405 | 41.75530805778298 | 51.03048613788248 4 | 10.564064682281362 | 49.909283547301825 |
| SE_417_3_20 | SE_417 | SE_417_3 | CATACTAACAGACTATAGCA | 668.6903113216804 | 754.5413258985916 | 720.8946097232172 | 615.5189155649074 | 791.2035064961374 | 594.2887353697284 | 569.2889641524985 | 496.3105825419971 | 672.2048553136535 | 421.36601410994393 | 595.10897710185 | 1318.5336304589505 |
| SE_417_3_25 | SE_417 | SE_417_3 | GGGCCCTGTTCCAAATGCAG | 30.34575156727237 | 72.95565137827442 | 85.91483704920533 | 45.545797134824014 | 82.72637199289373 | 66.70587845986748 | 29.056336544645482 | 52.93979547114636 | 24.863587779104755 | 58.320555586151414 | 69.25331291717781 | 84.7297139291403 |
| SE_417_3_26 | SE_417 | SE_417_3 | TGTCCCCTGCATTTGGAACA | 198.3311620289587 | 242.85237376603678 | 263.66967232342324 | 219.92113473672168 | 302.2694361278809 | 126.48127604078768 | 164.65257375299106 | 263.7536238651756 | 258.40371584712443 | 134.13727784814824 | 225.36671322200237 | 365.61451900930405 |
| SE_417_3_29 | SE_417 | SE_417_3 | GGTAAGCTATGGGGGGAGAA | 282.86575568064603 | 406.75274124599576 | 430.5617121086612 | 366.96899940577249 | 361.6627288407277 | 260.759343070391 | 219.53676500398808 | 302.5131169779792 | 321.4506705727115 | 156.00748619295504 | 257.05890726884644 | 337.7581747038333 |
| SE_417_3_5 | SE_417 | SE_417_3 | ATACCTGCTGAGCAATTGAA | 128.96944416090756 | 217.86756165018937 | 254.78193055971235 | 131.43215744620645 | 170.75571654943448 | 90.09625142631451 | 125.91079169346375 | 58.61191641448347 | 113.66211556162173 | 46.65644446812913 | 66.90574298778195 | 53.39132658548567 |
| SE_418_1_16 | SE_418 | SE_418_1 | GCTCCCAGTTCTCCAGTGCA | 92.12103154350541 | 156.9046200875217 | 157.0167711588925 | 134.03477442533924 | 85.90815553108195 | 161.133680435524 | 59.18883370205561 | 104.93423745173654 | 161.6133205641809 | 19.683187510326103 | 118.55228143449084 | 40.62383544547823 |
| SE_418_1_21 | SE_418 | SE_418_1 | GTTTTACAGCATCTGGCACG | 136.55588205272565 | 161.90158251069118 | 156.02924429625796 | 111.91253010271043 | 167.57933310124625 | 152.47057933683993 | 166.80489497852037 | 90.57449833816732 | 149.44642368951298 | 143.20176569314734 | 107.94333418369929 |  |
| SE_418_1_3 | SE_418 | SE_418_1 | GATTTAATCACCATCTCAAA | 16.256652625324485 | 20.98724217731182 | 15.800429802152705 | 39.039254686699201 | 24.393673792776355 | 0.8663101098684087 | 6.456963676587884 | 17.961716320567515 | 57.71904305863604 | 13.851131951710961 | 22.30191432926065 | 0.0 |
| SE_418_1_33 | SE_418 | SE_418_1 | TGTGGAATAGGACTGTGTGC | 4539.941189832284 | 4129.489746507259 | 4023.1844383731323 | 4289.112781610856 | 3682.3841481965 | 3677.486416391395 | 4390.735300079762 | 3777.6325482625152 | 3799.6890038139013 | 4089.728960478868 | 3225.561082989909 | 4005.5101749241535 |
| SE_418_1_36 | SE_418 | SE_418_1 | GGTTCTTACTCGGCCAGCAG | 8.670214733506391 | 7.9951398770711695 | 1.9750537252690888 | 0.0 | 14.848323178211695 | 1.7326202197368175 | 2.152321225529295 | 3.78141396222474 | 0.0 | 0.0 | 18.780559435166865 | 1.1606810127279494 |
| SE_418_2_26 | SE_418 | SE_418_2 | TGTCCACAGAGTGCATCGGA | 913.623877543236 | 943.426505494398 | 977.6515940081986 | 952.5578143626051 | 1160.290396925971 | 756.2887259151208 | 1001.9055304838868 | 1109.8449979129612 | 986.5516436637637 | 828.15188932335 | 1074.0132426986052 | 712.6581418149609 |
| SE_418_2_27 | SE_418 | SE_418_2 | AAGCCATCCGATGCACTCTG | 2183.8103360019 22 | 2003.7819316909618 | 2029.367702713988 | 2108.1197530975687 | 2182.703507197119 | 2427.4009278512813 | 2033.9435581251837 | 2278.3019122404057 | 2126.7247403912816 | 2060.9026330256256 | 1976.6538805513126 | 1809.501698842873 |
| SE_418_2_29 | SE_418 | SE_418_2 | GGATGGCTTCTGGAAATCTG | 603.6637008203825 | 703.5723091822629 | 761.3832110912334 | 674.0777975953954 | 652.2656253285852 | 737.2299034980158 | 602.6499431482026 | 683.4905736721217 | 740.5797217061915 | 1070.9112019507054 | 762.9602270536539 | 501.41419749847415 |
| SE_418_2_40 | SE_418 | SE_418_2 | CGGCCTCCTGGATGAAATAT | 26.010644200519174 | 52.9678016855965 | 48.38881626909266 | 62.46280749918722 | 77.42339942924669 | 18.192512307236584 | 21.52321225529295 | 105.87959094229272 | 39.07135222430747 | 86.75182643440023 | 102.11929192871983 | 16.249534178191293 |
| SE_418_2_41 | SE_418 | SE_418_2 | GCGGCCTCCTGGATGAAATA | 190.74472413714062 | 268.8365783665181 | 261.69461859815 42 | 279.78132552567761 | 253.48208852423282 | 228.70586900552599 | 247.51694093586892 | 259.97220990295085 | 288.59521529318016 | 139.96933340676338 | 268.79675691582577 | 148.56716962917753 |
| SE_419_1_10 | SE_419 | SE_419_1 | TTTCAGTGTGGAGAAGAGGG | 558.1450734694739 | 718.5631964517713 | 594.4911713059955 | 638.9424683771026 | 654.386814354044 | 716.438460861174 | 545.6134306716763 | 497.25593603255334 | 729.0359130944644 | 640.0680975580118 | 564.5905680197038 | 680.1590734585784 |
| SE_419_1_12 | SE_419 | SE_419_1 | ATTGGGATGCTAGCTTGCCA | 1220.3327237410247 | 1074.3469209814384 | 1083.3169683100948 | 1125.6318434749364 | 1082.8669974967243 | 912.2245456914344 | 1295.6973777686355 | 1007.7468209328932 | 887.0972925473446 | 880.6403893508864 | 1171.437394768533 | 1038.8095063915148 |
| SE_419_1_14 | SE_419 | SE_419_1 | CCTGAGGCTTGATTGCTACA | 1180.2329805985576 | 1175.285561929462 | 1159.3565367329547 | 1095.7017482149092 | 1221.8048786642767 | 1065.5614351381428 | 1109.5215917603516 | 1461.516496399862 | 1080.6780831132317 | 1144.5409033782214 | 1537.658303754287 | 821.7621570113882 |
| SE_419_1_16 | SE_419 | SE_419_1 | GAGCTCCCAGGACTAGGGGA | 166.90163361999802 | 193.88214201897586 | 272.55741408713413 | 182.18318853929605 | 226.96722572409305 | 72.77004922894633 | 120.52998862964051 | 162.60080037566382 | 212.22848140021557 | 14.580138896537854 | 48.12518355261509 | 22.05293924183104 |
| SE_419_1_3 | SE_419 | SE_419_1 | GGGTCTGCCTTATTTTATAG | 4.335107366753196 | 22.986027146579612 | 32.58838646693995 | 16.917010364363207 | 22.272484767317543 | 0.8663101098684087 | 20.44705164325283 | 12.289595377230405 | 23.087617223454416 | 23.328222234460565 | 26.997054188052367 | 0.0 |
| SE_419_2_0 | SE_419 | SE_419_2 | CCAAAAGTATTATTCTCACA | 311.04395356454177 | 452.72479553915497 | 458.21246426242845 | 420.3226421299473 | 493.17644841917416 | 540.5775085578871 | 383.1131781442145 | 396.1031125430415 | 455.5364475243121 | 362.31645157896565 | 385.00146842092073 | 347.04362280565687 |
| SE_419_2_10 | SE_419 | SE_419_2 | ATGGTGGCCGAAGAGAACTG | 3030.2400493604837 | 2809.292274305286 | 2955.66789986519 | 3008.625227875718 | 2851.9386447293746 | 2809.4436630324997 | 3103.6472072132433 | 3244.453179588827 | 3020.0379298834023 | 3019.5465674529786 | 2979.0662404033437 | 3413.5628584328992 |
| SE_419_2_7 | SE_419 | SE_419_2 | ATAATGACTATAGCAGACAG | 110.54523785220648 | 120.92649064070145 | 103.69032057662713 | 117.11776406097604 | 126.21074701479941 | 129.0802063703929 | 116.22534617858193 | 101.15282348951179 | 84.35860139339113 | 182.98074315155006 | 83.33873249355297 | 73.12290380186082 |
| SE_419_2_8 | SE_419 | SE_419_2 | GGAAATACGGAGTTTCTAGT | 30.34575156727237 | 34.97873696218637 | 35.55096705484358 | 31.23140374959361 | 30.757240869152795 | 33.78609428486794 | 9.685445514881827 | 10.398888396118036 | 53.27911666951019 | 37.908361130998415 | 15.259204541073078 | 61.51609367458132 |
| SE_419_2_9 | SE_419 | SE_419_2 | CACTACCTAACATATGACCA | 1232.2542689995958 | 1163.2928521138551 | 1156.393956145051 | 1242.7496075359124 | 1138.0179121586534 | 957.2726714045916 | 1107.3692705348224 | 1103.227523479068 | 1304.4503731251743 | 1292.5293131780807 | 1227.7790730740337 | 1462.4580760372162 |
| SE_419_3_3 | SE_419 | SE_419_3 | CAGATGCATGACAAATACAG | 1989.814281339717 | 1784.9149775561386 | 1845.6877062639628 | 1897.3077777878118 | 1901.6459613238262 | 1727.422359077607 | 2096.3608736655333 | 2109.0836374308487 | 1580.6137945288021 | 2148.3834664048527 | 1989.5655151629896 | 1817.6264659319688 |
| SE_419_3_4 | SE_419 | SE_419_3 | AGTCCCAACACACCACCCAG | 378.2381177492163 | 379.76914416088056 | 400.9359062296249 | 377.3794619742561 | 424.2378050917627 | 613.3475577868334 | 317.4673807655751 | 489.6931608108384 | 342.76231724051553 | 342.63320460863956 | 328.65979011542015 | 489.80738737119464 |
| SE_419_3_5 | SE_419 | SE_419_3 | GATTTCTGACTTAGCTAACA | 4.335107366753196 | 25.984204600481302 | 25.675698428498144 | 6.506542447832002 | 5.3029725636470335 | 35.51871450460476 | 6.456963676587884 | 0.945353490556185 | 5.327911666951019 | 13.851131951710961 | 0.0 | 10.446129114551544 |
| SE_419_3_6 | SE_419 | SE_419_3 | AATGGTGGAGAGAATTCCAA | 107.2939073271416 | 154.90583511825392 | 138.25376076883617 | 144.44524234187045 | 132.57431409117584 | 136.8769973592086 | 168.95721620404964 | 127.62272122508497 | 129.64585056247478 | 87.48083337922712 | 169.0250349165018 | 293.6522962201712 |
| SE_419_3_9 | SE_419 | SE_419_3 | ATTACTTCCTAAATACAGAA | 0.0 | 0.9993924846338962 | 3.950107450538176 | 6.506542447832002 | 1.0605945127294067 | 0.0 | 0.0 | 31.196665188354107 | 0.0 | 0.7290069448268927 | 4.695139858791716 | 0.0 |
| SE_42_1_10 | SE_42 | SE_42_1 | ACCGCGATGAGAGGCGCTCG | 1909.6147950547827 | 2287.6093973269885 | 2150.8335068180368 | 1901.211703256511 | 2208.1577550626246 | 2215.154950933521 | 1940.3175848146593 | 2566.634726860042 | 2036.1502420531144 | 2274.5016675995 | 2425.0397370659216 | 2473.4112381232603 |
| SE_42_1_11 | SE_42 | SE_42_1 | GCCGCGAGCGCCTCTCATCG | 469.27537245103343 | 509.6901671632871 | 612.2666548334173 | 553.0561080657202 | 559.9939027211268 | 355.1871450460476 | 510.1001304504429 | 432.97189867473276 | 494.6077997486196 | 1069.4531880610516 | 573.9808477372873 | 600.0720835803498 |
| SE_42_1_14 | SE_42 | SE_42_1 | CAGAACGCAGAGAAAATCCC | 2168.637460218286 | 2340.577199012585 | 2345.376298757042 | 2119.8315295036664 | 2201.7942084262486 | 2025.4330368723397 | 2116.8079253080614 | 2060.870609412483 | 2132.940637336058 | 2285.4367720323085 | 1905.0529977047388 | 1640.0422709845925 |
| SE_42_1_7 | SE_42 | SE_42_1 | CTCATCGCGGTAGCTGCGAT | 331.63571355661946 | 333.7970898677213 | 371.3101003505886 | 381.2833874429553 | 313.9359757679044 | 317.935810320 77 | 440.1496906207408 4 | 516.1630058436771 | 407.58524252175295 | 371.79354186171525 | 460.1237061615882 | 312.2231924238184 |
| SE_42_1_9 | SE_42 | SE_42_1 | ATCGCAGCTACCGCGATGAG | 1991.9818350230935 | 1643.0012447381253 | 1794.3363094069666 | 1909.0195541939095 | 1657.7092233960627 | 1966.5239494012878 | 1985.5163305507747 | 1828.313650735662 | 1582.3897650844526 | 1638.8076119708546 | 1593.9999820597877 | 1842.0007671992557 |
| SE_42_2_0 | SE_42 | SE_42_2 | CCAGAACTAGATGCTCATGG | 932.048083851937 | 1053.3596788041266 | 1008.2649267498695 | 1005.9114624348275 | 1019.2313267329599 | 927.8181276690658 | 927.6504482031261 | 996.402579046219 | 1025.622995888071 | 1301.2773965160034 | 1338.1148597556391 | 1439.2444557826573 |
| SE_42_2_12 | SE_42 | SE_42_2 | ATACCACCTGCTAAGTGCAG | 74.78060207649263 | 51.968409200962604 | 40.488601368016305 | 72.87327541571842 | 58.33269820011737 | 67.57218856973589 | 111.92070372752333 | 36.868786131691216 | 63.93494000341222 | 84.56480559991955 | 29.344624117448227 | 69.64086076367697 |
| SE_42_2_5 | SE_42 | SE_42_2 | AATCGCCCACAACACCCATG | 1282.1080037172576 | 1259.2345306387092 | 1231.4459977052763 | 1289.596713160303 | 1333.1673025008643 | 1763.80738369208 | 1510.929500321565 | 1112.6810583846298 | 1171.252581451399 | 2610.5738694251027 | 1495.4020450251617 | 1537.902341864533 |
| SE_42_2_7 | SE_42 | SE_42_2 | GAGCTCCCCATAGTAGAACA | 81.28326312662242 | 135.9173779102099 | 119.49075037877982 | 84.58505182181602 | 154.84679885849337 | 83.16577054736723 | 74.25508228706067 | 120.0598933006355 | 68.37486639253807 | 240.57229179287658 | 132.63770101086598 | 167.1380658328247 |
| SE_42_2_9 | SE_42 | SE_42_2 | GGAACTAGACAAAATAATGG | 93.2048083851937 | 168.89732990312845 | 166.89203978523796 | 175.67664609146405 | 151.66501532030517 | 124.74865582105086 | 109.76838250199404 | 124.78666075341643 | 261.0676716805999 | 252.23640291010486 | 68.07952795247988 | 356.32907090748046 |
| SE_42_3_10 | SE_42 | SE_42_3 | AGATTAAAAGGCACCGAAGG | 89.95347786012881 | 115.92952821753195 | 114.55311606560711 | 62.46280749918722 | 158.02858293966816 | 47.64706504276248 | 115.14918556581728 | 139.91231660231537 | 52.39113139168501 5 | 96.95792366197672 | 57.515463270198524 | 6.964086076367696 |
| SE_42_3_11 | SE_42 | SE_42_3 | GAATGTGTAAGTGCCTCCTT | 473.6104798177866 | 600.6348832649716 | 553.0150430753447 | 636.3398513979698 | 562.1150917465856 | 835.122945913146 | 497.18620309726714 | 691.9987550871274 | 479.5120500255917 | 536.549111392593 | 527.0294491493702 | 461.95104306572387 |
| SE_42_3_13 | SE_42 | SE_42_3 | GGCAGCATTGACGTCAGAAT | 3321.776019774636 | 3319.9818339538033 | 3111.6971441614483 | 3267.5856173012317 | 3241.176830901067 | 3485.1655720006083 | 3254.309693000294 | 3147.08177006154 | 3420.519290182554 | 2966.3292585006 26 | 3853.536039103301 | 3464.632822992929 |
| SE_42_3_8 | SE_42 | SE_42_3 | CTTCATCTAACGAGATTAAA | 16.256652625324485 | 15.990279754142339 | 25.675698428498144 | 18.218318853929606 | 28.63605184369398 | 3.465240439473635 | 1.0761606127646475 | 8.508181415005666 | 22.199631945629246 | 2.1870208344806 78 | 10.564064682281362 | 0.0 |
| SE_42_3_9 | SE_42 | SE_42_3 | ACGAGATTAAAAGGCACCGA | 5906.583787201229 | 7193.6271043974845 | 7576.3060901322215 | 6775.913305172247 | 7499.4637995096355 | 6454.010318519645 | 6324.595921217833 | 7140.254914170865 | 7539.8829940135165 | 7773.401052689156 | 6816.169290000874 | 7335.50400044064 |
| SE_42_4_10 | SE_42 | SE_42_4 | AATTTGGTCAAGATGTGAAG | 17.340429467012783 | 11.992709815606755 | 20.738064115325425 | 11.711776406097604 | 15.9089176978411 | 0.0 | 7.533124289352532 | 12.289595377230405 | 0.8879852778251698 | 0.0 | 0.0 | 6.964086076367696 |
| SE_42_4_11 | SE_42 | SE_42_4 | TGGCTAGGAGTGGGGAAGGT | 110.54523785220648 | 160.9021900260573 | 151.09160998308525 | 85.88636031138243 | 147.42263726938754 | 152.47057933683993 | 67.79811860417279 | 173.94504226233803 | 133.19779167377547 | 183.70975009637695 | 116.20471150509498 | 37.14179240729438 |
| SE_42_4_13 | SE_42 | SE_42_4 | CCTGGGGTAAACCCAAATGA | 308.8763998811652 | 396.7588163996568 | 393.0356913285485 | 342.24413275596334 | 452.87385693546767 | 617.6791083361754 | 322.8481838293942 6 | 408.4732277995781 | 333.8851807307168 5 | 411.9985226089731 | 698.7299696622256 |  |
| SE_42_4_21 | SE_42 | SE_42_4 | AGAGTAGCGGAAGATGAGAT | 1480.4391657462163 | 1672.9830192771421 | 1549.4296474735995 | 1608.417293104071 | 1649.2244672942275 | 1492.6523193032683 | 1770.284207997845 | 1636.4068921527562 | 1511.3509428584389 | 1555.7008202060589 | 1782.9793613761542 | 1592.4543494627467 |
| SE_42_4_4 | SE_42 | SE_42_4 | GCCAATTGAAAGGAAGATAG | 453.01871982570896 | 522.6822694635277 | 474.01289406458113 | 442.44488645257616 | 536.6608234410799 | 483.4010413065721 | 455.21593919944587 | 513.3269453720085 | 388.9375516874244 | 264.629520972162 | 453.0809963734006 | 302.9377443219948 |
| SE_42_5_10 | SE_42 | SE_42_5 | CTAGAATTACTGAAAGGAAG | 87.78592417675222 | 99.93924846338962 | 105.66537430189621 | 131.43215744620645 | 145.30094832878972 | 129.94651648026132 | 90.39749147223039 | 134.2401956589782 8 | 186.47690834328566 | 76.54572920682372 | 181.936669528179 | 177.58419494737626 |
| SE_42_5_11 | SE_42 | SE_42_5 | TTAGCTTCCCTAAGGCAAAG | 212.42026097090658 | 222.86452407335884 | 245.89418879600146 | 230.33160265325287 | 214.24009157134017 | 250.36362175197013 | 260.43086828904467 | 217.43130282792256 | 206.90056973326455 | 179.33570842741 56 | 190.1531642810645 | 263.4745898892445 |
| SE_42_5_12 | SE_42 | SE_42_5 | GAGAATGCTTTGAAACCGGG | 186.4096167703874 | 228.86087898116222 | 240.95655448282875 | 257.6590809341473 | 221.664253160446 | 356.05345551591597 | 264.7355107401033 | 256.1907959407261 6 | 327.66656751748764 | 748.6901323372188 | 251.1899824453568 | 393.4708633147749 |
| SE_42_5_13 | SE_42 | SE_42_5 | TTTCTAGGAAAACAAACAGT | 0.0 | 0.0 | 0.0 | 0.0 | 0.0 | 10.395721318420904 | 2.152321225529295 | 0.945353490556185 | 0.0 | 0.0 | 0.0 | 0.0 |
| SE_42_5_9 | SE_42 | SE_42_5 | GGATGACTAGAATTACTGAA | 195.07983150389381 | 365.777649376006 | 320.94623052622 68 | 338.34020727876 41 | 331.96608284843043 | 229.57217911512 83 | 237.8314954209871 | 243.90120056349573 | 236.20408390149515 | 245.67354040666283 | 193.67451917515828 | 395.79222534023074 |
| SE_420_1_0 | SE_420 | SE_420_1 | GTGAGCTGATTTGCTAGTGA | 682.7794102636283 | 702.572916697629 | 710.0318142342371 | 762.5667748859106 | 666.0533539940675 | 469.5400795486775 | 778.0641230288401 | 801.6597599916449 | 718.3800897605623 | 653.1902225648959 | 657.3195802308403 | 777.6562785277262 |
| SE_420_1_18 | SE_420 | SE_420_1 | CTCTGCAAAATGAGGACACC | 212.42026097090658 | 193.88214201897586 | 155.0417174336234 | 195.19627343496006 | 134.69550311663465 | 110.88769406315632 | 189.40426784657797 | 186.23463763956843 | 226.43624584541828 | 63.42360419993966 | 271.1443268452216 | 124.19286836189059 |
| SE_420_1_2 | SE_420 | SE_420_1 | GCAAATCAGCTCACCAAGTT | 1353.6372752686852 | 1639.0036747995898 | 1732.1221170609901 | 1426.2341064754775 | 1752.10213502898 | 1410.3528588657693 | 1408.6942421089236 | 1986.1876836585448 | 1402.1287536859431 | 1448.5367993718506 | 1961.3946760102394 | 1681.8267874427988 |
| SE_420_1_24 | SE_420 | SE_420_1 | ATGACGCATCATGTCTTCCC | 8585.680139854703 | 6444.082740919363 | 6201.668697344937 | 7783.126076096641 | 6397.506100783781 | 6299.807118963068 | 8306.883769930313 | 6792.36482964619 | 6797.527301751675 | 5320.292683346663 | 7144.829080116294 | 6123.753023152662 |
| SE_420_1_4 | SE_420 | SE_420_1 | TCCTGACCGGGAATCGAACC | 717.4602691976539 | 694.5777768205578 | 709.0442873716026 | 481.4841411395682 | 551.5091466192915 | 406.2994415282837 | 506.8716486612149 | 622.9879502765259 | 562.9826661411577 | 616.0108683787242 | 465.9926309850778 | 410.8810785056941 |

| | | | | | | | | | | | | | | | |
|---|---|---|---|---|---|---|---|---|---|---|---|---|---|---|---|
| SE_420_2_16 | SE_420 | SE_420_2 | GCAGCTGCAGAAAAATGTAT | 243.84978937986725 | 77.9526138014439 | 205.40558742798515 | 231.63291114281927 | 75.30221040378788 | 846.3849773414353 | 340.0667536336286 | 200.41493999791123 | 764.5553242074711 | 26.244250013768134 | 117.3784964697929 | 230.97552153286193 |
| SE_420_2_18 | SE_420 | SE_420_2 | TTTGTTTTGCCTATCTGGGA | 31.429528408960667 | 18.98845720804403 | 14.81290293951816 | 15.615701874796805 | 9.545350614564661 | 12.128341538157722 | 41.970263897821255 | 26.46989773557318 | 2.6639558334755096 | 5.832055558615141 | 4.695139858791716 | 0.0 |
| SE_420_2_6 | SE_420 | SE_420_2 | GGGGACTTCTGACCTAGTTA | 104.0425768020767 | 166.89854493386065 | 160.96687860943067 | 167.86879515406565 | 179.24047265126973 | 206.1818061486813 | 102.23525821264151 | 205.14170745069214 | 69.26285167036325 | 237.656264013567 | 197.19587406925208 | 109.10401519642724 |
| SE_420_2_7 | SE_420 | SE_420_2 | ACTAGGTCAGAAGTCCCCAC | 657.8525429047975 | 671.5917496739783 | 620.1668697344936 | 728.7327541571842 | 545.145579542915 | 803.0694718480149 | 714.570646875726 | 437.69866612751366 | 762.7793536518209 | 1053.41503527486 | 495.33725510252606 | 589.6259544657983 |
| SE_420_2_8 | SE_420 | SE_420_2 | CCTGTCTTAGAGCACCTGAA | 526.7155450605133 | 514.6871295864565 | 531.2894520973847 | 510.112927910029 | 580.1451984629855 | 259.89303296052265 | 499.33852432279645 | 469.84068480642395 | 488.3919028038434 | 626.2169656063008 | 373.26361877394146 | 370.25724306021584 |
| SE_421_1_12 | SE_421 | SE_421_1 | GTTAGCCACAAGCTCAGCGC | 809.5813007411593 | 723.5601588749408 | 781.1337483439244 | 784.6890192085394 | 853.7785827471724 | 1056.0320239295902 | 604.8022643737319 | 720.3593598038129 | 825.8263083774079 | 272.64859736525784 | 762.9602270536539 | 1272.1063899498326 |
| SE_421_1_17 | SE_421 | SE_421_1 | ACTCAAGGGAGGAGAAAGGA | 274.19554094713965 | 363.7788644067382 | 319.9587034935923 | 311.0127290063697 | 323.48132638246904 | 197.51870504999718 | 265.8116713528679 | 225.93948424292822 | 368.51389029744547 | 521.2399655512282 | 334.5287149389098 | 369.0965620474879 |
| SE_421_1_25 | SE_421 | SE_421_1 | TGAGTCAAAATAAAGATGGT | 1156.389890081415 | 1349.17985425576 | 1249.2214812326981 | 1181.5881085262915 | 1269.5316317370998 | 1428.545371173006 | 1221.4422954878748 | 1400.06851951371 | 1234.299536176986 | 1571.0099661019538 | 1584.6097023422042 | 1143.2707975370301 |
| SE_421_1_29 | SE_421 | SE_421_1 | GAGTTGCTGGTGTTCTATAC | 466.0240419259685 | 483.7059625628058 | 532.2769789600193 | 458.06058832737295 | 446.5102898590802 | 427.9571942749939 | 376.6562144676266 | 430.13583820306417 | 340.09836140704004 | 371.0645349168836 | 260.5802621629402 4 | 271.59935697834015 |
| SE_421_1_30 | SE_421 | SE_421_1 | CTGGTGTTCTATACTGGAAG | 92.12103154350541 | 83.94896870924728 | 115.54064292824165 | 59.86019052005442 | 94.3929116329172 | 41.582885273683615 | 76.40740350628997 | 100.20746999895562 | 89.68651306034215 | 75.81672226199683 | 137.3328408696577 | 66.15881772549312 |
| SE_421_2_23 | SE_421 | SE_421_2 | GTACCCCCTCCTTTTAGGTG | 14.089098941947885 | 13.991494784874547 | 11.850322351614528 | 6.506542447832002 | 1.0605945127294067 | 29.454543735525895 | 9.685445514881827 | 11.34424188667422 | 7.991867500426528 | 56.86254169649763 | 0.0 | 1.1606810127279494 |
| SE_421_2_26 | SE_421 | SE_421_2 | GCTGACCCCACCTAAAAGGA | 664.3552039549272 | 855.4799668466152 | 881.8614883326478 | 797.7021041042035 | 957.7168449946543 | 791.8074404197256 | 671.5242223651401 | 780.8619831994088 | 1117.0854795040636 | 802.6366462544088 | 819.3019053591545 | 1291.8379671662078 |
| SE_421_2_30 | SE_421 | SE_421_2 | AGGTGGGGTCAGCAGACACA | 1846.7557382368614 | 1590.0334430525288 | 1631.3943770722667 | 2045.6569455983815 | 1614.224848374157 | 1575.8180898506355 | 2099.5893555038274 | 1599.538106021065 | 1457.1838409111035 | 1381.4681604469615 | 2099.901301844595 | 1938.3372912556756 |
| SE_421_2_31 | SE_421 | SE_421_2 | GGTGGGGTCAGCAGACACAG | 740.2195828731082 | 944.4258979790319 | 960.8636373434114 | 897.9028578008163 | 924.8384151000427 | 642.8021015223593 | 766.226356288429 | 726.9768342377063 | 860.4577342125896 | 965.2051949508059 | 666.7098599484237 | 597.750721554894 |
| SE_421_2_32 | SE_421 | SE_421_2 | GCAGACACAGGGGCGCCATG | 1529.2091236221897 | 1979.7965120597482 | 2055.0434011424863 | 1672.1814090928247 | 2220.8849096553777 | 1836.5774329210265 | 1329.0583567643396 | 2054.25313497859 | 2408.2160734618606 | 1783.1509870465795 | 2000.129579845271 | 1717.8078988373652 |
| SE_421_3_22 | SE_421 | SE_421_3 | TTACTAACAGCCAACGGGCA | 200.4987157123353 | 201.87728189604704 | 198.49289938954334 | 203.00412437235846 | 156.9679878839522 | 224.37431845591786 | 238.90765603375175 | 287.38746112908024 | 202.4606433441387 | 299.62185432385286 | 214.80264853972102 | 183.38760001101602 |
| SE_421_3_32 | SE_421 | SE_421_3 | TTAGTAAGGGATATCCTGGG | 632.9256755459666 | 766.5340357141984 | 696.2064381573535 | 559.5626505135522 | 672.4169210704439 | 531.914407459203 | 544.5372700589116 | 603.135526974846 | 711.276207537961 | 354.29737518586984 | 508.24888971420324 | 614.0002557330853 |
| SE_421_3_4 | SE_421 | SE_421_3 | CTTGTGTTGAGATTTCTTGT | 359.81391144051526 | 409.75091869989745 | 412.7862285812394 | 406.0082487447169 | 509.0853661101152 | 551.8395399861763 | 385.2654993697438 | 465.11391735364305 | 417.3530805778298 | 326.59511128244793 | 436.6480068676296 | 379.5426911620395 |
| SE_421_3_8 | SE_421 | SE_421_3 | AGACCATCCCAAATTCCTGT | 1153.13855955635 | 1122.3177602438655 | 1193.9199769251638 | 1155.5619387349636 | 1185.7446652314768 | 1090.6844283243265 | 1246.1939895814617 | 1187.3639841385684 | 1222.755727565259 | 1288.8842784539463 | 1041.147263687063 | 1305.766139318943 |
| SE_421_3_9 | SE_421 | SE_421_3 | TGTTGGAAGGAAGAAGCAGA | 502.8724545433707 | 548.666474064009 | 512.5264417073283 | 511.4142363995954 | 459.2374240118331 | 423.62564372565186 | 406.78871162503674 | 417.84624282583377 | 545.2229605846543 | 247.8623612411435 | 362.69955409166005 | 529.270541803945 |
| SE_422_1_13 | SE_422 | SE_422_1 | GCAGGACAGGATGGCAGGCG | 15884.917168625398 | 16133.192879444987 | 16179.64011740437 | 16167.45667437296 | 15435.892538263786 | 14386.811994584663 | 16316.747210737585 | 16283.713874830286 | 15415.424423044948 | 14504.322174275856 | 16775.7347154628 | 15381.344780670786 |
| SE_422_1_23 | SE_422 | SE_422_1 | GCAGTCCTGGCTGAGCCCAG | 882.1943491342753 | 997.3936996646283 | 909.5122404864151 | 884.8897729051523 | 988.4740858638071 | 1147.8608955756415 | 964.2399090371241 | 1344.292663570895 | 1268.0429767343423 | 933.1288893784226 | 1074.0132426986052 | 934.3482152459993 |
| SE_422_1_31 | SE_422 | SE_422_1 | GGTGCAGCCTGGAGCTTCAG | 346.80858934025565 | 544.6689041254734 | 582.640848954381 | 395.5977808281857 | 491.05525939371535 | 542.3101287776238 | 337.9144324080993 | 451.87896848585643 | 301.02700918273257 | 387.10268770308 | 389.69660827971245 | 352.84702786929665 |
| SE_422_1_36 | SE_422 | SE_422_1 | TCAGGGGTGTGACGCTTTGG | 19804.938005011973 | 18591.69839164437 | 19193.572102165 | 17503.900493157653 | 17988.743530403466 | 17585.22892021883 | 18347.462287024475 | 16932.22636935183 | 17575.00461871576 | 19609.557808898586 | 15505.699383659643 | 17792.079244106735 |
| SE_422_1_41 | SE_422 | SE_422_1 | AGCCTCTTGGAAACACCCTC | 79.11570944324582 | 126.92284554850481 | 106.65290116453076 | 74.17458390528482 | 121.96836896388177 | 14.727271867762948 | 81.7882065701132 | 49.15838150892162 | 116.32607139509724 | 87.48083337922712 | 56.341678305500594 | 20.89225822910309 |
| SE_422_2_19 | SE_422 | SE_422_2 | TCAGAAAGCTGCAGACTGGT | 395.5785472162291 | 85.94775367851507 | 227.13117840594512 | 374.7768449951233 | 89.08993906927017 | 917.4224063506448 | 584.3552127312036 | 247.68261452572048 | 775.2111475413732 | 40.82438891030599 | 181.936669528179 | 133.47831646371418 |
| SE_422_2_38 | SE_422 | SE_422_2 | GCAGGAAACAGGCTCACAAG | 7291.650590878875 | 7657.345217264912 | 7388.6759862316585 | 7224.864734072656 | 7468.706558640482 | 6830.855216312403 | 7110.193168536026 | 8430.662428780059 | 7506.13955345616 | 6951.081218924422 | 7859.664123617333 | 7334.343319427912 |
| SE_422_2_4 | SE_422 | SE_422_2 | TTAGGTGATGGAGCCAAACG | 1299.4484331842705 | 1296.2120525701635 | 1297.610297501791 | 1228.435214150682 | 1233.4714183043 | 1051.7004733802482 | 1169.7865860751717 | 1541.8715430971376 | 1086.8939800580079 | 1373.4490840538658 | 1453.1457862960362 | 1109.6110481679198 |
| SE_422_2_44 | SE_422 | SE_422_2 | CTGACAGGCCCCCCAGACCG | 355.47880407376203 | 467.7156828086634 | 524.3767640589429 | 413.81609968211535 | 436.96493924451556 | 419.29409317630984 | 348.67603853574576 | 395.15775905248535 | 498.15974085992025 | 422.0950210547708 | 319.2695103978367 | 732.3897190313361 |
| SE_422_2_7 | SE_422 | SE_422_2 | CGTTTGGCTCCATCACCTAA | 1651.6759067329676 | 1940.8202051590265 | 1873.33845841773 | 1832.2423533094918 | 2161.491616942531 | 2457.7217816966754 | 1686.3436802022027 | 1861.4010229051282 | 2030.8223303861632 | 2197.9559386530814 | 1743.0706725764246 | 2367.789265965017 |
| SE_422_3_38 | SE_422 | SE_422_3 | GGGGGAAGACGGGGACCGTA | 549.4748587359676 | 695.5771693051918 | 545.1148281742683 | 514.0168533787282 | 618.3266009212441 | 645.4010318519645 | 518.7094153525601 | 619.2065363143012 | 501.7116819712209 | 290.14476404110326 | 714.8350435010387 | 509.5389645875698 |
| SE_422_3_43 | SE_422 | SE_422_3 | GTCTTCCCCCATCCACGAAG | 107.2939073271416 | 135.9173779102099 | 141.2163413567398 | 156.15701874796804 | 164.39214947305805 | 132.54544680986655 | 179.71882233169612 | 110.60635839507364 | 243.3079661240965 | 104.24799311024564 | 104.46686185811568 | 255.34982280014887 |
| SE_422_3_48 | SE_422 | SE_422_3 | GCTTCTCCTCATCCTCTTCG | 17.340429467012783 | 0.0 | 3.950107450538176 | 19.519627343496005 | 1.0605945127294067 | 0.0 | 5.380803063823238 | 0.0 | 0.8879852778251698 | 0.0 | 0.0 | 0.0 |
| SE_422_3_49 | SE_422 | SE_422_3 | GGATGAGGAGAAGCTCTCCG | 472.5267029760983 | 701.5735242129952 | 674.4808471793935 | 473.67629020216975 | 772.1128052670081 | 670.5240250381484 | 504.7193273866197 | 682.5452201815656 | 830.2662347665338 | 669.9573822959144 | 811.085410606269 | 551.323481045776 |
| SE_422_3_50 | SE_422 | SE_422_3 | GGAGAAGCTCTCCGAGGACG | 1127.127915355831 | 1404.146440910624 | 1414.138467292667 | 1212.8195122758852 | 1441.3479427992638 | 1283.8715828249817 | 1162.2534617858194 | 1238.4130726286023 | 1234.299536176986 | 1673.7999453225455 | 993.022080134448 | 1127.021263358839 |
| SE_422_4_15 | SE_422 | SE_422_4 | TCCTCAGCTGAACTTTGACC | 913.623877543236 | 894.456273747337 | 900.6244987227042 | 975.9813671748003 | 936.5049547400662 | 787.4758898703835 | 817.8820657011321 | 710.9058248982511 | 964.3520117181343 | 804.8236670888895 | 693.706914136476 | 542.0380329439524 |
| SE_422_4_16 | SE_422 | SE_422_4 | TCCAGGTCAAAGTTCAGCTG | 1918.285009788289 | 2067.743050707531 | 2045.1681325161408 | 1919.4300221104406 | 1852.8586137382736 | 2216.0212610433896 | 1906.9566058189553 | 1997.5319255452189 | 1971.327316771877 | 2130.1582927841805 | 1798.2385659172273 | 1959.2295494847785 |
| SE_422_4_19 | SE_422 | SE_422_4 | CTGAACTTTGACCTGGACCG | 295.8710777809056 | 424.7418059694059 | 440.4369807350066 | 314.9166544750689 | 401.96532032444514 | 385.5079988914419 | 293.79184728474877 | 202.30564697902358 | 342.76231724051553 | 371.0645349168836 | 273.4918967746175 | 263.4745898892445 |
| SE_422_4_21 | SE_422 | SE_422_4 | TGACTACTGGAAACACGCCC | 2243.418062294779 | 2054.7509484072907 | 2112.3199591752896 | 2195.3074218985175 | 2197.5518303753306 | 2557.3474443315426 | 2042.552843027301 | 1990.9144511113257 | 2236.8349148416028 | 1679.6320008811606 | 1518.8777443191202 | 1778.1633114992185 |
| SE_422_4_28 | SE_422 | SE_422_4 | GCTGTGGTGGACGAAGGAGA | 2395.1468201311404 | 2480.49214686133 | 2508.318231091742 | 2367.0801425212826 | 2386.337653641165 | 2137.1870410453644 | 2629.0603769840336 | 2119.4825258269666 | 2147.1484017812604 | 3508.7104254518345 | 2251.319562290628 | 2764.7421723179755 |
| SE_423_1_10 | SE_423 | SE_423_1 | AGGTGGCCAGCCTGACCCCG | 959.1425048941445 | 1023.3779042651097 | 1050.7285818431549 | 992.8983775391636 | 1009.6859761183953 | 955.5400511848549 | 913.6603602371857 | 1002.074699989556 | 892.4252042142956 | 1639.5366189156816 | 1235.9955678269193 | 1102.646962091552 |
| SE_423_1_13 | SE_423 | SE_423_1 | GGAGAGAGCTCCAGATCAGG | 1235.5055995246607 | 1575.0425557830204 | 1461.539756699125 | 1404.1118602421461 | 1569.679878839522 | 2020.235176213129 | 1308.6113051218113 | 1567.3960873421547 | 1480.271458134558 | 1872.8188412602872 | 1422.62737721389 | 1851.2862153010794 |
| SE_423_1_27 | SE_423 | SE_423_1 | TTTGGCAGACAGGGAAACCG | 1072.939073271416 | 1158.2958896906857 | 1110.967720463862 | 1137.3436198810339 | 1086.0487810349125 | 909.6256153618292 | 1065.399006637001 | 801.6597599916449 | 1027.3989664437215 | 672.1444031303951 | 902.6406378527074 | 1202.4655291861557 |
| SE_423_1_3 | SE_423 | SE_423_1 | GAGAAAGCAAATCAGTTCGT | 3160.2932703630795 | 3304.9909466842946 | 3356.6038060948154 | 3227.245054124673 | 3259.206937617467 | 2649.1763159775937 | 3162.836040915299 | 2923.032992799724 | 3144.3558687789264 | 3477.363126824278 | 3340.592009530306 | 2585.997296357871 |
| SE_423_1_31 | SE_423 | SE_423_1 | AACAGAATACCGTTCCCACC | 237.34712832973747 | 264.8390084279825 | 194.54279193900518 | 239.44076208021767 | 212.11890254588135 | 347.3903540572319 | 277.64943809327906 | 174.89039575289422 | 243.3079661240965 | 569.3544239098031 | 233.58320797488787 | 107.94333418369929 |
| SE_423_2_10 | SE_423 | SE_423_2 | CCAGGAAGAGCACTGCAGGC | 228.67691359623106 | 135.9173779102099 | 154.05419057098888 | 153.55440176883525 | 75.30221040378788 | 459.1443582302566 | 264.7355107401033 | 106.82494443284891 | 522.1353433611998 | 17.496166675845423 | 80.99116256415711 | 119.5501443109788 |
| SE_423_2_15 | SE_423 | SE_423_2 | AGAAGGAGGAATCGGCCCTG | 2244.501839136467 | 2361.5644411898966 | 2199.2223230871296 | 2320.233036896892 | 2527.396723834176 | 2437.796649169702 | 2527.901279384157 | 2355.820898466013 | 2526.318115412608 | 2192.852890039293 | 2464.948425865651 | 2378.235395079568 |
| SE_423_2_27 | SE_423 | SE_423_2 | GTACTGGCCCGGGACTTGGG | 282.86575568064603 | 502.6944197708498 | 510.5513879820593 | 400.80301478645134 | 446.5102898590802 | 295.41174746512735 | 352.9806809868044 | 387.59493112803585 | 262.8436422362503 | 382.72864603411864 | 316.9219404684408 | 348.20430381838486 |
| SE_423_2_30 | SE_423 | SE_423_2 | CTCCCTTCCTCCCAAGTCCC | 106.21013048545329 | 116.92892070216585 | 123.440857829318 | 136.63739140447206 | 129.39253055298764 | 57.176467251314975 | 99.00677637434757 | 129.51342820619735 | 71.92680750383876 | 126.84720839987932 | 151.41826044603283 | 39.46315443275028 |
| SE_423_2_6 | SE_423 | SE_423_2 | CCAGTGCGTGCAAAGGCCCA | 2196.8156581021817 | 1474.1039148349969 | 1505.9784655176798 | 2078.1896578375413 | 1982.2511442912612 | 1888.556039513131 | 2285.7651415121113 | 2073.1602047897136 | 1716.4755420360532 | 1858.9677093085763 | 2030.6479889274171 | 1229.1611924788986 |
| SE_423_3_12 | SE_423 | SE_423_3 | GGAGCTACATGCCCTATGCT | 1774.1426898437453 | 1566.0480234213153 | 1656.0825486381304 | 1698.2075788841526 | 1616.3460373996159 | 1298.5988546927447 | 1580.8799401512672 | 1623.1719432849698 | 1417.224503408971 | 1215.25457702643 | 1348.6789244379204 | 1094.5221950024563 |
| SE_423_3_20 | SE_423 | SE_423_3 | GTAAATAAACCAGGGCAGGA | 177.73940203688102 | 294.8207829669994 | 328.8464452573032 | 191.29234796626088 | 329.8448934588455 | 257.2941026309174 | 205.54667703804768 | 344.1086705624513 | 221.99631945629244 | 213.59903483427954 | 302.83652089206566 | 262.31390887651656 |
| SE_423_3_28 | SE_423 | SE_423_3 | AGACCAATACAGCAACCTAT | 1532.4604541472547 | 1587.0352655986271 | 1540.5419057098886 | 1519.9283158135556 | 1606.8006867850513 | 2248.9410452183893 | 1670.2012710107329 | 1831.1497112073303 | 1538.8784864710192 | 1078.930278343801 | 1407.3681726728169 | 1708.5224507355415 |
| SE_423_3_32 | SE_423 | SE_423_3 | GGATGGTGAGTATTTCTGGT | 910.3725470181711 | 909.4471610168455 | 870.0111659810333 | 910.9159426964803 | 764.6886436779023 | 900.9625142631451 | 868.4616145010705 | 915.1021788583871 | 651.7811939236747 | 645.1711461718 | 1098.6627269572616 | 566.4123342112393 |
| SE_423_3_4 | SE_423 | SE_423_3 | TGGGTGCTGGGGATACAGAG | 113.79656837727138 | 146.91069524118274 | 125.4159115545871 | 130.13084895664005 | 125.15015250207 | 110.88769406315632 | 117.30150679134657 | 104.93423745173654 | 94.126439449468 | 249.3203751307973 | 160.8085401636162 7 | 123.03218734916264 |
| SE_423_4_16 | SE_423 | SE_423_4 | GTTTACTGATGAGCAGGCAG | 54.188842084414944 | 90.94471610168455 | 81.96472959866715 | 72.87327541571842 | 56.21150917465856 | 35.51871450460476 | 77.48356411905462 | 93.58999556506231 | 59.49501361428638 | 69.98466670338169 | 159.63475519891836 | 13.928172152735392 |
| SE_423_4_17 | SE_423 | SE_423_4 | CTTCCGACCCACACTGCAGA | 96.4561389102586 | 31.980559508284678 | 54.31397744489992 | 88.48897729051524 | 16.969512203670508 | 331.79677207960054 | 96.85445514881827 | 66.17474433893295 | 206.90056973326455 | 8.748083337922711 | 23.47569929395858 | 59.19473164912542 |
| SE_423_4_18 | SE_423 | SE_423_4 | GGTTGAGTACATCTCTCTGA | 214.5878146542832 | 227.86148649652833 | 269.5948334992305 | 205.60674135149128 | 207.87652449496372 | 218.310147686839 | 237.8314954209871 | 393.267052071373 | 182.92496723198497 | 255.15243068941243 | 313.4005855743471 | 224.01143545649424 |
| SE_423_4_19 | SE_423 | SE_423_4 | GGCTCAAATGACATGTTAAG | 322.9654988231131 | 387.7642840379517 | 426.61160465812304 | 337.0388987976977 | 343.6326221243278 | 536.245958008545 | 336.83827179533466 | 423.5183637691709 | 585.1822980867869 | 303.2668890479873 | 462.471276090984 | 299.455701283811 |
| SE_423_4_2 | SE_423 | SE_423_4 | AAGACAAAACCGGAAGTCGA | 1313.5375321262184 | 1697.9678313929896 | 1584.980614528443 | 1513.4217733657238 | 1587.709985555922 | 1264.8127604078768 | 1350.5815690196325 | 1698.8002225294645 | 1407.456665352894 | 2164.421619191044 | 1531.7893789307973 | 1087.5581089260886 |
| SE_424_1_22 | SE_424 | SE_424_1 | GGGAGGTTGGACCCCAAAGT | 405.3325387914238 | 522.6822694635277 | 495.73848504254113 | 498.4011515039314 | 519.6913112374093 | 749.3582450361736 | 529.4710214802066 | 534.1247221642445 | 397.8174044656761 | 468.751465523692 | 788.7834962770083 | 521.1457747148493 |
| SE_424_1_27 | SE_424 | SE_424_1 | GCCGTGCCAAGACTGGTCAA | 2513.2784958751654 | 2370.558973551602 | 2300.9375899384877 | 2653.3680102258904 | 2581.487043983376 | 2498.438356860491 | 2586.0139524734477 | 2870.093197328578 | 2788.273772371033 | 2396.974834590823 | 3030.712778850053 | 2810.0087318143655 |
| SE_424_1_30 | SE_424 | SE_424_1 | GGAGTGCTCTGTTACCAGTG | 920.1265385933658 | 1109.3256579436247 | 1022.090302826753 | 1005.9114624348275 | 1134.8361286204652 | 861.9785593190667 | 955.630624135007 | 1251.648021496389 | 1019.407098943295 | 869.7052851784829 | 1368.6332688377852 | 973.8113696787495 |
| SE_424_1_38 | SE_424 | SE_424_1 | CTGAATGTGACCAGGCGCAG | 1013.3313469785595 | 1054.3590712887606 | 1058.6287967442313 | 1028.0337067574565 | 1071.2004578567007 | 1322.8555377690602 | 1221.4422954878748 | 987.8943976312133 | 1245.8433447887132 | 1479.155091053765 | 943.7231116171349 | 1378.889043120804 |
| SE_424_1_42 | SE_424 | SE_424_1 | GAATGCAGAGGCTCAGGATG | 921.2103154350541 | 937.4301505865947 | 1013.2025610630421 | 908.3133257173475 | 946.0503053546308 | 977.1978039315651 | 860.928490211718 | 979.3862162162077 | 1250.283271177839 | 1092.7814102955122 | 1105.705436745449 | 865.8680354950503 |
| SE_424_2_1 | SE_424 | SE_424_2 | AGCCTCATTTAATCCCCACG | 247.10111990493215 | 281.8286806667587 | 253.79440369707783 | 227.72898567412008 | 311.81478674244556 | 232.17110944473353 | 231.3745317443992 | 346.9447310341199 | 304.57895029403323 | 391.47672937204135 | 323.96465025662843 | 473.5578531930034 |
| SE_424_2_28 | SE_424 | SE_424_2 | TCACTTTTGTTTCTGAACCG | 70.44549470973944 | 111.93195827899638 | 127.39096527985618 | 79.37981786355043 | 86.96875004381135 | 187.12298373157628 | 51.655709412703075 | 33.08737216946648 | 118.10204195074758 | 115.91210422747594 | 138.50662583435562 | 83.5690329164123 5 |
| SE_424_2_32 | SE_424 | SE_424_2 | ATCTGGGCGCAGATCTGCAG | 30.34575156727237 | 16.989672238776237 | 32.58838646693995 | 23.423552812195208 | 8.484756101835254 | 62.37432791052543 | 24.75169409358689 | 29.305958207241734 | 66.59889583688773 | 2.9160277793075706 | 11.73784964697929 | 0.0 |
| SE_424_2_39 | SE_424 | SE_424_2 | CTTCCTGCAGACGCAGATGT | 45.51862735090855 | 10.993317330972857 | 5.925161175807264 | 27.32747828089441 | 9.545350614564661 | 173.26202197368175 | 54.88419125099702 | 30.25131169779792 | 83.47061611556596 | 1.4580138896537853 | 22.30191432926065 | 12.767491140007444 |
| SE_424_2_4 | SE_424 | SE_424_2 | ATAATATACAAATAGACGGG | 13.005322100259587 | 8.994532361705065 | 13.825376076883616 | 1.3013084895664004 | 12.727134152752882 | 12.128341538157722 | 4.30464245105859 | 3.78141396222474 | 20.423661389978903 | 3.6450347241344634 | 8.216494752885502 | 31.338387343654635 |
| SE_425_1_18 | SE_425 | SE_425_1 | AACCACAGAATGCAGCCAGA | 846.4297133585615 | 974.4076725180488 | 1080.3543877221912 | 972.0774417061011 | 993.7770584274541 | 932.1496782184078 | 839.405277956425 | 715.632592351032 | 875.5534839356174 | 898.1365560267318 | 1061.1016080869279 | 1500.7605494572385 |
| SE_425_1_32 | SE_425 | SE_425_1 | TCAGCCGTAACCAGCCCCTG | 2151.2970307512733 | 2471.4976144996253 | 2524.1186608938947 | 2177.089103044588 | 2425.5796506121533 | 2594.5987790558843 | 2273.9273747717 | 2515.5856383700084 | 2398.4482354057836 | 2812.5087931421517 | 2115.160506385668 | 2922.594790048977 |
| SE_425_1_37 | SE_425 | SE_425_1 | AGGAGAGGCCTCATGAGCCG | 2369.1361759306214 | 2663.3809715493335 | 2610.0334979431 | 2262.9754633559705 | 2641.9409312089524 | 2086.074744563128 | 2213.66238045688 | 2487.225033653323 | 2553.845659025188 | 2510.6999179838185 | 2598.759911841215 | 3180.2659748745814 |
| SE_425_1_39 | SE_425 | SE_425_1 | CCAGGGTTGGGACCACATGG | 4273.332086776963 | 4207.4423603087025 | 4203.901854235254 | 4014.5366903123454 | 4565.859377300096 | 4435.507762526253 | 4024.8406917397815 | 4171.844953824444 | 3844.08826770516 | 3268.1381336589598 | 3385.195838188827 | 3475.0789521074807 |
| SE_425_1_52 | SE_425 | SE_425_1 | GGAGGGGTCAGAGAGGAGCA | 2265.0935991285446 | 3004.173808809492 | 3059.3582204418176 | 1957.1679683078662 | 2151.9462663279664 | 1556.7592674335306 | 1696.0291257170845 | 1100.3914630073994 | 2407.328088184035 | 891.5754935232898 | 707.7923337128512 | 825.2442000495721 |
| SE_425_2_10 | SE_425 | SE_425_2 | GTTAGTTCCTCCTAAACCAC | 203.7500462374002 | 205.8748518345826 | 189.60515762583245 | 191.29234796626088 | 158.0285823966816 | 90.09625142631451 | 192.6327496848719 | 397.0484660335977 | 194.4687758437122 | 305.453909882468 | 242.9734876924713 | 68.48017975094902 |
| SE_425_2_11 | SE_425 | SE_425_2 | GGTTAGTTCCTCCTAAACCA | 54.188842084414944 | 79.95139877071169 | 65.1767729338799 | 61.16149900962082 | 51.96913112374093 | 29.454543735525895 | 54.88419125099702 | 46.32232103725307 | 80.80666028209045 | 74.35870837234305 | 32.86597901154201 | 32.49906835638259 |
| SE_425_2_23 | SE_425 | SE_425_2 | CCGGCCCATAAACATGCCCG | 579.82061030324 | 454.72358050842274 | 462.1625717129666 | 573.8770438987826 | 460.2980185245625 | 394.171099990126 | 747.93162587143 | 455.6603824480812 | 695.292472537108 | 541.6521600063812 | 389.69660827971245 | 550.162800033048 |
| SE_425_2_26 | SE_425 | SE_425_2 | TCAGAACCAGTAAGAGGACA | 1433.8367615536195 | 1504.0856893740138 | 1373.6498659246508 | 1505.6139224283254 | 1174.0781255914533 | 1380.0320050203752 | 1560.4328885087389 | 1501.2213430032218 | 1245.8433447887132 | 1664.322855039796 | 1613.9543264596525 | 1109.6110481679198 |
| SE_425_2_28 | SE_425 | SE_425_2 | CAGCCGCTGTCCCTGCTGGA | 1863.0123908621858 | 1838.882171726369 | 1753.8477080389503 | 1935.0457239852374 | 2064.9775162841547 | 2101.6683265407596 | 1910.1850876572494 | 2387.017563654367 | 2148.036387059086 | 1813.769278729309 | 2567.0677177943708 | 1833.87600011016 |
| SE_425_3_0 | SE_425 | SE_425_3 | TCAGAAGCACATACAACCAC | 9610.933032091834 | 8110.070012804067 | 8434.466933761641 | 8612.059583950439 | 8166.5777480164315 | 9203.678607241975 | 9147.365208499503 | 7622.38519435452 | 7546.986876236118 | 8632.900240640063 | 8738.829062176083 | 6972.210843456793 |

| | | | | | | | | | | | | | | | |
|---|---|---|---|---|---|---|---|---|---|---|---|---|---|---|---|
| SE_425_3_12 | SE_425 | SE_425_3 | TTGCTGGTGGGAAAGTAACA | 408.5838693164887 | 395.75942391502286 | 434.51181955919935 | 400.80301478645134 | 393.4805642226099 | 420.1604032861782 | 417.5503177526832 | 379.0867497130302 | 541.6710194733536 | 361.58744463413876 | 642.0603756897672 | 359.8111139456643 |
| SE_425_3_14 | SE_425 | SE_425_3 | TCTTTTGGCCGGAAATGGAT | 136.55588205272565 | 178.8912547494674 | 175.77978154894885 | 171.77272062276487 | 176.05868911308153 | 142.07485801841904 | 133.4439159828163 | 167.32756782844476 | 168.71720278678225 | 174.23265981362735 | 147.89690555193906 | 185.70896203647192 |
| SE_425_3_18 | SE_425 | SE_425_3 | GATCTGAGAGCACCTGCTAT | 539.7208671607729 | 498.6968498323142 | 552.0275162127101 | 653.2568617623331 | 614.0842228703265 | 466.9411492190723 | 643.5440464332592 | 709.960471407695 | 775.2111475413732 | 790.9725351371785 | 983.6318004168645 | 399.2742683784146 |
| SE_425_3_2 | SE_425 | SE_425_3 | GTATCTTTTCACGTACTCAT | 29.26197472558407 | 49.96962423169481 | 42.463655093285396 | 36.43663770785921 | 84.84756101835254 | 3.465240439473635 | 41.970263897821255 | 9.45353490556185 | 29.303514168230603 | 73.62970142751615 | 18.780559435166865 | 11.606810127279495 |
| SE_426_1_0 | SE_426 | SE_426_1 | ATGAACTGATACAGTCCCCG | 348.97614302363223 | 346.78919216796197 | 325.88386466939954 | 411.21348270298256 | 351.0567837134336 | 465.2085289993355 | 517.6332547397955 | 518.9990663153455 | 154.50943834157954 | 198.2898889929148 | 604.4992568194334 | 461.95104306572387 |
| SE_426_1_13 | SE_426 | SE_426_1 | GATGTGGGTGACCCCAGCAA | 527.7993219022015 | 704.5717016668968 | 641.8924607124536 | 553.0561080657202 | 505.903582571927 | 355.1871450460476 | 442.3020118462701 | 318.58412631743437 | 634.0214883671712 | 225.26314595150984 | 267.6229719511278 | 493.2894304093785 |
| SE_426_1_18 | SE_426 | SE_426_1 | ATTTATATACAGAAAAGAGA | 0.0 | 0.9993924846338962 | 0.987526862634544 | 0.0 | 0.0 | 0.0 | 0.0 | 0.0 | 0.0 | 0.0 | 0.0 | 0.0 |
| SE_426_1_5 | SE_426 | SE_426_1 | GCTCATACAGTCACTCCCCA | 3274.089838740351 | 3326.97758134624 | 3367.466601583795 | 3268.8869257907977 | 3492.5377304179365 | 3888.866083199287 | 3317.8031691534084 | 4253.145354012277 | 3202.9628971153875 | 3953.404661796239 | 5068.403477565657 | 5041.998319290213 |
| SE_426_1_6 | SE_426 | SE_426_1 | ATACCTCCTTCCCAGCCACT | 240.59845885480235 | 178.8912547494674 | 165.9045129226034 | 230.33160265325287 | 135.75609762936406 | 179.3261927427606 | 185.09962539551935 | 108.71565141396128 | 294.81111223795637 | 118.09912506195661 | 140.85419576375148 | 240.26096963468552 |
| SE_426_2_2 | SE_426 | SE_426_2 | TGTAGGATCCCTTCTTTAGA | 154.98008836142674 | 227.86148649652833 | 225.15612468067604 | 201.70281588279207 | 199.39176839312847 | 113.48662439276154 | 162.50025252746178 | 161.65544688510764 | 214.00445195586593 | 115.18309728264904 | 98.59793703462604 | 129.99627342553035 |
| SE_426_2_24 | SE_426 | SE_426_2 | ACATTGAGTGTGACAGGGGA | 1624.58148569076 | 1868.863946265386 | 1734.0971707862593 | 1798.4083325807653 | 1628.0125770396394 | 1917.1442731387885 | 1724.0093016489652 | 2009.8215209224493 | 1371.0492689620621 | 2162.9636053013905 | 2263.057411937607 | 1826.9119140337923 |
| SE_426_2_30 | SE_426 | SE_426_2 | ATGCGGTGAAAATTCTTGGG | 494.2022398098643 | 783.5237079529746 | 806.8094467724225 | 628.5320004605715 | 718.0224851178084 | 691.3154676749901 | 491.8054000334439 | 737.3757226338244 | 790.3068972644011 | 855.8541532267719 | 818.1281203944566 | 770.6921924513584 |
| SE_426_2_32 | SE_426 | SE_426_2 | CTGGGAGTCCTGGTGCCGGG | 279.6144251555811 | 423.742413484772 | 388.0980570153758 | 361.7637600994593 | 462.4192075500213 | 492.93045251512456 | 307.7819352506892 | 256.19079594072616 | 557.6547544742066 | 865.3312435095215 | 370.91604884454557 | 419.0058455947897 |
| SE_426_2_40 | SE_426 | SE_426_2 | GGGATTAACTACCAAGGTGG | 26.010644200519174 | 21.986634661945715 | 57.27655803280356 | 27.32747828089441 | 36.06021343279983 | 137.743307469077 | 17.21856980423436 | 42.54090707502832 | 38.1833669464823 | 123.20217367574486 | 53.994108376104734 | 85.89039494186825 |
| SE_427_1_15 | SE_427 | SE_427_1 | TCTGTGGCTGCTTCCAAGGA | 688.1982944720698 | 896.4550587166049 | 913.4623479369533 | 870.5753795199219 | 856.9603662853607 | 804.8020920677517 | 733.9415379054896 | 797.8783460294202 | 805.402646987429 | 952.0830699439218 | 1000.0647899226356 | 1144.4314785497581 |
| SE_427_1_17 | SE_427 | SE_427_1 | CTGCTCATTTTAGTTCCACT | 183.15828624532253 | 233.8578414043317 | 210.3432217411579 | 161.36225270623365 | 212.11890254588135 | 121.28341538157723 | 134.52007659558095 | 133.2948421684221 | 202.4606433441387 | 439.5911877306163 | 53.994108376104734 | 104.46129114551545 |
| SE_427_1_19 | SE_427 | SE_427_1 | AAGCTGGAAGATGCACCAAA | 1014.4151238202478 | 1100.3311255819197 | 1094.1797637990749 | 1003.3088454556947 | 1086.0487810349125 | 1230.1603560131405 | 1022.3525821264151 | 1116.4624723468544 | 776.0991328191984 | 1087.6783616817238 | 1007.1074997108232 | 1334.783164637142 |
| SE_427_1_27 | SE_427 | SE_427_1 | CCTCCCCCAACACAGCCGGA | 447.59983561726744 | 430.7381608772093 | 521.4141834710392 | 493.1959175456658 | 530.2972563647033 | 365.5828663644685 | 444.4543330717994 | 487.80240112699147 | 327.66656751748764 | 430.11409744786664 | 385.00146842092073 | 235.61824558377373 |
| SE_427_1_4 | SE_427 | SE_427_1 | GTACAAAAGACATCCAGCGG | 318.63039145635986 | 487.70353250134133 | 523.3892371963084 | 511.4142363995954 | 469.84336913912716 | 749.3582450361736 | 427.23576326756506 | 480.239573202542 | 616.2617828106678 | 559.8773336270535 | 401.4344579266917 | 601.2327645930778 |
| SE_427_2_13 | SE_427 | SE_427_2 | TGCTGGGCAGGGACGCTGCG | 4568.1193877161795 | 3842.664103417331 | 4062.685512878514 | 3985.9079035418845 | 3456.4775169851364 | 3200.149545853902 | 3954.8902519100793 | 3269.977723833844 | 3305.9691893431072 | 3650.1377727482513 | 2821.779055133821 | 2974.8254356217344 |
| SE_427_2_25 | SE_427 | SE_427_2 | GAGCAAGAGTGAACAGGAAG | 949.3885133189499 | 1095.3341631587502 | 1080.3543877221912 | 1059.26511050705 | 1194.229421333312 | 960.7379118440653 | 911.5080390116564 | 1096.6100490451747 | 1030.950907555022 | 1016.2356810886884 | 1072.8394577339072 | 1279.0704760262004 |
| SE_427_2_26 | SE_427 | SE_427_2 | AGAGCAAGAGTGAACAGGAA | 1400.239679461282 | 1792.9101174332097 | 1726.196955885183 | 1466.5746677413333 | 1593.012958119569 | 1961.3260887420774 | 1456.045309070568 | 1421.8116497965023 | 1670.3003075891443 | 1511.2313966261486 | 1287.642106273628 | 2127.528296330331 |
| SE_427_2_28 | SE_427 | SE_427_2 | TCTATAGAGCAAGAGTGAAC | 147.39365046960864 | 204.87545934994873 | 168.86709351050703 | 124.92561499837444 | 185.60403972764618 | 183.65774329210265 | 179.71882233169612 | 108.71565141396128 | 183.81295250981015 | 252.23640291010486 | 79.81737759945918 | 17.410215190919242 |
| SE_427_2_30 | SE_427 | SE_427_2 | ACGCTGCAAAGCAGATGTGC | 18381.939011875238 | 15123.806469964751 | 15110.148525171158 | 17554.65152425074 | 15672.405114602443 | 15621.303901147146 | 18272.13104413095 | 15613.458250025951 | 14824.91421329121 | 15083.153688468408 | 15877.789217468886 | 15329.114135098029 |
| SE_427_3_25 | SE_427 | SE_427_3 | GGGATGATGTTACAGTCCAG | 555.9775197860973 | 576.649463633758 | 498.70106563044476 | 615.5189155649074 | 411.5106709390098 | 706.9090496526215 | 677.9811860417279 | 574.7749222581605 | 540.7830341955284 | 424.2820418892515 | 558.7216431962142 | 659.2668152294752 |
| SE_427_3_31 | SE_427 | SE_427_3 | CCACTGAGCTCTAGTTACTG | 505.0400082267473 | 567.654931272053 | 510.5513879820593 | 649.3529362936338 | 618.3266009212441 | 485.9999716361773 | 615.5638705013783 | 667.4195643326666 | 483.95197641471754 | 408.24388910305987 | 563.4167830550059 | 685.9624785222181 |
| SE_427_3_7 | SE_427 | SE_427_3 | AATGGGACACAGCTCATATG | 1046.9284290708968 | 1080.3432758892418 | 1060.6038504695002 | 1145.1514708184325 | 1109.3818603149596 | 745.0266944868315 | 1010.514815386004 | 1072.97621178127 | 1078.0141272797562 | 1027.8997922059186 | 1098.6627269572616 | 863.5466734695943 |
| SE_427_3_8 | SE_427 | SE_427_3 | TGCCAAATAATGTTCCCAGG | 719.6278228810305 | 872.4696390853913 | 949.0133149917968 | 753.4576154589458 | 1030.8978663729833 | 642.8021015223593 | 535.9279851567944 | 986.9490441406572 | 979.4477614411622 | 802.6366462544088 | 1036.4521238282714 | 762.5674253622627 |
| SE_427_3_9 | SE_427 | SE_427_3 | GTAACTCTACTGTGTAGCCA | 238.43090517142576 | 240.85358879676897 | 222.1935440927724 | 210.81197530975686 | 257.72446659324584 | 136.8769973592086 | 203.39435581251837 | 229.72089820515296 | 187.36489362111084 | 129.7632361791869 | 267.6229719511278 | 139.28172152735394 |
| SE_428_1_16 | SE_428 | SE_428_1 | GGTGGGTGGGAGGTTCTCCG | 5.4188842084414945 | 5.9963549078033775 | 9.87526862634544 | 3.9039254686992013 | 7.4241615891058474 | 0.0 | 15.066248578705064 | 1.89070698111237 | 12.431793889552377 | 0.7290069448268927 | 8.216494752885502 | 0.0 |
| SE_428_1_17 | SE_428 | SE_428_1 | GCATCAGCACTGGAAGGACA | 7877.973862232245 | 6583.997688768108 | 6411.0243922234595 | 7453.895028236342 | 6478.111283751216 | 6700.042389722273 | 8693.225429912822 | 7474.910049827755 | 6280.719870057426 | 9283.903442370478 | 7570.913022301642 | 7278.630630816971 |
| SE_428_1_25 | SE_428 | SE_428_1 | ACTTCCTCATGGTGACCCAA | 274.19554094713965 | 345.78979968332806 | 258.73203801025056 | 247.2486130176161 | 352.11737822616305 | 330.0641518598637 | 225.99372868057597 | 335.60048914744567 | 253.07580418017338 | 96.22891671714983 | 240.62591776307545 | 271.59935697834015 |
| SE_428_1_26 | SE_428 | SE_428_1 | TCTCCCTGTTGAAACTCAAG | 105.126353643765 | 139.91494784874547 | 151.09160998308525 | 165.26617817493286 | 131.51371957844643 | 134.27806702960336 | 110.8445431147587 | 172.99968877178185 | 62.15896944776188 | 128.3052222895331 | 99.77172199932397 | 31.338387343654635 |
| SE_428_1_4 | SE_428 | SE_428_1 | TAAAAGGGACGTCTACCCAG | 587.407048195058 | 681.5856745203172 | 693.2438575694499 | 701.4052758762898 | 585.4481710266325 | 759.7539663545945 | 661.8387768502582 | 707.1244109360264 | 730.8118836501147 | 577.373500302899 | 575.1546327019852 | 721.9435899167845 |
| SE_429_1_1 | SE_429 | SE_429_1 | ACACTCATCAGTGTGAACCA | 1507.5335867884237 | 1491.093587073773 | 1446.726853759607 | 1568.0767299275126 | 1469.9839946429577 | 1412.0854790855062 | 1525.9957489002702 | 1404.795286966491 | 1396.8008420189922 | 1759.822764812119 | 1341.636214649733 | 1782.8060355501302 |
| SE_429_1_11 | SE_429 | SE_429_1 | TGTGGTTACAACAAAGGGGA | 267.69287989700985 | 315.8080251443112 | 316.00859604305407 | 227.72898567412008 | 293.7846800260457 | 145.54009845789267 | 194.7850709104012 | 402.72058697693484 | 234.42811334584482 | 189.5418056549921 | 282.88217649220087 | 167.1380658328247 |
| SE_429_1_12 | SE_429 | SE_429_1 | AAGAAGGACCAAGACTCCAG | 937.4669680603786 | 1041.3669689885198 | 1120.8429890902075 | 880.9858474364531 | 979.9893297619718 | 1245.7539379907716 | 840.4814385691897 | 1146.7137840446524 | 1078.0141272797562 | 852.9381254474644 | 1139.7452007216891 | 841.4937342277633 |
| SE_429_1_14 | SE_429 | SE_429_1 | ATATTTTCTAAAGAAATGGA | 1.083776841688299 | 3.9975699385355847 | 4.93763431317272 | 1.3013084895664004 | 1.0605945127294067 | 0.0 | 6.456963676587884 | 0.0 | 8.879852778251697 | 22.599215289633673 | 0.0 | 0.0 |
| SE_429_1_18 | SE_429 | SE_429_1 | TCTCTTTGAGCTATGTCAGG | 9.75399157519469 | 41.97448435462364 | 53.32645058226538 | 13.013084895664004 | 13.787728665482287 | 34.65240439473635 | 9.685445514881827 | 42.54090707502832 | 20.423661389978903 | 18.225173620672315 | 199.54344399864794 | 0.0 |
| SE_429_2_0 | SE_429 | SE_429_2 | CTGAGGTGAGAAAGTTAGTG | 1309.202424759465 | 1349.17985425576 | 1317.3608347544816 | 1398.9066262838805 | 1343.7732476281583 | 1198.9731920578777 | 1531.3765519640933 | 1809.406580924538 | 1155.2688464505459 | 1584.8610980536646 | 1532.9631638954954 | 1684.1481494682546 |
| SE_429_2_1 | SE_429 | SE_429_2 | GGGAAGGGCTCTGTGTTCAG | 966.7289427859627 | 1176.2849544140959 | 1073.4416996837494 | 1041.0467916531204 | 1098.7759151876653 | 1259.6148997486664 | 997.6008880328282 | 1120.2438863090792 | 983.8876878302881 | 1074.5562366748397 | 1342.8099996144308 | 1587.811625411835 |
| SE_429_2_13 | SE_429 | SE_429_2 | AGGATCAGAGGAAGCCCAGG | 1169.3952121816744 | 1603.0255453527695 | 1410.1883598421289 | 1419.7275621169429 | 1643.9214947305804 | 1305.529335571692 | 1041.7234731561787 | 1259.2108494208385 | 1403.0167389637681 | 1328.2506534745985 | 1379.1973335200666 | 959.8831975260142 |
| SE_429_2_22 | SE_429 | SE_429_2 | TTTAAAGGCACAGTTCTCAT | 66.11038734298623 | 101.9380334326574 | 98.75268626345441 | 87.18766880094883 | 109.24123481112889 | 77.96790988815678 | 64.56963676587885 | 140.85767009287156 | 98.56636583859384 | 74.35870837234305 | 225.36671322200237 | 69.64086076367697 |
| SE_429_2_29 | SE_429 | SE_429_2 | CTAGATGTGGTAAATCAGAT | 498.5373471766175 | 515.6865220710904 | 565.8528922895937 | 486.68937509783376 | 626.8113570230794 | 429.68981449473074 | 506.871648612149 | 708.0697644265825 | 463.5283150247386 | 328.0531251721017 | 652.6244403720485 | 643.0172810512839 |
| SE_429_3_17 | SE_429 | SE_429_3 | CCAGGGTTAATGTTCACTGA | 1143.3845679811554 | 1423.1348981186682 | 1259.0967498590437 | 1220.6273632132836 | 1265.2892536861823 | 1143.5293450262996 | 1000.8293698711221 | 1090.9379281018375 | 1025.622995888071 | 1406.9834035159029 | 1014.1502094990107 | 1234.9645975425383 |
| SE_429_3_2 | SE_429 | SE_429_3 | TCAGTGTGCATTCTCCTGGA | 3108.271981962041 | 2863.2594684761125 | 2907.2790835960977 | 2866.78260251478 | 3087.3906265553032 | 3112.6522247571925 | 3008.9450732899545 | 3256.7427749660574 | 2617.7805990286006 | 2540.589202721721 | 3513.1383993409017 | 2396.8062912832156 |
| SE_429_3_3 | SE_429 | SE_429_3 | TCATTCCTCATTCAGTCCCT | 460.60515771752705 | 510.68955964792093 | 551.0399893500755 | 504.90769395176335 | 610.9024393321383 | 517.18713559144 | 601.5737825354379 | 364.90644735468743 | 428.896889189557 | 381.27063214446486 | 650.2768704426527 | 500.2535164857462 |
| SE_429_3_4 | SE_429 | SE_429_3 | TGGAGCGCAGATGTAAGTGG | 1327.6266310681663 | 1281.221165300655 | 1390.437822589438 | 1331.2385848264275 | 1283.319360402582 | 970.2673230526178 | 1272.0218442878133 | 1072.97621178127 | 1440.3121206324254 | 922.9227921508461 | 808.7378406768731 | 986.578860818757 |
| SE_429_3_5 | SE_429 | SE_429_3 | ACTGAATGAGGAATGAACTG | 1649.508353049591 | 1833.8852093031994 | 1798.2864168575047 | 1676.0853345615237 | 1833.7679125091443 | 1617.400975124319 | 1792.8835808659028 | 1836.8218321506674 | 1661.4204548108926 | 2082.0438344256054 | 1860.4491690462175 | 1381.2104051462597 |
| SE_43_1_16 | SE_43 | SE_43_1 | CCGCCATCTTCTTCCACTAG | 693.6171786805113 | 646.6069375581309 | 661.6429979651446 | 814.6191144685666 | 718.0224851178084 | 861.1122492091982 | 947.0213392328898 | 731.7036016904872 | 664.212987813227 | 779.3084240199482 | 665.5360749837257 | 1056.219721582434 |
| SE_43_1_18 | SE_43 | SE_43_1 | TGGAAGAAGATGGCGGAAGG | 281.78197883895774 | 275.83232575895534 | 306.13332741670865 | 334.4362818185649 | 429.5407776554097 | 280.68447559736444 | 323.9243444421589 | 261.8629168840632 | 335.6584350179142 | 167.67159731018532 | 320.4432953625346 | 331.95476964019355 |
| SE_43_1_19 | SE_43 | SE_43_1 | GAAGATGGCGGAAGGCGGAG | 2491.602959041399 | 2843.271618783435 | 2597.1956487288508 | 2845.961666681718 | 2908.1501539040332 | 3025.1549036604833 | 2572.0238645075074 | 2451.301601012188 | 3252.690072673597 | 1830.5364384603274 | 2649.232665323226 | 2464.125790021437 |
| SE_43_1_23 | SE_43 | SE_43_1 | CGTCGCGATGTCAGACCGCT | 17885.569218381996 | 13847.582267087266 | 14209.524026448455 | 15740.627489795179 | 14380.600998098025 | 15097.186284676758 | 16431.896396303404 | 13690.609250234671 | 13432.553297661343 | 13039.747222118629 | 13172.21487384016 | 13648.448028667957 |
| SE_43_1_30 | SE_43 | SE_43_1 | CGCTCCGGAAAGGGGACACC | 2628.158841094125 | 2865.2582534453804 | 2882.590912030234 | 2315.0278029386263 | 2824.36318739841 | 2713.283264107856 | 2350.33477827799 | 2290.591507617636 | 2561.837526525615 | 2710.447820866387 | 2919.2032072037496 | 1883.7852836574618 |
| SE_43_2_0 | SE_43 | SE_43_2 | GAGTTCTTGTAGCTTTGCAC | 39.01596630077876 | 69.95747392437274 | 85.91483704920533 | 54.65495656178882 | 53.02972563647034 | 64.97325824013066 | 36.58946083399801 | 77.51898622560716 | 70.15083694818841 | 87.48083337922712 | 66.90574298778195 | 34.820430381838484 |
| SE_43_2_12 | SE_43 | SE_43_2 | TCTCTAGCCAGGCTTTTGGG | 9.75399157519469 | 17.98906472341013 | 35.55096705484358 | 13.013084895664004 | 31.8178353818822 | 1.7326202197368175 | 0.0 | 8.508181415005666 | 28.415528890405433 | 13.122125006884067 | 0.0 | 9.285448101823595 |
| SE_43_2_13 | SE_43 | SE_43_2 | AGCAGCACACTGAAAGGCAT | 4410.971745671377 | 3663.7728486678634 | 3739.7642287970184 | 4156.379315675083 | 3873.2911604877936 | 3265.989114203901 | 4430.553242752054 | 3835.2991111864426 | 4220.594025503032 | 4645.23225243696 | 3966.219395714302 | 2731.082422948865 |
| SE_43_2_4 | SE_43 | SE_43_2 | TGAAGAAGAGTGATGTCAGC | 2398.3981506562054 | 2569.438077993747 | 2617.933712844176 | 2484.1979065822584 | 2517.8513732196116 | 2609.326050923647 | 2396.60968462687 | 2505.18674997389 | 2237.722900119428 | 2262.108549797848 | 2439.1251566422966 | 2254.042526717678 |
| SE_43_2_5 | SE_43 | SE_43_2 | ACTAAGCTAAGACTCCTTGG | 551.6424124193442 | 617.6245555037478 | 596.4662250312646 | 579.0822778570482 | 646.9626527649381 | 531.914407459203 | 509.02396983767824 | 486.85704763643525 | 568.3105778081086 | 683.0795073027984 | 501.2061799260157 | 916.93800005508 |
| SE_430_1_1 | SE_430 | SE_430_1 | AGTTTTTTCTCTGGTCTGGA | 0.0 | 0.9993924846338962 | 0.987526862634544 | 0.0 | 0.0 | 0.0 | 0.0 | 0.945353490556185 | 0.0 | 0.0 | 0.0 | 0.0 |
| SE_430_1_12 | SE_430 | SE_430_1 | AAGGTAGGAAAAGAATAGCA | 1292.9457721341405 | 1498.0893344662104 | 1558.3173892373104 | 1379.3869989403845 | 1467.862805617499 | 1781.1335858894483 | 1291.392735317577 | 1538.090129134913 | 1522.006766192341 | 1183.1782714540468 | 1778.2842215173625 | 1982.4431697393377 |
| SE_430_1_19 | SE_430 | SE_430_1 | CGAGTGTTCCCTTCTGATAA | 1692.859426717123 | 1886.853010988796 | 1758.7853423521228 | 1664.373558155426 | 1596.194741657757 | 1484.8555283144526 | 1678.81055591285 | 1772.5377947928469 | 1601.037455918781 | 1790.4410564948485 | 1916.7908473517182 | 1910.4809469502047 |
| SE_430_1_7 | SE_430 | SE_430_1 | GGGTATCTGTGATGTAAAGG | 47.686181034285156 | 58.96415659339988 | 46.41376254382357 | 85.88636031138243 | 34.999618920070425 | 13.86096175789454 | 94.70213392328898 | 63.33868386726439 | 50.615160836034676 | 115.91210422747594 | 30.518409082146157 | 27.856344305470785 |
| SE_430_1_8 | SE_430 | SE_430_1 | TTTGTGCTGCCAGTTCTGCA | 92.12103154350541 | 14.990887269508443 | 38.51354764274722 | 40.34056317655841 | 27.575457330964575 | 208.7807364782865 | 82.86436718287786 | 13.23494886778659 | 103.89427750554486 | 4.374041668961356 | 17.606774470468935 | 44.10587848366208 |
| SE_430_2_14 | SE_430 | SE_430_2 | ATACCGATGTTCGCTGACCA | 872.4403575590807 | 774.5291755912696 | 796.934178146077 | 654.5581702518995 | 750.90091501242 | 572.6309826230182 | 839.405277956425 | 811.1132948972067 | 602.0540183654651 | 989.2624241300933 | 1172.6111797332312 | 598.911402567622 |
| SE_430_2_3 | SE_430 | SE_430_2 | TTCTGTAGAGAGGCCTAAGG | 1.083776841688299 | 2.9981774539016888 | 0.0 | 0.0 | 0.0 | 0.0 | 0.0 | 0.945353490556185 | 6.215896944776189 | 0.7290069448268927 | 0.0 | 16.249534178191293 |
| SE_430_2_37 | SE_430 | SE_430_2 | GTTACTAAGGCACGTGACCG | 263.3577725302566 | 260.8414384894469 | 262.6821454607887 | 275.8773997880769 | 251.3608995168694 | 238.2352802138124 | 241.05997725928103 | 204.19635396013595 | 271.72349501450196 | 300.35086126867975 | 232.40942301018995 | 302.9377443219948 |
| SE_430_2_5 | SE_430 | SE_430_2 | AAGTGTGTGTGTAAGCGCGG | 1236.5893763663491 | 1271.227240454316 | 1458.5771761112214 | 1370.2778395134196 | 1562.2557172504162 | 1350.5774612848493 | 1276.3264867388718 | 1346.1833705520075 | 1395.912856741167 | 1234.937764536756 | 1552.9175082953602 | 610.5182126949014 |
| SE_430_3_1 | SE_430 | SE_430_3 | CTTACGATGGACTTCTCCGG | 714.2089386725889 | 762.5364657756628 | 790.0214901076353 | 746.9510730111139 | 814.5365857761844 | 882.7700019559085 | 660.7626162374936 | 856.4902624439036 | 887.0972925473446 | 1256.807972881563 | 920.2474123231764 | 820.6014759986602 |
| SE_430_3_10 | SE_430 | SE_430_3 | CAGTTTTGAGCACTAAACAG | 398.829877741294 | 550.6652590332768 | 550.0524624874411 | 433.33572702561133 | 534.539634415621 | 481.6684210868353 | 448.758975522858 | 599.3541130126213 | 597.6140919763393 | 821.590826819908 | 469.5139858791716 | 699.8906506749535 |
| SE_430_3_13 | SE_430 | SE_430_3 | GTTTTGCTCTACAAACAACT | 4.335107366753196 | 16.989672238776237 | 19.75053725269088 | 13.013084895664004 | 25.454268305505764 | 0.8663101098684087 | 1.0761606127646475 | 0.0 | 16.871720278678225 | 25.515243068941242 | 0.0 | 0.0 |
| SE_430_3_2 | SE_430 | SE_430_3 | ACTTACGATGGACTTCTCCG | 515.8777766436302 | 521.6828769788938 | 526.351817784212 | 510.112927910029 | 532.4184453901622 | 472.13900987828276 | 373.4277326293327 | 253.35473546905757 | 589.6222244759127 | 191.72882648947277 | 255.88512230414852 | 560.6089291475996 |
| SE_430_3_8 | SE_430 | SE_430_3 | ACAGGAAGAGGAAATGAAAG | 1051.26353643765 | 1321.1968646860107 | 1253.1715886832365 | 1298.7058725872675 | 1219.6836896388177 | 1180.7806797506412 | 1235.4323834538152 | 1176.019742251894 | 1149.0529495057697 | 1340.6437715366555 | 1010.6288546049169 | 1150.2348836133979 |
| SE_430_4_19 | SE_430 | SE_430_4 | CAGAAATGTTCCATAAATGA | 337.054597765061 | 374.7721817377111 | 405.87354054279757 | 321.4231969229009 | 391.3593751971511 | 479.06949075723 | 274.42095625498513 | 366.7971543357998 | 408.4732277995781 | 297.4348334893722 | 388.52282331501453 | 203.11917722739116 |
| SE_430_4_2 | SE_430 | SE_430_4 | AGAAGCGTCTGTTCACACAT | 1476.1040583794631 | 1645.999422192027 | 1556.3423355120415 | 1565.4741129483798 | 1581.3464184795455 | 1771.6041746808958 | 1318.296750636693 | 1755.5214319628355 | 1532.662589526243 | 1347.2048340400977 | 2106.944011632783 | 2277.256146972237 |
| SE_430_4_3 | SE_430 | SE_430_4 | GCTAATATAGGCCAGCATGG | 630.75812186259 | 726.5583363288425 | 707.0692336463335 | 791.1955616563714 | 766.809832703361 | 621.1443487756491 | 601.5737825354379 | 731.7036016904872 | 578.9664011420107 | 994.3654727438816 | 710.1399036422471 | 662.7488582676591 |
| SE_430_4_5 | SE_430 | SE_430_4 | ACTGCAAGTGAGAAATGGCA | 2593.477982160099 | 2344.5747689511204 | 2226.873075240897 | 2666.3810951215546 | 2433.0038122012593 | 2235.0800834604947 | 2612.917967792564 | 2610.120987425627 | 2370.0327065153783 | 2292.7268414805776 | 2542.418233535714 | 2249.399802666766 |
| SE_430_4_7 | SE_430 | SE_430_4 | CTTGGACTGCAAGTGAGAAA | 508.2913387518122 | 539.6719417023039 | 526.351817784212 | 610.3136816066418 | 608.7812503066795 | 518.9197558111769 | 514.4047729015015 | 688.2173411249026 | 608.2699153102413 | 542.3811669512081 | 836.9086798296235 | 739.3538051077038 |
| SE_430_5_15 | SE_430 | SE_430_5 | GAAGTGAGCTCGGGAAAGTA | 703.371170255706 | 888.4599188395337 | 901.6120255853388 | 869.2740710303555 | 1002.2618145292894 | 912.2245456914344 | 857.700008373424 | 852.7088484816788 | 725.4839719831638 | 768.3733198475448 | 1010.6288546049169 | 1015.5958861369558 |
| SE_430_5_17 | SE_430 | SE_430_5 | AATGTCTTAGAAGTGAGCTC | 114.88034521895969 | 138.91555536411158 | 154.05419057098888 | 101.50206218617923 | 158.0285823966816 | 188.8556039513131 | 79.63588534458391 | 111.55171188562983 | 164.2772763976564 | 252.23640291010486 | 130.29013108147012 | 174.10215190919243 |
| SE_430_5_27 | SE_430 | SE_430_5 | GGAGAGCGCCCCTTATTAGC | 6091.909627129929 | 6839.842164834386 | 6835.660943156314 | 5734.866513519127 | 6170.5388750596885 | 5679.529080297288 | 5403.402436691295 | 5817.705380882762 | 6468.084763678537 | 5393.193377829351 | 5755.067681913946 | 6381.424207978266 |

| | | | | | | | | | | | | | | | |
|---|---|---|---|---|---|---|---|---|---|---|---|---|---|---|---|
| SE_430_5_28 | SE_430 | SE_430_5 | AGGAGAGCGCCCCTTATTAG | 2584.807767426593 | 3010.1701637172955 | 3020.8446727990704 | 2780.896242203398 | 3277.237044333867 | 3129.1121168446925 | 2509.606548967158 | 3369.2398403422435 | 3001.390239049074 | 3288.5503281141127 | 3872.316598538468 | 3262.674326778266 |
| SE_430_5_5 | SE_430 | SE_430_5 | TTTTTTACCGCTTTCCCTAG | 0.0 | 0.9993924846338962 | 0.0 | 0.0 | 0.0 | 0.0 | 0.0 | 0.0 | 0.0 | 0.0 | 0.0 | 0.0 |
| SE_431_1_0 | SE_431 | SE_431_1 | GTAGGCTGCAGGGCTAAGAG | 704.4549470973943 | 274.8329332743215 | 477.9630015151193 | 536.139097701357 | 191.9676068040226 | 2086.9410546729964 | 924.4219663648322 | 561.5399733903739 | 1618.7971614752846 | 70.71367364820858 | 241.7997027277734 | 648.8206861149238 |
| SE_431_1_13 | SE_431 | SE_431_1 | AGATTCATGTCCCATCCCAG | 211.3364841292183 | 300.81713787480277 | 334.77160643311043 | 262.8643148924129 | 238.63376536411653 | 326.5989114203901 | 229.2222105188699 | 121.00524679119168 | 313.4588030722849 | 247.8623612411435 | 224.19292825730446 | 500.2535164857462 |
| SE_431_1_27 | SE_431 | SE_431_1 | TTAGTTCCTCATCTGTACAA | 37.93218945909046 | 45.972054293159225 | 60.23913862070719 | 35.13532921829281 | 32.87842989461161 | 19.05882241710499 | 23.675533480822246 | 34.97807915057884 | 29.303514168230603 | 38.63736807582531 | 31.692194046844083 | 18.57089620364719 |
| SE_431_1_29 | SE_431 | SE_431_1 | GACTCTTACTGACACAACAG | 131.13699784428417 | 153.90644263362 | 154.05419057098888 | 175.67664609146405 | 170.75571654943448 | 119.5507951618404 | 164.65257375299106 | 231.61160518626534 | 198.90870223283804 | 179.3357084274156 | 118.55228143449084 | 80.08698987822851 |
| SE_431_1_3 | SE_431 | SE_431_1 | ACCTACATATAAATGCACCA | 1724.2889551260835 | 1964.8056247902398 | 2075.7814652578118 | 1957.1679683078662 | 1918.6154735274968 | 2151.914312913127 | 1721.856980423436 | 1975.7887952624267 | 1783.9624231507662 | 2066.7346885842408 | 2312.3563804549203 | 2055.5660735411984 |
| SE_431_2_12 | SE_431 | SE_431_2 | TGATGTAATTCAGTCTTAGA | 503.95623138505897 | 643.6087601042292 | 659.6679442398754 | 683.1869570223603 | 594.9935216411972 | 637.6042408631488 | 581.1267308929097 | 772.3538017844031 | 749.4595744844434 | 764.7282851234104 | 640.8865907250693 | 530.4312228166729 |
| SE_431_2_3 | SE_431 | SE_431_2 | ATTGGCTTACATGATTCTGC | 257.9388883218151 | 390.7624614918534 | 373.28515407585763 | 320.1218884333345 | 222.72484767317542 | 294.54543735525897 | 224.91756806781132 | 255.24544245016995 | 307.2429061275087 | 230.3661945652981 | 254.7113373394506 | 384.1854152129513 |
| SE_431_2_4 | SE_431 | SE_431_2 | GCTTGCAGATGGCACACTGT | 8536.91018197873 | 6917.794778635829 | 6874.1744907990615 | 7816.960096825367 | 7362.647107367542 | 6338.791073907147 | 8422.032955496132 | 7898.428413596926 | 6955.588681204555 | 7266.012219089639 | 8724.743642599706 | 6704.0935295166355 |
| SE_431_2_5 | SE_431 | SE_431_2 | AGCTTGCAGATGGCACACTG | 1877.1014898041337 | 1791.9107249485758 | 1675.8330858908212 | 1914.224788152175 | 1730.8902447743917 | 1423.3475105137954 | 1892.966517853015 | 1730.9422412083748 | 1549.5343098049213 | 2037.574410791165 | 1757.1560921527998 | 1385.8531291971717 |
| SE_431_2_6 | SE_431 | SE_431_2 | ACAGTGTGCCATCTGCAAGC | 1986.562950814652 | 2232.642810672124 | 2292.049848174777 | 2130.2419974201975 | 2215.5819370917307 | 2326.908955106546 | 1957.5361546188938 | 2473.04473129498 | 2209.3073712290225 | 2499.0358068665882 | 2423.8659521012232 | 2709.029483707034 |
| SE_432_1_12 | SE_432 | SE_432_1 | TGTCTAGCCCAGAAGAACTG | 379.3218945909046 | 378.76975167624664 | 341.6842944715522 | 396.89908931775216 | 316.0571647933632 | 176.7272624131554 | 331.4574687315114 | 256.19079594072616 | 305.4669355718584 | 117.37011811712972 | 284.05596145689884 | 162.4953417819129 |
| SE_432_1_15 | SE_432 | SE_432_1 | TCTGGGCTAGACAAGCTAGA | 797.659755482588 | 731.555298752012 | 704.1066530584299 | 679.283031553661 | 800.7488571107021 | 1095.8822889835371 | 821.110547539426 | 719.4140063132568 | 798.2987647648276 | 731.9229726062002 | 798.1737759945918 | 1330.14044058623 |
| SE_432_1_20 | SE_432 | SE_432_1 | TGGCTTCCTGAAGATAGGGA | 5.4188842084414945 | 1.9987849692677924 | 2.962580587903632 | 2.602616979132801 | 0.0 | 0.0 | 0.0 | 0.0 | 16.871720278678225 | 0.0 | 0.0 | 0.0 |
| SE_432_1_3 | SE_432 | SE_432_1 | CTGAGCAGAGAAGGCCTGAG | 1651.6759067329676 | 1673.9824117617761 | 1779.5234064674482 | 1713.8232807589493 | 1709.6783545198036 | 1588.8127414986616 | 1754.1417988063754 | 1687.4559806427903 | 1965.1114198271007 | 1845.8455843016923 | 1670.296004765153 | 1749.1462861810198 |
| SE_432_1_38 | SE_432 | SE_432_1 | CTTTTGCTAGAAAGGCTGGG | 10.837768416882989 | 12.992102300240651 | 13.825376076883616 | 16.917010364363207 | 9.545350614564661 | 3.465240439473635 | 12.913927353175769 | 0.945353490556185 | 41.73530805778298 | 0.7290069448268927 | 1.173784964697929 | 1.1606810127279494 |
| SE_433_1_13 | SE_433 | SE_433_1 | GCCCTCATTAAGTTTTACAG | 57.440172609479845 | 112.93135076363026 | 98.75268626345441 | 67.66804145745283 | 103.93826224748186 | 67.57218856973589 | 66.72195799140815 | 82.2457536783881 | 53.27911666951019 | 123.20217367574486 | 104.46686185811568 | 82.4083519036844 |
| SE_433_1_16 | SE_433 | SE_433_1 | TGGCCTGTAATTCAAGGGCA | 40.09974314246706 | 50.9690167163287 | 57.27655803280356 | 31.23140374959361 | 56.21150917465856 | 40.71657516381521 | 52.73187002546773 | 58.61191641448347 | 65.71091055906257 | 103.51898616541875 | 42.25625872912545 | 35.98111139456643 |
| SE_433_1_19 | SE_433 | SE_433_1 | GGGATCAGCTCATTAAAAGG | 187.49339361207572 | 238.8548038275012 | 226.1436515433106 | 179.58057156016326 | 159.089176909411 | 163.73261076512924 | 203.39435581251837 | 191.90675858290555 | 244.1959514019217 | 589.0376114201292 | 90.38144228174053 | 363.2931569838482 |
| SE_433_1_21 | SE_433 | SE_433_1 | TCCCCTGGCCTGTAATTCAA | 40.09974314246706 | 71.95625889364052 | 54.31397744489992 | 32.53271223916001 | 41.363185996446866 | 72.77004922894633 | 13.990087965940416 | 13.23494886778659 | 143.85361500767752 | 8.748083337922711 | 55.167893340802664 | 35.98111139456643 |
| SE_433_1_29 | SE_433 | SE_433_1 | AGAAGCAGAAGGCAAATCCG | 1713.4511867092006 | 1756.9319879863895 | 1815.074373522292 | 1737.2468335711446 | 1765.8898636944623 | 1977.785980829577 | 1666.972789172439 | 1960.6631394135277 | 1806.1620550963953 | 2268.66961230129 | 2142.1575605737203 | 1580.8475393354672 |
| SE_433_2_1 | SE_433 | SE_433_2 | GGCATCTGTGCACATCAAAT | 441.09717456713764 | 573.6512861798564 | 497.7135387678102 | 525.7286297848258 | 503.7823935464682 | 414.9625426269678 | 460.5967422632691 | 574.7749222581605 | 535.4551225285774 | 550.4002433443039 | 551.6789334080266 | 555.9662050966878 |
| SE_433_2_2 | SE_433 | SE_433_2 | ACTTATTTCCAATATAGTGA | 9.75399157519469 | 21.986634661945715 | 20.738064115325425 | 2.602616979132801 | 4.242378050917627 | 42.44919538355203 | 12.913927353175769 | 10.398888396118036 | 7.991867500426528 | 0.0 | 0.0 | 17.410215190919242 |
| SE_433_2_3 | SE_433 | SE_433_2 | GAAAAGCTGGAGAAGAAGAG | 120.29922942740117 | 168.89732990312845 | 158.9918248841616 | 127.52823197750725 | 139.9984756802817 | 248.6310015322333 | 130.21543414452233 | 152.2019119795458 | 226.43624584541828 | 209.95400011014507 | 187.80559435166865 | 291.3309341947153 |
| SE_433_2_5 | SE_433 | SE_433_2 | ACTACAGGCACATTTTATGA | 824.7541765247955 | 1085.3402383124112 | 1154.418902419782 | 903.108091759082 | 1179.3810981551003 | 569.1657421835446 | 694.1235952331976 | 1228.9595377230405 | 942.1523797725051 | 741.4000628889498 | 976.589090628677 | 846.1364582786751 |
| SE_433_2_6 | SE_433 | SE_433_2 | GATTCATCGTCACCAAACTC | 282.86575568064603 | 236.8560188582334 | 279.470102125576 | 244.64599603848328 | 308.63300320425736 | 250.36362175197013 | 288.41104422092553 | 297.78634952519826 | 301.02700918273257 | 360.129430744485 | 239.45213279837753 | 647.6600051021958 |
| SE_433_3_10 | SE_433 | SE_433_3 | TCTAGAATAGAAGCTGCCCA | 331.63571355661946 | 429.73876839257537 | 436.48687328446846 | 317.5192714542017 | 501.6612045210094 | 477.3368705374932 | 343.2952354719225 | 462.27785688197446 | 429.78487446738217 | 317.11802099969833 | 299.3151659979719 | 598.911402567622 |
| SE_433_3_11 | SE_433 | SE_433_3 | TCTTGCCTGTTCTTGCAGGA | 801.9948628493412 | 758.5388958371273 | 819.6472959866716 | 879.6845389468867 | 887.7176071545134 | 732.8983529486737 | 811.4251020245442 | 882.9601601794768 | 670.4288847580032 | 868.976278233656 | 738.3107427949974 | 651.1420481403796 |
| SE_433_3_13 | SE_433 | SE_433_3 | GGTCTAATGTTCTTTAAAAG | 40.09974314246706 | 51.968409200962604 | 48.38881626909266 | 31.23140374959361 | 48.78734758555271 | 20.791442636841808 | 40.8941032850566 | 37.8141396222474 | 62.15896944776188 | 131.22125006884067 | 9.390279717583432 | 46.42724050911798 |
| SE_433_3_2 | SE_433 | SE_433_3 | CATTAAAAAAATCTAAGGCA | 52.02128840103835 | 111.93195827899638 | 125.4159115545871 | 91.09159426964803 | 138.9378811675523 | 165.46523098486605 | 36.58946083399801 | 109.66100490451745 | 61.27098416993672 | 32.076305572383276 | 80.99116256415711 | 85.89039494186825 |
| SE_433_3_22 | SE_433 | SE_433_3 | ATTTTTGTAGAGATGAGGCC | 7.586437891818092 | 9.993924846338961 | 4.93763431317272 | 7.807850937398403 | 4.242378050917627 | 0.8663101098684087 | 24.75169409358689 | 15.12565584889896 | 11.543808611727208 | 8.748083337922711 | 14.085419576375148 | 2.321362025455899 |
| SE_434_1_15 | SE_434 | SE_434_1 | TGGTGAGGAGAGAGGCTTAT | 47.686181034285156 | 33.97934447755247 | 41.47612823065085 | 53.35364807222242 | 27.575457330964575 | 95.29411208552496 | 47.35106696164449 | 35.92343264113503 | 41.73530805778298 | 171.31663203431978 | 31.692194046844083 | 145.08512659099367 |
| SE_434_1_32 | SE_434 | SE_434_1 | ATTGGAGTTGGAGCTGGGCA | 473.6104798177866 | 557.661006425714 | 554.0025699379792 | 595.9992882214113 | 579.0846039502561 | 563.1015714144656 | 674.752704203434 | 428.2451312219518 | 550.5508722516053 | 1162.0370700540668 | 584.5449124195686 | 930.8661722078155 |
| SE_434_1_36 | SE_434 | SE_434_1 | GAGTGGAGCATGGGCCCCGG | 1026.336669078819 | 1338.1865369247869 | 1318.3483616171163 | 1159.4658642036627 | 1393.6211897264404 | 1526.4384135881362 | 1046.0281156072374 | 1693.1281015861273 | 1479.3834728567329 | 1605.2732925088176 | 1500.0971848839533 | 1912.8023089756607 |
| SE_434_1_48 | SE_434 | SE_434_1 | TCGCACAGATGGCTCTTGAT | 4171.457063658262 | 3968.587556481202 | 4005.4089548457105 | 4205.829038278606 | 4492.6783559217665 | 5001.2082642703235 | 4729.725893100625 | 5014.154913910005 | 3966.6302360450336 | 7559.802017854877 | 4758.524246885404 | 4863.253443330108 |
| SE_434_1_51 | SE_434 | SE_434_1 | CTTGGGCTAGTCCTTCCCAC | 3217.7334429725593 | 2548.4508358164353 | 2629.7840351957907 | 2895.411389285241 | 2727.849086740034 | 2965.3795060795633 | 3445.866282072401 | 2855.912894970235 | 2416.207940962287 | 3037.042932148835 | 2487.2503401949116 | 3378.742428051061 |
| SE_434_2_10 | SE_434 | SE_434_2 | GGACTCTGGCAATTGCCCTG | 99.7074694353235 | 190.88396456507417 | 177.75483527421792 | 169.17010364363205 | 127.2713415275288 | 129.94651648026132 | 137.74855843387488 | 213.64988886569782 | 109.22218917249589 | 206.3089653860106 | 119.72606639918877 | 176.42351393464833 |
| SE_434_2_11 | SE_434 | SE_434_2 | GGGAATGTGGATTTTTACTG | 118.13167574402458 | 266.83779339725027 | 244.90666193336693 | 232.9342196323857 | 220.6036586477166 | 202.71656570920763 | 157.11944946363852 | 260.91756339350707 | 137.6377180629013 | 182.25173620672317 | 476.5566956673592 | 75.44426582731671 |
| SE_434_2_36 | SE_434 | SE_434_2 | ATTGTGATGGTCCAAGCAGG | 249.26867358830876 | 305.81410029797223 | 322.9212840814959 | 260.2616979132801 | 247.11852146595177 | 304.0748485638115 | 342.2190748591579 | 410.2834149013843 | 400.48136029915156 | 439.5911877306163 | 355.6568443034725 | 287.84889115653147 |
| SE_434_2_57 | SE_434 | SE_434_2 | ACTGCGTGTGGAGCTGGGCG | 3744.4489880330725 | 3438.909539625237 | 3498.8076743141896 | 3408.126934174403 | 3523.294971287089 | 5001.2082642703235 | 3940.900163944139 | 3629.2120502451944 | 3828.104532704307 | 4933.189995643583 | 3577.6965723992876 | 4208.6293521515445 |
| SE_434_2_6 | SE_434 | SE_434_2 | TCTCGCCCGCACCTTCCTCA | 5573.864296802922 | 5136.877371018227 | 5167.728072166569 | 5291.120318576985 | 5273.275917290611 | 5889.1761268854425 | 5822.028915056743 | 5600.27407805484 | 5317.255843617117 | 7817.870476323596 | 6584.933651955382 | 6690.1653573639005 |
| SE_435_1_15 | SE_435 | SE_435_1 | GGTCAGGGAGGCCACAAAAA | 626.4230144958368 | 895.455666231971 | 726.8197708990244 | 609.0123731170754 | 760.4462656269847 | 675.7218856973589 | 533.7756639312652 | 799.7690530105325 | 812.5065292100304 | 542.3811669512081 | 669.0574298778196 | 940.151620309639 |
| SE_435_1_16 | SE_435 | SE_435_1 | GAGTGGGTCAGGGAGAGCCT | 1269.102681616998 | 1322.1962571706447 | 1301.560404952329 | 1327.3346593577285 | 1305.5918451698997 | 1302.0640951322184 | 1333.3629992153983 | 1225.1781237608159 | 1165.9246697844478 | 1490.8192021709954 | 1584.6097023422042 | 1078.2726608242651 |
| SE_435_1_26 | SE_435 | SE_435_1 | GGAGAGGTTTACCAGCAGAA | 766.2302270736274 | 1132.3116850902043 | 1149.4812681066091 | 960.3656653000036 | 1149.684451798677 | 892.299413164461 | 901.8225934967746 | 1315.9320588542096 | 1218.315801176133 | 1015.5066741438615 | 1126.8335661100118 | 986.578860818757 |
| SE_435_1_28 | SE_435 | SE_435_1 | GTGAGGGGAAAGCACTGTGG | 5940.1808692935665 | 6186.239479883818 | 6028.851496383892 | 6293.127855543113 | 6567.201222820487 | 6494.72689368346 | 6347.195294085891 | 7105.276835020287 | 6058.723550601134 | 6493.264857573133 | 6957.023485764626 | 7029.084213080462 |
| SE_435_1_45 | SE_435 | SE_435_1 | GGCCTTGTGAAATCTTCTAG | 37.93218945909046 | 46.97144677779312 | 55.301504307534465 | 45.545797134824014 | 69.99923784014085 | 30.320853845394307 | 37.665621446762664 | 114.38777235729839 | 73.70277805948909 | 26.244250013768134 | 79.81737759945918 | 171.7807898837365 |
| SE_436_1_11 | SE_436 | SE_436_1 | CAACAGTTCACAGCAAAGGG | 720.7115997227188 | 885.461741385632 | 871.9862197063023 | 802.9073380624691 | 867.5663114126547 | 846.3849773414353 | 667.2195799140815 | 813.0040018783191 | 794.7468236535269 | 1280.8652020608504 | 726.5728931480181 | 869.3500785332342 |
| SE_436_1_32 | SE_436 | SE_436_1 | GGTCCGGCCAGCAGGAACCA | 457.3538271924621 | 439.7326932389143 | 481.9131089656575 | 460.66320530650574 | 462.4192075500213 | 265.0908936197331 | 469.2060271653863 | 315.7480658457658 | 396.9294191878509 | 460.7323891305962 | 311.0530156449512 | 236.7789265965017 |
| SE_436_1_35 | SE_436 | SE_436_1 | TTCTGAGCCCCAATTCTGAG | 4.335107366753196 | 7.9951398770711695 | 6.912688038441808 | 3.9039254686992013 | 10.605945127294067 | 0.8663101098684087 | 0.0 | 15.12565584889896 | 0.0 | 0.7290069448268927 | 50.47275348201095 | 0.0 |
| SE_436_1_5 | SE_436 | SE_436_1 | GGGCGTGCGGAGGTCAGCGA | 4982.1221412411105 | 4696.145285294679 | 4490.284644399272 | 4920.2473990505605 | 4912.673782962612 | 4820.149451307826 | 5299.014857253124 | 5310.050556454091 | 4741.841383586407 | 5199.277530505398 | 5340.721589375577 | 5520.198896534128 |
| SE_436_1_9 | SE_436 | SE_436_1 | ACCTGTGTATGACACTAGAA | 311.04395356454177 | 445.7290481467177 | 452.2873030866212 | 288.8904846837409 | 597.114710666656 | 669.6577149282799 | 327.1528262804528 | 644.7310805593182 | 384.49762529829854 | 524.8850002753627 | 565.7643529844017 | 523.4671367403052 |
| SE_437_1_11 | SE_437 | SE_437_1 | AGGGACTTGAGCATCCACAA | 2477.5138600994514 | 2600.4192450173978 | 2611.021024805734 | 2587.001277258004 | 2387.3982481538947 | 2829.368818830223 | 2675.3352833329136 | 2300.9903960137544 | 2712.795023755894 | 2768.768376452538 | 2503.6833297006824 | 3211.604362218236 |
| SE_437_1_12 | SE_437 | SE_437_1 | GCTACTCAAGAGGCCAAGAC | 270.9442104220747 | 312.8098476904095 | 348.59698250999406 | 318.8205799437681 | 308.63300320425736 | 336.1283226289426 | 273.34479564222045 | 276.9885727329622 | 388.0495664095992 | 353.56836824104295 | 248.84241251596094 | 412.04175951842205 |
| SE_437_1_23 | SE_437 | SE_437_1 | GGTGGGGGCATGGGCTGCAG | 318.63039145635986 | 102.9374259172913 | 225.15612468067604 | 235.53683661151848 | 88.02934455654076 | 573.4972927328865 | 443.3781724590348 | 292.11422858186114 | 759.2274125405202 | 34.26332640686395 | 136.15905590495976 | 232.1362025455899 |
| SE_437_1_3 | SE_437 | SE_437_1 | TGTATTCTTTTTTTGCCCCT | 83.45081680999901 | 106.9349958558269 | 120.47827724141437 | 89.79028578008163 | 104.99885676021127 | 122.14972549144564 | 76.40740350628997 | 142.74837707398393 | 97.67838056076867 | 189.5418056549921 | 115.03092654039705 | 74.28358481458876 |
| SE_437_1_31 | SE_437 | SE_437_1 | AGATTGATTCCCACGAGCAG | 123.55055995246607 | 196.88031947287755 | 179.729888899487 | 188.68973098712806 | 186.6646342403756 | 206.1818061486813 | 195.86123152316583 | 189.070698111237 | 188.252878898936 | 330.9691529514093 | 302.83652089206566 | 136.96035950189804 |
| SE_437_2_10 | SE_437 | SE_437_2 | AGGACCCAAGAAACTACAAG | 565.7315113612921 | 767.5334281988323 | 712.9943948221407 | 661.0647126997314 | 704.2347564523261 | 663.5935441592011 | 599.4214613099086 | 846.0913740477856 | 662.4370172575767 | 673.6024170200488 | 771.1767218065394 | 551.323481045776 |
| SE_437_2_12 | SE_437 | SE_437_2 | CTTTGATCCTGGATACTCAG | 81.28326312662242 | 54.96658665486429 | 59.25161175807264 | 67.66804145745283 | 51.96913112374093 | 90.96256153618292 | 125.91079169346375 | 149.36585150787724 | 7.991867500426528 | 48.114458358574915 | 93.90279717583432 | 20.89225822910309 |
| SE_437_2_18 | SE_437 | SE_437_2 | CATCTGCCAACAGAGAGCAC | 1.083776841688299 | 0.0 | 0.0 | 1.3013084895664004 | 0.0 | 0.0 | 0.0 | 0.0 | 7.103882222601358 | 0.0 | 0.0 | 0.0 |
| SE_437_2_5 | SE_437 | SE_437_2 | GTGCACTCTTTACCACACTG | 761.8951197068741 | 756.5401108678594 | 810.7595542229607 | 730.0340626467506 | 804.9912351616197 | 665.3261643789378 | 732.8653772927249 | 774.2445087655155 | 602.0540183654651 | 618.9268961580318 | 1061.1016080869279 | 802.0305797950131 |
| SE_437_2_71 | SE_437 | SE_437_2 | AGGATTTTAGCTTAGCTCCC | 89.95347786012881 | 132.9192004563082 | 106.65290116453076 | 110.61122161314404 | 137.87728665482288 | 87.49732109670929 | 59.18883370205561 | 150.31120499843342 | 194.4687758437122 | 145.80138896537852 | 149.070690516637 | 131.15695443825828 |
| SE_438_1_10 | SE_438 | SE_438_1 | TCTCTTCCAGAGGAAGTCTG | 17.340429467012783 | 19.987849692677923 | 14.81290293951816 | 29.93009526002721 | 12.727134152752882 | 0.8663101098684087 | 13.990087965940416 | 46.32232103725307 | 20.423661389978903 | 61.23658336545898 | 0.0 | 45.26655949639003 |
| SE_438_1_13 | SE_438 | SE_438_1 | GTTTGAGTGGAAGGAGCCCG | 124.63433679415438 | 141.91373281801324 | 111.59053547770347 | 127.52823197750725 | 161.21036593486983 | 134.27806702960336 | 164.65257375299106 | 126.67736773452879 | 97.67838056076867 | 289.41575709627637 | 181.936669528179 | 90.53311899278006 |
| SE_438_1_21 | SE_438 | SE_438_1 | ATGAGGCCAAAGGTTCTGAG | 2111.197287608806 | 2467.5000445610895 | 2328.5883420922546 | 2135.447231378463 | 2490.275915888647 | 1799.3260981966848 | 2215.814701682409 | 2633.7548246895312 | 2554.7336443030135 | 1990.917966322244 | 2812.388775416238 | 2317.879982417715 |
| SE_438_1_5 | SE_438 | SE_438_1 | TGCTGTGGACAGATACCATG | 2804.8144662893174 | 2615.4101322869064 | 2422.4033940425365 | 2702.8177328294137 | 2682.2435226926696 | 3194.0853750848228 | 2895.9482089496664 | 2528.820587237795 | 2387.7924120718817 | 2263.566563687502 | 2380.4359084074 | 3331.154506529215 |
| SE_438_1_9 | SE_438 | SE_438_1 | CTCCAAACTGAAGCTCACCA | 160.39897256986825 | 179.89064723410132 | 193.55526507637063 | 180.88188004972966 | 258.7850611059753 | 64.97325824013066 | 189.40426784657797 | 181.50787018678753 | 179.3730261206843 | 58.320555586151414 | 186.63180938697073 | 103.3006101327875 |
| SE_438_2_18 | SE_438 | SE_438_2 | TCACCCAAGCTGAACTGAAC | 702.2873934140177 | 659.5990398583715 | 641.8924607124536 | 614.217607075341 | 594.9935216411972 | 656.6630632802538 | 802.815817122427 | 528.4526012209075 | 566.5346072524583 | 406.7858752134061 | 527.0294491493702 | 597.750721554894 |
| SE_438_2_19 | SE_438 | SE_438_2 | CACCCAAGCTGAACTGAACT | 1062.1013048545328 | 1061.3548186811977 | 924.3251434259332 | 1151.6580132662643 | 1000.1406255038305 | 1211.9678437059038 | 1181.6243528155828 | 1011.528234895118 | 1054.9265100563016 | 1175.1591950609509 | 829.8659700414358 | 1425.3162836299218 |
| SE_438_2_20 | SE_438 | SE_438_2 | AGCCCAGTTCAGTTCAGCTT | 684.9469639470049 | 763.5358582602967 | 720.8946097232172 | 624.6280749918722 | 881.354040078137 | 624.6095892151227 | 750.0839470969593 | 680.6545132004532 | 603.8299889211155 | 603.6177503166671 | 570.4594928431935 | 662.7488582676591 |
| SE_438_2_21 | SE_438 | SE_438_2 | GAGCCCAGTTCAGTTCAGCT | 958.0587280524562 | 856.479359331249 | 838.4103063767279 | 953.8591228521715 | 912.1112809472899 | 628.9411397644648 | 891.0609873691282 | 1274.3365052697375 | 639.3494000341223 | 823.7778476543887 | 1135.0500608628975 | 844.9757772659472 |
| SE_438_2_28 | SE_438 | SE_438_2 | ATCTGTAGTCCTAGCTGCTC | 285.0333093640226 | 232.85844891969782 | 268.607306636596 | 339.6415157768305 | 252.42149402959882 | 142.07485801841904 | 370.1992507910387 | 368.6878613169122 | 310.79484723880944 | 347.0073057376009 | 375.6111887033373 | 217.04734938012655 |
| SE_439_1_13 | SE_439 | SE_439_1 | GTCAAGTCAGAACTCAGAAA | 218.92292202103638 | 247.84933618920624 | 335.75913329574496 | 312.3140374959361 | 164.39214947305805 | 402.83420108881006 | 347.59987792298114 | 193.79746556401793 | 337.4344055735645 | 465.8354377443844 | 66.90574298778195 | 206.601220265575 |
| SE_439_1_16 | SE_439 | SE_439_1 | AGACTGGGCCTCCGGAGTGA | 277.4468714722045 | 359.78129446820265 | 416.73633603177757 | 364.3663770785921 | 452.8738569354567 | 510.2566547124927 | 262.583189514574 | 377.1960427319178 | 328.5545527953128 | 424.2820418892515 | 451.90721140870266 | 384.1854152129513 |
| SE_439_1_17 | SE_439 | SE_439_1 | GGAAATGCAATTTTACCAAG | 41.18351998415536 | 42.973876839257535 | 57.27655803280356 | 32.53271223916001 | 32.87842989461161 | 33.78609428486794 | 43.0464245105859 | 34.03272566002266 | 62.15896944776188 | 68.5266528137279 | 73.94845277596953 | 29.017025318198737 |
| SE_439_1_4 | SE_439 | SE_439_1 | TATACAGTAGCATGCACAGG | 796.5759786408997 | 935.4313656173268 | 889.7617032337241 | 835.4400503016291 | 880.2934455654076 | 1032.641650963143 | 776.9879624160754 | 831.9110716894428 | 1014.9671725541691 | 917.090736592231 | 807.5640557121752 | 918.098681067808 |
| SE_439_1_8 | SE_439 | SE_439_1 | GCTGTTGTGCGCCGGCTAGA | 675.1929723718102 | 764.5352507449306 | 858.1608436294188 | 745.6497645215475 | 668.1745430195263 | 1003.1871072276173 | 689.818952782139 | 931.1731881978423 | 811.6185439322052 | 978.32731995769 | 949.5920364406246 | 795.0664937186453 |
| SE_439_2_10 | SE_439 | SE_439_2 | AGTCAGGTGCAGCACAGTGA | 1618.0788246406303 | 1727.9496059320065 | 1768.6606109784684 | 1884.2946928921479 | 1565.4375007886044 | 1831.379572261816 | 1322.6013930877518 | 1505.9481104560027 | 1760.8748059273116 | 2006.2271121636086 | 1265.3401919443675 | 1482.1896532535914 |
| SE_439_2_23 | SE_439 | SE_439_2 | TAGGTGGGTTTCCCACAGCA | 409.667646158177 | 478.70900013963626 | 540.1771938610956 | 465.8684392647714 | 521.8125002628681 | 586.4919443809127 | 496.11004248450246 | 541.687550088694 | 356.97008168571824 | 889.3884726888091 | 912.0309175702909 | 526.949179778489 |
| SE_439_2_27 | SE_439 | SE_439_2 | GATCCTTTATGGGGTTACCT | 19.50798315038938 | 24.984812115847404 | 30.613332741670867 | 32.53271223916001 | 26.51486281823517 | 10.395721318420904 | 11.837766740411123 | 9.45353490556185 | 81.69464555991561 | 44.46942363444045 | 1.173784964697929 | 35.98111139456643 |
| SE_439_2_3 | SE_439 | SE_439_2 | TCTTCTCTTTGGTCTTGCCA | 43.351073667531956 | 87.94653864778286 | 71.10193410968716 | 57.25757354092162 | 43.48437502190568 | 53.71122681184134 | 78.55972473181926 | 85.08181415005664 | 45.28724916908366 | 112.99607644816837 | 76.29602270536539 | 34.820430381838484 |

| | | | | | | | | | | | | | | | |
|---|---|---|---|---|---|---|---|---|---|---|---|---|---|---|---|
| SE_439_2_7 | SE_439 | SE_439_2 | TTACTACTCACAGGTCTGGA | 68.27794102636283 | 73.95504386290831 | 77.02709528549444 | 75.47589239485123 | 51.96913112374093 | 285.8823362565749 | 101.15909759987686 | 177.72645622456278 | 69.26285167036325 | 223.80513206185606 | 145.5493356225432 | 141.60308355280984 |
| SE_439_3_0 | SE_439 | SE_439_3 | CTGCTTTCTAAGACCCTGCG | 122.46678311077778 | 98.93985597875572 | 127.39096527985618 | 130.13084895664005 | 143.1802592184699 | 120.41710527170882 | 110.8445431147587 | 161.65544688510764 | 27.527543612580263 | 126.11820145505243 | 197.19587406925208 | 87.05107595459621 |
| SE_439_3_12 | SE_439 | SE_439_3 | CTGTGTCCTGTGTCCCCGAA | 534.3019829523314 | 449.7266180852533 | 479.9380552403884 | 464.56713077520493 | 512.2671496483034 | 622.0106588855175 | 507.9478092249136 | 524.6711872586827 | 488.3919028038434 | 554.7742850132653 | 693.706914136476 | 302.9377443219948 |
| SE_439_3_26 | SE_439 | SE_439_3 | GATGGTGAATAAATGCTCAT | 2322.5337717380244 | 2330.5832741662457 | 2434.253716394151 | 2407.4207056978407 | 2488.1547268631884 | 2950.6522342118 | 2436.427627299162 | 2507.0774569550026 | 2380.6885298492803 | 2330.635202611576 | 2731.397612852081 | 2769.3848963688874 |
| SE_439_3_33 | SE_439 | SE_439_3 | GTGGGAGGTCCGCAGCGCCA | 1226.8353847911544 | 1306.2059774165023 | 1350.9367480840563 | 1462.670742272634 | 1320.4401683481115 | 1341.9143601861651 | 1197.7667620070526 | 1477.587505739317 | 1398.5768125746424 | 1180.2622436747392 | 1155.0044052627622 | 1529.7775747754374 |
| SE_439_4_12 | SE_439 | SE_439_4 | TGGACCTGAGGCTCTCCCAG | 86.70214733506391 | 104.9362108865591 | 105.66537430189621 | 115.81645557140963 | 64.69626527649382 | 70.17111889934111 | 100.08293698711222 | 117.22383282896693 | 103.0062922277197 | 89.6678542137078 | 72.7746678112716 | 62.67677468730927 |
| SE_439_4_21 | SE_439 | SE_439_4 | ACTTAGGAAAGAAAGCCTCA | 570.0666187280452 | 625.619695380819 | 564.8653654269592 | 644.1477023353682 | 550.4485521065621 | 628.9411397644648 | 657.5341343991996 | 672.1463317854475 | 758.339427262695 | 804.0946601440626 | 741.8320976890911 | 1101.486281078824 |
| SE_439_4_3 | SE_439 | SE_439_4 | GCCAGTGTCTCTCATGAGGA | 622.0879071290835 | 697.5759542744595 | 694.2313844320845 | 735.2392966050162 | 721.2042686559965 | 492.93045251512456 | 525.166379029148 | 630.5507782009754 | 707.7242664266603 | 833.2549379371383 | 626.8011711486942 | 612.8395747203573 |
| SE_439_4_31 | SE_439 | SE_439_4 | AGCCAGGACTCAGTTTTCAA | 149.56120415298525 | 225.86270152726053 | 252.80687683444327 | 165.26617817493286 | 191.9676068040226 | 263.3582733999963 | 129.1392735317577 | 307.2398844307601 | 328.5545527953128 | 209.95400011014507 | 204.23858385743966 | 413.20244053115 |
| SE_439_4_4 | SE_439 | SE_439_4 | GGGCAGGCTGAGAAGCCGGT | 547.307305052591 | 579.6476410876597 | 596.4662250312646 | 654.5581702518995 | 687.2652442486556 | 723.3689417401213 | 520.8617365780893 | 675.9277457476722 | 813.3945144878555 | 687.4535489717598 | 627.9749561133921 | 586.1439114276145 |
| SE_439_5_15 | SE_439 | SE_439_5 | TGGAGTCTGTTCATGCCGCG | 274.19554094713965 | 311.8104552057756 | 357.48472427370496 | 406.0082487447169 | 355.29916176435125 | 260.759343070391 | 321.7720232166296 | 303.4584704685354 | 504.37563780469645 | 249.3203751307973 | 271.1443268452216 | 271.59935697834015 |
| SE_439_5_17 | SE_439 | SE_439_5 | ATGACATCATACCAGCCACG | 2128.5377170758193 | 1940.8202051590265 | 1898.0266299835937 | 2291.604250126431 | 2042.7050315168374 | 1406.8876184262958 | 2419.2090574949275 | 1837.7671856412237 | 1900.2884945458634 | 2786.2645431283836 | 2065.8615378683553 | 1637.7209089591368 |
| SE_439_5_20 | SE_439 | SE_439_5 | AGGCCAGTAGAGGGAGCTGT | 3478.9236618194395 | 3800.689619062707 | 4031.084653274209 | 3777.6985452112604 | 3646.3239347637004 | 3970.2992335269173 | 3609.4426952126278 | 3611.250333924627 | 4116.699747997487 | 3612.229411617253 | 3577.6965723992876 | 3622.48544072393 |
| SE_439_5_30 | SE_439 | SE_439_5 | GCTCTTTGCAGGGCAGGCAG | 508.2913387518122 | 523.6816619481616 | 529.3143983721156 | 446.34881192127534 | 428.48018314268035 | 672.2566452578852 | 455.21593919944587 | 566.2667408431548 | 414.6891247443543 | 319.305041834179 | 676.1001396660072 | 392.3101823020469 |
| SE_439_5_9 | SE_439 | SE_439_5 | GCATTTACCATGCAGATGGA | 619.920353445707 | 614.6263780498462 | 521.4141834710392 | 625.9293834814387 | 623.6295734848911 | 529.3154771295978 | 733.9415379054896 | 712.7965318793634 | 609.1579005880665 | 548.9422294546501 | 605.6730417841313 | 805.5126228331969 |
| SE_439_6_18 | SE_439 | SE_439_6 | GTGCAGCCTCAACCTAAGGG | 980.8180417279106 | 864.4744992083203 | 902.5995524479732 | 991.5970690495972 | 858.0209607980901 | 931.2833681085394 | 1070.7798097008242 | 883.905513670033 | 715.7161339270868 | 1650.471723088085 | 705.4447637834553 | 1540.2237038899889 |
| SE_439_6_20 | SE_439 | SE_439_6 | GAGATGATTAAAGGACCCCA | 324.04927566480137 | 425.74119845403976 | 430.5617121086612 | 360.4624516098929 | 389.2381861716923 | 414.0962325170994 | 348.67603853574576 | 383.8135171658111 | 403.1453161326271 | 423.5530349444246 | 515.2915995023908 | 593.1079975039821 |
| SE_439_6_24 | SE_439 | SE_439_6 | TGCAGTTCACTCTTCAACAG | 669.7740881633687 | 719.5625889364053 | 718.919555997948 | 707.9118183241219 | 757.2644820887964 | 667.9250947085432 | 746.8554652586654 | 761.0095598977289 | 725.4839719831638 | 911.9876879784428 | 697.2282690305699 | 927.3841291696316 |
| SE_439_6_29 | SE_439 | SE_439_6 | GGCTGCCTTGACATGTTAAA | 396.6623240579174 | 309.81167023650784 | 318.9711766309577 | 344.8467497350961 | 421.05602155357445 | 372.51334724341575 | 285.1825623826316 | 541.687550088694 | 450.2085358573611 | 276.2936320893923 | 373.26361877394146 | 511.8603266130257 |
| SE_439_6_32 | SE_439 | SE_439_6 | TGTCAAGGCAGCCATCCAAA | 2339.8742012050375 | 2521.46723873132 | 2411.5405985535567 | 2540.1541716336137 | 2785.1211904274223 | 1718.759257978923 | 2401.990487690693 | 3052.5464210059213 | 2693.25934764374 | 2986.0124460109523 | 3098.7923068025325 | 2330.6474735577226 |
| SE_44_1_12 | SE_44 | SE_44_1 | TGACACCTCCTCCACCAGAG | 1407.8261173531002 | 1500.0881194354781 | 1533.629217671447 | 1368.9765310238533 | 1670.4363575488155 | 1301.19778502235 | 1071.8559703135888 | 1803.734459981201 | 1217.4278158983077 | 1189.010327012662 | 1601.0426918479752 | 1347.5506557771494 |
| SE_44_1_13 | SE_44 | SE_44_1 | AGCTACCTCTCTGGTGGAGG | 102.95879996038839 | 150.9082651797183 | 127.39096527985618 | 109.30991312357764 | 102.87766773475245 | 9.529411208552496 | 74.25508228076067 | 206.08706094124832 | 113.66211556162173 | 4.374041668961356 | 254.7113373394506 | 30.177706330926686 |
| SE_44_1_14 | SE_44 | SE_44_1 | TCCTCCACCAGAGAGGTAGC | 5.4188842084414945 | 0.0 | 1.975053725269088 | 11.711776406097604 | 7.4241615891058474 | 0.0 | 19.370891029763655 | 26.46989773557318 | 0.0 | 0.0 | 31.692194046844083 | 0.0 |
| SE_44_1_40 | SE_44 | SE_44_1 | CGGTGCCCAGGTCCAGCAGG | 981.9018185695988 | 1071.3487435275367 | 971.7264328323913 | 1017.6232388409252 | 1026.6554883220658 | 662.7272340493327 | 1026.6572245774737 | 1064.4680303662644 | 784.0910003196249 | 798.9916115302743 | 886.2076483469364 | 510.69964560029774 |
| SE_44_1_41 | SE_44 | SE_44_1 | TGGCGGTGCCCAGGTCCAGC | 158.23141888649164 | 132.9192004563082 | 175.77978154894885 | 191.29234796626088 | 136.81669214209347 | 132.54544680986655 | 190.4804284593426 | 165.43686084733238 | 124.31793889552377 | 216.5150626135871 | 119.72606639918877 | 58.034050636397474 |
| SE_44_2_13 | SE_44 | SE_44_2 | GAGTTTGCTGGCGGAGGCCG | 147.39365046960864 | 232.85844891969782 | 191.58021135110155 | 167.86879515406565 | 238.63376536411653 | 150.73795911710312 | 108.6922218892294 | 325.20160075132765 | 215.78042251151626 | 83.10679171026577 | 318.0957254331388 | 159.01329874372908 |
| SE_44_2_14 | SE_44 | SE_44_2 | GCTCATGAGTTTGCTGGCGG | 688.1982944720698 | 1011.3851944495029 | 1049.7410549805204 | 779.4837852502739 | 976.8075462237836 | 876.7058311868296 | 786.6734079309573 | 881.0694531983644 | 893.3131894921208 | 1307.1094520746185 | 883.8600784175405 | 691.7658835858579 |
| SE_44_2_15 | SE_44 | SE_44_2 | CCGGCTCATGAGTTTGCTGG | 280.6982019972694 | 278.830503212857 | 294.28300506509413 | 253.7551554654481 | 291.6634910005869 | 187.12298373157628 | 161.42409191469713 | 323.3108937702153 | 252.18781890234823 | 294.5188057100646 | 180.76288456348107 | 255.34982280014887 |
| SE_44_2_21 | SE_44 | SE_44_2 | GGCCGGCCCAGGACTCGGCG | 183.15828624532253 | 202.87667438068092 | 193.55526507637063 | 217.31851775758886 | 162.27096044759924 | 100.4919727447354 | 257.20238645075074 | 185.28928414901227 | 191.80482001023668 | 178.6067014825887 | 115.03092654039705 | 90.53311899278006 |
| SE_44_2_5 | SE_44 | SE_44_2 | TCGGTTCAGTCCAAGTAGGG | 446.51605877557915 | 432.73694584647706 | 464.1376254382357 | 347.4493667142289 | 426.3589941172215 | 495.5293828447298 | 413.2456753016246 | 384.7588706563673 | 404.0333014104522 | 494.2667085926332 | 278.1870366334092 | 430.61265572206923 |
| SE_44_3_10 | SE_44 | SE_44_3 | GGCCTCTTGGTCAAGCACCA | 43.351073667531956 | 41.97448435462364 | 28.63827901640178 | 23.423552812195208 | 33.939024407341016 | 13.86096175789454 | 47.35106696164449 | 46.32232103725307 | 32.855455279531284 | 32.80531251721017 | 58.68924823489645 | 2.321362025455899 |
| SE_44_3_17 | SE_44 | SE_44_3 | GAGGGTTCAAAAGGAATGAG | 701.2036165723293 | 969.4107100948793 | 969.7513791071223 | 882.2871559260195 | 902.5659303327251 | 768.4170674532785 | 707.0375225863734 | 759.1188529166166 | 951.0322325507568 | 807.0106879233701 | 894.4241430998219 | 689.444521560402 |
| SE_44_3_18 | SE_44 | SE_44_3 | AGACTTGTAAGAGAGGCCGT | 333.80326723999605 | 321.8043800521146 | 335.75913329574496 | 330.5323563498657 | 267.2698172078105 | 436.620295373678 | 417.5503177526832 | 277.93392622351837 | 237.09206917932033 | 473.12550719265334 | 407.30338275018136 | 261.15322786378863 |
| SE_44_3_26 | SE_44 | SE_44_3 | ATAGGGTCCTAGGTGAGATG | 709.8738313058358 | 654.602077435202 | 669.5432128662209 | 723.5275201989186 | 638.4778966631029 | 623.7432791052543 | 792.0542109947805 | 640.9496665970935 | 705.0603105931848 | 618.1978892132049 | 665.5360749837257 | 756.764020298623 |
| SE_44_3_6 | SE_44 | SE_44_3 | GGATGAGAAGAGAGTGATGG | 993.82336382817 | 1145.303787390445 | 1149.4812681066091 | 974.6800586852339 | 991.6558694019953 | 920.0213366802501 | 1042.7996337689433 | 980.3315697067638 | 792.9708530978767 | 1401.8803549021145 | 858.0368091941862 | 891.4030177750651 |
| SE_44_4_15 | SE_44 | SE_44_4 | AGCCAAAGCAGGGTTAAAAG | 486.6158019180462 | 578.6482486030259 | 622.1419234597628 | 512.7155448891617 | 558.9333082083973 | 604.6844566881493 | 548.8419125099703 | 795.9876390483078 | 574.5264747528848 | 536.549111392593 | 821.6494752885503 | 471.23649116754746 |
| SE_44_4_18 | SE_44 | SE_44_4 | ACTGCAATGCTGAATCTCAT | 2580.4726600598397 | 2388.548038275012 | 2516.218445992818 | 2584.3986602788714 | 2658.9104434126225 | 2391.8822133466765 | 2792.63679012426 | 2026.8378837524606 | 2600.908878749922 | 2520.906015211395 | 2569.4152877237666 | 1819.9478279574248 |
| SE_44_4_26 | SE_44 | SE_44_4 | GTACCAGGCTGACCTGGCTC | 873.524134400769 | 854.4805743619812 | 849.2731018657079 | 841.946592749461 | 843.1726376198784 | 1197.2405718381408 | 929.8027694286554 | 899.031169518932 | 741.4677069840168 | 1011.861639419727 | 781.7407864888207 | 970.3293266405657 |
| SE_44_4_5 | SE_44 | SE_44_4 | CTACATGGTGTGCTCAGCTT | 1453.344744704009 | 1416.1391507262308 | 1316.3733078918472 | 1275.2823197750724 | 1358.62157080637 | 946.0106399763023 | 1308.6113051218113 | 1331.0577147031086 | 1012.3032167206935 | 1403.3383687917683 | 1785.32693130555 | 1038.8095063915148 |
| SE_44_4_51 | SE_44 | SE_44_4 | TAACCTGCCTCCCTAACATG | 2622.7399568856836 | 2366.561403613066 | 2390.8025344382313 | 2497.2109914779226 | 2350.2774402083655 | 3116.117465196666 | 2518.2158338692752 | 2126.1000002608603 | 2579.5972320821184 | 2415.9290151563223 | 2351.091284289952 | 2956.254539418087 |
| SE_44_5_27 | SE_44 | SE_44_5 | ACCTCAGGACAAAGAATGAA | 463.8564882425919 | 543.6695116408396 | 598.4412787565337 | 543.9469486387554 | 633.1749240994558 | 688.716537345385 | 449.83513613562263 | 383.8135171658111 | 659.7730614241011 | 401.68282659961784 | 584.5449124195686 | 660.4274962422032 |
| SE_44_5_3 | SE_44 | SE_44_5 | TTAAACTCTTACAACAAAAA | 2.167553683376598 | 3.9975699385355847 | 5.925161175807264 | 5.205233958265602 | 6.363567076376441 | 46.78074593289407 | 3.228481838293942 | 6.617474433893295 | 49.72717555820951 | 18.954180565499207 | 16.432989505771005 | 54.55200759821362 |
| SE_44_5_34 | SE_44 | SE_44_5 | ACTCCTGGAGCAAAGCCACT | 239.51468201311405 | 248.84872867384016 | 265.64472604869235 | 236.83814510108488 | 363.7839178661865 | 280.68447559736444 | 305.62961402515987 | 385.7042241469235 | 293.9231269601312 | 231.09520151012498 | 451.90721140870266 | 225.1721164692222 |
| SE_44_5_4 | SE_44 | SE_44_5 | TCTAGTCCCAACATTTCGGG | 152.81253467805016 | 170.89611487239625 | 178.74236213685248 | 189.99103947669445 | 109.24123481112889 | 157.66843999605038 | 181.87114355722542 | 155.98332594177052 | 209.56452556674006 | 192.45783343429966 | 160.80854016361627 | 257.67118482560477 |
| SE_44_5_7 | SE_44 | SE_44_5 | CGTATGTTTTCACTTTTGAA | 816.0839617912891 | 1279.222380331387 | 1239.3462126063528 | 1052.7585680592179 | 1263.1680646607235 | 1452.8020542493214 | 902.8987541095393 | 1263.9376168736194 | 1340.8577695160063 | 1450.7238202055164 | 1367.4594838730873 | 1286.0345621025679 |
| SE_44_6_24 | SE_44 | SE_44_6 | CTAGGCAGTGTGCAGAGTGC | 8597.601685113275 | 6536.026849505681 | 6412.011919086094 | 7315.956328342303 | 6704.01791496258 | 6363.91406709333 | 8409.119028142955 | 7098.659360586394 | 5712.409292249317 | 5514.937537615443 | 7059.142777693345 | 5295.026780064905 |
| SE_44_6_36 | SE_44 | SE_44_6 | AGAGGGACACACTAAGCCTG | 7329.582780337965 | 6827.849455018779 | 6526.5650351517015 | 6883.921909806259 | 6695.533158860745 | 6571.828493461749 | 7442.726797880302 | 6712.95513643947 | 6342.878839505188 | 7831.7216082753075 | 7631.949840465934 | 6093.575316821734 |
| SE_44_6_38 | SE_44 | SE_44_6 | GATTATATCCTGGCTACCCC | 34.680858934025565 | 57.964764108765976 | 52.338923719630834 | 50.751031093089615 | 14.848323178211695 | 110.02138395328791 | 36.58946083399801 | 2.836060471668555 | 34.63142583518162 | 84.56480559991955 | 9.390279717583432 | 52.23064557275772 |
| SE_44_6_8 | SE_44 | SE_44_6 | CTTGTTATCTGTGTAAACAT | 265.5253262136332 | 193.88214201897586 | 203.43053370271608 | 182.18318853929605 | 236.5125763386577 | 194.91977472039196 | 306.70577463792455 | 162.60080037566382 | 126.98189472899928 | 236.92725706874012 | 277.01325166871123 | 197.3157721637514 |
| SE_44_6_9 | SE_44 | SE_44_6 | GCTATGCTAATTTTAGCTGA | 260.10644200519175 | 362.77947192210434 | 306.13332741670865 | 342.24413275596334 | 381.81402458258646 | 453.94649757104617 | 329.3051475059821 | 300.62240999686685 | 429.78487446738217 | 927.2968338198075 | 320.4432953625346 | 538.5559899057686 |
| SE_44_7_11 | SE_44 | SE_44_7 | CTCTCATGCCTAGCCCATCC | 99.7074694353235 | 110.93256579436247 | 102.70279371399258 | 115.81645557140963 | 96.51410065837601 | 269.4224441690751 | 105.46374005093546 | 149.36585150787724 | 107.44621861684554 | 167.67159731018532 | 43.43004369382337 | 189.19100507465575 |
| SE_44_7_12 | SE_44 | SE_44_7 | TGAATGCACCAGGATGGGCT | 1320.040193176348 | 1265.2308855465126 | 1339.0864257324417 | 1329.9372763368613 | 1408.4695129046522 | 1217.1657043651142 | 1508.7771790960358 | 1485.1503336637666 | 1308.002314236475 | 1318.7735631918488 | 1095.1413720631679 | 1165.3237367788613 |
| SE_44_7_13 | SE_44 | SE_44_7 | AATCATGAATGCACCAGGAT | 548.3910818942793 | 536.6737642484022 | 530.3019252347501 | 495.7985345247986 | 463.47980206275076 | 498.99462328420344 | 561.755839863146 | 481.1849266930982 | 631.3575325336957 | 653.9192295097228 | 545.810008584537 | 393.4708633147749 |
| SE_44_7_14 | SE_44 | SE_44_7 | AAATCATGAATGCACCAGGA | 48.769957875973454 | 71.95625889364052 | 58.264084895438096 | 68.96934994701923 | 57.27210368738796 | 96.16042219539337 | 77.48356411905462 | 65.22939084837677 | 87.9105425046918 | 156.00748619295504 | 39.90868879972959 | 110.26469620915519 |
| SE_44_7_9 | SE_44 | SE_44_7 | TGGGCTAGGCATGAGAGCCT | 293.703524097529 | 241.8529812814029 | 284.40773643874866 | 229.03029416368648 | 270.45160074599875 | 275.486614938154 | 252.89774399969215 | 278.8792797140746 | 272.61148029232714 | 239.8432848480477 | 289.9248862803885 | 222.85075444376628 |
| SE_44_8_1 | SE_44 | SE_44_8 | TACCTCACATGAGCACCCAT | 1773.0589130020571 | 1531.069286459129 | 1557.329862374676 | 1728.1376741441798 | 1733.0114337998507 | 1949.1977472039196 | 1612.088597921442 | 1839.657892622336 | 1460.7357820224042 | 1802.1051676120787 | 1234.8217828622214 | 1799.0555697283216 |
| SE_44_8_19 | SE_44 | SE_44_8 | GCAACTCTGGAGTACAGAAG | 263.3577725302566 | 387.7642840379517 | 368.3475197626849 | 356.55852614119374 | 357.4203507898101 | 545.7753692170975 | 262.583189514574 | 521.8351267870141 | 361.4100080748441 | 218.7020834480678 | 348.61413451528495 | 298.295020271083 |
| SE_44_8_23 | SE_44 | SE_44_8 | GAGTTGTGAAGAATGAAACA | 211.3364841292183 | 248.84872867384016 | 272.55741408713413 | 222.52375171585447 | 244.99733244049295 | 158.5347501059188 | 273.34479564222045 | 161.65544688510764 | 221.1083341784673 | 164.75556953087775 | 212.45507861032516 | 254.18914178742094 |
| SE_44_8_25 | SE_44 | SE_44_8 | AAGTGATGAGATCTTGTGGA | 240.59845885480235 | 281.8286806667587 | 283.42020957611413 | 292.7944101524401 | 310.7541922297162 | 315.33687999210076 | 264.7355107401033 | 296.8409960346421 | 391.6015075208999 | 232.55321539977876 | 262.9278320923361 | 393.4708633147749 |
| SE_44_8_5 | SE_44 | SE_44_8 | GGAGGCCTGTGAATTTGGAT | 1184.5680879653107 | 1222.257008707255 | 1243.2963200568909 | 1234.941756598514 | 1356.5003817809113 | 1337.5828096368232 | 1147.1872132071142 | 1366.0357938536874 | 1151.7169053392452 | 1559.3458549847235 | 1167.9160398744393 | 1664.4165722518794 |
| SE_440_1_15 | SE_440 | SE_440_1 | TGAAAGGACAGCCCAAACAG | 1848.923291920238 | 1933.8244577665891 | 1989.8666282086062 | 1862.172448569519 | 1835.889101534603 | 1750.812732044054 | 1916.6420513338371 | 2274.520498278181 | 2148.036387059086 | 2117.765174722123 | 2646.88509539383 | 1753.7890102319316 |
| SE_440_1_2 | SE_440 | SE_440_1 | GCAGACATTAAAATTGGCAA | 1519.455132046995 | 1592.0322280217965 | 1517.8287878692943 | 1647.456547791063 | 1811.4954277418267 | 1467.5293261170843 | 1563.6613703470327 | 1781.0459762078526 | 1693.3879248125988 | 1821.059348177578 | 1784.1531463408521 | 1695.754959595534 |
| SE_440_1_3 | SE_440 | SE_440_1 | TTAGGATTATAATACAACAT | 69.36171786805113 | 80.95079125534559 | 68.13935352178353 | 94.99551973834723 | 85.90815553108195 | 29.454543735525895 | 45.19874573611519 | 33.08737216946648 | 86.13457194904147 | 41.553395855132884 | 76.29602270536539 | 124.19286836189059 |
| SE_440_1_5 | SE_440 | SE_440_1 | GGTTCCTTAGGAAGGAAGTT | 0.0 | 1.9987849692677924 | 1.975053725269088 | 0.0 | 0.0 | 0.0 | 0.0 | 8.508181415005666 | 0.0 | 0.0 | 0.0 | 0.0 |
| SE_440_1_8 | SE_440 | SE_440_1 | GCGTTGTGAAAATTTTGCAT | 3139.701510371002 | 3902.6276524953646 | 3861.2300329010673 | 3670.9912490668157 | 3930.5632641751813 | 3571.7965829874493 | 3545.9492190595133 | 4004.5173859959996 | 4011.9174852141173 | 4115.244203547809 | 4135.244430630804 | 3382.2244710892446 |
| SE_440_2_16 | SE_440 | SE_440_2 | ACACTGCACCCAGAGAAGGG | 2282.4340285955577 | 2453.508549776215 | 2466.842102861091 | 2322.835653876025 | 2653.6074708489755 | 2166.6415847808903 | 2193.2153288143513 | 3255.7974214755013 | 1965.999405104926 | 2803.0317028594022 | 3777.2400163979355 | 2299.309086214068 |
| SE_440_2_17 | SE_440 | SE_440_2 | TGCACACTGCACCCAGAGAA | 29.26197472558407 | 55.96597913949819 | 48.38881626909266 | 28.62878677046081 | 34.999618920070425 | 67.57218856973589 | 31.208657770174778 | 5.67212094333711 | 49.72717555820951 | 40.0953819654791 | 0.0 | 106.78265317097134 |
| SE_440_2_20 | SE_440 | SE_440_2 | TAAATGGAGAACCGATACCG | 415.0865303666185 | 549.665866548643 | 560.915257976421 | 490.59330056653295 | 564.2362807720444 | 590.8234949302547 | 425.08344204203576 | 530.3433082020198 | 650.0052233680243 | 738.4840351096423 | 812.2591955709669 | 454.9869569893562 |
| SE_440_2_6 | SE_440 | SE_440_2 | GCACAGAGACAGGTCTAACA | 3870.1671016689156 | 4019.5565731975303 | 4165.3883065925065 | 4057.4798704680366 | 4017.5320142189926 | 4701.464966255854 | 3583.614840506276 | 3900.528502034819 | 4760.489074420735 | 3292.9243697830743 | 4047.2105582784593 | 4560.315699008113 |
| SE_440_2_8 | SE_440 | SE_440_2 | GCAACCTGAAGCACAGAGAC | 2284.601582278934 | 2390.5468232442795 | 2294.0249019000457 | 2236.949293564642 | 2289.8235529827894 | 1781.9998959993168 | 2440.7322697502204 | 2282.083326202631 | 2236.8349148416028 | 3133.271848865985 | 1840.4948246463528 | 3027.056081194492 |
| SE_441_1_13 | SE_441 | SE_441_1 | TGTCCTCAAGGCCAAGGCAG | 642.6796671211613 | 651.6038999813003 | 549.0649356248065 | 573.8770438987826 | 492.1158539064447 | 598.6202859190704 | 525.166379029148 | 546.4143175414749 | 833.8181758778344 | 360.85843768931187 | 555.2002883021204 | 590.7866354785263 |
| SE_441_1_14 | SE_441 | SE_441_1 | GCATGTTGTCCTCAAGGCCA | 4858.571581288644 | 4244.419882240157 | 4336.230453828282 | 4627.45298889812 | 4167.0758405138395 | 3930.4489684729706 | 5027.822382836433 | 4191.697377126125 | 3745.521901866566 | 4460.064488450929 | 4191.586108936304 | 4227.200248355191 |
| SE_441_1_21 | SE_441 | SE_441_1 | AGAGGGCATGAGGTTCCACA | 3286.011383998922 | 3152.0838965353087 | 3101.8218755351027 | 3154.3717787089545 | 2935.725611234998 | 3027.7538339900884 | 2857.206426890139 | 3272.8137843055124 | 2898.383946821354 | 2443.631279059744 | 3199.7378137665546 | 2966.7006685326387 |
| SE_441_1_6 | SE_441 | SE_441_1 | GGCATCTGTGTTCTTCCAGG | 99.7074694353235 | 199.87849692677923 | 182.69246958739063 | 176.97795458103047 | 145.30144824392872 | 213.11228702762855 | 140.9770402721688 | 188.1253446206808 | 237.09206917932033 | 181.52272926189627 | 103.29307689341775 | 127.67491140007444 |
| SE_441_1_8 | SE_441 | SE_441_1 | TCTGTGTTCTTCCAGGAGGA | 118.13167574402458 | 140.91434033337936 | 158.9918248841616 | 105.40598765487843 | 148.48323178211695 | 106.55614351381428 | 153.8909676253446 | 172.05433528122566 | 118.10204195074758 | 260.98448624802757 | 117.3784964697929 | 139.28172152735394 |
| SE_442_1_38 | SE_442 | SE_442_1 | ATCCGAGAAGAGCAGAGCGG | 197.2473851872704 | 236.8560188582334 | 237.00644703229057 | 258.9603894237137 | 279.9969513605634 | 257.2941026309174 | 272.2686350294558 | 324.25624726077143 | 202.4606433441387 | 250.04938207562418 | 395.5655331032021 | 219.36871140558245 |
| SE_442_1_39 | SE_442 | SE_442_1 | GGAATCCGAGAAGAGCAGAG | 97.53991575194691 | 162.90097499532507 | 154.05419057098888 | 118.41907255054244 | 90.15053358199957 | 140.34223779868222 | 52.73187002546773 | 76.57363273505099 | 183.81295250981015 | 58.320555586151414 | 17.606774470468935 | 109.10401519642724 |
| SE_442_1_42 | SE_442 | SE_442_1 | GTCTTCTCCACTGTTCCTGG | 18.42420630870108 | 47.97083926242702 | 45.42623568118903 | 20.820935833062407 | 41.363185996446866 | 7.796790988815679 | 13.990087965940416 | 0.0 | 0.8879852778251698 | 44.46942363444045 | 1.173784964697929 | 12.767491140007444 |
| SE_442_1_45 | SE_442 | SE_442_1 | AACAGTGGAGAAGACAAGGC | 871.3565807173924 | 952.4210378561031 | 1018.1401953762149 | 922.6277191025779 | 923.7778205873133 | 935.6149186578814 | 893.2133085946574 | 1292.2982215903048 | 1115.3095089484132 | 1166.4111117230282 | 1396.8041079905356 | 748.6392532095274 |
| SE_442_1_46 | SE_442 | SE_442_1 | GCTCTCAGAATTCCACACCA | 132.22077468597246 | 158.9034050567895 | 195.5303188016397 | 114.51514708184324 | 126.21074701479941 | 151.60426922697152 | 123.75847046793446 | 176.7811027340066 | 245.97192195757202 | 120.28614589643729 | 278.1870366334092 | 262.31390887651656 |
| SE_443_1_11 | SE_443 | SE_443_1 | TGTGTGCTAGTTGGTCACCA | 3644.7415185977493 | 2945.209652216092 | 2829.2644614479686 | 3466.6858162048907 | 3076.7846814280088 | 2494.106806311149 | 3576.0817162169237 | 3583.8350826984974 | 2734.994655701523 | 2346.6733553977674 | 3024.843854026563 | 2758.9387672543357 |

| | | | | | | | | | | | | | | | |
|---|---|---|---|---|---|---|---|---|---|---|---|---|---|---|---|
| SE_443_1_15 | SE_443 | SE_443_1 | AGACTTCACTTCTAGCCAAA | 82.36703996831072 | 134.91798542557598 | 110.60300861506893 | 119.72038104010883 | 133.63490860390525 | 14.727271867762948 | 95.77829453605362 | 131.40413518730972 | 109.22218917249589 | 89.6678542137078 | 71.60088284657367 | 20.89225822910309 |
| SE_443_1_16 | SE_443 | SE_443_1 | GTACCCCAGATGCTGCAACC | 140.89098941947887 | 171.89550735703014 | 144.17892194464343 | 160.06094421666725 | 151.66501532030517 | 252.96255208157535 | 165.7287343657557 | 149.36585150787724 | 171.38115862025776 | 176.41968064810803 | 361.52576912696213 | 184.54828102374395 |
| SE_443_1_18 | SE_443 | SE_443_1 | CCTTGTAATCCACCCCCGCC | 2365.8848454055565 | 2889.243673076594 | 2977.39349084315 | 2449.062577363966 | 2939.9679892859153 | 3157.70035047035 | 2323.430762958874 | 3586.671143170166 | 3289.097469064429 | 2983.8254251764715 | 3232.6037927780967 | 2811.1694128270933 |
| SE_443_1_19 | SE_443 | SE_443_1 | TGAAAGTCATCCTAGGAAGA | 244.93356622155557 | 271.8347558204198 | 296.25805879036324 | 275.8773997880769 | 326.6631099206573 | 235.63634988420716 | 208.7751588763416 | 305.34917744964775 | 420.0170364113053 | 226.72115984116363 | 460.1237061615882 | 198.47645317647937 |
| SE_443_2_14 | SE_443 | SE_443_2 | CTGGAAGTAGAGAAAGGACT | 1057.7661974877797 | 1268.2290630004143 | 1205.7702992767784 | 1242.7496075359124 | 1302.4100616317114 | 1104.545390082221 | 1103.0646280837636 | 1390.6149846081482 | 1379.9291217403138 | 919.2777574267117 | 1255.9499122267841 | 1063.1838076588017 |
| SE_443_2_20 | SE_443 | SE_443_2 | TTTGTATTTGGATTGCTGAG | 16.256652625324485 | 28.982382054382988 | 30.613332741670867 | 14.314393385230405 | 4.242378050917627 | 6.064170769078861 | 13.990087965940416 | 17.01636283001133 | 8.879852778251697 | 0.7290069448268927 | 7.042709788187574 | 37.14179240729438 |
| SE_443_2_26 | SE_443 | SE_443_2 | ACAGAGGCAGAAAGGAGGGT | 6454.974869095508 | 6805.862820356833 | 6656.918581019461 | 5884.516989819263 | 6400.68788432197 | 5937.689493038073 | 5996.366934324616 | 5192.826723625124 | 5739.048850584072 | 5184.697391608861 | 4135.244430630804 | 4151.755982527875 |
| SE_443_2_4 | SE_443 | SE_443_2 | TTAGGATTACCTTAAATGTA | 36.84841261740216 | 78.9520062860778 | 65.1767729338799 | 57.25757354092162 | 39.24199697098805 | 54.57753692170975 | 104.3875794381708 | 36.868786131691216 | 119.87801250639792 | 215.05704872393332 | 63.384388093688166 | 34.820430381838484 |
| SE_443_2_9 | SE_443 | SE_443_2 | GCAGTCTACCATTTACCTTA | 1817.4937635112772 | 2040.759453622416 | 1908.8894254725735 | 1863.4737570590855 | 1902.7065558365557 | 2104.2672568703647 | 1790.7312596403733 | 2020.2204093185674 | 2061.901815110044 | 2096.6239733221432 | 2139.8099906443244 | 1751.4676482064756 |
| SE_443_3_15 | SE_443 | SE_443_3 | AGCTTAACAGTCCAAGTGAG | 539.7208671607729 | 669.5929647047104 | 650.7802024761645 | 567.3705014509505 | 634.2355186121853 | 625.4758993249911 | 537.004145769559 | 657.9660294271048 | 544.334975306829 | 570.08343085463 | 664.3622900190278 | 1153.7169266515818 |
| SE_443_3_19 | SE_443 | SE_443_3 | TGGAGTCTTTCACAGAGCCA | 74.78060207649263 | 93.94289355558624 | 93.81505195028168 | 67.66804145745283 | 88.02934455654076 | 82.29946043749882 | 71.02660044246673 | 67.12009782948914 | 104.78226278337003 | 35.72134029651774 | 129.1163461167722 | 30.177706330926686 |
| SE_443_3_21 | SE_443 | SE_443_3 | ACACCAGATAGAAGGAACAG | 733.7169218229784 | 849.4836119388118 | 878.8989077447442 | 878.3832304573203 | 1029.8372718602538 | 739.8288338276211 | 680.1335072672572 | 1090.9379281018375 | 828.4902642108834 | 821.590826819908 | 1207.824728674169 | 776.4955975149982 |
| SE_443_3_22 | SE_443 | SE_443_3 | ACAGTAAAAGCTTTCCAGCG | 354.39502723207374 | 567.654931272053 | 529.3143983721156 | 499.70245999349777 | 515.4489331864917 | 351.72190460657396 | 420.77879959097714 | 556.813205937593 | 459.0883886356128 | 418.44998633063636 | 319.2695103978367 | 268.1173139401563 |
| SE_443_3_5 | SE_443 | SE_443_3 | AACAAGCTCTTAAGAAGCAC | 124.63433679415438 | 118.92770567143364 | 128.37849214249073 | 124.92561499837444 | 113.48361286204653 | 68.4384986796043 | 108.6922218892294 | 58.61191641448347 | 199.7966875106632 | 114.45409033782215 | 62.210603128990236 | 84.7297139291403 |
| SE_443_4_19 | SE_443 | SE_443_4 | CTTCTCCTCGGGAGAAAGCA | 4.335107366753196 | 0.9993924846338962 | 0.987526862634544 | 6.506542447832002 | 3.1817835381882205 | 0.0 | 5.380803063823238 | 0.0 | 31.96747000170611 | 0.0 | 0.0 | 1.1606810127279494 |
| SE_443_4_23 | SE_443 | SE_443_4 | CTTCTAAGTCAAGCAGACCT | 75.86437891818092 | 95.94167852485404 | 81.96472959866715 | 78.07850937398402 | 89.08993906927017 | 97.02673230526177 | 107.61606127646475 | 146.52979103620868 | 85.2465866712163 | 92.58388199301537 | 85.68630242294881 | 16.249534178191293 |
| SE_443_4_32 | SE_443 | SE_443_4 | CTGATTAGTAAGGAGTTTGG | 2422.241241173348 | 2706.3548483885907 | 2681.135432052787 | 2714.5295092355113 | 2602.6989342379643 | 2311.3153731289144 | 2434.2753060736327 | 2851.1861275174538 | 2738.5465968128237 | 2619.3219527630254 | 2973.197315579854 | 2055.5660735411984 |
| SE_443_4_37 | SE_443 | SE_443_4 | AGCGCAAGATGAAATTAACA | 1346.0508373768673 | 1711.959326177864 | 1618.5565278580177 | 1594.1028997188405 | 1510.2865861266753 | 1820.983850943395 | 1644.3734163043814 | 1877.4720322445835 | 1527.334677859292 | 2103.9140427704124 | 2351.091284289952 | 1211.7509772879791 |
| SE_443_4_4 | SE_443 | SE_443_4 | GCATCAGTGGACTAGAGAGG | 2285.6853591206223 | 2857.263113568309 | 2716.6863991076307 | 2423.0364075726375 | 2701.334223921799 | 2656.106796856541 | 2219.043183520703 | 2742.4704761034927 | 3202.9628971153875 | 2636.818119438871 | 2409.7805325248482 | 2168.1521317758097 |
| SE_444_1_12 | SE_444 | SE_444_1 | TAGACCTGAAGTAATCACTG | 1290.778218450764 | 1352.1780317096616 | 1360.8120167104016 | 1339.0464357638261 | 1607.8612812977806 | 1263.08014018814 | 1431.2936149769812 | 1636.4068921527562 | 1051.374568945001 | 1468.9489938261886 | 1494.2282600604638 | 1183.8946329825085 |
| SE_444_1_7 | SE_444 | SE_444_1 | GTTTTTTGTGGACCTTTAGG | 2.167553683376598 | 1.9987849692677924 | 2.962580587903632 | 7.807850937398403 | 3.1817835381882205 | 0.0 | 1.0761606127646475 | 47.26767452780925 | 7.991867500426528 | 0.7290069448268927 | 22.30191432926065 | 2.321362025455899 |
| SE_444_1_9 | SE_444 | SE_444_1 | ATCTAGCTCTGGAAGCAGAA | 35.76463577571386 | 69.95747392437274 | 54.31397744489992 | 54.65495656178882 | 40.30259148371746 | 10.395721318420904 | 62.417315540349556 | 87.9178746217252 | 31.07948472388094 | 87.48083337922712 | 78.64359263476125 | 70.80154177640492 |
| SE_444_2_1 | SE_444 | SE_444_2 | ATCCCCTGAGAGTAGAGAAA | 20.59175999207768 | 26.983597085115196 | 27.650752153767232 | 40.34056317655841 | 31.8178353818822 | 0.8663101098684087 | 15.066248578705064 | 17.961716320567515 | 27.527543612580263 | 8.019076393095819 | 16.432989505771005 | 2.321362025455899 |
| SE_444_2_10 | SE_444 | SE_444_2 | TGGATGGGCAGAAAAAGCTG | 201.5824925540236 | 282.82807315139263 | 258.73203801025056 | 219.92113473672168 | 307.572408691528 | 194.91977472039196 | 177.56650110616684 | 209.86847490347307 | 208.6765402889149 | 175.69067370328113 | 431.95286700883787 | 237.93960760922963 |
| SE_444_2_12 | SE_444 | SE_444_2 | TAGGATGCATTACCTCAGGG | 5127.348238027342 | 5800.4739808151335 | 6027.863969521257 | 5121.950214933352 | 5754.785826069761 | 4964.823239655851 | 4430.553242752054 | 5406.476612490822 | 5270.1926238923825 | 4501.6178843060625 | 5422.886536904432 | 6193.393883916338 |
| SE_444_2_21 | SE_444 | SE_444_2 | ATACAAGCTACTTAGAAGCT | 88.86970101844051 | 88.94593113241676 | 96.77763253818532 | 102.80337067574564 | 110.3018293238583 | 51.978606592104526 | 87.16900963393644 | 59.557269905039654 | 126.09390945117411 | 145.07238202055163 | 57.515463270198524 | 111.42537722188314 |
| SE_444_2_25 | SE_444 | SE_444_2 | GCTTGTATCAAGGAGATAAA | 668.6903113216804 | 854.4805743619812 | 853.2232093162461 | 667.5712551475634 | 825.1425309034785 | 560.5026410848604 | 737.1700197437835 | 747.7746110299423 | 761.0033830961705 | 745.0450976130843 | 924.942552181968 | 697.5692886494976 |
| SE_444_3_0 | SE_444 | SE_444_3 | TTCGTTACTTAGCTTAGCTA | 30.34575156727237 | 64.96051150120326 | 62.21419234597627 | 49.44972260352321 | 40.30259148371746 | 38.98395494407839 | 50.57954879993843 | 82.2457536783881 | 41.73530805778298 | 45.92743752409424 | 110.33578668160533 | 62.67677468730927 |
| SE_444_3_10 | SE_444 | SE_444_3 | ATCCAGCTCCCATAGGAAGA | 54.188842084414944 | 83.94896870924728 | 62.21419234597627 | 62.46280749918722 | 110.3018293238583 | 79.7005301078936 | 72.10276105523138 | 66.17474433893295 | 107.44621861684554 | 16.038152786191638 | 55.167893340802664 | 49.909283547301825 |
| SE_444_3_21 | SE_444 | SE_444_3 | CTCCTGAACTGTGTACACAG | 54.188842084414944 | 35.97812944682026 | 54.31397744489992 | 42.943180155691216 | 26.51486281823517 | 138.6096175789454 | 24.75169409358689 | 8.508181415005666 | 105.67024806119521 | 31.347298627556384 | 18.780559435166865 | 106.78265317097134 |
| SE_444_3_3 | SE_444 | SE_444_3 | GATGATGTCTGTCTGAGGTC | 1468.517620487645 | 1726.9502134473726 | 1840.75007195079 | 1503.0113054491926 | 1550.5891776103927 | 1435.4758520519533 | 1547.5189611555631 | 1290.4075146091925 | 1844.3454220428775 | 2103.9140427704124 | 1109.2267916395429 | 1628.435460857313 |
| SE_444_3_8 | SE_444 | SE_444_3 | TATTGGCTGTGATTAACAGC | 483.36447139298133 | 526.6798394020633 | 554.9900968006137 | 442.44488645257616 | 485.7522868300683 | 402.83420108881006 | 425.08344204203576 | 408.3927079202719 | 539.0070636398781 | 567.8964100201493 | 316.9219404684408 | 639.5352380131002 |
| SE_445_1_10 | SE_445 | SE_445_1 | GCAGGAGGAGGAGCTGGGCG | 13315.28227698244 | 13259.939486122534 | 13309.887054588384 | 12979.250874935278 | 13508.792308634454 | 12212.373618814958 | 13411.113556273036 | 14168.9581164561 | 12739.036795679885 | 11384.172450416756 | 15630.120589917624 | 14094.14953755549 |
| SE_445_1_20 | SE_445 | SE_445_1 | GCAGTAGCAGAGCAGTGAAA | 4291.756293085664 | 4510.258283152773 | 4476.459268322388 | 4653.479158689448 | 4767.372334718683 | 5318.277764482161 | 4660.8516138836885 | 5512.356203433114 | 5226.681345278949 | 4723.965002478264 | 5541.438818338923 | 5271.813159810346 |
| SE_445_1_21 | SE_445 | SE_445_1 | AAGAGATTTGGACAACACGG | 295.8710777809056 | 425.74119845403976 | 446.3621419108139 | 339.6415157768305 | 475.14634170277424 | 537.9785782282818 | 222.76524684228204 | 352.61685197745703 | 428.00890391173186 | 225.99215289633673 | 433.1266519735358 | 636.0531949749163 |
| SE_445_1_30 | SE_445 | SE_445_1 | AGGCATTTAGGAGGATTGGG | 811.7488544245359 | 892.4574887780693 | 889.7617032337241 | 853.6583691555587 | 977.868140736513 | 745.8930045966999 | 811.4251020245442 | 830.9657181988866 | 781.4270444861494 | 866.7892573991753 | 737.1369578302995 | 755.6033392858951 |
| SE_445_1_8 | SE_445 | SE_445_1 | TGTCCTCCAGACTTTCCTTG | 41.18351998415536 | 49.96962423169481 | 60.23913862070719 | 41.641871666124814 | 75.30221040378788 | 66.70587845986748 | 33.36097899570407 | 76.57363273505099 | 47.95120500255917 | 9.477090282749604 | 79.81737759945918 | 190.3516860873837 |
| SE_445_2_15 | SE_445 | SE_445_2 | TGTTTAGGCTCACACAGCTA | 514.7939998019419 | 546.6676890947413 | 519.4391297457702 | 507.5103109308962 | 518.6307167246799 | 378.57751801249464 | 561.755839863146 | 349.78079150578844 | 476.84809419211615 | 570.8124377994569 | 367.39469395045177 | 630.2497899112766 |
| SE_445_2_19 | SE_445 | SE_445_2 | GGACACTCATCCCATTTGTG | 975.399157519469 | 990.3979522721911 | 1028.0154640025603 | 955.1604313417379 | 911.0506864345604 | 1020.5133094249854 | 938.4120543307725 | 808.2772344255382 | 1174.8045225626997 | 670.6863892407413 | 786.4359263476125 | 1132.8246684224787 |
| SE_445_2_29 | SE_445 | SE_445_2 | GAACACCCCTTGACTGAAGG | 273.1117641054513 | 305.81410029797223 | 275.51999467503776 | 274.5760912985105 | 263.02743915689285 | 70.17111889934111 | 261.50702890180935 | 197.57887952624267 | 256.62774529147407 | 143.61436813089784 | 57.515463270198524 | 41.78451645820618 |
| SE_445_2_30 | SE_445 | SE_445_2 | GATATGTCCTTCCCTGGGCA | 237.34712832973747 | 279.82989569749094 | 296.25805879036324 | 321.4231969229009 | 301.2088416151515 | 389.8395494407839 | 255.05006522522146 | 374.35998226024924 | 340.9863466848652 | 423.5530349444246 | 458.9499211968903 | 283.20616710561967 |
| SE_445_2_39 | SE_445 | SE_445_2 | CTATAGGGCAGACAGAAGCC | 293.703524097529 | 358.78190198356873 | 358.4722511363395 | 300.6022610898385 | 338.32964956068076 | 228.7058690052599 | 257.20238645075074 | 483.07563367421056 | 451.09652113518626 | 480.41557664092227 | 538.7672987963495 | 256.51050381287683 |
| SE_445_3_15 | SE_445 | SE_445_3 | AGGATAACCTCATTGCCTTA | 1108.7037090471297 | 1186.2788792604347 | 1166.2692247713965 | 1094.4004397253427 | 1045.7461895511951 | 1180.7806797506412 | 1164.4057830113486 | 1166.5662073463322 | 1248.5073006221887 | 1035.1898616541876 | 752.3961623713725 | 1158.3596507024936 |
| SE_445_3_6 | SE_445 | SE_445_3 | GTCAACCTTTTCATATTAGA | 43.351073667531956 | 60.962941562667666 | 69.12688038441809 | 52.05233958265602 | 62.575076251035 | 65.83956834999907 | 23.675533480822246 | 20.79777679223607 | 53.27911666951019 | 34.992333351690846 | 12.911634611677218 | 197.3157721637514 |
| SE_445_3_7 | SE_445 | SE_445_3 | TCTAATATGAAAAGGTTGAC | 49.85373471766175 | 102.9374259172913 | 80.97720273603261 | 58.55888203048802 | 82.72637199289373 | 79.7005301078936 | 71.02660044246673 | 77.51898622560716 | 66.59889583688773 | 77.27473615165061 | 76.29602270536539 | 49.909283547301825 |
| SE_445_3_9 | SE_445 | SE_445_3 | AGGTTGACAGGATAGTTTGA | 942.88585226882 | 1051.3608938348589 | 1052.703635568424 | 968.1735162374019 | 948.1714943800896 | 1628.6630065526085 | 905.0510753350685 | 830.0203647083305 | 1081.566068391057 | 1010.4036255300732 | 1037.6259087929693 | 1497.2785064190548 |
| SE_445_4_11 | SE_445 | SE_445_4 | CTTGACTGACCAAAACCCAG | 914.7076543849242 | 1053.3596788041266 | 1131.7057845791874 | 1158.1645557140964 | 1023.4737047838776 | 1003.1871072276173 | 979.3061576158292 | 1059.7412629134833 | 904.856998103848 | 1425.208577136575 | 1012.9764245343127 | 912.2952760041683 |
| SE_445_4_12 | SE_445 | SE_445_4 | CCTTGACTGACCAAAACCCA | 2590.2266516350346 | 2463.502474622554 | 2603.120809904658 | 2627.3418404345625 | 2692.8494678199636 | 3121.315325855877 | 2631.212698209563 | 2743.415829594049 | 2579.5972320821184 | 2763.6653278387503 | 3018.9749292030733 | 4092.56125087875 |
| SE_445_4_13 | SE_445 | SE_445_4 | GTGCCTCAAATGAGGAGCAG | 144.14231994454374 | 161.90158251069118 | 173.80472782367974 | 188.68973098712806 | 187.725228753105 | 112.62031428289313 | 108.6922218892294 | 222.15807028070347 | 180.26101139850945 | 104.97700005507254 | 245.32105762186717 | 140.44240254008187 |
| SE_445_4_21 | SE_445 | SE_445_4 | TTTGAGGCACTGGGGTACCA | 195.07983150389381 | 204.87545934994873 | 228.11870526857967 | 249.85122999674888 | 183.48285070218736 | 226.97324878552308 | 168.95721620404964 | 176.7811027340066 | 286.81924473752986 | 486.2476321995374 | 113.85714157569912 | 193.83372912556754 |
| SE_445_4_23 | SE_445 | SE_445_4 | ATGATAGGTGAGCATTGCCA | 2541.4566937590607 | 2649.389476764459 | 2561.644681674007 | 2576.590809341473 | 2649.365092798058 | 2787.7859335565395 | 2598.9278798266237 | 2427.667763748283 | 2740.3225673684738 | 2557.3563624527396 | 2779.522796404696 | 2951.6118153671755 |
| SE_445_5_13 | SE_445 | SE_445_5 | CTGCCCCTGGGACTGAGAAT | 131.13699784428417 | 240.85358879676897 | 184.66752331265974 | 173.07402911233126 | 181.36166167672854 | 219.1764577967074 | 217.3844437784588 | 150.31120499843342 | 359.6340375191938 | 273.37760431008473 | 91.55522724643846 | 70.80154177640492 |
| SE_445_5_22 | SE_445 | SE_445_5 | AACTTGGGACTCAGACTCCA | 266.60910305532155 | 266.83779339725027 | 295.27053192772865 | 218.6198262471553 | 295.9058690515045 | 128.21389626052448 | 161.42409191469713 | 187.17999113012462 | 330.3305233509632 | 118.09912506195661 | 279.3608215981071 | 197.3157721637514 |
| SE_445_5_3 | SE_445 | SE_445_5 | CTAGCCACGTTTCCAATCCT | 1158.5574437647915 | 1058.356641227296 | 1099.1173981122474 | 1188.0946509741236 | 1281.1981713771233 | 1456.2672946887951 | 1154.7203374964668 | 1251.648021496389 | 927.9446153273025 | 949.1670421646143 | 1419.1060223197962 | 858.9039494186826 |
| SE_445_5_4 | SE_445 | SE_445_5 | TCTTACAGCAATGAGCACAG | 2222.826302302701 | 2272.61851005748 | 2383.8898463997893 | 2178.3904115341543 | 2541.1844524996586 | 2818.973097511802 | 2261.0134474185243 | 2418.2142288427212 | 2169.3480337268898 | 1885.9409662671712 | 2589.3696321236316 | 2078.7796937957573 |
| SE_445_5_9 | SE_445 | SE_445_5 | TTGTCAATCTCAGGTCAGGA | 180.99073256194592 | 201.87728189604704 | 178.74236213685248 | 184.78580551842887 | 307.572408691528 | 111.75400417302473 | 201.24203458698906 | 305.34917744964775 | 269.0595391810264 | 131.95025701366757 | 143.20176569314734 | 156.69193671827318 |
| SE_446_1_10 | SE_446 | SE_446_1 | GGATTCTGTGGTGAATCATA | 24.926867358830876 | 40.97509186998975 | 25.675698428498144 | 29.93009526002721 | 60.45388722557618 | 0.8663101098684087 | 19.370891029763655 | 36.868786131691216 | 3.551941111300679 | 32.076305572383276 | 42.25625872912545 | 25.534982280014887 |
| SE_446_1_2 | SE_446 | SE_446_1 | ATATAAGACCAAATCACAAA | 71.52927155142773 | 85.94775367851507 | 84.92731018657079 | 59.86019052005442 | 67.87804881468203 | 102.22459296447224 | 76.40740350628997 | 78.46433971616335 | 42.62329333560815 | 76.54572920682372 | 21.128129364562724 | 139.28172152735394 |
| SE_446_1_3 | SE_446 | SE_446_1 | AATATAAGACCAAATCACAA | 34.680858934025565 | 42.973876839257535 | 34.56344019220904 | 27.32747828089441 | 62.575076251035 | 1.7326202197368175 | 19.370891029763655 | 42.54090707502832 | 92.35046889381766 | 40.82438891030599 | 28.170839152750297 | 16.249534178191293 |
| SE_446_1_8 | SE_446 | SE_446_1 | TTGGATATATATCCAGACGT | 16.256652625324485 | 14.990887269508443 | 16.78795666478725 | 3.9039254686992013 | 7.4241615891058474 | 136.01068724934018 | 11.837766740411123 | 17.01636283001133 | 0.8879852778251698 | 61.965590310285876 | 10.564064682281362 | 34.820430381838484 |
| SE_446_1_9 | SE_446 | SE_446_1 | ACTGAGTTCTACAAGCAGTT | 268.77665673869814 | 377.7703591916128 | 364.39741231214674 | 337.0388987976977 | 333.02667699703375 | 220.04276790657582 | 407.86487223780136 | 307.2398844307601 | 278.8273772371033 | 608.7207989304553 | 570.4594928431935 | 270.4386759656122 |
| SE_446_2_14 | SE_446 | SE_446_2 | TGTAGAAAAGAATATCCCAG | 185.3258399286991 | 300.81713787480277 | 280.4576289882105 | 196.49758192452646 | 304.3906251533397 | 87.49732109670929 | 207.69899826357695 | 152.2019119795458 | 221.99631945629244 | 487.7056460891912 | 194.84830413985622 | 277.40276204197994 |
| SE_446_2_16 | SE_446 | SE_446_2 | ATATTATTCCCCAGTTGCCC | 26.010644200519174 | 36.977521931454156 | 35.55096705484358 | 31.23140374959361 | 33.939024407341016 | 41.582885273683615 | 31.208657770174778 | 20.79777679223607 | 2.6639558334755096 | 8.019076393095819 | 5.868924823489645 | 8.124767089095647 |
| SE_446_2_17 | SE_446 | SE_446_2 | GCTATGGAAAAATTAGACAA | 201.5824925540236 | 279.82989569749094 | 272.55741408713413 | 270.6721658298113 | 253.4820885423282 | 327.4652215302585 | 229.2222105188699 | 267.53503782740034 | 357.8580669635434 | 441.0492016202701 | 153.7658303754287 | 316.86591647473017 |
| SE_446_2_2 | SE_446 | SE_446_2 | AGTTGGAGACATGAGTGCCA | 469.27537245103343 | 478.70900013963626 | 456.23741053715935 | 421.6239506195137 | 493.17644841917416 | 676.5881958072272 | 462.7490634887984 | 510.4908849003399 | 516.8074316942489 | 417.72097938580947 | 323.96465025662843 | 714.9795038404169 |
| SE_446_2_21 | SE_446 | SE_446_2 | GTTTATCTACAGCTTGTGAG | 11.921545258571289 | 21.986634661945715 | 13.825376076883616 | 15.615701874796805 | 32.87842989461161 | 13.86096175789454 | 6.456963676587884 | 5.67212094333711 | 7.991867500426528 | 10.206097227576498 | 1.173784964697929 | 0.0 |
| SE_446_3_12 | SE_446 | SE_446_3 | TGAAGGCTCTGAGGCAGGAA | 1598.5708414902408 | 1776.9198376790673 | 1838.775018225521 | 1614.923835551903 | 1664.0727904724392 | 1407.7539285361643 | 1377.4855843387488 | 1354.691551967013 | 1561.9661036944738 | 1473.32303549515 | 1423.801162178588 | 1533.2596178136212 |
| SE_446_3_14 | SE_446 | SE_446_3 | ATGTCTGAAGGCTCTGAGGC | 214.5878146542832 | 243.85176625067066 | 241.94408134546327 | 225.12636869498726 | 241.81554890230473 | 225.24062856578627 | 206.6228376508123 | 193.79746556401793 | 204.23661389978906 | 145.80138896537852 | 349.78791947998286 | 116.06810127279495 |
| SE_446_3_15 | SE_446 | SE_446_3 | CAGCATGTCTGAAGGCTCTG | 1456.5960752290737 | 1627.010964983983 | 1560.2924429625796 | 1665.6748666449926 | 1589.8311745813808 | 1445.871573370374 | 1532.452712576858 | 1305.5331704580915 | 1308.002314236475 | 1708.7922786742363 | 1459.0147111195258 | 1529.7775747754374 |
| SE_446_3_16 | SE_446 | SE_446_3 | ACAGGAAGCAGCATGTCTGA | 8948.745381820285 | 8002.135624463606 | 8078.957263213205 | 9185.93662784922 | 8400.969135329631 | 9182.887164605132 | 9863.012015987993 | 8013.76153944478 | 8987.298996868543 | 9767.964053735535 | 7312.680330068098 | 8166.551605553852 |
| SE_446_3_8 | SE_446 | SE_446_3 | GGAAGGGTCTTGGTAATCTG | 1209.4949553241415 | 1489.0948021045053 | 1428.9513702321851 | 1366.3739140447206 | 1328.9249244499467 | 1036.9732015124853 | 1113.82623421141 | 1584.412450172166 | 1426.9923414650477 | 1410.6284382400372 | 1997.7820099158753 | 1245.4107266570898 |
| SE_447_1_14 | SE_447 | SE_447_1 | GGAGGCTAGTTAATAGTCAG | 1075.1066269547925 | 1127.3147226670349 | 1246.2589006447945 | 1085.291280298378 | 1174.0781255914533 | 1289.9357535940605 | 912.5841996244211 | 800.7144065010887 | 1062.030392278903 | 1104.4455214127424 | 766.4815819477477 | 1027.2026962642353 |
| SE_447_1_15 | SE_447 | SE_447_1 | AATGGAGCTCTTGATCCCCG | 844.2621596751849 | 933.432580648059 | 920.375035975395 | 907.0120172277811 | 940.7473327909838 | 1052.5667834901167 | 852.3192053096008 | 889.57763461337 | 798.2987647648276 | 759.6252365096221 | 859.2105941588841 | 1356.8361038789728 |
| SE_447_1_16 | SE_447 | SE_447_1 | ATAAAGCAATTGTGGCCCCG | 242.76601253817896 | 293.8213904823655 | 330.8214989825723 | 255.0564639550145 | 374.38986299348056 | 325.73260131052166 | 255.05006522522146 | 242.01049358238336 | 250.41184834669787 | 388.5607015927338 | 318.0957254331388 | 399.2742683784146 |
| SE_447_1_3 | SE_447 | SE_447_1 | GCAGCTTCTGTATGAAAGAA | 326.21682934817795 | 428.73937590794145 | 386.12300329010674 | 339.6415157768305 | 428.48018314268035 | 386.3743090013103 | 274.42095625498513 | 342.21796358133895 | 369.40187557527065 | 314.93100016521765 | 248.84241251596094 | 466.59376711663566 |
| SE_447_1_5 | SE_447 | SE_447_1 | GTTTCCATATCATTAGGTGA | 13.005322100259587 | 6.9957473924372735 | 10.862795488979984 | 5.205233958265602 | 7.4241615891058474 | 6.064170769078861 | 15.066248578705064 | 8.508181415005666 | 1.7759705556503396 | 2.187020834480678 | 15.259204541073078 | 8.124767089095647 |
| SE_447_2_12 | SE_447 | SE_447_2 | GCTGCAACATCTGTCCCGCT | 5362.527812673703 | 4952.9891538455895 | 5004.786139831869 | 5313.242562899613 | 5037.823935464682 | 5281.02642975782 | 5693.96580213775 | 4748.510583063718 | 5301.272108616264 | 5116.899745739959 | 5677.597874243883 | 5096.550326888426 |
| SE_447_2_14 | SE_447 | SE_447_2 | GGAGCAACGCTTGAACCCAG | 1374.229035260763 | 1497.0899419815764 | 1579.0554533526358 | 1421.0288706065094 | 1536.8014489449104 | 1176.449129201299 | 1592.7177068916783 | 1145.7684305540963 | 1231.6355803435106 | 1504.6703341227064 | 1039.9734787223651 | 1252.3748127334575 |
| SE_447_2_19 | SE_447 | SE_447_2 | AACCCCCTCCCCCACCCAAG | 28.17819788389577 | 26.983597085115196 | 30.613332741670867 | 35.13532921829281 | 50.90853661101153 | 50.2459863723677 | 24.75169409358689 | 27.415251226129364 | 23.975602501279585 | 34.26332640686395 | 23.47569929395858 | 31.338387343654635 |
| SE_447_2_5 | SE_447 | SE_447_2 | CTCTGTCTCCCTAGGCTGGA | 63.94283365960963 | 81.9501837399795 | 94.80257881291622 | 94.99551973834723 | 109.24123481112889 | 6.064170769078861 | 62.417315540349556 | 81.30040018783191 | 34.63142583518162 | 136.32429868262892 | 125.5949912226784 | 1.1606810127279494 |
| SE_447_2_9 | SE_447 | SE_447_2 | ATGTAATTACAGGTTACAGG | 57.440172609479845 | 104.9362108865591 | 87.88989077447442 | 75.47589239485123 | 93.3323171201878 | 13.86096175789454 | 80.71204595734856 | 96.42605603673087 | 72.81479278166393 | 117.37011811712972 | 64.5581730583861 | 51.069964560029774 |

| | | | | | | | | | | | | | | | |
|---|---|---|---|---|---|---|---|---|---|---|---|---|---|---|---|
| SE_447_3_1 | SE_447 | SE_447_3 | TGTGGGTGACAGTAAGGACC | 3795.3864995924228 | 3127.099084419461 | 3364.5040209958916 | 3477.096284121422 | 3257.0857485920083 | 3143.8393887124553 | 3704.1448291359166 | 3312.518630908872 | 3472.022436296414 | 3429.248668465703 | 3642.254745457674 | 2888.9350406798662 |
| SE_447_3_2 | SE_447 | SE_447_3 | GGGTGACAGTAAGGACCTGG | 420.50541457506 | 602.6336682342394 | 489.81332386673387 | 482.7854496291346 | 496.35823195736236 | 714.7058406414372 | 425.08344204203576 | 378.141396222474 | 508.8155641938223 | 766.9153059578911 | 326.31222018602426 | 625.6070658603647 |
| SE_447_3_4 | SE_447 | SE_447_3 | CCTCCACAAAGGGATCCACC | 1118.4577006223244 | 1113.3232278821604 | 1176.144493397742 | 938.2434209773747 | 1053.170351140301 | 798.7379212986729 | 921.1934845265382 | 653.2392619743239 | 941.26439449468 | 1226.1896811988333 | 626.8011711486942 | 1302.2840962807593 |
| SE_447_3_5 | SE_447 | SE_447_3 | GCAAAGGAGACCTCCACAAA | 1684.1892119836166 | 2086.731507915575 | 2069.8563040820045 | 1782.7926307059686 | 2070.280488847802 | 2431.732478400623 | 1606.7077948576186 | 2283.974033183743 | 2405.552117628385 | 2133.074320563488 | 2498.9881898418907 | 2998.0390558762933 |
| SE_447_3_6 | SE_447 | SE_447_3 | AGCAAAGGAGACCTCCACAA | 222.1742525461013 | 371.7740042838094 | 352.5470899605322 | 240.7420705697841 | 317.11775930609264 | 390.70585955065235 | 238.90765603375175 | 267.53503782740034 | 269.0595391810264 | 314.20199322039076 | 232.40942301018995 | 337.7581747038333 |
| SE_447_4_1 | SE_447 | SE_447_4 | ACACAGAGAGGTGTTAGTAT | 870.272803875704 | 912.4453384707472 | 1029.9905177278295 | 994.19968602873 | 1069.079268831242 | 814.3315032763043 | 843.7099204074837 | 943.4627835750726 | 955.4721589398827 | 1324.605618750464 | 885.0338633822385 | 727.7469949804243 |
| SE_447_4_13 | SE_447 | SE_447_4 | AGAGTTCTTAGCACATACAT | 108.37768416882989 | 127.92223803313871 | 93.81505195028168 | 122.32299801924164 | 94.3929116329172 | 117.81817494210358 | 118.37766740411122 | 174.89039575289422 | 82.5826308377408 | 104.97700005507254 | 139.68041079905356 | 84.7297139291403 |
| SE_447_4_16 | SE_447 | SE_447_4 | TCTTTCCCATAAGCAGAACC | 178.8231788785693 | 156.9046200875217 | 207.38064115325426 | 180.88188004972966 | 162.27096044759924 | 192.32084439078673 | 157.11944946363852 | 174.89039575289422 | 216.66840778934142 | 121.01515284126418 | 233.58320797488787 | 175.26283292192036 |
| SE_447_4_17 | SE_447 | SE_447_4 | TATGGTTTTCTCCCACCACC | 484.4482482346696 | 505.69259722475147 | 571.778053465401 | 510.112927910029 | 429.5407776554097 | 544.0427489973607 | 373.4277326293327 | 461.3325033914183 | 642.901341145423 | 166.94259036535843 | 498.85860999661986 | 414.3631215438779 |
| SE_447_4_21 | SE_447 | SE_447_4 | ATTTCACCAGAAGTTGTGGG | 9.75399157519469 | 13.991494784874547 | 11.850322351614528 | 10.410467916531204 | 3.1817835381882205 | 76.23528966841997 | 21.52321225529295 | 0.945353490556185 | 15.983735000853056 | 1.4580138896537853 | 1.173784964697929 | 31.338387343654635 |
| SE_448_1_11 | SE_448 | SE_448_1 | TTGGCTAATGAGAAATAGGA | 49.85373471766175 | 68.95808143973883 | 56.28903117016901 | 53.35364807222242 | 69.99923784014085 | 77.10159977828837 | 43.0464245105859 | 16.071009339455145 | 126.09390945117411 | 92.58388199301537 | 9.390279717583432 | 34.820430381838484 |
| SE_448_1_15 | SE_448 | SE_448_1 | CTAAGGAACAGAATGTAGCT | 859.4350354588211 | 909.4471610168455 | 949.0133149917968 | 800.3047210833363 | 764.6886436779023 | 604.6844566881493 | 821.110547539426 | 791.2608715955269 | 645.5652969788985 | 1041.7509241576297 | 845.1251745825089 | 601.2327645930778 |
| SE_448_1_16 | SE_448 | SE_448_1 | TGGCAGTGAAATCAGCTGCG | 316.4628377729833 | 314.8086326596773 | 337.73418702101407 | 353.95590916206095 | 343.6326221243278 | 562.2352613045973 | 307.7819352506892 | 291.168875091305 | 245.08393667974687 | 295.2478126548915 | 224.19292825730446 | 589.6259544657983 |
| SE_448_1_2 | SE_448 | SE_448_1 | TAGACTGGAGGGACAAAGGG | 1237.6731532080373 | 1513.0802217357189 | 1386.4877151388998 | 1322.129425399463 | 1352.2580037299936 | 1523.839483258531 | 1255.8794350963435 | 1090.9379281018375 | 1458.959811466754 | 1896.8760704395747 | 1208.9985136388668 | 1371.9249570444363 |
| SE_448_1_5 | SE_448 | SE_448_1 | GTAAAATGCCCGTTGACATG | 904.9536628097296 | 980.4040274258522 | 902.5995524479732 | 780.7850937398402 | 950.2926834055485 | 1024.8448599743276 | 688.7427921693744 | 813.0040018783191 | 869.3375869908413 | 724.6329031579313 | 811.085410606269 | 943.6336633478229 |
| SE_45_1_1 | SE_45 | SE_45_1 | AGCCGGTAAGTGGCAGAGCT | 536.469536635708 | 644.608152588863 | 683.3685889431044 | 637.6411598875362 | 581.2057929757149 | 921.7539568999869 | 500.41468493556107 | 628.6600712198631 | 772.5471917078977 | 653.9192295097228 | 582.1973424901728 | 385.3460962256792 |
| SE_45_1_3 | SE_45 | SE_45_1 | GAGGTCACACAGCCGGTAAG | 5681.158204130063 | 5709.529264713449 | 5723.705695829817 | 5963.896807682813 | 5792.967228528019 | 5420.502357446633 | 5803.734184639744 | 5929.257092768393 | 6167.057754495804 | 5856.112787794429 | 6729.309202613227 | 6127.235066190845 |
| SE_45_1_4 | SE_45 | SE_45_1 | TACCGGCTGTGTGACCTCAG | 6501.577273288105 | 6079.304484027991 | 6223.3942883228965 | 6483.118895019807 | 6362.506481863711 | 6541.507639616354 | 6510.7717072261175 | 6400.988484555929 | 5726.61705669452 | 7416.916656668805 | 6920.636151858989 | 6720.343063694827 |
| SE_45_1_5 | SE_45 | SE_45_1 | CTCCACTGAGGTCACACAGC | 304.541292514412 | 209.8724217731182 | 259.7195648728851 | 308.4101120272369 | 318.178353818822 | 246.8983813124965 | 275.49711686774975 | 187.17999113012462 | 328.5545527953128 | 139.96933340676338 | 166.67746498710594 | 307.5804683729066 |
| SE_45_1_6 | SE_45 | SE_45_1 | GAGGTTAAGAGACTCCACTG | 421.5891914167483 | 545.6682966101073 | 612.2666548334173 | 415.11740817168175 | 473.0251526773154 | 152.47057933683993 | 266.8878319656326 | 194.7428190545741 | 584.2943128089618 | 123.93118062057175 | 134.98527094026184 | 87.05107595459621 |
| SE_45_2_0 | SE_45 | SE_45_2 | GTAAAGGATTCAGACTTGCA | 291.5359704141524 | 335.7958748369891 | 354.5221436858013 | 420.3226421299473 | 469.84336913912716 | 358.6523854855212 | 258.2785470635154 | 377.1960427319178 | 292.1471564044809 | 207.0379723308375 | 478.90426559675507 | 525.7884987657611 |
| SE_45_2_18 | SE_45 | SE_45_2 | GTAATTGGCCCCAGACCACA | 2191.3967738937404 | 2612.4119548330045 | 2460.9169416852837 | 2296.8094840846966 | 2438.3067847649063 | 2134.5881107157593 | 2152.321225529295 | 2402.143219503266 | 2502.3425129113284 | 2224.2001886668495 | 2203.194378738013 | 2483.857367237812 |
| SE_45_2_19 | SE_45 | SE_45_2 | GCACCCATCCATGCTAACTA | 1731.8753930179016 | 1762.9283428941928 | 1739.034805099432 | 1784.093939195535 | 1713.9207325707214 | 1565.4223685322145 | 1852.0724145679583 | 2003.204046488556 | 1935.8079056588701 | 1519.2504730192443 | 2182.06624937345 | 1558.794600093636 |
| SE_45_2_2 | SE_45 | SE_45_2 | TGGCACCTGTAGGCAAACAG | 15.172875783636185 | 31.980559508284678 | 33.5759133295745 | 48.14841411395682 | 24.393673792776355 | 37.251334724341575 | 40.8941032850566 | 14.180302358342775 | 21.311646667804077 | 29.8892847379026 | 23.47569929395858 | 38.30247342002233 |
| SE_45_2_5 | SE_45 | SE_45_2 | GTCACTAGACAAGAAAACAG | 256.8551114801268 | 312.8098476904095 | 287.3703170266523 | 326.62843088116654 | 376.5110520189394 | 246.8983813124965 | 207.69899826357695 | 246.7372610351643 | 405.8092719661026 | 221.61811122737538 | 396.7393180679 | 217.04734938012655 |
| SE_45_3_12 | SE_45 | SE_45_3 | TAGTTCCAGACCAGCCTGAG | 354.39502723207374 | 412.74909615379914 | 419.6989166196812 | 366.9689940577249 | 397.7229422735275 | 274.62030482828555 | 453.06361797391656 | 350.72614499634466 | 498.15974085992025 | 853.6671323922913 | 289.9248862803885 | 532.7525848421288 |
| SE_45_3_14 | SE_45 | SE_45_3 | TTTTGCTGTGCCACTCAGGC | 61.77527997623304 | 71.95625889364052 | 78.01462214812898 | 91.09159426964803 | 57.27210368738796 | 111.75400417302473 | 75.33124289352533 | 66.17474433893295 | 20.423661389978903 | 18.954180565499207 | 112.68335661100119 | 92.85448101823596 |
| SE_45_3_15 | SE_45 | SE_45_3 | GGTGTTTTGCTGTGCCACTC | 100.7912462770118 | 142.91312530264716 | 156.02924429625796 | 128.82954046707366 | 166.51333849851687 | 143.80747823815585 | 143.1293614976981 | 70.90151179171387 | 232.6521427901945 | 122.47316673091797 | 76.29602270536539 | 169.4594278582806 |
| SE_45_3_4 | SE_45 | SE_45_3 | TGAAATCCCAACGCTTTAGG | 225.42558307116616 | 216.86816916555549 | 216.26838291696515 | 262.8643148924129 | 190.90701229129323 | 125.61496593091927 | 214.15596194016484 | 294.95028905352973 | 209.56452556674006 | 252.23640291010486 | 440.1693617617234 | 89.37243798005211 |
| SE_45_3_9 | SE_45 | SE_45_3 | GCTTTAGGAGGCTGAGGCTG | 430.2594061502547 | 676.5887120971477 | 625.1045040476664 | 489.29199207696655 | 656.5080033795027 | 304.0748485638115 | 303.47729279963056 | 510.4908849003399 | 503.4876525268713 | 623.3009378269933 | 728.920463077414 | 434.0946987602531 |
| SE_45_4_10 | SE_45 | SE_45_4 | CAGCAAGATGGAATTCACCA | 1311.3699784428418 | 1504.0856893740138 | 1486.2279282649888 | 1340.3477442533924 | 1599.3765251959453 | 1447.604193590111 | 1126.7401615645858 | 1376.4346822498053 | 1823.9217606528987 | 1030.8158199852262 | 1665.6008649063613 | 1418.352197553554 |
| SE_45_4_23 | SE_45 | SE_45_4 | TCTAGACTGGAGGCTATGGT | 622.0879071290835 | 760.537680806395 | 697.1939650199881 | 658.4620957205987 | 667.1139485067969 | 689.5828474552534 | 492.8815606462085 | 736.4303691432681 | 673.9808258693039 | 576.644493358072 | 993.022080134448 | 545.5200759821363 |
| SE_45_4_28 | SE_45 | SE_45_4 | GTAGAGGTCAGCTGATAGGG | 3111.523312487106 | 3829.67200111709 | 3708.1633691927127 | 3522.642081256246 | 3557.23399569443 | 3662.759144523632 | 3074.5908706685977 | 3244.453179588827 | 3583.9085813023853 | 3289.2793350589395 | 3544.8305933877455 | 4523.173906600819 |
| SE_45_4_7 | SE_45 | SE_45_4 | GCTGCTCAGTAAATATGCGT | 1778.4777972104985 | 1706.9623637546947 | 1809.1492123464848 | 1616.2251440414693 | 1833.7679125091443 | 1581.0159505098459 | 1705.7145712319661 | 1596.7020455493964 | 1718.2515125917034 | 1900.521105163709 | 1597.5213369538815 | 1634.2388659209528 |
| SE_45_4_8 | SE_45 | SE_45_4 | ATATATACCAAGCAGCGTGT | 104.0425768020767 | 131.9198079716743 | 149.11655625781614 | 111.91253010271043 | 128.33193604025823 | 193.18715450065514 | 160.34793130193248 | 50.10373499947781 | 162.50130584200608 | 199.74790288256858 | 71.60088284657367 | 241.4216506474135 |
| SE_45_5_1 | SE_45 | SE_45_5 | CATCTGCTAAACAGAAAAGA | 101.8750231187001 | 161.90158251069118 | 186.64257703792882 | 122.32299801924164 | 150.60442080757576 | 302.34222834407467 | 118.37766740411122 | 126.67736773452879 | 148.29354139680336 | 254.42342374458553 | 73.94845277596953 | 239.1002886219576 |
| SE_45_5_18 | SE_45 | SE_45_5 | GTTGACAGACAATTCTTCAT | 372.8192335407748 | 438.73330075428044 | 422.66149720758483 | 511.4142363995954 | 368.02629591710416 | 395.03741009999436 | 519.7855759653247 | 404.61129395804716 | 524.7992991946753 | 1365.43000766077 | 622.1060312899024 | 596.590040542166 |
| SE_45_5_2 | SE_45 | SE_45_5 | TAGTGTCTCCTTCCAGAGCA | 690.3658481554464 | 744.5474010522527 | 617.20428914659 | 728.7327541571842 | 788.0217229579492 | 771.0159977828838 | 703.8090407480795 | 929.2824812167298 | 830.2662347665338 | 895.9495351922511 | 619.7584613605065 | 463.1117240784518 |
| SE_45_5_20 | SE_45 | SE_45_5 | ACTTGTACCCCAAGACCCAC | 1460.9311825958268 | 1310.203547355038 | 1319.3358884797508 | 1553.7623365422821 | 1389.378811675523 | 1752.5453522637908 | 1419.45584823657 | 1426.5384172492832 | 1604.5893970300817 | 1474.052042439977 | 1613.9543264596525 | 1028.363377276963 |
| SE_45_5_8 | SE_45 | SE_45_5 | CTACATCATTGACCACCCCC | 1625.6652625324484 | 1593.0316205064305 | 1560.2924429625796 | 1639.6486968536647 | 1479.5293452575224 | 1295.9999243631394 | 1946.7745484912473 | 1817.9147623395438 | 1670.3003075891443 | 1819.6013342879241 | 2264.2311969023053 | 994.7036279078527 |
| SE_46_1_25 | SE_46 | SE_46_1 | GCTGGGCAAGGCTCATGCCG | 4473.830802489298 | 3905.625829949266 | 3747.6644436980946 | 4162.885858122915 | 4065.258767291816 | 3686.1495174900792 | 4491.894397679638 | 4216.276567880585 | 3781.041312979573 | 4473.91562040264 | 3969.740750608396 | 4482.55007115534 |
| SE_46_1_3 | SE_46 | SE_46_1 | CCAGCGTCACCAACATAGCG | 1556.3035446643971 | 1078.344490919974 | 1185.032235161453 | 1288.2954046707364 | 1194.229421333312 | 1031.7753408532749 | 1428.065133138687 | 1398.1778125325977 | 1055.8144953341268 | 601.4307294821864 | 1370.980838767181 | 904.1705089150726 |
| SE_46_1_30 | SE_46 | SE_46_1 | AGCACCAGCAGCATGCCCGG | 2727.866310529448 | 2950.2066146392617 | 3200.574561798557 | 2963.0794307426936 | 3018.4519832278916 | 4037.005111986785 | 2947.6039183623693 | 3417.4528683606086 | 3002.278224326899 | 4245.736446671823 | 3698.5964237631742 | 3361.3322128601417 |
| SE_46_1_32 | SE_46 | SE_46_1 | GATGCTTTGATGGTCCAAAC | 552.7261892610325 | 458.72115044695835 | 544.1273013116338 | 549.152182597021 | 564.2362807720444 | 683.5186766861744 | 669.3719011396107 | 509.54553140978373 | 599.3900625319897 | 856.5831601715988 | 415.5198775030669 | 831.0476051132118 |
| SE_46_1_41 | SE_46 | SE_46_1 | GCGGGCTGTGGCGAGCCAAG | 3286.011383998922 | 3308.98851662283 | 3370.429182171699 | 3534.3538576623437 | 3259.206937617467 | 3132.577357284166 | 3651.412959110449 | 3272.8137843055124 | 3380.5599526804212 | 3679.298050541327 | 3247.8629973191696 | 3384.5458331147006 |
| SE_46_2_10 | SE_46 | SE_46_2 | GGAACTCCCCAGCAAGGCAG | 0.0 | 0.0 | 0.987526862634544 | 0.0 | 0.0 | 0.0 | 0.0 | 0.0 | 0.0 | 0.0 | 0.0 | 0.0 |
| SE_46_2_18 | SE_46 | SE_46_2 | GTCTGAGTAGAGGCAGGAAG | 461.68893455921534 | 491.70110243987693 | 461.1750448503321 | 590.7940542631458 | 550.4485521065621 | 460.87697844999343 | 693.047434620433 | 448.0975545236317 | 293.035141682306 | 1056.3310630541675 | 832.2135399708317 | 610.5182126949014 |
| SE_46_2_27 | SE_46 | SE_46_2 | CTTTATCCAGGGGGCCACAG | 34.680858934025565 | 61.96233404730156 | 55.301504307534465 | 58.55888203048802 | 60.45388722557618 | 45.91443582302566 | 21.52321225529295 | 15.12565584889896 | 26.639558334755094 | 22.599215289633673 | 4.695139858791716 | 80.08698987822851 |
| SE_46_2_38 | SE_46 | SE_46_2 | AGGACCATCCCCAAACAGCG | 4549.695181407479 | 4926.005556760474 | 5060.087644139404 | 4908.535622644463 | 4967.824697624541 | 4970.021100315061 | 4514.493770547696 | 5109.63561645618 | 5941.509493928211 | 4536.610217657753 | 4513.203189263537 | 4444.247597735319 |
| SE_46_2_53 | SE_46 | SE_46_2 | GGTTCTGCCACCACCACAGA | 32.51330525064897 | 37.97691441608806 | 28.63827901640178 | 36.43663770785921 | 44.54496953463509 | 16.459892087499767 | 17.21856980423436 | 68.06545132004533 | 159.83735000853056 | 18.954180565499207 | 196.02208910455414 | 37.14179240729438 |
| SE_46_3_16 | SE_46 | SE_46_3 | TCTCATCACAGGAGGCCGTG | 2580.4726600598397 | 2419.529205298663 | 2548.806832459758 | 2397.0102377813096 | 2608.0019068016113 | 2360.6950493914137 | 2443.9607515885145 | 2700.8749225190204 | 2163.132136782114 | 2219.097140053061 | 3107.0088015554184 | 2439.7514887541497 |
| SE_46_3_33 | SE_46 | SE_46_3 | TTAACTCAAGCTGTCAAGGG | 61.77527997623304 | 83.94896870924728 | 83.93978332393624 | 53.35364807222242 | 112.42301834931712 | 32.919784174999535 | 73.17892166799602 | 126.67736773452879 | 70.15083694818841 | 115.18309728264904 | 111.50957164630326 | 64.99813671276517 |
| SE_46_3_34 | SE_46 | SE_46_3 | TCTCCTCCAGGGCCTCCCTC | 32.51330525064897 | 59.96354907803377 | 73.07698783495626 | 53.35364807222242 | 36.06021343279983 | 21.65775274671022 | 40.8941032850566 | 24.57919075446081 | 47.95120500255917 | 54.67552086201695 | 21.128129364562724 | 3.482043038183848 |
| SE_46_3_38 | SE_46 | SE_46_3 | CCAGTGATCCACACATGCCA | 1065.3526353795978 | 820.5012298844288 | 847.2980481404388 | 920.0251021234451 | 839.9908540816901 | 823.8609144848567 | 1238.6608652921093 | 721.3047132943692 | 700.620384204059 | 991.449444964574 | 757.0913022301643 | 775.3349165022702 |
| SE_46_3_9 | SE_46 | SE_46_3 | ACTCCCCCCTCGGCAACGGT | 327.30060618986624 | 336.795267321623 | 422.66149720758483 | 335.7375903081313 | 293.7846800260457 | 537.9785782282818 | 313.1627383145124 | 326.1469542418838 | 451.09652113518626 | 602.1597364270133 | 461.2974911262861 | 302.9377443219948 |
| SE_46_4_1 | SE_46 | SE_46_4 | ACACATAGCTCTCCCTGAGA | 972.1478269944041 | 1113.3232278821604 | 1060.6038504695002 | 988.9944520704644 | 1081.806402983995 | 1334.9838793072179 | 949.173660458419 | 1176.9650957424503 | 1005.1993344980922 | 1407.7124104607296 | 995.3696500638438 | 957.5618355005582 |
| SE_46_4_11 | SE_46 | SE_46_4 | CACTTCAGAGACTTCCGCTG | 324.04927566480137 | 371.7740042838094 | 326.87139153203407 | 389.09123838035373 | 380.753430069857 | 449.61494702170415 | 378.80853569315593 | 462.27785688197446 | 358.7460522413686 | 615.2818614338973 | 462.471276090984 | 399.2742683784146 |
| SE_46_4_13 | SE_46 | SE_46_4 | GATGAGTCCTGAATTTATGT | 565.7315113612921 | 719.5625889364053 | 769.2834259923098 | 692.296116449325 | 708.4771345032437 | 1053.433093599985 | 625.2493160162602 | 833.8017786705552 | 665.9889583688773 | 509.57585443399796 | 1105.705436745449 | 1310.408863369855 |
| SE_46_4_15 | SE_46 | SE_46_4 | TTGAATGGAATATTCCAGAA | 15.172875783636185 | 23.98541963121351 | 19.75053725269088 | 24.724861301761607 | 27.575457330964575 | 2.598930329605226 | 13.990087965940416 | 38.75949311280358 | 18.647690834328564 | 8.019076393095819 | 63.384388093688166 | 1.1606810127279494 |
| SE_46_4_30 | SE_46 | SE_46_4 | GGTAGGAAATGAGCTCAGGG | 227.59313675454277 | 391.7618539764873 | 334.77160643311043 | 278.4800167672097 | 334.0872715097631 | 252.09624197170695 | 238.90765603375175 | 370.57856829802455 | 351.6421700187672 | 303.9958959928142 | 380.306328562129 | 126.51423038734649 |
| SE_46_5_0 | SE_46 | SE_46_5 | AATAAACGCTGAGTGACAGT | 180.99073256194592 | 128.9216305177726 | 139.2412876314707 | 119.72038104010883 | 144.2408537311993 | 188.8556039513131 | 170.0333768168143 | 199.46958650735505 | 82.5826308377408 | 69.98466670338169 | 255.88512230414852 | 32.49906835638259 |
| SE_46_5_20 | SE_46 | SE_46_5 | GATAGGAGCTGACGTCCATC | 1989.814281339717 | 1944.817775097562 | 1972.0911446811845 | 2144.556390805428 | 2029.9778973640846 | 1868.6309069861577 | 2038.2482005762424 | 1311.2052914014287 | 1831.9136281533254 | 2528.196084659664 | 1561.1340030482456 | 1872.1784735301824 |
| SE_46_5_22 | SE_46 | SE_46_5 | CGTCAGCTCCTATCTCTCCT | 9024.609760738465 | 9127.451562161374 | 9250.164122297774 | 8734.38258196968 | 9794.590325056071 | 10640.020769403796 | 8557.629192704477 | 8932.645132265392 | 9489.010678839764 | 10647.146429196768 | 9180.172208902502 | 10836.117934828137 |
| SE_46_5_26 | SE_46 | SE_46_5 | CTATCTCTCCTGGGGAGTGG | 10.837768416882989 | 30.98116702365078 | 33.5759133295745 | 28.62878677046081 | 36.06021343279983 | 6.93048087894727 | 15.066248578705064 | 18.9070698111237 | 19.535676112153734 | 14.580138896537854 | 5.868924823489645 | 3.482043038183848 |
| SE_46_5_34 | SE_46 | SE_46_5 | GGAGTGGGGCCAAGATCGGG | 850.7648207253146 | 1253.2381757309058 | 1243.2963200568909 | 956.4617398313043 | 1266.3498481989116 | 768.4170674532785 | 704.8852013608441 | 1121.1892397996355 | 1457.1838409111035 | 1017.6936949783421 | 1146.7879105098766 | 1156.0382886770376 |
| SE_46_6_12 | SE_46 | SE_46_6 | TGCCACTGTTCAAACATTGA | 548.3910818942793 | 618.6239479883817 | 549.0649356248065 | 556.9600335344194 | 740.294969885126 | 498.128313174335 | 501.4908455483257 | 503.8734104664466 | 373.8418019643965 | 496.4537294271139 | 504.7275348201095 | 637.2138759876442 |
| SE_46_6_13 | SE_46 | SE_46_6 | TCCAGGAGCTTCCAGTCCCT | 376.07056406583973 | 391.7618539764873 | 455.2498836745248 | 394.2964723386193 | 471.964558164586 | 563.1015714144656 | 330.3813081187468 | 494.41987556088475 | 494.6077997486196 | 432.3011182823473 | 556.3740732668184 | 481.68262028209904 |
| SE_46_6_28 | SE_46 | SE_46_6 | GCAGGGCGCATTTCCAGGCA | 26949.19494542124 | 25200.680892528326 | 25326.113919125517 | 26689.837121006873 | 25909.263351466678 | 25006.04132135162 | 26642.508290214377 | 26466.116321610956 | 25645.90280886873 | 24633.144665700704 | 25498.130788133112 | 23504.951188753705 |
| SE_46_6_36 | SE_46 | SE_46_6 | CTAATTAAAACACCACTCTG | 251.43622727168534 | 268.8365783665181 | 263.66967232342324 | 273.2747828089441 | 203.6341464440461 | 160.2673703256556 | 331.4574687315114 | 305.34917744964775 | 221.99631945629244 | 316.38901405487144 | 167.85124995180385 | 255.34982280014887 |
| SE_46_6_37 | SE_46 | SE_46_6 | ACAGTGCTGCACATCTGTGA | 4801.131408679164 | 4201.4460054009 | 4348.080776179897 | 4770.596922750424 | 4378.134148546991 | 4264.844670882177 | 4962.17658545779 | 4219.1126283522535 | 3487.118186019442 | 4457.8774676164485 | 4712.746633262185 | 3773.3739723785634 |
| SE_46_7_3 | SE_46 | SE_46_7 | GCCCTTGGAAGGCTTCCCAG | 54.188842084414944 | 89.94532361705066 | 91.8399982250126 | 37.737946197425615 | 86.96875004381135 | 4.331550549342044 | 26.904015319116187 | 94.5353490556185 | 93.23845417164283 | 21.87020834480678 | 48.12518355261509 | 20.89225822910309 |
| SE_46_7_31 | SE_46 | SE_46_7 | AAGTCTGGAGACCACGACTG | 242.76601253817896 | 216.86816916555549 | 200.46795311481245 | 288.8904846837409 | 163.33155496032865 | 246.8983813124965 | 232.45069235716386 | 238.22907962015861 | 209.56452556674006 | 103.51898616541875 | 206.5861537868355 | 264.6352709019725 |
| SE_46_7_41 | SE_46 | SE_46_7 | AGGCCCTAGTAAGTGCCCGG | 162.56652625324483 | 203.8760668653148 | 230.09375899384875 | 153.55440176883525 | 238.63376536411653 | 116.95186483223517 | 176.4903404934022 | 282.66069367629933 | 118.10204195074758 | 368.1485071375808 | 485.9469753849426 | 85.89039494186825 |
| SE_46_7_44 | SE_46 | SE_46_7 | TTGCCGGGAAAGAGGCAACA | 164.73407993662144 | 202.87667438068092 | 198.49289938954334 | 290.1917931733073 | 203.6341464440461 | 218.310147686839 | 124.83463108069911 | 185.28928414901227 | 142.07764445202716 | 95.49990977232294 | 89.2076573170426 | 114.907420260067 |
| SE_46_7_5 | SE_46 | SE_46_7 | CTTCCTGTGAGCGTGCCAAG | 146.30987362792035 | 213.8699917116538 | 212.31827546642697 | 137.93869989403845 | 99.69588419656424 | 120.41710527170882 | 91.47365208499504 | 147.47514452676486 | 126.98189472899928 | 183.70975009637695 | 69.25331291717781 | 139.28172152735394 |
| SE_47_1_1 | SE_47 | SE_47_1 | CAGATAATAGACTGGTCATG | 1143.3845679811554 | 1390.1549461257496 | 1432.9014776827235 | 1004.6101539452611 | 1196.3506103587708 | 537.9785782282818 | 783.4449260926633 | 431.0811916936204 | 876.4414692134426 | 385.6446738134262 | 223.01914329260651 | 305.2591063474507 |
| SE_47_1_13 | SE_47 | SE_47_1 | GCAGGAGCCATAGAGACCCA | 3534.196280745543 | 4130.489138991893 | 4048.8601368016307 | 2912.3283996496043 | 3442.6897883196543 | 1723.9571186381334 | 2561.262258379861 | 1334.8391286653332 | 2984.5185187703955 | 680.8924864683178 | 788.7834962770083 | 821.7621570113882 |
| SE_47_1_2 | SE_47 | SE_47_1 | TCCGTATAGCAAGTGTACCC | 2434.1627864319194 | 2526.4642011544897 | 2612.9960785310036 | 1647.456547791063 | 2123.3102144842724 | 1612.2031144651087 | 1335.5153204409276 | 804.4958204633134 | 1845.2334073207028 | 417.72097938580947 | 515.2915995023908 | 320.34795951291403 |
| SE_47_1_22 | SE_47 | SE_47_1 | TGGGTTTATGAGCCACCCCA | 17.340429467012783 | 22.986027146579612 | 12.837849214249072 | 6.506542447832002 | 2.1211890254588135 | 0.0 | 13.990087965940416 | 0.945353490556185 | 20.423661389978903 | 0.0 | 1.173784964697929 | 0.0 |
| SE_47_1_9 | SE_47 | SE_47_1 | ATGAAGGGAGAATTCTACAT | 483.36447139298133 | 591.6403509032665 | 636.9548263992809 | 644.1477023353682 | 625.75076251035 | 610.7486274572282 | 563.9081610886752 | 704.2883504643578 | 641.1253705897726 | 833.9839448819652 | 848.6465294766027 | 562.9302911730555 |
| SE_47_2_0 | SE_47 | SE_47_2 | AGACGACACAAAGACTCCAG | 1457.679852070762 | 2004.7813241755957 | 1914.8145866483808 | 1620.1290695101686 | 1766.9504582071916 | 1313.3261265605076 | 1290.3165747048124 | 1742.2864830950489 | 1999.7428456622824 | 1267.0140701091395 | 1686.728994270924 | 2136.813744432155 |

| | | | | | | | | | | | | | | |
|---|---|---|---|---|---|---|---|---|---|---|---|---|---|---|
| SE_47_2_14 | SE_47 | SE_47_2 | GCATCTTCCAGCTAAAGGAA | 274.19554094713965 | 387.7642840379517 | 430.5617121086612 | 318.8205799437681 | 295.9058690515045 | 335.2620125190742 | 231.3745317443992 | 415.9555358447214 | 244.1959514019217 | 220.1600973377216 | 169.0250349165018 | 509.5389645875698 |
| SE_47_2_19 | SE_47 | SE_47_2 | TGTGAAGTGGTGAAGAACCC | 2616.2372958355536 | 2862.2600759914785 | 2910.2416641840014 | 2478.9972572623993 | 2803.151297143822 | 2494.106806311149 | 2292.222105188699 | 3074.2895512887135 | 2826.4571393175156 | 2370.7305845770547 | 3186.8261791548775 | 2794.9198786489023 |
| SE_47_2_21 | SE_47 | SE_47_2 | ACAAGGTGTTAGTGGTCAGG | 2.167553683376598 | 4.996962423169481 | 2.962580587903632 | 6.506542447832002 | 4.242378050917627 | 0.0 | 1.0761606127646475 | 0.0 | 11.543808611727208 | 10.206097227576498 | 1.173784964697929 | 0.0 |
| SE_47_2_8 | SE_47 | SE_47_2 | AGAAGGAAGAAGTGAGACCA | 1610.4923867488121 | 1577.041340752288 | 1600.7810443305 96 | 1505.6139224283254 | 1596.194741657757 | 1074.224536236827 | 1658.3635042703218 | 1759.3028459256023 | 1626.789028975711 | 1312.2125006884069 | 1576.3932075893188 | 1221.0364253898028 |
| SE_47_3_11 | SE_47 | SE_47_3 | CAGTTTAGAACACTTCCCAG | 345.72481249856736 | 405.75334876136185 | 428.5866583833921 | 221.22244322628808 | 269.3910062332693 | 428.8235043848623 | 195.86123152316583 | 69.95615830115769 | 260.17968640277473 | 64.88161808959345 | 41.08247376442752 | 12.767491140007444 |
| SE_47_3_2 | SE_47 | SE_47_3 | GTTCTCATTCAGGAGCTCCA | 2.167553683376598 | 2.998177453901 6888 | 5.925161175807264 | 3.9039254686992013 | 0.0 | 0.0 | 0.0 | 0.0 | 14.207764445202717 | 0.0 | 5.868924823489645 | 0.0 |
| SE_47_3_28 | SE_47 | SE_47_3 | GGGCCTTCCGCTAAAGGCAA | 34.680858934025565 | 47.97083962242702 | 58.264084895438096 | 41.641871666124814 | 31.8178353818822 | 31.187163955262715 | 24.75169409358689 | 13.23494886778659 | 35.51941111300679 | 9.477090282749604 | 3.521354894093787 | 6.964086076367696 |
| SE_47_3_29 | SE_47 | SE_47_3 | AGACCCTTGCCTTTAGCGGA | 234.09579780467257 | 322.80377253674845 | 359.459777998974 | 208.20935833062407 | 313.9359757679044 | 235.63634988420716 | 215.2321225529295 | 146.52979103620868 | 151.84548250810403 | 186.62577787568452 | 104.46686185811568 | 46.42724050911798 |
| SE_47_3_4 | SE_47 | SE_47_3 | CTTTATCTCTTGCCCTGCCT | 46.60240419259685 | 60.962941562667666 | 50.36386999436174 | 88.48897729051524 | 39.24199697098805 | 58.90908747105179 | 79.63588534458391 | 6.617474433893295 | 32.855455279531284 | 196.831875103261 | 15.259204541073078 | 35.98111139456643 |
| SE_47_4_20 | SE_47 | SE_47_4 | GCTTTTTCAGAATGGCCTGG | 59.60772629285644 | 86.94714616314897 | 107.6404280271653 | 100.20075369661284 | 146.36204275665813 | 149.0053388973663 | 66.72195799140815 | 119.11453981007931 | 155.39742361940472 | 125.38919451022554 | 184.28423945757487 | 150.88853165463343 |
| SE_47_4_32 | SE_47 | SE_47_4 | AGACACTCCCACATTCTTCA | 908.2049933347945 | 851.4823969080795 | 910.4997673490496 | 852.3570606659923 | 919.5354425363956 | 960.7379118440653 | 893.2133085946574 | 853.6542019722351 | 1001.6473933867915 | 995.0944796887085 | 748.8748074772788 | 1193.180081084332 |
| SE_47_4_4 | SE_47 | SE_47_4 | TAGAAGACAGAACAAATACA | 1471.76895101271 | 1975.7989421212128 | 2036.2803907524299 | 1484.7929865952628 | 1965.2816320875907 | 2312.181683238783 | 1344.1246053430448 | 1218.5606493269224 | 1846.121392598528 | 1330.437674309079 | 1010.6288546049169 | 1829.2332760592483 |
| SE_47_4_8 | SE_47 | SE_47_4 | GCAAGAGGCGGTTTGCCAAT | 5376.616911615651 | 5524.641655056178 | 5542.988279967695 | 5211.740500713434 | 5458.879957018256 | 4528.202944282172 | 5043.964792027903 | 4831.701690232661 | 4959.397776653574 | 5054.205148484847 | 4225.625872912545 | 4251.574549622479 |
| SE_47_4_9 | SE_47 | SE_47_4 | GGTAGATGCAAAAAGATGGG | 352.22747354869716 | 526.6798394020633 | 558.940204251152 | 352.6546006724945 | 440.1467227827038 | 523.2513063605189 | 426.1596026548004 | 231.61160518626534 | 478.6240647477665 | 219.4310903928947 | 199.54344399864794 | 399.2742683784146 |
| SE_47_5_10 | SE_47 | SE_47_5 | TCTCACCACATCAAGCAGGG | 830.173060733237 | 918.4416933785506 | 861.1234242173224 | 909.6146342069139 | 905.7477138709133 | 1110.6095608513 | 790.9780503820159 | 640.9496665970935 | 672.2048553136535 | 486.2476321995374 | 770.0029368418415 | 819.4407949859323 |
| SE_47_5_17 | SE_47 | SE_47_5 | TACAGGTTCCTCCTTGCCAT | 625.3392376541485 | 563.6573613335174 | 585.6034295422846 | 616.8202240544738 | 635.2961131249147 | 912.2245456914344 | 755.4647501607825 | 485.9116941458791 | 656.2211203128005 | 602.1597364270133 | 340.3976397623994 | 863.5466734695943 |
| SE_47_5_2 | SE_47 | SE_47_5 | CTAACACACGCAAAACCGTC | 4249.48899625982 | 3709.7449029610225 | 3735.81412134648 | 3650.1703132337534 | 3493.598324930666 | 2432.598788510492 | 3667.5553683019184 | 3360.7316589272377 | 3198.5229707262615 | 3241.164876700365 | 3104.661231626022 | 2685.815863452475 |
| SE_47_5_6 | SE_47 | SE_47_5 | TACTGGTTGCTCATTTACAA | 740.2195828731082 | 788.520670376144 | 817.6722422614025 | 745.6497645215475 | 757.2644820887964 | 614.2138678967018 | 736.0938591310189 | 745.88390404883 | 946.592306161631 | 773.4763684613331 | 545.810008584537 | 1252.3748127334575 |
| SE_47_5_7 | SE_47 | SE_47_5 | GGTTGGTTACAACCATGTCC | 67.19416418467453 | 89.94532361705066 | 99.74021312608895 | 59.86019052005442 | 72.12042686559965 | 128.21389626052448 | 69.95043982970209 | 42.54090707502832 | 90.57449833816732 | 45.92743752409424 | 97.42415206992811 | 161.33466076918498 |
| SE_48_1_10 | SE_48 | SE_48_1 | TTGAAAGAAAAAAATAACAG | 2.167553683376598 | 8.994532361705065 | 30.613332741670867 | 9.109159426964803 | 28.63605184369398 | 1.7326202197368175 | 7.533124289352532 | 15.12565584889896 | 1.7759705556503396 | 51.030486137882484 | 4.695139858791716 | 47.58792152184593 |
| SE_48_1_12 | SE_48 | SE_48_1 | ATAACAGAGGAAGAGAGGAA | 171.23674098675122 | 168.89732990312845 | 194.54279193900518 | 187.38842249756166 | 165.45274398578746 | 178.4598826328922 | 239.98381664651637 | 196.63352603566865 | 166.05324695330674 | 27.702263903421922 | 123.24742129328254 | 84.7297139291403 |
| SE_48_1_18 | SE_48 | SE_48_1 | TAAAAATACACACATTAGCA | 106.21013048545329 | 143.91251778728105 | 154.05419057098888 | 122.32299801924164 | 144.2408537311993 | 126.48127604078768 | 89.32133085946575 | 108.71565141396128 | 118.10204195074758 | 356.4843960203505 | 190.1531642810645 | 287.84889115653147 |
| SE_48_1_19 | SE_48 | SE_48_1 | ATACACACATTAGCAGGGCG | 649.182328171291 | 638.6117976810597 | 696.2064381573535 | 567.3705014509065 | 721.2042686559965 | 594.2887353697284 | 601.5737825354379 | 599.3541130126213 | 690.8525461479821 | 673.6024170200488 | 589.2400522783604 | 778.816959540454 |
| SE_48_1_8 | SE_48 | SE_48_1 | TTATGTTGTTATTTTGGCAG | 91.0372547018171 | 102.9374259172913 | 111.59053547770347 | 74.17458390528482 | 71.05983235287025 | 97.89304241513018 | 46.27490634887984 | 106.82494443284891 | 127.86988000682445 | 38.63736807582531 | 110.33578668160533 | 25.534982280014887 |
| SE_49_1_14 | SE_49 | SE_49_1 | TCTCGCAGAGCCGGGACCCC | 460.60515771752705 | 512.6883446171887 | 504.626226806252 | 445.04750343170895 | 688.325838761385 | 679.1871261368325 | 487.5007575823853 | 655.1299689554362 | 556.7667691963815 | 361.58744463413876 | 856.8630242294882 | 567.5730152239672 |
| SE_49_1_16 | SE_49 | SE_49_1 | AAGCGCTGCTCTCGCAGAGC | 33.59708209233727 | 77.95261380144339 | 53.32645058226538 | 18.218318853929606 | 57.27210368738796 | 67.57218856973589 | 19.370891029763655 | 20.79777679223607 | 39.07135222430747 | 48.11445835857 4915 | 32.86597901154201 | 132.31763545098624 |
| SE_49_1_17 | SE_49 | SE_49_1 | GCAAATGGAGTGGGTGCTCG | 2852.5006473236026 | 3026.1604434714377 | 2931.9672551616961 | 3222.0398201664075 | 2846.6356721657276 | 3372.5452577177152 | 2906.7098150773127 | 3114.9397513826298 | 3209.1787940601635 | 3431.4356893001836 | 2626.9307509939654 | 3638.7349749021214 |
| SE_49_1_20 | SE_49 | SE_49_1 | GAGCCTGCGCGGACGCCCCG | 403.1649851080472 | 502.6944197708498 | 582.640884954381 | 428.13049306734575 | 505.903585571927 | 379.4438282122363 | 411.09335057609535 | 676.8730992382285 | 520.3593728055495 | 681.6214934321941 | 762.9602270536539 | 349.3649848311128 |
| SE_49_1_9 | SE_49 | SE_49_1 | GGACCCCCGGGCCCTGCTTG | 9.75399157519469 | 45.972054293159225 | 37.526020780112674 | 22.12224432262881 | 24.393673792776355 | 18.192512307236584 | 16.142409191469714 | 17.01636283001133 | 55.94307250298569 | 42.282402799959776 | 16.432989505771005 | 134.63899747644214 |
| SE_49_2_10 | SE_49 | SE_49_2 | AACTGCCTAACAGTCTGGAG | 350.0599198653205 | 378.76975167624664 | 375.26020780112674 | 415.11740817168175 | 381.81402458258646 | 461.74328855986187 | 322.84818382939426 | 398.9391730147101 | 380.9456841869978 | 494.2667085926332 | 416.6936624677648 | 373.7392860983997 |
| SE_49_2_20 | SE_49 | SE_49_2 | AGGTTCTCTCCCATGAACCC | 1.083776841688299 | 1.9987849692677924 | 3.950107450538176 | 2.602616979132801 | 4.242378050917627 | 0.0 | 0.0 | 0.0 | 0.0 | 0.0 | 0.0 | 0.0 |
| SE_49_2_24 | SE_49 | SE_49_2 | AGGCTGATGCAAGAGCTCCA | 1545.4657762475142 | 1639.0036747995898 | 1567.2051310010213 | 1837.4475872677574 | 1599.3765251959453 | 1576.684399960504 | 1577.651458312973 | 1987.1330371491008 | 1604.5893970300817 | 1991.6469732670707 | 2016.5625693510422 | 2039.3165393630072 |
| SE_49_2_8 | SE_49 | SE_49_2 | GCAACTGCCTAACAGTCTGG | 915.7914312266125 | 787.5212778915102 | 704.1066530584299 | 779.4837852502739 | 759.3856711142553 | 1308.9945760111657 | 850.1668840840715 | 933.1581591789546 | 675.7567964249542 | 994.3654724738876 | 1045.8424035458547 | 949.4370684114626 |
| SE_49_3_12 | SE_49 | SE_49_3 | TTCATGCTGAGGGAACCGCA | 56.35639576779154 | 66.95929647047105 | 66.16429979651446 | 55.95626505135522 | 107.12004578567009 | 51.978606592104526 | 44.12258512335055 | 92.64464207450614 | 76.3667338929646 | 50.30147919305559 | 11.73784964697929 | 54.55200759821362 |
| SE_49_3_15 | SE_49 | SE_49_3 | TCTAGGAAGCCGGTACAGGG | 15.172875783636185 | 15.990279754142339 | 12.837849214249072 | 16.917010364363207 | 43.48437502190568 | 0.8663101098684087 | 11.837766740411123 | 32.14201867891029 | 17.759705556503395 | 87.48083337922712 | 14.085419576375148 | 1.1606810127279494 |
| SE_49_3_16 | SE_49 | SE_49_3 | TCCCTGTACCGGCTTCCTAG | 8.670214733506391 | 8.994532361705065 | 11.850322351614528 | 16.917010364363207 | 4.242378050917627 | 43.31550549342044 | 4.30464245105859 | 35.92343264113503 | 14.207764445202717 | 31.347298627556384 | 24.64948425865651 | 22.05293924183104 |
| SE_49_3_4 | SE_49 | SE_49_3 | CCTCTCTTGTCCCTATTTGG | 27.094421042207472 | 59.96354907803377 | 67.151826659149 | 36.43663770785921 | 77.42339942924669 | 20.791442636841808 | 35.513300221233365 | 77.51898622560716 | 125.20592417334893 | 9.477090282749604 | 130.29013108147012 | 107.94333418369929 |
| SE_49_3_7 | SE_49 | SE_49_3 | TGTGCAGCCTGCCCAGTTAG | 7151.843378301084 | 5909.407761640228 | 5808.633006016388 | 7033.572386106394 | 6536.443981951334 | 5869.250994358469 | 7193.057535718904 | 6648.671099081649 | 5974.364949207742 | 6565.436545110995 | 6819.690644894968 | 6115.6282560635655 |
| SE_49_4_10 | SE_49 | SE_49_4 | GGAAAGTCAGGGTTACGCAA | 568.9828418863569 | 620.6227329576495 | 586.5909564049192 | 650.6542447832002 | 633.1749240994558 | 627.2085195447279 | 529.4710214802066 | 634.3321921632001 | 811.6185439322052 | 463.64841690990374 | 714.8350435010387 | 437.57674179843696 |
| SE_49_4_19 | SE_49 | SE_49_4 | GTGTTCTGACTGCAAACACG | 208.0851536041534 | 168.89732990312845 | 151.09160998308525 | 213.41459228888968 | 225.90663121136365 | 311.00532944275875 | 163.5764131402264 | 197.57887952624267 | 300.1390239049074 | 169.858618144666 | 268.79675691582577 | 191.51236710011165 |
| SE_49_4_20 | SE_49 | SE_49_4 | TGTTCTGACTGCAAACACGG | 199.414938870647 | 216.86816916555549 | 185.6550501752943 | 160.06094421666725 | 156.9679878839522 | 131.67913669999814 | 205.54667703804768 | 254.30008895961376 | 259.29170112494955 | 147.9884097998592 | 273.4918967746175 | 77.76562785277261 |
| SE_49_4_21 | SE_49 | SE_49_4 | GTTCTGACTGCAAACACGGG | 40.09974314246706 | 61.96233404730156 | 51.35139685699629 | 49.44972260352321 | 80.60518296743491 | 53.71122681184134 | 30.13249751741013 | 70.90151179171387 | 74.59076333731426 | 149.44642368951298 | 109.1620017169074 | 114.907420260067 |
| SE_49_4_25 | SE_49 | SE_49_4 | CTTGGAAACTCTTCAAACCA | 434.5945135170 0784 | 438.73330075428044 | 450.3122493613521 | 432.03441853604494 | 498.4794209828212 | 128.21389626052448 | 462.7490634887584 | 654.18461546488 | 417.3530805778498 | 413.3469377168381 | 584.5449124195686 | 322.66932153836996 |
| SE_49_5_11 | SE_49 | SE_49_5 | ATCTGCTTGTCCATACAACG | 273.1117641054513 | 217.86756165018937 | 237.00644703229057 | 300.6022610898385 | 250.30030500414 | 386.3743090013103 | 273.34479564222045 | 293.05958207241736 | 360.5220227970189 | 171.31663203431978 | 355.6568443034725 | 358.6504329329364 |
| SE_49_5_12 | SE_49 | SE_49_5 | TGCACATTAGGTTCACGCAT | 10917.967903167922 | 8432.873785340817 | 8392.003278668355 | 9772.826756643668 | 8069.003052845326 | 8009.903275843307 | 10154.651542047213 | 8263.334860951612 | 8154.368806268534 | 7892.22918469594 | 9524.091203558995 | 7030.24489409319 |
| SE_49_5_24 | SE_49 | SE_49_5 | GCCAAATAGTCAGCAAGGCA | 1518.3713552053068 | 1662.9890944308033 | 1674.8455590281867 | 1588.897665760575 | 1590.89176909411 | 1300.3314749124816 | 1550.747442993857 | 1764.9749668683974 | 1561.0781184166485 | 1742.3265981362736 | 1423.801162178588 | 1689.9515545318943 |
| SE_49_5_25 | SE_49 | SE_49_5 | ATACATCACCATCATTCTGT | 641.595890279473 | 642.6093676195952 | 657.6928905146063 | 627.2306919710051 | 540.9032014919975 | 643.6684116322277 | 455.21593919944587 | 449.98826150474406 | 768.1072653187719 | 891.5754935232898 | 562.242998090308 | 678.9983924458504 |
| SE_49_5_7 | SE_49 | SE_49_5 | GGACTCCCGTCCGCTCAGCA | 190.74472413714062 | 202.87667438068092 | 188.61763076319792 | 225.12636869498726 | 207.87652449496372 | 211.37966680789174 | 279.80175931880837 | 227.8301912240406 | 107.44621861684554 | 346.278298792774 | 259.4064771982423 | 307.5804683729066 |
| SE_49_6_12 | SE_49 | SE_49_6 | TCAGTGCTAGACCTGGTCTT | 5998.704818744734 | 5561.619176987632 | 5803.695371703216 | 5629.460525864249 | 5961.601756051995 | 5384.11733283216 | 5753.154635839805 | 5600.27407805484 | 5379.414813064878 | 6712.695947966027 | 5702.247358502539 | 5885.813415543432 |
| SE_49_6_17 | SE_49 | SE_49_6 | CTAGAGCTGGGATAAGGGAC | 617.7527997623304 | 666.5947872508087 | 663.6180516904136 | 650.6542447832002 | 605.5996676684912 | 492.93045251512456 | 474.5868302292095 | 521.8351267870141 | 572.7505041972345 | 295.9768195997184 | 476.5566956673592 | 571.0550582621511 |
| SE_49_6_2 | SE_49 | SE_49_6 | CTACGTGCCTTGCATCACTG | 4655.905311892932 | 3651.7801388522566 | 3602.4979948908167 | 4130.353145883755 | 3604.9607487672533 | 3465.240439473635 | 4718.96428697298 | 3649.064473546874 | 3519.0856560211478 | 3277.6152239417092 | 3474.40349550587 | 3075.804683729066 |
| SE_49_6_23 | SE_49 | SE_49_6 | GGAGAAGATTAGGTGCTCAG | 6023.631686103566 | 7238.59976620391 | 7037.116423133761 | 6533.869926128897 | 7407.192076902176 | 6867.240420926876 | 6283.701817932777 | 7288.675412188187 | 6958.25263703803 | 7508.042524772168 | 7742.28562714754 | 6309.461985189133 |
| SE_49_6_8 | SE_49 | SE_49_6 | GGTCACTGTTCAATGGTAAA | 461.68893455921534 | 633.6148352578901 | 590.5410638554573 | 536.139097701357 | 626.8113570230794 | 492.93045251512456 | 501.4908455483257 | 677.8184527287847 | 673.0928405914786 | 399.49580576513716 | 509.4226746789012 | 560.6089291475996 |
| SE_49_7_12 | SE_49 | SE_49_7 | AAGCTGTTGCCCGAGCCCGG | 171.23674098675122 | 250.84751364310793 | 225.15612468067604 | 180.88188004972966 | 236.5125763386577 | 263.3582733999963 | 175.41417988063753 | 206.08706094124832 | 210.45251084456524 | 414.8049516065019 | 138.50662583435562 | 212.40462532921475 |
| SE_49_7_22 | SE_49 | SE_49_7 | AGTAGCGGCCCGTCCTCGCG | 6096.244734496681 | 6814.857352718538 | 6659.881161607365 | 6235.870282002191 | 6771.895963777262 | 6083.229591495966 | 5910.274085303444 | 6493.631266430435 | 6817.062977863829 | 7669.153059578911 | 7202.344543386493 | 6546.240911785635 |
| SE_49_7_27 | SE_49 | SE_49_7 | AGTGCTGGTCCAATGTGGCG | 1333.0455152766076 | 1490.0941945891393 | 1452.6520149354142 | 1495.203454511794 | 1477.4081562320637 | 1270.0106210670872 | 1283.8596110282244 | 1481.368919701542 | 1617.0211909196341 | 1207.2355006333341 | 1280.5993964854406 | 1497.2785064190548 |
| SE_49_7_35 | SE_49 | SE_49_7 | CAGCGTCTCCACGCGCTCCG | 976.4829343611573 | 808.508520068822 | 821.6223497119406 | 1025.4310897783234 | 843.1726376198784 | 984.1282848105124 | 969.6207121009473 | 860.2716764061283 | 750.3475597622685 | 914.9037157577503 | 743.0058826537891 | 1028.363377276963 |
| SE_49_7_36 | SE_49 | SE_49_7 | GGAGGCATCCGCGGAGCGCG | 909.2887701764828 | 931.4337956787913 | 972.7139596950259 | 1009.8153879035267 | 1166.6539640023475 | 887.967862615119 | 872.7662569521291 | 948.1895510278536 | 965.2399969959596 | 1187.5523131230082 | 934.3328318995515 | 735.87176206952 |
| SE_49_8_12 | SE_49 | SE_49_8 | ACTGTGCCTGGCACTTAATG | 2536.0378095506194 | 2570.437470478381 | 2476.7173714874366 | 2695.0098818920155 | 2770.2728672492103 | 2899.539937729564 | 2884.1104422092553 | 2786.902090159633 | 2536.97393874651 | 4215.84716193392 | 2537.7230936769224 | 3075.804683729066 |
| SE_49_8_15 | SE_49 | SE_49_8 | TATAATGTCAGAAGAAGTCC | 99.70746943533235 | 111.93195827899638 | 142.20386821973433 | 81.98243484268323 | 135.75609762936406 | 58.04277736118338 | 61.3411549275849 | 139.91231660231537 | 147.40555615897818 | 64.88161808959345 | 30.518409082146157 | 34.820430381838484 |
| SE_49_8_2 | SE_49 | SE_49_8 | TCTAGGGAAAGTTTTTCCTG | 187.49339361207572 | 215.86877668092157 | 245.89418879600146 | 197.79889041409285 | 212.11890254588135 | 407.16575163815213 | 185.09962539551935 | 156.9286794323267 | 303.69096501620805 | 152.36245146882055 | 179.58909959978315 | 424.8092506584295 |
| SE_49_8_4 | SE_49 | SE_49_8 | TCTTGAAATCCAGTGTTCTA | 293.703524097529 | 378.76975167624664 | 332.79655270784133 | 309.7114205168033 | 366.9657014043747 | 531.914407459203 | 253.9793906124568 | 258.08150292183853 | 413.8011394665291 | 673.6024170200488 | 274.6656817393154 | 521.1457747148493 |
| SE_49_8_7 | SE_49 | SE_49_8 | AGTGTCTACAAGCTTAGAAA | 83.45081680999901 | 120.92649064070145 | 115.54064292824165 | 80.68112635311682 | 77.42339942924669 | 263.3582733999963 | 72.10276105523138 | 80.35504669727572 | 143.85361500767752 | 67.79764586890101 | 24.64948425865651 | 46.42724050911798 |
| SE_49_9_11 | SE_49 | SE_49_9 | TATTAAGGAAACAACCTCAG | 124.63433679415438 | 165.89915244922576 | 163.9294591973343 | 180.88188004972966 | 153.7862043457 64 | 123.88234571118245 | 112.99686434028798 | 147.47514452676486 | 123.42995361769 86 | 136.32429868262892 | 64.5581730583861 | 87.05107595459621 |
| SE_49_9_2 | SE_49 | SE_49_9 | GTTCTATTCAAAACATTCGA | 2.167553683376598 | 0.9993924846338962 | 4.93763431317272 | 0.0 | 5.3029725636470335 | 0.8663101098684087 | 0.0 | 0.945353490556185 | 4.439926389125849 | 0.7290069448268927 | 0.0 | 0.0 |
| SE_49_9_23 | SE_49 | SE_49_9 | TAAGAATGATAAAAGCAGAG | 226.50935991285448 | 336.795267321623 | 295.27053192772865 | 236.83814510108488 | 305.45121966606916 | 540.5775085578871 | 242.13613787204568 | 166.38221433788857 | 275.2754361258026 | 399.49580576513716 | 238.2783478336796 | 338.9188557165612 |
| SE_49_9_24 | SE_49 | SE_49_9 | GGAGGAAAGAGATTTCACAT | 3097.43421354515 84 | 3689.757053268345 | 3842.467022511011 | 3322.2405738630205 | 3580.5670749744772 | 3543.2083493617915 | 3184.3592531705917 | 3120.6118723259665 | 3404.5355551817 01 | 3308.233515624439 | 3320.6376651304413 | 4082.115121764198 |
| SE_49_9_26 | SE_49 | SE_49_9 | GACCTATTTTTGCAGAAACA | 54.188842084414944 | 100.93864094802352 | 66.16429979651446 | 65.06542447832003 | 56.21150917465856 | 208.7807364782865 | 79.63588534458391 | 103.98888396118035 | 90.57449833816732 | 81.64877782061198 | 78.64359263476125 | 13.928172152735392 |
| SE_5_1_18 | SE_5 | SE_5_1 | GCTGACACTGGTACAGGGAA | 2066.762437099586 | 2254.6294453340697 | 2358.2141479712914 | 2128.940688930631 | 2503.0030500414 | 2082.6095041236545 | 2056.5429309932415 | 2318.952112334322 | 2002.406801495758 | 2337.9252720598447 | 1955.5257511867499 | 1782.8060355501302 |
| SE_5_1_20 | SE_5 | SE_5_1 | GCTCTCCCGCCCATGAACAC | 9.75399157519469 | 29.98177453901 6886 | 14.81290293951816 | 18.218318853929606 | 19.090701229129323 | 6.064170769078861 | 7.533124289352532 | 23.633837263904624 | 17.759705556503395 | 38.63736807582531 | 23.47569929395858 | 1.1606810127279494 |
| SE_5_1_26 | SE_5 | SE_5_1 | TTTAGTTTACCAGTGGAGGA | 2.167553683376598 | 12.992102300240651 | 10.862795488979984 | 9.109159426964803 | 7.4241615891058474 | 0.8663101098684087 | 3.228481838293942 | 1.89070698111237 | 0.0 | 17.496166675845423 | 9.390279717583432 | 1.1606810127279494 |
| SE_5_1_29 | SE_5 | SE_5_1 | CTTCCTCTCTGTTAATGAGG | 33.59708209233727 | 55.96597913498914 | 28.63827901640178 | 40.340563171655 81 | 74.24161589105847 | 54.57753692170975 | 25.82785474061531 38 | 41.59555358447214 | 30.19149944605577 2 | 61.23658336545898 | 39.90868879972959 | 18.57089620364719 |
| SE_5_1_8 | SE_5 | SE_5_1 | CCTGCCCTCACCCAAGTCAG | 3047.5804788274963 | 2584.4289652632556 | 2540.906661755868 2 | 2754.87007241207 | 2821.181403860222 | 2906.4704186085114 | 3177.902289494004 | 2593.1046245956154 | 2458.831234297895 | 3957.7787034652 | 3407.497752518088 | 2526.802564708746 |
| SE_5_2_10 | SE_5 | SE_5_2 | GCCGCCGCGTAAGTGCCGCG | 281.78197883895774 | 263.8396159433486 | 295.27053192772865 | 288.8904846837409 | 336.20846053522195 | 151.60426922567152 | 174.33801926787288 | 294.95028905352973 | 248.63587779104753 | 200.47690982739547 | 237.10456286898167 | 234.4575645710458 |
| SE_5_2_12 | SE_5 | SE_5_2 | GCACTTACGCGGCGGCTCGG | 488.7833556014228 | 809.5079125534559 | 809.7720273603261 | 586.8901287944466 | 864.3845278744665 | 595.1550454795968 | 386.3416599825084 | 519.9444198059017 | 725.4839719831638 | 809.1977087578508 | 581.0235575254749 | 1098.00423804064 |
| SE_5_2_13 | SE_5 | SE_5_2 | TCTGGAACTGCCTTGCACA | 1405.6585636697237 | 1253.2381757309058 | 1274.8971796611963 | 1461.3694337830677 | 1369.227515933664 | 1192.0427111789304 | 1529.224230738564 | 1156.1673189502142 | 1242.2914036774125 | 1807.9372231706939 | 1190.2179542037 | 1301.1234152680313 |
| SE_5_2_18 | SE_5 | SE_5_2 | GCGAGCGGCCAGGAGGGCTC | 3810.559375376059 | 3793.69387167027 | 3619.285951555604 | 3833.6548102626157 | 3400.266007810478 | 4129.700293742704 | 4163.6654107864215 | 3870.2771903370212 | 3687.80285880793 | 3556.824883810409 | 3818.322490162363 | 3350.88608374559 |
| SE_5_2_9 | SE_5 | SE_5_2 | CCGCCGCGTAAGTGCCGCGC | 561.3964039945388 | 452.72479553915497 | 476.97547465248476 | 476.27890718130254 | 429.5407776554097 | 336.994632738811 | 493.9577212589732 | 513.3269453720085 | 361.4100080748441 | 318.5760348893521 | 397.9131030325979 | 244.90369368559732 |
| SE_5_3_11 | SE_5 | SE_5_3 | ACGTGTTTGGATGTTTTGGT | 17152.936073400706 | 16836.7651882725 | 17316.28353629673 | 17153.84850946429 | 17018.29955125606 | 17181.52840902015 | 17345.556756540587 | 15202.22948163401 | 18369.7516434236929 | 18214.9675244474 | 15103.091140768252 | 15000.641408496018 |
| SE_5_3_12 | SE_5 | SE_5_3 | GAGTTCTGCTTCCTAGAATG | 55.27261892610324 | 63.96111901656935 6 | 47.40128940645811 | 80.68112635311682 | 31.8178353818822 | 6.93048087894727 | 52.73187002546773 | 53.88514896170255 | 31.07948472388094 | 129.03422923436 | 65.73195802308402 | 63.83745570003722 |
| SE_5_3_14 | SE_5 | SE_5_3 | TGGACAGGTTCCCGAACAAT | 441.09717456713764 | 410.7503111845313 | 423.6490240702194 | 394.2964723386193 | 404.08650934990396 | 702.5774991032795 | 460.5967422632691 | 399.88452650526625 | 543.4469900290039 | 291.60277793075704 | 494.16347013782814 | 529.270541803945 |

| | | | | | | | | | | | | | | | |
|---|---|---|---|---|---|---|---|---|---|---|---|---|---|---|---|
| SE_5_3_17 | SE_5 | SE_5_3 | GAGTGGAGCCATTAGCTAGG | 1251.7622521499852 | 1345.1822843172242 | 1339.0864257324417 | 1289.596713160303 | 1266.3498481989116 | 1148.72720568551 | 1389.32335107916 | 1271.5004447980689 | 1448.3039881328518 | 1224.0026603643528 | 1497.7496149545575 | 1081.7547038624489 |
| SE_5_3_5 | SE_5 | SE_5_3 | TACATGCTCTTTGTCAACAA | 2258.590938078415 | 2144.6962720243414 | 2215.0227528892824 | 2229.141442627244 | 2389.519437179353 | 2379.753871808519 | 2195.367650039881 | 2629.9734107273066 | 2160.4681809486383 | 2138.906376122103 | 1851.058889328634 | 2964.3793065071827 |
| SE_5_4_52 | SE_5 | SE_5_4 | GCCAAGGAAGAGGCTCTTGG | 309.9601767228535 | 623.6209104115512 | 600.4163324818028 | 343.54544124552973 | 514.3883386737623 | 394.171099990126 | 199.0897133614598 | 270.37109829906893 | 506.15160836034676 | 423.5530349444246 | 266.4491869864299 | 356.32907090748046 |
| SE_5_4_53 | SE_5 | SE_5_4 | TCCACCAAGAGCCTCTTCCT | 138.72343573610226 | 142.91312530264716 | 198.49289938954334 | 130.13084895664005 | 216.36128059679896 | 63.24063802039384 | 117.30150679134657 | 179.61716320567515 | 119.87801250639792 | 147.9884097998592 | 123.24742129328254 | 257.67118482560477 |
| SE_5_4_54 | SE_5 | SE_5_4 | GGAAGAGGCTCTTGGTGGAG | 11684.19813024155 | 16120.200777144746 | 15519.972173164493 | 13567.442312219291 | 15815.585373820913 | 14850.287903364262 | 12523.281050742202 | 17120.35171397251 | 14973.207754688014 | 15367.466396950897 | 17991.775938889856 | 17147.901282042723 |
| SE_5_4_55 | SE_5 | SE_5_4 | GGAGAGGACTGTGAAGCCGT | 4833.644713929813 | 5194.842135126993 | 5123.289363348014 | 5246.875829931727 | 5164.0346824794815 | 5622.352613045972 | 4922.358642785498 | 4972.559360325533 | 4901.6787335949375 | 4910.5907803539485 | 4503.8129095459535 | 5812.690511741571 |
| SE_5_4_56 | SE_5 | SE_5_4 | AGGACTGTGAAGCCGTCGGC | 2294.3555738541286 | 2934.216334885119 | 2984.306178881592 | 2611.726138559766 | 2736.3338428418692 | 2172.70575549969 | 2432.1229848481034 | 2421.05028931439 | 3201.186926559737 | 2232.948272004772 | 2223.1487231378774 | 2534.9273317978414 |
| SE_5_5_1 | SE_5 | SE_5_5 | ACTAGCAGCTTCCTTCCCTA | 1691.7756498754345 | 1721.9532510242032 | 1714.3466335335684 | 1761.9716948729063 | 1641.8003057051217 | 1728.2886691874753 | 1562.5852097342681 | 1805.6251669623134 | 1676.5162045339205 | 2077.6697927566443 | 1900.3578578459471 | 2377.0747140668404 |
| SE_5_5_15 | SE_5 | SE_5_5 | AATTTCCCACAGTCTTTCAA | 0.0 | 3.9975699385355847 | 1.975053725269088 | 2.602616979132801 | 5.3029725636470335 | 0.0 | 0.0 | 0.0 | 0.0 | 0.0 | 0.0 | 0.0 |
| SE_5_5_19 | SE_5 | SE_5_5 | GGAAATGCAAACATGTCTGG | 748.8897976066146 | 913.4447309553811 | 839.3978332393624 | 670.1738721266962 | 847.415015670796 | 686.1176070157798 | 726.4084136161371 | 1021.927123291236 | 786.7549561531005 | 801.178632364755 | 988.3269402756563 | 1019.0779291751396 |
| SE_5_5_2 | SE_5 | SE_5_5 | AATGACACTACGGTGCAACG | 877.8592417675221 | 759.5382883217611 | 712.9943948221407 | 782.0864022294066 | 671.3563265577145 | 866.3101098684087 | 881.3755418542463 | 1002.074699989556 | 818.7224261548065 | 753.793180951007 | 678.447709595403 | 611.6788937076293 |
| SE_5_5_3 | SE_5 | SE_5_5 | CTAGCAGCTTCCTTCCCTAG | 1451.1771910206323 | 1319.198079716743 | 1250.2090080953328 | 1242.7496075359124 | 1220.7442841515472 | 1707.4972265506335 | 1373.18094188769 | 1046.5063140456969 | 1482.0474286902083 | 1139.4378547644333 | 916.7260574290826 | 1159.5203317152216 |
| SE_5_6_17 | SE_5 | SE_5_6 | CTTCCAGTCACCACCCCCCA | 1209.4949553241415 | 1520.0759691281562 | 1544.492013160427 | 1300.007181076834 | 1516.6501532030516 | 1216.299394255246 | 1204.2237256836406 | 1503.1120499843341 | 1227.1956539543846 | 1019.1517088679959 | 1766.5463718703832 | 1412.5487924899144 |
| SE_5_6_23 | SE_5 | SE_5_6 | AGGAGGCCAATCAGATAGAA | 2529.53514850049 | 2999.1768463863223 | 2996.1565012332067 | 2632.547074392828 | 2807.3936751947394 | 3264.256493984164 | 2341.7254933758727 | 2952.338951006966 | 2885.0641676539767 | 2001.1240635498204 | 2563.546362900277 | 4003.1888128986975 |
| SE_5_6_27 | SE_5 | SE_5_6 | AATCAGATAGAATGGAGAGG | 18.42420630870108 | 30.98116702365078 | 60.23913862070719 | 37.737946197425615 | 66.81745430195262 | 0.8663101098684087 | 20.4470516425283 | 85.08181415005664 | 6.215896944776189 | 29.160277793075707 | 50.47275348201095 | 58.034050636397474 |
| SE_5_6_4 | SE_5 | SE_5_6 | CCTATCAGCGACAGTCACCA | 1880.3528203291985 | 1859.8694139036809 | 1893.088995670421 | 1752.8625354459414 | 2141.3403212006724 | 1607.0052538058983 | 1745.5325139042582 | 2088.2858606386126 | 1756.434879538186 | 1676.715973101853 | 1981.3490204101042 | 1826.9119140337923 |
| SE_5_6_7 | SE_5 | SE_5_6 | CTTTAGAGGCAAATATCCAG | 53.105065242726646 | 50.9690167163287 | 60.23913862070719 | 46.847105624390416 | 96.51410065837601 | 25.989303296052263 | 27.980175931880833 | 116.27847933841076 | 55.05508722516053 | 63.42360419993966 | 157.2871852695225 | 30.177706330926686 |
| SE_50_1_22 | SE_50 | SE_50_1 | GCGGTGGTGTGCTAGATGCA | 1711.283633025824 | 1593.0316205064305 | 1545.4795400230614 | 1624.0329949788677 | 1738.3144063634977 | 1237.0908368920877 | 1652.9827012064984 | 1280.0086262130744 | 1725.3553948143049 | 1115.3806255851457 | 976.589090628677 | 1091.0401519642726 |
| SE_50_1_23 | SE_50 | SE_50_1 | GCTTCAACTCTCACGCCTGG | 909.2887701764828 | 1077.34509843534 | 1120.8429890902075 | 938.2434209773747 | 1156.0480188750535 | 848.9839076710406 | 957.7829453605362 | 947.2441975372974 | 989.2155994972392 | 1491.5482091158224 | 907.3357777114992 | 1091.0401519642726 |
| SE_50_1_26 | SE_50 | SE_50_1 | CTGAAGCCACCGCAAGGCGA | 6.5026610501297935 | 5.9963549078033775 | 9.87526862634544 | 11.711776406097604 | 11.666539640023474 | 0.0 | 0.0 | 0.945353490556185 | 0.8879852778251698 | 0.7290069448268927 | 1.173784964697929 | 0.0 |
| SE_50_1_4 | SE_50 | SE_50_1 | GCTGCAGGCACTCCAGACCG | 1018.750231187001 | 313.8092401750434 | 569.8029997401319 | 913.5185596756131 | 242.87614341503414 | 2309.5827529091775 | 1422.684330074864 | 464.16856386308683 | 1717.3635273138784 | 80.91977087578509 | 180.76288456348107 | 475.87921521845925 |
| SE_50_1_7 | SE_50 | SE_50_1 | GGCCAAGTCTATAAACTGGG | 122.46678311077778 | 169.89672238776234 | 180.71741586212156 | 170.47141213319847 | 168.63452752397566 | 200.9839454894708 | 123.75847046793446 | 111.55171188562983 | 103.89427750554486 | 105.70600699989943 | 161.98232512831422 | 62.67677468730927 |
| SE_50_2_12 | SE_50 | SE_50_2 | AATGAGATTTATCCTCACAA | 42.26729682584366 | 63.961119016569356 | 50.36386999436174 | 52.05233958265602 | 80.60518296743491 | 75.36897955855156 | 61.3411549275849 | 108.71565141396128 | 71.92680750383876 | 106.43501394472632 | 118.55228143449084 | 123.03218734916264 |
| SE_50_2_14 | SE_50 | SE_50_2 | GCTGCTGCAATGGGAAGCCT | 2279.1826980704927 | 2464.501867107188 | 2292.049848174777 | 2562.2764159562425 | 2596.335367161588 | 1821.8501610532635 | 2426.74218178428 | 2509.913517426671 | 2187.107739283393 | 2281.0627303633473 | 2298.270960878545 | 1513.528040597246 |
| SE_50_2_18 | SE_50 | SE_50_2 | TAGCTTATGATTATTCAGAG | 87.78592417675222 | 137.91616287947767 | 116.52816979087619 | 148.34916781056964 | 128.33193604025823 | 176.7272624131554 | 108.6922218892294 | 152.2019119795458 | 168.71720278678225 | 69.98466670338169 | 107.98821675220947 | 121.87150633643469 |
| SE_50_2_3 | SE_50 | SE_50_2 | CCAAATTTAAAATCTGCCAG | 20.59175999207768 | 28.982382054382988 | 28.63827901640178 | 15.615701874796805 | 51.96913112374093 | 37.251334724341575 | 1.0761606127646475 | 33.08737216946648 | 38.1833669464823 | 6.561062503442034 | 75.12223774066746 | 84.7297139291403 |
| SE_50_2_7 | SE_50 | SE_50_2 | AATATTGGAGAAGTCATCCT | 33.59708209233727 | 45.972054293159225 | 53.32645058226538 | 41.641871666124814 | 45.60556404736449 | 136.8769973592086 | 40.8941032850566 | 44.431614056140695 | 29.303514168230603 | 5.832055558615141 | 73.94845277596953 | 31.338387343654635 |
| SE_50_3_1 | SE_50 | SE_50_3 | AGCCTCCCAGAAGTGCTCCA | 32.51330525064897 | 45.972054293159225 | 37.526020780112674 | 37.737946197425615 | 33.939024407341016 | 74.50266944868315 | 35.513300221233365 | 36.868786131691216 | 38.1833669464823 | 46.65644446892113 | 5.868924823489645 | 19.73157721637514 |
| SE_50_3_11 | SE_50 | SE_50_3 | ATTGAAGTGCTTGATTCTAA | 106.21013048545329 | 153.90644263362 | 170.8421472357761 | 93.69421124878083 | 130.45312506571702 | 222.64169823618104 | 88.2451702467011 | 100.20746999895562 | 276.1634214036278 | 79.4617569861313 | 123.24742129328254 | 206.601220265575 |
| SE_50_3_3 | SE_50 | SE_50_3 | ATGGCCTCTTCCAGAAAGCA | 607.9988081871356 | 512.6883446171887 | 539.189666998461 | 534.8377892117906 | 486.8128813427977 | 367.3154865842053 | 450.9112967483873 | 582.33775018261 | 752.1235303179188 | 343.36227101346645 | 527.0294491493702 | 581.5011873767027 |
| SE_50_3_4 | SE_50 | SE_50_3 | TCTCCACGCAGAAATGGACT | 809.5813007411593 | 776.5279605605373 | 782.1212752065588 | 757.361540927645 | 677.7198936340909 | 509.39034460262434 | 772.6833199650168 | 894.304402066151 | 737.0277805948909 | 924.3808060404999 | 786.4359263476125 | 694.0872456113137 |
| SE_50_3_5 | SE_50 | SE_50_3 | CAGCCGAGTCCATTTCTGCG | 1076.1904037964807 | 1227.2539711304246 | 1266.0094378974854 | 1102.208290662741 | 1396.8029732646287 | 1671.9785120460288 | 1132.1209646284092 | 1126.8613607429725 | 1118.861450059714 | 1573.9259938812613 | 1498.9233999192554 | 1752.6283292192036 |
| SE_51_1_10 | SE_51 | SE_51_1 | ACTAAGAGCATTGTCACCCA | 2220.6587486193243 | 2078.736368038504 | 2024.4300684008153 | 2046.9582540879478 | 2083.0076230005548 | 2315.6469236782564 | 2075.913822023005 | 2483.443619691098 | 1687.1720278678226 | 2083.5018483152594 | 2522.4638891358495 | 1940.6586532811314 |
| SE_51_1_15 | SE_51 | SE_51_1 | CGAGTACTATGGGGTCCACC | 2851.4168704819144 | 3459.8967818025485 | 3269.7014421829754 | 2942.2584949096313 | 3259.206937617467 | 3539.7431089223182 | 2657.0405529159148 | 2872.9292578002464 | 3073.3170465529124 | 3240.435869755538 | 3011.932219414886 | 2801.88396472527 |
| SE_51_1_20 | SE_51 | SE_51_1 | ACAGTGGCCAAACTACACAC | 9642.362560500795 | 8284.963697615 | 8331.764140047648 | 8850.19903754109 | 8122.032778481796 | 7850.50221562752 | 9305.560818575907 | 8240.646377178266 | 8178.3444087698135 | 8003.038240309627 | 8635.535985282664 | 8207.17544099933 |
| SE_51_1_25 | SE_51 | SE_51_1 | CTATGAGACATACTCCAAGT | 178.8231788785693 | 217.86756165018937 | 198.49289938954334 | 158.75963572710086 | 299.0876525896927 | 77.10159977828837 | 150.66248578705066 | 175.8357492434504 | 217.5563930671666 | 188.0837917653383 | 228.88806811609615 | 206.601220265575 |
| SE_51_1_5 | SE_51 | SE_51_1 | GCATGGAAACCTCTAAGGAG | 234.09579780467257 | 385.76549906868394 | 388.0980570153758 | 204.30543286192486 | 415.75304898992744 | 285.8823362565749 | 275.49711686774975 | 339.3819031096704 | 435.1127861343332 | 247.8623612411435 | 423.7363722559524 | 237.93960760922963 |
| SE_51_2_27 | SE_51 | SE_51_2 | AACTCGCTAATTCATCTGAG | 55.27261892610324 | 66.95929647047105 | 71.10193410968716 | 87.18766880094883 | 64.69626527649382 | 129.94651648026132 | 77.48356411905462 | 58.61191641448347 | 71.92680750383876 | 132.67926395849446 | 28.170839152750297 | 32.49906835638259 |
| SE_51_2_3 | SE_51 | SE_51_2 | ACCCATTTCAACCACCCCAG | 193.9960546622055 | 279.82989569749094 | 289.3453707519214 | 231.63291114281927 | 373.3292684807512 | 178.4598826328922 | 204.47051642528302 | 253.35473546905757 | 142.96562972985234 | 505.9308197098635 | 226.5404981867003 | 392.3101823020469 |
| SE_51_2_30 | SE_51 | SE_51_2 | TCCAAGGCCACCTCTCCCCT | 1631.0841467408898 | 1440.1245703574443 | 1412.163413567398 | 1562.871495969247 | 1537.8620434576399 | 1051.7004733802482 | 1683.1151983639086 | 1592.9206315871718 | 1419.0004739646213 | 1869.1738065361528 | 2115.160506385668 | 1168.805779817045 |
| SE_51_2_6 | SE_51 | SE_51_2 | GGACTGTATTTGGTAGCCAT | 365.2327956489567 | 487.70353250134133 | 448.337195636083 | 413.81609968211535 | 540.9032014919975 | 194.05346461052355 | 355.1330022123337 | 356.3982659396817 | 468.85622669168964 | 560.6063405718804 | 380.306328562129 | 314.5445544492743 |
| SE_51_2_7 | SE_51 | SE_51_2 | TTAGGAGAAGGGAACCCCTG | 676.2767492134985 | 697.5759542744595 | 734.7199858001007 | 631.1346174397042 | 743.4767534233141 | 616.812798226307 | 617.7161917269076 | 697.6708760304646 | 624.2536503110944 | 608.7207989304553 | 766.4815819477477 | 408.5597164802382 |
| SE_51_3_18 | SE_51 | SE_51_3 | CCCCTCACTTAGACACCCGT | 128.96944416090756 | 144.91191027191493 | 117.51569665351074 | 141.84262536273764 | 130.45312506571702 | 370.78072702367893 | 88.2451702467011 | 103.04353047062416 | 204.23661389978906 | 165.48457647570464 | 153.7658303754287 | 646.4993240894678 |
| SE_51_3_19 | SE_51 | SE_51_3 | AATGGAGTGGTAACTAGAGA | 112.71279153558308 | 222.86452407335884 | 218.24343664223423 | 157.45832723753446 | 260.90625013143404 | 309.27270922302193 | 102.23525821264151 | 172.99968877178185 | 238.86803973497067 | 206.3089653860106 | 268.79675691582577 | 234.4575645710458 |
| SE_51_3_2 | SE_51 | SE_51_3 | AGAGAAACCTGAGCAGCATG | 2240.166731769714 | 2207.6579985562767 | 2198.234796224495 | 2551.8659480397114 | 2414.9737054848592 | 2935.9249623440373 | 2436.427627299162 | 2378.5093822393615 | 2304.321795956316 | 2839.482050100747 | 2133.941065820835 | 2710.190164719762 |
| SE_51_3_30 | SE_51 | SE_51_3 | GTATCTGGTGTTGGGAAGAT | 209.16893044584168 | 242.85237376603678 | 241.94408134546327 | 231.63291114281927 | 167.57393301124625 | 296.2780575749958 | 176.4903404934022 | 294.0049355629735 | 256.62774529147407 | 202.66393066187615 | 180.76288456348107 | 119.5501443109788 |
| SE_51_3_6 | SE_51 | SE_51_3 | TAAATTAAGGAGGTTAAGCA | 22.759313675454276 | 60.962941562667666 | 41.47612823065085 | 27.32747828089441 | 77.42339942924669 | 5.197860659210452 | 6.456963676587884 | 17.961716320567515 | 68.37486639253807 | 19.683187510326103 | 11.73784964697929 | 2.321362025455899 |
| SE_51_4_10 | SE_51 | SE_51_4 | CTACCTGCCATGTTTCTGGG | 56.35639576779154 | 22.986027146579612 | 68.13935352178353 | 58.55888203048802 | 46.6661585600939 | 67.57218856973589 | 66.72195799140815 | 80.35504669727572 | 106.55823333902038 | 129.7632361791869 | 71.60088284657367 | 76.60494684004466 |
| SE_51_4_15 | SE_51 | SE_51_4 | GCCACCAATAAGATGCCAGA | 611.2501387122006 | 632.6154427732563 | 622.1419234597628 | 577.7809693674818 | 707.4165399905143 | 662.7272340493327 | 501.4908455483257 | 901.8672299906005 | 621.5896944776189 | 720.9878684337968 | 922.5949822525722 | 598.911402567622 |
| SE_51_4_17 | SE_51 | SE_51_4 | AAAGCCACACAGCTGTCCCG | 900.6185554429763 | 977.4058499719505 | 1060.6038504695002 | 956.4617398313043 | 916.3536589982074 | 1109.7432507414317 | 959.9352665860655 | 1251.648021496389 | 1078.9021125575812 | 878.4533685164057 | 1493.0544750957658 | 1142.1101165243022 |
| SE_51_4_2 | SE_51 | SE_51_4 | CACACCCTCAAGCTGAAGGG | 848.597267041938 | 632.6154427732563 | 650.7802024761645 | 812.0164974894338 | 828.3243144416666 | 988.4598353598543 | 851.2430446968361 | 1248.8119610247204 | 420.9050216891305 | 1060.705104723129 | 1144.4403405804808 | 1328.9797595735022 |
| SE_51_4_7 | SE_51 | SE_51_4 | GTTGATTGACGATGGCCTCA | 169.06918730337463 | 242.85237376603678 | 223.18107095540694 | 235.53683661151848 | 249.2397104914106 | 160.2673703256556 | 222.76524684228204 | 155.03797245121433 | 249.5238630688727 | 235.46924317908633 | 198.36965903395 | 320.34795951291403 |
| SE_51_5_12 | SE_51 | SE_51_5 | TGGAGCTAGATGGCCAGGCG | 475.77803350116324 | 423.742413484772 | 494.75095817990655 | 473.67629020216975 | 460.2980185245625 | 380.31013823223145 | 436.92120878244685 | 314.8027123552096 | 573.6384894750597 | 454.17132662715414 | 307.5316607508574 | 354.0077088820246 |
| SE_51_5_23 | SE_51 | SE_51_5 | GTTTCCATCGTAACACACAG | 88.86970101844051 | 58.96415659339988 | 46.41376254382357 | 84.58505182181602 | 59.39329271284678 | 27.72192351578908 | 106.5399006637001 | 69.95615830115769 | 139.41368861855165 | 104.24799311024564 | 119.72606639918877 | 92.85448101823596 |
| SE_51_5_26 | SE_51 | SE_51_5 | AGAGGCCACAGCCTGCCTGA | 1708.032302500759 | 1631.0085349225185 | 1630.4068502096322 | 1668.2774836241254 | 1660.891006934251 | 1632.128246992082 | 1767.055726159551 | 1585.3578036627223 | 1532.662589526243 | 1920.933299618862 | 1684.3814243415281 | 1122.3785393079272 |
| SE_51_5_3 | SE_51 | SE_51_5 | TACATTTAATTAAATGACCA | 7.586437891818092 | 12.992102300240651 | 17.77548352742179 | 11.711776406097604 | 38.181402458258646 | 4.331550549342044 | 11.837766740411123 | 14.180302358342775 | 28.415528890405433 | 31.347298627556384 | 15.259204541073078 | 0.0 |
| SE_51_5_40 | SE_51 | SE_51_5 | CCTGTTGGTGACTCTGTTGA | 896.2834480762232 | 981.403419910486 | 1046.7784743926168 | 910.9159426964803 | 1014.9889486820423 | 644.534721742096 | 863.0808114372472 | 949.1349045184097 | 984.7756731081133 | 1000.1975283024967 | 899.1192829586137 | 883.2782506859695 |
| SE_51_6_0 | SE_51 | SE_51_6 | AGGCTGTGAGCAAAGCCCTG | 1765.472475110239 | 1997.7855767831584 | 2039.2429713403335 | 1972.7836701826632 | 1989.675305880367 | 2105.1335669802334 | 1594.8700281172075 | 1687.4559806427903 | 1965.1114198271007 | 2133.803327508315 | 1963.7422459396353 | 1773.5205874483067 |
| SE_51_6_12 | SE_51 | SE_51_6 | ACACCAAGCTCATTCCACAA | 5677.906873604998 | 5435.695723923762 | 5526.200323302908 | 5719.25081164433 | 5623.272106491315 | 5361.593269975581 | 5760.687760129158 | 5355.427524000788 | 5451.341620568717 | 5313.73162084322 | 4413.431467264213 | 5844.028899085225 |
| SE_51_6_16 | SE_51 | SE_51_6 | GGTCATTTCCTTTAAGACCT | 54.188842084414944 | 52.9678016855965 | 42.463655093285396 | 65.06542447832003 | 68.93864332741144 | 42.44919538355203 | 40.8941032850566 | 55.775855942814914 | 82.5826308377408 | 66.33963197924723 | 39.90868879972959 | 33.659749369110536 |
| SE_51_6_2 | SE_51 | SE_51_6 | GTATGTGTCTGTGATGGAGA | 4214.808137325795 | 4438.302024259133 | 4235.502713839559 | 4360.684748537008 | 4540.405108994591 | 4327.218998792701 | 4363.831284760646 | 4672.882303819222 | 4400.855036901541 | 4881.430502560873 | 4872.381388461104 | 4156.398706578787 |
| SE_51_6_4 | SE_51 | SE_51_6 | AATCAGGAGATAATTCCAAA | 9.75399157519469 | 14.990887269508443 | 15.800429802152705 | 29.93009526002721 | 15.9089176909411 | 34.65240439473635 | 27.980175931880833 | 8.508181415005666 | 0.8879852778251698 | 48.84346530340181 | 8.216494752885502 | 37.14179240729438 |
| SE_51_7_10 | SE_51 | SE_51_7 | CTTACTGCTACCCCACACGA | 1265.851351091933 | 1363.1713490406344 | 1331.1862108313653 | 1171.1776406097604 | 1289.6829274789586 | 1522.1068630387942 | 1222.5184561006395 | 1416.139528853165 | 1426.9923414650477 | 1318.044556247022 | 1550.5699383659642 | 1481.0289722408634 |
| SE_51_7_20 | SE_51 | SE_51_7 | GCCTCGACTCCAAAGAAAGG | 494.2022398098643 | 740.5498311137171 | 712.9943948221407 | 606.4097561379426 | 764.6886436779023 | 417.561472956573 | 563.9081610886752 | 757.2281459355042 | 691.7405314258073 | 949.8960491094412 | 847.4727445119048 | 381.86405318749536 |
| SE_51_7_3 | SE_51 | SE_51_7 | TTAGCAAAGAAACAATTCCC | 408.5838693164887 | 469.7144677779312 | 479.9380552403884 | 429.43180155691215 | 363.7839178661865 | 562.2352613045973 | 392.79862365909634 | 604.0808804654022 | 430.67285974520735 | 492.80869470297944 | 617.4108914311107 | 708.0154177640492 |
| SE_51_7_8 | SE_51 | SE_51_7 | TCTTACTGCTACCCCACACG | 2087.3541970916635 | 1969.8025872134094 | 2142.9332919169606 | 2015.7268503383543 | 2206.036586477166 | 1905.015931600631 | 1863.9101813083694 | 2000.3679860168875 | 1934.03193510322 | 1934.7844315705731 | 1982.5228053748021 | 1756.1103722573876 |
| SE_51_7_9 | SE_51 | SE_51_7 | TCCACCTCTGTCACTTTCCA | 513.7102229602536 | 452.72479553915497 | 488.8257970040993 | 550.4534910865874 | 370.147484942563 | 422.7593336157835 | 554.2227155737935 | 329.92836820410855 | 635.7974589228215 | 972.4952643990748 | 629.14874107809 | 827.565562075028 |
| SE_51_8_18 | SE_51 | SE_51_8 | GTTTTCTCCAGGTAACCAAG | 4.335107366753196 | 7.9951398770711695 | 4.93763431317272 | 2.602616979132801 | 1.0605945127294067 | 28.588233625657487 | 0.0 | 0.945353490556185 | 0.0 | 1.4580138896537853 | 0.0 | 0.0 |
| SE_51_8_2 | SE_51 | SE_51_8 | TGAGCTTTATCTGACTGCAA | 744.5546902398613 | 749.5443634754222 | 815.6971885361334 | 825.0295823850979 | 736.0525918342083 | 1035.2405812927484 | 729.6368954544309 | 687.2719876343465 | 959.0241000511834 | 578.1025072477258 | 1078.7083825573968 | 1120.0571772824712 |
| SE_51_8_35 | SE_51 | SE_51_8 | GTTCTACTCTTGACCTTGGT | 10.837768416882989 | 11.992709815606755 | 24.688171565863602 | 5.205233958265602 | 4.242378050917627 | 4.331550549342044 | 0.0 | 3.78141396222474 | 10.655823333902038 | 2.187020834480678 | 0.0 | 88.21175696732416 |
| SE_51_8_9 | SE_51 | SE_51_8 | GGAGGGCCGAGTGAGCACAG | 8713.565807173924 | 8173.031739336003 | 8247.824356723711 | 8787.736230041903 | 8743.541162941228 | 8256.801657155804 | 8893.391303887047 | 8787.06069471974 | 8618.785106571098 | 9100.922699218929 | 9471.27088014759 | 8471.810711901302 |
| SE_52_1_17 | SE_52 | SE_52_1 | GGTTTGGTCCTGGTGACCAA | 15.172875783636185 | 51.968409200962604 | 40.488601368016305 | 26.02616979132801 | 55.15091466192915 | 68.4384986796043 | 35.513300221233365 | 86.97252113116902 | 33.74344055735645 | 29.8892847379026 | 63.384388093688166 | 46.42724050911798 |
| SE_52_1_18 | SE_52 | SE_52_1 | TATGGAAAGTGGAAACAGGC | 963.4776122608978 | 1470.1063448964612 | 1512.8911535561215 | 567.3705014509505 | 651.2050308158557 | 268.5561340592067 | 371.2754114038034 | 321.4201867891029 | 723.7080014275134 | 182.25173620672317 | 86.86008738764674 | 133.47831646371418 |
| SE_52_1_38 | SE_52 | SE_52_1 | AGAATCCTGAAATAGGATGA | 16.256652625324485 | 34.97873696218637 | 27.650752153767232 | 20.820935833062407 | 19.090701229129323 | 52.844916701972934 | 10.761606127646475 | 34.03272566002266 | 31.07948472388094 | 1.4580138896537853 | 46.95139858791716 | 1.1606810127279494 |
| SE_52_1_4 | SE_52 | SE_52_1 | TCCACTCCCAAACCCAACCA | 901.7023322846646 | 1230.2521485843263 | 1250.2090080953328 | 421.6239506195137 | 506.9641770846564 | 576.0962230624918 | 447.6828149100933 | 155.03797245121433 | 432.4488303008577 | 118.09912506195661 | 50.47275348201095 | 84.7297139291403 |
| SE_52_1_9 | SE_52 | SE_52_1 | AGTCTTTCAATTACAGCGAT | 29.26197472558407 | 43.97326932389143 | 34.56344019220904 | 16.917010364363207 | 38.181402458258646 | 0.8663101098684087 | 15.066248578705064 | 6.617474433893295 | 56.831057780810866 | 5.103048613788249 | 4.695139858791716 | 3.482043038183848 |
| SE_52_2_10 | SE_52 | SE_52_2 | ATGCAAACGTAGAGACCACT | 1904.1959108463411 | 1689.9726915159185 | 1762.7354498026612 | 1828.3384278407925 | 1759.5262966180858 | 1756.8769028131328 | 1844.5392902786057 | 1415.194175362609 | 1782.1864525951157 | 1554.9718133157621 | 1834.6258998228632 | 2100.8326330375885 |
| SE_52_2_23 | SE_52 | SE_52_2 | TGGCGTTCTCTCTCACCTGA | 44.434850509220254 | 56.96537162413208 | 42.463655093285396 | 45.545797134824014 | 26.51486281823517 | 126.48127604078768 | 33.36097899570407 | 59.557269905039654 | 50.615160836034676 | 8.019076393095819 | 77.46980767006332 | 69.64086076367697 |
| SE_52_2_24 | SE_52 | SE_52_2 | GCAGCAGTCTCCTTGACAGA | 3396.556621851129 | 3602.8099071051956 | 3615.335844105066 | 3668.3886320876827 | 3541.325078003489 | 3165.4971414591655 | 3357.6211118257 | 3531.8406407179073 | 3614.988066026266 | 3545.889779638006 | 3598.8247017638505 | 3819.8012128876817 |
| SE_52_2_7 | SE_52 | SE_52_2 | ATTTACTAACCAATATAGCA | 3.2513305250648967 | 6.9957473924372735 | 4.93763431317272 | 0.0 | 18.030106716399914 | 0.0 | 0.0 | 0.0 | 7.103882222601358 | 0.0 | 1.173784964697929 | 0.0 |
| SE_52_2_9 | SE_52 | SE_52_2 | ATATTATCACCCCCACTCAC | 298.0386314642822 | 305.81410029797223 | 250.8318231091742 | 290.1917931733073 | 322.42073186973965 | 191.45453428091832 | 271.19247441669114 | 211.75918188458544 | 239.75602501279585 | 303.9958959928142 | 309.8792306802533 | 191.51236710011165 |

| | | | | | | | | | | | | | | | |
|---|---|---|---|---|---|---|---|---|---|---|---|---|---|---|---|
| SE_52_3_16 | SE_52 | SE_52_3 | AAGAGCCACGGCCTAAAGCA | 592.8259324034995 | 992.3967372414589 | 864.086004805226 | 420.3226421299473 | 397.7229422735275 | 344.7914237276267 | 280.877919931573 | 190.96140509234937 | 648.2292528123739 | 68.5266528137279 | 37.56111887033373 | 222.85075444376628 |
| SE_52_3_19 | SE_52 | SE_52_3 | AGGCTGATTGATCTTCTATG | 1030.6717764455723 | 1593.0316205064305 | 1535.604271396716 | 831.5361248329299 | 954.5350614564661 | 994.5240061289333 | 714.570646875726 | 469.84068480642395 | 1308.002314236475 | 242.03030568252836 | 545.810008584537 | 292.49161520744326 |
| SE_52_3_28 | SE_52 | SE_52_3 | GCAGCGCAAACCCTTAGCCT | 25300.770369213336 | 25086.750149280062 | 24871.851562313626 | 20779.29396139628 | 20359.172266353693 | 17011.73162748594 | 20023.04436109903 | 15770.386929458278 | 21249.48769835631 | 11174.218450306611 | 13078.312076664326 | 14047.722297046372 |
| SE_52_3_34 | SE_52 | SE_52_3 | GCAACTACACAGGACGGTGT | 2620.5724032023068 | 3536.8500031193585 | 3329.9405808036827 | 1531.6400922196533 | 1460.4386440283931 | 859.3796289894615 | 1233.280062228286 | 392.32169858081676 | 1689.8359837012981 | 250.04938207562418 | 250.0161974806589 | 250.70709874923708 |
| SE_52_3_6 | SE_52 | SE_52_3 | ACTCAGCCCTTGGCCTCCCA | 390.15966300778763 | 520.6834844942599 | 504.626226806252 | 239.44076208021767 | 213.17949705861076 | 77.10159977828837 | 168.95721620404964 | 89.80858160283758 | 246.8599072353972 | 44.46942363444045 | 64.5581730583861 | 11.606810127279495 |
| SE_52_4_10 | SE_52 | SE_52_4 | ATTAACTCCTGGCCATTTGG | 2.167553683376598 | 9.993924846338961 | 6.912688038441808 | 2.602616979132801 | 7.4241615891058474 | 0.0 | 1.0761606127646475 | 1.89070698111237 | 15.095749723027886 | 0.0 | 0.0 | 0.0 |
| SE_52_4_11 | SE_52 | SE_52_4 | GCTGCAAAACAAAGCCAGTG | 2081.935312883222 | 2331.5826666508797 | 2316.73801974064 | 2451.6651943430984 | 2507.2454280923175 | 3264.256493984164 | 2217.9670229079384 | 2776.5032017635153 | 2436.631602352266 | 2289.081806756443 | 3355.851214071379 | 2909.827298908969 |
| SE_52_4_16 | SE_52 | SE_52_4 | ATAGTTAAAACAACTTGTAG | 8.670214733506391 | 14.990887269508443 | 23.700644703229056 | 23.423552812195208 | 45.60556404736449 | 32.919784174999535 | 4.30464245105859 | 29.305958207241734 | 15.095749723027886 | 4.374041668961356 | 23.47569929395858 | 54.55200759821362 |
| SE_52_4_5 | SE_52 | SE_52_4 | TCTCACGTGATTGGATCAGA | 654.6012123797325 | 692.5789918512901 | 733.7324589374662 | 771.6759343128755 | 634.2355186121853 | 614.2138678967018 | 708.113683199138 | 804.4958204633134 | 679.3087375362549 | 763.9992781785835 | 800.5213459239876 | 750.9606152349833 |
| SE_52_4_9 | SE_52 | SE_52_4 | TTTGAGCTCACCCTGATGGG | 75.86437891818092 | 111.93195827899638 | 66.16429979651446 | 91.09159426964803 | 58.33269820011737 | 12.994651648026132 | 101.15909759987686 | 90.75393509339376 | 55.05508722516053 | 142.15635424124406 | 83.33873249355297 | 13.928172152735392 |
| SE_52_5_1 | SE_52 | SE_52_5 | AAACTCTGTTGATTACTAAA | 3.2513305250648967 | 4.996962423169481 | 5.925161175807264 | 0.0 | 0.0 | 0.0 | 5.380803063823238 | 0.0 | 2.6639558334755096 | 0.7290069448268927 | 12.911634611677218 | 0.0 |
| SE_52_5_17 | SE_52 | SE_52_5 | ATTTCAGTGTTAGAAAAGCG | 4.335107366753196 | 12.992102300240651 | 4.93763431317272 | 3.9039254686992013 | 22.272484767317543 | 4.331550549342044 | 3.228481838293942 | 24.57919075446081 | 1.7759705556503396 | 1.4580138896537853 | 0.0 | 0.0 |
| SE_52_5_23 | SE_52 | SE_52_5 | TCGGCCCTGTTAGCATTGGG | 2445.0005548488025 | 2505.4769589771777 | 2417.465759729364 | 2161.473401169791 | 2274.9752298045773 | 2350.299328072993 | 2245.947198839819 | 2209.2911074298045 | 2538.7499093021606 | 1858.2387023637493 | 1832.2783298934671 | 2079.9403748084856 |
| SE_52_5_37 | SE_52 | SE_52_5 | CCAGTGCCCTGACAAATTTG | 568.9828418863569 | 645.6075450734969 | 646.8300950256263 | 631.1346174397042 | 679.8410826595497 | 660.9946138295959 | 498.26236371003176 | 586.1191641448347 | 835.5941464334848 | 555.5032919580922 | 487.12076034964053 | 894.885060813249 |
| SE_52_5_5 | SE_52 | SE_52_5 | AACAGCCTCTGAGAATCCTG | 450.8511661423323 | 692.5789918512901 | 760.395684228599 | 346.1480582246625 | 418.9348325281157 | 508.5240344927559 | 311.0104170889831 | 262.80827037461944 | 828.4902642108834 | 262.44250013768135 | 125.5949912226784 | 283.20616710561967 |
| SE_53_1_27 | SE_53 | SE_53_1 | GACAATTGTGAACAGAGTTA | 502.8724545433707 | 493.6998874091447 | 485.86321641619566 | 556.9600335344194 | 478.32812524096244 | 453.0801874611778 | 586.5075339567329 | 706.1790574454702 | 502.5996672490461 | 612.3658336545898 | 667.8836449131217 | 683.6411164967623 |
| SE_53_1_29 | SE_53 | SE_53_1 | GGTGACTGAACGGAGCTGGG | 492.0346861264877 | 611.6282005959445 | 660.65547110251 | 506.20900244132974 | 760.4462656269847 | 509.39034460262434 | 441.22585123350547 | 710.9058248982511 | 705.0603105931848 | 442.50721550992387 | 823.9970452179462 | 737.0324430822479 |
| SE_53_1_30 | SE_53 | SE_53_1 | ACATGTTTAGCATTCCCTCA | 1317.8726394929715 | 1286.2181277238244 | 1446.726853759607 | 1463.9720507622005 | 1481.6505342829812 | 1187.7111606295884 | 1368.8762994366316 | 1310.2599379108724 | 1475.8315317454321 | 1615.479389736394 | 2270.100121725795 | 1245.4107266570898 |
| SE_53_1_33 | SE_53 | SE_53_1 | GGAAAAGCAAAGCCGCCCTG | 406.4163156331121 | 697.5759542744595 | 723.8571903111208 | 508.8116194204626 | 631.053735073997 | 699.1122586638058 | 426.1596026548004 | 627.7147177293068 | 467.0802561360393 | 1007.4875977507656 | 867.4270889117696 | 812.4767089095646 |
| SE_53_1_9 | SE_53 | SE_53_1 | GTGGCCTGAGTGTCTGGGCA | 156.06386520311503 | 222.86452407335884 | 227.13117840594512 | 175.67664609146405 | 215.30068608406958 | 152.47057933683993 | 136.67239782111022 | 250.518674997389 | 243.3079661240965 | 135.59529173780203 | 251.1899824453568 | 480.52193926937105 |
| SE_53_2_13 | SE_53 | SE_53_2 | ATTCCAGACTCAGGGCCCCG | 5.4188842084414945 | 12.992102300240651 | 16.78795666478725 | 5.205233958265602 | 22.272484767317543 | 1.7326202197368175 | 1.0761606127646475 | 0.945353490556185 | 39.07135222430747 | 0.0 | 0.0 | 19.73157721637514 |
| SE_53_2_14 | SE_53 | SE_53_2 | GAGATTGCCCCTAACAGAAA | 334.88704408168434 | 357.7825094989348 | 400.9359062296249 | 366.9689940577249 | 400.90472581171576 | 373.3796573532842 | 304.55345341239524 | 284.5514006574117 | 382.7216547426482 | 285.7707223721419 | 199.54344399864794 | 442.21946584934875 |
| SE_53_2_17 | SE_53 | SE_53_2 | TCCAGAGTCACAGAGAATGA | 1404.5747868280355 | 1429.1312530264715 | 1439.8141657211652 | 1346.8542867012245 | 1430.7419976719698 | 1565.4223685322145 | 1354.8862114706913 | 1349.9647845142322 | 1869.2090098219824 | 1479.884097998592 | 1395.6303230258377 | 1414.8701545153704 |
| SE_53_2_27 | SE_53 | SE_53_2 | CCTGCAAAGCCACATAGGAA | 723.9629302477837 | 813.5054824919915 | 847.2980481404388 | 812.0164974894338 | 863.3239333617371 | 1073.3582261269585 | 783.4449260926633 | 884.8508671605891 | 988.3276142194139 | 820.8618198750811 | 717.1826134304347 | 665.070220293115 |
| SE_53_2_9 | SE_53 | SE_53_2 | TGAGAGGCTTCAGGTTACCT | 2492.6867358830873 | 2541.455088423998 | 2505.355650503838 | 2602.6169791328007 | 2773.454650787399 | 2636.181664329568 | 2422.4375393332216 | 2469.2633173327554 | 2808.6974337610122 | 2781.8905014594225 | 2993.151659979719 | 2173.9555368394495 |
| SE_53_3_1 | SE_53 | SE_53_3 | AGGGTTTCACTGGTACCAAG | 382.57322511596954 | 532.6761943098667 | 480.9255821030229 | 422.9252591090801 | 421.05602155357445 | 223.50800834604945 | 465.97754532709234 | 296.8409960346421 | 522.1353433611998 | 309.09894460660246 | 502.3799648907136 | 412.04175951842205 |
| SE_53_3_14 | SE_53 | SE_53_3 | CCTTCCTGCTACCTTCTCCC | 1649.508353049591 | 1701.9654013315253 | 1780.510933330083 | 1595.404208208407 | 1990.7359003930965 | 1950.9303674236564 | 1711.0953742957895 | 1991.8598046018817 | 1571.7339417505505 | 1896.1470634947477 | 1862.7967389756134 | 1449.6905848972087 |
| SE_53_3_16 | SE_53 | SE_53_3 | TTTAGTTGAGAAGCTGCAGA | 0.0 | 0.0 | 1.975053725269088 | 0.0 | 0.0 | 0.0 | 0.0 | 0.0 | 17.759705556503395 | 0.0 | 0.0 | 0.0 |
| SE_53_3_17 | SE_53 | SE_53_3 | GCAGAGGGAGTTTATTTCTG | 2785.3064831389283 | 3378.945990547203 | 3425.730686479233 | 2882.398304389577 | 3516.9314042107126 | 2596.331399275621 | 2745.285723162616 | 3589.5072036418346 | 3951.5344863220057 | 3464.970008762221 | 3920.441782091083 | 3461.1507799547453 |
| SE_53_3_4 | SE_53 | SE_53_3 | TCTGGGAACATGGCATAAGA | 862.6863659838859 | 767.5334281988323 | 786.071382657097 | 808.1125720207347 | 719.0830796305378 | 876.7058311868296 | 874.9185781776584 | 771.408448293847 | 868.449601713016 | 572.9994586339376 | 791.1310662064042 | 896.045741825977 |
| SE_53_4_12 | SE_53 | SE_53_4 | TTTACTTAATAAAGGCCTGG | 1.083776841688299 | 1.9987849692677924 | 0.987526862634544 | 0.0 | 0.0 | 0.0 | 0.0 | 0.0 | 0.0 | 0.0 | 0.0 | 0.0 |
| SE_53_4_15 | SE_53 | SE_53_4 | CCCACATCAGAGATGTGAAA | 27.094421042207472 | 37.97691441608806 | 46.41376254382357 | 27.32747828089441 | 68.93864332741144 | 82.29946043749882 | 16.142409191469714 | 27.415251226129364 | 71.03882222601358 | 6.561062503442034 | 17.606774470468935 | 42.945197470934126 |
| SE_53_4_18 | SE_53 | SE_53_4 | CTGATGTGGGAGATGAAGAG | 407.5000924748004 | 428.73937590794145 | 407.8485942680667 | 351.3532921829281 | 375.45045750621 | 450.48125713157253 | 333.60978995704073 | 460.3871499008621 | 394.26546335437536 | 194.64485426878034 | 453.0809963734006 | 367.93588103476 |
| SE_53_4_6 | SE_53 | SE_53_4 | TTAAGATTTGAGCACACTGG | 96.4561389102586 | 77.9526138014439 | 88.87741763710896 | 68.96934994701923 | 90.15053358199957 | 13.86096175789454 | 110.8445431147587 | 76.57363273505099 | 35.51941111300679 | 86.02281948957334 | 193.67451917515828 | 78.92630886550056 |
| SE_53_5_12 | SE_53 | SE_53_5 | GCATGTGTCCCTCTGCTTCA | 8112.069660036917 | 6827.849455018779 | 7525.94222013786 | 7906.750382605449 | 7302.193220141966 | 7610.5343151939705 | 8794.384527512699 | 6158.032637482989 | 7720.144005412026 | 8371.18674744721 | 7812.712725029416 | 7298.362208033346 |
| SE_53_5_13 | SE_53 | SE_53_5 | AAAAAATTGTCAAAGAGCCA | 9.75399157519469 | 5.9963549078033775 | 1.975053725269088 | 13.013084895664004 | 3.1817835381882205 | 0.0 | 5.380803063823238 | 0.0 | 0.0 | 0.0 | 0.0 | 0.0 |
| SE_53_5_3 | SE_53 | SE_53_5 | TGAATAAGTGAATGAACGCA | 819.335292316354 | 722.560766390307 | 788.0464363823661 | 767.7720088441763 | 679.8410826595497 | 933.0159883282762 | 776.9879624160754 | 591.7912850881718 | 735.2518100392406 | 991.449444964574 | 495.33725510252606 | 1161.8416937406773 |
| SE_53_5_5 | SE_53 | SE_53_5 | TGTGGGTGAGATGGCTCAGA | 2084.102866566599 | 2118.71206742386 | 2054.0558742798517 | 2030.0412437235848 | 2106.340702280602 | 1969.9891898407614 | 2199.6722924909395 | 2103.4115164875116 | 1933.1439498253947 | 2013.5171816118775 | 1910.9219225282284 | 2179.758941903089 |
| SE_53_5_7 | SE_53 | SE_53_5 | CAGCCCATAATCCAAAACCA | 1273.4377889837513 | 1306.2059774165023 | 1400.3130912157835 | 1313.020265972498 | 1510.2865861266753 | 1535.9678247966888 | 1222.5184561006395 | 1560.7786129082615 | 1069.1342745015045 | 1418.6475146331331 | 1142.092770651085 | 1485.6716962917753 |
| SE_53_6_10 | SE_53 | SE_53_6 | TCATGATATTTACTGCGTGG | 640.5121134377847 | 769.5322131681 | 796.934178146077 | 681.8856485327939 | 845.2938266453372 | 808.2673325072253 | 640.3155645949653 | 913.2114718772747 | 1000.7594081089663 | 448.339271068539 | 1055.2326832634383 | 1131.6639874097507 |
| SE_53_6_12 | SE_53 | SE_53_6 | CTAAAACCCAGACCACACAG | 664.3552039549272 | 848.4842194541778 | 767.3083722670407 | 724.828828688485 | 797.5670735725139 | 748.4919349263051 | 683.3619891055512 | 745.88390404883 | 843.5860139339113 | 901.0525838060394 | 684.3166344188926 | 708.0154177640492 |
| SE_53_6_15 | SE_53 | SE_53_6 | CAGGAAGCCATGAGTCCAGG | 2761.4633926217857 | 2780.309892251499 | 2647.5595187232125 | 2632.547074392828 | 2470.1246201467884 | 2901.272557949301 | 2961.5940063283097 | 2960.8471324219713 | 2512.9983362452303 | 3388.424279555397 | 2486.0765552302137 | 2811.1694128270933 |
| SE_53_6_20 | SE_53 | SE_53_6 | TCATGAGCCCCACATGCAGC | 3308.7706976743766 | 3532.852433180823 | 3509.6704698031695 | 3372.9916049561098 | 3705.717227476547 | 2754.86614938154 | 3171.4453258174162 | 3654.7365944902112 | 3882.271634651642 | 3860.8207798032236 | 3598.8247017638505 | 3115.267838161816 |
| SE_53_6_23 | SE_53 | SE_53_6 | ATCCAGTCTGAGTCCCCTGT | 171.23674098675122 | 281.8286806667587 | 241.94408134546327 | 266.7682403611121 | 286.3605184369398 | 290.2138868059169 | 196.93739213593048 | 311.96665188354103 | 198.02071695501286 | 466.5644446892113 | 201.8910139280438 | 155.53125570554522 |
| SE_54_1_11 | SE_54 | SE_54_1 | GCATGAAGAGGAAGCCAATC | 12802.655830863874 | 12080.656354254537 | 12126.8298731522 | 12846.517408999505 | 11543.510676546863 | 12506.919056170216 | 13840.501640766131 | 12827.501513356874 | 12191.149879261757 | 18124.570662286205 | 12577.105896738309 | 15244.384421168888 |
| SE_54_1_12 | SE_54 | SE_54_1 | GAAGAGGAAGCCAATCAGGC | 1956.2171992473795 | 2157.6883743245817 | 2293.037375037411 | 2154.966858721959 | 2274.9752298045773 | 2252.4062856578626 | 2231.9571108738787 | 2595.940685067284 | 2248.3787234533297 | 2999.1345710178366 | 3047.1457683558237 | 2576.7118482560477 |
| SE_54_1_6 | SE_54 | SE_54_1 | CGTCTACATTTACACAGCTG | 3086.5964451282753 | 4036.546245436307 | 4107.124221697069 | 3638.4585368276557 | 3944.350992840664 | 3810.89817331113 | 3118.7134557919485 | 4356.188884482901 | 3946.2065746550547 | 4731.98407887136 | 5388.846772928192 | 5152.263015499368 |
| SE_54_1_7 | SE_54 | SE_54_1 | CTCAGCTGTGTAAATGTAGA | 798.7435323242763 | 868.4720691468558 | 824.5849302998442 | 834.1387418120627 | 940.7473327909838 | 894.8983434940662 | 809.2727807990149 | 983.1676301784324 | 840.9220581004358 | 1175.8882020057779 | 1298.2061709559096 | 1015.5958861369558 |
| SE_54_1_9 | SE_54 | SE_54_1 | TGGCTTCCTCTTCATGCTGA | 22.759313675454276 | 13.991494784874547 | 11.850322351614528 | 28.62878677046081 | 13.787728665482287 | 6.93048087894727 | 4.30464245105859 | 18.9070698111237 | 7.991867500426528 | 0.7290069448268927 | 8.216494752885502 | 6.964086076367696 |
| SE_54_2_11 | SE_54 | SE_54_2 | GGCCTGAGAGAGCAAGAGAA | 230.84446727960767 | 287.8250355745621 | 303.170746828805 | 255.0564639550145 | 301.2088416151515 | 398.502650539468 | 297.0203291230427 | 518.9990663153455 | 331.21850862878836 | 321.4920626686597 | 516.4653844670888 | 243.74301267286938 |
| SE_54_2_12 | SE_54 | SE_54_2 | TTACCCTCATTCTTTTAAGT | 20.59175999207768 | 23.98541963121351 | 21.725590977959968 | 14.314393385230405 | 21.211890254588134 | 0.8663101098684087 | 5.380803063823238 | 7.56282792444948 | 10.655823333902038 | 13.851131951710961 | 44.6038286585213 | 0.0 |
| SE_54_2_14 | SE_54 | SE_54_2 | ACACCCTATTCAGTGTCAGC | 464.9402650842802 | 412.74909615379914 | 418.7113897570467 | 465.8684392647714 | 353.17797273889244 | 370.78072702367893 | 520.8617365780893 | 496.3105825419971 | 335.6584350179142 | 408.97289604788676 | 583.3711274548707 | 393.4708633147749 |
| SE_54_2_5 | SE_54 | SE_54_2 | TTTCTGCTATTTACAAGCAA | 7.586437891818092 | 4.996962423169481 | 1.975053725269088 | 1.3013084895664004 | 7.4241615891058474 | 0.0 | 15.066248578705064 | 0.0 | 15.095749723027886 | 0.7290069448268927 | 0.0 | 8.124767089095647 |
| SE_54_2_7 | SE_54 | SE_54_2 | GATGTGGAACACAAAAGGAA | 666.5227576383038 | 749.5443634754222 | 759.4081573659644 | 796.4007956146371 | 688.325838761385 | 688.716537345385 | 736.0938591310189 | 542.6329035792502 | 831.1542200443589 | 916.3617296474041 | 671.4049998072154 | 808.9946658713808 |
| SE_54_3_1 | SE_54 | SE_54_3 | GACAAAAGCAGACAGATCAA | 1780.645350893875 | 2053.7515559226567 | 2100.469636823675 | 1841.3515127364567 | 2088.310595564202 | 2046.2244795091815 | 2027.486594448596 | 1912.4501113951624 | 2073.4456237217714 | 2192.852890039293 | 1782.9793613761542 | 2286.5415950740603 |
| SE_54_3_12 | SE_54 | SE_54_3 | ATTGAAGTATAACATACACG | 156.06386520311503 | 213.8699917116538 | 188.61763076319792 | 147.04785932100324 | 161.21036593486983 | 210.51335669802333 | 111.92070372752333 | 166.38221433788857 | 157.17339417505505 | 150.17543063433988 | 122.07363632858463 | 388.8281392638631 |
| SE_54_3_14 | SE_54 | SE_54_3 | TTTTTTGAGAGGGCAAGGGG | 1.083776841688299 | 2.9981774539016888 | 0.987526862634544 | 3.9039254686992013 | 12.727134152752882 | 6.93048087894727 | 7.533124289352532 | 1.89070698111237 | 0.0 | 34.992333351690846 | 1.173784964697929 | 1.1606810127279494 |
| SE_54_3_16 | SE_54 | SE_54_3 | TCTTTTTTGAGAGGGCAAGG | 11.921545258571289 | 20.98724217731182 | 15.800429802152705 | 16.917010364363207 | 8.484756101835254 | 0.0 | 13.990087965940416 | 17.01636283001133 | 14.207764445202717 | 18.225173620672315 | 43.43004369382337 | 1.1606810127279494 |
| SE_54_3_8 | SE_54 | SE_54_3 | ATACGTCGTGTTGAGACAGC | 2428.743902223478 | 2087.730900400209 | 2217.985333477186 | 1987.0980635678934 | 1776.4958088217563 | 2329.507885436151 | 2095.2847130527684 | 1904.8872834707129 | 2109.853020112603 | 1970.5057718670907 | 1558.7864331188498 | 1640.0422709845925 |
| SE_54_4_10 | SE_54 | SE_54_4 | CTGAGGTGAGTTTTCATGCA | 997.074694353235 | 1161.2940671445874 | 1100.1049249748821 | 1050.155951080085 | 1303.470656144441 | 1146.994585465773 | 1026.6572245774737 | 1138.2056026296468 | 1219.2037864539582 | 1368.3460354400775 | 1235.9955678269193 | 1041.1308684169705 |
| SE_55_1_1 | SE_55 | SE_55_1 | TCACCGTTGCCTATCTGCCA | 3908.099291128006 | 3401.9320176937827 | 3342.7784300179314 | 3681.401716983347 | 3279.358233359326 | 3687.882137709816 | 3668.6315289146833 | 3688.769320150234 | 3628.3078451936435 | 3534.225668520776 | 3695.0750688690805 | 3916.1377369441016 |
| SE_55_1_19 | SE_55 | SE_55_1 | GTGCTGGTGGAATAAACCCG | 476.8618103428515 | 663.596609796907 | 594.4911713059955 | 524.4273212952594 | 456.0556404736449 | 436.620295373678 | 506.871648612149 | 414.064828863609 | 511.4795200272978 | 807.0106879233701 | 393.21796317380625 | 410.8810785056941 |
| SE_55_1_21 | SE_55 | SE_55_1 | TCGCCTCATTAGAGAAGCAA | 621.0041302873952 | 627.6184803500868 | 628.06708463557 | 602.5058306692434 | 695.7500003504908 | 518.9197558111769 | 577.8982490546157 | 610.6983548992955 | 553.2148280850807 | 354.29737518586984 | 771.1767218065394 | 515.3423696512095 |
| SE_55_1_23 | SE_55 | SE_55_1 | AGCTCTCCAGACATCTTAGG | 32.51330525064897 | 36.977521931454156 | 41.47612823065085 | 28.62878677046081 | 36.06021343279983 | 48.51336615263089 | 34.43713960846872 | 51.04908849003399 | 15.983735000853056 | 44.46942363444045 | 113.85714157569912 | 15.088853165463343 |
| SE_55_1_28 | SE_55 | SE_55_1 | GCATTTTTACTCAAGAGGTG | 49.85373471766175 | 78.9520062860778 | 64.18924607124536 | 46.847105624390416 | 68.93864332741144 | 77.10159977828837 | 74.25508228076067 | 171.10898179066947 | 103.89427750554486 | 53.94651391719005 | 201.8910139280438 | 49.909283547301825 |
| SE_55_2_21 | SE_55 | SE_55_2 | AGGAGCGCCGTGGTGATGGG | 8304.981937857434 | 8400.893225832531 | 8392.99080553099 | 7083.022108709918 | 7723.24924169554 | 6409.828502916356 | 7343.720021505955 | 6047.426279087916 | 6915.629343702422 | 5651.261836298072 | 5890.052952854208 | 5722.157392748791 |
| SE_55_2_25 | SE_55 | SE_55_2 | ATATGCAAAGAGCCCCAAGA | 74.78060207649263 | 106.9349958558269 | 138.25376076883617 | 87.18766880094883 | 153.786204345764 | 183.65774329210265 | 44.12258512335055 | 121.00524679119168 | 190.02884945458632 | 99.1449444964574 | 32.86597901154201 | 333.1154506529215 |
| SE_55_2_26 | SE_55 | SE_55_2 | GATATGCAAAGAGCCCCAAG | 87.78592417675222 | 162.90097499532507 | 158.00429802152703 | 106.70729614444484 | 151.66501532030517 | 147.27271867762948 | 108.6922218892294 | 143.69373056454012 | 218.44437834499178 | 158.19450702743572 | 153.7658303754287 | 131.15695443825828 |
| SE_55_2_3 | SE_55 | SE_55_2 | AACATGCTAGTGCTAGGCCT | 1455.5122983873855 | 1302.2084074779668 | 1247.2464275074292 | 1324.7320423785957 | 1283.319360402582 | 1150.4598259052468 | 1324.753714313281 | 1675.16638526556 | 1190.7882575635526 | 884.2854240750208 | 1503.618539778047 | 1600.5791165518422 |
| SE_55_2_40 | SE_55 | SE_55_2 | GCCACAGTGGAGCAGCCGCA | 5486.078372626169 | 5011.953310438989 | 5116.376675309572 | 5365.294902482269 | 5193.731328835905 | 5896.10660776439 | 5488.419125099702 | 5606.891552488733 | 4814.65617636807 | 5058.579190153808 | 7149.5242199750855 | 5649.03448894693 |
| SE_55_3_18 | SE_55 | SE_55_3 | AGTGTGGGATGCATTTTAGG | 1121.7090311473894 | 1162.2934596292212 | 1288.72255573808 | 1366.3739140447206 | 1231.3502292788412 | 960.7379118440653 | 1156.872658721996 | 1528.6365942293512 | 1252.0592417334894 | 1330.437674309079 | 1939.0927616809788 | 987.739541831485 |
| SE_55_3_33 | SE_55 | SE_55_3 | AAGGGTTGTGTGGATTACGT | 2830.825110489837 | 2705.355455903957 | 2605.095863629927 | 2269.482005803802 | 2473.3064036849764 | 2277.5292788440465 | 2437.5037879119263 | 2229.143530731484 | 2334.5132954023716 | 1782.4219801017525 | 1608.0854016361627 | 3042.1449343599556 |
| SE_55_3_41 | SE_55 | SE_55_3 | CATACACAGAGAGGAAACGG | 2649.8343779278907 | 3018.1653035943664 | 2924.067040260885 | 2386.5997698647784 | 2573.0022878815407 | 1939.6683359953672 | 2066.2283765081233 | 1753.6307249817232 | 2497.0146012443774 | 1658.4907994811808 | 1710.2046935648825 | 2025.3883672102718 |
| SE_55_3_5 | SE_55 | SE_55_3 | GACTTTGTAGGGACTTGCAA | 59.60772629285644 | 96.94107100948793 | 102.70279371399258 | 67.66804145745283 | 103.93826224748186 | 84.03208065723565 | 38.74178205952731 | 201.36029348846742 | 184.70093778763533 | 37.17935418617152 | 356.8306292681704 | 99.81856709460365 |
| SE_55_3_6 | SE_55 | SE_55_3 | GCATAAGGACTGAGTAGAGC | 3201.476790347235 | 3332.9739362540436 | 3268.7139153203407 | 3172.5900975628842 | 3204.0560229555376 | 3149.037249371666 | 2848.597141988022 | 3103.5955094959554 | 2631.100378195978 | 3182.8443211142135 | 2736.0927527108724 | 3235.978663485523 |
| SE_55_4_17 | SE_55 | SE_55_4 | GCAGTTGTAGACAGATGGGC | 2025.5789171154306 | 2318.590564350639 | 2353.2765136581183 | 2074.285732368842 | 2301.4900926228124 | 2298.3207214808886 | 2124.341049597414 | 1873.6906182823586 | 2524.5421448569577 | 2692.2226472457146 | 2334.6582947841807 | 2401.4490153341276 |
| SE_55_4_26 | SE_55 | SE_55_4 | AGAATCAGCTTGATTTCAGA | 193.9960546622055 | 208.8730292884843 | 198.49289938954334 | 179.58057156016326 | 193.02820131675202 | 146.40640856776108 | 186.175786008284 | 204.19635396013595 | 206.90056973326455 | 139.96933340676338 | 296.96759606857603 | 75.44426582731671 |
| SE_55_4_33 | SE_55 | SE_55_4 | CGAGAATTCAGGTACCAAGA | 2229.3289633528307 | 2538.4569109700965 | 2442.1539312952273 | 2355.368366115185 | 2510.4272116305056 | 2917.7324500368004 | 2042.552843027301 | 2303.826456485423 | 2479.254895687874 | 2100.2690080462776 | 2416.8232423130357 | 2234.3109495013027 |
| SE_55_4_4 | SE_55 | SE_55_4 | CTACTAATAACTCAAAAGAG | 16.256652625324485 | 32.97995199291857 | 30.613332741670867 | 32.53271223916001 | 27.575457330964575 | 90.96256153618292 | 17.21856980423436 | 3.78141396222474 | 21.311646667804077 | 67.06863892407412 | 1.173784964697929 | 111.42537722188314 |
| SE_55_4_5 | SE_55 | SE_55_4 | GCACTGGAAGTGGCTCTCAG | 731.5493681396017 | 784.5231004376085 | 915.4374016622223 | 753.4576154589458 | 784.839939419761 | 641.9357914124909 | 693.047434620433 | 925.5010672545051 | 651.7811939236747 | 853.6671323922913 | 815.7805504650606 | 495.6107924348344 |
| SE_55_5_11 | SE_55 | SE_55_5 | GAGCCTGGTATCCACAAGAG | 254.68755779675024 | 344.7904071986942 | 313.04601545515044 | 269.37085734024487 | 331.9660824843043 | 467.8074593289407 | 323.9243444421589 | 328.0376612229962 | 392.48949279872505 | 316.38901405487144 | 277.01325166871123 | 297.13433925835506 |

| | | | | | | | | | | | | | | | |
|---|---|---|---|---|---|---|---|---|---|---|---|---|---|---|---|
| SE_55_5_13 | SE_55 | SE_55_5 | TTAACTACAAATGTACTCAG | 28.17819788389577 | 67.95868895510495 | 49.376343131727204 | 29.93009526002721 | 31.8178353818822 | 72.77004922894633 | 30.13249715741013 | 61.44797688615203 | 48.83919028038434 | 25.515243068941242 | 10.564064682281362 | 45.26655949639003 |
| SE_55_5_18 | SE_55 | SE_55_5 | GGGAATGGATGAACAAGCTG | 272.027987263763 | 321.8043800521146 | 317.9836497683232 | 251.15253848631528 | 344.6932166370572 | 164.59892087499765 | 313.1627383145124 | 407.44735442971574 | 332.1064939066135 | 271.91959042043095 | 471.86155580856746 | 293.6522962201712 |
| SE_55_5_27 | SE_55 | SE_55_5 | CTGCTACCCGGGCCTCCCGA | 8.670214733506391 | 44.97266180852533 | 53.32645058226538 | 20.820935833062407 | 48.78734758555271 | 96.16042219539337 | 3.228481838293942 | 20.79777679223607 | 11.543808611727208 | 12.393118062057175 | 8.216494752885502 | 66.15881772549312 |
| SE_55_5_28 | SE_55 | SE_55_5 | CAGTGGAAAGGAGACAAGCA | 3421.4834892099598 | 3166.075391320183 | 3110.7096172988136 | 3694.414801879011 | 3345.115093148549 | 3176.759172887455 | 3943.052485169668 | 3769.1243668475095 | 3313.9610568435337 | 2978.722376562683 | 3968.566965643698 | 3494.810529323856 |
| SE_56_1_1 | SE_56 | SE_56_1 | GATAGTAAACTCAGACCCAT | 772.7328881237571 | 812.5060900073576 | 771.2584797175789 | 835.4400503016291 | 906.8083083836427 | 897.4972738236714 | 955.630624135007 | 921.7196532922803 | 912.8488656042746 | 1199.9454311850652 | 902.6406378527074 | 696.4086076367697 |
| SE_56_1_12 | SE_56 | SE_56_1 | AGAGGATGCATTTTGCAAAG | 838.8432754667433 | 1133.3110775748382 | 1168.2442784966656 | 1076.1821208714132 | 1269.5316317370998 | 1393.8929667782697 | 965.3160696498887 | 1199.6535795157988 | 1549.5343098049213 | 1132.8767922609911 | 1305.248880744097 | 1572.7227722463715 |
| SE_56_1_20 | SE_56 | SE_56_1 | GCCCTATTTGCCAAGCCACA | 351.14369670700887 | 344.7904071986942 | 383.1604227022031 | 316.2179629646353 | 356.3597562770807 | 234.77003977433876 | 316.39122015280634 | 292.11422858186114 | 226.43624584541828 | 204.12194455152994 | 282.8821764922087 | 344.722260780201 |
| SE_56_1_23 | SE_56 | SE_56_1 | TTTGGTCTCCACCTAAAGTG | 53.105065242726646 | 26.983597085115196 | 15.800429802152705 | 40.34056317655841 | 28.63605184369398 | 1.7326202197368175 | 46.27490634887984 | 26.46989773557318 | 35.51941111300679 | 78.73275004130441 | 61.03681816429231 | 1.1606810127279494 |
| SE_56_1_24 | SE_56 | SE_56_1 | GCTGTTGGAGTGCTAGCTGA | 1632.1679235825782 | 1835.8839942724674 | 1819.0244809728301 | 1552.4610280527156 | 1971.6451991639672 | 2005.5079043453661 | 1429.141293751452 | 1751.740018000611 | 1879.8648331558845 | 1265.5560562194858 | 1732.5066078941431 | 2254.042526717678 |
| SE_56_2_0 | SE_56 | SE_56_2 | GATTGCACTTGTTCAGGGGA | 325.13305250648966 | 313.8092401750434 | 308.10838114197776 | 347.4493667142289 | 312.875381255175 | 161.133680435524 | 406.78871162503674 | 317.63877282687815 | 292.1471564044809 | 540.1941461167274 | 353.30927437407667 | 185.70896203647192 |
| SE_56_2_1 | SE_56 | SE_56_2 | GAATAAACTATCATATGTAA | 62.859056817921335 | 81.9501837399795 | 110.60300861506893 | 117.11776406097604 | 45.60556404736449 | 49.379676262499295 | 89.32133085946575 | 91.69928858394995 | 140.30167389637683 | 52.48850002753627 | 61.03681816429231 | 131.15695443825828 |
| SE_56_2_5 | SE_56 | SE_56_2 | TCATCTTTGACAAACCACAG | 164.73407993662144 | 257.8432610355452 | 228.11870526857967 | 173.07402911233126 | 257.72446659324584 | 150.73795911710312 | 165.7287343657557 | 349.78079150578844 | 248.63587779104753 | 150.90443757916677 | 220.67157336321065 | 302.9377443219948 |
| SE_56_2_7 | SE_56 | SE_56_2 | GGCTTTCCCTTTGAAAAGAG | 3.2513305250648967 | 2.9981774539016888 | 5.925161175807264 | 3.9039254686992013 | 8.484756101835254 | 36.38502461447317 | 4.30464245105859 | 9.45353490556185 | 0.8879852778251698 | 0.7290069448268927 | 7.042709788187574 | 134.63899747644214 |
| SE_56_3_1 | SE_56 | SE_56_3 | TCAATTTCCTCAATAAGTTG | 124.63433679415438 | 120.92649064070145 | 167.87956664787248 | 140.54131687317124 | 117.72599091296415 | 61.50801780065702 | 131.291594757287 | 112.49706537618601 | 123.4299536176986 | 40.0953819654791 | 82.16494752885504 | 61.51609367458132 |
| SE_56_3_11 | SE_56 | SE_56_3 | GGAAGTGGCATGCAGAGTGG | 2695.353005278799 | 3088.122777518739 | 3013.9319847606284 | 2702.8177328294137 | 3002.5430655369505 | 2678.6308597131197 | 2669.9544802690903 | 2822.8255228007683 | 2845.992815429669 | 2189.2078553151587 | 2673.8821495818825 | 2835.5437140943804 |
| SE_56_3_13 | SE_56 | SE_56_3 | TCTTTCCCTGTAGCTCCTAT | 115.96412206064798 | 84.94836119388117 | 78.01462214812898 | 104.10467916531204 | 117.72599091296415 | 45.91443582302566 | 87.16900963393644 | 97.37140952728706 | 166.94123223113192 | 80.91977087578509 | 64.5581730583861 | 77.76562785277261 |
| SE_56_3_2 | SE_56 | SE_56_3 | GTAAACACCTCAACTTATTG | 76.94815575986922 | 70.95686640900662 | 62.21419234597627 | 72.87327541571842 | 72.12042686559965 | 25.12299318618385 | 72.10276105523138 | 45.37696754669688 | 148.29354139680336 | 21.141201399979888 | 76.29602270536539 | 20.89225822910309 |
| SE_56_3_7 | SE_56 | SE_56_3 | GCAAGCCAAATAATGTGGAT | 1033.9231069706373 | 999.3924846338962 | 1002.3397655740622 | 1055.3611850383509 | 1031.9584608857128 | 1351.4437713947177 | 1160.1011405602899 | 1285.6807471564116 | 994.5435111641901 | 989.9914310749202 | 1288.815891238326 | 1203.6262101988837 |
| SE_56_4_17 | SE_56 | SE_56_4 | AATGGGGGGAAAAAAACCCG | 126.80189047753097 | 239.8541963121351 | 258.73203801025056 | 157.45832723753446 | 278.93635684783396 | 248.6310015322333 | 193.70891029763655 | 201.36029348846742 | 247.74789251322238 | 465.10643079955753 | 198.36965903395 | 232.1362025455899 |
| SE_56_4_4 | SE_56 | SE_56_4 | ACTATCATATGTAAGGGCAG | 1361.2237131605034 | 1631.0085349225185 | 1642.2571725612468 | 1310.4176489933652 | 1555.8921501740397 | 1637.3261076512924 | 1416.227366398276 | 1759.3028459250602 | 1518.4548250810403 | 1369.8040493297312 | 1851.058889328634 | 1288.3559241280238 |
| SE_56_4_5 | SE_56 | SE_56_4 | TCATCACAGCCTCCACAAAA | 6418.126456478106 | 7715.309981373679 | 7735.297915016384 | 6824.061719286204 | 8174.001909605538 | 8125.988830565674 | 6228.81762668178 | 8100.734060575949 | 8859.429116861718 | 7464.302108082554 | 8320.961614743619 | 7796.294362493636 |
| SE_56_4_6 | SE_56 | SE_56_4 | GCTGATGTCATTGAGAAAGG | 1198.6571869072586 | 1345.1822843172242 | 1279.8348139743691 | 1251.8587669628773 | 1179.3810981551003 | 1166.0534078828782 | 1292.4688959303417 | 1091.8832815923936 | 1091.3339064471336 | 1223.2736534195258 | 1192.565524133096 | 1416.0308355280984 |
| SE_56_4_9 | SE_56 | SE_56_4 | GGTTGAAATTGAAATCTGGA | 31.429528408960667 | 78.9520062860778 | 95.79010567555078 | 83.28374333224963 | 64.69626527649382 | 67.57218856973589 | 29.056336544645482 | 16.071009339455145 | 40.84732277995781 | 49.5724722482287 | 62.210603128990236 | 249.54641773650914 |
| SE_57_1_10 | SE_57 | SE_57_1 | CACATTGCCCCCGACTACCG | 947.2209596355732 | 1192.275234168238 | 1222.5582559415654 | 1160.7671726932292 | 1126.35137251863 | 1273.475861506561 | 1056.7897217348839 | 948.1895510278536 | 1253.8352122891397 | 1570.2809591571267 | 881.5125084881447 | 1525.1348507245254 |
| SE_57_1_11 | SE_57 | SE_57_1 | ACATTGCCCCCGACTACCGT | 6999.030843623034 | 6548.0195593212875 | 6544.340518679123 | 7064.803789855988 | 6447.354042882063 | 6819.593184884114 | 7239.332442067784 | 6400.043131065372 | 6849.030447865534 | 7657.488948461681 | 5934.656781512729 | 7791.651638442724 |
| SE_57_1_35 | SE_57 | SE_57_1 | GGTTGTAGTAGTTGTGGCGA | 4328.6047057030655 | 5074.915036970925 | 5165.7530184413 | 4463.488119212753 | 4891.461892708024 | 4871.261747790062 | 4423.020118462701 | 4297.576968068417 | 4859.943425537154 | 4714.487912195515 | 4913.463862225531 | 4504.603010397172 |
| SE_57_1_37 | SE_57 | SE_57_1 | GGTGGTGGTGGCGGCAGCAG | 134.38832836934907 | 221.86513158872495 | 210.3432217411579 | 84.58505182181602 | 95.45350614564661 | 57.176467251314975 | 53.808030638232374 | 21.743130282792254 | 142.96562972985234 | 24.78623612411435 | 9.390279717583432 | 10.446129114551544 |
| SE_57_1_7 | SE_57 | SE_57_1 | TTCGTGTATCTTAGTCCCGG | 92.12103154350541 | 102.9374259172913 | 103.69032057662713 | 93.6942112487808 | 146.36204275665813 | 95.29411208552496 | 109.76838250199404 | 69.95615830115769 | 118.10204195074758 | 115.18309728264904 | 63.38438809368166 | 68.48017975094902 |
| SE_57_2_22 | SE_57 | SE_57_2 | CCTGGGATCTGCAATGTGTT | 2680.1801294951633 | 2617.408917256174 | 2624.846400882618 | 2619.533989497164 | 2379.9740865647886 | 3104.8554337683768 | 2759.275811128556 | 2537.3287686528006 | 2598.2449229164467 | 2283.978758142655 | 2264.2311969023053 | 2750.81400016524 |
| SE_57_2_23 | SE_57 | SE_57_2 | CCCTGGGATCTGCAATGTGT | 725.046707089472 | 732.5546912366459 | 667.5681591409518 | 733.9379881154498 | 654.386814354044 | 748.4919349263051 | 789.9018897692513 | 566.2667408431548 | 689.0765755923318 | 599.9727155925326 | 723.0515382539243 | 690.6052025731299 |
| SE_57_2_24 | SE_57 | SE_57_2 | ACTTCAGTGGGCGCAGCCCT | 30.34575156727237 | 51.968409200962604 | 59.25161175807264 | 33.834020728726415 | 94.3929116329172 | 73.63635933881474 | 33.36097899570407 | 125.7320142439726 | 48.83919028038434 | 5.832055558615141 | 105.64064682281361 | 35.9811139456643 |
| SE_57_2_27 | SE_57 | SE_57_2 | GACTCGGACTTTAACTTCAG | 246.01734306324386 | 313.8092401750434 | 320.9462303562268 | 257.6590809341473 | 221.664253160446 | 297.1443676848642 | 180.79498294446077 | 243.90120056349573 | 331.21850862878836 | 550.4002433443039 | 350.9617044446808 | 272.76003799106815 |
| SE_57_3_1 | SE_57 | SE_57_3 | AAGACGGAGAGAACTGTAGA | 522.3804376937601 | 806.5097350995542 | 765.3333185417716 | 612.9162985857746 | 709.5377290159731 | 407.16575163815213 | 629.5539584673188 | 789.3701646144145 | 652.6691792014998 | 1187.5523131230082 | 775.871861665331 | 530.4312228166729 |
| SE_57_3_11 | SE_57 | SE_57_3 | AAAATCGCATGGCTAGGAAG | 74.78060207649263 | 128.9216305177726 | 108.62795488979984 | 83.28374333224963 | 115.60480188750533 | 140.34223779868222 | 114.07302495305264 | 105.87959094229272 | 133.19779167377547 | 128.3052222895331 | 208.93372371623136 | 135.79967848917008 |
| SE_57_3_29 | SE_57 | SE_57_3 | TGGTTGCACGAGAATTTGCC | 0.0 | 0.0 | 0.987526862634544 | 0.0 | 0.0 | 0.0 | 0.0 | 0.0 | 0.0 | 6.561062503442034 | 0.0 | 0.0 |
| SE_57_3_4 | SE_57 | SE_57_3 | GGGTCCCCACTCAGTGAAAA | 8.670214733506391 | 7.9951398770711695 | 13.825376076883616 | 16.917010364363207 | 6.363567076376441 | 34.65240439473635 | 16.142409191469714 | 17.961716320567515 | 24.863587779104755 | 19.683187510326103 | 11.73784964697929 | 13.928172152735392 |
| SE_57_3_6 | SE_57 | SE_57_3 | TTGGGCAACTCATTGCACTG | 1438.1718689203726 | 1404.146440910624 | 1722.2468484346448 | 1405.4131687317124 | 1321.5007628608407 | 1456.2672946887951 | 1539.9858368662105 | 1441.6640730981821 | 1644.5487345322144 | 960.1021463370176 | 1199.6082339212835 | 1399.781301349907 |
| SE_57_4_29 | SE_57 | SE_57_4 | GCTTCTCCACAGACCCCACC | 8.670214733506391 | 7.9951398770711695 | 0.987526862634544 | 7.807850937398403 | 4.242378050917627 | 0.8663101098684087 | 22.599372868057596 | 0.945353490556185 | 0.8879852778251698 | 9.477090282749604 | 0.0 | 0.0 |
| SE_57_4_36 | SE_57 | SE_57_4 | CAGGATCTACCGGATGCCCG | 714.2089386725889 | 722.560766390307 | 708.0567605089681 | 572.5757354092162 | 799.6882625979727 | 596.8876656993336 | 659.6864556247289 | 812.0586483877629 | 648.2292528123739 | 914.9037157577503 | 863.9057340176757 | 853.1005443550429 |
| SE_57_4_44 | SE_57 | SE_57_4 | GGTGGCCAGAAGGGGCACCA | 139.80721257779055 | 185.8870021419047 | 185.6550501752943 | 115.81645557140963 | 186.6646342403756 | 105.68983340394587 | 100.08293698711222 | 52.93979547114636 | 205.12459917761421 | 25.515243068941242 | 28.170839152750297 | 42.945197470934126 |
| SE_57_4_47 | SE_57 | SE_57_4 | CCGACACTCCAGCTGCACCG | 5068.824288576174 | 4439.301416743767 | 4618.6631365417625 | 5311.941254410047 | 5121.610901970305 | 4608.769784499935 | 5394.7931517891775 | 5674.011650318223 | 4891.91089553886 | 5458.804002863772 | 6391.259132780224 | 5856.796390225233 |
| SE_57_4_7 | SE_57 | SE_57_4 | AACCTGTTTCCAATATAGAG | 1.083776841688299 | 3.9975699385355847 | 7.900214901076352 | 5.205233958265602 | 15.9089176909411 | 0.0 | 1.0761606127646475 | 0.0 | 5.327911666951019 | 0.0 | 0.0 | 4.642724050911798 |
| SE_57_5_3 | SE_57 | SE_57_5 | CAGGACGCAGGTGGGCTCTG | 13420.408630626205 | 11725.872022209503 | 11975.738263169116 | 13177.049765349371 | 11502.147490550416 | 10786.427177971556 | 13500.434887132502 | 11526.695110351564 | 12010.000882585422 | 11306.168707320277 | 11808.276744861167 | 10592.374922155266 |
| SE_57_5_4 | SE_57 | SE_57_5 | CTCTGGGGAAGCCGTCACAG | 69.36171786805113 | 60.962941562667666 | 74.0645146975908 | 37.737946197425615 | 46.6661585600939 | 58.04277736118338 | 102.23525821264151 | 97.37140952728706 | 14.207764445202717 | 53.94651391719005 | 183.11045449287693 | 92.85448101823596 |
| SE_57_5_5 | SE_57 | SE_57_5 | TGCTGGTGACTCCTCTGTGA | 1186.7356416486873 | 1130.3129001209365 | 1224.5333096668346 | 1151.6580132662643 | 1387.257622650064 | 1212.8341538157722 | 1071.8559703135888 | 1170.347621308557 | 1001.6473933867915 | 1151.8309728264903 | 1105.705436745449 | 1318.5336304589505 |
| SE_57_5_6 | SE_57 | SE_57_5 | TCACCAGCACCCCCGAATGA | 4035.984958447225 | 4106.5037193606795 | 4257.2283048175195 | 3931.2529469800957 | 4361.16463634332 | 4320.288517913755 | 3802.0754448974994 | 3812.6106274130943 | 5126.339008884705 | 4971.098356774581 | 4330.09273477066 | 4551.030250906289 |
| SE_57_5_7 | SE_57 | SE_57_5 | CTACCTTCATTCGGGGGTGC | 450.8511661423323 | 508.69077467865316 | 547.0898818995374 | 510.112927910029 | 418.9348325281157 | 501.59355361380864 | 398.1794267229196 | 350.72614499634466 | 389.8255369652495 | 770.5603406820255 | 507.0751047495053 | 689.444521560402 |
| SE_57_6_17 | SE_57 | SE_57_6 | ACTCGGACTTTAACTTCAGT | 1025.2528922371307 | 1107.326872974357 | 1024.0653565520222 | 979.8852926434995 | 1075.4428359076185 | 1234.4919065624824 | 1007.28633354771 | 992.6211650839942 | 848.0259403230372 | 1000.9265352473236 | 1118.6170713571264 | 808.9946658713808 |
| SE_57_6_20 | SE_57 | SE_57_6 | GCACAGCCATCAGCCTTCAG | 688.1982944720698 | 878.4659939931947 | 817.6722422614025 | 756.0602324380786 | 877.1116620272194 | 1024.8448599743276 | 679.0573466544926 | 830.9657181988866 | 847.1379550452119 | 956.4571116128832 | 693.706914136476 | 955.2404734751024 |
| SE_57_6_24 | SE_57 | SE_57_6 | GGACCGTGTCGGAGTAGACG | 2045.08690026582 | 2177.6762240172598 | 2058.00598173039 | 1899.910394766946 | 2036.3414644404609 | 2144.1175219243114 | 1975.8308850358928 | 1957.827078941859 | 2136.4925784473585 | 2495.3907721424534 | 2389.8261881249837 | 2233.1502684885745 |
| SE_57_6_26 | SE_57 | SE_57_6 | GTCTACTCCGACACGGTCCA | 167.98541046168634 | 167.89793741849456 | 206.3931142906197 | 137.93869989403845 | 159.089176909411 | 109.1550738434195 | 116.22534617858193 | 204.19635396013595 | 139.41368861855165 | 86.75182643440023 | 260.58026216294024 | 114.907420260067 |
| SE_57_6_27 | SE_57 | SE_57_6 | GAGCTGGCCCTGGACCGTGT | 3585.1337923048927 | 3356.959355885257 | 3293.4020868862044 | 3505.7250708918828 | 3029.0579283551856 | 3427.9891047492933 | 3796.694641833676 | 2685.7492666701214 | 3422.2952607382044 | 4090.4579674236948 | 2319.399090243108 | 4203.9866281006325 |
| SE_58_1_13 | SE_58 | SE_58_1 | AAATACTTACTTGACAGAGA | 2.167553683376598 | 6.9957473924372735 | 4.93763431317272 | 7.807850937398403 | 10.605945127294067 | 0.8663101098684087 | 0.0 | 0.0 | 0.0 | 0.7290069448268927 | 0.0 | 0.0 |
| SE_58_1_15 | SE_58 | SE_58_1 | AGAGAGTGAGAGCGCTCCTG | 979.7342648862223 | 1157.2964972060518 | 1087.267075760633 | 1057.9638020174837 | 1178.3205036423708 | 819.5293639355147 | 1081.5414158284707 | 1130.6427747051973 | 1066.470318668029 | 873.3503199026175 | 1286.4683213089302 | 899.5277848641608 |
| SE_58_1_19 | SE_58 | SE_58_1 | CTCACTCTCTCGGATAAGAA | 91.0372547018171 | 90.94471610168455 | 78.01462214812898 | 75.47589239485123 | 71.05983235287025 | 88.3636312065777 | 118.37766740411122 | 63.33868386726439 | 132.3098063959503 | 139.2403264619365 | 51.646538446708874 | 62.67677468730927 |
| SE_58_1_23 | SE_58 | SE_58_1 | AAGAAATGCTTAGCAAACAG | 220.00669886272468 | 229.86027146579613 | 234.04386644438694 | 313.61534598550253 | 199.39176839312847 | 246.0320712026281 | 266.8878319656326 | 294.95028905352973 | 232.6521427901945 | 194.64485426878034 | 203.06479889274172 | 307.5804683729066 |
| SE_58_1_27 | SE_58 | SE_58_1 | TGGCAAAATCCCCAGAGCCA | 105.126353643765 | 130.9204154870404 | 112.57806234033802 | 102.80337067574564 | 144.2408537311993 | 86.63101098684088 | 130.21543414452233 | 109.66100490451745 | 101.23032167206935 | 272.64859736525784 | 72.7746678112716 | 10.446129114551544 |
| SE_58_2_10 | SE_58 | SE_58_2 | CAGACACAGGACAAGACCTT | 6154.768683947849 | 5835.45271777732 | 5497.5620442865065 | 5723.154737113029 | 5819.4820913462545 | 5449.090591072291 | 6055.555768026671 | 6058.77052097459 | 5154.75453777511 | 5525.14363484302 | 5181.086834176659 | 6129.556428216301 |
| SE_58_2_13 | SE_58 | SE_58_2 | GGCAGACAAATGCCTCACAG | 948.3047364772616 | 1097.332948128018 | 1021.1027759641186 | 843.2479012390274 | 1263.1680646607235 | 725.9678720697265 | 969.6207121009473 | 809.2225879160943 | 1137.5091408940425 | 1165.6821047782014 | 933.1590469348536 | 879.7962076477856 |
| SE_58_2_26 | SE_58 | SE_58_2 | TGAGAGGCCAAGGTGACTTG | 5132.767122235784 | 4769.100936672952 | 4689.76507065145 | 4973.601047122783 | 4121.470276466474 | 4556.7911779078295 | 5067.640325508725 | 4592.527257121947 | 3965.742250767208 | 4743.64818998859 | 4602.410846580579 | 5235.83204841578 |
| SE_58_2_36 | SE_58 | SE_58_2 | ACAGCTTTCAGGAGCGGAGG | 741.3033597147964 | 869.4714616314897 | 836.4352526514588 | 775.5798597815747 | 953.4744669437366 | 781.4117191013047 | 750.0839470969593 | 657.0206759365486 | 695.292472537108 | 662.6673128476455 | 933.1590469348536 | 681.3197544713063 |
| SE_58_2_4 | SE_58 | SE_58_2 | AGGTTACACTGGCAATTTAA | 606.9150313454473 | 794.5170252839474 | 804.8343930471534 | 687.0908824910595 | 825.1425309034785 | 787.4758898703835 | 698.4282376842563 | 844.2006670666732 | 647.3412675345488 | 974.6822852335555 | 845.1251745825089 | 708.0154177640492 |
| SE_58_3_13 | SE_58 | SE_58_3 | GACACAGTACAAGAAGGGAA | 1580.1466351815398 | 1948.8153450360976 | 1939.5027582142445 | 1437.9458809708724 | 1515.5895586903223 | 1554.1603371039253 | 1489.406288066272 | 1543.7622500782502 | 1757.322864816011 | 1306.3804451297917 | 1615.1281114243504 | 1686.4695114937106 |
| SE_58_3_14 | SE_58 | SE_58_3 | AAAGGATTCTCTGTATGCCC | 88.86970101844051 | 167.89793741849456 | 158.9918248841616 | 87.18766880094883 | 42.42378050917627 | 8.663101098684088 | 38.74178205952731 | 68.06545132004533 | 143.85361500767752 | 22.599215289633673 | 46.95139858791716 | 11.606810127279495 |
| SE_58_3_2 | SE_58 | SE_58_3 | TTGTGGTTTCAGCCTTTTTC | 107.2939073271416 | 111.93195827899638 | 97.76515940081985 | 102.80337067574564 | 82.72637199289373 | 100.4919727447354 | 131.291594757287 | 73.73757226338243 | 94.126439449468 | 315.66000711004455 | 132.63770101086598 | 63.83745570003722 |
| SE_58_3_5 | SE_58 | SE_58_3 | TCAACTTTCCTAAAATTCAC | 197.2473851872704 | 247.84933618920624 | 252.80687683444327 | 234.2355281219521 | 194.08879582948143 | 446.1497065822305 | 151.73864639981528 | 411.2287683919405 | 364.0739639083196 | 269.7325695859503 | 296.96759606857603 | 312.2231924238184 |
| SE_58_3_7 | SE_58 | SE_58_3 | TGGACTTTCTCTAAGAAGTG | 60.69150313454474 | 90.94471610168455 | 57.27655803280356 | 40.34056317655841 | 50.90853661101153 | 0.8663101098684087 | 87.16900963393644 | 68.06545132004533 | 42.62329333560815 | 108.622034779207 | 69.25331291717781 | 24.37430126728694 |
| SE_58_4_0 | SE_58 | SE_58_4 | CTCTGCGGATAGCATTTGGT | 220.00669886272468 | 237.85541134286728 | 246.881715658636 | 275.8773997880769 | 249.2397104914106 | 199.251325269734 | 301.3249715741013 | 270.37109829906893 | 325.89059696183733 | 322.2210696134866 | 316.9219404684408 | 275.081400016524 |
| SE_58_4_1 | SE_58 | SE_58_4 | TTAACTCTGCGGATAGCATT | 102.95879996038839 | 122.92527560996923 | 101.71526685135804 | 110.61122161314404 | 63.6356707637644 | 155.06950966644516 | 89.32133085946575 | 115.33312584785457 | 99.45435111641902 | 26.973256958595027 | 165.503680022408 | 52.23064557275772 |
| SE_58_4_3 | SE_58 | SE_58_4 | TACTTAATCCTTTAACTCTG | 147.39365046960864 | 169.89672238776234 | 184.66752331265974 | 156.15701874796804 | 220.6036586477166 | 31.187163955262715 | 149.586325174286 | 220.2673632995911 | 216.66840778934142 | 83.83579865509266 | 431.95286700883787 | 83.56903291641235 |
| SE_58_4_5 | SE_58 | SE_58_4 | ACACAAAACTTAGTTCAGCA | 2118.7837255006243 | 2127.706599785565 | 2202.184903675033 | 2313.72649444906 | 2320.580793851942 | 2208.224470054574 | 2592.470916150036 | 2582.7057361994976 | 2123.172799279981 | 2581.413591632027 | 3170.3931896491063 | 1942.9800153065873 |
| SE_58_4_6 | SE_58 | SE_58_4 | TCCAAAATAAAACAATCAAA | 94.288585226882 | 135.9173779102099 | 163.9294591973343 | 136.63739140447206 | 165.45274398578746 | 273.7539947184172 | 119.45382801687587 | 223.10342377125966 | 158.94936473070538 | 207.7669792756644 | 144.37555065784528 | 190.3516860873837 |
| SE_59_1_10 | SE_59 | SE_59_1 | TGGGCTATCCCACTGAAGAG | 92.12103154350541 | 117.92831318679976 | 79.00214901076352 | 57.25757354092162 | 139.9984756802817 | 163.73261076512924 | 71.02660044246673 | 185.28928414901227 | 175.8210850093836 | 45.92743752409424 | 145.5493356225432 | 235.61824558377373 |
| SE_59_1_6 | SE_59 | SE_59_1 | AAGCACCTTAGACATGCACG | 1496.6958183715408 | 1417.1385432108648 | 1411.1758867047633 | 1353.3608291490564 | 1273.7740097880176 | 1670.245891826292 | 1382.866387402572 | 1589.139217624947 | 1298.2344761803981 | 737.0260212199885 | 1119.7908563218243 | 1109.6110481679198 |
| SE_59_1_7 | SE_59 | SE_59_1 | TTCTTCCTCGTGCATGTCTA | 2.167553683376598 | 3.9975699385355847 | 4.93763431317272 | 3.9039254686992013 | 4.242378050917627 | 39.8502650539468 | 1.0761606127646475 | 6.617474433893295 | 24.863587779104755 | 2.187020834480678 | 5.868924823489645 | 25.534982280014887 |
| SE_59_1_8 | SE_59 | SE_59_1 | AAATATACACCTCTCTTCAG | 35.76463577571386 | 42.973876839257535 | 31.60085960430541 | 32.53271223916001 | 41.363185996446866 | 31.187163955262715 | 32.28481838293943 | 32.14201867891029 | 30.191499446055772 | 13.851131951710961 | 63.38438809368166 | 4.642724050911798 |
| SE_59_1_9 | SE_59 | SE_59_1 | AATATACACCTCTCTTCAGT | 113.79656837727138 | 82.94957622461338 | 137.26623390620162 | 106.70729614444484 | 123.02896347661118 | 100.4919727447354 | 132.36775537005164 | 104.93423745173654 | 201.57265806631355 | 293.7897987652377 | 197.19587406925208 | 155.53125570554522 |
| SE_59_2_0 | SE_59 | SE_59_2 | TCACGCAGACTTTAAATCAG | 6974.103976264203 | 7348.532939513038 | 7572.355982681684 | 7076.5155662620855 | 7439.009912284059 | 7737.015591234758 | 6431.135821881533 | 6964.419164927415 | 8038.0427348734365 | 7120.210830124261 | 6597.845286567059 | 7879.863395410049 |
| SE_59_2_1 | SE_59 | SE_59_2 | AGTTTTGCTATAAAGACAAA | 2.167553683376598 | 5.9963549078033775 | 14.81290293951816 | 2.602616979132801 | 24.393673792776355 | 0.0 | 6.456963676587884 | 22.68848377334844 | 9.767838056076867 | 3.6450347241344634 | 5.868924823489645 | 1.1606810127279494 |
| SE_59_2_11 | SE_59 | SE_59_2 | AAGAAAATATGACTGATAGG | 31.429528408960667 | 56.96537162413208 | 66.16429979651446 | 53.35364807222242 | 45.60556404736449 | 28.588233625657487 | 38.74178205952731 | 75.6282792444948 | 40.84732277995781 | 62.69459725511277 | 26.997054188052367 | 66.15881772549312 |
| SE_59_2_17 | SE_59 | SE_59_2 | TATCCAGTTGAGAATCAGAG | 80.19948628493412 | 129.9210230024065 | 141.2163413567398 | 102.80337067574564 | 123.02896347661118 | 28.588233625657487 | 66.72195799140815 | 55.775855942814914 | 118.10204195074758 | 69.98466670338169 | 112.68335661100119 | 105.6219721582434 |

| | | | | | | | | | | | | | | | |
|---|---|---|---|---|---|---|---|---|---|---|---|---|---|---|---|
| SE_59_2_8 | SE_59 | SE_59_2 | ATTGCAGGAACAGTCAAACG | 877.8592417675221 | 950.4222528868353 | 976.6640671455641 | 1056.6624935279171 | 986.3528968383483 | 1113.2084911809052 | 998.6770486455929 | 926.4464207450613 | 1104.6536856145112 | 825.9648684888693 | 1059.92782312223 | 658.1061342167474 |
| SE_6_1_23 | SE_6 | SE_6_1 | AGCAGTACCTGGTGACCGCC | 1204.0760711157002 | 1017.3815493573063 | 1082.3294414474603 | 1201.1077358697876 | 1174.0781255914533 | 976.3314938216967 | 1199.919083232582 | 1035.1620721590225 | 1356.8415045168595 | 1471.8650216054962 | 1505.966109707443 | 1732.8967520028284 |
| SE_6_1_24 | SE_6 | SE_6_1 | GCAGTACCTGGTGACCGCCG | 2719.196095795942 | 2403.53892554452 | 2507.330704229107 | 2650.765393246758 | 2715.121952587281 | 2851.8928816868015 | 2792.63679012426 | 2858.7489554419035 | 2703.915170977642 | 2065.2766746945867 | 2987.2827351562296 | 2403.770377359583 |
| SE_6_1_25 | SE_6 | SE_6_1 | CAGTACCTGGTGACCGCCGG | 980.8180417279106 | 905.4495910783099 | 1006.2898730246004 | 1000.7062284765619 | 812.4153967507256 | 884.5026221756453 | 995.4485668072989 | 916.9928858394994 | 1101.1017445032105 | 813.5717504268122 | 1165.5684699450435 | 1034.166782340603 |
| SE_6_1_26 | SE_6 | SE_6_1 | TACCTGGTGACCGCCGGGGG | 188.577170453764 | 282.82807315139263 | 288.35784388928687 | 281.0826337463425 | 255.60327756778702 | 207.0481162585497 | 202.31819519975372 | 396.1031125430415 | 339.21037612921486 | 406.7858752134061 | 365.04712402105594 | 178.74487596010422 |
| SE_6_1_27 | SE_6 | SE_6_1 | ACCTGGTGACCGCCGGGGGT | 612.3339155538889 | 950.4222528868353 | 912.4748210743187 | 688.3921909806259 | 826.2031254162079 | 1101.946459752616 | 525.166379029148 | 823.4028902744371 | 1001.6473933867915 | 511.76287526847864 | 616.2371064664128 | 871.67144055869 |
| SE_6_2_1 | SE_6 | SE_6_2 | AATCTTGGAGCTAACCACAG | 36.84841261740216 | 32.97995199291857 | 63.20171920861082 | 27.32747828089441 | 28.63605184369398 | 60.641707690788614 | 35.513300221233365 | 14.180302358342775 | 9.767838056076867 | 115.18309728264904 | 12.911634611677218 | 100.9792481073316 |
| SE_6_2_14 | SE_6 | SE_6_2 | CCAGATGCAGAAAGGCACAC | 2972.799876751004 | 2392.5456082135474 | 2352.288986795484 | 2717.132126214644 | 2610.12309582707 | 2598.0640194953576 | 2970.203291230427 | 2724.508759782925 | 2104.525108445652 | 2818.340848700767 | 3217.3445882370233 | 2633.585217879717 |
| SE_6_2_18 | SE_6 | SE_6_2 | GCAGTCCCTTGGAGAGAAGG | 671.9416418467454 | 766.5340357141984 | 738.670093250639 | 718.322286240653 | 846.3544211580665 | 676.5881958072272 | 630.6301190800834 | 726.0314807471501 | 606.493944754591 | 515.4079099926131 | 435.4742219029317 | 855.4219063804987 |
| SE_6_2_19 | SE_6 | SE_6_2 | ATGAGCCTCCTTCTCTCCAA | 1162.8925511315447 | 1190.2764491989703 | 1169.2318053593 | 1267.474468837674 | 1092.412348111289 | 1118.4063518401156 | 1270.9456836750487 | 1249.7573145152767 | 1237.8514772882868 | 1589.235139722626 | 1173.7849646979291 | 1722.450622888277 |
| SE_6_2_3 | SE_6 | SE_6_2 | GTTTCTTGCCCAAAGTCACA | 6.5026610501297935 | 15.990279754142339 | 5.925161175807264 | 15.615701874796805 | 0.0 | 23.390372966447035 | 13.990087965940416 | 5.67212094333711 | 20.423661389978903 | 0.7290069448268927 | 0.0 | 34.820430381838484 |
| SE_6_3_1 | SE_6 | SE_6_3 | TCACATTCGACAGTAACAAG | 1147.7196753479086 | 999.3924846338962 | 1087.267075760633 | 1090.4965142566436 | 1005.4435980674776 | 1131.401003488142 | 1048.1804368327666 | 1134.424188667422 | 985.6636583859384 | 1136.5218269851257 | 1349.8527094026183 | 766.0494684004466 |
| SE_6_3_2 | SE_6 | SE_6_3 | ATAGAGCAGTGAAAAGTCAA | 742.3871365564847 | 945.4252904636658 | 975.6765402829295 | 862.7675285825235 | 898.3235522818076 | 1246.62024810064 | 795.2826928330745 | 941.5720765939602 | 695.292472537108 | 1431.0406326951902 | 1225.4315031446379 | 1750.3069671937478 |
| SE_6_3_5 | SE_6 | SE_6_3 | TATGTAGTCAGGGTAGACAA | 922.2940922767424 | 1029.374259172913 | 1078.379333996922 | 991.5970690495972 | 1056.3521346784892 | 852.4491481105142 | 956.7067847477716 | 1196.8175190441302 | 1122.4133911710146 | 1544.7657160881856 | 1199.6082339212835 | 956.4011544878304 |
| SE_6_3_6 | SE_6 | SE_6_3 | ATGTAGTCAGGGTAGACAAG | 696.8685092055762 | 725.5589438442087 | 658.6804173772408 | 732.6366796258834 | 656.5080033795027 | 599.4865960289388 | 864.156972050012 | 807.331880934982 | 671.3168700358284 | 1338.456750702175 | 1017.6715643931045 | 829.8869241004838 |
| SE_6_3_8 | SE_6 | SE_6_3 | CTCCTATTGAGATCTCCCTA | 37.93218945909046 | 8.994532361705065 | 18.763010390056337 | 32.53271223916001 | 29.69664635642339 | 92.69518175591973 | 21.52321225529295 | 34.03272566002266 | 3.551941111300679 | 5.103048613788249 | 14.085419576375148 | 51.069964560029774 |
| SE_6_4_15 | SE_6 | SE_6_4 | GCCGGCCCCCCACCTCCCCC | 37.93218945909046 | 30.98116702365078 | 50.36386999436174 | 54.65495656178882 | 38.181402458258646 | 54.57753692170975 | 37.665621446762664 | 40.650200093915956 | 31.07948472388094 | 62.69459725511277 | 50.47275348201095 | 94.01516203096391 |
| SE_6_4_24 | SE_6 | SE_6_4 | GAAACCAGCCTGGCTCCGGG | 277.4468714722045 | 490.701709955243 | 499.6885924930793 | 396.89908931775216 | 462.4192075500213 | 315.33687999210076 | 285.1825623826316 | 518.9990663153455 | 404.9212866882774 | 659.7512850683379 | 550.5051484433287 | 495.6107924348344 |
| SE_6_4_29 | SE_6 | SE_6_4 | GGGCATTTCCGAAACCAGCC | 485.5320250763579 | 651.6038999813003 | 606.34149365761 | 598.6019052005443 | 566.3574697975032 | 590.8234949302547 | 558.527358024852 | 512.3815918814523 | 752.1235303179188 | 624.7589511764171 | 879.1649385587489 | 620.964341809453 |
| SE_6_4_30 | SE_6 | SE_6_4 | GGAAATGCCCTTACAGCAGC | 1722.1214014427069 | 1941.8195976436602 | 1842.7251256760592 | 1707.3167383111174 | 2091.4923791023903 | 1947.465126984183 | 1760.5987624829634 | 2236.706358655934 | 1600.149470640956 | 2233.677278949599 | 2199.673023843919 | 1766.556501371939 |
| SE_6_4_6 | SE_6 | SE_6_4 | GGCTGGAGAGGGGGCCGGA | 354.39502723207374 | 456.7223654776906 | 409.8236479933358 | 370.87291952642414 | 474.0857471900448 | 300.60960812433785 | 349.75219914851044 | 354.5075589585694 | 589.6222244759127 | 370.33552797206147 | 368.56847891514974 | 544.3593949694083 |
| SE_60_1_12 | SE_60 | SE_60_1 | TCTTCAGTCAGGCCTCCTGA | 748.8897976066146 | 951.4216453714691 | 986.5393357719095 | 748.2523815006803 | 871.8086894635724 | 877.5721412966981 | 589.7360157950268 | 967.0966208389773 | 1054.0385247784766 | 907.6136463094814 | 934.3328318995515 | 1209.4296152625234 |
| SE_60_1_25 | SE_60 | SE_60_1 | CAAAGGTGTTCTACACCCCG | 279.6144251555811 | 281.8286806667587 | 286.38279016401776 | 363.0650685890257 | 289.54230197512805 | 329.1978417499953 | 283.0302411571023 | 331.8190751852209 | 274.38745084797745 | 185.16776398603074 | 286.4035313862947 | 518.8244126893934 |
| SE_60_1_27 | SE_60 | SE_60_1 | CCTGGAACAGTGCTGTTGCG | 7542.0030413088725 | 6063.314204273848 | 6442.625251827765 | 6874.812750379294 | 6591.594896613263 | 6020.85526358544 | 8230.476366424024 | 6514.430903422671 | 6643.90584868792 | 6918.275906407212 | 5994.519814712324 | 5743.049650977894 |
| SE_60_1_29 | SE_60 | SE_60_1 | ACTGCACTGCCGATTCCCCC | 5840.473399858242 | 5297.779561044284 | 5220.0669958862 | 5930.062786954087 | 5865.08765539362 | 5502.801817884132 | 6116.896922954256 | 6271.475056349731 | 5353.663240007949 | 6222.074274097529 | 6422.9513268270675 | 6156.252091509044 |
| SE_60_1_4 | SE_60 | SE_60_1 | GCATACGCGGAGCTACGGGT | 2037.500462374002 | 2516.4702763081505 | 2498.4429624653962 | 2132.84461439933 | 2608.0019068016113 | 2231.614843021021 | 2072.685340184711 | 2233.8702981842653 | 2079.661520666548 | 2375.833633190843 | 2494.293049983099 | 2378.235395079568 |
| SE_60_2_11 | SE_60 | SE_60_2 | CAGAGTCTCAGTCAGCCCAT | 1533.544230988943 | 1577.041340752288 | 1648.1823337370354 | 1549.8584110735828 | 1453.0144824392873 | 1111.4758709611683 | 1478.6446819386256 | 1427.4837707398394 | 1426.9923414650477 | 1782.4219801017525 | 1602.2164768126731 | 1540.2237038899889 |
| SE_60_2_12 | SE_60 | SE_60_2 | GTCAGCCCATCGGCTCAACG | 772.7328881237571 | 738.5510461444493 | 653.7427830640681 | 649.3529362956338 | 698.9317888886791 | 760.6202764644629 | 737.1700197437585 | 508.8155641938223 | 776.3923962406407 | 626.8011711486942 | 552.4841620585039 |
| SE_60_2_13 | SE_60 | SE_60_2 | TCAGCCCATCGGCTCAACGA | 1324.3753005431013 | 1294.2132676008955 | 1241.321266331622 | 1357.2647546177557 | 1383.0152445991464 | 1011.8502083263014 | 1119.2070372752335 | 1213.8338818741415 | 1324.8740345151534 | 1385.113195171096 | 1086.9248773102822 | 821.7621570113882 |
| SE_60_2_8 | SE_60 | SE_60_2 | TTATGTTCTTAAACACAGGA | 19.50798315038938 | 19.987849692677923 | 33.5759133295745 | 18.218318853929606 | 6.363567076376441 | 12.128341538157722 | 4.30464245105859 | 7.56282792444948 | 7.103882222601358 | 1.4580138896537853 | 38.73490383503166 | 24.37430126728694 |
| SE_60_2_9 | SE_60 | SE_60_2 | ACTGAGACTCTGGAACCCAC | 2177.3076749517927 | 2104.7205726389852 | 2215.0227528892824 | 2451.6651943430984 | 2120.128430946084 | 2313.0479933486513 | 2351.4109388907546 | 2352.9843017940346 | 2398.4482354057836 | 3726.6835019550754 | 2417.9970272777337 | 2471.0898760978043 |
| SE_60_3_12 | SE_60 | SE_60_3 | TCCCCCATATTGAACCCATA | 35.76463577571386 | 34.97873696218637 | 26.66322529113269 | 26.02616979132801 | 31.8178353818822 | 18.192512307236584 | 19.370891029763655 | 54.83050245225873 | 34.63142583518162 | 21.87020834480678 | 22.30191432926065 | 3.482043038183848 |
| SE_60_3_15 | SE_60 | SE_60_3 | ATACCGTATGGGTTCAATAT | 140.89098941947887 | 164.89975996459287 | 190.592684488467 | 114.51514708184324 | 163.33155496032865 | 219.17645797767074 | 152.81480701257993 | 155.98332594177052 | 205.12459917761421 | 53.94651391179005 | 235.93077790428373 | 176.42351393464833 |
| SE_60_3_17 | SE_60 | SE_60_3 | ACAAATTACCTTAAGCTTTG | 124.63433679415438 | 147.91008772581662 | 142.20386821937433 | 115.81645557140963 | 168.63452752397566 | 92.69518175591973 | 117.30150679134657 | 117.22383282896693 | 126.09390945117411 | 122.47316673091797 | 136.15905590495976 | 142.76376456553777 |
| SE_60_3_2 | SE_60 | SE_60_3 | GTGCGGACCCAAAGAGTGAG | 632.9256755459666 | 656.6008624044698 | 669.5432128662209 | 646.750319314501 | 826.2031254162079 | 673.1229553677535 | 504.7193273866197 | 714.6872388604759 | 689.9645608701569 | 788.0565073578709 | 964.8512409816976 | 1030.684739302419 |
| SE_60_3_3 | SE_60 | SE_60_3 | CAATCTTCAGTAGACATACA | 149.56120415298525 | 193.88214201897586 | 194.54279193900518 | 130.13084895664005 | 222.72484767317542 | 140.34223779868222 | 164.65257375299106 | 142.74837707398393 | 261.067617605999 | 293.7897987652377 | 217.15021846911688 | 208.9225822910309 |
| SE_60_4_1 | SE_60 | SE_60_4 | TAAAAGTAGAATAAACACAA | 48.769957875973454 | 80.95079125534559 | 67.151826659149 | 75.47589239485123 | 171.8163110621639 | 39.8502650539468 | 48.42722757440914 | 92.64464207450614 | 89.68651306034215 | 89.6678542137078 | 69.25331291717781 | 215.88666836739858 |
| SE_60_4_2 | SE_60 | SE_60_4 | GCACGTAATAGAATTGATTT | 6964.349984689009 | 8038.113753910427 | 7814.300064027147 | 7793.536544013172 | 8658.693601922876 | 8877.079695821585 | 7228.570835940137 | 8199.99617708435 | 8581.489724902442 | 7818.599483268424 | 8367.913013331536 | 9367.85645372728 |
| SE_60_4_3 | SE_60 | SE_60_4 | AATAATACTAGTCTTATAAT | 4.335107366753196 | 11.992709815606755 | 7.900214901076352 | 6.506542447832002 | 10.605945127294067 | 0.0 | 3.228481838293942 | 0.0 | 12.431793889552377 | 16.038152786191638 | 0.0 | 1.1606810127279494 |
| SE_60_4_4 | SE_60 | SE_60_4 | TATAAGACTAGTATTATTCT | 13.005322100259587 | 18.98845720804403 | 9.87526862634544 | 20.820935833062407 | 5.3029725636470335 | 6.064170769078861 | 31.208657770174778 | 1.89070698111237 | 2.6639558334755096 | 1.4580138896537853 | 0.0 | 26.695663292742836 |
| SE_60_4_5 | SE_60 | SE_60_4 | AGAAACAAGCAGCCTCCTAG | 67.19416418467453 | 120.92649064070145 | 115.54064292824165 | 96.29682822791364 | 160.14977142214042 | 32.05347406513112 | 57.03651247652632 | 291.168875091305 | 103.0062922277197 | 53.21750697236316 | 206.5861537868355 | 74.28358481458876 |
| SE_60_5_1 | SE_60 | SE_60_5 | GCACTAATTTCTCTAAGCCT | 409.667646158177 | 579.6476410876597 | 542.1522475863646 | 508.8116194204626 | 566.3574697975032 | 736.3635933881474 | 517.6332547397955 | 485.9116941458791 | 458.2004033577876 | 487.7056460891912 | 633.8438809368816 | 409.72039749296613 |
| SE_60_5_10 | SE_60 | SE_60_5 | TTCTACATATAAGGTGACAT | 3.2513305250648967 | 9.993924846338961 | 7.900214901076352 | 0.0 | 3.181783581882205 | 0.0 | 0.0 | 0.0 | 0.8879852778251698 | 0.0 | 0.0 | 0.0 |
| SE_60_5_12 | SE_60 | SE_60_5 | GACCTTATATGTAGAATCCA | 120.29922942740117 | 202.87667438068092 | 161.95440547206522 | 128.82954046707366 | 193.02820131675202 | 80.566840217762 | 137.74855843387488 | 96.42605603673087 | 159.83735000853056 | 46.65644446892113 | 173.7201747752935 | 68.48017975094902 |
| SE_60_5_4 | SE_60 | SE_60_5 | GGCTTAGAGAAATTAGTGCG | 140.89098941947887 | 207.87363680385042 | 208.36816801588878 | 154.85571025840164 | 251.3608995168594 | 189.7219140611815 | 117.30150679134657 | 298.7317030157545 | 160.72533528635574 | 127.57621534470621 | 242.9734876924713 | 257.67118482560477 |
| SE_60_5_8 | SE_60 | SE_60_5 | ACAAGTATTGATTAGTACCG | 295.8710777809056 | 238.8548038275012 | 238.98150075755964 | 212.11328379932328 | 304.3906251533397 | 315.33687999210076 | 220.61292561675273 | 310.07594490242866 | 192.69280528806183 | 185.89677093085763 | 203.06479889274172 | 306.41978736017865 |
| SE_60_6_13 | SE_60 | SE_60_6 | GAAGGTGTCCAAACGCAGAA | 1831.582862453225 | 2352.5699088281917 | 2262.4240422957405 | 1871.281607996484 | 2124.370808997002 | 1786.3314465486587 | 1837.0061659892533 | 2487.225033653323 | 2414.4319704066365 | 2082.0438344256054 | 2315.877735349014 | 1750.3069671937478 |
| SE_60_6_17 | SE_60 | SE_60_6 | GTTCTTCTGCATTCCTGCAA | 9.75399157519469 | 23.98541963121351 | 12.837849214249072 | 19.519627343496005 | 23.33307928004695 | 0.0 | 22.599372868057596 | 42.54090707502823 | 0.0 | 3.6450347241344634 | 18.780559435166865 | 0.0 |
| SE_60_6_18 | SE_60 | SE_60_6 | CACTTATTAGACAAACAAGT | 120.29922942740117 | 179.89064723410132 | 151.09160998308525 | 165.26617817493286 | 194.08879568948143 | 170.66309164407653 | 167.88105569128502 | 175.8357492434504 | 209.56452556674006 | 265.3585279169889 | 111.50957164630326 | 302.9377443219948 |
| SE_60_6_2 | SE_60 | SE_60_6 | AGATATATGAGCAGAAACAG | 163.65030309493312 | 246.84994370457235 | 271.56988772244996 | 175.67664609146405 | 251.3608995168594 | 375.112277573021 | 208.7751588763416 | 162.60080037566382 | 285.0432741818795 | 236.19825012391323 | 210.1075068609293 | 511.8603266130257 |
| SE_60_6_3 | SE_60 | SE_60_6 | TATTCCTGCTGTATAACAGA | 44.434850509220254 | 55.96597913949819 | 76.03956842285989 | 44.24448864525762 | 95.45350614564661 | 116.08555472236677 | 51.655709412703075 | 102.09817698006798 | 37.29538166865713 | 51.759493082709376 | 65.73195802308402 | 127.67491140007444 |
| SE_60_7_1 | SE_60 | SE_60_7 | AATAGACCACGAATTCCACA | 63.94283365960963 | 118.92770567143364 | 122.45330966683447 | 102.80337067574564 | 75.30221040378788 | 75.36897955855156 | 111.92070372752333 | 154.09261896065814 | 106.55823333902038 | 231.82420845495187 | 99.77172199932397 | 170.62010887100857 |
| SE_60_7_11 | SE_60 | SE_60_7 | AGACATGTCGTGTAGAAACG | 857.2674817754445 | 1054.3590712887606 | 1020.115249101484 | 953.8591228521715 | 984.2317078128895 | 686.1176070157798 | 820.0343869266613 | 934.954602160067 | 1155.2688464505459 | 783.6824656889096 | 597.4565470312459 | 834.5296481513957 |
| SE_60_7_4 | SE_60 | SE_60_7 | GATAGAATATTACCGCTAGG | 48.769957875973454 | 80.95079125534559 | 91.8399982250126 | 94.99551973834723 | 77.42339942924669 | 229.5721791151283 | 66.72195799140815 | 115.33312584785457 | 69.26285167036325 | 89.6678542137078 | 51.646538446708874 | 85.89039494186825 |
| SE_60_7_5 | SE_60 | SE_60_7 | ATAGCTAGTCAGCCCGCGCG | 591.7421555618112 | 681.5856745203172 | 605.3539667949755 | 684.4882655119267 | 622.5688792121617 | 525.8502366901241 | 581.1267308929097 | 435.8079591464013 | 533.679151972927 | 647.3581670062807 | 393.21796317380625 | 659.2668152294752 |
| SE_60_7_7 | SE_60 | SE_60_7 | AGTTTCATCTCTGCCACGCG | 62.859056817921335 | 79.95139877071169 | 100.72773998872348 | 46.847105624390416 | 20.15129574185873 | 77.96790988815678 | 48.42722757440914 | 44.431614056140695 | 100.34233639424419 | 25.515243068941242 | 35.21354894093787 | 98.6578860818757 |
| SE_60_8_2 | SE_60 | SE_60_8 | TTACTTTCTTGTCCACTCTC | 133.30455152766078 | 133.9185929409421 | 116.52816979087619 | 92.39290275923144 | 79.5445884547055 | 84.89839076710406 | 129.1392735317577 | 26.46989773557318 | 85.24656867712163 | 9.477090282749604 | 36.3873339056358 | 61.51609367458132 |
| SE_60_8_3 | SE_60 | SE_60_8 | AGCTCTTTGCATCCGGAGAG | 22.759313675454276 | 38.97630690072195 | 39.50107450538176 | 37.737946197425615 | 89.08993906927017 | 25.989303296052263 | 13.990087965940416 | 58.61191641448347 | 6.215896944776189 | 3.6450347241344634 | 7.042709788187574 | 27.856344305470785 |
| SE_60_8_4 | SE_60 | SE_60_8 | AGAGAACAGCTCTTTGCATC | 4195.300154175405 | 4165.467875954079 | 4209.827015411061 | 4445.269800358824 | 4319.801450346874 | 4775.101325594669 | 4075.42024053972 | 3868.386483355909 | 4549.148578298345 | 4804.88477335405 | 4330.09273477066 | 4559.155017995386 |
| SE_60_8_6 | SE_60 | SE_60_8 | ACTTGACCTGAGATTAACTA | 512.6264461185654 | 604.6324532035072 | 654.7303099267027 | 581.684894836181 | 711.65989180414319 | 838.5881863526197 | 501.4908455483257 | 706.1790574454702 | 641.1253705897726 | 580.2895280822065 | 770.0029368418415 | 1328.9797595735022 |
| SE_60_8_7 | SE_60 | SE_60_8 | GACTTGACCTGAGATTAACT | 441.09717456713764 | 480.7077851089041 | 558.940204251152 | 562.165267492685 | 494.23704293190355 | 677.4545059170956 | 521.9378971908541 | 605.9715874465146 | 744.1316628174923 | 636.4230628338773 | 554.0265033374225 | 675.5163494076666 |
| SE_61_1_0 | SE_61 | SE_61_1 | GAGCTGGTGATGGTGCAGGA | 1292.9457721341405 | 1447.1203177498817 | 1554.3672817867723 | 1466.5746677413333 | 1553.770961148581 | 1282.1389626052448 | 1281.7072898026952 | 1490.8224546071037 | 1627.6770142535363 | 1862.6127440327107 | 1698.4668439179034 | 1216.393701338891 |
| SE_61_1_28 | SE_61 | SE_61_1 | CTACTGCTGTCCCTGCTGTG | 520.2128840103835 | 584.6446035108293 | 537.214613273192 | 558.2613420239858 | 659.689786917691 | 229.5721791151283 | 556.3750367993227 | 757.2281459355042 | 589.6222244759127 | 234.74023623425944 | 1106.879221710147 | 456.14763800208414 |
| SE_61_1_4 | SE_61 | SE_61_1 | GCAGATTCCAGAGCTGGTGA | 1449.0096373372558 | 1633.0073198917864 | 1573.1302921768286 | 1732.0415996127879 | 1518.7713422285105 | 1985.5827718183928 | 1594.8700281172075 | 1376.4346822498053 | 1752.8829384268852 | 1565.9069174881654 | 1397.9778929552335 | 1680.6661064300708 |
| SE_61_1_6 | SE_61 | SE_61_1 | TCTGCAGCTCATACCCCGCG | 676.2767492134985 | 154.90583511825392 | 279.470102125576 | 467.1697477543378 | 137.87728665482288 | 1373.101524141428 | 852.3192053096008 | 544.5236105603626 | 997.2074669976656 | 46.65644446892113 | 164.32989505771008 | 373.7392860983997 |
| SE_61_1_9 | SE_61 | SE_61_1 | CTAAATGCTGGAGGACCACG | 1312.45375528453 | 1237.2478959767634 | 1189.9698694746255 | 1210.2168952967525 | 1262.107470147994 | 1377.43307469077 | 1304.3066626707528 | 1436.9373056454012 | 1196.1161692305036 | 1097.8844589093003 | 1842.8423945757486 | 1240.768002606178 |
| SE_61_2_11 | SE_61 | SE_61_2 | TTCAGAAAGTGGACTTTGCA | 19.50798315038938 | 32.97995199291857 | 38.51354764274722 | 33.834020728726415 | 16.969512203670508 | 19.9251325269734 | 25.827854706351538 | 33.08737216946648 | 60.382998892111544 | 32.076305572383276 | 0.0 | 30.177706330926686 |
| SE_61_2_17 | SE_61 | SE_61_2 | GGTCTGGGCTGGAGAGGGAG | 328.3843830315546 | 471.713252747199 | 409.8236479933358 | 318.8205799437681 | 407.2682928880922 | 135.14437713947177 | 308.8580958634538 | 204.19635396013595 | 318.78671473923595 | 391.47672937204135 | 73.94845277596953 | 147.40648861644956 |
| SE_61_2_25 | SE_61 | SE_61_2 | AGGATCTTTCTCTTACGTGG | 29.26197472558407 | 67.95868895510495 | 84.92731016857079 | 94.99551973834723 | 95.45350614564661 | 78.83421999802519 | 67.79811860417279 | 41.59555358447214 | 27.4868811147129 | 111.53806255851458 | 160.80854016361627 | 24.37430126728694 |
| SE_61_2_3 | SE_61 | SE_61_2 | GAGTGAGAAAGAAAAGCACA | 1079.4417343215457 | 1259.2345306387092 | 1239.3462126063528 | 1115.2213755584053 | 1321.5007628608407 | 968.534702832881 | 1107.3692705348224 | 1292.2982215903048 | 1126.8533175601406 | 1589.964146667453 | 1361.5905590495977 | 1098.00423804064 |
| SE_61_2_31 | SE_61 | SE_61_2 | GTTCACCAAGGGGAAACAGA | 3.2513305250648967 | 7.9951398770711695 | 18.763010390056337 | 9.109159426964803 | 21.211890254588134 | 3.465240439473635 | 0.0 | 0.945353490556185 | 0.0 | 0.0 | 0.0 | 1.1606810127279494 |
| SE_61_3_12 | SE_61 | SE_61_3 | AAGCTTGTGGTGAGAACAAG | 359.81391144051526 | 370.7746117991755 | 394.02321819118305 | 387.78992989078733 | 386.05640263350404 | 209.64704658185492 | 289.48720483369016 | 292.11422858186114 | 336.5464202957393 | 322.2210696134866 | 272.3181118099195 | 374.8999671111277 |
| SE_61_3_18 | SE_61 | SE_61_3 | GAGCGGGTCGATCTCTTGAG | 2889.3490599410047 | 3129.097869388729 | 3141.3229500404846 | 2873.289144962612 | 3180.722943675491 | 2651.7752463071993 | 2832.454732796552 | 2737.7437086507116 | 2733.2186851458728 | 3100.4665363487743 | 2703.2267736993304 | 2278.4168279849646 |
| SE_61_3_20 | SE_61 | SE_61_3 | AGTGGAGTTCAAGACAGCCT | 1497.7795952132292 | 1594.0310129910635 | 1459.5647029738561 | 1447.0550403978373 | 1628.0125770396394 | 932.1496782184078 | 1471.1155764927732 | 1425.5930637587269 | 1353.2895634055587 | 2046.3224941290878 | 1801.759920811321 | 1348.7113367898774 |
| SE_61_3_21 | SE_61 | SE_61_3 | TGTTGCTCAGACAAGGCAAG | 1697.194534083876 | 1630.0091424378847 | 1605.7186786437685 | 1674.7840260719574 | 1518.7713422285105 | 2336.4383663150984 | 1872.5194662104866 | 1536.1994221538007 | 1618.7971614752846 | 2145.467438625545 | 1561.1340030482456 | 1672.5413393409751 |
| SE_61_3_32 | SE_61 | SE_61_3 | TTATAAACTCCAGGCAATGA | 62.859056817921335 | 90.94471610168455 | 85.91483704920533 | 68.96934994701923 | 63.63567076376441 | 109.1550738434195 | 63.49347651383142 | 120.05989330066355 | 49.72717555829155 | 115.91210422747594 | 68.07952795247988 | 123.03218734916264 |
| SE_61_4_1 | SE_61 | SE_61_4 | GGATTTCTCGGCTGAAGACG | 28.17819788389577 | 42.973876839257535 | 36.53849391747813 | 37.737946197425615 | 43.48437502190568 | 54.57753692170975 | 23.675533480822246 | 96.42605603673087 | 36.40739639083196 | 38.63736807582531 | 159.63475519891836 | 39.46315443275028 |
| SE_61_4_25 | SE_61 | SE_61_4 | TTAGCAGTGGGGGTTCTGTA | 182.0745094036342 | 240.85358879676897 | 257.744511147616 | 196.49758192452646 | 253.4820885423282 | 390.70585955065235 | 132.36775537005164 | 217.43130282792256 | 194.4687758437122 | 140.69834035159028 | 167.85124995180385 | 174.10215190919243 |
| SE_61_4_28 | SE_61 | SE_61_4 | GGGTTCTGTAGGGTCAGGAA | 21.675536833765978 | 11.992709815606755 | 16.78795666478725 | 9.109159426964803 | 14.848323171221195 | 0.0 | 11.837766740411123 | 7.56282792444948 | 0.0 | 0.7290069448268927 | 8.216494752885502 | 0.0 |
| SE_61_4_29 | SE_61 | SE_61_4 | GGTTCTGTAGGGTCAGGAAG | 13.005322100259587 | 16.989672238776237 | 24.688171565863602 | 14.314393385230405 | 13.787728665482287 | 2.598930329605226 | 18.294730416999005 | 19.852423301679885 | 0.8879852778251698 | 18.954180565499207 | 4.695139858791716 | 0.0 |
| SE_61_4_30 | SE_61 | SE_61_4 | TCTGTAGGGTCAGGAAGGGG | 607.9988081871356 | 774.5291755912696 | 760.395684228599 | 510.112927910029 | 597.114710666656 | 406.2994415282837 | 398.1794267229196 | 209.86847490347307 | 595.8381214206889 | 229.6371876204712 | 130.29013108147012 | 133.47831646371418 |

| | | | | | | | | | | | | | | | |
|---|---|---|---|---|---|---|---|---|---|---|---|---|---|---|---|
| SE_61_5_15 | SE_61 | SE_61_5 | GCCCAGTCATTCCTAAACCA | 662.1876502715506 | 719.5625889364053 | 756.4455767780607 | 625.9293834814387 | 712.7195125541614 | 1063.828814918406 | 654.3056525609056 | 755.3374389543918 | 695.292472537108 | 1019.8807158128228 | 651.4506554073506 | 1499.5998684445108 |
| SE_61_5_18 | SE_61 | SE_61_5 | TCTCGTTGATACAATACGGA | 260.10644200519175 | 292.82199799773156 | 240.95655448282875 | 251.15253848631528 | 207.87652449496372 | 257.2941026309174 | 170.0333768168143 | 245.7919075446081 | 169.60518806460743 | 345.54929184794713 | 185.45802442227279 | 304.0984253347228 |
| SE_61_5_19 | SE_61 | SE_61_5 | GCCCTTAGATCACTCAGAGA | 54.188842084414944 | 75.95382883217611 | 77.02709528549444 | 50.751031093089615 | 54.09032014919974 | 69.3048087894727 | 13.990087965940416 | 54.83050245225873 | 43.51127861343332 | 38.63736807582531 | 53.994108376104734 | 152.0492126673614 |
| SE_61_5_27 | SE_61 | SE_61_5 | CGTGACCGAGAGCCCGAGAA | 7141.0056098842015 | 7837.235864499014 | 7920.952965191677 | 7364.10474245626 | 8376.575461536855 | 8701.218743518297 | 7182.295929591257 | 8028.887195293679 | 6990.220107039737 | 8481.995803060896 | 7789.237025735457 | 8605.289028365018 |
| SE_61_5_9 | SE_61 | SE_61_5 | TAGGACAGACTCCACCCCCG | 8661.544518772886 | 10341.713430991558 | 10709.72882527163 | 9615.368429406133 | 10751.246575537996 | 10254.512770512354 | 8118.555662696501 | 11410.416631013153 | 11214.36607365407 | 9537.597859170237 | 10553.50061759908 | 10708.44302342806 |
| SE_62_1_13 | SE_62 | SE_62_1 | AGGCGTATACTCTTCTTCCT | 2387.5603822393223 | 2742.332977835411 | 2684.0980126406907 | 2685.9007224650504 | 2732.0914647909517 | 2721.080055096672 | 2523.596636933098 | 2754.760071480723 | 2468.599072353972 | 2364.169522073613 | 2956.7643260740833 | 3479.7216761583923 |
| SE_62_1_15 | SE_62 | SE_62_1 | CCTCAGGCTCACGTGTGCAG | 1396.9883489362173 | 1155.297712236784 | 1148.4937412439747 | 1506.9152309178917 | 1244.0773634315942 | 1376.5667645809015 | 1401.161117819571 | 1166.5662073463322 | 1331.9779167377546 | 1230.5637228677947 | 1318.1605153557744 | 1410.2274304644586 |
| SE_62_1_16 | SE_62 | SE_62_1 | CACTGCACACGTGAGCCTGA | 6185.114435515122 | 5103.8974190253075 | 5059.100117276769 | 5676.307631488638 | 4929.643295166283 | 4497.015780326909 | 5713.336693167514 | 4597.254024574728 | 4891.91089553886 | 4900.384683126373 | 4591.846781898298 | 4272.4668078515815 |
| SE_62_1_17 | SE_62 | SE_62_1 | GCTGGAACACAGCTACCCTC | 4745.858789753061 | 4022.554750651432 | 3943.194762499734 | 4525.950926711941 | 4514.9508406890845 | 4237.122747366387 | 4885.769181951499 | 4891.258960137701 | 4423.942654124996 | 4623.362044092153 | 6250.404937016472 | 4111.132147082397 |
| SE_62_1_22 | SE_62 | SE_62_1 | GTTTGGGAGAATGAAGCCAG | 120.29922942740117 | 133.9185929409421 | 125.4159115545871 | 100.20075369661284 | 79.5445884547055 | 145.54009845789267 | 81.7882065701132 | 173.94504226233803 | 93.23845417164283 | 180.79372231706938 | 136.15905590495976 | 59.19473164912542 |
| SE_62_2_19 | SE_62 | SE_62_2 | CCTGAAATCAAGGCATCAGA | 1531.3766773055663 | 1658.9915244922677 | 1772.6107184290065 | 1653.963090238895 | 1613.1642538614276 | 1994.245872917077 | 1642.221095078852 | 1548.489017531031 | 2140.9325048364844 | 1196.3003964609309 | 1464.8836359430154 | 1831.5546380847043 |
| SE_62_2_25 | SE_62 | SE_62_2 | AGAGAGAGAAATGACAAACG | 1036.0906606540138 | 1371.1664889177055 | 1365.7496510235744 | 1052.7585680592179 | 1342.712653115429 | 1464.9303957874793 | 966.3922302626535 | 1282.844686684743 | 1549.5343098049213 | 1100.8004866886079 | 1291.1634611677218 | 1709.6831317482695 |
| SE_62_2_3 | SE_62 | SE_62_2 | GGCCACTGTGATGTACTGCT | 768.3977807570039 | 828.4963697615 | 847.2980481404388 | 873.1779964990546 | 839.9908540816901 | 909.6256153618292 | 885.6801843053049 | 782.7526901805212 | 858.6817636569392 | 617.468882268378 | 680.7952795247988 | 702.2120127004094 |
| SE_62_2_8 | SE_62 | SE_62_2 | TCTGGGATAGAGTGAAGCCT | 614.5014692372655 | 664.596002281541 | 590.5410638554573 | 622.0254580127394 | 778.4763723433846 | 295.41174746512735 | 701.6567195225501 | 588.9552246165033 | 571.8625189194094 | 810.6557226475046 | 541.1148687257453 | 352.84702786929665 |
| SE_62_3_10 | SE_62 | SE_62_3 | GCAACAAGATTGATTAGAAA | 205.91759992077678 | 406.75274124599576 | 371.3101003505886 | 304.5061865585377 | 336.20846053522195 | 239.1015903236808 | 311.0104170889831 | 260.91756339350707 | 467.96824141386446 | 672.873410075222 | 191.32694924576242 | 629.0891088985486 |
| SE_62_3_11 | SE_62 | SE_62_3 | ATTTTCCTGCAGATTTCACA | 0.0 | 0.0 | 0.987526862634544 | 0.0 | 0.0 | 0.0 | 0.0 | 0.0 | 0.0 | 0.0 | 0.0 | 0.0 |
| SE_62_3_12 | SE_62 | SE_62_3 | AATATACAAATACAAATCAA | 17.340429467012783 | 25.984204600481302 | 28.63827901640178 | 13.013084895664004 | 18.030106716399914 | 24.256683076315444 | 5.380803063823238 | 13.23494886778659 | 33.74344055735645 | 6.561062503442034 | 16.432989505771005 | 1.1606810127279494 |
| SE_62_3_31 | SE_62 | SE_62_3 | GAGAACATGGGTTATGCCGG | 1298.364656342582 | 1677.9799817003118 | 1733.1096439236248 | 1488.6969120639621 | 1713.9207325707214 | 1259.6148997486664 | 1311.8397869601054 | 1532.4180081915758 | 1780.4104820394655 | 1375.6361048883464 | 1642.1251656124027 | 2009.1388330320806 |
| SE_62_4_0 | SE_62 | SE_62_4 | AAACCTTACACACTTCCCAG | 63.94283365960963 | 87.94653864778286 | 92.82752508764715 | 63.76411598875362 | 98.63528968383483 | 112.62031428289313 | 80.71204595734856 | 220.2673632995911 | 83.47061611556596 | 39.36637502065221 | 262.9278320923361 | 94.01516203096391 |
| SE_62_4_15 | SE_62 | SE_62_4 | GTTCTGTGCTGGGCTTCCAA | 10.837768416882989 | 14.990887269508443 | 8.887741763710896 | 11.711776406097604 | 11.666539640023474 | 1.7326202197368175 | 11.837766740411123 | 10.398888396118036 | 18.647690834328564 | 2.187020834480678 | 4.695139858791716 | 4.642724050911798 |
| SE_62_4_2 | SE_62 | SE_62_4 | GTCTTTTGAAAGGCAGCAGG | 22.759313675454276 | 34.97873696218637 | 46.41376254382357 | 16.917010364363207 | 29.69664635642339 | 89.2299413164461 | 12.913927353175769 | 30.25131169779792 | 27.527543612580263 | 48.84346530340181 | 9.390279717583432 | 128.83559241280238 |
| SE_62_4_3 | SE_62 | SE_62_4 | GTGTCAAGTAAGGCTCTGCA | 171.23674098675122 | 60.962941562667666 | 138.25376076883617 | 170.47141213319847 | 67.87804881468203 | 643.6684116322277 | 265.8116713528679 | 143.69373056454012 | 524.7992991946753 | 24.78623612411435 | 115.03092654039705 | 167.1380658328247 |
| SE_62_4_33 | SE_62 | SE_62_4 | GTGAGTACACGAGCTCAAGG | 757.5600123401209 | 921.4398708324522 | 928.2752508764714 | 830.2348163433635 | 835.7484760307725 | 729.4331125092001 | 723.1799317778431 | 1036.1074256495788 | 1027.3989664437215 | 772.7473615165062 | 917.8998423937805 | 676.6770304203945 |
| SE_62_5_14 | SE_62 | SE_62_5 | AGATGCCTAAAGAGAACAAA | 383.6570019576578 | 541.6707266715717 | 543.1397744489992 | 502.30507697263056 | 569.5392533356915 | 505.05879405328227 | 401.4079085612135 | 410.2834149013843 | 602.9420036432903 | 605.8047711511477 | 462.471276090984 | 454.9869569893562 |
| SE_62_5_19 | SE_62 | SE_62_5 | CAGAAAAGAGACTAAGAGCG | 571.1503955697335 | 677.5881045817816 | 708.0567605089681 | 611.6149900962082 | 693.628811325032 | 859.3796289894615 | 509.02396983767824 | 604.0808804654022 | 716.604119204912 | 891.5754935232898 | 677.2739246307051 | 1044.6129114551545 |
| SE_62_5_5 | SE_62 | SE_62_5 | TTGATTGCGCGAGGACTCCA | 450.8511661423323 | 519.684092009626 | 542.1522475863646 | 458.06058832737295 | 588.6299545648208 | 338.7272529585478 | 460.5967422632691 | 463.2232103725307 | 388.9375516874244 | 581.7475419718603 | 516.4653844670888 | 240.26096963468552 |
| SE_62_5_6 | SE_62 | SE_62_5 | GCATAAACATTGATTGCGCG | 11934.550580671548 | 12269.541533850343 | 12327.297826267013 | 12443.11177723392 | 12312.441698275683 | 12119.678437059038 | 11907.717180240825 | 12014.497511478556 | 12793.203897627222 | 12788.968833098177 | 13080.659646593722 | 12338.039165298103 |
| SE_62_5_7 | SE_62 | SE_62_5 | ATCATGACATGTCGAATGGG | 838.8432754667433 | 692.5789918512901 | 827.5475108877479 | 925.2303360817107 | 575.9028204120679 | 775.3475483322258 | 888.9086661435988 | 784.6433971616335 | 767.2192800409467 | 842.003021275061 | 822.8232602532482 | 890.2423367623372 |
| SE_63_1_15 | SE_63 | SE_63_1 | TATTTGAGAAGACAGACCCA | 165.81785677830973 | 189.88457208044028 | 166.89203978523796 | 157.45832723753446 | 217.42187510952837 | 229.5721791151283 | 128.06311291899306 | 292.11422858186114 | 253.96378945799856 | 121.74415978609107 | 219.49778839851274 | 135.79967848917008 |
| SE_63_1_22 | SE_63 | SE_63_1 | TTTGACTAGTCAGGAAGCCA | 118.13167574402458 | 113.93074324826416 | 113.56558920297256 | 114.51514708184324 | 80.60518296743491 | 116.95186483223517 | 77.48356411905462 | 76.57363273505099 | 139.41368861855165 | 51.759493082709376 | 41.08247376442752 | 47.58792152184593 |
| SE_63_1_23 | SE_63 | SE_63_1 | GCTGCCCTGTGGATGTTGGA | 1783.89668141894 | 1937.8220277051248 | 1970.1160909559153 | 1918.1287136208744 | 1900.585366811097 | 2326.908955106546 | 1886.509554176427 | 2022.1111162996797 | 2156.9162398373373 | 2037.574410791165 | 1531.7893789307973 | 2420.0199115377745 |
| SE_63_1_30 | SE_63 | SE_63_1 | AGACCCACAGACGCAGGCAG | 2051.58956131595 | 2233.642203156758 | 2212.0601723013788 | 2220.032283200279 | 2269.6722572409303 | 1995.9784931368138 | 2210.4338986185858 | 2093.9579815819498 | 2102.749137890002 | 2559.54338328722 | 1974.3063106219167 | 2222.7041393740233 |
| SE_63_1_7 | SE_63 | SE_63_1 | GGACAGACTCCCAGGACAAG | 715.2927155142772 | 709.5686640900662 | 715.9569754100444 | 752.1563069693794 | 749.8403204996906 | 884.5026221756453 | 879.223220628717 | 744.9385505582737 | 749.4595744844434 | 1076.0142505644935 | 557.5478582315163 | 838.0116911895794 |
| SE_63_2_18 | SE_63 | SE_63_2 | TCATGGTCAGAAGCAACATG | 5238.977252721237 | 4159.4715210462755 | 4402.394753624797 | 5002.229833893243 | 4363.285825368779 | 4748.245712188748 | 4912.673197270616 | 5214.569853907917 | 4692.114208028197 | 3880.5039673135498 | 6020.343083935678 | 4918.96613194105 |
| SE_63_2_19 | SE_63 | SE_63_2 | ACAGTAAATAGGCAGAAAGG | 1494.5282646881642 | 1670.9842343078744 | 1651.1449143249577 | 1653.963090238895 | 1752.10213502898 | 1475.3261171059 | 1753.0656381936108 | 1949.3188975268536 | 1353.2895634055587 | 2243.154369232349 | 2061.166398009563 | 1658.6131671882397 |
| SE_63_2_24 | SE_63 | SE_63_2 | ACTTCCCTGCCCTACAGCTG | 444.3485050922026 | 450.7260105698872 | 483.88816269092655 | 434.63703551517773 | 607.7206557939501 | 323.99998109078484 | 363.74228711445085 | 406.50200093915953 | 502.5996672490461 | 347.7363126824278 | 467.16641594977574 | 423.64856964570157 |
| SE_63_2_37 | SE_63 | SE_63_2 | CTGCCTTGCAGAGAACCAGC | 239.51468201311405 | 254.84508358164354 | 274.53246781240324 | 201.70281588279207 | 217.42187510952837 | 267.6898239493383 | 208.7751588763416 | 304.4038239590916 | 237.9800544571455 | 166.94259036535843 | 403.7820278560876 | 354.0077088820246 |
| SE_63_2_5 | SE_63 | SE_63_2 | GCTCAGCCAGTTCAGCCACA | 2710.5258810624355 | 2518.4690612774184 | 2622.8713471573487 | 2645.5601592884923 | 2410.7313274339417 | 2289.657620382204 | 2639.8219831116803 | 2736.7983551601556 | 2529.8700565239087 | 2272.3146470254246 | 2387.478618195588 | 2572.0691242051357 |
| SE_63_3_1 | SE_63 | SE_63_3 | GGGCCGTTCCGTGCACTGCA | 189.6609472954523 | 47.97083926242702 | 89.8649444997435 | 140.54131687317124 | 40.30259148371746 | 464.34221888946706 | 300.24881096133663 | 102.09817698006798 | 301.91499446055775 | 16.038152786191638 | 77.46980767006332 | 134.63899747644214 |
| SE_63_3_2 | SE_63 | SE_63_3 | GAGACTTGCTTGAACCCTAG | 338.13837460674927 | 360.7806869528365 | 398.9608525043558 | 409.91217421341617 | 412.57126545173924 | 294.54543735525897 | 341.1429142463933 | 427.29977773139564 | 335.6584350179142 | 314.20199322039076 | 450.73342644400475 | 557.1268861094158 |
| SE_63_3_21 | SE_63 | SE_63_3 | GCAGGAAGCAGACTCCTGTG | 4656.98908873462 | 5338.754652914274 | 5387.946562534073 | 4924.151324519259 | 5333.729804516187 | 5669.133358978866 | 4417.639315398877 | 5073.712183815045 | 5290.616285282362 | 5318.834669457009 | 5307.855610364035 | 5346.096744624935 |
| SE_63_3_24 | SE_63 | SE_63_3 | AAGCTCTTAGGACAACGCAG | 172.32051782843953 | 213.8699917116538 | 186.64257703792882 | 287.5891761941745 | 252.42149402959882 | 256.42779252104896 | 242.13613787204568 | 318.58412631743437 | 229.1002016788938 | 468.751465523692 | 531.7245890081618 | 120.71082532370674 |
| SE_63_3_8 | SE_63 | SE_63_3 | GGACAGAGTGAACCAGAGGA | 980.8180417279106 | 1218.2594387687195 | 1242.3087931942564 | 1176.382874568026 | 1218.6230951260884 | 1378.2993848006383 | 873.8424175648937 | 1323.494886778659 | 1297.3464909025731 | 1321.6895909711563 | 1239.516922721013 | 1635.3995469336808 |
| SE_63_4_1 | SE_63 | SE_63_4 | TGAAACTGAGAGGCTACAGT | 231.92824412129596 | 336.795267321623 | 332.79655270784133 | 299.3009526002721 | 334.0872715097631 | 350.8555944967055 | 207.69899826357695 | 165.43686084733238 | 288.59521529318016 | 406.7858752134061 | 104.46686185811568 | 573.3764202876071 |
| SE_63_4_34 | SE_63 | SE_63_4 | GGACTGTCCTCAACACAGCA | 1028.5042227621957 | 958.4173927639065 | 941.1131000907204 | 1024.1297812887572 | 887.7176071545134 | 1192.0427111789304 | 993.2962455817697 | 882.0148066889207 | 846.2499697673868 | 809.1977087578508 | 1172.6111797332312 | 999.3463519587644 |
| SE_63_4_4 | SE_63 | SE_63_4 | GGTTGAATAGGAGTTTTCCA | 14.089098941947885 | 36.977521931454156 | 24.688171565863602 | 37.737946197425615 | 37.120807945529236 | 23.390372966447035 | 26.904015319116187 | 41.59555358447214 | 33.74344055735645 | 61.965590310285876 | 132.63770101086598 | 4.642724050911798 |
| SE_63_4_41 | SE_63 | SE_63_4 | GCAGCCTCAAAAGCTAAAGT | 2469.9274222076333 | 2437.518270022073 | 2482.642532663244 | 2706.7216582981127 | 2604.820123263423 | 2779.122832457855 | 2672.1068014946195 | 2779.339262235184 | 2226.179091507701 | 3421.2295920726074 | 3092.923381979043 | 2472.250557110532 |
| SE_63_4_6 | SE_63 | SE_63_4 | GGCTGACTCGGCTCGCACAG | 955.8911743690796 | 969.4107100948793 | 965.801271656584 | 1100.9069821731748 | 1127.4119670313594 | 1031.7753408532749 | 1060.0182035731777 | 1173.1836817802257 | 1038.9427750554487 | 987.8044102404396 | 1115.0957164630327 | 1225.6791494407146 |
| SE_64_1_15 | SE_64 | SE_64_1 | GCATTTCTCAGAAAAGGAAG | 82.36703996831072 | 98.93985597875572 | 85.91483704920533 | 93.69421124878083 | 54.09032014919974 | 44.18181560328885 | 71.02660044246673 | 88.86322811228139 | 103.0062922277197 | 123.93118062057175 | 38.73490383503166 | 48.74860253457388 |
| SE_64_1_19 | SE_64 | SE_64_1 | TTTTATCTCCATCGGTTCCA | 3.2513305250648967 | 0.9993924846338962 | 0.0 | 15.615701874796805 | 10.605945127294067 | 0.0 | 9.685445514881827 | 2.836060471668555 | 0.0 | 1.4580138896537853 | 0.0 | 0.0 |
| SE_64_1_23 | SE_64 | SE_64_1 | GGTTATTTCAGGAAAGAGGT | 10.837768416882989 | 29.981774539016886 | 28.63827901640178 | 32.53271223916001 | 23.33307928004695 | 27.72192351578908 | 4.30464245105859 | 1.89070698111237 | 12.431793889552377 | 2.187020834480678 | 2.347569929395858 | 69.64086076367697 |
| SE_64_1_5 | SE_64 | SE_64_1 | GTTTGCTGCCAAACATACTG | 411.83519984155356 | 546.6676890947413 | 532.2769789600193 | 422.9252591090801 | 448.63147888453904 | 382.04275845196827 | 359.43764466339223 | 450.9336149953003 | 412.91315418870397 | 418.44998633063636 | 423.7363722559524 | 509.5389645875698 |
| SE_64_1_8 | SE_64 | SE_64_1 | AGTCACCCAAGAGGAAGGAA | 176.65562519519273 | 284.8268581206604 | 280.4576289882105 | 140.54131687317124 | 203.6341464440461 | 94.42780197565655 | 129.1392735317577 | 96.42605603673087 | 214.00445195586593 | 78.73275004130441 | 93.90279717583432 | 126.51423038734649 |
| SE_64_2_0 | SE_64 | SE_64_2 | TGCAGCAGCCGAGAGCCGAA | 3603.557998613594 | 3955.595454180961 | 3912.5814297580637 | 3900.021543230502 | 4039.80449898631 | 4396.523807582174 | 3695.5355442337996 | 3722.8020458102565 | 4113.147806886186 | 4057.6526549064847 | 4899.378442649156 | 3577.2188812275404 |
| SE_64_2_1 | SE_64 | SE_64_2 | ATACTTTTGTGGAACTCTAT | 1.083776841688299 | 6.9957473924372735 | 1.975053725269088 | 15.615701874796805 | 2.1211890254588135 | 0.0 | 0.0 | 0.945353490556185 | 0.0 | 26.244250013768134 | 0.0 | 0.0 |
| SE_64_2_15 | SE_64 | SE_64_2 | GCCCTGAATCTAAGTCAGAA | 118.13167574402458 | 180.8900397187352 | 152.07913684571977 | 105.40598765487843 | 89.08993906927017 | 142.07485801841904 | 112.99686434028798 | 102.09817698006798 | 198.02071695501286 | 131.95025701366757 | 124.42120625798047 | 89.37243798005211 |
| SE_64_2_2 | SE_64 | SE_64_2 | GATACTTTTGTGGAACTCTA | 7.586437891818092 | 8.994532361705065 | 13.825376076883616 | 13.013084895664004 | 28.63605184369398 | 0.0 | 0.0 | 7.56282792444948 | 0.0 | 1.4580138896537853 | 1.173784964697929 | 1.1606810127279494 |
| SE_64_2_30 | SE_64 | SE_64_2 | CAGAGGCTCCTCCCTCTGGA | 895.1996712345349 | 1056.3578562580283 | 1024.0653565520222 | 921.3264106130115 | 1193.1688268205826 | 848.1175975611721 | 823.2628687649553 | 1166.5662073463322 | 952.8082031064072 | 1044.6669519369373 | 800.5213459239876 | 1159.5203317152216 |
| SE_64_3_11 | SE_64 | SE_64_3 | AACTCACTCACTGGAAGCCA | 275.27931778882794 | 249.84812115847404 | 286.38279016401776 | 279.7813252567761 | 239.69435987684594 | 126.48127604078768 | 316.39122015280634 | 194.7428190545741 | 304.57895029403323 | 197.5608820480879 | 224.19292825730446 | 182.22691899828806 |
| SE_64_3_14 | SE_64 | SE_64_3 | TTTTCTGGTGAGTAGGGTAG | 17.340429467012783 | 16.989672238776237 | 7.900214901076352 | 13.013084895664004 | 9.545350614564661 | 1.7326202197368175 | 32.28481838293943 | 16.071009339455145 | 14.207764445202717 | 69.2556597585548 | 3.521354894093787 | 1.1606810127279494 |
| SE_64_3_19 | SE_64 | SE_64_3 | GGAAGCACATGGAGAATGTG | 1844.5881845534848 | 1863.8669838422163 | 1899.0141568462282 | 1851.7619806529879 | 1664.0727904724392 | 1932.73785511642 | 1850.9962539551937 | 1828.313650735662 | 1999.7428456622824 | 1649.7427161432581 | 1807.6288456348107 | 1741.0215190919241 |
| SE_64_3_3 | SE_64 | SE_64_3 | AGCCCACTCCGAAAAGCGGG | 132.22077468597246 | 142.91312530264716 | 115.54064292824165 | 206.90804984105768 | 99.69588419656424 | 123.01603560131404 | 235.6791741954578 | 190.96140509234937 | 123.4299536176986 | 129.7632361791869 | 219.49778839851274 | 78.92630886550056 |
| SE_64_3_45 | SE_64 | SE_64_3 | GAATGCCGAGTCCATCCCCG | 859.4350354588211 | 801.5127726763848 | 881.8614883326478 | 901.8067832695154 | 799.6882625979727 | 805.6684021776201 | 689.818952782139 | 807.331880934982 | 981.2237319968126 | 596.3276808683981 | 852.1678843706965 | 782.2990025786379 |
| SE_65_1_10 | SE_65 | SE_65_1 | GCGTCTCAACGTTATTTGAA | 341.38970513181414 | 618.6239479883817 | 618.1918160092246 | 419.02133364038093 | 615.144817383056 | 401.9678909789416 | 347.59987792298114 | 514.2722988625646 | 656.2211203128005 | 459.2743752409424 | 361.52576912696213 | 377.22132913658356 |
| SE_65_1_11 | SE_65 | SE_65_1 | GCATTTTGTAGATGCCAAAG | 95.37236206857031 | 105.935603371193 | 134.303653318298 | 98.89944520704643 | 132.57431409117584 | 171.52940175394494 | 78.55972473181926 | 112.49706537618601 | 136.74973278507613 | 139.96933340676338 | 199.54344399864794 | 124.19286836189059 |
| SE_65_1_16 | SE_65 | SE_65_1 | AAAAATGCGAAGATATCGCA | 36.84841261740216 | 35.97812944682026 | 50.36386999436174 | 37.737946197425615 | 29.69664635642339 | 15.593581977631358 | 18.294730416999005 | 99.26211650839943 | 26.639558334755094 | 8.019076393095819 | 38.73490383503166 | 83.56903291641235 |
| SE_65_1_19 | SE_65 | SE_65_1 | TCTTGAGCTCCTGAGTTCAA | 584.1557176699931 | 542.6701191562056 | 554.9900968006137 | 614.217607075341 | 484.6916923173389 | 548.3742995467027 | 722.1037711650785 | 589.9005781070595 | 628.6935767002202 | 980.5143407921706 | 585.7186973842665 | 561.7696101603275 |
| SE_65_1_5 | SE_65 | SE_65_1 | GGTAGTCCCAGTTTCTCAGG | 55.27261892610324 | 62.96172653193546 | 64.18924607124536 | 49.44972260352321 | 30.757240869152795 | 7.796790988815679 | 57.03651247652632 | 42.54090707502832 | 55.94307250298569 | 69.98466670338169 | 10.564064682281362 | 38.30247342002233 |
| SE_65_2_2 | SE_65 | SE_65_2 | AGCGGCAGGGCCTGGCCGGC | 1168.3114353399862 | 1413.1409732723291 | 1420.0636284684742 | 1164.6710981619283 | 1269.5316317370998 | 1819.2512307236584 | 1049.2565974455313 | 1218.5606493269224 | 1427.880326742873 | 1638.0786050260278 | 943.7231116171349 | 1308.087501344399 |
| SE_65_2_31 | SE_65 | SE_65_2 | TGAGGAGGTGGGCGAGGGGC | 4666.743080309815 | 5637.573005819809 | 5545.950860555599 | 4585.811117231995 | 4797.068981075106 | 4393.058567142701 | 4480.056630939227 | 4166.172832881108 | 4818.208117479371 | 4445.484349554391 | 3651.645025175257 | 3592.3077343930036 |
| SE_65_2_32 | SE_65 | SE_65_2 | GCGCTGAGGAGGTGGGCGAG | 6363.937614393692 | 6004.350047680448 | 5975.525045801626 | 5798.63062950788 | 5711.301451047855 | 5355.529099206503 | 6407.460288400711 | 5798.798311071639 | 5146.762670274684 | 6037.635517056325 | 6029.733363653261 | 4880.663658521027 |
| SE_65_2_39 | SE_65 | SE_65_2 | TCCTCAGCGCCTTCTCCTCG | 820.4190691580422 | 854.4805743619812 | 824.5849302998442 | 867.9727625407891 | 845.2938266453372 | 1143.5293450262996 | 954.5544635222423 | 667.4195643326666 | 795.6348089313522 | 809.9267157026777 | 868.6008738764675 | 773.0135544768143 |
| SE_65_2_45 | SE_65 | SE_65_2 | AGCCCCCCGCTCGCTGCCTC | 128.96944416090756 | 201.87728189604704 | 215.2808560543306 | 226.4276771845537 | 206.8159299822343 | 128.21389626052448 | 162.50025252746178 | 182.45322367734371 | 224.66027528976795 | 203.39293760670304 | 79.81737759945918 | 186.86964304919985 |
| SE_65_3_13 | SE_65 | SE_65_3 | GCAAAAGTAGGTGTACTATC | 2268.3449296536096 | 2098.724217731182 | 2129.107915840077 | 2283.796399189033 | 2037.4020589531904 | 2306.1175124697043 | 2334.1923690865206 | 2009.8215209224493 | 2416.207940962287 | 2819.798862590421 | 1710.2046935648825 | 1984.7645317647934 |
| SE_65_3_14 | SE_65 | SE_65_3 | GCCTAAGCCACCTCACCAAA | 2058.0922223660796 | 2158.6877668092156 | 2220.9479140650897 | 2091.2027427332055 | 2340.7320895938005 | 2729.743156195356 | 2030.7150762868898 | 2522.2031128039016 | 1965.999405104926 | 1740.1395773017928 | 2315.877735349014 | 2312.076577354075 |
| SE_65_3_2 | SE_65 | SE_65_3 | CTTCTCAGACACACACCCTG | 198.3311620289587 | 291.8226055130977 | 264.6571991860578 | 206.90804984105768 | 327.7237044333867 | 356.9197652657844 | 205.54667703804768 | 182.45322367734371 | 261.9556569584251 | 540.1941461167274 | 230.0618530807941 | 85.89039494186825 |
| SE_65_3_3 | SE_65 | SE_65_3 | ATTTTGCAGTCAAAGCCCCA | 29.26197472558407 | 37.97691441608806 | 46.41376254382357 | 32.53271223916001 | 48.78734758555271 | 31.187163955262715 | 35.513300221233365 | 20.79777679223607 | 104.78226278337003 | 48.114458358574915 | 4.695139858791716 | 17.410215190919242 |
| SE_65_3_9 | SE_65 | SE_65_3 | GAACCCTCATCTCTACTAGG | 26.010644200519174 | 42.973876839257535 | 53.32645058226538 | 35.13532921829281 | 14.848323178211695 | 25.12299318618385 | 37.665621446762664 | 0.945353490556185 | 83.47061611556596 | 81.64877782061198 | 9.390279717583432 | 24.37430126728694 |
| SE_65_4_11 | SE_65 | SE_65_4 | GATTGGTGTGACAGGGCCTA | 516.9615534853186 | 558.660398910348 | 548.0774087621719 | 564.7678844718178 | 451.8132624227273 | 548.3742995467027 | 588.6598551822622 | 551.1410849942558 | 626.0296208667447 | 928.0258407646344 | 545.810008584537 | 532.7525848421288 |
| SE_65_4_14 | SE_65 | SE_65_4 | ACTTACAAAGACTAACTCTG | 544.055974527526 | 602.6336682342394 | 643.8675144377227 | 646.750319314501 | 654.386814354044 | 744.1603843769631 | 548.8419125099703 | 527.5072477303512 | 626.9176061445698 | 736.2970142751616 | 572.8070627725893 | 580.3405063639747 |
| SE_65_4_24 | SE_65 | SE_65_4 | AGATCCGGTCGTTGAAGCCA | 901.7023322846646 | 1012.3845869341368 | 1065.541484782673 | 1028.0337067574565 | 1023.4737047838776 | 1140.0641045868258 | 893.2133085946574 | 1280.0086262130744 | 908.4089392151487 | 1308.5674659642723 | 879.1649385587489 | 1028.363377276963 |

| | | | | | | | | | | | | | | | |
|---|---|---|---|---|---|---|---|---|---|---|---|---|---|---|---|
| SE_65_4_29 | SE_65 | SE_65_4 | GCCTTTCATGTAGTGACTGG | 267.69287989700985 | 340.7928372601586 | 304.15827369143955 | 309.7114205168033 | 274.6939787969163 | 210.51335669802333 | 261.50702890180935 | 459.4417964103059 | 216.66840778934142 | 317.11802099969833 | 657.3195802308403 | 221.69007343103834 |
| SE_65_4_5 | SE_65 | SE_65_4 | TCAATCATGGGGTCAGCTGG | 888.6970101844051 | 1057.3572487426623 | 1157.1814830076856 | 841.946592749461 | 1093.4729426240183 | 947.7432601960392 | 690.8951133949037 | 421.6276567880585 | 959.9120852380286 | 667.7703614614337 | 713.6612585363408 | 381.86405318749536 |
| SE_65_5_1 | SE_65 | SE_65_5 | GTGGACAGGCATATTTACGG | 404.24876194973547 | 513.6877371018227 | 474.01289406458113 | 369.57161103685775 | 414.69245447719805 | 301.47591823420623 | 397.1032661101549 | 380.03210320358636 | 491.94384391514404 | 646.6291600614538 | 309.8792306802533 | 388.8281392636831 |
| SE_65_5_2 | SE_65 | SE_65_5 | AGTGGACAGGCATATTTACG | 1117.3739237806362 | 1221.2576162226212 | 1215.6455679031237 | 1240.1469905567797 | 1157.1086133877827 | 929.5507478888026 | 1125.6640009518212 | 1091.8832815923936 | 1025.622995888071 | 1300.5483895711766 | 1464.8836359430154 | 1078.2726608242651 |
| SE_65_5_3 | SE_65 | SE_65_5 | AAGTGGACAGGCATATTTAC | 716.3764923559655 | 659.5990398583715 | 681.3935352178354 | 748.2523815006803 | 527.1154728265152 | 810.8662628368305 | 602.6499431482026 | 782.7526901805212 | 626.9176061445698 | 790.2435281923516 | 537.5935138316515 | 465.4330861039077 |
| SE_65_5_4 | SE_65 | SE_65_5 | AAAGTGGACAGGCATATTTA | 118.13167574402458 | 181.8894322033691 | 176.76730841158337 | 137.93869989403845 | 155.90739337122278 | 124.74865582105086 | 103.31141882540615 | 141.80302358342774 | 146.517570841153 | 297.4348334893722 | 152.59204541073078 | 118.38946329825085 |
| SE_65_5_5 | SE_65 | SE_65_5 | GGTATTGCTGTGCATTTCTG | 211.3364841292183 | 207.87363680385042 | 203.43053370271608 | 195.19627343496006 | 211.05830803315195 | 148.1390287874979 | 234.60301358269314 | 287.38746112908024 | 280.6033477925365 | 86.75182643440023 | 207.75993875153344 | 155.53125570554522 |
| SE_66_1_0 | SE_66 | SE_66_1 | GAGCTGGCAGAGAAATGGGA | 790.0733175907699 | 1221.2576162226212 | 1127.7556771286493 | 843.2479012390274 | 984.2317078128895 | 829.0587751440671 | 845.8622416330129 | 726.0314807471501 | 961.6880558846589 | 905.4266254750007 | 727.746678112716 | 630.2497899112766 |
| SE_66_1_10 | SE_66 | SE_66_1 | TCTGGGGGTCAGGAGCACCA | 2319.28244121296 | 2398.541963121351 | 2393.765115026135 | 2352.765749136052 | 2191.1882632989546 | 1779.4009656697115 | 2228.728629035585 | 2223.471409788147 | 2489.022733743951 | 2003.311084384301 | 2020.083924245136 | 2026.5490482229998 |
| SE_66_1_2 | SE_66 | SE_66_1 | CCTTGAGCTTTCTAGGCCAT | 479.0293640262281 | 327.800734959918 | 346.62192878472496 | 374.7768449951233 | 364.8445123789159 | 493.796762624993 | 492.8815606462085 | 644.7310805593182 | 628.6935767002202 | 299.62185432385286 | 613.8895365370169 | 381.86405318749536 |
| SE_66_1_29 | SE_66 | SE_66_1 | GTGCCCAAGCCACACTCACA | 655.6849892214209 | 613.6269855652123 | 679.4184814925663 | 611.6149900962082 | 637.4173021503734 | 627.2085195447279 | 620.9446735652016 | 488.7477546175476 | 589.6222244759127 | 754.5221878958339 | 413.172307573671 | 767.2101494131746 |
| SE_66_1_47 | SE_66 | SE_66_1 | GTAGGAGGCTTCAGGGCTGA | 2102.5270728753 | 2213.65435346408 | 2330.563395817524 | 1920.7313306000071 | 1981.1905497785317 | 1152.1924461249837 | 1813.3306325084309 | 1522.0191197954578 | 1844.3454220428775 | 1523.6245146882056 | 1390.9351831670458 | 1284.87388108984 |
| SE_66_2_2 | SE_66 | SE_66_2 | AAAGTCACTGAGATGGTGCA | 385.8245556410344 | 315.8080251443112 | 387.11053015274126 | 425.52787608821296 | 355.29916176435125 | 286.74864636644327 | 350.82835976127507 | 171.10898179066947 | 408.4732277995781 | 94.04189588266915 | 119.72606639918877 | 179.90555697283216 |
| SE_66_2_25 | SE_66 | SE_66_2 | ACTGTAATCAGCCTGCGGGG | 233.01202096298425 | 387.7642840379517 | 339.70924074628317 | 296.6983356211393 | 390.2987806844217 | 260.759343070391 | 238.90765603375175 | 284.5514006574117 | 192.69280528806183 | 201.93492371704926 | 251.1899824453568 | 116.06810127279495 |
| SE_66_2_26 | SE_66 | SE_66_2 | AGCAACCTGAGCCGAAAGCA | 721.7953765644071 | 916.4429084092828 | 857.1733167667842 | 640.243776866669 | 746.6585369615024 | 444.4170863624937 | 595.11681885885 | 484.9663406553229 | 704.1723253153597 | 355.02638213069673 | 306.35787578615947 | 280.88480508016374 |
| SE_66_2_5 | SE_66 | SE_66_2 | AATTGGAGTGATGATGGCCG | 163.65030309493312 | 154.90583511825392 | 193.55526507637063 | 204.30543286192486 | 177.11928362581094 | 129.94651648026132 | 161.42409191469713 | 130.45878169675353 | 274.38745084797745 | 185.89677093085763 | 110.33578668160533 | 168.29874684555267 |
| SE_66_2_9 | SE_66 | SE_66_2 | TGTGATACCTGGAGTCCCCT | 549.4748587359676 | 623.6209104115512 | 590.5410638554573 | 573.8770438987826 | 470.9039636518566 | 551.8395399861763 | 510.1001304504429 | 434.86260565584513 | 704.1723253153597 | 454.90033357198104 | 518.8129543964847 | 289.0095721692594 |
| SE_66_3_13 | SE_66 | SE_66_3 | AAGGAGGTGGTAGATGGCCA | 2175.140121268416 | 2291.606967265524 | 2359.2016748339256 | 1980.5915211200615 | 2064.9775162841547 | 1963.9250190716825 | 2041.4766824145363 | 2350.148777522676 | 2433.9676465187904 | 2175.3567233634476 | 2103.4226567386886 | 2088.065141897581 |
| SE_66_3_21 | SE_66 | SE_66_3 | CTCCAGATATTTGCTCCTGA | 13.005322100259587 | 35.97812944682026 | 28.63827901640178 | 26.02616979132801 | 18.030106716399914 | 12.994651648026132 | 11.837766740411123 | 14.180302358342775 | 58.607028336461205 | 6.561062503442034 | 1.173784964697929 | 61.51609367458132 |
| SE_66_3_25 | SE_66 | SE_66_3 | TCGCAACCACTTGTGTAATA | 1867.347498228939 | 1877.858478627091 | 1825.9371690112719 | 1786.6965561746679 | 2007.705412596767 | 1506.5132810611628 | 1826.2445598616068 | 1792.3902180945267 | 1929.592008714094 | 1406.9834035159029 | 1389.761398202348 | 1749.1462861810198 |
| SE_66_3_3 | SE_66 | SE_66_3 | GAGTTTGCAGGTAAGGATGG | 3394.389068167752 | 3589.817804804955 | 3472.144449023057 | 3564.2839529223706 | 3361.02401083949 | 3384.673599255873 | 3668.6315289146833 | 3747.3812365647173 | 3644.291580194497 | 5135.124919360632 | 3800.7157156918943 | 3080.4474077799778 |
| SE_66_3_9 | SE_66 | SE_66_3 | ATAGCCCTGGAGATCAGGC | 102.95879996038839 | 155.9052276028878 | 122.45333096668347 | 106.70729614444484 | 160.14977142214042 | 151.60426922697152 | 67.79811860417279 | 208.92312141291688 | 120.76599778422309 | 184.43875704120384 | 115.03092654039705 | 141.60308355280984 |
| SE_66_4_10 | SE_66 | SE_66_4 | GTTACGCTAGCCTGGAAGCT | 985.1531490946637 | 980.4040274258522 | 1068.5040653705767 | 913.5185596756131 | 1011.807165143854 | 1662.4491008374764 | 994.3724061945343 | 815.8400623499876 | 705.94829587101 | 1154.747000605798 | 1178.4801045567208 | 913.4559570168961 |
| SE_66_4_11 | SE_66 | SE_66_4 | ACGCTAGCCTGGAAGCTTGG | 10188.586088711698 | 9705.100418279766 | 9598.761104807769 | 10100.7564960144 | 9885.8014531508 | 10542.994037098535 | 10288.09545803003 | 10105.828814045617 | 8959.771453255964 | 9587.170331418465 | 11048.837872701606 | 11749.573891845032 |
| SE_66_4_18 | SE_66 | SE_66_4 | TTAAAGTGAGAATTCTATAA | 4.335107366753196 | 22.986027146579612 | 7.900214901076352 | 3.9039254686992013 | 4.242378050917627 | 23.390372966447035 | 7.533124289352532 | 29.305958207241734 | 2.6639558334755096 | 6.561062503442034 | 9.390279717583432 | 0.0 |
| SE_66_4_8 | SE_66 | SE_66_4 | GTAACATGCTCAAATGAAGT | 1509.7011404718003 | 1498.0893344662104 | 1430.9264239574543 | 1449.65765737697 | 1586.6493910431925 | 1776.8020353401064 | 1668.0489497852036 | 1637.3522456433125 | 1353.2895634055587 | 1564.4489035985116 | 1514.1826044603285 | 1705.0404076973578 |
| SE_66_4_9 | SE_66 | SE_66_4 | CTTCTGATACATGGGTCAGG | 79.11570944324582 | 76.95322131681 | 70.11440724705263 | 72.87327541571842 | 100.75647870929365 | 41.582885273683615 | 54.88419125099702 | 84.13646065950047 | 114.5501008394469 | 24.057229179287457 | 32.86597901154201 | 27.856344305470785 |
| SE_66_5_22 | SE_66 | SE_66_5 | GCCACAGCCATGGTGGGCGA | 973.2316038360924 | 852.4817893927135 | 872.9737465689369 | 873.1779964990546 | 886.657012641784 | 836.8555661328828 | 902.8987541095393 | 888.6322861128139 | 840.9221085004358 | 611.6368267076929 | 889.7290032410302 | 661.5881772549312 |
| SE_66_5_23 | SE_66 | SE_66_5 | TGCTGGGCCACAGCCATGGT | 974.3153806777807 | 822.5000148536966 | 843.3479406899006 | 1024.1297812887572 | 818.778963827102 | 1030.9090307434064 | 985.763121292417 | 1003.0200534801123 | 873.7775133799671 | 1024.2547574817843 | 1056.4064682281362 | 793.9058127059175 |
| SE_66_5_26 | SE_66 | SE_66_5 | TGGCTGTGGCCCAGCAGCTG | 49.85373471766175 | 80.95079125534559 | 84.92731018657079 | 52.05233958265602 | 39.24199697098805 | 180.19250285262902 | 43.0464245105859 | 25.524544245016994 | 90.57449833816732 | 45.198430579267345 | 16.432989505771005 | 97.49720506914775 |
| SE_66_5_27 | SE_66 | SE_66_5 | GGCTGTGGCCCAGCAGCTGC | 80.19948628493412 | 95.94167852485404 | 124.42838469195254 | 158.75963572710086 | 69.99923784104085 | 30.320853845394307 | 88.2451702467011 | 134.24019565897828 | 84.35860139339113 | 153.82046535847434 | 143.20176569314734 | 34.820430381838484 |
| SE_66_6_12 | SE_66 | SE_66_6 | ATCGCTGCTGTATCACTAGG | 2815.6522347062005 | 2702.357278450055 | 2763.100161651454 | 2702.8177328294137 | 2506.184833579588 | 2408.342105434176 | 2827.073929732729 | 2675.3503782740036 | 2655.963965975083 | 3240.435869755538 | 2795.955785910467 | 2516.3564355941944 |
| SE_66_6_13 | SE_66 | SE_66_6 | AACCCTGTACCCCAGAAAGC | 76.94815575986922 | 114.93013573289807 | 110.60300861506893 | 61.16149900962082 | 121.96836896388177 | 201.85025559933922 | 58.112673089290965 | 126.67736773452879 | 161.6133205641809 | 42.82402799959776 | 106.81443178751154 | 127.67491140007444 |
| SE_66_6_16 | SE_66 | SE_66_6 | CTATTGAGCACCAGGACCAA | 165.81785677830973 | 224.86330904262664 | 216.26838291696515 | 166.56748666449926 | 150.60442080757576 | 170.66309164407653 | 172.1856980423436 | 139.91231660231537 | 138.5257033407265 | 544.5681877856888 | 75.12223774066746 | 459.629681040268 |
| SE_66_6_28 | SE_66 | SE_66_6 | GGTCTGAGCTCTAAGCTGCG | 246.01734306324386 | 263.8396159433486 | 215.2808560543306 | 242.04337905935049 | 158.0285823966816 | 268.5561340592067 | 209.85131948910626 | 242.01049358238336 | 227.32423112324346 | 318.5760348893521 | 268.79675691582577 | 450.3442329384444 |
| SE_66_6_5 | SE_66 | SE_66_6 | CTACGTGGAGGCCACTCAGG | 824.7541765247955 | 777.5273530451713 | 870.0111659810333 | 753.4576154589458 | 817.7183693143726 | 753.6897955855156 | 729.6368954544309 | 858.3809694425016 | 713.0521780936114 | 586.8505905856485 | 746.5272375478829 | 620.964341809453 |
| SE_66_7_11 | SE_66 | SE_66_7 | TTCCTAATCCCGTAAAGCCT | 31.429528408960667 | 26.983597085115196 | 30.613332741670867 | 32.53271223916001 | 37.120807945529236 | 12.994651648026132 | 29.056336544645482 | 20.79777679223607 | 84.35860139339113 | 1.4580138896537853 | 30.518409082146157 | 12.767491140007444 |
| SE_66_7_15 | SE_66 | SE_66_7 | ATTTAGGGCCTAGCACACAG | 108.37768416882989 | 86.94714616314897 | 121.46580410404891 | 132.73346593577284 | 84.84756101835254 | 113.48662439276154 | 178.6426617189315 | 224.99413075237203 | 175.8210850093836 | 168.4006042550122 | 267.6229719511278 | 167.1380658328247 |
| SE_66_7_2 | SE_66 | SE_66_7 | GTACAGCAGTTATTAAGTGT | 1193.238302698817 | 1348.180461771126 | 1355.874382397229 | 1195.902501911522 | 1409.5301074173815 | 1212.8341538157722 | 1150.4156950454083 | 1100.3914630073994 | 1564.6300595279492 | 1080.3882922334549 | 796.9999910298939 | 1534.4202988263492 |
| SE_66_7_6 | SE_66 | SE_66_7 | CAGCAGTGTGAGCTACCTGA | 1998.4844960732232 | 2140.6987020858055 | 2135.0330770158844 | 2044.3556371088815 | 2006.6448180840375 | 2151.914312913127 | 1931.7082999125423 | 2288.700800636524 | 2006.8467278848836 | 2070.379723308375 | 2469.643565724443 | 1738.7001570664684 |
| SE_66_7_7 | SE_66 | SE_66_7 | TGACACAGCGAGGGCACACG | 15144.697585752288 | 12219.57190961865 | 12154.480625305969 | 13648.123438572407 | 11558.358999725075 | 11812.138348055752 | 15329.907928832403 | 10867.783727433904 | 10901.79525585961 | 11854.381929830102 | 10307.005775012514 | 10731.65664368262 |
| SE_67_1_11 | SE_67 | SE_67_1 | AAGCAAAGTCCAGGGTGCCA | 1434.9205383953079 | 1694.969653939088 | 1634.3569576601703 | 1543.351868625751 | 1565.4375007886044 | 1585.347501059188 | 1467.883075810979 | 1807.5158739434257 | 1625.0130584200608 | 1640.2656258605084 | 1968.437385798427 | 1883.7852836574618 |
| SE_67_1_12 | SE_67 | SE_67_1 | GTGCCACCTGAGGTTTTGTC | 320.7979451397365 | 330.79891241381966 | 343.6593481968213 | 325.3271229160001 | 375.45045750621 | 288.4812665861801 | 323.9243444721589 | 305.34917744964775 | 382.7216547426482 | 149.44642368951298 | 399.0868879972959 | 116.06810127279495 |
| SE_67_1_18 | SE_67 | SE_67_1 | GCTCCTCACTCCTCGCTGTG | 169.06918730337463 | 290.8232130284638 | 266.6322529113269 | 192.59365645582727 | 260.90625013143404 | 412.3636122973626 | 145.28168272322742 | 218.37665631847872 | 285.0432741818795 | 102.78997922059186 | 228.88806811609615 | 579.1798253512468 |
| SE_67_1_28 | SE_67 | SE_67_1 | AGGCAACAAGCATCTCTAGG | 1119.5414774640128 | 1311.202939839672 | 1323.2859593022891 | 1207.6142783176197 | 1246.198552457053 | 1055.1657138197218 | 1053.5612398965898 | 880.1240997078082 | 1296.4585056247479 | 1124.1287089230684 | 919.0736273584785 | 1138.6280734861184 |
| SE_67_1_9 | SE_67 | SE_67_1 | CAAAACCTCAGGTGGCACAG | 576.569279778175 | 570.6531087259547 | 613.2541816960519 | 527.0299382743922 | 604.5388722557618 | 670.5240250381484 | 718.8752893267845 | 554.9224989564806 | 544.334975306829 | 478.9575627512685 | 411.9985226089731 | 351.68634685656866 |
| SE_67_2_15 | SE_67 | SE_67_2 | CCCCCAGCATCCTTTAATAA | 55.27261892610324 | 95.94167852485404 | 67.151826659149 | 48.14841411395682 | 108.18064029839948 | 55.44384703157816 | 24.75169409358689 | 33.08737216946648 | 63.93494000341222 | 128.3052222895331 | 36.3873339056358 | 90.53311899278006 |
| SE_67_2_24 | SE_67 | SE_67_2 | ATGACTGCAGAGGTGAACAA | 390.15966300778763 | 128.9216305172765 | 239.9690276201942 | 355.25721765162734 | 100.75647870929365 | 859.3796289846515 | 566.0604823142046 | 216.48594933736638 | 952.8080201064072 | 32.076305572383276 | 185.45802442227279 | 280.88480508016374 |
| SE_67_2_4 | SE_67 | SE_67_2 | CTGGGTCTCTAAGATTCAGT | 134.38832863934907 | 94.94228604022014 | 100.72773998872348 | 140.54131687317124 | 78.4839939419761 | 58.90908747105179 | 159.27177068916782 | 99.26211650839943 | 141.189659174202 | 107.16402088955321 | 78.64359263476125 | 32.49906835638259 |
| SE_67_2_7 | SE_67 | SE_67_2 | CTTAGATCTGAACTAAGAGA | 48.769957875973454 | 45.972054293159225 | 57.276558032803556 | 40.34056317655841 | 41.36318599646866 | 33.78609428486794 | 53.808030638232374 | 55.775855942814914 | 100.34233639424419 | 43.74041668961356 | 71.60088284657367 | 68.48017975094902 |
| SE_67_2_9 | SE_67 | SE_67_2 | TGCACTGGTGACAAGATCCC | 3504.9343060199585 | 3477.8858465259586 | 3496.8326205889202 | 3465.384507715324 | 3394.963035246831 | 3657.5612838644215 | 3525.5021674169852 | 3263.3602493999506 | 3305.081204065282 | 3997.8740854306793 | 3199.7378137665546 | 3372.939022987421 |
| SE_67_3_10 | SE_67 | SE_67_3 | ATAGCTGAGAAGCCCGAAGG | 10.837768416882989 | 19.987849926677923 | 26.66322529113269 | 13.013084895664004 | 10.605945127294067 | 8.663101098684088 | 12.91932735317769 | 0.945353490556185 | 15.983735000853056 | 13.122125006884067 | 0.0 | 20.89225822910309 |
| SE_67_3_12 | SE_67 | SE_67_3 | AGAGACCGCTGAAGCGCCCC | 72.61304839311603 | 97.94046349412183 | 72.08946097232172 | 113.21383859227684 | 49.84794209828212 | 74.50266944868315 | 78.55972473181926 | 101.15282348951179 | 73.70277805948909 | 147.98840979988592 | 111.50957164630326 | 71.96222278913287 |
| SE_67_3_16 | SE_67 | SE_67_3 | ACGTCGGGCGGCCCGAGCCG | 3303.3518134659353 | 4212.439322731872 | 4109.099275422338 | 3578.5983463076013 | 4206.317837484828 | 3696.5452388085 | 3162.836040915299 | 3581.944375717385 | 4124.6916154979135 | 3749.282717244709 | 4048.384343243157 | 3413.5628584328992 |
| SE_67_3_19 | SE_67 | SE_67_3 | GCCGCCCGACGTCTACCCAC | 1243.092037416479 | 1439.1251778728106 | 1291.6851363259836 | 1138.6449283706004 | 1370.2881104463936 | 1400.823447657217 | 1061.0943641859424 | 1210.052467911917 | 1412.784577019845 | 720.9878684337968 | 1162.0471150509497 | 935.5088962587272 |
| SE_67_3_8 | SE_67 | SE_67_3 | TGAGAGATCATGGCCGCCTT | 3166.7959314132095 | 3183.0650635589595 | 3056.395639853914 | 2976.092515638358 | 3171.177593060926 | 3160.299280799955 | 2872.272675468844 | 3371.1305473233556 | 2923.247534600459 | 3504.336383782873 | 3448.5802262825155 | 4253.895911647935 |
| SE_67_4_15 | SE_67 | SE_67_4 | GCAGATGGCAGGAAGGCAGA | 8940.075167086778 | 8271.971595314759 | 8108.583069092241 | 8910.059228061144 | 7711.582702055516 | 8928.191990330282 | 9579.981774830892 | 8310.602535479422 | 7719.256020134201 | 8955.12131025355 | 9380.88943786585 | 10731.65664368262 |
| SE_67_4_19 | SE_67 | SE_67_4 | GAGGACTCTGACCCAGGCAG | 1301.615986867647 | 1594.0310129910645 | 1638.3070651107087 | 1501.709996959626 | 1526.1955038176163 | 1132.26731359801 | 1501.2440548066832 | 1408.5767009287156 | 1655.2045578661164 | 1145.2699103230484 | 1361.5905590495977 | 1537.902341864533 |
| SE_67_4_2 | SE_67 | SE_67_4 | TGCACAAAATGGTCACCAGA | 1553.0522141393324 | 1635.0061048610542 | 1634.3569576601703 | 1687.7971109676214 | 1553.770961148581 | 1972.5881201703667 | 1572.27065524915 | 2105.302223468624 | 1362.1694161838104 | 1741.5975911914466 | 1699.6406288826013 | 1429.9590076808338 |
| SE_67_4_27 | SE_67 | SE_67_4 | GCAGGTGCGACACCTTTGTG | 13064.929826552443 | 10489.623518771375 | 10129.063030042518 | 11934.300157813459 | 10486.097947355645 | 13799.453740093883 | 12648.115681822901 | 11211.892397996355 | 10797.909078354064 | 10314.719262355704 | 11192.039638394754 | 12498.21314505456 |
| SE_67_4_7 | SE_67 | SE_67_4 | TTACAGAGAACAGCTGGGGA | 195.07983150389381 | 222.86452407335884 | 235.0313933070215 | 200.40150739322567 | 181.36166167672854 | 308.4063991131535 | 165.7287343657557 | 142.74837707398393 | 177.59705556503397 | 207.0379723308375 | 187.80559435166865 | 175.26283292192036 |
| SE_68_1_12 | SE_68 | SE_68_1 | GCAGAGAGGACAGGTCCGAG | 7190.8593446401863 | 7260.586400865256 | 7226.721580759593 | 7274.314456766178 | 7556.735903197023 | 7008.448788835427 | 7144.63008144045 | 8152.72850255654 | 7411.125128728867 | 6688.63871878674 | 8633.188415353268 | 8145.659347324749 |
| SE_68_1_15 | SE_68 | SE_68_1 | ACTCTCCTCTTCCGTCCCCT | 96.4561389102586 | 137.91616287947767 | 136.27870704356707 | 149.65047630013606 | 178.17987813854035 | 355.1871450460476 | 135.59623720834557 | 193.79746556401793 | 77.25471917078977 | 227.45016678599052 | 292.2724562097843 | 109.10401519642724 |
| SE_68_1_18 | SE_68 | SE_68_1 | CTTGCCTAGTGAGAAAGCTG | 682.7794102636283 | 693.5783843359239 | 656.7053636519718 | 723.5275201989186 | 758.3250766015258 | 650.598892511175 | 638.1632433694359 | 709.960471407695 | 444.88062419041006 | 540.1941461167274 | 631.4963110074858 | 833.3689671386677 |
| SE_68_1_25 | SE_68 | SE_68_1 | GGGCCAGTGGTGAGAATGGA | 487.6995787597345 | 652.6032924659343 | 716.9445022726679 | 383.88600442208815 | 441.2073172954332 | 319.66843054144283 | 292.7156866719841 | 233.5023121673777 | 448.4325653017107 | 182.98074315155006 | 66.90574298778195 | 51.069964560029774 |
| SE_68_1_33 | SE_68 | SE_68_1 | GGTCCAGAGGAATAGGAGAT | 223.25802938778958 | 289.82382054382987 | 294.28300506509413 | 240.7420705697841 | 360.6021343279983 | 314.4705698822324 | 217.3844437784588 | 407.44735442971574 | 350.75418474094204 | 257.3394515238931 | 446.03828658521303 | 398.11358736568667 |
| SE_68_2_10 | SE_68 | SE_68_2 | CCCCAGGTAACCAAACACCT | 83.45081680999901 | 110.93256579436247 | 138.25376076883617 | 94.99551973834723 | 117.72599091296415 | 59.7753975809202 | 88.2451702467011 | 107.7702979234051 | 122.54196833987343 | 126.84720839987932 | 36.3873339056358 | 320.34795951291403 |
| SE_68_2_13 | SE_68 | SE_68_2 | TCTCAGGCTGAAGGAGAGGC | 1749.2158224849145 | 2216.652530917982 | 2201.1973768123985 | 1738.548142060711 | 1803.0106716399914 | 1608.737874025635 | 1405.4657602706295 | 1206.271053949692 | 2005.0707573292334 | 1119.754667254107 | 1023.5404892165941 | 842.6544152404913 |
| SE_68_2_2 | SE_68 | SE_68_2 | AGGAGAAGCTTATTTGCCAG | 2516.52982640023 | 2945.209652216092 | 2926.0420939861538 | 2739.2543705372727 | 2869.9687514457746 | 2629.2511834506204 | 2850.7494632135513 | 3274.7044912866525 | 3187.8671473923596 | 3276.8862169968825 | 4095.3357418310743 | 3554.005260972981 |
| SE_68_2_20 | SE_68 | SE_68_2 | TAGGGACTCCTAGATGAGTG | 1036.0906606540138 | 1161.2940671445874 | 1175.1569665351074 | 961.66697378957 | 1034.0796499111716 | 697.379638444069 | 717.7991287140198 | 853.6542019722351 | 1189.9002722857274 | 1116.1096325299727 | 1112.7481465336368 | 930.8661722078155 |
| SE_68_2_30 | SE_68 | SE_68_2 | AGCCAAGTAGGGACGCAGAG | 1005.7449090867414 | 1062.3542111658317 | 1126.7681502660148 | 1108.7148331105732 | 1021.3525157584187 | 965.0694623934073 | 1057.8658823476485 | 1148.6044910257647 | 955.4721589398827 | 1477.6970771641113 | 819.3019053591545 | 1525.1348507245254 |
| SE_68_3_10 | SE_68 | SE_68_3 | GGAGGAGATAATTTGCCAGG | 726.1304839311603 | 1031.3730441418308 | 1006.2898730246004 | 956.4617398313043 | 965.1410065837601 | 995.3903612368016 | 996.5247274200635 | 929.2824812167298 | 1163.2607139509723 | 952.8120768887487 | 793.4786361358 | 1058.54108360789 |
| SE_68_3_13 | SE_68 | SE_68_3 | GGTCAACCCGAACAGTCACC | 1019.8340080286893 | 1003.3900545724317 | 888.7741763710897 | 956.4617398313043 | 1098.7759151876653 | 1154.7913764545888 | 1035.266509479591 | 984.1129836689886 | 1009.639260887218 | 1374.1780909986926 | 893.250358135124 | 572.2157392748791 |
| SE_68_3_14 | SE_68 | SE_68_3 | TCTCCTGGATGAGTAGACCT | 67.19416418467453 | 60.962941562667666 | 84.92731018657079 | 106.70729614444484 | 121.96836896388177 | 62.37432791052543 | 83.94052779564251 | 193.79746556401793 | 33.74344055755645 | 51.03408612387484 | 110.33578668160533 | 61.51609367458132 |
| SE_68_3_2 | SE_68 | SE_68_3 | TCACTTTAGGAGAACCTCAG | 94.288585226882 | 106.9349958558269 | 134.303653318298 | 71.57196692615203 | 69.99923784104085 | 258.1604127407858 | 79.63588534458391 | 51.994441980590175 | 95.90241000511834 | 159.6525209170895 | 37.56111887033373 | 171.7807898837365 |
| SE_68_3_3 | SE_68 | SE_68_3 | AAGAGGACAACAGACCACTG | 3742.281434349696 | 4010.5620408358254 | 3857.2799250450529 | 3905.2267771887678 | 3712.0807945529236 | 3820.4275845196826 | 3862.340439212232 | 4527.29786627357 | 3871.6158113177403 | 4100.6640646512715 | 3655.166380069351 | 4152.916663540603 |
| SE_68_4_22 | SE_68 | SE_68_4 | CCTGAGGGCCGGAGATAAGA | 4.335107366753196 | 8.994532361705065 | 10.862795488979984 | 3.9039254686992013 | 5.3029725636470335 | 0.0 | 6.456963676587884 | 21.743130282792254 | 0.0 | 52.48850002753627 | 17.606774470468935 | 37.14179240729438 |
| SE_68_4_23 | SE_68 | SE_68_4 | AAAGGAAGACAAATAGCAGA | 606.9150313454473 | 778.5267455298051 | 817.6722422614025 | 772.9772428024419 | 709.5377290159731 | 842.0534267920933 | 721.0276105523138 | 620.1518898048573 | 800.9627205983031 | 842.003021275061 | 721.8777532892263 | 503.7355595239301 |
| SE_68_4_29 | SE_68 | SE_68_4 | GGGTCACTCTAGCATCAGGG | 167.98541046168634 | 191.88335704970808 | 211.33071486037924 | 225.12636869498726 | 229.08841474955184 | 170.66309164407653 | 260.43086828904467 | 194.7428195045741 | 216.66840778934142 | 326.5951128244793 | 272.3181118099195 | 309.9018303983625 |
| SE_68_4_36 | SE_68 | SE_68_4 | TGTGGACTATTGTGGAGGCA | 2119.8675023423125 | 2126.7072073009313 | 2295.0124287626804 | 2248.66106997074 | 2014.0689796731433 | 1626.9303863328716 | 2113.5794434697677 | 2251.832014504833 | 2061.901815110044 | 1879.3799037637293 | 2637.4948156762466 | 1988.2465748029774 |
| SE_68_4_56 | SE_68 | SE_68_4 | ACATCAGGAAGGGAGCCCCA | 2907.773266249706 | 3039.1525457716784 | 2984.306178881592 | 2743.1582960059723 | 3218.9043461337496 | 2546.951723013122 | 2641.9743043372096 | 2429.5584707293956 | 2646.196127919006 | 2093.7079455428357 | 2013.0412144569482 | 2408.413101410495 |

| | | | | | | | | | | | | | | | |
|---|---|---|---|---|---|---|---|---|---|---|---|---|---|---|---|
| SE_68_5_10 | SE_68 | SE_68_5 | GCATTTCCAGATTGGCAGAA | 253.60378095506195 | 346.78919216796197 | 322.9212840814959 | 385.18731291165454 | 307.572408691528 | 312.73794966249557 | 370.1992507910387 | 438.6440196180698 | 335.6584350179142 | 476.7705419167878 | 291.0986712450864 | 460.79036205299593 |
| SE_68_5_11 | SE_68 | SE_68_5 | TTTCCAGATTGGCAGAAGGG | 14.089098941947885 | 22.986027146579612 | 12.837849214249072 | 9.109159426964803 | 29.69664635642339 | 0.0 | 5.380803063823238 | 20.79777679223607 | 16.871720278678225 | 6.561062503442034 | 0.0 | 1.1606810127279494 |
| SE_68_5_12 | SE_68 | SE_68_5 | CAGAGAAGTGAGGTATGTGT | 13349.963135916467 | 11488.016610866636 | 11244.968384819553 | 12350.718874474707 | 11351.54306974284 | 11006.469945878132 | 12306.972767576508 | 13612.144910518507 | 11169.078824484986 | 11543.824971333845 | 13298.983650027536 | 11134.412955099218 |
| SE_68_5_17 | SE_68 | SE_68_5 | GCAGGCTGATTGTACCCAAC | 14392.556457620609 | 12184.593172656463 | 12205.832022162964 | 13452.927165137447 | 12351.683695246671 | 11348.662439276155 | 13700.600761106727 | 12153.464474590315 | 11221.46995587667 | 12836.354284511926 | 12396.343012174828 | 11692.700522221363 |
| SE_68_5_7 | SE_68 | SE_68_5 | GCTCCATTCAGTTTGTGGCA | 753.2249049733678 | 864.4744992083203 | 809.7720273603261 | 746.9510730111139 | 823.0213418780196 | 1014.4491386559066 | 684.4381497183158 | 689.1626946154589 | 692.6285167036324 | 752.3351670613532 | 692.5331291717781 | 602.3934456058057 |
| SE_68_6_11 | SE_68 | SE_68_6 | GAGGAAGCAGAGCTGCGGCT | 4955.027720198903 | 5277.791711351606 | 5430.410217627357 | 5322.351722326578 | 5395.244286254492 | 6531.978228407802 | 5642.310092725047 | 5538.826101168688 | 5378.526827787054 | 5598.773336270536 | 5149.394640129814 | 5604.928610463267 |
| SE_68_6_2 | SE_68 | SE_68_6 | AGATCCTGGTAAGTCAGGCA | 520.2128840103835 | 694.5777768205578 | 695.218911294719 | 545.2482571283218 | 562.1150917465856 | 660.1283037197275 | 447.6828149100933 | 920.7742998017242 | 630.4695472558706 | 638.610083668358 | 772.3505067712373 | 590.7866354785263 |
| SE_68_6_3 | SE_68 | SE_68_6 | CAGATCCTGGTAAGTCAGGC | 740.2195828731082 | 756.5401108678594 | 800.8842855966152 | 785.9903276981058 | 873.9298784890311 | 772.7486180026206 | 727.4845742289017 | 704.2883504643578 | 750.3475597622685 | 600.7017225373595 | 521.1605243258805 | 798.5485367568292 |
| SE_68_6_4 | SE_68 | SE_68_6 | GCCTGACTTACCAGGATCTG | 1641.921915157773 | 1514.0796142203528 | 1476.3526596386432 | 1553.7623365422821 | 1309.8342232208174 | 1424.2138206236639 | 1589.4892250533844 | 1535.2540686632444 | 1816.8178784302975 | 1599.4412369502024 | 1375.675978625973 | 901.8491468896167 |
| SE_68_6_6 | SE_68 | SE_68_6 | TACCAGGATCTGAGGTCCCG | 548.3910818942793 | 605.6318456881411 | 584.61590267965 | 651.9555532727666 | 518.6307167246799 | 587.3582544907811 | 608.0307462120259 | 477.40351273087344 | 865.7856458795405 | 791.7015420820054 | 467.16641594977574 | 515.3423696512095 |
| SE_68_7_56 | SE_68 | SE_68_7 | GTCGCAGAGACCACTCACCG | 3011.815843051783 | 3318.9824414691693 | 3587.6850919512985 | 3365.1837540187116 | 3644.2027457382414 | 3760.6521869387625 | 3035.8490886090703 | 3093.1966210998376 | 3675.3710649183777 | 3492.6722726656426 | 3000.1943697679067 | 3449.543969827466 |
| SE_68_7_57 | SE_68 | SE_68_7 | CCAGCATGATCCTCGGTGAG | 1828.3315319281603 | 1490.0941945891393 | 1541.5294325725233 | 1754.1638439355079 | 1416.9542690064875 | 1653.7859997387923 | 1706.790731844731 | 1286.6261006469679 | 1419.8884592424465 | 1537.4756466399167 | 1354.5478492614102 | 1290.6772861534798 |
| SE_68_7_58 | SE_68 | SE_68_7 | CACTCACCGAGGATCATGCT | 1146.6358985062202 | 1048.3627163809572 | 1096.1548175243438 | 1185.4920339949908 | 1029.8372718602538 | 1077.6897766763004 | 1244.0416683559324 | 1045.5609605551406 | 1078.9021125575812 | 1720.4563897914666 | 666.7098599484237 | 1064.3444886715297 |
| SE_68_7_59 | SE_68 | SE_68_7 | ACTCACCGAGGATCATGCTG | 1292.9457721341405 | 1048.3627163809572 | 1105.0425592880547 | 1249.2561499837443 | 1129.5331560568181 | 665.3261643789378 | 1131.0448040156446 | 983.1676301784324 | 1034.5028486663227 | 1443.4337507572475 | 904.9882077821032 | 1048.0949544933383 |
| SE_68_7_60 | SE_68 | SE_68_7 | TGGGTCCCCAGCATGATCCT | 22.759313675454276 | 17.98906472341013 | 13.825376076883616 | 31.23140374959361 | 22.272484767317543 | 0.8663101098684087 | 48.42722757440914 | 24.57919075446081 | 4.439926389125849 | 24.057229179287457 | 24.64948425865651 | 3.482043038183848 |
| SE_69_1_1 | SE_69 | SE_69_1 | TGTTGAGTGTTTCCACGCGA | 1049.0959827542733 | 1093.3353781894825 | 1126.7681502660148 | 960.3656653000036 | 1215.4413115879001 | 730.2994226190685 | 952.402142296713 | 1422.7570032870585 | 777.8751033748488 | 821.590826819908 | 1700.8144138472992 | 704.5333747258653 |
| SE_69_1_10 | SE_69 | SE_69_1 | GTAGGGAGGAAGAAAGAGGC | 1766.5562519519272 | 1949.8147375207313 | 1890.1264150825173 | 2014.4255418487878 | 1948.31211988392 | 1865.165666546684 | 1983.3640093252452 | 1907.7233439423812 | 1948.2396995484225 | 2853.3331820524577 | 1944.9616865044684 | 2193.6871140558246 |
| SE_69_1_16 | SE_69 | SE_69_1 | CAGAGTCCCTGACAACACCA | 1297.280879500894 | 1159.2952821753195 | 1029.9905177278295 | 1333.8412018055606 | 1283.319360402582 | 1687.5720940236602 | 1094.4553431816464 | 1126.8613607429725 | 1305.3383584029996 | 1155.476007550625 | 799.3475609592897 | 2100.8326330375885 |
| SE_69_1_3 | SE_69 | SE_69_1 | TGATACCTGGAGCCCTTCAT | 203.7500462374002 | 269.83597085115196 | 319.9587034935923 | 251.15253848631528 | 356.3597562770807 | 220.04276790657582 | 192.6327496848719 | 270.37109829906893 | 309.90686196098426 | 386.3736807582531 | 390.87039324441037 | 200.79781520193526 |
| SE_69_1_6 | SE_69 | SE_69_1 | GCAGTGGCTTAAACAAGACA | 2919.6948115082773 | 3162.0778213816475 | 2938.879943200403 | 2916.2323251183034 | 2787.242379452881 | 2501.037287190096 | 2799.093753800848 | 3135.7375281748655 | 3035.13367960643 | 2919.672814031705 | 3315.9425252716496 | 2892.41708371805 |
| SE_69_2_11 | SE_69 | SE_69_2 | GCAGGGTGTAGGGAACACCA | 10267.701798154943 | 8519.820931503966 | 8634.934886876454 | 10112.468272420498 | 9132.779349112921 | 8850.224082415663 | 10814.337997671943 | 9782.517920275403 | 8859.429116861718 | 8958.766344977685 | 9370.325373183567 | 8577.432684059546 |
| SE_69_2_12 | SE_69 | SE_69_2 | ACTTGTACAAAGGTGCCTGT | 934.2156375353137 | 928.4356182248896 | 828.5350377503825 | 858.8636031138243 | 918.4748480236663 | 1484.8555283144526 | 921.1934845265382 | 838.5285461233361 | 931.4965564386031 | 1175.1591950609509 | 1063.4491780163237 | 860.0646304314105 |
| SE_69_2_14 | SE_69 | SE_69_2 | GGGGTGGCAGGGGAACAGAG | 374.98678722415144 | 433.73633833111097 | 367.3599929000504 | 464.56713077520493 | 454.9950459609155 | 412.3636122973626 | 446.6066542973287 | 402.72058697693484 | 398.70538974350126 | 247.1333542963166 | 201.8910139280438 | 344.722260780201 |
| SE_69_2_21 | SE_69 | SE_69_2 | TCACCACCCCTGCCAGGAGA | 1339.5481763267373 | 1403.1470484259903 | 1376.6124465125545 | 1342.9503612325252 | 1345.894436653617 | 1826.1817116026057 | 1303.2305020579881 | 1344.292663570895 | 1371.0492689620621 | 1422.2925493572675 | 1407.3681726728169 | 2085.7437798721253 |
| SE_69_2_36 | SE_69 | SE_69_2 | GCTGAGTGTGGCAAGAGCTG | 4888.917332855916 | 4447.296556620838 | 4526.8231383167495 | 4878.605527384435 | 4706.918447493107 | 4130.566603852572 | 5183.8656716873065 | 4637.904224668644 | 4297.848744673822 | 4078.7938563064645 | 3997.9115897611464 | 4213.272076202456 |
| SE_69_3_3 | SE_69 | SE_69_3 | TGCCGGAGCCGGGAAAATGG | 238.43090517142576 | 324.8025575060163 | 306.13332741670865 | 201.70281588279207 | 292.72408551331625 | 243.43314087302286 | 250.74542277416285 | 153.14726547010198 | 238.86803973497067 | 286.4997293169688 | 140.85419576375148 | 448.0228709129885 |
| SE_69_3_5 | SE_69 | SE_69_3 | CCCTGCCGGAGCCGGGAAAA | 22.759313675454276 | 24.984812115847404 | 37.526020780112674 | 11.711776406097604 | 19.090701229129323 | 42.44919538355203 | 12.913927353175769 | 4.726767452780925 | 60.382998892111544 | 15.309145841364746 | 2.347569929395858 | 88.21175696732416 |
| SE_69_3_7 | SE_69 | SE_69_3 | TCCCGGCTCCGGCAGGGCGA | 275.27931778882794 | 336.795267321623 | 296.25805879036324 | 236.83814510108488 | 346.814405662516 | 299.7432980144694 | 260.43086828904467 | 285.49675414796786 | 228.21221640106864 | 312.0149723859101 | 277.01325166871123 | 305.2591063474507 |
| SE_69_3_8 | SE_69 | SE_69_3 | GCCCTCGCCCTGCCGGAGCC | 114.88034521895969 | 170.89611487239625 | 172.81720096104522 | 102.80337067574564 | 142.1196647057405 | 85.76470087697247 | 110.8445431147587 | 363.9610938641312 | 198.02071695501286 | 188.0837917653383 | 277.01325166871123 | 149.7278506419055 |
| SE_69_3_9 | SE_69 | SE_69_3 | TGCCCTCGCCCTGCCGGAGC | 252.52000411337366 | 234.8572338889656 | 278.4825752629414 | 243.34468754891688 | 304.3906251533397 | 445.2833964723621 | 325.00050505492356 | 353.5622054680132 | 190.9168347324115 | 359.4004237996581 | 369.74226387984766 | 333.1154506529215 |
| SE_69_4_44 | SE_69 | SE_69_4 | ATCATCAGCGACATGAAGGT | 1682.02165830024 | 1607.023115291305 | 1548.442120610965 | 1557.6662620109814 | 1542.1044215085574 | 1355.7753219440597 | 1683.1151983639086 | 1955.9363719607468 | 1672.0762781447947 | 1713.8953272880246 | 2376.914553513306 | 1649.3277190864162 |
| SE_69_4_47 | SE_69 | SE_69_4 | TCATGTCGCTGATGATGGTC | 6637.049378499142 | 5415.707874231083 | 5174.64076020501 | 5827.259416278341 | 5257.366999599669 | 4845.27244449401 | 5880.141588146033 | 4958.37905796719 | 5597.859191409871 | 5454.429961194811 | 5506.225269397985 | 5531.805706661407 |
| SE_69_4_48 | SE_69 | SE_69_4 | CGCTGATGATGGTCTGGAAG | 4741.523682386308 | 4547.235805084228 | 4701.615393003064 | 4641.76738228335 | 4651.767532831178 | 4984.748372182824 | 4497.275200743462 | 4498.937261556885 | 4574.900151355275 | 4410.4920162027 | 4563.675942745548 | 4448.89032178623 |
| SE_69_4_49 | SE_69 | SE_69_4 | AAGGAATGCAGCGCGCTCGG | 3823.5646974763185 | 3854.6568132329376 | 3866.16766721424 | 3991.11313750015 | 4580.707700478308 | 3905.3259752867866 | 3877.406687791025 | 4342.953935615114 | 3814.784753536929 | 4027.763370168582 | 5478.0544302452345 | 4696.115377497284 |
| SE_69_4_50 | SE_69 | SE_69_4 | GAGAAGGAATGCAGCGCGCT | 3656.6630638563206 | 3257.0201074218676 | 3336.853268842124 | 3616.336292505027 | 3362.0846053522196 | 2897.807317509827 | 3840.817226957027 | 3407.0539799644907 | 3477.350347963365 | 3900.9161617687028 | 3943.9174813850414 | 2580.1938912942314 |
| SE_7_1_12 | SE_7 | SE_7_1 | GGTCCTGCAGGAATGTAGAA | 34.680858934025565 | 6.9957473924372735 | 16.78795666478725 | 55.95626505135522 | 2.1211890254588135 | 78.83421999802519 | 85.01668840840715 | 32.14201867891029 | 29.303514168230603 | 1.4580138896537853 | 5.868924823489645 | 1.1606810127279494 |
| SE_7_1_17 | SE_7 | SE_7_1 | TCCTTGATACAAGACACACA | 278.5306483138928 | 438.73330075428044 | 381.185368976934 | 329.2310478602993 | 389.2381861716923 | 246.8983813124965 | 262.583189514574 | 301.567763487423 | 600.2780478098148 | 267.5455487514696 | 329.83357508011807 | 237.93960760922963 |
| SE_7_1_20 | SE_7 | SE_7_1 | GCACACGAGTTGCTGAGCAA | 6050.726107145772 | 5601.594876372988 | 5558.7887097698485 | 5814.246331382677 | 5816.3003078080665 | 5038.459598994666 | 5802.65802402698 | 5790.290129656633 | 5594.30725029857 | 5439.849822298273 | 6851.382838941812 | 5609.571334514179 |
| SE_7_1_28 | SE_7 | SE_7_1 | TGCCTTAGACTACTGCCGCA | 959.1425048941445 | 964.4137476717099 | 960.8636373434114 | 897.9028578008163 | 1051.0491621148421 | 1087.219187884853 | 924.4219663648322 | 1200.5989330063549 | 741.4677069840168 | 831.0679171026576 | 1014.1502094990107 | 1174.6091848806848 |
| SE_7_1_32 | SE_7 | SE_7_1 | GCAACTCCTCAACTCCTCCA | 759.7275660234975 | 1048.3627163809572 | 1035.9156789036367 | 899.2041662903827 | 1058.473323703948 | 953.807430965118 | 662.9149374630229 | 722.2500667849254 | 911.9608803264493 | 796.8045906957936 | 1000.0647899226356 | 887.9209747368814 |
| SE_7_2_11 | SE_7 | SE_7_2 | GCATCTATCTCCTAAGAATG | 259.02266516350346 | 361.7800794374704 | 340.6967676089177 | 333.1349733289985 | 339.39024407341014 | 341.32618328815306 | 235.6791741954578 | 452.8243219764126 | 267.2835686253761 | 267.5455487514696 | 578.675987596079 | 479.3612582566431 |
| SE_7_2_16 | SE_7 | SE_7_2 | TTTGCTTTGTGTATTTTGGG | 43.351073667531956 | 50.9690167163287 | 59.25161175807264 | 61.16149900962082 | 81.66577748016432 | 64.10694813026224 | 78.55972473181926 | 33.08737216946648 | 31.07948472388094 | 470.93848635817267 | 43.43004369382337 | 31.338387343654635 |
| SE_7_2_23 | SE_7 | SE_7_2 | GTTCATTGTTGAATCTGCAG | 9.75399157519469 | 3.9975699385355847 | 1.975053725269088 | 9.109159426964803 | 2.1211890254588135 | 1.7326202197368175 | 6.456963676587884 | 1.89070698111237 | 20.423661389978903 | 0.0 | 0.0 | 0.0 |
| SE_7_2_4 | SE_7 | SE_7_2 | TTAAATGGCTATTAGAGGAA | 28.17819788389577 | 56.96537162413208 | 49.376343131727204 | 42.943180155691216 | 30.757240869152795 | 118.68448505197199 | 51.655709412703075 | 34.97807915057884 | 23.975602501279585 | 123.93118062057175 | 25.823269223354437 | 112.5860582346111 |
| SE_7_2_9 | SE_7 | SE_7_2 | TTAACGGTACTGCAGTTCAT | 246.01734306324386 | 36.977521931454156 | 81.96472959866715 | 144.44524234187045 | 24.393673792776355 | 265.9572037296015 | 394.95094488462564 | 70.90151179171387 | 357.8580669635434 | 6.561062503442034 | 50.47275348201095 | 81.24767089095646 |
| SE_7_3_1 | SE_7 | SE_7_3 | AGAATCAAAGGAAGCAACCA | 320.7979451397365 | 332.79769738308744 | 321.93375721886133 | 448.95142890040813 | 416.8136435026569 | 368.1817966940737 | 455.21593919944587 | 560.5946198998178 | 456.4244328021373 | 317.8470279445252 | 514.1178145376929 | 330.7940886274656 |
| SE_7_3_18 | SE_7 | SE_7_3 | CTCCCACCAGACAGAGGCAG | 115.96412206064798 | 157.9040125721556 | 119.49075037877982 | 150.95178478970246 | 139.9984756802817 | 261.62565318025946 | 165.7287343657557 | 161.65544688510764 | 119.87801250639792 | 72.17168753786237 | 95.07658214053225 | 99.81856709460365 |
| SE_7_3_21 | SE_7 | SE_7_3 | GCATCAATGTTCCTGGACCA | 3447.4941334104788 | 3476.8864540413247 | 3552.134124896455 | 3376.8955304248093 | 3401.326602323207 | 3881.9356023203395 | 3672.9361713657418 | 3719.020631848032 | 3251.8020873957717 | 4210.744113320132 | 3761.9808118568626 | 3601.593182494827 |
| SE_7_3_3 | SE_7 | SE_7_3 | CCTAAAGGTAGAAGAATCAA | 3056.250693561003 | 3342.967861100383 | 3223.2876796391515 | 3080.19719480367 | 3350.418065712196 | 3739.8607443019205 | 2820.616966056141 | 3455.2670079828563 | 3452.48676018426 | 2630.257056935429 | 3399.2812577652026 | 4237.646377469743 |
| SE_7_3_7 | SE_7 | SE_7_3 | ACAGGATTTATCATTCTGTA | 296.9548546225939 | 281.8286806667587 | 295.27053192772865 | 275.8773997880769 | 312.875381255175 | 231.30479933486512 | 256.1262258379861 | 275.09786575184984 | 167.8292175089571 | 359.4004237996581 | 210.1075086809293 | 259.9925468510607 |
| SE_7_4_15 | SE_7 | SE_7_4 | TGGAAGTGCCCAGAGATAGG | 140.89098941947887 | 132.9192004563082 | 103.69032057662713 | 137.93869989403845 | 129.39253055298764 | 44.18181560328885 | 110.8445431147587 | 98.31676301784324 | 139.41368861855165 | 126.11820145505243 | 112.68335661100119 | 89.37243798005211 |
| SE_7_4_18 | SE_7 | SE_7_4 | TCAAGGGGCTTATACTGTAA | 822.5866228414188 | 927.4362257402556 | 935.1879389149132 | 940.8460379565075 | 838.9302595689608 | 726.8341821795949 | 766.226356288429 | 1072.97621178127 | 773.4351769857229 | 894.4915213025973 | 1159.699545121554 | 815.9587519477484 |
| SE_7_4_21 | SE_7 | SE_7_4 | GTAAGGGGGCCACAGACAGC | 3370.5459776506095 | 3612.803831951535 | 3598.5478874402784 | 3112.72990704283 | 3531.7797273889246 | 3123.914256185482 | 3291.9753144470565 | 3414.6168078889405 | 2910.8157407109065 | 2679.8295291836575 | 3669.2517996457264 | 3025.895400181764 |
| SE_7_4_30 | SE_7 | SE_7_4 | GCAGTTACTGGAACCTTGAC | 1286.4431110840108 | 1328.192612078448 | 1273.9096527985619 | 1173.7802575888932 | 1262.107470147994 | 1535.1015146868203 | 1154.7203374964668 | 1176.019742251894 | 1176.58049311835 | 1354.4949034883666 | 1314.6391604616806 | 1571.5620912336435 |
| SE_7_4_44 | SE_7 | SE_7_4 | TGATTTAGCATAGGGGGAAA | 85.61837049337561 | 128.9216305177726 | 143.19139508200888 | 78.07850937398402 | 158.0285823966816 | 145.54009845789267 | 101.15909759987686 | 114.38777235729839 | 87.02255722686664 | 72.90069448268926 | 139.68041079905356 | 145.08512659099367 |
| SE_7_5_14 | SE_7 | SE_7_5 | GATGCTTCTGGAAGTCTTGT | 330.5519367149312 | 394.760031430389 | 407.8485942680667 | 387.78992989078733 | 398.78353678625695 | 408.89837185788895 | 398.1794267229196 | 442.42543358029457 | 344.5382877961659 | 347.7363126824278 | 245.32105762186717 | 646.4993240894678 |
| SE_7_5_24 | SE_7 | SE_7_5 | TTTTGCACCGAACCTCACTA | 93.2048083851937 | 96.94107100948793 | 91.8399982250126 | 87.18766880094883 | 43.48437502190568 | 13.86096175789454 | 87.16900963393644 | 86.97252113116902 | 76.3667338929646 | 72.17168753786237 | 63.384388093688166 | 15.088853165463343 |
| SE_7_5_27 | SE_7 | SE_7_5 | TGGAGCTGAAACAGGAGGGA | 399.9136545829823 | 397.7582088842907 | 448.337195636083 | 402.10432327601774 | 321.3601373570102 | 259.0267228506542 | 313.1627383145124 | 273.20715877073746 | 264.6196127919006 | 333.88518073071685 | 197.19587406925208 | 181.06623798556012 |
| SE_7_5_28 | SE_7 | SE_7_5 | CACTAACTCCCACAGCCCCA | 2862.2546388987976 | 2857.263113568309 | 2908.266610458732 | 2724.9399771520425 | 2780.8788123765044 | 2546.951723013122 | 2765.732774805144 | 2869.1478438380213 | 2670.1717304202857 | 2217.6391261634076 | 2994.325444944417 | 2307.4338533031637 |
| SE_7_5_8 | SE_7 | SE_7_5 | AAGGACTCAGACTTGAGCAG | 1294.029548975829 | 1466.1087749579258 | 1442.7767463090688 | 1484.7929865952628 | 1607.8612812977806 | 1794.9945476473429 | 1457.1214696833326 | 1221.396709798591 | 1505.1350459136627 | 1449.9948132606894 | 1617.4756813537463 | 1934.8552482174916 |
| SE_7_6_0 | SE_7 | SE_7_6 | GGGAATTCAGGAAGAAAGGA | 31.429528408960667 | 44.97266180852533 | 50.36386999436174 | 48.14841411395682 | 67.87804881468203 | 5.197860659210452 | 48.42722757440914 | 109.66100490451745 | 44.39926389125849 | 43.74041668961356 | 298.141381033274 | 5.803405063639747 |
| SE_7_6_12 | SE_7 | SE_7_6 | TCTCACAGAGGGCCTGAAAT | 450.8511661423323 | 579.6476410876597 | 564.8653654269592 | 456.75927983780656 | 598.1753051793854 | 613.3475577868334 | 407.86487223780136 | 649.4578480120991 | 735.2518100392406 | 457.0873544064617 | 833.3873249355296 | 566.4123342112393 |
| SE_7_6_18 | SE_7 | SE_7_6 | TCTTTTCAGAAGCTTGTGAG | 6.5026610501297935 | 26.983597085115196 | 25.675698428498144 | 14.314393385230405 | 26.51486281823517 | 0.0 | 1.0761606127646475 | 9.45353490556185 | 15.095749723027886 | 0.7290069448268927 | 1.173784964697929 | 1.1606810127279494 |
| SE_7_6_3 | SE_7 | SE_7_6 | TTAACAGCTAGTGGGAACGC | 1168.3114353399862 | 968.4113176102454 | 936.1754657775477 | 917.4224851443123 | 808.1730186998079 | 1013.5828285460382 | 987.9154425179464 | 849.8727880100104 | 618.9257386441434 | 949.8960491094412 | 791.1310662064042 | 550.162800033048 |
| SE_7_6_4 | SE_7 | SE_7_6 | ATAGCATGTAAGAGACATGG | 603.6637008203825 | 621.6221254422834 | 579.6782683664774 | 586.8901287944466 | 640.5990856885617 | 714.7058406414372 | 552.0703943482641 | 617.3158293331888 | 675.7567964249542 | 490.62167386849876 | 558.7216431962142 | 473.5578531930034 |
| SE_7_7_18 | SE_7 | SE_7_7 | TATTTAACTGGCTTACAGTG | 88.86970101844051 | 152.90705014898612 | 171.82967409841066 | 102.80337067574564 | 78.4839939419761 | 134.27806702960336 | 118.37766740411122 | 86.02716764061283 | 128.75786528464963 | 106.43501394472632 | 123.24742129328254 | 110.26469620915519 |
| SE_7_7_2 | SE_7 | SE_7_7 | GTATGGCTCAAAGGAGTGGG | 10171.245659244685 | 10944.347099225797 | 11213.367525215248 | 10780.039527568062 | 11181.847947706136 | 10785.560867861688 | 10846.622816054882 | 12495.682438171654 | 12087.255601756211 | 13063.804451297916 | 12516.069078574017 | 11713.592780450466 |
| SE_7_7_6 | SE_7 | SE_7_7 | GCCATGCTAACTAAAATCAG | 1894.4419192711464 | 1930.8262803126875 | 1920.739747824188 | 2083.3948917958073 | 1950.433308909379 | 2409.208415544045 | 2032.867397512419 | 2420.104935823834 | 1939.3598467701709 | 2222.7421747771955 | 2191.4565290910336 | 1715.4865368119092 |
| SE_7_7_7 | SE_7 | SE_7_7 | TGGTTTGGCATTGTGACAAG | 0.0 | 0.0 | 2.962580587903632 | 1.3013084895664004 | 0.0 | 6.064170769078861 | 0.0 | 0.0 | 0.8879852778251698 | 0.0 | 0.0 | 10.446129114551544 |
| SE_7_7_9 | SE_7 | SE_7_7 | AAGCCTTCTCTCTCTCGCTG | 148.47742731129696 | 212.87059922701988 | 205.40558742798515 | 210.81197530975686 | 264.0880336696223 | 172.39571186381335 | 139.90087965940418 | 255.24544245016995 | 127.86988000682445 | 87.48083337922712 | 196.02208910455414 | 262.31390887651656 |
| SE_7_8_13 | SE_7 | SE_7_8 | CTTGCCCCCCATGCTCGCCC | 2774.468714722045 | 2582.4301802939876 | 2597.1956487288508 | 2652.066701736324 | 2601.638339725235 | 3175.8928627775863 | 2916.3952605921945 | 2431.4491777105077 | 2421.535852629238 | 2934.252952928243 | 2236.060357749555 | 3130.35669132728 |
| SE_7_8_14 | SE_7 | SE_7_8 | GCTTGCCCCCCATGCTCGCC | 992.7395869864818 | 884.4623489009981 | 931.2378314643751 | 1093.0991312357764 | 797.5670735725139 | 928.6844377789341 | 1151.4918556581729 | 795.9876390483078 | 895.9771453255963 | 987.8044102404396 | 775.871861665331 | 1197.822805135244 |
| SE_7_8_15 | SE_7 | SE_7_8 | CATGGGGGGCAAGCAGAGCA | 14002.396794612821 | 13815.60170757898 | 14195.69865037157 | 14536.91713694626 | 13749.54726302403 | 13422.608842301124 | 14871.463507794664 | 14971.563229938301 | 13900.521539075207 | 14424.131410344899 | 14819.035179311353 | 15059.836140145144 |
| SE_7_8_22 | SE_7 | SE_7_8 | TGTCATCGGTGGAGACCCCC | 448.68361245895574 | 478.70900013963626 | 409.8236479933358 | 510.112927910029 | 518.6307167246799 | 436.620295373678 | 540.232627607853 | 411.2287683919405 | 462.64032974691344 | 710.0527642613935 | 318.0957254331388 | 262.31390887651656 |
| SE_7_8_40 | SE_7 | SE_7_8 | TTCGGGCACTACCGGACGGG | 227.59313675454277 | 258.8426535201791 | 270.5823603618651 | 271.9734743193777 | 300.1482471024221 | 88.3636312065777 | 233.5268529699285 | 155.03797245121433 | 340.09836140704004 | 338.2592223996782 | 235.93077790428373 | 270.4386759656122 |
| SE_7_9_0 | SE_7 | SE_7_9 | AGGCGCAGCTCGGGAAGCCG | 2693.185451595423 | 3188.062025982129 | 3100.8343486724684 | 2845.961666681718 | 3012.088416151515 | 2749.668288722329 | 2943.299275911311 | 2856.858248460791 | 3265.121866563149 | 3497.046314334604 | 2676.2297195112783 | 2713.6722077579457 |
| SE_7_9_12 | SE_7 | SE_7_9 | GGACCCCTCCAGGCCTCGGG | 5.4188842084414945 | 16.989672238776237 | 20.738064115325425 | 7.807850937398403 | 9.545350614564661 | 21.65775274671022 | 9.685445514881827 | 10.398888396118036 | 3.551941111300679 | 6.561062503442034 | 46.95139858791716 | 0.0 |
| SE_7_9_26 | SE_7 | SE_7_9 | GGTCGCGGCGTCCTGAGGCG | 2880.6788452074984 | 2811.29105927515 | 2947.767684964114 | 2601.3156706432346 | 3425.720276115984 | 2918.598760146669 | 2880.881960370961 | 3519.551045340677 | 2684.379494865488 | 3213.4626127969427 | 3341.765794495004 | 2971.3433925835507 |
| SE_7_9_4 | SE_7 | SE_7_9 | CGCCTGGCCGCCCAGCGCCG | 3.2513305250648967 | 1.9987849692677924 | 4.93763431317272 | 0.0 | 4.242378050917627 | 0.0 | 0.0 | 0.0 | 4.439926389125849 | 0.0 | 0.0 | 0.0 |
| SE_7_9_7 | SE_7 | SE_7_9 | TCCAGGCCTCGGGCGGGCCG | 220.00669886272468 | 251.84690612774185 | 260.7070917355196 | 217.31851775758886 | 249.2397104914106 | 198.3850151598656 | 248.59310154863357 | 293.05958207241736 | 241.53199556844618 | 312.0149723859101 | 342.74520969179525 | 176.42351393464833 |
| SE_70_1_10 | SE_70 | SE_70_1 | GCTGCGCGTGGTCATCTGGG | 1944.2956539888082 | 1922.8311404356164 | 1804.211578033312 | 1987.0980635678934 | 1827.4043454327677 | 1876.4276979749734 | 1857.4532176317816 | 1857.6196089429036 | 1891.4086417676117 | 2202.3299803220425 | 1915.6170623870203 | 1548.3484709790846 |

| | | | | | | | | | | | | | | |
|---|---|---|---|---|---|---|---|---|---|---|---|---|---|---|
| SE_70_1_17 | SE_70 | SE_70_1 | TCCCACTTCTCAATACAGGG | 81.28326312662242 | 115.92952821753195 | 82.9522564613017 | 88.48897729051524 | 141.0590701930111 | 16.459892087499767 | 87.16900963393644 | 140.85767009287156 | 69.26285167036325 | 123.20217367574486 | 102.11929192871983 | 307.5804683729066 |
| SE_70_1_19 | SE_70 | SE_70_1 | GCAGTTAGCTAGCTGCACCC | 4583.292263499816 | 4089.5140471219033 | 3979.7332564172125 | 4243.566984476031 | 3947.532776378852 | 4293.432904507834 | 4220.701923262947 | 3904.3099159970443 | 4542.932681353569 | 3175.5542516659443 | 4049.558128207855 | 3688.6442584494234 |
| SE_70_1_21 | SE_70 | SE_70_1 | CTTTTCAAAGAAGGACACTG | 30.34575156727237 | 25.984204600481302 | 31.60085960430541 | 29.93009526002721 | 39.24199697098805 | 12.994651648026132 | 12.913927353175769 | 24.57919075446081 | 79.91867500426528 | 73.62970142751615 | 52.820323411406804 | 32.49906835638259 |
| SE_70_1_3 | SE_70 | SE_70_1 | AGTTCGTGCAAGTAAGGTAG | 619.920353445707 | 519.684092009626 | 484.87568955356113 | 571.2744269196498 | 534.539634415621 | 655.7967531703854 | 506.871648612149 | 453.7696754669688 | 505.26362308252163 | 626.2169656063008 | 615.0633215017148 | 448.0228709129885 |
| SE_70_2_29 | SE_70 | SE_70_2 | GGTTCCAGTGGGCAGAGCGA | 67.19416418467453 | 103.93681840192521 | 73.07698783495626 | 57.25757354092162 | 92.27172260745839 | 77.96790988815678 | 67.79811860417279 | 29.305958207241734 | 101.23032167206935 | 129.7632361791869 | 49.29896851731302 | 109.10401519642724 |
| SE_70_2_30 | SE_70 | SE_70_2 | CTCTGCCCACTGGAACCACT | 61.77527997623304 | 52.9678016855965 | 66.16429979651446 | 71.57196692615203 | 72.12042686559965 | 71.90373911907793 | 53.808030638232374 | 63.33868386726439 | 89.68651306034215 | 18.954180565499207 | 75.12223774066746 | 330.7940886274656 |
| SE_70_2_31 | SE_70 | SE_70_2 | GGCAACCGAGTGGTTCCAGT | 88.86970101844051 | 123.92466809460312 | 122.45333096668347 | 75.47589239485123 | 126.21074701479941 | 136.01068724934018 | 59.18883370205561 | 101.15282348951179 | 120.76599778422309 | 108.622034779207 | 70.42709788187574 | 128.83559241280238 |
| SE_70_2_33 | SE_70 | SE_70_2 | GCAGACACAGGGCAACCGAG | 7610.2809823352345 | 6796.8682879951275 | 6964.039435298804 | 7115.554820949078 | 7191.8913908181075 | 7110.673381799899 | 8067.976113896562 | 7750.007915579605 | 6799.303272307325 | 9153.411199246464 | 7803.322445311832 | 7674.422856157202 |
| SE_70_2_5 | SE_70 | SE_70_2 | TGCCTCTTCGGGTTGAGGGA | 147.39365046960864 | 155.9052276028878 | 157.0167711588925 | 187.38842249756166 | 177.11928362581094 | 149.0053388973663 | 120.52998862964051 | 125.7320142439726 | 82.5826308377408 | 69.98466670338169 | 105.64064682281361 | 73.12290380186082 |
| SE_70_3_29 | SE_70 | SE_70_3 | GGAAACCTCCCACCCGGGCA | 141.97476626116716 | 216.86816916555549 | 193.55526507637063 | 183.48449702886245 | 206.8159299822343 | 286.74864636644327 | 143.1293614976981 | 245.7919075446081 | 206.0125844554394 | 286.4997293169688 | 245.32105762186717 | 600.0720835803498 |
| SE_70_3_33 | SE_70 | SE_70_3 | CCATCTGAAGAATGAGGAGG | 480.1131408679164 | 605.6318456881411 | 609.3040742455137 | 545.2482571283218 | 587.5693600520914 | 665.3261643789378 | 612.3353886630844 | 680.6545132004532 | 633.133503089346 | 912.7166949232696 | 691.3593442070802 | 466.59376711663566 |
| SE_70_3_34 | SE_70 | SE_70_3 | CTGCCATCTGAAGAATGAGG | 261.19021884688004 | 232.85844891969782 | 223.18107095540694 | 203.00412437235846 | 291.6634910005869 | 103.09090307434064 | 167.88105559128502 | 308.1852379213163 | 305.4669355718584 | 30.61829168272949 | 358.0044142328683 | 156.69193671827318 |
| SE_70_3_4 | SE_70 | SE_70_3 | GTGTCTGCATTTGAAGCCAG | 326.21682934817795 | 415.74727360770083 | 427.59913152075757 | 385.18731291165454 | 479.3887197536919 | 563.1015714144656 | 358.3614840506276 | 437.69866612751366 | 412.91315418870397 | 1064.3501394472632 | 302.83652089206566 | 453.8262759766282 |
| SE_70_3_8 | SE_70 | SE_70_3 | GGAGAAGCTGTGGGACCTCA | 2507.8596116667236 | 2957.2023620316986 | 2938.879943200403 | 2879.7956874104443 | 2892.241236213092 | 3130.844737064429 | 3116.5611345664192 | 3166.93419336322 | 2610.676716805999 | 3836.763550623936 | 3995.5640198317506 | 3352.0467647583177 |
| SE_70_4_11 | SE_70 | SE_70_4 | GCACCTGGCAAGATCTACTG | 3302.2680366242466 | 3206.051090705539 | 3267.7263884577064 | 3383.402072872641 | 3385.417684632266 | 3606.4489873821854 | 3573.9293949913945 | 3719.020631848032 | 2978.3026218256196 | 3984.7519604237955 | 4158.720129924763 | 3278.9238609564572 |
| SE_70_4_14 | SE_70 | SE_70_4 | GTATTATTAGTAGAGACGGC | 22.759313675454276 | 30.98116702365078 | 36.53849391747813 | 44.24448864525762 | 48.78734758555271 | 25.989303296052263 | 24.75169409358689 | 7.56282792444948 | 52.391131391685015 | 83.10679171026577 | 38.73490383503166 | 90.53311899278006 |
| SE_70_4_15 | SE_70 | SE_70_4 | GATCTACTGGGGAATTCCCG | 34.680858934025565 | 60.962941562667666 | 73.07698783495626 | 42.943180155691216 | 58.33269820011737 | 43.31550549342044 | 41.970263897821255 | 80.35504669727572 | 91.46248361599248 | 64.88161808959345 | 57.515463270198524 | 31.338387343654635 |
| SE_70_4_39 | SE_70 | SE_70_4 | AGTCTGGACAAATCCGCCCA | 280.6982019972694 | 348.78797713722975 | 363.4098854495122 | 288.8904846837409 | 340.4508385861396 | 295.41174746512735 | 272.2686350294558 | 461.3325033914183 | 325.00261168401215 | 295.2478126548915 | 422.56258729125443 | 218.20803039285448 |
| SE_70_4_7 | SE_70 | SE_70_4 | GAGGTCTCAGTAGAACCAGA | 287.2008630473992 | 399.75699385355847 | 384.14794956483763 | 324.0258139020337 | 462.4192075500213 | 352.58821471644234 | 344.3713960846872 | 583.2831036731661 | 449.3205505795359 | 335.34319462037064 | 473.0353407732654 | 305.2591063474507 |
| SE_70_5_1 | SE_70 | SE_70_5 | TATAGGCAGACTGAGCCCAG | 1575.8115278147866 | 1546.0601737286374 | 1503.0158849297761 | 1486.0942950848294 | 1676.799924625192 | 1518.6416225993205 | 1619.6217222107944 | 1936.083948659067 | 1861.217142321556 | 1795.5441051086366 | 1698.4668439179034 | 1905.838222899293 |
| SE_70_5_11 | SE_70 | SE_70_5 | GTGTGTTGTTGGGTGCTGAG | 718.5440460393422 | 718.5631964517713 | 826.5599840251134 | 728.7327541571842 | 940.7473327909838 | 730.2994226190685 | 784.521086705428 | 921.7196532922803 | 733.4758394835902 | 535.0910975029392 | 909.683347640895 | 396.95290635295873 |
| SE_70_5_18 | SE_70 | SE_70_5 | CCTTTCATCAGGAAACCTTG | 319.71416829804815 | 316.8074176289451 | 343.6593481968213 | 294.0957186420065 | 318.178353818822 | 749.3582450361736 | 316.39122015280634 | 285.49675414796786 | 262.8436422362503 | 844.1900421095417 | 154.93961534012664 | 420.1665266075177 |
| SE_70_5_35 | SE_70 | SE_70_5 | GCACGGGAGGAGGCTTACAG | 8812.189499767559 | 8715.701858492208 | 8154.996831636065 | 8951.701099727268 | 9095.658541167391 | 9629.903181297232 | 9522.945262354366 | 10795.936862151633 | 8648.976606017153 | 13665.235180780102 | 12035.991028012564 | 11245.838332321102 |
| SE_70_5_6 | SE_70 | SE_70_5 | CGATTATCTCAGACACAGAG | 832.3406144166136 | 829.4957622461338 | 919.3875091127605 | 827.6321993642307 | 872.8692839763017 | 1027.4437903039327 | 816.8059050883675 | 830.9657181988866 | 1088.669950613658 | 889.3884726888091 | 663.1885050543299 | 1122.3785393079272 |
| SE_70_6_13 | SE_70 | SE_70_6 | GCCATCCCTAGCCATCACCT | 3359.7082092337264 | 2797.2995644902753 | 2812.4765047831816 | 3142.6600023028573 | 2890.120047187633 | 3275.5185254124535 | 3548.1015402850426 | 2834.1697646874427 | 2733.2186851458728 | 3990.5840159824106 | 2653.9278051820174 | 3116.4285191745444 |
| SE_70_6_27 | SE_70 | SE_70_6 | TTGTGGTAACTGCCAGGAAG | 351.14369670700887 | 437.7339082696465 | 420.6864434823158 | 442.44488645257616 | 395.6017532480687 | 392.43847977038916 | 447.6828149100933 | 437.69866612751366 | 436.0007714121584 | 468.751465523692 | 505.9013197848074 | 472.39717218027545 |
| SE_70_6_34 | SE_70 | SE_70_6 | GGCTGGAAAGAACATCTCCT | 1080.5255111632341 | 1181.2819168372653 | 1242.3087931942564 | 1215.422129255018 | 1272.713415275288 | 1363.5721129328754 | 1145.034891981585 | 1355.6369054575694 | 1158.8207875618466 | 1174.430188116124 | 2331.136939890087 | 1137.4673924733904 |
| SE_70_6_37 | SE_70 | SE_70_6 | CACATCCATTCTCCTCACAA | 2337.7066475216607 | 2371.558366036236 | 2343.401245031773 | 2334.5474302821226 | 2580.4264494706467 | 3203.6147862933753 | 2608.6133253415055 | 1936.083948659067 | 2549.4057326360626 | 2204.5170011565233 | 2194.9778839851274 | 2794.9198786489023 |
| SE_70_6_9 | SE_70 | SE_70_6 | CCCCAAATAACAAAATCCCA | 126.80189047753097 | 197.87971195751143 | 186.64257703792882 | 209.51066682019047 | 211.05830803315195 | 191.45453428091832 | 114.07302495305264 | 200.41493999791123 | 286.81924473752986 | 123.93118062057175 | 127.94256115207426 | 293.6522962201712 |
| SE_70_7_17 | SE_70 | SE_70_7 | ACTAGATACCAGTCTCAGAA | 105.126353643765 | 191.88335704970808 | 173.80472782367974 | 122.32299801924164 | 154.84679885849337 | 151.60426922697152 | 117.30150679134657 | 190.01605160179318 | 182.92496723198497 | 83.10679171026577 | 183.11045449287693 | 127.67491140007444 |
| SE_70_7_19 | SE_70 | SE_70_7 | GGGTTCTAAGAGATATTCAA | 2.167553683376598 | 12.992102300240651 | 1.975053725269088 | 3.9039254686992013 | 12.727134152752882 | 0.0 | 0.0 | 19.852423301679885 | 10.655823333902038 | 0.0 | 1.173784964697929 | 0.0 |
| SE_70_7_3 | SE_70 | SE_70_7 | ATACATGGGATGCTACCTGA | 715.2927155142772 | 656.6008624044698 | 731.7574052121971 | 812.0164974894338 | 710.5983235287025 | 849.8502177809089 | 849.0907234713069 | 557.7585594281492 | 821.3863819882821 | 438.1331738409625 | 667.8836449131217 | 964.525921576926 |
| SE_70_7_33 | SE_70 | SE_70_7 | GGAGAGAGAATCACAACCAA | 1959.4685297724445 | 2422.5273827525643 | 2296.9874824879494 | 2134.1459228888966 | 2330.1261444665065 | 2053.1549603881285 | 1947.850709104012 | 2579.869675727829 | 2395.784279572308 | 2224.2001886668495 | 2780.6965813693937 | 2192.5264330430964 |
| SE_70_7_6 | SE_70 | SE_70_7 | TACTCAGGACCATATAGTGA | 366.31657249064506 | 383.76671409941616 | 388.0980570153758 | 314.9166544750689 | 361.6627288407277 | 351.72190460657396 | 301.3249715741013 | 411.2287683919405 | 325.00261168401215 | 261.71349319285446 | 372.0898338092435 | 167.1380658328247 |
| SE_71_1_10 | SE_71 | SE_71_1 | CTTCTGATGACGGAAGGCAG | 115.96412206064798 | 102.9374259172913 | 116.52816979087619 | 102.80337067574564 | 103.93826224748186 | 173.26202197368175 | 122.6823098551698 | 52.93979547114636 | 113.66211556162173 | 54.67552086201695 | 59.86303319959438 | 583.8225494021585 |
| SE_71_1_32 | SE_71 | SE_71_1 | GCTCACTGCACAGGCAGCAG | 2227.1614096694543 | 2171.6798691094564 | 2063.9311429061972 | 2339.752664240388 | 2241.0362053972362 | 2420.470446972334 | 2392.305042175811 | 2411.596754408828 | 2243.938797064204 | 2828.5469459283436 | 2900.4226477685825 | 2331.8081545704504 |
| SE_71_1_34 | SE_71 | SE_71_1 | TCTTTCTCTGAGTGAACAGA | 29.26197472558407 | 20.98724217731182 | 21.725590977959968 | 20.820935833062407 | 15.9089176909411 | 0.0 | 48.42722757440914 | 29.305958207241734 | 1.7759705556503396 | 2.9160277793075706 | 26.997054188052367 | 5.803405063639747 |
| SE_71_1_46 | SE_71 | SE_71_1 | TTTGCCTCACCACTTCACGT | 524.5479913771367 | 485.70474753207355 | 474.01289406458113 | 597.3005967109779 | 493.17644841917416 | 596.8876656993336 | 515.4809335142661 | 300.62240999686685 | 615.3737975328427 | 567.1674030753225 | 456.6023512674944 | 581.5011873767027 |
| SE_71_1_9 | SE_71 | SE_71_1 | GGTTGTCAGGAAACAACCAT | 110.54523785220648 | 143.91251778728105 | 136.27870704356707 | 110.61122161314404 | 82.72637199289373 | 141.20854790855063 | 125.91079169346375 | 147.47514452676486 | 162.50130584200608 | 381.27063214446486 | 274.6656817393154 | 229.814840520134 |
| SE_71_2_10 | SE_71 | SE_71_2 | GCATCTGGAATAGGCCGCAG | 641.595890279473 | 825.4981923075983 | 753.4829961901571 | 759.9641579067778 | 721.2042686559965 | 1041.3047520618272 | 729.6368954544309 | 872.5612717833587 | 667.7649289245277 | 936.7739241025571 | 859.2105941588841 | 914.6166380296241 |
| SE_71_2_17 | SE_71 | SE_71_2 | CCAACTACCAGCCTCTCCCA | 241.68223569649066 | 359.78129446820265 | 377.23526152639585 | 324.0258139020337 | 389.2381861716923 | 489.4652120756509 | 269.0401531911619 | 381.92281018469873 | 384.49762529829854 | 427.19806966855907 | 505.9013197848074 | 355.1683898947525 |
| SE_71_2_24 | SE_71 | SE_71_2 | GATCAGATAGAAGGAAACCT | 243.84978937986725 | 293.8213904823655 | 261.6946185981542 | 205.60674135149128 | 169.69512203670507 | 177.5935725230238 | 166.80489497852037 | 138.96696311175918 | 290.3711858488305 | 246.40434735148972 | 198.36965903395 | 213.5653063419427 |
| SE_71_2_43 | SE_71 | SE_71_2 | GCAAGGGGTATAGGCCCAGG | 1853.258399286991 | 2221.6494933411514 | 2217.985333477186 | 1945.4561919017688 | 2246.3391779608833 | 2040.1603087401027 | 1889.738036014721 | 2342.5859495982263 | 2100.9731673343517 | 2270.8566331357706 | 2448.51543635988 | 1961.5509115102345 |
| SE_71_2_9 | SE_71 | SE_71_2 | GGAAGCACGTTTAGCCCCTG | 1930.2065550468603 | 1873.8609086888554 | 1845.6877062639628 | 1819.2292684138279 | 1882.555260094697 | 1333.251259087481 | 1901.5758027551321 | 1556.9971989460366 | 1651.6526167548159 | 1880.837917653383 | 1574.045637659923 | 1764.235139346483 |
| SE_71_3_2 | SE_71 | SE_71_3 | TCCTGCGAGGAGGACAACAG | 521.2966608520718 | 423.742413484772 | 374.2726809384922 | 400.80301478645134 | 381.81402458258646 | 223.50800834604945 | 530.5471820929712 | 478.3488662214296 | 583.4063275311365 | 215.7860556687602 | 585.7186973842665 | 270.4386759656122 |
| SE_71_3_22 | SE_71 | SE_71_3 | TCCAAGGTCACACAGTGAGA | 621.0041302873952 | 711.5674490593341 | 728.7948246242935 | 594.697979731845 | 556.8121191829385 | 1077.6897766763004 | 588.6598551822622 | 479.2942197119858 | 1012.3032167206935 | 595.5986739235713 | 680.7952795247988 | 1022.5599722133235 |
| SE_71_3_3 | SE_71 | SE_71_3 | TGGGCAGCAAATGAAGGCAT | 1627.832816215825 | 1363.1713490406344 | 1384.5126614136307 | 1483.4916781056966 | 1354.3791927554523 | 1356.641632053928 | 1549.6712823810924 | 1423.7023567776146 | 1371.9372542398874 | 1632.9755564122395 | 1307.5964506734929 | 1150.2348836133979 |
| SE_71_3_31 | SE_71 | SE_71_3 | CACCGATGTTTCCTACCTTG | 4573.538271924622 | 4224.432032547479 | 4304.629594223978 | 4540.265320097171 | 4451.31516992532 | 4821.015761417694 | 4435.934045815877 | 4478.139484764648 | 3842.3122971495095 | 3518.9165226794107 | 4521.419684016422 | 4531.298673689915 |
| SE_71_3_4 | SE_71 | SE_71_3 | GAATCTGTCCACAGAGAAAG | 42.26729682584366 | 70.95686640900662 | 64.18924607124536 | 53.35364807222242 | 60.45388722557618 | 30.320853845394307 | 54.88419125099702 | 18.9070698111237 | 50.615160836034676 | 58.320555586151414 | 53.994108376104734 | 47.58792152184593 |
| SE_72_1_17 | SE_72 | SE_72_1 | TATAGCCTGGATCACACGGA | 697.9522860472645 | 686.5826369434867 | 625.1045040476664 | 611.6149900962082 | 622.5689789721617 | 864.5774896486719 | 578.9744096673803 | 625.8240107481945 | 659.7730614241011 | 1007.4875977507656 | 537.5935138316515 | 799.7092177695572 |
| SE_72_1_22 | SE_72 | SE_72_1 | CATAGTCTGGGAGCTGTGGG | 712.0413849892124 | 778.5267455298051 | 821.6223497119406 | 823.7282738955315 | 1007.5647870929364 | 866.3101098684087 | 827.5675112160139 | 1214.7792353646978 | 802.7386911539535 | 1171.5141603368165 | 1491.8806901310677 | 1137.4673924733904 |
| SE_72_1_23 | SE_72 | SE_72_1 | ATGTCAGCAGAGGAACTGTG | 3191.7227987720403 | 2900.2369904075667 | 2828.276934585334 | 3266.284308811665 | 3146.78391926815 | 3930.4489684729706 | 3188.6638956216507 | 3408.9446869456033 | 3422.2952607382044 | 3614.4164324517337 | 3407.497752518088 | 3914.9770559313733 |
| SE_72_1_3 | SE_72 | SE_72_1 | GTAGATAACCCGCAGCATTG | 568.9828418863569 | 666.5947872508087 | 578.6907415038428 | 653.2568617623331 | 741.3555643978553 | 495.5293828447298 | 691.9712740076683 | 778.0259227277403 | 559.430725029857 | 1022.7967435921304 | 764.1340120183518 | 1033.006101327875 |
| SE_72_1_6 | SE_72 | SE_72_1 | GCAGCATTGAGGTTTCCTTT | 896.2834480762232 | 1292.2144826316278 | 1321.31094220502 | 1099.6056736836083 | 1145.4420737477592 | 1206.7699830466934 | 944.8690180073605 | 1089.9925746112813 | 1612.5812645305084 | 1806.47920928104 | 1304.0750957793991 | 1317.3729494462227 |
| SE_72_2_0 | SE_72 | SE_72_2 | ATAGTCACTCAAAAGCTCTG | 125.71811363584267 | 182.888824688003 | 193.55526507637063 | 136.63739140447206 | 146.36204275665813 | 168.0641613144713 | 159.27177068916782 | 233.5023121673777 | 190.9168347324115 | 406.0568682685792 | 230.0618530807941 | 284.3668481183476 |
| SE_72_2_1 | SE_72 | SE_72_2 | AGAGACAGTAAAAAGATTGG | 574.4017260947984 | 644.608152588863 | 644.8550413003572 | 553.0561080657202 | 789.0823174706786 | 831.6577054736724 | 620.9446735652016 | 742.1024900866053 | 650.0052233680243 | 796.0755837509668 | 611.541966607621 | 1203.6262101988837 |
| SE_72_2_19 | SE_72 | SE_72_2 | GGGGAGCTCAGCTATGCTGA | 37.93218945909046 | 37.97691441608806 | 42.463655093285396 | 81.98243484268323 | 47.72675307282331 | 82.29946043749882 | 40.8941032850566 | 24.57919075446081 | 55.05508722516053 | 67.06863892407412 | 51.646538446708874 | 29.017025318198737 |
| SE_72_2_3 | SE_72 | SE_72_2 | GGTCTTACATCCTTCTCAAC | 134.38832836934907 | 178.8912547494674 | 164.91698605996885 | 182.18318853929605 | 185.60403972764618 | 142.94116812828744 | 175.41417988063753 | 135.18554914953447 | 100.34233639424419 | 171.31663203431978 | 120.8998513638867 | 154.37057469281729 |
| SE_72_2_9 | SE_72 | SE_72_2 | CACAGACTAATGTTTATCAA | 628.5905681792134 | 737.5516536598154 | 609.3040742455137 | 571.2744269196498 | 679.8410826595497 | 742.4277641572263 | 666.1434193013168 | 554.9224989564806 | 789.418911986576 | 659.7512850683379 | 528.2032341140681 | 1160.6810127279493 |
| SE_72_3_0 | SE_72 | SE_72_3 | GGTGGAGGGAGTTAGTTATG | 10251.44514552962 | 11219.180032500119 | 11304.219996577625 | 11152.21375558405 | 11644.267155256157 | 12910.619567368894 | 11086.606632701398 | 12236.655581759258 | 11970.929530361114 | 15134.91318155112 | 12233.186902081816 | 13021.680281794865 |
| SE_72_3_1 | SE_72 | SE_72_3 | TGTCAAGTAAAGGAATAATA | 302.3737388310354 | 383.76671409941616 | 401.9234330922594 | 342.24413275596334 | 452.8738569354567 | 383.7753786717051 | 261.50702890180935 | 431.0811916936204 | 326.77858223966246 | 291.60277793075704 | 586.8924823489646 | 240.26096963468552 |
| SE_72_3_13 | SE_72 | SE_72_3 | TACCTCTCTGTCTTCAGGCA | 309.9601767228535 | 365.777649376006 | 374.2726809384922 | 441.14357796300976 | 361.6627288407277 | 387.2406191111787 | 392.79862365909634 | 328.0376612229962 | 347.2022436296414 | 571.5414447442838 | 334.5287149389098 | 489.80738737119464 |
| SE_72_3_6 | SE_72 | SE_72_3 | GATAGGCACCAAAACTCTGA | 1083.7768416882989 | 1241.245465915299 | 1257.1216961337745 | 1198.5051188906548 | 1193.1688268205826 | 1361.8394927131385 | 1175.167389138995 | 1319.7134728164342 | 1355.9535192390342 | 1229.105708978141 | 1332.2459349321496 | 1691.1122355446223 |
| SE_72_3_8 | SE_72 | SE_72_3 | TTATATTAGAATCATCCTGT | 10.837768416882989 | 5.9963549078033775 | 8.887741763710896 | 13.013084895664004 | 15.9089176909411 | 0.0 | 26.904015319116187 | 17.01636283001133 | 5.327911666951019 | 13.851131951710961 | 26.997054188052367 | 0.0 |
| SE_73_1_1 | SE_73 | SE_73_1 | ACTTAAAGTGTACAAATCAA | 383.6570019576578 | 507.69138219401924 | 515.489022295232 | 424.22656759864657 | 520.7519057501387 | 414.0962325170994 | 386.3416599825084 | 364.90644735468743 | 531.0151961394515 | 645.9001531166269 | 239.45213279837753 | 429.4519747093413 |
| SE_73_1_2 | SE_73 | SE_73_1 | TTAATATACAACAGAGTAGA | 10.837768416882989 | 10.993317330972857 | 23.700644703229056 | 18.218318853929606 | 11.666539640023474 | 37.251334724341575 | 4.30464245105859 | 8.508181415005666 | 0.0 | 0.0 | 0.0 | 0.0 |
| SE_73_1_25 | SE_73 | SE_73_1 | GCCCATTCTTGGCAGCAAGG | 269.8604335803864 | 237.85541134286728 | 266.6322529113269 | 275.8773997880769 | 244.99733244049295 | 315.33687999210076 | 290.56336544645484 | 263.7536238561756 | 209.56452556674006 | 149.44642368951298 | 181.936669528179 | 212.40462532921475 |
| SE_73_1_3 | SE_73 | SE_73_1 | TTTAATATACAACAGAGTAG | 0.0 | 0.0 | 4.93763431317272 | 2.602616979132801 | 3.1817835381882205 | 0.0 | 0.0 | 6.617474433893295 | 0.0 | 0.0 | 3.521354894093787 | 0.0 |
| SE_73_2_12 | SE_73 | SE_73_2 | TGCTTAACTGGTCTCCCCAG | 1043.6770985458318 | 1268.2290630004143 | 1278.8472871117344 | 1129.5357689436355 | 1159.2298024132415 | 1231.8929762328771 | 871.6900963393645 | 1171.2929747991132 | 1615.245220363984 | 645.1711461718 | 1189.044169239002 | 1027.2026962642353 |
| SE_73_2_13 | SE_73 | SE_73_2 | ATAGGTATAGCAGAGGTTGG | 3558.039371262685 | 3931.6100345497475 | 3811.85368976934 | 3616.336292505027 | 3985.7141788371105 | 3151.636179701271 | 3644.955995433861 | 3407.0539799644907 | 3601.6682868588887 | 3581.6111199345237 | 3861.7525338561863 | 2959.736582456271 |
| SE_73_2_17 | SE_73 | SE_73_2 | TTTTTGGATAGGTATAGCAG | 0.0 | 3.9975699385355847 | 1.975053725269088 | 3.9039254686992013 | 6.363567076376441 | 0.0 | 0.0 | 0.0 | 18.647690834328564 | 12.393118062057175 | 1.173784964697929 | 0.0 |
| SE_73_2_3 | SE_73 | SE_73_2 | TGTTTGCTACAGCACCTAAT | 177.73940203688102 | 163.900367479959 | 169.85462037314159 | 214.71590077845607 | 216.36128059679896 | 191.45453428091832 | 217.3844437784588 | 225.93948424292822 | 172.26914389808294 | 121.01515284126418 | 235.93077790428373 | 188.03032406192781 |
| SE_73_2_9 | SE_73 | SE_73_2 | TTTGTCACTTGTTGATTCGA | 23.843090517142578 | 83.94896870924728 | 61.22666548334173 | 33.834020728726415 | 48.78734758555271 | 29.454543735525895 | 36.58946083399801 | 15.12565584889896 | 60.382998892111544 | 35.72134029651774 | 43.43004369382337 | 8.124767089095647 |
| SE_73_3_0 | SE_73 | SE_73_3 | TTTTCTTCTCTCAAGCTTCT | 0.0 | 0.0 | 0.0 | 7.807850937398403 | 0.0 | 0.0 | 2.152321225529295 | 0.0 | 0.0 | 0.0 | 0.0 | 0.0 |
| SE_73_3_1 | SE_73 | SE_73_3 | GTAGTTTAACTGGAGAGTGG | 288.2846398890875 | 388.7636765225856 | 403.8984868175285 | 360.4624516098929 | 293.7846800260457 | 330.0641518598637 | 328.2289868932175 | 516.1630058436771 | 343.6503025183407 | 364.50347241344633 | 488.2945453143385 | 185.70896203647192 |
| SE_73_3_10 | SE_73 | SE_73_3 | GGAAGTTTAACTCCAACACT | 343.5572588151908 | 454.72358050842274 | 404.88601368016305 | 395.5977808281857 | 337.2690550479513 | 390.70585955065235 | 331.4574687315114 | 438.6440196180698 | 526.5752697503257 | 277.0226390342192 | 301.66273592736775 | 710.336779789505 |
| SE_73_3_2 | SE_73 | SE_73_3 | AGTTGACTCCAAGGAAAGGA | 54.188842084414944 | 57.964764108765976 | 49.376343131727204 | 68.96934994701923 | 31.8178353818822 | 10.395721318420904 | 59.18883370205561 | 16.071009339455145 | 20.423661389978903 | 11.664111117230282 | 43.43004369382337 | 75.44426582731671 |
| SE_73_3_3 | SE_73 | SE_73_3 | ACTGAAGACAGGCTTAAGCA | 1553.0522141393324 | 1538.065033851566 | 1516.8412610066596 | 1486.0942950848294 | 1595.1341471450278 | 1412.0854790855062 | 1510.929500321565 | 1444.5001335698507 | 1808.8260109298708 | 1465.3039591020543 | 1439.060366719661 | 1542.5450659154449 |
| SE_73_4_12 | SE_73 | SE_73_4 | TGTCTCTGTTCCTCTCATCG | 86.70214733506391 | 107.93438834046079 | 177.75483527421792 | 123.62430650880805 | 138.9378811675523 | 99.625662634867 | 116.22534617858193 | 33.08737216946648 | 164.2772763976564 | 128.3052222895331 | 111.50957164630326 | 12.767491140007444 |
| SE_73_4_15 | SE_73 | SE_73_4 | GTTGATAGTCAGTTGCTTAA | 61.77527997623304 | 90.94471610168455 | 82.9522564613017 | 68.96934994701923 | 63.6356707637644 | 110.02138395328791 | 66.72195799140815 | 35.92343264113503 | 196.24474639936253 | 53.21750697236316 | 11.73784964697929 | 78.92630886550056 |
| SE_73_4_4 | SE_73 | SE_73_4 | ATATTGGTCAATATTTACCT | 24.926867358830876 | 48.970231747060915 | 38.51354764274722 | 65.06542447832003 | 45.60556404736449 | 19.9251325269734 | 55.960351863761666 | 18.9070698111237 | 19.535676112153734 | 238.3852709583939 | 4.695139858791716 | 20.89225822910309 |

| | | | | | | | | | | | | | | | |
|---|---|---|---|---|---|---|---|---|---|---|---|---|---|---|---|
| SE_73_4_5 | SE_73 | SE_73_4 | GGGAGGGTCACATATCAAGG | 1208.4111784824534 | 1465.1093824732918 | 1397.3505106278799 | 1331.2385848264275 | 1316.1977902971937 | 1424.2138206236639 | 1150.4156950454083 | 933.0638951789546 | 1714.699571480403 | 1530.1855771916478 | 954.2871762994163 | 1185.0553139952365 |
| SE_74_1_30 | SE_74 | SE_74_1 | TCGCTCCATGGCCGCCCGCG | 679.5280797385634 | 662.5972173122732 | 773.2335334428479 | 612.9162985857746 | 761.506860139714 | 693.048087894727 | 691.9712740076683 | 765.7363273505099 | 846.2499697673868 | 484.06061136505673 | 723.0515382539243 | 862.3859924568665 |
| SE_74_1_32 | SE_74 | SE_74_1 | GCGGCCATGGAGCGAGCCTA | 2418.989910648283 | 2366.561403613066 | 2360.1892016965603 | 2486.8005235613914 | 2593.1535836233993 | 2491.5078759815433 | 2360.020223792872 | 2877.656025253027 | 2696.8112887550406 | 3002.779605741971 | 3184.4786092254813 | 2581.3545723069597 |
| SE_74_1_33 | SE_74 | SE_74_1 | ATCCCAGCGCTCGCTCGGCG | 249.26867358830876 | 243.85176625067066 | 247.86924252127056 | 249.85122999674888 | 197.27057936766965 | 380.31013823223145 | 231.3745317443992 | 404.61129395804716 | 248.63587779104753 | 454.90033357198104 | 192.50073421046037 | 280.88480508016374 |
| SE_74_1_34 | SE_74 | SE_74_1 | CGGGCCCTAGGCTCGCTCCA | 109.46146101051819 | 131.9198079716743 | 144.17892194464343 | 88.48897729051524 | 126.21074701479941 | 232.17110944473353 | 77.48356411905462 | 198.52423301679886 | 226.43624584541828 | 94.77090282749604 | 151.41826044603283 | 347.04362280565687 |
| SE_74_1_39 | SE_74 | SE_74_1 | CTAGCGTAGCGCCTGCCCGA | 233.01202096298425 | 199.87849692677923 | 229.10623213121423 | 260.2616979132801 | 252.42149402959882 | 259.89303296052265 | 346.5237173102165 | 266.5896843368442 | 205.12459917761421 | 309.09894460660246 | 251.1899824453568 | 142.76376456553777 |
| SE_74_2_1 | SE_74 | SE_74_2 | GCTCTGTCAGTGACTTAAGA | 81.28326312662245 | 106.9349958558269 | 138.25376076883617 | 119.72038104010883 | 143.1802592184699 | 116.08555472236677 | 96.85445514881827 | 160.71009339455145 | 142.07764452202716 | 118.07199061651661 | 109.1620017169074 | 51.069964560029774 |
| SE_74_2_23 | SE_74 | SE_74_2 | TAAGGCTGAGTATAGAGAGG | 2098.1919655085467 | 2343.5753764664864 | 2353.2765136581183 | 2048.2595625775143 | 2479.669970761353 | 2376.288631369045 | 1961.8407970699523 | 1851.0021345090101 | 2259.922532065057 | 1633.7045633570665 | 1520.0515292838181 | 2793.759197636174 |
| SE_74_2_37 | SE_74 | SE_74_2 | TGTGTCCACTCCCAGCAACA | 1264.767574250245 | 1330.1913970477158 | 1345.011586908249 | 1391.0987753464822 | 1202.7141774351473 | 924.3528872295922 | 1329.0583567643396 | 1541.8715430971376 | 1344.409710627307 | 1070.1821950058784 | 1251.2547723679922 | 1034.166782340603 |
| SE_74_2_4 | SE_74 | SE_74_2 | CCCTGGATGTTTTAAAGCTG | 163.65030309493312 | 207.87363680385042 | 239.9690276201942 | 182.18318853929605 | 267.2698172078105 | 178.4598826328922 | 117.30150679134657 | 303.4584704685354 | 232.6521427901945 | 239.8432848480477 | 262.9278320923361 | 661.5881772549312 |
| SE_74_2_9 | SE_74 | SE_74_2 | GTCCTGCTCTCAGAGATACA | 176.65562519519273 | 197.87971195751143 | 196.51784566427426 | 184.78580551842887 | 195.14939034221084 | 245.16576109275968 | 154.96712823810924 | 226.8848377334844 | 215.78042251151626 | 120.28614589643729 | 76.29602270536539 | 299.455701283811 |
| SE_74_3_1 | SE_74 | SE_74_3 | GCAATTCACAAGTAAATGCA | 636.1770060710314 | 635.613620227158 | 633.0047189487427 | 573.8770438987826 | 692.5682168123026 | 773.614928112489 | 584.3552127312036 | 862.1623833872408 | 644.6773117010732 | 500.09876415124836 | 1340.462429685035 | 543.1987139566803 |
| SE_74_3_17 | SE_74 | SE_74_3 | TCAGGCTGCATTAGGTACAG | 8846.870358701584 | 7318.551164974022 | 7531.8673813136675 | 8370.016204891088 | 8142.184074223655 | 7550.7589176130505 | 8557.629192704477 | 8000.526590576994 | 6784.207522584297 | 8873.472532432937 | 8002.86588931048 | 6757.484856102122 |
| SE_74_3_25 | SE_74 | SE_74_3 | TGACGTTAAAGCTTCCCCGA | 2396.230596972829 | 2849.2679736912382 | 2936.9084889475134 | 2577.8921178310393 | 2771.33346176194 | 2577.272576858516 | 2247.023359452584 | 3305.9011564749797 | 2661.291877642034 | 2390.413772087381 | 3256.079492072055 | 2482.696686225084 |
| SE_74_3_37 | SE_74 | SE_74_3 | GTTGAAACCAACAGGAGCTG | 23.843090517142578 | 20.98724217731182 | 31.60085960430541 | 41.641871666124814 | 49.84794209828212 | 42.44919538355203 | 23.675533480822246 | 7.56282792444948 | 0.8879852778251698 | 0.0 | 9.390279717583432 | 1.1606810127279494 |
| SE_74_3_5 | SE_74 | SE_74_3 | TCCATGGCACTAACTTCTGG | 2012.5735950151711 | 1953.812307459267 | 1987.891574483337 | 2294.206867105564 | 1850.7374247128148 | 2790.3848638861446 | 2049.009806703889 | 1712.9805248878072 | 2011.2866542740096 | 2146.196445570372 | 1461.3622810489217 | 2724.1183368724974 |
| SE_75_1_16 | SE_75 | SE_75_1 | TCTCTCCTCTAAGAGTGAAA | 5.4188842084414945 | 8.994532361705065 | 4.93763431317272 | 9.109159426964803 | 6.363567076376441 | 0.0 | 10.761606127646475 | 0.945353490556185 | 15.095749723027886 | 22.599215289633673 | 0.0 | 0.0 |
| SE_75_1_18 | SE_75 | SE_75_1 | GTCTTTTCCAGACTAAACAT | 9.75399157519469 | 13.991494784874547 | 34.56344019220904 | 2.602616979132801 | 24.393673792776355 | 49.379676262499295 | 0.0 | 11.34424188667422 | 5.327911666951019 | 4.374041668961356 | 3.521354894093787 | 51.069964560029774 |
| SE_75_1_29 | SE_75 | SE_75_1 | GACTGGTGAAGACTTCCGAA | 507.2075619101239 | 628.6178728347207 | 536.2270864105574 | 593.3966712422786 | 543.0243905174563 | 414.0962325170994 | 486.42459696962067 | 375.30533575080545 | 439.5521252345904 | 400.95381965479095 | 389.69660827971245 | 297.13433925835506 |
| SE_75_1_34 | SE_75 | SE_75_1 | CGAATGGGATGACTCCCAGG | 988.4044796197286 | 1173.2867769601942 | 1220.5832022162965 | 990.2957605600308 | 1177.2599091296415 | 1091.550738434195 | 964.2399090371241 | 1420.866296305946 | 1087.781965335833 | 1162.7660769988938 | 1584.6097023422042 | 1046.9342734806105 |
| SE_75_1_9 | SE_75 | SE_75_1 | CTGCCAGCCTTACAGACCAG | 223.25802938778958 | 158.9034050567895 | 133.31612645566344 | 163.96486968536647 | 197.27057936766965 | 89.2299413164461 | 146.35784333599204 | 204.19635396013595 | 269.9475244588516 | 187.3547848205114 | 179.58909959878315 | 170.62010887100857 |
| SE_75_2_1 | SE_75 | SE_75_2 | GGTGATATCACGAGGTACAG | 151.72875783636184 | 227.86148649652833 | 210.3432221741579 | 173.07402911233126 | 222.72484767317542 | 177.5935725230238 | 186.175786008284 | 199.46958650735505 | 269.9475244588516 | 201.20591677222237 | 127.94256115207426 | 161.33466076918498 |
| SE_75_2_2 | SE_75 | SE_75_2 | GCACCTGTGGTGATATCACG | 601.4961471370059 | 612.6275930805783 | 562.89031170169 | 654.5581702518995 | 557.8772136956679 | 858.513318879593 | 714.570646875726 | 606.9169409370708 | 797.4107794870025 | 926.5678268948496 | 496.511040067224 | 602.3934456058057 |
| SE_75_2_3 | SE_75 | SE_75_2 | GATTTCCTGGCAGTAAGAAG | 29.26197472558407 | 54.96658665486429 | 26.66322529113269 | 16.917010364363207 | 5.3029725636470335 | 97.02673230526177 | 54.88419125099702 | 55.775855942814914 | 21.311646667804077 | 39.36637502065221 | 42.25625872912545 | 29.017025318198737 |
| SE_75_2_40 | SE_75 | SE_75_2 | GATTATCTCCTCGGACTGTG | 49.85373471766175 | 101.9380334326574 | 82.9522564613017 | 87.18766880094883 | 99.69588419656424 | 68.4384986796043 | 94.70213392328898 | 34.97807915057884 | 95.01442472729316 | 29.160277793075707 | 68.07952795247988 | 37.14179240729438 |
| SE_75_2_8 | SE_75 | SE_75_2 | GTGTGACTACTTAGACAGGA | 306.7088461977886 | 443.7302631774499 | 418.7113897570467 | 312.3140374959361 | 532.4184453901622 | 386.3743090013103 | 365.89460833998015 | 438.6440196180698 | 417.3530805778298 | 525.6140072201896 | 342.74520969179525 | 387.6674582511351 |
| SE_75_3_0 | SE_75 | SE_75_3 | ACTGGGCTGAAACATCCCTT | 2399.4819274978936 | 2276.6160799960153 | 2196.259742499226 | 2281.1937822099 | 2614.365473877988 | 2524.427660156543 | 2549.42449163945 | 2455.0830149744124 | 2609.788731528174 | 2149.1124733496795 | 2453.2105762186716 | 2475.7326001487163 |
| SE_75_3_1 | SE_75 | SE_75_3 | GATCACGTGGAAATGAAATC | 525.631768218825 | 643.6087601042292 | 571.778053465401 | 491.89460905609934 | 506.9641770846564 | 388.1069292210471 | 509.02396983767824 | 480.239573202542 | 577.1904305863603 | 344.82028490312024 | 565.7643529844017 | 427.1306126838854 |
| SE_75_3_10 | SE_75 | SE_75_3 | GAGAAGAAACCAACACCCCT | 1845.671961395173 | 1990.7898293907213 | 1993.8167356591443 | 2024.836009765319 | 2007.705412596767 | 1706.6309164407653 | 2008.1157034188323 | 1737.559715642268 | 2209.3073712290225 | 1813.769278729309 | 1750.113382364612 | 1644.6849950355042 |
| SE_75_3_15 | SE_75 | SE_75_3 | TACTTCTTCAGGTTTTGGCG | 345.72481249856736 | 389.7630690072195 | 397.97332564172126 | 377.3794619742561 | 365.90510689164535 | 505.05879405328227 | 264.7355107401033 | 259.97220990295085 | 420.9050216891305 | 371.79354186171525 | 336.8762848683056 | 508.3782835748419 |
| SE_75_3_18 | SE_75 | SE_75_3 | GCAATAAGCAGATTTGGAGA | 975.399157519469 | 1137.308647513374 | 1218.6081484910274 | 1030.6363237365892 | 1267.410442171641 | 971.1336331624861 | 988.991603130711 | 1204.3803469685797 | 1399.4647978524677 | 1229.105708978141 | 1435.5390118255673 | 1400.941982362635 |
| SE_75_4_11 | SE_75 | SE_75_4 | TCAGGAGCTGAGCCAAGTCA | 6708.57865005057 | 5926.397433879004 | 6163.155149702189 | 6678.315168454767 | 6024.17683230303 | 5320.010384701898 | 6817.477481864042 | 6218.535260878585 | 6498.276263124592 | 7334.538871903367 | 5895.921877677698 | 6282.76632189639 |
| SE_75_4_14 | SE_75 | SE_75_4 | AAGGAGCGTGATGGAGTAAG | 5758.106359889932 | 4928.004341729742 | 4826.043777695017 | 5500.630985397174 | 4502.223706536315 | 4895.5184308663775 | 6079.231301507493 | 4684.226545705897 | 4681.458384694295 | 4752.396273326513 | 4783.173731144061 | 5486.539147165017 |
| SE_75_4_18 | SE_75 | SE_75_4 | GTAAGCGGCAGAGGGCCTGA | 1941.0443234637432 | 2232.642810672124 | 2269.336730334182 | 2115.9276040349673 | 1936.6455802438968 | 2295.721791151283 | 2157.702028593118 | 2150.679191015321 | 2392.232338461007 | 2298.5588970391927 | 2144.5051305031166 | 2358.503817863193 |
| SE_75_4_4 | SE_75 | SE_75_4 | TTATATATGTGTGTGGCCAG | 65.02661050129794 | 96.94107100948793 | 93.81505195028168 | 63.76411598875362 | 143.1802592184699 | 26.85561340592067 | 46.27490634887984 | 86.97252113116902 | 35.51941111300679 | 47.38545141374802 | 285.22974642159676 | 19.73157721637514 |
| SE_75_4_6 | SE_75 | SE_75_4 | TGAGCAGCACCAGCTCTGGT | 3320.692242932948 | 3067.1355353414274 | 2947.767684964114 | 3197.314958864646 | 3067.239330813444 | 3261.657563654559 | 3312.422366089585 | 3525.223166284014 | 2901.935887932655 | 2599.638765252699 | 3792.499220939009 | 3296.3340761473764 |
| SE_75_5_17 | SE_75 | SE_75_5 | TTTGCCAGAGCCTAGATGAG | 208.0851536041534 | 175.89307729556572 | 168.86709351050703 | 231.63291114281927 | 212.11890254588135 | 61.50801780065702 | 242.13613787204568 | 171.10898179066947 | 135.86174750725098 | 144.34337507572474 | 343.9189946564932 | 221.69007343103834 |
| SE_75_5_20 | SE_75 | SE_75_5 | TGACTGCTCTGGCAAATTAA | 449.767389300644 | 562.6579688488836 | 612.2666548334173 | 533.5364807222242 | 488.9340703682565 | 639.3368610828857 | 450.9112967483873 | 548.3050245225874 | 408.4732277995781 | 687.4535489717598 | 619.7584613605065 | 538.5559899057686 |
| SE_75_5_23 | SE_75 | SE_75_5 | GCAAGAACACTGAGACTACG | 6119.004048172135 | 5383.727314722799 | 5546.938387418234 | 5777.809693674818 | 5629.635673576691 | 6153.400710395307 | 5723.022138682395 | 5803.52507852442 | 5543.692089462535 | 4495.785828747447 | 6718.745137930946 | 5526.002301597768 |
| SE_75_5_6 | SE_75 | SE_75_5 | ATAGCCACATTAAATAAGCA | 344.64103565687907 | 540.6713341869379 | 487.83827014146476 | 510.112927910029 | 501.6612045210094 | 578.695153392097 | 354.056841599569 | 501.037349994778 | 602.9420036432903 | 438.1331738409625 | 208.93372371623136 | 567.5730152239672 |
| SE_75_5_7 | SE_75 | SE_75_5 | AGAAATCCCAGCTATTCAGG | 39.01596630077876 | 78.9520062860778 | 79.98967587339807 | 53.35364807222242 | 92.27172260745839 | 18.192512307236584 | 38.74178205952731 | 83.19110716894428 | 49.72717558820951 | 97.68693060680361 | 89.2076573170426 | 49.909283547301825 |
| SE_75_6_1 | SE_75 | SE_75_6 | GCTGCACTAGGAGTTTCACA | 1201.9085174323234 | 1389.1555536411156 | 1300.5728780896945 | 1391.0987753464822 | 1353.318598242723 | 1381.7646252401119 | 1152.5680162709375 | 1582.5217431910537 | 1187.236316452252 | 1112.4645978058381 | 1183.1752444155125 | 1244.2500456443618 |
| SE_75_6_19 | SE_75 | SE_75_6 | CCATTCCATTAGACGTGCAG | 1941.0443234637432 | 1730.9477833859082 | 1994.3780268405899 | 2062.5739559627445 | 1931.3426076820498 | 1542.898305675636 | 2072.685340184711 | 1799.00769252842 | 1643.6607492543892 | 1758.3647509224652 | 1997.7820099158753 | 1860.571663402903 |
| SE_75_6_2 | SE_75 | SE_75_6 | GAGCTGCACGATAGTCAAAA | 908.2049933347945 | 1155.297712236784 | 1028.0154640025603 | 916.1211766547459 | 952.4138724310072 | 1494.384939523005 | 943.7928573945958 | 986.9490441406572 | 1333.7538872934051 | 776.3923962406407 | 998.8910049579376 | 1448.529903884481 |
| SE_75_6_23 | SE_75 | SE_75_6 | CTTCTATCTTATGGAAAGCG | 2.167553683376598 | 12.992102400481302 | 12.837849214249072 | 1.3013084895664004 | 5.3029725636470335 | 0.0 | 16.142409191469714 | 0.945353490556185 | 28.415528890405433 | 26.244250013768134 | 1.173784964697929 | 0.0 |
| SE_75_6_8 | SE_75 | SE_75_6 | TTAGGCTTGAGGCTGCCACC | 1222.5002774244012 | 908.4477685322116 | 969.7513791071223 | 1214.1208207654515 | 884.5358236163253 | 622.0106588855175 | 1118.1308766624688 | 904.703290462269 | 925.2806594938269 | 1598.7122300053757 | 920.2474123231764 | 912.2952760041683 |
| SE_75_7_11 | SE_75 | SE_75_7 | TTCAGCTTACAAATGTGCAA | 46.60240419259685 | 44.97266180852533 | 44.43870881855448 | 50.751031093089615 | 63.6356707637644 | 2.598930329605226 | 62.417315540349556 | 49.15838150892162 | 42.62329333560815 | 62.69459725511277 | 18.780559435166865 | 2.321362025455899 |
| SE_75_7_14 | SE_75 | SE_75_7 | AGATGAGGAACTAAGACCCG | 1659.2623446247856 | 2097.724825246548 | 2033.3178101645262 | 1945.4561919017688 | 2113.7648628996708 | 2155.379553352601 | 1632.5356496839703 | 2227.252823750372 | 2088.5413734447993 | 1996.021014936032 | 2840.5596145689883 | 1732.8967520028284 |
| SE_75_7_23 | SE_75 | SE_75_7 | CTCTGGGGTTTCTTTCACAA | 21.675536833765978 | 39.975699385355846 | 26.66322529113269 | 40.34056317655841 | 51.96913112374093 | 32.05347406513112 | 19.370891029763655 | 27.415251226129364 | 12.431793889552377 | 12.393118062057175 | 28.170839152750297 | 17.410215190919242 |
| SE_75_7_3 | SE_75 | SE_75_7 | GCTTTTGGCCTTGACAACCA | 150.64498099467355 | 160.9021900260573 | 160.9687860943067 | 147.04785932100324 | 170.75571654943448 | 208.7807364782865 | 188.3281072338133 | 86.02716764061283 | 245.97192195757202 | 378.35460435651573 | 167.85124995180385 | 129.99627342553035 |
| SE_75_7_5 | SE_75 | SE_75_7 | ACTGTGGAGGAAGTCAAAGG | 1033.9231069706373 | 1126.3153301824009 | 1087.267075760633 | 1188.0946509741236 | 1240.895579893406 | 1338.4491197466914 | 1133.1971252411738 | 1027.599244234573 | 1347.9616517386078 | 958.6441324473639 | 785.2621413829145 | 1576.2048152845553 |
| SE_76_1_11 | SE_76 | SE_76_1 | CTAACTGTAGGGCAGGCCCC | 346.80858934025565 | 384.7661065840 5 | 385.13547642747216 | 353.95590916206095 | 399.8441312989863 | 331.79677207960054 | 328.2289868932175 | 501.98270348533424 | 402.2573308548019 | 344.09127795829335 | 442.5169316911192 | 334.2761316656494 |
| SE_76_1_17 | SE_76 | SE_76_1 | GCACTGCTCCCTAACAGGTT | 5752.6874756814905 | 5995.3555153187435 | 5838.2588118954245 | 5816.84894836181 | 5534.182167422045 | 6054.641357870309 | 5408.783239755118 | 6311.179902953091 | 5833.1752900335405 | 5251.037023588108 | 6019.16929897098 | 6399.995104181913 |
| SE_76_1_26 | SE_76 | SE_76_1 | GAGGAAACCCTTAACCGTGA | 1684.1892119836166 | 2121.7102448777614 | 2119.2326472137315 | 1780.1900137268358 | 2107.4012967933313 | 1941.4009562151039 | 1781.0458141254915 | 1899.2151625273757 | 2059.2378592765685 | 2325.5321539977876 | 1849.885104363936 | 2140.295787470339 |
| SE_76_1_37 | SE_76 | SE_76_1 | GGTGGGCAACCGCAGCTCCG | 935.299414377002 | 1084.3408587277775 | 1067.5165385079422 | 1064.4703444653155 | 1039.3826224748186 | 886.2356423953821 | 985.7631212924 17 | 922.6650067828366 | 1202.3320661752798 | 584.6635697511679 | 731.2680330068098 | 1024.8813342387793 |
| SE_76_1_43 | SE_76 | SE_76_1 | GCAACAGAGGGCATTCCTGG | 3437.7401418352842 | 4140.483063838232 | 4205.876907960523 | 3724.344897139038 | 3811.776678749488 | 2886.545286081538 | 3118.7134557919485 | 3490.245087133435 | 3806.7928860365028 | 2364.169522073613 | 2982.5875952974375 | 2791.4378356107186 |
| SE_76_2_21 | SE_76 | SE_76_2 | ATACAGCAGAGAAAGAGGGC | 572.2341724114218 | 562.6579688488836 | 640.904933849819 | 690.9948079597586 | 716.9618906050789 | 666.1924744888063 | 591.8883370205762 | 717.5232993321445 | 508.8155641938223 | 520.5109586064013 | 666.7098599484237 | 746.3178911840715 |
| SE_76_2_24 | SE_76 | SE_76_2 | AGCCACGGACAGTCTCCACA | 896.2834480762232 | 1073.3475284968044 | 1032.953098315733 | 1017.6232388409252 | 970.4439791474072 | 745.8930045966999 | 883.5278630797756 | 780.8619831994088 | 1164.1486992287976 | 1071.6402088955322 | 902.6406378527074 | 991.2215848696688 |
| SE_76_2_36 | SE_76 | SE_76_2 | AGCATAGTACGACAGAACCC | 1498.8633720549174 | 1415.139758241597 | 1385.5001882762654 | 1452.2602743561029 | 1555.8921501740397 | 1309.860886121034 | 1459.2737909088619 | 1589.139217624947 | 1411.0086064641948 | 1496.6512577296107 | 1687.902779235622 | 1326.6583975480462 |
| SE_76_2_46 | SE_76 | SE_76_2 | CTGAAATCACAGACTGCAGG | 72.61304839311603 | 25.984204600481302 | 39.50107450538176 | 75.47589239485123 | 20.151259574185873 | 51.978606592104526 | 86.0928490217458 | 71.84686522827006 | 99.45435111641902 | 2.9160277793075706 | 22.30191432926065 | 39.46315443275028 |
| SE_76_2_5 | SE_76 | SE_76_2 | CACTTCCCGTCAGCTCCCTG | 1302.6997637093352 | 1592.0322280217965 | 1492.15308944 0796 | 1302.6097980559668 | 1661.9516014469805 | 1230.1603560131405 | 1214.985331811287 | 1421.8116497965023 | 1546.8703539714459 | 1188.281320067835 | 1424.9749471432858 | 1175.7698658934128 |
| SE_76_3_24 | SE_76 | SE_76_3 | GGATCCCAATCCAACCACAT | 248.18489674662044 | 292.82199799773156 | 301.1956931035359 | 232.9342196323857 | 191.96760840540226 | 552.7058509060447 | 241.05997725928103 | 180.56251669623134 | 261.0676716805999 | 286.4997293169688 | 181.936669528179 | 557.1268861094158 |
| SE_76_3_25 | SE_76 | SE_76_3 | GGGATCCCAATCCAACCACA | 80.19948628493412 | 155.9052276028878 | 150.1040831204507 | 53.35364807222242 | 111.36242383658771 | 63.24063802039384 | 23.675533480822246 | 41.59555358447214 | 78.14270444861494 | 35.72134029651774 | 180.76288456348107 | 271.59935697834015 |
| SE_76_3_26 | SE_76 | SE_76_3 | GTGCTCATATCTGCTAAACG | 300.2061851476588 | 319.80559508284676 | 331.8090258452068 | 314.91665447506089 | 276.81516782237514 | 228.70586900525 99 | 272.2686350294558 | 278.8792797140746 | 192.69280528806183 | 267.5455487514696 | 353.30927437407667 | 135.79967848917008 |
| SE_76_3_41 | SE_76 | SE_76_3 | TTCGCTTCAAGAAAGCCCAT | 50.93751155935005 | 64.96051150120326 | 42.463655093285396 | 33.834020728726415 | 49.84794209828212 | 105.68983340394587 | 41.970263897821255 | 81.30040018783191 | 62.15896944776188 | 7.290069448268927 | 29.344624117448227 | 45.26655949639003 |
| SE_76_3_5 | SE_76 | SE_76_3 | TGACTTTCAGCCCAGCAGGT | 415.0865303666185 | 444.7296556620838 | 432.5367658339303 | 400.80301478645134 | 398.78353678625695 | 671.3903351480168 | 467.053705939857 | 628.6600712198631 | 555.8787839185563 | 537.2781183374199 | 604.4992568194334 | 714.9795038404169 |
| SE_76_4_3 | SE_76 | SE_76_4 | GCGTGCCGCTCCAAGTGTCG | 1705.8647488173824 | 1569.0462008752 17 | 1593.868356292154 | 1452.2602743561029 | 1480.5899397 0752 | 1328.9197856838139 | 1297.8496989941648 | 1532.4180081915758 | 1740.4511445373328 | 838.3579865509265 | 1302.9013108147012 | 1051.5769975315222 |
| SE_76_4_4 | SE_76 | SE_76_4 | CGTGCCGCTCCAAGTGTCGA | 2908.857043091394 | 3059.140395464356 | 3087.996499458219 | 2464.6782792387626 | 2433.0038122012593 | 2018.5025559933924 | 2282.5366596738 17 | 1965.3899068663086 | 2540.5258798578107 | 1205.0484797988536 | 1532.9631638954954 | 1287.1952431152959 |
| SE_76_4_5 | SE_76 | SE_76_4 | CTAACCCTCGACACTTGGAG | 394.4947703745408 | 445.7290481467177 | 454.2623568118 9024 | 387.78992989078733 | 353.17797277889244 | 251.22993186183854 | 318.54354137833565 | 367.74250782635596 | 534.5671372507522 | 115.91210422747594 | 127.94256115207426 | 126.51423038734649 |
| SE_76_4_6 | SE_76 | SE_76_4 | CATTTCTAACCCTCGACACT | 257.9388883218151 | 318.8062025982129 | 321.93375721886133 | 329.2310478602993 | 324.5419208951985 | 250.36362175197013 | 295.9441685102781 | 340.32725660022 66 | 393.37747807655023 | 287.9577432066226 | 433.1266519735358 | 308.74114938563457 |
| SE_76_4_9 | SE_76 | SE_76_4 | AAAGTTTTGATAAGCAAATA | 2.167553683376598 | 0.0 | 6.912688038441808 | 7.807850937398403 | 0.0 | 0.0 | 0.0 | 0.0 | 0.8879852778251698 | 0.0 | 1.173784964697929 | 8.124767089095647 |
| SE_76_5_17 | SE_76 | SE_76_5 | TTTCTCCTGCTGCCAAGTGT | 8.670214733506391 | 4.996962423169481 | 10.862795488979984 | 29.93009526002721 | 5.3029725636470335 | 10.395721318420904 | 16.142409191469714 | 1.89070698111237 | 3.551941111300679 | 11.664111117230282 | 0.0 | 1.1606810127279494 |
| SE_76_5_18 | SE_76 | SE_76_5 | TGTTATAGCCAGAAAGAGGC | 515.8777766436302 | 640.6105826503275 | 695.218911294719 | 554.3574165552866 | 631.053735073997 | 680.9197463565692 | 500.41468493556107 | 566.2667408431548 | 460.86435919126313 | 746.5031115027381 | 438.9955767970255 | 620.964341809453 |
| SE_76_5_30 | SE_76 | SE_76_5 | AGAATACCCCCAGGGAGACA | 170.15296414506292 | 200.87788941141312 | 183.6799964550252 | 165.26617817493286 | 153.7862043457 64 | 121.28341538157723 | 147.43400398487567 | 151.25655848889896 | 167.82921750895 71 | 137.78231257228 27 | 129.1163461167722 | 188.03032406192781 |
| SE_76_5_36 | SE_76 | SE_76_5 | GCTGAGTGCTGGTCGGAAGT | 4506.344107739947 | 4572.220617200075 | 4658.164211047144 | 4416.641013588363 | 4476.7694382308255 | 3618.5773289203435 | 4616.729028760337 | 4600.090085046396 | 4151.331173832668 | 3701.168258886134 | 4905.247367472645 | 4877.181615482844 |
| SE_76_5_42 | SE_76 | SE_76_5 | TGTGGAGCTAGAATCCCCAG | 2104.6946265586766 | 2502.478781523276 | 2489.5552207016854 | 2311.1238774699273 | 2503.0030500414 | 2049.6897199486552 | 2078.0661432485344 | 2639.4269456328684 | 2507.6704245782794 | 3307.5045086796 12 | 2886.3372281922075 | 3602.7538635075553 |
| SE_77_1_1 | SE_77 | SE_77_1 | TTTACAGGCAGCACAGACAC | 1176.9816500734926 | 1044.3651464424215 | 1015.1776147883113 | 1029.3350152470227 | 1086.0487810349125 | 782.2780292111731 | 979.3061576158292 | 891.4683415944825 | 1022.9590400545956 | 890.1174796336 36 | 406.12959778548344 | 1111.9324101933755 |
| SE_77_1_11 | SE_77 | SE_77_1 | TCATCACTGCTAGAAATCAT | 2240.166731769714 | 2193.666503771402 | 2295.0124287626804 | 2316.329111428193 | 2356.6410072847416 | 1994.245872917077 | 2253.4803231291717 | 2477.7714987477607 | 2466.8231017983217 | 2563.9174249561815 | 2724.3549030638933 | 2329.4867925449944 |
| SE_77_1_14 | SE_77 | SE_77_1 | CGTGAAGTCTGGAGGAAACC | 20959.160341410014 | 22003.624334184493 | 21480.684316026603 | 20537.250582336932 | 21055.98286121691 | 22553.5174003141 55 | 20326.52165389866 | 21116.360918553506 | 21443.0684889222 | 18104.158467831054 | 20109.28401520492 | 22988.448138089767 |
| SE_77_1_5 | SE_77 | SE_77_1 | TTGCGTTGTACAGATTCCCA | 626.4230144958368 | 751.5431484446899 | 732.7449320748317 | 666.269946657997 | 760.4462656269847 | 706.9090496526215 | 598.345300697144 | 608.8076479181832 | 731.6998689279399 | 524.1559933305358 | 429.60529707944204 | 623.2857038349089 |
| SE_77_1_6 | SE_77 | SE_77_1 | TGTGAGAAGCACATTGCAGG | 2155.6321381180264 | 2105.719965123619 | 2243.661031905684 | 2123.7354549723655 | 2131.7949705861074 | 2542.6201724637795 | 2221.1955047462325 | 2198.892219033686 | 2479.254895687874 | 2517.2609804872604 | 2676.2297195112783 | 2031.1917722739115 |

| | | | | | | | | | | | | | | | |
|---|---|---|---|---|---|---|---|---|---|---|---|---|---|---|---|
| SE_77_2_14 | SE_77 | SE_77_2 | TGAGTGTGTACCCAGTGCTA | 3067.088461977886 | 2749.3287252278483 | 2838.1522032116795 | 2942.2584949096313 | 2794.666541041987 | 3272.919595082848 | 2989.574182260191 | 2802.027746008532 | 2807.8094484831868 | 2840.211057045574 | 2920.3769921684475 | 2862.239377387123 |
| SE_77_2_15 | SE_77 | SE_77_2 | TATGGGGCAGTTGTGCCTAG | 5862.148936692009 | 5318.766803221595 | 5556.813656044579 | 5930.062786954087 | 5359.184072821692 | 5947.218904246626 | 5713.336693167514 | 4809.013206459314 | 5593.419265020744 | 5662.196940470475 | 5323.114814905108 | 5816.172554779755 |
| SE_77_2_6 | SE_77 | SE_77_2 | TTATCTATACAGCCAAAGAA | 7.586437891818092 | 14.990887269508443 | 14.81290293951816 | 31.23140374959361 | 21.211890254588134 | 3.465240439473635 | 11.837766740411123 | 0.0 | 0.0 | 78.00374309647752 | 1.173784964697929 | 1.1606810127279494 |
| SE_77_3_10 | SE_77 | SE_77_3 | ACCCTCATAGTAGGTTAGCA | 250.35245042999705 | 262.8402234587147 | 294.28300506509413 | 294.0957186420065 | 295.9058690515045 | 320.5347406513112 | 270.1163138039265 | 290.2235216007488 | 242.41998084627136 | 660.4802920131648 | 311.0530156449512 | 330.7940886274656 |
| SE_77_3_11 | SE_77 | SE_77_3 | TCCATGCTAACCTACTATGA | 726.1304839311603 | 818.5024449151609 | 803.8468661845188 | 862.7675285825235 | 798.6276680852433 | 951.2085006355128 | 738.2461803565482 | 781.807336689965 | 798.2987647648276 | 811.3847295923315 | 758.2650871948622 | 516.5030506639375 |
| SE_77_3_12 | SE_77 | SE_77_3 | GTCCATGCTAACCTACTATG | 538.6370903190846 | 573.6512861798564 | 569.8029997401319 | 527.0299382743922 | 571.6604423611502 | 866.3101098684087 | 485.348436356856 | 554.9224989564806 | 542.5590047511787 | 569.3544239098031 | 447.21207154991095 | 571.0550582621511 |
| SE_77_3_8 | SE_77 | SE_77_3 | CTACTATGAGGGTCGAGCAG | 276.36309463051623 | 346.78919216796197 | 307.1208542793432 | 335.7375903081313 | 238.63376536411653 | 272.02137449868036 | 344.3713960846872 | 220.2673632995911 | 278.8273772371033 | 332.42716684106307 | 235.93077790428373 | 249.54641773650914 |
| SE_77_3_9 | SE_77 | SE_77_3 | CTCTGCTCGACCCTCATAGT | 89.95347786012881 | 83.94896870924728 | 83.93978332393624 | 118.41907255054244 | 109.24123481112889 | 73.63635933881474 | 114.07302495305264 | 171.10898179066947 | 79.91867500426528 | 34.26332640686395 | 172.54638981059557 | 89.37243798005211 |
| SE_77_4_1 | SE_77 | SE_77_4 | CTAATTTACTTAAATGCACA | 50.93751155935005 | 68.95808143973883 | 42.463655093285396 | 68.96934994701923 | 38.181402458258646 | 58.90908747105179 | 76.40740350628997 | 22.68848377334844 | 86.13457194904147 | 89.6678542137078 | 25.823269223354437 | 97.49720506914775 |
| SE_77_4_11 | SE_77 | SE_77_4 | ACAACCCACAGATTTGCCCA | 1421.9152162950481 | 1515.0790067049866 | 1512.8911535561215 | 1311.7189574829317 | 1465.7416165920401 | 1245.7539379907716 | 1261.2602381601669 | 1681.7838596994532 | 1372.8252395177126 | 1006.7585908059388 | 1878.0559435166865 | 1254.6961747589132 |
| SE_77_4_12 | SE_77 | SE_77_4 | ATAGATAACAAACCTTGCAA | 298.0386314642822 | 572.6518936952225 | 446.3621419108139 | 417.72002515081454 | 482.5705032918801 | 468.67376943880913 | 373.4277326293327 | 319.5294798079905 | 567.4225925302835 | 572.9994586339376 | 363.873339056358 | 299.455701283811 |
| SE_77_4_4 | SE_77 | SE_77_4 | GGGTTCCTAACACCTCACAG | 56.35639576779154 | 30.98116702365078 | 47.40128940645811 | 40.34056317655841 | 33.939024407341016 | 57.176467251314975 | 66.72195799140815 | 46.32232103725307 | 39.07135222430747 | 8.748083337922711 | 42.25625872912545 | 45.26655949639003 |
| SE_77_4_5 | SE_77 | SE_77_4 | TTTACAAGTGAAGAAACTAG | 20.59175999207768 | 11.992709815606755 | 24.688171565863602 | 19.519627343496005 | 22.272484767317543 | 8.663101098684088 | 16.142409191469714 | 21.743130282792254 | 3.551941111300679 | 0.0 | 14.085419576375148 | 119.5501443109788 |
| SE_78_1_4 | SE_78 | SE_78_1 | AACAGAATAAAGGATCCAGT | 213.50403781259487 | 262.8402234587147 | 269.5948334992305 | 264.1656233819793 | 222.72484767317542 | 75.36897955855156 | 285.1825623826316 | 160.71009339455145 | 229.1002016788938 | 395.1217640961758 | 213.62886357502308 | 203.11917722739116 |
| SE_78_1_5 | SE_78 | SE_78_1 | ATACTAGGTGTACCTTTTGA | 58.52394945116814 | 62.96172653193546 | 60.23913862070719 | 93.69421124878083 | 66.81745430195262 | 40.71657516381521 | 71.02660044246673 | 102.09817698006798 | 60.382998892111544 | 75.08771531716994 | 28.170839152750297 | 33.659749369110536 |
| SE_78_1_6 | SE_78 | SE_78_1 | TCAACTTTCCAACAGAATAA | 57.440172609479845 | 110.93256579436247 | 96.77763253818532 | 102.80337067574564 | 135.75609762936406 | 84.03208065723565 | 131.291594757287 | 123.84130726286024 | 94.126439449468 | 225.99215289633673 | 122.07363632858463 | 46.42724050911798 |
| SE_78_1_8 | SE_78 | SE_78_1 | GAATAAGGTCCCTTTGATAA | 33.59708209233727 | 56.96537162413208 | 47.40128940645811 | 67.66804145745283 | 55.15091466192915 | 113.48662439276154 | 26.904015319116187 | 32.14201867891029 | 82.5826308377408 | 66.33963197924723 | 15.259204541073078 | 67.31949873822107 |
| SE_78_1_9 | SE_78 | SE_78_1 | GTCAATCTACCCTTATCAAA | 52.02128840103835 | 66.95929647047105 | 81.96472959866715 | 72.87327541571842 | 101.81707322202305 | 37.251334724341575 | 45.19874573611519 | 187.17999113012462 | 121.65398306204825 | 56.13353475167074 | 127.94256115207426 | 16.249534178191293 |
| SE_79_1_0 | SE_79 | SE_79_1 | TGTGCAGGCAGAGCCTGGTT | 597.1610397702527 | 528.6786243713311 | 523.3892371963084 | 540.0430231700561 | 505.903582571927 | 583.8930140513074 | 624.1731554034956 | 540.7421965981379 | 535.4551225285774 | 411.88892382719433 | 543.4624386551411 | 566.4123342112393 |
| SE_79_1_1 | SE_79 | SE_79_1 | GTGCAGGCAGAGCCTGGTTC | 211.3364841292183 | 200.87788941141312 | 186.64257703792882 | 192.59365645582727 | 104.99885676021127 | 113.48662439276154 | 195.86123152316583 | 102.09817698006798 | 196.24474639936253 | 46.65644446892113 | 56.341678305500594 | 162.4953417819129 |
| SE_79_1_2 | SE_79 | SE_79_1 | ATATCTTTGTCTCCCGAACC | 63.94283365960963 | 47.97083926242702 | 74.06451469759081 | 68.96934994701923 | 42.42378050917627 | 19.05882241710499 | 75.33124289352533 | 38.75949311280358 | 10.655823333902038 | 127.57621534470621 | 34.03976397623994 | 11.606810127279495 |
| SE_79_1_3 | SE_79 | SE_79_1 | TTCGGGAGACAAAGATATTC | 172.32051782843953 | 184.8876096572708 | 227.13117840594512 | 130.13084895664005 | 154.84679885849337 | 229.5721791151283 | 119.45382801687587 | 140.85767009287156 | 244.1959514019217 | 115.91210422747594 | 225.36671322200237 | 129.99627342553035 |
| SE_79_1_4 | SE_79 | SE_79_1 | GCTGAAGCATCAGAAACTCA | 3045.41292514412 | 3427.916222294264 | 3384.2545582485823 | 3245.463372978603 | 3260.2675321301963 | 3205.347406513112 | 3275.832905255587 | 2870.093197328578 | 3565.2608904680565 | 3816.3513561687832 | 3268.9911266837325 | 3255.7102407018983 |
| SE_79_2_10 | SE_79 | SE_79_2 | AGGAAAGGAGGAAGGCCTGA | 6693.405774266934 | 6593.991613614447 | 6347.822673014849 | 6782.419847620079 | 6254.325841565312 | 6751.154686204509 | 6602.245359311112 | 7640.346910675087 | 6535.57164479325 | 6342.360419993966 | 7332.634674467963 | 7357.556939682471 |
| SE_79_2_15 | SE_79 | SE_79_2 | ATATTTCTGACCCTGTATGT | 18.42420630870108 | 19.987849692677923 | 17.77548352742179 | 15.615701874796805 | 15.9089176909411 | 1.7326202197368175 | 21.52321225529295 | 18.9070698111237 | 31.07948472388094 | 56.13353475167074 | 12.911634611677218 | 1.1606810127279494 |
| SE_79_2_19 | SE_79 | SE_79_2 | TGACAGTTCTAGGCACAGAG | 2109.02973392543 | 2418.529812814029 | 2444.1289850204967 | 2320.233036896892 | 2354.519818259283 | 2229.882222801284 | 2199.6722924909395 | 2723.563406292369 | 2192.435650950344 | 2733.7760431008473 | 2469.643565724443 | 2170.4734938012652 |
| SE_79_2_2 | SE_79 | SE_79_2 | GAGACACTATACATACAAAT | 499.6211240183058 | 690.5802068820223 | 765.3333185417716 | 547.8508741074546 | 783.7793449070316 | 902.6951344828819 | 443.3781724590348 | 650.4032015026553 | 880.8813956025684 | 516.8659238822669 | 442.5169316911192 | 644.1779620640119 |
| SE_79_2_8 | SE_79 | SE_79_2 | TTCAATAATTTCCTTCCTGA | 1.083776841688299 | 1.9987849692677924 | 2.962580587903632 | 0.0 | 15.9089176909411 | 0.0 | 0.0 | 5.67212094433711 | 0.0 | 0.7290069448268927 | 3.521354894093787 | 0.0 |
| SE_79_3_10 | SE_79 | SE_79_3 | GCAGCAATTATCGTCCGAGC | 1833.7504161366016 | 2392.5456082135474 | 2529.0562952070673 | 2264.2767718455366 | 2133.9161596115664 | 2200.427679065758 | 2012.4203458698908 | 2436.175945163289 | 2763.4101845919286 | 2528.9250916044907 | 1863.9705239403113 | 2030.0310912611835 |
| SE_79_3_11 | SE_79 | SE_79_3 | GCTGCTTCTCCTGCTTCCAC | 35.76463577571386 | 19.987849692677923 | 14.81290293951816 | 32.53271223916001 | 16.969512203670508 | 48.51336615263089 | 61.3411549275849 | 29.305958207241734 | 81.69464555991561 | 8.019076393095819 | 23.47569929395858 | 1.1606810127279494 |
| SE_79_3_13 | SE_79 | SE_79_3 | TGCTTCCACTGGTTTATCTG | 29.26197472558407 | 38.97630690072195 | 40.488601368016305 | 22.12224432262881 | 69.99923784014085 | 2.598930329605226 | 35.513300221233365 | 34.03272566002266 | 19.535676112153734 | 59.7785694758052 | 21.128129364562724 | 1.1606810127279494 |
| SE_79_3_14 | SE_79 | SE_79_3 | CAAAGCCACAGATAAACCAG | 144.14231994454374 | 154.90583511825392 | 186.64257703792882 | 126.22692348794084 | 184.54344521491677 | 111.75400417302473 | 117.30150679134657 | 276.043219242406 | 198.02071695501286 | 53.21750697236316 | 220.67157336321065 | 185.70896203647192 |
| SE_79_3_8 | SE_79 | SE_79_3 | TCCTGCAGAACTCACCTGCT | 540.8046440024611 | 158.9034050567895 | 322.9212840814959 | 450.2527373899745 | 114.54420737477592 | 1271.743241286824 | 726.4084136161371 | 416.9008893352776 | 1277.8108147904193 | 58.320555586151414 | 226.5404981867003 | 367.93588103476 |
| SE_8_1_11 | SE_8 | SE_8_1 | GCCCGCGACTTGAAGAACGG | 725.046707089472 | 908.4477685322116 | 968.7638522444877 | 800.3047210833363 | 825.1425309034785 | 649.7325824013066 | 747.93162587143 | 1076.7576257434948 | 874.6654986577922 | 1070.9112019507054 | 1365.1119139436914 | 734.711081056792 |
| SE_8_1_20 | SE_8 | SE_8_1 | CGAATGCGTGGCGGAGATCG | 4515.014322473453 | 4457.290481467177 | 4282.904003246017 | 4546.771862545003 | 4660.252288933013 | 4214.598684509809 | 4758.782229645271 | 5252.383993530164 | 4322.712332452927 | 3696.7942172171724 | 5891.226737818906 | 5011.820612959285 |
| SE_8_1_22 | SE_8 | SE_8_1 | TACGAATGCGTGGCGGAGAT | 19157.92323052406 | 16636.88669170047 | 16703.029354600676 | 18508.510647102914 | 16381.942843618417 | 16323.015090140558 | 19183.639083142607 | 17177.07292340588 | 15990.838883075658 | 17142.59830760438 | 16158.323824031691 | 14654.75846670309 |
| SE_8_1_30 | SE_8 | SE_8_1 | CAACGTGGTCAGGTGAGCCA | 9250.035343809632 | 7351.53111696694 | 7305.723729770357 | 8599.046499054773 | 7322.344515883824 | 6633.336511262406 | 9036.520665384745 | 8300.203647083305 | 7127.857825102637 | 6440.7763575455965 | 7776.32539112378 | 6759.806218127577 |
| SE_8_1_7 | SE_8 | SE_8_1 | CGCGCTTCAACGCCACGAAA | 1392.6532415694642 | 1338.1865369247869 | 1353.89932867196 | 1289.596713160303 | 1238.774390867947 | 1543.7646157855042 | 1466.8069151982145 | 1060.6866164040396 | 1525.5587073036418 | 1966.1317301981294 | 1118.6170713571264 | 1261.660260835281 |
| SE_8_2_18 | SE_8 | SE_8_2 | AATTTACTGGTCCCTTTGAA | 6.5026610501297935 | 12.992102300240651 | 4.93763431317272 | 0.0 | 6.363567076376441 | 15.593581977631358 | 0.0 | 8.508181415005666 | 13.319779167377547 | 1.4580138896537853 | 0.0 | 0.0 |
| SE_8_2_30 | SE_8 | SE_8_2 | TTGCTGAGAGCAGAGAAGAG | 213.50403781259487 | 201.87728189604704 | 177.75483527421792 | 191.29234796626088 | 260.90625013143404 | 126.48127604078768 | 289.48720483369016 | 249.57332150683285 | 158.06137945288023 | 328.0531251721017 | 89.2076573170426 | 175.26283292192036 |
| SE_8_2_6 | SE_8 | SE_8_2 | GCACATACTGTGCTTTGGAG | 473.6104798177866 | 552.6640440025446 | 535.2395595479229 | 527.0299382743922 | 503.7823935464682 | 580.4277736118338 | 484.27227574409136 | 556.813205937593 | 595.8381214206889 | 236.19825012391323 | 462.471276090984 | 376.06064812385563 |
| SE_8_2_8 | SE_8 | SE_8_2 | GGAAGGTCTGGGAGCCAACA | 1238.7569300497257 | 1591.0328355371628 | 1529.6791102209088 | 1214.1208207654515 | 1392.5605952137112 | 1775.0694151203695 | 1238.6608652921093 | 1517.2923523426768 | 1146.3889936722942 | 1033.7318477645338 | 1530.6155939660994 | 1305.766139318943 |
| SE_8_2_9 | SE_8 | SE_8_2 | TCCCCTCCAGACGTTCCCCG | 113.79656837727138 | 142.91312530264716 | 112.57806234033802 | 102.80337067574564 | 96.51410065837601 | 76.23528966841997 | 63.4934761531142 | 110.60635839507364 | 98.56636583859384 | 75.81672226199683 | 237.10456286898167 | 62.67677468730927 |
| SE_8_3_11 | SE_8 | SE_8_3 | ACTGACACTCGCAGCCCCCT | 4260.326764676703 | 4641.178698639814 | 4659.151737909779 | 4931.959175456657 | 4932.825078704471 | 4922.374044272298 | 4455.30493684564 | 4535.806047688576 | 5416.710194733536 | 5148.976051312343 | 4490.901274934276 | 6338.479010507332 |
| SE_8_3_12 | SE_8 | SE_8_3 | AGGGGGCTGCGAGTGTCAGT | 6993.611959414593 | 6649.957592753945 | 6635.192990041502 | 7232.672585010054 | 6628.715704558792 | 6706.106560491352 | 6950.921397846858 | 6679.867764270003 | 6457.428940344635 | 6121.471315711417 | 6984.020539952678 | 5976.346534536212 |
| SE_8_3_17 | SE_8 | SE_8_3 | TTACCTGCTCCGCGAGGCCG | 39.01596630077876 | 48.970231747060915 | 76.03956842285989 | 28.62878677046081 | 78.4839939419761 | 39.8502650539468 | 38.74178205952731 | 75.6282792444948 | 45.28724916908366 | 91.12586810336158 | 21.128129364562724 | 29.017025318198737 |
| SE_8_3_6 | SE_8 | SE_8_3 | AGTCGCGTCGGGCCTCCCGA | 179.90695572025763 | 252.84629861237573 | 291.3204244771905 | 226.4276771845537 | 237.57317085138712 | 316.2031901019692 | 176.4903404934022 | 227.8301912240406 | 300.1390239049074 | 372.52254880654215 | 247.66862755126303 | 116.06810127279495 |
| SE_8_4_2 | SE_8 | SE_8_4 | TGACCAGTAAGGCCAAGTAA | 1189.9869721737523 | 1162.2934596292212 | 1217.6206216283929 | 1197.2038104010885 | 1219.6836896388177 | 1447.604193590111 | 1275.2503261261072 | 1189.2546911196807 | 1426.9923414650477 | 945.5220074404798 | 1064.6229629810216 | 1452.0119469226647 |
| SE_8_4_3 | SE_8 | SE_8_4 | CTGCGTTACTGAGAAAAGGA | 332.71949039830776 | 387.7642840379517 | 356.4971974110704 | 270.6721658298113 | 410.4500764262804 | 169.79678153420812 | 287.33488360816085 | 351.6714984869008 | 273.49946557015227 | 259.5264723583738 | 257.05890726884644 | 215.88666836739858 |
| SE_8_4_4 | SE_8 | SE_8_4 | TTAACTAATTAAAAACAGAA | 3.2513305250648967 | 1.9987849692677924 | 4.93763431317272 | 5.205233958265602 | 0.0 | 0.0 | 2.152321225529295 | 0.0 | 16.871720278678225 | 1.4580138896537853 | 0.0 | 4.642724050911798 |
| SE_8_4_8 | SE_8 | SE_8_4 | TTACGTTCACAGCAGCAGTG | 3109.3557588037297 | 2820.285591636855 | 2696.93586185494 | 2793.9093270990616 | 2860.42340083121 | 2663.0372777354883 | 2819.5408054433765 | 2839.84188563078 | 2701.2512151441665 | 2882.4934598455334 | 2851.1236792512696 | 2493.1428153396355 |
| SE_8_4_9 | SE_8 | SE_8_4 | GAAACTCTCAGTTTGAACCA | 74.78060207649263 | 46.97144677779312 | 52.338923719630834 | 45.545797134824014 | 42.42378050917627 | 66.70587845986748 | 52.73187002546773 | 66.17474433893295 | 32.855455279531284 | 43.74041668961356 | 64.5581730583861 | 80.08698987822851 |
| SE_80_1_3 | SE_80 | SE_80_1 | TTACAGCAGAAGAATTCAGT | 293.703524097529 | 328.80012744455183 | 331.8090258452068 | 290.1917931733073 | 314.9965702806338 | 194.91977472039196 | 289.48720483369016 | 406.50200093915953 | 525.6872844725006 | 343.36227101346645 | 383.8276834562228 | 406.2383544547823 |
| SE_80_1_4 | SE_80 | SE_80_1 | CAGTTAGAGCAAGCATTGAG | 1623.4977088490718 | 1541.0632113054678 | 1497.0907237539689 | 1643.5526223223637 | 1596.194741657757 | 2178.769926319048 | 1693.876804491555 | 1836.8218321506674 | 1668.524337033494 | 1699.3151883914868 | 1767.720156835081 | 1889.5886887211018 |
| SE_80_1_7 | SE_80 | SE_80_1 | ATGCTTGCTCTAACTGTAGA | 333.80326723999605 | 310.8110627211417 | 315.02106918041954 | 392.9951638490529 | 371.20807945529236 | 175.86095230328698 | 403.5602297867428 | 269.4257448085127 | 370.2898608530958 | 315.66000711004455 | 356.8306292681704 | 220.52939241831038 |
| SE_80_1_8 | SE_80 | SE_80_1 | GCTATTGCAAAACTTAAGCA | 198.3311620289587 | 290.8232130284638 | 319.9587034935923 | 256.35777244458086 | 331.9660824843043 | 267.6898239493383 | 167.88105559128502 | 274.1525122612937 | 286.81924473752986 | 311.2859654410832 | 248.84241251596094 | 287.84889115653147 |
| SE_80_1_9 | SE_80 | SE_80_1 | TATCTCTCCCTTGCTCAACA | 170.15296414506292 | 151.90765766435223 | 157.0167711588925 | 147.04785932100324 | 177.11928362581094 | 109.1550738434195 | 229.2222105188699 | 216.48594933736638 | 214.00445195586593 | 200.47690982739547 | 221.8453583279086 | 133.47831646371418 |
| SE_80_2_1 | SE_80 | SE_80_2 | CGTGTTCCTTGACTTGTAGA | 1052.3473132793383 | 1388.1561611564819 | 1230.4584708426419 | 1258.3653094107092 | 1541.043826995828 | 1663.3154109473448 | 1136.4256070794677 | 1179.801156214119 | 1304.4503731251743 | 1252.4339312126017 | 1028.2356290753858 | 1922.0877570774842 |
| SE_80_2_10 | SE_80 | SE_80_2 | GTGAATAAAGAGGCTGGGAG | 1270.1864584586863 | 1759.9301654402911 | 1690.6459888303393 | 1473.0812101891654 | 1660.891006934251 | 1925.8073742374727 | 1454.9691484578034 | 1537.1447756443567 | 1826.5857164863742 | 1903.4371329430166 | 1187.8703842743041 | 1457.8153519863044 |
| SE_80_2_16 | SE_80 | SE_80_2 | AGCCTCATCCTCCTCCCCTG | 618.8365766040187 | 687.5820294281206 | 603.3789130697064 | 648.0516278040674 | 548.3273630811033 | 1004.0534173374857 | 615.5638705013783 | 505.764117447559 | 515.0314611385985 | 440.3201946754432 | 680.7952795247988 | 923.9020861314477 |
| SE_80_2_17 | SE_80 | SE_80_2 | TAGTAAACGACCCTGAAAAG | 203.7500462374002 | 295.82017545163325 | 283.42020957611413 | 238.13945359065127 | 334.0872715097631 | 216.5775274671022 | 156.0432888508739 | 282.66069367629933 | 198.90870223283804 | 306.1829168272949 | 523.5080942552763 | 239.1002886219576 |
| SE_80_2_3 | SE_80 | SE_80_2 | CCTCCTATCTCAGAACCCTG | 176.65562519519273 | 204.87545934994873 | 250.8318231091742 | 152.25309327926885 | 213.17949705861076 | 164.59892087499765 | 164.65257375299106 | 264.6989773557318 | 217.5563930671666 | 247.1333542963166 | 179.58909959878315 | 80.08698987822851 |
| SE_81_1_10 | SE_81 | SE_81_1 | TCCCAGCTGCAGCACACAGA | 161.48274941155654 | 40.97509186998975 | 97.76515940081985 | 143.14393385230406 | 55.15091466192915 | 204.44918592894447 | 224.91756806781132 | 92.64464207450614 | 361.4100080748441 | 16.038152786191638 | 69.25331291717781 | 140.44240254008187 |
| SE_81_1_11 | SE_81 | SE_81_1 | GCAGATGGTGAATAGCCCTG | 5648.644898879414 | 5346.749792791345 | 5255.6179629410435 | 5637.268376801647 | 5357.062883796233 | 5011.603985588745 | 5554.064922478346 | 5420.656914849164 | 5047.3083191582655 | 5544.826822353346 | 4756.176676956008 | 5422.701691464979 |
| SE_81_1_13 | SE_81 | SE_81_1 | AGGGCCTGCATTCAACCACA | 3789.9676153839814 | 3762.712704646619 | 3797.0407868298216 | 4080.903423280232 | 3756.6257640875588 | 3853.3473686946822 | 3833.2841026676742 | 4135.92152118331 | 3967.5182213228586 | 2993.3025154592215 | 4851.253259096541 | 3428.6517115983625 |
| SE_81_1_8 | SE_81 | SE_81_1 | TTCCTTGATCCCAAACCAGA | 11.921545258571289 | 1.9987849692677924 | 2.962580587903632 | 2.602616979132801 | 0.0 | 0.0 | 1.0761606127646475 | 0.945353490556185 | 2.6639558334755096 | 0.0 | 0.0 | 0.0 |
| SE_81_1_9 | SE_81 | SE_81_1 | GCCGTCTGTGTGCTGCAGCT | 757.5600123401209 | 220.86573910409106 | 362.42235858687764 | 638.9424683771026 | 219.5430641349872 | 1712.6950872098441 | 1043.8757943871081 | 527.5072477303512 | 1551.3102803605716 | 75.81672226199683 | 245.32105762186717 | 396.95290635295873 |
| SE_81_2_1 | SE_81 | SE_81_2 | CCAGCTGGTAGAAAGTCAGT | 790.0733175907699 | 699.5747392437273 | 705.0941799210644 | 801.6060295729027 | 759.3856711142553 | 1127.0694529387997 | 816.8059050883675 | 801.6597599916449 | 743.2436775396671 | 567.1674030753225 | 802.8689158533834 | 1230.3218734916263 |
| SE_81_2_13 | SE_81 | SE_81_2 | TTAACCCTGTGACTTTGCAC | 211.3364841292183 | 198.87910444214535 | 176.76730841158337 | 127.52823197750725 | 167.57393301124625 | 165.46523098486605 | 151.73864639981528 | 130.45878169675353 | 160.72533528635574 | 357.9424099100043 | 107.98821675220947 | 248.38573672378118 |
| SE_81_2_14 | SE_81 | SE_81_2 | GGAAGGAAGAAGTTTCTGAA | 302.3737388310354 | 487.70353250134133 | 458.21246426242845 | 294.0957186420065 | 368.02629591710416 | 618.5454184460439 | 297.0203291230427 | 293.05958207241736 | 407.58524252175295 | 497.9117433167677 | 291.0986712450864 | 555.9662050966878 |
| SE_81_2_15 | SE_81 | SE_81_2 | GTAGCTGTGAATAATTGGAA | 312.12773040623006 | 422.74302100013807 | 403.8984868175285 | 369.57161103685775 | 409.389481913551 | 294.54543735525897 | 396.02710549739027 | 362.07038688301884 | 461.7523444690883 | 381.27063214446486 | 396.7393180679 | 420.1665266075177 |
| SE_81_2_26 | SE_81 | SE_81_2 | GGGAGAATGAGGGCACACGG | 2145.878146542832 | 2311.594816958202 | 2338.4636107186 | 2191.4034964298185 | 2547.548019576035 | 1994.245872917077 | 2021.029630772008 | 1945.5374835646287 | 2209.3073712290225 | 1368.3460354400775 | 1547.0485834718704 | 1651.649081111872 |
| SE_82_1_15 | SE_82 | SE_82_1 | CTGAAAGCTCTCCACCTCCA | 486.6158019180462 | 545.6682966101073 | 558.940204251152 | 533.5364807222242 | 566.3574697975032 | 438.3529155934148 | 471.3583483909156 | 556.813205937593 | 528.351240305976 | 438.1331738409625 | 767.6553669124456 | 576.8584633257909 |
| SE_82_1_16 | SE_82 | SE_82_1 | ACTGAAAGCTCTCCACCTCC | 1612.6599404321887 | 1694.969653939088 | 1633.3694307975359 | 1726.8363656546135 | 1735.1326228253095 | 2235.0800834604947 | 1625.0025252746177 | 1997.5319255452189 | 1503.3590753580124 | 1969.047757977437 | 2104.596441703387 | 1521.6528076863417 |
| SE_82_1_18 | SE_82 | SE_82_1 | ATCACTGAACTGCACGCTCC | 1884.6879276959519 | 1587.0352655986271 | 1635.344484522805 | 1745.054684508543 | 1480.589939770252 | 1620.8662155637928 | 1877.90026927431 | 1947.428190545741 | 1821.2578048194232 | 1678.1739869915068 | 2250.14577732593 | 1541.3843849027169 |
| SE_82_1_19 | SE_82 | SE_82_1 | GATCACTGAACTGCACGCTC | 2109.02973392543 | 1734.9453533244439 | 1803.2240511706775 | 1997.5085314844246 | 1912.2519064511205 | 2097.3367759914177 | 2186.7583651377636 | 2072.2148512991575 | 1821.2578048194232 | 1917.2882648947277 | 2083.468312338824 | 2538.4093748360256 |
| SE_82_1_20 | SE_82 | SE_82_1 | GTGCAGTTCAGTGATCGTAC | 856.1837049337562 | 825.4981923075983 | 838.4103063767279 | 913.5185596756131 | 750.90091501242 | 728.5668023993318 | 846.9384022457775 | 1051.2330814984778 | 766.3312947631215 | 651.7322086752421 | 714.8350435010387 | 793.9058127059175 |
| SE_82_2_13 | SE_82 | SE_82_2 | TAAAAGTGAGTGCAGAGACG | 373.90301038246315 | 495.6986723784125 | 453.2748299492557 | 432.03441853604494 | 475.14634170277424 | 570.8983624032813 | 385.2654993697438 | 445.26149405196315 | 550.5508722516053 | 384.1866599237724 | 392.0441782091083 | 488.6467063584667 |
| SE_82_2_17 | SE_82 | SE_82_2 | TTTACAGCAAAACTACTCAA | 79.11570944324582 | 97.94046349412183 | 101.71526685135804 | 62.46280749918722 | 85.90815553108195 | 110.88769406315632 | 50.57954879993843 | 68.06545132004533 | 167.8292175089571 | 64.15261114476655 | 52.820323411406804 | 154.37057469281729 |
| SE_82_2_19 | SE_82 | SE_82_2 | CAGCCAGTAGGACCTTCCAG | 939.6345217437552 | 1085.3402383124112 | 1143.556106930802 | 1031.9376322261555 | 1126.35137251863 | 776.2138584420942 | 863.0808114372472 | 804.4958204633134 | 863.121690046065 | 1439.0597090882861 | 821.6494752885503 | 1151.3955646261259 |
| SE_82_2_22 | SE_82 | SE_82_2 | ATTGGTCTACACTAGCCACA | 161.48274941155654 | 147.91008772581662 | 158.9918248841616 | 167.86879515406565 | 145.30144824392872 | 132.54544680986655 | 159.27177068916782 | 148.42049801732105 | 144.74160028550267 | 234.01122928943255 | 122.07363632858463 | 31.338387343654635 |

| | | | | | | | | | | | | | | | |
|---|---|---|---|---|---|---|---|---|---|---|---|---|---|---|---|
| SE_82_2_29 | SE_82 | SE_82_2 | GCCCCAAGAGGATGTGGCAG | 200.4987157123353 | 213.8699917116538 | 274.53246781240324 | 219.92113473672168 | 277.8757623351046 | 292.81281713552215 | 145.28168272322742 | 296.8409960346421 | 242.41998084627136 | 94.77090282749604 | 198.36965903395 | 479.3612582566431 |
| SE_83_1_16 | SE_83 | SE_83_1 | CCTGCACTGCGTTCCCAACA | 2985.8051988512634 | 2736.3366229276076 | 2560.6571548113725 | 2938.354569440932 | 2627.0926080307404 | 2527.026590486148 | 2752.8188474519684 | 2401.1978660127097 | 2275.90626706591 | 2431.9671679425137 | 2786.5655061928837 | 2417.6985495123186 |
| SE_83_1_2 | SE_83 | SE_83_1 | CAGGAGATTGAACTTAGAGG | 977.5667112028456 | 1479.1008772581663 | 1466.4773910122979 | 1159.4658642036627 | 1367.1063269082053 | 1658.9838603980027 | 955.630624135007 | 1314.9867053636533 | 1692.4999395347736 | 1146.7279242127022 | 1592.8261970950898 | 1746.8249241555638 |
| SE_83_1_5 | SE_83 | SE_83_1 | CAGAGTGACATCGCTCTAGG | 2362.6335148804915 | 2030.765528776077 | 2117.2575934884626 | 2377.4906104378138 | 2402.246571332106 | 2089.539985002602 | 2418.132896882163 | 2430.5038242199516 | 2330.9613542910706 | 2103.185035825585 | 2294.749605984451 | 2303.9518102649795 |
| SE_83_1_6 | SE_83 | SE_83_1 | TGGCAGCCATGTCCACTGCA | 59.60772629285644 | 18.98845720804403 | 24.688171565863602 | 33.834020728726415 | 21.211890254588134 | 112.62031428289313 | 58.112673089290965 | 35.92343264113503 | 202.4606433441387 | 4.374041668961356 | 9.390279717583432 | 76.60494684004466 |
| SE_83_1_8 | SE_83 | SE_83_1 | GCAGTGGACATGGCTGCCAA | 5111.091585402018 | 4406.3214647508485 | 4644.338834970261 | 5140.168533787281 | 4942.370429319036 | 5352.063858767029 | 5053.650237542784 | 5310.995909944648 | 4243.681642726487 | 3990.5840159824106 | 5103.617026506596 | 4618.349749644511 |
| SE_83_2_10 | SE_83 | SE_83_2 | AAGAGATCAGGATATTTAAA | 74.78060207649263 | 142.91312530264716 | 131.34107273039436 | 115.81645557140963 | 133.63490860390525 | 295.41174746512735 | 136.67239782111022 | 97.37140952728706 | 124.31793889552377 | 303.9958959928142 | 37.56111887033373 | 348.20430381838486 |
| SE_83_2_26 | SE_83 | SE_83_2 | CCATCCTGCCGGGGTCCTGG | 153.89631151973845 | 253.84569109700962 | 291.3204244771905 | 226.4276771845537 | 260.90625013143404 | 247.7646914223649 | 124.83463108069911 | 250.518674997389 | 253.96378945799856 | 153.09145841364744 | 144.37555065784528 | 369.0965620474879 |
| SE_83_2_3 | SE_83 | SE_83_2 | AAGGCTCTACTTTTTTTAAG | 42.26729682584366 | 58.96415659339988 | 57.27655803280356 | 62.46280749918722 | 60.45388722557618 | 76.23528966841997 | 27.980175931880833 | 43.48626056558451 | 140.30167389637683 | 57.59154864132452 | 53.994108376104734 | 40.62383544547823 |
| SE_83_2_44 | SE_83 | SE_83_2 | CTGCCCCTCCTCGGAGTAAA | 84.53459365168732 | 100.93864094802352 | 97.76515940081985 | 61.16149900962082 | 121.96836896388177 | 100.4919727447354 | 61.3411549275849 | 32.14201867891029 | 157.17339417505505 | 71.44268059303548 | 36.3873339056358 | 314.5445544492743 |
| SE_83_2_7 | SE_83 | SE_83_2 | AATGAGATCACTTTTCCAAA | 31.429528408960667 | 54.96658665486429 | 78.01462214812898 | 44.24448864525762 | 59.39329271284678 | 100.4919727447354 | 38.74178205952731 | 94.5353490556185 | 38.1833669464823 | 48.114458358574915 | 127.94256115207426 | 45.26655949639003 |
| SE_83_3_15 | SE_83 | SE_83_3 | GTGAATGGAAAATACAAAAT | 197.2473851872704 | 237.85541134286728 | 237.00644703229057 | 231.63291114281927 | 302.2694361278809 | 161.99999054539242 | 201.24203458698906 | 335.60048914744567 | 275.2754361258026 | 258.7974654135469 | 221.8453583279086 | 248.38573672378118 |
| SE_83_3_18 | SE_83 | SE_83_3 | ATCTAACAGAGGGTTAGGGG | 67.19416418467453 | 113.93074324826416 | 134.303653318298 | 70.27065843658562 | 62.575076251035 | 113.48662439276154 | 71.02660044246673 | 44.431614056140695 | 88.79852778251698 | 23.328222234460565 | 51.646538446708874 | 71.96222278913287 |
| SE_83_3_22 | SE_83 | SE_83_3 | AGCTCAATTTATCTAACAGA | 55.27261892610324 | 113.93074324826416 | 83.93978332393624 | 55.95626505135522 | 93.3323171201878 | 162.86630065526083 | 36.58946083399801 | 44.431614056140695 | 107.44621861684554 | 41.553395855132884 | 37.56111887033373 | 253.02846077469297 |
| SE_83_3_23 | SE_83 | SE_83_3 | TTACGTGTCTCATCCAGCCC | 4962.614158090721 | 4154.4745586231065 | 3986.6459444556544 | 4744.570752959096 | 4019.6532032444516 | 4341.079960550596 | 4798.600172317563 | 3988.4463766565445 | 4114.923777441837 | 4266.148641126976 | 4487.379920040183 | 4726.29308382821 |
| SE_83_3_4 | SE_83 | SE_83_3 | GCCGGCACAGGCCCTGCTGG | 56.35639576779154 | 58.96415659339988 | 57.27655803280356 | 111.91253010271043 | 53.02972563647034 | 189.7219140611815 | 122.6823098551698 | 70.90151179171387 | 86.13457194904147 | 85.29381254474644 | 145.5493356225432 | 32.49906835638259 |
| SE_83_4_2 | SE_83 | SE_83_4 | CCAGCCCTGTGCAGCGACTG | 1178.065426915181 | 1021.3791192958419 | 1098.129871249613 | 1188.0946509741236 | 968.3227901219484 | 1065.5614351381428 | 1265.5648806112254 | 989.7851046123257 | 1120.6374206153644 | 1151.1019658816635 | 977.7628755933749 | 990.0609038569409 |
| SE_83_4_24 | SE_83 | SE_83_4 | TGGGGGCTCAGGCTTCCCAT | 303.4575156727237 | 281.8286806667587 | 320.9462303562268 | 262.8643148924129 | 309.69359771698674 | 225.24062856578627 | 238.90765603375175 | 289.2781681101926 | 226.43624584541828 | 353.56836824104295 | 183.11045449287693 | 363.2931569838482 |
| SE_83_4_25 | SE_83 | SE_83_4 | TCAGAGGTGAACAGTCCCCA | 5412.381547391365 | 5240.814189420152 | 5441.273013116338 | 5427.757709981456 | 5336.911588054375 | 5618.887372606499 | 5196.779599040483 | 4909.220676458269 | 5193.825889999418 | 5119.08676657444 | 5061.36076777747 | 4206.307990126089 |
| SE_83_4_32 | SE_83 | SE_83_4 | GTCCCCTGAAGCCAATTCCC | 21.675536833765978 | 42.973876839257535 | 53.32645058226538 | 14.314393385230405 | 46.6661585600939 | 1.7326202197368175 | 15.066248578705064 | 16.071009339455145 | 20.423661389978903 | 37.17935418617152 | 34.03976397623994 | 20.89225822910309 |
| SE_83_4_40 | SE_83 | SE_83_4 | CGTGTCAGATGAGACCCAGC | 12127.462858492065 | 11914.75720180531 | 12017.214391399766 | 11670.13453443148 | 11955.021347485872 | 11076.641064777474 | 11101.672881280103 | 12035.29528827079 | 11592.647802007592 | 10373.768824886682 | 10801.169245150342 | 12263.755580483514 |
| SE_83_5_1 | SE_83 | SE_83_5 | TCAGCCGCAGTCGCTGCACA | 3489.7614302363227 | 3278.0073495991796 | 3400.054988050735 | 3301.4196380299577 | 3018.4519832278916 | 3119.58270563614 | 3620.204301340274 | 3771.960427319178 | 3597.2283604697627 | 2949.5620987696075 | 3693.9012839043826 | 3739.7142230094532 |
| SE_83_5_10 | SE_83 | SE_83_5 | TCGACCCTAAGCATCCTCGT | 3756.370533291644 | 3409.927157570854 | 3242.050690029208 | 3274.0921597490633 | 2751.1821660200812 | 3110.9196045374556 | 3161.7598803025344 | 2874.8199647813585 | 2718.122935422845 | 3194.5084322314437 | 2621.0618261704753 | 2429.305359639598 |
| SE_83_5_12 | SE_83 | SE_83_5 | GTGCGTGATCTCCAATGGCA | 3706.516798573982 | 3350.9630009774537 | 3208.4747766996334 | 3387.3059983413405 | 3437.3868157560073 | 3298.9088983789006 | 3721.363398940151 | 3190.5680306271242 | 3092.8527226650663 | 2791.367591742172 | 3121.0942211317933 | 3278.9238609564572 |
| SE_83_5_46 | SE_83 | SE_83_5 | TTCTGGCTCCTCCTCCACCA | 28.17819788389577 | 25.984204600481302 | 25.675698428498144 | 16.917010364363207 | 4.242378050917627 | 67.57218856973589 | 5.380803063823238 | 41.59555358447214 | 6.215896944776189 | 0.0 | 46.95139858791716 | 63.83745570003722 |
| SE_83_5_50 | SE_83 | SE_83_5 | GTCTGACCCACAGTCCCTGG | 248.18489674662044 | 225.86270152726053 | 197.50537252690881 | 242.04337905935049 | 232.27019828774007 | 252.09624197170695 | 289.48720483369016 | 267.53503782740034 | 465.304285580389 | 238.3852709583939 | 385.00146842092073 | 341.24021774201714 |
| SE_83_6_15 | SE_83 | SE_83_6 | CCAAGAACCAAAGTAAGCCT | 1579.0628583398516 | 1520.0759691281562 | 1540.5419057098886 | 1460.0681252935012 | 1630.1337660650981 | 1353.1763916144544 | 1374.2571025004547 | 1362.2543798914626 | 1463.3997378558797 | 1402.6093618469415 | 1429.6700870020775 | 1133.9853494352067 |
| SE_83_6_20 | SE_83 | SE_83_6 | AAGCAGTCGCTCAAAGCAGC | 721.7953765644071 | 615.62577053448 | 618.1918160092246 | 707.9118183241219 | 830.4455034671255 | 996.25662634867 | 841.5575991819543 | 1041.779546592916 | 710.3882222601359 | 801.9076393095819 | 1164.3946849803456 | 587.3045924403424 |
| SE_83_6_22 | SE_83 | SE_83_6 | GCAGCAGGAAGTGATGTGGC | 11763.313839684797 | 10294.741984213764 | 10402.607970992287 | 11325.287784696384 | 10664.277825494184 | 11787.015354869569 | 11449.272759203084 | 11041.728769696241 | 10274.87764971504 | 11212.855818382435 | 11332.893834158505 | 11744.93116779412 |
| SE_83_6_23 | SE_83 | SE_83_6 | CATTTTCTGCGTTTCCTTCT | 70.44549470973944 | 89.94532361705066 | 82.9522564613017 | 84.58505182181602 | 106.05945127294068 | 88.3636312065777 | 88.2451702467011 | 94.5353490556185 | 112.77413028379657 | 72.17168753786237 | 98.59793703462604 | 89.37243798005211 |
| SE_83_6_31 | SE_83 | SE_83_6 | CAACGGCTGAGTTTCGGTGG | 6304.329888100835 | 6386.117976810597 | 6797.147395513566 | 6624.961520382544 | 6142.9634177287235 | 7541.229506404498 | 6360.1092214390665 | 6590.059182667165 | 7031.955415097519 | 7206.9626565586605 | 6306.746615321973 | 8066.733038459249 |
| SE_83_7_0 | SE_83 | SE_83_7 | GTGTGAGGAAGTGTGAGAAG | 14850.99406165476 | 14727.047653565094 | 15115.086159484332 | 15266.95119959301 | 15292.712279045316 | 14335.699698102428 | 14937.109305173308 | 14699.30142465812 | 14570.062438555386 | 14870.283660578956 | 14615.970380418612 | 15888.5623832329 |
| SE_83_7_13 | SE_83 | SE_83_7 | TGTCAACTTTCACATCAACG | 886.5294565010286 | 977.4058499719505 | 948.0257881291623 | 939.5447294669411 | 920.596037049125 | 823.8609144848567 | 997.6008880328282 | 759.1188529166166 | 808.0666028209044 | 820.8618198750811 | 835.7348948649254 | 1079.433341836993 |
| SE_83_7_16 | SE_83 | SE_83_7 | GAGGCAGCATCCATACCTGT | 24076.10253810556 | 20689.42321689092 | 20197.886921464327 | 22916.04250126431 | 20621.139110997854 | 19955.453380818795 | 24415.931982404323 | 22640.270745330075 | 20589.71463693221 | 20200.05343420837 | 22029.596217450733 | 19630.597968267808 |
| SE_83_7_2 | SE_83 | SE_83_7 | GGCAATGGTGATTCACAGGC | 197.2473851872704 | 403.7545637920941 | 365.38493917478127 | 266.7682403611121 | 358.48094530253945 | 749.3582450361736 | 210.9274801018709 | 300.62240999686685 | 283.26730362622914 | 339.717236289332 | 299.3151659979719 | 607.0361696567176 |
| SE_83_7_8 | SE_83 | SE_83_7 | GCGCACTGTATCCTGCCTGA | 10365.241713906891 | 9438.262624882516 | 9183.999822501259 | 10415.673150489469 | 9802.014486645177 | 10847.068885662346 | 10236.439748617327 | 10500.041219607547 | 8604.577342125895 | 10204.639213686844 | 10426.731841411703 | 11972.424646288799 |
| SE_84_1_10 | SE_84 | SE_84_1 | GGAAGGTCTGTTTAACAAAA | 389.0758861660993 | 507.69138219401924 | 546.1023550369028 | 389.09123838035373 | 535.6002289283504 | 454.8128076809146 | 435.84504816968223 | 584.2284571637223 | 576.3024453085352 | 405.3278613237523 | 453.0809963734006 | 406.2383544547823 |
| SE_84_1_12 | SE_84 | SE_84_1 | TGTTAAACAGACCTTCCAGA | 494.2022398098643 | 631.6160502886224 | 591.5285907180919 | 461.96451379607214 | 521.8125002628681 | 513.7218951519664 | 454.13977858668125 | 608.8076479181832 | 531.0151961394515 | 587.5795975304754 | 651.4506554073506 | 495.6107924348344 |
| SE_84_1_17 | SE_84 | SE_84_1 | GGACAAGAGCCACCTTCAGG | 898.4510017595998 | 1165.2916370831229 | 1201.82019182624 | 1212.8195122758852 | 1354.3791927554523 | 1295.9999243631394 | 1068.627488475295 | 1376.4346822498053 | 1590.381632584879 | 1233.4797506471023 | 1599.8689068832773 | 1288.3559241280238 |
| SE_84_1_2 | SE_84 | SE_84_1 | ACCCTTATCCTGAGTGAACG | 230.84446727960767 | 261.8408309740808 | 236.018920169656 | 227.72898567412008 | 220.6036586477166 | 300.60960812433785 | 174.33801926787288 | 296.8409960346421 | 182.92496723198497 | 214.32804177910643 | 248.84241251596094 | 419.0058455947897 |
| SE_84_1_25 | SE_84 | SE_84_1 | GGTGGAGTCCTCAGGACAAG | 479.0293640262281 | 519.684092009626 | 477.9630015151193 | 553.0561080657202 | 509.0853661101152 | 431.42243471446756 | 516.5570941270308 | 673.0916852760038 | 465.304285580389 | 316.38901405487144 | 712.4874735716429 | 535.0739468675847 |
| SE_84_2_1 | SE_84 | SE_84_2 | AATGGAACTGAGGAAGCTCA | 244.93356622155557 | 308.8122777518739 | 316.00859604305407 | 278.4800167672097 | 240.75495438957532 | 353.4545248263108 | 214.15596194016484 | 292.11422858186114 | 253.07580418017338 | 296.7058265445453 | 395.5655331032021 | 200.79781520193526 |
| SE_84_2_23 | SE_84 | SE_84_2 | CAGGCAAGACACGGTCTGAT | 21531.394513821433 | 16502.968098759527 | 16036.44872232236 | 19185.19106167744 | 17075.571654943447 | 19296.191387208935 | 22120.48139537733 | 17625.170477929514 | 16857.51251423302 | 15830.385806915974 | 18037.553552513076 | 16753.26973771522 |
| SE_84_2_26 | SE_84 | SE_84_2 | TGTCTTGCCTGCCAACTCGA | 781.4031028572635 | 624.6203028961851 | 580.6657952291118 | 610.3136816066418 | 554.6909301574798 | 726.8341821795949 | 647.8486888843178 | 598.4087595220651 | 706.8362811488352 | 726.0909170475851 | 584.5449124195686 | 540.8773519312244 |
| SE_84_2_27 | SE_84 | SE_84_2 | CTCTCGGCCATCGAGTTGGC | 5.4188842084414945 | 4.996962423169481 | 6.912688038441808 | 7.807850937398403 | 4.242378050917627 | 0.8663101098684087 | 0.0 | 0.0 | 0.0 | 0.0 | 0.0 | 2.321362025455899 |
| SE_84_2_28 | SE_84 | SE_84_2 | ACGGCTCTCGGCCATCGAGT | 2366.9686222472446 | 2234.641595641392 | 2068.86877721937 | 2266.8793888246696 | 2225.127287706295 | 2164.9089645611534 | 2433.199145460868 | 2247.1052470520517 | 2206.643415395547 | 2386.7687373632466 | 3134.0058557434704 | 2637.067260917901 |
| SE_85_1_10 | SE_85 | SE_85_1 | GGCAGCTCAGAGAATCCATG | 679.5280797385634 | 805.5103426149203 | 769.2834259923098 | 805.5099550416019 | 804.9912351616197 | 648.8662722914381 | 736.0938591310189 | 1071.0855048001577 | 802.7386911539535 | 1374.1780909986926 | 708.9661186775492 | 422.4878886329736 |
| SE_85_1_18 | SE_85 | SE_85_1 | TGGCCAGGAGATAAGGGACA | 154.98008836142674 | 193.88214201897586 | 170.8421472357761 | 201.70281588279207 | 202.5735519313167 | 61.50801780065702 | 194.7850709104012 | 349.78079150578844 | 185.58892306546048 | 452.71331273750036 | 380.306328562129 | 103.3006101327875 |
| SE_85_1_20 | SE_85 | SE_85_1 | ACACCCCTTCCCACGAACAA | 1261.51624372518 | 1161.2940671445874 | 1207.7453530020473 | 1158.1645557140964 | 1110.4424548276888 | 1082.0213272256426 | 1148.2633738198788 | 1231.795598194709 | 986.5516436637637 | 1302.7354104056571 | 1448.4506464372444 | 1485.6716962917753 |
| SE_85_1_25 | SE_85 | SE_85_1 | GTACCGCCCCAGGCAGGAAA | 514.7939998019419 | 680.5862820356833 | 703.1191261957954 | 611.6149900962082 | 673.4775155831733 | 489.4652120756509 | 546.6895912844409 | 549.2503780131435 | 770.7712211522473 | 707.8657434269128 | 516.4653844670888 | 468.9151291420916 |
| SE_85_1_5 | SE_85 | SE_85_1 | GTTGAAATGGAGCAAGATGC | 406.4163156331121 | 477.7096076550024 | 445.37461504817935 | 370.87291952642414 | 395.6017532480687 | 269.4224441690751 | 301.3249715741013 | 518.9990663153455 | 257.51573056929925 | 436.67515995130873 | 615.0633215017148 | 639.5352380131002 |
| SE_85_2_11 | SE_85 | SE_85_2 | TGATCCACATGGAATCTCAA | 493.118462968176 | 582.6458185415614 | 593.5036444433609 | 503.60638546219695 | 643.7808692267499 | 738.0962136078842 | 499.33852432279645 | 482.13028018365435 | 564.758636696808 | 575.1864794684183 | 381.4801135268269 | 630.2497899112766 |
| SE_85_2_13 | SE_85 | SE_85_2 | TCAGTTTCTTTCTCTGCAAG | 54.188842084414944 | 72.95565137827442 | 94.80257881291622 | 130.13084895664005 | 59.39329271284678 | 122.14972549144564 | 100.08293698711222 | 51.994441980590175 | 46.17523444690883 | 96.95792366197672 | 9.390279717583432 | 18.57089620364719 |
| SE_85_2_14 | SE_85 | SE_85_2 | TTAGGGGCAGATGAGGGGGT | 941.8020754271317 | 1302.2084074779668 | 1318.3483616171163 | 1076.1821208714132 | 1331.0461134754055 | 635.0053105335436 | 787.7495685437219 | 853.6542019722351 | 1148.1649642279444 | 599.2437086477057 | 697.2282690305699 | 550.162800033048 |
| SE_85_2_27 | SE_85 | SE_85_2 | CAACACTTGGGGACTCCACA | 602.5799239786942 | 698.5753467590935 | 716.944502272679 | 688.3921909806259 | 739.2343753723965 | 404.5668213085469 | 561.755839863146 | 789.3701646144145 | 754.7874861513943 | 551.8582572339577 | 881.5125084881447 | 631.4104709240045 |
| SE_85_2_6 | SE_85 | SE_85_2 | ATGCAGAGTGGTAATTCAGG | 1366.642597368945 | 1533.0680714283967 | 1539.5543788472542 | 1519.9283158135556 | 1546.3467995594751 | 1393.8929667782697 | 1501.2440548066832 | 1308.36923092976 | 1419.8884592424465 | 1707.3342647845825 | 1603.390261777371 | 1884.9459646701898 |
| SE_85_3_17 | SE_85 | SE_85_3 | GCCATCCAGGAAAGGAAGAG | 644.8472208045379 | 683.584459489585 | 725.8322440363899 | 720.9249032197858 | 758.3250766015258 | 753.6897955855156 | 685.5143103310804 | 854.5995554627913 | 511.4795200272978 | 505.9308197098635 | 796.9999910298939 | 625.6070658603647 |
| SE_85_3_20 | SE_85 | SE_85_3 | TTTCCTGGATGGCACCAGCA | 30.34575156727237 | 44.97266180852533 | 46.41376254382357 | 37.737946197425615 | 12.727134152752882 | 12.128341538157722 | 65.6457973786435 | 52.93979547114636 | 42.62329333560815 | 172.77464592397357 | 30.518409082146157 | 46.42724050911798 |
| SE_85_3_25 | SE_85 | SE_85_3 | CCAGGGTTTACCACTCCAAT | 1258.264913200115 | 1270.227847969682 | 1341.061479457711 | 1379.3869989403845 | 1373.4698939845816 | 969.4010129427494 | 1252.6509532580496 | 1353.746198476457 | 1163.2607139509723 | 1460.200910488266 | 1140.918985686387 | 1258.1782177970972 |
| SE_85_3_26 | SE_85 | SE_85_3 | CGATTGGAGTGGTAAACCCT | 3210.1470050807416 | 3281.005527053081 | 3450.418858045097 | 3361.2798285500126 | 3757.6863586002883 | 2867.486463664433 | 3128.3989013068303 | 3407.999333455047 | 3650.507477139273 | 3187.9473697280014 | 3749.0691772451855 | 3811.676445798586 |
| SE_85_3_32 | SE_85 | SE_85_3 | GGGCCGGAAGGCATGCCAAA | 252.52000411337366 | 240.85358879676897 | 251.81934997180872 | 262.8643148924129 | 286.3605184369398 | 468.67376943880913 | 316.39122015280634 | 277.93392622351837 | 353.4181405744176 | 285.041715427315 | 375.6111887033373 | 495.6107924348344 |
| SE_85_4_17 | SE_85 | SE_85_4 | AAACCTCTCTTCCACTCCAG | 34.680858934025565 | 16.989672238776237 | 25.675698428498144 | 27.32747828089441 | 27.575457330964575 | 28.588233625657487 | 54.88419125099702 | 7.56282792444948 | 1.7759705556503396 | 9.477090282749604 | 1.173784964697929 | 0.0 |
| SE_85_4_22 | SE_85 | SE_85_4 | GGTGGGGCCCTAAGACCCAT | 414.0027535249302 | 432.73694584647706 | 407.8485942680667 | 374.7768449951233 | 358.48094530253945 | 446.1497065822305 | 426.1596026548004 | 264.6989773557318 | 424.45696280043114 | 486.2476321995374 | 289.9248862803885 | 665.070220293115 |
| SE_85_4_39 | SE_85 | SE_85_4 | TCTCACTGTGTCACCATGGG | 1043.6770985458318 | 1123.3171527284994 | 1238.3586857437183 | 1147.7540877975653 | 1256.804497584347 | 986.7272151401176 | 997.6008880328282 | 1364.145086872575 | 740.5797217061915 | 699.846667033817 | 1272.3829017325552 | 841.4937342277633 |
| SE_85_4_7 | SE_85 | SE_85_4 | TTATCAGACATTCTGAACCT | 241.68223569649066 | 202.87667438068092 | 254.78193055971235 | 235.53683661151848 | 199.39176839312847 | 162.86630065526083 | 224.91756806781132 | 169.21827480955713 | 402.2573308548019 | 185.89677093085763 | 338.0500698330036 | 141.60308355280984 |
| SE_85_4_9 | SE_85 | SE_85_4 | GCAGCCACTTTAAAACCAAG | 1393.7370184111523 | 1467.1081674425595 | 1390.437822589438 | 1293.500638629002 | 1379.8334610609581 | 1573.2191595210302 | 1225.7469379389336 | 1661.9314363977733 | 1162.3727286731473 | 1369.8040493297312 | 1532.9631638954954 | 1055.059040569706 |
| SE_86_1_11 | SE_86 | SE_86_1 | AGTTTCCTAGTCATTCACCA | 21.675536833765978 | 32.97995199291857 | 53.32645058226538 | 20.820935833062407 | 22.272484767317543 | 0.8663101098684087 | 8.60928490211718 | 17.961716320567515 | 139.41368861855165 | 2.187020834480678 | 70.42709788187574 | 20.89225822910309 |
| SE_86_1_18 | SE_86 | SE_86_1 | TTACAATGCCCTAGAAACAA | 392.3272166911642 | 383.76671409941616 | 350.5720362352631 | 329.2310478602993 | 310.7541922297162 | 356.9197652657844 | 362.6661265016862 | 332.76442867577714 | 353.4181405744176 | 260.2554793032007 | 315.7481555037429 | 543.1987139566803 |
| SE_86_1_2 | SE_86 | SE_86_1 | GCTGATGTAATATAGTGTTG | 3728.192335407748 | 3842.664103417331 | 4088.361211307012 | 3989.8118290105836 | 3851.018675720476 | 4052.598693964416 | 3814.9893722506754 | 3877.840018261471 | 4235.68977522606 | 4137.114411892616 | 4157.546344960065 | 5133.692119295721 |
| SE_86_1_3 | SE_86 | SE_86_1 | ATATAGTGTTGGGGGAGCAG | 133.30455152766078 | 145.91130275654885 | 143.19139508200888 | 100.20075369661284 | 195.14939034221084 | 56.31015714144657 | 202.31819519975372 | 123.84130726286024 | 103.89427750554486 | 39.36637502065221 | 149.070690516637 | 80.08698987822851 |
| SE_86_1_4 | SE_86 | SE_86_1 | TCAAACTGTCCTTTCACAGG | 917.9589849099892 | 883.4629564163642 | 976.6640671455641 | 831.5361248329299 | 942.8685218164426 | 1195.5079516184041 | 772.6833199650168 | 933.0638951789546 | 1116.1974942262384 | 864.6022365646946 | 1205.477158744773 | 1751.4676482064756 |
| SE_86_2_13 | SE_86 | SE_86_2 | CTCAATTGTTAATTGCCTGG | 54.188842084414944 | 83.94896870924728 | 83.93978332393624 | 68.96934994701923 | 104.99885676021127 | 77.10159977828837 | 46.27490634887984 | 126.67736773452879 | 77.25471917078977 | 53.94651391719005 | 49.29896851731302 | 95.17584304369186 |
| SE_86_2_18 | SE_86 | SE_86_2 | ACTATGTTGTCCCTGTGCTG | 390.15966300778763 | 463.7181128701278 | 439.4494538723721 | 392.9951638490529 | 417.87423801538625 | 308.4063991131535 | 340.0667536336286 | 479.2942197119858 | 371.17784613092095 | 310.55695849625624 | 546.983793549235 | 299.455701283811 |
| SE_86_2_20 | SE_86 | SE_86_2 | GCCTTCTCATCCACAGCACA | 858.3512586171328 | 884.4623489009981 | 880.8739614700132 | 856.2609861346915 | 727.567835732373 | 963.3368421736706 | 874.9185781776584 | 882.9601601794768 | 933.2725269942534 | 1207.9645075781611 | 630.3225260427879 | 1477.5469292026796 |
| SE_86_2_21 | SE_86 | SE_86_2 | TAGCACTTCATAGTGCACCA | 169.06918730337463 | 139.91494784874547 | 112.57806234033802 | 162.66356119580004 | 184.54344521491677 | 142.94116812828744 | 156.0432888508739 | 149.36585150787724 | 156.28540889722987 | 94.77090282749604 | 185.45802442227279 | 467.7544481293636 |
| SE_86_2_5 | SE_86 | SE_86_2 | AGACACCCAAGGCTAAGAAA | 241.68223569649066 | 298.81835290553494 | 385.13547642747216 | 217.31851775758886 | 374.38986299348056 | 376.8448977927578 | 161.42409191469713 | 364.90644735468743 | 380.9456841869978 | 103.51898616541875 | 720.7039683245284 | 451.50491395117234 |
| SE_86_3_11 | SE_86 | SE_86_3 | ACCAACTCTGACAAGTCTGA | 389.0758861660993 | 407.7521337306296 | 426.61160465812304 | 416.41871666124814 | 389.2381861716923 | 524.9839265802557 | 394.95094488462564 | 640.0043131065372 | 544.334975306829 | 266.0875348618158 | 765.3077969830497 | 398.11358736568667 |
| SE_86_3_17 | SE_86 | SE_86_3 | TAGCACATAGAGATATGAAC | 683.8631871053166 | 801.5127726763848 | 788.0464363823661 | 632.4359259292706 | 771.0522107542787 | 636.7379307532805 | 524.0902184163833 | 608.8076479181832 | 822.2743672661072 | 438.8621807857894 | 491.81590020843225 | 768.3708304259026 |
| SE_86_3_6 | SE_86 | SE_86_3 | AGATCCAGCAGAGACAGCAG | 2684.5152368619165 | 3225.039547913583 | 3145.273057491023 | 3160.8783211567866 | 3201.934833930079 | 3551.0051403506072 | 3367.306557340582 | 3325.7535797766586 | 3606.1082132480146 | 3682.943085265462 | 3304.20467562467 | 4136.667129362412 |
| SE_86_3_8 | SE_86 | SE_86_3 | GTCTCTGCTGGATCTCCACT | 60.69150313454474 | 115.92952821753195 | 90.85247136237805 | 63.76411598875362 | 156.9679878839522 | 147.27271867762948 | 32.28481838293943 | 97.37140952728706 | 111.88614500597139 | 64.15261114476655 | 31.692194046844083 | 11.606810127279495 |
| SE_86_3_9 | SE_86 | SE_86_3 | GGAGATTGTGAAAACAACCT | 479.0293640262281 | 532.6761943098667 | 490.8008507293684 | 569.9731184300834 | 549.3879575938327 | 686.1176070157798 | 714.570646875726 | 771.408448293847 | 443.9926389125849 | 771.2893476268524 | 658.4933651955382 | 583.8225494021585 |

| | | | | | | | | | | | | | | | |
|---|---|---|---|---|---|---|---|---|---|---|---|---|---|---|---|
| SE_87_1_29 | SE_87 | SE_87_1 | GGTTGAGAACCACTGCCACC | 1008.9962396118062 | 1056.3578562580283 | 900.6244987227042 | 947.3525804043395 | 1027.716082834795 | 801.336851628278 | 1092.3030219561172 | 1315.9320588542096 | 1023.8470253324208 | 1304.922431240138 | 1250.0809874032943 | 839.1723722023074 |
| SE_87_1_4 | SE_87 | SE_87_1 | GCACTGGCTTCTAGTTTGAG | 1279.940450033881 | 1436.127000418909 | 1450.6769612101452 | 1540.7492516466182 | 1420.1360525446755 | 1221.4972549144563 | 1153.6441768837021 | 1496.494575550441 | 1059.3664364454276 | 1260.4530076056974 | 1471.926345731203 | 1398.6206203371792 |
| SE_87_1_6 | SE_87 | SE_87_1 | TACTTCATAGAGATGTTGCG | 327.30060618986624 | 368.7758268299077 | 414.7612823065085 | 286.2878677046081 | 404.08650934990396 | 382.90906856183665 | 290.56336544645484 | 239.1744331107148 | 336.5464202957393 | 672.873410075222 | 302.83652089206566 | 302.9377443219948 |
| SE_87_1_8 | SE_87 | SE_87_1 | AGCGTGTACTGGCACCTAGA | 3482.1749923445045 | 3392.9374853320774 | 3320.065312177337 | 3268.8869257907977 | 3210.419590031914 | 3404.5987317828462 | 2970.203291230427 | 2988.2623836481007 | 3166.5555007245553 | 3477.363126824278 | 3030.712778850053 | 3399.634686280164 |
| SE_87_1_9 | SE_87 | SE_87_1 | GATTGATCTGATGGGCACGA | 280.6982019972694 | 361.7800794374704 | 354.5221436858013 | 308.4101120272369 | 363.7839178661865 | 206.1818061486813 | 368.04692956550946 | 389.4856381091482 | 538.1190783620528 | 1056.3310630541675 | 470.68777084386954 | 549.00211902032 |
| SE_87_2_13 | SE_87 | SE_87_2 | TGTTTCTGCTGTGAAGTCAC | 31.429528408960667 | 45.972054293159225 | 53.32645058226538 | 36.43663770785921 | 40.30259148371746 | 77.96790988815678 | 1.0761606127646475 | 12.289595377230405 | 44.39926389125849 | 2.187020834480678 | 2.347569929395858 | 82.4083519036844 |
| SE_87_2_16 | SE_87 | SE_87_2 | CTCAGTGCTCAACTTCCTGA | 574.4017260947984 | 516.6859145557244 | 518.4516028831356 | 459.36189681693935 | 564.2362807720444 | 652.3315127309118 | 619.868512952437 | 665.5288573515543 | 626.9176061445698 | 884.2854240750208 | 956.6347462288122 | 542.0380329439524 |
| SE_87_2_17 | SE_87 | SE_87_2 | CATCAGGAAGTTGAGCACTG | 3241.576533489702 | 3182.0656710743256 | 3297.3521943367427 | 3076.293269334971 | 3305.873096177561 | 3486.0318821104765 | 3222.0248746173547 | 3287.9394401544114 | 3527.077523521574 | 3138.3748974797727 | 3767.849736680352 | 3237.139344498251 |
| SE_87_2_18 | SE_87 | SE_87_2 | GCACTGAGGTTAATACACAC | 1581.2304120232282 | 1686.9745140620169 | 1770.6356647037373 | 1480.8890611265638 | 1630.1337660650981 | 2196.0961285164162 | 1558.2805672832096 | 1290.4075146091925 | 1636.556867031788 | 1646.8266883639506 | 1378.0235485553687 | 1908.159584924749 |
| SE_87_2_19 | SE_87 | SE_87_2 | ACTGTTGAGCTAGAGGAAGA | 571.1503955697335 | 627.6184803500868 | 576.7156877785737 | 494.4972260352322 | 849.5362046962548 | 436.620295373678 | 463.82522410156304 | 777.080569237184 | 491.94384391514404 | 387.10268770308 | 807.5640557121752 | 761.4067443495348 |
| SE_87_3_13 | SE_87 | SE_87_3 | GCCCAATGACATTTACAGGG | 466.0240419259685 | 746.5461860215205 | 708.0567605089681 | 573.8770438987826 | 724.3860521941848 | 616.812798226307 | 572.5174459907925 | 708.0697644265825 | 723.7080014275134 | 673.6024170200488 | 775.871861665331 | 699.890650674953 5 |
| SE_87_3_19 | SE_87 | SE_87_3 | ATCAACAGCGACTCAGGCAT | 836.6757217833667 | 882.4635639317303 | 921.3625628380296 | 871.8766880094882 | 926.9596041255015 | 651.4652026210433 | 897.517951045716 | 903.7579369717129 | 864.8976606017154 | 637.8810767235311 | 751.2223774066746 | 1484.5110152790473 |
| SE_87_3_21 | SE_87 | SE_87_3 | ACAGCGACTCAGGCATAGGT | 13245.92055911439 | 12064.666074500396 | 11994.501273559172 | 13303.276688837312 | 12166.079655519025 | 10192.138442601829 | 14304.326864867695 | 13220.768565428247 | 11771.132842850451 | 12525.79732601567 | 12409.254646786505 | 12006.084395657908 |
| SE_87_3_6 | SE_87 | SE_87_3 | CTGTCCCCTGAAAGACCGCA | 227.59313675454277 | 262.8402234587147 | 259.7195648728851 | 317.5192714542017 | 310.7541922297162 | 431.42243471446756 | 258.2785470635154 | 150.31120499843342 | 269.9475244588516 | 387.8316946479069 | 178.4153146340852 | 522.3064557275773 |
| SE_87_3_8 | SE_87 | SE_87_3 | AGTAACACGTAAACACCTCA | 429.1756293085664 | 355.78372452966704 | 419.6989166196812 | 379.9820789533889 | 424.2378050917627 | 550.1069197664395 | 429.38808449309437 | 290.2235216007488 | 356.08209640789306 | 321.4920626686597 | 442.5169316911192 | 507.2176025621139 |
| SE_88_1_1 | SE_88 | SE_88_1 | GCAATTGCTGGAAGAACCCT | 1560.6386520311505 | 1652.9951695844643 | 1623.4941621711903 | 1518.6270073239893 | 1731.9508392871212 | 1977.785980829577 | 1647.6018981426753 | 1640.188306114981 | 1662.3084400887178 | 1767.112834260388 | 1736.0279627882371 | 1708.5224507355415 |
| SE_88_1_2 | SE_88 | SE_88_1 | ACTTCCCCAGAGGTCCAGCA | 134.38832836934907 | 151.90765766435223 | 145.16644880727796 | 123.62430650880805 | 139.9984756802817 | 124.74865582105086 | 180.79498294446077 | 107.7702979234051 | 25.751573056929924 | 106.43501394472632 | 76.29602270536539 | 92.85448101823596 |
| SE_88_1_25 | SE_88 | SE_88_1 | ACACACCATACAGATAACCA | 2048.338230790885 | 2125.7078148162973 | 2146.883399367499 | 2036.5477861714166 | 2197.5518303753306 | 2131.1228702762855 | 1841.3108084403118 | 1840.6032461128923 | 1731.571291759081 | 2891.241543183456 | 1655.03680022408 | 2296.987724188612 |
| SE_88_1_27 | SE_88 | SE_88_1 | GCCTGCAACAGTCCAAACTG | 2801.563135764253 | 2713.350595781028 | 2729.52424832188 | 2710.6255837668123 | 2977.088797231445 | 2506.2351478493065 | 2657.0405529159148 | 3005.278746478112 | 2806.9214632053618 | 3102.653557183255 | 3298.3357508011804 | 2551.176865976033 |
| SE_88_1_3 | SE_88 | SE_88_1 | CATGTCTGAGGAAGGCTGGT | 3019.4022809436005 | 2846.2697962373363 | 2761.125107926185 | 3115.332524021963 | 2828.605565449328 | 3409.796592442057 | 2694.7061743626773 | 2260.340195919838 | 2698.587259310691 | 3018.0887515833356 | 1967.263600833729 | 2801.88396472527 |
| SE_88_2_17 | SE_88 | SE_88_2 | AGCTAGACTTTGGGAACAAG | 414.0027535249302 | 538.67254921767 | 506.60128053152107 | 443.74619494214255 | 575.9028204120679 | 291.08019691578534 | 305.62961402515987 | 523.7258337681264 | 411.1371836330536 | 319.305041834179 | 482.4256204908488 | 371.4179240729438 3 |
| SE_88_2_18 | SE_88 | SE_88_2 | GAGCTGACTGTCACTGCAGG | 278.5306483138928 | 72.95565137827442 | 141.2163413567398 | 203.00412437235846 | 84.84756101835254 | 696.5133283342007 | 408.94103285056605 | 207.0324144318045 | 535.4551225285774 | 27.702263903421922 | 76.29602270536539 | 181.06623798556012 |
| SE_88_2_31 | SE_88 | SE_88_2 | CTAAGATTTAGAAGCAAAGA | 28.17819788389577 | 36.977521931454156 | 39.50107450538176 | 31.23140374959361 | 22.272484767317543 | 11.262031428289314 | 15.066248578705064 | 30.25131169779792 | 59.49501361428638 | 14.580138896537854 | 4.695139858791716 | 5.803405063639747 |
| SE_88_2_33 | SE_88 | SE_88_2 | TGGCAAGGGAGCACACACAG | 296.9548546225939 | 326.80134247528406 | 287.3703170266523 | 283.68525072547527 | 274.6939787969163 | 249.49731164210172 | 269.0401531911619 | 279.82463320463074 | 202.4606433441387 | 325.86610433762104 | 288.75110131569056 | 458.4690002754 |
| SE_88_2_8 | SE_88 | SE_88_2 | GCACAAAGCCCATGATAACA | 185.3258399286991 | 183.8882171726369 | 124.42838469195254 | 169.17010364363205 | 172.8769055748933 | 248.6310015322333 | 171.10953742957895 | 83.19110716894428 | 110.11017445032105 | 51.030486137882484 | 112.68335661100119 | 55.71268861094157 |
| SE_88_3_12 | SE_88 | SE_88_3 | GTGGCACCTGAGCGGCAACC | 178.8231788785693 | 216.86816916555549 | 195.5303188016397 | 150.95178478970246 | 199.39176839312847 | 144.67378834802426 | 156.0432888508739 | 154.09261896065814 | 256.62774529147407 | 119.5571389516104 | 52.820323411406804 | 306.41978736017865 |
| SE_88_3_25 | SE_88 | SE_88_3 | CTAGACACCTCATTCAAGCC | 895.1996712345349 | 793.5176327993136 | 849.2731018657079 | 788.5929446772386 | 944.9897108419015 | 826.4598448144619 | 772.6833199650168 | 832.856425179999 | 814.2824997656807 | 966.6632088404597 | 1349.8527094026183 | 925.0627671441757 |
| SE_88_3_26 | SE_88 | SE_88_3 | TTTGGGGAAGGCAGGAAATG | 215.67159149597148 | 253.84569109700962 | 268.607306636596 | 258.9603894237137 | 274.6939787969163 | 155.93581977631356 | 214.15596194016484 | 181.50787018678753 | 218.44437834499178 | 376.1675835306766 | 190.1531642810645 | 268.1173139401563 |
| SE_88_3_28 | SE_88 | SE_88_3 | GCAGGAAATGAGGCTCAGAT | 4771.8694339535805 | 5001.9593855926505 | 4996.8859249307925 | 4947.574877331454 | 4944.491618344494 | 5299.218942065057 | 4472.523506649875 | 4419.527568350165 | 4651.266885248239 | 4282.186793913167 | 4500.29155465186 | 5574.750904132341 |
| SE_88_3_42 | SE_88 | SE_88_3 | CCTTTGTTGTTCAAGCCGGG | 83.45081680999901 | 117.92831318679976 | 137.26623390620162 | 84.58505182181602 | 116.66539640023474 | 223.50800834604945 | 94.70213392328898 | 122.89595377230405 | 97.67838056076867 | 88.20984032405401 | 230.0618530807941 | 97.49720506914775 |
| SE_88_4_11 | SE_88 | SE_88_4 | AGCTGTGCCAGTCTTTCCAA | 468.19159560934514 | 510.68955964792093 | 502.6511730809829 | 472.37498171260336 | 440.1467227827038 | 553.5721602059132 | 401.4079085612135 | 536.0154291453568 | 460.86435919126313 | 556.961305847746 | 849.8203144413006 | 826.4048810623 |
| SE_88_4_24 | SE_88 | SE_88_4 | GCTGGGGGTTTCCACTGCCT | 1330.877961593231 | 1424.1342906033021 | 1560.2924429625796 | 1535.5440176883526 | 1406.3483238791935 | 1381.7646252401119 | 1366.7239782111023 | 1547.5436640404748 | 1459.8477967445792 | 1365.43000766077 | 1443.7555065784527 | 1246.5714076698177 |
| SE_88_4_25 | SE_88 | SE_88_4 | GCCAAGCACAGGCCTCCCCA | 398.829877741294 | 597.6367058110699 | 593.5036444433609 | 497.099843014365 | 628.9325460485383 | 570.0320522934129 | 379.88469630592056 | 805.4411739538697 | 533.679151972927 | 701.3046809234708 | 455.42856630279647 | 224.01143545649424 |
| SE_88_4_4 | SE_88 | SE_88_4 | ATTCTGTGAGAGGGGCATAC | 13.005322100259587 | 23.98541963121351 | 25.675698428498144 | 6.506542447832002 | 36.06021343279983 | 25.12299318618385 | 2.152321225529295 | 4.726767452780925 | 4.439926389125849 | 0.7290069448268927 | 1.173784964697929 | 0.0 |
| SE_88_4_7 | SE_88 | SE_88_4 | CCTGGCACAGTGAGGCCAGG | 1370.9777047356981 | 1226.2545786457906 | 1174.169439672473 | 1245.3522245150452 | 1176.199314616912 | 834.2566358032776 | 1320.4490718622224 | 1227.0688307419282 | 1499.8071342467117 | 1382.1971673917885 | 1122.1384262512202 | 1091.0401519642726 |
| SE_88_5_11 | SE_88 | SE_88_5 | TTAAGAGAAGAGCAGCCGCG | 393.4109935328525 | 492.7004949245108 | 491.7883775920029 | 425.52787608821296 | 523.933689288327 | 613.3475577868334 | 331.4574687315114 | 575.7202757487166 | 364.0739639083196 | 463.64841690990374 | 585.7186973842665 | 358.6504329329364 |
| SE_88_5_16 | SE_88 | SE_88_5 | AAACAGGCAGTGGAAGACAG | 662.1876502715506 | 577.648856118392 | 586.5909564049192 | 675.3791060849618 | 675.5987046086321 | 1030.042720633538 | 663.9910980757875 | 829.0750112177742 | 768.9952505965971 | 885.0144310198477 | 880.3387235234468 | 685.9624785222181 |
| SE_88_5_2 | SE_88 | SE_88_5 | TCACCCCCGAGACCAGCCCA | 868.1052501923274 | 730.5559062673781 | 810.7595542229607 | 886.1910813947187 | 662.8715704558792 | 784.0106494309099 | 888.9086661435988 | 852.7088484816788 | 588.7342391980876 | 891.5754935232898 | 591.5876222077562 | 596.590040542166 |
| SE_88_5_20 | SE_88 | SE_88_5 | CCAGCAGGCTGGGAGCACGG | 1137.9656837727139 | 957.4180002792725 | 1062.5789041947694 | 1067.0729614444483 | 1144.38147923503 | 1237.0908368920877 | 988.991603130711 | 1094.7193420640622 | 1203.220051453105 | 902.5105976956931 | 1379.1973335200666 | 1534.4202988263492 |
| SE_88_5_7 | SE_88 | SE_88_5 | GAGACCAGCCCAGGGCCTCG | 638.344559754408 | 785.5224929222423 | 842.360413827266 | 743.0471475424147 | 722.264863168726 | 738.9625237177527 | 615.5638705013783 | 659.8567364082171 | 827.6022789330582 | 620.3849100476856 | 813.4329805356648 | 878.6355266350578 |
| SE_88_6_10 | SE_88 | SE_88_6 | GTCACATGTCAGCTCCCAAG | 1043.6770985458318 | 1343.1834993479565 | 1241.321266331622 | 1064.4703444653155 | 1250.4409305079705 | 1333.251259087481 | 1018.0479396753565 | 1183.5825701763436 | 1007.8632903315677 | 1625.6854869639706 | 1453.1457862960362 | 883.2782506859695 |
| SE_88_6_20 | SE_88 | SE_88_6 | TCACTCCTTTGCTCAGCACG | 693.6171786805113 | 659.5990398583715 | 633.9922458113773 | 625.9293834814387 | 800.7488571107021 | 694.7807081144638 | 695.1997558459623 | 903.7579369717129 | 577.1904305863603 | 762.5412642889297 | 953.1133913347184 | 958.7225165132862 |
| SE_88_6_43 | SE_88 | SE_88_6 | GATGTTGCCCTCAGACCACA | 3308.7706976743766 | 2765.319004981991 | 2726.5616677339763 | 3153.0704702193884 | 2736.3338428418692 | 3080.5987506920615 | 3331.7932571193487 | 2861.585015913572 | 3186.9791621145346 | 2168.066653915179 | 2826.4741949926133 | 2739.2071900379606 |
| SE_88_6_46 | SE_88 | SE_88_6 | GTGAGTCAGTTGTCTCCCTG | 1080.5255111632341 | 1026.3760817190114 | 995.4270775356204 | 972.0774417061011 | 1046.8067840639244 | 1059.497264369064 | 1086.922218892294 | 936.8453091411793 | 1186.348331174427 | 1180.2622436747392 | 974.2415206992811 | 1140.9494355115744 |
| SE_88_6_9 | SE_88 | SE_88_6 | AGTCACATGTCAGCTCCCAA | 457.3538271924621 | 606.6312381727749 | 642.8799875750882 | 458.06058832737295 | 547.2667685683739 | 582.1603938315707 | 456.29209981221055 | 660.8020898987734 | 570.086548363759 | 469.4804724685189 | 657.3195802308403 | 600.0720835803498 |
| SE_88_7_13 | SE_88 | SE_88_7 | GTATGTGCAAAACAGACCTC | 1120.625254305701 | 1021.3791192958419 | 1062.5789041947694 | 1057.9638020174837 | 1074.382241394889 | 1267.4116907374819 | 1123.511679726292 | 1410.4674079098281 | 1068.2462892236792 | 986.3463963507858 | 1071.66567276920 9 | 932.0268532205434 |
| SE_88_7_14 | SE_88 | SE_88_7 | GGAAGGCTAATTGAGGGCAT | 3886.42375429424 | 3591.816589774223 | 3531.3960607811296 | 3745.1658329721004 | 3481.9317852906424 | 3638.502461447317 | 4099.095774020542 | 3682.1518457163406 | 3142.579898223276 | 4052.5496062926964 | 3454.449151106005 | 3461.1507799547453 |
| SE_88_7_23 | SE_88 | SE_88_7 | TGAAGATCGGAGACTGCACG | 2072.1813213080277 | 2086.731507915575 | 2182.4343664223425 | 2164.076018148924 | 2093.613568127849 | 2065.283301926286 | 2159.8543498186473 | 2324.6242332776587 | 2124.060784557806 | 2859.165237611073 | 2015.3887843863442 | 1770.038544410123 |
| SE_88_7_5 | SE_88 | SE_88_7 | TTGAGTTGACGCTACCCACG | 680.6118565802517 | 684.5838519742189 | 694.2313844320845 | 610.3136816066418 | 755.1432930633376 | 866.3101098684087 | 569.2889641524985 | 553.9771454659244 | 625.1416355889195 | 433.0301252271742 | 471.86155580856746 | 638.3745570003722 |
| SE_88_7_6 | SE_88 | SE_88_7 | GCAGGAAGCACACTATGACA | 18552.0919760203 | 15807.390929454335 | 16296.168287195245 | 17238.433561286107 | 15781.646349413573 | 16359.40011475503 | 18817.744474802625 | 17608.154115099504 | 15024.710900801872 | 18256.520919299874 | 17138.434269554462 | 17551.81827447205 |
| SE_88_8_13 | SE_88 | SE_88_8 | TGTGTCACTGGGCAGCTATA | 825.8379533664838 | 914.444123440015 | 976.6640671455641 | 917.4224851443123 | 968.3227901219484 | 1039.5721318420906 | 772.6833199650168 | 836.6378391422237 | 1077.126142001931 | 1031.5448269300532 | 881.5125084881447 | 1384.6924481844437 |
| SE_88_8_21 | SE_88 | SE_88_8 | CGTGATTCTCCGGGCAACCA | 253.60378095506195 | 302.81592284407054 | 316.99612290568865 | 262.8643148924129 | 286.3605184369398 | 406.2994415282837 | 269.0401531911619 | 199.46958650735505 | 232.6521427901945 | 252.23640291010486 | 260.5802621629402 4 | 302.9377443219948 |
| SE_88_8_22 | SE_88 | SE_88_8 | GTCTCCGGTCATACACCAGG | 99.7074694353235 | 113.93074324826416 | 141.2163413567398 | 131.43215744620645 | 107.12004578567009 | 123.01603560131404 | 81.7882065701132 | 57.66656292392729 | 168.71720278678225 | 62.69459725511277 | 88.03387235234467 | 222.85075444376628 |
| SE_88_8_35 | SE_88 | SE_88_8 | CCTCTTCCAGGTGTGAACCG | 224.34180622947787 | 200.87788941141312 | 243.91913507073238 | 231.63291114281927 | 290.60289648785744 | 251.22993186183854 | 177.56650110616684 | 352.61685197745703 | 244.1959514019217 | 194.64485426878034 | 590.4138372430583 | 140.44240254008187 |
| SE_89_1_13 | SE_89 | SE_89_1 | TCCTAATTCAGCACAAGGAA | 320.7979451397365 | 326.80134247528406 | 359.459777998974 | 368.27030254729135 | 369.08689042983355 | 318.8021204315744 | 245.3646197103396 | 380.9774566941426 | 420.0170364113053 | 335.34319462037064 | 341.5714247270973 4 | 459.629681040268 |
| SE_89_1_17 | SE_89 | SE_89_1 | GTGGACACCCTGATGATGCA | 358.73013459882696 | 262.8402234587147 | 209.35569487852334 | 264.1656233819793 | 341.51143309886896 | 205.31549603881288 | 334.68595056980536 | 457.55108942919355 | 231.7641575123693 | 391.47672937204135 | 490.64211524373434 | 213.5653063419427 |
| SE_89_1_18 | SE_89 | SE_89_1 | TATGGTTTTCTTATGAACCA | 61.77527997623304 | 69.95747392437274 | 77.02709528549444 | 53.35364807222242 | 60.45388722557618 | 44.18181560328885 | 48.42722757440914 | 82.2457536783881 | 43.51127861343332 | 8.019076393095819 | 23.47569929395858 | 34.820430381838484 |
| SE_89_1_20 | SE_89 | SE_89_1 | CAGGGTGTCCACCTTTAGAA | 2097.1081886668585 | 2139.6993096011715 | 2137.0081307411533 | 2220.032283200279 | 2215.5819370917307 | 2388.4169729072028 | 2019.9534701592434 | 2302.8811029948665 | 2324.7454573462946 | 2127.9712719496997 | 2220.8011532084815 | 2958.575901443543 |
| SE_89_1_5 | SE_89 | SE_89_1 | TGCCTGGCCCTTATTCCCAA | 869.1890270340157 | 866.473284177588 | 906.5496598985114 | 920.0251021234451 | 985.2923023256188 | 1104.545390082221 | 733.9415379054896 | 1036.1074256495788 | 998.983437553316 | 808.4687018130239 | 996.5434350285417 | 1664.4165722518794 |
| SE_89_2_0 | SE_89 | SE_89_2 | AGTGGGGCAGGAGAGATGAC | 2491.602959041399 | 2318.590564350639 | 2218.9728603398203 | 2273.3859312725017 | 2054.3715711568607 | 1789.7966869881325 | 2425.6660211715152 | 2060.870609412483 | 2052.1339770539676 | 2259.1925220185403 | 2241.9292825730445 | 1929.051843153852 |
| SE_89_2_11 | SE_89 | SE_89_2 | CATAGAGGAGGAAAGGTGCA | 5043.897421217343 | 4721.130097410525 | 4368.818840295223 | 4566.291489888499 | 4546.768676070967 | 4055.1976242940214 | 4770.6199963856825 | 4740.002401648711 | 5205.369698611145 | 4024.1183354444474 | 4888.814377966874 | 5213.779109173949 |
| SE_89_2_19 | SE_89 | SE_89_2 | AGGAGGGGATGTAAATACAA | 3251.330525064897 | 3434.911969686701 | 3564.9719741107037 | 3375.594221935243 | 3789.5041939821704 | 3342.224403872321 | 3467.389494327694 | 3939.287995147623 | 3846.7522235386355 | 3319.1686197968424 | 4421.6479620170985 | 3485.525081222032 |
| SE_89_2_6 | SE_89 | SE_89_2 | CATCTTTCAGTGCATGTGCA | 1466.3500668042684 | 1327.193219593814 | 1324.2735227929236 | 1408.0157857108452 | 1200.5929884096884 | 1561.0908179828725 | 1445.2837029429215 | 1291.3528680997488 | 1831.0256428755001 | 1184.6362853437006 | 1286.4683213089302 | 1821.1085089701526 |
| SE_89_3_17 | SE_89 | SE_89_3 | TCTCGCGGGGCTGGCTATGC | 18093.65437198615 | 16824.772478811643 | 17251.106763362848 | 16724.416707907378 | 15806.04002320635 | 16075.250398718192 | 16483.552105716106 | 15947.168032192285 | 16760.722118950078 | 15566.48529288864 | 15593.733256011987 | 13554.432866636993 |
| SE_89_3_18 | SE_89 | SE_89_3 | CTCGCGGGGCTGGCTATGCC | 9041.950190205478 | 7985.145952224831 | 8093.770166152723 | 8856.705579988922 | 8084.911970536267 | 7900.748201999888 | 9079.56708989533 | 8416.482126421715 | 8101.977674876849 | 8439.713400260936 | 8718.874717776216 | 8094.58938276472 |
| SE_89_3_19 | SE_89 | SE_89_3 | GCGGGGCTGGCTATGCCGGG | 3848.4915648351493 | 5811.467298146106 | 6032.801603834429 | 4959.286653737552 | 5674.180643102326 | 6325.796422259121 | 4141.066037918364 | 5755.312050506054 | 6033.859962822029 | 5778.838051642778 | 5758.58903680804 | 7558.354754884407 |
| SE_89_3_20 | SE_89 | SE_89_3 | GGGCTGGCTATGCCGGGTGG | 681.69563342194 | 880.4647789624626 | 946.0507344038932 | 832.8374333224963 | 773.1733997797376 | 1538.5667551262939 | 807.1204595734856 | 525.6165407492389 | 996.3194817198405 | 1224.0026603643528 | 430.77908204413995 | 1453.1726279353927 |
| SE_89_3_21 | SE_89 | SE_89_3 | TATGCCGGGTGGCGGCTCCC | 1443.5907531288142 | 2251.6312678801683 | 2269.336730334182 | 1625.3343034684342 | 2230.430260269942 | 2089.539985002602 | 1258.031756321873 | 2514.6402848794523 | 2438.4075729079163 | 1955.1966260257261 | 2234.886572784857 | 2117.08216721578 |
| SE_89_4_14 | SE_89 | SE_89_4 | GAAGCCTGAGACATGAAAGC | 682.7794102636283 | 773.5297831066356 | 720.8946097232172 | 722.2262117093522 | 673.4775155831733 | 926.9518175591974 | 653.229491948141 | 500.09199650422187 | 689.0765755923318 | 997.2815005231892 | 503.5537498554116 | 871.67144055869 |
| SE_89_4_18 | SE_89 | SE_89_4 | GCAACACCAAGTGTAGCCAT | 890.8645638677817 | 872.4696390853913 | 913.4623479369533 | 918.7237936338787 | 852.717988234443 | 898.3635839335399 | 973.925354552006 | 952.9163184806345 | 792.0828678200514 | 814.3007573716391 | 596.28276206654 8 | 456.14763800208414 |
| SE_89_4_2 | SE_89 | SE_89_4 | TCTTAGGGAAAGGAAACTAG | 496.3697934932409 | 492.7004949245108 | 436.48687328446846 | 355.25721765162734 | 564.2362807720444 | 341.32618328815306 | 412.16951468886 | 538.8514896170254 | 462.64032974691344 | 191.72882648947277 | 579.849772560777 | 184.54828102374395 |
| SE_89_4_4 | SE_89 | SE_89_4 | GGTGCTGGCCAAAGAGACAG | 803.0786396910295 | 1008.3870169956012 | 919.3875091127605 | 975.9813671748003 | 1099.8365097003948 | 848.1175975611721 | 835.1006355053664 | 966.1512673484211 | 1161.484743395322 | 1035.1898616541876 | 1210.1722986035647 | 1260.4995798225532 |
| SE_89_4_9 | SE_89 | SE_89_4 | TGACCACCTGAAGCCCTGGG | 538.6370903190846 | 667.5941797354426 | 659.6679442398754 | 545.2482571283218 | 609.8418448194088 | 626.3422094348595 | 544.5372700589116 | 469.84068480642395 | 809.8425733765548 | 822.3198337647349 | 308.70544571555536 | 403.9169924293264 |
| SE_89_6_0 | SE_89 | SE_89_6 | AATTTATTAGACATCTACTG | 1.083776841688299 | 5.9963549078033775 | 4.93763431317272 | 2.602616979132801 | 4.242378050917627 | 2.598930329605226 | 2.152321225529295 | 0.0 | 5.327911666951019 | 0.0 | 0.0 | 0.0 |
| SE_89_6_11 | SE_89 | SE_89_6 | TATGGTCTCCTAATTCCAGC | 380.4056714325929 | 418.7454510616025 | 406.86106740543215 | 438.54096098387697 | 429.5407776554097 | 280.68447559736444 | 320.69586260386495 | 360.17967990190647 | 496.3837703042699 | 333.15617378588996 | 488.2945453143385 | 357.48975192020845 |
| SE_89_6_23 | SE_89 | SE_89_6 | TTTTAACAGGAGATTTGGTG | 9.75399157519469 | 10.993317330972857 | 15.800429802152705 | 5.205233958265602 | 3.1817835381882205 | 33.78609428486794 | 0.0 | 7.56282792444948 | 25.751573056929924 | 0.7290069448268927 | 0.0 | 10.446129114551544 |
| SE_89_6_27 | SE_89 | SE_89_6 | GCAAGAGAGAAAGTGGGGGG | 739.1358060314199 | 1363.1713490406344 | 1344.0240600456145 | 913.5185596756131 | 1318.3189793226527 | 962.4705320638021 | 783.4449260926633 | 1422.7570032870585 | 1419.8884592424465 | 1239.3118062057174 | 1422.62737721389 | 1145.5921595624861 |
| SE_89_6_30 | SE_89 | SE_89_6 | CTTCCGGGAGTAATGCTGTA | 319.71416829804815 | 293.8213904823655 | 304.15827369143955 | 337.0388987976977 | 298.02705807696327 | 157.66843999605038 | 341.1429142463933 | 397.9938195241539 | 215.78042251151626 | 341.1752501789858 | 473.0353407732654 | 96.3365240564198 |
| SE_9_1_11 | SE_9 | SE_9_1 | ATAAGGTTTTAGTTGGCCAG | 128.96944416090756 | 189.88457208044028 | 224.1685978180415 | 156.15701874796804 | 129.39253055298764 | 130.81282659012973 | 102.23525821264151 | 115.33312584785457 | 147.40555611897818 | 110.0800486688608 | 174.89395973999143 | 229.814840520134 |
| SE_9_1_23 | SE_9 | SE_9_1 | GCAACCCTGGAGCATTAGCC | 1747.048268801538 | 1782.9161925868707 | 1679.7831933413595 | 1683.893185498922 | 1582.4070129922748 | 2133.7218006058906 | 1987.668651776304 | 1660.040729416661 | 1622.349102586585 | 1285.2392437298117 | 1660.9057250475696 | 1895.3920937847415 |
| SE_9_1_26 | SE_9 | SE_9_1 | CTGCTGTTGCCACTGGTGGA | 112.71279153558308 | 182.888824688003 | 172.81720096104522 | 183.48449702886245 | 127.2713415275288 | 278.9518553776276 | 111.92070372752333 | 242.01049358238336 | 177.59705556503397 | 119.5571389516104 | 106.81443178751154 | 169.4594278582806 |

| | | | | | | | | | | | | | | | |
|---|---|---|---|---|---|---|---|---|---|---|---|---|---|---|---|
| SE_9_1_32 | SE_9 | SE_9_1 | TGACATTACTGTTATTACTA | 405.3325387914238 | 539.6719417023039 | 527.3393446468465 | 426.82918457777936 | 533.4790399028916 | 422.7593336157835 | 476.7391514547388 | 524.6711872586827 | 420.0170364113053 | 497.1827363719408 | 446.03828658521303 | 461.95104306572387 |
| SE_9_1_9 | SE_9 | SE_9_1 | TTAGAAATTAGGCCTGAGAG | 31.42952840896067 | 58.96415659339988 | 50.36386999436174 | 28.62878677046081 | 58.33269820011737 | 54.57753692170975 | 25.827854706515374 | 71.84686528227006 | 114.5501008394469 | 8.748083337922711 | 23.47569929395858 | 62.67677468730927 |
| SE_9_2_10 | SE_9 | SE_9_2 | GCAGTACAGATACCATCACC | 10665.44789905455 | 9038.505631028956 | 9244.238961121966 | 10668.126997465351 | 9280.20198638231 | 9653.293554263679 | 10747.616039680534 | 9270.13632839395 | 8963.323394367264 | 10761.60051953459 | 10681.443178751155 | 9531.51247652192 |
| SE_9_2_11 | SE_9 | SE_9_2 | GGATGGACATGGAAGAGAAA | 732.6331449812901 | 893.4566881267032 | 917.4124553874914 | 808.1125720207347 | 937.5655492525796 | 809.1336426170938 | 790.9780503820159 | 951.9709649900783 | 867.5616164351909 | 694.0146114752018 | 1001.2385748873335 | 1178.0912279188688 |
| SE_9_2_13 | SE_9 | SE_9_2 | GTTGGGAGCAAGCAGAGAAT | 700.1198397306411 | 815.5042674612592 | 814.7096616734988 | 704.0078928554226 | 794.3852900343256 | 654.930443060517 | 738.2461803565482 | 761.9549133882851 | 990.1035847750643 | 849.2930907233299 | 859.2105941588841 | 1246.5714076698177 |
| SE_9_2_25 | SE_9 | SE_9_2 | GATCCTAACCCCTAAGGTGA | 117.04789890233629 | 96.94107100948793 | 124.42838469195254 | 128.82954046707366 | 117.72599091296415 | 139.4759276888138 | 117.30150679134657 | 90.75393509339376 | 149.18152667462851 | 102.06097227576497 | 72.7746678112716 | 92.85448101823596 |
| SE_9_2_6 | SE_9 | SE_9_2 | AGGACTAATCCCCTTCATGA | 1178.065426915181 | 1304.2071924472345 | 1422.0386821937434 | 1311.7189574829317 | 1472.1051836684167 | 1462.3314654578474 | 1323.6775537005165 | 1401.9592264948224 | 1393.2489009076914 | 1475.5100563296307 | 1484.8379803428802 | 1604.0611595900261 |
| SE_9_3_0 | SE_9 | SE_9_3 | ATATGTATACAAATCATGTG | 50.93751155935005 | 35.97812944682026 | 52.338923719630834 | 44.24448864525762 | 14.848323178211695 | 51.11229648223612 | 32.28481838293943 | 44.431614056140695 | 86.13457194904147 | 5.103048613788249 | 31.692194046844083 | 45.26655949639003 |
| SE_9_3_10 | SE_9 | SE_9_3 | GTAGTATTGGATTGGCTGGT | 807.4137470577826 | 844.4866495156423 | 840.385360101997 | 961.66697378957 | 748.7797259869611 | 622.8769689953858 | 838.3291173436604 | 984.1129836689886 | 884.4333367138692 | 1134.3348061506645 | 820.4756903238524 | 728.9076759931522 |
| SE_9_3_12 | SE_9 | SE_9_3 | GGCAATATGAAATCACCAGT | 597.1610397702527 | 762.5364657756628 | 598.4412787565337 | 661.0647126997314 | 746.6585369615024 | 723.3689417401213 | 724.2560923906077 | 796.9329925388639 | 727.259942538814 | 853.6671323922913 | 746.5272375478829 | 677.8377114331224 |
| SE_9_3_5 | SE_9 | SE_9_3 | ATATTGGCCTCCCTGTGAGG | 140.89098941947887 | 148.90948021045054 | 133.31612645566344 | 149.65047630013606 | 180.30106716399914 | 45.048125713157255 | 162.50025252746178 | 127.62272122508497 | 264.6196127919006 | 177.14868759293492 | 130.29013108147012 | 61.51609367458132 |
| SE_9_3_9 | SE_9 | SE_9_3 | TCTGGGATCTTCTCTCCAAA | 212.42026097090658 | 282.82807315139263 | 260.7070917355196 | 255.0564639550145 | 301.2088416151515 | 220.90907801644423 | 222.76524684228204 | 263.7536238651756 | 288.59521529318016 | 210.68300705497197 | 151.41826044603283 | 349.3649848311128 |
| SE_9_4_1 | SE_9 | SE_9_4 | TAGCTCTCAGAGATTGAGCC | 477.9455871845398 | 371.7740042838094 | 378.22278838903037 | 494.4972260352322 | 470.9039636518566 | 234.77003977433876 | 537.004145769559 | 390.4309915997044 | 323.2266411283618 | 570.8124377994569 | 316.9219404684408 | 307.5804683729066 |
| SE_9_4_11 | SE_9 | SE_9_4 | AGAATGGGTCAGGAGGCAGA | 825.8379533664838 | 1046.3639314116892 | 993.4520238103513 | 918.7237936338787 | 1118.927210929524 | 984.1282848105124 | 874.9185781776584 | 818.6761228216562 | 1083.3420389467071 | 1011.861639419727 | 806.3902707474773 | 733.5504000440641 |
| SE_9_4_12 | SE_9 | SE_9_4 | CACAGGAAAGGCATGCCGCA | 4795.712524470723 | 3987.576013689246 | 4094.2863724828194 | 4469.994661660586 | 3906.169590382405 | 3868.940950672313 | 4939.577212589732 | 4278.669898257293 | 3995.045764935439 | 3954.8626756858926 | 4630.58168573333 | 4986.285630679271 |
| SE_9_4_16 | SE_9 | SE_9_4 | GGAATTAGCTAGCCCAGAGA | 207.0013767624651 | 300.81713787480277 | 324.89633780676496 | 287.5891761941745 | 327.7237044333867 | 211.37966680789174 | 272.2686350294558 | 419.73694980694614 | 217.5563930671666 | 423.5530349444246 | 478.90426559675507 | 456.14763800208414 |
| SE_9_4_19 | SE_9 | SE_9_4 | TTTAGGTTTTACAATCCAAG | 7.586437891818092 | 10.993317330972857 | 16.78795666478725 | 7.807850937398403 | 9.545350614564661 | 6.064170769078861 | 20.4470516425283 | 15.12565584889896 | 50.615160836034676 | 4.374041668961356 | 23.47569929395858 | 13.928172152735392 |
| SE_9_5_0 | SE_9 | SE_9_5 | TTAACTTCTGAACATTGCCA | 70.44549470973944 | 79.95139877071169 | 73.07698783495626 | 72.87327541571842 | 84.84756101835254 | 24.256683076315444 | 82.86436718287786 | 136.13090264009065 | 24.863587779104755 | 158.19450702743572 | 179.58909959878315 | 25.534982280014887 |
| SE_9_5_11 | SE_9 | SE_9_5 | GGTAACAATGAGGAAAAGCT | 193.9960546622055 | 258.8426535201735 | 263.6571991860578 | 217.31851775758886 | 218.48246962225778 | 170.66309164407653 | 194.7850709104012 | 116.27847933841076 | 300.1390239049074 | 150.90443757916677 | 152.59204541073078 | 266.95663292742836 |
| SE_9_5_18 | SE_9 | SE_9_5 | AGGCTTGGAAAGCTTCAACA | 298.0386314642822 | 293.8213904823655 | 340.6967676089177 | 342.24413275596334 | 303.33003064061035 | 294.54543735525897 | 319.61970199110027 | 343.16331707189516 | 347.2022436296414 | 309.09894460660246 | 340.3976397623994 | 321.50864052564197 |
| SE_9_5_19 | SE_9 | SE_9_5 | CTTGATTCTGTGCTAAACAA | 240.59845885480235 | 235.8566263735995 | 263.66967232342324 | 243.34468754891688 | 284.239329411481 | 318.8021204315744 | 215.2321225529295 | 338.43654961911426 | 329.4425380713 | 197.5608820480879 | 333.35492997421187 | 395.79222534023074 |
| SE_9_5_8 | SE_9 | SE_9_5 | GGGCAATTCCCAGAACTGAG | 237.34712832973747 | 304.8147078133383 | 265.64472604869235 | 296.6983356211393 | 266.2092226950811 | 482.53473119670366 | 364.81844772721547 | 294.0049355629735 | 457.3124180799624 | 200.47690982739547 | 501.2061799260157 | 278.5634430547079 |
| SE_9_6_18 | SE_9 | SE_9_6 | CAGCAGATATCAAAAAGGAA | 402.0812082663589 | 542.6701191562056 | 519.4391297457702 | 592.0953627527122 | 617.2660064085147 | 472.13900987828276 | 459.5205816505045 | 624.8786572576383 | 624.2536503110944 | 519.0529447167476 | 496.511040067224 | 503.7355595239301 |
| SE_9_6_4 | SE_9 | SE_9_6 | CTTGGGAATGAAAAGTGGGG | 384.7407787993461 | 519.684092009626 | 531.2894520973847 | 430.73311004647854 | 352.11737822616305 | 590.8234949302547 | 331.4574687315114 | 366.7971543357998 | 694.4044872592827 | 344.09127795829335 | 372.0898338092435 | 713.8188228276889 |
| SE_9_6_7 | SE_9 | SE_9_6 | GGCGAGTTCTTGGGTCCCAG | 1185.651864806999 | 1574.0431632983864 | 1527.7040564956396 | 1318.2254999307636 | 1537.8620434576399 | 1810.5881296249743 | 1047.104276220002 | 1496.494575550441 | 1911.8323031575906 | 1295.4453409573882 | 1318.1605153557744 | 1716.6472178246372 |
| SE_9_6_8 | SE_9 | SE_9_6 | GGACTTTCGTGAACAAATAA | 28.17819788389577 | 43.97326932389143 | 61.22666543834173 | 44.24448864525762 | 25.454268305505764 | 4.331550549342044 | 18.294730416999005 | 34.97807915057884 | 76.3667338929646 | 32.076305723832676 | 32.86597901154201 | 1.1606810127279494 |
| SE_9_6_9 | SE_9 | SE_9_6 | CTAAGGATACTGTTCTCTGA | 293.703524097529 | 339.7934447755247 | 307.1208542793432 | 305.8074950481041 | 371.20807945529236 | 499.8609333940718 | 265.8116713528679 | 187.17999113012462 | 305.4669355718584 | 239.1142779032208 | 217.15021846911688 | 356.32907090748046 |
| SE_9_7_14 | SE_9 | SE_9_7 | GAAGCTGTCTCCAAAGGCAT | 2262.9260454451683 | 2341.576591497219 | 2156.758667993844 | 2308.5212604907942 | 1950.433308909379 | 1651.187069409187 | 2231.9571108738787 | 2334.077768183221 | 2302.5458253060653 | 2203.7879942116965 | 2186.761389232242 | 2407.2524203977673 |
| SE_9_7_17 | SE_9 | SE_9_7 | GCAGTTGCTTTACATAGCAA | 1078.3579574798573 | 1273.2260254235837 | 1227.4958902547382 | 1233.6404481089476 | 1061.6551072421362 | 905.2940648124871 | 1235.4323834538152 | 1081.4843931962757 | 1183.6843753409514 | 1304.922431240138 | 908.5095626761971 | 1720.129260862821 |
| SE_9_7_21 | SE_9 | SE_9_7 | AATGCACTCATTACTGGAGG | 303.4575156727237 | 403.7545637920941 | 353.53461682316674 | 376.0781534846897 | 429.5407776554097 | 242.56683076315446 | 283.0302411571023 | 338.43654961911426 | 229.98818695671898 | 216.5150626135871 | 379.1325435974311 | 194.9944101382955 |
| SE_9_7_4 | SE_9 | SE_9_7 | TTTTCACCCTAAGCTGCAAA | 8.670214733506391 | 6.9957473924273756 | 12.837849214249072 | 5.205233958265602 | 11.666539640023474 | 0.0 | 0.0 | 0.945353490556185 | 0.0 | 0.0 | 1.173784964697929 | 0.0 |
| SE_9_7_6 | SE_9 | SE_9_7 | GAGTAACAGGACACAAGCCA | 1779.5615740521869 | 1658.9915244922677 | 1703.4838380445885 | 1696.9062703945863 | 1801.9500771272621 | 1434.609541942085 | 1968.2977607465402 | 1999.4226325263312 | 1328.425975626454 | 1245.8728687091595 | 2227.8438629966695 | 1519.3314456608857 |
| SE_9_8_10 | SE_9 | SE_9_8 | AACGTTTCAGAAGAGCTCAG | 554.893742944409 | 644.608152588863 | 663.6180516904136 | 683.1869570223603 | 552.5697411320209 | 635.0053105335436 | 637.0870827566713 | 702.3976434832455 | 898.6411011590718 | 803.3656531992357 | 791.1310662064042 | 569.8943772494232 |
| SE_9_8_12 | SE_9 | SE_9_8 | AGAGCTGCAGAGGAGTGAGG | 270.9442104220747 | 90.94471610168455 | 168.86709351050703 | 171.77272062276487 | 83.78696650562313 | 748.4919349263051 | 292.7156866719841 | 161.65544688510764 | 599.3900625319897 | 32.80531251721017 | 98.59793703462604 | 249.54641773650914 |
| SE_9_8_4 | SE_9 | SE_9_8 | AGCCCCATGAAGACAAAGCA | 260.10644200519175 | 325.80194999065014 | 262.6821454607887 | 416.41871666124814 | 361.6627288407277 | 269.4224441690751 | 250.74542277416285 | 342.21796358133895 | 169.60518806460743 | 276.2936320893923 | 257.05890726884644 | 294.81297723289913 |
| SE_9_8_5 | SE_9 | SE_9_8 | CTACGAAAGAGAGCAAGAGA | 475.77803350116324 | 597.6367058110699 | 567.8279460148628 | 474.97759869173615 | 606.6600612812207 | 572.6309826230182 | 472.43450900368026 | 934.954602160067 | 615.3737975328427 | 552.5872641787846 | 899.1192829586137 | 648.8206861149238 |
| SE_9_8_7 | SE_9 | SE_9_8 | TCCTCAACACCCCAGCCTGA | 506.1237850684356 | 697.5759542744595 | 615.2292354213209 | 567.3705014509505 | 674.5381100959027 | 609.8823173473597 | 449.83513613562263 | 635.2775456537563 | 403.1453161326271 | 538.7361322270737 | 521.1605243258805 | 920.4200430932639 |
| SE_90_1_10 | SE_90 | SE_90_1 | CTATGCTCATTTCTACACAG | 85.61837049337561 | 114.93013573289807 | 88.87741763710896 | 81.98243484268323 | 81.66577748016432 | 65.83956834999907 | 68.87427921693744 | 73.73757226338243 | 54.167101947335354 | 124.66018756539864 | 113.85714157569912 | 37.14179240729438 |
| SE_90_1_14 | SE_90 | SE_90_1 | TGTCAAGGAGAAAGTACAAA | 505.0400082267473 | 590.6409584186326 | 644.8550413003572 | 605.1084476483762 | 521.8125002628681 | 390.70585955065235 | 499.33852432279645 | 410.2834149013843 | 587.8462539202624 | 605.8047711511477 | 589.2400522783604 | 641.856600038556 |
| SE_90_1_17 | SE_90 | SE_90_1 | GGAAGTGGCCCCGATGAAGC | 512.6264461185654 | 631.6160502886224 | 665.5931054156827 | 597.3005967109779 | 733.9314028087495 | 512.8555850420979 | 534.8518245440298 | 640.0043131065372 | 536.3431078064026 | 980.5143407921706 | 653.7982253367464 | 674.3556683949387 |
| SE_90_1_22 | SE_90 | SE_90_1 | CGAGGCTGAGAGTGGAGAGG | 164132.58378948443 | 150924.25545947248 | 151179.49987385972 | 168473.90360171403 | 162987.9223382043 | 175705.88279362052 | 178773.95331368878 | 183618.8445311995 | 156316.48838195376 | 225684.51196561978 | 194517.2967798114 | 215461.85911674018 |
| SE_90_1_6 | SE_90 | SE_90_1 | ATGACAGCCATGCCAAGAGA | 5289.914764280587 | 5355.744325153049 | 5257.593016666312 | 5414.744625085792 | 5141.7621977121635 | 5479.411444917685 | 5860.77069711627 | 6117.382437389073 | 5287.064344171061 | 5649.8038224084185 | 6326.7009597218375 | 4007.8315369496095 |
| SE_90_2_1 | SE_90 | SE_90_2 | GGGTTTCTGCAACACAGCGA | 46.60240419259685 | 52.9678016855965 | 57.27655803280356 | 26.02616979132801 | 30.757240896152194 | 20.791442636841808 | 21.52321225529295 | 85.08181415005664 | 58.607028336461205 | 121.01515284126418 | 82.16494752885504 | 5.803405063639747 |
| SE_90_2_10 | SE_90 | SE_90_2 | GCTTGCCCTGCAGGACTACG | 405.3325387914238 | 92.94350107095235 | 167.87956664787248 | 269.37085734024487 | 71.05983235287025 | 1112.3421810710367 | 436.92120878244685 | 165.43686084733238 | 600.2780478098148 | 48.114458358574915 | 72.72901221113639 | 183.38760001101602 |
| SE_90_2_13 | SE_90 | SE_90_2 | ATGTTTCTAACATGACCACA | 96.45613891025 86 | 68.95808143973883 | 61.22666548334173 | 79.37981786355043 | 68.93864332741144 | 25.98930329605 2263 | 118.37766740411122 | 41.59555358447214 | 63.04695472558706 | 24.78623612411435 | 83.33873249355297 | 2.321362025455899 |
| SE_90_2_16 | SE_90 | SE_90_2 | GAGTCTCCAGGACTGACAGC | 290.4521935724641 | 302.81592284407054 | 359.459777998974 | 311.0127290063697 | 265.1486281823517 | 335.2620125190742 | 275.49711686774975 | 170.1636283001133 | 323.2266411283618 | 222.34711817220227 | 165.503680022408 | 420.1665266075177 |
| SE_90_2_2 | SE_90 | SE_90_2 | GGTAGTCGATGGAGGAGTTA | 2063.511106574521 | 2827.281339029292 | 2929.992201436692 | 2213.5257407524473 | 2392.7012207175417 | 2060.9517513769442 | 1938.16526358913 | 1441.6640730981821 | 2830.897065706641 | 1939.1584732395345 | 1409.7157426022127 | 1214.0723393134351 |
| SE_90_3_0 | SE_90 | SE_90_3 | TGAATTCATGCAGTACTAAG | 140.89098941947887 | 157.9040125271556 | 162.94193233469977 | 144.44542234187045 | 144.24085371311993 | 111.75400417302473 | 143.12936516976981 | 178.67180971511897 | 133.19779167377547 | 255.15243068941243 | 150.24447548133492 | 77.76562785277261 |
| SE_90_3_1 | SE_90 | SE_90_3 | TGTGCAGAATGGCATGAGAA | 3113.690866170483 | 3245.027397606261 | 3223.2876796391515 | 3313.1314144360554 | 3264.509910181114 | 3506.8233247473186 | 3403.89601817458 | 3570.6001338307105 | 2883.2881970983262 | 3486.1112101622007 | 3779.5875863273313 | 3328.833144503759 |
| SE_90_3_2 | SE_90 | SE_90_3 | TGTTGCTTCTGGGCACAGCA | 873.524134400769 | 933.4325860484579 | 895.6868644095315 | 878.3832304573203 | 873.9298784890311 | 763.2192067940681 | 888.9086661435988 | 996.402579046219 | 978.5597761633371 | 1464.5749521572272 | 795.8262060651958 | 846.1364582786751 |
| SE_90_3_5 | SE_90 | SE_90_3 | TTATTATTCTCATAGTACAG | 6.5026610501297935 | 2.9981774539016888 | 0.0 | 2.602616979132801 | 0.0 | 0.0 | 5.380803063823238 | 5.67212094333711 | 0.0 | 0.0 | 0.0 | 0.0 |
| SE_90_3_9 | SE_90 | SE_90_3 | CATAGGGCAACAATGCCCAA | 278.5306483138928 | 347.7885846525959 | 340.6967676089177 | 321.4231969229009 | 370.147484942563 | 220.90907801644423 | 262.5831895145574 | 217.43130282792256 | 355.19411113006794 | 112.99607644816837 | 99.77172199932397 | 847.2971392914031 |
| SE_90_4_18 | SE_90 | SE_90_4 | GGGCTGGCAAAGCCTGCAGG | 13.005322100259587 | 3.9975699385355847 | 10.862795488979984 | 3.9039254686992013 | 0.0 | 38.98395494407839 | 12.913927353175769 | 14.180302358342775 | 61.27098416993672 | 0.0 | 17.606774470468935 | 5.803405063639747 |
| SE_90_4_22 | SE_90 | SE_90_4 | GTCAGGTCGTCAGGTCTGGA | 4364.3693414787795 | 4505.2613207296035 | 4271.0536808944025 | 4304.728483485653 | 4004.80488006624 | 4248.384778794676 | 4403.649227432938 | 4278.669898257293 | 3955.086427433306 | 4607.323891305961 | 3726.7672629159247 | 4621.831792682695 |
| SE_90_4_27 | SE_90 | SE_90_4 | GCGCCTTTGCCATTTCCTGC | 301.2899619893471 | 445.7290481467177 | 400.9359062296249 | 348.7506752037953 | 466.6615856003896 | 285.8823362565749 | 329.3051475059821 | 537.9061361264693 | 467.0802561360393 | 394.3927571513489 | 285.22974642159676 | 465.4330861039077 |
| SE_90_4_28 | SE_90 | SE_90_4 | TGAAGTGGTCTTCCAAGCAG | 1669.0163361999803 | 1629.0097499532508 | 1640.2821188359776 | 1755.4651524250742 | 1736.193217338039 | 1631.2619368822136 | 1743.380192678729 | 2287.7554471459675 | 1688.0600131456476 | 1426.6665910262288 | 2286.5331112315657 | 1305.766139318943 |
| SE_90_4_8 | SE_90 | SE_90_4 | GCATTCCACAGTTCAGAACA | 1732.95916985959 | 1815.8961445797893 | 1909.8769523352082 | 1651.3604732592762 | 1968.463415625779 | 1940.5346461052347 | 1727.237783487259 | 2146.897777053096 | 1773.3065998186864 | 2085.6888691497397 | 2590.5434170883295 | 1838.518724161072 |
| SE_90_5_0 | SE_90 | SE_90_5 | ATAGTTGGAGAGTTACAGCC | 134.38832836934907 | 194.88153450360974 | 176.76730841158337 | 150.95178478970246 | 142.1196647057405 | 91.82887164605133 | 181.87114355722542 | 172.99968877178185 | 110.11017445032105 | 147.2594028550323 | 177.2415296693873 | 306.41978736017865 |
| SE_90_5_14 | SE_90 | SE_90_5 | TTCCATATAGTTCTTAGTGA | 1.083776841688299 | 7.9951398770711695 | 1.975053725269088 | 2.602616979132801 | 0.0 | 4.331550549342044 | 0.0 | 0.0 | 0.0 | 0.0 | 2.347569929395858 | 0.0 |
| SE_90_5_19 | SE_90 | SE_90_5 | TCTGTGTCTTTACACAACAA | 209.16893044584168 | 254.84508358164354 | 250.83182310917 42 | 214.71590778457607 | 201.5129574185873 | 200.98394548947 08 | 189.40426784657797 | 183.3985771678999 | 207.78855501108973 | 197.5608820480879 | 106.81443178751154 | 181.06623798556012 |
| SE_90_5_2 | SE_90 | SE_90_5 | GCATTTGTAAAGCAAATAGT | 340.30592829012585 | 398.75760136892455 | 392.048164465914 | 347.4493667142289 | 359.5415398152689 | 229.5721791151283 | 385.2654993697438 | 353.5622054680132 | 388.0495664095992 | 280.66767375835366 | 416.6936624677648 | 305.2591063474507 |
| SE_90_5_4 | SE_90 | SE_90_5 | ATACAAACAGAGTGAACAAG | 231.92824412129596 | 246.84994370457235 | 278.48257526219414 | 261.5630064028465 | 269.3910062332693 | 313.60425977236395 | 251.8215833869275 | 516.1630058436771 | 310.7948472380894 | 189.5418056549921 | 543.4624386551411 | 306.41978736017865 |
| SE_91_1_11 | SE_91 | SE_91_1 | ATGCCCCCGAGTGTACCCCA | 457.3538271924621 | 672.5911421586121 | 662.630524827779 | 586.8901287944466 | 667.1139485067969 | 503.32617383354545 | 525.166379029148 | 613.5344153709641 | 554.102813362906 | 688.9115628614136 | 331.007360044816 | 669.7129443440268 |
| SE_91_1_2 | SE_91 | SE_91_1 | GGCTCAGGGAGGGGAAAACA | 276.36309463051623 | 362.77947192210434 | 351.5595630978977 | 271.9734743193277 | 243.93673792776355 | 286.74864636644327 | 375.58005385486194 | 337.49119612855804 | 277.051406681453 | 441.778208565097 | 267.6229719511278 | 174.10215190919243 |
| SE_91_1_24 | SE_91 | SE_91_1 | TTTCCCCCTTGTAAATCTCA | 1.083776841688299 | 0.0 | 2.962580587903632 | 16.917010364363207 | 0.0 | 5.197860659210452 | 9.685445514881827 | 11.34424188667422 | 7.103882226601358 | 4.374041668961356 | 17.606774470468935 | 0.0 |
| SE_91_1_30 | SE_91 | SE_91_1 | TCAGCTCGGCACCAGACGCC | 6894.9882668209575 | 5676.54931272053 | 5897.510423653497 | 6104.438124555984 | 5525.6974113202095 | 5773.090572163076 | 7019.795677063796 | 5301.5423750390855 | 5740.824821139723 | 6894.947684172751 | 4458.035295922735 | 5972.864491498028 |
| SE_91_1_5 | SE_91 | SE_91_1 | GGGCATGTCACCTTGCGCCA | 2757.1282852550326 | 2326.5857042277103 | 2210.0851185761094 | 2642.9575423093593 | 2294.065931033707 | 2219.4865014828633 | 2877.6534785632 | 2511.8042244077837 | 2158.692210392988 | 2587.9746541354 69 | 2469.643565724443 | 2235.4716305140305 |
| SE_91_2_21 | SE_91 | SE_91_2 | TGAGCTCCCCTGCTGATGGG | 1123.876584830766 | 1081.3426683738758 | 1139.6059994802638 | 1154.2606302453971 | 1151.8056408241357 | 1005.7860375572226 | 1018.0479396753565 | 1094.7193420640622 | 1009.639260887218 | 1156.2050144954517 | 883.8600784175405 | 930.8661722078155 |
| SE_91_2_23 | SE_91 | SE_91_2 | TGGGACAACCAGAAGCTCAG | 118.13167574402458 | 174.89368481093183 | 136.27870704356707 | 117.11776406097604 | 142.1196647057405 | 184.52405340197106 | 103.31141882540615 | 113.44241886667422 | 178.48504084285912 | 94.04189588266915 | 173.7201747752935 | 250.70709874923708 |
| SE_91_2_47 | SE_91 | SE_91_2 | GGCAAGTAACGGATGCAGGG | 1148.8034521895968 | 1062.3542111658317 | 1110.967720463862 | 1070.9768869131476 | 1223.9260676897354 | 1043.037372281564 | 1086.922218892294 | 710.9058248982511 | 989.2155994972392 | 946.2510143853067 | 447.21207154991095 | 1198.9834861479717 |
| SE_91_2_6 | SE_91 | SE_91_2 | GGACTGCAGGAGGAGGCTGG | 331.63571355661946 | 103.93681840192521 | 216.26838291696515 | 279.7813252567761 | 98.63528968383483 | 590.8234949302547 | 496.11004248450246 | 183.3985771678999 | 380.05769890917264 | 30.61829168272949 | 42.25625872912545 | 100.9792481073316 |
| SE_91_2_8 | SE_91 | SE_91_2 | TGGTAAAGCAGGCTGCCCGC | 37.93218945909046 | 31.980559508284678 | 44.43870881855448 | 28.62878677046081 | 61.51448173830559 | 30.320853845349207 | 36.58946083399801 | 16.071009339455145 | 33.74344055735645 | 118.09912506195661 | 49.29896851731302 | 48.74860253457388 |
| SE_91_3_1 | SE_91 | SE_91_3 | TACTGACCCCATATACCCCT | 3209.063228239053 | 3198.0559508284678 | 3208.4747766996334 | 3495.3146029753516 | 3358.902821814031 | 3182.823343656534 | 3145.617471111065 | 2923.97834629028 | 2890.3920793209277 | 3469.344050431182 | 3480.2724203293596 | 2792.5985166234464 |
| SE_91_3_10 | SE_91 | SE_91_3 | ACAATTATGGACATGAGCCA | 654.6012123797325 | 600.6348832649716 | 573.7531071906701 | 631.1346174397042 | 712.7195125561416 | 781.4117191013047 | 742.5508228076068 | 683.4905736721217 | 699.7323989262338 | 944.0639953550826 | 571.6332778078914 | 1029.524058289691 |
| SE_91_3_18 | SE_91 | SE_91_3 | TGGAGAAAGGGTCCATACAG | 306.7088461977886 | 379.76914416088056 | 397.97332564172126 | 351.3532921829281 | 363.7839178661865 | 184.52405340197106 | 216.30828316569415 | 246.7372610351643 | 383.60964002047336 | 291.60277793075704 | 257.05890726884644 | 362.13247597112024 |
| SE_91_3_38 | SE_91 | SE_91_3 | GTACAAGGAGGGTCAAGGGA | 360.89768828220355 | 441.7314782081821 | 506.6012805315 2107 | 400.80301478645134 | 443.328506320892 | 437.4866054853464 | 355.1330022123337 | 398.9391730147101 | 428.896889189557 | 235.46924317908633 | 615.0633215017148 | 456.14763800208414 |
| SE_91_3_6 | SE_91 | SE_91_3 | TCCATAATTGTTGAACCCAG | 628.5905681792134 | 699.5747392437273 | 680.4060083552608 | 605.1084476483762 | 642.7202747140204 | 781.4117191013047 | 526.2425396419126 | 598.4087595220651 | 769.8832358744222 | 648.0871739511076 | 426.08394218534823 | 554.8055240839599 |
| SE_91_4_10 | SE_91 | SE_91_4 | CAGCATAGCTTTCTGTTGGG | 604.7474776620708 | 664.596002281541 | 669.5432128662209 | 629.8333089501378 | 907.8689028963722 | 777.9464786618311 | 562.8320004759106 | 983.1676301784324 | 698.8444136484086 | 575.9154864132452 | 889.7290032410302 | 636.0531949749163 |
| SE_91_4_12 | SE_91 | SE_91_4 | ACAGGGCTGGAAGGGTGGAA | 65420.021494830784 | 69506.747913802284 | 68674.59308133146 | 65552.11385341786 | 64970.95925529073 | 64182.3171098208 | 63939.006664797876 | 61678.643137847735 | 67894.46635723466 | 57076.869738267335 | 58848.88299009537 | 64350.476707662 97 |
| SE_91_4_15 | SE_91 | SE_91_4 | CCACCAGGGCTGCCTCCACG | 347.89236618194394 | 429.73876839257537 | 428.5866583833921 | 352.6546006724945 | 353.17797273889244 | 343.92511361775826 | 295.9441685102781 | 402.72058697693484 | 421.79300696695566 | 290.87377098593015 | 403.7820278560876 | 340.0795367292892 |
| SE_91_4_27 | SE_91 | SE_91_4 | AGGAAGGCCCCATCAGCAGA | 2451.503215898932 | 3022.1628735329023 | 2990.2313400573994 | 2642.9575423093593 | 3140.4203521917734 | 2432.598788510492 | 2272.8512141589354 | 3328.5896402483 27 | 2594.6929818051462 | 2805.94773063871 | 3210.301878448836 | 3144.284863480015 |

| | | | | | | | | | | | | | | | |
|---|---|---|---|---|---|---|---|---|---|---|---|---|---|---|---|
| SE_91_4_3 | SE_91 | SE_91_4 | ACAGCCTTGTGACCAATGAT | 3029.1562725187955 | 2685.367606211279 | 2660.3973679374617 | 3117.9351410010954 | 2957.9980960023154 | 3314.5024803565316 | 3015.4020369665423 | 2476.8261452572046 | 2634.652319307279 | 1885.2119593223445 | 2383.957263301494 | 2707.868802694306 |
| SE_91_5_12 | SE_91 | SE_91_5 | GGGAGGAAACGCTTACCAAT | 1915.0336792632243 | 2067.743050707531 | 2117.2575934884626 | 1997.5085314844246 | 2198.61242488806 | 1990.7806324776031 | 2086.6754281506514 | 2510.8588709172273 | 1765.3147323164376 | 2592.3486958044305 | 2838.2120446395925 | 2105.4753570885005 |
| SE_91_5_13 | SE_91 | SE_91_5 | GCACCGTTGGTCAGATGGAA | 1745.9644919598495 | 1644.0006372227592 | 1681.7582470666284 | 1784.093939195535 | 1633.3155496032864 | 1295.1336142532712 | 1732.6185865510824 | 1713.9258783783634 | 1409.2326359085444 | 1524.3535216330326 | 1494.2282600604638 | 1335.94384564987 |
| SE_91_5_23 | SE_91 | SE_91_5 | GGAGACAGAATCTTTCATGG | 312.12773040623006 | 604.6324532035072 | 568.8154728774973 | 467.1697477543378 | 534.539634415621 | 485.1336615263089 | 404.63639039950743 | 428.2451312219518 | 601.1660330389566 | 694.0146114752018 | 496.511040067224 | 659.2668152294752 |
| SE_91_5_26 | SE_91 | SE_91_5 | TGGGCATGAGAAGGAAGGGG | 200.4987157123353 | 254.84508358164354 | 246.881715658636 | 124.92561499837444 | 168.63452752397566 | 113.48662439276154 | 140.9770402721688 | 155.03797245121433 | 198.02071695501286 | 43.01140974478667 | 23.47569929395858 | 55.71268861094157 |
| SE_91_5_8 | SE_91 | SE_91_5 | TCAGGAGTCCTTAAGACAGA | 1017.6664543453127 | 1185.279486775801 | 1100.1049249748821 | 1005.9114624348275 | 972.565168712866 | 829.0587751440671 | 958.8591059733009 | 1076.7576257434948 | 998.983437553316 | 995.0944796887085 | 914.3784874996867 | 935.5088962587272 |
| SE_91_6_0 | SE_91 | SE_91_6 | CGATCTTGGTGACCAGCTCC | 6858.139854203556 | 6726.910814070755 | 6743.380592933087 | 7054.014104163284 | 6900.160025101875 | 7056.385137897793 | 6721.4633178544755 | 6494.724322201391 | 7203.317621834526 | 7461.751020584735 | 6391.870337092818 | |
| SE_91_6_15 | SE_91 | SE_91_6 | GCTGTGGCAGGCAGATTGGG | 2698.604335803864 | 2547.4514433318013 | 2607.0709173551963 | 2934.450643972233 | 2564.5175317797057 | 3013.0265621223257 | 2821.6931266689057 | 2605.394219972846 | 2433.0796612409654 | 2739.6080986594625 | 2672.7083646171845 | 3211.604362218236 |
| SE_91_6_17 | SE_91 | SE_91_6 | CTTTGCTGTGGCAGGCAGAT | 214.5878146542832 | 243.85176625067066 | 223.18107095540694 | 265.4669318715457 | 270.45160074599875 | 348.2566641671003 | 209.85131948910626 | 320.47483329854674 | 402.2573308548019 | 232.55321539977876 | 196.02208910455414 | 277.40276204197994 |
| SE_91_6_18 | SE_91 | SE_91_6 | TGCCTGCCACAGCAAAGTGC | 2076.516428674781 | 1919.8329629817147 | 1854.5754480276737 | 1963.6745107556983 | 1966.3422266003201 | 1448.4705036999794 | 2098.5131948910625 | 1818.8601158300999 | 1918.0482001023668 | 3177.741272500425 | 1798.2385659172273 | 1924.4091191029402 |
| SE_91_6_25 | SE_91 | SE_91_6 | CGCATCCTCCAGGGGGCCCA | 1399.155902619594 | 1723.9520359934709 | 1735.0846976488938 | 1482.19036961613 | 1667.2545740106275 | 1483.1229080947157 | 1235.4323834538152 | 1717.707292340588 | 1902.9524503793389 | 2003.311084384301 | 2020.083924245136 | 1882.624602644734 |
| SE_92_1_10 | SE_92 | SE_92_1 | TTCAACATCGGACATTCAGG | 9.75399157519469 | 32.97995199291857 | 20.738064115325425 | 20.820935833062407 | 15.9089176909411 | 6.93048087894727 | 12.913927353175769 | 42.54090707502832 | 5.327911666951019 | 9.477090282749604 | 95.07658214053225 | 1.1606810127279494 |
| SE_92_1_11 | SE_92 | SE_92_1 | AAGCACATACATGATAGAGG | 783.5706565406401 | 754.5413258985916 | 677.4434277672972 | 705.309201344989 | 825.1425309034785 | 557.0374006453868 | 801.7396565096624 | 1141.0416631013154 | 868.449601713016 | 950.625056054268 | 1015.3239944637086 | 500.2535164857462 |
| SE_92_1_22 | SE_92 | SE_92_1 | TCTGCAGGAGGAGGAAGTGG | 311.04395356454177 | 110.93256579436247 | 209.35569487852334 | 242.04337905935049 | 32.87842989461161 | 358.6523854855212 | 377.73237508039125 | 121.00524679119168 | 546.9989311403045 | 16.76715973101853 | 30.518409082146157 | 126.51423038734649 |
| SE_92_1_5 | SE_92 | SE_92_1 | TATGCATGTGAGAAAGAGGG | 1398.0721257779055 | 1605.0243303220373 | 1569.1801847262905 | 1371.5791480029861 | 1395.7423787518992 | 1128.8020731585366 | 1179.4720315900536 | 1190.200044610237 | 1614.3572350861587 | 867.5182643440022 | 869.7746588411654 | 937.8302582841832 |
| SE_92_1_9 | SE_92 | SE_92_1 | TCAAGCACATACATGATAGA | 1592.068180440111 | 1533.0680714283967 | 1626.456742759094 | 1525.1335497718212 | 1604.6794977595923 | 1986.4490819282612 | 1520.6149458364469 | 1636.4068921527562 | 1667.6363517556688 | 1718.269368956986 | 1356.895419190806 | 1364.9608709680685 |
| SE_92_2_17 | SE_92 | SE_92_2 | GTAGTAGCGTTAGTAATTGT | 1055.598643804403 | 1052.3602863194926 | 1081.3419145848256 | 1098.304365194042 | 987.4134913510777 | 1329.7860186480075 | 992.2200849690049 | 938.7360161222917 | 914.6248361599248 | 929.4838546542882 | 951.9396063700204 | 1871.0177925174544 |
| SE_92_2_7 | SE_92 | SE_92_2 | TGTAGCTGCAAGTTCTACCT | 1608.3248330654355 | 1372.1658814023394 | 1437.839111995896 | 1400.2079347734468 | 1472.1051836684167 | 1354.0427017243228 | 1489.406288066272 | 1869.909204320134 | 1864.7690834328566 | 1476.2390632744575 | 1532.9631638954954 | 1695.754959595534 |
| SE_92_2_8 | SE_92 | SE_92_2 | TTGACCTTTGACATTTTCAA | 58.5239494511 6814 | 84.94836119388117 | 76.03956842285989 | 83.28374333224963 | 50.90853661101153 | 231.30479933486512 | 53.80803063823 2374 | 37.8141396224274 | 56.831057780810866 | 39.366375020652 21 | 34.03976397623994 | 175.26283292192036 |
| SE_92_3_11 | SE_92 | SE_92_3 | AACTTTCTCGGCTCCTTGCA | 43.351073667531956 | 101.9380334326574 | 88.87741763710896 | 79.37981786355043 | 67.87804881468203 | 52.844916701972934 | 65.6457973786435 | 79.40969320671954 | 91.46248361599248 | 74.35870837234305 | 5.868924823489645 | 153.20989368008932 |
| SE_92_3_13 | SE_92 | SE_92_3 | TATTGTGGACTTCTTCACCA | 522.3804376937601 | 644.608152588863 | 638.92988012455 | 631.1346174397042 | 664.9927594811338 | 432.28874482433594 | 509.02396983767824 | 710.9058248982511 | 713.9401633714365 | 628.4039864407815 | 685.4904193835905 | 610.5182126949014 |
| SE_92_3_16 | SE_92 | SE_92_3 | GGTATTTCTGAGAACAGGGT | 26.010644200519174 | 66.95929647047105 | 54.31397744489992 | 57.25757354092162 | 92.27172260745839 | 25.989303296052263 | 29.056336544645482 | 88.86322811228139 | 66.59889583688773 | 25.515243068941242 | 30.518409082146157 | 38.30247342002233 |
| SE_92_3_3 | SE_92 | SE_92_3 | CAGCTGCAGAACAAAAGCCA | 312.12773040623006 | 113.93074324826416 | 202.44300684008152 | 255.0564639550145 | 86.96875004381135 | 783.1443393210415 | 400.3317479488424 | 242.95584707293955 | 835.5941464334848 | 32.076305572383276 | 116.20471150509498 | 178.74487596010422 |
| SE_93_1_12 | SE_93 | SE_93_1 | AATTTGGCATCAAGGCCACC | 34.680858934025565 | 48.970231747060915 | 42.463655093285396 | 42.943180155691216 | 24.393673792776355 | 3.465240439473635 | 29.056336544645482 | 73.73757226338243 | 157.17339417505505 | 40.8243889103059 9 | 21.128129364562724 | 1.1606810127279494 |
| SE_93_1_19 | SE_93 | SE_93_1 | CCCTGAGAAGCAGTTCCCAA | 436.7620672003845 | 447.7278331159855 | 423.6490240702194 | 424.22656759864657 | 482.5705032918801 | 412.3636122973626 | 350.82835976127507 | 572.8842152770482 | 558.5427397520318 | 356.4843960203505 | 657.3195802308403 | 422.4878886329736 |
| SE_93_1_27 | SE_93 | SE_93_1 | CTGAGGAGTGATTTGCACAA | 2433.079009590231 | 2463.502474622554 | 2399.690276201942 | 2684.599147395484 | 2764.9698946855633 | 2483.711084992728 | 2675.3352833329136 | 2732.071587707375 | 2430.41570540749 | 2513.615945763126 | 2953.2429711799896 | 3548.2018559093412 |
| SE_93_1_29 | SE_93 | SE_93_1 | ATATTACTAACCTCACTAAG | 24.926867358830876 | 23.98541963121351 | 16.78795666478725 | 20.820935833062407 | 31.8178353818822 | 20.791442636841808 | 2.152321225529295 | 18.9070698111237 | 64.82292528123739 | 1.4580138896537853 | 44.6038286585213 | 16.249534178191293 |
| SE_93_1_43 | SE_93 | SE_93_1 | GCAAACCGTGACTGTGGTGG | 4541.024966673973 | 4255.41319957113 | 4163.413252867238 | 4281.304930673457 | 4106.621955288263 | 4382.662845824279 | 4624.26251304969 | 4781.579552331884 | 4200.170364113053 | 4056.1946410168307 | 4785.521301073457 | 4335.143582538891 |
| SE_93_2_19 | SE_93 | SE_93_2 | AGTCTCCTCTCTGTGCCAGG | 26.010644200519174 | 51.968409200962604 | 37.526020780112674 | 59.86019052005442 | 37.120807945529236 | 76.23528966841997 | 50.57954879993843 | 46.32232103725307 | 111.88614500597139 | 19.683187510326103 | 15.259204541073078 | 20.89225822910309 |
| SE_93_2_23 | SE_93 | SE_93_2 | CTGTGCCAGGAGGCAGTCAG | 160.39897256986825 | 180.8900397187352 | 181.7049427247561 | 148.34916781056964 | 152.72560983303458 | 51.11229648223612 | 124.83463108069911 | 135.18554914953447 | 141.189659174202 | 194.64485426878034 | 97.42415206992811 | 64.99813671276517 |
| SE_93_2_25 | SE_93 | SE_93_2 | GCCACTCCCGGGAACAACAG | 295.8710777809056 | 425.74119845403976 | 406.86106740543215 | 395.5977808281857 | 460.2980185245625 | 521.518686140782 | 338.99059302086397 | 397.9938195241539 | 523.023328639025 | 429.38509050303975 | 473.0353407732654 | 537.3953088930406 |
| SE_93_2_43 | SE_93 | SE_93_2 | AGCCTCCCTGACTTCCCCCA | 213.50403781259487 | 283.8274656360265 | 247.86924252127056 | 217.31851775758886 | 242.87614341503414 | 278.9518553776276 | 212.00364071463557 | 253.35473546905757 | 344.5382877961659 | 384.1866599237724 | 230.0618530807941 | 221.69007343103834 |
| SE_93_3_21 | SE_93 | SE_93_3 | CATGCAGCGCTCACCAGACA | 17701.327155294985 | 15903.33260797919 | 15848.818618421798 | 17835.734157997085 | 16879.361670088507 | 16943.293128068507 | 18321.634432318122 | 18064.75985103814 | 15935.783795850497 | 20344.396809284095 | 18314.566804181788 | 18703.213839098178 |
| SE_93_3_35 | SE_93 | SE_93_3 | GCTATCTCCCTGGACTTCCG | 231.92824412129596 | 288.824428059196 | 306.13332741670865 | 189.99103947669445 | 249.2397104914106 | 256.42779252104896 | 133.4439159828163 | 117.22383282896693 | 132.3098063959503 | 153.82046535847434 | 116.20471150509498 | 140.44240254008187 |
| SE_93_3_36 | SE_93 | SE_93_3 | GAGACTGCAGAGCAGCACGG | 183.15828624532253 | 38.97630690072195 | 89.8649444997435 | 191.29234796626088 | 24.393673792776355 | 245.16576109275968 | 230.29837113163455 | 138.96696311175918 | 277.93939195927817 | 7.290069448268927 | 49.29896851731302 | 171.7807898837365 |
| SE_93_3_37 | SE_93 | SE_93_3 | GGAGACTGCAGAGCAGCACG | 199.414938870647 | 53.96719417023039 | 82.9522564613017 | 258.9603894237137 | 67.87804881468203 | 761.4865865743312 | 274.42095625498513 | 94.5353490556185 | 260.17968640277473 | 23.328222234460565 | 62.210603128990236 | 110.26469620915519 |
| SE_93_3_44 | SE_93 | SE_93_3 | GAGAGGAGACTGCCCCCAGG | 5164.196650644744 | 7243.5967286264795 | 7043.041584309568 | 6159.093081117773 | 7214.163875585425 | 6818.7268747742455 | 5488.419125099702 | 8323.837484347208 | 7436.876701785797 | 7371.718226089539 | 8704.789298199841 | 7796.294362493636 |
| SE_93_4_1 | SE_93 | SE_93_4 | GAGAGTGGACCCAGAGAGAG | 891.94834070947 | 1030.373651657547 | 1008.2649267498695 | 985.0905266017651 | 1058.473323703948 | 1138.3314843670892 | 955.630624135007 | 1324.440240269215 | 1202.3320661752798 | 1763.4677995362533 | 1308.7702356381908 | 1176.9305469061408 |
| SE_93_4_12 | SE_93 | SE_93_4 | GCCGGGGCTGGGACATCACA | 2591.310428476723 | 2549.4502283010693 | 2625.8339277452524 | 2862.878677046081 | 2748.0003824818928 | 3113.518534867061 | 2861.5110693411975 | 3131.956114212641 | 2579.5972320821184 | 3392.7983212243585 | 3279.555191366014 | 3880.156625549535 |
| SE_93_4_23 | SE_93 | SE_93_4 | GCAATCTGTTAGGCAGCCAG | 1418.6638857699834 | 1441.1239628420783 | 1536.591982593505 | 1534.242709198786 | 1749.9809460035212 | 1151.3261360151153 | 1430.2174543642166 | 1764.0296133778413 | 1677.4041898117457 | 1606.0022994536446 | 1921.4859872105098 | 1592.4543494627467 |
| SE_93_4_31 | SE_93 | SE_93_4 | GAGTCCCCAGAGGCTGGGAG | 33.59708209233727 | 60.962941562667666 | 65.1767729338799 | 20.820935833062407 | 44.54496953463509 | 55.44384703157816 | 16.142409191469714 | 13.23494886778659 | 31.96747000170611 | 2.187020834480678 | 2.347569929395858 | 41.78451645820618 |
| SE_93_4_7 | SE_93 | SE_93_4 | TAATGAATGGAACATGGCAC | 2568.5511148012683 | 2104.7205726389852 | 2141.945765054326 | 2311.1238774699273 | 2171.0369675570955 | 2384.951732467729 | 2608.6133253415055 | 2510.8588709172273 | 2390.456367905357 | 2144.7384316807183 | 2987.2827351562296 | 3063.0371925890586 |
| SE_93_5_11 | SE_93 | SE_93_5 | GTACATTTCAGCAGCCCTGC | 894.1158943928466 | 907.4483760475778 | 932.2255383207295 | 903.108091759665 | 835.7484760307725 | 1030.042720633538 | 898.5941156584806 | 868.779857821134 | 801.8570581861284 | 1258.9949931160436 | 890.9027882057281 | 802.0305797950131 |
| SE_93_5_24 | SE_93 | SE_93_5 | TGTACTCCCAGCCACCCAGG | 403.1649851080472 | 544.6689041254734 | 498.70106563044476 | 545.2482571283218 | 414.69245447719805 | 456.5454279006514 | 366.9707689527448 | 488.7477546175476 | 534.5671372507522 | 713.6977989855279 | 264.10161705703405 | 326.1513645765538 |
| SE_93_5_26 | SE_93 | SE_93_5 | GCAAACACCCAACAGTGCAC | 108.37768416882989 | 112.93135076363026 | 106.65290116453076 | 92.39290275921444 | 83.78696650562313 | 40.71657516381521 | 89.32133085946575 | 150.31120499843342 | 84.35860139339113 | 173.50365286880046 | 96.25036710523018 | 82.4083519036844 |
| SE_93_5_3 | SE_93 | SE_93_5 | ACTGTCTCCTTGCAAAGCCA | 554.893742944409 | 527.6792318866972 | 533.2645058226537 | 543.9469486387554 | 523.933689288327 | 1075.0908463466953 | 479.96763329303275 | 506.70947093811515 | 710.3882222601359 | 472.39650024782645 | 482.4256204908488 | 504.896240536658 |
| SE_93_5_32 | SE_93 | SE_93_5 | TGAAGGCCCAAGAACTTGCA | 2915.359704141524 | 2945.209652216092 | 2779.8881183162416 | 2907.1231656913387 | 3066.1787363007147 | 3102.2565034387717 | 3048.7630159622463 | 3782.3593157152964 | 2855.760653485746 | 2809.592765362844 | 3734.98375766881 | 3220.8898103200595 |
| SE_93_6_23 | SE_93 | SE_93_6 | TAGGGGCCCAGTCCAGAATG | 6606.7036269318705 | 6879.817864219742 | 7089.455346853391 | 7053.09201344989 | 7188.7096077209756 | 7477.9888683841045 | 6748.603202647104 | 7695.177413127346 | 7305.454880667672 | 7069.180349366378 | 7769.282681335592 | 8563.50451190681 |
| SE_93_6_31 | SE_93 | SE_93_6 | CTGGCAAGACAGGAGCGGCA | 1157.4736669231033 | 1100.3311255819197 | 1224.5333096668346 | 1232.3391396193813 | 1163.4721804641592 | 971.1336331624861 | 1344.1246053430448 | 1253.5387284775013 | 1088.669950613658 | 1127.044736702376 | 1135.0500608628975 | 820.6014759986602 |
| SE_93_6_32 | SE_93 | SE_93_6 | GAGCGCGCCCCTCTTCTCGG | 2015.8249255402359 | 2166.682906686287 | 2419.440813454633 | 2203.115272859316 | 2531.639101885924 | 2496.705736640754 | 2137.25497695059 | 2314.225344881541 | 2580.4852173599434 | 2431.967167942 5137 | 2628.1045359586633 | 2355.0217748250093 |
| SE_93_6_33 | SE_93 | SE_93_6 | CTGGTTTAGTCCTGGATGGG | 62.859056817921335 | 56.96537162413208 | 75.05204156022535 | 55.95626505135522 | 90.15053358199957 | 24.256683076315444 | 60.26499431482026 | 23.633837263904624 | 89.68651306034215 | 60.50757642063209 | 49.29896851731302 | 27.856344305470785 |
| SE_93_6_38 | SE_93 | SE_93_6 | ACACACGCACTCCTGTCTGC | 16109.258974854874 | 12899.158799169098 | 13014.616522660655 | 14723.004250954255 | 13363.490860390524 | 12195.04741661759 | 15448.285596236514 | 12612.90627100062 | 13609.262367948551 | 11554.766075506249 | 12200.320923070274 | 11525.562456388538 |
| SE_94_1_2 | SE_94 | SE_94_1 | GAGAAGCCAAGAGGAAGACG | 1563.8899825562153 | 1748.9368481093184 | 1605.7186786437685 | 1380.6883074299508 | 1283.319360402582 | 1876.4276979749734 | 1296.7735383814002 | 1305.5331704580915 | 1432.320253131999 | 894.4915213025973 | 1000.0647899226356 | 1202.4655291861557 |
| SE_94_1_20 | SE_94 | SE_94_1 | GTATGATGCTCTATGAAGGG | 229.76069043791938 | 222.86452407335884 | 244.90666193336693 | 218.6198262471553 | 254.5426830550576 | 191.45453428091832 | 150.66248578705066 | 356.3982659396817 | 207.78855501108973 | 244.21732651700904 | 325.13843522132635 | 178.74487596010422 |
| SE_94_1_3 | SE_94 | SE_94_1 | AGCCTCTGTAGCCCTTTGAG | 102.95879996038839 | 116.92892070216585 | 122.45333096668347 | 54.65495656178882 | 85.90815553108195 | 83.16577054736723 | 144.20552211046277 | 114.38777235729839 | 127.86988000682445 | 269.7325695859503 | 163.15611009301213 | 88.21175696732416 |
| SE_94_1_5 | SE_94 | SE_94_1 | ACAGGGCTAGAATTTACCAA | 1283.191780558946 | 1776.9198376790673 | 1737.059751374163 | 1551.1597195631493 | 1723.466083185286 | 1771.6041746808958 | 1332.2868386026337 | 1836.8218321506674 | 1991.7509781618558 | 1568.822945267473 | 1571.6980677305269 | 2337.61155963409 |
| SE_94_1_9 | SE_94 | SE_94_1 | TTACAGCACTTTGAGCCACT | 896.2834480762232 | 800.5133801917508 | 764.345791679137 | 869.2740710303555 | 826.2031254162079 | 719.9037013006476 | 873.8424176648937 | 1262.046909892507 | 618.9257386441434 | 688.1825559165867 | 1282.9469664148364 | 754.4426582371671 |
| SE_94_2_12 | SE_94 | SE_94_2 | TATCTCATCTTCTTGGACCC | 172.32051782843953 | 186.8863946265386 | 162.94193233469977 | 195.19627343496066 | 248.17911597868118 | 98.75935252499859 | 207.69899826357695 | 222.15807028070347 | 178.48504084285912 | 300.35086126867975 | 296.96759606857603 | 502.5748785112021 |
| SE_94_2_20 | SE_94 | SE_94_2 | TTTCAAAAGCTGCCTCACCC | 140.89098941947887 | 150.9082651797183 | 153.06666370835433 | 110.61122161314404 | 164.39214947305805 | 153.33688944670834 | 99.00677637434757 | 89.80858160283758 | 157.17339417505505 | 70.71367364820858 | 123.24742129328254 | 230.97552153286193 |
| SE_94_2_22 | SE_94 | SE_94_2 | CTAGGGGCCGGAAGTAATGA | 518.0453303270069 | 498.6968498323142 | 541.1647207237302 | 642.8463938458018 | 622.5689789721617 | 440.9518459 2302 | 520.8617365780893 | 809.2225879160943 | 502.5996672490461 | 683.8085142476253 | 795.8262060651958 | 529.270541803945 |
| SE_94_2_5 | SE_94 | SE_94_2 | TCTTGGTGTATAGTTCTGTG | 784.6544333823284 | 870.4708541161235 | 807.796973635057 | 780.7850937398402 | 788.0217229579492 | 584.7593241611759 | 898.5941116584806 | 706.1790574454702 | 670.4288847580032 | 801.9076393059519 | 949.5920364406246 | 1005.1497570224042 |
| SE_94_2_9 | SE_94 | SE_94_2 | GGTTCTGCCAGGCATCTGGA | 9.75399157519469 | 8.994532361705065 | 20.738064115325425 | 13.013084895664004 | 16.969512203670508 | 0.8663016098684087 | 38.74178205952731 | 28.36060471668555 | 13.319779167377547 | 10.93510417240339 | 39.90868879972959 | 0.0 |
| SE_95_1_1 | SE_95 | SE_95_1 | GTGCCATGGAGATAGAGCCA | 701.2036165723293 | 595.6379208418022 | 624.1169771850318 | 638.9424683771026 | 533.4790399028916 | 537.9785782282818 | 781.2926048671341 | 518.0537128247894 | 527.4632550281508 | 533.6330836132854 | 469.5139858791716 | 535.0739468675847 |
| SE_95_1_11 | SE_95 | SE_95_1 | ATTTTGGGCCAGCCAGAGCA | 23.843090517142578 | 11.992709815606755 | 23.700644703229056 | 31.23140374959361 | 15.9089176909411 | 1.7326202197368175 | 25.827854706351538 | 40.650200093915956 | 10.655823333902038 | 5.103048613788249 | 102.11929192871983 | 11.606810127279495 |
| SE_95_1_12 | SE_95 | SE_95_1 | TATCCAGGAAACATCTGTGG | 682.7794102636283 | 782.5243154683407 | 790.0214901076353 | 756.0602324380786 | 1022.4131102711481 | 1014.4491386559066 | 708.1136831991138 | 975.6048022539829 | 733.4758394835902 | 691.8275906407212 | 981.2842304874687 | 930.8661722078155 |
| SE_95_1_13 | SE_95 | SE_95_1 | ACAGGAGGTCCCATTAACCC | 1546.5495530892026 | 1621.0146100761797 | 1606.7062055064032 | 1592.8015912292742 | 1712.860138057992 | 1905.015931600631 | 1425.9128119131578 | 1482.3142731920982 | 1885.1927448228355 | 1379.281139612481 | 1687.902779235622 | 1678.3447444046149 |
| SE_95_1_3 | SE_95 | SE_95_1 | GTTATTGGCCACTAAAGGAA | 120.29922942740117 | 127.92223803313871 | 149.11655625781614 | 126.22692348794084 | 141.0590701930111 | 103.95721318420905 | 104.38757943381708 | 143.69373056450412 | 119.87801250639792 | 190.99981954464587 | 244.14727265716925 | 329.6334076147376 |
| SE_95_2_0 | SE_95 | SE_95_2 | ACATTTCTGAAGGCTACAGT | 466.0240419259685 | 624.6203028961851 | 616.2167622839555 | 514.0168533787282 | 643.7808692267499 | 518.9197558111769 | 447.6828149100933 | 491.5838150892162 | 627.8055914223951 | 778.5794170751213 | 651.4506554073506 | 774.1742354895423 |
| SE_95_2_1 | SE_95 | SE_95_2 | GAGCATGCATATAACATACG | 1884.6879276959519 | 1811.8985746412538 | 1789.3986750937938 | 1907.7182457043443 | 1783.9199704810862 | 2123.32607928747 | 1783.198135351021 | 1989.0237441302133 | 2053.0219623317926 | 1691.2961119983909 | 2211.410873490898 | 2331.8081545704504 |
| SE_95_2_2 | SE_95 | SE_95_2 | CATGCATATAACATACGTGG | 4397.966423571117 | 4409.31964220475 | 4419.182710289585 | 4451.7763428066555 | 4290.10480399045 | 4667.678871970987 | 3922.60543352714 | 4044.2222325993594 | 4161.986997166571 | 4297.495939754533 | 4188.064754042211 | 4814.504840795535 |
| SE_95_2_3 | SE_95 | SE_95_2 | CAGAAGCCTCAAAAGAGAGA | 53.105065242726646 | 93.94289355558624 | 83.93978332393624 | 118.41907255054244 | 71.05983235287025 | 95.29411208552496 | 27.980175931880833 | 84.13646065950047 | 108.33420389467071 | 23.328222234460565 | 25.823269223354437 | 95.17584304369186 |
| SE_95_2_4 | SE_95 | SE_95_2 | TATAACATACGTGGAGGTTT | 107.29390752141416 | 136.91677039484378 | 147.14150253254707 | 98.89944520704643 | 151.66501532030517 | 76.23528966841997 | 101.15909759987686 | 83.19110716894428 | 63.04695472558706 | 169.85861814466 | 170.19881988 11997 | 109.10401519642724 |
| SE_96_1_12 | SE_96 | SE_96_1 | CTTCAGGGCCCAGGATCTGG | 99.7074694353235 | 131.9198079716743 | 139.2412876314707 | 109.30991312357764 | 98.63528968383483 | 96.16042219539337 | 79.63588534458391 | 34.97807915057884 | 72.81479278166393 | 18.954180565499207 | 23.47569929395858 | 58.034050636397474 |
| SE_96_1_19 | SE_96 | SE_96_1 | GCAACCTTGCTCTTGGCAAT | 2759.295838938409 | 2294.6051447194254 | 2199.2223208712296 | 2473.7874386657627 | 2278.157013342766 | 2191.7645779670743 | 2569.8715432811978 | 2439.9573591255134 | 2242.1628265085537 | 2596.7227374733916 | 2314.703950384316 | 2780.991706496167 |
| SE_96_1_20 | SE_96 | SE_96_1 | TTATCCCCATCTGTGAAATG | 31.429528408960667 | 33.97934447755247 | 43.45118195591 9935 | 28.62878677046081 | 26.51486281823517 | 14.727271867762948 | 26.904015319116187 | 20.79777679223607 | 46.17523444690883 | 20.412194455152996 | 2.347569929395858 | 40.62383544547823 |
| SE_96_1_33 | SE_96 | SE_96_1 | GAAGAGTTGGAGGCTGAAAA | 295.8710777809056 | 446.7284406313516 | 439.4494538723721 | 304.5061865585377 | 348.9355946879748 | 297.1443676848642 | 179.71882233169612 | 193.79746565640179 3 | 364.0739639083196 | 88.9388472688 0909 | 99.77172199932397 | 114.9074202600967 |
| SE_96_1_8 | SE_96 | SE_96_1 | GACACTATTGGAAAGTACAG | 225.42558307116616 | 292.82199799773156 | 316.99612290568865 | 226.4276771845537 | 275.75457330964576 | 410.63099207762576 | 223.84140745504666 | 345.99937754 35637 | 281.49133307057883 | 258.7974654135469 | 320.4432953625346 | 236.7789265965017 |
| SE_96_2_11 | SE_96 | SE_96_2 | ATATGTATTATCGCTGGGCA | 135.47210521103736 | 165.89915244922676 | 171.82967409841066 | 145.74655083143685 | 130.45312506571702 | 341.32618328815306 | 136.67239782111022 | 163.54615386622 | 172.26914389808294 | 117.37011811712972 | 118.55228143449084 | 263.4745898892445 |
| SE_96_2_22 | SE_96 | SE_96_2 | TGATTTACTAGTCTCTAAAA | 5.418884208441 4945 | 13.99149478487 4547 | 8.887741763710896 | 9.109159426964803 | 6.363567076354461 | 23.390372966447035 | 5.380803063823238 | 8.508181415005666 | 5.327911666951019 | 5.832055558651 41 | 9.390279717583432 | 63.83745570003722 |
| SE_96_2_24 | SE_96 | SE_96_2 | AGTAAATCAAGACTCCAAAA | 15.172875783636185 | 18.98845720804403 | 13.825376076883616 | 11.711776406097604 | 12.727134152752882 | 1.7326202197368175 | 5.380803063823238 | 19.852423301679885 | 21.311646667804077 | 1.4580138896537853 | 19.954344399864794 | 37.14179240729438 |
| SE_96_2_9 | SE_96 | SE_96_2 | GCATTGCTGGCCAGGCGCAG | 3203.6443440306116 | 3680.7625209066396 | 3801.9784211429946 | 3449.7688058405274 | 3758.7469531130173 | 3389.005149805215 | 3266.147459740705 | 3853.26082750701 | 3612.3241101927906 | 4230.427300830458 | 4242.058862418316 | 3904.530926816822 |

| | | | | | | | | | | | | | | | |
|---|---|---|---|---|---|---|---|---|---|---|---|---|---|---|---|
| SE_96_3_13 | SE_96 | SE_96_3 | GCTGAAAAGGGAGAAGAAAG | 702.2873934140177 | 972.408887548781 | 977.6515940081986 | 638.9424683771026 | 714.8407015796201 | 374.24596746315257 | 513.3286122887368 | 476.45815924031723 | 754.7874861513943 | 400.95381965479095 | 385.00146842092073 | 184.54828102374395 |
| SE_96_3_15 | SE_96 | SE_96_3 | GGTAAGAAATGGACTAAGAG | 97.53991575194691 | 201.87728189604704 | 197.50537252690881 | 123.62430650880805 | 167.57393301124625 | 63.24063802039384 | 139.90087965940418 | 104.93423745173654 | 150.95749723027888 | 142.15635424124406 | 83.33873249355297 | 156.69193671827318 |
| SE_96_3_18 | SE_96 | SE_96_3 | ATGAATCCACTTCTGTACAT | 97.53991575194691 | 160.9021900260573 | 129.36601900512528 | 117.11776406097604 | 170.75571654943448 | 174.99464219341857 | 79.63588534458391 | 60.50262339559584 | 124.31793889552377 | 126.84720839987932 | 34.03976397623994 | 106.78265317097134 |
| SE_96_3_19 | SE_96 | SE_96_3 | AGTGGATTCATCATAATCCT | 148.47742731129696 | 184.8876096572708 | 156.02924429625796 | 130.13084895664005 | 164.39214947305805 | 55.44384703157816 | 109.76838250199404 | 65.22939084837677 | 133.19779167377547 | 55.404527806843845 | 9.390279717583432 | 16.249534178191293 |
| SE_97_1_13 | SE_97 | SE_97_1 | CCAAGGCCACCATCCAGCCG | 4.335107366753196 | 3.9975699385355847 | 5.925161175807264 | 14.314393385230405 | 5.3029725636470335 | 0.0 | 0.0 | 0.0 | 0.8879852778251698 | 0.0 | 0.0 | 0.0 |
| SE_97_1_18 | SE_97 | SE_97_1 | ACTTGTGCCCAGAAGAGTCA | 1213.8300626908947 | 1328.192612078448 | 1222.5582559415654 | 1482.19036961613 | 1184.6840707187473 | 1418.149649854585 | 1505.5486972577419 | 1111.7357048940735 | 1674.7402339782702 | 1650.471723088085 | 1237.1693527916173 | 1231.4825545043543 |
| SE_97_1_32 | SE_97 | SE_97_1 | CACGTCCCAGAGCAGGGCCA | 52600.0252344999 | 44214.1229126882 | 44246.14108034075 | 49466.63961388758 | 45839.955434677686 | 42450.0616936619 | 51309.185695392865 | 41797.85923145116 | 43177.39614897106 | 38512.70788825991 | 36264.086484342515 | 35208.09784008962 |
| SE_97_1_49 | SE_97 | SE_97_1 | CTATCAGACCCTGGGGACAG | 119.21545258571288 | 219.86634661945715 | 185.6550501752943 | 140.54131687317124 | 153.786204345764 | 103.09090307434064 | 63.4934761531142 | 183.3985771678999 | 285.0432741818795 | 155.27847924812812 | 93.90279717583432 | 207.76190127830296 |
| SE_97_1_9 | SE_97 | SE_97_1 | GCATGCCTGCTGACCCCGGC | 2643.331716877761 | 2871.254608353184 | 2922.091986535616 | 2898.0140062643736 | 2837.090321551163 | 2814.64154696246 | 2541.8913673500974 | 2897.5084485547072 | 2753.6423465358516 | 3383.321230941609 | 2727.876257957987 | 3177.9446128491254 |
| SE_97_2_16 | SE_97 | SE_97_2 | GAAGCCCTCAAGAACCCCCC | 176.65562519519273 | 284.8268581206604 | 256.75698428498146 | 245.94730452804967 | 216.36128059679896 | 270.28875427894354 | 163.5764131402264 | 272.2618052801813 | 215.78042251151626 | 386.3736807582531 | 273.4918967746175 | 253.02846077469297 |
| SE_97_2_19 | SE_97 | SE_97_2 | AGACACACACAGCTCCTGGG | 1719.9538477593303 | 2040.759453622416 | 2188.3595275981497 | 1786.6965561746679 | 2156.188644378884 | 2725.411605646014 | 1512.0056609343296 | 1970.1166743190895 | 2164.908107337764 | 1733.5785147983506 | 1859.2753840815196 | 1861.732344415631 |
| SE_97_2_20 | SE_97 | SE_97_2 | CAGACACACACAGCTCCTGG | 3935.1937121702135 | 4276.400441748442 | 4423.132817740123 | 4129.051837394189 | 4487.3753833581195 | 4042.202972645995 | 4437.010206428641 | 4501.773322028553 | 4118.475718553137 | 4205.641064706344 | 4236.189937594826 | 3887.1207116259025 |
| SE_97_2_28 | SE_97 | SE_97_2 | AGATATGGCCCCCATCAGAC | 932.048083851937 | 1255.2369607001735 | 1203.7952455515092 | 1076.1821208714132 | 1282.2587658898528 | 1449.3368138098479 | 1006.2101729349454 | 1495.5492220598846 | 1276.922829512594 | 1447.0787854813818 | 1916.7908473517182 | 1435.7624127444735 |
| SE_97_2_29 | SE_97 | SE_97_2 | GTGGAGCTCCAGTCTGATGG | 189.6609472954523 | 223.86391655799275 | 256.75698428498146 | 264.1656233819793 | 238.63376536411653 | 306.6737788934167 | 213.0798013274002 | 186.23463763956843 | 304.57895029403323 | 312.743979330737 | 212.45507861032516 | 229.814840520134 |
| SE_97_3_1 | SE_97 | SE_97_3 | GCATCCCTCAACATTCTTCC | 1662.5136751498505 | 1640.0030672842236 | 1574.1178190394633 | 1566.775421437946 | 1588.7705800686513 | 1558.4918876532672 | 1676.6582346873208 | 1358.472965929238 | 1561.0781184166485 | 2291.9978345357504 | 1984.870375304198 | 1486.8323773045033 |
| SE_97_3_11 | SE_97 | SE_97_3 | TGACTAGCCACACTCACATG | 4213.724360484106 | 3316.9836564999014 | 3250.938431792919 | 3896.117617761803 | 3439.5080047814663 | 3646.2992524361325 | 3693.3832230082703 | 3361.677012417794 | 3259.793954896198 | 3347.599890645091 | 3231.430007813399 | 3463.4721419802013 |
| SE_97_3_2 | SE_97 | SE_97_3 | TCTTTGGGACCAGGGTCCAG | 15.172875783636185 | 24.984812115847404 | 11.850322351614528 | 23.423552812195208 | 27.575457330964575 | 0.0 | 7.533124289352532 | 5.67212094333711 | 74.59076333731426 | 5.832055558615141 | 0.0 | 24.37430126728694 |
| SE_97_3_22 | SE_97 | SE_97_3 | AGAGACAGAAATAAACTGCC | 1315.705085809595 | 1673.9824117617761 | 1623.4941621711903 | 1323.4307338890292 | 1631.1943605778276 | 1916.27796302892 | 1422.684330074864 | 1571.1775013043796 | 1823.0337753750737 | 1244.4148548195058 | 1925.0073421046036 | 1657.4524861755117 |
| SE_97_3_34 | SE_97 | SE_97_3 | CACAGCCCAGATGTTCTCCC | 173.40429467012783 | 167.89793741849456 | 163.9294591973343 | 178.27926307059687 | 187.725228753105 | 199.251325269734 | 94.70213392328898 | 259.0268564123947 | 230.87617223454416 | 170.5876250894929 | 358.0044142328683 | 189.19100507465575 |
| SE_97_4_11 | SE_97 | SE_97_4 | TGGAGCCACAGGATTAATCG | 277.4468714722045 | 286.8256430899282 | 303.170746828805 | 226.4276771845537 | 305.45121966606916 | 283.28340592696964 | 238.90765603375175 | 288.33281461963645 | 286.81924473752986 | 343.36227101346645 | 504.7275348201095 | 316.86591647473017 |
| SE_97_4_12 | SE_97 | SE_97_4 | GGGCTGACATATGGGTGAGA | 361.98146512389184 | 467.7156828086634 | 447.34966877344846 | 400.80301478645134 | 658.6291924049616 | 479.9358008670984 | 307.7819352506892 | 571.9388617864919 | 415.57711002217945 | 261.71349319285446 | 573.9808477372873 | 314.5445544492743 |
| SE_97_4_16 | SE_97 | SE_97_4 | GCAGTAGACAGGCCAGAGCT | 4901.922654956176 | 4730.124629772231 | 4926.77151768374 | 4980.107589570614 | 5020.854423261011 | 5502.801817884132 | 5389.412348725355 | 5286.416719190187 | 4143.339306332242 | 5724.162530780761 | 5087.184037000824 | 5660.641299074209 |
| SE_97_4_20 | SE_97 | SE_97_4 | CAGGGAGGATTCCCAGGCAG | 3607.893105980347 | 4983.97032086924 | 4959.35990415068 | 3995.017062968849 | 4732.372715798613 | 4708.3954471348015 | 3612.6711770509214 | 4573.620187310823 | 4655.706811637365 | 5030.147919305559 | 4515.550759192933 | 4302.644514182509 |
| SE_97_4_33 | SE_97 | SE_97_4 | TCACCTGTCCAAGGCAGCCG | 1400.239679461282 | 1577.041340752288 | 1634.3569576601703 | 1266.1731603481076 | 1445.5903208501813 | 1347.1122208453755 | 1104.1407886965283 | 1347.1287240425636 | 1267.1549914565173 | 1047.5829797162448 | 1014.1502094990107 | 1983.6038507520657 |
| SE_97_5_11 | SE_97 | SE_97_5 | GAGAAGGGCCCGATAGTACA | 505.0400082267473 | 665.5953947661749 | 636.9548263992809 | 595.9992882214113 | 738.1737808596671 | 434.02136504407275 | 648.9248494970824 | 584.2284571637223 | 616.2617828106678 | 1129.9607644816836 | 487.12076034964053 | 572.2157392748791 |
| SE_97_5_23 | SE_97 | SE_97_5 | ATTAAGAGCAGGGAGTTGCA | 186.4096167703874 | 148.90948021045054 | 150.1040831204507 | 162.66356119580004 | 193.02820131675202 | 208.7807364782865 | 217.3844437784588 | 167.32756782844476 | 342.76231724051553 | 145.80138896537852 | 279.3608215981071 | 175.26283292192036 |
| SE_97_5_25 | SE_97 | SE_97_5 | GCAGTCTGACTCCAGAGCAG | 1852.174622445303 | 1969.8025872134094 | 1971.10361781855 | 1894.705160808679 | 1946.1909308584613 | 2225.550672251942 | 1818.7114355722542 | 1703.5269899822454 | 2411.768014573161 | 1525.8115355226864 | 1673.817359659247 | 2140.295787470339 |
| SE_97_5_36 | SE_97 | SE_97_5 | GTAACCTTAACAAGCCTTAT | 74.78060207649263 | 93.94289355558624 | 104.67784743926167 | 48.14841411395682 | 85.90815553108195 | 227.8395588953915 | 31.208657770174778 | 52.93979547114636 | 53.27911666951019 | 113.72508339299526 | 196.02208910455414 | 68.48017975094902 |
| SE_97_5_9 | SE_97 | SE_97_5 | TCGGGCCCTTCTCACCCACA | 2400.565704339582 | 2118.71206742386 | 2240.6984513177804 | 2333.246121792556 | 2359.82279082293 | 1825.3154014927372 | 2510.6827095799226 | 2337.8591821454456 | 2421.535852629238 | 3166.077161383195 | 2611.671546452892 | 1551.8305140172683 |
| SE_97_6_12 | SE_97 | SE_97_6 | CAGAGGTCCCTCTCTCTGAA | 747.8060207649263 | 848.4842194541778 | 785.0838557944625 | 796.4007956146371 | 734.9919973214788 | 776.2138584420942 | 851.2430446968361 | 813.9493553688753 | 876.4414692134426 | 653.1902225648959 | 787.6097113123104 | 732.3897190313361 |
| SE_97_6_18 | SE_97 | SE_97_6 | AAATTGGCAGGGCAACCAAG | 62.859056817921335 | 72.95565137827442 | 64.18924607124536 | 106.70729614444484 | 118.78658542569356 | 198.3850151598656 | 125.91079169346375 | 96.42605603673087 | 63.04695472558706 | 74.35870837234305 | 58.68924823489645 | 155.53125570554522 |
| SE_97_6_22 | SE_97 | SE_97_6 | GGGTGCTTCCCAGAGATGGG | 55.27261892610324 | 86.94714616314897 | 95.79010567555078 | 79.37981786355043 | 143.1802592184699 | 35.51871450460476 | 79.63588534458391 | 175.8357492434504 | 152.7334677859292 | 82.37778476543888 | 139.68041079905356 | 109.10401519642724 |
| SE_97_6_37 | SE_97 | SE_97_6 | CCATCAGCCCATTTTCCATT | 1005.7449090867414 | 1100.3311255819197 | 1104.0550324254202 | 1018.9245473304916 | 1094.5335371367478 | 996.25662634867 | 1060.0182035731777 | 1045.5609605551406 | 875.5534839356174 | 1257.5369798263898 | 1069.3181028398133 | 1053.898359556978 |
| SE_97_6_6 | SE_97 | SE_97_6 | TCAGGCAGATGTTTGAGGCA | 12722.45634457894 | 11783.83678631827 | 11705.15590280725 | 12240.107652861563 | 11159.575462938818 | 10688.534135556427 | 12418.893471304033 | 11782.88590629229 | 11570.448170061962 | 13014.960985994514 | 11558.260547380507 | 11712.432099437738 |
| SE_97_7_1 | SE_97 | SE_97_7 | TTTAACCAGCTACCCCAGTG | 45.51862735090855 | 65.95990398583714 | 48.38881626909266 | 70.27065843658562 | 50.90853661101153 | 33.78609428486794 | 55.960351863761666 | 28.36060471668555 | 125.20592417334893 | 115.91210422747594 | 19.954344399864794 | 32.49906835638259 |
| SE_97_7_10 | SE_97 | SE_97_7 | ACACTGACCCAACAGCAGAC | 2391.895489606076 | 2814.2892367290515 | 2996.1565012332067 | 2443.8573434057002 | 2668.4557940271875 | 2533.9570713650955 | 2296.5267476397576 | 2897.5084485547072 | 2869.968417930949 | 2535.4861541079326 | 3034.2341337441467 | 2772.866939407071 |
| SE_97_7_13 | SE_97 | SE_97_7 | GACTAGAGAGATCCTGACCT | 557.0612966277856 | 666.5947872508087 | 622.1419234597628 | 602.5058306692434 | 650.1444363031263 | 508.5240344927559 | 580.0505702801449 | 735.4850156527119 | 733.4758394835902 | 784.4114726337365 | 663.1885050543299 | 552.4841620585039 |
| SE_97_7_20 | SE_97 | SE_97_7 | GGTTGAACAGAAGCCACCAC | 752.1411281316795 | 742.5486160829848 | 749.5328887396189 | 801.6060295729027 | 922.7172260745839 | 483.4010413065721 | 831.8721536670726 | 717.5232993321445 | 798.2987647648276 | 711.5107781510472 | 743.0058826537891 | 450.3442329384444 |
| SE_97_7_27 | SE_97 | SE_97_7 | GCCCCTTTTCTCGTTTGTGA | 5.4188842084414945 | 12.992102300240651 | 10.862795488979984 | 14.314393385230405 | 8.484756101835254 | 0.0 | 3.228481838293942 | 0.945353490556185 | 32.855455279531284 | 0.7290069448268927 | 0.0 | 0.0 |
| SE_97_8_15 | SE_97 | SE_97_8 | GTGTCTTTACCACACATAGA | 35.76463577571386 | 52.9678016855965 | 35.55096705484358 | 37.737946197425615 | 29.69664635642339 | 127.34758615065608 | 22.599372868057596 | 19.852423301679885 | 47.063219724734 | 36.45034724134463 | 3.521354894093787 | 42.945197470934126 |
| SE_97_8_16 | SE_97 | SE_97_8 | GCTGGGATAGGAAGTGGAGC | 1038.2582143373904 | 1273.2260254235837 | 1211.6954604525856 | 1017.6232388409252 | 1229.2290402533824 | 1354.0427017243228 | 903.9749147223039 | 1209.1071144213606 | 1141.9490672831682 | 1159.8500492195863 | 930.8114770054577 | 810.1553468841087 |
| SE_97_8_18 | SE_97 | SE_97_8 | CTATGTGTGGTAAAGACACT | 427.00807562518975 | 418.7454510616025 | 433.5242926965648 | 512.7155448891617 | 399.8441312989863 | 656.6630632802538 | 465.97754532709234 | 543.5782570698063 | 348.09022890746655 | 498.6407502615946 | 548.1575785139329 | 587.3045924403424 |
| SE_97_8_26 | SE_97 | SE_97_8 | GTTGTACTGGGACCATCCGG | 246.01734306324386 | 361.7800794374704 | 304.15827369143955 | 281.0826337463425 | 317.11775930609264 | 539.7111984480186 | 223.84140745504666 | 245.7919075446081 | 423.5689775226059 | 115.18309728264904 | 208.93372371623136 | 467.7544481293636 |
| SE_97_8_36 | SE_97 | SE_97_8 | CAGAGCTAGAAGGTGCCTTA | 1004.6611322450531 | 956.4186077946387 | 1044.8034206673476 | 949.9551973834723 | 1103.018293238583 | 1097.6149092032738 | 979.3061576158292 | 1262.9922633830631 | 1007.8632903315677 | 926.5678268749806 | 1359.2429891202019 | 1422.994921604466 |
| SE_98_1_21 | SE_98 | SE_98_1 | TTAAGATTCTGCTATGCAAA | 13.005322100259587 | 12.992102300240651 | 11.850322351614528 | 13.013084895664004 | 23.33307928004695 | 1.7326202197368175 | 19.370891029763655 | 13.23494886778659 | 7.103882222601358 | 13.851131951710961 | 55.167893340802664 | 1.1606810127279494 |
| SE_98_1_27 | SE_98 | SE_98_1 | GCACGCGCTCGGGAGCAGCT | 8705.979369282106 | 7269.580933226961 | 7234.62179566067 | 8744.793049886212 | 7965.064790597845 | 8449.988811656458 | 9295.875373061024 | 8368.269098403349 | 7261.943602054239 | 6837.356135531426 | 9183.693563796596 | 7767.277337175437 |
| SE_98_1_30 | SE_98 | SE_98_1 | CGGGCCTGCAGGATGAGCCA | 535.3857597940197 | 135.9173779102099 | 222.1935440927724 | 442.44488645257616 | 126.21074701479941 | 1329.7860186480075 | 710.2660044246674 | 277.93392622351837 | 907.5209539373235 | 40.82438891030599 | 166.67746498710594 | 376.06064812385563 |
| SE_98_1_41 | SE_98 | SE_98_1 | GGTGAGCTCCTCCTGCCATG | 2644.4154937194494 | 2492.484856676937 | 2432.278662668882 | 2575.2895008519063 | 2368.307546924765 | 2006.3742144552346 | 2658.116713528679 | 2189.4386841281244 | 2381.5765151271053 | 2959.0391890523574 | 1832.2783298934671 | 1819.9478279574248 |
| SE_98_1_46 | SE_98 | SE_98_1 | GGAGGGGATTCCACACTCCA | 7031.544148873683 | 7918.18665575436 | 7890.339632450007 | 7453.895028236342 | 7352.041162240248 | 7740.480831674232 | 6960.60684336174 | 7498.54388709166 | 7748.5595343024315 | 8481.26679611607 | 7776.32539112378 | 8958.136056234314 |
| SE_98_2_12 | SE_98 | SE_98_2 | CCACTGCAGCGGTTGGCCAG | 26.010644200519174 | 12.992102300240651 | 31.60085960430541 | 24.724861301761607 | 15.9089176909411 | 197.51870504999718 | 10.761606127646475 | 11.34424188667422 | 117.21405667292241 | 11.664111117230282 | 10.564064682281362 | 46.42724050911798 |
| SE_98_2_15 | SE_98 | SE_98_2 | AGGGTCCCTGTCCATCCCTG | 1571.4764204480334 | 1908.8396456507417 | 1927.65243586263 | 1448.3563488874038 | 1544.2256105340161 | 939.080159097355 | 1020.2002609008858 | 765.7363273505099 | 1469.615634800656 | 617.468882268378 | 674.9263547013092 | 586.1439114276145 |
| SE_98_2_21 | SE_98 | SE_98_2 | TAACCCCTCTCAACACCGCA | 757.5600123401209 | 917.4423008939167 | 921.3625628380296 | 688.3921909806259 | 762.5674546524434 | 604.6844566881493 | 576.8220884418511 | 280.76998669518696 | 879.1054250469181 | 195.37386121360723 | 163.15611009301213 | 181.06623798556012 |
| SE_98_2_33 | SE_98 | SE_98_2 | GCCCCAGGCCAGGTTCAGAG | 31.429528408960667 | 24.984812115847404 | 43.451181955919935 | 48.14841411395682 | 43.48437502190568 | 91.82887164605133 | 16.142409191469714 | 20.79777679223607 | 23.975602501279585 | 0.0 | 3.521354894093787 | 62.67677468730927 |
| SE_98_3_12 | SE_98 | SE_98_3 | CTGCCTTCCCCCAAAACACA | 542.9721976858377 | 553.6634364871785 | 503.63869994361744 | 536.139097701357 | 675.5987046086321 | 602.085526358544 | 558.527358024852 | 765.7363273505099 | 753.8995008735692 | 292.33178487558393 | 679.6214945601009 | 526.949179778489 |
| SE_98_3_20 | SE_98 | SE_98_3 | ATTTTACAGGAGAGAAACGG | 8.670214733506391 | 17.98906472341013 | 23.700644703229056 | 27.32747828089441 | 18.030106716399914 | 1.7326202197368175 | 19.370891029763655 | 19.852423301679885 | 13.319779167377547 | 107.8930278343801 | 15.259204541073078 | 0.0 |
| SE_98_3_23 | SE_98 | SE_98_3 | GGGAAGCTCCCGGAACAACC | 425.92429878350146 | 537.6731567330362 | 546.1023550369028 | 420.3226421299473 | 466.66158560093896 | 453.94649757104617 | 477.8153120675035 | 551.1410849942558 | 572.7505041972345 | 526.3430141650165 | 534.0721589375577 | 683.6411164967623 |
| SE_98_3_26 | SE_98 | SE_98_3 | GCTGCACCTCCCGTGTTGGG | 2125.2863865507543 | 2276.6160799960153 | 1992.8292087965099 | 2158.8707841906585 | 2235.733232833589 | 2482.8447748828594 | 2130.798013274002 | 1856.6742554523473 | 2424.1998084627135 | 1948.635563522284 | 2227.8438629966695 | 2193.6871140558246 |
| SE_98_3_9 | SE_98 | SE_98_3 | TGTGGGGTGCAGGCTGGGAA | 3244.827864014767 | 3613.8032244361684 | 3518.5582115668803 | 3477.096284121422 | 3226.328507722855 | 3235.668260358507 | 3596.528767859452 | 3364.5130728894624 | 4006.589573547166 | 2911.653737638609 | 3312.421170377556 | 3312.583610325568 |
| SE_98_4_14 | SE_98 | SE_98_4 | GGGCAATTGACTTAACATCT | 347.89236618194394 | 437.7339082696465 | 435.49934642183393 | 377.3794619742561 | 479.3887197536919 | 577.8288432822286 | 362.6661265016862 | 483.07563367421056 | 528.351240305976 | 545.2971947305157 | 605.6730417841313 | 547.8414380075922 |
| SE_98_4_15 | SE_98 | SE_98_4 | AGTCAATTGCCCAAATACCC | 707.7062776224592 | 762.5364657756628 | 851.2481555909769 | 749.5536899902467 | 726.5072412196437 | 628.9411397644648 | 778.0641230288401 | 751.5560249921671 | 951.0322325507568 | 968.1212227301135 | 944.8968965818328 | 779.977640553182 |
| SE_98_4_18 | SE_98 | SE_98_4 | TACCCAGGATCAAACAGCAG | 1126.0441385141426 | 1232.250933553594 | 1147.5062143813402 | 1178.9854915471587 | 1138.0179121586534 | 1282.1389626052448 | 985.763121292417 | 1213.8338818741415 | 982.999702552463 | 1522.1665007985519 | 1059.92782312223 | 1250.0534507080015 |
| SE_98_4_19 | SE_98 | SE_98_4 | ACCCAGGATCAAACAGCAGA | 280.6982019972694 | 316.8074176289451 | 258.73203801025056 | 415.11740817168175 | 363.7839178661865 | 101.35828285460381 | 445.5304936845641 | 380.03210320358636 | 251.29983362452305 | 313.47298627556387 | 524.6818792199742 | 294.81297723289913 |
| SE_98_4_20 | SE_98 | SE_98_4 | TCCCTCTGCTGTTTGATCCT | 826.9217302081721 | 735.5528686905476 | 724.8447171737553 | 864.0688370720899 | 788.0217229579492 | 730.2994226190685 | 716.7229681012552 | 949.1349045184097 | 776.0991328191984 | 853.6671323922913 | 632.6700959721837 | 1028.363377276963 |
| SE_98_5_22 | SE_98 | SE_98_5 | GAAATGGACGAGATAGGCAG | 644.8472208045379 | 788.520670376144 | 718.919555997948 | 594.697979731845 | 679.8410826595497 | 604.6844566881493 | 659.6864556247289 | 622.9879502765259 | 792.9708530978767 | 621.1139169925125 | 819.3019053591545 | 538.5559899057686 |
| SE_98_5_26 | SE_98 | SE_98_5 | GGTCTGTGCAGAGGCCGCTG | 111.62901469389479 | 188.8851795958064 | 213.30580232906152 | 145.74655083143685 | 190.90701229129323 | 256.42779252104896 | 108.6922218892294 | 225.93948424292822 | 326.77858223966246 | 160.3815278619164 | 211.28129364562722 | 172.94147089646447 |
| SE_98_5_30 | SE_98 | SE_98_5 | ACTCTGCACCTCCGGACACG | 793.3246481158347 | 867.4726766622219 | 898.6494449974351 | 851.0557521764259 | 801.8094516234315 | 673.989265477622 | 743.6269834203714 | 1049.3423745173654 | 864.0096753238902 | 605.8047711511477 | 915.5522724643846 | 1059.7017646206177 |
| SE_98_5_41 | SE_98 | SE_98_5 | GCTCCCACTCATTTGACAGG | 277.4468714722045 | 286.8256430899282 | 374.2726809384922 | 244.64599603848328 | 345.7538111497866 | 386.3743090013103 | 284.1064017698669 | 542.6329035792502 | 281.49133307057883 | 387.8316946479069 | 720.7039683245284 | 425.96993167115744 |
| SE_98_5_8 | SE_98 | SE_98_5 | GGCGCTGATGGAGCGATTAG | 303.4575156727237 | 421.7436285155042 | 424.63655093285394 | 303.2048780689713 | 376.5110520189394 | 421.02671339604666 | 291.63952605921946 | 373.4146287696931 | 375.6177725200468 | 390.74772242721446 | 665.5360749837257 | 515.3423696512095 |
| SE_98_6_11 | SE_98 | SE_98_6 | GTGGATCTCTGTGTTGGCCT | 19.50798315038938 | 37.97691441608806 | 18.763010390056337 | 36.43663770785921 | 22.272484767317543 | 17.326202197368175 | 20.4470516425283 | 11.34424188667422 | 18.647690834328564 | 47.38545141374802 | 39.90868879972959 | 18.57089620364719 |
| SE_98_6_12 | SE_98 | SE_98_6 | AGGCCAACACAGAGATCCAC | 5858.897606166944 | 5201.8378825194295 | 5282.281188232176 | 5698.429875811267 | 5428.122716149103 | 5716.780415021629 | 6300.920387737011 | 5356.372877491344 | 5558.787839185563 | 6020.8683573253065 | 5914.702437112865 | 5644.391764896018 |
| SE_98_6_3 | SE_98 | SE_98_6 | GGGCAGCGTCCTCAGCCAGC | 3185.2201377219103 | 3005.1732012941256 | 2786.800806354683 | 3322.2405738630205 | 3076.7846814280088 | 2826.7698885006175 | 3191.8923774599443 | 3258.63348194717 | 2670.1717304202857 | 3131.084828031504 | 2805.3460656280504 | 3417.044901471083 |
| SE_98_6_5 | SE_98 | SE_98_6 | GCTGCCCCAGGCCCTGACTG | 1020.9177848703775 | 1000.39187711853 | 949.0133149917968 | 1060.5664189966164 | 1095.5941316494773 | 810.8662628368305 | 1157.9488193347606 | 1136.3148956485343 | 1030.950907555022 | 1102.2585005782616 | 1106.879221710147 | 1013.2745241114999 |
| SE_98_6_9 | SE_98 | SE_98_6 | GTTGGCCTCGGCCTCAGTCA | 265.5253262136332 | 266.83779339725027 | 292.307951339825 | 346.1480582246625 | 249.2397104914106 | 161.99999054539242 | 309.9342564762185 | 275.09786575184984 | 211.34049612239042 | 494.2667085926332 | 214.80264853972102 | 343.561579767473 |
| SE_98_7_15 | SE_98 | SE_98_7 | ACGAGGCCACCAGCTCATGG | 21635.437090623513 | 21197.114599084936 | 21516.235283081445 | 21482.000545762137 | 20484.32241885576 | 17564.43747758199 | 20842.002587412928 | 17879.470566889126 | 20119.082439684873 | 17959.81509275533 | 16450.596280241476 | 15515.983778147229 |
| SE_98_7_22 | SE_98 | SE_98_7 | GGAGAAGAGAGCAGTCGTGG | 2603.231973735294 | 3012.168948686563 | 2871.728116541254 | 2680.695488506785 | 2751.1821660200812 | 3533.6789381532394 | 2545.119849188391 | 2856.858248460791 | 2769.6260815367045 | 2844.585098714535 | 2334.6582947841807 | 3450.7046508401936 |
| SE_98_7_32 | SE_98 | SE_98_7 | GCAGAGGAAGCCGGGTACAG | 1071.8552964297276 | 1473.104522350363 | 1552.3922280615031 | 1395.0027008151812 | 1782.8593758981328 | 2445.5934401585178 | 1267.7172018367546 | 1902.0512229990443 | 1785.7383937064164 | 2180.459771977236 | 2004.8247197040628 | 3115.267838161816 |
| SE_98_7_38 | SE_98 | SE_98_7 | ACGGCTCAGGGGCAGACAGA | 7341.504325596537 | 7164.644722340402 | 7356.087599764718 | 7515.056527245963 | 7139.922259694366 | 7127.999583997267 | 7556.7998228333545 | 7322.708137848209 | 7634.897418740809 | 7569.279108137626 | 7095.530111598981 | 6892.123853578564 |
| SE_98_7_43 | SE_98 | SE_98_7 | AGAAAGGGCTTGGCTGAAGT | 525.631768218825 | 757.5395033524933 | 696.2064381573535 | 602.5058306692434 | 739.2343753723965 | 540.5775085578871 | 583.2790521184389 | 722.2500667849254 | 613.5978269771923 | 605.0757642063209 | 613.8895365370169 | 769.5315114386304 |
| SE_99_1_29 | SE_99 | SE_99_1 | AGCCTTCCTGACCAACAGAC | 340.30592829012585 | 313.8092401750434 | 309.0959080046123 | 357.85983463076013 | 308.63300320425736 | 311.87163955262713 | 259.35470767628004 | 361.1250333924627 | 303.69096501620805 | 219.4310903928947 | 268.79675691582577 | 342.40089875474507 |
| SE_99_1_30 | SE_99 | SE_99_1 | TCTGTTGGTCAGGAAGGCTG | 645.9309976462262 | 577.648856118392 | 572.7655803280355 | 550.4534910865874 | 562.1150917465856 | 405.43313141841526 | 487.5007575823853 | 335.60048914744567 | 721.0440455940378 | 215.05704872393332 | 288.75110131569056 | 495.6107924348344 |
| SE_99_1_31 | SE_99 | SE_99_1 | CTGTTGGTCAGGAAGGCTGA | 560.3126271528505 | 473.7120377164668 | 485.86321641619566 | 481.4841411395682 | 551.5091466192915 | 560.5026410848604 | 581.1267308929097 | 624.8786572576383 | 490.16787335949374 | 262.44250013768135 | 584.5449124195686 | 261.15322786378863 |

| | | | | | | | | | | | | | | | |
|---|---|---|---|---|---|---|---|---|---|---|---|---|---|---|---|
| SE_99_1_32 | SE_99 | SE_99_1 | TTTGGGAAGCGAGACGGCTA | 458.4376040341504 | 473.7120377164668 | 516.4765491578665 | 448.95142890040813 | 528.1760673392446 | 535.3796478986766 | 455.21593919944587 | 407.44735442971574 | 300.1390239049074 | 413.3469377168481 | 280.53460656280504 | 785.7810456168218 |
| SE_99_1_33 | SE_99 | SE_99_1 | TGGAAGTTTGGGAAGCGAGA | 37.93218945909046 | 80.95079125534559 | 74.06451469759081 | 50.751031093089615 | 111.36242383658771 | 19.9251325269734 | 36.58946083399801 | 80.35504669727572 | 53.27911666951019 | 61.23658336545898 | 116.20471150509498 | 39.46315443275028 |
| SE_99_3_14 | SE_99 | SE_99_3 | GCTGCGAAGCTGTCAGCACA | 3196.0579061387934 | 2813.289844244418 | 2849.0149987006594 | 2996.91345147142 | 2693.910062332693 | 2643.9784553183836 | 2908.862136302842 | 2748.14259704683 | 2890.3920793209277 | 2592.3486958044305 | 3014.2797893442817 | 2899.3811697944175 |
| SE_99_3_15 | SE_99 | SE_99_3 | TTTTTGATGTCACAACAGGC | 4.335107366753196 | 5.9963549078033775 | 3.950107450538176 | 0.0 | 3.1817835381882205 | 0.8663101098684087 | 9.685445514881827 | 0.0 | 0.0 | 1.4580138896537853 | 0.0 | 0.0 |
| SE_99_3_19 | SE_99 | SE_99_3 | AATGTCCCCAAGAAAGAGTG | 309.9601767228535 | 382.76732161478225 | 406.86106740543215 | 368.27030254729135 | 290.60289648785744 | 326.5989114203901 | 305.62961402515987 | 380.03210320358636 | 260.17968640277473 | 248.5913681859704 | 288.75110131569056 | 409.72039749296613 |
| SE_99_3_4 | SE_99 | SE_99_3 | CTAACCACAGTAACCCCAGG | 733.7169218229784 | 795.5164177685814 | 845.3229944151697 | 799.0034125937699 | 904.687119358184 | 1042.1710621716957 | 867.3854538883058 | 1224.2327702702596 | 1015.8551578319942 | 1027.1707852610919 | 1436.7127967902652 | 1277.9097950134724 |
| SE_99_3_6 | SE_99 | SE_99_3 | TTAACAGTACTTACCACCTG | 291.5359704141524 | 314.8086326596773 | 293.2954782024596 | 340.94282426639694 | 287.42111294966924 | 183.65774329210265 | 285.1825623826316 | 266.5896843368442 | 204.23661389978906 | 396.5797779858296 | 374.4374037386394 | 275.081400016524 |
| SE_99_4_0 | SE_99 | SE_99_4 | CTTTGAATCATTCTTTTACA | 49.85373471766175 | 74.95443634754221 | 60.23913862070719 | 63.76411598875362 | 97.57469517110542 | 18.192512307236584 | 29.056336544645482 | 151.2565584889896 | 76.3667338929646 | 121.74415978609107 | 69.25331291717781 | 91.69380000550801 |
| SE_99_4_18 | SE_99 | SE_99_4 | ACGGAGTTCCAGGAATCGTA | 4535.606082465531 | 4858.046867805369 | 4755.929370447964 | 4446.57110884839 | 4552.071648634614 | 4237.989057476256 | 4595.205816505045 | 5096.400667588393 | 4839.519764147175 | 5265.617162484646 | 5381.8040631400045 | 4208.6293521515445 |
| SE_99_4_34 | SE_99 | SE_99_4 | GGGAGAGGCTGGGCTCTCCG | 46.60240419259685 | 93.94289355558624 | 128.37849214249073 | 100.20075369661284 | 75.30221040378788 | 229.5721791151283 | 59.18883370205561 | 63.33868386726439 | 62.15896944776188 | 116.64111117230283 | 99.77172199932397 | 104.46129114551545 |
| SE_99_4_42 | SE_99 | SE_99_4 | CTCCCCAGCCCAAAACCTCA | 117.04789890233629 | 139.91494784874547 | 163.9294591973343 | 122.32299801924164 | 133.63490860390525 | 114.35293450262995 | 129.1392735317577 | 167.32756782844476 | 136.74973278507613 | 169.858618144666 | 118.55228143449084 | 134.63899747644214 |
| SE_99_4_5 | SE_99 | SE_99_4 | TGTCACACCCACAGGCTCCA | 775.984218648822 | 882.4635639317303 | 844.3354675525352 | 831.5361248329299 | 815.5971802889138 | 705.1764294328847 | 714.570646875726 | 933.0638951789546 | 856.0178078234637 | 712.968792040701 | 799.3475609592897 | 1026.0420152515073 |
| SE_99_5_2 | SE_99 | SE_99_5 | GTGTTCCTAGTCCTGAACTG | 71.52927155142773 | 88.94593113241676 | 63.20171920861082 | 104.10467916531204 | 89.08993906927017 | 5.197860659210452 | 111.92070372752333 | 98.31676301784324 | 150.95749723027888 | 55.404527806843845 | 42.25625872912545 | 33.659749369110536 |
| SE_99_5_33 | SE_99 | SE_99_5 | TGGACCCCTCTAGGATCCTG | 10.837768416882989 | 25.984204600481302 | 26.66322529113269 | 15.615701874796805 | 30.757240869152795 | 36.38502461447317 | 6.456963676587884 | 0.0 | 24.863587779104755 | 2.9160277793075706 | 0.0 | 4.642724050911798 |
| SE_99_5_34 | SE_99 | SE_99_5 | CCTGCCCCAGGATCCTAGAG | 1098.9497174719352 | 1200.2703740453094 | 1105.0425592880547 | 1085.291280298378 | 1146.5026682604887 | 1371.368903921691 | 1042.7996337689433 | 1483.2596266826542 | 1082.454053668882 | 1510.5023896813216 | 1369.807053802483 | 1483.3503342663194 |
| SE_99_5_5 | SE_99 | SE_99_5 | GTTTTCATACCCCTGGCGAT | 4.335107366753196 | 7.9951398770711695 | 5.925161175807264 | 9.109159426964803 | 3.1817835381882205 | 5.197860659210452 | 0.0 | 64.28403735782058 | 30.191499446055772 | 0.7290069448268927 | 89.2076573170426 | 75.44426582731671 |
| SE_99_5_8 | SE_99 | SE_99_5 | CCAAGTGAGACACTGTCAGG | 506.1237850684356 | 693.5783843359239 | 584.61590267965 | 612.9162985857746 | 625.75076251035 | 468.67376943880913 | 465.97754532709234 | 441.4800800897384 | 657.1091055906256 | 519.0529447167476 | 388.52282331501453 | 342.40089875474507 |
| SE_99_6_2 | SE_99 | SE_99_6 | TTTCTCCCTCGGCTTTATTG | 0.0 | 0.0 | 0.987526862634544 | 2.602616979132801 | 0.0 | 0.0 | 0.0 | 0.945353490556185 | 0.0 | 0.0 | 0.0 | 0.0 |
| SE_99_6_23 | SE_99 | SE_99_6 | GGAAGGGCAGTCCAGGCACA | 15229.232179403976 | 13253.943131214732 | 13296.061678511502 | 15049.632681835421 | 13202.280494455656 | 12994.65164802613 | 15869.064395827492 | 13777.58177136584 | 12596.959151227858 | 13347.388152835578 | 14071.334156798774 | 12971.770998247563 |
| SE_99_6_25 | SE_99 | SE_99_6 | TAAGCCCCTGAATCCCACCA | 1349.302167901932 | 1541.0632113054678 | 1520.791368457198 | 1558.9675705005477 | 1459.3780495156636 | 882.7700019559085 | 1284.935771640989 | 1884.0895066784767 | 1348.849637016433 | 2543.5052305010286 | 1738.375532717633 | 1355.6754228662448 |
| SE_99_6_31 | SE_99 | SE_99_6 | CTAAATCCCCAAGGTTCGGG | 82.36703996831072 | 64.96051150120326 | 77.02709528549444 | 104.10467916531204 | 77.42339942924669 | 115.21924461249836 | 102.23525821264151 | 67.12009782948914 | 70.15083694818841 | 5.103048613788249 | 46.95139858791716 | 76.60494684004466 |
| SE_99_6_32 | SE_99 | SE_99_6 | AGTCAGCCACCCAAACCTTG | 1202.9922942740118 | 1339.1859294094209 | 1314.398254166578 | 1249.2561499837443 | 1416.9542690064875 | 1417.2833397447166 | 1309.6874657345759 | 1252.593374986945 | 1284.0267117351955 | 1175.8882020057779 | 940.2017567230412 | 1609.8645646536659 |
| SE_99_7_2 | SE_99 | SE_99_7 | CTCGTTGAGGCTCATCACCT | 2601.0644200519173 | 2356.567478766727 | 2415.4907060040946 | 2403.5167802291417 | 2531.639101885094 | 2738.40625729404 | 2574.1761857330366 | 2623.3559362934134 | 2276.7942523437355 | 2279.6047164736933 | 2196.1516689498253 | 2649.8347520579086 |
| SE_99_7_3 | SE_99 | SE_99_7 | CCTGCATGGAGTGCTCGTTG | 12131.797965858817 | 10754.462527145357 | 10831.19462937568 | 11569.933780734866 | 11007.910447618513 | 10887.78546082616 | 11965.829853330115 | 11006.750690545661 | 10134.575975818663 | 10905.214887665486 | 10389.17072254137 | 12563.211281767324 |
| SE_99_7_4 | SE_99 | SE_99_7 | ACTCCATGCAGGCGCTGTCC | 1230.0867153162192 | 1475.1033073196309 | 1435.864058270627 | 1162.0684811827955 | 1551.6497721231221 | 1133.1336237078785 | 1071.8559703135888 | 1612.7730548888517 | 1301.7864172916989 | 1382.1971673917885 | 1547.0485834718704 | 1327.8190785607742 |
| SE_99_7_5 | SE_99 | SE_99_7 | GCGCCAGGACAGCGCCTGCA | 17.340429467012783 | 1.9987849692677924 | 0.987526862634544 | 3.9039254686992013 | 2.1211890254588135 | 18.192512307236584 | 21.52321225529295 | 15.12565584889896 | 5.327911666951019 | 0.7290069448268927 | 0.0 | 1.1606810127279494 |
| SE_99_7_6 | SE_99 | SE_99_7 | GCTCAAGTAGAGCTTGCGCC | 1633.2517004242663 | 1432.1294304803732 | 1574.1178190394633 | 1729.4389826337463 | 1498.6200464866517 | 1130.5346933782735 | 1457.1214696833326 | 1200.5989330063549 | 1446.5280175772016 | 1232.0217367574485 | 1051.7113283693443 | 1106.1290051297358 |
