## Supplementary material for "Screening of clustered regulatory elements reveals functional cooperating dependencies in Leukemia": Table S2

**TableS2:Genesets negatively correlated with CRISPRi inhibition of SEKIT**

| GENESET_NAME | SIZE | ES | NES | NOM_p-val | FDR_q-val |
|---|---|---|---|---|---|
| KEGG_CYTOKINE_CYTOKINE_RECEPTOR_INTERACTION | 80 | -0.5101066 | -1.858465 | 0.006024096 | 0.15128882 |
| GO_CYTOKINE_BINDING | 45 | -0.5284298 | -1.7387868 | 0.019543974 | 0.18277322 |
| AP1_Q4 | 156 | -0.4322358 | -1.7331688 | 0.002853067 | 0.18397884 |
| GO_PLATELET_DERIVED_GROWTH_FACTOR_RECEPTOR_BINDING | 9 | -0.7660931 | -1.6965766 | 0.02097902 | 0.18942948 |
| REACTOME_SIGNALING_BY_PDGF | 81 | -0.46986297 | -1.6964734 | 0.016897082 | 0.18859418 |
| AP1_Q6 | 139 | -0.43164703 | -1.6909294 | 0.007587254 | 0.18844871 |
| GO_CYTOKINE_ACTIVITY | 51 | -0.48939356 | -1.6587768 | 0.02724359 | 0.18603332 |
| TGANTCA_AP1_C | 554 | -0.35058564 | -1.5819211 | 0.0001 | 0.20607616 |
| REACTOME_SIGNALING_BY_SCF_KIT | 68 | -0.4470207 | -1.5715805 | 0.042944785 | 0.20909369 |
| PDGF_UP.V1_DN | 44 | -0.47405526 | -1.5650235 | 0.04715447 | 0.21040736 |
| AP1_01 | 138 | -0.4016879 | -1.5597069 | 0.017595308 | 0.21429409 |
| GO_CYTOKINE_RECEPTOR_BINDING | 114 | -0.4026892 | -1.5483524 | 0.016467066 | 0.21501972 |
| REACTOME_PIP3_ACTIVATES_AKT_SIGNALING | 25 | -0.53417414 | -1.5436361 | 0.042016808 | 0.21708472 |
| REACTOME_PI3K_AKT_ACTIVATION | 32 | -0.502016 | -1.5235479 | 0.0461285 | 0.22794746 |
| AP1_Q2 | 143 | -0.3835286 | -1.5055175 | 0.023988007 | 0.2345746 |
| AP1_Q2_01 | 143 | -0.3823649 | -1.502686 | 0.021770682 | 0.23683275 |
| AP1_Q6_01 | 141 | -0.3752396 | -1.4760147 | 0.023809524 | 0.25127 |
| GO_CYTOKINE_RECEPTOR_ACTIVITY | 36 | -0.45751223 | -1.4679023 | 0.044701986 | 0.25599676 |
| AP1_C | 138 | -0.3734915 | -1.4593756 | 0.038402457 | 0.26026753 |
| REACTOME_REGULATION_OF_KIT_SIGNALING | 13 | -0.56535697 | -1.4172556 | 0.10175438 | 0.2858632 |
| PID_KIT_PATHWAY | 45 | -0.41782665 | -1.3744195 | 0.080314964 | 0.31436348 |
