## Supplementary material for "Screening of clustered regulatory elements reveals functional cooperating dependencies in Leukemia": Table S3

**Table S3: sequences of sgRNA used in single CRISPRi experiments**

| sgRNA ID | 20nt-sgRNA guide | 73mer sgRNA PCR oligos for cloning in pU6-sgRNA-EF1Alpha-puro-T2A-BFP using XhoI and BstXI | cloning in pLV-hU6-sgRNA-hUbC-dCas9 | |
|---|---|---|---|---|
| sgRNA2 | CAGATAATAGACTGGTCATG | GGAGAACCACCTTGTTGGCAGATAATAGACTGGTCATGGTTTTAGAGCTAGAAATAGCAAGTTAAAATAAGGC | Forward | CACCCAGATAATAGACTGGTCATG |
| | | | Reverse | AAACCATGACCAGTCTATTATCTG |
| sgRNA5 | TCCGTATAGCAAGTGTACCC | GGAGAACCACCTTGTTGGTCCGTATAGCAAGTGTACCCGTTTTAGAGCTAGAAATAGCAAGTTAAAATAAGGC | Forward | CACCTCCGTATAGCAAGTGTACCC |
| | | | Reverse | AAACGGGTACACTTGCTATACGGA |
| sgRNA7 | AAGGGTCTTATCTAACTACT | GGAGAACCACCTTGTTGGAAGGGTCTTATCTAACTACTGTTTTAGAGCTAGAAATAGCAAGTTAAAATAAGGC | Forward | CACCAAGGGTCTTATCTAACTACT |
| | | | Reverse | AAACAGTAGTTAGATAAGACCCTT |
| sgRNA8 | AGGGCCTTCCGCTAAAGGCA | GGAGAACCACCTTGTTGGAGGGCCTTCCGCTAAAGGCAGTTTTAGAGCTAGAAATAGCAAGTTAAAATAAGGC | Forward | CACCAGGGCCTTCCGCTAAAGGCA |
| | | | Reverse | AAACTGCCTTTAGCGGAAGGCCCT |
| sgRenilla | GGAACACGGCCGTATTAGGG | GGAGAACCACCTTGTTGGGGAACACGGCCGTATTAGGGGTTTTAGAGCTAGAAATAGCAAGTTAAAATAAGGC | Forward | CACCGGAACACGGCCGTATTAGGG |
| | | | Reverse | AAACCCCTAATACGGCCGTGTTCC |
| sgRNA3 | AGACCCTTGCCTTTAGCGGA | GGAGAACCACCTTGTTGGAGACCCTTGCCTTTAGCGGAGTTTTAGAGCTAGAAATAGCAAGTTAAAATAAGGC | | |
| sgGFP | GACCAGGATGGGCACCACCC | GGAGAACCACCTTGTTGGGACCAGGATGGGCACCACCCGTTTTAGAGCTAGAAATAGCAAGTTAAAATAAGGC | | |
