## Supplementary material for "Screening of clustered regulatory elements reveals functional cooperating dependencies in Leukemia": Table S4

**Table S4: sequences of shRNA used in gene knock-down experiments**

| Gene Symbol | shRNA ID | 22nt-shRNA guide | mRNA target site | 97mer oligo |
|---|---|---|---|---|
| KIT | shKIT1 | TTCTGCTCAGACATCGTCGTGC | GCACGACGATGTCTGAGCAGAA | TGCTGTTGACAGTGAGCGACACGACGATGTCTGAGCAGAATAGTGAAGCCACAGATGTATTCTGCTCAGACATCGTCGTGCTGCCTACTGCCTCGGA |
| KIT | shKIT3 | TTATCTACTACTTCCAAGGTTG | CAACCTTGGAAGTAGTAGATAA | TGCTGTTGACAGTGAGCGAAACCTTGGAAGTAGTAGATAATAGTGAAGCCACAGATGTATTATCTACTACTTCCAAGGTTGTGCCTACTGCCTCGGA |
| PDGFRA | shPDGFRA1 | TTAAACATCAACCAAGTCCTCC | GGAGGACTTGGTTGATGTTTAA | TGCTGTTGACAGTGAGCGAGAGGACTTGGTTGATGTTTAATAGTGAAGCCACAGATGTATTAAACATCAACCAAGTCCTCCTGCCTACTGCCTCGGA |
| PDGFRA | shPDGFRA3 | ATACTGTGTAGTATCAGCCTGC | GCAGGCTGATACTACACAGTAT | TGCTGTTGACAGTGAGCGACAGGCTGATACTACACAGTATTAGTGAAGCCACAGATGTAATACTGTGTAGTATCAGCCTGCTGCCTACTGCCTCGGA |
