## Supplementary material for "Screening of clustered regulatory elements reveals functional cooperating dependencies in Leukemia": Table S5

**Table S5: sequences of primers used in quantitative PCR experiments**

| Gene Symbol | Primer ID | Primer sequence |
|---|---|---|
| KIT | hKIT fwd | CAATTCTGACGTCAATGCTGCCA |
| | hKIT rev | GAGCATGCCATTCACGAGCC |
| PDGFRA | hPDGFRA fwd | CCTGTCCCTGAGGAGGAGGAC |
| | hPDGFRA rev | GAAGGTGGAACTGCTGGAACCC |
| CSF2RA | hCSF2RA fwd | TCCCGCCAGTTCCACAGATC |
| | hCSF2RA rev | GACCTCTTCGCGGTAGCCTT |
| CSF2RB | hCSF2RB fwd | AACGGGATCTGGAGCGAGTG |
| | hCSF2RB rev | AGATCACGATGAGGGCCAGC |
| GAPDH | hGAPDH fwd | CTTTTGCGTCGCCAGCCGAG |
| | hGAPDH rev | CCAGGCGCCCAATACGACCA |
